# Full-length 16S rRNA gene amplicon analysis of human gut microbiota using MinION™ nanopore sequencing confers species-level resolution

**DOI:** 10.1101/2020.05.06.078147

**Authors:** Yoshiyuki Matsuo, Shinnosuke Komiya, Yoshiaki Yasumizu, Yuki Yasuoka, Katsura Mizushima, Tomohisa Takagi, Kirill Kryukov, Aisaku Fukuda, Yoshiharu Morimoto, Yuji Naito, Hidetaka Okada, Hidemasa Bono, So Nakagawa, Kiichi Hirota

## Abstract

**Background:** Species-level genetic characterization of complex bacterial communities has important clinical applications in both diagnosis and treatment. Amplicon sequencing of the 16S ribosomal RNA (rRNA) gene has proven to be a powerful strategy for the taxonomic classification of bacteria. This study aims to improve the method for full-length 16S rRNA gene analysis using the nanopore long-read sequencer MinION™. We compared it to the conventional short-read sequencing method in both a mock bacterial community and human fecal samples.

**Results:** We modified our existing protocol for full-length 16S amplicon sequencing by MinION™. A new strategy for library construction with an optimized primer set overcame PCR-associated bias and enabled taxonomic classification across a broad range of bacterial species. We compared the performance of full-length and short-read 16S amplicon sequencing for the characterization of human gut microbiota with a complex bacterial composition. The relative abundance of dominant bacterial genera was highly similar between full-length and short-read sequencing. At the species level, MinION™ long-read sequencing had better resolution for discriminating between members of particular taxa such as *Bifidobacterium*, allowing an accurate representation of the sample bacterial composition.

**Conclusions:** Our present microbiome study, comparing the discriminatory power of full-length and short-read sequencing, clearly illustrated the analytical advantage of sequencing the full-length 16S rRNA gene, which provided the requisite species-level resolution and accuracy in clinical settings.

## Background

Recent advances in DNA sequencing technology have had a revolutionary impact on clinical microbiology [1]. Next-generation sequencing (NGS) technology enables parallel sequencing of DNA on a massive scale to generate vast quantities of accurate data. NGS platforms are now increasingly used in the field of clinical research [2]. Metagenomic sequencing offers numerous advantages over traditional culture-based techniques that have long been the standard test for detecting pathogenic bacteria. This method is particularly useful for characterizing uncultured bacteria and novel pathogens [3].

Among sequence-based bacterial analyses, amplicon sequencing of the 16S ribosomal RNA (rRNA) gene has proven to be a reliable and efficient option for taxonomic classification [4, 5]. The bacterial 16S rRNA gene contains nine variable regions (V1 to V9) that are separated by highly conserved sequences across different taxa. For bacterial identification, the 16S rRNA gene is first amplified by polymerase chain reaction (PCR) with primers annealing to conserved regions and then sequenced. The sequencing data are subjected to bioinformatic analysis in which the variable regions are used to discriminate between bacterial taxa [6].

Since the conventional parallel-type short-read sequencer cannot yield reads covering the full length of the 16S rRNA gene [7], several regions of it have been targeted for sequencing, which often causes ambiguity in taxonomic classification [8]. New sequencing platforms have overcome these technical restrictions, particularly those affecting read length. A prime example is the MinION™ sequencer from Oxford Nanopore Technologies, which is capable of producing long sequences with no theoretical read length limit [9-11]. MinION™ sequencing targets the entire 16S rRNA gene, allowing the identification of bacteria with more accuracy and sensitivity [12, 13]. Furthermore, MinION™ produces sequencing data in real time, which reduces turnaround time for data processing [14, 15].

Given these features of MinION™ sequencing, we had previously conducted full-length 16S amplicon sequencing analyses using the MinION™ platform coupled to a bioinformatics pipeline, which allowed us to identify bacterial pathogens with a total analysis time of under two hours [16]. However, we also found that the approach of using the commercial 16S Barcoding Kit (SQK-RAB204) available from Oxford Nanopore Technologies has a limited ability to detect particular taxa such as *Bifidobacterium* [16]. This is probably due to sequence mismatches in the primer used for 16S gene amplification [17]. Deviations or aberrancies in the *Bifidobacterium* composition in the human gut have been reported in several diseases including obesity, allergy, and inflammatory disorders [18]. Based on their putative health-promoting effects, several strains of *Bifidobacterium* have been utilized as probiotics [19]. Within these contexts, the species-level characterization of *Bifidobacterium* diversity in human gut microbiota is potentially important in clinical practice.

Our 16S rRNA gene sequence analysis using MinION™ has been tested only with pre-characterized mock bacterial DNA and a limited number of pathogenic bacteria from a patient-derived sample [20]. Its applicability to highly complex bacterial communities has not yet been thoroughly investigated. Therefore, in this study we modified our existing protocol for 16S amplicon sequencing by MinION™ and applied it to human gut microbiota with a complex bacterial composition [21], including *Bifidobacterium*, to determine whether full-length 16S rRNA gene sequencing with MinION™ is an effective characterization tool.

## Results

### Classification of the mock bacterial community

The 16S rRNA gene sequence of *Bifidobacterium* has three base mismatches with the 27F forward primer provided in the commercial sequencing kit (16S Barcoding Kit, SQK-RAB204, Oxford Nanopore Technologies; Additional File 2: Supplementary Fig. S1a), which biases amplification toward underrepresentation of *Bifidobacterium* species (Additional File 2: Supplementary Fig S2, Additional File 3: Supplementary Table S1-S3). To overcome this drawback, we introduced three degenerate bases to the 16S rRNA gene-specific sequences of the primer (Additional File 2: Supplementary Fig. S1b). The competence of the modified primer set was then evaluated by 16S rRNA gene sequence analysis of a ten-species mock community. The V1-V9 region of the 16S rRNA gene was amplified by the four-primer PCR method with the rapid adapter attachment chemistry and sequenced (Fig. 1a). MinION™ sequencing generated 8651 pass reads (Table 1). Following adapter trimming and size selection, 6972 reads (80.6% of pass reads with an average lead length of 1473 bp) were retained for bacterial identification. Full-length 16S amplicon sequencing with the modified primer set led to the successful identification of all the ten bacterial genera, including *Bifidobacterium* (Fig. 1b, Additional File 3: Supplementary Table S4). At the species level, 92.5% of analyzed reads were correctly assigned to each bacterial taxon included in the mock community, demonstrating the excellent discriminatory power of this full-length sequencing method for species identification (Fig. 1c). The only exception was *Bacillus cereus*. Discrimination of *Bacillus cereus* from the closely related species such as *Bacillus anthracis* and *Bacillus thuringiensis* was not fully achieved with the 16S gene analysis using MinION™ (Additional File 4).

**Table 1.**
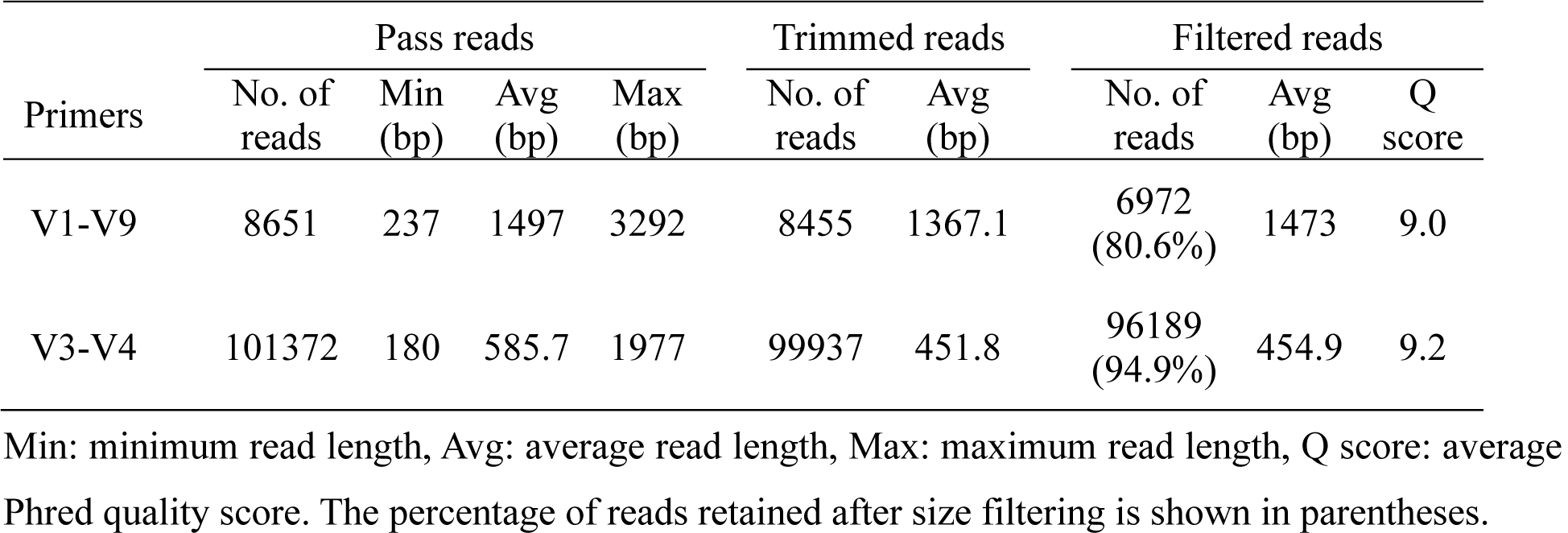
MinION™ sequencing statistics for the mock community sample

**Fig. 1.**
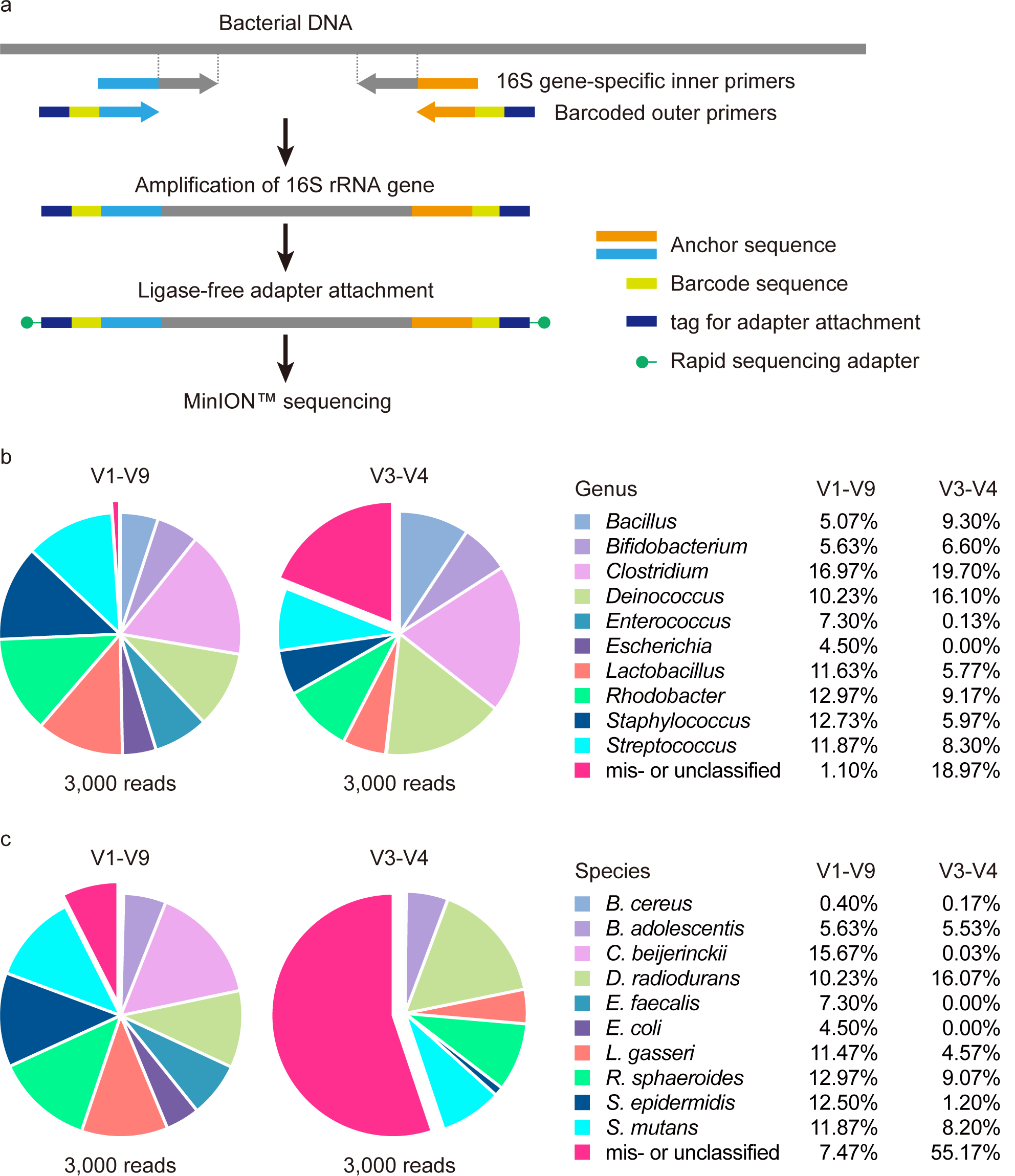
16S rRNA gene sequence analysis using the MinION™ nanopore sequencer. **a** Workflow of 16S rRNA amplicon sequencing on the MinION™ platform. Sequencing libraries are generated by the four-primer PCR-based strategy, enabling simplified post-PCR adapter attachment. At the initial stage of PCR, the 16S rRNA gene is amplified with the inner primer pairs. The resulting PCR products are targeted for amplification with the outer primers to introduce the barcode and tag sequences at both ends, to which adapter molecules can be attached in a single-step reaction. **b, c** Taxonomic assignments of a mock community analyzed by MinION™ sequencing. The V1-V9 or V3-V4 region of the 16S rRNA gene was amplified from a pre-characterized mock community sample comprising ten bacterial species and sequenced on the MinION™ platform. Three thousand reads were randomly selected from the processed data set and aligned directly to the reference genome database of 5850 representative bacterial species. The pie charts represent taxonomic profiles at the (b) genus and (c) species levels. Slices corresponding to misclassified (assigned to bacteria not present in the mock community) or unclassified (not classified at the species level but placed in a higher taxonomic rank) reads are exploded. The relative abundance (%) of each taxon is shown.

We compared the resolution of full-length and short-read 16S amplicon sequencing for the taxonomic classification of bacteria. The V3-V4 region was amplified by four-primer PCR from the ten-species mock community DNA, and the samples were sequenced on MinION™. After removing the adapter/barcode sequences and filtering reads by length, 96189 reads with an average length of 454.9 bp for downstream analysis were yielded (Table 1). In contrast to full-length sequencing with the highest resolution, a significant number of V3-V4 reads could not be classified down to the genus level, but could only be assigned to a higher taxonomic rank (Fig. 1b, 1c). Most reads derived from *Enterococcus faecalis* and *Escherichia coli* were not assigned down to each taxon, as more than two species produced the same similarity score for the V3-V4 sequence read queries (Additional File 5). Such reads were classified to the parent taxon, as more specific classification was impossible (Additional File 3: Supplementary Table S5). The classifications were not affected by increasing the number of analyzed reads to 10000 (Additional File 2: Supplementary Fig. S3, Additional File 3: Supplementary Table S6).

For eight of the ten bacterial strains constituting the mock community, each subset of V1-V9 sequencing reads classified to the specific genus were assigned with almost complete accuracy (98.2-100%) to the corresponding species (Fig. 2). V3-V4 short-read sequencing showed a discriminatory power comparable to that of V1-V9 full-length sequencing in the classification of three genera (*Deinococcus, Rhodobacter*, and *Streptococcus*) with more than 98% of reads correctly assigned. However, the V3-V4 region was not suitable for species-level identification of other taxa, such as *Clostridium* and *Staphylococcus*. Only 0.2% of the V3-V4 reads belonging to the genus *Clostridium* were assigned to *Clostridium beijerinckii*, a component of the mock community. In contrast, 92.3% of *Clostridium* reads obtained from V1-V9 full-length sequencing were correctly classified at the species level. These results suggest a lower resolution of the V3-V4 region for species-level classification, emphasizing the advantage of long-read sequencing for obtaining an accurate representation of the sample bacterial composition.

**Fig. 2.**
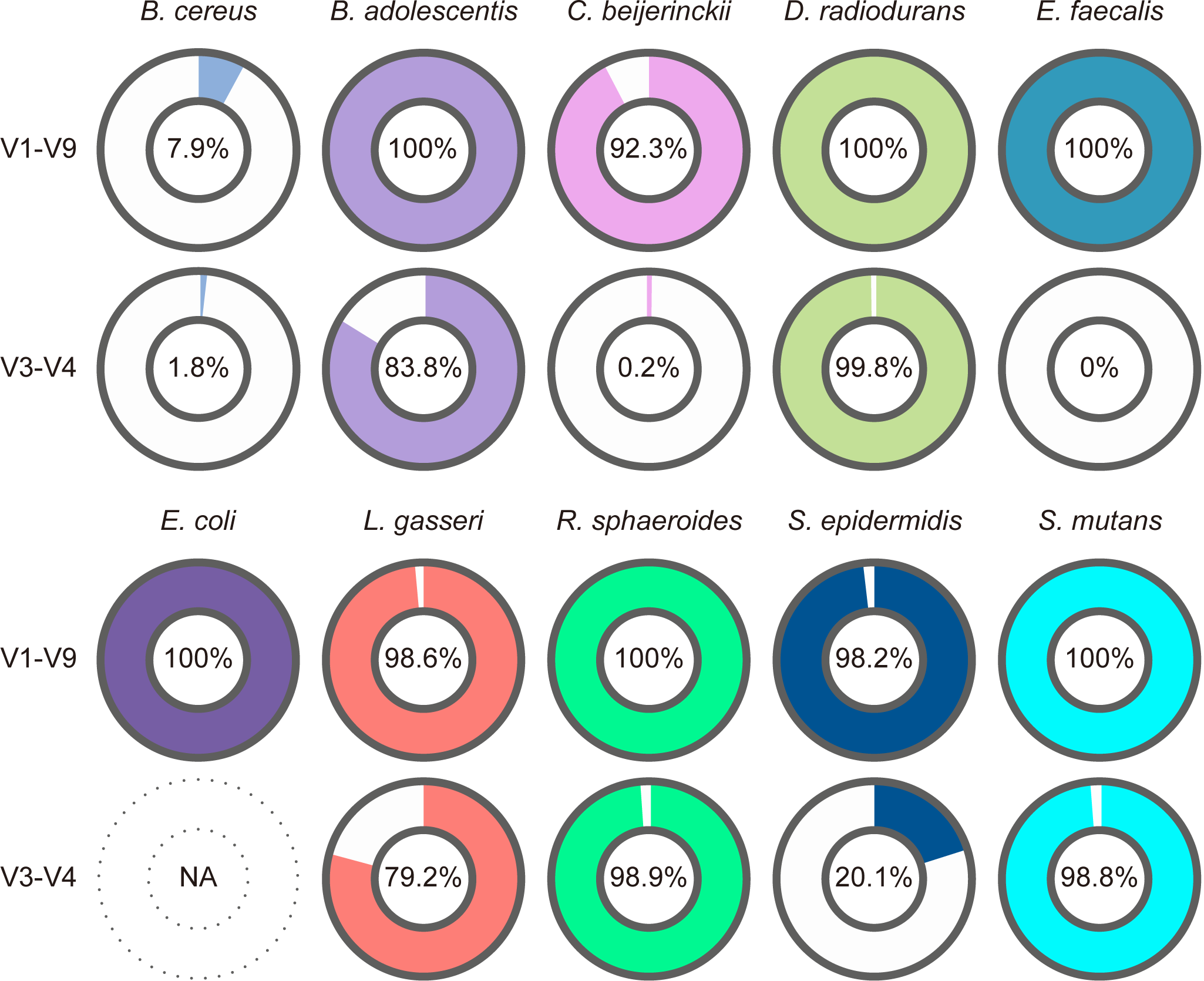
Accurate taxonomic assignment afforded by full-length MinION™ sequencing of the 16S rRNA gene. Classification accuracy compared between full-length (V1-V9) and partial (V3-V4) 16S sequencing data obtained from composition profiling of the ten-species mock community. The donut charts show the proportions of reads correctly assigned to the species constituting the mock community. The percentage of correctly classified reads is shown in the center hole. NA: not assigned (no reads were classified in *Escherichia* genus).

### Classification of human fecal bacteria

We assessed the performance of our full-length 16S amplicon sequencing approach in the context of a highly complex bacterial community. The V1-V9 region was amplified by four-primer PCR from six human fecal samples (F1-F6) and analyzed by MinION™ sequencing. (Table 2). In Fig. 3, the numbers of species detected are plotted against the numbers of reads analyzed. The curve started to plateau at around 20000 reads. There was a highly significant correlation between the read numbers 20000 and 30000 (Pearson’s correlation coefficient *r* > 0.999, Additional File 6: Supplementary Table S7). Based on these observations, randomly sampled 20000 reads were used in further analysis to determine the bacterial composition of the human gut.

**Table 2.**
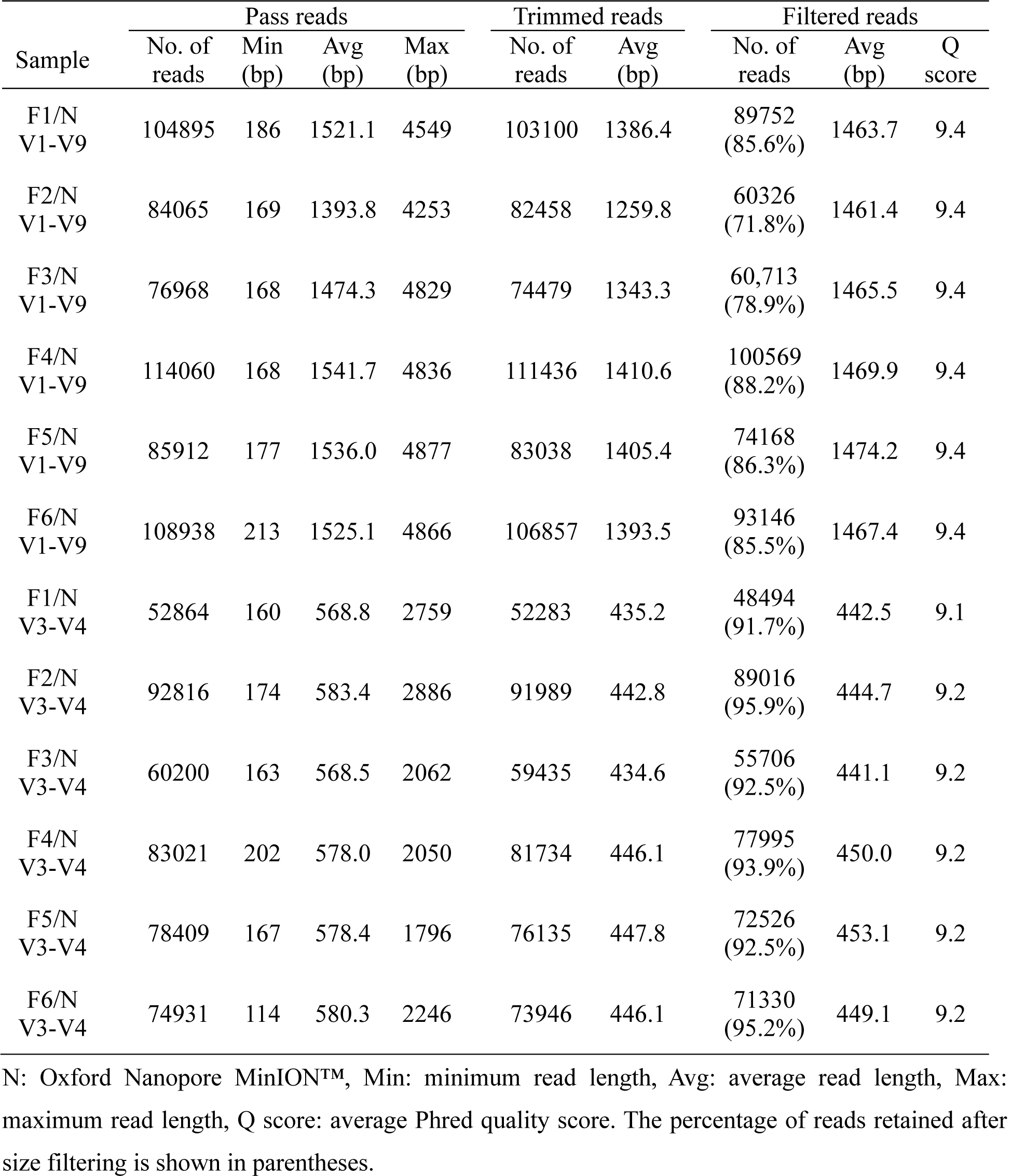
Statistics of MinION™ sequencing data for human fecal samples

**Fig. 3.**
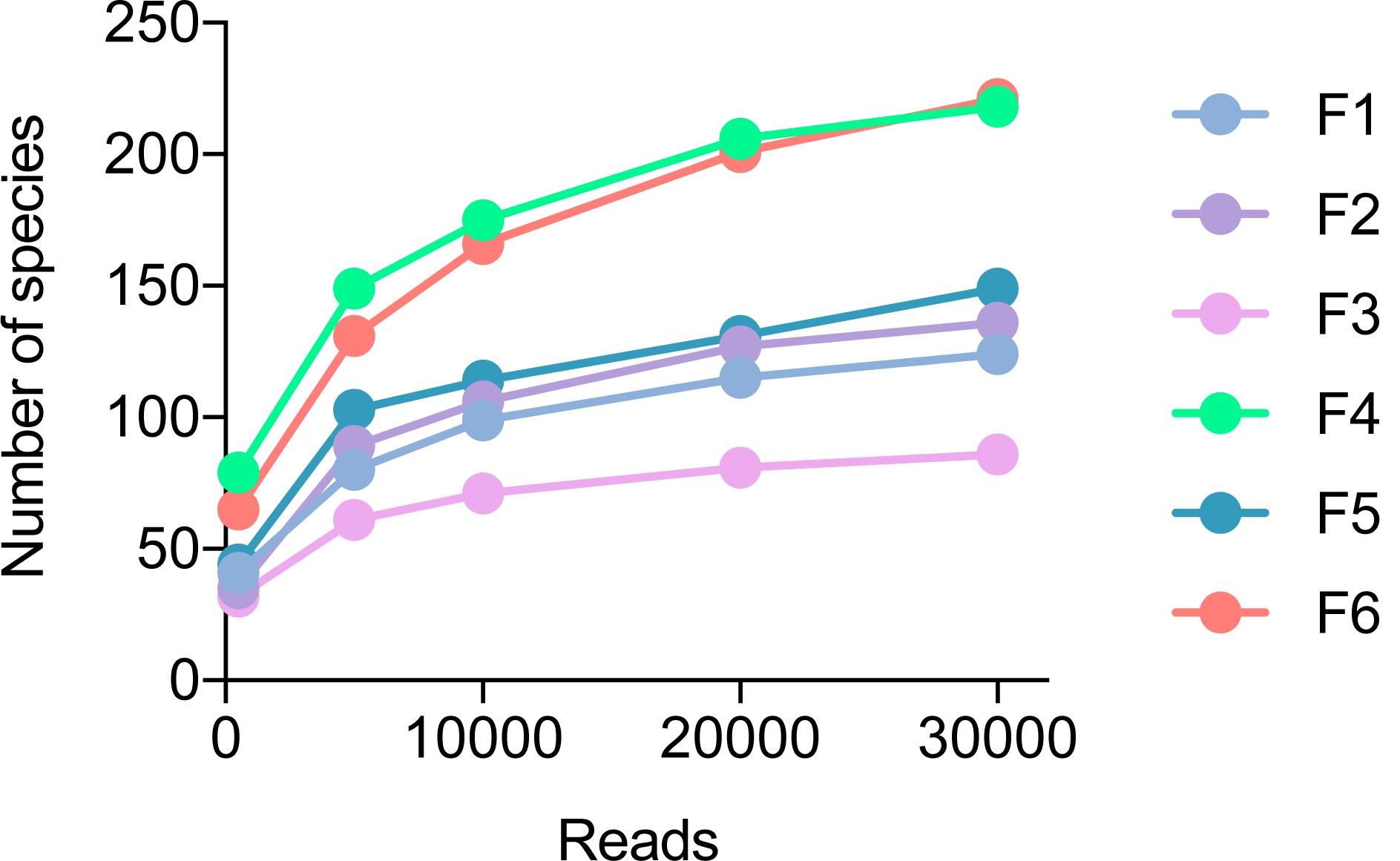
16S rRNA gene sequence analysis of human gut microbiota. Six human fecal samples (F1-F6) were subjected to full-length 16S rRNA amplicon sequencing via MinION™. Numbers of detected species are plotted against numbers of reads used for taxonomic classification.

For comparison, amplicon sequencing of the V3-V4 region was also conducted using the MinION™ (Table 2) and the Illumina MiSeq™ platform (Table 3). The processed reads from each data set were allocated to the reference bacterial genome using our bioinformatics pipeline to determine the bacterial compositions (Additional File 7 for V1-V9 MinION™ sequencing, Additional File 8 for V3-V4 MinION™ sequencing, and Additional File 9 for V3-V4 MiSeq™ sequencing). From MiSeq™ sequencing data, the bacterial composition was also analyzed by the operational taxonomic unit (OTU)-based approach using the QIIME 2 (ver. 2019.7) pipeline (Additional File 2: Supplementary Fig. S4, Additional File 10) [22, 23]. Although *Bacteroides* was underrepresented in the OTU-based analysis, the two analytical methods (our bioinformatics pipeline and OTU-based method) produced similar taxonomic profiles in the dominant phylotypes for the MiSeq™ data. This result confirmed the validity of our method for the taxonomic classification of the bacterial community.

**Table 3.**
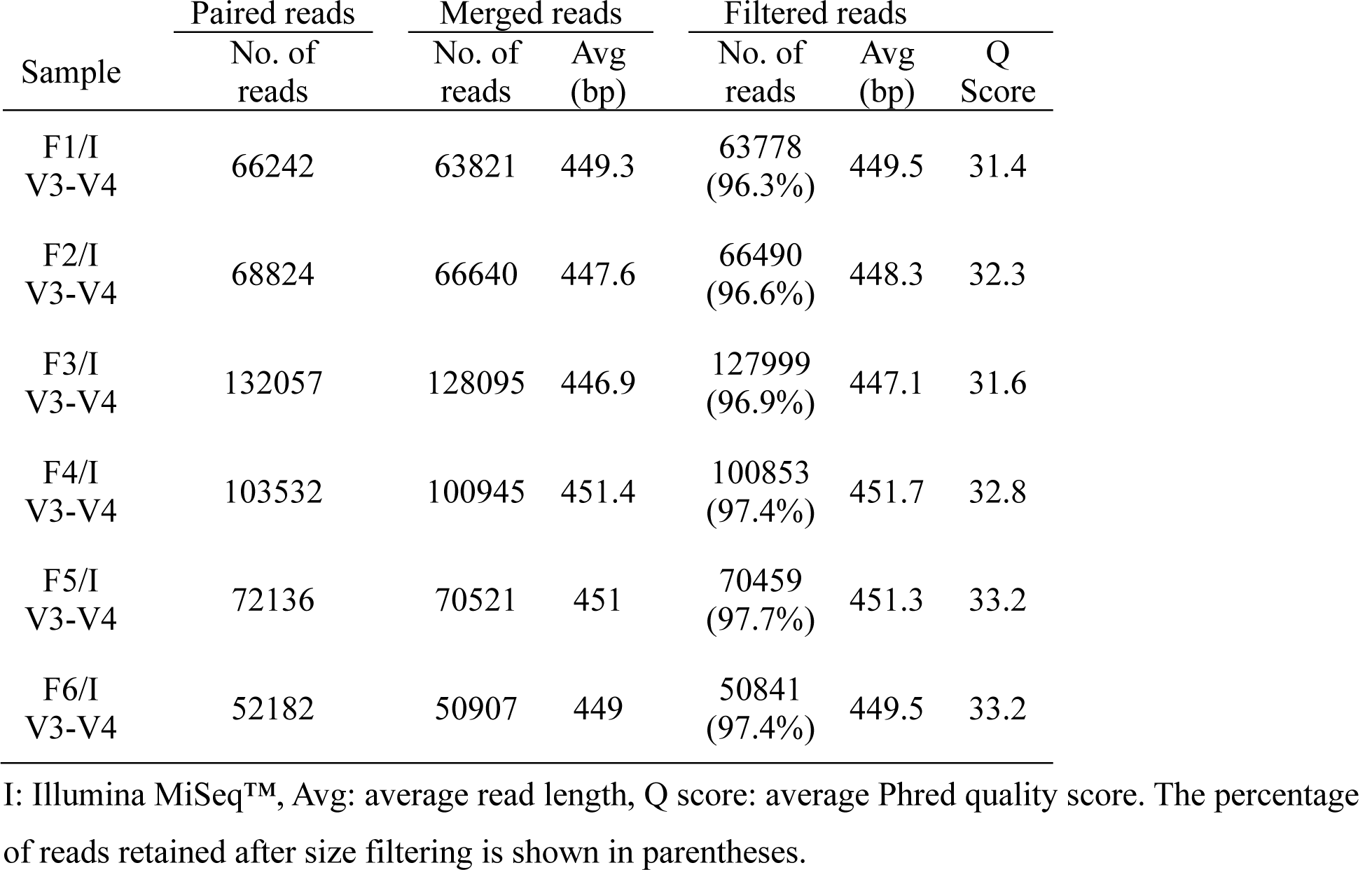
Statistics of MiSeq™ sequencing data for human fecal samples

The three sequencing methods (V1-V9 MinION™ sequencing, V3-V4 MinION™ sequencing, and V3-V4 MiSeq™ sequencing) revealed similar profiles for the six fecal samples at the genus level (Fig. 4). Statistically significant similarities have been found in the relative genus abundances across these sequencing methods. Thus, at the genus level, V1-V9 full-length MinION™ sequencing exhibited a discriminatory power comparable to that of high-quality short-read sequencing with MiSeq™ technology.

**Fig. 4.**
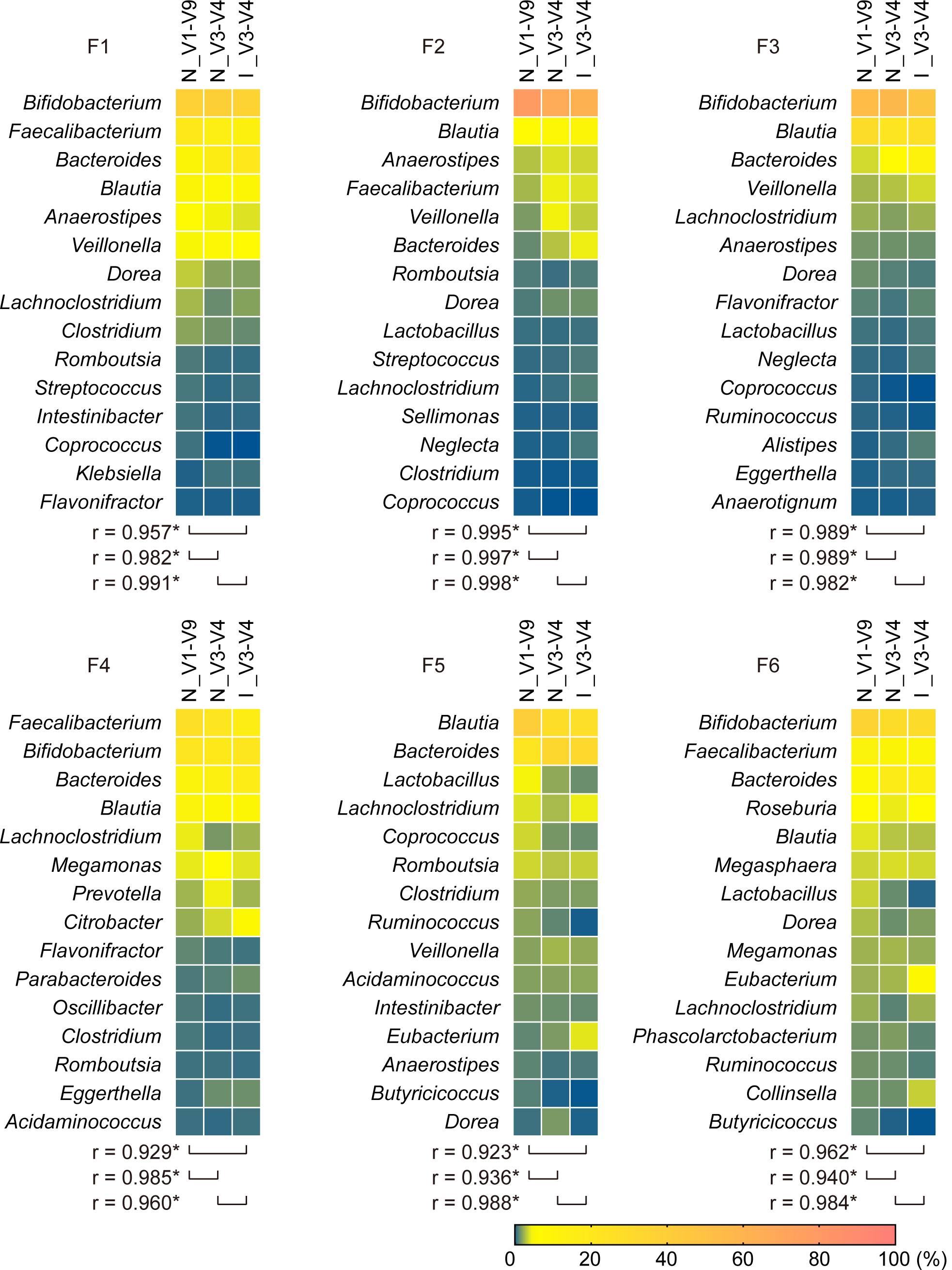
Comparison of taxonomic profiles of human gut microbiota between sequencing methodologies. Six fecal samples (F1-F6) were analyzed by sequencing the entire 16S rRNA gene using MinION™ (N_V1-V9). For comparison, the V3-V4 region was sequenced on MinION™ (N_V3-V4) or MiSeq™ platforms (I_V1-V9). Randomly sampled 20000 reads from each data set were allocated to the reference genome database of 5850 representative bacterial species. A heat map shows the relative genus abundance (%) of classified reads. The 15 most abundant taxa are shown. The Pearson correlation coefficient (*r*) between sequencing methods was computed. Asterisks indicate significant correlations at *P* < 0.05.

### The species-level taxonomic resolution achieved by full-length sequencing of the 16S rRNA gene using MinION™

While genus classification using long versus short reads was relatively comparable, we observed considerable differences across amplified regions in the species-level profiling of human gut microbiota. As shown in Fig. 5, the number of ambiguous reads that were not assigned to species but could be classified at a higher level was significantly greater in the V3-V4 data set in comparison than in the V1-V9 data set.

**Fig. 5.**
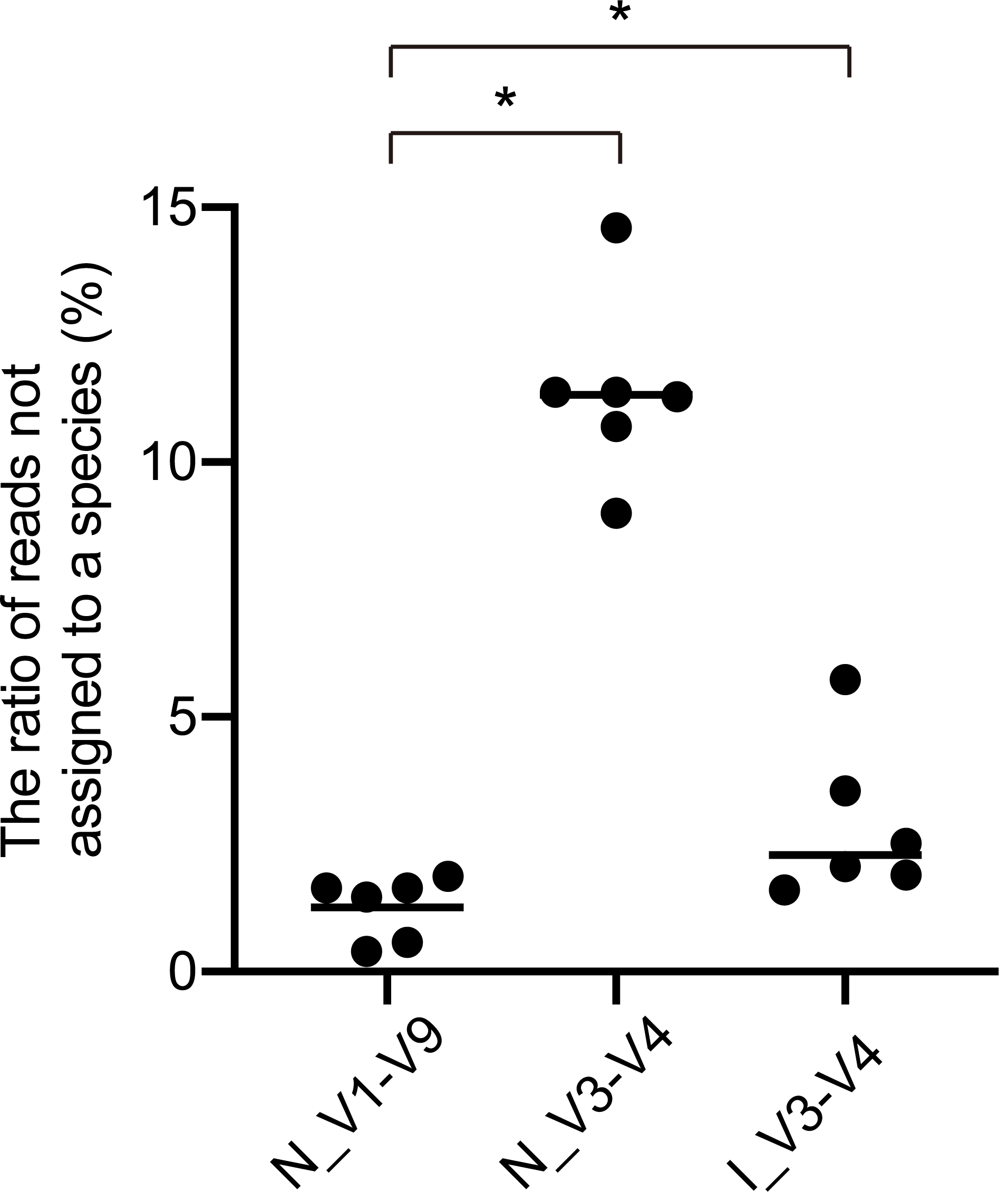
Comparison of taxonomic resolution. The percentages of ambiguous reads not assigned to the species level are plotted for six fecal samples analyzed by MinION™ (N_V1-V9 and N_V3-V4) or MiSeq™ (I_V3-V4). Horizontal bars represent mean values. * *P* < 0.05 (statistically significant).

When species compositions of the dominant taxa (*Bifidobacterium, Blautia*, and *Bacteroides*) were analyzed, the V1-V9 and V3-V4 sequencing produced comparable results for *Blautia* (Additional File 2: Supplementary Fig. S5, Additional File 6: Supplementary Table S8) and *Bacteroides* genus (Additional File 2: Supplementary Fig. S6, Additional File 6: Supplementary Table S9) in most of the fecal samples. For *Bifidobacterium*, there appeared to be considerable deviations in the relative abundances of some species depending on the sequencing method used (Fig. 6, Additional File 6: Supplementary Table S10). Notably, most of the *Bifidobacterium* reads generated by V1-V9 MinION™ sequencing were classified into the *Bifidobacterium* species that were isolated from human sources [18, 24]. A significant number of the V3-V4 reads, however, were assigned to *Bifidobacterium* species of non-human origin (Additional File 2: Supplementary Fig. S7). From V3-V4 MiSeq™ sequencing data, the OTU-based classification analysis using the QIIME 2 pipeline also revealed a lower resolution of short-read sequencing for taxonomic separation of *Bifidobacterium* genus. Except for *Bifidobacterium longum, Bifidobacterium* species could not be reliably identified by the V3-V4 sequencing strategy (Additional File 2: Supplementary Fig. S8). These results suggest that species classification of *Bifidobacterium* based on V3-V4 sequencing can potentially lead to misidentification and biased community profiling that lacks biological significance.

**Fig. 6.**
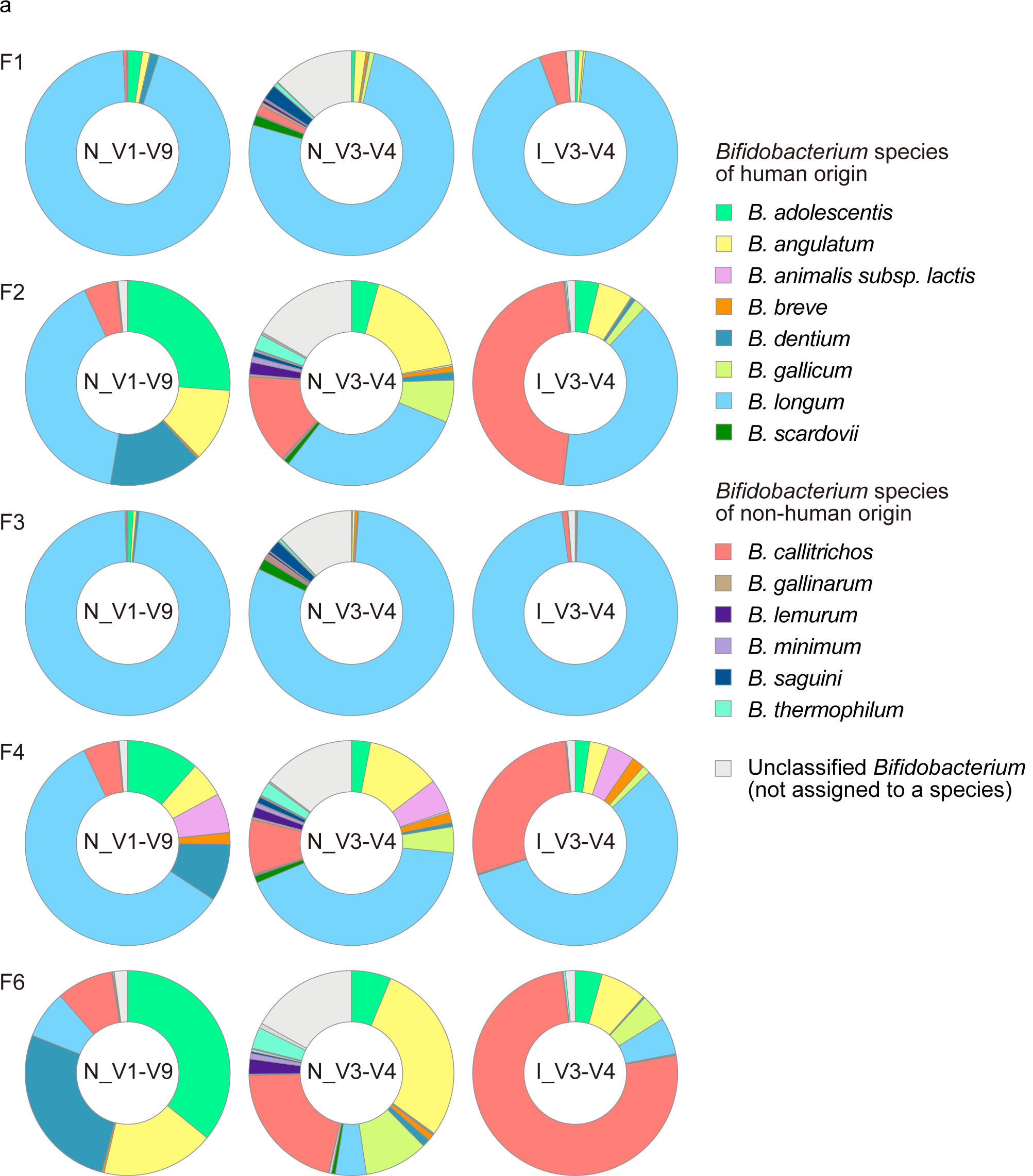
Species composition of *Bifidobacterium* in six fecal samples. MinION™ V1-V9 sequencing confers species-level resolution for bacterial composition profiling. Results obtained by the three sequencing methods are shown. The legends show the 14 most abundant *Bifidobacterium* species.

## Discussion

16S rRNA amplicon sequencing is a powerful strategy for taxonomic classification of bacteria and has been extensively employed for analyzing samples from environmental and clinical sources [5, 25, 26]. We assessed the performance of MinION™ sequencing by comparing the resolution of the V1-V9 and V3-V4 reads for the taxonomic classification of bacteria. Due to the error-prone nature of MinION™ sequencing, the existing OTU-based approach, requiring at least 97% sequence identity threshold, has been considered unsuitable for taxonomic classification of MinION™ reads [27, 28].

Instead, the reads were analyzed by the direct read mapping method that assigns sequences to taxonomic bins based on the similarity to a reference database [14, 15]. Long-read MinION™ sequencing with the optimized primer set successfully identified *Bifidobacterium* species leading to a better representation of the species composition of the mock community. For improving the classification results, the reads were filtered by length to eliminate those outside the expected size range. Typically, extremely short reads possess only one primer-binding site, suggesting that they are derived from incomplete sequencing. There also exist unexpectedly longer reads with a continuous sequence structure in which two 16S amplicons are linked end-to-end. Because these reads can potentially result in unclassified reads or misclassification, they were eliminated before alignment to the reference sequences of the bacterial genome. We also modified library construction for MinION™ sequencing with a four-primer PCR strategy, which enabled ligase-free adapter attachment to occur in a single-step reaction. The four-primer PCR generates amplicons with particular chemical modifications at the 5’ ends to which adapter molecules can be attached non-enzymatically. Unlike the ligation-based approach, the PCR products amplified by the four-primer method are subjected directly to the adapter attachment reaction without repairing their 5’ ends, substantially reducing the time required for sample preparation. Furthermore, because the protocol is free of Nanopore’s transposase-based technology (e.g. Rapid Sequencing Kit, SQK-RAD004) that cleaves DNA molecules to produce chemically modified ends for library construction, the PCR products are kept intact, enabling sequencing of the entire amplified region. Thus, the four-primer PCR-based method allowed us to perform amplicon sequencing on the MinION™ platform with user-defined arbitrary primer pairs, taking advantage of the rapid adapter attachment chemistry. This method can be applied to a wide range of sequence-based analyses, including detection of functional genetic markers like antimicrobial resistance genes and identification of genetic variations in targeted loci [11, 29, 30].

Our present microbiome study, comparing the discriminatory power of the V1-V9 and V3-V4 reads sequenced on the MinION™ platform, clearly illustrated the advantage of sequencing the entire 16S rRNA gene. The full-length 16S gene sequencing provided better resolution than short-read sequencing for discriminating between members of certain bacterial taxa, including *Clostridium, Enterococcus, Escherichia*, and *Staphylococcus*. Consistently, comprehensive *in silico* experiments using sequencing a sequencing data set consisting of different regions of the 16S rRNA gene have shown that the choice of the regions to be sequenced substantially affects the classification results [6, 31]. As shown here and in previous publications, short-read sequencing of the 16S rRNA gene may be a reasonable option for providing a rough estimation of bacterial diversity. However, it was not suited for analysis requiring species-level resolution and accuracy, which was afforded by sequencing the entire 16S rRNA gene. It is important to note that even full-length 16S gene analysis fails to discriminate some closely related species such as members of *Bacillus cereus* group, indicating the limitations of the 16S amplicon sequencing alone in species allocation. Long read sequencing targeting other phylogenetic markers may be an alternative to 16S rRNA amplicon sequencing and provide better resolution for bacterial identification. In the analysis of the human fecal samples, we used the taxonomic resolution of the V3-V4 region sequenced with MiSeq™, which generates highly accurate reads, as a benchmark for the taxonomic resolution of the full-length 16S gene sequenced with MinION™. The relative abundance of dominant bacterial taxa was highly similar at the genus level between full-length MinION™ and short-read MiSeq™ sequencing. Despite the lower read quality, the full-length sequencing by MinION™ enabled reliable identification of bacterial genera with an accuracy comparable to MiSeq™ technology. The MiSeq™ platform enables 16S rRNA gene sequencing on a massive scale with reduced cost (approximately 20 USD per sample). Considering a low equipment price (1000 USD) and an affordable per-run cost (approximately 50 USD per sample), the MinION™ sequencer could be a viable option for practical applications in clinical microbiology.

At the species level, MinION™ long-read sequencing had better resolution for accurate identification of the composition of human gut microbiota. Composition profiles of *Bifidobacterium*, one of the dominant genera present in the human gut [32], appeared to differ considerably between the two sequencing platforms. While most MinION™ V1-V9 reads were assigned to *Bifidobacterium* species of human origin, a significant number of the MiSeq™ V3-V4 reads were assigned to non-human *Bifidobacterium* species [24]. Such improbable errors in species classification may be attributed to the lower resolution provided by the V3-V4 region, but in fact, 16S rRNA gene sequence analysis of mouse gut microbiota has revealed that V3-V4 reads sequenced on MiSeq™ are not well-suited for classifying *Bifidobacterium* species, consistent with our findings [33]. Since the taxonomic assignment is a critical step for analyzing the bacterial diversity and community composition, the reference database quality and the alignment algorithm must be further evaluated for each sequencing data set.

### Conclusions

Our modified protocol for 16S amplicon sequencing overcame known limitations, such as the primer-associated bias toward the underrepresentation of *Bifidobacterium*, and enabled taxonomic classification across a broad range of bacterial species. Benchmarking with MiSeq™ sequencing technology demonstrated the analytical advantage of sequencing the full-length 16S gene with MinION™, which provided the requisite species-level resolution and accuracy. With the recent progress in nanopore sequencing chemistry and base-calling algorithms, sequencing accuracy is continuously improving [34, 35]. This will soon enable us to exploit the full potential of MinION™ long-read sequencing technology. High-quality long sequences will allow better discrimination between closely related species, and even bacterial strains, in sequence-based bacterial analyses.

## Methods

### Mock bacterial community DNA

A mixture of bacterial DNA (10 Strain Even Mix Genomic Material, MSA-1000) was obtained from the American Type Culture Collection (ATCC, Manassas, VA, USA), comprising genomic DNA prepared from the following ten bacterial strains: *Bacillus cereus* (ATCC 10987), *Bifidobacterium adolescentis* (ATCC 15703), *Clostridium beijerinckii* (ATCC 35702), *Deinococcus radiodurans* (ATCC BAA-816), *Enterococcus faecalis* (ATCC 47077), *Escherichia coli* (ATCC 700926), *Lactobacillus gasseri* (ATCC 33323), *Rhodobacter sphaeroides* (ATCC 17029), *Staphylococcus epidermidis* (ATCC 12228), and *Streptococcus mutans* (ATCC 700610).

### Fecal DNA

DNA was extracted from six human fecal samples using the NucleoSpin^®^ Microbial DNA Kit (Macherey-Nagel, Düren, Germany), as described previously [36]. Briefly, human feces stored using the Feces Collection Kit (Techno Suruga Lab, Shizuoka, Japan) were subjected to mechanical disruption by bead-beating, and DNA was isolated using silica membrane spin columns. Extracted DNA was purified with the Agencourt AMPure^®^ XP (Beckman Coulter, Brea, CA, USA).

### 16S rRNA gene sequencing on the MinION™ platform

Four-primer PCR with rapid adapter attachment chemistry generated 16S gene amplicons with modified 5′ ends for simplified post-PCR adapter attachment following the manufacturer’s instructions with slight modifications. For amplification of the V1-V9 region of the 16S rRNA gene, the following inner primers were used, with 16S rRNA gene-specific sequences underlined: forward primer (S-D-Bact-0008-c-S-20 [37]) with anchor sequence 5′-TTTCTGTTGGTGCTGATATTGC<u>AGRGTTYGATYMTGGCTCAG</u>-3′ and reverse primer (1492R) with anchor sequence 5′-ACTTGCCTGTCGCTCTATCTTC<u>CGGYTACCTTGTTACGACTT</u>-3′. For amplification of the V3-V4 region, the following inner primers were used, with 16S rRNA gene-specific sequences underlined: 341F with anchor sequence 5′-TTTCTGTTGGTGCTGATATTGC<u>CCTACGGGNGGCWGCAG</u>-3′ and 806R with anchor sequence 5′-ACTTGCCTGTCGCTCTATCTTC<u>GGACTACHVGGGTWTCTAAT</u>-3′. PCR amplification of 16S rRNA genes was conducted using the KAPA2G™ Robust HotStart ReadyMix PCR Kit (Kapa Biosystems, Wilmington, MA, USA) in a total volume of 25 μl containing inner primer pairs (50 nM each) and the barcoded outer primer mixture (3%) from the PCR Barcoding Kit (SQK-PBK004; Oxford Nanopore Technologies, Oxford, UK). Amplification was performed with the following PCR conditions: initial denaturation at 95 °C for 3 min, 5 cycles of 95 °C for 15 sec, 55 °C for 15 sec, and 72 °C for 30 sec, 30 cycles of 95 °C for 15 sec, 62 °C for 15 sec, and 72 °C for 30 sec, followed by a final extension at 72 °C for 1 min. Amplified DNA was purified using AMPure^®^ XP (Beckman Coulter) and quantified by a NanoDrop^®^ 1000 (Thermo Fischer Scientific, Waltham, MA, USA). A total of 100 ng of DNA was incubated with 1 μl of Rapid Adapter at room temperature for 5 min. The prepared DNA library (11 μl) was mixed with 34 μl of Sequencing Buffer, 25.5 μl of Loading Beads, and 4.5 μl of water, loaded onto the R9.4 flow cell (FLO-MIN106; Oxford Nanopore Technologies), and sequenced on the MinION™ Mk1B. MINKNOW software ver. 1.11.5 (Oxford Nanopore Technologies) was used for data acquisition.

### 16S rRNA gene sequencing on the MiSeq™ platform

Sequencing libraries were constructed as described previously [36]. Briefly, the V3-V4 regions of the 16S rRNA gene were amplified using a 16S (V3–V4) Metagenomic Library Construction Kit for NGS (Takara Bio Inc, Kusatsu, Japan). The following primers were used (16S rRNA gene-specific sequences are underlined): 341F with overhang adapter 5′-TCGTCGGCAGCGTCAGATGTGTATAAGAGACAG<u>CCTACGGGNGGCWGCAG</u>-3′ and 806R with overhang adapter 5′-GTCTCGTGGGCTCGGAGATGTGTATAAGAGACAG<u>GGACTACHVGGGTWTCT AAT</u>-3′. The second PCR was performed using the Nextera^®^ XT Index Kit (Illumina, San Diego, CA, USA) for sample multiplexing with index adapters. The libraries were sequenced on the MiSeq™ platform using the MiSeq™ Reagent Kit v3 (2 × 250 bp; Illumina).

### Bioinformatics analysis

Albacore software ver. 2.3.4 (Oxford Nanopore Technologies) was used for basecalling the MinION™ sequencing data (FAST5 files) to generate pass reads (FASTQ format) with a mean quality score > 7. The adapter and barcode sequences were trimmed using the EPI2ME Fastq Barcoding workflow ver. 3.10.4 (Oxford Nanopore Technologies). The reads were filtered by size using SeqKit software ver. 0.10.0 [38], retaining 1300-1950 bp sequences for the V1-V9 region and 350-600 bp sequences for the V3-V4 region, based on the size distribution of 16S rRNA gene sequences in the SILVA database ver. 132 [39, 40]. The average Phred quality score was assessed using NanoPlot ver. 1.27.0 [41]. The processed reads from each set were analyzed using our bioinformatics pipeline [42], as described previously [14, 15]. Briefly, FASTQ files were converted to FASTA files. Simple repetitive sequences were masked using the TANTAN program ver. 18 with default parameters [43]. To remove reads derived from human DNA, we searched each read against the human genome (GRCh38) using minimap2 ver. 2.14 with a map-ont-option [44]. Then, unmatched reads were regarded as reads derived from bacteria. For each read, a minimap2 search with 5850 representative bacterial genome sequences stored in the GenomeSync database (Additional File 1) [45] was performed. For each read, the species showing the highest minimap2 score were assigned to the query sequence. When more than two species showed the same similarity score, the reads were classified at any higher taxonomic rank covering all the identified species. Taxa were determined based on the NCBI taxonomy database [46]. Low-abundance taxa with less than 0.01% of total reads were discarded from the analysis.

### Statistical analyses

Differences between groups were evaluated by one-way analysis of variance (ANOVA) followed by Dunnett’s test for multiple comparisons. The Pearson correlation coefficient was computed to compare the bacterial compositions analyzed by different sequencing methods. Statistical significance was defined by a *P*-value < 0.05. Statistical analyses were performed with Prism8 (GraphPad Software, Inc. La Jolla, CA, USA).

### Statement of ethics

The Sunkaky Institutional Review Board approved this study (No. 2017-27). All participants provided written informed consent.

## Abbreviations

NGS: next-generation sequencing;
OTU: operational taxonomic unit;
PCR: polymerase chain reaction;
rRNA: ribosomal RNA

## Additional files

**Additional File 1:** Representative bacterial genomes stored in the GenomeSync database.

**Additional File 2: Fig. S1**. Sequence heterogeneities of the 27F primer-annealing site in 16S rRNA genes. **Fig. S2**. Evaluation of 16S rRNA PCR primers for identification of bacterial species. **Fig. S3**. Effect of read number on taxonomic classification. **Fig. S4**. Rarefaction curves of observed OTUs in 16S V3-V4 amplicon sequencing of human fecal samples using the MiSeq™ platform. **Fig. S5**. Species composition of *Blautia* in human fecal samples. **Fig. S6**. Species composition of *Bacteroides* in human fecal samples. **Fig. S7**. Deviations in the relative abundances of *Bifidobacterium* species in human fecal samples. **Fig. S8**. Comparison of species composition of fecal *Bifidobacterium* between classification methods.

**Additional File 3: Tables S1-S6**. Taxonomic assignment of the mock community analyzed by MinION™ sequencing.

**Additional File 4:** Alignment search results for V1-V9 amplicon sequencing of the mock community.

**Additional File 5:** Alignment search results for V3-V4 amplicon sequencing of the mock community.

**Additional File 6: Table S7**. Correlations between numbers of reads and numbers of detected species in 16S rRNA gene sequencing of human fecal samples. **Table S8**. Comparison of species composition of fecal *Blautia* between sequencing methods. **Table S9**. Comparison of species composition of fecal *Bacteroides* between sequencing methods. **Table S10**. Comparison of species composition of fecal *Bifidobacterium* between sequencing methods.

**Additional File 7:** Taxonomic profile of human fecal samples from MinION™ sequencing (amplicons: V1-V9).

**Additional File 8:** Taxonomic profile of human fecal samples from MinION™ sequencing (amplicons: V3-V4).

**Additional File 9:** Taxonomic profile of human fecal samples from MiSeq™ sequencing (amplicons: V3-V4).

**Additional File 10:** Taxonomic profiles of human fecal samples from MiSeq™ sequencing (amplicons: V3-V4, taxonomic classification by OTU-based analysis using the QIIME 2 pipeline).

## Declarations

### Ethics approval and consent to participate

This study was approved by the Sunkaky Institutional Review Board (No. 2017-27).

### Consent for publication

All participants provided written informed consent.

### Availability of data and materials

The sequence datasets supporting the conclusions of this article are available in the DDBJ DRA database (https://www.ddbj.nig.ac.jp/dra/index-e.html) under accession numbers DRR225043 to DRR225065.

### Competing interests

The authors declare that they have no competing interests.

### Funding

This work was supported by Japan Society for the Promotion of Science KAKENHI Grant Number JP19K09339 to YMa and the branding program as a world-leading research university on intractable immune and allergic diseases supported by the Ministry of Education, Culture, Sports, Science and Technology of Japan.

### Authors’ contributions

YMa, KK, AF, YMo, YN, SN and KH designed and supervised the study. SK, AF, YMo, and HO contributed to sample collection. YMa, KM, and TT conducted the experiments. YMa, YYasumizu, YYasuoka, and SN analyzed the data. YMa wrote the manuscript. YYasumizu, HB, SN, and KH contributed to editing the manuscript. All authors read and approved the final manuscript.

## Acknowledgements

We are grateful to Tadashi Imanishi (Tokai University School of Medicine) and Shino Matsukawa (Kyoto University Hospital) for helpful comments and discussion. We would like to thank Editage (www.editage.com) for English language editing.

## Notes

### Competing Interest Statement

The authors have declared no competing interest.

