## Additional File 2 for "Full-length 16S rRNA gene amplicon analysis of human gut microbiota using MinION™ nanopore sequencing confers species-level resolution"

a

| Species | Sequence |
|---|---|
| *Bacillus cereus* | AGAGTTTGATCCTGGCTCAG |
| *Bifidobacterium adolescentis* | AGGGTTCGATTCTGGCTCAG |
| *Clostridium beijerinckii* | AGAGTTTGATCCTGGCTCAG |
| *Deinococcus radiodurans* | AGAGTTTGATCCTGGCTCAG |
| *Enterococcus faecalis* | AGAGTTTGATCCTGGCTCAG |
| *Escherichia coli* | AGAGTTTGATCATGGCTCAG |
| *Lactobacillus gasseri* | AGAGTTTGATCCTGGCTCAG |
| *Rhodobacter sphaeroides* | AGAGTTTGATCCTGGCTCAG |
| *Staphylococcus epidermidis* | AGAGTTTGATCCTGGCTCAG |
| *Streptococcus mutans* | AGAGTTTGATCCTGGCTCAG |

b

| Forward primer | Sequence |
|---|---|
| Original 27F (ONT, SQK-RAB204) | AGAGTTTGATCMTGGCTCAG |
| This study (S-D-Bact-0008-c-S-20) | AGRGTTYGATYMTGGCTCAG |
| 27F-I | AGAGTTTGATCATGGCTCAG |
| 27F-II | AGGGTTCGATTCTGGCTCAG |
| 27F-III | AGAGTTTGATCCTGGCTCAG |

**Supplementary Fig. S1** Sequence heterogeneities of the 27F primer-annealing site in 16S rRNA genes. **a** Multiple sequence alignment for the ten bacterial species constituting the mock community. Variable nucleotides are shaded. Three mismatched bases in *Bifidobacterium* are shown in red. **b** 16S rRNA gene-specific sequences of the original 27F (Oxford Nanopore Technologies, ONT) and the primers used in this study. 27F-**I**, 27F-**II**, and 27F-**III** correspond to the sequences for *Escherichia coli*, *Bifidobacterium adolescentis*, and the other eight species, respectively.

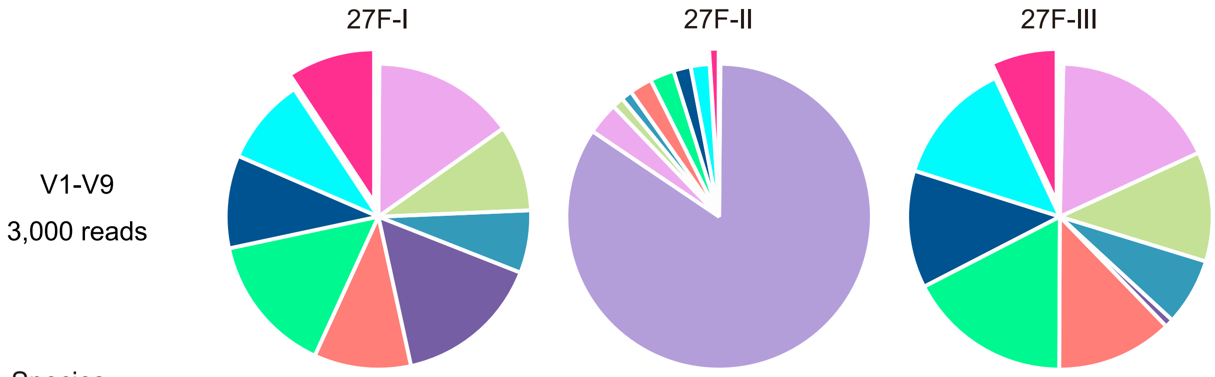


| Species | 27F-I | 27F-II | 27F-III |
|---|---|---|---|
| *B. cereus* | 0.10% | 0.13% | 0.37% |
| *B. adolescentis* | 0.00% | 84.30% | 0.00% |
| *C. beijerinckii* | 15.10% | 3.40% | 17.77% |
| *D. radiodurans* | 9.13% | 1.20% | 11.60% |
| *E. faecalis* | 6.63% | 1.17% | 7.10% |
| *E. coli* | 15.60% | 0.00% | 0.90% |
| *L. gasseri* | 10.27% | 2.40% | 12.33% |
| *R. sphaeroides* | 14.83% | 2.57% | 17.37% |
| *S. epidermidis* | 9.83% | 1.83% | 12.43% |
| *S. mutans* | 9.20% | 2.00% | 13.17% |
| mis- or unclassified | 9.30% | 1.00% | 6.97% |

**Supplementary Fig. S2** Evaluation of 16S rRNA PCR primers for identification of bacterial species. The V1-V9 region of the 16S rRNA gene was amplified from the ten-species mock community sample using the indicated 27F variant and 1492R primer and sequenced on MinION™. The pie charts represent taxonomic profiles at the species level, and the relative abundances (%) of each taxon are shown.

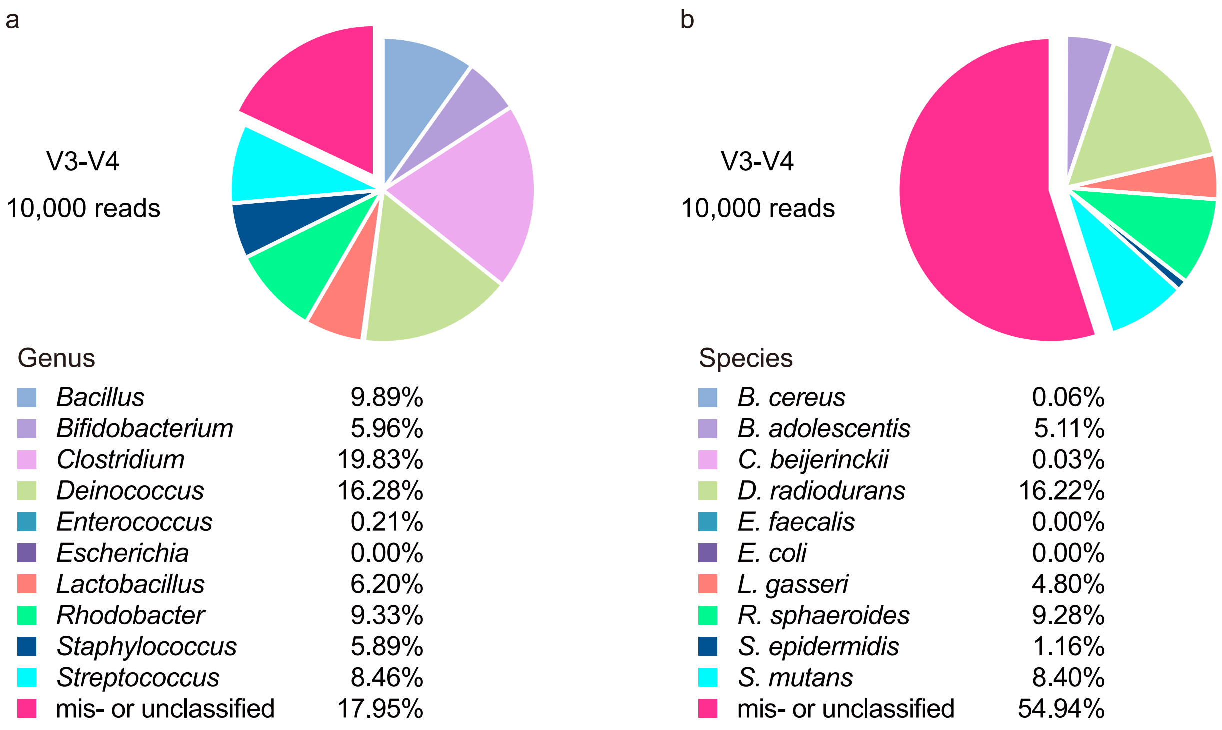


**Supplementary Fig. S3** Effect of read number on taxonomic classification. **a, b** The V3-V4 region of the 16S rRNA gene was amplified from a 10-species mock community sample and sequenced on MinION™ as in Fig. 1. Ten thousand reads were used for taxonomic profiling. The pie charts represent taxonomic profiles at the (a) genus and (b) species level. The relative abundances (%) of each taxon are shown.

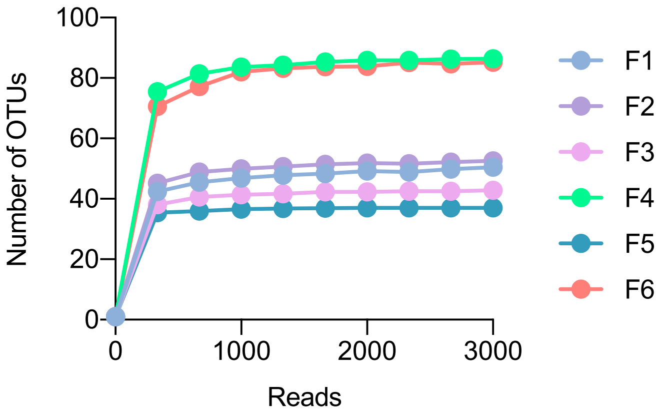


**Supplementary Fig. S4** Rarefaction curves of observed OTUs in 16S V3-V4 amplicon sequencing of human fecal samples using the MiSeq™ platform.

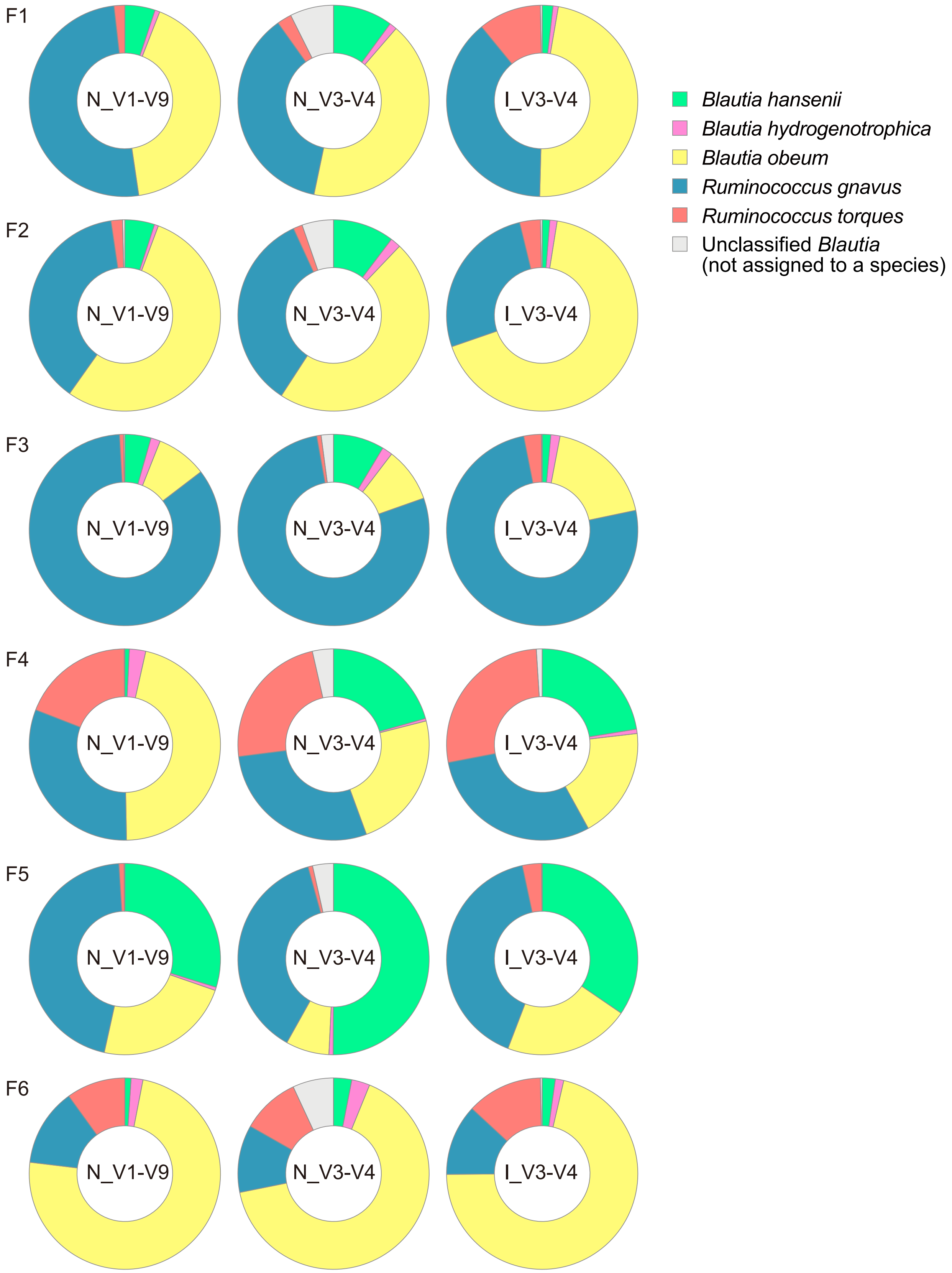


**Supplementary Fig. S5** Species composition of *Blautia* in human fecal samples. Results obtained by the three sequencing methods are shown. N_V1-V9: sequencing of the 16S V1-V9 region using Oxford Nanopore MinION™. N_V3-V4: sequencing of the 16S V3-V4 region using Oxford Nanopore MinION™. I_V3-V4: sequencing of the 16S V3-V4 region using Illumina MiSeq™.

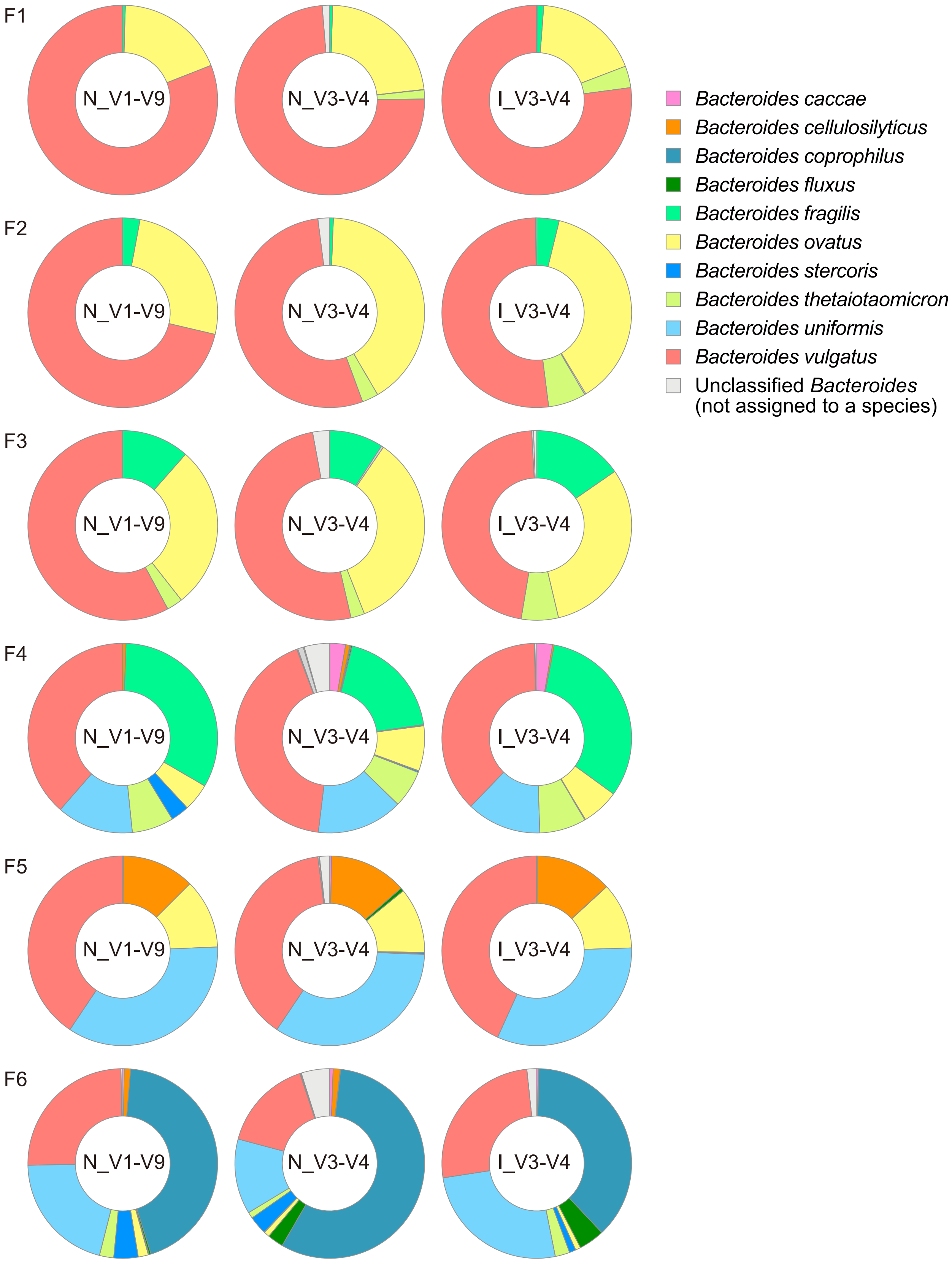


**Supplementary Fig. S6** Species composition of *Bacteroides* in human fecal samples. Results obtained by the three sequencing methods are shown. N_V1-V9: sequencing of the 16S V1-V9 region using Oxford Nanopore MinION™. N_V3-V4: sequencing of the 16S V3-V4 region using Oxford Nanopore MinION™. I_V3-V4: sequencing of the 16S V3-V4 region using Illumina MiSeq™. The legends show the ten most abundant species.

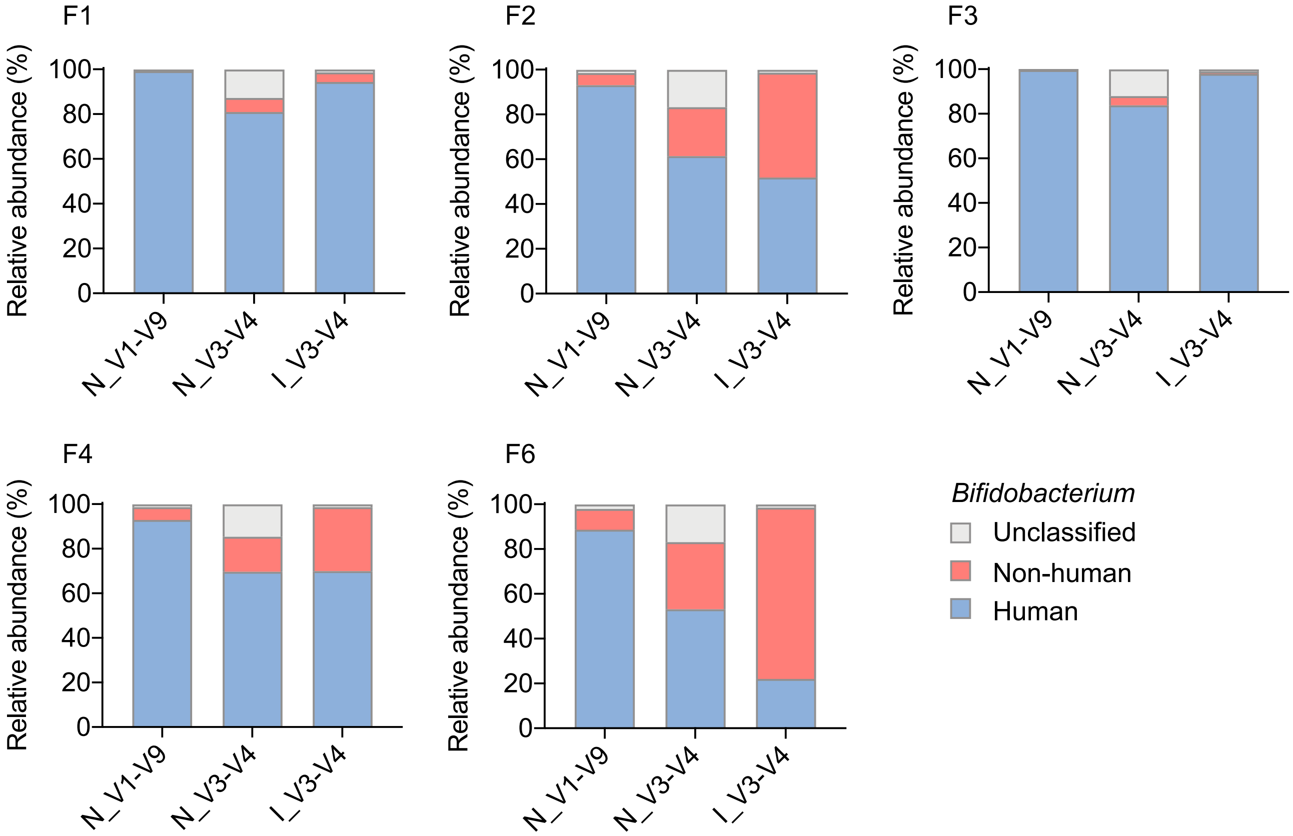


**Supplementary Fig. S7** Deviations in the relative abundances of *Bifidobacterium* species in human fecal samples. The proportions (%) of *Bifidobacterium* species originating from human or non-human hosts are shown. N_V1-V9: sequencing of the 16S V1-V9 region using Oxford Nanopore MinION™. N_V3-V4: sequencing of the 16S V3-V4 region using Oxford Nanopore MinION™. I_V3-V4: sequencing of the 16S V3-V4 region using Illumina MiSeq™.

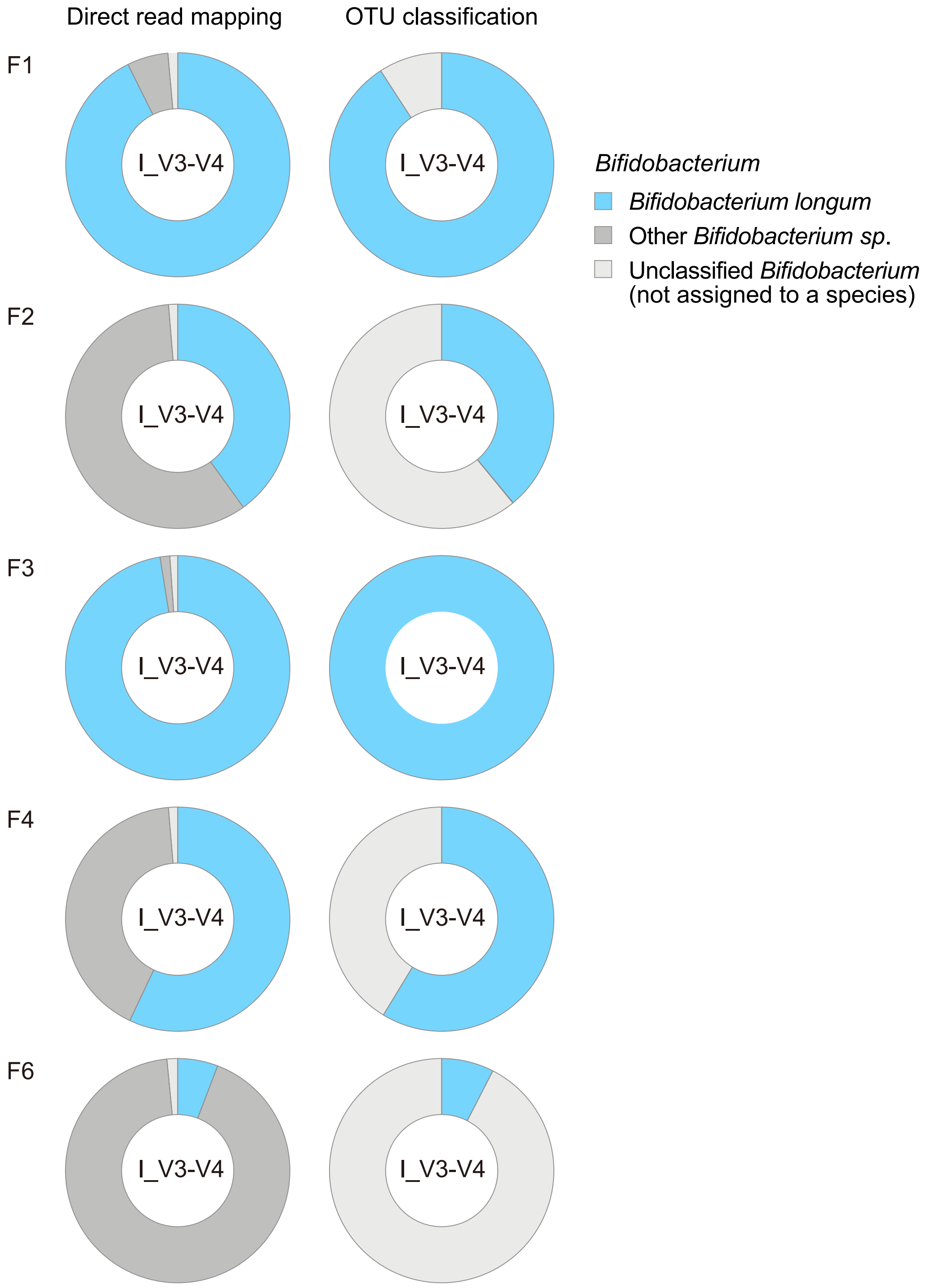


**Supplementary Fig. S8** Comparison of species composition of fecal *Bifidobacterium* between classification methods. MiSeq™ V3-V4 reads (I_V3-V4) from the human fecal samples (F1-F6) were either mapped directly to the reference bacterial genome (Direct read mapping) or clustered into OTUs followed by taxonomic annotation using the QIIME 2 pipeline (OTU classification). Taxonomic profiles for reads assigned to *Bifidobacterium* are shown. The OTU-based method identified *Bifidobacterium longum,* and other reads were not assigned to species level, categorized as "Unclassified *Bifidobacterium*" in the charts (right). For clarity, reads assigned to *Bifidobacterium* species other than *Bifidobacterium longum* in our direct mapping approach were shown as "Other *Bifidobacterium*" (left).
