## Additional File 4 for "Full-length 16S rRNA gene amplicon analysis of human gut microbiota using MinION™ nanopore sequencing confers species-level resolution"

Alignment search results for V1–V9 amplicon sequencing of the mock community.
The top three hits for 3,000 query sequences are shown. The values represent the bit score of alignment.

1
735 Escherichia coli str. K-12 substr. MG1655 GCA_000005845.2
733 Escherichia coli O104_3AH4 str. 2011C-3493 GCA_000299455.1
733 Shigella flexneri 2a str. 301 GCA_000006925.2
720 Escherichia coli O157_3AH7 str. Sakai GCA_000008865.1
2
399 Deinococcus radiodurans R1 GCA_000008565.1
240 Deinococcus hopiensis KR-140 GCA_900176165.1
239 Deinococcus deserti VCD115 GCA_000020685.1
3
823 Clostridium beijerinckii GCA_000833105.2
786 Clostridium saccharoperbutylacetonicum N1-4_28HMT_29 GCA_000340885.1
757 Clostridium puniceum GCA_002006345.1
4
716 Clostridium beijerinckii GCA_000833105.2
656 Clostridium saccharobutylicum DSM 13864 GCA_000473995.1
653 Clostridium saccharoperbutylacetonicum N1-4_28HMT_29 GCA_000340885.1
5
474 Staphylococcus epidermidis ATCC 12228 GCA_000007645.1
448 Staphylococcus capitis subsp. capitis GCA_001028645.1
386 Megasphaera cerevisiae DSM 20462 GCA_001045675.1
386 Staphylococcus warneri SG1 GCA_000332735.1
6
544 Staphylococcus epidermidis ATCC 12228 GCA_000007645.1
513 Staphylococcus capitis subsp. capitis GCA_001028645.1
448 Staphylococcus hominis subsp. hominis C80 GCA_000183685.1
7
773 Clostridium beijerinckii GCA_000833105.2
762 Clostridium saccharoperbutylacetonicum N1-4_28HMT_29 GCA_000340885.1
721 Clostridium puniceum GCA_002006345.1
8
680 Lactobacillus gasseri ATCC 33323 = JCM 1131 GCA_000014425.1
609 Lactobacillus hominis DSM 23910 = CRBIP 24.179 GCA_000296835.1
446 Lactobacillus iners DSM 13335 GCA_000160875.1
9
765 Lactobacillus gasseri ATCC 33323 = JCM 1131 GCA_000014425.1
701 Lactobacillus hominis DSM 23910 = CRBIP 24.179 GCA_000296835.1
499 Lactobacillus psittaci DSM 15354 GCA_000425905.1
a
675 Clostridium beijerinckii GCA_000833105.2
635 Clostridium saccharoperbutylacetonicum N1-4_28HMT_29 GCA_000340885.1
594 Clostridium puniceum GCA_002006345.1
b
462 Clostridium beijerinckii GCA_000833105.2
440 Clostridium saccharobutylicum DSM 13864 GCA_000473995.1
438 Clostridium butyricum GCA_001456065.2
c
676 Lactobacillus gasseri ATCC 33323 = JCM 1131 GCA_000014425.1
645 Lactobacillus hominis DSM 23910 = CRBIP 24.179 GCA_000296835.1
480 Lactobacillus iners DSM 13335 GCA_000160875.1
d
753 Clostridium beijerinckii GCA_000833105.2
724 Clostridium saccharoperbutylacetonicum N1-4_28HMT_29 GCA_000340885.1
697 Clostridium puniceum GCA_002006345.1
e
742 Streptococcus mutans UA159 GCA_000007465.2
546 Streptococcus ratti FA-1 = DSM 20564 GCA_000286075.1
490 Streptococcus halotolerans GCA_001598035.1
f
384 Enterococcus faecalis V583 GCA_000007785.1
384 Streptomyces cinnamoneus GCA_001885705.1
351 Enterococcus faecium DO GCA_000174395.2
338 Enterococcus casseliflavus EC20 GCA_000157355.2
338 Enterococcus dispar ATCC 51266 GCA_000406945.1
338 Enterococcus hirae ATCC 9790 GCA_000271405.2
338 Enterococcus massiliensis GCA_001050095.1
338 Enterococcus mundtii QU 25 GCA_000504125.1
338 Enterococcus saccharolyticus subsp. saccharolyticus ATCC 43076 GCA_000407285.1
g
824 Deinococcus radiodurans R1 GCA_000008565.1
529 Deinococcus deserti VCD115 GCA_000020685.1
486 Deinococcus hopiensis KR-140 GCA_900176165.1

h
787 Staphylococcus epidermidis ATCC 12228 GCA_000007645.1
776 Staphylococcus capitis subsp. capitis GCA_001028645.1
704 Staphylococcus hominis subsp. hominis C80 GCA_000183685.1
i
668 Staphylococcus epidermidis ATCC 12228 GCA_000007645.1
627 Staphylococcus capitis subsp. capitis GCA_001028645.1
573 Staphylococcus hominis subsp. hominis C80 GCA_000183685.1
j
736 Enterococcus faecalis V583 GCA_000007785.1
689 Streptomyces cinnamoneus GCA_001885705.1
564 Enterococcus rivorum GCA_001742285.1
k
672 Rhodobacter sphaeroides 2.4.1 GCA_000012905.2
624 Rhodobacter sphaeroides ATCC 17025 GCA_000016405.1
520 Gemmobacter aquatilis GCA_900110025.1
l
753 Rhodobacter sphaeroides 2.4.1 GCA_000012905.2
688 Rhodobacter sphaeroides ATCC 17025 GCA_000016405.1
561 Gemmobacter aquatilis GCA_900110025.1
m
717 Bacillus anthracis str. Ames GCA_000007845.1
717 Bacillus anthracis str. Sterne GCA_000008165.1
717 Bacillus cereus ATCC 14579 GCA_000007825.1
717 Bacillus thuringiensis YBT-1518 GCA_000497525.2
717 _5BBacillus thuringiensis_5D serovar konkukian str. 97-27 GCA_000008505.1
704 Bacillus pseudomycoides DSM 12442 GCA_000161455.1
697 Bacillus mycoides GCA_000832605.1
n
536 Lactobacillus gasseri ATCC 33323 = JCM 1131 GCA_000014425.1
522 Lactobacillus hominis DSM 23910 = CRBIP 24.179 GCA_000296835.1
382 Lactobacillus iners DSM 13335 GCA_000160875.1
o
522 Escherichia coli str. K-12 substr. MG1655 GCA_000005845.2
513 Escherichia coli O104_3AH4 str. 2011C-3493 GCA_000299455.1
513 Shigella flexneri 2a str. 301 GCA_000006925.2
506 Escherichia coli O157_3AH7 str. Sakai GCA_000008865.1
p
833 Clostridium beijerinckii GCA_000833105.2
774 Clostridium saccharoperbutylacetonicum N1-4_28HMT_29 GCA_000340885.1
770 Clostridium puniceum GCA_002006345.1
q
514 Staphylococcus epidermidis ATCC 12228 GCA_000007645.1
505 Staphylococcus capitis subsp. capitis GCA_001028645.1
479 Staphylococcus condimenti GCA_001618885.1
479 Staphylococcus simulans GCA_001559115.1
r
510 Escherichia coli O157_3AH7 str. Sakai GCA_000008865.1
510 Escherichia coli str. K-12 substr. MG1655 GCA_000005845.2
501 Escherichia coli IAI39 GCA_000026345.1
495 Escherichia coli UMN026 GCA_000026325.2
s
476 Staphylococcus epidermidis ATCC 12228 GCA_000007645.1
440 Staphylococcus capitis subsp. capitis GCA_001028645.1
383 Staphylococcus lugdunensis HKU09-01 GCA_000025085.1
t
640 Staphylococcus epidermidis ATCC 12228 GCA_000007645.1
618 Staphylococcus capitis subsp. capitis GCA_001028645.1
582 Staphylococcus aureus subsp. aureus NCTC 8325 GCA_000013425.1
u
593 Rhodobacter sphaeroides 2.4.1 GCA_000012905.2
524 Rhodobacter sphaeroides ATCC 17025 GCA_000016405.1
443 Gemmobacter aquatilis GCA_900110025.1
v
788 Deinococcus radiodurans R1 GCA_000008565.1
518 Deinococcus deserti VCD115 GCA_000020685.1
489 Deinococcus gobiensis I-0 GCA_000252445.1
w
1035 Enterococcus faecalis V583 GCA_000007785.1
989 Streptomyces cinnamoneus GCA_001885705.1
929 Enterococcus rivorum GCA_001742285.1
x
519 Enterococcus faecalis V583 GCA_000007785.1
473 Enterococcus rivorum GCA_001742285.1
472 Streptomyces cinnamoneus GCA_001885705.1
y

518 Staphylococcus epidermidis ATCC 12228 GCA_000007645.1
478 Staphylococcus capitis subsp. capitis GCA_001028645.1
450 Staphylococcus warneri SG1 GCA_000332735.1
z
492 Escherichia coli str. K-12 substr. MG1655 GCA_000005845.2
482 Shigella flexneri 2a str. 301 GCA_000006925.2
478 Escherichia coli UMN026 GCA_000026325.2
A
876 Staphylococcus epidermidis ATCC 12228 GCA_000007645.1
868 Staphylococcus capitis subsp. capitis GCA_001028645.1
800 Staphylococcus haemolyticus JCSC1435 GCA_000009865.1
800 Staphylococcus hominis subsp. hominis C80 GCA_000183685.1
B
639 Escherichia coli O157_3AH7 str. Sakai GCA_000008865.1
635 Escherichia coli UMN026 GCA_000026325.2
635 Escherichia coli str. K-12 substr. MG1655 GCA_000005845.2
631 Escherichia coli IAI39 GCA_000026345.1
631 Escherichia coli O83_3AH1 str. NRG 857C GCA_000183345.1
631 Shigella flexneri 2a str. 301 GCA_000006925.2
C
660 Deinococcus radiodurans R1 GCA_000008565.1
452 Deinococcus gobiensis I-0 GCA_000252445.1
434 Deinococcus deserti VCD115 GCA_000020685.1
D
571 Escherichia coli str. K-12 substr. MG1655 GCA_000005845.2
558 Escherichia coli O157_3AH7 str. Sakai GCA_000008865.1
558 Escherichia coli O83_3AH1 str. NRG 857C GCA_000183345.1
558 Escherichia coli UMN026 GCA_000026325.2
558 Shigella flexneri 2a str. 301 GCA_000006925.2
554 Escherichia coli O104_3AH4 str. 2011C-3493 GCA_000299455.1
E
709 Rhodobacter sphaeroides 2.4.1 GCA_000012905.2
615 Rhodobacter sphaeroides ATCC 17025 GCA_000016405.1
544 Gemmobacter aquatilis GCA_900110025.1
F
943 Streptococcus mutans UA159 GCA_000007465.2
662 Streptococcus ratti FA-1 = DSM 20564 GCA_000286075.1
592 Streptococcus macacae NCTC 11558 GCA_000187995.3
G
549 Clostridium saccharoperbutylacetonicum N1-4_28HMT_29 GCA_000340885.1
543 Clostridium beijerinckii GCA_000833105.2
513 Clostridium puniceum GCA_002006345.1
H
806 Streptococcus mutans UA159 GCA_000007465.2
581 Streptococcus ratti FA-1 = DSM 20564 GCA_000286075.1
560 Streptococcus gallolyticus subsp. gallolyticus DSM 16831 GCA_002000985.1
I
711 Enterococcus faecalis V583 GCA_000007785.1
665 Streptomyces cinnamoneus GCA_001885705.1
621 Enterococcus rivorum GCA_001742285.1
J
796 Clostridium beijerinckii GCA_000833105.2
776 Clostridium saccharoperbutylacetonicum N1-4_28HMT_29 GCA_000340885.1
762 Clostridium saccharobutylicum DSM 13864 GCA_000473995.1
K
818 Streptococcus mutans UA159 GCA_000007465.2
653 Streptococcus ratti FA-1 = DSM 20564 GCA_000286075.1
640 Streptococcus merionis DSM 19192 GCA_000380085.1
L
586 Clostridium beijerinckii GCA_000833105.2
563 Clostridium saccharoperbutylacetonicum N1-4_28HMT_29 GCA_000340885.1
538 Clostridium puniceum GCA_002006345.1
M
707 Deinococcus radiodurans R1 GCA_000008565.1
470 Deinococcus deserti VCD115 GCA_000020685.1
426 Deinococcus soli Cha et al. 2016 GCA_001007995.1
N
738 Bifidobacterium adolescentis ATCC 15703 GCA_000010425.1
697 Bifidobacterium dentium JCM 1195 = DSM 20436 GCA_001042595.1
660 Bifidobacterium angulatum DSM 20098 = JCM 7096 GCA_001025155.1
O
462 Staphylococcus epidermidis ATCC 12228 GCA_000007645.1
456 Staphylococcus lugdunensis HKU09-01 GCA_000025085.1
449 Staphylococcus lutrae GCA_002101335.1
P
830 Enterococcus faecalis V583 GCA_000007785.1

783 Streptomyces cinnamoneus GCA_001885705.1
778 Enterococcus haemoperoxidus ATCC BAA-382 GCA_000407165.1
Q
742 Clostridium beijerinckii GCA_000833105.2
696 Clostridium saccharoperbutylacetonicum N1-4_28HMT_29 GCA_000340885.1
679 Clostridium puniceum GCA_002006345.1
R
789 Clostridium beijerinckii GCA_000833105.2
788 Clostridium saccharoperbutylacetonicum N1-4_28HMT_29 GCA_000340885.1
772 Clostridium puniceum GCA_002006345.1
S
619 Clostridium beijerinckii GCA_000833105.2
593 Clostridium saccharoperbutylacetonicum N1-4_28HMT_29 GCA_000340885.1
592 Clostridium puniceum GCA_002006345.1
T
533 Escherichia coli O157_3AH7 str. Sakai GCA_000008865.1
519 Escherichia coli str. K-12 substr. MG1655 GCA_000005845.2
519 Shigella flexneri 2a str. 301 GCA_000006925.2
518 Escherichia coli UMN026 GCA_000026325.2
U
738 Clostridium beijerinckii GCA_000833105.2
705 Clostridium puniceum GCA_002006345.1
705 Clostridium saccharoperbutylacetonicum N1-4_28HMT_29 GCA_000340885.1
693 Clostridium saccharobutylicum DSM 13864 GCA_000473995.1
V
606 Bifidobacterium adolescentis ATCC 15703 GCA_000010425.1
520 Bifidobacterium dentium JCM 1195 = DSM 20436 GCA_001042595.1
476 Bifidobacterium angulatum DSM 20098 = JCM 7096 GCA_001025155.1
W
685 Deinococcus radiodurans R1 GCA_000008565.1
472 Deinococcus deserti VCD115 GCA_000020685.1
430 Deinococcus puniceus GCA_001644565.1
X
901 Lactobacillus gasseri ATCC 33323 = JCM 1131 GCA_000014425.1
834 Lactobacillus hominis DSM 23910 = CRBIP 24.179 GCA_000296835.1
592 Lactobacillus iners DSM 13335 GCA_000160875.1
Y
658 Bifidobacterium adolescentis ATCC 15703 GCA_000010425.1
501 Bifidobacterium angulatum DSM 20098 = JCM 7096 GCA_001025155.1
476 Bifidobacterium dentium JCM 1195 = DSM 20436 GCA_001042595.1
Z
819 Lactobacillus gasseri ATCC 33323 = JCM 1131 GCA_000014425.1
764 Lactobacillus hominis DSM 23910 = CRBIP 24.179 GCA_000296835.1
576 Lactobacillus iners DSM 13335 GCA_000160875.1
10
803 Lactobacillus gasseri ATCC 33323 = JCM 1131 GCA_000014425.1
709 Lactobacillus hominis DSM 23910 = CRBIP 24.179 GCA_000296835.1
594 Lactobacillus iners DSM 13335 GCA_000160875.1
11
523 Deinococcus radiodurans R1 GCA_000008565.1
375 Deinococcus deserti VCD115 GCA_000020685.1
370 Deinococcus hopiensis KR-140 GCA_900176165.1
12
652 Deinococcus radiodurans R1 GCA_000008565.1
384 Deinococcus deserti VCD115 GCA_000020685.1
340 Deinococcus proteolyticus MRP GCA_000190555.1
13
558 Staphylococcus epidermidis ATCC 12228 GCA_000007645.1
527 Staphylococcus haemolyticus JCSC1435 GCA_000009865.1
518 Staphylococcus capitis subsp. capitis GCA_001028645.1
14
880 Clostridium beijerinckii GCA_000833105.2
822 Clostridium saccharoperbutylacetonicum N1-4_28HMT_29 GCA_000340885.1
797 Clostridium butyricum GCA_001456065.2
15
786 Clostridium beijerinckii GCA_000833105.2
753 Clostridium saccharoperbutylacetonicum N1-4_28HMT_29 GCA_000340885.1
730 Clostridium puniceum GCA_002006345.1
16
704 Escherichia coli O157_3AH7 str. Sakai GCA_000008865.1
704 Escherichia coli str. K-12 substr. MG1655 GCA_000005845.2
680 Shigella flexneri 2a str. 301 GCA_000006925.2
674 Escherichia coli UMN026 GCA_000026325.2
17
499 Staphylococcus epidermidis ATCC 12228 GCA_000007645.1
481 Staphylococcus capitis subsp. capitis GCA_001028645.1

422 Staphylococcus warneri SG1 GCA_000332735.1
18
516 Clostridium beijerinckii GCA_000833105.2
516 Clostridium saccharobutylicum DSM 13864 GCA_000473995.1
501 Clostridium saccharoperbutylacetonicum N1-4_28HMT_29 GCA_000340885.1
482 Clostridium puniceum GCA_002006345.1
19
636 Rhodobacter sphaeroides 2.4.1 GCA_000012905.2
563 Rhodobacter sphaeroides ATCC 17025 GCA_000016405.1
488 Pseudorhodobacter ferrugineus DSM 5888 GCA_000420745.1
1a
712 Bifidobacterium adolescentis ATCC 15703 GCA_000010425.1
550 Bifidobacterium callitrichos DSM 23973 GCA_000741175.1
539 Bifidobacterium angulatum DSM 20098 = JCM 7096 GCA_001025155.1
1b
781 Clostridium beijerinckii GCA_000833105.2
777 Clostridium puniceum GCA_002006345.1
777 Clostridium saccharoperbutylacetonicum N1-4_28HMT_29 GCA_000340885.1
747 Clostridium saccharobutylicum DSM 13864 GCA_000473995.1
1c
586 Clostridium beijerinckii GCA_000833105.2
547 Clostridium saccharoperbutylacetonicum N1-4_28HMT_29 GCA_000340885.1
540 Clostridium saccharobutylicum DSM 13864 GCA_000473995.1
1d
525 Streptococcus mutans UA159 GCA_000007465.2
357 Streptococcus ratti FA-1 = DSM 20564 GCA_000286075.1
353 Streptococcus sobrinus DSM 20742 = ATCC 33478 GCA_000686605.1
1e
811 Clostridium beijerinckii GCA_000833105.2
778 Clostridium saccharoperbutylacetonicum N1-4_28HMT_29 GCA_000340885.1
738 Clostridium saccharobutylicum DSM 13864 GCA_000473995.1
1f
470 Clostridium beijerinckii GCA_000833105.2
441 Clostridium saccharoperbutylacetonicum N1-4_28HMT_29 GCA_000340885.1
421 Clostridium saccharobutylicum DSM 13864 GCA_000473995.1
1g
963 Staphylococcus epidermidis ATCC 12228 GCA_000007645.1
916 Staphylococcus capitis subsp. capitis GCA_001028645.1
853 Staphylococcus warneri SG1 GCA_000332735.1
1h
785 Rhodobacter sphaeroides 2.4.1 GCA_000012905.2
753 Rhodobacter sphaeroides ATCC 17025 GCA_000016405.1
675 Gemmobacter aquatilis GCA_900110025.1
1i
688 Lactobacillus gasseri ATCC 33323 = JCM 1131 GCA_000014425.1
629 Lactobacillus hominis DSM 23910 = CRBIP 24.179 GCA_000296835.1
459 Lactobacillus psittaci DSM 15354 GCA_000425905.1
1j
742 Staphylococcus epidermidis ATCC 12228 GCA_000007645.1
732 Staphylococcus capitis subsp. capitis GCA_001028645.1
679 Megasphaera cerevisiae DSM 20462 GCA_001045675.1
679 Staphylococcus warneri SG1 GCA_000332735.1
1k
608 Bacillus anthracis str. Ames GCA_000007845.1
608 Bacillus anthracis str. Sterne GCA_000008165.1
608 _5BBacillus thuringiensis_5D serovar konkukian str. 97-27 GCA_000008505.1
607 Bacillus cereus ATCC 14579 GCA_000007825.1
602 Bacillus thuringiensis YBT-1518 GCA_000497525.2
1l
621 Clostridium beijerinckii GCA_000833105.2
577 Clostridium saccharoperbutylacetonicum N1-4_28HMT_29 GCA_000340885.1
550 Clostridium puniceum GCA_002006345.1
550 Clostridium saccharobutylicum DSM 13864 GCA_000473995.1
1m
705 Deinococcus radiodurans R1 GCA_000008565.1
399 Deinococcus deserti VCD115 GCA_000020685.1
376 Deinococcus gobiensis I-0 GCA_000252445.1
1n
526 Lactobacillus gasseri ATCC 33323 = JCM 1131 GCA_000014425.1
461 Lactobacillus hominis DSM 23910 = CRBIP 24.179 GCA_000296835.1
399 Lactobacillus iners DSM 13335 GCA_000160875.1
1o
863 Deinococcus radiodurans R1 GCA_000008565.1
561 Deinococcus deserti VCD115 GCA_000020685.1
550 Deinococcus hopiensis KR-140 GCA_900176165.1
1p

696 Staphylococcus epidermidis ATCC 12228 GCA_000007645.1
692 Staphylococcus capitis subsp. capitis GCA_001028645.1
628 Staphylococcus warneri SG1 GCA_000332735.1
1q
473 Lactobacillus gasseri ATCC 33323 = JCM 1131 GCA_000014425.1
420 Lactobacillus hominis DSM 23910 = CRBIP 24.179 GCA_000296835.1
332 Lactobacillus iners DSM 13335 GCA_000160875.1
1r
776 Deinococcus radiodurans R1 GCA_000008565.1
461 Deinococcus marmoris DSM 12784 GCA_000701405.1
457 Deinococcus puniceus GCA_001644565.1
1s
806 Rhodobacter sphaeroides 2.4.1 GCA_000012905.2
768 Rhodobacter sphaeroides ATCC 17025 GCA_000016405.1
608 Gemmobacter aquatilis GCA_900110025.1
1t
778 Staphylococcus epidermidis ATCC 12228 GCA_000007645.1
728 Staphylococcus capitis subsp. capitis GCA_001028645.1
670 Staphylococcus aureus subsp. aureus NCTC 8325 GCA_000013425.1
1u
620 Enterococcus faecalis V583 GCA_000007785.1
606 Enterococcus massiliensis GCA_001050095.1
596 Enterococcus mundtii QU 25 GCA_000504125.1
1v
708 Staphylococcus epidermidis ATCC 12228 GCA_000007645.1
677 Staphylococcus capitis subsp. capitis GCA_001028645.1
645 Staphylococcus lugdunensis HKU09-01 GCA_000025085.1
1w
768 Deinococcus radiodurans R1 GCA_000008565.1
539 Deinococcus marmoris DSM 12784 GCA_000701405.1
539 Deinococcus swuensis GCA_000800395.1
536 Deinococcus deserti VCD115 GCA_000020685.1
1x
742 Rhodobacter sphaeroides 2.4.1 GCA_000012905.2
691 Rhodobacter sphaeroides ATCC 17025 GCA_000016405.1
544 Gemmobacter aquatilis GCA_900110025.1
1y
520 Clostridium beijerinckii GCA_000833105.2
493 Clostridium saccharobutylicum DSM 13864 GCA_000473995.1
493 Clostridium saccharoperbutylacetonicum N1-4_28HMT_29 GCA_000340885.1
487 Clostridium chromiireducens GCA_002029255.1
1z
900 Streptococcus mutans UA159 GCA_000007465.2
679 Streptococcus ratti FA-1 = DSM 20564 GCA_000286075.1
645 Streptococcus gordonii str. Challis substr. CH1 GCA_000017005.1
1A
769 Clostridium beijerinckii GCA_000833105.2
735 Clostridium saccharoperbutylacetonicum N1-4_28HMT_29 GCA_000340885.1
688 Clostridium puniceum GCA_002006345.1
1B
695 Rhodobacter sphaeroides 2.4.1 GCA_000012905.2
596 Rhodobacter sphaeroides ATCC 17025 GCA_000016405.1
555 Gemmobacter aquatilis GCA_900110025.1
1C
526 Rhodobacter sphaeroides 2.4.1 GCA_000012905.2
455 Rhodobacter sphaeroides ATCC 17025 GCA_000016405.1
333 Pseudorhodobacter psychrotolerans GCA_001294535.1
1D
625 Lactobacillus gasseri ATCC 33323 = JCM 1131 GCA_000014425.1
575 Lactobacillus hominis DSM 23910 = CRBIP 24.179 GCA_000296835.1
475 Lactobacillus iners DSM 13335 GCA_000160875.1
1E
495 Lactobacillus gasseri ATCC 33323 = JCM 1131 GCA_000014425.1
457 Lactobacillus hominis DSM 23910 = CRBIP 24.179 GCA_000296835.1
337 Lactobacillus iners DSM 13335 GCA_000160875.1
1F
784 Escherichia coli str. K-12 substr. MG1655 GCA_000005845.2
761 Escherichia coli O104_3AH4 str. 2011C-3493 GCA_000299455.1
752 Shigella flexneri 2a str. 301 GCA_000006925.2
1G
542 Clostridium saccharoperbutylacetonicum N1-4_28HMT_29 GCA_000340885.1
540 Clostridium beijerinckii GCA_000833105.2
513 Clostridium butyricum GCA_001456065.2
1H
648 Rhodobacter sphaeroides 2.4.1 GCA_000012905.2
608 Rhodobacter sphaeroides ATCC 17025 GCA_000016405.1

502 Gemmobacter aquatilis GCA_900110025.1
1I
745 Clostridium beijerinckii GCA_000833105.2
692 Clostridium saccharobutylicum DSM 13864 GCA_000473995.1
692 Clostridium saccharoperbutylacetonicum N1-4_28HMT_29 GCA_000340885.1
677 Clostridium puniceum GCA_002006345.1
1J
824 Lactobacillus gasseri ATCC 33323 = JCM 1131 GCA_000014425.1
771 Lactobacillus hominis DSM 23910 = CRBIP 24.179 GCA_000296835.1
517 Lactobacillus iners DSM 13335 GCA_000160875.1
1K
818 Staphylococcus epidermidis ATCC 12228 GCA_000007645.1
773 Staphylococcus capitis subsp. capitis GCA_001028645.1
702 Staphylococcus warneri SG1 GCA_000332735.1
1L
777 Staphylococcus epidermidis ATCC 12228 GCA_000007645.1
739 Staphylococcus capitis subsp. capitis GCA_001028645.1
661 Staphylococcus haemolyticus JCSC1435 GCA_000009865.1
661 Staphylococcus hominis subsp. hominis C80 GCA_000183685.1
1M
380 Lactobacillus gasseri ATCC 33323 = JCM 1131 GCA_000014425.1
380 Lactobacillus hominis DSM 23910 = CRBIP 24.179 GCA_000296835.1
358 Lactobacillus kimchicus JCM 15530 GCA_001433995.1
356 Enterococcus faecalis V583 GCA_000007785.1
1N
414 Deinococcus radiodurans R1 GCA_000008565.1
254 Deinococcus deserti VCD115 GCA_000020685.1
244 Deinococcus hopiensis KR-140 GCA_900176165.1
1O
592 Bifidobacterium adolescentis ATCC 15703 GCA_000010425.1
457 Bifidobacterium dentium JCM 1195 = DSM 20436 GCA_001042595.1
450 Bifidobacterium angulatum DSM 20098 = JCM 7096 GCA_001025155.1
450 Bifidobacterium callitrichos DSM 23973 GCA_000741175.1
1P
645 Enterococcus faecalis V583 GCA_000007785.1
598 Streptomyces cinnamoneus GCA_001885705.1
577 Enterococcus rivorum GCA_001742285.1
1Q
439 Enterococcus faecalis V583 GCA_000007785.1
399 Enterococcus haemoperoxidus ATCC BAA-382 GCA_000407165.1
393 Enterococcus rivorum GCA_001742285.1
1R
490 Clostridium beijerinckii GCA_000833105.2
487 Clostridium saccharoperbutylacetonicum N1-4_28HMT_29 GCA_000340885.1
482 Clostridium saccharobutylicum DSM 13864 GCA_000473995.1
1S
780 Escherichia coli O104_3AH4 str. 2011C-3493 GCA_000299455.1
780 Escherichia coli str. K-12 substr. MG1655 GCA_000005845.2
773 Shigella flexneri 2a str. 301 GCA_000006925.2
758 Escherichia coli O157_3AH7 str. Sakai GCA_000008865.1
758 Escherichia coli UMN026 GCA_000026325.2
1T
417 Clostridium beijerinckii GCA_000833105.2
379 Clostridium saccharoperbutylacetonicum N1-4_28HMT_29 GCA_000340885.1
369 Clostridium saccharobutylicum DSM 13864 GCA_000473995.1
1U
780 Streptococcus mutans UA159 GCA_000007465.2
586 Streptococcus ratti FA-1 = DSM 20564 GCA_000286075.1
537 Streptococcus gordonii str. Challis substr. CH1 GCA_000017005.1
1V
720 Staphylococcus epidermidis ATCC 12228 GCA_000007645.1
659 Staphylococcus capitis subsp. capitis GCA_001028645.1
635 Staphylococcus aureus subsp. aureus NCTC 8325 GCA_000013425.1
1W
485 Bifidobacterium adolescentis ATCC 15703 GCA_000010425.1
451 Bifidobacterium dentium JCM 1195 = DSM 20436 GCA_001042595.1
440 Bifidobacterium angulatum DSM 20098 = JCM 7096 GCA_001025155.1
1X
723 Enterococcus faecalis V583 GCA_000007785.1
676 Streptomyces cinnamoneus GCA_001885705.1
629 Enterococcus saccharolyticus subsp. saccharolyticus ATCC 43076 GCA_000407285.1
1Y
793 Staphylococcus epidermidis ATCC 12228 GCA_000007645.1
755 Staphylococcus capitis subsp. capitis GCA_001028645.1
708 Staphylococcus haemolyticus JCSC1435 GCA_000009865.1
1Z

736 Staphylococcus epidermidis ATCC 12228 GCA_000007645.1
673 Staphylococcus capitis subsp. capitis GCA_001028645.1
637 Staphylococcus hominis subsp. hominis C80 GCA_000183685.1
20
634 Deinococcus radiodurans R1 GCA_000008565.1
393 Deinococcus deserti VCD115 GCA_000020685.1
341 Deinococcus hopiensis KR-140 GCA_900176165.1
21
445 Lactobacillus gasseri ATCC 33323 = JCM 1131 GCA_000014425.1
432 Lactobacillus hominis DSM 23910 = CRBIP 24.179 GCA_000296835.1
334 Lactobacillus hamsteri DSM 5661 = JCM 6256 GCA_000615445.1
22
767 Clostridium beijerinckii GCA_000833105.2
743 Clostridium saccharoperbutylacetonicum N1-4_28HMT_29 GCA_000340885.1
706 Clostridium saccharobutylicum DSM 13864 GCA_000473995.1
23
778 Bacillus anthracis str. Ames GCA_000007845.1
778 Bacillus anthracis str. Sterne GCA_000008165.1
778 _5BBacillus thuringiensis_5D serovar konkukian str. 97-27 GCA_000008505.1
763 Bacillus cereus ATCC 14579 GCA_000007825.1
763 Bacillus pseudomycoides DSM 12442 GCA_000161455.1
748 Bacillus thuringiensis YBT-1518 GCA_000497525.2
24
616 Staphylococcus epidermidis ATCC 12228 GCA_000007645.1
605 Staphylococcus capitis subsp. capitis GCA_001028645.1
551 Staphylococcus warneri SG1 GCA_000332735.1
25
710 Staphylococcus epidermidis ATCC 12228 GCA_000007645.1
660 Staphylococcus capitis subsp. capitis GCA_001028645.1
605 Staphylococcus haemolyticus JCSC1435 GCA_000009865.1
26
502 Clostridium beijerinckii GCA_000833105.2
457 Clostridium saccharoperbutylacetonicum N1-4_28HMT_29 GCA_000340885.1
435 Clostridium butyricum GCA_001456065.2
27
996 Streptococcus mutans UA159 GCA_000007465.2
768 Streptococcus ratti FA-1 = DSM 20564 GCA_000286075.1
732 Streptococcus gordonii str. Challis substr. CH1 GCA_000017005.1
28
586 Clostridium beijerinckii GCA_000833105.2
548 Clostridium saccharoperbutylacetonicum N1-4_28HMT_29 GCA_000340885.1
529 Clostridium butyricum GCA_001456065.2
29
865 Lactobacillus gasseri ATCC 33323 = JCM 1131 GCA_000014425.1
794 Lactobacillus hominis DSM 23910 = CRBIP 24.179 GCA_000296835.1
633 Lactobacillus iners DSM 13335 GCA_000160875.1
2a
623 Deinococcus radiodurans R1 GCA_000008565.1
402 Deinococcus gobiensis I-0 GCA_000252445.1
367 Deinococcus pimensis DSM 21231 GCA_000519345.1
2b
676 Staphylococcus epidermidis ATCC 12228 GCA_000007645.1
659 Staphylococcus capitis subsp. capitis GCA_001028645.1
629 Staphylococcus warneri SG1 GCA_000332735.1
2c
777 Clostridium beijerinckii GCA_000833105.2
769 Clostridium saccharoperbutylacetonicum N1-4_28HMT_29 GCA_000340885.1
748 Clostridium saccharobutylicum DSM 13864 GCA_000473995.1
2d
856 Staphylococcus epidermidis ATCC 12228 GCA_000007645.1
829 Staphylococcus capitis subsp. capitis GCA_001028645.1
761 Staphylococcus warneri SG1 GCA_000332735.1
2e
890 Streptococcus mutans UA159 GCA_000007465.2
683 Streptococcus ratti FA-1 = DSM 20564 GCA_000286075.1
642 Streptococcus gordonii str. Challis substr. CH1 GCA_000017005.1
2f
812 Lactobacillus gasseri ATCC 33323 = JCM 1131 GCA_000014425.1
757 Lactobacillus hominis DSM 23910 = CRBIP 24.179 GCA_000296835.1
548 Lactobacillus psittaci DSM 15354 GCA_000425905.1
2g
421 Enterococcus faecalis V583 GCA_000007785.1
374 Streptomyces cinnamoneus GCA_001885705.1
338 Enterococcus faecium DO GCA_000174395.2
2h
766 Rhodobacter sphaeroides 2.4.1 GCA_000012905.2

712 Rhodobacter sphaeroides ATCC 17025 GCA_000016405.1
556 Defluviimonas alba GCA_001620265.1
556 Pseudorhodobacter ferrugineus DSM 5888 GCA_000420745.1
2i
745 Lactobacillus gasseri ATCC 33323 = JCM 1131 GCA_000014425.1
701 Lactobacillus hominis DSM 23910 = CRBIP 24.179 GCA_000296835.1
457 Lactobacillus iners DSM 13335 GCA_000160875.1
2j
588 Escherichia coli str. K-12 substr. MG1655 GCA_000005845.2
563 Escherichia coli O104_3AH4 str. 2011C-3493 GCA_000299455.1
563 Escherichia coli O157_3AH7 str. Sakai GCA_000008865.1
563 Shigella flexneri 2a str. 301 GCA_000006925.2
558 Escherichia coli UMN026 GCA_000026325.2
2k
857 Streptococcus mutans UA159 GCA_000007465.2
630 Streptococcus ratti FA-1 = DSM 20564 GCA_000286075.1
591 Streptococcus ferus DSM 20646 GCA_000372425.1
2l
780 Staphylococcus epidermidis ATCC 12228 GCA_000007645.1
757 Staphylococcus capitis subsp. capitis GCA_001028645.1
703 Staphylococcus warneri SG1 GCA_000332735.1
2m
701 Lactobacillus gasseri ATCC 33323 = JCM 1131 GCA_000014425.1
686 Lactobacillus hominis DSM 23910 = CRBIP 24.179 GCA_000296835.1
550 Lactobacillus iners DSM 13335 GCA_000160875.1
2n
713 Rhodobacter sphaeroides 2.4.1 GCA_000012905.2
611 Rhodobacter sphaeroides ATCC 17025 GCA_000016405.1
472 Gemmobacter aquatilis GCA_900110025.1
2o
702 Escherichia coli str. K-12 substr. MG1655 GCA_000005845.2
688 Escherichia coli O157_3AH7 str. Sakai GCA_000008865.1
688 Escherichia coli UMN026 GCA_000026325.2
684 Escherichia coli IAI39 GCA_000026345.1
2p
712 Enterococcus faecalis V583 GCA_000007785.1
665 Streptomyces cinnamoneus GCA_001885705.1
599 Enterococcus rivorum GCA_001742285.1
2q
541 Staphylococcus epidermidis ATCC 12228 GCA_000007645.1
514 Staphylococcus capitis subsp. capitis GCA_001028645.1
514 Staphylococcus haemolyticus JCSC1435 GCA_000009865.1
499 Staphylococcus aureus subsp. aureus NCTC 8325 GCA_000013425.1
499 Staphylococcus hominis subsp. hominis C80 GCA_000183685.1
499 Staphylococcus lugdunensis HKU09-01 GCA_000025085.1
2r
826 Clostridium beijerinckii GCA_000833105.2
767 Clostridium saccharoperbutylacetonicum N1-4_28HMT_29 GCA_000340885.1
755 Clostridium butyricum GCA_001456065.2
2s
845 Clostridium beijerinckii GCA_000833105.2
803 Clostridium saccharoperbutylacetonicum N1-4_28HMT_29 GCA_000340885.1
775 Clostridium butyricum GCA_001456065.2
775 Clostridium saccharobutylicum DSM 13864 GCA_000473995.1
2t
921 Staphylococcus epidermidis ATCC 12228 GCA_000007645.1
865 Staphylococcus capitis subsp. capitis GCA_001028645.1
780 Staphylococcus aureus subsp. aureus NCTC 8325 GCA_000013425.1
2u
916 Lactobacillus gasseri ATCC 33323 = JCM 1131 GCA_000014425.1
813 Lactobacillus hominis DSM 23910 = CRBIP 24.179 GCA_000296835.1
593 Lactobacillus iners DSM 13335 GCA_000160875.1
2v
589 Lactobacillus gasseri ATCC 33323 = JCM 1131 GCA_000014425.1
555 Lactobacillus hominis DSM 23910 = CRBIP 24.179 GCA_000296835.1
457 Lactobacillus iners DSM 13335 GCA_000160875.1
2w
408 Clostridium beijerinckii GCA_000833105.2
388 Clostridium saccharobutylicum DSM 13864 GCA_000473995.1
375 Clostridium saccharoperbutylacetonicum N1-4_28HMT_29 GCA_000340885.1
2x
584 Rhodobacter sphaeroides 2.4.1 GCA_000012905.2
549 Rhodobacter sphaeroides ATCC 17025 GCA_000016405.1
503 Gemmobacter aquatilis GCA_900110025.1
2y
700 Deinococcus radiodurans R1 GCA_000008565.1

478 Deinococcus gobiensis I-0 GCA_000252445.1
463 Deinococcus deserti VCD115 GCA_000020685.1
2z
903 Clostridium beijerinckii GCA_000833105.2
895 Clostridium saccharoperbutylacetonicum N1-4_28HMT_29 GCA_000340885.1
888 Clostridium saccharobutylicum DSM 13864 GCA_000473995.1
2A
578 Rhodobacter sphaeroides 2.4.1 GCA_000012905.2
501 Rhodobacter sphaeroides ATCC 17025 GCA_000016405.1
426 Gemmobacter aquatilis GCA_900110025.1
426 Rhodovulum sulfidophilum DSM 1374 GCA_001633165.1
2B
516 Rhodobacter sphaeroides 2.4.1 GCA_000012905.2
425 Rhodobacter sphaeroides ATCC 17025 GCA_000016405.1
350 Pseudorhodobacter psychrotolerans GCA_001294535.1
2C
755 Clostridium beijerinckii GCA_000833105.2
736 Clostridium saccharoperbutylacetonicum N1-4_28HMT_29 GCA_000340885.1
712 Clostridium puniceum GCA_002006345.1
2D
442 Rhodobacter sphaeroides 2.4.1 GCA_000012905.2
389 Rhodobacter sphaeroides ATCC 17025 GCA_000016405.1
292 Gemmobacter aquatilis GCA_900110025.1
2E
859 Lactobacillus gasseri ATCC 33323 = JCM 1131 GCA_000014425.1
801 Lactobacillus hominis DSM 23910 = CRBIP 24.179 GCA_000296835.1
617 Lactobacillus iners DSM 13335 GCA_000160875.1
2F
581 Enterococcus faecalis V583 GCA_000007785.1
552 Enterococcus canis NBRC 100695 GCA_001544375.1
547 Enterococcus phoeniculicola ATCC BAA-412 GCA_000407505.1
2G
718 Lactobacillus gasseri ATCC 33323 = JCM 1131 GCA_000014425.1
639 Lactobacillus hominis DSM 23910 = CRBIP 24.179 GCA_000296835.1
419 Lactobacillus iners DSM 13335 GCA_000160875.1
2H
740 Bifidobacterium adolescentis ATCC 15703 GCA_000010425.1
623 Bifidobacterium dentium JCM 1195 = DSM 20436 GCA_001042595.1
594 Bifidobacterium angulatum DSM 20098 = JCM 7096 GCA_001025155.1
2I
544 Lactobacillus gasseri ATCC 33323 = JCM 1131 GCA_000014425.1
465 Lactobacillus hominis DSM 23910 = CRBIP 24.179 GCA_000296835.1
348 Lactobacillus iners DSM 13335 GCA_000160875.1
2J
325 Staphylococcus epidermidis ATCC 12228 GCA_000007645.1
305 Staphylococcus capitis subsp. capitis GCA_001028645.1
274 Staphylococcus lugdunensis HKU09-01 GCA_000025085.1
2K
788 Bifidobacterium adolescentis ATCC 15703 GCA_000010425.1
642 Bifidobacterium dentium JCM 1195 = DSM 20436 GCA_001042595.1
601 Bifidobacterium angulatum DSM 20098 = JCM 7096 GCA_001025155.1
2L
787 Staphylococcus epidermidis ATCC 12228 GCA_000007645.1
726 Staphylococcus capitis subsp. capitis GCA_001028645.1
684 Staphylococcus aureus subsp. aureus NCTC 8325 GCA_000013425.1
2M
604 Bacillus thuringiensis YBT-1518 GCA_000497525.2
588 Bacillus anthracis str. Ames GCA_000007845.1
588 Bacillus anthracis str. Sterne GCA_000008165.1
588 Bacillus cereus ATCC 14579 GCA_000007825.1
588 _5BBacillus thuringiensis_5D serovar konkukian str. 97-27 GCA_000008505.1
565 Bacillus pseudomycoides DSM 12442 GCA_000161455.1
2N
896 Enterococcus faecalis V583 GCA_000007785.1
849 Streptomyces cinnamoneus GCA_001885705.1
783 Enterococcus rivorum GCA_001742285.1
2O
680 Lactobacillus gasseri ATCC 33323 = JCM 1131 GCA_000014425.1
600 Lactobacillus hominis DSM 23910 = CRBIP 24.179 GCA_000296835.1
407 Lactobacillus jensenii GCA_001936235.1
407 Lactobacillus psittaci DSM 15354 GCA_000425905.1
2P
923 Lactobacillus gasseri ATCC 33323 = JCM 1131 GCA_000014425.1
860 Lactobacillus hominis DSM 23910 = CRBIP 24.179 GCA_000296835.1
627 Lactobacillus iners DSM 13335 GCA_000160875.1
2Q

706 Lactobacillus gasseri ATCC 33323 = JCM 1131 GCA_000014425.1
660 Lactobacillus hominis DSM 23910 = CRBIP 24.179 GCA_000296835.1
449 Lactobacillus iners DSM 13335 GCA_000160875.1
2R
591 Clostridium beijerinckii GCA_000833105.2
578 Clostridium saccharobutylicum DSM 13864 GCA_000473995.1
574 Clostridium puniceum GCA_002006345.1
2S
776 Enterococcus faecalis V583 GCA_000007785.1
729 Streptomyces cinnamoneus GCA_001885705.1
653 Enterococcus rivorum GCA_001742285.1
2T
609 Clostridium beijerinckii GCA_000833105.2
587 Clostridium saccharoperbutylacetonicum N1-4_28HMT_29 GCA_000340885.1
547 Clostridium puniceum GCA_002006345.1
2U
858 Enterococcus faecalis V583 GCA_000007785.1
814 Streptomyces cinnamoneus GCA_001885705.1
773 Enterococcus rivorum GCA_001742285.1
2V
1085 Streptococcus mutans UA159 GCA_000007465.2
807 Streptococcus ratti FA-1 = DSM 20564 GCA_000286075.1
757 Streptococcus gordonii str. Challis substr. CH1 GCA_000017005.1
2W
816 Clostridium beijerinckii GCA_000833105.2
771 Clostridium saccharoperbutylacetonicum N1-4_28HMT_29 GCA_000340885.1
758 Clostridium puniceum GCA_002006345.1
2X
464 Bifidobacterium adolescentis ATCC 15703 GCA_000010425.1
385 Bifidobacterium dentium JCM 1195 = DSM 20436 GCA_001042595.1
370 Bifidobacterium angulatum DSM 20098 = JCM 7096 GCA_001025155.1
2Y
376 Lactobacillus gasseri ATCC 33323 = JCM 1131 GCA_000014425.1
332 Lactobacillus hominis DSM 23910 = CRBIP 24.179 GCA_000296835.1
253 Lactobacillus iners DSM 13335 GCA_000160875.1
2Z
705 Clostridium beijerinckii GCA_000833105.2
676 Clostridium saccharoperbutylacetonicum N1-4_28HMT_29 GCA_000340885.1
648 Clostridium saccharobutylicum DSM 13864 GCA_000473995.1
30
792 Enterococcus faecalis V583 GCA_000007785.1
748 Streptomyces cinnamoneus GCA_001885705.1
710 Enterococcus rivorum GCA_001742285.1
31
839 Enterococcus faecalis V583 GCA_000007785.1
792 Streptomyces cinnamoneus GCA_001885705.1
712 Enterococcus rivorum GCA_001742285.1
32
841 Enterococcus faecalis V583 GCA_000007785.1
794 Streptomyces cinnamoneus GCA_001885705.1
735 Enterococcus haemoperoxidus ATCC BAA-382 GCA_000407165.1
33
595 Streptococcus mutans UA159 GCA_000007465.2
441 Streptococcus ratti FA-1 = DSM 20564 GCA_000286075.1
419 Streptococcus equinus GCA_000964315.1
419 Streptococcus gallolyticus subsp. gallolyticus DSM 16831 GCA_002000985.1
34
830 Lactobacillus gasseri ATCC 33323 = JCM 1131 GCA_000014425.1
759 Lactobacillus hominis DSM 23910 = CRBIP 24.179 GCA_000296835.1
525 Lactobacillus iners DSM 13335 GCA_000160875.1
35
754 Staphylococcus epidermidis ATCC 12228 GCA_000007645.1
702 Staphylococcus capitis subsp. capitis GCA_001028645.1
633 Staphylococcus aureus subsp. aureus NCTC 8325 GCA_000013425.1
36
777 Streptococcus mutans UA159 GCA_000007465.2
581 Streptococcus ratti FA-1 = DSM 20564 GCA_000286075.1
532 Streptococcus gordonii str. Challis substr. CH1 GCA_000017005.1
37
737 Deinococcus radiodurans R1 GCA_000008565.1
423 Deinococcus deserti VCD115 GCA_000020685.1
419 Deinococcus puniceus GCA_001644565.1
38
753 Streptococcus mutans UA159 GCA_000007465.2
501 Streptococcus ratti FA-1 = DSM 20564 GCA_000286075.1
457 Streptococcus gordonii str. Challis substr. CH1 GCA_000017005.1

39
684 Clostridium beijerinckii GCA_000833105.2
635 Clostridium saccharoperbutylacetonicum N1-4_28HMT_29 GCA_000340885.1
628 Clostridium puniceum GCA_002006345.1
3a
527 Clostridium beijerinckii GCA_000833105.2
482 Clostridium puniceum GCA_002006345.1
482 Clostridium saccharoperbutylacetonicum N1-4_28HMT_29 GCA_000340885.1
463 Clostridium saccharobutylicum DSM 13864 GCA_000473995.1
3b
648 Streptococcus mutans UA159 GCA_000007465.2
446 Streptococcus ratti FA-1 = DSM 20564 GCA_000286075.1
433 Streptococcus ferus DSM 20646 GCA_000372425.1
3c
366 Staphylococcus epidermidis ATCC 12228 GCA_000007645.1
358 Staphylococcus haemolyticus JCSC1435 GCA_000009865.1
343 Staphylococcus hominis subsp. hominis C80 GCA_000183685.1
343 Staphylococcus lugdunensis HKU09-01 GCA_000025085.1
3d
711 Bifidobacterium adolescentis ATCC 15703 GCA_000010425.1
561 Bifidobacterium dentium JCM 1195 = DSM 20436 GCA_001042595.1
542 Bifidobacterium angulatum DSM 20098 = JCM 7096 GCA_001025155.1
3e
722 Deinococcus radiodurans R1 GCA_000008565.1
444 Deinococcus deserti VCD115 GCA_000020685.1
403 Deinococcus gobiensis I-0 GCA_000252445.1
3f
570 Clostridium beijerinckii GCA_000833105.2
570 Clostridium saccharoperbutylacetonicum N1-4_28HMT_29 GCA_000340885.1
549 Clostridium puniceum GCA_002006345.1
527 Clostridium butyricum GCA_001456065.2
3g
643 Deinococcus radiodurans R1 GCA_000008565.1
468 Deinococcus gobiensis I-0 GCA_000252445.1
433 Deinococcus soli Cha et al. 2016 GCA_001007995.1
3h
617 Clostridium beijerinckii GCA_000833105.2
595 Clostridium saccharoperbutylacetonicum N1-4_28HMT_29 GCA_000340885.1
572 Clostridium saccharobutylicum DSM 13864 GCA_000473995.1
3i
782 Enterococcus faecalis V583 GCA_000007785.1
735 Streptomyces cinnamoneus GCA_001885705.1
693 Enterococcus rivorum GCA_001742285.1
3j
716 Clostridium beijerinckii GCA_000833105.2
691 Clostridium saccharoperbutylacetonicum N1-4_28HMT_29 GCA_000340885.1
672 Clostridium saccharobutylicum DSM 13864 GCA_000473995.1
3k
711 Bifidobacterium adolescentis ATCC 15703 GCA_000010425.1
615 Bifidobacterium dentium JCM 1195 = DSM 20436 GCA_001042595.1
565 Bifidobacterium angulatum DSM 20098 = JCM 7096 GCA_001025155.1
3l
893 Lactobacillus gasseri ATCC 33323 = JCM 1131 GCA_000014425.1
826 Lactobacillus hominis DSM 23910 = CRBIP 24.179 GCA_000296835.1
653 Lactobacillus iners DSM 13335 GCA_000160875.1
3m
729 Streptococcus mutans UA159 GCA_000007465.2
503 Streptococcus ratti FA-1 = DSM 20564 GCA_000286075.1
498 Streptococcus macacae NCTC 11558 GCA_000187995.3
3n
732 Escherichia coli O104_3AH4 str. 2011C-3493 GCA_000299455.1
732 Escherichia coli str. K-12 substr. MG1655 GCA_000005845.2
724 Escherichia coli IAI39 GCA_000026345.1
717 Escherichia coli O157_3AH7 str. Sakai GCA_000008865.1
717 Shigella flexneri 2a str. 301 GCA_000006925.2
3o
462 Streptococcus mutans UA159 GCA_000007465.2
360 Streptococcus ferus DSM 20646 GCA_000372425.1
355 Streptococcus ratti FA-1 = DSM 20564 GCA_000286075.1
3p
815 Enterococcus faecalis V583 GCA_000007785.1
768 Streptomyces cinnamoneus GCA_001885705.1
700 Enterococcus asini ATCC 700915 GCA_000407365.1
3q
551 Streptococcus mutans UA159 GCA_000007465.2
418 Streptococcus ratti FA-1 = DSM 20564 GCA_000286075.1

361 Streptococcus equinus GCA_000964315.1
3r
724 Streptococcus mutans UA159 GCA_000007465.2
528 Streptococcus gordonii str. Challis substr. CH1 GCA_000017005.1
508 Streptococcus ratti FA-1 = DSM 20564 GCA_000286075.1
3s
644 Lactobacillus gasseri ATCC 33323 = JCM 1131 GCA_000014425.1
582 Lactobacillus hominis DSM 23910 = CRBIP 24.179 GCA_000296835.1
447 Lactobacillus iners DSM 13335 GCA_000160875.1
3t
837 Deinococcus radiodurans R1 GCA_000008565.1
533 Deinococcus deserti VCD115 GCA_000020685.1
509 Deinococcus gobiensis I-0 GCA_000252445.1
3u
804 Staphylococcus epidermidis ATCC 12228 GCA_000007645.1
789 Staphylococcus capitis subsp. capitis GCA_001028645.1
703 Staphylococcus aureus subsp. aureus NCTC 8325 GCA_000013425.1
3v
670 Escherichia coli str. K-12 substr. MG1655 GCA_000005845.2
659 Escherichia coli O104_3AH4 str. 2011C-3493 GCA_000299455.1
644 Shigella flexneri 2a str. 301 GCA_000006925.2
3w
795 Bifidobacterium adolescentis ATCC 15703 GCA_000010425.1
699 Bifidobacterium angulatum DSM 20098 = JCM 7096 GCA_001025155.1
668 Bifidobacterium longum NCC2705 GCA_000007525.1
3x
708 Bifidobacterium adolescentis ATCC 15703 GCA_000010425.1
633 Bifidobacterium dentium JCM 1195 = DSM 20436 GCA_001042595.1
631 Bifidobacterium angulatum DSM 20098 = JCM 7096 GCA_001025155.1
3y
416 Clostridium beijerinckii GCA_000833105.2
398 Clostridium saccharoperbutylacetonicum N1-4_28HMT_29 GCA_000340885.1
374 Clostridium butyricum GCA_001456065.2
3z
752 Clostridium beijerinckii GCA_000833105.2
714 Clostridium saccharoperbutylacetonicum N1-4_28HMT_29 GCA_000340885.1
696 Clostridium puniceum GCA_002006345.1
3A
800 Bacillus thuringiensis YBT-1518 GCA_000497525.2
789 Bacillus anthracis str. Ames GCA_000007845.1
789 Bacillus anthracis str. Sterne GCA_000008165.1
789 Bacillus cereus ATCC 14579 GCA_000007825.1
789 _5BBacillus thuringiensis_5D serovar konkukian str. 97-27 GCA_000008505.1
784 Bacillus pseudomycoides DSM 12442 GCA_000161455.1
3B
605 Deinococcus radiodurans R1 GCA_000008565.1
396 Deinococcus deserti VCD115 GCA_000020685.1
355 Deinococcus proteolyticus MRP GCA_000190555.1
3C
657 Rhodobacter sphaeroides 2.4.1 GCA_000012905.2
622 Rhodobacter sphaeroides ATCC 17025 GCA_000016405.1
551 Pseudorhodobacter psychrotolerans GCA_001294535.1
3D
720 Staphylococcus epidermidis ATCC 12228 GCA_000007645.1
633 Staphylococcus capitis subsp. capitis GCA_001028645.1
578 Staphylococcus hominis subsp. hominis C80 GCA_000183685.1
3E
738 Staphylococcus epidermidis ATCC 12228 GCA_000007645.1
715 Staphylococcus capitis subsp. capitis GCA_001028645.1
629 Staphylococcus lugdunensis HKU09-01 GCA_000025085.1
3F
793 Deinococcus radiodurans R1 GCA_000008565.1
494 Deinococcus deserti VCD115 GCA_000020685.1
449 Deinococcus gobiensis I-0 GCA_000252445.1
3G
662 Clostridium beijerinckii GCA_000833105.2
611 Clostridium puniceum GCA_002006345.1
611 Clostridium saccharobutylicum DSM 13864 GCA_000473995.1
611 Clostridium saccharoperbutylacetonicum N1-4_28HMT_29 GCA_000340885.1
582 Clostridium butyricum GCA_001456065.2
3H
669 Enterococcus faecalis V583 GCA_000007785.1
622 Streptomyces cinnamoneus GCA_001885705.1
600 Enterococcus faecium DO GCA_000174395.2
3I
775 Deinococcus radiodurans R1 GCA_000008565.1

468 Deinococcus soli Cha et al. 2016 GCA_001007995.1
436 Deinococcus deserti VCD115 GCA_000020685.1
3J
586 Streptococcus mutans UA159 GCA_000007465.2
440 Streptococcus ratti FA-1 = DSM 20564 GCA_000286075.1
424 Streptococcus halotolerans GCA_001598035.1
3K
761 Staphylococcus epidermidis ATCC 12228 GCA_000007645.1
747 Staphylococcus capitis subsp. capitis GCA_001028645.1
672 Staphylococcus warneri SG1 GCA_000332735.1
3L
857 Staphylococcus epidermidis ATCC 12228 GCA_000007645.1
808 Staphylococcus capitis subsp. capitis GCA_001028645.1
782 Megasphaera cerevisiae DSM 20462 GCA_001045675.1
782 Staphylococcus warneri SG1 GCA_000332735.1
3M
548 Streptococcus mutans UA159 GCA_000007465.2
347 Streptococcus ratti FA-1 = DSM 20564 GCA_000286075.1
330 Streptococcus macacae NCTC 11558 GCA_000187995.3
3N
503 Escherichia coli IAI39 GCA_000026345.1
503 Escherichia coli O157_3AH7 str. Sakai GCA_000008865.1
503 Escherichia coli str. K-12 substr. MG1655 GCA_000005845.2
495 Escherichia coli UMN026 GCA_000026325.2
477 Escherichia coli O83_3AH1 str. NRG 857C GCA_000183345.1
3O
438 Clostridium beijerinckii GCA_000833105.2
406 Clostridium saccharoperbutylacetonicum N1-4_28HMT_29 GCA_000340885.1
385 Clostridium saccharobutylicum DSM 13864 GCA_000473995.1
3P
654 Staphylococcus epidermidis ATCC 12228 GCA_000007645.1
605 Staphylococcus capitis subsp. capitis GCA_001028645.1
552 Megasphaera cerevisiae DSM 20462 GCA_001045675.1
552 Staphylococcus warneri SG1 GCA_000332735.1
3Q
642 Bacillus anthracis str. Ames GCA_000007845.1
642 Bacillus anthracis str. Sterne GCA_000008165.1
642 Bacillus cereus ATCC 14579 GCA_000007825.1
642 _5BBacillus thuringiensis_5D serovar konkukian str. 97-27 GCA_000008505.1
621 Bacillus thuringiensis YBT-1518 GCA_000497525.2
620 Bacillus mycoides GCA_000832605.1
3R
784 Streptococcus mutans UA159 GCA_000007465.2
575 Streptococcus ratti FA-1 = DSM 20564 GCA_000286075.1
531 Streptococcus gordonii str. Challis substr. CH1 GCA_000017005.1
3S
784 Deinococcus radiodurans R1 GCA_000008565.1
531 Deinococcus deserti VCD115 GCA_000020685.1
485 Deinococcus gobiensis I-0 GCA_000252445.1
3T
752 Lactobacillus gasseri ATCC 33323 = JCM 1131 GCA_000014425.1
730 Lactobacillus hominis DSM 23910 = CRBIP 24.179 GCA_000296835.1
553 Lactobacillus iners DSM 13335 GCA_000160875.1
3U
454 Clostridium saccharoperbutylacetonicum N1-4_28HMT_29 GCA_000340885.1
453 Clostridium beijerinckii GCA_000833105.2
451 Clostridium saccharobutylicum DSM 13864 GCA_000473995.1
3V
750 Staphylococcus epidermidis ATCC 12228 GCA_000007645.1
733 Staphylococcus capitis subsp. capitis GCA_001028645.1
673 Staphylococcus warneri SG1 GCA_000332735.1
3W
598 Clostridium beijerinckii GCA_000833105.2
588 Clostridium saccharoperbutylacetonicum N1-4_28HMT_29 GCA_000340885.1
559 Clostridium puniceum GCA_002006345.1
3X
723 Enterococcus faecalis V583 GCA_000007785.1
677 Streptomyces cinnamoneus GCA_001885705.1
622 Enterococcus haemoperoxidus ATCC BAA-382 GCA_000407165.1
3Y
613 Clostridium beijerinckii GCA_000833105.2
576 Clostridium puniceum GCA_002006345.1
576 Clostridium saccharoperbutylacetonicum N1-4_28HMT_29 GCA_000340885.1
561 Clostridium butyricum GCA_001456065.2
3Z
768 Deinococcus radiodurans R1 GCA_000008565.1

507 Deinococcus deserti VCD115 GCA_000020685.1
456 Deinococcus murrayi DSM 11303 GCA_000482805.1
40
683 Staphylococcus epidermidis ATCC 12228 GCA_000007645.1
626 Staphylococcus capitis subsp. capitis GCA_001028645.1
518 Staphylococcus aureus subsp. aureus NCTC 8325 GCA_000013425.1
41
468 Rhodobacter sphaeroides 2.4.1 GCA_000012905.2
403 Rhodobacter sphaeroides ATCC 17025 GCA_000016405.1
318 Defluviimonas indica GCA_900106675.1
42
738 Escherichia coli O157_3AH7 str. Sakai GCA_000008865.1
738 Escherichia coli str. K-12 substr. MG1655 GCA_000005845.2
719 Escherichia coli IAI39 GCA_000026345.1
709 Escherichia coli UMN026 GCA_000026325.2
43
475 Escherichia coli O157_3AH7 str. Sakai GCA_000008865.1
475 Escherichia coli str. K-12 substr. MG1655 GCA_000005845.2
471 Escherichia coli IAI39 GCA_000026345.1
470 Escherichia coli UMN026 GCA_000026325.2
44
690 Enterococcus faecalis V583 GCA_000007785.1
643 Streptomyces cinnamoneus GCA_001885705.1
620 Enterococcus rivorum GCA_001742285.1
45
851 Bacillus cereus ATCC 14579 GCA_000007825.1
851 Bacillus thuringiensis YBT-1518 GCA_000497525.2
849 Bacillus anthracis str. Ames GCA_000007845.1
849 Bacillus anthracis str. Sterne GCA_000008165.1
849 _5BBacillus thuringiensis_5D serovar konkukian str. 97-27 GCA_000008505.1
834 Bacillus pseudomycoides DSM 12442 GCA_000161455.1
46
349 Clostridium saccharoperbutylacetonicum N1-4_28HMT_29 GCA_000340885.1
346 Clostridium beijerinckii GCA_000833105.2
344 Clostridium butyricum GCA_001456065.2
344 Clostridium saccharobutylicum DSM 13864 GCA_000473995.1
47
794 Streptococcus mutans UA159 GCA_000007465.2
550 Streptococcus ratti FA-1 = DSM 20564 GCA_000286075.1
510 Streptococcus gordonii str. Challis substr. CH1 GCA_000017005.1
48
728 Clostridium beijerinckii GCA_000833105.2
700 Clostridium saccharoperbutylacetonicum N1-4_28HMT_29 GCA_000340885.1
681 Clostridium puniceum GCA_002006345.1
49
494 Rhodobacter sphaeroides 2.4.1 GCA_000012905.2
389 Gemmobacter aquatilis GCA_900110025.1
383 Rhodobacter sphaeroides ATCC 17025 GCA_000016405.1
4a
432 Clostridium beijerinckii GCA_000833105.2
415 Clostridium saccharoperbutylacetonicum N1-4_28HMT_29 GCA_000340885.1
393 Clostridium butyricum GCA_001456065.2
4b
774 Streptococcus mutans UA159 GCA_000007465.2
599 Streptococcus ratti FA-1 = DSM 20564 GCA_000286075.1
537 Streptococcus ferus DSM 20646 GCA_000372425.1
4c
746 Streptococcus mutans UA159 GCA_000007465.2
488 Streptococcus gordonii str. Challis substr. CH1 GCA_000017005.1
486 Streptococcus ratti FA-1 = DSM 20564 GCA_000286075.1
4d
1002 Streptococcus mutans UA159 GCA_000007465.2
762 Streptococcus ratti FA-1 = DSM 20564 GCA_000286075.1
666 Streptococcus gordonii str. Challis substr. CH1 GCA_000017005.1
4e
780 Lactobacillus gasseri ATCC 33323 = JCM 1131 GCA_000014425.1
719 Lactobacillus hominis DSM 23910 = CRBIP 24.179 GCA_000296835.1
554 Lactobacillus iners DSM 13335 GCA_000160875.1
4f
708 Staphylococcus epidermidis ATCC 12228 GCA_000007645.1
661 Staphylococcus capitis subsp. capitis GCA_001028645.1
655 Staphylococcus haemolyticus JCSC1435 GCA_000009865.1
4g
702 Rhodobacter sphaeroides 2.4.1 GCA_000012905.2
621 Rhodobacter sphaeroides ATCC 17025 GCA_000016405.1
500 Defluviimonas alba GCA_001620265.1

4h
774 Enterococcus faecalis V583 GCA_000007785.1
731 Enterococcus canis NBRC 100695 GCA_001544375.1
727 Streptomyces cinnamoneus GCA_001885705.1
4i
926 Clostridium beijerinckii GCA_000833105.2
898 Clostridium saccharoperbutylacetonicum N1-4_28HMT_29 GCA_000340885.1
876 Clostridium puniceum GCA_002006345.1
4j
929 Staphylococcus epidermidis ATCC 12228 GCA_000007645.1
877 Staphylococcus capitis subsp. capitis GCA_001028645.1
836 Staphylococcus haemolyticus JCSC1435 GCA_000009865.1
4k
434 Escherichia coli O104_3AH4 str. 2011C-3493 GCA_000299455.1
434 Escherichia coli O157_3AH7 str. Sakai GCA_000008865.1
434 Escherichia coli O83_3AH1 str. NRG 857C GCA_000183345.1
434 Escherichia coli UMN026 GCA_000026325.2
434 Escherichia coli str. K-12 substr. MG1655 GCA_000005845.2
434 Shigella flexneri 2a str. 301 GCA_000006925.2
409 Escherichia coli IAI39 GCA_000026345.1
406 Salmonella enterica subsp. enterica serovar Typhimurium str. LT2 GCA_000006945.2
4l
598 Staphylococcus epidermidis ATCC 12228 GCA_000007645.1
559 Staphylococcus capitis subsp. capitis GCA_001028645.1
527 Staphylococcus hominis subsp. hominis C80 GCA_000183685.1
4m
591 Staphylococcus epidermidis ATCC 12228 GCA_000007645.1
515 Staphylococcus capitis subsp. capitis GCA_001028645.1
513 Staphylococcus haemolyticus JCSC1435 GCA_000009865.1
513 Staphylococcus hominis subsp. hominis C80 GCA_000183685.1
4n
613 Lactobacillus gasseri ATCC 33323 = JCM 1131 GCA_000014425.1
576 Lactobacillus hominis DSM 23910 = CRBIP 24.179 GCA_000296835.1
475 Lactobacillus iners DSM 13335 GCA_000160875.1
4o
681 Streptococcus mutans UA159 GCA_000007465.2
501 Streptococcus ratti FA-1 = DSM 20564 GCA_000286075.1
469 Streptococcus macacae NCTC 11558 GCA_000187995.3
4p
488 Bacillus thuringiensis YBT-1518 GCA_000497525.2
480 Bacillus anthracis str. Ames GCA_000007845.1
480 Bacillus anthracis str. Sterne GCA_000008165.1
480 Bacillus pseudomycoides DSM 12442 GCA_000161455.1
480 _5BBacillus thuringiensis_5D serovar konkukian str. 97-27 GCA_000008505.1
479 Bacillus cereus ATCC 14579 GCA_000007825.1
4q
599 Streptococcus mutans UA159 GCA_000007465.2
464 Streptococcus ratti FA-1 = DSM 20564 GCA_000286075.1
432 Streptococcus gordonii str. Challis substr. CH1 GCA_000017005.1
4r
644 Bifidobacterium adolescentis ATCC 15703 GCA_000010425.1
552 Bifidobacterium dentium JCM 1195 = DSM 20436 GCA_001042595.1
529 Bifidobacterium angulatum DSM 20098 = JCM 7096 GCA_001025155.1
4s
717 Bacillus anthracis str. Ames GCA_000007845.1
717 Bacillus anthracis str. Sterne GCA_000008165.1
717 Bacillus cereus ATCC 14579 GCA_000007825.1
717 _5BBacillus thuringiensis_5D serovar konkukian str. 97-27 GCA_000008505.1
712 Bacillus thuringiensis YBT-1518 GCA_000497525.2
701 Bacillus pseudomycoides DSM 12442 GCA_000161455.1
4t
807 Clostridium beijerinckii GCA_000833105.2
755 Clostridium saccharobutylicum DSM 13864 GCA_000473995.1
747 Clostridium puniceum GCA_002006345.1
4u
750 Escherichia coli str. K-12 substr. MG1655 GCA_000005845.2
739 Escherichia coli O104_3AH4 str. 2011C-3493 GCA_000299455.1
739 Escherichia coli O157_3AH7 str. Sakai GCA_000008865.1
739 Shigella flexneri 2a str. 301 GCA_000006925.2
735 Escherichia coli IAI39 GCA_000026345.1
735 Shigella dysenteriae Sd197 GCA_000012005.1
4v
483 Streptococcus mutans UA159 GCA_000007465.2
389 Lactobacillus gasseri ATCC 33323 = JCM 1131 GCA_000014425.1
374 Lactobacillus hominis DSM 23910 = CRBIP 24.179 GCA_000296835.1
4w

765 Deinococcus radiodurans R1 GCA_000008565.1
452 Deinococcus deserti VCD115 GCA_000020685.1
413 Deinococcus soli Cha et al. 2016 GCA_001007995.1
4x
799 Clostridium beijerinckii GCA_000833105.2
775 Clostridium saccharoperbutylacetonicum N1-4_28HMT_29 GCA_000340885.1
743 Clostridium puniceum GCA_002006345.1
4y
509 Streptococcus mutans UA159 GCA_000007465.2
415 Streptococcus equinus GCA_000964315.1
402 Streptococcus anginosus C238 GCA_000463505.1
4z
758 Deinococcus radiodurans R1 GCA_000008565.1
481 Deinococcus deserti VCD115 GCA_000020685.1
477 Deinococcus gobiensis I-0 GCA_000252445.1
4A
899 Staphylococcus epidermidis ATCC 12228 GCA_000007645.1
858 Staphylococcus capitis subsp. capitis GCA_001028645.1
788 Staphylococcus aureus subsp. aureus NCTC 8325 GCA_000013425.1
4B
733 Staphylococcus epidermidis ATCC 12228 GCA_000007645.1
688 Staphylococcus capitis subsp. capitis GCA_001028645.1
654 Staphylococcus aureus subsp. aureus NCTC 8325 GCA_000013425.1
4C
737 Escherichia coli O157_3AH7 str. Sakai GCA_000008865.1
737 Escherichia coli str. K-12 substr. MG1655 GCA_000005845.2
731 Escherichia coli O83_3AH1 str. NRG 857C GCA_000183345.1
723 Shigella flexneri 2a str. 301 GCA_000006925.2
4D
825 Rhodobacter sphaeroides 2.4.1 GCA_000012905.2
780 Rhodobacter sphaeroides ATCC 17025 GCA_000016405.1
594 Gemmobacter aquatilis GCA_900110025.1
594 Pseudorhodobacter ferrugineus DSM 5888 GCA_000420745.1
4E
677 Deinococcus radiodurans R1 GCA_000008565.1
419 Deinococcus deserti VCD115 GCA_000020685.1
410 Deinococcus puniceus GCA_001644565.1
4F
801 Streptococcus mutans UA159 GCA_000007465.2
606 Streptococcus ratti FA-1 = DSM 20564 GCA_000286075.1
589 Streptococcus equinus GCA_000964315.1
4G
652 Staphylococcus epidermidis ATCC 12228 GCA_000007645.1
623 Staphylococcus capitis subsp. capitis GCA_001028645.1
593 Staphylococcus aureus subsp. aureus NCTC 8325 GCA_000013425.1
4H
686 Bifidobacterium adolescentis ATCC 15703 GCA_000010425.1
558 Bifidobacterium dentium JCM 1195 = DSM 20436 GCA_001042595.1
541 Bifidobacterium angulatum DSM 20098 = JCM 7096 GCA_001025155.1
4I
614 Staphylococcus epidermidis ATCC 12228 GCA_000007645.1
556 Staphylococcus capitis subsp. capitis GCA_001028645.1
526 Staphylococcus warneri SG1 GCA_000332735.1
4J
600 Rhodobacter sphaeroides 2.4.1 GCA_000012905.2
503 Rhodobacter sphaeroides ATCC 17025 GCA_000016405.1
459 Gemmobacter megaterium GCA_900156815.1
4K
535 Lactobacillus gasseri ATCC 33323 = JCM 1131 GCA_000014425.1
528 Clostridium beijerinckii GCA_000833105.2
515 Clostridium saccharoperbutylacetonicum N1-4_28HMT_29 GCA_000340885.1
4L
658 Streptococcus mutans UA159 GCA_000007465.2
505 Streptococcus ratti FA-1 = DSM 20564 GCA_000286075.1
470 Streptococcus gordonii str. Challis substr. CH1 GCA_000017005.1
4M
454 Bifidobacterium adolescentis ATCC 15703 GCA_000010425.1
395 Bifidobacterium dentium JCM 1195 = DSM 20436 GCA_001042595.1
356 Bifidobacterium stellenboschense GCA_000741785.1
4N
662 Clostridium beijerinckii GCA_000833105.2
618 Clostridium saccharoperbutylacetonicum N1-4_28HMT_29 GCA_000340885.1
590 Clostridium puniceum GCA_002006345.1
590 Clostridium saccharobutylicum DSM 13864 GCA_000473995.1
4O
703 Staphylococcus epidermidis ATCC 12228 GCA_000007645.1

698 Staphylococcus capitis subsp. capitis GCA_001028645.1
659 Staphylococcus aureus subsp. aureus NCTC 8325 GCA_000013425.1
4P
578 Rhodobacter sphaeroides 2.4.1 GCA_000012905.2
519 Rhodobacter sphaeroides ATCC 17025 GCA_000016405.1
432 Gemmobacter aquatilis GCA_900110025.1
432 Gemmobacter megaterium GCA_900156815.1
4Q
835 Bacillus anthracis str. Ames GCA_000007845.1
835 Bacillus anthracis str. Sterne GCA_000008165.1
835 _5BBacillus thuringiensis_5D serovar konkukian str. 97-27 GCA_000008505.1
824 Bacillus pseudomycoides DSM 12442 GCA_000161455.1
820 Bacillus cereus ATCC 14579 GCA_000007825.1
4R
594 Bacillus anthracis str. Ames GCA_000007845.1
594 Bacillus anthracis str. Sterne GCA_000008165.1
594 _5BBacillus thuringiensis_5D serovar konkukian str. 97-27 GCA_000008505.1
586 Bacillus cereus ATCC 14579 GCA_000007825.1
576 Bacillus pseudomycoides DSM 12442 GCA_000161455.1
4S
627 Lactobacillus gasseri ATCC 33323 = JCM 1131 GCA_000014425.1
561 Lactobacillus hominis DSM 23910 = CRBIP 24.179 GCA_000296835.1
475 Lactobacillus crispatus ST1 GCA_000091765.1
4T
453 Lactobacillus gasseri ATCC 33323 = JCM 1131 GCA_000014425.1
426 Lactobacillus hominis DSM 23910 = CRBIP 24.179 GCA_000296835.1
325 Lactobacillus iners DSM 13335 GCA_000160875.1
4U
647 Lactobacillus gasseri ATCC 33323 = JCM 1131 GCA_000014425.1
561 Lactobacillus hominis DSM 23910 = CRBIP 24.179 GCA_000296835.1
438 Lactobacillus iners DSM 13335 GCA_000160875.1
4V
353 Rhodobacter sphaeroides 2.4.1 GCA_000012905.2
316 Rhodobacter sphaeroides ATCC 17025 GCA_000016405.1
281 Rhodobacter vinaykumarii GCA_900156695.1
4W
720 Rhodobacter sphaeroides 2.4.1 GCA_000012905.2
608 Rhodobacter sphaeroides ATCC 17025 GCA_000016405.1
511 Gemmobacter megaterium GCA_900156815.1
4X
805 Clostridium beijerinckii GCA_000833105.2
794 Clostridium saccharoperbutylacetonicum N1-4_28HMT_29 GCA_000340885.1
751 Clostridium puniceum GCA_002006345.1
4Y
768 Streptococcus mutans UA159 GCA_000007465.2
579 Streptococcus ratti FA-1 = DSM 20564 GCA_000286075.1
565 Streptococcus gordonii str. Challis substr. CH1 GCA_000017005.1
4Z
767 Staphylococcus epidermidis ATCC 12228 GCA_000007645.1
754 Staphylococcus capitis subsp. capitis GCA_001028645.1
693 Staphylococcus aureus subsp. aureus NCTC 8325 GCA_000013425.1
50
870 Enterococcus faecalis V583 GCA_000007785.1
823 Streptomyces cinnamoneus GCA_001885705.1
783 Enterococcus rivorum GCA_001742285.1
51
818 Bifidobacterium adolescentis ATCC 15703 GCA_000010425.1
750 Bifidobacterium dentium JCM 1195 = DSM 20436 GCA_001042595.1
710 Bifidobacterium longum NCC2705 GCA_000007525.1
52
834 Lactobacillus gasseri ATCC 33323 = JCM 1131 GCA_000014425.1
771 Lactobacillus hominis DSM 23910 = CRBIP 24.179 GCA_000296835.1
586 Lactobacillus iners DSM 13335 GCA_000160875.1
53
884 Staphylococcus epidermidis ATCC 12228 GCA_000007645.1
840 Staphylococcus capitis subsp. capitis GCA_001028645.1
812 Staphylococcus aureus subsp. aureus NCTC 8325 GCA_000013425.1
54
700 Bifidobacterium adolescentis ATCC 15703 GCA_000010425.1
564 Bifidobacterium dentium JCM 1195 = DSM 20436 GCA_001042595.1
547 Bifidobacterium callitrichos DSM 23973 GCA_000741175.1
55
655 Streptococcus mutans UA159 GCA_000007465.2
432 Streptococcus gordonii str. Challis substr. CH1 GCA_000017005.1
415 Streptococcus ratti FA-1 = DSM 20564 GCA_000286075.1
56

721 Deinococcus radiodurans R1 GCA_000008565.1
403 Deinococcus deserti VCD115 GCA_000020685.1
394 Deinococcus hopiensis KR-140 GCA_900176165.1
57
772 Deinococcus radiodurans R1 GCA_000008565.1
441 Deinococcus deserti VCD115 GCA_000020685.1
441 Deinococcus gobiensis I-0 GCA_000252445.1
434 Deinococcus soli Cha et al. 2016 GCA_001007995.1
58
640 Deinococcus radiodurans R1 GCA_000008565.1
398 Deinococcus deserti VCD115 GCA_000020685.1
390 Deinococcus soli Cha et al. 2016 GCA_001007995.1
59
905 Deinococcus radiodurans R1 GCA_000008565.1
586 Deinococcus deserti VCD115 GCA_000020685.1
560 Deinococcus murrayi DSM 11303 GCA_000482805.1
5a
603 Rhodobacter sphaeroides 2.4.1 GCA_000012905.2
547 Rhodobacter sphaeroides ATCC 17025 GCA_000016405.1
446 Defluviimonas alba GCA_001620265.1
5b
435 Streptococcus mutans UA159 GCA_000007465.2
302 Streptococcus ratti FA-1 = DSM 20564 GCA_000286075.1
285 Streptococcus gordonii str. Challis substr. CH1 GCA_000017005.1
5c
606 Deinococcus radiodurans R1 GCA_000008565.1
403 Deinococcus deserti VCD115 GCA_000020685.1
391 Deinococcus soli Cha et al. 2016 GCA_001007995.1
5d
764 Lactobacillus gasseri ATCC 33323 = JCM 1131 GCA_000014425.1
679 Lactobacillus hominis DSM 23910 = CRBIP 24.179 GCA_000296835.1
523 Lactobacillus hamsteri DSM 5661 = JCM 6256 GCA_000615445.1
5e
462 Escherichia coli O157_3AH7 str. Sakai GCA_000008865.1
462 Escherichia coli str. K-12 substr. MG1655 GCA_000005845.2
462 Shigella flexneri 2a str. 301 GCA_000006925.2
461 Escherichia coli O104_3AH4 str. 2011C-3493 GCA_000299455.1
454 Escherichia coli IAI39 GCA_000026345.1
454 Escherichia coli UMN026 GCA_000026325.2
5f
626 Bifidobacterium adolescentis ATCC 15703 GCA_000010425.1
507 Bifidobacterium angulatum DSM 20098 = JCM 7096 GCA_001025155.1
485 Bifidobacterium callitrichos DSM 23973 GCA_000741175.1
5g
832 Deinococcus radiodurans R1 GCA_000008565.1
517 Deinococcus gobiensis I-0 GCA_000252445.1
497 Deinococcus deserti VCD115 GCA_000020685.1
5h
1023 Streptococcus mutans UA159 GCA_000007465.2
770 Streptococcus ratti FA-1 = DSM 20564 GCA_000286075.1
702 Streptococcus sobrinus DSM 20742 = ATCC 33478 GCA_000686605.1
5i
741 Clostridium beijerinckii GCA_000833105.2
711 Clostridium puniceum GCA_002006345.1
697 Clostridium saccharoperbutylacetonicum N1-4_28HMT_29 GCA_000340885.1
5j
413 Bifidobacterium adolescentis ATCC 15703 GCA_000010425.1
363 Bifidobacterium angulatum DSM 20098 = JCM 7096 GCA_001025155.1
346 Bifidobacterium breve DSM 20213 = JCM 1192 GCA_001025175.1
5k
645 Staphylococcus epidermidis ATCC 12228 GCA_000007645.1
619 Staphylococcus capitis subsp. capitis GCA_001028645.1
534 Staphylococcus haemolyticus JCSC1435 GCA_000009865.1
5l
783 Staphylococcus epidermidis ATCC 12228 GCA_000007645.1
752 Staphylococcus capitis subsp. capitis GCA_001028645.1
671 Staphylococcus aureus subsp. aureus NCTC 8325 GCA_000013425.1
671 Staphylococcus warneri SG1 GCA_000332735.1
5m
566 Enterococcus faecalis V583 GCA_000007785.1
539 Enterococcus rivorum GCA_001742285.1
523 Streptomyces cinnamoneus GCA_001885705.1
5n
740 Clostridium beijerinckii GCA_000833105.2
712 Clostridium saccharoperbutylacetonicum N1-4_28HMT_29 GCA_000340885.1
666 Clostridium saccharobutylicum DSM 13864 GCA_000473995.1

5o
737 Clostridium beijerinckii GCA_000833105.2
686 Clostridium saccharoperbutylacetonicum N1-4_28HMT_29 GCA_000340885.1
666 Clostridium puniceum GCA_002006345.1
5p
923 Lactobacillus gasseri ATCC 33323 = JCM 1131 GCA_000014425.1
852 Lactobacillus hominis DSM 23910 = CRBIP 24.179 GCA_000296835.1
656 Lactobacillus iners DSM 13335 GCA_000160875.1
5q
733 Staphylococcus epidermidis ATCC 12228 GCA_000007645.1
669 Staphylococcus capitis subsp. capitis GCA_001028645.1
626 Staphylococcus warneri SG1 GCA_000332735.1
5r
532 Rhodobacter sphaeroides 2.4.1 GCA_000012905.2
511 Rhodobacter sphaeroides ATCC 17025 GCA_000016405.1
416 Defluviimonas alba GCA_001620265.1
416 Thioclava indica GCA_000714545.1
5s
813 Streptococcus mutans UA159 GCA_000007465.2
613 Streptococcus ratti FA-1 = DSM 20564 GCA_000286075.1
580 Streptococcus merionis DSM 19192 GCA_000380085.1
5t
709 Clostridium beijerinckii GCA_000833105.2
660 Clostridium saccharoperbutylacetonicum N1-4_28HMT_29 GCA_000340885.1
643 Clostridium saccharobutylicum DSM 13864 GCA_000473995.1
5u
818 Streptococcus mutans UA159 GCA_000007465.2
621 Streptococcus ratti FA-1 = DSM 20564 GCA_000286075.1
589 Streptococcus gordonii str. Challis substr. CH1 GCA_000017005.1
5v
681 Deinococcus radiodurans R1 GCA_000008565.1
421 Deinococcus deserti VCD115 GCA_000020685.1
417 Deinococcus puniceus GCA_001644565.1
5w
578 Lactobacillus gasseri ATCC 33323 = JCM 1131 GCA_000014425.1
530 Lactobacillus hominis DSM 23910 = CRBIP 24.179 GCA_000296835.1
415 Lactobacillus jensenii GCA_001936235.1
415 Lactobacillus psittaci DSM 15354 GCA_000425905.1
5x
511 Staphylococcus epidermidis ATCC 12228 GCA_000007645.1
505 Streptomyces cinnamoneus GCA_001885705.1
480 Staphylococcus capitis subsp. capitis GCA_001028645.1
5y
746 Streptococcus mutans UA159 GCA_000007465.2
524 Streptococcus ferus DSM 20646 GCA_000372425.1
513 Streptococcus gordonii str. Challis substr. CH1 GCA_000017005.1
5z
664 Escherichia coli str. K-12 substr. MG1655 GCA_000005845.2
658 Escherichia coli IAI39 GCA_000026345.1
658 Escherichia coli O157_3AH7 str. Sakai GCA_000008865.1
658 Escherichia coli O83_3AH1 str. NRG 857C GCA_000183345.1
658 Escherichia coli UMN026 GCA_000026325.2
658 Shigella flexneri 2a str. 301 GCA_000006925.2
643 Shigella dysenteriae Sd197 GCA_000012005.1
5A
879 Bacillus anthracis str. Ames GCA_000007845.1
879 Bacillus anthracis str. Sterne GCA_000008165.1
879 _5BBacillus thuringiensis_5D serovar konkukian str. 97-27 GCA_000008505.1
853 Bacillus cereus ATCC 14579 GCA_000007825.1
842 Bacillus pseudomycoides DSM 12442 GCA_000161455.1
5B
787 Lactobacillus gasseri ATCC 33323 = JCM 1131 GCA_000014425.1
736 Lactobacillus hominis DSM 23910 = CRBIP 24.179 GCA_000296835.1
578 Lactobacillus iners DSM 13335 GCA_000160875.1
5C
1005 _5BBacillus thuringiensis_5D serovar konkukian str. 97-27 GCA_000008505.1
995 Bacillus anthracis str. Ames GCA_000007845.1
995 Bacillus anthracis str. Sterne GCA_000008165.1
983 Bacillus cereus ATCC 14579 GCA_000007825.1
5D
735 Lactobacillus gasseri ATCC 33323 = JCM 1131 GCA_000014425.1
702 Lactobacillus hominis DSM 23910 = CRBIP 24.179 GCA_000296835.1
512 Lactobacillus iners DSM 13335 GCA_000160875.1
5E
426 Streptococcus mutans UA159 GCA_000007465.2
320 Streptococcus ratti FA-1 = DSM 20564 GCA_000286075.1

309 Streptococcus sobrinus DSM 20742 = ATCC 33478 GCA_000686605.1
5F
771 Enterococcus faecalis V583 GCA_000007785.1
724 Streptomyces cinnamoneus GCA_001885705.1
661 Enterococcus faecium DO GCA_000174395.2
5G
694 Clostridium beijerinckii GCA_000833105.2
680 Clostridium saccharoperbutylacetonicum N1-4_28HMT_29 GCA_000340885.1
657 Clostridium puniceum GCA_002006345.1
5H
664 Rhodobacter sphaeroides 2.4.1 GCA_000012905.2
618 Rhodobacter sphaeroides ATCC 17025 GCA_000016405.1
469 Pseudorhodobacter ferrugineus DSM 5888 GCA_000420745.1
5I
599 Rhodobacter sphaeroides 2.4.1 GCA_000012905.2
516 Rhodobacter sphaeroides ATCC 17025 GCA_000016405.1
475 Gemmobacter aquatilis GCA_900110025.1
5J
537 Deinococcus radiodurans R1 GCA_000008565.1
384 Deinococcus deserti VCD115 GCA_000020685.1
372 Deinococcus soli Cha et al. 2016 GCA_001007995.1
5K
681 Clostridium beijerinckii GCA_000833105.2
652 Clostridium saccharoperbutylacetonicum N1-4_28HMT_29 GCA_000340885.1
620 Clostridium puniceum GCA_002006345.1
5L
903 Clostridium beijerinckii GCA_000833105.2
873 Clostridium saccharoperbutylacetonicum N1-4_28HMT_29 GCA_000340885.1
838 Clostridium puniceum GCA_002006345.1
5M
763 Bifidobacterium adolescentis ATCC 15703 GCA_000010425.1
634 Bifidobacterium callitrichos DSM 23973 GCA_000741175.1
618 Bifidobacterium asteroides PRL2011 GCA_000304215.1
5N
624 Bifidobacterium adolescentis ATCC 15703 GCA_000010425.1
537 Bifidobacterium dentium JCM 1195 = DSM 20436 GCA_001042595.1
519 Bifidobacterium longum NCC2705 GCA_000007525.1
5O
840 Streptococcus mutans UA159 GCA_000007465.2
697 Streptococcus ratti FA-1 = DSM 20564 GCA_000286075.1
651 Streptococcus sobrinus DSM 20742 = ATCC 33478 GCA_000686605.1
5P
473 Rhodobacter sphaeroides 2.4.1 GCA_000012905.2
420 Rhodobacter sphaeroides ATCC 17025 GCA_000016405.1
379 Gemmobacter aquatilis GCA_900110025.1
5Q
845 Rhodobacter sphaeroides 2.4.1 GCA_000012905.2
764 Rhodobacter sphaeroides ATCC 17025 GCA_000016405.1
635 Gemmobacter aquatilis GCA_900110025.1
5R
587 Enterococcus faecalis V583 GCA_000007785.1
546 Streptomyces cinnamoneus GCA_001885705.1
521 Enterococcus saccharolyticus subsp. saccharolyticus ATCC 43076 GCA_000407285.1
5S
853 Streptococcus mutans UA159 GCA_000007465.2
631 Streptococcus ratti FA-1 = DSM 20564 GCA_000286075.1
600 Streptococcus gordonii str. Challis substr. CH1 GCA_000017005.1
5T
601 Deinococcus radiodurans R1 GCA_000008565.1
431 Deinococcus deserti VCD115 GCA_000020685.1
404 Deinococcus gobiensis I-0 GCA_000252445.1
5U
659 Lactobacillus gasseri ATCC 33323 = JCM 1131 GCA_000014425.1
622 Lactobacillus hominis DSM 23910 = CRBIP 24.179 GCA_000296835.1
474 Lactobacillus psittaci DSM 15354 GCA_000425905.1
5V
806 Bacillus anthracis str. Ames GCA_000007845.1
806 Bacillus anthracis str. Sterne GCA_000008165.1
806 _5BBacillus thuringiensis_5D serovar konkukian str. 97-27 GCA_000008505.1
793 Bacillus cereus ATCC 14579 GCA_000007825.1
793 Bacillus mycoides GCA_000832605.1
781 Bacillus thuringiensis YBT-1518 GCA_000497525.2
5W
746 Deinococcus radiodurans R1 GCA_000008565.1
448 Deinococcus deserti VCD115 GCA_000020685.1
445 Deinococcus hopiensis KR-140 GCA_900176165.1

5X
784 Rhodobacter sphaeroides 2.4.1 GCA_000012905.2
721 Rhodobacter sphaeroides ATCC 17025 GCA_000016405.1
586 Gemmobacter megaterium GCA_900156815.1
5Y
468 Deinococcus radiodurans R1 GCA_000008565.1
348 Deinococcus deserti VCD115 GCA_000020685.1
315 Deinococcus soli Cha et al. 2016 GCA_001007995.1
5Z
772 Lactobacillus gasseri ATCC 33323 = JCM 1131 GCA_000014425.1
732 Lactobacillus hominis DSM 23910 = CRBIP 24.179 GCA_000296835.1
558 Lactobacillus iners DSM 13335 GCA_000160875.1
60
458 Rhodobacter sphaeroides 2.4.1 GCA_000012905.2
398 Rhodobacter sphaeroides ATCC 17025 GCA_000016405.1
338 Defluviimonas alba GCA_001620265.1
61
630 Rhodobacter sphaeroides 2.4.1 GCA_000012905.2
592 Rhodobacter sphaeroides ATCC 17025 GCA_000016405.1
463 Pseudorhodobacter psychrotolerans GCA_001294535.1
62
579 Deinococcus radiodurans R1 GCA_000008565.1
419 Deinococcus deserti VCD115 GCA_000020685.1
407 Deinococcus hopiensis KR-140 GCA_900176165.1
407 Deinococcus marmoris DSM 12784 GCA_000701405.1
63
798 Lactobacillus gasseri ATCC 33323 = JCM 1131 GCA_000014425.1
718 Lactobacillus hominis DSM 23910 = CRBIP 24.179 GCA_000296835.1
530 Lactobacillus iners DSM 13335 GCA_000160875.1
64
396 Deinococcus radiodurans R1 GCA_000008565.1
228 Deinococcus deserti VCD115 GCA_000020685.1
195 Deinococcus marmoris DSM 12784 GCA_000701405.1
65
741 Escherichia coli IAI39 GCA_000026345.1
741 Escherichia coli O157_3AH7 str. Sakai GCA_000008865.1
741 Escherichia coli str. K-12 substr. MG1655 GCA_000005845.2
740 Escherichia coli UMN026 GCA_000026325.2
729 Escherichia coli O83_3AH1 str. NRG 857C GCA_000183345.1
66
721 Deinococcus radiodurans R1 GCA_000008565.1
459 Deinococcus gobiensis I-0 GCA_000252445.1
448 Deinococcus hopiensis KR-140 GCA_900176165.1
67
739 Staphylococcus epidermidis ATCC 12228 GCA_000007645.1
693 Staphylococcus capitis subsp. capitis GCA_001028645.1
628 Staphylococcus haemolyticus JCSC1435 GCA_000009865.1
68
875 Staphylococcus epidermidis ATCC 12228 GCA_000007645.1
828 Staphylococcus capitis subsp. capitis GCA_001028645.1
771 Staphylococcus warneri SG1 GCA_000332735.1
69
730 Deinococcus radiodurans R1 GCA_000008565.1
449 Deinococcus deserti VCD115 GCA_000020685.1
422 Deinococcus gobiensis I-0 GCA_000252445.1
422 Deinococcus soli Cha et al. 2016 GCA_001007995.1
6a
878 Staphylococcus epidermidis ATCC 12228 GCA_000007645.1
805 Staphylococcus capitis subsp. capitis GCA_001028645.1
762 Staphylococcus aureus subsp. aureus NCTC 8325 GCA_000013425.1
6b
679 Bifidobacterium adolescentis ATCC 15703 GCA_000010425.1
567 Bifidobacterium dentium JCM 1195 = DSM 20436 GCA_001042595.1
542 Bifidobacterium angulatum DSM 20098 = JCM 7096 GCA_001025155.1
6c
692 Rhodobacter sphaeroides 2.4.1 GCA_000012905.2
596 Rhodobacter sphaeroides ATCC 17025 GCA_000016405.1
523 Gemmobacter aquatilis GCA_900110025.1
6d
624 Lactobacillus gasseri ATCC 33323 = JCM 1131 GCA_000014425.1
567 Lactobacillus hominis DSM 23910 = CRBIP 24.179 GCA_000296835.1
434 Lactobacillus iners DSM 13335 GCA_000160875.1
6e
537 Rhodobacter sphaeroides 2.4.1 GCA_000012905.2
494 Rhodobacter sphaeroides ATCC 17025 GCA_000016405.1
454 Defluviimonas alba GCA_001620265.1

6f
862 Bifidobacterium adolescentis ATCC 15703 GCA_000010425.1
747 Bifidobacterium dentium JCM 1195 = DSM 20436 GCA_001042595.1
713 Bifidobacterium stellenboschense GCA_000741785.1
6g
586 Bifidobacterium adolescentis ATCC 15703 GCA_000010425.1
508 Bifidobacterium dentium JCM 1195 = DSM 20436 GCA_001042595.1
495 Bifidobacterium angulatum DSM 20098 = JCM 7096 GCA_001025155.1
6h
775 Deinococcus radiodurans R1 GCA_000008565.1
475 Deinococcus deserti VCD115 GCA_000020685.1
423 Deinococcus soli Cha et al. 2016 GCA_001007995.1
6i
801 Streptococcus mutans UA159 GCA_000007465.2
547 Streptococcus ratti FA-1 = DSM 20564 GCA_000286075.1
487 Streptococcus criceti HS-6 GCA_000187975.3
6j
526 Enterococcus faecalis V583 GCA_000007785.1
479 Streptomyces cinnamoneus GCA_001885705.1
472 Enterococcus canis NBRC 100695 GCA_001544375.1
6k
558 Streptococcus mutans UA159 GCA_000007465.2
370 Streptococcus ratti FA-1 = DSM 20564 GCA_000286075.1
348 Streptococcus macacae NCTC 11558 GCA_000187995.3
6l
453 Clostridium beijerinckii GCA_000833105.2
440 Clostridium saccharoperbutylacetonicum N1-4_28HMT_29 GCA_000340885.1
439 Clostridium chromiireducens GCA_002029255.1
6m
755 Rhodobacter sphaeroides 2.4.1 GCA_000012905.2
674 Rhodobacter sphaeroides ATCC 17025 GCA_000016405.1
573 Gemmobacter megaterium GCA_900156815.1
6n
386 Rhodobacter sphaeroides 2.4.1 GCA_000012905.2
321 Rhodobacter sphaeroides ATCC 17025 GCA_000016405.1
239 Rhodobacter capsulatus SB 1003 GCA_000021865.1
6o
646 Staphylococcus epidermidis ATCC 12228 GCA_000007645.1
609 Staphylococcus capitis subsp. capitis GCA_001028645.1
578 Staphylococcus aureus subsp. aureus NCTC 8325 GCA_000013425.1
6p
555 Lactobacillus gasseri ATCC 33323 = JCM 1131 GCA_000014425.1
555 Lactobacillus hominis DSM 23910 = CRBIP 24.179 GCA_000296835.1
436 Lactobacillus iners DSM 13335 GCA_000160875.1
383 Lactobacillus hamsteri DSM 5661 = JCM 6256 GCA_000615445.1
6q
506 Enterococcus faecalis V583 GCA_000007785.1
462 Streptomyces cinnamoneus GCA_001885705.1
404 Enterococcus asini ATCC 700915 GCA_000407365.1
6r
633 Clostridium beijerinckii GCA_000833105.2
604 Clostridium saccharobutylicum DSM 13864 GCA_000473995.1
601 Clostridium saccharoperbutylacetonicum N1-4_28HMT_29 GCA_000340885.1
6s
916 Clostridium beijerinckii GCA_000833105.2
913 Clostridium saccharoperbutylacetonicum N1-4_28HMT_29 GCA_000340885.1
880 Clostridium puniceum GCA_002006345.1
880 Clostridium saccharobutylicum DSM 13864 GCA_000473995.1
6t
326 Staphylococcus epidermidis ATCC 12228 GCA_000007645.1
311 Staphylococcus capitis subsp. capitis GCA_001028645.1
279 Staphylococcus haemolyticus JCSC1435 GCA_000009865.1
279 Staphylococcus lugdunensis HKU09-01 GCA_000025085.1
6u
673 Clostridium beijerinckii GCA_000833105.2
655 Clostridium saccharoperbutylacetonicum N1-4_28HMT_29 GCA_000340885.1
613 Clostridium saccharobutylicum DSM 13864 GCA_000473995.1
6v
523 Clostridium beijerinckii GCA_000833105.2
476 Clostridium saccharoperbutylacetonicum N1-4_28HMT_29 GCA_000340885.1
467 Clostridium puniceum GCA_002006345.1
6w
620 Bifidobacterium adolescentis ATCC 15703 GCA_000010425.1
515 Bifidobacterium angulatum DSM 20098 = JCM 7096 GCA_001025155.1
489 Bifidobacterium dentium JCM 1195 = DSM 20436 GCA_001042595.1
6x

730 Enterococcus faecalis V583 GCA_000007785.1
683 Streptomyces cinnamoneus GCA_001885705.1
639 Enterococcus rivorum GCA_001742285.1
6y
505 Rhodobacter sphaeroides 2.4.1 GCA_000012905.2
421 Rhodobacter sphaeroides ATCC 17025 GCA_000016405.1
418 Gemmobacter aquatilis GCA_900110025.1
6z
776 Staphylococcus epidermidis ATCC 12228 GCA_000007645.1
766 Staphylococcus capitis subsp. capitis GCA_001028645.1
708 Staphylococcus aureus subsp. aureus NCTC 8325 GCA_000013425.1
6A
883 Staphylococcus epidermidis ATCC 12228 GCA_000007645.1
827 Staphylococcus capitis subsp. capitis GCA_001028645.1
781 Staphylococcus haemolyticus JCSC1435 GCA_000009865.1
6B
959 Enterococcus faecalis V583 GCA_000007785.1
912 Streptomyces cinnamoneus GCA_001885705.1
884 Enterococcus rivorum GCA_001742285.1
6C
834 Clostridium beijerinckii GCA_000833105.2
825 Clostridium saccharoperbutylacetonicum N1-4_28HMT_29 GCA_000340885.1
793 Clostridium puniceum GCA_002006345.1
6D
559 Enterococcus faecalis V583 GCA_000007785.1
512 Streptomyces cinnamoneus GCA_001885705.1
431 Enterococcus rivorum GCA_001742285.1
6E
608 Staphylococcus epidermidis ATCC 12228 GCA_000007645.1
573 Staphylococcus capitis subsp. capitis GCA_001028645.1
540 Staphylococcus haemolyticus JCSC1435 GCA_000009865.1
6F
864 Escherichia coli O104_3AH4 str. 2011C-3493 GCA_000299455.1
864 Escherichia coli str. K-12 substr. MG1655 GCA_000005845.2
849 Shigella flexneri 2a str. 301 GCA_000006925.2
810 Escherichia coli O157_3AH7 str. Sakai GCA_000008865.1
6G
881 Bacillus anthracis str. Ames GCA_000007845.1
881 Bacillus anthracis str. Sterne GCA_000008165.1
881 Bacillus pseudomycoides DSM 12442 GCA_000161455.1
881 _5BBacillus thuringiensis_5D serovar konkukian str. 97-27 GCA_000008505.1
873 Bacillus cereus ATCC 14579 GCA_000007825.1
854 Bacillus thuringiensis YBT-1518 GCA_000497525.2
6H
701 Rhodobacter sphaeroides 2.4.1 GCA_000012905.2
607 Rhodobacter sphaeroides ATCC 17025 GCA_000016405.1
459 Gemmobacter aquatilis GCA_900110025.1
6I
802 Clostridium beijerinckii GCA_000833105.2
779 Clostridium saccharoperbutylacetonicum N1-4_28HMT_29 GCA_000340885.1
758 Clostridium puniceum GCA_002006345.1
6J
602 Clostridium beijerinckii GCA_000833105.2
602 Clostridium saccharoperbutylacetonicum N1-4_28HMT_29 GCA_000340885.1
569 Clostridium puniceum GCA_002006345.1
562 Clostridium butyricum GCA_001456065.2
6K
828 Bifidobacterium adolescentis ATCC 15703 GCA_000010425.1
719 Bifidobacterium dentium JCM 1195 = DSM 20436 GCA_001042595.1
625 Bifidobacterium asteroides PRL2011 GCA_000304215.1
6L
761 Streptococcus mutans UA159 GCA_000007465.2
544 Streptococcus gordonii str. Challis substr. CH1 GCA_000017005.1
536 Streptococcus ratti FA-1 = DSM 20564 GCA_000286075.1
6M
879 Enterococcus faecalis V583 GCA_000007785.1
832 Streptomyces cinnamoneus GCA_001885705.1
742 Enterococcus asini ATCC 700915 GCA_000407365.1
6N
747 Lactobacillus gasseri ATCC 33323 = JCM 1131 GCA_000014425.1
694 Lactobacillus hominis DSM 23910 = CRBIP 24.179 GCA_000296835.1
505 Lactobacillus jensenii GCA_001936235.1
505 Lactobacillus psittaci DSM 15354 GCA_000425905.1
6O
714 Clostridium beijerinckii GCA_000833105.2
690 Clostridium saccharobutylicum DSM 13864 GCA_000473995.1

680 Clostridium saccharoperbutylacetonicum N1-4_28HMT_29 GCA_000340885.1
6P
956 Lactobacillus gasseri ATCC 33323 = JCM 1131 GCA_000014425.1
878 Lactobacillus hominis DSM 23910 = CRBIP 24.179 GCA_000296835.1
642 Lactobacillus iners DSM 13335 GCA_000160875.1
6Q
684 Rhodobacter sphaeroides 2.4.1 GCA_000012905.2
640 Rhodobacter sphaeroides ATCC 17025 GCA_000016405.1
597 Gemmobacter aquatilis GCA_900110025.1
6R
599 Lactobacillus gasseri ATCC 33323 = JCM 1131 GCA_000014425.1
556 Lactobacillus hominis DSM 23910 = CRBIP 24.179 GCA_000296835.1
409 Lactobacillus iners DSM 13335 GCA_000160875.1
6S
523 Clostridium beijerinckii GCA_000833105.2
508 Clostridium puniceum GCA_002006345.1
508 Clostridium saccharobutylicum DSM 13864 GCA_000473995.1
508 Clostridium saccharoperbutylacetonicum N1-4_28HMT_29 GCA_000340885.1
484 Clostridium butyricum GCA_001456065.2
6T
849 Rhodobacter sphaeroides 2.4.1 GCA_000012905.2
723 Rhodobacter sphaeroides ATCC 17025 GCA_000016405.1
631 Defluviimonas alba GCA_001620265.1
6U
817 Escherichia coli str. K-12 substr. MG1655 GCA_000005845.2
797 Shigella flexneri 2a str. 301 GCA_000006925.2
790 Escherichia coli UMN026 GCA_000026325.2
6V
808 Bacillus anthracis str. Ames GCA_000007845.1
808 Bacillus anthracis str. Sterne GCA_000008165.1
808 _5BBacillus thuringiensis_5D serovar konkukian str. 97-27 GCA_000008505.1
796 Bacillus cereus ATCC 14579 GCA_000007825.1
783 Bacillus pseudomycoides DSM 12442 GCA_000161455.1
6W
450 Bifidobacterium adolescentis ATCC 15703 GCA_000010425.1
351 Bifidobacterium callitrichos DSM 23973 GCA_000741175.1
346 Bifidobacterium longum NCC2705 GCA_000007525.1
6X
682 Streptococcus mutans UA159 GCA_000007465.2
525 Streptococcus ratti FA-1 = DSM 20564 GCA_000286075.1
497 Streptococcus halotolerans GCA_001598035.1
6Y
568 Enterococcus faecalis V583 GCA_000007785.1
525 Streptomyces cinnamoneus GCA_001885705.1
451 Enterococcus massiliensis GCA_001050095.1
6Z
642 Streptococcus mutans UA159 GCA_000007465.2
456 Streptococcus ratti FA-1 = DSM 20564 GCA_000286075.1
442 Streptococcus gordonii str. Challis substr. CH1 GCA_000017005.1
70
602 Shigella flexneri 2a str. 301 GCA_000006925.2
599 Escherichia coli O104_3AH4 str. 2011C-3493 GCA_000299455.1
599 Escherichia coli O157_3AH7 str. Sakai GCA_000008865.1
599 Escherichia coli str. K-12 substr. MG1655 GCA_000005845.2
590 Escherichia coli O83_3AH1 str. NRG 857C GCA_000183345.1
590 Escherichia coli UMN026 GCA_000026325.2
71
752 Enterococcus faecalis V583 GCA_000007785.1
732 Enterococcus faecium DO GCA_000174395.2
709 Enterococcus canis NBRC 100695 GCA_001544375.1
72
681 Lactobacillus gasseri ATCC 33323 = JCM 1131 GCA_000014425.1
616 Lactobacillus hominis DSM 23910 = CRBIP 24.179 GCA_000296835.1
486 Lactobacillus iners DSM 13335 GCA_000160875.1
73
688 Enterococcus faecalis V583 GCA_000007785.1
647 Streptomyces cinnamoneus GCA_001885705.1
618 Enterococcus faecium DO GCA_000174395.2
74
380 Bifidobacterium adolescentis ATCC 15703 GCA_000010425.1
330 Bifidobacterium dentium JCM 1195 = DSM 20436 GCA_001042595.1
304 Bifidobacterium tsurumiense GCA_000741765.1
75
636 Streptococcus mutans UA159 GCA_000007465.2
469 Streptococcus equinus GCA_000964315.1
467 Streptococcus ratti FA-1 = DSM 20564 GCA_000286075.1

76
673 Staphylococcus epidermidis ATCC 12228 GCA_000007645.1
662 Staphylococcus warneri SG1 GCA_000332735.1
660 Staphylococcus capitis subsp. capitis GCA_001028645.1
77
878 Enterococcus faecalis V583 GCA_000007785.1
832 Streptomyces cinnamoneus GCA_001885705.1
770 Enterococcus canis NBRC 100695 GCA_001544375.1
78
746 Streptococcus mutans UA159 GCA_000007465.2
560 Streptococcus ratti FA-1 = DSM 20564 GCA_000286075.1
501 Streptococcus gordonii str. Challis substr. CH1 GCA_000017005.1
79
816 Clostridium beijerinckii GCA_000833105.2
760 Clostridium saccharoperbutylacetonicum N1-4_28HMT_29 GCA_000340885.1
739 Clostridium butyricum GCA_001456065.2
7a
603 Rhodobacter sphaeroides 2.4.1 GCA_000012905.2
566 Rhodobacter sphaeroides ATCC 17025 GCA_000016405.1
474 Gemmobacter aquatilis GCA_900110025.1
7b
783 Staphylococcus epidermidis ATCC 12228 GCA_000007645.1
748 Staphylococcus haemolyticus JCSC1435 GCA_000009865.1
731 Staphylococcus capitis subsp. capitis GCA_001028645.1
7c
689 Deinococcus radiodurans R1 GCA_000008565.1
383 Deinococcus deserti VCD115 GCA_000020685.1
362 Deinococcus gobiensis I-0 GCA_000252445.1
7d
577 Streptococcus mutans UA159 GCA_000007465.2
450 Streptococcus ratti FA-1 = DSM 20564 GCA_000286075.1
411 Streptococcus ferus DSM 20646 GCA_000372425.1
7e
759 Staphylococcus epidermidis ATCC 12228 GCA_000007645.1
728 Staphylococcus capitis subsp. capitis GCA_001028645.1
667 Staphylococcus warneri SG1 GCA_000332735.1
7f
731 Lactobacillus gasseri ATCC 33323 = JCM 1131 GCA_000014425.1
652 Lactobacillus hominis DSM 23910 = CRBIP 24.179 GCA_000296835.1
500 Lactobacillus jensenii GCA_001936235.1
7g
675 Deinococcus radiodurans R1 GCA_000008565.1
458 Deinococcus gobiensis I-0 GCA_000252445.1
420 Deinococcus deserti VCD115 GCA_000020685.1
7h
716 Deinococcus radiodurans R1 GCA_000008565.1
491 Deinococcus deserti VCD115 GCA_000020685.1
475 Deinococcus hopiensis KR-140 GCA_900176165.1
7i
422 Rhodobacter sphaeroides 2.4.1 GCA_000012905.2
392 Rhodobacter sphaeroides ATCC 17025 GCA_000016405.1
287 Pseudorhodobacter wandonensis GCA_001202035.1
7j
805 Lactobacillus gasseri ATCC 33323 = JCM 1131 GCA_000014425.1
743 Lactobacillus hominis DSM 23910 = CRBIP 24.179 GCA_000296835.1
543 Lactobacillus iners DSM 13335 GCA_000160875.1
7k
739 Bacillus cereus ATCC 14579 GCA_000007825.1
724 Bacillus anthracis str. Ames GCA_000007845.1
724 Bacillus anthracis str. Sterne GCA_000008165.1
724 _5BBacillus thuringiensis_5D serovar konkukian str. 97-27 GCA_000008505.1
719 Bacillus thuringiensis YBT-1518 GCA_000497525.2
7l
618 Enterococcus faecalis V583 GCA_000007785.1
586 Streptomyces cinnamoneus GCA_001885705.1
518 Enterococcus rivorum GCA_001742285.1
7m
763 Rhodobacter sphaeroides 2.4.1 GCA_000012905.2
681 Rhodobacter sphaeroides ATCC 17025 GCA_000016405.1
552 Pseudorhodobacter psychrotolerans GCA_001294535.1
7n
783 Rhodobacter sphaeroides 2.4.1 GCA_000012905.2
733 Rhodobacter sphaeroides ATCC 17025 GCA_000016405.1
588 Pseudorhodobacter ferrugineus DSM 5888 GCA_000420745.1
7o
541 Enterococcus faecalis V583 GCA_000007785.1

494 Streptomyces cinnamoneus GCA_001885705.1
454 Enterococcus canis NBRC 100695 GCA_001544375.1
7p
460 Streptococcus mutans UA159 GCA_000007465.2
340 Streptococcus equinus GCA_000964315.1
338 Streptococcus sobrinus DSM 20742 = ATCC 33478 GCA_000686605.1
7q
843 Staphylococcus epidermidis ATCC 12228 GCA_000007645.1
789 Staphylococcus capitis subsp. capitis GCA_001028645.1
708 Staphylococcus hominis subsp. hominis C80 GCA_000183685.1
7r
939 Lactobacillus gasseri ATCC 33323 = JCM 1131 GCA_000014425.1
823 Lactobacillus hominis DSM 23910 = CRBIP 24.179 GCA_000296835.1
615 Lactobacillus iners DSM 13335 GCA_000160875.1
7s
657 Escherichia coli str. K-12 substr. MG1655 GCA_000005845.2
654 Escherichia coli O157_3AH7 str. Sakai GCA_000008865.1
654 Shigella flexneri 2a str. 301 GCA_000006925.2
646 Escherichia coli UMN026 GCA_000026325.2
7t
878 Staphylococcus epidermidis ATCC 12228 GCA_000007645.1
835 Staphylococcus capitis subsp. capitis GCA_001028645.1
743 Staphylococcus hominis subsp. hominis C80 GCA_000183685.1
743 Staphylococcus warneri SG1 GCA_000332735.1
7u
798 Clostridium beijerinckii GCA_000833105.2
770 Clostridium saccharoperbutylacetonicum N1-4_28HMT_29 GCA_000340885.1
749 Clostridium puniceum GCA_002006345.1
7v
641 Lactobacillus gasseri ATCC 33323 = JCM 1131 GCA_000014425.1
612 Lactobacillus hominis DSM 23910 = CRBIP 24.179 GCA_000296835.1
454 Lactobacillus psittaci DSM 15354 GCA_000425905.1
7w
837 Bacillus anthracis str. Ames GCA_000007845.1
837 Bacillus anthracis str. Sterne GCA_000008165.1
837 _5BBacillus thuringiensis_5D serovar konkukian str. 97-27 GCA_000008505.1
822 Bacillus cereus ATCC 14579 GCA_000007825.1
802 Bacillus thuringiensis YBT-1518 GCA_000497525.2
7x
638 Deinococcus radiodurans R1 GCA_000008565.1
430 Deinococcus deserti VCD115 GCA_000020685.1
430 Deinococcus hopiensis KR-140 GCA_900176165.1
402 Deinococcus murrayi DSM 11303 GCA_000482805.1
7y
651 Deinococcus radiodurans R1 GCA_000008565.1
406 Deinococcus deserti VCD115 GCA_000020685.1
390 Deinococcus puniceus GCA_001644565.1
7z
857 Lactobacillus gasseri ATCC 33323 = JCM 1131 GCA_000014425.1
763 Lactobacillus hominis DSM 23910 = CRBIP 24.179 GCA_000296835.1
593 Lactobacillus iners DSM 13335 GCA_000160875.1
7A
519 Escherichia coli O104_3AH4 str. 2011C-3493 GCA_000299455.1
514 Escherichia coli str. K-12 substr. MG1655 GCA_000005845.2
514 Shigella flexneri 2a str. 301 GCA_000006925.2
488 Escherichia coli O157_3AH7 str. Sakai GCA_000008865.1
7B
668 Rhodobacter sphaeroides 2.4.1 GCA_000012905.2
637 Rhodobacter sphaeroides ATCC 17025 GCA_000016405.1
562 Pseudorhodobacter psychrotolerans GCA_001294535.1
7C
609 Staphylococcus epidermidis ATCC 12228 GCA_000007645.1
565 Staphylococcus capitis subsp. capitis GCA_001028645.1
513 Staphylococcus warneri SG1 GCA_000332735.1
7D
844 Deinococcus radiodurans R1 GCA_000008565.1
591 Deinococcus deserti VCD115 GCA_000020685.1
560 Deinococcus proteolyticus MRP GCA_000190555.1
7E
836 Staphylococcus epidermidis ATCC 12228 GCA_000007645.1
791 Staphylococcus capitis subsp. capitis GCA_001028645.1
736 Megasphaera cerevisiae DSM 20462 GCA_001045675.1
736 Staphylococcus warneri SG1 GCA_000332735.1
7F
539 Bifidobacterium adolescentis ATCC 15703 GCA_000010425.1
432 Bifidobacterium dentium JCM 1195 = DSM 20436 GCA_001042595.1

360 Bifidobacterium asteroides PRL2011 GCA_000304215.1
7G
713 Escherichia coli str. K-12 substr. MG1655 GCA_000005845.2
704 Escherichia coli O104_3AH4 str. 2011C-3493 GCA_000299455.1
688 Shigella flexneri 2a str. 301 GCA_000006925.2
7H
707 Deinococcus radiodurans R1 GCA_000008565.1
456 Deinococcus deserti VCD115 GCA_000020685.1
440 Deinococcus marmoris DSM 12784 GCA_000701405.1
7I
678 Clostridium beijerinckii GCA_000833105.2
642 Clostridium saccharoperbutylacetonicum N1-4_28HMT_29 GCA_000340885.1
616 Clostridium saccharobutylicum DSM 13864 GCA_000473995.1
7J
480 Bifidobacterium adolescentis ATCC 15703 GCA_000010425.1
403 Bifidobacterium dentium JCM 1195 = DSM 20436 GCA_001042595.1
388 Bifidobacterium breve DSM 20213 = JCM 1192 GCA_001025175.1
7K
721 Streptococcus mutans UA159 GCA_000007465.2
490 Streptococcus ratti FA-1 = DSM 20564 GCA_000286075.1
481 Streptococcus criceti HS-6 GCA_000187975.3
7L
657 Lactobacillus gasseri ATCC 33323 = JCM 1131 GCA_000014425.1
609 Lactobacillus hominis DSM 23910 = CRBIP 24.179 GCA_000296835.1
494 Lactobacillus psittaci DSM 15354 GCA_000425905.1
7M
685 Staphylococcus epidermidis ATCC 12228 GCA_000007645.1
673 Staphylococcus capitis subsp. capitis GCA_001028645.1
602 Staphylococcus warneri SG1 GCA_000332735.1
7N
618 Escherichia coli IAI39 GCA_000026345.1
618 Escherichia coli O157_3AH7 str. Sakai GCA_000008865.1
618 Escherichia coli str. K-12 substr. MG1655 GCA_000005845.2
609 Escherichia coli UMN026 GCA_000026325.2
606 Escherichia coli O83_3AH1 str. NRG 857C GCA_000183345.1
7O
738 Staphylococcus epidermidis ATCC 12228 GCA_000007645.1
707 Staphylococcus capitis subsp. capitis GCA_001028645.1
664 Staphylococcus hominis subsp. hominis C80 GCA_000183685.1
7P
876 Streptococcus mutans UA159 GCA_000007465.2
612 Streptococcus gordonii str. Challis substr. CH1 GCA_000017005.1
606 Streptococcus ratti FA-1 = DSM 20564 GCA_000286075.1
7Q
344 Rhodobacter sphaeroides 2.4.1 GCA_000012905.2
292 Rhodobacter sphaeroides ATCC 17025 GCA_000016405.1
272 Defluviimonas alba GCA_001620265.1
7R
483 Staphylococcus epidermidis ATCC 12228 GCA_000007645.1
440 Staphylococcus capitis subsp. capitis GCA_001028645.1
401 Megasphaera cerevisiae DSM 20462 GCA_001045675.1
401 Staphylococcus warneri SG1 GCA_000332735.1
7S
705 Deinococcus radiodurans R1 GCA_000008565.1
395 Deinococcus deserti VCD115 GCA_000020685.1
368 Deinococcus proteolyticus MRP GCA_000190555.1
7T
629 Bacillus thuringiensis YBT-1518 GCA_000497525.2
628 Bacillus anthracis str. Ames GCA_000007845.1
628 Bacillus anthracis str. Sterne GCA_000008165.1
628 Bacillus cereus ATCC 14579 GCA_000007825.1
628 Bacillus pseudomycoides DSM 12442 GCA_000161455.1
628 _5BBacillus thuringiensis_5D serovar konkukian str. 97-27 GCA_000008505.1
618 Bacillus mycoides GCA_000832605.1
7U
824 Staphylococcus epidermidis ATCC 12228 GCA_000007645.1
798 Staphylococcus capitis subsp. capitis GCA_001028645.1
724 Staphylococcus hominis subsp. hominis C80 GCA_000183685.1
7V
756 Staphylococcus epidermidis ATCC 12228 GCA_000007645.1
729 Staphylococcus capitis subsp. capitis GCA_001028645.1
696 Staphylococcus aureus subsp. aureus NCTC 8325 GCA_000013425.1
7W
581 Bifidobacterium adolescentis ATCC 15703 GCA_000010425.1
454 Bifidobacterium dentium JCM 1195 = DSM 20436 GCA_001042595.1
453 Bifidobacterium longum NCC2705 GCA_000007525.1

7X
593 Lactobacillus gasseri ATCC 33323 = JCM 1131 GCA_000014425.1
523 Lactobacillus hominis DSM 23910 = CRBIP 24.179 GCA_000296835.1
379 Lactobacillus jensenii GCA_001936235.1
379 Lactobacillus psittaci DSM 15354 GCA_000425905.1
7Y
782 Escherichia coli IAI39 GCA_000026345.1
782 Escherichia coli O157_3AH7 str. Sakai GCA_000008865.1
782 Escherichia coli str. K-12 substr. MG1655 GCA_000005845.2
771 Escherichia coli O83_3AH1 str. NRG 857C GCA_000183345.1
759 Shigella flexneri 2a str. 301 GCA_000006925.2
7Z
902 Streptococcus mutans UA159 GCA_000007465.2
657 Streptococcus ratti FA-1 = DSM 20564 GCA_000286075.1
649 Streptococcus gordonii str. Challis substr. CH1 GCA_000017005.1
80
777 Clostridium beijerinckii GCA_000833105.2
752 Clostridium saccharoperbutylacetonicum N1-4_28HMT_29 GCA_000340885.1
721 Clostridium saccharobutylicum DSM 13864 GCA_000473995.1
81
501 Staphylococcus epidermidis ATCC 12228 GCA_000007645.1
451 Staphylococcus capitis subsp. capitis GCA_001028645.1
440 Staphylococcus aureus subsp. aureus NCTC 8325 GCA_000013425.1
82
949 Rhodobacter sphaeroides 2.4.1 GCA_000012905.2
866 Rhodobacter sphaeroides ATCC 17025 GCA_000016405.1
711 Pseudorhodobacter psychrotolerans GCA_001294535.1
83
811 Deinococcus radiodurans R1 GCA_000008565.1
494 Deinococcus deserti VCD115 GCA_000020685.1
492 Deinococcus gobiensis I-0 GCA_000252445.1
84
524 Clostridium beijerinckii GCA_000833105.2
495 Clostridium saccharobutylicum DSM 13864 GCA_000473995.1
485 Clostridium puniceum GCA_002006345.1
485 Clostridium saccharoperbutylacetonicum N1-4_28HMT_29 GCA_000340885.1
85
660 Bifidobacterium adolescentis ATCC 15703 GCA_000010425.1
522 Bifidobacterium dentium JCM 1195 = DSM 20436 GCA_001042595.1
503 Bifidobacterium angulatum DSM 20098 = JCM 7096 GCA_001025155.1
86
736 Rhodobacter sphaeroides 2.4.1 GCA_000012905.2
662 Rhodobacter sphaeroides ATCC 17025 GCA_000016405.1
552 Gemmobacter aquatilis GCA_900110025.1
87
775 Deinococcus radiodurans R1 GCA_000008565.1
461 Deinococcus proteolyticus MRP GCA_000190555.1
460 Deinococcus deserti VCD115 GCA_000020685.1
88
803 Streptococcus mutans UA159 GCA_000007465.2
565 Streptococcus ratti FA-1 = DSM 20564 GCA_000286075.1
561 Streptococcus gordonii str. Challis substr. CH1 GCA_000017005.1
89
635 Deinococcus radiodurans R1 GCA_000008565.1
448 Deinococcus deserti VCD115 GCA_000020685.1
408 Deinococcus gobiensis I-0 GCA_000252445.1
8a
419 Bifidobacterium adolescentis ATCC 15703 GCA_000010425.1
348 Bifidobacterium angulatum DSM 20098 = JCM 7096 GCA_001025155.1
326 Bifidobacterium dentium JCM 1195 = DSM 20436 GCA_001042595.1
8b
448 Clostridium beijerinckii GCA_000833105.2
427 Clostridium saccharoperbutylacetonicum N1-4_28HMT_29 GCA_000340885.1
415 Clostridium saccharobutylicum DSM 13864 GCA_000473995.1
8c
696 Clostridium beijerinckii GCA_000833105.2
680 Clostridium saccharoperbutylacetonicum N1-4_28HMT_29 GCA_000340885.1
651 Clostridium saccharobutylicum DSM 13864 GCA_000473995.1
8d
690 Rhodobacter sphaeroides 2.4.1 GCA_000012905.2
589 Rhodobacter sphaeroides ATCC 17025 GCA_000016405.1
482 Gemmobacter aquatilis GCA_900110025.1
8e
525 Enterococcus faecalis V583 GCA_000007785.1
492 Enterococcus thailandicus GCA_001652875.1
491 Enterococcus canis NBRC 100695 GCA_001544375.1

8f
591 Streptococcus mutans UA159 GCA_000007465.2
417 Streptococcus ratti FA-1 = DSM 20564 GCA_000286075.1
398 Streptococcus sobrinus DSM 20742 = ATCC 33478 GCA_000686605.1
8g
439 Enterococcus faecalis V583 GCA_000007785.1
395 Streptomyces cinnamoneus GCA_001885705.1
390 Enterococcus faecium DO GCA_000174395.2
8h
773 Lactobacillus gasseri ATCC 33323 = JCM 1131 GCA_000014425.1
692 Lactobacillus hominis DSM 23910 = CRBIP 24.179 GCA_000296835.1
577 Lactobacillus iners DSM 13335 GCA_000160875.1
8i
697 Deinococcus radiodurans R1 GCA_000008565.1
425 Deinococcus deserti VCD115 GCA_000020685.1
406 Deinococcus gobiensis I-0 GCA_000252445.1
406 Deinococcus murrayi DSM 11303 GCA_000482805.1
8j
332 Rhodobacter sphaeroides 2.4.1 GCA_000012905.2
253 Rhodobacter sphaeroides ATCC 17025 GCA_000016405.1
250 Pseudorhodobacter psychrotolerans GCA_001294535.1
8k
607 Staphylococcus epidermidis ATCC 12228 GCA_000007645.1
521 Staphylococcus capitis subsp. capitis GCA_001028645.1
509 Staphylococcus haemolyticus JCSC1435 GCA_000009865.1
509 Staphylococcus hominis subsp. hominis C80 GCA_000183685.1
8l
611 Deinococcus radiodurans R1 GCA_000008565.1
379 Deinococcus deserti VCD115 GCA_000020685.1
365 Deinococcus hopiensis KR-140 GCA_900176165.1
8m
769 Escherichia coli O104_3AH4 str. 2011C-3493 GCA_000299455.1
768 Escherichia coli str. K-12 substr. MG1655 GCA_000005845.2
768 Shigella flexneri 2a str. 301 GCA_000006925.2
765 Escherichia coli O157_3AH7 str. Sakai GCA_000008865.1
8n
466 Clostridium saccharobutylicum DSM 13864 GCA_000473995.1
462 Clostridium beijerinckii GCA_000833105.2
451 Clostridium puniceum GCA_002006345.1
451 Clostridium saccharoperbutylacetonicum N1-4_28HMT_29 GCA_000340885.1
8o
1002 Streptococcus mutans UA159 GCA_000007465.2
777 Streptococcus ratti FA-1 = DSM 20564 GCA_000286075.1
756 Streptococcus gordonii str. Challis substr. CH1 GCA_000017005.1
8p
770 Lactobacillus gasseri ATCC 33323 = JCM 1131 GCA_000014425.1
713 Lactobacillus hominis DSM 23910 = CRBIP 24.179 GCA_000296835.1
543 Lactobacillus iners DSM 13335 GCA_000160875.1
8q
864 Enterococcus faecalis V583 GCA_000007785.1
817 Streptomyces cinnamoneus GCA_001885705.1
744 Enterococcus rivorum GCA_001742285.1
8r
615 Enterococcus faecalis V583 GCA_000007785.1
568 Streptomyces cinnamoneus GCA_001885705.1
509 Enterococcus rivorum GCA_001742285.1
8s
531 Lactobacillus gasseri ATCC 33323 = JCM 1131 GCA_000014425.1
476 Lactobacillus hominis DSM 23910 = CRBIP 24.179 GCA_000296835.1
374 Lactobacillus amylolyticus GCA_002075105.1
374 Lactobacillus hamsteri DSM 5661 = JCM 6256 GCA_000615445.1
8t
466 Bacillus anthracis str. Ames GCA_000007845.1
466 Bacillus anthracis str. Sterne GCA_000008165.1
466 Bacillus cereus ATCC 14579 GCA_000007825.1
466 Bacillus mycoides GCA_000832605.1
466 Bacillus pseudomycoides DSM 12442 GCA_000161455.1
466 _5BBacillus thuringiensis_5D serovar konkukian str. 97-27 GCA_000008505.1
451 Bacillus thuringiensis YBT-1518 GCA_000497525.2
391 Bacillus cytotoxicus NVH 391-98 GCA_000017425.1
8u
818 Bacillus anthracis str. Ames GCA_000007845.1
818 Bacillus anthracis str. Sterne GCA_000008165.1
818 Bacillus cereus ATCC 14579 GCA_000007825.1
818 Bacillus thuringiensis YBT-1518 GCA_000497525.2
818 _5BBacillus thuringiensis_5D serovar konkukian str. 97-27 GCA_000008505.1

803 Bacillus pseudomycoides DSM 12442 GCA_000161455.1
778 Bacillus mycoides GCA_000832605.1
8v
350 Enterococcus faecalis V583 GCA_000007785.1
312 Enterococcus rivorum GCA_001742285.1
303 Streptomyces cinnamoneus GCA_001885705.1
8w
848 Bacillus cereus ATCC 14579 GCA_000007825.1
846 Bacillus anthracis str. Ames GCA_000007845.1
846 Bacillus anthracis str. Sterne GCA_000008165.1
846 _5BBacillus thuringiensis_5D serovar konkukian str. 97-27 GCA_000008505.1
811 Bacillus mycoides GCA_000832605.1
811 Bacillus thuringiensis YBT-1518 GCA_000497525.2
8x
648 Bifidobacterium adolescentis ATCC 15703 GCA_000010425.1
574 Bifidobacterium angulatum DSM 20098 = JCM 7096 GCA_001025155.1
543 Bifidobacterium dentium JCM 1195 = DSM 20436 GCA_001042595.1
8y
620 Clostridium beijerinckii GCA_000833105.2
577 Clostridium puniceum GCA_002006345.1
577 Clostridium saccharoperbutylacetonicum N1-4_28HMT_29 GCA_000340885.1
565 Clostridium butyricum GCA_001456065.2
8z
819 Rhodobacter sphaeroides 2.4.1 GCA_000012905.2
731 Rhodobacter sphaeroides ATCC 17025 GCA_000016405.1
607 Pseudorhodobacter psychrotolerans GCA_001294535.1
8A
749 Deinococcus radiodurans R1 GCA_000008565.1
466 Deinococcus deserti VCD115 GCA_000020685.1
444 Deinococcus soli Cha et al. 2016 GCA_001007995.1
8B
731 Clostridium beijerinckii GCA_000833105.2
690 Clostridium puniceum GCA_002006345.1
690 Clostridium saccharoperbutylacetonicum N1-4_28HMT_29 GCA_000340885.1
682 Clostridium saccharobutylicum DSM 13864 GCA_000473995.1
8C
861 Deinococcus radiodurans R1 GCA_000008565.1
592 Deinococcus gobiensis I-0 GCA_000252445.1
560 Deinococcus deserti VCD115 GCA_000020685.1
8D
737 Clostridium beijerinckii GCA_000833105.2
676 Clostridium saccharoperbutylacetonicum N1-4_28HMT_29 GCA_000340885.1
661 Clostridium puniceum GCA_002006345.1
8E
928 Enterococcus faecalis V583 GCA_000007785.1
881 Streptomyces cinnamoneus GCA_001885705.1
795 Enterococcus rivorum GCA_001742285.1
8F
714 Lactobacillus gasseri ATCC 33323 = JCM 1131 GCA_000014425.1
662 Lactobacillus hominis DSM 23910 = CRBIP 24.179 GCA_000296835.1
464 Lactobacillus jensenii GCA_001936235.1
464 Lactobacillus psittaci DSM 15354 GCA_000425905.1
8G
887 Staphylococcus epidermidis ATCC 12228 GCA_000007645.1
823 Staphylococcus capitis subsp. capitis GCA_001028645.1
720 Staphylococcus warneri SG1 GCA_000332735.1
8H
510 Bifidobacterium adolescentis ATCC 15703 GCA_000010425.1
405 Bifidobacterium dentium JCM 1195 = DSM 20436 GCA_001042595.1
403 Bifidobacterium longum NCC2705 GCA_000007525.1
8I
526 Rhodobacter sphaeroides 2.4.1 GCA_000012905.2
439 Rhodobacter sphaeroides ATCC 17025 GCA_000016405.1
372 Gemmobacter megaterium GCA_900156815.1
8J
683 Clostridium beijerinckii GCA_000833105.2
655 Clostridium saccharoperbutylacetonicum N1-4_28HMT_29 GCA_000340885.1
642 Clostridium saccharobutylicum DSM 13864 GCA_000473995.1
8K
579 Clostridium beijerinckii GCA_000833105.2
529 Clostridium saccharobutylicum DSM 13864 GCA_000473995.1
524 Clostridium butyricum GCA_001456065.2
524 Clostridium saccharoperbutylacetonicum N1-4_28HMT_29 GCA_000340885.1
8L
715 Staphylococcus epidermidis ATCC 12228 GCA_000007645.1
678 Staphylococcus capitis subsp. capitis GCA_001028645.1

583 Staphylococcus haemolyticus JCSC1435 GCA_000009865.1
8M
737 Streptococcus mutans UA159 GCA_000007465.2
553 Streptococcus ratti FA-1 = DSM 20564 GCA_000286075.1
486 Streptococcus halotolerans GCA_001598035.1
8N
433 Staphylococcus epidermidis ATCC 12228 GCA_000007645.1
415 Staphylococcus capitis subsp. capitis GCA_001028645.1
372 Staphylococcus hominis subsp. hominis C80 GCA_000183685.1
8O
568 Enterococcus faecalis V583 GCA_000007785.1
524 Streptomyces cinnamoneus GCA_001885705.1
480 Enterococcus rivorum GCA_001742285.1
8P
724 Clostridium beijerinckii GCA_000833105.2
680 Clostridium saccharoperbutylacetonicum N1-4_28HMT_29 GCA_000340885.1
674 Clostridium butyricum GCA_001456065.2
8Q
810 Rhodobacter sphaeroides 2.4.1 GCA_000012905.2
726 Rhodobacter sphaeroides ATCC 17025 GCA_000016405.1
600 Gemmobacter aquatilis GCA_900110025.1
8R
681 Rhodobacter sphaeroides 2.4.1 GCA_000012905.2
595 Rhodobacter sphaeroides ATCC 17025 GCA_000016405.1
512 Gemmobacter aquatilis GCA_900110025.1
8S
797 Streptococcus mutans UA159 GCA_000007465.2
593 Streptococcus ratti FA-1 = DSM 20564 GCA_000286075.1
542 Streptococcus gordonii str. Challis substr. CH1 GCA_000017005.1
8T
670 Lactobacillus gasseri ATCC 33323 = JCM 1131 GCA_000014425.1
599 Lactobacillus hominis DSM 23910 = CRBIP 24.179 GCA_000296835.1
473 Lactobacillus hamsteri DSM 5661 = JCM 6256 GCA_000615445.1
8U
724 Deinococcus radiodurans R1 GCA_000008565.1
373 Deinococcus deserti VCD115 GCA_000020685.1
373 Deinococcus proteolyticus MRP GCA_000190555.1
361 Deinococcus murrayi DSM 11303 GCA_000482805.1
8V
683 Rhodobacter sphaeroides 2.4.1 GCA_000012905.2
645 Rhodobacter sphaeroides ATCC 17025 GCA_000016405.1
527 Defluviimonas alba GCA_001620265.1
527 Pseudorhodobacter psychrotolerans GCA_001294535.1
8W
886 Rhodobacter sphaeroides 2.4.1 GCA_000012905.2
846 Rhodobacter sphaeroides ATCC 17025 GCA_000016405.1
688 Gemmobacter aquatilis GCA_900110025.1
8X
731 Lactobacillus gasseri ATCC 33323 = JCM 1131 GCA_000014425.1
676 Lactobacillus hominis DSM 23910 = CRBIP 24.179 GCA_000296835.1
496 Lactobacillus iners DSM 13335 GCA_000160875.1
8Y
789 Deinococcus radiodurans R1 GCA_000008565.1
474 Deinococcus deserti VCD115 GCA_000020685.1
425 Deinococcus gobiensis I-0 GCA_000252445.1
8Z
654 Rhodobacter sphaeroides 2.4.1 GCA_000012905.2
574 Rhodobacter sphaeroides ATCC 17025 GCA_000016405.1
503 Gemmobacter megaterium GCA_900156815.1
90
375 Lactobacillus gasseri ATCC 33323 = JCM 1131 GCA_000014425.1
375 Lactobacillus hominis DSM 23910 = CRBIP 24.179 GCA_000296835.1
313 Lactobacillus jensenii GCA_001936235.1
313 Lactobacillus psittaci DSM 15354 GCA_000425905.1
296 Lactobacillus mellis GCA_000967245.1
91
391 Deinococcus radiodurans R1 GCA_000008565.1
222 Deinococcus gobiensis I-0 GCA_000252445.1
202 Deinococcus maricopensis DSM 21211 GCA_000186385.1
92
750 Streptococcus mutans UA159 GCA_000007465.2
478 Streptococcus gordonii str. Challis substr. CH1 GCA_000017005.1
478 Streptococcus ratti FA-1 = DSM 20564 GCA_000286075.1
467 Streptococcus gallolyticus subsp. gallolyticus DSM 16831 GCA_002000985.1
93
682 Streptococcus mutans UA159 GCA_000007465.2

494 Streptococcus ratti FA-1 = DSM 20564 GCA_000286075.1
473 Streptococcus gordonii str. Challis substr. CH1 GCA_000017005.1
94
777 Bifidobacterium adolescentis ATCC 15703 GCA_000010425.1
683 Bifidobacterium dentium JCM 1195 = DSM 20436 GCA_001042595.1
641 Bifidobacterium thermophilum GCA_000741495.1
95
642 Staphylococcus epidermidis ATCC 12228 GCA_000007645.1
582 Staphylococcus capitis subsp. capitis GCA_001028645.1
553 Staphylococcus hominis subsp. hominis C80 GCA_000183685.1
96
350 Deinococcus radiodurans R1 GCA_000008565.1
155 Deinococcus deserti VCD115 GCA_000020685.1
155 Deinococcus proteolyticus MRP GCA_000190555.1
141 Deinococcus frigens DSM 12807 GCA_000701425.1
141 Deinococcus puniceus GCA_001644565.1
97
477 Escherichia coli O104_3AH4 str. 2011C-3493 GCA_000299455.1
477 Escherichia coli str. K-12 substr. MG1655 GCA_000005845.2
471 Escherichia coli IAI39 GCA_000026345.1
471 Shigella flexneri 2a str. 301 GCA_000006925.2
464 Escherichia coli O157_3AH7 str. Sakai GCA_000008865.1
464 Shigella dysenteriae Sd197 GCA_000012005.1
98
419 Clostridium beijerinckii GCA_000833105.2
390 Clostridium saccharoperbutylacetonicum N1-4_28HMT_29 GCA_000340885.1
384 Clostridium puniceum GCA_002006345.1
99
886 Clostridium beijerinckii GCA_000833105.2
847 Clostridium saccharoperbutylacetonicum N1-4_28HMT_29 GCA_000340885.1
788 Clostridium saccharobutylicum DSM 13864 GCA_000473995.1
9a
641 Enterococcus faecalis V583 GCA_000007785.1
594 Streptomyces cinnamoneus GCA_001885705.1
552 Enterococcus asini ATCC 700915 GCA_000407365.1
9b
628 Enterococcus faecalis V583 GCA_000007785.1
581 Streptomyces cinnamoneus GCA_001885705.1
555 Enterococcus asini ATCC 700915 GCA_000407365.1
9c
639 Streptococcus mutans UA159 GCA_000007465.2
462 Streptococcus ratti FA-1 = DSM 20564 GCA_000286075.1
441 Streptococcus halotolerans GCA_001598035.1
9d
720 Staphylococcus epidermidis ATCC 12228 GCA_000007645.1
686 Staphylococcus capitis subsp. capitis GCA_001028645.1
640 Staphylococcus haemolyticus JCSC1435 GCA_000009865.1
9e
829 Rhodobacter sphaeroides 2.4.1 GCA_000012905.2
695 Rhodobacter sphaeroides ATCC 17025 GCA_000016405.1
562 Defluviimonas alba GCA_001620265.1
9f
645 Lactobacillus gasseri ATCC 33323 = JCM 1131 GCA_000014425.1
601 Lactobacillus hominis DSM 23910 = CRBIP 24.179 GCA_000296835.1
449 Lactobacillus iners DSM 13335 GCA_000160875.1
9g
674 Enterococcus faecalis V583 GCA_000007785.1
627 Streptomyces cinnamoneus GCA_001885705.1
576 Enterococcus faecium DO GCA_000174395.2
9h
726 Streptococcus mutans UA159 GCA_000007465.2
523 Streptococcus ratti FA-1 = DSM 20564 GCA_000286075.1
494 Streptococcus salivarius GCA_000785515.1
9i
893 Lactobacillus gasseri ATCC 33323 = JCM 1131 GCA_000014425.1
781 Lactobacillus hominis DSM 23910 = CRBIP 24.179 GCA_000296835.1
613 Lactobacillus psittaci DSM 15354 GCA_000425905.1
9j
799 Clostridium beijerinckii GCA_000833105.2
778 Clostridium saccharoperbutylacetonicum N1-4_28HMT_29 GCA_000340885.1
738 Clostridium puniceum GCA_002006345.1
9k
710 Bacillus anthracis str. Ames GCA_000007845.1
710 Bacillus anthracis str. Sterne GCA_000008165.1
710 _5BBacillus thuringiensis_5D serovar konkukian str. 97-27 GCA_000008505.1
697 Bacillus cereus ATCC 14579 GCA_000007825.1

689 Bacillus thuringiensis YBT-1518 GCA_000497525.2
9l
790 Staphylococcus epidermidis ATCC 12228 GCA_000007645.1
763 Staphylococcus capitis subsp. capitis GCA_001028645.1
678 Staphylococcus hominis subsp. hominis C80 GCA_000183685.1
9m
752 Rhodobacter sphaeroides 2.4.1 GCA_000012905.2
681 Rhodobacter sphaeroides ATCC 17025 GCA_000016405.1
523 Pseudorhodobacter psychrotolerans GCA_001294535.1
9n
708 Rhodobacter sphaeroides 2.4.1 GCA_000012905.2
631 Rhodobacter sphaeroides ATCC 17025 GCA_000016405.1
461 Gemmobacter aquatilis GCA_900110025.1
9o
666 Staphylococcus epidermidis ATCC 12228 GCA_000007645.1
626 Staphylococcus capitis subsp. capitis GCA_001028645.1
590 Staphylococcus haemolyticus JCSC1435 GCA_000009865.1
590 Staphylococcus warneri SG1 GCA_000332735.1
9p
754 Clostridium beijerinckii GCA_000833105.2
745 Clostridium saccharoperbutylacetonicum N1-4_28HMT_29 GCA_000340885.1
728 Clostridium saccharobutylicum DSM 13864 GCA_000473995.1
9q
475 Clostridium beijerinckii GCA_000833105.2
468 Clostridium puniceum GCA_002006345.1
468 Clostridium saccharoperbutylacetonicum N1-4_28HMT_29 GCA_000340885.1
453 Clostridium chromiireducens GCA_002029255.1
9r
736 Lactobacillus gasseri ATCC 33323 = JCM 1131 GCA_000014425.1
640 Lactobacillus hominis DSM 23910 = CRBIP 24.179 GCA_000296835.1
485 Lactobacillus iners DSM 13335 GCA_000160875.1
9s
274 Staphylococcus epidermidis ATCC 12228 GCA_000007645.1
237 Staphylococcus capitis subsp. capitis GCA_001028645.1
236 Staphylococcus haemolyticus JCSC1435 GCA_000009865.1
9t
960 Bacillus anthracis str. Ames GCA_000007845.1
960 Bacillus anthracis str. Sterne GCA_000008165.1
952 Bacillus cereus ATCC 14579 GCA_000007825.1
952 _5BBacillus thuringiensis_5D serovar konkukian str. 97-27 GCA_000008505.1
937 Bacillus thuringiensis YBT-1518 GCA_000497525.2
9u
834 Escherichia coli str. K-12 substr. MG1655 GCA_000005845.2
808 Escherichia coli UMN026 GCA_000026325.2
805 Shigella flexneri 2a str. 301 GCA_000006925.2
9v
884 Streptococcus mutans UA159 GCA_000007465.2
652 Streptococcus ratti FA-1 = DSM 20564 GCA_000286075.1
606 Streptococcus ferus DSM 20646 GCA_000372425.1
9w
559 Deinococcus radiodurans R1 GCA_000008565.1
313 Deinococcus soli Cha et al. 2016 GCA_001007995.1
306 Deinococcus deserti VCD115 GCA_000020685.1
9x
638 Enterococcus faecalis V583 GCA_000007785.1
591 Streptomyces cinnamoneus GCA_001885705.1
573 Enterococcus rivorum GCA_001742285.1
9y
585 Rhodobacter sphaeroides 2.4.1 GCA_000012905.2
522 Rhodobacter sphaeroides ATCC 17025 GCA_000016405.1
428 Defluviimonas alba GCA_001620265.1
9z
896 Streptococcus mutans UA159 GCA_000007465.2
641 Streptococcus gordonii str. Challis substr. CH1 GCA_000017005.1
636 Streptococcus ratti FA-1 = DSM 20564 GCA_000286075.1
9A
448 Lactobacillus gasseri ATCC 33323 = JCM 1131 GCA_000014425.1
443 Lactobacillus hominis DSM 23910 = CRBIP 24.179 GCA_000296835.1
323 Lactobacillus kefiranofaciens ZW3 GCA_000214785.1
9B
809 Enterococcus faecalis V583 GCA_000007785.1
769 Streptomyces cinnamoneus GCA_001885705.1
687 Enterococcus rivorum GCA_001742285.1
9C
455 Enterococcus faecalis V583 GCA_000007785.1
425 Enterococcus dispar ATCC 51266 GCA_000406945.1

424 Enterococcus phoeniculicola ATCC BAA-412 GCA_000407505.1
9D
568 Staphylococcus epidermidis ATCC 12228 GCA_000007645.1
545 Staphylococcus capitis subsp. capitis GCA_001028645.1
474 Staphylococcus aureus subsp. aureus NCTC 8325 GCA_000013425.1
9E
396 Lactobacillus gasseri ATCC 33323 = JCM 1131 GCA_000014425.1
360 Lactobacillus hominis DSM 23910 = CRBIP 24.179 GCA_000296835.1
310 Lactobacillus iners DSM 13335 GCA_000160875.1
9F
935 Escherichia coli O157_3AH7 str. Sakai GCA_000008865.1
935 Escherichia coli UMN026 GCA_000026325.2
935 Escherichia coli str. K-12 substr. MG1655 GCA_000005845.2
924 Escherichia coli O83_3AH1 str. NRG 857C GCA_000183345.1
921 Shigella flexneri 2a str. 301 GCA_000006925.2
9G
661 Streptococcus mutans UA159 GCA_000007465.2
458 Streptococcus agalactiae 2603V_2FR GCA_000007265.1
455 Streptococcus gordonii str. Challis substr. CH1 GCA_000017005.1
9H
414 Deinococcus radiodurans R1 GCA_000008565.1
177 Deinococcus gobiensis I-0 GCA_000252445.1
167 Deinococcus deserti VCD115 GCA_000020685.1
9I
913 Lactobacillus gasseri ATCC 33323 = JCM 1131 GCA_000014425.1
848 Lactobacillus hominis DSM 23910 = CRBIP 24.179 GCA_000296835.1
589 Lactobacillus psittaci DSM 15354 GCA_000425905.1
9J
800 Staphylococcus epidermidis ATCC 12228 GCA_000007645.1
781 Staphylococcus capitis subsp. capitis GCA_001028645.1
738 Staphylococcus aureus subsp. aureus NCTC 8325 GCA_000013425.1
9K
540 Bacillus pseudomycoides DSM 12442 GCA_000161455.1
525 Bacillus anthracis str. Ames GCA_000007845.1
525 Bacillus anthracis str. Sterne GCA_000008165.1
525 _5BBacillus thuringiensis_5D serovar konkukian str. 97-27 GCA_000008505.1
500 Bacillus cereus ATCC 14579 GCA_000007825.1
9L
823 Lactobacillus gasseri ATCC 33323 = JCM 1131 GCA_000014425.1
764 Lactobacillus hominis DSM 23910 = CRBIP 24.179 GCA_000296835.1
596 Lactobacillus iners DSM 13335 GCA_000160875.1
9M
775 Clostridium beijerinckii GCA_000833105.2
710 Clostridium saccharoperbutylacetonicum N1-4_28HMT_29 GCA_000340885.1
709 Clostridium saccharobutylicum DSM 13864 GCA_000473995.1
9N
687 Enterococcus faecalis V583 GCA_000007785.1
640 Streptomyces cinnamoneus GCA_001885705.1
555 Enterococcus massiliensis GCA_001050095.1
9O
731 Staphylococcus epidermidis ATCC 12228 GCA_000007645.1
713 Staphylococcus capitis subsp. capitis GCA_001028645.1
643 Staphylococcus haemolyticus JCSC1435 GCA_000009865.1
9P
580 Lactobacillus gasseri ATCC 33323 = JCM 1131 GCA_000014425.1
557 Lactobacillus hominis DSM 23910 = CRBIP 24.179 GCA_000296835.1
441 Lactobacillus iners DSM 13335 GCA_000160875.1
9Q
744 Rhodobacter sphaeroides 2.4.1 GCA_000012905.2
677 Rhodobacter sphaeroides ATCC 17025 GCA_000016405.1
545 Pseudorhodobacter psychrotolerans GCA_001294535.1
9R
541 Bacillus anthracis str. Ames GCA_000007845.1
541 Bacillus anthracis str. Sterne GCA_000008165.1
541 _5BBacillus thuringiensis_5D serovar konkukian str. 97-27 GCA_000008505.1
525 Bacillus cereus ATCC 14579 GCA_000007825.1
525 Bacillus thuringiensis YBT-1518 GCA_000497525.2
516 Bacillus pseudomycoides DSM 12442 GCA_000161455.1
9S
793 Clostridium beijerinckii GCA_000833105.2
748 Clostridium puniceum GCA_002006345.1
748 Clostridium saccharoperbutylacetonicum N1-4_28HMT_29 GCA_000340885.1
719 Clostridium saccharobutylicum DSM 13864 GCA_000473995.1
9T
985 Deinococcus radiodurans R1 GCA_000008565.1
622 Deinococcus deserti VCD115 GCA_000020685.1

581 Deinococcus gobiensis I-0 GCA_000252445.1
9U
582 Deinococcus radiodurans R1 GCA_000008565.1
372 Deinococcus gobiensis I-0 GCA_000252445.1
363 Deinococcus marmoris DSM 12784 GCA_000701405.1
363 Deinococcus swuensis GCA_000800395.1
9V
675 Streptococcus mutans UA159 GCA_000007465.2
478 Streptococcus gordonii str. Challis substr. CH1 GCA_000017005.1
471 Streptococcus ratti FA-1 = DSM 20564 GCA_000286075.1
9W
498 Clostridium beijerinckii GCA_000833105.2
496 Clostridium saccharobutylicum DSM 13864 GCA_000473995.1
485 Clostridium saccharoperbutylacetonicum N1-4_28HMT_29 GCA_000340885.1
9X
869 Streptococcus mutans UA159 GCA_000007465.2
658 Streptococcus ratti FA-1 = DSM 20564 GCA_000286075.1
619 Streptococcus gordonii str. Challis substr. CH1 GCA_000017005.1
9Y
330 Bifidobacterium adolescentis ATCC 15703 GCA_000010425.1
287 Bifidobacterium breve DSM 20213 = JCM 1192 GCA_001025175.1
286 Bifidobacterium lemurum GCA_001895165.1
286 Bifidobacterium mongoliense DSM 21395 GCA_000741285.1
9Z
598 Rhodobacter sphaeroides 2.4.1 GCA_000012905.2
551 Rhodobacter sphaeroides ATCC 17025 GCA_000016405.1
439 Pseudorhodobacter psychrotolerans GCA_001294535.1
a0
822 Lactobacillus gasseri ATCC 33323 = JCM 1131 GCA_000014425.1
783 Lactobacillus hominis DSM 23910 = CRBIP 24.179 GCA_000296835.1
567 Lactobacillus hamsteri DSM 5661 = JCM 6256 GCA_000615445.1
567 Lactobacillus psittaci DSM 15354 GCA_000425905.1
a1
833 Enterococcus faecalis V583 GCA_000007785.1
786 Streptomyces cinnamoneus GCA_001885705.1
748 Enterococcus rivorum GCA_001742285.1
a2
767 Deinococcus radiodurans R1 GCA_000008565.1
495 Deinococcus deserti VCD115 GCA_000020685.1
466 Deinococcus hopiensis KR-140 GCA_900176165.1
a3
768 Lactobacillus gasseri ATCC 33323 = JCM 1131 GCA_000014425.1
709 Lactobacillus hominis DSM 23910 = CRBIP 24.179 GCA_000296835.1
533 Lactobacillus iners DSM 13335 GCA_000160875.1
a4
553 Bifidobacterium adolescentis ATCC 15703 GCA_000010425.1
454 Bifidobacterium dentium JCM 1195 = DSM 20436 GCA_001042595.1
432 Bifidobacterium callitrichos DSM 23973 GCA_000741175.1
a5
569 Rhodobacter sphaeroides 2.4.1 GCA_000012905.2
539 Rhodobacter sphaeroides ATCC 17025 GCA_000016405.1
460 Pseudorhodobacter psychrotolerans GCA_001294535.1
a6
623 Rhodobacter sphaeroides 2.4.1 GCA_000012905.2
561 Rhodobacter sphaeroides ATCC 17025 GCA_000016405.1
460 Defluviimonas alba GCA_001620265.1
a7
520 Clostridium beijerinckii GCA_000833105.2
507 Clostridium saccharoperbutylacetonicum N1-4_28HMT_29 GCA_000340885.1
500 Clostridium saccharobutylicum DSM 13864 GCA_000473995.1
a8
754 Clostridium beijerinckii GCA_000833105.2
753 Clostridium saccharoperbutylacetonicum N1-4_28HMT_29 GCA_000340885.1
719 Clostridium saccharobutylicum DSM 13864 GCA_000473995.1
a9
734 Rhodobacter sphaeroides 2.4.1 GCA_000012905.2
688 Rhodobacter sphaeroides ATCC 17025 GCA_000016405.1
598 Gemmobacter aquatilis GCA_900110025.1
aa
388 Rhodobacter sphaeroides 2.4.1 GCA_000012905.2
358 Rhodobacter sphaeroides ATCC 17025 GCA_000016405.1
320 Gemmobacter aquatilis GCA_900110025.1
ab
393 Rhodobacter sphaeroides 2.4.1 GCA_000012905.2
320 Rhodobacter sphaeroides ATCC 17025 GCA_000016405.1
269 Gemmobacter megaterium GCA_900156815.1

269 Pseudorhodobacter psychrotolerans GCA_001294535.1
ac
726 Rhodobacter sphaeroides 2.4.1 GCA_000012905.2
679 Rhodobacter sphaeroides ATCC 17025 GCA_000016405.1
591 Defluviimonas alba GCA_001620265.1
ad
528 Bacillus anthracis str. Ames GCA_000007845.1
528 Bacillus anthracis str. Sterne GCA_000008165.1
528 Bacillus cereus ATCC 14579 GCA_000007825.1
528 Bacillus pseudomycoides DSM 12442 GCA_000161455.1
528 Bacillus thuringiensis YBT-1518 GCA_000497525.2
528 _5BBacillus thuringiensis_5D serovar konkukian str. 97-27 GCA_000008505.1
521 Bacillus mycoides GCA_000832605.1
472 Bacillus cytotoxicus NVH 391-98 GCA_000017425.1
ae
649 Enterococcus faecalis V583 GCA_000007785.1
602 Streptomyces cinnamoneus GCA_001885705.1
560 Enterococcus rivorum GCA_001742285.1
af
795 Deinococcus radiodurans R1 GCA_000008565.1
522 Deinococcus deserti VCD115 GCA_000020685.1
454 Deinococcus hopiensis KR-140 GCA_900176165.1
ag
574 Bacillus thuringiensis YBT-1518 GCA_000497525.2
564 Bacillus anthracis str. Ames GCA_000007845.1
564 Bacillus anthracis str. Sterne GCA_000008165.1
564 Bacillus cereus ATCC 14579 GCA_000007825.1
564 _5BBacillus thuringiensis_5D serovar konkukian str. 97-27 GCA_000008505.1
549 Bacillus mycoides GCA_000832605.1
ah
802 Rhodobacter sphaeroides 2.4.1 GCA_000012905.2
745 Rhodobacter sphaeroides ATCC 17025 GCA_000016405.1
613 Pseudorhodobacter psychrotolerans GCA_001294535.1
ai
851 Streptococcus mutans UA159 GCA_000007465.2
603 Streptococcus ratti FA-1 = DSM 20564 GCA_000286075.1
564 Streptococcus halotolerans GCA_001598035.1
aj
832 Streptococcus mutans UA159 GCA_000007465.2
592 Streptococcus ratti FA-1 = DSM 20564 GCA_000286075.1
537 Streptococcus ferus DSM 20646 GCA_000372425.1
ak
660 Streptococcus mutans UA159 GCA_000007465.2
466 Streptococcus ratti FA-1 = DSM 20564 GCA_000286075.1
451 Streptococcus iniae GCA_000831485.1
al
750 Streptococcus mutans UA159 GCA_000007465.2
545 Streptococcus ratti FA-1 = DSM 20564 GCA_000286075.1
494 Streptococcus sobrinus DSM 20742 = ATCC 33478 GCA_000686605.1
am
863 Lactobacillus gasseri ATCC 33323 = JCM 1131 GCA_000014425.1
802 Lactobacillus hominis DSM 23910 = CRBIP 24.179 GCA_000296835.1
562 Lactobacillus iners DSM 13335 GCA_000160875.1
an
569 Escherichia coli str. K-12 substr. MG1655 GCA_000005845.2
557 Escherichia coli O104_3AH4 str. 2011C-3493 GCA_000299455.1
543 Shigella flexneri 2a str. 301 GCA_000006925.2
ao
645 Rhodobacter sphaeroides 2.4.1 GCA_000012905.2
552 Rhodobacter sphaeroides ATCC 17025 GCA_000016405.1
475 Gemmobacter megaterium GCA_900156815.1
ap
842 Staphylococcus epidermidis ATCC 12228 GCA_000007645.1
765 Megasphaera cerevisiae DSM 20462 GCA_001045675.1
765 Staphylococcus warneri SG1 GCA_000332735.1
754 Staphylococcus capitis subsp. capitis GCA_001028645.1
aq
742 Staphylococcus epidermidis ATCC 12228 GCA_000007645.1
700 Staphylococcus capitis subsp. capitis GCA_001028645.1
631 Megasphaera cerevisiae DSM 20462 GCA_001045675.1
631 Staphylococcus warneri SG1 GCA_000332735.1
ar
527 Clostridium saccharoperbutylacetonicum N1-4_28HMT_29 GCA_000340885.1
512 Clostridium puniceum GCA_002006345.1
509 Clostridium beijerinckii GCA_000833105.2
as

633 Clostridium beijerinckii GCA_000833105.2
616 Clostridium saccharoperbutylacetonicum N1-4_28HMT_29 GCA_000340885.1
607 Clostridium saccharobutylicum DSM 13864 GCA_000473995.1
at
215 Lactobacillus gasseri ATCC 33323 = JCM 1131 GCA_000014425.1
203 Lactobacillus hominis DSM 23910 = CRBIP 24.179 GCA_000296835.1
108 Lactobacillus delbrueckii subsp. bulgaricus ATCC 11842 = JCM 1002 GCA_000056065.1
au
826 Deinococcus radiodurans R1 GCA_000008565.1
497 Deinococcus gobiensis I-0 GCA_000252445.1
485 Deinococcus deserti VCD115 GCA_000020685.1
av
641 Enterococcus faecalis V583 GCA_000007785.1
594 Streptomyces cinnamoneus GCA_001885705.1
559 Enterococcus rivorum GCA_001742285.1
aw
570 Streptococcus mutans UA159 GCA_000007465.2
376 Streptococcus gordonii str. Challis substr. CH1 GCA_000017005.1
373 Streptococcus equinus GCA_000964315.1
373 Streptococcus gallolyticus subsp. gallolyticus DSM 16831 GCA_002000985.1
ax
755 Lactobacillus gasseri ATCC 33323 = JCM 1131 GCA_000014425.1
687 Lactobacillus hominis DSM 23910 = CRBIP 24.179 GCA_000296835.1
442 Lactobacillus iners DSM 13335 GCA_000160875.1
ay
796 Staphylococcus epidermidis ATCC 12228 GCA_000007645.1
781 Staphylococcus capitis subsp. capitis GCA_001028645.1
742 Staphylococcus hominis subsp. hominis C80 GCA_000183685.1
az
401 Staphylococcus epidermidis ATCC 12228 GCA_000007645.1
363 Staphylococcus haemolyticus JCSC1435 GCA_000009865.1
347 Staphylococcus hominis subsp. hominis C80 GCA_000183685.1
aA
534 Clostridium beijerinckii GCA_000833105.2
497 Clostridium saccharoperbutylacetonicum N1-4_28HMT_29 GCA_000340885.1
474 Clostridium saccharobutylicum DSM 13864 GCA_000473995.1
aB
453 Streptococcus mutans UA159 GCA_000007465.2
333 Streptococcus ratti FA-1 = DSM 20564 GCA_000286075.1
319 Streptococcus sobrinus DSM 20742 = ATCC 33478 GCA_000686605.1
aC
655 Clostridium beijerinckii GCA_000833105.2
601 Clostridium saccharoperbutylacetonicum N1-4_28HMT_29 GCA_000340885.1
567 Clostridium puniceum GCA_002006345.1
aD
552 Clostridium beijerinckii GCA_000833105.2
536 Clostridium saccharoperbutylacetonicum N1-4_28HMT_29 GCA_000340885.1
529 Clostridium saccharobutylicum DSM 13864 GCA_000473995.1
aE
730 Bifidobacterium adolescentis ATCC 15703 GCA_000010425.1
637 Bifidobacterium dentium JCM 1195 = DSM 20436 GCA_001042595.1
596 Bifidobacterium angulatum DSM 20098 = JCM 7096 GCA_001025155.1
aF
548 Clostridium puniceum GCA_002006345.1
534 Clostridium beijerinckii GCA_000833105.2
533 Clostridium saccharoperbutylacetonicum N1-4_28HMT_29 GCA_000340885.1
aG
843 Deinococcus radiodurans R1 GCA_000008565.1
519 Deinococcus deserti VCD115 GCA_000020685.1
505 Deinococcus hopiensis KR-140 GCA_900176165.1
aH
510 Streptococcus mutans UA159 GCA_000007465.2
375 Streptococcus ratti FA-1 = DSM 20564 GCA_000286075.1
365 Streptococcus macacae NCTC 11558 GCA_000187995.3
aI
712 Bifidobacterium adolescentis ATCC 15703 GCA_000010425.1
612 Bifidobacterium dentium JCM 1195 = DSM 20436 GCA_001042595.1
553 Bifidobacterium angulatum DSM 20098 = JCM 7096 GCA_001025155.1
aJ
628 Clostridium beijerinckii GCA_000833105.2
606 Clostridium saccharoperbutylacetonicum N1-4_28HMT_29 GCA_000340885.1
597 Clostridium saccharobutylicum DSM 13864 GCA_000473995.1
aK
658 Staphylococcus epidermidis ATCC 12228 GCA_000007645.1
602 Staphylococcus capitis subsp. capitis GCA_001028645.1
601 Staphylococcus simulans GCA_001559115.1

aL
746 Streptococcus mutans UA159 GCA_000007465.2
572 Streptococcus ratti FA-1 = DSM 20564 GCA_000286075.1
506 Streptococcus ferus DSM 20646 GCA_000372425.1
aM
608 Rhodobacter sphaeroides 2.4.1 GCA_000012905.2
553 Rhodobacter sphaeroides ATCC 17025 GCA_000016405.1
460 Gemmobacter megaterium GCA_900156815.1
aN
612 Rhodobacter sphaeroides 2.4.1 GCA_000012905.2
582 Rhodobacter sphaeroides ATCC 17025 GCA_000016405.1
507 Pseudorhodobacter psychrotolerans GCA_001294535.1
aO
707 Staphylococcus epidermidis ATCC 12228 GCA_000007645.1
684 Staphylococcus capitis subsp. capitis GCA_001028645.1
659 Staphylococcus warneri SG1 GCA_000332735.1
aP
362 Streptococcus mutans UA159 GCA_000007465.2
266 Streptococcus equinus GCA_000964315.1
262 Streptococcus gallolyticus subsp. gallolyticus DSM 16831 GCA_002000985.1
262 Streptococcus gordonii str. Challis substr. CH1 GCA_000017005.1
aQ
900 Clostridium beijerinckii GCA_000833105.2
868 Clostridium saccharoperbutylacetonicum N1-4_28HMT_29 GCA_000340885.1
838 Clostridium puniceum GCA_002006345.1
aR
741 Enterococcus faecalis V583 GCA_000007785.1
694 Streptomyces cinnamoneus GCA_001885705.1
621 Enterococcus massiliensis GCA_001050095.1
aS
736 Bacillus anthracis str. Ames GCA_000007845.1
736 Bacillus anthracis str. Sterne GCA_000008165.1
736 _5BBacillus thuringiensis_5D serovar konkukian str. 97-27 GCA_000008505.1
721 Bacillus cereus ATCC 14579 GCA_000007825.1
711 Bacillus thuringiensis YBT-1518 GCA_000497525.2
aT
863 Lactobacillus gasseri ATCC 33323 = JCM 1131 GCA_000014425.1
820 Lactobacillus hominis DSM 23910 = CRBIP 24.179 GCA_000296835.1
656 Lactobacillus iners DSM 13335 GCA_000160875.1
aU
745 Rhodobacter sphaeroides 2.4.1 GCA_000012905.2
666 Rhodobacter sphaeroides ATCC 17025 GCA_000016405.1
543 Gemmobacter aquatilis GCA_900110025.1
aV
692 Streptococcus mutans UA159 GCA_000007465.2
476 Streptococcus ratti FA-1 = DSM 20564 GCA_000286075.1
468 Streptococcus equinus GCA_000964315.1
aW
791 Clostridium beijerinckii GCA_000833105.2
715 Clostridium saccharoperbutylacetonicum N1-4_28HMT_29 GCA_000340885.1
681 Clostridium saccharobutylicum DSM 13864 GCA_000473995.1
aX
425 Staphylococcus epidermidis ATCC 12228 GCA_000007645.1
392 Staphylococcus capitis subsp. capitis GCA_001028645.1
388 Staphylococcus aureus subsp. aureus NCTC 8325 GCA_000013425.1
388 Staphylococcus haemolyticus JCSC1435 GCA_000009865.1
aY
813 Bacillus cereus ATCC 14579 GCA_000007825.1
813 _5BBacillus thuringiensis_5D serovar konkukian str. 97-27 GCA_000008505.1
812 Bacillus anthracis str. Ames GCA_000007845.1
812 Bacillus anthracis str. Sterne GCA_000008165.1
807 Bacillus thuringiensis YBT-1518 GCA_000497525.2
aZ
649 Clostridium beijerinckii GCA_000833105.2
617 Clostridium saccharoperbutylacetonicum N1-4_28HMT_29 GCA_000340885.1
612 Clostridium puniceum GCA_002006345.1
b0
642 Deinococcus radiodurans R1 GCA_000008565.1
375 Deinococcus gobiensis I-0 GCA_000252445.1
332 Deinococcus deserti VCD115 GCA_000020685.1
332 Deinococcus hopiensis KR-140 GCA_900176165.1
b1
543 Deinococcus radiodurans R1 GCA_000008565.1
299 Deinococcus deserti VCD115 GCA_000020685.1
282 Deinococcus gobiensis I-0 GCA_000252445.1
b2

615 Staphylococcus epidermidis ATCC 12228 GCA_000007645.1
559 Staphylococcus capitis subsp. capitis GCA_001028645.1
513 Staphylococcus aureus subsp. aureus NCTC 8325 GCA_000013425.1
b3
687 Clostridium beijerinckii GCA_000833105.2
641 Clostridium saccharoperbutylacetonicum N1-4_28HMT_29 GCA_000340885.1
605 Clostridium puniceum GCA_002006345.1
b4
920 Staphylococcus epidermidis ATCC 12228 GCA_000007645.1
891 Staphylococcus capitis subsp. capitis GCA_001028645.1
813 Staphylococcus aureus subsp. aureus NCTC 8325 GCA_000013425.1
b5
848 Streptococcus mutans UA159 GCA_000007465.2
640 Streptococcus ratti FA-1 = DSM 20564 GCA_000286075.1
619 Streptococcus gordonii str. Challis substr. CH1 GCA_000017005.1
b6
787 Staphylococcus epidermidis ATCC 12228 GCA_000007645.1
743 Staphylococcus capitis subsp. capitis GCA_001028645.1
714 Staphylococcus aureus subsp. aureus NCTC 8325 GCA_000013425.1
b7
705 Staphylococcus epidermidis ATCC 12228 GCA_000007645.1
678 Staphylococcus capitis subsp. capitis GCA_001028645.1
642 Staphylococcus warneri SG1 GCA_000332735.1
b8
607 Staphylococcus epidermidis ATCC 12228 GCA_000007645.1
549 Staphylococcus capitis subsp. capitis GCA_001028645.1
520 Staphylococcus warneri SG1 GCA_000332735.1
b9
750 Streptococcus mutans UA159 GCA_000007465.2
543 Streptococcus ratti FA-1 = DSM 20564 GCA_000286075.1
531 Streptococcus equinus GCA_000964315.1
ba
786 Bifidobacterium adolescentis ATCC 15703 GCA_000010425.1
624 Bifidobacterium dentium JCM 1195 = DSM 20436 GCA_001042595.1
609 Bifidobacterium breve DSM 20213 = JCM 1192 GCA_001025175.1
bb
656 Streptococcus mutans UA159 GCA_000007465.2
531 Streptococcus ratti FA-1 = DSM 20564 GCA_000286075.1
499 Streptococcus ferus DSM 20646 GCA_000372425.1
bc
526 Bifidobacterium adolescentis ATCC 15703 GCA_000010425.1
456 Bifidobacterium dentium JCM 1195 = DSM 20436 GCA_001042595.1
434 Bifidobacterium thermophilum GCA_000741495.1
bd
463 Streptococcus mutans UA159 GCA_000007465.2
352 Streptococcus ratti FA-1 = DSM 20564 GCA_000286075.1
345 Streptococcus equinus GCA_000964315.1
345 Streptococcus gallolyticus subsp. gallolyticus DSM 16831 GCA_002000985.1
be
533 Escherichia coli O157_3AH7 str. Sakai GCA_000008865.1
533 Escherichia coli str. K-12 substr. MG1655 GCA_000005845.2
524 Escherichia coli UMN026 GCA_000026325.2
508 Escherichia coli O104_3AH4 str. 2011C-3493 GCA_000299455.1
508 Escherichia coli O83_3AH1 str. NRG 857C GCA_000183345.1
508 Shigella flexneri 2a str. 301 GCA_000006925.2
bf
687 Lactobacillus gasseri ATCC 33323 = JCM 1131 GCA_000014425.1
616 Lactobacillus hominis DSM 23910 = CRBIP 24.179 GCA_000296835.1
450 Lactobacillus hamsteri DSM 5661 = JCM 6256 GCA_000615445.1
bg
743 Clostridium beijerinckii GCA_000833105.2
708 Clostridium saccharoperbutylacetonicum N1-4_28HMT_29 GCA_000340885.1
657 Clostridium puniceum GCA_002006345.1
bh
683 Streptococcus mutans UA159 GCA_000007465.2
471 Streptococcus ratti FA-1 = DSM 20564 GCA_000286075.1
443 Streptococcus ferus DSM 20646 GCA_000372425.1
bi
765 Staphylococcus epidermidis ATCC 12228 GCA_000007645.1
749 Staphylococcus capitis subsp. capitis GCA_001028645.1
660 Staphylococcus aureus subsp. aureus NCTC 8325 GCA_000013425.1
bj
716 Streptococcus mutans UA159 GCA_000007465.2
573 Streptococcus ratti FA-1 = DSM 20564 GCA_000286075.1
501 Streptococcus merionis DSM 19192 GCA_000380085.1
bk

539 Clostridium beijerinckii GCA_000833105.2
518 Clostridium saccharoperbutylacetonicum N1-4_28HMT_29 GCA_000340885.1
482 Clostridium saccharobutylicum DSM 13864 GCA_000473995.1
bl
831 Lactobacillus gasseri ATCC 33323 = JCM 1131 GCA_000014425.1
743 Lactobacillus hominis DSM 23910 = CRBIP 24.179 GCA_000296835.1
563 Lactobacillus iners DSM 13335 GCA_000160875.1
bm
658 Deinococcus radiodurans R1 GCA_000008565.1
413 Deinococcus gobiensis I-0 GCA_000252445.1
373 Deinococcus deserti VCD115 GCA_000020685.1
bn
662 Lactobacillus gasseri ATCC 33323 = JCM 1131 GCA_000014425.1
597 Lactobacillus hominis DSM 23910 = CRBIP 24.179 GCA_000296835.1
452 Lactobacillus iners DSM 13335 GCA_000160875.1
bo
622 Streptococcus mutans UA159 GCA_000007465.2
506 Streptococcus ratti FA-1 = DSM 20564 GCA_000286075.1
485 Streptococcus ovis DSM 16829 GCA_000380125.1
485 Streptococcus pyogenes M1 GAS GCA_000006785.2
485 Streptococcus uberis 0140J GCA_000009545.1
bp
237 Rhodobacter sphaeroides 2.4.1 GCA_000012905.2
184 Rhodobacter sphaeroides ATCC 17025 GCA_000016405.1
172 Rhodobacter vinaykumarii GCA_900156695.1
bq
536 Lactobacillus gasseri ATCC 33323 = JCM 1131 GCA_000014425.1
491 Lactobacillus hominis DSM 23910 = CRBIP 24.179 GCA_000296835.1
408 Lactobacillus iners DSM 13335 GCA_000160875.1
br
459 Staphylococcus epidermidis ATCC 12228 GCA_000007645.1
453 Staphylococcus capitis subsp. capitis GCA_001028645.1
398 Staphylococcus warneri SG1 GCA_000332735.1
bs
728 Escherichia coli str. K-12 substr. MG1655 GCA_000005845.2
706 Shigella flexneri 2a str. 301 GCA_000006925.2
704 Escherichia coli O104_3AH4 str. 2011C-3493 GCA_000299455.1
bt
748 Lactobacillus gasseri ATCC 33323 = JCM 1131 GCA_000014425.1
698 Lactobacillus hominis DSM 23910 = CRBIP 24.179 GCA_000296835.1
532 Lactobacillus iners DSM 13335 GCA_000160875.1
bu
684 Clostridium beijerinckii GCA_000833105.2
652 Clostridium saccharoperbutylacetonicum N1-4_28HMT_29 GCA_000340885.1
645 Clostridium saccharobutylicum DSM 13864 GCA_000473995.1
bv
878 Staphylococcus epidermidis ATCC 12228 GCA_000007645.1
816 Staphylococcus capitis subsp. capitis GCA_001028645.1
765 Staphylococcus hominis subsp. hominis C80 GCA_000183685.1
bw
529 Lactobacillus gasseri ATCC 33323 = JCM 1131 GCA_000014425.1
498 Lactobacillus hominis DSM 23910 = CRBIP 24.179 GCA_000296835.1
405 Lactobacillus iners DSM 13335 GCA_000160875.1
bx
606 Clostridium beijerinckii GCA_000833105.2
580 Clostridium saccharoperbutylacetonicum N1-4_28HMT_29 GCA_000340885.1
558 Clostridium saccharobutylicum DSM 13864 GCA_000473995.1
by
796 Clostridium beijerinckii GCA_000833105.2
768 Clostridium saccharoperbutylacetonicum N1-4_28HMT_29 GCA_000340885.1
755 Clostridium saccharobutylicum DSM 13864 GCA_000473995.1
bz
470 Streptococcus mutans UA159 GCA_000007465.2
279 Streptococcus gordonii str. Challis substr. CH1 GCA_000017005.1
274 Streptococcus anginosus C238 GCA_000463505.1
274 Streptococcus ratti FA-1 = DSM 20564 GCA_000286075.1
bA
593 Lactobacillus gasseri ATCC 33323 = JCM 1131 GCA_000014425.1
541 Lactobacillus hominis DSM 23910 = CRBIP 24.179 GCA_000296835.1
434 Lactobacillus iners DSM 13335 GCA_000160875.1
bB
752 Clostridium beijerinckii GCA_000833105.2
711 Clostridium saccharobutylicum DSM 13864 GCA_000473995.1
710 Clostridium saccharoperbutylacetonicum N1-4_28HMT_29 GCA_000340885.1
bC
772 Staphylococcus epidermidis ATCC 12228 GCA_000007645.1

747 Staphylococcus capitis subsp. capitis GCA_001028645.1
652 Staphylococcus aureus subsp. aureus NCTC 8325 GCA_000013425.1
bD
607 Deinococcus radiodurans R1 GCA_000008565.1
396 Deinococcus deserti VCD115 GCA_000020685.1
372 Deinococcus gobiensis I-0 GCA_000252445.1
bE
683 Deinococcus radiodurans R1 GCA_000008565.1
442 Deinococcus gobiensis I-0 GCA_000252445.1
426 Deinococcus deserti VCD115 GCA_000020685.1
bF
721 Streptococcus mutans UA159 GCA_000007465.2
575 Streptococcus ratti FA-1 = DSM 20564 GCA_000286075.1
567 Streptococcus gordonii str. Challis substr. CH1 GCA_000017005.1
bG
510 Clostridium beijerinckii GCA_000833105.2
510 Clostridium saccharoperbutylacetonicum N1-4_28HMT_29 GCA_000340885.1
495 Clostridium puniceum GCA_002006345.1
483 Clostridium taeniosporum GCA_001735765.1
bH
789 Clostridium beijerinckii GCA_000833105.2
789 Clostridium saccharoperbutylacetonicum N1-4_28HMT_29 GCA_000340885.1
773 Clostridium puniceum GCA_002006345.1
748 Clostridium butyricum GCA_001456065.2
bI
741 Clostridium beijerinckii GCA_000833105.2
692 Clostridium saccharoperbutylacetonicum N1-4_28HMT_29 GCA_000340885.1
672 Clostridium saccharobutylicum DSM 13864 GCA_000473995.1
bJ
538 Bifidobacterium adolescentis ATCC 15703 GCA_000010425.1
443 Bifidobacterium angulatum DSM 20098 = JCM 7096 GCA_001025155.1
406 Bifidobacterium gallicum DSM 20093 = LMG 11596 GCA_000741205.1
bK
616 Deinococcus radiodurans R1 GCA_000008565.1
367 Deinococcus deserti VCD115 GCA_000020685.1
366 Deinococcus gobiensis I-0 GCA_000252445.1
bL
683 Escherichia coli O157_3AH7 str. Sakai GCA_000008865.1
683 Escherichia coli str. K-12 substr. MG1655 GCA_000005845.2
678 Escherichia coli UMN026 GCA_000026325.2
650 Escherichia coli IAI39 GCA_000026345.1
650 Shigella flexneri 2a str. 301 GCA_000006925.2
bM
833 Streptococcus mutans UA159 GCA_000007465.2
608 Streptococcus ratti FA-1 = DSM 20564 GCA_000286075.1
568 Streptococcus criceti HS-6 GCA_000187975.3
bN
675 Lactobacillus gasseri ATCC 33323 = JCM 1131 GCA_000014425.1
648 Lactobacillus hominis DSM 23910 = CRBIP 24.179 GCA_000296835.1
467 Lactobacillus iners DSM 13335 GCA_000160875.1
bO
784 Rhodobacter sphaeroides 2.4.1 GCA_000012905.2
706 Rhodobacter sphaeroides ATCC 17025 GCA_000016405.1
579 Gemmobacter aquatilis GCA_900110025.1
bP
550 Bacillus anthracis str. Ames GCA_000007845.1
550 Bacillus anthracis str. Sterne GCA_000008165.1
550 _5BBacillus thuringiensis_5D serovar konkukian str. 97-27 GCA_000008505.1
533 Bacillus cereus ATCC 14579 GCA_000007825.1
533 Bacillus thuringiensis YBT-1518 GCA_000497525.2
508 Bacillus mycoides GCA_000832605.1
bQ
765 Clostridium beijerinckii GCA_000833105.2
698 Clostridium saccharoperbutylacetonicum N1-4_28HMT_29 GCA_000340885.1
693 Clostridium saccharobutylicum DSM 13864 GCA_000473995.1
bR
640 Clostridium beijerinckii GCA_000833105.2
635 Clostridium saccharoperbutylacetonicum N1-4_28HMT_29 GCA_000340885.1
630 Clostridium puniceum GCA_002006345.1
bS
797 Deinococcus radiodurans R1 GCA_000008565.1
530 Deinococcus gobiensis I-0 GCA_000252445.1
517 Deinococcus deserti VCD115 GCA_000020685.1
bT
550 Deinococcus radiodurans R1 GCA_000008565.1
351 Deinococcus deserti VCD115 GCA_000020685.1

351 Deinococcus hopiensis KR-140 GCA_900176165.1
323 Deinococcus puniceus GCA_001644565.1
bU
681 Rhodobacter sphaeroides 2.4.1 GCA_000012905.2
609 Rhodobacter sphaeroides ATCC 17025 GCA_000016405.1
496 Gemmobacter aquatilis GCA_900110025.1
bV
893 Clostridium beijerinckii GCA_000833105.2
865 Clostridium saccharoperbutylacetonicum N1-4_28HMT_29 GCA_000340885.1
817 Clostridium saccharobutylicum DSM 13864 GCA_000473995.1
bW
658 Deinococcus radiodurans R1 GCA_000008565.1
449 Deinococcus deserti VCD115 GCA_000020685.1
422 Deinococcus soli Cha et al. 2016 GCA_001007995.1
bX
677 Lactobacillus gasseri ATCC 33323 = JCM 1131 GCA_000014425.1
622 Lactobacillus hominis DSM 23910 = CRBIP 24.179 GCA_000296835.1
497 Lactobacillus iners DSM 13335 GCA_000160875.1
bY
624 Streptococcus mutans UA159 GCA_000007465.2
474 Streptococcus ratti FA-1 = DSM 20564 GCA_000286075.1
445 Streptococcus macacae NCTC 11558 GCA_000187995.3
bZ
392 Rhodobacter sphaeroides 2.4.1 GCA_000012905.2
352 Rhodobacter sphaeroides ATCC 17025 GCA_000016405.1
276 Defluviimonas alba GCA_001620265.1
c0
851 Streptococcus mutans UA159 GCA_000007465.2
636 Streptococcus ratti FA-1 = DSM 20564 GCA_000286075.1
617 Streptococcus ferus DSM 20646 GCA_000372425.1
c1
668 Deinococcus radiodurans R1 GCA_000008565.1
433 Deinococcus deserti VCD115 GCA_000020685.1
427 Deinococcus soli Cha et al. 2016 GCA_001007995.1
c2
714 Enterococcus faecalis V583 GCA_000007785.1
674 Streptomyces cinnamoneus GCA_001885705.1
632 Enterococcus rivorum GCA_001742285.1
c3
616 Bifidobacterium adolescentis ATCC 15703 GCA_000010425.1
489 Bifidobacterium angulatum DSM 20098 = JCM 7096 GCA_001025155.1
488 Bifidobacterium thermophilum GCA_000741495.1
c4
421 Rhodobacter sphaeroides 2.4.1 GCA_000012905.2
348 Rhodobacter sphaeroides ATCC 17025 GCA_000016405.1
319 Gemmobacter nectariphilus DSM 15620 GCA_000429765.1
319 Rhodobacter vinaykumarii GCA_900156695.1
c5
667 Staphylococcus epidermidis ATCC 12228 GCA_000007645.1
636 Staphylococcus capitis subsp. capitis GCA_001028645.1
566 Staphylococcus hominis subsp. hominis C80 GCA_000183685.1
c6
625 Bifidobacterium adolescentis ATCC 15703 GCA_000010425.1
540 Bifidobacterium dentium JCM 1195 = DSM 20436 GCA_001042595.1
483 Bifidobacterium stellenboschense GCA_000741785.1
c7
460 Streptococcus mutans UA159 GCA_000007465.2
345 Streptococcus ratti FA-1 = DSM 20564 GCA_000286075.1
338 Streptococcus gordonii str. Challis substr. CH1 GCA_000017005.1
c8
737 Staphylococcus epidermidis ATCC 12228 GCA_000007645.1
705 Staphylococcus capitis subsp. capitis GCA_001028645.1
656 Staphylococcus warneri SG1 GCA_000332735.1
c9
644 Streptococcus mutans UA159 GCA_000007465.2
431 Streptococcus ratti FA-1 = DSM 20564 GCA_000286075.1
377 Streptococcus halotolerans GCA_001598035.1
ca
399 Staphylococcus epidermidis ATCC 12228 GCA_000007645.1
374 Staphylococcus capitis subsp. capitis GCA_001028645.1
342 Staphylococcus lugdunensis HKU09-01 GCA_000025085.1
cb
738 Enterococcus faecalis V583 GCA_000007785.1
691 Streptomyces cinnamoneus GCA_001885705.1
618 Enterococcus canis NBRC 100695 GCA_001544375.1
cc

755 Rhodobacter sphaeroides 2.4.1 GCA_000012905.2
709 Rhodobacter sphaeroides ATCC 17025 GCA_000016405.1
609 Defluviimonas alba GCA_001620265.1
cd
583 Deinococcus radiodurans R1 GCA_000008565.1
277 Deinococcus deserti VCD115 GCA_000020685.1
266 Deinococcus marmoris DSM 12784 GCA_000701405.1
266 Deinococcus proteolyticus MRP GCA_000190555.1
ce
618 Clostridium beijerinckii GCA_000833105.2
589 Clostridium saccharoperbutylacetonicum N1-4_28HMT_29 GCA_000340885.1
579 Clostridium puniceum GCA_002006345.1
cf
306 Streptococcus mutans UA159 GCA_000007465.2
256 Streptococcus ratti FA-1 = DSM 20564 GCA_000286075.1
245 Streptococcus sanguinis SK36 GCA_000014205.1
cg
479 Rhodobacter sphaeroides 2.4.1 GCA_000012905.2
426 Rhodobacter sphaeroides ATCC 17025 GCA_000016405.1
339 Gemmobacter aquatilis GCA_900110025.1
ch
602 Rhodobacter sphaeroides 2.4.1 GCA_000012905.2
573 Rhodobacter sphaeroides ATCC 17025 GCA_000016405.1
459 Defluviimonas indica GCA_900106675.1
ci
406 Clostridium saccharoperbutylacetonicum N1-4_28HMT_29 GCA_000340885.1
403 Clostridium beijerinckii GCA_000833105.2
376 Clostridium puniceum GCA_002006345.1
376 Clostridium saccharobutylicum DSM 13864 GCA_000473995.1
cj
715 Clostridium beijerinckii GCA_000833105.2
673 Clostridium saccharoperbutylacetonicum N1-4_28HMT_29 GCA_000340885.1
668 Clostridium butyricum GCA_001456065.2
ck
780 Streptococcus mutans UA159 GCA_000007465.2
560 Streptococcus ratti FA-1 = DSM 20564 GCA_000286075.1
533 Streptococcus equinus GCA_000964315.1
cl
742 Clostridium beijerinckii GCA_000833105.2
705 Clostridium saccharoperbutylacetonicum N1-4_28HMT_29 GCA_000340885.1
669 Clostridium puniceum GCA_002006345.1
cm
920 Clostridium beijerinckii GCA_000833105.2
889 Clostridium saccharoperbutylacetonicum N1-4_28HMT_29 GCA_000340885.1
872 Clostridium saccharobutylicum DSM 13864 GCA_000473995.1
cn
701 Rhodobacter sphaeroides 2.4.1 GCA_000012905.2
627 Rhodobacter sphaeroides ATCC 17025 GCA_000016405.1
587 Gemmobacter aquatilis GCA_900110025.1
co
742 Bacillus anthracis str. Ames GCA_000007845.1
742 Bacillus anthracis str. Sterne GCA_000008165.1
742 _5BBacillus thuringiensis_5D serovar konkukian str. 97-27 GCA_000008505.1
740 Bacillus pseudomycoides DSM 12442 GCA_000161455.1
726 Bacillus cereus ATCC 14579 GCA_000007825.1
cp
700 Lactobacillus gasseri ATCC 33323 = JCM 1131 GCA_000014425.1
653 Lactobacillus hominis DSM 23910 = CRBIP 24.179 GCA_000296835.1
495 Lactobacillus jensenii GCA_001936235.1
cq
701 Lactobacillus gasseri ATCC 33323 = JCM 1131 GCA_000014425.1
629 Lactobacillus hominis DSM 23910 = CRBIP 24.179 GCA_000296835.1
473 Lactobacillus kalixensis DSM 16043 GCA_001434335.1
cr
523 Bacillus anthracis str. Ames GCA_000007845.1
523 Bacillus anthracis str. Sterne GCA_000008165.1
523 _5BBacillus thuringiensis_5D serovar konkukian str. 97-27 GCA_000008505.1
514 Bacillus cereus ATCC 14579 GCA_000007825.1
499 Bacillus thuringiensis YBT-1518 GCA_000497525.2
cs
701 Lactobacillus gasseri ATCC 33323 = JCM 1131 GCA_000014425.1
655 Lactobacillus hominis DSM 23910 = CRBIP 24.179 GCA_000296835.1
471 Lactobacillus iners DSM 13335 GCA_000160875.1
ct
683 Streptococcus mutans UA159 GCA_000007465.2
454 Streptococcus ratti FA-1 = DSM 20564 GCA_000286075.1

429 Streptococcus equinus GCA_000964315.1
cu
487 Rhodobacter sphaeroides 2.4.1 GCA_000012905.2
450 Rhodobacter sphaeroides ATCC 17025 GCA_000016405.1
379 Gemmobacter aquatilis GCA_900110025.1
cv
674 Rhodobacter sphaeroides 2.4.1 GCA_000012905.2
592 Rhodobacter sphaeroides ATCC 17025 GCA_000016405.1
560 Gemmobacter aquatilis GCA_900110025.1
cw
683 Staphylococcus epidermidis ATCC 12228 GCA_000007645.1
623 Staphylococcus capitis subsp. capitis GCA_001028645.1
542 Staphylococcus aureus subsp. aureus NCTC 8325 GCA_000013425.1
cx
677 Staphylococcus epidermidis ATCC 12228 GCA_000007645.1
653 Staphylococcus capitis subsp. capitis GCA_001028645.1
634 Staphylococcus haemolyticus JCSC1435 GCA_000009865.1
cy
785 Clostridium saccharoperbutylacetonicum N1-4_28HMT_29 GCA_000340885.1
784 Clostridium beijerinckii GCA_000833105.2
760 Clostridium saccharobutylicum DSM 13864 GCA_000473995.1
cz
574 Lactobacillus gasseri ATCC 33323 = JCM 1131 GCA_000014425.1
548 Lactobacillus hominis DSM 23910 = CRBIP 24.179 GCA_000296835.1
457 Lactobacillus iners DSM 13335 GCA_000160875.1
cA
697 Lactobacillus gasseri ATCC 33323 = JCM 1131 GCA_000014425.1
664 Lactobacillus hominis DSM 23910 = CRBIP 24.179 GCA_000296835.1
532 Lactobacillus iners DSM 13335 GCA_000160875.1
cB
770 Enterococcus faecalis V583 GCA_000007785.1
724 Streptomyces cinnamoneus GCA_001885705.1
672 Enterococcus rivorum GCA_001742285.1
cC
752 Streptococcus mutans UA159 GCA_000007465.2
587 Streptococcus ratti FA-1 = DSM 20564 GCA_000286075.1
510 Streptococcus equinus GCA_000964315.1
cD
598 Clostridium beijerinckii GCA_000833105.2
583 Clostridium saccharoperbutylacetonicum N1-4_28HMT_29 GCA_000340885.1
570 Clostridium puniceum GCA_002006345.1
cE
506 Escherichia coli O157_3AH7 str. Sakai GCA_000008865.1
506 Escherichia coli UMN026 GCA_000026325.2
506 Escherichia coli str. K-12 substr. MG1655 GCA_000005845.2
495 Escherichia coli O104_3AH4 str. 2011C-3493 GCA_000299455.1
495 Escherichia coli O83_3AH1 str. NRG 857C GCA_000183345.1
495 Shigella flexneri 2a str. 301 GCA_000006925.2
475 Escherichia coli IAI39 GCA_000026345.1
cF
716 Bacillus cereus ATCC 14579 GCA_000007825.1
702 Bacillus anthracis str. Ames GCA_000007845.1
702 Bacillus anthracis str. Sterne GCA_000008165.1
702 _5BBacillus thuringiensis_5D serovar konkukian str. 97-27 GCA_000008505.1
700 Bacillus thuringiensis YBT-1518 GCA_000497525.2
cG
650 Bifidobacterium adolescentis ATCC 15703 GCA_000010425.1
493 Bifidobacterium dentium JCM 1195 = DSM 20436 GCA_001042595.1
464 Bifidobacterium angulatum DSM 20098 = JCM 7096 GCA_001025155.1
cH
768 Clostridium beijerinckii GCA_000833105.2
755 Clostridium saccharoperbutylacetonicum N1-4_28HMT_29 GCA_000340885.1
727 Clostridium saccharobutylicum DSM 13864 GCA_000473995.1
cI
646 Streptococcus mutans UA159 GCA_000007465.2
506 Streptococcus ratti FA-1 = DSM 20564 GCA_000286075.1
471 Streptococcus equinus GCA_000964315.1
cJ
453 Staphylococcus epidermidis ATCC 12228 GCA_000007645.1
433 Staphylococcus capitis subsp. capitis GCA_001028645.1
420 Staphylococcus warneri SG1 GCA_000332735.1
cK
602 Enterococcus faecalis V583 GCA_000007785.1
555 Streptomyces cinnamoneus GCA_001885705.1
524 Enterococcus sulfureus ATCC 49903 GCA_000407605.1
cL

675 Rhodobacter sphaeroides 2.4.1 GCA_000012905.2
624 Rhodobacter sphaeroides ATCC 17025 GCA_000016405.1
490 Pseudorhodobacter wandonensis GCA_001202035.1
cM
701 Staphylococcus epidermidis ATCC 12228 GCA_000007645.1
657 Staphylococcus capitis subsp. capitis GCA_001028645.1
639 Staphylococcus lugdunensis HKU09-01 GCA_000025085.1
cN
762 Deinococcus radiodurans R1 GCA_000008565.1
474 Deinococcus gobiensis I-0 GCA_000252445.1
465 Deinococcus deserti VCD115 GCA_000020685.1
cO
658 Enterococcus faecalis V583 GCA_000007785.1
611 Streptomyces cinnamoneus GCA_001885705.1
579 Enterococcus hirae ATCC 9790 GCA_000271405.2
cP
726 Deinococcus radiodurans R1 GCA_000008565.1
475 Deinococcus deserti VCD115 GCA_000020685.1
442 Deinococcus gobiensis I-0 GCA_000252445.1
442 Deinococcus soli Cha et al. 2016 GCA_001007995.1
cQ
617 Clostridium saccharoperbutylacetonicum N1-4_28HMT_29 GCA_000340885.1
601 Clostridium beijerinckii GCA_000833105.2
586 Clostridium saccharobutylicum DSM 13864 GCA_000473995.1
cR
612 Rhodobacter sphaeroides 2.4.1 GCA_000012905.2
581 Rhodobacter sphaeroides ATCC 17025 GCA_000016405.1
473 Pseudorhodobacter wandonensis GCA_001202035.1
cS
569 Bifidobacterium adolescentis ATCC 15703 GCA_000010425.1
432 Bifidobacterium dentium JCM 1195 = DSM 20436 GCA_001042595.1
391 Bifidobacterium angulatum DSM 20098 = JCM 7096 GCA_001025155.1
cT
803 Clostridium beijerinckii GCA_000833105.2
787 Clostridium puniceum GCA_002006345.1
786 Clostridium saccharoperbutylacetonicum N1-4_28HMT_29 GCA_000340885.1
cU
621 Lactobacillus gasseri ATCC 33323 = JCM 1131 GCA_000014425.1
586 Lactobacillus hominis DSM 23910 = CRBIP 24.179 GCA_000296835.1
455 Lactobacillus iners DSM 13335 GCA_000160875.1
cV
359 Rhodobacter sphaeroides 2.4.1 GCA_000012905.2
324 Rhodobacter sphaeroides ATCC 17025 GCA_000016405.1
274 Pseudorhodobacter wandonensis GCA_001202035.1
cW
356 Bifidobacterium adolescentis ATCC 15703 GCA_000010425.1
301 Bifidobacterium dentium JCM 1195 = DSM 20436 GCA_001042595.1
289 Bifidobacterium callitrichos DSM 23973 GCA_000741175.1
cX
848 Lactobacillus gasseri ATCC 33323 = JCM 1131 GCA_000014425.1
777 Lactobacillus hominis DSM 23910 = CRBIP 24.179 GCA_000296835.1
610 Lactobacillus iners DSM 13335 GCA_000160875.1
cY
513 Rhodobacter sphaeroides 2.4.1 GCA_000012905.2
487 Rhodobacter sphaeroides ATCC 17025 GCA_000016405.1
361 Pseudorhodobacter wandonensis GCA_001202035.1
cZ
700 Deinococcus radiodurans R1 GCA_000008565.1
393 Deinococcus deserti VCD115 GCA_000020685.1
386 Deinococcus hopiensis KR-140 GCA_900176165.1
d0
752 Bacillus anthracis str. Ames GCA_000007845.1
752 Bacillus anthracis str. Sterne GCA_000008165.1
752 _5BBacillus thuringiensis_5D serovar konkukian str. 97-27 GCA_000008505.1
735 Bacillus pseudomycoides DSM 12442 GCA_000161455.1
733 Bacillus cereus ATCC 14579 GCA_000007825.1
d1
695 Rhodobacter sphaeroides 2.4.1 GCA_000012905.2
633 Rhodobacter sphaeroides ATCC 17025 GCA_000016405.1
539 Gemmobacter aquatilis GCA_900110025.1
d2
601 Staphylococcus epidermidis ATCC 12228 GCA_000007645.1
567 Staphylococcus capitis subsp. capitis GCA_001028645.1
536 Staphylococcus haemolyticus JCSC1435 GCA_000009865.1
d3
608 Streptococcus mutans UA159 GCA_000007465.2

431 Streptococcus ratti FA-1 = DSM 20564 GCA_000286075.1
387 Streptococcus ovis DSM 16829 GCA_000380125.1
d4
795 Staphylococcus epidermidis ATCC 12228 GCA_000007645.1
737 Staphylococcus capitis subsp. capitis GCA_001028645.1
710 Staphylococcus hominis subsp. hominis C80 GCA_000183685.1
d5
720 Clostridium beijerinckii GCA_000833105.2
691 Clostridium saccharoperbutylacetonicum N1-4_28HMT_29 GCA_000340885.1
676 Clostridium puniceum GCA_002006345.1
d6
788 Bacillus anthracis str. Ames GCA_000007845.1
788 Bacillus anthracis str. Sterne GCA_000008165.1
788 _5BBacillus thuringiensis_5D serovar konkukian str. 97-27 GCA_000008505.1
768 Bacillus cereus ATCC 14579 GCA_000007825.1
762 Bacillus thuringiensis YBT-1518 GCA_000497525.2
d7
756 Clostridium beijerinckii GCA_000833105.2
732 Clostridium saccharoperbutylacetonicum N1-4_28HMT_29 GCA_000340885.1
727 Clostridium saccharobutylicum DSM 13864 GCA_000473995.1
d8
694 Bifidobacterium adolescentis ATCC 15703 GCA_000010425.1
561 Bifidobacterium dentium JCM 1195 = DSM 20436 GCA_001042595.1
543 Bifidobacterium angulatum DSM 20098 = JCM 7096 GCA_001025155.1
d9
652 Enterococcus faecalis V583 GCA_000007785.1
605 Streptomyces cinnamoneus GCA_001885705.1
586 Enterococcus faecium DO GCA_000174395.2
da
651 Clostridium beijerinckii GCA_000833105.2
629 Clostridium saccharoperbutylacetonicum N1-4_28HMT_29 GCA_000340885.1
621 Clostridium saccharobutylicum DSM 13864 GCA_000473995.1
db
591 Rhodobacter sphaeroides 2.4.1 GCA_000012905.2
513 Rhodobacter sphaeroides ATCC 17025 GCA_000016405.1
375 Defluviimonas alba GCA_001620265.1
dc
667 Rhodobacter sphaeroides 2.4.1 GCA_000012905.2
575 Rhodobacter sphaeroides ATCC 17025 GCA_000016405.1
487 Defluviimonas alba GCA_001620265.1
dd
746 Rhodobacter sphaeroides 2.4.1 GCA_000012905.2
666 Rhodobacter sphaeroides ATCC 17025 GCA_000016405.1
591 Gemmobacter aquatilis GCA_900110025.1
de
624 Deinococcus radiodurans R1 GCA_000008565.1
290 Deinococcus deserti VCD115 GCA_000020685.1
289 Deinococcus soli Cha et al. 2016 GCA_001007995.1
df
941 Lactobacillus gasseri ATCC 33323 = JCM 1131 GCA_000014425.1
837 Lactobacillus hominis DSM 23910 = CRBIP 24.179 GCA_000296835.1
647 Lactobacillus iners DSM 13335 GCA_000160875.1
dg
935 Bacillus anthracis str. Ames GCA_000007845.1
935 Bacillus anthracis str. Sterne GCA_000008165.1
935 Bacillus cereus ATCC 14579 GCA_000007825.1
935 _5BBacillus thuringiensis_5D serovar konkukian str. 97-27 GCA_000008505.1
904 Bacillus mycoides GCA_000832605.1
904 Bacillus thuringiensis YBT-1518 GCA_000497525.2
887 Bacillus pseudomycoides DSM 12442 GCA_000161455.1
dh
603 Enterococcus faecalis V583 GCA_000007785.1
556 Streptomyces cinnamoneus GCA_001885705.1
547 Enterococcus asini ATCC 700915 GCA_000407365.1
di
411 Deinococcus radiodurans R1 GCA_000008565.1
273 Deinococcus gobiensis I-0 GCA_000252445.1
266 Deinococcus frigens DSM 12807 GCA_000701425.1
dj
407 Clostridium beijerinckii GCA_000833105.2
389 Clostridium puniceum GCA_002006345.1
367 Clostridium saccharoperbutylacetonicum N1-4_28HMT_29 GCA_000340885.1
dk
697 Deinococcus radiodurans R1 GCA_000008565.1
486 Deinococcus deserti VCD115 GCA_000020685.1
466 Deinococcus soli Cha et al. 2016 GCA_001007995.1

dl
823 Staphylococcus epidermidis ATCC 12228 GCA_000007645.1
776 Staphylococcus capitis subsp. capitis GCA_001028645.1
722 Staphylococcus warneri SG1 GCA_000332735.1
dm
621 Bacillus anthracis str. Ames GCA_000007845.1
621 Bacillus anthracis str. Sterne GCA_000008165.1
620 _5BBacillus thuringiensis_5D serovar konkukian str. 97-27 GCA_000008505.1
601 Bacillus cereus ATCC 14579 GCA_000007825.1
dn
249 Bifidobacterium adolescentis ATCC 15703 GCA_000010425.1
201 Bifidobacterium dentium JCM 1195 = DSM 20436 GCA_001042595.1
188 Bifidobacterium callitrichos DSM 23973 GCA_000741175.1
do
312 Clostridium beijerinckii GCA_000833105.2
311 Clostridium saccharoperbutylacetonicum N1-4_28HMT_29 GCA_000340885.1
305 Clostridium puniceum GCA_002006345.1
dp
445 Bifidobacterium adolescentis ATCC 15703 GCA_000010425.1
355 Bifidobacterium dentium JCM 1195 = DSM 20436 GCA_001042595.1
314 Bifidobacterium reuteri DSM 23975 GCA_000741695.1
dq
600 Clostridium saccharoperbutylacetonicum N1-4_28HMT_29 GCA_000340885.1
591 Clostridium beijerinckii GCA_000833105.2
579 Clostridium puniceum GCA_002006345.1
dr
673 Rhodobacter sphaeroides 2.4.1 GCA_000012905.2
614 Rhodobacter sphaeroides ATCC 17025 GCA_000016405.1
518 Gemmobacter megaterium GCA_900156815.1
ds
633 Rhodobacter sphaeroides 2.4.1 GCA_000012905.2
579 Rhodobacter sphaeroides ATCC 17025 GCA_000016405.1
473 Thioclava atlantica GCA_000737065.1
dt
647 Bifidobacterium adolescentis ATCC 15703 GCA_000010425.1
522 Bifidobacterium angulatum DSM 20098 = JCM 7096 GCA_001025155.1
521 Bifidobacterium dentium JCM 1195 = DSM 20436 GCA_001042595.1
du
632 Streptococcus mutans UA159 GCA_000007465.2
465 Streptococcus macacae NCTC 11558 GCA_000187995.3
460 Streptococcus ratti FA-1 = DSM 20564 GCA_000286075.1
dv
589 Staphylococcus capitis subsp. capitis GCA_001028645.1
587 Staphylococcus epidermidis ATCC 12228 GCA_000007645.1
579 Megasphaera cerevisiae DSM 20462 GCA_001045675.1
579 Staphylococcus warneri SG1 GCA_000332735.1
dw
471 Escherichia coli O157_3AH7 str. Sakai GCA_000008865.1
471 Escherichia coli str. K-12 substr. MG1655 GCA_000005845.2
471 Shigella flexneri 2a str. 301 GCA_000006925.2
469 Escherichia coli O83_3AH1 str. NRG 857C GCA_000183345.1
469 Escherichia coli UMN026 GCA_000026325.2
456 Shigella dysenteriae Sd197 GCA_000012005.1
dx
675 Escherichia coli IAI39 GCA_000026345.1
675 Escherichia coli O157_3AH7 str. Sakai GCA_000008865.1
675 Escherichia coli UMN026 GCA_000026325.2
675 Escherichia coli str. K-12 substr. MG1655 GCA_000005845.2
670 Escherichia coli O83_3AH1 str. NRG 857C GCA_000183345.1
662 Shigella flexneri 2a str. 301 GCA_000006925.2
dy
640 Staphylococcus epidermidis ATCC 12228 GCA_000007645.1
586 Staphylococcus capitis subsp. capitis GCA_001028645.1
546 Staphylococcus haemolyticus JCSC1435 GCA_000009865.1
dz
878 Rhodobacter sphaeroides 2.4.1 GCA_000012905.2
761 Rhodobacter sphaeroides ATCC 17025 GCA_000016405.1
625 Gemmobacter aquatilis GCA_900110025.1
dA
672 Lactobacillus gasseri ATCC 33323 = JCM 1131 GCA_000014425.1
582 Lactobacillus hominis DSM 23910 = CRBIP 24.179 GCA_000296835.1
420 Lactobacillus hamsteri DSM 5661 = JCM 6256 GCA_000615445.1
dB
648 Bifidobacterium adolescentis ATCC 15703 GCA_000010425.1
550 Bifidobacterium dentium JCM 1195 = DSM 20436 GCA_001042595.1
513 Bifidobacterium angulatum DSM 20098 = JCM 7096 GCA_001025155.1

dC
660 Clostridium beijerinckii GCA_000833105.2
634 Clostridium saccharoperbutylacetonicum N1-4_28HMT_29 GCA_000340885.1
600 Clostridium butyricum GCA_001456065.2
dD
817 Bifidobacterium adolescentis ATCC 15703 GCA_000010425.1
686 Bifidobacterium dentium JCM 1195 = DSM 20436 GCA_001042595.1
646 Bifidobacterium stellenboschense GCA_000741785.1
dE
433 Deinococcus radiodurans R1 GCA_000008565.1
228 Deinococcus gobiensis I-0 GCA_000252445.1
218 Deinococcus murrayi DSM 11303 GCA_000482805.1
dF
709 Clostridium beijerinckii GCA_000833105.2
636 Clostridium puniceum GCA_002006345.1
635 Clostridium saccharoperbutylacetonicum N1-4_28HMT_29 GCA_000340885.1
dG
844 Staphylococcus epidermidis ATCC 12228 GCA_000007645.1
790 Staphylococcus capitis subsp. capitis GCA_001028645.1
743 Staphylococcus aureus subsp. aureus NCTC 8325 GCA_000013425.1
dH
799 Streptococcus mutans UA159 GCA_000007465.2
605 Streptococcus ratti FA-1 = DSM 20564 GCA_000286075.1
568 Streptococcus gordonii str. Challis substr. CH1 GCA_000017005.1
dI
729 Clostridium beijerinckii GCA_000833105.2
682 Clostridium saccharoperbutylacetonicum N1-4_28HMT_29 GCA_000340885.1
671 Clostridium saccharobutylicum DSM 13864 GCA_000473995.1
dJ
610 Clostridium beijerinckii GCA_000833105.2
573 Clostridium saccharoperbutylacetonicum N1-4_28HMT_29 GCA_000340885.1
571 Clostridium saccharobutylicum DSM 13864 GCA_000473995.1
dK
801 Rhodobacter sphaeroides 2.4.1 GCA_000012905.2
730 Rhodobacter sphaeroides ATCC 17025 GCA_000016405.1
664 Pseudorhodobacter psychrotolerans GCA_001294535.1
dL
809 Lactobacillus gasseri ATCC 33323 = JCM 1131 GCA_000014425.1
713 Lactobacillus hominis DSM 23910 = CRBIP 24.179 GCA_000296835.1
566 Lactobacillus iners DSM 13335 GCA_000160875.1
dM
503 Clostridium beijerinckii GCA_000833105.2
501 Clostridium puniceum GCA_002006345.1
501 Clostridium saccharoperbutylacetonicum N1-4_28HMT_29 GCA_000340885.1
486 Clostridium saccharobutylicum DSM 13864 GCA_000473995.1
dN
776 Clostridium beijerinckii GCA_000833105.2
734 Clostridium saccharobutylicum DSM 13864 GCA_000473995.1
733 Clostridium saccharoperbutylacetonicum N1-4_28HMT_29 GCA_000340885.1
dO
768 Deinococcus radiodurans R1 GCA_000008565.1
431 Deinococcus deserti VCD115 GCA_000020685.1
427 Deinococcus gobiensis I-0 GCA_000252445.1
dP
674 Bacillus anthracis str. Ames GCA_000007845.1
674 Bacillus anthracis str. Sterne GCA_000008165.1
674 _5BBacillus thuringiensis_5D serovar konkukian str. 97-27 GCA_000008505.1
665 Bacillus cereus ATCC 14579 GCA_000007825.1
661 Bacillus pseudomycoides DSM 12442 GCA_000161455.1
dQ
942 Enterococcus faecalis V583 GCA_000007785.1
895 Streptomyces cinnamoneus GCA_001885705.1
808 Enterococcus rivorum GCA_001742285.1
dR
858 Clostridium beijerinckii GCA_000833105.2
824 Clostridium saccharoperbutylacetonicum N1-4_28HMT_29 GCA_000340885.1
762 Clostridium butyricum GCA_001456065.2
762 Clostridium puniceum GCA_002006345.1
dS
756 Bacillus cereus ATCC 14579 GCA_000007825.1
754 Bacillus anthracis str. Ames GCA_000007845.1
754 Bacillus anthracis str. Sterne GCA_000008165.1
754 _5BBacillus thuringiensis_5D serovar konkukian str. 97-27 GCA_000008505.1
746 Bacillus mycoides GCA_000832605.1
dT
807 Enterococcus faecalis V583 GCA_000007785.1

764 Streptomyces cinnamoneus GCA_001885705.1
705 Enterococcus faecium DO GCA_000174395.2
dU
421 Bifidobacterium adolescentis ATCC 15703 GCA_000010425.1
295 Bifidobacterium dentium JCM 1195 = DSM 20436 GCA_001042595.1
275 Bifidobacterium asteroides PRL2011 GCA_000304215.1
dV
698 Clostridium beijerinckii GCA_000833105.2
664 Clostridium saccharobutylicum DSM 13864 GCA_000473995.1
657 Clostridium saccharoperbutylacetonicum N1-4_28HMT_29 GCA_000340885.1
dW
602 Clostridium beijerinckii GCA_000833105.2
587 Clostridium saccharobutylicum DSM 13864 GCA_000473995.1
587 Clostridium saccharoperbutylacetonicum N1-4_28HMT_29 GCA_000340885.1
555 Clostridium chromiireducens GCA_002029255.1
dX
577 Lactobacillus gasseri ATCC 33323 = JCM 1131 GCA_000014425.1
540 Lactobacillus hominis DSM 23910 = CRBIP 24.179 GCA_000296835.1
420 Lactobacillus iners DSM 13335 GCA_000160875.1
dY
705 Clostridium beijerinckii GCA_000833105.2
682 Clostridium saccharoperbutylacetonicum N1-4_28HMT_29 GCA_000340885.1
652 Clostridium puniceum GCA_002006345.1
dZ
368 Staphylococcus epidermidis ATCC 12228 GCA_000007645.1
353 Staphylococcus condimenti GCA_001618885.1
340 Staphylococcus lugdunensis HKU09-01 GCA_000025085.1
e0
724 Clostridium beijerinckii GCA_000833105.2
707 Clostridium saccharoperbutylacetonicum N1-4_28HMT_29 GCA_000340885.1
661 Clostridium butyricum GCA_001456065.2
e1
752 Enterococcus faecalis V583 GCA_000007785.1
706 Streptomyces cinnamoneus GCA_001885705.1
642 Enterococcus faecium DO GCA_000174395.2
e2
550 Bifidobacterium adolescentis ATCC 15703 GCA_000010425.1
475 Bifidobacterium dentium JCM 1195 = DSM 20436 GCA_001042595.1
448 Bifidobacterium bifidum PRL2010 GCA_000165905.1
e3
713 Enterococcus faecalis V583 GCA_000007785.1
667 Streptomyces cinnamoneus GCA_001885705.1
617 Enterococcus asini ATCC 700915 GCA_000407365.1
e4
685 Clostridium beijerinckii GCA_000833105.2
645 Clostridium saccharoperbutylacetonicum N1-4_28HMT_29 GCA_000340885.1
640 Clostridium saccharobutylicum DSM 13864 GCA_000473995.1
e5
546 Streptococcus mutans UA159 GCA_000007465.2
347 Streptococcus equinus GCA_000964315.1
347 Streptococcus gordonii str. Challis substr. CH1 GCA_000017005.1
346 Streptococcus gallolyticus subsp. gallolyticus DSM 16831 GCA_002000985.1
e6
739 Staphylococcus epidermidis ATCC 12228 GCA_000007645.1
709 Staphylococcus capitis subsp. capitis GCA_001028645.1
678 Staphylococcus hominis subsp. hominis C80 GCA_000183685.1
e7
570 Enterococcus faecalis V583 GCA_000007785.1
524 Enterococcus thailandicus GCA_001652875.1
523 Enterococcus faecium DO GCA_000174395.2
523 Streptomyces cinnamoneus GCA_001885705.1
e8
230 Streptococcus mutans UA159 GCA_000007465.2
176 Streptococcus ferus DSM 20646 GCA_000372425.1
172 Streptococcus urinalis 2285-97 GCA_000188055.3
e9
876 Staphylococcus epidermidis ATCC 12228 GCA_000007645.1
828 Staphylococcus capitis subsp. capitis GCA_001028645.1
806 Staphylococcus aureus subsp. aureus NCTC 8325 GCA_000013425.1
ea
669 Deinococcus radiodurans R1 GCA_000008565.1
387 Deinococcus deserti VCD115 GCA_000020685.1
369 Deinococcus gobiensis I-0 GCA_000252445.1
eb
702 Enterococcus faecalis V583 GCA_000007785.1
656 Streptomyces cinnamoneus GCA_001885705.1

612 Enterococcus faecium DO GCA_000174395.2
ec
617 Clostridium beijerinckii GCA_000833105.2
609 Clostridium saccharoperbutylacetonicum N1-4_28HMT_29 GCA_000340885.1
600 Clostridium saccharobutylicum DSM 13864 GCA_000473995.1
ed
763 Clostridium beijerinckii GCA_000833105.2
738 Clostridium saccharoperbutylacetonicum N1-4_28HMT_29 GCA_000340885.1
696 Clostridium saccharobutylicum DSM 13864 GCA_000473995.1
ee
942 Lactobacillus gasseri ATCC 33323 = JCM 1131 GCA_000014425.1
852 Lactobacillus hominis DSM 23910 = CRBIP 24.179 GCA_000296835.1
639 Lactobacillus hamsteri DSM 5661 = JCM 6256 GCA_000615445.1
639 Lactobacillus iners DSM 13335 GCA_000160875.1
ef
706 Escherichia coli O157_3AH7 str. Sakai GCA_000008865.1
706 Escherichia coli UMN026 GCA_000026325.2
706 Escherichia coli str. K-12 substr. MG1655 GCA_000005845.2
702 Escherichia coli IAI39 GCA_000026345.1
702 Shigella flexneri 2a str. 301 GCA_000006925.2
701 Escherichia coli O83_3AH1 str. NRG 857C GCA_000183345.1
eg
402 Rhodobacter sphaeroides 2.4.1 GCA_000012905.2
342 Rhodobacter sphaeroides ATCC 17025 GCA_000016405.1
295 Defluviimonas alba GCA_001620265.1
eh
682 Deinococcus radiodurans R1 GCA_000008565.1
386 Deinococcus deserti VCD115 GCA_000020685.1
356 Deinococcus marmoris DSM 12784 GCA_000701405.1
ei
683 Streptococcus mutans UA159 GCA_000007465.2
479 Streptococcus ratti FA-1 = DSM 20564 GCA_000286075.1
449 Streptococcus sobrinus DSM 20742 = ATCC 33478 GCA_000686605.1
ej
643 Clostridium beijerinckii GCA_000833105.2
618 Clostridium saccharoperbutylacetonicum N1-4_28HMT_29 GCA_000340885.1
605 Clostridium saccharobutylicum DSM 13864 GCA_000473995.1
ek
420 Streptococcus mutans UA159 GCA_000007465.2
291 Streptococcus gordonii str. Challis substr. CH1 GCA_000017005.1
283 Streptococcus ratti FA-1 = DSM 20564 GCA_000286075.1
el
797 Staphylococcus epidermidis ATCC 12228 GCA_000007645.1
734 Staphylococcus capitis subsp. capitis GCA_001028645.1
711 Staphylococcus aureus subsp. aureus NCTC 8325 GCA_000013425.1
em
673 Clostridium beijerinckii GCA_000833105.2
643 Clostridium saccharoperbutylacetonicum N1-4_28HMT_29 GCA_000340885.1
639 Clostridium puniceum GCA_002006345.1
en
586 Clostridium beijerinckii GCA_000833105.2
551 Clostridium puniceum GCA_002006345.1
551 Clostridium saccharoperbutylacetonicum N1-4_28HMT_29 GCA_000340885.1
529 Clostridium saccharobutylicum DSM 13864 GCA_000473995.1
eo
907 Enterococcus faecalis V583 GCA_000007785.1
860 Streptomyces cinnamoneus GCA_001885705.1
836 Enterococcus rivorum GCA_001742285.1
ep
689 Staphylococcus epidermidis ATCC 12228 GCA_000007645.1
666 Staphylococcus capitis subsp. capitis GCA_001028645.1
629 Staphylococcus warneri SG1 GCA_000332735.1
eq
695 Staphylococcus epidermidis ATCC 12228 GCA_000007645.1
665 Staphylococcus capitis subsp. capitis GCA_001028645.1
603 Staphylococcus lugdunensis HKU09-01 GCA_000025085.1
er
608 Staphylococcus capitis subsp. capitis GCA_001028645.1
608 Staphylococcus epidermidis ATCC 12228 GCA_000007645.1
548 Staphylococcus haemolyticus JCSC1435 GCA_000009865.1
545 Staphylococcus aureus subsp. aureus NCTC 8325 GCA_000013425.1
es
668 Clostridium beijerinckii GCA_000833105.2
658 Clostridium saccharoperbutylacetonicum N1-4_28HMT_29 GCA_000340885.1
635 Clostridium puniceum GCA_002006345.1
et

704 Deinococcus radiodurans R1 GCA_000008565.1
444 Deinococcus deserti VCD115 GCA_000020685.1
404 Deinococcus hopiensis KR-140 GCA_900176165.1
eu
759 Lactobacillus gasseri ATCC 33323 = JCM 1131 GCA_000014425.1
662 Lactobacillus hominis DSM 23910 = CRBIP 24.179 GCA_000296835.1
478 Lactobacillus iners DSM 13335 GCA_000160875.1
ev
656 Lactobacillus gasseri ATCC 33323 = JCM 1131 GCA_000014425.1
580 Lactobacillus hominis DSM 23910 = CRBIP 24.179 GCA_000296835.1
454 Lactobacillus hamsteri DSM 5661 = JCM 6256 GCA_000615445.1
ew
294 Escherichia coli O104_3AH4 str. 2011C-3493 GCA_000299455.1
294 Escherichia coli str. K-12 substr. MG1655 GCA_000005845.2
294 Shigella flexneri 2a str. 301 GCA_000006925.2
269 Escherichia coli O157_3AH7 str. Sakai GCA_000008865.1
269 Escherichia coli O83_3AH1 str. NRG 857C GCA_000183345.1
269 Escherichia coli UMN026 GCA_000026325.2
233 Pluralibacter gergoviae GCA_000757785.1
ex
890 Staphylococcus epidermidis ATCC 12228 GCA_000007645.1
872 Staphylococcus capitis subsp. capitis GCA_001028645.1
829 Staphylococcus haemolyticus JCSC1435 GCA_000009865.1
ey
660 Deinococcus radiodurans R1 GCA_000008565.1
420 Deinococcus deserti VCD115 GCA_000020685.1
382 Deinococcus soli Cha et al. 2016 GCA_001007995.1
ez
567 Rhodobacter sphaeroides 2.4.1 GCA_000012905.2
535 Rhodobacter sphaeroides ATCC 17025 GCA_000016405.1
441 Defluviimonas alba GCA_001620265.1
eA
813 Rhodobacter sphaeroides 2.4.1 GCA_000012905.2
683 Rhodobacter sphaeroides ATCC 17025 GCA_000016405.1
548 Gemmobacter aquatilis GCA_900110025.1
eB
736 Lactobacillus gasseri ATCC 33323 = JCM 1131 GCA_000014425.1
652 Lactobacillus hominis DSM 23910 = CRBIP 24.179 GCA_000296835.1
462 Lactobacillus iners DSM 13335 GCA_000160875.1
eC
637 Bacillus anthracis str. Ames GCA_000007845.1
637 Bacillus anthracis str. Sterne GCA_000008165.1
637 Bacillus cereus ATCC 14579 GCA_000007825.1
637 _5BBacillus thuringiensis_5D serovar konkukian str. 97-27 GCA_000008505.1
622 Bacillus thuringiensis YBT-1518 GCA_000497525.2
605 Bacillus pseudomycoides DSM 12442 GCA_000161455.1
eD
785 Escherichia coli IAI39 GCA_000026345.1
785 Escherichia coli O83_3AH1 str. NRG 857C GCA_000183345.1
770 Escherichia coli O157_3AH7 str. Sakai GCA_000008865.1
770 Escherichia coli UMN026 GCA_000026325.2
770 Escherichia coli str. K-12 substr. MG1655 GCA_000005845.2
754 Shigella dysenteriae Sd197 GCA_000012005.1
eE
587 Rhodobacter sphaeroides 2.4.1 GCA_000012905.2
564 Rhodobacter sphaeroides ATCC 17025 GCA_000016405.1
485 Gemmobacter megaterium GCA_900156815.1
eF
419 Clostridium beijerinckii GCA_000833105.2
398 Clostridium saccharobutylicum DSM 13864 GCA_000473995.1
385 Clostridium saccharoperbutylacetonicum N1-4_28HMT_29 GCA_000340885.1
eG
734 Deinococcus radiodurans R1 GCA_000008565.1
424 Deinococcus deserti VCD115 GCA_000020685.1
420 Deinococcus gobiensis I-0 GCA_000252445.1
eH
703 Clostridium beijerinckii GCA_000833105.2
676 Clostridium saccharoperbutylacetonicum N1-4_28HMT_29 GCA_000340885.1
654 Clostridium saccharobutylicum DSM 13864 GCA_000473995.1
eI
540 Rhodobacter sphaeroides 2.4.1 GCA_000012905.2
459 Rhodobacter sphaeroides ATCC 17025 GCA_000016405.1
393 Defluviimonas alba GCA_001620265.1
eJ
474 Lactobacillus gasseri ATCC 33323 = JCM 1131 GCA_000014425.1
463 Lactobacillus hominis DSM 23910 = CRBIP 24.179 GCA_000296835.1

384 Lactobacillus iners DSM 13335 GCA_000160875.1
eK
480 Deinococcus radiodurans R1 GCA_000008565.1
322 Deinococcus proteolyticus MRP GCA_000190555.1
311 Deinococcus soli Cha et al. 2016 GCA_001007995.1
eL
485 Bifidobacterium adolescentis ATCC 15703 GCA_000010425.1
381 Bifidobacterium angulatum DSM 20098 = JCM 7096 GCA_001025155.1
378 Bifidobacterium dentium JCM 1195 = DSM 20436 GCA_001042595.1
eM
821 Clostridium beijerinckii GCA_000833105.2
790 Clostridium saccharoperbutylacetonicum N1-4_28HMT_29 GCA_000340885.1
753 Clostridium puniceum GCA_002006345.1
eN
899 Staphylococcus epidermidis ATCC 12228 GCA_000007645.1
844 Staphylococcus capitis subsp. capitis GCA_001028645.1
797 Megasphaera cerevisiae DSM 20462 GCA_001045675.1
797 Staphylococcus warneri SG1 GCA_000332735.1
eO
698 Clostridium beijerinckii GCA_000833105.2
655 Clostridium saccharoperbutylacetonicum N1-4_28HMT_29 GCA_000340885.1
644 Clostridium puniceum GCA_002006345.1
eP
703 Clostridium beijerinckii GCA_000833105.2
678 Clostridium saccharoperbutylacetonicum N1-4_28HMT_29 GCA_000340885.1
677 Clostridium saccharobutylicum DSM 13864 GCA_000473995.1
eQ
517 Rhodobacter sphaeroides 2.4.1 GCA_000012905.2
490 Rhodobacter sphaeroides ATCC 17025 GCA_000016405.1
441 Pseudorhodobacter ferrugineus DSM 5888 GCA_000420745.1
eR
879 Lactobacillus gasseri ATCC 33323 = JCM 1131 GCA_000014425.1
808 Lactobacillus hominis DSM 23910 = CRBIP 24.179 GCA_000296835.1
605 Lactobacillus hamsteri DSM 5661 = JCM 6256 GCA_000615445.1
eS
763 Lactobacillus gasseri ATCC 33323 = JCM 1131 GCA_000014425.1
696 Lactobacillus hominis DSM 23910 = CRBIP 24.179 GCA_000296835.1
508 Lactobacillus iners DSM 13335 GCA_000160875.1
eT
466 Streptococcus mutans UA159 GCA_000007465.2
353 Streptococcus sobrinus DSM 20742 = ATCC 33478 GCA_000686605.1
340 Streptococcus ratti FA-1 = DSM 20564 GCA_000286075.1
eU
613 Lactobacillus gasseri ATCC 33323 = JCM 1131 GCA_000014425.1
528 Lactobacillus hominis DSM 23910 = CRBIP 24.179 GCA_000296835.1
400 Lactobacillus hamsteri DSM 5661 = JCM 6256 GCA_000615445.1
eV
554 Bifidobacterium adolescentis ATCC 15703 GCA_000010425.1
455 Bifidobacterium dentium JCM 1195 = DSM 20436 GCA_001042595.1
435 Bifidobacterium angulatum DSM 20098 = JCM 7096 GCA_001025155.1
eW
541 Bifidobacterium adolescentis ATCC 15703 GCA_000010425.1
387 Bifidobacterium angulatum DSM 20098 = JCM 7096 GCA_001025155.1
385 Bifidobacterium callitrichos DSM 23973 GCA_000741175.1
eX
617 Streptococcus mutans UA159 GCA_000007465.2
413 Streptococcus ratti FA-1 = DSM 20564 GCA_000286075.1
408 Streptococcus ferus DSM 20646 GCA_000372425.1
eY
643 Streptococcus mutans UA159 GCA_000007465.2
547 Streptococcus ratti FA-1 = DSM 20564 GCA_000286075.1
499 Streptococcus gallolyticus subsp. gallolyticus DSM 16831 GCA_002000985.1
eZ
596 Lactobacillus gasseri ATCC 33323 = JCM 1131 GCA_000014425.1
523 Lactobacillus hominis DSM 23910 = CRBIP 24.179 GCA_000296835.1
408 Lactobacillus iners DSM 13335 GCA_000160875.1
f0
833 Lactobacillus gasseri ATCC 33323 = JCM 1131 GCA_000014425.1
765 Lactobacillus hominis DSM 23910 = CRBIP 24.179 GCA_000296835.1
558 Lactobacillus iners DSM 13335 GCA_000160875.1
558 Lactobacillus psittaci DSM 15354 GCA_000425905.1
f1
849 Staphylococcus epidermidis ATCC 12228 GCA_000007645.1
818 Staphylococcus capitis subsp. capitis GCA_001028645.1
795 Staphylococcus lugdunensis HKU09-01 GCA_000025085.1
f2

786 Enterococcus faecalis V583 GCA_000007785.1
739 Streptomyces cinnamoneus GCA_001885705.1
682 Enterococcus rivorum GCA_001742285.1
f3
760 Clostridium beijerinckii GCA_000833105.2
708 Clostridium saccharoperbutylacetonicum N1-4_28HMT_29 GCA_000340885.1
671 Clostridium puniceum GCA_002006345.1
f4
610 Escherichia coli str. K-12 substr. MG1655 GCA_000005845.2
599 Shigella flexneri 2a str. 301 GCA_000006925.2
590 Escherichia coli IAI39 GCA_000026345.1
f5
439 Clostridium beijerinckii GCA_000833105.2
375 Clostridium puniceum GCA_002006345.1
375 Clostridium saccharoperbutylacetonicum N1-4_28HMT_29 GCA_000340885.1
364 Clostridium saccharobutylicum DSM 13864 GCA_000473995.1
f6
657 Escherichia coli O157_3AH7 str. Sakai GCA_000008865.1
657 Escherichia coli str. K-12 substr. MG1655 GCA_000005845.2
656 Escherichia coli UMN026 GCA_000026325.2
642 Shigella flexneri 2a str. 301 GCA_000006925.2
f7
665 Clostridium beijerinckii GCA_000833105.2
640 Clostridium saccharoperbutylacetonicum N1-4_28HMT_29 GCA_000340885.1
636 Clostridium chromiireducens GCA_002029255.1
f8
936 Streptococcus mutans UA159 GCA_000007465.2
673 Streptococcus gordonii str. Challis substr. CH1 GCA_000017005.1
655 Streptococcus ratti FA-1 = DSM 20564 GCA_000286075.1
f9
823 Escherichia coli str. K-12 substr. MG1655 GCA_000005845.2
802 Escherichia coli O104_3AH4 str. 2011C-3493 GCA_000299455.1
791 Shigella flexneri 2a str. 301 GCA_000006925.2
fa
518 Deinococcus radiodurans R1 GCA_000008565.1
329 Deinococcus deserti VCD115 GCA_000020685.1
325 Deinococcus hopiensis KR-140 GCA_900176165.1
fb
804 Clostridium beijerinckii GCA_000833105.2
772 Clostridium saccharoperbutylacetonicum N1-4_28HMT_29 GCA_000340885.1
714 Clostridium puniceum GCA_002006345.1
fc
705 Bacillus anthracis str. Ames GCA_000007845.1
705 Bacillus anthracis str. Sterne GCA_000008165.1
705 _5BBacillus thuringiensis_5D serovar konkukian str. 97-27 GCA_000008505.1
694 Bacillus pseudomycoides DSM 12442 GCA_000161455.1
693 Bacillus cereus ATCC 14579 GCA_000007825.1
fd
614 Lactobacillus gasseri ATCC 33323 = JCM 1131 GCA_000014425.1
571 Lactobacillus hominis DSM 23910 = CRBIP 24.179 GCA_000296835.1
452 Lactobacillus iners DSM 13335 GCA_000160875.1
fe
469 Bacillus mycoides GCA_000832605.1
465 Bacillus cereus ATCC 14579 GCA_000007825.1
465 Bacillus thuringiensis YBT-1518 GCA_000497525.2
463 Bacillus anthracis str. Ames GCA_000007845.1
463 Bacillus anthracis str. Sterne GCA_000008165.1
463 _5BBacillus thuringiensis_5D serovar konkukian str. 97-27 GCA_000008505.1
ff
739 Bacillus anthracis str. Ames GCA_000007845.1
739 Bacillus anthracis str. Sterne GCA_000008165.1
739 Bacillus cereus ATCC 14579 GCA_000007825.1
739 Bacillus thuringiensis YBT-1518 GCA_000497525.2
739 _5BBacillus thuringiensis_5D serovar konkukian str. 97-27 GCA_000008505.1
716 Bacillus mycoides GCA_000832605.1
715 Bacillus pseudomycoides DSM 12442 GCA_000161455.1
fg
739 Streptococcus mutans UA159 GCA_000007465.2
528 Streptococcus ratti FA-1 = DSM 20564 GCA_000286075.1
479 Streptococcus gordonii str. Challis substr. CH1 GCA_000017005.1
fh
680 Deinococcus radiodurans R1 GCA_000008565.1
403 Deinococcus deserti VCD115 GCA_000020685.1
393 Deinococcus hopiensis KR-140 GCA_900176165.1
fi
689 Lactobacillus gasseri ATCC 33323 = JCM 1131 GCA_000014425.1

603 Lactobacillus hominis DSM 23910 = CRBIP 24.179 GCA_000296835.1
426 Lactobacillus iners DSM 13335 GCA_000160875.1
fj
563 Rhodobacter sphaeroides 2.4.1 GCA_000012905.2
511 Rhodobacter sphaeroides ATCC 17025 GCA_000016405.1
394 Defluviimonas alba GCA_001620265.1
394 Pseudorhodobacter wandonensis GCA_001202035.1
fk
693 Rhodobacter sphaeroides 2.4.1 GCA_000012905.2
620 Rhodobacter sphaeroides ATCC 17025 GCA_000016405.1
448 Defluviimonas alba GCA_001620265.1
fl
841 Enterococcus faecalis V583 GCA_000007785.1
794 Streptomyces cinnamoneus GCA_001885705.1
725 Enterococcus asini ATCC 700915 GCA_000407365.1
fm
540 Deinococcus radiodurans R1 GCA_000008565.1
348 Deinococcus deserti VCD115 GCA_000020685.1
317 Deinococcus soli Cha et al. 2016 GCA_001007995.1
fn
758 Staphylococcus epidermidis ATCC 12228 GCA_000007645.1
724 Staphylococcus capitis subsp. capitis GCA_001028645.1
677 Staphylococcus haemolyticus JCSC1435 GCA_000009865.1
fo
815 Staphylococcus epidermidis ATCC 12228 GCA_000007645.1
765 Staphylococcus capitis subsp. capitis GCA_001028645.1
702 Staphylococcus haemolyticus JCSC1435 GCA_000009865.1
fp
625 Streptococcus mutans UA159 GCA_000007465.2
469 Streptococcus pseudoporcinus LQ 940-04 GCA_000188035.3
467 Streptococcus iniae GCA_000831485.1
fq
657 Rhodobacter sphaeroides 2.4.1 GCA_000012905.2
588 Rhodobacter sphaeroides ATCC 17025 GCA_000016405.1
548 Gemmobacter aquatilis GCA_900110025.1
fr
528 Clostridium beijerinckii GCA_000833105.2
511 Clostridium saccharoperbutylacetonicum N1-4_28HMT_29 GCA_000340885.1
475 Clostridium saccharobutylicum DSM 13864 GCA_000473995.1
fs
825 Deinococcus radiodurans R1 GCA_000008565.1
535 Deinococcus deserti VCD115 GCA_000020685.1
505 Deinococcus gobiensis I-0 GCA_000252445.1
ft
691 Streptococcus mutans UA159 GCA_000007465.2
541 Streptococcus gordonii str. Challis substr. CH1 GCA_000017005.1
525 Streptococcus ratti FA-1 = DSM 20564 GCA_000286075.1
fu
527 Clostridium saccharoperbutylacetonicum N1-4_28HMT_29 GCA_000340885.1
523 Clostridium beijerinckii GCA_000833105.2
502 Clostridium puniceum GCA_002006345.1
fv
576 Staphylococcus epidermidis ATCC 12228 GCA_000007645.1
561 Staphylococcus capitis subsp. capitis GCA_001028645.1
546 Staphylococcus condimenti GCA_001618885.1
fw
962 Enterococcus faecalis V583 GCA_000007785.1
915 Streptomyces cinnamoneus GCA_001885705.1
853 Enterococcus massiliensis GCA_001050095.1
fx
736 Lactobacillus gasseri ATCC 33323 = JCM 1131 GCA_000014425.1
672 Lactobacillus hominis DSM 23910 = CRBIP 24.179 GCA_000296835.1
462 Lactobacillus iners DSM 13335 GCA_000160875.1
fy
862 Streptococcus mutans UA159 GCA_000007465.2
582 Streptococcus ratti FA-1 = DSM 20564 GCA_000286075.1
566 Streptococcus halotolerans GCA_001598035.1
fz
925 Staphylococcus epidermidis ATCC 12228 GCA_000007645.1
842 Staphylococcus capitis subsp. capitis GCA_001028645.1
771 Staphylococcus lugdunensis HKU09-01 GCA_000025085.1
fA
443 Bifidobacterium adolescentis ATCC 15703 GCA_000010425.1
385 Bifidobacterium dentium JCM 1195 = DSM 20436 GCA_001042595.1
355 Bifidobacterium angulatum DSM 20098 = JCM 7096 GCA_001025155.1
fB

795 Lactobacillus gasseri ATCC 33323 = JCM 1131 GCA_000014425.1
712 Lactobacillus hominis DSM 23910 = CRBIP 24.179 GCA_000296835.1
522 Lactobacillus iners DSM 13335 GCA_000160875.1
fC
443 Staphylococcus epidermidis ATCC 12228 GCA_000007645.1
409 Staphylococcus capitis subsp. capitis GCA_001028645.1
355 Staphylococcus warneri SG1 GCA_000332735.1
fD
715 Bacillus anthracis str. Ames GCA_000007845.1
715 Bacillus anthracis str. Sterne GCA_000008165.1
715 Bacillus cereus ATCC 14579 GCA_000007825.1
715 Bacillus thuringiensis YBT-1518 GCA_000497525.2
715 _5BBacillus thuringiensis_5D serovar konkukian str. 97-27 GCA_000008505.1
699 Bacillus pseudomycoides DSM 12442 GCA_000161455.1
697 Bacillus mycoides GCA_000832605.1
fE
635 Streptococcus mutans UA159 GCA_000007465.2
444 Streptococcus ratti FA-1 = DSM 20564 GCA_000286075.1
422 Streptococcus ferus DSM 20646 GCA_000372425.1
422 Streptococcus gordonii str. Challis substr. CH1 GCA_000017005.1
fF
620 Rhodobacter sphaeroides 2.4.1 GCA_000012905.2
566 Rhodobacter sphaeroides ATCC 17025 GCA_000016405.1
527 Gemmobacter megaterium GCA_900156815.1
fG
858 Shigella flexneri 2a str. 301 GCA_000006925.2
856 Escherichia coli O83_3AH1 str. NRG 857C GCA_000183345.1
847 Escherichia coli O157_3AH7 str. Sakai GCA_000008865.1
847 Escherichia coli UMN026 GCA_000026325.2
847 Escherichia coli str. K-12 substr. MG1655 GCA_000005845.2
fH
583 Streptococcus mutans UA159 GCA_000007465.2
405 Streptococcus ratti FA-1 = DSM 20564 GCA_000286075.1
356 Streptococcus ferus DSM 20646 GCA_000372425.1
fI
747 Streptococcus mutans UA159 GCA_000007465.2
586 Streptococcus ferus DSM 20646 GCA_000372425.1
583 Streptococcus ratti FA-1 = DSM 20564 GCA_000286075.1
fJ
676 Enterococcus faecalis V583 GCA_000007785.1
629 Streptomyces cinnamoneus GCA_001885705.1
556 Enterococcus faecium DO GCA_000174395.2
556 Enterococcus rivorum GCA_001742285.1
fK
673 Deinococcus radiodurans R1 GCA_000008565.1
390 Deinococcus gobiensis I-0 GCA_000252445.1
377 Deinococcus deserti VCD115 GCA_000020685.1
fL
766 Bifidobacterium adolescentis ATCC 15703 GCA_000010425.1
658 Bifidobacterium dentium JCM 1195 = DSM 20436 GCA_001042595.1
645 Bifidobacterium angulatum DSM 20098 = JCM 7096 GCA_001025155.1
fM
581 Staphylococcus epidermidis ATCC 12228 GCA_000007645.1
559 Staphylococcus capitis subsp. capitis GCA_001028645.1
542 Staphylococcus haemolyticus JCSC1435 GCA_000009865.1
fN
499 Clostridium beijerinckii GCA_000833105.2
498 Clostridium saccharobutylicum DSM 13864 GCA_000473995.1
489 Clostridium puniceum GCA_002006345.1
489 Clostridium saccharoperbutylacetonicum N1-4_28HMT_29 GCA_000340885.1
fO
562 Clostridium beijerinckii GCA_000833105.2
544 Clostridium saccharoperbutylacetonicum N1-4_28HMT_29 GCA_000340885.1
534 Clostridium puniceum GCA_002006345.1
fP
901 Deinococcus radiodurans R1 GCA_000008565.1
555 Deinococcus deserti VCD115 GCA_000020685.1
513 Deinococcus murrayi DSM 11303 GCA_000482805.1
fQ
652 Enterococcus faecalis V583 GCA_000007785.1
620 Streptomyces cinnamoneus GCA_001885705.1
577 Enterococcus rivorum GCA_001742285.1
fR
771 Staphylococcus epidermidis ATCC 12228 GCA_000007645.1
748 Staphylococcus capitis subsp. capitis GCA_001028645.1
703 Staphylococcus aureus subsp. aureus NCTC 8325 GCA_000013425.1

fS
614 Enterococcus faecalis V583 GCA_000007785.1
567 Streptomyces cinnamoneus GCA_001885705.1
509 Enterococcus rivorum GCA_001742285.1
fT
712 Enterococcus faecalis V583 GCA_000007785.1
666 Streptomyces cinnamoneus GCA_001885705.1
641 Enterococcus dispar ATCC 51266 GCA_000406945.1
fU
473 Bifidobacterium adolescentis ATCC 15703 GCA_000010425.1
390 Bifidobacterium callitrichos DSM 23973 GCA_000741175.1
388 Bifidobacterium angulatum DSM 20098 = JCM 7096 GCA_001025155.1
fV
693 Rhodobacter sphaeroides 2.4.1 GCA_000012905.2
599 Rhodobacter sphaeroides ATCC 17025 GCA_000016405.1
507 Pseudorhodobacter psychrotolerans GCA_001294535.1
fW
846 Rhodobacter sphaeroides 2.4.1 GCA_000012905.2
757 Rhodobacter sphaeroides ATCC 17025 GCA_000016405.1
662 Defluviimonas alba GCA_001620265.1
fX
810 Streptococcus mutans UA159 GCA_000007465.2
625 Streptococcus ratti FA-1 = DSM 20564 GCA_000286075.1
621 Streptococcus ferus DSM 20646 GCA_000372425.1
fY
647 Enterococcus faecalis V583 GCA_000007785.1
600 Streptomyces cinnamoneus GCA_001885705.1
545 Enterococcus rivorum GCA_001742285.1
fZ
615 Rhodobacter sphaeroides 2.4.1 GCA_000012905.2
513 Rhodobacter sphaeroides ATCC 17025 GCA_000016405.1
428 Gemmobacter megaterium GCA_900156815.1
g0
723 Streptococcus mutans UA159 GCA_000007465.2
576 Streptococcus gordonii str. Challis substr. CH1 GCA_000017005.1
549 Streptococcus salivarius GCA_000785515.1
549 Streptococcus thermophilus JIM 8232 GCA_000253395.1
g1
668 Lactobacillus gasseri ATCC 33323 = JCM 1131 GCA_000014425.1
635 Lactobacillus hominis DSM 23910 = CRBIP 24.179 GCA_000296835.1
470 Lactobacillus iners DSM 13335 GCA_000160875.1
g2
494 Lactobacillus gasseri ATCC 33323 = JCM 1131 GCA_000014425.1
446 Lactobacillus hominis DSM 23910 = CRBIP 24.179 GCA_000296835.1
355 Staphylococcus epidermidis ATCC 12228 GCA_000007645.1
g3
513 Bifidobacterium adolescentis ATCC 15703 GCA_000010425.1
393 Bifidobacterium dentium JCM 1195 = DSM 20436 GCA_001042595.1
342 Bifidobacterium animalis subsp. lactis DSM 10140 GCA_000022965.1
342 Bifidobacterium thermophilum RBL67 GCA_000347695.1
g4
720 Rhodobacter sphaeroides 2.4.1 GCA_000012905.2
622 Rhodobacter sphaeroides ATCC 17025 GCA_000016405.1
494 Gemmobacter aquatilis GCA_900110025.1
g5
666 Clostridium beijerinckii GCA_000833105.2
646 Clostridium saccharoperbutylacetonicum N1-4_28HMT_29 GCA_000340885.1
640 Clostridium puniceum GCA_002006345.1
g6
566 Rhodobacter sphaeroides 2.4.1 GCA_000012905.2
545 Rhodobacter sphaeroides ATCC 17025 GCA_000016405.1
466 Pseudorhodobacter psychrotolerans GCA_001294535.1
g7
449 Staphylococcus epidermidis ATCC 12228 GCA_000007645.1
400 Staphylococcus capitis subsp. capitis GCA_001028645.1
358 Staphylococcus lutrae GCA_002101335.1
g8
796 Deinococcus radiodurans R1 GCA_000008565.1
439 Deinococcus gobiensis I-0 GCA_000252445.1
429 Deinococcus deserti VCD115 GCA_000020685.1
g9
528 Clostridium beijerinckii GCA_000833105.2
495 Clostridium saccharoperbutylacetonicum N1-4_28HMT_29 GCA_000340885.1
481 Clostridium saccharobutylicum DSM 13864 GCA_000473995.1
ga
493 Rhodobacter sphaeroides 2.4.1 GCA_000012905.2

462 Rhodobacter sphaeroides ATCC 17025 GCA_000016405.1
393 Dinoroseobacter shibae DFL 12 = DSM 16493 GCA_000018145.1
gb
689 Lactobacillus gasseri ATCC 33323 = JCM 1131 GCA_000014425.1
659 Lactobacillus hominis DSM 23910 = CRBIP 24.179 GCA_000296835.1
513 Lactobacillus iners DSM 13335 GCA_000160875.1
gc
731 Staphylococcus epidermidis ATCC 12228 GCA_000007645.1
701 Staphylococcus capitis subsp. capitis GCA_001028645.1
651 Staphylococcus lugdunensis HKU09-01 GCA_000025085.1
gd
694 Streptococcus mutans UA159 GCA_000007465.2
522 Streptococcus ratti FA-1 = DSM 20564 GCA_000286075.1
510 Streptococcus gordonii str. Challis substr. CH1 GCA_000017005.1
ge
609 Deinococcus radiodurans R1 GCA_000008565.1
405 Deinococcus deserti VCD115 GCA_000020685.1
375 Deinococcus frigens DSM 12807 GCA_000701425.1
gf
663 Clostridium beijerinckii GCA_000833105.2
644 Clostridium saccharoperbutylacetonicum N1-4_28HMT_29 GCA_000340885.1
622 Clostridium puniceum GCA_002006345.1
gg
499 Lactobacillus gasseri ATCC 33323 = JCM 1131 GCA_000014425.1
471 Lactobacillus hominis DSM 23910 = CRBIP 24.179 GCA_000296835.1
416 Lactobacillus iners DSM 13335 GCA_000160875.1
gh
326 Bifidobacterium adolescentis ATCC 15703 GCA_000010425.1
294 Bifidobacterium longum NCC2705 GCA_000007525.1
275 Bifidobacterium subtile GCA_000741775.1
gi
963 Bacillus cereus ATCC 14579 GCA_000007825.1
961 Bacillus anthracis str. Ames GCA_000007845.1
961 Bacillus anthracis str. Sterne GCA_000008165.1
961 _5BBacillus thuringiensis_5D serovar konkukian str. 97-27 GCA_000008505.1
935 Bacillus thuringiensis YBT-1518 GCA_000497525.2
gj
916 Lactobacillus gasseri ATCC 33323 = JCM 1131 GCA_000014425.1
906 Lactobacillus hominis DSM 23910 = CRBIP 24.179 GCA_000296835.1
681 Lactobacillus jensenii GCA_001936235.1
gk
736 Enterococcus faecalis V583 GCA_000007785.1
689 Streptomyces cinnamoneus GCA_001885705.1
649 Enterococcus rivorum GCA_001742285.1
gl
520 Deinococcus radiodurans R1 GCA_000008565.1
274 Deinococcus deserti VCD115 GCA_000020685.1
256 Deinococcus gobiensis I-0 GCA_000252445.1
gm
545 Escherichia coli str. K-12 substr. MG1655 GCA_000005845.2
531 Escherichia coli O83_3AH1 str. NRG 857C GCA_000183345.1
527 Escherichia coli O157_3AH7 str. Sakai GCA_000008865.1
527 Escherichia coli UMN026 GCA_000026325.2
527 Shigella flexneri 2a str. 301 GCA_000006925.2
gn
639 Lactobacillus gasseri ATCC 33323 = JCM 1131 GCA_000014425.1
609 Lactobacillus hominis DSM 23910 = CRBIP 24.179 GCA_000296835.1
429 Lactobacillus iners DSM 13335 GCA_000160875.1
go
736 Staphylococcus epidermidis ATCC 12228 GCA_000007645.1
686 Staphylococcus capitis subsp. capitis GCA_001028645.1
644 Staphylococcus hominis subsp. hominis C80 GCA_000183685.1
gp
744 Bacillus anthracis str. Ames GCA_000007845.1
744 Bacillus anthracis str. Sterne GCA_000008165.1
744 _5BBacillus thuringiensis_5D serovar konkukian str. 97-27 GCA_000008505.1
728 Bacillus pseudomycoides DSM 12442 GCA_000161455.1
724 Bacillus cereus ATCC 14579 GCA_000007825.1
gq
720 Bacillus anthracis str. Ames GCA_000007845.1
720 Bacillus anthracis str. Sterne GCA_000008165.1
720 Bacillus cereus ATCC 14579 GCA_000007825.1
720 _5BBacillus thuringiensis_5D serovar konkukian str. 97-27 GCA_000008505.1
702 Bacillus thuringiensis YBT-1518 GCA_000497525.2
699 Bacillus mycoides GCA_000832605.1
gr

546 Rhodobacter sphaeroides 2.4.1 GCA_000012905.2
504 Rhodobacter sphaeroides ATCC 17025 GCA_000016405.1
396 Gemmobacter megaterium GCA_900156815.1
gs
683 Bifidobacterium adolescentis ATCC 15703 GCA_000010425.1
556 Bifidobacterium dentium JCM 1195 = DSM 20436 GCA_001042595.1
527 Bifidobacterium angulatum DSM 20098 = JCM 7096 GCA_001025155.1
527 Bifidobacterium stellenboschense GCA_000741785.1
gt
680 Rhodobacter sphaeroides 2.4.1 GCA_000012905.2
612 Rhodobacter sphaeroides ATCC 17025 GCA_000016405.1
532 Defluviimonas alba GCA_001620265.1
gu
504 Escherichia coli O157_3AH7 str. Sakai GCA_000008865.1
504 Escherichia coli str. K-12 substr. MG1655 GCA_000005845.2
496 Escherichia coli O104_3AH4 str. 2011C-3493 GCA_000299455.1
496 Shigella flexneri 2a str. 301 GCA_000006925.2
491 Escherichia coli UMN026 GCA_000026325.2
gv
952 Bacillus anthracis str. Ames GCA_000007845.1
952 Bacillus anthracis str. Sterne GCA_000008165.1
952 _5BBacillus thuringiensis_5D serovar konkukian str. 97-27 GCA_000008505.1
935 Bacillus cereus ATCC 14579 GCA_000007825.1
935 Bacillus pseudomycoides DSM 12442 GCA_000161455.1
909 Bacillus thuringiensis YBT-1518 GCA_000497525.2
gw
608 Streptococcus mutans UA159 GCA_000007465.2
455 Streptococcus ratti FA-1 = DSM 20564 GCA_000286075.1
445 Streptococcus ferus DSM 20646 GCA_000372425.1
gx
839 Deinococcus radiodurans R1 GCA_000008565.1
443 Deinococcus deserti VCD115 GCA_000020685.1
439 Deinococcus soli Cha et al. 2016 GCA_001007995.1
gy
492 Clostridium beijerinckii GCA_000833105.2
465 Clostridium saccharoperbutylacetonicum N1-4_28HMT_29 GCA_000340885.1
435 Clostridium butyricum GCA_001456065.2
gz
540 Clostridium beijerinckii GCA_000833105.2
522 Clostridium saccharobutylicum DSM 13864 GCA_000473995.1
504 Clostridium saccharoperbutylacetonicum N1-4_28HMT_29 GCA_000340885.1
gA
852 Bifidobacterium adolescentis ATCC 15703 GCA_000010425.1
715 Bifidobacterium dentium JCM 1195 = DSM 20436 GCA_001042595.1
695 Bifidobacterium angulatum DSM 20098 = JCM 7096 GCA_001025155.1
gB
604 Lactobacillus gasseri ATCC 33323 = JCM 1131 GCA_000014425.1
565 Lactobacillus hominis DSM 23910 = CRBIP 24.179 GCA_000296835.1
380 Lactobacillus iners DSM 13335 GCA_000160875.1
gC
690 Lactobacillus gasseri ATCC 33323 = JCM 1131 GCA_000014425.1
646 Lactobacillus hominis DSM 23910 = CRBIP 24.179 GCA_000296835.1
424 Lactobacillus iners DSM 13335 GCA_000160875.1
gD
478 Staphylococcus epidermidis ATCC 12228 GCA_000007645.1
458 Megasphaera cerevisiae DSM 20462 GCA_001045675.1
458 Staphylococcus warneri SG1 GCA_000332735.1
453 Staphylococcus capitis subsp. capitis GCA_001028645.1
gE
737 Lactobacillus gasseri ATCC 33323 = JCM 1131 GCA_000014425.1
651 Lactobacillus hominis DSM 23910 = CRBIP 24.179 GCA_000296835.1
510 Lactobacillus psittaci DSM 15354 GCA_000425905.1
gF
752 Deinococcus radiodurans R1 GCA_000008565.1
449 Deinococcus deserti VCD115 GCA_000020685.1
441 Deinococcus proteolyticus MRP GCA_000190555.1
gG
834 Escherichia coli str. K-12 substr. MG1655 GCA_000005845.2
833 Escherichia coli IAI39 GCA_000026345.1
833 Escherichia coli O157_3AH7 str. Sakai GCA_000008865.1
817 Escherichia coli UMN026 GCA_000026325.2
gH
836 Lactobacillus gasseri ATCC 33323 = JCM 1131 GCA_000014425.1
749 Lactobacillus hominis DSM 23910 = CRBIP 24.179 GCA_000296835.1
543 Lactobacillus iners DSM 13335 GCA_000160875.1
gI

934 Staphylococcus epidermidis ATCC 12228 GCA_000007645.1
852 Staphylococcus capitis subsp. capitis GCA_001028645.1
802 Staphylococcus hominis subsp. hominis C80 GCA_000183685.1
gJ
845 Bacillus anthracis str. Ames GCA_000007845.1
845 Bacillus anthracis str. Sterne GCA_000008165.1
845 _5BBacillus thuringiensis_5D serovar konkukian str. 97-27 GCA_000008505.1
830 Bacillus cereus ATCC 14579 GCA_000007825.1
822 Bacillus pseudomycoides DSM 12442 GCA_000161455.1
gK
578 Clostridium beijerinckii GCA_000833105.2
546 Clostridium saccharoperbutylacetonicum N1-4_28HMT_29 GCA_000340885.1
527 Clostridium butyricum GCA_001456065.2
527 Clostridium neonatale GCA_001458595.1
gL
821 Streptococcus mutans UA159 GCA_000007465.2
558 Streptococcus ratti FA-1 = DSM 20564 GCA_000286075.1
530 Streptococcus macacae NCTC 11558 GCA_000187995.3
gM
737 Enterococcus faecalis V583 GCA_000007785.1
690 Streptomyces cinnamoneus GCA_001885705.1
652 Enterococcus massiliensis GCA_001050095.1
gN
554 Staphylococcus epidermidis ATCC 12228 GCA_000007645.1
504 Staphylococcus capitis subsp. capitis GCA_001028645.1
488 Staphylococcus aureus subsp. aureus NCTC 8325 GCA_000013425.1
gO
690 Clostridium beijerinckii GCA_000833105.2
669 Clostridium saccharoperbutylacetonicum N1-4_28HMT_29 GCA_000340885.1
662 Clostridium saccharobutylicum DSM 13864 GCA_000473995.1
gP
809 Bacillus thuringiensis YBT-1518 GCA_000497525.2
800 Bacillus anthracis str. Ames GCA_000007845.1
800 Bacillus anthracis str. Sterne GCA_000008165.1
800 Bacillus cereus ATCC 14579 GCA_000007825.1
800 _5BBacillus thuringiensis_5D serovar konkukian str. 97-27 GCA_000008505.1
785 Bacillus pseudomycoides DSM 12442 GCA_000161455.1
gQ
802 Bifidobacterium adolescentis ATCC 15703 GCA_000010425.1
657 Bifidobacterium dentium JCM 1195 = DSM 20436 GCA_001042595.1
612 Bifidobacterium angulatum DSM 20098 = JCM 7096 GCA_001025155.1
gR
562 Lactobacillus gasseri ATCC 33323 = JCM 1131 GCA_000014425.1
478 Lactobacillus hominis DSM 23910 = CRBIP 24.179 GCA_000296835.1
301 Lactobacillus psittaci DSM 15354 GCA_000425905.1
gS
660 Rhodobacter sphaeroides 2.4.1 GCA_000012905.2
616 Rhodobacter sphaeroides ATCC 17025 GCA_000016405.1
495 Pseudorhodobacter psychrotolerans GCA_001294535.1
gT
756 Enterococcus faecalis V583 GCA_000007785.1
716 Streptomyces cinnamoneus GCA_001885705.1
668 Enterococcus canis NBRC 100695 GCA_001544375.1
gU
871 Clostridium beijerinckii GCA_000833105.2
848 Clostridium saccharoperbutylacetonicum N1-4_28HMT_29 GCA_000340885.1
844 Clostridium puniceum GCA_002006345.1
gV
639 Clostridium beijerinckii GCA_000833105.2
597 Clostridium saccharoperbutylacetonicum N1-4_28HMT_29 GCA_000340885.1
590 Clostridium puniceum GCA_002006345.1
gW
452 Clostridium beijerinckii GCA_000833105.2
428 Clostridium saccharoperbutylacetonicum N1-4_28HMT_29 GCA_000340885.1
423 Clostridium chromiireducens GCA_002029255.1
gX
872 Bacillus anthracis str. Ames GCA_000007845.1
872 Bacillus anthracis str. Sterne GCA_000008165.1
872 _5BBacillus thuringiensis_5D serovar konkukian str. 97-27 GCA_000008505.1
871 Bacillus cereus ATCC 14579 GCA_000007825.1
871 Bacillus thuringiensis YBT-1518 GCA_000497525.2
843 Bacillus pseudomycoides DSM 12442 GCA_000161455.1
gY
721 Streptococcus mutans UA159 GCA_000007465.2
545 Streptococcus ratti FA-1 = DSM 20564 GCA_000286075.1
489 Streptococcus gordonii str. Challis substr. CH1 GCA_000017005.1

gZ
602 Deinococcus radiodurans R1 GCA_000008565.1
394 Deinococcus deserti VCD115 GCA_000020685.1
367 Deinococcus hopiensis KR-140 GCA_900176165.1
h0
475 Streptococcus mutans UA159 GCA_000007465.2
355 Streptococcus ratti FA-1 = DSM 20564 GCA_000286075.1
340 Streptococcus macacae NCTC 11558 GCA_000187995.3
h1
542 Streptococcus mutans UA159 GCA_000007465.2
401 Streptococcus gordonii str. Challis substr. CH1 GCA_000017005.1
388 Streptococcus salivarius GCA_000785515.1
h2
502 Deinococcus radiodurans R1 GCA_000008565.1
341 Deinococcus deserti VCD115 GCA_000020685.1
341 Deinococcus puniceus GCA_001644565.1
339 Deinococcus murrayi DSM 11303 GCA_000482805.1
339 Deinococcus phoenicis GCA_000599865.1
h3
845 Bacillus anthracis str. Ames GCA_000007845.1
845 Bacillus anthracis str. Sterne GCA_000008165.1
845 _5BBacillus thuringiensis_5D serovar konkukian str. 97-27 GCA_000008505.1
835 Bacillus cereus ATCC 14579 GCA_000007825.1
810 Bacillus thuringiensis YBT-1518 GCA_000497525.2
h4
739 Deinococcus radiodurans R1 GCA_000008565.1
535 Deinococcus puniceus GCA_001644565.1
528 Deinococcus gobiensis I-0 GCA_000252445.1
h5
540 Streptococcus mutans UA159 GCA_000007465.2
418 Streptococcus ratti FA-1 = DSM 20564 GCA_000286075.1
409 Streptococcus macacae NCTC 11558 GCA_000187995.3
h6
979 _5BBacillus thuringiensis_5D serovar konkukian str. 97-27 GCA_000008505.1
963 Bacillus anthracis str. Ames GCA_000007845.1
963 Bacillus anthracis str. Sterne GCA_000008165.1
943 Bacillus cereus ATCC 14579 GCA_000007825.1
943 Bacillus thuringiensis YBT-1518 GCA_000497525.2
h7
797 Lactobacillus gasseri ATCC 33323 = JCM 1131 GCA_000014425.1
761 Lactobacillus hominis DSM 23910 = CRBIP 24.179 GCA_000296835.1
579 Lactobacillus psittaci DSM 15354 GCA_000425905.1
h8
573 Rhodobacter sphaeroides 2.4.1 GCA_000012905.2
558 Rhodobacter sphaeroides ATCC 17025 GCA_000016405.1
513 Gemmobacter aquatilis GCA_900110025.1
h9
562 Bacillus thuringiensis YBT-1518 GCA_000497525.2
542 Bacillus anthracis str. Ames GCA_000007845.1
542 Bacillus anthracis str. Sterne GCA_000008165.1
542 _5BBacillus thuringiensis_5D serovar konkukian str. 97-27 GCA_000008505.1
541 Bacillus cereus ATCC 14579 GCA_000007825.1
ha
401 Bifidobacterium adolescentis ATCC 15703 GCA_000010425.1
308 Bifidobacterium dentium JCM 1195 = DSM 20436 GCA_001042595.1
293 Bifidobacterium stellenboschense GCA_000741785.1
hb
973 Enterococcus faecalis V583 GCA_000007785.1
926 Streptomyces cinnamoneus GCA_001885705.1
842 Enterococcus saccharolyticus subsp. saccharolyticus ATCC 43076 GCA_000407285.1
hc
714 Escherichia coli O104_3AH4 str. 2011C-3493 GCA_000299455.1
714 Escherichia coli str. K-12 substr. MG1655 GCA_000005845.2
707 Shigella flexneri 2a str. 301 GCA_000006925.2
685 Escherichia coli O157_3AH7 str. Sakai GCA_000008865.1
hd
561 Rhodobacter sphaeroides 2.4.1 GCA_000012905.2
506 Rhodobacter sphaeroides ATCC 17025 GCA_000016405.1
480 Gemmobacter aquatilis GCA_900110025.1
he
572 Deinococcus radiodurans R1 GCA_000008565.1
353 Deinococcus gobiensis I-0 GCA_000252445.1
329 Deinococcus deserti VCD115 GCA_000020685.1
hf
695 Enterococcus faecalis V583 GCA_000007785.1
648 Streptomyces cinnamoneus GCA_001885705.1

634 Enterococcus rivorum GCA_001742285.1
hg
615 Clostridium beijerinckii GCA_000833105.2
578 Clostridium saccharoperbutylacetonicum N1-4_28HMT_29 GCA_000340885.1
564 Clostridium puniceum GCA_002006345.1
hh
631 Lactobacillus gasseri ATCC 33323 = JCM 1131 GCA_000014425.1
611 Lactobacillus hominis DSM 23910 = CRBIP 24.179 GCA_000296835.1
432 Lactobacillus iners DSM 13335 GCA_000160875.1
hi
828 Staphylococcus epidermidis ATCC 12228 GCA_000007645.1
753 Staphylococcus capitis subsp. capitis GCA_001028645.1
713 Staphylococcus warneri SG1 GCA_000332735.1
hj
684 Staphylococcus epidermidis ATCC 12228 GCA_000007645.1
643 Staphylococcus capitis subsp. capitis GCA_001028645.1
623 Staphylococcus aureus subsp. aureus NCTC 8325 GCA_000013425.1
hk
915 Staphylococcus epidermidis ATCC 12228 GCA_000007645.1
852 Staphylococcus capitis subsp. capitis GCA_001028645.1
798 Staphylococcus aureus subsp. aureus NCTC 8325 GCA_000013425.1
hl
698 Clostridium beijerinckii GCA_000833105.2
685 Clostridium saccharoperbutylacetonicum N1-4_28HMT_29 GCA_000340885.1
675 Clostridium saccharobutylicum DSM 13864 GCA_000473995.1
hm
678 Rhodobacter sphaeroides 2.4.1 GCA_000012905.2
586 Rhodobacter sphaeroides ATCC 17025 GCA_000016405.1
509 Jannaschia donghaensis GCA_001403795.1
hn
656 Rhodobacter sphaeroides 2.4.1 GCA_000012905.2
562 Rhodobacter sphaeroides ATCC 17025 GCA_000016405.1
500 Pseudorhodobacter psychrotolerans GCA_001294535.1
ho
773 Clostridium beijerinckii GCA_000833105.2
751 Clostridium saccharoperbutylacetonicum N1-4_28HMT_29 GCA_000340885.1
722 Clostridium saccharobutylicum DSM 13864 GCA_000473995.1
hp
631 Staphylococcus epidermidis ATCC 12228 GCA_000007645.1
606 Staphylococcus capitis subsp. capitis GCA_001028645.1
543 Staphylococcus lugdunensis HKU09-01 GCA_000025085.1
hq
709 Bifidobacterium adolescentis ATCC 15703 GCA_000010425.1
569 Bifidobacterium dentium JCM 1195 = DSM 20436 GCA_001042595.1
543 Bifidobacterium angulatum DSM 20098 = JCM 7096 GCA_001025155.1
hr
769 Deinococcus radiodurans R1 GCA_000008565.1
490 Deinococcus deserti VCD115 GCA_000020685.1
465 Deinococcus gobiensis I-0 GCA_000252445.1
hs
624 Lactobacillus gasseri ATCC 33323 = JCM 1131 GCA_000014425.1
596 Lactobacillus hominis DSM 23910 = CRBIP 24.179 GCA_000296835.1
447 Lactobacillus hamsteri DSM 5661 = JCM 6256 GCA_000615445.1
ht
743 Clostridium beijerinckii GCA_000833105.2
718 Clostridium saccharoperbutylacetonicum N1-4_28HMT_29 GCA_000340885.1
692 Clostridium saccharobutylicum DSM 13864 GCA_000473995.1
hu
543 Escherichia coli str. K-12 substr. MG1655 GCA_000005845.2
525 Escherichia coli O157_3AH7 str. Sakai GCA_000008865.1
525 Shigella flexneri 2a str. 301 GCA_000006925.2
521 Escherichia coli O104_3AH4 str. 2011C-3493 GCA_000299455.1
hv
572 Deinococcus radiodurans R1 GCA_000008565.1
374 Deinococcus deserti VCD115 GCA_000020685.1
374 Deinococcus hopiensis KR-140 GCA_900176165.1
364 Deinococcus puniceus GCA_001644565.1
hw
892 Streptococcus mutans UA159 GCA_000007465.2
660 Streptococcus ratti FA-1 = DSM 20564 GCA_000286075.1
592 Streptococcus sobrinus DSM 20742 = ATCC 33478 GCA_000686605.1
hx
659 Staphylococcus epidermidis ATCC 12228 GCA_000007645.1
609 Staphylococcus capitis subsp. capitis GCA_001028645.1
590 Staphylococcus aureus subsp. aureus NCTC 8325 GCA_000013425.1
hy

907 Deinococcus radiodurans R1 GCA_000008565.1
541 Deinococcus deserti VCD115 GCA_000020685.1
517 Deinococcus gobiensis I-0 GCA_000252445.1
hz
697 Lactobacillus gasseri ATCC 33323 = JCM 1131 GCA_000014425.1
651 Lactobacillus hominis DSM 23910 = CRBIP 24.179 GCA_000296835.1
461 Lactobacillus iners DSM 13335 GCA_000160875.1
hA
396 Rhodobacter sphaeroides 2.4.1 GCA_000012905.2
355 Rhodobacter sphaeroides ATCC 17025 GCA_000016405.1
316 Defluviimonas alba GCA_001620265.1
hB
449 Escherichia coli str. K-12 substr. MG1655 GCA_000005845.2
433 Escherichia coli O104_3AH4 str. 2011C-3493 GCA_000299455.1
433 Escherichia coli O157_3AH7 str. Sakai GCA_000008865.1
433 Shigella flexneri 2a str. 301 GCA_000006925.2
432 Escherichia coli O83_3AH1 str. NRG 857C GCA_000183345.1
432 Escherichia coli UMN026 GCA_000026325.2
hC
508 Rhodobacter sphaeroides 2.4.1 GCA_000012905.2
442 Rhodobacter sphaeroides ATCC 17025 GCA_000016405.1
428 Gemmobacter aquatilis GCA_900110025.1
hD
659 Enterococcus faecalis V583 GCA_000007785.1
632 Enterococcus rivorum GCA_001742285.1
620 Enterococcus massiliensis GCA_001050095.1
hE
691 Clostridium beijerinckii GCA_000833105.2
681 Clostridium saccharoperbutylacetonicum N1-4_28HMT_29 GCA_000340885.1
664 Clostridium saccharobutylicum DSM 13864 GCA_000473995.1
hF
644 Staphylococcus epidermidis ATCC 12228 GCA_000007645.1
621 Megasphaera cerevisiae DSM 20462 GCA_001045675.1
621 Staphylococcus warneri SG1 GCA_000332735.1
604 Staphylococcus capitis subsp. capitis GCA_001028645.1
hG
895 Deinococcus radiodurans R1 GCA_000008565.1
605 Deinococcus deserti VCD115 GCA_000020685.1
563 Deinococcus murrayi DSM 11303 GCA_000482805.1
hH
713 Staphylococcus epidermidis ATCC 12228 GCA_000007645.1
696 Staphylococcus capitis subsp. capitis GCA_001028645.1
651 Staphylococcus warneri SG1 GCA_000332735.1
hI
747 Staphylococcus epidermidis ATCC 12228 GCA_000007645.1
712 Staphylococcus capitis subsp. capitis GCA_001028645.1
654 Staphylococcus warneri SG1 GCA_000332735.1
hJ
642 Clostridium beijerinckii GCA_000833105.2
605 Clostridium saccharoperbutylacetonicum N1-4_28HMT_29 GCA_000340885.1
592 Clostridium puniceum GCA_002006345.1
hK
633 Deinococcus radiodurans R1 GCA_000008565.1
413 Deinococcus deserti VCD115 GCA_000020685.1
409 Deinococcus hopiensis KR-140 GCA_900176165.1
hL
728 Lactobacillus gasseri ATCC 33323 = JCM 1131 GCA_000014425.1
676 Lactobacillus hominis DSM 23910 = CRBIP 24.179 GCA_000296835.1
462 Lactobacillus acetotolerans GCA_001042405.1
462 Lactobacillus iners DSM 13335 GCA_000160875.1
hM
297 Rhodobacter sphaeroides 2.4.1 GCA_000012905.2
286 Rhodobacter sphaeroides ATCC 17025 GCA_000016405.1
210 Gemmobacter aquatilis GCA_900110025.1
hN
508 Escherichia coli IAI39 GCA_000026345.1
508 Escherichia coli O104_3AH4 str. 2011C-3493 GCA_000299455.1
508 Escherichia coli UMN026 GCA_000026325.2
508 Escherichia coli str. K-12 substr. MG1655 GCA_000005845.2
508 Shigella flexneri 2a str. 301 GCA_000006925.2
503 Escherichia coli O157_3AH7 str. Sakai GCA_000008865.1
494 Escherichia coli O83_3AH1 str. NRG 857C GCA_000183345.1
hO
658 Bacillus anthracis str. Ames GCA_000007845.1
658 Bacillus anthracis str. Sterne GCA_000008165.1
658 _5BBacillus thuringiensis_5D serovar konkukian str. 97-27 GCA_000008505.1

643 Bacillus cereus ATCC 14579 GCA_000007825.1
633 Bacillus thuringiensis YBT-1518 GCA_000497525.2
hP
871 Rhodobacter sphaeroides 2.4.1 GCA_000012905.2
836 Rhodobacter sphaeroides ATCC 17025 GCA_000016405.1
723 Gemmobacter aquatilis GCA_900110025.1
hQ
706 Streptococcus mutans UA159 GCA_000007465.2
517 Streptococcus ratti FA-1 = DSM 20564 GCA_000286075.1
487 Streptococcus gordonii str. Challis substr. CH1 GCA_000017005.1
hR
754 Rhodobacter sphaeroides 2.4.1 GCA_000012905.2
710 Rhodobacter sphaeroides ATCC 17025 GCA_000016405.1
617 Pseudorhodobacter psychrotolerans GCA_001294535.1
hS
838 Escherichia coli str. K-12 substr. MG1655 GCA_000005845.2
828 Escherichia coli O104_3AH4 str. 2011C-3493 GCA_000299455.1
812 Shigella flexneri 2a str. 301 GCA_000006925.2
hT
852 Deinococcus radiodurans R1 GCA_000008565.1
531 Deinococcus deserti VCD115 GCA_000020685.1
522 Deinococcus gobiensis I-0 GCA_000252445.1
hU
656 Lactobacillus gasseri ATCC 33323 = JCM 1131 GCA_000014425.1
636 Lactobacillus hominis DSM 23910 = CRBIP 24.179 GCA_000296835.1
482 Lactobacillus hamsteri DSM 5661 = JCM 6256 GCA_000615445.1
hV
870 Bifidobacterium adolescentis ATCC 15703 GCA_000010425.1
729 Bifidobacterium dentium JCM 1195 = DSM 20436 GCA_001042595.1
708 Bifidobacterium angulatum DSM 20098 = JCM 7096 GCA_001025155.1
hW
759 Enterococcus faecalis V583 GCA_000007785.1
716 Streptomyces cinnamoneus GCA_001885705.1
648 Enterococcus asini ATCC 700915 GCA_000407365.1
hX
758 Bifidobacterium adolescentis ATCC 15703 GCA_000010425.1
612 Bifidobacterium dentium JCM 1195 = DSM 20436 GCA_001042595.1
574 Bifidobacterium angulatum DSM 20098 = JCM 7096 GCA_001025155.1
hY
687 Streptococcus mutans UA159 GCA_000007465.2
494 Streptococcus ratti FA-1 = DSM 20564 GCA_000286075.1
485 Streptococcus macacae NCTC 11558 GCA_000187995.3
hZ
664 Deinococcus radiodurans R1 GCA_000008565.1
367 Deinococcus deserti VCD115 GCA_000020685.1
310 Deinococcus soli Cha et al. 2016 GCA_001007995.1
i0
513 Rhodobacter sphaeroides 2.4.1 GCA_000012905.2
462 Rhodobacter sphaeroides ATCC 17025 GCA_000016405.1
369 Defluviimonas alba GCA_001620265.1
i1
598 Rhodobacter sphaeroides 2.4.1 GCA_000012905.2
493 Rhodobacter sphaeroides ATCC 17025 GCA_000016405.1
438 Gemmobacter megaterium GCA_900156815.1
i2
881 Deinococcus radiodurans R1 GCA_000008565.1
542 Deinococcus deserti VCD115 GCA_000020685.1
537 Deinococcus gobiensis I-0 GCA_000252445.1
i3
750 Clostridium beijerinckii GCA_000833105.2
685 Clostridium saccharoperbutylacetonicum N1-4_28HMT_29 GCA_000340885.1
664 Clostridium puniceum GCA_002006345.1
i4
619 Rhodobacter sphaeroides 2.4.1 GCA_000012905.2
543 Rhodobacter sphaeroides ATCC 17025 GCA_000016405.1
447 Pseudorhodobacter wandonensis GCA_001202035.1
i5
466 Streptococcus mutans UA159 GCA_000007465.2
324 Streptococcus gordonii str. Challis substr. CH1 GCA_000017005.1
313 Streptococcus equinus GCA_000964315.1
i6
656 Clostridium beijerinckii GCA_000833105.2
655 Clostridium saccharoperbutylacetonicum N1-4_28HMT_29 GCA_000340885.1
634 Clostridium saccharobutylicum DSM 13864 GCA_000473995.1
i7
902 Streptococcus mutans UA159 GCA_000007465.2

647 Streptococcus ratti FA-1 = DSM 20564 GCA_000286075.1
616 Streptococcus gordonii str. Challis substr. CH1 GCA_000017005.1
i8
595 Bifidobacterium adolescentis ATCC 15703 GCA_000010425.1
540 Bifidobacterium dentium JCM 1195 = DSM 20436 GCA_001042595.1
511 Bifidobacterium angulatum DSM 20098 = JCM 7096 GCA_001025155.1
i9
653 Bifidobacterium adolescentis ATCC 15703 GCA_000010425.1
556 Bifidobacterium angulatum DSM 20098 = JCM 7096 GCA_001025155.1
543 Bifidobacterium dentium JCM 1195 = DSM 20436 GCA_001042595.1
ia
466 Rhodobacter sphaeroides 2.4.1 GCA_000012905.2
442 Rhodobacter sphaeroides ATCC 17025 GCA_000016405.1
384 Pseudorhodobacter psychrotolerans GCA_001294535.1
ib
439 Rhodobacter sphaeroides 2.4.1 GCA_000012905.2
392 Rhodobacter sphaeroides ATCC 17025 GCA_000016405.1
346 Thioclava indica GCA_000714545.1
ic
194 Deinococcus radiodurans R1 GCA_000008565.1
97 Deinococcus gobiensis I-0 GCA_000252445.1
85 Deinococcus deserti VCD115 GCA_000020685.1
85 Deinococcus hopiensis KR-140 GCA_900176165.1
85 Deinococcus indicus GCA_002198095.1
85 Deinococcus soli Cha et al. 2016 GCA_001007995.1
id
562 Clostridium beijerinckii GCA_000833105.2
556 Clostridium saccharoperbutylacetonicum N1-4_28HMT_29 GCA_000340885.1
534 Bacillus anthracis str. Ames GCA_000007845.1
534 Bacillus anthracis str. Sterne GCA_000008165.1
534 Clostridium saccharobutylicum DSM 13864 GCA_000473995.1
534 _5BBacillus thuringiensis_5D serovar konkukian str. 97-27 GCA_000008505.1
ie
422 Bifidobacterium adolescentis ATCC 15703 GCA_000010425.1
360 Bifidobacterium angulatum DSM 20098 = JCM 7096 GCA_001025155.1
333 Bifidobacterium dentium JCM 1195 = DSM 20436 GCA_001042595.1
if
607 Escherichia coli O104_3AH4 str. 2011C-3493 GCA_000299455.1
607 Escherichia coli str. K-12 substr. MG1655 GCA_000005845.2
607 Shigella flexneri 2a str. 301 GCA_000006925.2
606 Escherichia coli UMN026 GCA_000026325.2
599 Escherichia coli O83_3AH1 str. NRG 857C GCA_000183345.1
ig
644 Staphylococcus epidermidis ATCC 12228 GCA_000007645.1
580 Megasphaera cerevisiae DSM 20462 GCA_001045675.1
580 Staphylococcus capitis subsp. capitis GCA_001028645.1
580 Staphylococcus warneri SG1 GCA_000332735.1
553 Staphylococcus aureus subsp. aureus NCTC 8325 GCA_000013425.1
ih
846 Staphylococcus epidermidis ATCC 12228 GCA_000007645.1
794 Staphylococcus capitis subsp. capitis GCA_001028645.1
737 Staphylococcus hominis subsp. hominis C80 GCA_000183685.1
737 Staphylococcus lugdunensis HKU09-01 GCA_000025085.1
ii
620 Streptococcus mutans UA159 GCA_000007465.2
471 Streptococcus ratti FA-1 = DSM 20564 GCA_000286075.1
432 Streptococcus criceti HS-6 GCA_000187975.3
ij
842 Streptococcus mutans UA159 GCA_000007465.2
653 Streptococcus ratti FA-1 = DSM 20564 GCA_000286075.1
609 Streptococcus gordonii str. Challis substr. CH1 GCA_000017005.1
ik
775 Clostridium beijerinckii GCA_000833105.2
725 Clostridium saccharoperbutylacetonicum N1-4_28HMT_29 GCA_000340885.1
718 Clostridium puniceum GCA_002006345.1
il
805 Clostridium beijerinckii GCA_000833105.2
775 Clostridium saccharoperbutylacetonicum N1-4_28HMT_29 GCA_000340885.1
749 Clostridium saccharobutylicum DSM 13864 GCA_000473995.1
im
613 Lactobacillus gasseri ATCC 33323 = JCM 1131 GCA_000014425.1
597 Lactobacillus hominis DSM 23910 = CRBIP 24.179 GCA_000296835.1
549 Staphylococcus epidermidis ATCC 12228 GCA_000007645.1
in
739 Rhodobacter sphaeroides 2.4.1 GCA_000012905.2
665 Rhodobacter sphaeroides ATCC 17025 GCA_000016405.1

525 Rhodobacter capsulatus SB 1003 GCA_000021865.1
io
759 Streptococcus mutans UA159 GCA_000007465.2
560 Streptococcus ratti FA-1 = DSM 20564 GCA_000286075.1
491 Streptococcus ferus DSM 20646 GCA_000372425.1
ip
570 Lactobacillus gasseri ATCC 33323 = JCM 1131 GCA_000014425.1
513 Lactobacillus hominis DSM 23910 = CRBIP 24.179 GCA_000296835.1
379 Lactobacillus iners DSM 13335 GCA_000160875.1
iq
747 Rhodobacter sphaeroides 2.4.1 GCA_000012905.2
689 Rhodobacter sphaeroides ATCC 17025 GCA_000016405.1
534 Pseudorhodobacter psychrotolerans GCA_001294535.1
ir
604 Staphylococcus epidermidis ATCC 12228 GCA_000007645.1
549 Staphylococcus capitis subsp. capitis GCA_001028645.1
514 Staphylococcus hominis subsp. hominis C80 GCA_000183685.1
is
680 Shigella flexneri 2a str. 301 GCA_000006925.2
678 Escherichia coli O104_3AH4 str. 2011C-3493 GCA_000299455.1
677 Escherichia coli str. K-12 substr. MG1655 GCA_000005845.2
it
764 Lactobacillus gasseri ATCC 33323 = JCM 1131 GCA_000014425.1
717 Lactobacillus hominis DSM 23910 = CRBIP 24.179 GCA_000296835.1
584 Lactobacillus psittaci DSM 15354 GCA_000425905.1
iu
596 Streptococcus mutans UA159 GCA_000007465.2
482 Streptococcus sobrinus DSM 20742 = ATCC 33478 GCA_000686605.1
465 Streptococcus ratti FA-1 = DSM 20564 GCA_000286075.1
iv
607 Streptococcus mutans UA159 GCA_000007465.2
462 Streptococcus gordonii str. Challis substr. CH1 GCA_000017005.1
425 Streptococcus ratti FA-1 = DSM 20564 GCA_000286075.1
iw
787 Escherichia coli O83_3AH1 str. NRG 857C GCA_000183345.1
779 Escherichia coli str. K-12 substr. MG1655 GCA_000005845.2
773 Escherichia coli O104_3AH4 str. 2011C-3493 GCA_000299455.1
773 Escherichia coli O157_3AH7 str. Sakai GCA_000008865.1
773 Shigella flexneri 2a str. 301 GCA_000006925.2
ix
595 Clostridium beijerinckii GCA_000833105.2
564 Clostridium butyricum GCA_001456065.2
553 Clostridium puniceum GCA_002006345.1
553 Clostridium saccharoperbutylacetonicum N1-4_28HMT_29 GCA_000340885.1
iy
754 Enterococcus faecalis V583 GCA_000007785.1
713 Streptomyces cinnamoneus GCA_001885705.1
650 Enterococcus canis NBRC 100695 GCA_001544375.1
650 Enterococcus rivorum GCA_001742285.1
iz
560 Clostridium beijerinckii GCA_000833105.2
549 Clostridium saccharoperbutylacetonicum N1-4_28HMT_29 GCA_000340885.1
534 Clostridium puniceum GCA_002006345.1
534 Clostridium saccharobutylicum DSM 13864 GCA_000473995.1
iA
886 Enterococcus faecalis V583 GCA_000007785.1
839 Streptomyces cinnamoneus GCA_001885705.1
756 Enterococcus haemoperoxidus ATCC BAA-382 GCA_000407165.1
iB
847 Clostridium beijerinckii GCA_000833105.2
792 Clostridium saccharoperbutylacetonicum N1-4_28HMT_29 GCA_000340885.1
784 Clostridium saccharobutylicum DSM 13864 GCA_000473995.1
iC
694 Lactobacillus gasseri ATCC 33323 = JCM 1131 GCA_000014425.1
661 Lactobacillus hominis DSM 23910 = CRBIP 24.179 GCA_000296835.1
465 Lactobacillus iners DSM 13335 GCA_000160875.1
iD
782 Deinococcus radiodurans R1 GCA_000008565.1
552 Deinococcus deserti VCD115 GCA_000020685.1
489 Deinococcus hopiensis KR-140 GCA_900176165.1
iE
891 Streptococcus mutans UA159 GCA_000007465.2
667 Streptococcus ratti FA-1 = DSM 20564 GCA_000286075.1
641 Streptococcus anginosus C238 GCA_000463505.1
iF
800 Rhodobacter sphaeroides 2.4.1 GCA_000012905.2

734 Rhodobacter sphaeroides ATCC 17025 GCA_000016405.1
622 Defluviimonas alba GCA_001620265.1
iG
416 Bifidobacterium adolescentis ATCC 15703 GCA_000010425.1
321 Bifidobacterium dentium JCM 1195 = DSM 20436 GCA_001042595.1
304 Bifidobacterium breve DSM 20213 = JCM 1192 GCA_001025175.1
iH
506 Lactobacillus gasseri ATCC 33323 = JCM 1131 GCA_000014425.1
402 Lactobacillus hominis DSM 23910 = CRBIP 24.179 GCA_000296835.1
315 Lactobacillus acetotolerans GCA_001042405.1
iI
612 Clostridium beijerinckii GCA_000833105.2
595 Clostridium saccharoperbutylacetonicum N1-4_28HMT_29 GCA_000340885.1
581 Clostridium puniceum GCA_002006345.1
iJ
516 Clostridium beijerinckii GCA_000833105.2
495 Clostridium saccharoperbutylacetonicum N1-4_28HMT_29 GCA_000340885.1
479 Clostridium saccharobutylicum DSM 13864 GCA_000473995.1
iK
558 Deinococcus radiodurans R1 GCA_000008565.1
270 Deinococcus deserti VCD115 GCA_000020685.1
263 Deinococcus puniceus GCA_001644565.1
iL
464 Clostridium beijerinckii GCA_000833105.2
449 Clostridium saccharoperbutylacetonicum N1-4_28HMT_29 GCA_000340885.1
433 Clostridium puniceum GCA_002006345.1
iM
674 Clostridium beijerinckii GCA_000833105.2
634 Clostridium saccharoperbutylacetonicum N1-4_28HMT_29 GCA_000340885.1
619 Clostridium puniceum GCA_002006345.1
iN
619 Staphylococcus epidermidis ATCC 12228 GCA_000007645.1
594 Staphylococcus capitis subsp. capitis GCA_001028645.1
565 Staphylococcus haemolyticus JCSC1435 GCA_000009865.1
iO
749 Deinococcus radiodurans R1 GCA_000008565.1
555 Deinococcus gobiensis I-0 GCA_000252445.1
529 Deinococcus deserti VCD115 GCA_000020685.1
iP
708 _5BBacillus thuringiensis_5D serovar konkukian str. 97-27 GCA_000008505.1
707 Bacillus anthracis str. Ames GCA_000007845.1
707 Bacillus anthracis str. Sterne GCA_000008165.1
700 Bacillus cereus ATCC 14579 GCA_000007825.1
iQ
510 Deinococcus radiodurans R1 GCA_000008565.1
289 Deinococcus soli Cha et al. 2016 GCA_001007995.1
272 Deinococcus deserti VCD115 GCA_000020685.1
272 Deinococcus gobiensis I-0 GCA_000252445.1
iR
609 Bacillus anthracis str. Ames GCA_000007845.1
609 Bacillus anthracis str. Sterne GCA_000008165.1
608 Bacillus cereus ATCC 14579 GCA_000007825.1
608 _5BBacillus thuringiensis_5D serovar konkukian str. 97-27 GCA_000008505.1
593 Bacillus pseudomycoides DSM 12442 GCA_000161455.1
593 Bacillus thuringiensis YBT-1518 GCA_000497525.2
iS
689 Enterococcus faecalis V583 GCA_000007785.1
651 Enterococcus rivorum GCA_001742285.1
643 Streptomyces cinnamoneus GCA_001885705.1
iT
651 Deinococcus radiodurans R1 GCA_000008565.1
399 Deinococcus deserti VCD115 GCA_000020685.1
374 Deinococcus hopiensis KR-140 GCA_900176165.1
iU
755 Rhodobacter sphaeroides 2.4.1 GCA_000012905.2
698 Rhodobacter sphaeroides ATCC 17025 GCA_000016405.1
526 Gemmobacter aquatilis GCA_900110025.1
iV
552 Staphylococcus epidermidis ATCC 12228 GCA_000007645.1
510 Staphylococcus capitis subsp. capitis GCA_001028645.1
489 Staphylococcus aureus subsp. aureus NCTC 8325 GCA_000013425.1
iW
623 Rhodobacter sphaeroides 2.4.1 GCA_000012905.2
544 Rhodobacter sphaeroides ATCC 17025 GCA_000016405.1
489 Pseudorhodobacter ferrugineus DSM 5888 GCA_000420745.1
iX

896 Streptococcus mutans UA159 GCA_000007465.2
663 Streptococcus equinus GCA_000964315.1
663 Streptococcus ratti FA-1 = DSM 20564 GCA_000286075.1
646 Streptococcus pseudoporcinus LQ 940-04 GCA_000188035.3
iY
622 Lactobacillus gasseri ATCC 33323 = JCM 1131 GCA_000014425.1
561 Lactobacillus hominis DSM 23910 = CRBIP 24.179 GCA_000296835.1
382 Lactobacillus iners DSM 13335 GCA_000160875.1
iZ
821 Streptococcus mutans UA159 GCA_000007465.2
647 Streptococcus ratti FA-1 = DSM 20564 GCA_000286075.1
573 Streptococcus gordonii str. Challis substr. CH1 GCA_000017005.1
j0
795 Deinococcus radiodurans R1 GCA_000008565.1
517 Deinococcus deserti VCD115 GCA_000020685.1
488 Deinococcus hopiensis KR-140 GCA_900176165.1
j1
634 Clostridium beijerinckii GCA_000833105.2
577 Clostridium saccharoperbutylacetonicum N1-4_28HMT_29 GCA_000340885.1
573 Clostridium puniceum GCA_002006345.1
j2
734 Rhodobacter sphaeroides 2.4.1 GCA_000012905.2
694 Rhodobacter sphaeroides ATCC 17025 GCA_000016405.1
587 Pseudorhodobacter ferrugineus DSM 5888 GCA_000420745.1
j3
651 Staphylococcus epidermidis ATCC 12228 GCA_000007645.1
631 Staphylococcus capitis subsp. capitis GCA_001028645.1
576 Staphylococcus haemolyticus JCSC1435 GCA_000009865.1
576 Staphylococcus hominis subsp. hominis C80 GCA_000183685.1
j4
549 Bacillus anthracis str. Ames GCA_000007845.1
549 Bacillus anthracis str. Sterne GCA_000008165.1
549 Bacillus cereus ATCC 14579 GCA_000007825.1
549 Bacillus thuringiensis YBT-1518 GCA_000497525.2
549 _5BBacillus thuringiensis_5D serovar konkukian str. 97-27 GCA_000008505.1
534 Bacillus pseudomycoides DSM 12442 GCA_000161455.1
517 Bacillus cytotoxicus NVH 391-98 GCA_000017425.1
j5
512 Escherichia coli O83_3AH1 str. NRG 857C GCA_000183345.1
508 Escherichia coli O157_3AH7 str. Sakai GCA_000008865.1
508 Escherichia coli UMN026 GCA_000026325.2
508 Escherichia coli str. K-12 substr. MG1655 GCA_000005845.2
508 Shigella flexneri 2a str. 301 GCA_000006925.2
493 Escherichia coli IAI39 GCA_000026345.1
j6
796 Staphylococcus epidermidis ATCC 12228 GCA_000007645.1
765 Staphylococcus capitis subsp. capitis GCA_001028645.1
686 Staphylococcus warneri SG1 GCA_000332735.1
j7
770 Rhodobacter sphaeroides 2.4.1 GCA_000012905.2
707 Rhodobacter sphaeroides ATCC 17025 GCA_000016405.1
577 Pseudorhodobacter ferrugineus DSM 5888 GCA_000420745.1
j8
671 Staphylococcus epidermidis ATCC 12228 GCA_000007645.1
619 Staphylococcus capitis subsp. capitis GCA_001028645.1
568 Staphylococcus aureus subsp. aureus NCTC 8325 GCA_000013425.1
j9
528 Bifidobacterium adolescentis ATCC 15703 GCA_000010425.1
470 Bifidobacterium dentium JCM 1195 = DSM 20436 GCA_001042595.1
446 Bifidobacterium angulatum DSM 20098 = JCM 7096 GCA_001025155.1
ja
607 Rhodobacter sphaeroides 2.4.1 GCA_000012905.2
568 Rhodobacter sphaeroides ATCC 17025 GCA_000016405.1
500 Gemmobacter aquatilis GCA_900110025.1
jb
868 Staphylococcus epidermidis ATCC 12228 GCA_000007645.1
808 Staphylococcus capitis subsp. capitis GCA_001028645.1
795 Staphylococcus lugdunensis HKU09-01 GCA_000025085.1
jc
667 Deinococcus radiodurans R1 GCA_000008565.1
499 Deinococcus deserti VCD115 GCA_000020685.1
440 Deinococcus gobiensis I-0 GCA_000252445.1
jd
727 Clostridium beijerinckii GCA_000833105.2
695 Clostridium saccharoperbutylacetonicum N1-4_28HMT_29 GCA_000340885.1
688 Clostridium puniceum GCA_002006345.1

je
771 Bifidobacterium adolescentis ATCC 15703 GCA_000010425.1
618 Bifidobacterium angulatum DSM 20098 = JCM 7096 GCA_001025155.1
608 Bifidobacterium dentium JCM 1195 = DSM 20436 GCA_001042595.1
jf
834 Streptococcus mutans UA159 GCA_000007465.2
644 Streptococcus ratti FA-1 = DSM 20564 GCA_000286075.1
563 Streptococcus equinus GCA_000964315.1
jg
484 Deinococcus radiodurans R1 GCA_000008565.1
294 Deinococcus soli Cha et al. 2016 GCA_001007995.1
292 Deinococcus deserti VCD115 GCA_000020685.1
jh
558 Rhodobacter sphaeroides 2.4.1 GCA_000012905.2
486 Rhodobacter sphaeroides ATCC 17025 GCA_000016405.1
397 Gemmobacter aquatilis GCA_900110025.1
ji
494 Rhodobacter sphaeroides 2.4.1 GCA_000012905.2
442 Rhodobacter sphaeroides ATCC 17025 GCA_000016405.1
403 Gemmobacter nectariphilus DSM 15620 GCA_000429765.1
jj
627 Clostridium beijerinckii GCA_000833105.2
555 Clostridium saccharobutylicum DSM 13864 GCA_000473995.1
551 Clostridium saccharoperbutylacetonicum N1-4_28HMT_29 GCA_000340885.1
jk
964 Deinococcus radiodurans R1 GCA_000008565.1
619 Deinococcus deserti VCD115 GCA_000020685.1
577 Deinococcus marmoris DSM 12784 GCA_000701405.1
jl
863 Enterococcus faecalis V583 GCA_000007785.1
816 Streptomyces cinnamoneus GCA_001885705.1
727 Enterococcus rivorum GCA_001742285.1
jm
678 Lactobacillus gasseri ATCC 33323 = JCM 1131 GCA_000014425.1
642 Lactobacillus hominis DSM 23910 = CRBIP 24.179 GCA_000296835.1
425 Lactobacillus iners DSM 13335 GCA_000160875.1
jn
626 Clostridium beijerinckii GCA_000833105.2
607 Clostridium saccharoperbutylacetonicum N1-4_28HMT_29 GCA_000340885.1
598 Clostridium saccharobutylicum DSM 13864 GCA_000473995.1
jo
744 Enterococcus faecalis V583 GCA_000007785.1
703 Streptomyces cinnamoneus GCA_001885705.1
648 Enterococcus rivorum GCA_001742285.1
jp
728 Rhodobacter sphaeroides 2.4.1 GCA_000012905.2
636 Rhodobacter sphaeroides ATCC 17025 GCA_000016405.1
541 Pseudorhodobacter ferrugineus DSM 5888 GCA_000420745.1
jq
555 Clostridium saccharoperbutylacetonicum N1-4_28HMT_29 GCA_000340885.1
545 Clostridium beijerinckii GCA_000833105.2
538 Clostridium saccharobutylicum DSM 13864 GCA_000473995.1
jr
723 Clostridium beijerinckii GCA_000833105.2
696 Clostridium saccharoperbutylacetonicum N1-4_28HMT_29 GCA_000340885.1
672 Clostridium puniceum GCA_002006345.1
js
546 Clostridium saccharoperbutylacetonicum N1-4_28HMT_29 GCA_000340885.1
541 Clostridium beijerinckii GCA_000833105.2
516 Clostridium puniceum GCA_002006345.1
jt
824 Clostridium beijerinckii GCA_000833105.2
767 Clostridium saccharoperbutylacetonicum N1-4_28HMT_29 GCA_000340885.1
757 Clostridium saccharobutylicum DSM 13864 GCA_000473995.1
ju
714 Lactobacillus gasseri ATCC 33323 = JCM 1131 GCA_000014425.1
631 Lactobacillus hominis DSM 23910 = CRBIP 24.179 GCA_000296835.1
479 Lactobacillus iners DSM 13335 GCA_000160875.1
jv
470 Clostridium beijerinckii GCA_000833105.2
465 Clostridium saccharoperbutylacetonicum N1-4_28HMT_29 GCA_000340885.1
460 Staphylococcus epidermidis ATCC 12228 GCA_000007645.1
jw
850 Lactobacillus gasseri ATCC 33323 = JCM 1131 GCA_000014425.1
766 Lactobacillus hominis DSM 23910 = CRBIP 24.179 GCA_000296835.1
551 Lactobacillus iners DSM 13335 GCA_000160875.1

jx
672 Enterococcus faecalis V583 GCA_000007785.1
625 Streptomyces cinnamoneus GCA_001885705.1
595 Enterococcus phoeniculicola ATCC BAA-412 GCA_000407505.1
jy
565 Escherichia coli O104_3AH4 str. 2011C-3493 GCA_000299455.1
565 Escherichia coli O157_3AH7 str. Sakai GCA_000008865.1
565 Escherichia coli str. K-12 substr. MG1655 GCA_000005845.2
565 Shigella flexneri 2a str. 301 GCA_000006925.2
557 Escherichia coli O83_3AH1 str. NRG 857C GCA_000183345.1
557 Escherichia coli UMN026 GCA_000026325.2
520 Escherichia coli IAI39 GCA_000026345.1
jz
584 Staphylococcus epidermidis ATCC 12228 GCA_000007645.1
569 Staphylococcus capitis subsp. capitis GCA_001028645.1
514 Staphylococcus warneri SG1 GCA_000332735.1
jA
584 Enterococcus faecalis V583 GCA_000007785.1
538 Enterococcus rivorum GCA_001742285.1
538 Streptomyces cinnamoneus GCA_001885705.1
535 Enterococcus asini ATCC 700915 GCA_000407365.1
jB
729 Lactobacillus gasseri ATCC 33323 = JCM 1131 GCA_000014425.1
675 Lactobacillus hominis DSM 23910 = CRBIP 24.179 GCA_000296835.1
486 Lactobacillus hamsteri DSM 5661 = JCM 6256 GCA_000615445.1
jC
459 Escherichia coli str. K-12 substr. MG1655 GCA_000005845.2
417 Escherichia coli O104_3AH4 str. 2011C-3493 GCA_000299455.1
410 Escherichia coli UMN026 GCA_000026325.2
jD
675 Clostridium beijerinckii GCA_000833105.2
639 Clostridium saccharobutylicum DSM 13864 GCA_000473995.1
632 Clostridium saccharoperbutylacetonicum N1-4_28HMT_29 GCA_000340885.1
jE
752 Rhodobacter sphaeroides 2.4.1 GCA_000012905.2
700 Rhodobacter sphaeroides ATCC 17025 GCA_000016405.1
630 Gemmobacter aquatilis GCA_900110025.1
jF
347 Bacillus cereus ATCC 14579 GCA_000007825.1
333 Bacillus anthracis str. Ames GCA_000007845.1
333 Bacillus anthracis str. Sterne GCA_000008165.1
333 Bacillus thuringiensis YBT-1518 GCA_000497525.2
333 _5BBacillus thuringiensis_5D serovar konkukian str. 97-27 GCA_000008505.1
320 Bacillus mycoides GCA_000832605.1
320 Bacillus pseudomycoides DSM 12442 GCA_000161455.1
jG
768 Clostridium beijerinckii GCA_000833105.2
753 Clostridium saccharoperbutylacetonicum N1-4_28HMT_29 GCA_000340885.1
708 Clostridium puniceum GCA_002006345.1
jH
678 Clostridium beijerinckii GCA_000833105.2
643 Clostridium saccharoperbutylacetonicum N1-4_28HMT_29 GCA_000340885.1
600 Clostridium puniceum GCA_002006345.1
jI
676 Rhodobacter sphaeroides 2.4.1 GCA_000012905.2
613 Rhodobacter sphaeroides ATCC 17025 GCA_000016405.1
499 Pseudorhodobacter psychrotolerans GCA_001294535.1
jJ
697 Enterococcus faecalis V583 GCA_000007785.1
650 Streptomyces cinnamoneus GCA_001885705.1
642 Enterococcus rivorum GCA_001742285.1
jK
822 Enterococcus faecalis V583 GCA_000007785.1
775 Streptomyces cinnamoneus GCA_001885705.1
669 Enterococcus rivorum GCA_001742285.1
jL
763 Clostridium beijerinckii GCA_000833105.2
729 Clostridium saccharoperbutylacetonicum N1-4_28HMT_29 GCA_000340885.1
692 Clostridium saccharobutylicum DSM 13864 GCA_000473995.1
jM
904 Staphylococcus epidermidis ATCC 12228 GCA_000007645.1
837 Staphylococcus capitis subsp. capitis GCA_001028645.1
788 Staphylococcus aureus subsp. aureus NCTC 8325 GCA_000013425.1
jN
665 Bifidobacterium adolescentis ATCC 15703 GCA_000010425.1
487 Bifidobacterium angulatum DSM 20098 = JCM 7096 GCA_001025155.1

478 Bifidobacterium dentium JCM 1195 = DSM 20436 GCA_001042595.1
jO
720 Clostridium beijerinckii GCA_000833105.2
715 Clostridium saccharoperbutylacetonicum N1-4_28HMT_29 GCA_000340885.1
686 Clostridium saccharobutylicum DSM 13864 GCA_000473995.1
jP
815 Rhodobacter sphaeroides 2.4.1 GCA_000012905.2
737 Rhodobacter sphaeroides ATCC 17025 GCA_000016405.1
593 Gemmobacter aquatilis GCA_900110025.1
jQ
553 Staphylococcus epidermidis ATCC 12228 GCA_000007645.1
508 Staphylococcus capitis subsp. capitis GCA_001028645.1
498 Staphylococcus hominis subsp. hominis C80 GCA_000183685.1
jR
679 Clostridium beijerinckii GCA_000833105.2
654 Clostridium puniceum GCA_002006345.1
652 Clostridium saccharoperbutylacetonicum N1-4_28HMT_29 GCA_000340885.1
jS
715 Bacillus anthracis str. Ames GCA_000007845.1
715 Bacillus anthracis str. Sterne GCA_000008165.1
715 Bacillus cereus ATCC 14579 GCA_000007825.1
715 _5BBacillus thuringiensis_5D serovar konkukian str. 97-27 GCA_000008505.1
700 Bacillus thuringiensis YBT-1518 GCA_000497525.2
676 Bacillus mycoides GCA_000832605.1
jT
735 Escherichia coli O157_3AH7 str. Sakai GCA_000008865.1
735 Escherichia coli str. K-12 substr. MG1655 GCA_000005845.2
722 Escherichia coli IAI39 GCA_000026345.1
718 Escherichia coli UMN026 GCA_000026325.2
jU
935 Bacillus anthracis str. Ames GCA_000007845.1
935 Bacillus anthracis str. Sterne GCA_000008165.1
935 Bacillus cereus ATCC 14579 GCA_000007825.1
935 _5BBacillus thuringiensis_5D serovar konkukian str. 97-27 GCA_000008505.1
909 Bacillus thuringiensis YBT-1518 GCA_000497525.2
904 Bacillus pseudomycoides DSM 12442 GCA_000161455.1
jV
931 Rhodobacter sphaeroides 2.4.1 GCA_000012905.2
794 Rhodobacter sphaeroides ATCC 17025 GCA_000016405.1
668 Gemmobacter aquatilis GCA_900110025.1
668 Rhodobacter capsulatus SB 1003 GCA_000021865.1
jW
622 Streptococcus mutans UA159 GCA_000007465.2
455 Streptococcus gordonii str. Challis substr. CH1 GCA_000017005.1
436 Streptococcus orisratti DSM 15617 GCA_000380105.1
jX
828 Staphylococcus epidermidis ATCC 12228 GCA_000007645.1
813 Staphylococcus capitis subsp. capitis GCA_001028645.1
765 Staphylococcus aureus subsp. aureus NCTC 8325 GCA_000013425.1
jY
576 Bacillus anthracis str. Ames GCA_000007845.1
576 Bacillus anthracis str. Sterne GCA_000008165.1
576 Bacillus pseudomycoides DSM 12442 GCA_000161455.1
576 _5BBacillus thuringiensis_5D serovar konkukian str. 97-27 GCA_000008505.1
561 Bacillus cereus ATCC 14579 GCA_000007825.1
551 Bacillus thuringiensis YBT-1518 GCA_000497525.2
jZ
721 Rhodobacter sphaeroides 2.4.1 GCA_000012905.2
629 Rhodobacter sphaeroides ATCC 17025 GCA_000016405.1
551 Pseudorhodobacter ferrugineus DSM 5888 GCA_000420745.1
k0
795 Escherichia coli str. K-12 substr. MG1655 GCA_000005845.2
753 Escherichia coli O104_3AH4 str. 2011C-3493 GCA_000299455.1
739 Shigella flexneri 2a str. 301 GCA_000006925.2
k1
615 Escherichia coli O157_3AH7 str. Sakai GCA_000008865.1
615 Escherichia coli UMN026 GCA_000026325.2
615 Escherichia coli str. K-12 substr. MG1655 GCA_000005845.2
603 Escherichia coli O83_3AH1 str. NRG 857C GCA_000183345.1
599 Shigella flexneri 2a str. 301 GCA_000006925.2
k2
817 Enterococcus faecalis V583 GCA_000007785.1
770 Streptomyces cinnamoneus GCA_001885705.1
740 Enterococcus asini ATCC 700915 GCA_000407365.1
740 Enterococcus rivorum GCA_001742285.1
k3

709 Streptococcus mutans UA159 GCA_000007465.2
504 Streptococcus gordonii str. Challis substr. CH1 GCA_000017005.1
485 Streptococcus ratti FA-1 = DSM 20564 GCA_000286075.1
k4
719 Deinococcus radiodurans R1 GCA_000008565.1
499 Deinococcus gobiensis I-0 GCA_000252445.1
494 Deinococcus soli Cha et al. 2016 GCA_001007995.1
k5
595 Deinococcus radiodurans R1 GCA_000008565.1
361 Deinococcus gobiensis I-0 GCA_000252445.1
359 Deinococcus deserti VCD115 GCA_000020685.1
k6
695 Clostridium beijerinckii GCA_000833105.2
648 Clostridium butyricum GCA_001456065.2
645 Clostridium saccharobutylicum DSM 13864 GCA_000473995.1
k7
782 Bacillus cereus ATCC 14579 GCA_000007825.1
775 Bacillus anthracis str. Ames GCA_000007845.1
775 Bacillus anthracis str. Sterne GCA_000008165.1
775 Bacillus thuringiensis YBT-1518 GCA_000497525.2
775 _5BBacillus thuringiensis_5D serovar konkukian str. 97-27 GCA_000008505.1
766 Bacillus pseudomycoides DSM 12442 GCA_000161455.1
k8
636 Escherichia coli O157_3AH7 str. Sakai GCA_000008865.1
636 Escherichia coli str. K-12 substr. MG1655 GCA_000005845.2
628 Escherichia coli UMN026 GCA_000026325.2
614 Escherichia coli IAI39 GCA_000026345.1
k9
681 Streptococcus mutans UA159 GCA_000007465.2
547 Streptococcus ratti FA-1 = DSM 20564 GCA_000286075.1
506 Streptococcus macacae NCTC 11558 GCA_000187995.3
ka
793 Lactobacillus gasseri ATCC 33323 = JCM 1131 GCA_000014425.1
744 Lactobacillus hominis DSM 23910 = CRBIP 24.179 GCA_000296835.1
560 Lactobacillus iners DSM 13335 GCA_000160875.1
kb
595 Escherichia coli O104_3AH4 str. 2011C-3493 GCA_000299455.1
595 Escherichia coli O157_3AH7 str. Sakai GCA_000008865.1
595 Escherichia coli str. K-12 substr. MG1655 GCA_000005845.2
595 Shigella flexneri 2a str. 301 GCA_000006925.2
590 Escherichia coli IAI39 GCA_000026345.1
587 Escherichia coli O83_3AH1 str. NRG 857C GCA_000183345.1
587 Escherichia coli UMN026 GCA_000026325.2
kc
655 Escherichia coli O157_3AH7 str. Sakai GCA_000008865.1
655 Escherichia coli str. K-12 substr. MG1655 GCA_000005845.2
647 Escherichia coli UMN026 GCA_000026325.2
646 Shigella flexneri 2a str. 301 GCA_000006925.2
kd
533 Shigella flexneri 2a str. 301 GCA_000006925.2
528 Escherichia coli O104_3AH4 str. 2011C-3493 GCA_000299455.1
528 Escherichia coli O157_3AH7 str. Sakai GCA_000008865.1
528 Escherichia coli str. K-12 substr. MG1655 GCA_000005845.2
524 Escherichia coli IAI39 GCA_000026345.1
ke
615 Rhodobacter sphaeroides 2.4.1 GCA_000012905.2
501 Rhodobacter sphaeroides ATCC 17025 GCA_000016405.1
471 Gemmobacter megaterium GCA_900156815.1
kf
515 Lactobacillus gasseri ATCC 33323 = JCM 1131 GCA_000014425.1
490 Lactobacillus hominis DSM 23910 = CRBIP 24.179 GCA_000296835.1
329 Lactobacillus kefiranofaciens ZW3 GCA_000214785.1
kg
837 Escherichia coli O157_3AH7 str. Sakai GCA_000008865.1
837 Escherichia coli str. K-12 substr. MG1655 GCA_000005845.2
836 Escherichia coli UMN026 GCA_000026325.2
828 Shigella flexneri 2a str. 301 GCA_000006925.2
kh
761 Clostridium beijerinckii GCA_000833105.2
752 Clostridium saccharoperbutylacetonicum N1-4_28HMT_29 GCA_000340885.1
724 Clostridium puniceum GCA_002006345.1
ki
757 Rhodobacter sphaeroides 2.4.1 GCA_000012905.2
685 Rhodobacter sphaeroides ATCC 17025 GCA_000016405.1
545 Defluviimonas alba GCA_001620265.1
kj

746 Deinococcus radiodurans R1 GCA_000008565.1
438 Deinococcus deserti VCD115 GCA_000020685.1
422 Deinococcus proteolyticus MRP GCA_000190555.1
kk
491 Streptococcus mutans UA159 GCA_000007465.2
337 Streptococcus macacae NCTC 11558 GCA_000187995.3
323 Streptococcus ratti FA-1 = DSM 20564 GCA_000286075.1
kl
437 Staphylococcus epidermidis ATCC 12228 GCA_000007645.1
425 Staphylococcus hominis subsp. hominis C80 GCA_000183685.1
421 Megasphaera cerevisiae DSM 20462 GCA_001045675.1
421 Staphylococcus warneri SG1 GCA_000332735.1
km
522 Streptococcus mutans UA159 GCA_000007465.2
410 Streptococcus ratti FA-1 = DSM 20564 GCA_000286075.1
390 Streptococcus macacae NCTC 11558 GCA_000187995.3
kn
609 Rhodobacter sphaeroides 2.4.1 GCA_000012905.2
525 Rhodobacter sphaeroides ATCC 17025 GCA_000016405.1
477 Pseudorhodobacter ferrugineus DSM 5888 GCA_000420745.1
ko
761 Streptococcus mutans UA159 GCA_000007465.2
615 Streptococcus gordonii str. Challis substr. CH1 GCA_000017005.1
614 Streptococcus equinus GCA_000964315.1
kp
527 Lactobacillus gasseri ATCC 33323 = JCM 1131 GCA_000014425.1
521 Lactobacillus hominis DSM 23910 = CRBIP 24.179 GCA_000296835.1
391 Deinococcus radiodurans R1 GCA_000008565.1
kq
758 Bacillus anthracis str. Ames GCA_000007845.1
758 Bacillus anthracis str. Sterne GCA_000008165.1
758 _5BBacillus thuringiensis_5D serovar konkukian str. 97-27 GCA_000008505.1
733 Bacillus cereus ATCC 14579 GCA_000007825.1
727 Bacillus pseudomycoides DSM 12442 GCA_000161455.1
kr
710 Clostridium beijerinckii GCA_000833105.2
692 Clostridium saccharoperbutylacetonicum N1-4_28HMT_29 GCA_000340885.1
686 Clostridium puniceum GCA_002006345.1
ks
755 Bacillus cereus ATCC 14579 GCA_000007825.1
754 Bacillus thuringiensis YBT-1518 GCA_000497525.2
753 Bacillus anthracis str. Ames GCA_000007845.1
753 Bacillus anthracis str. Sterne GCA_000008165.1
753 _5BBacillus thuringiensis_5D serovar konkukian str. 97-27 GCA_000008505.1
kt
641 Deinococcus radiodurans R1 GCA_000008565.1
432 Deinococcus soli Cha et al. 2016 GCA_001007995.1
428 Deinococcus deserti VCD115 GCA_000020685.1
ku
492 Clostridium beijerinckii GCA_000833105.2
477 Clostridium saccharoperbutylacetonicum N1-4_28HMT_29 GCA_000340885.1
466 Clostridium puniceum GCA_002006345.1
kv
576 Bacillus thuringiensis YBT-1518 GCA_000497525.2
561 Bacillus cereus ATCC 14579 GCA_000007825.1
558 Bacillus anthracis str. Ames GCA_000007845.1
558 Bacillus anthracis str. Sterne GCA_000008165.1
558 _5BBacillus thuringiensis_5D serovar konkukian str. 97-27 GCA_000008505.1
kw
802 Bifidobacterium adolescentis ATCC 15703 GCA_000010425.1
658 Bifidobacterium angulatum DSM 20098 = JCM 7096 GCA_001025155.1
625 Bifidobacterium dentium JCM 1195 = DSM 20436 GCA_001042595.1
kx
694 Rhodobacter sphaeroides 2.4.1 GCA_000012905.2
605 Rhodobacter sphaeroides ATCC 17025 GCA_000016405.1
511 Gemmobacter aquatilis GCA_900110025.1
ky
617 Staphylococcus epidermidis ATCC 12228 GCA_000007645.1
515 Staphylococcus capitis subsp. capitis GCA_001028645.1
494 Megasphaera cerevisiae DSM 20462 GCA_001045675.1
494 Staphylococcus warneri SG1 GCA_000332735.1
kz
730 Deinococcus radiodurans R1 GCA_000008565.1
498 Deinococcus deserti VCD115 GCA_000020685.1
484 Deinococcus soli Cha et al. 2016 GCA_001007995.1
kA

692 Rhodobacter sphaeroides 2.4.1 GCA_000012905.2
656 Rhodobacter sphaeroides ATCC 17025 GCA_000016405.1
576 Gemmobacter megaterium GCA_900156815.1
kB
620 Bifidobacterium adolescentis ATCC 15703 GCA_000010425.1
468 Bifidobacterium callitrichos DSM 23973 GCA_000741175.1
466 Bifidobacterium dentium JCM 1195 = DSM 20436 GCA_001042595.1
kC
766 Bifidobacterium adolescentis ATCC 15703 GCA_000010425.1
617 Bifidobacterium dentium JCM 1195 = DSM 20436 GCA_001042595.1
600 Bifidobacterium callitrichos DSM 23973 GCA_000741175.1
kD
689 Streptococcus mutans UA159 GCA_000007465.2
500 Streptococcus equinus GCA_000964315.1
496 Streptococcus gallolyticus subsp. gallolyticus DSM 16831 GCA_002000985.1
kE
516 Streptococcus mutans UA159 GCA_000007465.2
361 Streptococcus ratti FA-1 = DSM 20564 GCA_000286075.1
350 Streptococcus sobrinus DSM 20742 = ATCC 33478 GCA_000686605.1
kF
950 Streptococcus mutans UA159 GCA_000007465.2
708 Streptococcus ratti FA-1 = DSM 20564 GCA_000286075.1
667 Streptococcus macacae NCTC 11558 GCA_000187995.3
kG
738 Deinococcus radiodurans R1 GCA_000008565.1
493 Deinococcus deserti VCD115 GCA_000020685.1
479 Deinococcus gobiensis I-0 GCA_000252445.1
kH
679 Enterococcus faecalis V583 GCA_000007785.1
632 Streptomyces cinnamoneus GCA_001885705.1
523 Enterococcus faecium DO GCA_000174395.2
kI
768 Rhodobacter sphaeroides 2.4.1 GCA_000012905.2
721 Rhodobacter sphaeroides ATCC 17025 GCA_000016405.1
604 Pseudorhodobacter psychrotolerans GCA_001294535.1
kJ
496 Streptococcus mutans UA159 GCA_000007465.2
401 Streptococcus ratti FA-1 = DSM 20564 GCA_000286075.1
378 Streptococcus sobrinus DSM 20742 = ATCC 33478 GCA_000686605.1
kK
988 Lactobacillus gasseri ATCC 33323 = JCM 1131 GCA_000014425.1
913 Lactobacillus hominis DSM 23910 = CRBIP 24.179 GCA_000296835.1
721 Lactobacillus iners DSM 13335 GCA_000160875.1
kL
722 Staphylococcus epidermidis ATCC 12228 GCA_000007645.1
681 Staphylococcus capitis subsp. capitis GCA_001028645.1
622 Staphylococcus aureus subsp. aureus NCTC 8325 GCA_000013425.1
kM
740 Clostridium beijerinckii GCA_000833105.2
726 Clostridium saccharoperbutylacetonicum N1-4_28HMT_29 GCA_000340885.1
682 Clostridium puniceum GCA_002006345.1
kN
655 Deinococcus radiodurans R1 GCA_000008565.1
439 Deinococcus deserti VCD115 GCA_000020685.1
432 Deinococcus gobiensis I-0 GCA_000252445.1
kO
348 Bifidobacterium adolescentis ATCC 15703 GCA_000010425.1
283 Bifidobacterium dentium JCM 1195 = DSM 20436 GCA_001042595.1
281 Bifidobacterium lemurum GCA_001895165.1
kP
749 Staphylococcus epidermidis ATCC 12228 GCA_000007645.1
727 Staphylococcus capitis subsp. capitis GCA_001028645.1
664 Staphylococcus aureus subsp. aureus NCTC 8325 GCA_000013425.1
kQ
734 Enterococcus faecalis V583 GCA_000007785.1
687 Streptomyces cinnamoneus GCA_001885705.1
647 Enterococcus rivorum GCA_001742285.1
kR
657 Clostridium beijerinckii GCA_000833105.2
643 Clostridium puniceum GCA_002006345.1
632 Clostridium saccharoperbutylacetonicum N1-4_28HMT_29 GCA_000340885.1
kS
533 Staphylococcus epidermidis ATCC 12228 GCA_000007645.1
493 Megasphaera cerevisiae DSM 20462 GCA_001045675.1
493 Staphylococcus warneri SG1 GCA_000332735.1
480 Staphylococcus aureus subsp. aureus NCTC 8325 GCA_000013425.1

kT
780 Bacillus anthracis str. Ames GCA_000007845.1
780 Bacillus anthracis str. Sterne GCA_000008165.1
780 Bacillus cereus ATCC 14579 GCA_000007825.1
780 Bacillus thuringiensis YBT-1518 GCA_000497525.2
780 _5BBacillus thuringiensis_5D serovar konkukian str. 97-27 GCA_000008505.1
732 Bacillus mycoides GCA_000832605.1
726 Bacillus pseudomycoides DSM 12442 GCA_000161455.1
kU
577 Streptococcus mutans UA159 GCA_000007465.2
446 Streptococcus ratti FA-1 = DSM 20564 GCA_000286075.1
420 Streptococcus gordonii str. Challis substr. CH1 GCA_000017005.1
kV
737 Lactobacillus gasseri ATCC 33323 = JCM 1131 GCA_000014425.1
658 Lactobacillus hominis DSM 23910 = CRBIP 24.179 GCA_000296835.1
461 Lactobacillus iners DSM 13335 GCA_000160875.1
kW
518 Deinococcus radiodurans R1 GCA_000008565.1
310 Deinococcus deserti VCD115 GCA_000020685.1
290 Deinococcus maricopensis DSM 21211 GCA_000186385.1
kX
950 Enterococcus faecalis V583 GCA_000007785.1
903 Streptomyces cinnamoneus GCA_001885705.1
831 Enterococcus canis NBRC 100695 GCA_001544375.1
kY
492 Bifidobacterium adolescentis ATCC 15703 GCA_000010425.1
364 Bifidobacterium angulatum DSM 20098 = JCM 7096 GCA_001025155.1
358 Bifidobacterium breve DSM 20213 = JCM 1192 GCA_001025175.1
kZ
847 Staphylococcus epidermidis ATCC 12228 GCA_000007645.1
811 Staphylococcus capitis subsp. capitis GCA_001028645.1
739 Staphylococcus aureus subsp. aureus NCTC 8325 GCA_000013425.1
l0
636 Rhodobacter sphaeroides 2.4.1 GCA_000012905.2
597 Rhodobacter sphaeroides ATCC 17025 GCA_000016405.1
480 Gemmobacter aquatilis GCA_900110025.1
l1
644 Deinococcus radiodurans R1 GCA_000008565.1
436 Deinococcus deserti VCD115 GCA_000020685.1
404 Deinococcus gobiensis I-0 GCA_000252445.1
l2
628 Rhodobacter sphaeroides 2.4.1 GCA_000012905.2
564 Rhodobacter sphaeroides ATCC 17025 GCA_000016405.1
507 Gemmobacter megaterium GCA_900156815.1
l3
633 Bacillus anthracis str. Ames GCA_000007845.1
633 Bacillus anthracis str. Sterne GCA_000008165.1
633 Bacillus pseudomycoides DSM 12442 GCA_000161455.1
633 _5BBacillus thuringiensis_5D serovar konkukian str. 97-27 GCA_000008505.1
614 Bacillus cereus ATCC 14579 GCA_000007825.1
614 Bacillus thuringiensis YBT-1518 GCA_000497525.2
599 Bacillus mycoides GCA_000832605.1
l4
752 Streptococcus mutans UA159 GCA_000007465.2
584 Streptococcus ratti FA-1 = DSM 20564 GCA_000286075.1
545 Streptococcus ferus DSM 20646 GCA_000372425.1
l5
541 Deinococcus radiodurans R1 GCA_000008565.1
341 Deinococcus hopiensis KR-140 GCA_900176165.1
341 Deinococcus puniceus GCA_001644565.1
332 Deinococcus phoenicis GCA_000599865.1
l6
363 Clostridium beijerinckii GCA_000833105.2
362 Clostridium saccharoperbutylacetonicum N1-4_28HMT_29 GCA_000340885.1
358 Clostridium saccharobutylicum DSM 13864 GCA_000473995.1
l7
415 Rhodobacter sphaeroides 2.4.1 GCA_000012905.2
382 Rhodobacter sphaeroides ATCC 17025 GCA_000016405.1
277 Gemmobacter aquatilis GCA_900110025.1
l8
522 Clostridium beijerinckii GCA_000833105.2
486 Clostridium saccharoperbutylacetonicum N1-4_28HMT_29 GCA_000340885.1
442 Clostridium butyricum GCA_001456065.2
l9
745 Bifidobacterium adolescentis ATCC 15703 GCA_000010425.1
574 Bifidobacterium dentium JCM 1195 = DSM 20436 GCA_001042595.1

535 Bifidobacterium bifidum PRL2010 GCA_000165905.1
la
441 Deinococcus radiodurans R1 GCA_000008565.1
278 Deinococcus soli Cha et al. 2016 GCA_001007995.1
266 Deinococcus deserti VCD115 GCA_000020685.1
lb
814 Escherichia coli str. K-12 substr. MG1655 GCA_000005845.2
791 Escherichia coli IAI39 GCA_000026345.1
780 Escherichia coli O157_3AH7 str. Sakai GCA_000008865.1
780 Shigella flexneri 2a str. 301 GCA_000006925.2
lc
278 Bifidobacterium adolescentis ATCC 15703 GCA_000010425.1
254 Bifidobacterium dentium JCM 1195 = DSM 20436 GCA_001042595.1
247 Bifidobacterium gallicum DSM 20093 = LMG 11596 GCA_000741205.1
ld
679 Staphylococcus epidermidis ATCC 12228 GCA_000007645.1
632 Staphylococcus haemolyticus JCSC1435 GCA_000009865.1
631 Staphylococcus capitis subsp. capitis GCA_001028645.1
le
735 Enterococcus faecalis V583 GCA_000007785.1
688 Streptomyces cinnamoneus GCA_001885705.1
632 Enterococcus faecium DO GCA_000174395.2
lf
671 Lactobacillus gasseri ATCC 33323 = JCM 1131 GCA_000014425.1
613 Lactobacillus hominis DSM 23910 = CRBIP 24.179 GCA_000296835.1
538 Lactobacillus iners DSM 13335 GCA_000160875.1
lg
323 Clostridium beijerinckii GCA_000833105.2
299 Clostridium saccharoperbutylacetonicum N1-4_28HMT_29 GCA_000340885.1
283 Clostridium saccharobutylicum DSM 13864 GCA_000473995.1
lh
738 Clostridium beijerinckii GCA_000833105.2
717 Clostridium saccharoperbutylacetonicum N1-4_28HMT_29 GCA_000340885.1
703 Clostridium puniceum GCA_002006345.1
li
905 Staphylococcus epidermidis ATCC 12228 GCA_000007645.1
860 Staphylococcus capitis subsp. capitis GCA_001028645.1
811 Staphylococcus lugdunensis HKU09-01 GCA_000025085.1
lj
488 Deinococcus radiodurans R1 GCA_000008565.1
281 Deinococcus deserti VCD115 GCA_000020685.1
278 Deinococcus gobiensis I-0 GCA_000252445.1
lk
643 Escherichia coli str. K-12 substr. MG1655 GCA_000005845.2
633 Escherichia coli O157_3AH7 str. Sakai GCA_000008865.1
618 Escherichia coli UMN026 GCA_000026325.2
ll
361 Clostridium beijerinckii GCA_000833105.2
346 Clostridium saccharoperbutylacetonicum N1-4_28HMT_29 GCA_000340885.1
339 Clostridium puniceum GCA_002006345.1
339 Clostridium saccharobutylicum DSM 13864 GCA_000473995.1
lm
640 Streptococcus mutans UA159 GCA_000007465.2
476 Streptococcus ratti FA-1 = DSM 20564 GCA_000286075.1
443 Streptococcus sobrinus DSM 20742 = ATCC 33478 GCA_000686605.1
ln
761 Rhodobacter sphaeroides 2.4.1 GCA_000012905.2
639 Rhodobacter sphaeroides ATCC 17025 GCA_000016405.1
528 Pseudorhodobacter psychrotolerans GCA_001294535.1
lo
504 Clostridium beijerinckii GCA_000833105.2
504 Clostridium saccharoperbutylacetonicum N1-4_28HMT_29 GCA_000340885.1
479 Clostridium butyricum GCA_001456065.2
471 Clostridium saccharobutylicum DSM 13864 GCA_000473995.1
lp
901 Streptococcus mutans UA159 GCA_000007465.2
659 Streptococcus ratti FA-1 = DSM 20564 GCA_000286075.1
625 Streptococcus gallolyticus subsp. gallolyticus DSM 16831 GCA_002000985.1
lq
664 Lactobacillus gasseri ATCC 33323 = JCM 1131 GCA_000014425.1
638 Lactobacillus hominis DSM 23910 = CRBIP 24.179 GCA_000296835.1
444 Lactobacillus psittaci DSM 15354 GCA_000425905.1
lr
600 Staphylococcus epidermidis ATCC 12228 GCA_000007645.1
555 Staphylococcus capitis subsp. capitis GCA_001028645.1
541 Staphylococcus aureus subsp. aureus NCTC 8325 GCA_000013425.1

541 Staphylococcus vitulinus F1028 GCA_000286335.1
ls
677 Rhodobacter sphaeroides 2.4.1 GCA_000012905.2
648 Rhodobacter sphaeroides ATCC 17025 GCA_000016405.1
507 Pseudorhodobacter psychrotolerans GCA_001294535.1
lt
387 Staphylococcus epidermidis ATCC 12228 GCA_000007645.1
379 Staphylococcus capitis subsp. capitis GCA_001028645.1
295 Staphylococcus lugdunensis HKU09-01 GCA_000025085.1
lu
539 Bifidobacterium adolescentis ATCC 15703 GCA_000010425.1
419 Bifidobacterium dentium JCM 1195 = DSM 20436 GCA_001042595.1
415 Bifidobacterium angulatum DSM 20098 = JCM 7096 GCA_001025155.1
lv
924 Enterococcus faecalis V583 GCA_000007785.1
877 Streptomyces cinnamoneus GCA_001885705.1
815 Enterococcus rivorum GCA_001742285.1
lw
562 Lactobacillus gasseri ATCC 33323 = JCM 1131 GCA_000014425.1
505 Lactobacillus hominis DSM 23910 = CRBIP 24.179 GCA_000296835.1
376 Lactobacillus iners DSM 13335 GCA_000160875.1
lx
888 Deinococcus radiodurans R1 GCA_000008565.1
535 Deinococcus deserti VCD115 GCA_000020685.1
498 Deinococcus hopiensis KR-140 GCA_900176165.1
ly
828 Streptococcus mutans UA159 GCA_000007465.2
597 Streptococcus ratti FA-1 = DSM 20564 GCA_000286075.1
554 Streptococcus ferus DSM 20646 GCA_000372425.1
554 Streptococcus gordonii str. Challis substr. CH1 GCA_000017005.1
lz
515 Rhodobacter sphaeroides 2.4.1 GCA_000012905.2
439 Rhodobacter sphaeroides ATCC 17025 GCA_000016405.1
373 Gemmobacter aquatilis GCA_900110025.1
lA
832 Lactobacillus gasseri ATCC 33323 = JCM 1131 GCA_000014425.1
794 Lactobacillus hominis DSM 23910 = CRBIP 24.179 GCA_000296835.1
552 Lactobacillus jensenii GCA_001936235.1
lB
870 Staphylococcus epidermidis ATCC 12228 GCA_000007645.1
813 Staphylococcus capitis subsp. capitis GCA_001028645.1
782 Staphylococcus warneri SG1 GCA_000332735.1
lC
699 Streptococcus mutans UA159 GCA_000007465.2
533 Streptococcus ratti FA-1 = DSM 20564 GCA_000286075.1
494 Streptococcus ferus DSM 20646 GCA_000372425.1
lD
826 Rhodobacter sphaeroides 2.4.1 GCA_000012905.2
732 Rhodobacter sphaeroides ATCC 17025 GCA_000016405.1
650 Pseudorhodobacter ferrugineus DSM 5888 GCA_000420745.1
lE
657 Streptococcus mutans UA159 GCA_000007465.2
480 Streptococcus gordonii str. Challis substr. CH1 GCA_000017005.1
479 Streptococcus ratti FA-1 = DSM 20564 GCA_000286075.1
lF
742 Streptococcus mutans UA159 GCA_000007465.2
527 Streptococcus ratti FA-1 = DSM 20564 GCA_000286075.1
517 Streptococcus gordonii str. Challis substr. CH1 GCA_000017005.1
lG
661 Deinococcus radiodurans R1 GCA_000008565.1
394 Deinococcus deserti VCD115 GCA_000020685.1
381 Deinococcus marmoris DSM 12784 GCA_000701405.1
lH
661 Staphylococcus epidermidis ATCC 12228 GCA_000007645.1
650 Staphylococcus capitis subsp. capitis GCA_001028645.1
606 Staphylococcus hominis subsp. hominis C80 GCA_000183685.1
lI
728 Lactobacillus gasseri ATCC 33323 = JCM 1131 GCA_000014425.1
658 Lactobacillus hominis DSM 23910 = CRBIP 24.179 GCA_000296835.1
525 Lactobacillus iners DSM 13335 GCA_000160875.1
lJ
873 Bacillus anthracis str. Ames GCA_000007845.1
873 Bacillus anthracis str. Sterne GCA_000008165.1
873 _5BBacillus thuringiensis_5D serovar konkukian str. 97-27 GCA_000008505.1
865 Bacillus cereus ATCC 14579 GCA_000007825.1
850 Bacillus thuringiensis YBT-1518 GCA_000497525.2

lK
643 Clostridium beijerinckii GCA_000833105.2
617 Clostridium saccharoperbutylacetonicum N1-4_28HMT_29 GCA_000340885.1
602 Clostridium saccharobutylicum DSM 13864 GCA_000473995.1
lL
552 Clostridium beijerinckii GCA_000833105.2
534 Clostridium saccharoperbutylacetonicum N1-4_28HMT_29 GCA_000340885.1
514 Clostridium butyricum GCA_001456065.2
514 Clostridium saccharobutylicum DSM 13864 GCA_000473995.1
lM
343 Clostridium beijerinckii GCA_000833105.2
328 Clostridium saccharoperbutylacetonicum N1-4_28HMT_29 GCA_000340885.1
323 Clostridium saccharobutylicum DSM 13864 GCA_000473995.1
lN
714 Lactobacillus gasseri ATCC 33323 = JCM 1131 GCA_000014425.1
638 Lactobacillus hominis DSM 23910 = CRBIP 24.179 GCA_000296835.1
463 Lactobacillus iners DSM 13335 GCA_000160875.1
lO
391 Streptococcus mutans UA159 GCA_000007465.2
286 Streptococcus ratti FA-1 = DSM 20564 GCA_000286075.1
279 Streptococcus ferus DSM 20646 GCA_000372425.1
lP
710 Staphylococcus epidermidis ATCC 12228 GCA_000007645.1
676 Staphylococcus capitis subsp. capitis GCA_001028645.1
659 Staphylococcus haemolyticus JCSC1435 GCA_000009865.1
lQ
860 Staphylococcus epidermidis ATCC 12228 GCA_000007645.1
808 Staphylococcus capitis subsp. capitis GCA_001028645.1
800 Staphylococcus aureus subsp. aureus NCTC 8325 GCA_000013425.1
lR
619 Escherichia coli str. K-12 substr. MG1655 GCA_000005845.2
612 Escherichia coli O104_3AH4 str. 2011C-3493 GCA_000299455.1
599 Shigella flexneri 2a str. 301 GCA_000006925.2
lS
686 Escherichia coli str. K-12 substr. MG1655 GCA_000005845.2
650 Escherichia coli UMN026 GCA_000026325.2
638 Shigella flexneri 2a str. 301 GCA_000006925.2
lT
653 Deinococcus radiodurans R1 GCA_000008565.1
369 Deinococcus deserti VCD115 GCA_000020685.1
349 Deinococcus gobiensis I-0 GCA_000252445.1
lU
515 Staphylococcus epidermidis ATCC 12228 GCA_000007645.1
501 Staphylococcus capitis subsp. capitis GCA_001028645.1
470 Staphylococcus simulans GCA_001559115.1
lV
664 Streptococcus mutans UA159 GCA_000007465.2
496 Streptococcus ratti FA-1 = DSM 20564 GCA_000286075.1
478 Streptococcus gordonii str. Challis substr. CH1 GCA_000017005.1
lW
582 Lactobacillus gasseri ATCC 33323 = JCM 1131 GCA_000014425.1
546 Lactobacillus hominis DSM 23910 = CRBIP 24.179 GCA_000296835.1
455 Lactobacillus helveticus GCA_001308285.1
lX
787 Clostridium beijerinckii GCA_000833105.2
735 Clostridium saccharoperbutylacetonicum N1-4_28HMT_29 GCA_000340885.1
718 Clostridium puniceum GCA_002006345.1
lY
769 Streptococcus mutans UA159 GCA_000007465.2
523 Streptococcus gordonii str. Challis substr. CH1 GCA_000017005.1
520 Streptococcus ratti FA-1 = DSM 20564 GCA_000286075.1
lZ
536 Deinococcus radiodurans R1 GCA_000008565.1
348 Deinococcus gobiensis I-0 GCA_000252445.1
340 Deinococcus soli Cha et al. 2016 GCA_001007995.1
m0
651 Streptococcus mutans UA159 GCA_000007465.2
473 Streptococcus ratti FA-1 = DSM 20564 GCA_000286075.1
440 Streptococcus gordonii str. Challis substr. CH1 GCA_000017005.1
m1
786 Enterococcus faecalis V583 GCA_000007785.1
740 Streptomyces cinnamoneus GCA_001885705.1
675 Enterococcus canis NBRC 100695 GCA_001544375.1
m2
765 Lactobacillus gasseri ATCC 33323 = JCM 1131 GCA_000014425.1
684 Lactobacillus hominis DSM 23910 = CRBIP 24.179 GCA_000296835.1

593 Lactobacillus psittaci DSM 15354 GCA_000425905.1
m3
755 Rhodobacter sphaeroides 2.4.1 GCA_000012905.2
662 Rhodobacter sphaeroides ATCC 17025 GCA_000016405.1
552 Gemmobacter aquatilis GCA_900110025.1
m4
388 Deinococcus radiodurans R1 GCA_000008565.1
292 Deinococcus deserti VCD115 GCA_000020685.1
274 Deinococcus gobiensis I-0 GCA_000252445.1
m5
854 Escherichia coli str. K-12 substr. MG1655 GCA_000005845.2
819 Shigella flexneri 2a str. 301 GCA_000006925.2
813 Escherichia coli O104_3AH4 str. 2011C-3493 GCA_000299455.1
m6
525 Deinococcus radiodurans R1 GCA_000008565.1
351 Deinococcus deserti VCD115 GCA_000020685.1
299 Deinococcus hopiensis KR-140 GCA_900176165.1
m7
955 Staphylococcus epidermidis ATCC 12228 GCA_000007645.1
909 Staphylococcus capitis subsp. capitis GCA_001028645.1
832 Staphylococcus aureus subsp. aureus NCTC 8325 GCA_000013425.1
m8
651 Deinococcus radiodurans R1 GCA_000008565.1
462 Deinococcus deserti VCD115 GCA_000020685.1
434 Deinococcus gobiensis I-0 GCA_000252445.1
m9
769 Streptococcus mutans UA159 GCA_000007465.2
598 Streptococcus ratti FA-1 = DSM 20564 GCA_000286075.1
564 Streptococcus sanguinis SK36 GCA_000014205.1
ma
506 Deinococcus radiodurans R1 GCA_000008565.1
300 Deinococcus deserti VCD115 GCA_000020685.1
296 Deinococcus puniceus GCA_001644565.1
mb
901 Bacillus anthracis str. Ames GCA_000007845.1
901 Bacillus anthracis str. Sterne GCA_000008165.1
893 _5BBacillus thuringiensis_5D serovar konkukian str. 97-27 GCA_000008505.1
880 Bacillus cereus ATCC 14579 GCA_000007825.1
880 Bacillus pseudomycoides DSM 12442 GCA_000161455.1
mc
693 Streptococcus mutans UA159 GCA_000007465.2
476 Streptococcus ferus DSM 20646 GCA_000372425.1
476 Streptococcus ratti FA-1 = DSM 20564 GCA_000286075.1
468 Streptococcus gordonii str. Challis substr. CH1 GCA_000017005.1
md
627 Clostridium beijerinckii GCA_000833105.2
609 Clostridium saccharoperbutylacetonicum N1-4_28HMT_29 GCA_000340885.1
572 Clostridium puniceum GCA_002006345.1
me
840 Bifidobacterium adolescentis ATCC 15703 GCA_000010425.1
684 Bifidobacterium dentium JCM 1195 = DSM 20436 GCA_001042595.1
643 Bifidobacterium angulatum DSM 20098 = JCM 7096 GCA_001025155.1
mf
567 Clostridium beijerinckii GCA_000833105.2
542 Clostridium saccharoperbutylacetonicum N1-4_28HMT_29 GCA_000340885.1
529 Clostridium saccharobutylicum DSM 13864 GCA_000473995.1
mg
612 Streptococcus mutans UA159 GCA_000007465.2
457 Streptococcus gordonii str. Challis substr. CH1 GCA_000017005.1
420 Streptococcus iniae GCA_000831485.1
mh
864 Lactobacillus gasseri ATCC 33323 = JCM 1131 GCA_000014425.1
782 Lactobacillus hominis DSM 23910 = CRBIP 24.179 GCA_000296835.1
544 Lactobacillus iners DSM 13335 GCA_000160875.1
mi
699 Clostridium beijerinckii GCA_000833105.2
645 Clostridium saccharoperbutylacetonicum N1-4_28HMT_29 GCA_000340885.1
629 Clostridium puniceum GCA_002006345.1
mj
749 Lactobacillus gasseri ATCC 33323 = JCM 1131 GCA_000014425.1
711 Lactobacillus hominis DSM 23910 = CRBIP 24.179 GCA_000296835.1
508 Lactobacillus iners DSM 13335 GCA_000160875.1
mk
655 Escherichia coli str. K-12 substr. MG1655 GCA_000005845.2
633 Shigella flexneri 2a str. 301 GCA_000006925.2
631 Escherichia coli O104_3AH4 str. 2011C-3493 GCA_000299455.1

631 Escherichia coli O157_3AH7 str. Sakai GCA_000008865.1
ml
569 Clostridium beijerinckii GCA_000833105.2
549 Clostridium saccharoperbutylacetonicum N1-4_28HMT_29 GCA_000340885.1
541 Clostridium saccharobutylicum DSM 13864 GCA_000473995.1
mm
512 Bacillus thuringiensis YBT-1518 GCA_000497525.2
496 Bacillus anthracis str. Ames GCA_000007845.1
496 Bacillus anthracis str. Sterne GCA_000008165.1
496 _5BBacillus thuringiensis_5D serovar konkukian str. 97-27 GCA_000008505.1
493 Bacillus pseudomycoides DSM 12442 GCA_000161455.1
mn
862 Bacillus anthracis str. Ames GCA_000007845.1
862 Bacillus anthracis str. Sterne GCA_000008165.1
862 _5BBacillus thuringiensis_5D serovar konkukian str. 97-27 GCA_000008505.1
851 Bacillus pseudomycoides DSM 12442 GCA_000161455.1
836 Bacillus cereus ATCC 14579 GCA_000007825.1
836 Bacillus thuringiensis YBT-1518 GCA_000497525.2
mo
627 Enterococcus faecalis V583 GCA_000007785.1
587 Streptomyces cinnamoneus GCA_001885705.1
538 Enterococcus massiliensis GCA_001050095.1
mp
596 Deinococcus radiodurans R1 GCA_000008565.1
368 Deinococcus soli Cha et al. 2016 GCA_001007995.1
360 Deinococcus deserti VCD115 GCA_000020685.1
mq
305 Rhodobacter sphaeroides 2.4.1 GCA_000012905.2
276 Rhodobacter sphaeroides ATCC 17025 GCA_000016405.1
255 Rhodobacter vinaykumarii GCA_900156695.1
mr
963 Staphylococcus epidermidis ATCC 12228 GCA_000007645.1
927 Staphylococcus capitis subsp. capitis GCA_001028645.1
845 Megasphaera cerevisiae DSM 20462 GCA_001045675.1
845 Staphylococcus warneri SG1 GCA_000332735.1
ms
714 Rhodobacter sphaeroides 2.4.1 GCA_000012905.2
599 Rhodobacter sphaeroides ATCC 17025 GCA_000016405.1
483 Defluviimonas alba GCA_001620265.1
mt
543 Staphylococcus epidermidis ATCC 12228 GCA_000007645.1
495 Staphylococcus capitis subsp. capitis GCA_001028645.1
477 Megasphaera cerevisiae DSM 20462 GCA_001045675.1
477 Staphylococcus warneri SG1 GCA_000332735.1
mu
739 Bifidobacterium adolescentis ATCC 15703 GCA_000010425.1
557 Bifidobacterium dentium JCM 1195 = DSM 20436 GCA_001042595.1
529 Bifidobacterium longum NCC2705 GCA_000007525.1
mv
866 Streptococcus mutans UA159 GCA_000007465.2
636 Streptococcus ratti FA-1 = DSM 20564 GCA_000286075.1
585 Streptococcus sanguinis SK36 GCA_000014205.1
mw
870 Bifidobacterium adolescentis ATCC 15703 GCA_000010425.1
744 Bifidobacterium dentium JCM 1195 = DSM 20436 GCA_001042595.1
657 Bifidobacterium callitrichos DSM 23973 GCA_000741175.1
mx
598 Rhodobacter sphaeroides 2.4.1 GCA_000012905.2
531 Rhodobacter sphaeroides ATCC 17025 GCA_000016405.1
468 Pseudorhodobacter ferrugineus DSM 5888 GCA_000420745.1
my
815 Staphylococcus epidermidis ATCC 12228 GCA_000007645.1
762 Staphylococcus capitis subsp. capitis GCA_001028645.1
683 Staphylococcus haemolyticus JCSC1435 GCA_000009865.1
mz
582 Streptococcus mutans UA159 GCA_000007465.2
444 Streptococcus ratti FA-1 = DSM 20564 GCA_000286075.1
401 Streptococcus henryi DSM 19005 GCA_000376985.1
mA
530 Streptococcus mutans UA159 GCA_000007465.2
378 Streptococcus ratti FA-1 = DSM 20564 GCA_000286075.1
345 Streptococcus ferus DSM 20646 GCA_000372425.1
mB
768 Clostridium beijerinckii GCA_000833105.2
738 Clostridium saccharoperbutylacetonicum N1-4_28HMT_29 GCA_000340885.1
673 Clostridium puniceum GCA_002006345.1

mC
628 Lactobacillus gasseri ATCC 33323 = JCM 1131 GCA_000014425.1
543 Lactobacillus hominis DSM 23910 = CRBIP 24.179 GCA_000296835.1
389 Lactobacillus iners DSM 13335 GCA_000160875.1
mD
798 Bacillus anthracis str. Ames GCA_000007845.1
798 Bacillus anthracis str. Sterne GCA_000008165.1
798 _5BBacillus thuringiensis_5D serovar konkukian str. 97-27 GCA_000008505.1
778 Bacillus cereus ATCC 14579 GCA_000007825.1
762 Bacillus thuringiensis YBT-1518 GCA_000497525.2
mE
731 Rhodobacter sphaeroides 2.4.1 GCA_000012905.2
609 Rhodobacter sphaeroides ATCC 17025 GCA_000016405.1
537 Gemmobacter megaterium GCA_900156815.1
mF
725 Escherichia coli O157_3AH7 str. Sakai GCA_000008865.1
725 Escherichia coli UMN026 GCA_000026325.2
725 Escherichia coli str. K-12 substr. MG1655 GCA_000005845.2
710 Shigella flexneri 2a str. 301 GCA_000006925.2
705 Escherichia coli O83_3AH1 str. NRG 857C GCA_000183345.1
mG
292 Streptococcus mutans UA159 GCA_000007465.2
248 Streptococcus salivarius GCA_000785515.1
248 Streptococcus thermophilus JIM 8232 GCA_000253395.1
247 Streptococcus criceti HS-6 GCA_000187975.3
247 Streptococcus gordonii str. Challis substr. CH1 GCA_000017005.1
mH
785 Deinococcus radiodurans R1 GCA_000008565.1
532 Deinococcus hopiensis KR-140 GCA_900176165.1
527 Deinococcus soli Cha et al. 2016 GCA_001007995.1
mI
675 Clostridium beijerinckii GCA_000833105.2
629 Clostridium saccharobutylicum DSM 13864 GCA_000473995.1
620 Clostridium saccharoperbutylacetonicum N1-4_28HMT_29 GCA_000340885.1
mJ
705 Staphylococcus epidermidis ATCC 12228 GCA_000007645.1
666 Staphylococcus capitis subsp. capitis GCA_001028645.1
642 Staphylococcus aureus subsp. aureus NCTC 8325 GCA_000013425.1
mK
587 Lactobacillus gasseri ATCC 33323 = JCM 1131 GCA_000014425.1
572 Lactobacillus hominis DSM 23910 = CRBIP 24.179 GCA_000296835.1
414 Lactobacillus jensenii GCA_001936235.1
414 Lactobacillus psittaci DSM 15354 GCA_000425905.1
mL
743 Clostridium beijerinckii GCA_000833105.2
724 Clostridium saccharoperbutylacetonicum N1-4_28HMT_29 GCA_000340885.1
693 Clostridium saccharobutylicum DSM 13864 GCA_000473995.1
mM
576 Rhodobacter sphaeroides 2.4.1 GCA_000012905.2
526 Rhodobacter sphaeroides ATCC 17025 GCA_000016405.1
408 Gemmobacter aquatilis GCA_900110025.1
mN
896 Staphylococcus epidermidis ATCC 12228 GCA_000007645.1
823 Staphylococcus capitis subsp. capitis GCA_001028645.1
759 Staphylococcus aureus subsp. aureus NCTC 8325 GCA_000013425.1
mO
639 Streptococcus mutans UA159 GCA_000007465.2
486 Streptococcus ratti FA-1 = DSM 20564 GCA_000286075.1
461 Streptococcus macacae NCTC 11558 GCA_000187995.3
mP
484 Bifidobacterium adolescentis ATCC 15703 GCA_000010425.1
422 Bifidobacterium dentium JCM 1195 = DSM 20436 GCA_001042595.1
375 Bifidobacterium stellenboschense GCA_000741785.1
mQ
589 Streptococcus mutans UA159 GCA_000007465.2
425 Streptococcus ratti FA-1 = DSM 20564 GCA_000286075.1
403 Streptococcus gordonii str. Challis substr. CH1 GCA_000017005.1
mR
660 Lactobacillus gasseri ATCC 33323 = JCM 1131 GCA_000014425.1
653 Lactobacillus hominis DSM 23910 = CRBIP 24.179 GCA_000296835.1
444 Lactobacillus hamsteri DSM 5661 = JCM 6256 GCA_000615445.1
mS
642 Rhodobacter sphaeroides 2.4.1 GCA_000012905.2
587 Rhodobacter sphaeroides ATCC 17025 GCA_000016405.1
492 Gemmobacter aquatilis GCA_900110025.1
mT

595 Bacillus thuringiensis YBT-1518 GCA_000497525.2
574 Bacillus anthracis str. Ames GCA_000007845.1
574 Bacillus anthracis str. Sterne GCA_000008165.1
574 Bacillus pseudomycoides DSM 12442 GCA_000161455.1
574 _5BBacillus thuringiensis_5D serovar konkukian str. 97-27 GCA_000008505.1
558 Bacillus cereus ATCC 14579 GCA_000007825.1
mU
616 Clostridium beijerinckii GCA_000833105.2
562 Clostridium saccharoperbutylacetonicum N1-4_28HMT_29 GCA_000340885.1
540 Clostridium butyricum GCA_001456065.2
mV
613 Clostridium beijerinckii GCA_000833105.2
595 Clostridium saccharoperbutylacetonicum N1-4_28HMT_29 GCA_000340885.1
580 Clostridium puniceum GCA_002006345.1
mW
585 Lactobacillus gasseri ATCC 33323 = JCM 1131 GCA_000014425.1
558 Lactobacillus hominis DSM 23910 = CRBIP 24.179 GCA_000296835.1
394 Lactobacillus gallinarum GCA_001314245.2
mX
606 Deinococcus radiodurans R1 GCA_000008565.1
414 Deinococcus deserti VCD115 GCA_000020685.1
363 Deinococcus hopiensis KR-140 GCA_900176165.1
mY
755 Clostridium beijerinckii GCA_000833105.2
720 Clostridium saccharoperbutylacetonicum N1-4_28HMT_29 GCA_000340885.1
693 Clostridium saccharobutylicum DSM 13864 GCA_000473995.1
mZ
411 Deinococcus radiodurans R1 GCA_000008565.1
259 Deinococcus deserti VCD115 GCA_000020685.1
236 Deinococcus murrayi DSM 11303 GCA_000482805.1
n0
429 Rhodobacter sphaeroides 2.4.1 GCA_000012905.2
393 Rhodobacter sphaeroides ATCC 17025 GCA_000016405.1
273 Defluviimonas alba GCA_001620265.1
n1
767 Staphylococcus epidermidis ATCC 12228 GCA_000007645.1
712 Staphylococcus capitis subsp. capitis GCA_001028645.1
618 Staphylococcus haemolyticus JCSC1435 GCA_000009865.1
n2
644 Staphylococcus epidermidis ATCC 12228 GCA_000007645.1
613 Staphylococcus capitis subsp. capitis GCA_001028645.1
590 Staphylococcus haemolyticus JCSC1435 GCA_000009865.1
n3
750 Staphylococcus epidermidis ATCC 12228 GCA_000007645.1
710 Staphylococcus capitis subsp. capitis GCA_001028645.1
639 Staphylococcus warneri SG1 GCA_000332735.1
n4
908 Lactobacillus gasseri ATCC 33323 = JCM 1131 GCA_000014425.1
826 Lactobacillus hominis DSM 23910 = CRBIP 24.179 GCA_000296835.1
633 Lactobacillus iners DSM 13335 GCA_000160875.1
n5
487 Enterococcus faecalis V583 GCA_000007785.1
455 Streptomyces cinnamoneus GCA_001885705.1
404 Enterococcus faecium DO GCA_000174395.2
n6
670 Clostridium beijerinckii GCA_000833105.2
610 Clostridium saccharoperbutylacetonicum N1-4_28HMT_29 GCA_000340885.1
600 Clostridium butyricum GCA_001456065.2
n7
558 Staphylococcus epidermidis ATCC 12228 GCA_000007645.1
524 Staphylococcus capitis subsp. capitis GCA_001028645.1
504 Staphylococcus pettenkoferi GCA_002208805.1
n8
649 Clostridium beijerinckii GCA_000833105.2
617 Clostridium saccharoperbutylacetonicum N1-4_28HMT_29 GCA_000340885.1
592 Clostridium puniceum GCA_002006345.1
n9
949 Streptococcus mutans UA159 GCA_000007465.2
707 Streptococcus ratti FA-1 = DSM 20564 GCA_000286075.1
655 Streptococcus gordonii str. Challis substr. CH1 GCA_000017005.1
na
590 Clostridium beijerinckii GCA_000833105.2
584 Clostridium saccharoperbutylacetonicum N1-4_28HMT_29 GCA_000340885.1
581 Clostridium saccharobutylicum DSM 13864 GCA_000473995.1
nb
678 Deinococcus radiodurans R1 GCA_000008565.1

423 Deinococcus deserti VCD115 GCA_000020685.1
409 Deinococcus hopiensis KR-140 GCA_900176165.1
nc
357 Streptococcus mutans UA159 GCA_000007465.2
214 Streptococcus ferus DSM 20646 GCA_000372425.1
199 Streptococcus dysgalactiae subsp. equisimilis AC-2713 GCA_000317855.1
nd
820 Bacillus cereus ATCC 14579 GCA_000007825.1
805 Bacillus anthracis str. Ames GCA_000007845.1
805 Bacillus anthracis str. Sterne GCA_000008165.1
805 Bacillus pseudomycoides DSM 12442 GCA_000161455.1
805 _5BBacillus thuringiensis_5D serovar konkukian str. 97-27 GCA_000008505.1
800 Bacillus mycoides GCA_000832605.1
ne
781 Deinococcus radiodurans R1 GCA_000008565.1
455 Deinococcus deserti VCD115 GCA_000020685.1
449 Deinococcus hopiensis KR-140 GCA_900176165.1
nf
481 Deinococcus radiodurans R1 GCA_000008565.1
309 Deinococcus deserti VCD115 GCA_000020685.1
305 Deinococcus puniceus GCA_001644565.1
ng
498 Staphylococcus epidermidis ATCC 12228 GCA_000007645.1
478 Staphylococcus capitis subsp. capitis GCA_001028645.1
442 Staphylococcus haemolyticus JCSC1435 GCA_000009865.1
442 Staphylococcus hominis subsp. hominis C80 GCA_000183685.1
nh
651 Lactobacillus gasseri ATCC 33323 = JCM 1131 GCA_000014425.1
598 Lactobacillus hominis DSM 23910 = CRBIP 24.179 GCA_000296835.1
466 Lactobacillus iners DSM 13335 GCA_000160875.1
ni
730 Clostridium beijerinckii GCA_000833105.2
720 Clostridium saccharoperbutylacetonicum N1-4_28HMT_29 GCA_000340885.1
702 Clostridium saccharobutylicum DSM 13864 GCA_000473995.1
nj
810 Deinococcus radiodurans R1 GCA_000008565.1
508 Deinococcus deserti VCD115 GCA_000020685.1
504 Deinococcus puniceus GCA_001644565.1
nk
648 Lactobacillus gasseri ATCC 33323 = JCM 1131 GCA_000014425.1
610 Lactobacillus hominis DSM 23910 = CRBIP 24.179 GCA_000296835.1
430 Lactobacillus iners DSM 13335 GCA_000160875.1
nl
603 Lactobacillus gasseri ATCC 33323 = JCM 1131 GCA_000014425.1
555 Lactobacillus hominis DSM 23910 = CRBIP 24.179 GCA_000296835.1
416 Lactobacillus hamsteri DSM 5661 = JCM 6256 GCA_000615445.1
nm
919 Streptococcus mutans UA159 GCA_000007465.2
673 Streptococcus ratti FA-1 = DSM 20564 GCA_000286075.1
619 Streptococcus criceti HS-6 GCA_000187975.3
nn
658 Lactobacillus gasseri ATCC 33323 = JCM 1131 GCA_000014425.1
598 Lactobacillus hominis DSM 23910 = CRBIP 24.179 GCA_000296835.1
468 Lactobacillus iners DSM 13335 GCA_000160875.1
no
641 Rhodobacter sphaeroides 2.4.1 GCA_000012905.2
523 Rhodobacter sphaeroides ATCC 17025 GCA_000016405.1
450 Gemmobacter aquatilis GCA_900110025.1
450 Gemmobacter megaterium GCA_900156815.1
np
642 Lactobacillus gasseri ATCC 33323 = JCM 1131 GCA_000014425.1
601 Lactobacillus hominis DSM 23910 = CRBIP 24.179 GCA_000296835.1
463 Lactobacillus hamsteri DSM 5661 = JCM 6256 GCA_000615445.1
nq
782 Clostridium beijerinckii GCA_000833105.2
738 Clostridium saccharoperbutylacetonicum N1-4_28HMT_29 GCA_000340885.1
735 Clostridium saccharobutylicum DSM 13864 GCA_000473995.1
nr
821 Staphylococcus epidermidis ATCC 12228 GCA_000007645.1
788 Staphylococcus capitis subsp. capitis GCA_001028645.1
698 Staphylococcus lugdunensis HKU09-01 GCA_000025085.1
ns
850 Deinococcus radiodurans R1 GCA_000008565.1
507 Deinococcus gobiensis I-0 GCA_000252445.1
488 Deinococcus proteolyticus MRP GCA_000190555.1
nt

855 Enterococcus faecalis V583 GCA_000007785.1
798 Enterococcus rivorum GCA_001742285.1
793 Streptomyces cinnamoneus GCA_001885705.1
nu
701 Rhodobacter sphaeroides 2.4.1 GCA_000012905.2
597 Rhodobacter sphaeroides ATCC 17025 GCA_000016405.1
476 Pseudorhodobacter ferrugineus DSM 5888 GCA_000420745.1
nv
606 Staphylococcus epidermidis ATCC 12228 GCA_000007645.1
559 Staphylococcus capitis subsp. capitis GCA_001028645.1
533 Staphylococcus hominis subsp. hominis C80 GCA_000183685.1
nw
777 Streptococcus mutans UA159 GCA_000007465.2
511 Streptococcus ratti FA-1 = DSM 20564 GCA_000286075.1
480 Streptococcus gordonii str. Challis substr. CH1 GCA_000017005.1
nx
541 Bifidobacterium adolescentis ATCC 15703 GCA_000010425.1
435 Bifidobacterium dentium JCM 1195 = DSM 20436 GCA_001042595.1
397 Bifidobacterium callitrichos DSM 23973 GCA_000741175.1
ny
518 Enterococcus faecalis V583 GCA_000007785.1
472 Streptomyces cinnamoneus GCA_001885705.1
458 Enterococcus rivorum GCA_001742285.1
nz
442 Escherichia coli str. K-12 substr. MG1655 GCA_000005845.2
412 Escherichia coli O104_3AH4 str. 2011C-3493 GCA_000299455.1
411 Escherichia coli UMN026 GCA_000026325.2
nA
821 Lactobacillus gasseri ATCC 33323 = JCM 1131 GCA_000014425.1
755 Lactobacillus hominis DSM 23910 = CRBIP 24.179 GCA_000296835.1
574 Lactobacillus iners DSM 13335 GCA_000160875.1
nB
818 Clostridium beijerinckii GCA_000833105.2
757 Clostridium saccharoperbutylacetonicum N1-4_28HMT_29 GCA_000340885.1
748 Clostridium saccharobutylicum DSM 13864 GCA_000473995.1
nC
728 Deinococcus radiodurans R1 GCA_000008565.1
440 Deinococcus deserti VCD115 GCA_000020685.1
437 Deinococcus soli Cha et al. 2016 GCA_001007995.1
nD
775 Enterococcus faecalis V583 GCA_000007785.1
732 Enterococcus rivorum GCA_001742285.1
728 Streptomyces cinnamoneus GCA_001885705.1
nE
733 Streptococcus mutans UA159 GCA_000007465.2
556 Streptococcus gordonii str. Challis substr. CH1 GCA_000017005.1
555 Streptococcus pseudoporcinus LQ 940-04 GCA_000188035.3
555 Streptococcus ratti FA-1 = DSM 20564 GCA_000286075.1
nF
764 Rhodobacter sphaeroides 2.4.1 GCA_000012905.2
711 Rhodobacter sphaeroides ATCC 17025 GCA_000016405.1
530 Pseudorhodobacter psychrotolerans GCA_001294535.1
nG
945 Streptococcus mutans UA159 GCA_000007465.2
669 Streptococcus ratti FA-1 = DSM 20564 GCA_000286075.1
661 Streptococcus gordonii str. Challis substr. CH1 GCA_000017005.1
nH
776 Streptococcus mutans UA159 GCA_000007465.2
570 Streptococcus ratti FA-1 = DSM 20564 GCA_000286075.1
536 Streptococcus gallolyticus subsp. gallolyticus DSM 16831 GCA_002000985.1
nI
928 Lactobacillus gasseri ATCC 33323 = JCM 1131 GCA_000014425.1
809 Lactobacillus hominis DSM 23910 = CRBIP 24.179 GCA_000296835.1
626 Lactobacillus iners DSM 13335 GCA_000160875.1
nJ
596 Rhodobacter sphaeroides 2.4.1 GCA_000012905.2
542 Rhodobacter sphaeroides ATCC 17025 GCA_000016405.1
484 Gemmobacter aquatilis GCA_900110025.1
nK
649 Bacillus anthracis str. Ames GCA_000007845.1
649 Bacillus anthracis str. Sterne GCA_000008165.1
649 _5BBacillus thuringiensis_5D serovar konkukian str. 97-27 GCA_000008505.1
641 Bacillus pseudomycoides DSM 12442 GCA_000161455.1
639 Bacillus thuringiensis YBT-1518 GCA_000497525.2
nL
788 Lactobacillus gasseri ATCC 33323 = JCM 1131 GCA_000014425.1

757 Lactobacillus hominis DSM 23910 = CRBIP 24.179 GCA_000296835.1
582 Lactobacillus iners DSM 13335 GCA_000160875.1
nM
769 Escherichia coli str. K-12 substr. MG1655 GCA_000005845.2
754 Escherichia coli O157_3AH7 str. Sakai GCA_000008865.1
754 Shigella flexneri 2a str. 301 GCA_000006925.2
751 Escherichia coli UMN026 GCA_000026325.2
nN
619 Escherichia coli O157_3AH7 str. Sakai GCA_000008865.1
619 Escherichia coli str. K-12 substr. MG1655 GCA_000005845.2
604 Shigella flexneri 2a str. 301 GCA_000006925.2
602 Escherichia coli UMN026 GCA_000026325.2
nO
696 Deinococcus radiodurans R1 GCA_000008565.1
492 Deinococcus deserti VCD115 GCA_000020685.1
467 Deinococcus gobiensis I-0 GCA_000252445.1
nP
733 Staphylococcus epidermidis ATCC 12228 GCA_000007645.1
668 Staphylococcus capitis subsp. capitis GCA_001028645.1
626 Staphylococcus aureus subsp. aureus NCTC 8325 GCA_000013425.1
nQ
613 Lactobacillus gasseri ATCC 33323 = JCM 1131 GCA_000014425.1
590 Lactobacillus hominis DSM 23910 = CRBIP 24.179 GCA_000296835.1
402 Lactobacillus iners DSM 13335 GCA_000160875.1
nR
718 Lactobacillus gasseri ATCC 33323 = JCM 1131 GCA_000014425.1
670 Lactobacillus hominis DSM 23910 = CRBIP 24.179 GCA_000296835.1
479 Lactobacillus jensenii GCA_001936235.1
479 Lactobacillus psittaci DSM 15354 GCA_000425905.1
nS
481 Staphylococcus epidermidis ATCC 12228 GCA_000007645.1
449 Staphylococcus capitis subsp. capitis GCA_001028645.1
424 Staphylococcus aureus subsp. aureus NCTC 8325 GCA_000013425.1
nT
859 Clostridium beijerinckii GCA_000833105.2
848 Clostridium saccharoperbutylacetonicum N1-4_28HMT_29 GCA_000340885.1
817 Clostridium saccharobutylicum DSM 13864 GCA_000473995.1
nU
601 Clostridium beijerinckii GCA_000833105.2
550 Clostridium saccharoperbutylacetonicum N1-4_28HMT_29 GCA_000340885.1
528 Clostridium chromiireducens GCA_002029255.1
nV
582 Clostridium saccharoperbutylacetonicum N1-4_28HMT_29 GCA_000340885.1
571 Clostridium beijerinckii GCA_000833105.2
570 Clostridium saccharobutylicum DSM 13864 GCA_000473995.1
nW
641 Clostridium beijerinckii GCA_000833105.2
639 Clostridium saccharoperbutylacetonicum N1-4_28HMT_29 GCA_000340885.1
622 Clostridium puniceum GCA_002006345.1
622 Clostridium saccharobutylicum DSM 13864 GCA_000473995.1
nX
779 Deinococcus radiodurans R1 GCA_000008565.1
466 Deinococcus deserti VCD115 GCA_000020685.1
412 Deinococcus proteolyticus MRP GCA_000190555.1
nY
711 Deinococcus radiodurans R1 GCA_000008565.1
412 Deinococcus deserti VCD115 GCA_000020685.1
393 Deinococcus soli Cha et al. 2016 GCA_001007995.1
nZ
739 Escherichia coli str. K-12 substr. MG1655 GCA_000005845.2
733 Escherichia coli IAI39 GCA_000026345.1
733 Shigella flexneri 2a str. 301 GCA_000006925.2
717 Escherichia coli O104_3AH4 str. 2011C-3493 GCA_000299455.1
717 Escherichia coli O157_3AH7 str. Sakai GCA_000008865.1
o0
863 Clostridium beijerinckii GCA_000833105.2
859 Clostridium saccharoperbutylacetonicum N1-4_28HMT_29 GCA_000340885.1
846 Clostridium puniceum GCA_002006345.1
o1
516 Clostridium beijerinckii GCA_000833105.2
515 Clostridium saccharoperbutylacetonicum N1-4_28HMT_29 GCA_000340885.1
496 Clostridium butyricum GCA_001456065.2
o2
518 Staphylococcus epidermidis ATCC 12228 GCA_000007645.1
488 Staphylococcus capitis subsp. capitis GCA_001028645.1
488 Staphylococcus haemolyticus JCSC1435 GCA_000009865.1

487 Staphylococcus hominis subsp. hominis C80 GCA_000183685.1
o3
807 Enterococcus faecalis V583 GCA_000007785.1
760 Streptomyces cinnamoneus GCA_001885705.1
719 Enterococcus massiliensis GCA_001050095.1
o4
519 Rhodobacter sphaeroides 2.4.1 GCA_000012905.2
504 Rhodobacter sphaeroides ATCC 17025 GCA_000016405.1
441 Pseudorhodobacter psychrotolerans GCA_001294535.1
o5
585 Escherichia coli str. K-12 substr. MG1655 GCA_000005845.2
580 Escherichia coli O157_3AH7 str. Sakai GCA_000008865.1
580 Shigella flexneri 2a str. 301 GCA_000006925.2
579 Escherichia coli O83_3AH1 str. NRG 857C GCA_000183345.1
579 Escherichia coli UMN026 GCA_000026325.2
o6
742 Deinococcus radiodurans R1 GCA_000008565.1
485 Deinococcus murrayi DSM 11303 GCA_000482805.1
480 Deinococcus deserti VCD115 GCA_000020685.1
o7
615 Staphylococcus epidermidis ATCC 12228 GCA_000007645.1
564 Staphylococcus capitis subsp. capitis GCA_001028645.1
556 Staphylococcus sciuri GCA_002209165.1
556 Staphylococcus warneri SG1 GCA_000332735.1
o8
726 Lactobacillus gasseri ATCC 33323 = JCM 1131 GCA_000014425.1
652 Lactobacillus hominis DSM 23910 = CRBIP 24.179 GCA_000296835.1
464 Lactobacillus iners DSM 13335 GCA_000160875.1
o9
720 Streptococcus mutans UA159 GCA_000007465.2
564 Streptococcus ratti FA-1 = DSM 20564 GCA_000286075.1
517 Streptococcus equinus GCA_000964315.1
517 Streptococcus gallolyticus subsp. gallolyticus DSM 16831 GCA_002000985.1
oa
815 Lactobacillus gasseri ATCC 33323 = JCM 1131 GCA_000014425.1
773 Lactobacillus hominis DSM 23910 = CRBIP 24.179 GCA_000296835.1
592 Lactobacillus hamsteri DSM 5661 = JCM 6256 GCA_000615445.1
ob
671 Escherichia coli O157_3AH7 str. Sakai GCA_000008865.1
671 Escherichia coli str. K-12 substr. MG1655 GCA_000005845.2
660 Escherichia coli O104_3AH4 str. 2011C-3493 GCA_000299455.1
660 Shigella flexneri 2a str. 301 GCA_000006925.2
651 Escherichia coli O83_3AH1 str. NRG 857C GCA_000183345.1
oc
570 Clostridium beijerinckii GCA_000833105.2
555 Clostridium saccharoperbutylacetonicum N1-4_28HMT_29 GCA_000340885.1
533 Clostridium saccharobutylicum DSM 13864 GCA_000473995.1
od
725 Lactobacillus gasseri ATCC 33323 = JCM 1131 GCA_000014425.1
660 Lactobacillus hominis DSM 23910 = CRBIP 24.179 GCA_000296835.1
478 Lactobacillus acetotolerans GCA_001042405.1
oe
769 Enterococcus faecalis V583 GCA_000007785.1
722 Streptomyces cinnamoneus GCA_001885705.1
703 Enterococcus phoeniculicola ATCC BAA-412 GCA_000407505.1
of
657 Lactobacillus gasseri ATCC 33323 = JCM 1131 GCA_000014425.1
576 Lactobacillus hominis DSM 23910 = CRBIP 24.179 GCA_000296835.1
441 Lactobacillus iners DSM 13335 GCA_000160875.1
og
792 Lactobacillus gasseri ATCC 33323 = JCM 1131 GCA_000014425.1
698 Lactobacillus hominis DSM 23910 = CRBIP 24.179 GCA_000296835.1
538 Lactobacillus iners DSM 13335 GCA_000160875.1
oh
626 Rhodobacter sphaeroides 2.4.1 GCA_000012905.2
554 Rhodobacter sphaeroides ATCC 17025 GCA_000016405.1
460 Gemmobacter megaterium GCA_900156815.1
oi
417 Rhodobacter sphaeroides 2.4.1 GCA_000012905.2
390 Rhodobacter sphaeroides ATCC 17025 GCA_000016405.1
283 Pseudorhodobacter wandonensis GCA_001202035.1
oj
838 Lactobacillus gasseri ATCC 33323 = JCM 1131 GCA_000014425.1
763 Lactobacillus hominis DSM 23910 = CRBIP 24.179 GCA_000296835.1
574 Lactobacillus iners DSM 13335 GCA_000160875.1
ok

695 Clostridium beijerinckii GCA_000833105.2
664 Clostridium saccharoperbutylacetonicum N1-4_28HMT_29 GCA_000340885.1
621 Clostridium puniceum GCA_002006345.1
ol
541 Streptococcus mutans UA159 GCA_000007465.2
371 Streptococcus ratti FA-1 = DSM 20564 GCA_000286075.1
335 Streptococcus gallolyticus subsp. gallolyticus DSM 16831 GCA_002000985.1
om
706 Clostridium beijerinckii GCA_000833105.2
668 Clostridium saccharobutylicum DSM 13864 GCA_000473995.1
665 Clostridium saccharoperbutylacetonicum N1-4_28HMT_29 GCA_000340885.1
on
775 Lactobacillus gasseri ATCC 33323 = JCM 1131 GCA_000014425.1
710 Lactobacillus hominis DSM 23910 = CRBIP 24.179 GCA_000296835.1
542 Lactobacillus iners DSM 13335 GCA_000160875.1
oo
691 Deinococcus radiodurans R1 GCA_000008565.1
423 Deinococcus deserti VCD115 GCA_000020685.1
390 Deinococcus hopiensis KR-140 GCA_900176165.1
op
822 Clostridium saccharoperbutylacetonicum N1-4_28HMT_29 GCA_000340885.1
817 Clostridium beijerinckii GCA_000833105.2
789 Clostridium saccharobutylicum DSM 13864 GCA_000473995.1
oq
513 Staphylococcus capitis subsp. capitis GCA_001028645.1
513 Staphylococcus epidermidis ATCC 12228 GCA_000007645.1
485 Megasphaera cerevisiae DSM 20462 GCA_001045675.1
485 Staphylococcus warneri SG1 GCA_000332735.1
458 Staphylococcus saprophyticus subsp. saprophyticus ATCC 15305 GCA_000010125.1
458 Staphylococcus xylosus GCA_000706685.1
or
1047 Lactobacillus gasseri ATCC 33323 = JCM 1131 GCA_000014425.1
930 Lactobacillus hominis DSM 23910 = CRBIP 24.179 GCA_000296835.1
692 Lactobacillus iners DSM 13335 GCA_000160875.1
os
717 Streptococcus mutans UA159 GCA_000007465.2
488 Streptococcus gordonii str. Challis substr. CH1 GCA_000017005.1
468 Streptococcus ferus DSM 20646 GCA_000372425.1
ot
527 Streptococcus mutans UA159 GCA_000007465.2
374 Streptococcus ratti FA-1 = DSM 20564 GCA_000286075.1
360 Streptococcus ovis DSM 16829 GCA_000380125.1
ou
706 Rhodobacter sphaeroides 2.4.1 GCA_000012905.2
598 Rhodobacter sphaeroides ATCC 17025 GCA_000016405.1
482 Gemmobacter aquatilis GCA_900110025.1
ov
666 Escherichia coli O104_3AH4 str. 2011C-3493 GCA_000299455.1
666 Escherichia coli str. K-12 substr. MG1655 GCA_000005845.2
659 Shigella flexneri 2a str. 301 GCA_000006925.2
646 Escherichia coli O157_3AH7 str. Sakai GCA_000008865.1
ow
621 Streptococcus mutans UA159 GCA_000007465.2
371 Streptococcus gordonii str. Challis substr. CH1 GCA_000017005.1
369 Streptococcus equinus GCA_000964315.1
ox
825 Staphylococcus epidermidis ATCC 12228 GCA_000007645.1
798 Staphylococcus capitis subsp. capitis GCA_001028645.1
704 Staphylococcus haemolyticus JCSC1435 GCA_000009865.1
704 Staphylococcus hominis subsp. hominis C80 GCA_000183685.1
oy
609 Staphylococcus epidermidis ATCC 12228 GCA_000007645.1
568 Staphylococcus capitis subsp. capitis GCA_001028645.1
524 Staphylococcus lutrae GCA_002101335.1
oz
705 Enterococcus faecalis V583 GCA_000007785.1
660 Streptomyces cinnamoneus GCA_001885705.1
623 Enterococcus rivorum GCA_001742285.1
oA
606 Bifidobacterium adolescentis ATCC 15703 GCA_000010425.1
576 Bifidobacterium dentium JCM 1195 = DSM 20436 GCA_001042595.1
539 Bifidobacterium angulatum DSM 20098 = JCM 7096 GCA_001025155.1
539 Bifidobacterium longum NCC2705 GCA_000007525.1
oB
644 Streptococcus mutans UA159 GCA_000007465.2
427 Streptococcus ratti FA-1 = DSM 20564 GCA_000286075.1

390 Streptococcus equinus GCA_000964315.1
oC
604 Streptococcus mutans UA159 GCA_000007465.2
500 Streptococcus ratti FA-1 = DSM 20564 GCA_000286075.1
481 Streptococcus macacae NCTC 11558 GCA_000187995.3
oD
687 Clostridium beijerinckii GCA_000833105.2
630 Clostridium saccharoperbutylacetonicum N1-4_28HMT_29 GCA_000340885.1
622 Clostridium puniceum GCA_002006345.1
oE
566 Streptococcus mutans UA159 GCA_000007465.2
386 Streptococcus gordonii str. Challis substr. CH1 GCA_000017005.1
371 Streptococcus ratti FA-1 = DSM 20564 GCA_000286075.1
oF
732 Lactobacillus gasseri ATCC 33323 = JCM 1131 GCA_000014425.1
666 Lactobacillus hominis DSM 23910 = CRBIP 24.179 GCA_000296835.1
486 Lactobacillus iners DSM 13335 GCA_000160875.1
oG
682 Deinococcus radiodurans R1 GCA_000008565.1
441 Deinococcus gobiensis I-0 GCA_000252445.1
440 Deinococcus deserti VCD115 GCA_000020685.1
oH
700 Lactobacillus gasseri ATCC 33323 = JCM 1131 GCA_000014425.1
632 Lactobacillus hominis DSM 23910 = CRBIP 24.179 GCA_000296835.1
471 Lactobacillus iners DSM 13335 GCA_000160875.1
oI
508 Staphylococcus epidermidis ATCC 12228 GCA_000007645.1
490 Staphylococcus capitis subsp. capitis GCA_001028645.1
435 Megasphaera cerevisiae DSM 20462 GCA_001045675.1
435 Staphylococcus warneri SG1 GCA_000332735.1
oJ
889 Deinococcus radiodurans R1 GCA_000008565.1
524 Deinococcus deserti VCD115 GCA_000020685.1
513 Deinococcus gobiensis I-0 GCA_000252445.1
oK
777 Clostridium beijerinckii GCA_000833105.2
768 Clostridium saccharoperbutylacetonicum N1-4_28HMT_29 GCA_000340885.1
744 Clostridium saccharobutylicum DSM 13864 GCA_000473995.1
oL
730 Enterococcus faecalis V583 GCA_000007785.1
683 Streptomyces cinnamoneus GCA_001885705.1
645 Enterococcus massiliensis GCA_001050095.1
oM
805 Streptococcus mutans UA159 GCA_000007465.2
575 Streptococcus ratti FA-1 = DSM 20564 GCA_000286075.1
527 Streptococcus halotolerans GCA_001598035.1
oN
724 Streptococcus mutans UA159 GCA_000007465.2
554 Streptococcus ratti FA-1 = DSM 20564 GCA_000286075.1
507 Streptococcus equinus GCA_000964315.1
oO
609 Streptococcus mutans UA159 GCA_000007465.2
464 Streptococcus ratti FA-1 = DSM 20564 GCA_000286075.1
422 Streptococcus macacae NCTC 11558 GCA_000187995.3
oP
798 Escherichia coli str. K-12 substr. MG1655 GCA_000005845.2
766 Escherichia coli O104_3AH4 str. 2011C-3493 GCA_000299455.1
766 Shigella flexneri 2a str. 301 GCA_000006925.2
740 Escherichia coli O157_3AH7 str. Sakai GCA_000008865.1
740 Escherichia coli O83_3AH1 str. NRG 857C GCA_000183345.1
oQ
456 Streptococcus mutans UA159 GCA_000007465.2
269 Streptococcus gordonii str. Challis substr. CH1 GCA_000017005.1
237 Streptococcus ratti FA-1 = DSM 20564 GCA_000286075.1
oR
416 Bifidobacterium adolescentis ATCC 15703 GCA_000010425.1
377 Bifidobacterium callitrichos DSM 23973 GCA_000741175.1
372 Bifidobacterium gallicum DSM 20093 = LMG 11596 GCA_000741205.1
oS
570 Enterococcus faecalis V583 GCA_000007785.1
527 Enterococcus haemoperoxidus ATCC BAA-382 GCA_000407165.1
527 Enterococcus rivorum GCA_001742285.1
524 Streptomyces cinnamoneus GCA_001885705.1
oT
777 Clostridium beijerinckii GCA_000833105.2
758 Clostridium saccharoperbutylacetonicum N1-4_28HMT_29 GCA_000340885.1

734 Clostridium saccharobutylicum DSM 13864 GCA_000473995.1
oU
660 Enterococcus faecalis V583 GCA_000007785.1
599 Streptomyces cinnamoneus GCA_001885705.1
537 Enterococcus haemoperoxidus ATCC BAA-382 GCA_000407165.1
oV
434 Escherichia coli O104_3AH4 str. 2011C-3493 GCA_000299455.1
434 Escherichia coli O157_3AH7 str. Sakai GCA_000008865.1
434 Escherichia coli str. K-12 substr. MG1655 GCA_000005845.2
434 Shigella flexneri 2a str. 301 GCA_000006925.2
433 Escherichia coli O83_3AH1 str. NRG 857C GCA_000183345.1
433 Escherichia coli UMN026 GCA_000026325.2
411 Escherichia coli IAI39 GCA_000026345.1
411 Shigella dysenteriae Sd197 GCA_000012005.1
oW
548 Clostridium saccharobutylicum DSM 13864 GCA_000473995.1
548 Clostridium saccharoperbutylacetonicum N1-4_28HMT_29 GCA_000340885.1
543 Clostridium beijerinckii GCA_000833105.2
531 Clostridium puniceum GCA_002006345.1
oX
746 Clostridium beijerinckii GCA_000833105.2
727 Clostridium saccharoperbutylacetonicum N1-4_28HMT_29 GCA_000340885.1
710 Clostridium saccharobutylicum DSM 13864 GCA_000473995.1
oY
517 Deinococcus radiodurans R1 GCA_000008565.1
332 Deinococcus gobiensis I-0 GCA_000252445.1
323 Deinococcus deserti VCD115 GCA_000020685.1
oZ
589 Bifidobacterium adolescentis ATCC 15703 GCA_000010425.1
523 Bifidobacterium dentium JCM 1195 = DSM 20436 GCA_001042595.1
450 Bifidobacterium tsurumiense GCA_000741765.1
p0
674 Staphylococcus epidermidis ATCC 12228 GCA_000007645.1
654 Staphylococcus capitis subsp. capitis GCA_001028645.1
619 Megasphaera cerevisiae DSM 20462 GCA_001045675.1
619 Staphylococcus warneri SG1 GCA_000332735.1
p1
383 Rhodobacter sphaeroides 2.4.1 GCA_000012905.2
359 Rhodobacter sphaeroides ATCC 17025 GCA_000016405.1
287 Gemmobacter aquatilis GCA_900110025.1
p2
650 Staphylococcus epidermidis ATCC 12228 GCA_000007645.1
624 Staphylococcus capitis subsp. capitis GCA_001028645.1
606 Staphylococcus hominis subsp. hominis C80 GCA_000183685.1
p3
653 Enterococcus faecalis V583 GCA_000007785.1
648 Streptomyces cinnamoneus GCA_001885705.1
596 Enterococcus saccharolyticus subsp. saccharolyticus ATCC 43076 GCA_000407285.1
p4
637 Enterococcus faecalis V583 GCA_000007785.1
590 Streptomyces cinnamoneus GCA_001885705.1
507 Enterococcus rivorum GCA_001742285.1
p5
725 Lactobacillus gasseri ATCC 33323 = JCM 1131 GCA_000014425.1
689 Lactobacillus hominis DSM 23910 = CRBIP 24.179 GCA_000296835.1
554 Lactobacillus iners DSM 13335 GCA_000160875.1
p6
601 Clostridium beijerinckii GCA_000833105.2
555 Clostridium saccharoperbutylacetonicum N1-4_28HMT_29 GCA_000340885.1
541 Clostridium chromiireducens GCA_002029255.1
p7
417 Enterococcus faecalis V583 GCA_000007785.1
387 Streptomyces cinnamoneus GCA_001885705.1
386 Enterococcus asini ATCC 700915 GCA_000407365.1
p8
493 Streptococcus mutans UA159 GCA_000007465.2
352 Streptococcus gallolyticus subsp. gallolyticus DSM 16831 GCA_002000985.1
350 Streptococcus equinus GCA_000964315.1
p9
619 Streptococcus mutans UA159 GCA_000007465.2
418 Streptococcus ratti FA-1 = DSM 20564 GCA_000286075.1
384 Streptococcus gordonii str. Challis substr. CH1 GCA_000017005.1
pa
815 Escherichia coli str. K-12 substr. MG1655 GCA_000005845.2
814 Escherichia coli O157_3AH7 str. Sakai GCA_000008865.1
806 Escherichia coli UMN026 GCA_000026325.2

pb
442 Deinococcus radiodurans R1 GCA_000008565.1
296 Deinococcus deserti VCD115 GCA_000020685.1
260 Deinococcus hopiensis KR-140 GCA_900176165.1
pc
884 Staphylococcus epidermidis ATCC 12228 GCA_000007645.1
825 Staphylococcus capitis subsp. capitis GCA_001028645.1
749 Megasphaera cerevisiae DSM 20462 GCA_001045675.1
749 Staphylococcus warneri SG1 GCA_000332735.1
pd
753 Deinococcus radiodurans R1 GCA_000008565.1
438 Deinococcus deserti VCD115 GCA_000020685.1
425 Deinococcus soli Cha et al. 2016 GCA_001007995.1
pe
761 Clostridium beijerinckii GCA_000833105.2
749 Clostridium saccharoperbutylacetonicum N1-4_28HMT_29 GCA_000340885.1
725 Clostridium puniceum GCA_002006345.1
pf
603 Streptococcus mutans UA159 GCA_000007465.2
398 Streptococcus gordonii str. Challis substr. CH1 GCA_000017005.1
395 Streptococcus ratti FA-1 = DSM 20564 GCA_000286075.1
pg
500 Streptococcus mutans UA159 GCA_000007465.2
353 Streptococcus ratti FA-1 = DSM 20564 GCA_000286075.1
334 Streptococcus macacae NCTC 11558 GCA_000187995.3
ph
795 Escherichia coli O104_3AH4 str. 2011C-3493 GCA_000299455.1
795 Escherichia coli str. K-12 substr. MG1655 GCA_000005845.2
771 Shigella flexneri 2a str. 301 GCA_000006925.2
746 Escherichia coli UMN026 GCA_000026325.2
pi
748 Rhodobacter sphaeroides 2.4.1 GCA_000012905.2
647 Rhodobacter sphaeroides ATCC 17025 GCA_000016405.1
529 Gemmobacter aquatilis GCA_900110025.1
pj
480 Rhodobacter sphaeroides 2.4.1 GCA_000012905.2
449 Rhodobacter sphaeroides ATCC 17025 GCA_000016405.1
358 Pseudorhodobacter ferrugineus DSM 5888 GCA_000420745.1
pk
789 Streptococcus mutans UA159 GCA_000007465.2
585 Streptococcus ratti FA-1 = DSM 20564 GCA_000286075.1
544 Streptococcus ferus DSM 20646 GCA_000372425.1
pl
731 Streptococcus mutans UA159 GCA_000007465.2
556 Streptococcus ratti FA-1 = DSM 20564 GCA_000286075.1
484 Streptococcus ferus DSM 20646 GCA_000372425.1
pm
893 Lactobacillus gasseri ATCC 33323 = JCM 1131 GCA_000014425.1
827 Lactobacillus hominis DSM 23910 = CRBIP 24.179 GCA_000296835.1
615 Lactobacillus psittaci DSM 15354 GCA_000425905.1
pn
568 Deinococcus radiodurans R1 GCA_000008565.1
426 Deinococcus gobiensis I-0 GCA_000252445.1
416 Deinococcus deserti VCD115 GCA_000020685.1
po
448 Streptococcus mutans UA159 GCA_000007465.2
406 Streptomyces cinnamoneus GCA_001885705.1
388 Enterococcus faecalis V583 GCA_000007785.1
pp
688 Staphylococcus epidermidis ATCC 12228 GCA_000007645.1
652 Staphylococcus capitis subsp. capitis GCA_001028645.1
592 Staphylococcus lugdunensis HKU09-01 GCA_000025085.1
pq
720 Lactobacillus gasseri ATCC 33323 = JCM 1131 GCA_000014425.1
633 Lactobacillus hominis DSM 23910 = CRBIP 24.179 GCA_000296835.1
457 Lactobacillus iners DSM 13335 GCA_000160875.1
pr
664 Deinococcus radiodurans R1 GCA_000008565.1
438 Deinococcus deserti VCD115 GCA_000020685.1
399 Deinococcus hopiensis KR-140 GCA_900176165.1
ps
619 Clostridium beijerinckii GCA_000833105.2
603 Clostridium saccharoperbutylacetonicum N1-4_28HMT_29 GCA_000340885.1
594 Clostridium saccharobutylicum DSM 13864 GCA_000473995.1
pt
465 Enterococcus faecalis V583 GCA_000007785.1

419 Enterococcus thailandicus GCA_001652875.1
419 Streptomyces cinnamoneus GCA_001885705.1
412 Enterococcus rivorum GCA_001742285.1
pu
887 Staphylococcus epidermidis ATCC 12228 GCA_000007645.1
859 Staphylococcus capitis subsp. capitis GCA_001028645.1
788 Staphylococcus hominis subsp. hominis C80 GCA_000183685.1
pv
430 Streptococcus mutans UA159 GCA_000007465.2
270 Streptococcus sobrinus DSM 20742 = ATCC 33478 GCA_000686605.1
259 Streptococcus equinus GCA_000964315.1
259 Streptococcus ratti FA-1 = DSM 20564 GCA_000286075.1
pw
404 Streptococcus mutans UA159 GCA_000007465.2
254 Streptococcus gordonii str. Challis substr. CH1 GCA_000017005.1
236 Streptococcus orisratti DSM 15617 GCA_000380105.1
px
363 Enterococcus faecalis V583 GCA_000007785.1
328 Enterococcus rivorum GCA_001742285.1
320 Streptomyces cinnamoneus GCA_001885705.1
py
724 Clostridium beijerinckii GCA_000833105.2
707 Clostridium saccharoperbutylacetonicum N1-4_28HMT_29 GCA_000340885.1
696 Clostridium saccharobutylicum DSM 13864 GCA_000473995.1
pz
735 Staphylococcus epidermidis ATCC 12228 GCA_000007645.1
681 Staphylococcus haemolyticus JCSC1435 GCA_000009865.1
673 Staphylococcus capitis subsp. capitis GCA_001028645.1
pA
720 Clostridium beijerinckii GCA_000833105.2
684 Clostridium saccharoperbutylacetonicum N1-4_28HMT_29 GCA_000340885.1
661 Clostridium butyricum GCA_001456065.2
pB
751 Staphylococcus epidermidis ATCC 12228 GCA_000007645.1
733 Staphylococcus capitis subsp. capitis GCA_001028645.1
688 Staphylococcus hominis subsp. hominis C80 GCA_000183685.1
pC
495 Rhodobacter sphaeroides 2.4.1 GCA_000012905.2
457 Rhodobacter sphaeroides ATCC 17025 GCA_000016405.1
350 Gemmobacter aquatilis GCA_900110025.1
pD
784 Lactobacillus gasseri ATCC 33323 = JCM 1131 GCA_000014425.1
738 Lactobacillus hominis DSM 23910 = CRBIP 24.179 GCA_000296835.1
556 Lactobacillus iners DSM 13335 GCA_000160875.1
pE
506 Deinococcus radiodurans R1 GCA_000008565.1
280 Deinococcus deserti VCD115 GCA_000020685.1
261 Deinococcus gobiensis I-0 GCA_000252445.1
pF
625 Clostridium beijerinckii GCA_000833105.2
600 Clostridium saccharoperbutylacetonicum N1-4_28HMT_29 GCA_000340885.1
559 Clostridium puniceum GCA_002006345.1
pG
496 Clostridium beijerinckii GCA_000833105.2
481 Clostridium puniceum GCA_002006345.1
481 Clostridium saccharoperbutylacetonicum N1-4_28HMT_29 GCA_000340885.1
468 Clostridium saccharobutylicum DSM 13864 GCA_000473995.1
pH
647 Rhodobacter sphaeroides 2.4.1 GCA_000012905.2
564 Rhodobacter sphaeroides ATCC 17025 GCA_000016405.1
468 Defluviimonas alba GCA_001620265.1
pI
801 Bacillus anthracis str. Ames GCA_000007845.1
801 Bacillus anthracis str. Sterne GCA_000008165.1
801 Bacillus cereus ATCC 14579 GCA_000007825.1
801 _5BBacillus thuringiensis_5D serovar konkukian str. 97-27 GCA_000008505.1
787 Bacillus pseudomycoides DSM 12442 GCA_000161455.1
783 Bacillus thuringiensis YBT-1518 GCA_000497525.2
pJ
495 Escherichia coli str. K-12 substr. MG1655 GCA_000005845.2
484 Escherichia coli O104_3AH4 str. 2011C-3493 GCA_000299455.1
469 Escherichia coli O83_3AH1 str. NRG 857C GCA_000183345.1
pK
740 Enterococcus faecalis V583 GCA_000007785.1
693 Streptomyces cinnamoneus GCA_001885705.1
616 Enterococcus rivorum GCA_001742285.1

pL
660 Enterococcus faecalis V583 GCA_000007785.1
613 Streptomyces cinnamoneus GCA_001885705.1
538 Enterococcus rivorum GCA_001742285.1
pM
694 Clostridium beijerinckii GCA_000833105.2
628 Clostridium saccharoperbutylacetonicum N1-4_28HMT_29 GCA_000340885.1
618 Clostridium puniceum GCA_002006345.1
pN
849 Rhodobacter sphaeroides 2.4.1 GCA_000012905.2
772 Rhodobacter sphaeroides ATCC 17025 GCA_000016405.1
640 Defluviimonas alba GCA_001620265.1
pO
748 Bacillus anthracis str. Ames GCA_000007845.1
748 Bacillus anthracis str. Sterne GCA_000008165.1
740 Bacillus cereus ATCC 14579 GCA_000007825.1
740 _5BBacillus thuringiensis_5D serovar konkukian str. 97-27 GCA_000008505.1
722 Bacillus pseudomycoides DSM 12442 GCA_000161455.1
pP
705 Shigella flexneri 2a str. 301 GCA_000006925.2
698 Escherichia coli O104_3AH4 str. 2011C-3493 GCA_000299455.1
698 Escherichia coli str. K-12 substr. MG1655 GCA_000005845.2
658 Escherichia coli O83_3AH1 str. NRG 857C GCA_000183345.1
pQ
712 Deinococcus radiodurans R1 GCA_000008565.1
497 Deinococcus gobiensis I-0 GCA_000252445.1
491 Deinococcus puniceus GCA_001644565.1
pR
827 Streptococcus mutans UA159 GCA_000007465.2
652 Streptococcus gordonii str. Challis substr. CH1 GCA_000017005.1
625 Streptococcus cristatus AS 1.3089 GCA_000385925.1
pS
703 Clostridium saccharoperbutylacetonicum N1-4_28HMT_29 GCA_000340885.1
699 Clostridium beijerinckii GCA_000833105.2
675 Clostridium saccharobutylicum DSM 13864 GCA_000473995.1
pT
631 Deinococcus radiodurans R1 GCA_000008565.1
445 Deinococcus gobiensis I-0 GCA_000252445.1
435 Deinococcus deserti VCD115 GCA_000020685.1
pU
591 Clostridium beijerinckii GCA_000833105.2
563 Clostridium puniceum GCA_002006345.1
563 Clostridium saccharoperbutylacetonicum N1-4_28HMT_29 GCA_000340885.1
541 Clostridium saccharobutylicum DSM 13864 GCA_000473995.1
pV
674 Bifidobacterium adolescentis ATCC 15703 GCA_000010425.1
570 Bifidobacterium dentium JCM 1195 = DSM 20436 GCA_001042595.1
533 Bifidobacterium angulatum DSM 20098 = JCM 7096 GCA_001025155.1
pW
760 Bacillus anthracis str. Ames GCA_000007845.1
760 Bacillus anthracis str. Sterne GCA_000008165.1
759 _5BBacillus thuringiensis_5D serovar konkukian str. 97-27 GCA_000008505.1
746 Bacillus cereus ATCC 14579 GCA_000007825.1
746 Bacillus thuringiensis YBT-1518 GCA_000497525.2
pX
640 Clostridium beijerinckii GCA_000833105.2
608 Clostridium saccharoperbutylacetonicum N1-4_28HMT_29 GCA_000340885.1
584 Clostridium saccharobutylicum DSM 13864 GCA_000473995.1
pY
768 Rhodobacter sphaeroides 2.4.1 GCA_000012905.2
698 Rhodobacter sphaeroides ATCC 17025 GCA_000016405.1
546 Gemmobacter aquatilis GCA_900110025.1
pZ
382 Clostridium beijerinckii GCA_000833105.2
349 Clostridium puniceum GCA_002006345.1
349 Clostridium saccharoperbutylacetonicum N1-4_28HMT_29 GCA_000340885.1
334 Clostridium saccharobutylicum DSM 13864 GCA_000473995.1
q0
883 Streptococcus mutans UA159 GCA_000007465.2
660 Streptococcus ratti FA-1 = DSM 20564 GCA_000286075.1
650 Streptococcus gordonii str. Challis substr. CH1 GCA_000017005.1
q1
486 Clostridium beijerinckii GCA_000833105.2
484 Clostridium puniceum GCA_002006345.1
470 Clostridium saccharoperbutylacetonicum N1-4_28HMT_29 GCA_000340885.1
q2

610 Staphylococcus epidermidis ATCC 12228 GCA_000007645.1
560 Staphylococcus capitis subsp. capitis GCA_001028645.1
548 Staphylococcus condimenti GCA_001618885.1
q3
537 Staphylococcus epidermidis ATCC 12228 GCA_000007645.1
507 Staphylococcus capitis subsp. capitis GCA_001028645.1
466 Megasphaera cerevisiae DSM 20462 GCA_001045675.1
466 Staphylococcus pettenkoferi GCA_002208805.1
466 Staphylococcus warneri SG1 GCA_000332735.1
q4
785 Clostridium beijerinckii GCA_000833105.2
747 Clostridium saccharoperbutylacetonicum N1-4_28HMT_29 GCA_000340885.1
739 Clostridium saccharobutylicum DSM 13864 GCA_000473995.1
q5
519 Clostridium beijerinckii GCA_000833105.2
470 Clostridium saccharoperbutylacetonicum N1-4_28HMT_29 GCA_000340885.1
464 Clostridium puniceum GCA_002006345.1
q6
723 Staphylococcus epidermidis ATCC 12228 GCA_000007645.1
696 Staphylococcus capitis subsp. capitis GCA_001028645.1
675 Staphylococcus aureus subsp. aureus NCTC 8325 GCA_000013425.1
q7
614 Streptococcus mutans UA159 GCA_000007465.2
400 Streptococcus macacae NCTC 11558 GCA_000187995.3
384 Streptococcus ratti FA-1 = DSM 20564 GCA_000286075.1
q8
754 Rhodobacter sphaeroides 2.4.1 GCA_000012905.2
729 Rhodobacter sphaeroides ATCC 17025 GCA_000016405.1
621 Pseudorhodobacter ferrugineus DSM 5888 GCA_000420745.1
q9
678 Enterococcus faecalis V583 GCA_000007785.1
631 Streptomyces cinnamoneus GCA_001885705.1
606 Enterococcus rivorum GCA_001742285.1
qa
565 Lactobacillus gasseri ATCC 33323 = JCM 1131 GCA_000014425.1
486 Lactobacillus hominis DSM 23910 = CRBIP 24.179 GCA_000296835.1
292 Lactobacillus acidophilus NCFM GCA_000011985.1
292 Lactobacillus gallinarum GCA_001314245.2
qb
636 Rhodobacter sphaeroides 2.4.1 GCA_000012905.2
573 Rhodobacter sphaeroides ATCC 17025 GCA_000016405.1
491 Pseudorhodobacter ferrugineus DSM 5888 GCA_000420745.1
qc
587 Staphylococcus epidermidis ATCC 12228 GCA_000007645.1
572 Staphylococcus capitis subsp. capitis GCA_001028645.1
564 Megasphaera cerevisiae DSM 20462 GCA_001045675.1
564 Staphylococcus aureus subsp. aureus NCTC 8325 GCA_000013425.1
564 Staphylococcus warneri SG1 GCA_000332735.1
qd
660 Lactobacillus gasseri ATCC 33323 = JCM 1131 GCA_000014425.1
601 Lactobacillus hominis DSM 23910 = CRBIP 24.179 GCA_000296835.1
479 Lactobacillus iners DSM 13335 GCA_000160875.1
qe
774 Staphylococcus epidermidis ATCC 12228 GCA_000007645.1
708 Staphylococcus capitis subsp. capitis GCA_001028645.1
673 Staphylococcus warneri SG1 GCA_000332735.1
qf
785 Streptococcus mutans UA159 GCA_000007465.2
579 Streptococcus ratti FA-1 = DSM 20564 GCA_000286075.1
535 Streptococcus criceti HS-6 GCA_000187975.3
qg
803 Bacillus anthracis str. Ames GCA_000007845.1
803 Bacillus anthracis str. Sterne GCA_000008165.1
803 Bacillus cereus ATCC 14579 GCA_000007825.1
803 Bacillus thuringiensis YBT-1518 GCA_000497525.2
803 _5BBacillus thuringiensis_5D serovar konkukian str. 97-27 GCA_000008505.1
776 Bacillus mycoides GCA_000832605.1
771 Bacillus pseudomycoides DSM 12442 GCA_000161455.1
qh
756 Rhodobacter sphaeroides 2.4.1 GCA_000012905.2
620 Rhodobacter sphaeroides ATCC 17025 GCA_000016405.1
508 Defluviimonas alba GCA_001620265.1
qi
558 Staphylococcus epidermidis ATCC 12228 GCA_000007645.1
531 Staphylococcus capitis subsp. capitis GCA_001028645.1
505 Staphylococcus haemolyticus JCSC1435 GCA_000009865.1

qj
521 Bifidobacterium adolescentis ATCC 15703 GCA_000010425.1
473 Bifidobacterium dentium JCM 1195 = DSM 20436 GCA_001042595.1
435 Bifidobacterium stellenboschense GCA_000741785.1
qk
705 Bifidobacterium adolescentis ATCC 15703 GCA_000010425.1
594 Bifidobacterium dentium JCM 1195 = DSM 20436 GCA_001042595.1
525 Bifidobacterium bifidum PRL2010 GCA_000165905.1
ql
576 Rhodobacter sphaeroides 2.4.1 GCA_000012905.2
543 Rhodobacter sphaeroides ATCC 17025 GCA_000016405.1
420 Gemmobacter aquatilis GCA_900110025.1
qm
532 Streptococcus mutans UA159 GCA_000007465.2
350 Streptococcus halotolerans GCA_001598035.1
337 Streptococcus ratti FA-1 = DSM 20564 GCA_000286075.1
qn
485 Clostridium beijerinckii GCA_000833105.2
470 Clostridium saccharoperbutylacetonicum N1-4_28HMT_29 GCA_000340885.1
465 Clostridium saccharobutylicum DSM 13864 GCA_000473995.1
qo
521 Escherichia coli O104_3AH4 str. 2011C-3493 GCA_000299455.1
520 Escherichia coli O157_3AH7 str. Sakai GCA_000008865.1
520 Escherichia coli str. K-12 substr. MG1655 GCA_000005845.2
520 Shigella flexneri 2a str. 301 GCA_000006925.2
503 Escherichia coli O83_3AH1 str. NRG 857C GCA_000183345.1
503 Escherichia coli UMN026 GCA_000026325.2
qp
531 Lactobacillus gasseri ATCC 33323 = JCM 1131 GCA_000014425.1
497 Lactobacillus hominis DSM 23910 = CRBIP 24.179 GCA_000296835.1
372 Lactobacillus psittaci DSM 15354 GCA_000425905.1
qq
610 Rhodobacter sphaeroides 2.4.1 GCA_000012905.2
516 Rhodobacter sphaeroides ATCC 17025 GCA_000016405.1
462 Defluviimonas alba GCA_001620265.1
qr
627 Streptococcus mutans UA159 GCA_000007465.2
454 Streptococcus gordonii str. Challis substr. CH1 GCA_000017005.1
429 Streptococcus macacae NCTC 11558 GCA_000187995.3
qs
698 Enterococcus faecalis V583 GCA_000007785.1
658 Streptomyces cinnamoneus GCA_001885705.1
593 Enterococcus asini ATCC 700915 GCA_000407365.1
qt
736 Rhodobacter sphaeroides 2.4.1 GCA_000012905.2
671 Rhodobacter sphaeroides ATCC 17025 GCA_000016405.1
563 Gemmobacter megaterium GCA_900156815.1
qu
852 Enterococcus faecalis V583 GCA_000007785.1
805 Streptomyces cinnamoneus GCA_001885705.1
729 Enterococcus rivorum GCA_001742285.1
qv
657 Deinococcus radiodurans R1 GCA_000008565.1
406 Deinococcus deserti VCD115 GCA_000020685.1
378 Deinococcus proteolyticus MRP GCA_000190555.1
qw
722 Rhodobacter sphaeroides 2.4.1 GCA_000012905.2
695 Rhodobacter sphaeroides ATCC 17025 GCA_000016405.1
564 Defluviimonas alba GCA_001620265.1
qx
793 Staphylococcus epidermidis ATCC 12228 GCA_000007645.1
783 Staphylococcus capitis subsp. capitis GCA_001028645.1
743 Staphylococcus warneri SG1 GCA_000332735.1
qy
721 Streptococcus mutans UA159 GCA_000007465.2
544 Streptococcus ratti FA-1 = DSM 20564 GCA_000286075.1
486 Streptococcus sobrinus DSM 20742 = ATCC 33478 GCA_000686605.1
qz
978 Streptococcus mutans UA159 GCA_000007465.2
734 Streptococcus ratti FA-1 = DSM 20564 GCA_000286075.1
720 Streptococcus gordonii str. Challis substr. CH1 GCA_000017005.1
qA
443 Escherichia coli O157_3AH7 str. Sakai GCA_000008865.1
443 Escherichia coli str. K-12 substr. MG1655 GCA_000005845.2
441 Escherichia coli UMN026 GCA_000026325.2
428 Escherichia coli O104_3AH4 str. 2011C-3493 GCA_000299455.1

428 Shigella flexneri 2a str. 301 GCA_000006925.2
qB
675 Lactobacillus gasseri ATCC 33323 = JCM 1131 GCA_000014425.1
642 Lactobacillus hominis DSM 23910 = CRBIP 24.179 GCA_000296835.1
474 Lactobacillus iners DSM 13335 GCA_000160875.1
qC
736 Clostridium beijerinckii GCA_000833105.2
710 Clostridium saccharoperbutylacetonicum N1-4_28HMT_29 GCA_000340885.1
708 Clostridium saccharobutylicum DSM 13864 GCA_000473995.1
qD
461 Staphylococcus epidermidis ATCC 12228 GCA_000007645.1
445 Staphylococcus capitis subsp. capitis GCA_001028645.1
407 Megasphaera cerevisiae DSM 20462 GCA_001045675.1
407 Staphylococcus warneri SG1 GCA_000332735.1
qE
627 Escherichia coli O157_3AH7 str. Sakai GCA_000008865.1
627 Escherichia coli UMN026 GCA_000026325.2
627 Escherichia coli str. K-12 substr. MG1655 GCA_000005845.2
627 Shigella flexneri 2a str. 301 GCA_000006925.2
621 Escherichia coli O104_3AH4 str. 2011C-3493 GCA_000299455.1
621 Escherichia coli O83_3AH1 str. NRG 857C GCA_000183345.1
582 Klebsiella oxytoca GCA_001022195.1
qF
587 Rhodobacter sphaeroides 2.4.1 GCA_000012905.2
580 Rhodobacter sphaeroides ATCC 17025 GCA_000016405.1
447 Defluviimonas alba GCA_001620265.1
qG
693 Deinococcus radiodurans R1 GCA_000008565.1
434 Deinococcus deserti VCD115 GCA_000020685.1
417 Deinococcus hopiensis KR-140 GCA_900176165.1
qH
649 Bacillus cereus ATCC 14579 GCA_000007825.1
643 Bacillus anthracis str. Ames GCA_000007845.1
643 Bacillus anthracis str. Sterne GCA_000008165.1
643 _5BBacillus thuringiensis_5D serovar konkukian str. 97-27 GCA_000008505.1
634 Bacillus pseudomycoides DSM 12442 GCA_000161455.1
qI
649 Staphylococcus epidermidis ATCC 12228 GCA_000007645.1
590 Staphylococcus capitis subsp. capitis GCA_001028645.1
545 Staphylococcus haemolyticus JCSC1435 GCA_000009865.1
545 Staphylococcus hominis subsp. hominis C80 GCA_000183685.1
qJ
590 Lactobacillus gasseri ATCC 33323 = JCM 1131 GCA_000014425.1
552 Lactobacillus hominis DSM 23910 = CRBIP 24.179 GCA_000296835.1
476 Streptococcus mutans UA159 GCA_000007465.2
qK
487 Enterococcus faecalis V583 GCA_000007785.1
441 Streptomyces cinnamoneus GCA_001885705.1
414 Enterococcus rivorum GCA_001742285.1
qL
697 Rhodobacter sphaeroides 2.4.1 GCA_000012905.2
644 Rhodobacter sphaeroides ATCC 17025 GCA_000016405.1
569 Gemmobacter aquatilis GCA_900110025.1
qM
438 Lactobacillus gasseri ATCC 33323 = JCM 1131 GCA_000014425.1
415 Lactobacillus hominis DSM 23910 = CRBIP 24.179 GCA_000296835.1
287 Lactobacillus iners DSM 13335 GCA_000160875.1
qN
471 Enterococcus faecalis V583 GCA_000007785.1
425 Streptomyces cinnamoneus GCA_001885705.1
401 Enterococcus asini ATCC 700915 GCA_000407365.1
401 Enterococcus massiliensis GCA_001050095.1
401 Enterococcus saccharolyticus subsp. saccharolyticus ATCC 43076 GCA_000407285.1
qO
617 Escherichia coli IAI39 GCA_000026345.1
617 Escherichia coli O83_3AH1 str. NRG 857C GCA_000183345.1
608 Escherichia coli O157_3AH7 str. Sakai GCA_000008865.1
608 Escherichia coli str. K-12 substr. MG1655 GCA_000005845.2
608 Shigella flexneri 2a str. 301 GCA_000006925.2
605 Escherichia coli O104_3AH4 str. 2011C-3493 GCA_000299455.1
qP
651 Staphylococcus epidermidis ATCC 12228 GCA_000007645.1
588 Staphylococcus capitis subsp. capitis GCA_001028645.1
571 Staphylococcus haemolyticus JCSC1435 GCA_000009865.1
qQ
693 Enterococcus faecalis V583 GCA_000007785.1

647 Streptomyces cinnamoneus GCA_001885705.1
589 Vagococcus lutrae LBD1 GCA_000498295.1
qR
794 Rhodobacter sphaeroides 2.4.1 GCA_000012905.2
694 Rhodobacter sphaeroides ATCC 17025 GCA_000016405.1
571 Gemmobacter aquatilis GCA_900110025.1
qS
791 Staphylococcus epidermidis ATCC 12228 GCA_000007645.1
752 Staphylococcus capitis subsp. capitis GCA_001028645.1
748 Staphylococcus hominis subsp. hominis C80 GCA_000183685.1
qT
751 Rhodobacter sphaeroides 2.4.1 GCA_000012905.2
693 Rhodobacter sphaeroides ATCC 17025 GCA_000016405.1
547 Pseudorhodobacter psychrotolerans GCA_001294535.1
qU
737 Streptococcus mutans UA159 GCA_000007465.2
561 Streptococcus ratti FA-1 = DSM 20564 GCA_000286075.1
517 Streptococcus gordonii str. Challis substr. CH1 GCA_000017005.1
qV
573 Streptococcus mutans UA159 GCA_000007465.2
430 Streptococcus ratti FA-1 = DSM 20564 GCA_000286075.1
396 Streptococcus gordonii str. Challis substr. CH1 GCA_000017005.1
qW
560 Streptococcus mutans UA159 GCA_000007465.2
350 Streptococcus gordonii str. Challis substr. CH1 GCA_000017005.1
344 Streptococcus ratti FA-1 = DSM 20564 GCA_000286075.1
qX
820 Clostridium beijerinckii GCA_000833105.2
812 Clostridium saccharobutylicum DSM 13864 GCA_000473995.1
806 Clostridium saccharoperbutylacetonicum N1-4_28HMT_29 GCA_000340885.1
qY
797 Enterococcus faecalis V583 GCA_000007785.1
750 Streptomyces cinnamoneus GCA_001885705.1
729 Enterococcus rivorum GCA_001742285.1
qZ
690 Lactobacillus gasseri ATCC 33323 = JCM 1131 GCA_000014425.1
667 Lactobacillus hominis DSM 23910 = CRBIP 24.179 GCA_000296835.1
488 Lactobacillus jensenii GCA_001936235.1
488 Lactobacillus psittaci DSM 15354 GCA_000425905.1
r0
678 Enterococcus faecalis V583 GCA_000007785.1
631 Streptomyces cinnamoneus GCA_001885705.1
624 Enterococcus rivorum GCA_001742285.1
r1
630 Bacillus anthracis str. Ames GCA_000007845.1
630 Bacillus anthracis str. Sterne GCA_000008165.1
630 Bacillus cereus ATCC 14579 GCA_000007825.1
630 _5BBacillus thuringiensis_5D serovar konkukian str. 97-27 GCA_000008505.1
623 Bacillus mycoides GCA_000832605.1
618 Bacillus thuringiensis YBT-1518 GCA_000497525.2
r2
729 Streptococcus mutans UA159 GCA_000007465.2
573 Streptococcus ratti FA-1 = DSM 20564 GCA_000286075.1
528 Streptococcus gordonii str. Challis substr. CH1 GCA_000017005.1
r3
815 Rhodobacter sphaeroides 2.4.1 GCA_000012905.2
732 Rhodobacter sphaeroides ATCC 17025 GCA_000016405.1
617 Gemmobacter aquatilis GCA_900110025.1
r4
857 Clostridium beijerinckii GCA_000833105.2
842 Clostridium saccharoperbutylacetonicum N1-4_28HMT_29 GCA_000340885.1
832 Clostridium saccharobutylicum DSM 13864 GCA_000473995.1
r5
661 Bacillus anthracis str. Ames GCA_000007845.1
661 Bacillus anthracis str. Sterne GCA_000008165.1
661 _5BBacillus thuringiensis_5D serovar konkukian str. 97-27 GCA_000008505.1
657 Bacillus thuringiensis YBT-1518 GCA_000497525.2
644 Bacillus cereus ATCC 14579 GCA_000007825.1
r6
981 Lactobacillus gasseri ATCC 33323 = JCM 1131 GCA_000014425.1
899 Lactobacillus hominis DSM 23910 = CRBIP 24.179 GCA_000296835.1
671 Lactobacillus iners DSM 13335 GCA_000160875.1
r7
760 Deinococcus radiodurans R1 GCA_000008565.1
511 Deinococcus deserti VCD115 GCA_000020685.1
494 Deinococcus murrayi DSM 11303 GCA_000482805.1

r8
801 Clostridium beijerinckii GCA_000833105.2
759 Clostridium saccharoperbutylacetonicum N1-4_28HMT_29 GCA_000340885.1
730 Clostridium saccharobutylicum DSM 13864 GCA_000473995.1
r9
696 Deinococcus radiodurans R1 GCA_000008565.1
472 Deinococcus gobiensis I-0 GCA_000252445.1
457 Deinococcus deserti VCD115 GCA_000020685.1
ra
766 Lactobacillus gasseri ATCC 33323 = JCM 1131 GCA_000014425.1
704 Lactobacillus hominis DSM 23910 = CRBIP 24.179 GCA_000296835.1
590 Lactobacillus iners DSM 13335 GCA_000160875.1
rb
426 Clostridium beijerinckii GCA_000833105.2
396 Clostridium saccharoperbutylacetonicum N1-4_28HMT_29 GCA_000340885.1
374 Clostridium puniceum GCA_002006345.1
rc
759 Clostridium beijerinckii GCA_000833105.2
709 Clostridium saccharoperbutylacetonicum N1-4_28HMT_29 GCA_000340885.1
680 Clostridium puniceum GCA_002006345.1
rd
627 Deinococcus radiodurans R1 GCA_000008565.1
312 Deinococcus deserti VCD115 GCA_000020685.1
302 Deinococcus gobiensis I-0 GCA_000252445.1
re
670 Clostridium beijerinckii GCA_000833105.2
651 Clostridium saccharoperbutylacetonicum N1-4_28HMT_29 GCA_000340885.1
614 Clostridium saccharobutylicum DSM 13864 GCA_000473995.1
rf
641 Rhodobacter sphaeroides 2.4.1 GCA_000012905.2
585 Rhodobacter sphaeroides ATCC 17025 GCA_000016405.1
451 Rhodobacter aestuarii GCA_900156655.1
rg
779 Lactobacillus gasseri ATCC 33323 = JCM 1131 GCA_000014425.1
714 Lactobacillus hominis DSM 23910 = CRBIP 24.179 GCA_000296835.1
503 Lactobacillus iners DSM 13335 GCA_000160875.1
rh
641 Staphylococcus epidermidis ATCC 12228 GCA_000007645.1
600 Staphylococcus capitis subsp. capitis GCA_001028645.1
591 Megasphaera cerevisiae DSM 20462 GCA_001045675.1
591 Staphylococcus aureus subsp. aureus NCTC 8325 GCA_000013425.1
591 Staphylococcus haemolyticus JCSC1435 GCA_000009865.1
591 Staphylococcus hominis subsp. hominis C80 GCA_000183685.1
591 Staphylococcus warneri SG1 GCA_000332735.1
ri
719 Clostridium beijerinckii GCA_000833105.2
691 Clostridium saccharoperbutylacetonicum N1-4_28HMT_29 GCA_000340885.1
666 Clostridium puniceum GCA_002006345.1
rj
769 Bifidobacterium adolescentis ATCC 15703 GCA_000010425.1
630 Bifidobacterium dentium JCM 1195 = DSM 20436 GCA_001042595.1
576 Bifidobacterium callitrichos DSM 23973 GCA_000741175.1
rk
942 Bacillus anthracis str. Ames GCA_000007845.1
942 Bacillus anthracis str. Sterne GCA_000008165.1
935 _5BBacillus thuringiensis_5D serovar konkukian str. 97-27 GCA_000008505.1
930 Bacillus pseudomycoides DSM 12442 GCA_000161455.1
rl
675 Escherichia coli str. K-12 substr. MG1655 GCA_000005845.2
644 Escherichia coli O104_3AH4 str. 2011C-3493 GCA_000299455.1
644 Shigella flexneri 2a str. 301 GCA_000006925.2
616 Escherichia coli O157_3AH7 str. Sakai GCA_000008865.1
rm
711 Bifidobacterium adolescentis ATCC 15703 GCA_000010425.1
608 Bifidobacterium dentium JCM 1195 = DSM 20436 GCA_001042595.1
580 Bifidobacterium thermophilum GCA_000741495.1
580 Bifidobacterium thermophilum RBL67 GCA_000347695.1
rn
613 Clostridium beijerinckii GCA_000833105.2
552 Clostridium saccharoperbutylacetonicum N1-4_28HMT_29 GCA_000340885.1
536 Clostridium butyricum GCA_001456065.2
ro
850 Staphylococcus epidermidis ATCC 12228 GCA_000007645.1
803 Staphylococcus capitis subsp. capitis GCA_001028645.1
731 Staphylococcus haemolyticus JCSC1435 GCA_000009865.1
rp

751 Rhodobacter sphaeroides 2.4.1 GCA_000012905.2
642 Rhodobacter sphaeroides ATCC 17025 GCA_000016405.1
558 Gemmobacter aquatilis GCA_900110025.1
rq
979 Lactobacillus gasseri ATCC 33323 = JCM 1131 GCA_000014425.1
909 Lactobacillus hominis DSM 23910 = CRBIP 24.179 GCA_000296835.1
730 Lactobacillus iners DSM 13335 GCA_000160875.1
rr
778 Clostridium saccharoperbutylacetonicum N1-4_28HMT_29 GCA_000340885.1
767 Clostridium beijerinckii GCA_000833105.2
757 Clostridium saccharobutylicum DSM 13864 GCA_000473995.1
rs
775 Bifidobacterium adolescentis ATCC 15703 GCA_000010425.1
659 Bifidobacterium callitrichos DSM 23973 GCA_000741175.1
653 Bifidobacterium angulatum DSM 20098 = JCM 7096 GCA_001025155.1
rt
460 Clostridium beijerinckii GCA_000833105.2
456 Clostridium saccharoperbutylacetonicum N1-4_28HMT_29 GCA_000340885.1
440 Clostridium puniceum GCA_002006345.1
ru
663 Clostridium beijerinckii GCA_000833105.2
640 Clostridium saccharoperbutylacetonicum N1-4_28HMT_29 GCA_000340885.1
613 Clostridium puniceum GCA_002006345.1
rv
661 Staphylococcus epidermidis ATCC 12228 GCA_000007645.1
655 Staphylococcus capitis subsp. capitis GCA_001028645.1
614 Staphylococcus haemolyticus JCSC1435 GCA_000009865.1
614 Staphylococcus hominis subsp. hominis C80 GCA_000183685.1
rw
613 Rhodobacter sphaeroides 2.4.1 GCA_000012905.2
555 Rhodobacter sphaeroides ATCC 17025 GCA_000016405.1
486 Gemmobacter aquatilis GCA_900110025.1
rx
535 Staphylococcus epidermidis ATCC 12228 GCA_000007645.1
533 Staphylococcus capitis subsp. capitis GCA_001028645.1
481 Staphylococcus pettenkoferi GCA_002208805.1
ry
637 Clostridium beijerinckii GCA_000833105.2
630 Clostridium saccharoperbutylacetonicum N1-4_28HMT_29 GCA_000340885.1
623 Clostridium puniceum GCA_002006345.1
rz
401 Escherichia coli str. K-12 substr. MG1655 GCA_000005845.2
386 Escherichia coli O104_3AH4 str. 2011C-3493 GCA_000299455.1
386 Shigella flexneri 2a str. 301 GCA_000006925.2
378 Escherichia coli O157_3AH7 str. Sakai GCA_000008865.1
378 Escherichia coli UMN026 GCA_000026325.2
rA
818 Clostridium beijerinckii GCA_000833105.2
793 Clostridium saccharoperbutylacetonicum N1-4_28HMT_29 GCA_000340885.1
763 Clostridium puniceum GCA_002006345.1
rB
764 Deinococcus radiodurans R1 GCA_000008565.1
510 Deinococcus deserti VCD115 GCA_000020685.1
486 Deinococcus gobiensis I-0 GCA_000252445.1
rC
802 Escherichia coli O157_3AH7 str. Sakai GCA_000008865.1
802 Escherichia coli str. K-12 substr. MG1655 GCA_000005845.2
800 Shigella flexneri 2a str. 301 GCA_000006925.2
799 Escherichia coli UMN026 GCA_000026325.2
rD
638 Lactobacillus gasseri ATCC 33323 = JCM 1131 GCA_000014425.1
564 Lactobacillus hominis DSM 23910 = CRBIP 24.179 GCA_000296835.1
459 Lactobacillus iners DSM 13335 GCA_000160875.1
rE
840 Staphylococcus epidermidis ATCC 12228 GCA_000007645.1
799 Staphylococcus capitis subsp. capitis GCA_001028645.1
762 Staphylococcus aureus subsp. aureus NCTC 8325 GCA_000013425.1
rF
784 Lactobacillus gasseri ATCC 33323 = JCM 1131 GCA_000014425.1
728 Lactobacillus hominis DSM 23910 = CRBIP 24.179 GCA_000296835.1
538 Lactobacillus iners DSM 13335 GCA_000160875.1
rG
537 Clostridium beijerinckii GCA_000833105.2
510 Clostridium saccharoperbutylacetonicum N1-4_28HMT_29 GCA_000340885.1
495 Clostridium puniceum GCA_002006345.1
rH

664 Escherichia coli O104_3AH4 str. 2011C-3493 GCA_000299455.1
664 Escherichia coli O157_3AH7 str. Sakai GCA_000008865.1
664 Escherichia coli O83_3AH1 str. NRG 857C GCA_000183345.1
664 Escherichia coli UMN026 GCA_000026325.2
664 Escherichia coli str. K-12 substr. MG1655 GCA_000005845.2
664 Shigella flexneri 2a str. 301 GCA_000006925.2
649 Escherichia coli IAI39 GCA_000026345.1
634 Kosakonia sacchari SP1 GCA_000300455.4
634 Shigella dysenteriae Sd197 GCA_000012005.1
rI
618 Bacillus anthracis str. Ames GCA_000007845.1
618 Bacillus anthracis str. Sterne GCA_000008165.1
618 _5BBacillus thuringiensis_5D serovar konkukian str. 97-27 GCA_000008505.1
608 Bacillus cereus ATCC 14579 GCA_000007825.1
594 Bacillus pseudomycoides DSM 12442 GCA_000161455.1
rJ
617 Streptococcus mutans UA159 GCA_000007465.2
437 Streptococcus gordonii str. Challis substr. CH1 GCA_000017005.1
429 Streptococcus ratti FA-1 = DSM 20564 GCA_000286075.1
rK
576 Rhodobacter sphaeroides 2.4.1 GCA_000012905.2
549 Rhodobacter sphaeroides ATCC 17025 GCA_000016405.1
459 Defluviimonas alba GCA_001620265.1
rL
793 Escherichia coli str. K-12 substr. MG1655 GCA_000005845.2
793 Shigella flexneri 2a str. 301 GCA_000006925.2
792 Escherichia coli O83_3AH1 str. NRG 857C GCA_000183345.1
792 Escherichia coli UMN026 GCA_000026325.2
778 Escherichia coli O157_3AH7 str. Sakai GCA_000008865.1
rM
740 Staphylococcus epidermidis ATCC 12228 GCA_000007645.1
709 Staphylococcus hominis subsp. hominis C80 GCA_000183685.1
703 Staphylococcus haemolyticus JCSC1435 GCA_000009865.1
rN
627 Staphylococcus epidermidis ATCC 12228 GCA_000007645.1
593 Staphylococcus capitis subsp. capitis GCA_001028645.1
561 Staphylococcus aureus subsp. aureus NCTC 8325 GCA_000013425.1
rO
747 Enterococcus faecalis V583 GCA_000007785.1
704 Streptomyces cinnamoneus GCA_001885705.1
620 Enterococcus faecium DO GCA_000174395.2
rP
710 Lactobacillus gasseri ATCC 33323 = JCM 1131 GCA_000014425.1
687 Lactobacillus hominis DSM 23910 = CRBIP 24.179 GCA_000296835.1
501 Lactobacillus iners DSM 13335 GCA_000160875.1
rQ
613 Staphylococcus epidermidis ATCC 12228 GCA_000007645.1
551 Staphylococcus warneri SG1 GCA_000332735.1
537 Staphylococcus capitis subsp. capitis GCA_001028645.1
rR
518 Rhodobacter sphaeroides 2.4.1 GCA_000012905.2
464 Rhodobacter sphaeroides ATCC 17025 GCA_000016405.1
376 Gemmobacter megaterium GCA_900156815.1
rS
452 Deinococcus radiodurans R1 GCA_000008565.1
292 Deinococcus deserti VCD115 GCA_000020685.1
270 Deinococcus gobiensis I-0 GCA_000252445.1
rT
752 Lactobacillus gasseri ATCC 33323 = JCM 1131 GCA_000014425.1
695 Lactobacillus hominis DSM 23910 = CRBIP 24.179 GCA_000296835.1
540 Lactobacillus iners DSM 13335 GCA_000160875.1
rU
762 Rhodobacter sphaeroides 2.4.1 GCA_000012905.2
650 Rhodobacter sphaeroides ATCC 17025 GCA_000016405.1
584 Gemmobacter megaterium GCA_900156815.1
rV
760 Escherichia coli O104_3AH4 str. 2011C-3493 GCA_000299455.1
760 Escherichia coli O157_3AH7 str. Sakai GCA_000008865.1
760 Escherichia coli O83_3AH1 str. NRG 857C GCA_000183345.1
760 Escherichia coli UMN026 GCA_000026325.2
760 Escherichia coli str. K-12 substr. MG1655 GCA_000005845.2
760 Shigella flexneri 2a str. 301 GCA_000006925.2
719 Escherichia coli IAI39 GCA_000026345.1
707 Shigella dysenteriae Sd197 GCA_000012005.1
rW
790 Staphylococcus epidermidis ATCC 12228 GCA_000007645.1

728 Staphylococcus capitis subsp. capitis GCA_001028645.1
693 Staphylococcus warneri SG1 GCA_000332735.1
rX
717 Deinococcus radiodurans R1 GCA_000008565.1
436 Deinococcus deserti VCD115 GCA_000020685.1
410 Deinococcus soli Cha et al. 2016 GCA_001007995.1
rY
650 Streptococcus mutans UA159 GCA_000007465.2
437 Streptococcus ratti FA-1 = DSM 20564 GCA_000286075.1
384 Streptococcus salivarius GCA_000785515.1
384 Streptococcus thermophilus JIM 8232 GCA_000253395.1
rZ
678 Rhodobacter sphaeroides 2.4.1 GCA_000012905.2
643 Rhodobacter sphaeroides ATCC 17025 GCA_000016405.1
543 Pseudorhodobacter psychrotolerans GCA_001294535.1
s0
750 Streptococcus mutans UA159 GCA_000007465.2
579 Streptococcus ratti FA-1 = DSM 20564 GCA_000286075.1
534 Streptococcus gordonii str. Challis substr. CH1 GCA_000017005.1
s1
386 Enterococcus faecalis V583 GCA_000007785.1
354 Enterococcus haemoperoxidus ATCC BAA-382 GCA_000407165.1
354 Enterococcus massiliensis GCA_001050095.1
354 Enterococcus rivorum GCA_001742285.1
347 Enterococcus canis NBRC 100695 GCA_001544375.1
347 Enterococcus dispar ATCC 51266 GCA_000406945.1
s2
528 Rhodobacter sphaeroides 2.4.1 GCA_000012905.2
491 Rhodobacter sphaeroides ATCC 17025 GCA_000016405.1
442 Gemmobacter megaterium GCA_900156815.1
s3
805 Enterococcus faecalis V583 GCA_000007785.1
758 Streptomyces cinnamoneus GCA_001885705.1
712 Enterococcus rivorum GCA_001742285.1
s4
824 Lactobacillus gasseri ATCC 33323 = JCM 1131 GCA_000014425.1
735 Lactobacillus hominis DSM 23910 = CRBIP 24.179 GCA_000296835.1
567 Lactobacillus iners DSM 13335 GCA_000160875.1
s5
794 Streptococcus mutans UA159 GCA_000007465.2
578 Streptococcus ratti FA-1 = DSM 20564 GCA_000286075.1
507 Streptococcus sobrinus DSM 20742 = ATCC 33478 GCA_000686605.1
s6
720 Enterococcus faecalis V583 GCA_000007785.1
684 Enterococcus canis NBRC 100695 GCA_001544375.1
677 Enterococcus rivorum GCA_001742285.1
s7
414 Escherichia coli O157_3AH7 str. Sakai GCA_000008865.1
414 Escherichia coli str. K-12 substr. MG1655 GCA_000005845.2
414 Shigella flexneri 2a str. 301 GCA_000006925.2
410 Escherichia coli O104_3AH4 str. 2011C-3493 GCA_000299455.1
399 Escherichia coli O83_3AH1 str. NRG 857C GCA_000183345.1
399 Escherichia coli UMN026 GCA_000026325.2
s8
882 Lactobacillus gasseri ATCC 33323 = JCM 1131 GCA_000014425.1
817 Lactobacillus hominis DSM 23910 = CRBIP 24.179 GCA_000296835.1
602 Lactobacillus iners DSM 13335 GCA_000160875.1
s9
473 Staphylococcus epidermidis ATCC 12228 GCA_000007645.1
447 Staphylococcus capitis subsp. capitis GCA_001028645.1
428 Staphylococcus haemolyticus JCSC1435 GCA_000009865.1
sa
374 Staphylococcus epidermidis ATCC 12228 GCA_000007645.1
355 Staphylococcus capitis subsp. capitis GCA_001028645.1
330 Staphylococcus pseudintermedius HKU10-03 GCA_000185885.1
sb
822 Enterococcus faecalis V583 GCA_000007785.1
776 Streptomyces cinnamoneus GCA_001885705.1
707 Enterococcus rivorum GCA_001742285.1
sc
328 Rhodobacter sphaeroides 2.4.1 GCA_000012905.2
310 Rhodobacter sphaeroides ATCC 17025 GCA_000016405.1
276 Defluviimonas alba GCA_001620265.1
sd
429 Deinococcus radiodurans R1 GCA_000008565.1
289 Deinococcus deserti VCD115 GCA_000020685.1

282 Deinococcus soli Cha et al. 2016 GCA_001007995.1
se
439 Rhodobacter sphaeroides 2.4.1 GCA_000012905.2
403 Rhodobacter sphaeroides ATCC 17025 GCA_000016405.1
345 Gemmobacter aquatilis GCA_900110025.1
sf
580 Enterococcus faecalis V583 GCA_000007785.1
533 Streptomyces cinnamoneus GCA_001885705.1
459 Enterococcus hirae ATCC 9790 GCA_000271405.2
459 Enterococcus phoeniculicola ATCC BAA-412 GCA_000407505.1
sg
462 Streptomyces cinnamoneus GCA_001885705.1
460 Bacillus anthracis str. Ames GCA_000007845.1
460 Bacillus anthracis str. Sterne GCA_000008165.1
460 Bacillus cereus ATCC 14579 GCA_000007825.1
460 Bacillus thuringiensis YBT-1518 GCA_000497525.2
460 _5BBacillus thuringiensis_5D serovar konkukian str. 97-27 GCA_000008505.1
448 Enterococcus faecalis V583 GCA_000007785.1
sh
542 Rhodobacter sphaeroides 2.4.1 GCA_000012905.2
438 Rhodobacter sphaeroides ATCC 17025 GCA_000016405.1
348 Pseudorhodobacter wandonensis GCA_001202035.1
si
699 Staphylococcus epidermidis ATCC 12228 GCA_000007645.1
642 Staphylococcus capitis subsp. capitis GCA_001028645.1
628 Staphylococcus hominis subsp. hominis C80 GCA_000183685.1
sj
379 Clostridium beijerinckii GCA_000833105.2
354 Clostridium saccharoperbutylacetonicum N1-4_28HMT_29 GCA_000340885.1
353 Clostridium saccharobutylicum DSM 13864 GCA_000473995.1
sk
763 Lactobacillus gasseri ATCC 33323 = JCM 1131 GCA_000014425.1
744 Lactobacillus hominis DSM 23910 = CRBIP 24.179 GCA_000296835.1
531 Lactobacillus iners DSM 13335 GCA_000160875.1
sl
541 Bifidobacterium adolescentis ATCC 15703 GCA_000010425.1
446 Bifidobacterium dentium JCM 1195 = DSM 20436 GCA_001042595.1
432 Bifidobacterium longum NCC2705 GCA_000007525.1
sm
844 Deinococcus radiodurans R1 GCA_000008565.1
556 Deinococcus deserti VCD115 GCA_000020685.1
552 Deinococcus gobiensis I-0 GCA_000252445.1
sn
716 Lactobacillus gasseri ATCC 33323 = JCM 1131 GCA_000014425.1
660 Lactobacillus hominis DSM 23910 = CRBIP 24.179 GCA_000296835.1
456 Lactobacillus iners DSM 13335 GCA_000160875.1
so
678 Lactobacillus gasseri ATCC 33323 = JCM 1131 GCA_000014425.1
626 Lactobacillus hominis DSM 23910 = CRBIP 24.179 GCA_000296835.1
416 Lactobacillus psittaci DSM 15354 GCA_000425905.1
sp
788 Enterococcus faecalis V583 GCA_000007785.1
741 Streptomyces cinnamoneus GCA_001885705.1
706 Enterococcus rivorum GCA_001742285.1
sq
706 Lactobacillus gasseri ATCC 33323 = JCM 1131 GCA_000014425.1
691 Lactobacillus hominis DSM 23910 = CRBIP 24.179 GCA_000296835.1
547 Lactobacillus hamsteri DSM 5661 = JCM 6256 GCA_000615445.1
sr
742 Rhodobacter sphaeroides 2.4.1 GCA_000012905.2
650 Rhodobacter sphaeroides ATCC 17025 GCA_000016405.1
590 Pseudorhodobacter ferrugineus DSM 5888 GCA_000420745.1
ss
657 Enterococcus faecalis V583 GCA_000007785.1
610 Streptomyces cinnamoneus GCA_001885705.1
535 Enterococcus faecium DO GCA_000174395.2
st
796 Deinococcus radiodurans R1 GCA_000008565.1
568 Deinococcus deserti VCD115 GCA_000020685.1
541 Deinococcus murrayi DSM 11303 GCA_000482805.1
su
880 Bacillus anthracis str. Ames GCA_000007845.1
880 Bacillus anthracis str. Sterne GCA_000008165.1
880 _5BBacillus thuringiensis_5D serovar konkukian str. 97-27 GCA_000008505.1
864 Bacillus cereus ATCC 14579 GCA_000007825.1
834 Bacillus pseudomycoides DSM 12442 GCA_000161455.1

sv
429 Rhodobacter sphaeroides 2.4.1 GCA_000012905.2
423 Rhodobacter sphaeroides ATCC 17025 GCA_000016405.1
396 Pseudorhodobacter ferrugineus DSM 5888 GCA_000420745.1
sw
796 Lactobacillus gasseri ATCC 33323 = JCM 1131 GCA_000014425.1
770 Lactobacillus hominis DSM 23910 = CRBIP 24.179 GCA_000296835.1
560 Lactobacillus iners DSM 13335 GCA_000160875.1
sx
713 Streptococcus mutans UA159 GCA_000007465.2
496 Streptococcus gordonii str. Challis substr. CH1 GCA_000017005.1
471 Streptococcus ratti FA-1 = DSM 20564 GCA_000286075.1
sy
713 Staphylococcus epidermidis ATCC 12228 GCA_000007645.1
653 Staphylococcus capitis subsp. capitis GCA_001028645.1
610 Staphylococcus haemolyticus JCSC1435 GCA_000009865.1
sz
589 Staphylococcus epidermidis ATCC 12228 GCA_000007645.1
546 Staphylococcus lugdunensis HKU09-01 GCA_000025085.1
542 Staphylococcus capitis subsp. capitis GCA_001028645.1
sA
680 Clostridium beijerinckii GCA_000833105.2
644 Clostridium saccharoperbutylacetonicum N1-4_28HMT_29 GCA_000340885.1
621 Clostridium puniceum GCA_002006345.1
sB
600 Lactobacillus gasseri ATCC 33323 = JCM 1131 GCA_000014425.1
561 Lactobacillus hominis DSM 23910 = CRBIP 24.179 GCA_000296835.1
430 Lactobacillus iners DSM 13335 GCA_000160875.1
sC
425 Staphylococcus epidermidis ATCC 12228 GCA_000007645.1
390 Staphylococcus capitis subsp. capitis GCA_001028645.1
389 Staphylococcus haemolyticus JCSC1435 GCA_000009865.1
sD
412 Rhodobacter sphaeroides 2.4.1 GCA_000012905.2
359 Rhodobacter sphaeroides ATCC 17025 GCA_000016405.1
281 Gemmobacter aquatilis GCA_900110025.1
sE
762 Streptococcus mutans UA159 GCA_000007465.2
609 Streptococcus ratti FA-1 = DSM 20564 GCA_000286075.1
513 Streptococcus halotolerans GCA_001598035.1
sF
601 Staphylococcus epidermidis ATCC 12228 GCA_000007645.1
573 Staphylococcus capitis subsp. capitis GCA_001028645.1
502 Staphylococcus pettenkoferi GCA_002208805.1
sG
721 Staphylococcus epidermidis ATCC 12228 GCA_000007645.1
693 Staphylococcus capitis subsp. capitis GCA_001028645.1
629 Staphylococcus aureus subsp. aureus NCTC 8325 GCA_000013425.1
sH
683 Bacillus anthracis str. Ames GCA_000007845.1
683 Bacillus anthracis str. Sterne GCA_000008165.1
683 _5BBacillus thuringiensis_5D serovar konkukian str. 97-27 GCA_000008505.1
663 Bacillus cereus ATCC 14579 GCA_000007825.1
663 Bacillus thuringiensis YBT-1518 GCA_000497525.2
659 Bacillus pseudomycoides DSM 12442 GCA_000161455.1
sI
693 Deinococcus radiodurans R1 GCA_000008565.1
416 Deinococcus proteolyticus MRP GCA_000190555.1
402 Deinococcus gobiensis I-0 GCA_000252445.1
sJ
505 Escherichia coli O157_3AH7 str. Sakai GCA_000008865.1
505 Escherichia coli str. K-12 substr. MG1655 GCA_000005845.2
497 Escherichia coli UMN026 GCA_000026325.2
493 Escherichia coli IAI39 GCA_000026345.1
493 Escherichia coli O104_3AH4 str. 2011C-3493 GCA_000299455.1
493 Shigella flexneri 2a str. 301 GCA_000006925.2
sK
531 Clostridium beijerinckii GCA_000833105.2
493 Clostridium saccharoperbutylacetonicum N1-4_28HMT_29 GCA_000340885.1
483 Clostridium puniceum GCA_002006345.1
sL
851 Enterococcus faecalis V583 GCA_000007785.1
809 Streptomyces cinnamoneus GCA_001885705.1
749 Enterococcus asini ATCC 700915 GCA_000407365.1
sM
762 Clostridium beijerinckii GCA_000833105.2

733 Clostridium saccharoperbutylacetonicum N1-4_28HMT_29 GCA_000340885.1
706 Clostridium puniceum GCA_002006345.1
sN
644 Rhodobacter sphaeroides 2.4.1 GCA_000012905.2
618 Rhodobacter sphaeroides ATCC 17025 GCA_000016405.1
497 Defluviimonas alba GCA_001620265.1
sO
935 Lactobacillus gasseri ATCC 33323 = JCM 1131 GCA_000014425.1
825 Lactobacillus hominis DSM 23910 = CRBIP 24.179 GCA_000296835.1
596 Lactobacillus iners DSM 13335 GCA_000160875.1
sP
754 Bifidobacterium adolescentis ATCC 15703 GCA_000010425.1
680 Bifidobacterium dentium JCM 1195 = DSM 20436 GCA_001042595.1
597 Bifidobacterium breve DSM 20213 = JCM 1192 GCA_001025175.1
sQ
798 Clostridium beijerinckii GCA_000833105.2
725 Clostridium saccharoperbutylacetonicum N1-4_28HMT_29 GCA_000340885.1
695 Clostridium puniceum GCA_002006345.1
sR
658 Rhodobacter sphaeroides 2.4.1 GCA_000012905.2
600 Rhodobacter sphaeroides ATCC 17025 GCA_000016405.1
492 Gemmobacter aquatilis GCA_900110025.1
sS
542 Rhodobacter sphaeroides 2.4.1 GCA_000012905.2
515 Rhodobacter sphaeroides ATCC 17025 GCA_000016405.1
450 Gemmobacter aquatilis GCA_900110025.1
sT
415 Clostridium beijerinckii GCA_000833105.2
395 Clostridium saccharoperbutylacetonicum N1-4_28HMT_29 GCA_000340885.1
392 Clostridium saccharobutylicum DSM 13864 GCA_000473995.1
sU
662 Clostridium saccharoperbutylacetonicum N1-4_28HMT_29 GCA_000340885.1
658 Clostridium beijerinckii GCA_000833105.2
641 Clostridium puniceum GCA_002006345.1
sV
498 Lactobacillus gasseri ATCC 33323 = JCM 1131 GCA_000014425.1
424 Lactobacillus hominis DSM 23910 = CRBIP 24.179 GCA_000296835.1
318 Lactobacillus acetotolerans GCA_001042405.1
sW
510 Rhodobacter sphaeroides 2.4.1 GCA_000012905.2
456 Rhodobacter sphaeroides ATCC 17025 GCA_000016405.1
381 Gemmobacter aquatilis GCA_900110025.1
sX
697 Clostridium beijerinckii GCA_000833105.2
643 Clostridium saccharoperbutylacetonicum N1-4_28HMT_29 GCA_000340885.1
601 Clostridium saccharobutylicum DSM 13864 GCA_000473995.1
sY
699 Deinococcus radiodurans R1 GCA_000008565.1
448 Deinococcus deserti VCD115 GCA_000020685.1
419 Deinococcus proteolyticus MRP GCA_000190555.1
sZ
822 Streptococcus mutans UA159 GCA_000007465.2
601 Streptococcus ratti FA-1 = DSM 20564 GCA_000286075.1
599 Streptococcus ferus DSM 20646 GCA_000372425.1
t0
820 Clostridium beijerinckii GCA_000833105.2
797 Clostridium saccharoperbutylacetonicum N1-4_28HMT_29 GCA_000340885.1
793 Clostridium puniceum GCA_002006345.1
t1
615 Clostridium beijerinckii GCA_000833105.2
608 Clostridium saccharoperbutylacetonicum N1-4_28HMT_29 GCA_000340885.1
582 Clostridium saccharobutylicum DSM 13864 GCA_000473995.1
t2
526 Rhodobacter sphaeroides 2.4.1 GCA_000012905.2
505 Rhodobacter sphaeroides ATCC 17025 GCA_000016405.1
381 Pseudorhodobacter psychrotolerans GCA_001294535.1
t3
533 Lactobacillus gasseri ATCC 33323 = JCM 1131 GCA_000014425.1
460 Lactobacillus hominis DSM 23910 = CRBIP 24.179 GCA_000296835.1
423 Lactobacillus psittaci DSM 15354 GCA_000425905.1
t4
658 Lactobacillus gasseri ATCC 33323 = JCM 1131 GCA_000014425.1
561 Lactobacillus hominis DSM 23910 = CRBIP 24.179 GCA_000296835.1
434 Lactobacillus psittaci DSM 15354 GCA_000425905.1
t5
654 Clostridium beijerinckii GCA_000833105.2

637 Clostridium saccharoperbutylacetonicum N1-4_28HMT_29 GCA_000340885.1
600 Clostridium saccharobutylicum DSM 13864 GCA_000473995.1
t6
261 Rhodobacter sphaeroides 2.4.1 GCA_000012905.2
208 Rhodobacter sphaeroides ATCC 17025 GCA_000016405.1
200 Gemmobacter aquatilis GCA_900110025.1
t7
699 Rhodobacter sphaeroides 2.4.1 GCA_000012905.2
597 Rhodobacter sphaeroides ATCC 17025 GCA_000016405.1
522 Defluviimonas alba GCA_001620265.1
t8
735 Enterococcus faecalis V583 GCA_000007785.1
688 Streptomyces cinnamoneus GCA_001885705.1
603 Enterococcus rivorum GCA_001742285.1
t9
584 Streptococcus mutans UA159 GCA_000007465.2
390 Streptococcus ratti FA-1 = DSM 20564 GCA_000286075.1
348 Streptococcus ferus DSM 20646 GCA_000372425.1
348 Streptococcus henryi DSM 19005 GCA_000376985.1
ta
887 Bacillus anthracis str. Ames GCA_000007845.1
887 Bacillus anthracis str. Sterne GCA_000008165.1
887 _5BBacillus thuringiensis_5D serovar konkukian str. 97-27 GCA_000008505.1
871 Bacillus cereus ATCC 14579 GCA_000007825.1
867 Bacillus thuringiensis YBT-1518 GCA_000497525.2
tb
761 Lactobacillus gasseri ATCC 33323 = JCM 1131 GCA_000014425.1
695 Lactobacillus hominis DSM 23910 = CRBIP 24.179 GCA_000296835.1
487 Lactobacillus hamsteri DSM 5661 = JCM 6256 GCA_000615445.1
tc
733 Clostridium beijerinckii GCA_000833105.2
709 Clostridium saccharoperbutylacetonicum N1-4_28HMT_29 GCA_000340885.1
661 Clostridium saccharobutylicum DSM 13864 GCA_000473995.1
td
357 Rhodobacter sphaeroides 2.4.1 GCA_000012905.2
296 Rhodobacter sphaeroides ATCC 17025 GCA_000016405.1
217 Rhodobacter capsulatus SB 1003 GCA_000021865.1
te
757 Escherichia coli str. K-12 substr. MG1655 GCA_000005845.2
742 Escherichia coli O157_3AH7 str. Sakai GCA_000008865.1
737 Escherichia coli UMN026 GCA_000026325.2
tf
891 Rhodobacter sphaeroides 2.4.1 GCA_000012905.2
786 Rhodobacter sphaeroides ATCC 17025 GCA_000016405.1
625 Gemmobacter megaterium GCA_900156815.1
tg
672 Deinococcus radiodurans R1 GCA_000008565.1
349 Deinococcus deserti VCD115 GCA_000020685.1
333 Deinococcus gobiensis I-0 GCA_000252445.1
th
718 Clostridium beijerinckii GCA_000833105.2
689 Clostridium saccharoperbutylacetonicum N1-4_28HMT_29 GCA_000340885.1
659 Clostridium puniceum GCA_002006345.1
ti
700 Bifidobacterium adolescentis ATCC 15703 GCA_000010425.1
655 Bifidobacterium dentium JCM 1195 = DSM 20436 GCA_001042595.1
611 Bifidobacterium animalis subsp. lactis DSM 10140 GCA_000022965.1
tj
804 Clostridium beijerinckii GCA_000833105.2
770 Clostridium saccharoperbutylacetonicum N1-4_28HMT_29 GCA_000340885.1
729 Clostridium puniceum GCA_002006345.1
tk
579 Enterococcus faecalis V583 GCA_000007785.1
532 Streptomyces cinnamoneus GCA_001885705.1
506 Enterococcus faecium DO GCA_000174395.2
tl
426 Rhodobacter sphaeroides 2.4.1 GCA_000012905.2
354 Gemmobacter megaterium GCA_900156815.1
354 Rhodobacter sphaeroides ATCC 17025 GCA_000016405.1
351 Pseudorhodobacter psychrotolerans GCA_001294535.1
tm
589 Deinococcus radiodurans R1 GCA_000008565.1
380 Deinococcus deserti VCD115 GCA_000020685.1
372 Deinococcus marmoris DSM 12784 GCA_000701405.1
tn
658 Staphylococcus epidermidis ATCC 12228 GCA_000007645.1

637 Staphylococcus haemolyticus JCSC1435 GCA_000009865.1
636 Staphylococcus hominis subsp. hominis C80 GCA_000183685.1
 to
519 Clostridium beijerinckii GCA_000833105.2
493 Clostridium saccharoperbutylacetonicum N1-4_28HMT_29 GCA_000340885.1
454 Clostridium saccharobutylicum DSM 13864 GCA_000473995.1
 tp
558 Deinococcus radiodurans R1 GCA_000008565.1
323 Deinococcus deserti VCD115 GCA_000020685.1
313 Deinococcus gobiensis I-0 GCA_000252445.1
 tq
823 Rhodobacter sphaeroides 2.4.1 GCA_000012905.2
729 Rhodobacter sphaeroides ATCC 17025 GCA_000016405.1
583 Rhodobacter vinaykumarii GCA_900156695.1
 tr
537 Staphylococcus epidermidis ATCC 12228 GCA_000007645.1
506 Staphylococcus capitis subsp. capitis GCA_001028645.1
455 Megasphaera cerevisiae DSM 20462 GCA_001045675.1
455 Staphylococcus warneri SG1 GCA_000332735.1
 ts
752 Bacillus anthracis str. Ames GCA_000007845.1
752 Bacillus anthracis str. Sterne GCA_000008165.1
752 _5BBacillus thuringiensis_5D serovar konkukian str. 97-27 GCA_000008505.1
737 Bacillus cereus ATCC 14579 GCA_000007825.1
727 Bacillus pseudomycoides DSM 12442 GCA_000161455.1
 tt
617 Lactobacillus gasseri ATCC 33323 = JCM 1131 GCA_000014425.1
570 Lactobacillus hominis DSM 23910 = CRBIP 24.179 GCA_000296835.1
415 Lactobacillus iners DSM 13335 GCA_000160875.1
 tu
647 Clostridium beijerinckii GCA_000833105.2
616 Clostridium saccharoperbutylacetonicum N1-4_28HMT_29 GCA_000340885.1
590 Clostridium puniceum GCA_002006345.1
 tv
393 Enterococcus faecalis V583 GCA_000007785.1
370 Enterococcus canis NBRC 100695 GCA_001544375.1
355 Enterococcus thailandicus GCA_001652875.1
 tw
840 Lactobacillus gasseri ATCC 33323 = JCM 1131 GCA_000014425.1
790 Lactobacillus hominis DSM 23910 = CRBIP 24.179 GCA_000296835.1
620 Lactobacillus iners DSM 13335 GCA_000160875.1
 tx
613 Staphylococcus epidermidis ATCC 12228 GCA_000007645.1
589 Staphylococcus capitis subsp. capitis GCA_001028645.1
537 Staphylococcus hominis subsp. hominis C80 GCA_000183685.1
 ty
735 Staphylococcus epidermidis ATCC 12228 GCA_000007645.1
707 Staphylococcus capitis subsp. capitis GCA_001028645.1
666 Staphylococcus aureus subsp. aureus NCTC 8325 GCA_000013425.1
 tz
777 Staphylococcus epidermidis ATCC 12228 GCA_000007645.1
715 Staphylococcus capitis subsp. capitis GCA_001028645.1
679 Staphylococcus hominis subsp. hominis C80 GCA_000183685.1
 tA
563 Escherichia coli str. K-12 substr. MG1655 GCA_000005845.2
545 Escherichia coli O157_3AH7 str. Sakai GCA_000008865.1
545 Shigella flexneri 2a str. 301 GCA_000006925.2
544 Escherichia coli O83_3AH1 str. NRG 857C GCA_000183345.1
544 Escherichia coli UMN026 GCA_000026325.2
 tB
384 Escherichia coli O157_3AH7 str. Sakai GCA_000008865.1
384 Escherichia coli UMN026 GCA_000026325.2
384 Escherichia coli str. K-12 substr. MG1655 GCA_000005845.2
377 Escherichia coli IAI39 GCA_000026345.1
377 Escherichia coli O104_3AH4 str. 2011C-3493 GCA_000299455.1
377 Shigella flexneri 2a str. 301 GCA_000006925.2
376 Enterobacter hormaechei subsp. steigerwaltii GCA_001729725.1
 tC
566 Clostridium beijerinckii GCA_000833105.2
560 Clostridium saccharoperbutylacetonicum N1-4_28HMT_29 GCA_000340885.1
548 Clostridium neonatale GCA_001458595.1
 tD
775 Bacillus anthracis str. Ames GCA_000007845.1
775 Bacillus anthracis str. Sterne GCA_000008165.1
775 Bacillus cereus ATCC 14579 GCA_000007825.1
775 Bacillus thuringiensis YBT-1518 GCA_000497525.2

775 _5BBacillus thuringiensis_5D serovar konkukian str. 97-27 GCA_000008505.1
734 Bacillus pseudomycoides DSM 12442 GCA_000161455.1
731 Bacillus mycoides GCA_000832605.1
tE
694 Staphylococcus epidermidis ATCC 12228 GCA_000007645.1
681 Staphylococcus capitis subsp. capitis GCA_001028645.1
633 Megasphaera cerevisiae DSM 20462 GCA_001045675.1
633 Staphylococcus warneri SG1 GCA_000332735.1
tF
896 Staphylococcus epidermidis ATCC 12228 GCA_000007645.1
856 Staphylococcus capitis subsp. capitis GCA_001028645.1
799 Staphylococcus aureus subsp. aureus NCTC 8325 GCA_000013425.1
tG
798 Deinococcus radiodurans R1 GCA_000008565.1
493 Deinococcus deserti VCD115 GCA_000020685.1
480 Deinococcus gobiensis I-0 GCA_000252445.1
tH
793 Streptococcus mutans UA159 GCA_000007465.2
612 Streptococcus ratti FA-1 = DSM 20564 GCA_000286075.1
558 Streptococcus gordonii str. Challis substr. CH1 GCA_000017005.1
tI
700 Clostridium beijerinckii GCA_000833105.2
668 Clostridium saccharoperbutylacetonicum N1-4_28HMT_29 GCA_000340885.1
630 Clostridium puniceum GCA_002006345.1
tJ
934 Clostridium beijerinckii GCA_000833105.2
921 Clostridium saccharoperbutylacetonicum N1-4_28HMT_29 GCA_000340885.1
885 Clostridium saccharobutylicum DSM 13864 GCA_000473995.1
tK
806 Streptococcus mutans UA159 GCA_000007465.2
618 Streptococcus ratti FA-1 = DSM 20564 GCA_000286075.1
584 Streptococcus gordonii str. Challis substr. CH1 GCA_000017005.1
tL
694 Rhodobacter sphaeroides 2.4.1 GCA_000012905.2
624 Rhodobacter sphaeroides ATCC 17025 GCA_000016405.1
534 Gemmobacter aquatilis GCA_900110025.1
tM
704 Enterococcus faecalis V583 GCA_000007785.1
657 Streptomyces cinnamoneus GCA_001885705.1
607 Enterococcus dispar ATCC 51266 GCA_000406945.1
tN
616 Lactobacillus gasseri ATCC 33323 = JCM 1131 GCA_000014425.1
557 Lactobacillus hominis DSM 23910 = CRBIP 24.179 GCA_000296835.1
398 Lactobacillus acetotolerans GCA_001042405.1
398 Lactobacillus hamsteri DSM 5661 = JCM 6256 GCA_000615445.1
tO
593 Bacillus mycoides GCA_000832605.1
592 Bacillus thuringiensis YBT-1518 GCA_000497525.2
591 Bacillus anthracis str. Ames GCA_000007845.1
591 Bacillus anthracis str. Sterne GCA_000008165.1
591 _5BBacillus thuringiensis_5D serovar konkukian str. 97-27 GCA_000008505.1
tP
673 Streptococcus mutans UA159 GCA_000007465.2
497 Streptococcus ratti FA-1 = DSM 20564 GCA_000286075.1
439 Streptococcus halotolerans GCA_001598035.1
tQ
665 Streptococcus mutans UA159 GCA_000007465.2
468 Streptococcus ratti FA-1 = DSM 20564 GCA_000286075.1
422 Streptococcus sobrinus DSM 20742 = ATCC 33478 GCA_000686605.1
tR
454 Enterococcus faecalis V583 GCA_000007785.1
412 Enterococcus haemoperoxidus ATCC BAA-382 GCA_000407165.1
407 Enterococcus aquimarinus GCA_001885765.1
407 Enterococcus rivorum GCA_001742285.1
407 Streptomyces cinnamoneus GCA_001885705.1
tS
641 Clostridium beijerinckii GCA_000833105.2
593 Clostridium saccharoperbutylacetonicum N1-4_28HMT_29 GCA_000340885.1
572 Clostridium saccharobutylicum DSM 13864 GCA_000473995.1
tT
918 Escherichia coli str. K-12 substr. MG1655 GCA_000005845.2
917 Escherichia coli UMN026 GCA_000026325.2
910 Escherichia coli O157_3AH7 str. Sakai GCA_000008865.1
tU
277 Bifidobacterium adolescentis ATCC 15703 GCA_000010425.1
262 Bifidobacterium asteroides PRL2011 GCA_000304215.1

244 Bifidobacterium coryneforme GCA_000737865.1
tV
606 Staphylococcus epidermidis ATCC 12228 GCA_000007645.1
572 Staphylococcus haemolyticus JCSC1435 GCA_000009865.1
567 Staphylococcus hominis subsp. hominis C80 GCA_000183685.1
tW
707 Rhodobacter sphaeroides 2.4.1 GCA_000012905.2
657 Rhodobacter sphaeroides ATCC 17025 GCA_000016405.1
527 Gemmobacter megaterium GCA_900156815.1
tX
560 Escherichia coli O157_3AH7 str. Sakai GCA_000008865.1
560 Escherichia coli str. K-12 substr. MG1655 GCA_000005845.2
552 Escherichia coli UMN026 GCA_000026325.2
532 Escherichia coli O83_3AH1 str. NRG 857C GCA_000183345.1
tY
688 Staphylococcus epidermidis ATCC 12228 GCA_000007645.1
623 Staphylococcus capitis subsp. capitis GCA_001028645.1
590 Staphylococcus haemolyticus JCSC1435 GCA_000009865.1
590 Staphylococcus hominis subsp. hominis C80 GCA_000183685.1
tZ
821 Rhodobacter sphaeroides 2.4.1 GCA_000012905.2
718 Rhodobacter sphaeroides ATCC 17025 GCA_000016405.1
588 Defluviimonas alba GCA_001620265.1
u0
712 Clostridium beijerinckii GCA_000833105.2
695 Clostridium saccharoperbutylacetonicum N1-4_28HMT_29 GCA_000340885.1
674 Clostridium saccharobutylicum DSM 13864 GCA_000473995.1
u1
627 Rhodobacter sphaeroides 2.4.1 GCA_000012905.2
554 Rhodobacter sphaeroides ATCC 17025 GCA_000016405.1
439 Defluviimonas alba GCA_001620265.1
439 Pseudorhodobacter ferrugineus DSM 5888 GCA_000420745.1
u2
501 Bacillus cereus ATCC 14579 GCA_000007825.1
501 _5BBacillus thuringiensis_5D serovar konkukian str. 97-27 GCA_000008505.1
500 Bacillus anthracis str. Ames GCA_000007845.1
500 Bacillus anthracis str. Sterne GCA_000008165.1
500 Bacillus thuringiensis YBT-1518 GCA_000497525.2
479 Bacillus pseudomycoides DSM 12442 GCA_000161455.1
u3
827 Streptococcus mutans UA159 GCA_000007465.2
645 Streptococcus ratti FA-1 = DSM 20564 GCA_000286075.1
605 Streptococcus halotolerans GCA_001598035.1
u4
561 Deinococcus radiodurans R1 GCA_000008565.1
380 Deinococcus gobiensis I-0 GCA_000252445.1
368 Deinococcus soli Cha et al. 2016 GCA_001007995.1
u5
935 Streptococcus mutans UA159 GCA_000007465.2
670 Streptococcus ratti FA-1 = DSM 20564 GCA_000286075.1
648 Streptococcus gordonii str. Challis substr. CH1 GCA_000017005.1
u6
640 Enterococcus faecalis V583 GCA_000007785.1
595 Streptomyces cinnamoneus GCA_001885705.1
589 Enterococcus haemoperoxidus ATCC BAA-382 GCA_000407165.1
u7
672 Rhodobacter sphaeroides 2.4.1 GCA_000012905.2
604 Rhodobacter sphaeroides ATCC 17025 GCA_000016405.1
542 Pseudorhodobacter psychrotolerans GCA_001294535.1
u8
704 Bacillus anthracis str. Ames GCA_000007845.1
704 Bacillus anthracis str. Sterne GCA_000008165.1
690 _5BBacillus thuringiensis_5D serovar konkukian str. 97-27 GCA_000008505.1
686 Bacillus thuringiensis YBT-1518 GCA_000497525.2
u9
758 Deinococcus radiodurans R1 GCA_000008565.1
510 Deinococcus puniceus GCA_001644565.1
498 Deinococcus soli Cha et al. 2016 GCA_001007995.1
ua
908 Staphylococcus epidermidis ATCC 12228 GCA_000007645.1
846 Staphylococcus capitis subsp. capitis GCA_001028645.1
785 Staphylococcus aureus subsp. aureus NCTC 8325 GCA_000013425.1
ub
661 Rhodobacter sphaeroides 2.4.1 GCA_000012905.2
593 Rhodobacter sphaeroides ATCC 17025 GCA_000016405.1
526 Defluviimonas alba GCA_001620265.1

uc
823 Deinococcus radiodurans R1 GCA_000008565.1
567 Deinococcus deserti VCD115 GCA_000020685.1
537 Deinococcus murrayi DSM 11303 GCA_000482805.1
ud
809 Rhodobacter sphaeroides 2.4.1 GCA_000012905.2
720 Rhodobacter sphaeroides ATCC 17025 GCA_000016405.1
594 Gemmobacter aquatilis GCA_900110025.1
ue
568 Lactobacillus gasseri ATCC 33323 = JCM 1131 GCA_000014425.1
522 Lactobacillus hominis DSM 23910 = CRBIP 24.179 GCA_000296835.1
502 Lactobacillus iners DSM 13335 GCA_000160875.1
uf
616 Shigella flexneri 2a str. 301 GCA_000006925.2
615 Escherichia coli O104_3AH4 str. 2011C-3493 GCA_000299455.1
615 Escherichia coli O157_3AH7 str. Sakai GCA_000008865.1
615 Escherichia coli str. K-12 substr. MG1655 GCA_000005845.2
614 Escherichia coli O83_3AH1 str. NRG 857C GCA_000183345.1
614 Escherichia coli UMN026 GCA_000026325.2
ug
784 Bacillus cereus ATCC 14579 GCA_000007825.1
784 Bacillus thuringiensis YBT-1518 GCA_000497525.2
784 _5BBacillus thuringiensis_5D serovar konkukian str. 97-27 GCA_000008505.1
783 Bacillus anthracis str. Ames GCA_000007845.1
783 Bacillus anthracis str. Sterne GCA_000008165.1
739 Bacillus pseudomycoides DSM 12442 GCA_000161455.1
uh
901 Clostridium beijerinckii GCA_000833105.2
873 Clostridium saccharoperbutylacetonicum N1-4_28HMT_29 GCA_000340885.1
869 Clostridium puniceum GCA_002006345.1
ui
440 Staphylococcus epidermidis ATCC 12228 GCA_000007645.1
406 Staphylococcus capitis subsp. capitis GCA_001028645.1
390 Staphylococcus aureus subsp. aureus NCTC 8325 GCA_000013425.1
uj
774 Staphylococcus epidermidis ATCC 12228 GCA_000007645.1
709 Staphylococcus capitis subsp. capitis GCA_001028645.1
661 Staphylococcus lugdunensis HKU09-01 GCA_000025085.1
uk
674 Bifidobacterium adolescentis ATCC 15703 GCA_000010425.1
576 Bifidobacterium dentium JCM 1195 = DSM 20436 GCA_001042595.1
574 Bifidobacterium angulatum DSM 20098 = JCM 7096 GCA_001025155.1
ul
568 Streptococcus mutans UA159 GCA_000007465.2
449 Streptococcus ratti FA-1 = DSM 20564 GCA_000286075.1
429 Streptococcus ovis DSM 16829 GCA_000380125.1
um
639 Rhodobacter sphaeroides 2.4.1 GCA_000012905.2
525 Rhodobacter sphaeroides ATCC 17025 GCA_000016405.1
455 Gemmobacter aquatilis GCA_900110025.1
un
920 Lactobacillus gasseri ATCC 33323 = JCM 1131 GCA_000014425.1
831 Lactobacillus hominis DSM 23910 = CRBIP 24.179 GCA_000296835.1
617 Lactobacillus iners DSM 13335 GCA_000160875.1
uo
844 Clostridium beijerinckii GCA_000833105.2
802 Clostridium saccharoperbutylacetonicum N1-4_28HMT_29 GCA_000340885.1
776 Clostridium puniceum GCA_002006345.1
up
455 Streptococcus mutans UA159 GCA_000007465.2
313 Streptococcus gordonii str. Challis substr. CH1 GCA_000017005.1
303 Streptococcus ratti FA-1 = DSM 20564 GCA_000286075.1
uq
594 Deinococcus radiodurans R1 GCA_000008565.1
405 Deinococcus deserti VCD115 GCA_000020685.1
385 Deinococcus hopiensis KR-140 GCA_900176165.1
ur
904 Staphylococcus epidermidis ATCC 12228 GCA_000007645.1
861 Staphylococcus capitis subsp. capitis GCA_001028645.1
811 Staphylococcus haemolyticus JCSC1435 GCA_000009865.1
811 Staphylococcus hominis subsp. hominis C80 GCA_000183685.1
us
696 Rhodobacter sphaeroides 2.4.1 GCA_000012905.2
647 Rhodobacter sphaeroides ATCC 17025 GCA_000016405.1
487 Gemmobacter aquatilis GCA_900110025.1
ut

765 Deinococcus radiodurans R1 GCA_000008565.1
472 Deinococcus deserti VCD115 GCA_000020685.1
459 Deinococcus gobiensis I-0 GCA_000252445.1
uu
562 Streptococcus mutans UA159 GCA_000007465.2
414 Streptococcus ratti FA-1 = DSM 20564 GCA_000286075.1
392 Streptococcus sobrinus DSM 20742 = ATCC 33478 GCA_000686605.1
uv
436 Lactobacillus gasseri ATCC 33323 = JCM 1131 GCA_000014425.1
414 Lactobacillus hominis DSM 23910 = CRBIP 24.179 GCA_000296835.1
308 Lactobacillus iners DSM 13335 GCA_000160875.1
uw
645 Streptococcus mutans UA159 GCA_000007465.2
495 Streptococcus ratti FA-1 = DSM 20564 GCA_000286075.1
437 Streptococcus merionis DSM 19192 GCA_000380085.1
ux
725 Lactobacillus gasseri ATCC 33323 = JCM 1131 GCA_000014425.1
694 Lactobacillus hominis DSM 23910 = CRBIP 24.179 GCA_000296835.1
528 Lactobacillus iners DSM 13335 GCA_000160875.1
uy
503 Rhodobacter sphaeroides 2.4.1 GCA_000012905.2
433 Rhodobacter sphaeroides ATCC 17025 GCA_000016405.1
387 Gemmobacter megaterium GCA_900156815.1
uz
818 Clostridium beijerinckii GCA_000833105.2
778 Clostridium saccharoperbutylacetonicum N1-4_28HMT_29 GCA_000340885.1
763 Clostridium puniceum GCA_002006345.1
uA
639 Staphylococcus epidermidis ATCC 12228 GCA_000007645.1
598 Staphylococcus capitis subsp. capitis GCA_001028645.1
538 Staphylococcus aureus subsp. aureus NCTC 8325 GCA_000013425.1
538 Staphylococcus simulans GCA_001559115.1
uB
447 Bifidobacterium adolescentis ATCC 15703 GCA_000010425.1
381 Bifidobacterium longum NCC2705 GCA_000007525.1
375 Bifidobacterium subtile GCA_000741775.1
uC
628 Streptococcus mutans UA159 GCA_000007465.2
460 Streptococcus ratti FA-1 = DSM 20564 GCA_000286075.1
460 Streptococcus sanguinis SK36 GCA_000014205.1
456 Streptococcus ferus DSM 20646 GCA_000372425.1
uD
851 Clostridium beijerinckii GCA_000833105.2
821 Clostridium saccharoperbutylacetonicum N1-4_28HMT_29 GCA_000340885.1
810 Clostridium puniceum GCA_002006345.1
uE
764 Clostridium beijerinckii GCA_000833105.2
732 Clostridium saccharoperbutylacetonicum N1-4_28HMT_29 GCA_000340885.1
711 Clostridium saccharobutylicum DSM 13864 GCA_000473995.1
uF
701 Lactobacillus gasseri ATCC 33323 = JCM 1131 GCA_000014425.1
685 Lactobacillus hominis DSM 23910 = CRBIP 24.179 GCA_000296835.1
546 Lactobacillus jensenii GCA_001936235.1
uG
688 Clostridium beijerinckii GCA_000833105.2
656 Clostridium saccharobutylicum DSM 13864 GCA_000473995.1
654 Clostridium saccharoperbutylacetonicum N1-4_28HMT_29 GCA_000340885.1
uH
644 Lactobacillus gasseri ATCC 33323 = JCM 1131 GCA_000014425.1
617 Lactobacillus hominis DSM 23910 = CRBIP 24.179 GCA_000296835.1
460 Lactobacillus iners DSM 13335 GCA_000160875.1
uI
707 Clostridium beijerinckii GCA_000833105.2
666 Clostridium saccharoperbutylacetonicum N1-4_28HMT_29 GCA_000340885.1
624 Clostridium saccharobutylicum DSM 13864 GCA_000473995.1
uJ
513 Clostridium beijerinckii GCA_000833105.2
513 Clostridium saccharoperbutylacetonicum N1-4_28HMT_29 GCA_000340885.1
476 Clostridium puniceum GCA_002006345.1
454 Clostridium saccharobutylicum DSM 13864 GCA_000473995.1
uK
850 _5BBacillus thuringiensis_5D serovar konkukian str. 97-27 GCA_000008505.1
849 Bacillus anthracis str. Ames GCA_000007845.1
849 Bacillus anthracis str. Sterne GCA_000008165.1
834 Bacillus cereus ATCC 14579 GCA_000007825.1
uL

558 Staphylococcus epidermidis ATCC 12228 GCA_000007645.1
541 Staphylococcus capitis subsp. capitis GCA_001028645.1
537 Staphylococcus haemolyticus JCSC1435 GCA_000009865.1
537 Staphylococcus hominis subsp. hominis C80 GCA_000183685.1
537 Staphylococcus lugdunensis HKU09-01 GCA_000025085.1
uM
656 Enterococcus faecalis V583 GCA_000007785.1
609 Streptomyces cinnamoneus GCA_001885705.1
600 Enterococcus rivorum GCA_001742285.1
uN
458 Clostridium beijerinckii GCA_000833105.2
439 Clostridium saccharoperbutylacetonicum N1-4_28HMT_29 GCA_000340885.1
425 Clostridium puniceum GCA_002006345.1
uO
331 Escherichia coli O157_3AH7 str. Sakai GCA_000008865.1
331 Escherichia coli UMN026 GCA_000026325.2
331 Escherichia coli str. K-12 substr. MG1655 GCA_000005845.2
331 Shigella flexneri 2a str. 301 GCA_000006925.2
325 Escherichia coli O83_3AH1 str. NRG 857C GCA_000183345.1
315 Escherichia coli O104_3AH4 str. 2011C-3493 GCA_000299455.1
uP
701 Lactobacillus gasseri ATCC 33323 = JCM 1131 GCA_000014425.1
638 Lactobacillus hominis DSM 23910 = CRBIP 24.179 GCA_000296835.1
501 Lactobacillus psittaci DSM 15354 GCA_000425905.1
uQ
697 Staphylococcus epidermidis ATCC 12228 GCA_000007645.1
676 Staphylococcus capitis subsp. capitis GCA_001028645.1
648 Staphylococcus warneri SG1 GCA_000332735.1
uR
767 Deinococcus radiodurans R1 GCA_000008565.1
445 Deinococcus deserti VCD115 GCA_000020685.1
423 Deinococcus marmoris DSM 12784 GCA_000701405.1
uS
1022 Enterococcus faecalis V583 GCA_000007785.1
977 Streptomyces cinnamoneus GCA_001885705.1
896 Enterococcus rivorum GCA_001742285.1
uT
546 Rhodobacter sphaeroides 2.4.1 GCA_000012905.2
494 Rhodobacter sphaeroides ATCC 17025 GCA_000016405.1
434 Thioclava atlantica GCA_000737065.1
uU
508 Rhodobacter sphaeroides ATCC 17025 GCA_000016405.1
495 Rhodobacter sphaeroides 2.4.1 GCA_000012905.2
444 Pseudorhodobacter wandonensis GCA_001202035.1
uV
637 Staphylococcus epidermidis ATCC 12228 GCA_000007645.1
587 Staphylococcus capitis subsp. capitis GCA_001028645.1
558 Staphylococcus aureus subsp. aureus NCTC 8325 GCA_000013425.1
558 Staphylococcus warneri SG1 GCA_000332735.1
uW
913 Escherichia coli str. K-12 substr. MG1655 GCA_000005845.2
891 Escherichia coli O83_3AH1 str. NRG 857C GCA_000183345.1
888 Escherichia coli O157_3AH7 str. Sakai GCA_000008865.1
uX
774 Rhodobacter sphaeroides 2.4.1 GCA_000012905.2
682 Rhodobacter sphaeroides ATCC 17025 GCA_000016405.1
554 Pseudorhodobacter ferrugineus DSM 5888 GCA_000420745.1
uY
679 Rhodobacter sphaeroides 2.4.1 GCA_000012905.2
620 Rhodobacter sphaeroides ATCC 17025 GCA_000016405.1
553 Gemmobacter aquatilis GCA_900110025.1
uZ
562 Lactobacillus gasseri ATCC 33323 = JCM 1131 GCA_000014425.1
466 Lactobacillus hominis DSM 23910 = CRBIP 24.179 GCA_000296835.1
269 Lactobacillus hamsteri DSM 5661 = JCM 6256 GCA_000615445.1
v0
527 Rhodobacter sphaeroides 2.4.1 GCA_000012905.2
433 Rhodobacter sphaeroides ATCC 17025 GCA_000016405.1
412 Gemmobacter megaterium GCA_900156815.1
v1
990 Rhodobacter sphaeroides 2.4.1 GCA_000012905.2
912 Rhodobacter sphaeroides ATCC 17025 GCA_000016405.1
804 Gemmobacter aquatilis GCA_900110025.1
v2
536 Streptococcus mutans UA159 GCA_000007465.2
342 Streptococcus criceti HS-6 GCA_000187975.3

341 Streptococcus macacae NCTC 11558 GCA_000187995.3
v3
616 Clostridium beijerinckii GCA_000833105.2
584 Clostridium saccharobutylicum DSM 13864 GCA_000473995.1
584 Clostridium saccharoperbutylacetonicum N1-4_28HMT_29 GCA_000340885.1
569 Clostridium puniceum GCA_002006345.1
v4
595 Deinococcus radiodurans R1 GCA_000008565.1
393 Deinococcus deserti VCD115 GCA_000020685.1
382 Deinococcus gobiensis I-0 GCA_000252445.1
v5
763 Clostridium beijerinckii GCA_000833105.2
735 Clostridium saccharoperbutylacetonicum N1-4_28HMT_29 GCA_000340885.1
714 Clostridium saccharobutylicum DSM 13864 GCA_000473995.1
v6
670 Rhodobacter sphaeroides 2.4.1 GCA_000012905.2
622 Rhodobacter sphaeroides ATCC 17025 GCA_000016405.1
507 Gemmobacter aquatilis GCA_900110025.1
v7
566 Bacillus anthracis str. Ames GCA_000007845.1
566 Bacillus anthracis str. Sterne GCA_000008165.1
566 _5BBacillus thuringiensis_5D serovar konkukian str. 97-27 GCA_000008505.1
562 Bacillus pseudomycoides DSM 12442 GCA_000161455.1
547 Bacillus cereus ATCC 14579 GCA_000007825.1
v8
815 Staphylococcus epidermidis ATCC 12228 GCA_000007645.1
758 Staphylococcus capitis subsp. capitis GCA_001028645.1
745 Staphylococcus aureus subsp. aureus NCTC 8325 GCA_000013425.1
v9
790 Clostridium beijerinckii GCA_000833105.2
725 Clostridium saccharoperbutylacetonicum N1-4_28HMT_29 GCA_000340885.1
706 Clostridium saccharobutylicum DSM 13864 GCA_000473995.1
va
888 Staphylococcus epidermidis ATCC 12228 GCA_000007645.1
835 Staphylococcus capitis subsp. capitis GCA_001028645.1
759 Staphylococcus aureus subsp. aureus NCTC 8325 GCA_000013425.1
vb
699 Staphylococcus epidermidis ATCC 12228 GCA_000007645.1
648 Staphylococcus capitis subsp. capitis GCA_001028645.1
593 Staphylococcus haemolyticus JCSC1435 GCA_000009865.1
593 Staphylococcus hominis subsp. hominis C80 GCA_000183685.1
vc
623 Lactobacillus gasseri ATCC 33323 = JCM 1131 GCA_000014425.1
533 Lactobacillus hominis DSM 23910 = CRBIP 24.179 GCA_000296835.1
426 Lactobacillus apis GCA_000970735.1
vd
540 Enterococcus faecalis V583 GCA_000007785.1
499 Streptomyces cinnamoneus GCA_001885705.1
478 Enterococcus canis NBRC 100695 GCA_001544375.1
ve
437 Staphylococcus epidermidis ATCC 12228 GCA_000007645.1
416 Staphylococcus capitis subsp. capitis GCA_001028645.1
380 Staphylococcus haemolyticus JCSC1435 GCA_000009865.1
vf
558 Streptococcus mutans UA159 GCA_000007465.2
423 Streptococcus ratti FA-1 = DSM 20564 GCA_000286075.1
404 Streptococcus henryi DSM 19005 GCA_000376985.1
vg
626 Lactobacillus gasseri ATCC 33323 = JCM 1131 GCA_000014425.1
552 Lactobacillus hominis DSM 23910 = CRBIP 24.179 GCA_000296835.1
469 Lactobacillus iners DSM 13335 GCA_000160875.1
vh
750 Enterococcus faecalis V583 GCA_000007785.1
703 Streptomyces cinnamoneus GCA_001885705.1
636 Enterococcus rivorum GCA_001742285.1
vi
573 Deinococcus radiodurans R1 GCA_000008565.1
329 Deinococcus gobiensis I-0 GCA_000252445.1
290 Deinococcus deserti VCD115 GCA_000020685.1
vj
659 Enterococcus faecalis V583 GCA_000007785.1
627 Streptomyces cinnamoneus GCA_001885705.1
620 Enterococcus faecium DO GCA_000174395.2
vk
626 Streptococcus mutans UA159 GCA_000007465.2
442 Streptococcus gordonii str. Challis substr. CH1 GCA_000017005.1

393 Streptococcus iniae GCA_000831485.1
vl
516 Bacillus thuringiensis YBT-1518 GCA_000497525.2
501 Bacillus anthracis str. Ames GCA_000007845.1
501 Bacillus anthracis str. Sterne GCA_000008165.1
501 Bacillus cereus ATCC 14579 GCA_000007825.1
501 _5BBacillus thuringiensis_5D serovar konkukian str. 97-27 GCA_000008505.1
475 Bacillus pseudomycoides DSM 12442 GCA_000161455.1
vm
481 Streptococcus mutans UA159 GCA_000007465.2
408 Streptococcus ratti FA-1 = DSM 20564 GCA_000286075.1
372 Streptococcus henryi DSM 19005 GCA_000376985.1
vn
525 Bifidobacterium adolescentis ATCC 15703 GCA_000010425.1
406 Bifidobacterium dentium JCM 1195 = DSM 20436 GCA_001042595.1
391 Bifidobacterium angulatum DSM 20098 = JCM 7096 GCA_001025155.1
vo
679 Clostridium beijerinckii GCA_000833105.2
671 Clostridium saccharoperbutylacetonicum N1-4_28HMT_29 GCA_000340885.1
648 Clostridium saccharobutylicum DSM 13864 GCA_000473995.1
vp
729 Streptococcus mutans UA159 GCA_000007465.2
538 Streptococcus ratti FA-1 = DSM 20564 GCA_000286075.1
518 Streptococcus gordonii str. Challis substr. CH1 GCA_000017005.1
vq
678 Deinococcus radiodurans R1 GCA_000008565.1
469 Deinococcus gobiensis I-0 GCA_000252445.1
459 Deinococcus deserti VCD115 GCA_000020685.1
vr
297 Streptococcus mutans UA159 GCA_000007465.2
185 Streptococcus gordonii str. Challis substr. CH1 GCA_000017005.1
181 Streptococcus equinus GCA_000964315.1
181 Streptococcus gallolyticus subsp. gallolyticus DSM 16831 GCA_002000985.1
181 Streptococcus macacae NCTC 11558 GCA_000187995.3
181 Streptococcus salivarius GCA_000785515.1
181 Streptococcus sanguinis SK36 GCA_000014205.1
181 Streptococcus thermophilus JIM 8232 GCA_000253395.1
vs
641 Bifidobacterium adolescentis ATCC 15703 GCA_000010425.1
553 Bifidobacterium dentium JCM 1195 = DSM 20436 GCA_001042595.1
543 Bifidobacterium tsurumiense GCA_000741765.1
vt
879 Enterococcus faecalis V583 GCA_000007785.1
832 Streptomyces cinnamoneus GCA_001885705.1
747 Enterococcus canis NBRC 100695 GCA_001544375.1
vu
574 Rhodobacter sphaeroides 2.4.1 GCA_000012905.2
510 Rhodobacter sphaeroides ATCC 17025 GCA_000016405.1
464 Gemmobacter aquatilis GCA_900110025.1
vv
781 Lactobacillus gasseri ATCC 33323 = JCM 1131 GCA_000014425.1
737 Lactobacillus hominis DSM 23910 = CRBIP 24.179 GCA_000296835.1
609 Lactobacillus iners DSM 13335 GCA_000160875.1
vw
612 Rhodobacter sphaeroides 2.4.1 GCA_000012905.2
538 Rhodobacter sphaeroides ATCC 17025 GCA_000016405.1
450 Gemmobacter aquatilis GCA_900110025.1
vx
698 Escherichia coli str. K-12 substr. MG1655 GCA_000005845.2
685 Escherichia coli O157_3AH7 str. Sakai GCA_000008865.1
685 Escherichia coli O83_3AH1 str. NRG 857C GCA_000183345.1
685 Escherichia coli UMN026 GCA_000026325.2
685 Shigella flexneri 2a str. 301 GCA_000006925.2
670 Escherichia coli IAI39 GCA_000026345.1
vy
591 Staphylococcus epidermidis ATCC 12228 GCA_000007645.1
556 Staphylococcus capitis subsp. capitis GCA_001028645.1
505 Staphylococcus warneri SG1 GCA_000332735.1
vz
539 Staphylococcus epidermidis ATCC 12228 GCA_000007645.1
520 Staphylococcus capitis subsp. capitis GCA_001028645.1
514 Staphylococcus haemolyticus JCSC1435 GCA_000009865.1
vA
267 Rhodobacter sphaeroides ATCC 17025 GCA_000016405.1
266 Rhodobacter sphaeroides 2.4.1 GCA_000012905.2
218 Rhodobacter aestuarii GCA_900156655.1

vB
634 Escherichia coli UMN026 GCA_000026325.2
634 Escherichia coli str. K-12 substr. MG1655 GCA_000005845.2
621 Escherichia coli O157_3AH7 str. Sakai GCA_000008865.1
619 Shigella flexneri 2a str. 301 GCA_000006925.2
vC
943 Lactobacillus gasseri ATCC 33323 = JCM 1131 GCA_000014425.1
902 Lactobacillus hominis DSM 23910 = CRBIP 24.179 GCA_000296835.1
678 Lactobacillus iners DSM 13335 GCA_000160875.1
vD
614 Staphylococcus epidermidis ATCC 12228 GCA_000007645.1
609 Staphylococcus capitis subsp. capitis GCA_001028645.1
568 Staphylococcus hominis subsp. hominis C80 GCA_000183685.1
vE
418 Streptococcus mutans UA159 GCA_000007465.2
327 Streptococcus merionis DSM 19192 GCA_000380085.1
321 Streptococcus gordonii str. Challis substr. CH1 GCA_000017005.1
vF
763 Lactobacillus gasseri ATCC 33323 = JCM 1131 GCA_000014425.1
729 Lactobacillus hominis DSM 23910 = CRBIP 24.179 GCA_000296835.1
518 Lactobacillus iners DSM 13335 GCA_000160875.1
vG
628 Escherichia coli str. K-12 substr. MG1655 GCA_000005845.2
623 Escherichia coli IAI39 GCA_000026345.1
617 Escherichia coli UMN026 GCA_000026325.2
vH
747 Clostridium beijerinckii GCA_000833105.2
703 Clostridium saccharoperbutylacetonicum N1-4_28HMT_29 GCA_000340885.1
675 Clostridium saccharobutylicum DSM 13864 GCA_000473995.1
vI
580 Rhodobacter sphaeroides 2.4.1 GCA_000012905.2
520 Rhodobacter sphaeroides ATCC 17025 GCA_000016405.1
374 Pseudorhodobacter ferrugineus DSM 5888 GCA_000420745.1
vJ
837 Escherichia coli str. K-12 substr. MG1655 GCA_000005845.2
832 Escherichia coli UMN026 GCA_000026325.2
830 Escherichia coli O104_3AH4 str. 2011C-3493 GCA_000299455.1
830 Shigella flexneri 2a str. 301 GCA_000006925.2
vK
838 Streptococcus mutans UA159 GCA_000007465.2
600 Streptococcus ratti FA-1 = DSM 20564 GCA_000286075.1
559 Streptococcus gallolyticus subsp. gallolyticus DSM 16831 GCA_002000985.1
vL
762 Clostridium beijerinckii GCA_000833105.2
725 Clostridium saccharoperbutylacetonicum N1-4_28HMT_29 GCA_000340885.1
703 Clostridium puniceum GCA_002006345.1
vM
863 Bifidobacterium adolescentis ATCC 15703 GCA_000010425.1
743 Bifidobacterium dentium JCM 1195 = DSM 20436 GCA_001042595.1
668 Bifidobacterium angulatum DSM 20098 = JCM 7096 GCA_001025155.1
vN
694 Escherichia coli O104_3AH4 str. 2011C-3493 GCA_000299455.1
693 Escherichia coli str. K-12 substr. MG1655 GCA_000005845.2
671 Shigella flexneri 2a str. 301 GCA_000006925.2
vO
646 Staphylococcus epidermidis ATCC 12228 GCA_000007645.1
601 Staphylococcus capitis subsp. capitis GCA_001028645.1
595 Staphylococcus aureus subsp. aureus NCTC 8325 GCA_000013425.1
vP
886 Clostridium beijerinckii GCA_000833105.2
872 Clostridium saccharoperbutylacetonicum N1-4_28HMT_29 GCA_000340885.1
851 Clostridium saccharobutylicum DSM 13864 GCA_000473995.1
vQ
471 Streptococcus mutans UA159 GCA_000007465.2
375 Streptococcus ratti FA-1 = DSM 20564 GCA_000286075.1
358 Streptococcus macacae NCTC 11558 GCA_000187995.3
vR
420 Rhodobacter sphaeroides 2.4.1 GCA_000012905.2
382 Rhodobacter sphaeroides ATCC 17025 GCA_000016405.1
316 Defluviimonas alba GCA_001620265.1
vS
642 Deinococcus radiodurans R1 GCA_000008565.1
437 Deinococcus deserti VCD115 GCA_000020685.1
414 Deinococcus puniceus GCA_001644565.1
vT
845 Escherichia coli str. K-12 substr. MG1655 GCA_000005845.2

821 Escherichia coli O104_3AH4 str. 2011C-3493 GCA_000299455.1
821 Shigella flexneri 2a str. 301 GCA_000006925.2
796 Escherichia coli O157_3AH7 str. Sakai GCA_000008865.1
796 Escherichia coli UMN026 GCA_000026325.2
vU
638 Staphylococcus epidermidis ATCC 12228 GCA_000007645.1
581 Staphylococcus capitis subsp. capitis GCA_001028645.1
524 Staphylococcus hominis subsp. hominis C80 GCA_000183685.1
vV
487 Clostridium beijerinckii GCA_000833105.2
463 Clostridium puniceum GCA_002006345.1
463 Clostridium saccharoperbutylacetonicum N1-4_28HMT_29 GCA_000340885.1
445 Clostridium saccharobutylicum DSM 13864 GCA_000473995.1
vW
329 Rhodobacter sphaeroides 2.4.1 GCA_000012905.2
291 Rhodobacter sphaeroides ATCC 17025 GCA_000016405.1
274 Paracoccus solventivorans GCA_900142875.1
vX
662 Enterococcus faecalis V583 GCA_000007785.1
615 Streptomyces cinnamoneus GCA_001885705.1
610 Staphylococcus epidermidis ATCC 12228 GCA_000007645.1
vY
508 Streptococcus mutans UA159 GCA_000007465.2
362 Streptococcus ratti FA-1 = DSM 20564 GCA_000286075.1
352 Streptococcus sobrinus DSM 20742 = ATCC 33478 GCA_000686605.1
vZ
489 Clostridium beijerinckii GCA_000833105.2
474 Clostridium saccharoperbutylacetonicum N1-4_28HMT_29 GCA_000340885.1
453 Clostridium puniceum GCA_002006345.1
w0
656 Rhodobacter sphaeroides 2.4.1 GCA_000012905.2
595 Rhodobacter sphaeroides ATCC 17025 GCA_000016405.1
519 Gemmobacter aquatilis GCA_900110025.1
w1
642 Enterococcus faecalis V583 GCA_000007785.1
602 Enterococcus rivorum GCA_001742285.1
595 Streptomyces cinnamoneus GCA_001885705.1
w2
855 Deinococcus radiodurans R1 GCA_000008565.1
535 Deinococcus deserti VCD115 GCA_000020685.1
511 Deinococcus gobiensis I-0 GCA_000252445.1
w3
589 Deinococcus radiodurans R1 GCA_000008565.1
400 Deinococcus deserti VCD115 GCA_000020685.1
391 Deinococcus gobiensis I-0 GCA_000252445.1
w4
699 Lactobacillus gasseri ATCC 33323 = JCM 1131 GCA_000014425.1
679 Lactobacillus hominis DSM 23910 = CRBIP 24.179 GCA_000296835.1
492 Lactobacillus iners DSM 13335 GCA_000160875.1
w5
768 Enterococcus faecalis V583 GCA_000007785.1
722 Streptomyces cinnamoneus GCA_001885705.1
674 Enterococcus rivorum GCA_001742285.1
w6
859 Enterococcus faecalis V583 GCA_000007785.1
812 Streptomyces cinnamoneus GCA_001885705.1
736 Enterococcus canis NBRC 100695 GCA_001544375.1
w7
801 Rhodobacter sphaeroides 2.4.1 GCA_000012905.2
708 Rhodobacter sphaeroides ATCC 17025 GCA_000016405.1
591 Pseudorhodobacter ferrugineus DSM 5888 GCA_000420745.1
w8
710 Escherichia coli O157_3AH7 str. Sakai GCA_000008865.1
710 Escherichia coli UMN026 GCA_000026325.2
710 Escherichia coli str. K-12 substr. MG1655 GCA_000005845.2
706 Escherichia coli O104_3AH4 str. 2011C-3493 GCA_000299455.1
706 Escherichia coli O83_3AH1 str. NRG 857C GCA_000183345.1
706 Shigella flexneri 2a str. 301 GCA_000006925.2
676 Escherichia coli IAI39 GCA_000026345.1
676 Shigella dysenteriae Sd197 GCA_000012005.1
w9
839 Enterococcus faecalis V583 GCA_000007785.1
792 Streptomyces cinnamoneus GCA_001885705.1
672 Enterococcus rivorum GCA_001742285.1
wa
915 Clostridium beijerinckii GCA_000833105.2

842 Clostridium saccharobutylicum DSM 13864 GCA_000473995.1
841 Clostridium saccharoperbutylacetonicum N1-4_28HMT_29 GCA_000340885.1
wb
566 Escherichia coli O104_3AH4 str. 2011C-3493 GCA_000299455.1
566 Escherichia coli str. K-12 substr. MG1655 GCA_000005845.2
559 Shigella flexneri 2a str. 301 GCA_000006925.2
557 Escherichia coli UMN026 GCA_000026325.2
wc
420 Enterococcus faecalis V583 GCA_000007785.1
373 Streptomyces cinnamoneus GCA_001885705.1
356 Enterococcus haemoperoxidus ATCC BAA-382 GCA_000407165.1
wd
766 Bacillus anthracis str. Ames GCA_000007845.1
766 Bacillus anthracis str. Sterne GCA_000008165.1
766 _5BBacillus thuringiensis_5D serovar konkukian str. 97-27 GCA_000008505.1
740 Bacillus cereus ATCC 14579 GCA_000007825.1
734 Bacillus pseudomycoides DSM 12442 GCA_000161455.1
we
685 Deinococcus radiodurans R1 GCA_000008565.1
390 Deinococcus deserti VCD115 GCA_000020685.1
382 Deinococcus murrayi DSM 11303 GCA_000482805.1
wf
747 Deinococcus radiodurans R1 GCA_000008565.1
436 Deinococcus deserti VCD115 GCA_000020685.1
421 Deinococcus gobiensis I-0 GCA_000252445.1
wg
700 Streptococcus mutans UA159 GCA_000007465.2
526 Streptococcus ratti FA-1 = DSM 20564 GCA_000286075.1
518 Streptococcus gordonii str. Challis substr. CH1 GCA_000017005.1
wh
808 Bifidobacterium adolescentis ATCC 15703 GCA_000010425.1
720 Bifidobacterium dentium JCM 1195 = DSM 20436 GCA_001042595.1
673 Bifidobacterium angulatum DSM 20098 = JCM 7096 GCA_001025155.1
wi
698 Streptococcus mutans UA159 GCA_000007465.2
479 Streptococcus ratti FA-1 = DSM 20564 GCA_000286075.1
449 Streptococcus equinus GCA_000964315.1
wj
772 Escherichia coli O104_3AH4 str. 2011C-3493 GCA_000299455.1
772 Escherichia coli str. K-12 substr. MG1655 GCA_000005845.2
758 Shigella flexneri 2a str. 301 GCA_000006925.2
741 Escherichia coli O157_3AH7 str. Sakai GCA_000008865.1
wk
705 Lactobacillus gasseri ATCC 33323 = JCM 1131 GCA_000014425.1
645 Lactobacillus hominis DSM 23910 = CRBIP 24.179 GCA_000296835.1
472 Lactobacillus iners DSM 13335 GCA_000160875.1
wl
761 Deinococcus radiodurans R1 GCA_000008565.1
445 Deinococcus deserti VCD115 GCA_000020685.1
436 Deinococcus puniceus GCA_001644565.1
wm
767 Lactobacillus gasseri ATCC 33323 = JCM 1131 GCA_000014425.1
682 Lactobacillus hominis DSM 23910 = CRBIP 24.179 GCA_000296835.1
546 Lactobacillus iners DSM 13335 GCA_000160875.1
wn
577 Clostridium beijerinckii GCA_000833105.2
534 Clostridium saccharoperbutylacetonicum N1-4_28HMT_29 GCA_000340885.1
517 Clostridium puniceum GCA_002006345.1
wo
503 Lactobacillus gasseri ATCC 33323 = JCM 1131 GCA_000014425.1
455 Lactobacillus hominis DSM 23910 = CRBIP 24.179 GCA_000296835.1
309 Lactobacillus hamsteri DSM 5661 = JCM 6256 GCA_000615445.1
wp
850 Deinococcus radiodurans R1 GCA_000008565.1
562 Deinococcus deserti VCD115 GCA_000020685.1
542 Deinococcus puniceus GCA_001644565.1
wq
711 Bifidobacterium adolescentis ATCC 15703 GCA_000010425.1
638 Bifidobacterium dentium JCM 1195 = DSM 20436 GCA_001042595.1
585 Bifidobacterium angulatum DSM 20098 = JCM 7096 GCA_001025155.1
wr
842 Staphylococcus epidermidis ATCC 12228 GCA_000007645.1
797 Staphylococcus capitis subsp. capitis GCA_001028645.1
736 Staphylococcus hominis subsp. hominis C80 GCA_000183685.1
ws
565 Staphylococcus epidermidis ATCC 12228 GCA_000007645.1

536 Staphylococcus capitis subsp. capitis GCA_001028645.1
518 Staphylococcus warneri SG1 GCA_000332735.1
wt
667 Deinococcus radiodurans R1 GCA_000008565.1
442 Deinococcus deserti VCD115 GCA_000020685.1
396 Deinococcus soli Cha et al. 2016 GCA_001007995.1
wu
615 Bifidobacterium adolescentis ATCC 15703 GCA_000010425.1
504 Bifidobacterium dentium JCM 1195 = DSM 20436 GCA_001042595.1
451 Bifidobacterium angulatum DSM 20098 = JCM 7096 GCA_001025155.1
wv
806 Deinococcus radiodurans R1 GCA_000008565.1
584 Deinococcus deserti VCD115 GCA_000020685.1
550 Deinococcus gobiensis I-0 GCA_000252445.1
ww
580 Staphylococcus epidermidis ATCC 12228 GCA_000007645.1
534 Staphylococcus capitis subsp. capitis GCA_001028645.1
483 Staphylococcus warneri SG1 GCA_000332735.1
wx
770 Staphylococcus epidermidis ATCC 12228 GCA_000007645.1
724 Staphylococcus capitis subsp. capitis GCA_001028645.1
696 Staphylococcus haemolyticus JCSC1435 GCA_000009865.1
wy
548 Staphylococcus epidermidis ATCC 12228 GCA_000007645.1
487 Staphylococcus capitis subsp. capitis GCA_001028645.1
419 Staphylococcus hominis subsp. hominis C80 GCA_000183685.1
wz
474 Streptococcus mutans UA159 GCA_000007465.2
344 Streptococcus ratti FA-1 = DSM 20564 GCA_000286075.1
296 Streptococcus gordonii str. Challis substr. CH1 GCA_000017005.1
wA
970 Deinococcus radiodurans R1 GCA_000008565.1
612 Deinococcus deserti VCD115 GCA_000020685.1
582 Deinococcus gobiensis I-0 GCA_000252445.1
wB
765 Enterococcus faecalis V583 GCA_000007785.1
718 Streptomyces cinnamoneus GCA_001885705.1
652 Enterococcus thailandicus GCA_001652875.1
wC
473 Escherichia coli str. K-12 substr. MG1655 GCA_000005845.2
458 Escherichia coli O157_3AH7 str. Sakai GCA_000008865.1
458 Shigella flexneri 2a str. 301 GCA_000006925.2
455 Escherichia coli O83_3AH1 str. NRG 857C GCA_000183345.1
455 Escherichia coli UMN026 GCA_000026325.2
wD
381 Lactobacillus gasseri ATCC 33323 = JCM 1131 GCA_000014425.1
316 Lactobacillus hominis DSM 23910 = CRBIP 24.179 GCA_000296835.1
276 Lactobacillus iners DSM 13335 GCA_000160875.1
wE
525 Clostridium beijerinckii GCA_000833105.2
473 Clostridium saccharoperbutylacetonicum N1-4_28HMT_29 GCA_000340885.1
467 Clostridium puniceum GCA_002006345.1
wF
688 Clostridium beijerinckii GCA_000833105.2
648 Clostridium saccharoperbutylacetonicum N1-4_28HMT_29 GCA_000340885.1
624 Clostridium saccharobutylicum DSM 13864 GCA_000473995.1
wG
817 Staphylococcus epidermidis ATCC 12228 GCA_000007645.1
779 Staphylococcus capitis subsp. capitis GCA_001028645.1
746 Staphylococcus pettenkoferi GCA_002208805.1
wH
711 Deinococcus radiodurans R1 GCA_000008565.1
514 Deinococcus deserti VCD115 GCA_000020685.1
498 Deinococcus gobiensis I-0 GCA_000252445.1
wI
760 Lactobacillus gasseri ATCC 33323 = JCM 1131 GCA_000014425.1
693 Lactobacillus hominis DSM 23910 = CRBIP 24.179 GCA_000296835.1
552 Lactobacillus iners DSM 13335 GCA_000160875.1
wJ
850 Rhodobacter sphaeroides 2.4.1 GCA_000012905.2
766 Rhodobacter sphaeroides ATCC 17025 GCA_000016405.1
666 Defluviimonas alba GCA_001620265.1
wK
735 Staphylococcus epidermidis ATCC 12228 GCA_000007645.1
686 Staphylococcus capitis subsp. capitis GCA_001028645.1
656 Staphylococcus warneri SG1 GCA_000332735.1

wL
611 Clostridium beijerinckii GCA_000833105.2
584 Clostridium saccharoperbutylacetonicum N1-4_28HMT_29 GCA_000340885.1
554 Clostridium puniceum GCA_002006345.1
wM
826 _5BBacillus thuringiensis_5D serovar konkukian str. 97-27 GCA_000008505.1
825 Bacillus anthracis str. Ames GCA_000007845.1
825 Bacillus anthracis str. Sterne GCA_000008165.1
825 Bacillus cereus ATCC 14579 GCA_000007825.1
825 Bacillus thuringiensis YBT-1518 GCA_000497525.2
809 Bacillus mycoides GCA_000832605.1
809 Bacillus pseudomycoides DSM 12442 GCA_000161455.1
wN
638 Streptococcus mutans UA159 GCA_000007465.2
454 Streptococcus sobrinus DSM 20742 = ATCC 33478 GCA_000686605.1
421 Streptococcus dysgalactiae subsp. equisimilis AC-2713 GCA_000317855.1
wO
743 Rhodobacter sphaeroides 2.4.1 GCA_000012905.2
619 Rhodobacter sphaeroides ATCC 17025 GCA_000016405.1
534 Thioclava indica GCA_000714545.1
wP
791 Streptococcus mutans UA159 GCA_000007465.2
554 Streptococcus ratti FA-1 = DSM 20564 GCA_000286075.1
522 Streptococcus gordonii str. Challis substr. CH1 GCA_000017005.1
wQ
694 _5BBacillus thuringiensis_5D serovar konkukian str. 97-27 GCA_000008505.1
683 Bacillus anthracis str. Ames GCA_000007845.1
683 Bacillus anthracis str. Sterne GCA_000008165.1
653 Bacillus cereus ATCC 14579 GCA_000007825.1
653 Bacillus thuringiensis YBT-1518 GCA_000497525.2
wR
458 Lactobacillus gasseri ATCC 33323 = JCM 1131 GCA_000014425.1
403 Lactobacillus hominis DSM 23910 = CRBIP 24.179 GCA_000296835.1
300 Lactobacillus amylolyticus GCA_002075105.1
300 Lactobacillus hamsteri DSM 5661 = JCM 6256 GCA_000615445.1
wS
784 Enterococcus faecalis V583 GCA_000007785.1
737 Streptomyces cinnamoneus GCA_001885705.1
659 Enterococcus asini ATCC 700915 GCA_000407365.1
wT
473 Clostridium beijerinckii GCA_000833105.2
437 Clostridium saccharoperbutylacetonicum N1-4_28HMT_29 GCA_000340885.1
423 Clostridium saccharobutylicum DSM 13864 GCA_000473995.1
wU
652 Streptococcus mutans UA159 GCA_000007465.2
486 Streptococcus ratti FA-1 = DSM 20564 GCA_000286075.1
462 Streptococcus gordonii str. Challis substr. CH1 GCA_000017005.1
wV
729 Clostridium beijerinckii GCA_000833105.2
695 Clostridium saccharobutylicum DSM 13864 GCA_000473995.1
679 Clostridium saccharoperbutylacetonicum N1-4_28HMT_29 GCA_000340885.1
wW
916 Streptococcus mutans UA159 GCA_000007465.2
769 Streptococcus ratti FA-1 = DSM 20564 GCA_000286075.1
667 Streptococcus ferus DSM 20646 GCA_000372425.1
wX
823 Staphylococcus epidermidis ATCC 12228 GCA_000007645.1
772 Staphylococcus capitis subsp. capitis GCA_001028645.1
700 Staphylococcus lugdunensis HKU09-01 GCA_000025085.1
wY
805 Rhodobacter sphaeroides 2.4.1 GCA_000012905.2
698 Rhodobacter sphaeroides ATCC 17025 GCA_000016405.1
585 Gemmobacter megaterium GCA_900156815.1
wZ
562 Escherichia coli str. K-12 substr. MG1655 GCA_000005845.2
551 Escherichia coli O104_3AH4 str. 2011C-3493 GCA_000299455.1
536 Escherichia coli IAI39 GCA_000026345.1
536 Escherichia coli UMN026 GCA_000026325.2
x0
571 Rhodobacter sphaeroides 2.4.1 GCA_000012905.2
548 Rhodobacter sphaeroides ATCC 17025 GCA_000016405.1
399 Gemmobacter megaterium GCA_900156815.1
x1
705 Deinococcus radiodurans R1 GCA_000008565.1
511 Deinococcus deserti VCD115 GCA_000020685.1
493 Deinococcus gobiensis I-0 GCA_000252445.1

x2
623 Rhodobacter sphaeroides 2.4.1 GCA_000012905.2
561 Rhodobacter sphaeroides ATCC 17025 GCA_000016405.1
515 Gemmobacter aquatilis GCA_900110025.1
x3
486 Rhodobacter sphaeroides 2.4.1 GCA_000012905.2
411 Rhodobacter sphaeroides ATCC 17025 GCA_000016405.1
377 Pseudorhodobacter psychrotolerans GCA_001294535.1
x4
652 Rhodobacter sphaeroides 2.4.1 GCA_000012905.2
602 Rhodobacter sphaeroides ATCC 17025 GCA_000016405.1
497 Gemmobacter aquatilis GCA_900110025.1
x5
380 Rhodobacter sphaeroides 2.4.1 GCA_000012905.2
364 Rhodobacter sphaeroides ATCC 17025 GCA_000016405.1
331 Defluviimonas alba GCA_001620265.1
x6
461 Clostridium beijerinckii GCA_000833105.2
460 Clostridium saccharoperbutylacetonicum N1-4_28HMT_29 GCA_000340885.1
436 Clostridium puniceum GCA_002006345.1
x7
676 Bifidobacterium adolescentis ATCC 15703 GCA_000010425.1
565 Bifidobacterium dentium JCM 1195 = DSM 20436 GCA_001042595.1
529 Bifidobacterium callitrichos DSM 23973 GCA_000741175.1
x8
756 Clostridium beijerinckii GCA_000833105.2
738 Clostridium saccharoperbutylacetonicum N1-4_28HMT_29 GCA_000340885.1
722 Clostridium puniceum GCA_002006345.1
x9
768 Clostridium beijerinckii GCA_000833105.2
725 Clostridium saccharoperbutylacetonicum N1-4_28HMT_29 GCA_000340885.1
712 Clostridium saccharobutylicum DSM 13864 GCA_000473995.1
xa
743 Clostridium beijerinckii GCA_000833105.2
691 Clostridium saccharoperbutylacetonicum N1-4_28HMT_29 GCA_000340885.1
663 Clostridium puniceum GCA_002006345.1
xb
697 Staphylococcus epidermidis ATCC 12228 GCA_000007645.1
651 Staphylococcus capitis subsp. capitis GCA_001028645.1
605 Staphylococcus aureus subsp. aureus NCTC 8325 GCA_000013425.1
xc
554 Staphylococcus epidermidis ATCC 12228 GCA_000007645.1
540 Staphylococcus capitis subsp. capitis GCA_001028645.1
479 Staphylococcus warneri SG1 GCA_000332735.1
xd
798 Clostridium beijerinckii GCA_000833105.2
783 Clostridium saccharoperbutylacetonicum N1-4_28HMT_29 GCA_000340885.1
768 Clostridium saccharobutylicum DSM 13864 GCA_000473995.1
xe
696 Clostridium beijerinckii GCA_000833105.2
676 Clostridium saccharoperbutylacetonicum N1-4_28HMT_29 GCA_000340885.1
647 Clostridium saccharobutylicum DSM 13864 GCA_000473995.1
xf
475 Rhodobacter sphaeroides 2.4.1 GCA_000012905.2
415 Rhodobacter sphaeroides ATCC 17025 GCA_000016405.1
351 Gemmobacter aquatilis GCA_900110025.1
xg
660 Deinococcus radiodurans R1 GCA_000008565.1
463 Deinococcus deserti VCD115 GCA_000020685.1
427 Deinococcus soli Cha et al. 2016 GCA_001007995.1
xh
902 Lactobacillus gasseri ATCC 33323 = JCM 1131 GCA_000014425.1
815 Lactobacillus hominis DSM 23910 = CRBIP 24.179 GCA_000296835.1
536 Lactobacillus psittaci DSM 15354 GCA_000425905.1
xi
449 Rhodobacter sphaeroides 2.4.1 GCA_000012905.2
379 Rhodobacter sphaeroides ATCC 17025 GCA_000016405.1
281 Thioclava indica GCA_000714545.1
xj
913 Rhodobacter sphaeroides 2.4.1 GCA_000012905.2
860 Rhodobacter sphaeroides ATCC 17025 GCA_000016405.1
739 Gemmobacter aquatilis GCA_900110025.1
xk
836 Escherichia coli O157_3AH7 str. Sakai GCA_000008865.1
836 Escherichia coli str. K-12 substr. MG1655 GCA_000005845.2
830 Escherichia coli UMN026 GCA_000026325.2

821 Shigella flexneri 2a str. 301 GCA_000006925.2
xl
839 Streptococcus mutans UA159 GCA_000007465.2
655 Streptococcus ratti FA-1 = DSM 20564 GCA_000286075.1
629 Streptococcus gordonii str. Challis substr. CH1 GCA_000017005.1
xm
532 Lactobacillus gasseri ATCC 33323 = JCM 1131 GCA_000014425.1
492 Lactobacillus hominis DSM 23910 = CRBIP 24.179 GCA_000296835.1
380 Lactobacillus iners DSM 13335 GCA_000160875.1
xn
705 Bacillus anthracis str. Ames GCA_000007845.1
705 Bacillus anthracis str. Sterne GCA_000008165.1
705 Bacillus cereus ATCC 14579 GCA_000007825.1
705 _5BBacillus thuringiensis_5D serovar konkukian str. 97-27 GCA_000008505.1
695 Bacillus thuringiensis YBT-1518 GCA_000497525.2
671 Bacillus pseudomycoides DSM 12442 GCA_000161455.1
xo
854 Staphylococcus epidermidis ATCC 12228 GCA_000007645.1
831 Staphylococcus capitis subsp. capitis GCA_001028645.1
788 Staphylococcus aureus subsp. aureus NCTC 8325 GCA_000013425.1
xp
738 Deinococcus radiodurans R1 GCA_000008565.1
518 Deinococcus gobiensis I-0 GCA_000252445.1
475 Deinococcus deserti VCD115 GCA_000020685.1
xq
617 Deinococcus radiodurans R1 GCA_000008565.1
384 Deinococcus gobiensis I-0 GCA_000252445.1
372 Deinococcus hopiensis KR-140 GCA_900176165.1
xr
807 Clostridium beijerinckii GCA_000833105.2
743 Clostridium saccharoperbutylacetonicum N1-4_28HMT_29 GCA_000340885.1
738 Clostridium puniceum GCA_002006345.1
xs
683 Enterococcus faecalis V583 GCA_000007785.1
636 Streptomyces cinnamoneus GCA_001885705.1
565 Enterococcus faecium DO GCA_000174395.2
xt
698 Clostridium beijerinckii GCA_000833105.2
621 Clostridium saccharoperbutylacetonicum N1-4_28HMT_29 GCA_000340885.1
607 Clostridium saccharobutylicum DSM 13864 GCA_000473995.1
xu
538 Rhodobacter sphaeroides 2.4.1 GCA_000012905.2
478 Rhodobacter sphaeroides ATCC 17025 GCA_000016405.1
367 Pseudorhodobacter ferrugineus DSM 5888 GCA_000420745.1
xv
514 Staphylococcus epidermidis ATCC 12228 GCA_000007645.1
493 Staphylococcus capitis subsp. capitis GCA_001028645.1
481 Megasphaera cerevisiae DSM 20462 GCA_001045675.1
481 Staphylococcus warneri SG1 GCA_000332735.1
xw
661 Bacillus anthracis str. Ames GCA_000007845.1
661 Bacillus anthracis str. Sterne GCA_000008165.1
661 _5BBacillus thuringiensis_5D serovar konkukian str. 97-27 GCA_000008505.1
651 Bacillus cereus ATCC 14579 GCA_000007825.1
645 Bacillus thuringiensis YBT-1518 GCA_000497525.2
xx
414 Rhodobacter sphaeroides 2.4.1 GCA_000012905.2
327 Rhodobacter sphaeroides ATCC 17025 GCA_000016405.1
313 Defluviimonas alba GCA_001620265.1
xy
645 Lactobacillus gasseri ATCC 33323 = JCM 1131 GCA_000014425.1
623 Lactobacillus hominis DSM 23910 = CRBIP 24.179 GCA_000296835.1
483 Lactobacillus iners DSM 13335 GCA_000160875.1
xz
744 Streptococcus mutans UA159 GCA_000007465.2
582 Streptococcus ratti FA-1 = DSM 20564 GCA_000286075.1
526 Streptococcus gordonii str. Challis substr. CH1 GCA_000017005.1
xA
823 Streptococcus mutans UA159 GCA_000007465.2
647 Streptococcus ratti FA-1 = DSM 20564 GCA_000286075.1
619 Streptococcus gordonii str. Challis substr. CH1 GCA_000017005.1
xB
814 Enterococcus faecalis V583 GCA_000007785.1
767 Streptomyces cinnamoneus GCA_001885705.1
728 Enterococcus rivorum GCA_001742285.1
xC

506 Rhodobacter sphaeroides 2.4.1 GCA_000012905.2
490 Rhodobacter sphaeroides ATCC 17025 GCA_000016405.1
391 Gemmobacter megaterium GCA_900156815.1
xD
443 Deinococcus radiodurans R1 GCA_000008565.1
305 Deinococcus gobiensis I-0 GCA_000252445.1
287 Deinococcus proteolyticus MRP GCA_000190555.1
xE
505 _5BBacillus thuringiensis_5D serovar konkukian str. 97-27 GCA_000008505.1
491 Bacillus anthracis str. Ames GCA_000007845.1
491 Bacillus anthracis str. Sterne GCA_000008165.1
491 Bacillus cereus ATCC 14579 GCA_000007825.1
491 Bacillus thuringiensis YBT-1518 GCA_000497525.2
476 Bacillus mycoides GCA_000832605.1
xF
336 Deinococcus radiodurans R1 GCA_000008565.1
239 Deinococcus gobiensis I-0 GCA_000252445.1
215 Deinococcus soli Cha et al. 2016 GCA_001007995.1
xG
738 Streptococcus mutans UA159 GCA_000007465.2
576 Streptococcus gordonii str. Challis substr. CH1 GCA_000017005.1
557 Streptococcus ratti FA-1 = DSM 20564 GCA_000286075.1
xH
652 Rhodobacter sphaeroides 2.4.1 GCA_000012905.2
584 Rhodobacter sphaeroides ATCC 17025 GCA_000016405.1
465 Gemmobacter megaterium GCA_900156815.1
xI
274 Deinococcus radiodurans R1 GCA_000008565.1
145 Deinococcus deserti VCD115 GCA_000020685.1
138 Deinococcus hopiensis KR-140 GCA_900176165.1
xJ
711 Staphylococcus epidermidis ATCC 12228 GCA_000007645.1
680 Staphylococcus capitis subsp. capitis GCA_001028645.1
642 Staphylococcus warneri SG1 GCA_000332735.1
xK
516 Deinococcus radiodurans R1 GCA_000008565.1
326 Deinococcus deserti VCD115 GCA_000020685.1
301 Deinococcus soli Cha et al. 2016 GCA_001007995.1
xL
753 Deinococcus radiodurans R1 GCA_000008565.1
508 Deinococcus deserti VCD115 GCA_000020685.1
482 Deinococcus soli Cha et al. 2016 GCA_001007995.1
xM
613 Clostridium beijerinckii GCA_000833105.2
600 Clostridium saccharoperbutylacetonicum N1-4_28HMT_29 GCA_000340885.1
584 Clostridium saccharobutylicum DSM 13864 GCA_000473995.1
xN
773 Staphylococcus epidermidis ATCC 12228 GCA_000007645.1
753 Staphylococcus capitis subsp. capitis GCA_001028645.1
717 Staphylococcus aureus subsp. aureus NCTC 8325 GCA_000013425.1
xO
430 Clostridium beijerinckii GCA_000833105.2
391 Clostridium saccharoperbutylacetonicum N1-4_28HMT_29 GCA_000340885.1
386 Clostridium neonatale GCA_001458595.1
xP
695 Lactobacillus gasseri ATCC 33323 = JCM 1131 GCA_000014425.1
621 Lactobacillus hominis DSM 23910 = CRBIP 24.179 GCA_000296835.1
486 Lactobacillus jensenii GCA_001936235.1
486 Lactobacillus psittaci DSM 15354 GCA_000425905.1
xQ
627 Clostridium beijerinckii GCA_000833105.2
579 Clostridium puniceum GCA_002006345.1
579 Clostridium saccharoperbutylacetonicum N1-4_28HMT_29 GCA_000340885.1
576 Clostridium butyricum GCA_001456065.2
xR
678 Clostridium beijerinckii GCA_000833105.2
671 Clostridium saccharoperbutylacetonicum N1-4_28HMT_29 GCA_000340885.1
655 Clostridium saccharobutylicum DSM 13864 GCA_000473995.1
xS
938 Bifidobacterium adolescentis ATCC 15703 GCA_000010425.1
800 Bifidobacterium angulatum DSM 20098 = JCM 7096 GCA_001025155.1
782 Bifidobacterium dentium JCM 1195 = DSM 20436 GCA_001042595.1
xT
800 Staphylococcus epidermidis ATCC 12228 GCA_000007645.1
737 Staphylococcus capitis subsp. capitis GCA_001028645.1
723 Staphylococcus warneri SG1 GCA_000332735.1

xU
837 Bacillus anthracis str. Ames GCA_000007845.1
837 Bacillus anthracis str. Sterne GCA_000008165.1
837 _5BBacillus thuringiensis_5D serovar konkukian str. 97-27 GCA_000008505.1
812 Bacillus cereus ATCC 14579 GCA_000007825.1
797 Bacillus pseudomycoides DSM 12442 GCA_000161455.1
xV
730 Staphylococcus epidermidis ATCC 12228 GCA_000007645.1
682 Staphylococcus capitis subsp. capitis GCA_001028645.1
634 Staphylococcus hominis subsp. hominis C80 GCA_000183685.1
xW
533 Clostridium beijerinckii GCA_000833105.2
508 Clostridium puniceum GCA_002006345.1
508 Clostridium saccharoperbutylacetonicum N1-4_28HMT_29 GCA_000340885.1
495 Clostridium butyricum GCA_001456065.2
xX
690 Clostridium beijerinckii GCA_000833105.2
644 Clostridium saccharoperbutylacetonicum N1-4_28HMT_29 GCA_000340885.1
632 Clostridium butyricum GCA_001456065.2
xY
703 Rhodobacter sphaeroides 2.4.1 GCA_000012905.2
643 Rhodobacter sphaeroides ATCC 17025 GCA_000016405.1
571 Gemmobacter aquatilis GCA_900110025.1
xZ
737 Clostridium beijerinckii GCA_000833105.2
729 Clostridium saccharobutylicum DSM 13864 GCA_000473995.1
726 Clostridium saccharoperbutylacetonicum N1-4_28HMT_29 GCA_000340885.1
y0
400 Rhodobacter sphaeroides 2.4.1 GCA_000012905.2
346 Rhodobacter sphaeroides ATCC 17025 GCA_000016405.1
337 Defluviimonas alba GCA_001620265.1
y1
460 Enterococcus faecalis V583 GCA_000007785.1
413 Streptomyces cinnamoneus GCA_001885705.1
392 Enterococcus haemoperoxidus ATCC BAA-382 GCA_000407165.1
y2
805 Clostridium beijerinckii GCA_000833105.2
799 Clostridium saccharoperbutylacetonicum N1-4_28HMT_29 GCA_000340885.1
743 Clostridium puniceum GCA_002006345.1
y3
679 Staphylococcus epidermidis ATCC 12228 GCA_000007645.1
640 Staphylococcus capitis subsp. capitis GCA_001028645.1
627 Megasphaera cerevisiae DSM 20462 GCA_001045675.1
627 Staphylococcus warneri SG1 GCA_000332735.1
y4
895 Lactobacillus gasseri ATCC 33323 = JCM 1131 GCA_000014425.1
818 Lactobacillus hominis DSM 23910 = CRBIP 24.179 GCA_000296835.1
558 Lactobacillus iners DSM 13335 GCA_000160875.1
y5
234 Staphylococcus epidermidis ATCC 12228 GCA_000007645.1
226 Staphylococcus capitis subsp. capitis GCA_001028645.1
214 Megasphaera cerevisiae DSM 20462 GCA_001045675.1
214 Staphylococcus aureus subsp. aureus NCTC 8325 GCA_000013425.1
214 Staphylococcus warneri SG1 GCA_000332735.1
y6
574 Clostridium beijerinckii GCA_000833105.2
527 Clostridium puniceum GCA_002006345.1
525 Clostridium saccharoperbutylacetonicum N1-4_28HMT_29 GCA_000340885.1
y7
564 Rhodobacter sphaeroides 2.4.1 GCA_000012905.2
518 Rhodobacter sphaeroides ATCC 17025 GCA_000016405.1
445 Defluviimonas alba GCA_001620265.1
y8
427 Rhodobacter sphaeroides 2.4.1 GCA_000012905.2
389 Rhodobacter sphaeroides ATCC 17025 GCA_000016405.1
335 Gemmobacter aquatilis GCA_900110025.1
y9
672 Lactobacillus gasseri ATCC 33323 = JCM 1131 GCA_000014425.1
651 Lactobacillus hominis DSM 23910 = CRBIP 24.179 GCA_000296835.1
497 Lactobacillus iners DSM 13335 GCA_000160875.1
ya
747 Enterococcus faecalis V583 GCA_000007785.1
701 Streptomyces cinnamoneus GCA_001885705.1
627 Enterococcus hirae ATCC 9790 GCA_000271405.2
yb
670 Rhodobacter sphaeroides 2.4.1 GCA_000012905.2

609 Rhodobacter sphaeroides ATCC 17025 GCA_000016405.1
487 Pseudorhodobacter psychrotolerans GCA_001294535.1
yc
411 Clostridium beijerinckii GCA_000833105.2
402 Clostridium saccharoperbutylacetonicum N1-4_28HMT_29 GCA_000340885.1
388 Clostridium puniceum GCA_002006345.1
yd
642 Clostridium beijerinckii GCA_000833105.2
619 Clostridium saccharoperbutylacetonicum N1-4_28HMT_29 GCA_000340885.1
600 Clostridium puniceum GCA_002006345.1
ye
654 Lactobacillus gasseri ATCC 33323 = JCM 1131 GCA_000014425.1
601 Lactobacillus hominis DSM 23910 = CRBIP 24.179 GCA_000296835.1
411 Lactobacillus psittaci DSM 15354 GCA_000425905.1
yf
923 Staphylococcus epidermidis ATCC 12228 GCA_000007645.1
864 Staphylococcus capitis subsp. capitis GCA_001028645.1
821 Staphylococcus hominis subsp. hominis C80 GCA_000183685.1
yg
784 Enterococcus faecalis V583 GCA_000007785.1
737 Streptomyces cinnamoneus GCA_001885705.1
656 Enterococcus haemoperoxidus ATCC BAA-382 GCA_000407165.1
656 Enterococcus rivorum GCA_001742285.1
yh
755 Streptococcus mutans UA159 GCA_000007465.2
593 Streptococcus gordonii str. Challis substr. CH1 GCA_000017005.1
575 Streptococcus ratti FA-1 = DSM 20564 GCA_000286075.1
yi
752 Clostridium beijerinckii GCA_000833105.2
748 Clostridium saccharoperbutylacetonicum N1-4_28HMT_29 GCA_000340885.1
727 Clostridium puniceum GCA_002006345.1
727 Clostridium saccharobutylicum DSM 13864 GCA_000473995.1
yj
465 Deinococcus radiodurans R1 GCA_000008565.1
280 Deinococcus marmoris DSM 12784 GCA_000701405.1
271 Deinococcus deserti VCD115 GCA_000020685.1
271 Deinococcus hopiensis KR-140 GCA_900176165.1
yk
788 Lactobacillus gasseri ATCC 33323 = JCM 1131 GCA_000014425.1
721 Lactobacillus hominis DSM 23910 = CRBIP 24.179 GCA_000296835.1
569 Lactobacillus iners DSM 13335 GCA_000160875.1
yl
651 Lactobacillus gasseri ATCC 33323 = JCM 1131 GCA_000014425.1
562 Lactobacillus hominis DSM 23910 = CRBIP 24.179 GCA_000296835.1
437 Lactobacillus iners DSM 13335 GCA_000160875.1
ym
839 Streptococcus mutans UA159 GCA_000007465.2
557 Streptococcus halotolerans GCA_001598035.1
542 Streptococcus gordonii str. Challis substr. CH1 GCA_000017005.1
yn
466 Escherichia coli O157_3AH7 str. Sakai GCA_000008865.1
466 Escherichia coli UMN026 GCA_000026325.2
466 Escherichia coli str. K-12 substr. MG1655 GCA_000005845.2
442 Shigella flexneri 2a str. 301 GCA_000006925.2
439 Escherichia coli O83_3AH1 str. NRG 857C GCA_000183345.1
yo
512 Bifidobacterium adolescentis ATCC 15703 GCA_000010425.1
421 Bifidobacterium angulatum DSM 20098 = JCM 7096 GCA_001025155.1
411 Bifidobacterium dentium JCM 1195 = DSM 20436 GCA_001042595.1
yp
869 Clostridium beijerinckii GCA_000833105.2
817 Clostridium saccharoperbutylacetonicum N1-4_28HMT_29 GCA_000340885.1
780 Clostridium saccharobutylicum DSM 13864 GCA_000473995.1
yq
714 Staphylococcus epidermidis ATCC 12228 GCA_000007645.1
662 Staphylococcus capitis subsp. capitis GCA_001028645.1
630 Staphylococcus hominis subsp. hominis C80 GCA_000183685.1
yr
723 Clostridium beijerinckii GCA_000833105.2
698 Clostridium saccharoperbutylacetonicum N1-4_28HMT_29 GCA_000340885.1
692 Clostridium puniceum GCA_002006345.1
ys
730 Rhodobacter sphaeroides 2.4.1 GCA_000012905.2
685 Rhodobacter sphaeroides ATCC 17025 GCA_000016405.1
558 Pseudorhodobacter psychrotolerans GCA_001294535.1
yt

637 Staphylococcus epidermidis ATCC 12228 GCA_000007645.1
610 Staphylococcus capitis subsp. capitis GCA_001028645.1
587 Staphylococcus aureus subsp. aureus NCTC 8325 GCA_000013425.1
yu
640 Streptococcus mutans UA159 GCA_000007465.2
447 Streptococcus ratti FA-1 = DSM 20564 GCA_000286075.1
423 Streptococcus ovis DSM 16829 GCA_000380125.1
yv
542 Deinococcus radiodurans R1 GCA_000008565.1
308 Deinococcus deserti VCD115 GCA_000020685.1
303 Deinococcus soli Cha et al. 2016 GCA_001007995.1
yw
625 Clostridium beijerinckii GCA_000833105.2
578 Clostridium saccharoperbutylacetonicum N1-4_28HMT_29 GCA_000340885.1
571 Clostridium puniceum GCA_002006345.1
yx
574 Shigella flexneri 2a str. 301 GCA_000006925.2
573 Escherichia coli O104_3AH4 str. 2011C-3493 GCA_000299455.1
573 Escherichia coli str. K-12 substr. MG1655 GCA_000005845.2
548 Escherichia coli UMN026 GCA_000026325.2
yy
724 Bacillus anthracis str. Ames GCA_000007845.1
724 Bacillus anthracis str. Sterne GCA_000008165.1
724 _5BBacillus thuringiensis_5D serovar konkukian str. 97-27 GCA_000008505.1
715 Bacillus cereus ATCC 14579 GCA_000007825.1
703 Bacillus pseudomycoides DSM 12442 GCA_000161455.1
yz
742 Staphylococcus epidermidis ATCC 12228 GCA_000007645.1
689 Staphylococcus capitis subsp. capitis GCA_001028645.1
625 Staphylococcus hominis subsp. hominis C80 GCA_000183685.1
yA
533 Staphylococcus epidermidis ATCC 12228 GCA_000007645.1
516 Staphylococcus haemolyticus JCSC1435 GCA_000009865.1
508 Staphylococcus capitis subsp. capitis GCA_001028645.1
yB
495 Rhodobacter sphaeroides 2.4.1 GCA_000012905.2
431 Rhodobacter sphaeroides ATCC 17025 GCA_000016405.1
389 Gemmobacter megaterium GCA_900156815.1
yC
717 Clostridium beijerinckii GCA_000833105.2
701 Clostridium saccharoperbutylacetonicum N1-4_28HMT_29 GCA_000340885.1
692 Clostridium saccharobutylicum DSM 13864 GCA_000473995.1
yD
608 Streptococcus mutans UA159 GCA_000007465.2
428 Streptococcus ratti FA-1 = DSM 20564 GCA_000286075.1
399 Streptococcus equinus GCA_000964315.1
399 Streptococcus gallolyticus subsp. gallolyticus DSM 16831 GCA_002000985.1
yE
905 Staphylococcus epidermidis ATCC 12228 GCA_000007645.1
861 Staphylococcus capitis subsp. capitis GCA_001028645.1
778 Staphylococcus warneri SG1 GCA_000332735.1
yF
742 Deinococcus radiodurans R1 GCA_000008565.1
489 Deinococcus deserti VCD115 GCA_000020685.1
487 Deinococcus gobiensis I-0 GCA_000252445.1
yG
652 Streptococcus mutans UA159 GCA_000007465.2
492 Streptococcus halotolerans GCA_001598035.1
487 Streptococcus ratti FA-1 = DSM 20564 GCA_000286075.1
yH
592 Streptococcus mutans UA159 GCA_000007465.2
471 Streptococcus ratti FA-1 = DSM 20564 GCA_000286075.1
453 Streptococcus sobrinus DSM 20742 = ATCC 33478 GCA_000686605.1
yI
414 Escherichia coli str. K-12 substr. MG1655 GCA_000005845.2
399 Escherichia coli IAI39 GCA_000026345.1
391 Escherichia coli UMN026 GCA_000026325.2
yJ
671 Clostridium beijerinckii GCA_000833105.2
668 Clostridium saccharoperbutylacetonicum N1-4_28HMT_29 GCA_000340885.1
632 Clostridium puniceum GCA_002006345.1
yK
554 Clostridium beijerinckii GCA_000833105.2
497 Clostridium saccharoperbutylacetonicum N1-4_28HMT_29 GCA_000340885.1
470 Clostridium saccharobutylicum DSM 13864 GCA_000473995.1
yL

842 Clostridium beijerinckii GCA_000833105.2
801 Clostridium saccharoperbutylacetonicum N1-4_28HMT_29 GCA_000340885.1
760 Clostridium saccharobutylicum DSM 13864 GCA_000473995.1
yM
573 Clostridium beijerinckii GCA_000833105.2
559 Clostridium saccharoperbutylacetonicum N1-4_28HMT_29 GCA_000340885.1
555 Clostridium saccharobutylicum DSM 13864 GCA_000473995.1
yN
468 Clostridium beijerinckii GCA_000833105.2
444 Clostridium saccharoperbutylacetonicum N1-4_28HMT_29 GCA_000340885.1
435 Clostridium butyricum GCA_001456065.2
yO
550 Escherichia coli O104_3AH4 str. 2011C-3493 GCA_000299455.1
535 Escherichia coli str. K-12 substr. MG1655 GCA_000005845.2
535 Shigella flexneri 2a str. 301 GCA_000006925.2
520 Escherichia coli O157_3AH7 str. Sakai GCA_000008865.1
520 Escherichia coli O83_3AH1 str. NRG 857C GCA_000183345.1
520 Escherichia coli UMN026 GCA_000026325.2
yP
404 Deinococcus radiodurans R1 GCA_000008565.1
252 Deinococcus gobiensis I-0 GCA_000252445.1
251 Deinococcus proteolyticus MRP GCA_000190555.1
yQ
657 Streptococcus mutans UA159 GCA_000007465.2
454 Streptococcus gordonii str. Challis substr. CH1 GCA_000017005.1
450 Streptococcus ratti FA-1 = DSM 20564 GCA_000286075.1
yR
733 Deinococcus radiodurans R1 GCA_000008565.1
488 Deinococcus deserti VCD115 GCA_000020685.1
479 Deinococcus puniceus GCA_001644565.1
yS
819 Bacillus cereus ATCC 14579 GCA_000007825.1
819 _5BBacillus thuringiensis_5D serovar konkukian str. 97-27 GCA_000008505.1
818 Bacillus anthracis str. Ames GCA_000007845.1
818 Bacillus anthracis str. Sterne GCA_000008165.1
803 Bacillus thuringiensis YBT-1518 GCA_000497525.2
yT
667 Streptococcus mutans UA159 GCA_000007465.2
472 Streptococcus ratti FA-1 = DSM 20564 GCA_000286075.1
426 Streptococcus ferus DSM 20646 GCA_000372425.1
yU
447 Staphylococcus epidermidis ATCC 12228 GCA_000007645.1
420 Staphylococcus capitis subsp. capitis GCA_001028645.1
419 Staphylococcus haemolyticus JCSC1435 GCA_000009865.1
yV
594 Bifidobacterium adolescentis ATCC 15703 GCA_000010425.1
503 Bifidobacterium dentium JCM 1195 = DSM 20436 GCA_001042595.1
408 Bifidobacterium stellenboschense GCA_000741785.1
yW
765 Lactobacillus gasseri ATCC 33323 = JCM 1131 GCA_000014425.1
707 Lactobacillus hominis DSM 23910 = CRBIP 24.179 GCA_000296835.1
527 Lactobacillus iners DSM 13335 GCA_000160875.1
yX
697 Lactobacillus gasseri ATCC 33323 = JCM 1131 GCA_000014425.1
625 Lactobacillus hominis DSM 23910 = CRBIP 24.179 GCA_000296835.1
474 Lactobacillus iners DSM 13335 GCA_000160875.1
yY
562 Deinococcus radiodurans R1 GCA_000008565.1
342 Deinococcus deserti VCD115 GCA_000020685.1
335 Deinococcus soli Cha et al. 2016 GCA_001007995.1
yZ
795 Rhodobacter sphaeroides 2.4.1 GCA_000012905.2
739 Rhodobacter sphaeroides ATCC 17025 GCA_000016405.1
547 Gemmobacter aquatilis GCA_900110025.1
z0
584 Deinococcus radiodurans R1 GCA_000008565.1
395 Deinococcus proteolyticus MRP GCA_000190555.1
386 Deinococcus deserti VCD115 GCA_000020685.1
z1
556 Rhodobacter sphaeroides 2.4.1 GCA_000012905.2
490 Rhodobacter sphaeroides ATCC 17025 GCA_000016405.1
443 Rhodobacter vinaykumarii GCA_900156695.1
z2
607 Staphylococcus epidermidis ATCC 12228 GCA_000007645.1
601 Staphylococcus capitis subsp. capitis GCA_001028645.1
519 Staphylococcus warneri SG1 GCA_000332735.1

z3
730 Enterococcus faecalis V583 GCA_000007785.1
683 Streptomyces cinnamoneus GCA_001885705.1
632 Enterococcus massiliensis GCA_001050095.1
632 Enterococcus rivorum GCA_001742285.1
z4
750 Staphylococcus epidermidis ATCC 12228 GCA_000007645.1
724 Staphylococcus capitis subsp. capitis GCA_001028645.1
712 Staphylococcus haemolyticus JCSC1435 GCA_000009865.1
z5
808 Deinococcus radiodurans R1 GCA_000008565.1
478 Deinococcus deserti VCD115 GCA_000020685.1
465 Deinococcus gobiensis I-0 GCA_000252445.1
z6
866 Streptococcus mutans UA159 GCA_000007465.2
590 Streptococcus ratti FA-1 = DSM 20564 GCA_000286075.1
572 Streptococcus gordonii str. Challis substr. CH1 GCA_000017005.1
z7
553 Clostridium beijerinckii GCA_000833105.2
524 Clostridium saccharoperbutylacetonicum N1-4_28HMT_29 GCA_000340885.1
521 Clostridium saccharobutylicum DSM 13864 GCA_000473995.1
z8
915 Clostridium beijerinckii GCA_000833105.2
898 Clostridium saccharoperbutylacetonicum N1-4_28HMT_29 GCA_000340885.1
878 Clostridium puniceum GCA_002006345.1
z9
674 Staphylococcus epidermidis ATCC 12228 GCA_000007645.1
668 Staphylococcus capitis subsp. capitis GCA_001028645.1
612 Staphylococcus hominis subsp. hominis C80 GCA_000183685.1
za
635 Streptococcus mutans UA159 GCA_000007465.2
444 Streptococcus ratti FA-1 = DSM 20564 GCA_000286075.1
422 Streptococcus halotolerans GCA_001598035.1
zb
625 Rhodobacter sphaeroides 2.4.1 GCA_000012905.2
547 Rhodobacter sphaeroides ATCC 17025 GCA_000016405.1
490 Pseudorhodobacter psychrotolerans GCA_001294535.1
zc
545 Enterococcus faecalis V583 GCA_000007785.1
504 Streptomyces cinnamoneus GCA_001885705.1
478 Enterococcus rivorum GCA_001742285.1
zd
488 Lactobacillus gasseri ATCC 33323 = JCM 1131 GCA_000014425.1
477 Lactobacillus hominis DSM 23910 = CRBIP 24.179 GCA_000296835.1
317 Lactobacillus hamsteri DSM 5661 = JCM 6256 GCA_000615445.1
ze
596 Clostridium beijerinckii GCA_000833105.2
586 Clostridium saccharobutylicum DSM 13864 GCA_000473995.1
571 Clostridium saccharoperbutylacetonicum N1-4_28HMT_29 GCA_000340885.1
zf
515 Rhodobacter sphaeroides 2.4.1 GCA_000012905.2
465 Rhodobacter sphaeroides ATCC 17025 GCA_000016405.1
418 Gemmobacter aquatilis GCA_900110025.1
zg
570 Streptococcus mutans UA159 GCA_000007465.2
421 Streptococcus ratti FA-1 = DSM 20564 GCA_000286075.1
367 Streptococcus gallolyticus subsp. gallolyticus DSM 16831 GCA_002000985.1
367 Streptococcus henryi DSM 19005 GCA_000376985.1
zh
725 Rhodobacter sphaeroides 2.4.1 GCA_000012905.2
664 Rhodobacter sphaeroides ATCC 17025 GCA_000016405.1
593 Gemmobacter aquatilis GCA_900110025.1
zi
782 Clostridium beijerinckii GCA_000833105.2
772 Clostridium saccharoperbutylacetonicum N1-4_28HMT_29 GCA_000340885.1
760 Clostridium puniceum GCA_002006345.1
zj
527 Rhodobacter sphaeroides 2.4.1 GCA_000012905.2
451 Rhodobacter sphaeroides ATCC 17025 GCA_000016405.1
397 Rhodobacter capsulatus SB 1003 GCA_000021865.1
zk
815 Clostridium beijerinckii GCA_000833105.2
776 Clostridium saccharoperbutylacetonicum N1-4_28HMT_29 GCA_000340885.1
766 Clostridium puniceum GCA_002006345.1
zl
724 Deinococcus radiodurans R1 GCA_000008565.1

428 Deinococcus deserti VCD115 GCA_000020685.1
417 Deinococcus soli Cha et al. 2016 GCA_001007995.1
zm
791 Streptococcus mutans UA159 GCA_000007465.2
581 Streptococcus ratti FA-1 = DSM 20564 GCA_000286075.1
524 Streptococcus gordonii str. Challis substr. CH1 GCA_000017005.1
zn
776 Escherichia coli O157_3AH7 str. Sakai GCA_000008865.1
776 Escherichia coli UMN026 GCA_000026325.2
776 Escherichia coli str. K-12 substr. MG1655 GCA_000005845.2
748 Shigella flexneri 2a str. 301 GCA_000006925.2
743 Escherichia coli O83_3AH1 str. NRG 857C GCA_000183345.1
zo
586 Clostridium beijerinckii GCA_000833105.2
557 Clostridium saccharoperbutylacetonicum N1-4_28HMT_29 GCA_000340885.1
523 Clostridium puniceum GCA_002006345.1
zp
935 Rhodobacter sphaeroides 2.4.1 GCA_000012905.2
822 Rhodobacter sphaeroides ATCC 17025 GCA_000016405.1
717 Gemmobacter megaterium GCA_900156815.1
zq
748 Bifidobacterium adolescentis ATCC 15703 GCA_000010425.1
618 Bifidobacterium dentium JCM 1195 = DSM 20436 GCA_001042595.1
598 Bifidobacterium angulatum DSM 20098 = JCM 7096 GCA_001025155.1
zr
628 Enterococcus faecalis V583 GCA_000007785.1
581 Streptomyces cinnamoneus GCA_001885705.1
576 Enterococcus thailandicus GCA_001652875.1
zs
632 Rhodobacter sphaeroides 2.4.1 GCA_000012905.2
542 Rhodobacter sphaeroides ATCC 17025 GCA_000016405.1
400 Gemmobacter megaterium GCA_900156815.1
zt
825 Lactobacillus gasseri ATCC 33323 = JCM 1131 GCA_000014425.1
784 Lactobacillus hominis DSM 23910 = CRBIP 24.179 GCA_000296835.1
521 Lactobacillus helveticus GCA_001308285.1
zu
548 Staphylococcus epidermidis ATCC 12228 GCA_000007645.1
507 Staphylococcus capitis subsp. capitis GCA_001028645.1
498 Staphylococcus aureus subsp. aureus NCTC 8325 GCA_000013425.1
zv
849 Enterococcus faecalis V583 GCA_000007785.1
806 Streptomyces cinnamoneus GCA_001885705.1
718 Enterococcus rivorum GCA_001742285.1
zw
884 Clostridium beijerinckii GCA_000833105.2
836 Clostridium saccharoperbutylacetonicum N1-4_28HMT_29 GCA_000340885.1
822 Clostridium puniceum GCA_002006345.1
zx
647 Staphylococcus epidermidis ATCC 12228 GCA_000007645.1
591 Staphylococcus capitis subsp. capitis GCA_001028645.1
542 Staphylococcus lentus F1142 GCA_000286395.1
zy
717 Lactobacillus gasseri ATCC 33323 = JCM 1131 GCA_000014425.1
708 Lactobacillus hominis DSM 23910 = CRBIP 24.179 GCA_000296835.1
530 Lactobacillus psittaci DSM 15354 GCA_000425905.1
zz
632 Clostridium beijerinckii GCA_000833105.2
596 Clostridium saccharoperbutylacetonicum N1-4_28HMT_29 GCA_000340885.1
580 Clostridium saccharobutylicum DSM 13864 GCA_000473995.1
zA
544 Staphylococcus epidermidis ATCC 12228 GCA_000007645.1
529 Staphylococcus capitis subsp. capitis GCA_001028645.1
504 Staphylococcus pettenkoferi GCA_002208805.1
zB
977 Staphylococcus epidermidis ATCC 12228 GCA_000007645.1
936 Staphylococcus capitis subsp. capitis GCA_001028645.1
844 Staphylococcus warneri SG1 GCA_000332735.1
zC
777 Clostridium beijerinckii GCA_000833105.2
744 Clostridium saccharoperbutylacetonicum N1-4_28HMT_29 GCA_000340885.1
719 Clostridium puniceum GCA_002006345.1
zD
483 Rhodobacter sphaeroides 2.4.1 GCA_000012905.2
446 Rhodobacter sphaeroides ATCC 17025 GCA_000016405.1
335 Pseudorhodobacter ferrugineus DSM 5888 GCA_000420745.1

zE
534 Clostridium beijerinckii GCA_000833105.2
514 Clostridium saccharobutylicum DSM 13864 GCA_000473995.1
493 Clostridium saccharoperbutylacetonicum N1-4_28HMT_29 GCA_000340885.1
zF
928 Lactobacillus gasseri ATCC 33323 = JCM 1131 GCA_000014425.1
871 Lactobacillus hominis DSM 23910 = CRBIP 24.179 GCA_000296835.1
640 Lactobacillus iners DSM 13335 GCA_000160875.1
zG
709 Deinococcus radiodurans R1 GCA_000008565.1
483 Deinococcus deserti VCD115 GCA_000020685.1
436 Deinococcus gobiensis I-0 GCA_000252445.1
zH
357 Enterococcus faecalis V583 GCA_000007785.1
338 Streptomyces cinnamoneus GCA_001885705.1
335 Enterococcus rivorum GCA_001742285.1
zI
677 Rhodobacter sphaeroides 2.4.1 GCA_000012905.2
633 Rhodobacter sphaeroides ATCC 17025 GCA_000016405.1
513 Rhodobacter capsulatus SB 1003 GCA_000021865.1
zJ
711 Rhodobacter sphaeroides 2.4.1 GCA_000012905.2
630 Rhodobacter sphaeroides ATCC 17025 GCA_000016405.1
575 Gemmobacter aquatilis GCA_900110025.1
zK
828 Clostridium beijerinckii GCA_000833105.2
780 Clostridium saccharoperbutylacetonicum N1-4_28HMT_29 GCA_000340885.1
747 Clostridium saccharobutylicum DSM 13864 GCA_000473995.1
zL
737 Lactobacillus gasseri ATCC 33323 = JCM 1131 GCA_000014425.1
683 Lactobacillus hominis DSM 23910 = CRBIP 24.179 GCA_000296835.1
531 Lactobacillus iners DSM 13335 GCA_000160875.1
zM
884 Streptococcus mutans UA159 GCA_000007465.2
668 Streptococcus ratti FA-1 = DSM 20564 GCA_000286075.1
628 Streptococcus gordonii str. Challis substr. CH1 GCA_000017005.1
zN
579 Rhodobacter sphaeroides 2.4.1 GCA_000012905.2
517 Rhodobacter sphaeroides ATCC 17025 GCA_000016405.1
460 Gemmobacter aquatilis GCA_900110025.1
zO
826 Rhodobacter sphaeroides 2.4.1 GCA_000012905.2
720 Rhodobacter sphaeroides ATCC 17025 GCA_000016405.1
618 Gemmobacter aquatilis GCA_900110025.1
zP
642 Clostridium beijerinckii GCA_000833105.2
602 Clostridium saccharobutylicum DSM 13864 GCA_000473995.1
602 Clostridium saccharoperbutylacetonicum N1-4_28HMT_29 GCA_000340885.1
575 Clostridium butyricum GCA_001456065.2
zQ
847 Rhodobacter sphaeroides 2.4.1 GCA_000012905.2
754 Rhodobacter sphaeroides ATCC 17025 GCA_000016405.1
662 Gemmobacter aquatilis GCA_900110025.1
zR
336 Staphylococcus epidermidis ATCC 12228 GCA_000007645.1
313 Staphylococcus warneri SG1 GCA_000332735.1
303 Staphylococcus capitis subsp. capitis GCA_001028645.1
zS
726 Clostridium beijerinckii GCA_000833105.2
677 Clostridium saccharoperbutylacetonicum N1-4_28HMT_29 GCA_000340885.1
676 Clostridium saccharobutylicum DSM 13864 GCA_000473995.1
zT
637 Clostridium beijerinckii GCA_000833105.2
604 Clostridium saccharoperbutylacetonicum N1-4_28HMT_29 GCA_000340885.1
573 Clostridium puniceum GCA_002006345.1
zU
789 Clostridium beijerinckii GCA_000833105.2
752 Clostridium saccharoperbutylacetonicum N1-4_28HMT_29 GCA_000340885.1
733 Clostridium saccharobutylicum DSM 13864 GCA_000473995.1
zV
717 Staphylococcus epidermidis ATCC 12228 GCA_000007645.1
699 Staphylococcus capitis subsp. capitis GCA_001028645.1
622 Staphylococcus warneri SG1 GCA_000332735.1
zW
700 Streptococcus mutans UA159 GCA_000007465.2
516 Streptococcus ratti FA-1 = DSM 20564 GCA_000286075.1

487 Streptococcus iniae GCA_000831485.1
zX
887 Clostridium beijerinckii GCA_000833105.2
831 Clostridium saccharoperbutylacetonicum N1-4_28HMT_29 GCA_000340885.1
797 Clostridium puniceum GCA_002006345.1
zY
485 Streptococcus mutans UA159 GCA_000007465.2
409 Streptococcus ratti FA-1 = DSM 20564 GCA_000286075.1
377 Streptococcus halotolerans GCA_001598035.1
377 Streptococcus sobrinus DSM 20742 = ATCC 33478 GCA_000686605.1
zZ
588 Rhodobacter sphaeroides 2.4.1 GCA_000012905.2
554 Rhodobacter sphaeroides ATCC 17025 GCA_000016405.1
440 Gemmobacter aquatilis GCA_900110025.1
440 Pseudorhodobacter psychrotolerans GCA_001294535.1
A0
801 Deinococcus radiodurans R1 GCA_000008565.1
573 Deinococcus deserti VCD115 GCA_000020685.1
529 Deinococcus gobiensis I-0 GCA_000252445.1
A1
783 Lactobacillus gasseri ATCC 33323 = JCM 1131 GCA_000014425.1
719 Lactobacillus hominis DSM 23910 = CRBIP 24.179 GCA_000296835.1
609 Lactobacillus psittaci DSM 15354 GCA_000425905.1
A2
752 Streptococcus mutans UA159 GCA_000007465.2
522 Streptococcus gordonii str. Challis substr. CH1 GCA_000017005.1
518 Streptococcus sanguinis SK36 GCA_000014205.1
A3
803 Enterococcus faecalis V583 GCA_000007785.1
756 Streptomyces cinnamoneus GCA_001885705.1
719 Enterococcus rivorum GCA_001742285.1
A4
540 Rhodobacter sphaeroides 2.4.1 GCA_000012905.2
503 Rhodobacter sphaeroides ATCC 17025 GCA_000016405.1
445 Pseudorhodobacter wandonensis GCA_001202035.1
A5
842 Bacillus anthracis str. Ames GCA_000007845.1
842 Bacillus anthracis str. Sterne GCA_000008165.1
842 Bacillus cereus ATCC 14579 GCA_000007825.1
842 _5BBacillus thuringiensis_5D serovar konkukian str. 97-27 GCA_000008505.1
817 Bacillus thuringiensis YBT-1518 GCA_000497525.2
802 Bacillus pseudomycoides DSM 12442 GCA_000161455.1
A6
675 Rhodobacter sphaeroides 2.4.1 GCA_000012905.2
624 Rhodobacter sphaeroides ATCC 17025 GCA_000016405.1
513 Gemmobacter aquatilis GCA_900110025.1
A7
809 Enterococcus faecalis V583 GCA_000007785.1
762 Streptomyces cinnamoneus GCA_001885705.1
716 Enterococcus rivorum GCA_001742285.1
A8
754 Staphylococcus epidermidis ATCC 12228 GCA_000007645.1
681 Staphylococcus capitis subsp. capitis GCA_001028645.1
630 Staphylococcus lugdunensis HKU09-01 GCA_000025085.1
A9
774 Staphylococcus epidermidis ATCC 12228 GCA_000007645.1
715 Staphylococcus capitis subsp. capitis GCA_001028645.1
672 Staphylococcus hominis subsp. hominis C80 GCA_000183685.1
Aa
764 Streptococcus mutans UA159 GCA_000007465.2
584 Streptococcus ratti FA-1 = DSM 20564 GCA_000286075.1
567 Streptococcus gordonii str. Challis substr. CH1 GCA_000017005.1
Ab
910 Lactobacillus gasseri ATCC 33323 = JCM 1131 GCA_000014425.1
810 Lactobacillus hominis DSM 23910 = CRBIP 24.179 GCA_000296835.1
635 Lactobacillus iners DSM 13335 GCA_000160875.1
Ac
753 Lactobacillus gasseri ATCC 33323 = JCM 1131 GCA_000014425.1
687 Lactobacillus hominis DSM 23910 = CRBIP 24.179 GCA_000296835.1
585 Lactobacillus jensenii GCA_001936235.1
Ad
419 Staphylococcus epidermidis ATCC 12228 GCA_000007645.1
405 Staphylococcus capitis subsp. capitis GCA_001028645.1
401 Staphylococcus aureus subsp. aureus NCTC 8325 GCA_000013425.1
Ae
890 Bacillus anthracis str. Ames GCA_000007845.1

890 Bacillus anthracis str. Sterne GCA_000008165.1
890 _5BBacillus thuringiensis_5D serovar konkukian str. 97-27 GCA_000008505.1
879 Bacillus pseudomycoides DSM 12442 GCA_000161455.1
874 Bacillus cereus ATCC 14579 GCA_000007825.1
Af
740 Enterococcus faecalis V583 GCA_000007785.1
693 Streptomyces cinnamoneus GCA_001885705.1
643 Enterococcus rivorum GCA_001742285.1
Ag
683 Lactobacillus gasseri ATCC 33323 = JCM 1131 GCA_000014425.1
657 Lactobacillus hominis DSM 23910 = CRBIP 24.179 GCA_000296835.1
503 Lactobacillus iners DSM 13335 GCA_000160875.1
503 Lactobacillus jensenii GCA_001936235.1
503 Lactobacillus psittaci DSM 15354 GCA_000425905.1
Ah
619 Deinococcus radiodurans R1 GCA_000008565.1
388 Deinococcus deserti VCD115 GCA_000020685.1
350 Deinococcus hopiensis KR-140 GCA_900176165.1
Ai
792 Streptococcus mutans UA159 GCA_000007465.2
595 Streptococcus ratti FA-1 = DSM 20564 GCA_000286075.1
561 Streptococcus criceti HS-6 GCA_000187975.3
Aj
727 Deinococcus radiodurans R1 GCA_000008565.1
389 Deinococcus gobiensis I-0 GCA_000252445.1
359 Deinococcus proteolyticus MRP GCA_000190555.1
Ak
680 Streptococcus mutans UA159 GCA_000007465.2
474 Streptococcus ratti FA-1 = DSM 20564 GCA_000286075.1
451 Streptococcus pyogenes M1 GAS GCA_000006785.2
Al
738 Deinococcus radiodurans R1 GCA_000008565.1
468 Deinococcus deserti VCD115 GCA_000020685.1
429 Deinococcus soli Cha et al. 2016 GCA_001007995.1
Am
699 Streptococcus mutans UA159 GCA_000007465.2
464 Streptococcus ferus DSM 20646 GCA_000372425.1
456 Streptococcus ratti FA-1 = DSM 20564 GCA_000286075.1
An
623 Escherichia coli O157_3AH7 str. Sakai GCA_000008865.1
623 Escherichia coli str. K-12 substr. MG1655 GCA_000005845.2
623 Shigella flexneri 2a str. 301 GCA_000006925.2
615 Escherichia coli O83_3AH1 str. NRG 857C GCA_000183345.1
615 Escherichia coli UMN026 GCA_000026325.2
609 Escherichia coli O104_3AH4 str. 2011C-3493 GCA_000299455.1
Ao
651 Escherichia coli str. K-12 substr. MG1655 GCA_000005845.2
646 Escherichia coli O157_3AH7 str. Sakai GCA_000008865.1
646 Shigella flexneri 2a str. 301 GCA_000006925.2
643 Escherichia coli O83_3AH1 str. NRG 857C GCA_000183345.1
643 Escherichia coli UMN026 GCA_000026325.2
Ap
440 Enterococcus faecalis V583 GCA_000007785.1
407 Enterococcus hirae ATCC 9790 GCA_000271405.2
393 Enterococcus faecium DO GCA_000174395.2
393 Streptomyces cinnamoneus GCA_001885705.1
Aq
551 Rhodobacter sphaeroides 2.4.1 GCA_000012905.2
493 Rhodobacter sphaeroides ATCC 17025 GCA_000016405.1
458 Pseudorhodobacter ferrugineus DSM 5888 GCA_000420745.1
Ar
757 Staphylococcus epidermidis ATCC 12228 GCA_000007645.1
751 Staphylococcus capitis subsp. capitis GCA_001028645.1
719 Staphylococcus aureus subsp. aureus NCTC 8325 GCA_000013425.1
As
979 Staphylococcus epidermidis ATCC 12228 GCA_000007645.1
903 Staphylococcus capitis subsp. capitis GCA_001028645.1
830 Staphylococcus haemolyticus JCSC1435 GCA_000009865.1
At
692 Lactobacillus gasseri ATCC 33323 = JCM 1131 GCA_000014425.1
599 Lactobacillus hominis DSM 23910 = CRBIP 24.179 GCA_000296835.1
345 Lactobacillus amylovorus GCA_000191545.1
Au
699 Deinococcus radiodurans R1 GCA_000008565.1
381 Deinococcus deserti VCD115 GCA_000020685.1
377 Deinococcus soli Cha et al. 2016 GCA_001007995.1

Av
822 Lactobacillus gasseri ATCC 33323 = JCM 1131 GCA_000014425.1
733 Lactobacillus hominis DSM 23910 = CRBIP 24.179 GCA_000296835.1
579 Lactobacillus hamsteri DSM 5661 = JCM 6256 GCA_000615445.1
Aw
587 Streptococcus mutans UA159 GCA_000007465.2
439 Streptococcus ratti FA-1 = DSM 20564 GCA_000286075.1
395 Streptococcus sobrinus DSM 20742 = ATCC 33478 GCA_000686605.1
Ax
513 Staphylococcus epidermidis ATCC 12228 GCA_000007645.1
491 Staphylococcus capitis subsp. capitis GCA_001028645.1
453 Staphylococcus warneri SG1 GCA_000332735.1
Ay
695 Escherichia coli O104_3AH4 str. 2011C-3493 GCA_000299455.1
687 Escherichia coli str. K-12 substr. MG1655 GCA_000005845.2
686 Shigella flexneri 2a str. 301 GCA_000006925.2
Az
782 Escherichia coli O157_3AH7 str. Sakai GCA_000008865.1
782 Escherichia coli str. K-12 substr. MG1655 GCA_000005845.2
781 Escherichia coli UMN026 GCA_000026325.2
767 Escherichia coli O104_3AH4 str. 2011C-3493 GCA_000299455.1
767 Shigella flexneri 2a str. 301 GCA_000006925.2
AA
399 Lactobacillus gasseri ATCC 33323 = JCM 1131 GCA_000014425.1
387 Lactobacillus hominis DSM 23910 = CRBIP 24.179 GCA_000296835.1
305 Lactobacillus psittaci DSM 15354 GCA_000425905.1
AB
814 Staphylococcus epidermidis ATCC 12228 GCA_000007645.1
786 Staphylococcus capitis subsp. capitis GCA_001028645.1
709 Staphylococcus hominis subsp. hominis C80 GCA_000183685.1
AC
704 Deinococcus radiodurans R1 GCA_000008565.1
479 Deinococcus deserti VCD115 GCA_000020685.1
464 Deinococcus marmoris DSM 12784 GCA_000701405.1
AD
403 Deinococcus radiodurans R1 GCA_000008565.1
294 Deinococcus deserti VCD115 GCA_000020685.1
264 Deinococcus hopiensis KR-140 GCA_900176165.1
AE
688 Clostridium beijerinckii GCA_000833105.2
658 Clostridium saccharoperbutylacetonicum N1-4_28HMT_29 GCA_000340885.1
626 Clostridium puniceum GCA_002006345.1
AF
868 Clostridium beijerinckii GCA_000833105.2
824 Clostridium saccharoperbutylacetonicum N1-4_28HMT_29 GCA_000340885.1
818 Clostridium butyricum GCA_001456065.2
AG
948 Lactobacillus gasseri ATCC 33323 = JCM 1131 GCA_000014425.1
874 Lactobacillus hominis DSM 23910 = CRBIP 24.179 GCA_000296835.1
687 Lactobacillus iners DSM 13335 GCA_000160875.1
AH
570 Lactobacillus gasseri ATCC 33323 = JCM 1131 GCA_000014425.1
516 Lactobacillus hominis DSM 23910 = CRBIP 24.179 GCA_000296835.1
377 Lactobacillus iners DSM 13335 GCA_000160875.1
AI
733 Escherichia coli O104_3AH4 str. 2011C-3493 GCA_000299455.1
720 Escherichia coli str. K-12 substr. MG1655 GCA_000005845.2
715 Shigella flexneri 2a str. 301 GCA_000006925.2
AJ
782 Rhodobacter sphaeroides 2.4.1 GCA_000012905.2
713 Rhodobacter sphaeroides ATCC 17025 GCA_000016405.1
638 Gemmobacter aquatilis GCA_900110025.1
AK
424 Streptococcus mutans UA159 GCA_000007465.2
297 Streptococcus ratti FA-1 = DSM 20564 GCA_000286075.1
287 Streptococcus sobrinus DSM 20742 = ATCC 33478 GCA_000686605.1
AL
431 Enterococcus faecalis V583 GCA_000007785.1
384 Streptomyces cinnamoneus GCA_001885705.1
353 Enterococcus rivorum GCA_001742285.1
AM
514 Streptococcus mutans UA159 GCA_000007465.2
346 Streptococcus ratti FA-1 = DSM 20564 GCA_000286075.1
333 Streptococcus gordonii str. Challis substr. CH1 GCA_000017005.1
AN
636 Streptococcus mutans UA159 GCA_000007465.2

425 Streptococcus gordonii str. Challis substr. CH1 GCA_000017005.1
417 Streptococcus ratti FA-1 = DSM 20564 GCA_000286075.1
AO
903 Escherichia coli O104_3AH4 str. 2011C-3493 GCA_000299455.1
903 Escherichia coli str. K-12 substr. MG1655 GCA_000005845.2
895 Shigella flexneri 2a str. 301 GCA_000006925.2
857 Escherichia coli IAI39 GCA_000026345.1
AP
533 Lactobacillus gasseri ATCC 33323 = JCM 1131 GCA_000014425.1
496 Lactobacillus hominis DSM 23910 = CRBIP 24.179 GCA_000296835.1
355 Lactobacillus kalixensis DSM 16043 GCA_001434335.1
AQ
702 Rhodobacter sphaeroides 2.4.1 GCA_000012905.2
648 Rhodobacter sphaeroides ATCC 17025 GCA_000016405.1
541 Pseudorhodobacter ferrugineus DSM 5888 GCA_000420745.1
AR
782 Clostridium beijerinckii GCA_000833105.2
748 Clostridium saccharoperbutylacetonicum N1-4_28HMT_29 GCA_000340885.1
707 Clostridium puniceum GCA_002006345.1
AS
706 Staphylococcus epidermidis ATCC 12228 GCA_000007645.1
670 Staphylococcus capitis subsp. capitis GCA_001028645.1
660 Staphylococcus aureus subsp. aureus NCTC 8325 GCA_000013425.1
AT
751 Staphylococcus epidermidis ATCC 12228 GCA_000007645.1
714 Staphylococcus capitis subsp. capitis GCA_001028645.1
638 Staphylococcus hominis subsp. hominis C80 GCA_000183685.1
AU
795 Staphylococcus epidermidis ATCC 12228 GCA_000007645.1
756 Staphylococcus capitis subsp. capitis GCA_001028645.1
730 Staphylococcus hominis subsp. hominis C80 GCA_000183685.1
AV
570 Rhodobacter sphaeroides 2.4.1 GCA_000012905.2
528 Rhodobacter sphaeroides ATCC 17025 GCA_000016405.1
431 Pseudorhodobacter wandonensis GCA_001202035.1
AW
769 Deinococcus radiodurans R1 GCA_000008565.1
486 Deinococcus deserti VCD115 GCA_000020685.1
467 Deinococcus gobiensis I-0 GCA_000252445.1
AX
864 Rhodobacter sphaeroides 2.4.1 GCA_000012905.2
770 Rhodobacter sphaeroides ATCC 17025 GCA_000016405.1
685 Defluviimonas alba GCA_001620265.1
AY
743 Streptococcus mutans UA159 GCA_000007465.2
547 Streptococcus ratti FA-1 = DSM 20564 GCA_000286075.1
515 Streptococcus gordonii str. Challis substr. CH1 GCA_000017005.1
AZ
512 Clostridium beijerinckii GCA_000833105.2
508 Clostridium saccharobutylicum DSM 13864 GCA_000473995.1
494 Clostridium saccharoperbutylacetonicum N1-4_28HMT_29 GCA_000340885.1
B0
733 Streptococcus mutans UA159 GCA_000007465.2
528 Streptococcus gordonii str. Challis substr. CH1 GCA_000017005.1
528 Streptococcus ratti FA-1 = DSM 20564 GCA_000286075.1
509 Streptococcus henryi DSM 19005 GCA_000376985.1
B1
639 Staphylococcus epidermidis ATCC 12228 GCA_000007645.1
616 Staphylococcus capitis subsp. capitis GCA_001028645.1
578 Staphylococcus aureus subsp. aureus NCTC 8325 GCA_000013425.1
B2
680 Lactobacillus gasseri ATCC 33323 = JCM 1131 GCA_000014425.1
622 Lactobacillus hominis DSM 23910 = CRBIP 24.179 GCA_000296835.1
472 Lactobacillus iners DSM 13335 GCA_000160875.1
B3
712 Rhodobacter sphaeroides 2.4.1 GCA_000012905.2
616 Rhodobacter sphaeroides ATCC 17025 GCA_000016405.1
526 Defluviimonas alba GCA_001620265.1
B4
767 Lactobacillus gasseri ATCC 33323 = JCM 1131 GCA_000014425.1
710 Lactobacillus hominis DSM 23910 = CRBIP 24.179 GCA_000296835.1
574 Lactobacillus iners DSM 13335 GCA_000160875.1
B5
611 Deinococcus radiodurans R1 GCA_000008565.1
412 Deinococcus deserti VCD115 GCA_000020685.1
390 Deinococcus gobiensis I-0 GCA_000252445.1

B6
699 Streptococcus mutans UA159 GCA_000007465.2
556 Streptococcus ratti FA-1 = DSM 20564 GCA_000286075.1
529 Streptococcus sobrinus DSM 20742 = ATCC 33478 GCA_000686605.1
B7
721 Deinococcus radiodurans R1 GCA_000008565.1
481 Deinococcus murrayi DSM 11303 GCA_000482805.1
473 Deinococcus deserti VCD115 GCA_000020685.1
B8
752 Escherichia coli str. K-12 substr. MG1655 GCA_000005845.2
748 Escherichia coli IAI39 GCA_000026345.1
748 Escherichia coli O157_3AH7 str. Sakai GCA_000008865.1
727 Escherichia coli O83_3AH1 str. NRG 857C GCA_000183345.1
B9
590 Rhodobacter sphaeroides 2.4.1 GCA_000012905.2
517 Rhodobacter sphaeroides ATCC 17025 GCA_000016405.1
459 Paracoccus solventivorans GCA_900142875.1
Ba
580 Clostridium beijerinckii GCA_000833105.2
577 Clostridium saccharoperbutylacetonicum N1-4_28HMT_29 GCA_000340885.1
573 Clostridium puniceum GCA_002006345.1
Bb
710 Enterococcus faecalis V583 GCA_000007785.1
663 Streptomyces cinnamoneus GCA_001885705.1
619 Enterococcus faecium DO GCA_000174395.2
Bc
610 Escherichia coli IAI39 GCA_000026345.1
610 Escherichia coli O157_3AH7 str. Sakai GCA_000008865.1
610 Escherichia coli str. K-12 substr. MG1655 GCA_000005845.2
610 Shigella flexneri 2a str. 301 GCA_000006925.2
609 Escherichia coli O83_3AH1 str. NRG 857C GCA_000183345.1
609 Escherichia coli UMN026 GCA_000026325.2
585 Escherichia coli O104_3AH4 str. 2011C-3493 GCA_000299455.1
Bd
631 Clostridium beijerinckii GCA_000833105.2
604 Clostridium saccharoperbutylacetonicum N1-4_28HMT_29 GCA_000340885.1
563 Clostridium puniceum GCA_002006345.1
563 Clostridium saccharobutylicum DSM 13864 GCA_000473995.1
Be
571 Rhodobacter sphaeroides 2.4.1 GCA_000012905.2
517 Rhodobacter sphaeroides ATCC 17025 GCA_000016405.1
385 Pseudorhodobacter ferrugineus DSM 5888 GCA_000420745.1
Bf
803 Staphylococcus epidermidis ATCC 12228 GCA_000007645.1
774 Staphylococcus lugdunensis HKU09-01 GCA_000025085.1
766 Staphylococcus haemolyticus JCSC1435 GCA_000009865.1
Bg
575 Enterococcus faecalis V583 GCA_000007785.1
529 Streptomyces cinnamoneus GCA_001885705.1
515 Enterococcus asini ATCC 700915 GCA_000407365.1
Bh
749 Enterococcus faecalis V583 GCA_000007785.1
705 Streptomyces cinnamoneus GCA_001885705.1
689 Enterococcus rivorum GCA_001742285.1
Bi
833 Staphylococcus epidermidis ATCC 12228 GCA_000007645.1
799 Staphylococcus capitis subsp. capitis GCA_001028645.1
744 Staphylococcus hominis subsp. hominis C80 GCA_000183685.1
Bj
802 Deinococcus radiodurans R1 GCA_000008565.1
492 Deinococcus deserti VCD115 GCA_000020685.1
463 Deinococcus hopiensis KR-140 GCA_900176165.1
Bk
789 Deinococcus radiodurans R1 GCA_000008565.1
503 Deinococcus deserti VCD115 GCA_000020685.1
475 Deinococcus hopiensis KR-140 GCA_900176165.1
Bl
616 Rhodobacter sphaeroides 2.4.1 GCA_000012905.2
571 Rhodobacter sphaeroides ATCC 17025 GCA_000016405.1
496 Pseudorhodobacter psychrotolerans GCA_001294535.1
Bm
637 Staphylococcus epidermidis ATCC 12228 GCA_000007645.1
596 Staphylococcus capitis subsp. capitis GCA_001028645.1
554 Staphylococcus simulans GCA_001559115.1
Bn
550 Deinococcus radiodurans R1 GCA_000008565.1

303 Deinococcus deserti VCD115 GCA_000020685.1
297 Deinococcus gobiensis I-0 GCA_000252445.1
Bo
694 Bacillus anthracis str. Ames GCA_000007845.1
694 Bacillus anthracis str. Sterne GCA_000008165.1
694 Bacillus cereus ATCC 14579 GCA_000007825.1
694 _5BBacillus thuringiensis_5D serovar konkukian str. 97-27 GCA_000008505.1
673 Bacillus thuringiensis YBT-1518 GCA_000497525.2
667 Bacillus pseudomycoides DSM 12442 GCA_000161455.1
Bp
674 Lactobacillus gasseri ATCC 33323 = JCM 1131 GCA_000014425.1
626 Lactobacillus hominis DSM 23910 = CRBIP 24.179 GCA_000296835.1
515 Lactobacillus iners DSM 13335 GCA_000160875.1
Bq
605 Enterococcus faecalis V583 GCA_000007785.1
558 Streptomyces cinnamoneus GCA_001885705.1
500 Enterococcus faecium DO GCA_000174395.2
500 Enterococcus rivorum GCA_001742285.1
Br
436 Bacillus pseudomycoides DSM 12442 GCA_000161455.1
434 Bacillus anthracis str. Ames GCA_000007845.1
434 Bacillus anthracis str. Sterne GCA_000008165.1
434 _5BBacillus thuringiensis_5D serovar konkukian str. 97-27 GCA_000008505.1
425 Bacillus cereus ATCC 14579 GCA_000007825.1
425 Bacillus thuringiensis YBT-1518 GCA_000497525.2
Bs
808 Staphylococcus epidermidis ATCC 12228 GCA_000007645.1
764 Staphylococcus capitis subsp. capitis GCA_001028645.1
744 Staphylococcus haemolyticus JCSC1435 GCA_000009865.1
Bt
843 Clostridium beijerinckii GCA_000833105.2
837 Clostridium saccharoperbutylacetonicum N1-4_28HMT_29 GCA_000340885.1
836 Clostridium butyricum GCA_001456065.2
Bu
595 Deinococcus radiodurans R1 GCA_000008565.1
399 Deinococcus deserti VCD115 GCA_000020685.1
398 Deinococcus soli Cha et al. 2016 GCA_001007995.1
Bv
562 Streptococcus mutans UA159 GCA_000007465.2
418 Streptococcus ratti FA-1 = DSM 20564 GCA_000286075.1
402 Streptococcus gordonii str. Challis substr. CH1 GCA_000017005.1
Bw
696 Streptococcus mutans UA159 GCA_000007465.2
520 Streptococcus ratti FA-1 = DSM 20564 GCA_000286075.1
490 Streptococcus henryi DSM 19005 GCA_000376985.1
Bx
556 Clostridium beijerinckii GCA_000833105.2
531 Clostridium saccharoperbutylacetonicum N1-4_28HMT_29 GCA_000340885.1
517 Clostridium butyricum GCA_001456065.2
By
630 Enterococcus faecalis V583 GCA_000007785.1
583 Streptomyces cinnamoneus GCA_001885705.1
500 Enterococcus rivorum GCA_001742285.1
Bz
780 Enterococcus faecalis V583 GCA_000007785.1
739 Streptomyces cinnamoneus GCA_001885705.1
653 Enterococcus rivorum GCA_001742285.1
BA
459 Rhodobacter sphaeroides 2.4.1 GCA_000012905.2
416 Rhodobacter sphaeroides ATCC 17025 GCA_000016405.1
338 Defluviimonas alba GCA_001620265.1
BB
811 Lactobacillus gasseri ATCC 33323 = JCM 1131 GCA_000014425.1
756 Lactobacillus hominis DSM 23910 = CRBIP 24.179 GCA_000296835.1
509 Lactobacillus iners DSM 13335 GCA_000160875.1
BC
531 Clostridium beijerinckii GCA_000833105.2
473 Clostridium saccharoperbutylacetonicum N1-4_28HMT_29 GCA_000340885.1
448 Clostridium butyricum GCA_001456065.2
BD
331 Clostridium beijerinckii GCA_000833105.2
272 Clostridium saccharoperbutylacetonicum N1-4_28HMT_29 GCA_000340885.1
267 Clostridium sartagoforme AAU1 GCA_000401215.1
BE
712 Rhodobacter sphaeroides 2.4.1 GCA_000012905.2
653 Rhodobacter sphaeroides ATCC 17025 GCA_000016405.1

566 Gemmobacter nectariphilus DSM 15620 GCA_000429765.1
BF
846 Rhodobacter sphaeroides 2.4.1 GCA_000012905.2
757 Rhodobacter sphaeroides ATCC 17025 GCA_000016405.1
610 Gemmobacter megaterium GCA_900156815.1
BG
731 Deinococcus radiodurans R1 GCA_000008565.1
498 Deinococcus deserti VCD115 GCA_000020685.1
484 Deinococcus gobiensis I-0 GCA_000252445.1
BH
578 Streptococcus mutans UA159 GCA_000007465.2
398 Streptococcus ratti FA-1 = DSM 20564 GCA_000286075.1
380 Streptococcus sobrinus DSM 20742 = ATCC 33478 GCA_000686605.1
BI
911 Lactobacillus gasseri ATCC 33323 = JCM 1131 GCA_000014425.1
878 Lactobacillus hominis DSM 23910 = CRBIP 24.179 GCA_000296835.1
608 Lactobacillus hamsteri DSM 5661 = JCM 6256 GCA_000615445.1
BJ
508 Clostridium beijerinckii GCA_000833105.2
464 Clostridium saccharoperbutylacetonicum N1-4_28HMT_29 GCA_000340885.1
453 Clostridium puniceum GCA_002006345.1
BK
776 Lactobacillus gasseri ATCC 33323 = JCM 1131 GCA_000014425.1
640 Lactobacillus hominis DSM 23910 = CRBIP 24.179 GCA_000296835.1
460 Lactobacillus delbrueckii subsp. bulgaricus ATCC 11842 = JCM 1002 GCA_000056065.1
BL
543 Bifidobacterium adolescentis ATCC 15703 GCA_000010425.1
419 Bifidobacterium angulatum DSM 20098 = JCM 7096 GCA_001025155.1
415 Bifidobacterium dentium JCM 1195 = DSM 20436 GCA_001042595.1
BM
732 Lactobacillus gasseri ATCC 33323 = JCM 1131 GCA_000014425.1
699 Lactobacillus hominis DSM 23910 = CRBIP 24.179 GCA_000296835.1
458 Lactobacillus psittaci DSM 15354 GCA_000425905.1
BN
662 Staphylococcus epidermidis ATCC 12228 GCA_000007645.1
654 Staphylococcus capitis subsp. capitis GCA_001028645.1
634 Staphylococcus haemolyticus JCSC1435 GCA_000009865.1
BO
809 Rhodobacter sphaeroides 2.4.1 GCA_000012905.2
751 Rhodobacter sphaeroides ATCC 17025 GCA_000016405.1
604 Defluviimonas alba GCA_001620265.1
BP
639 Clostridium beijerinckii GCA_000833105.2
585 Clostridium saccharoperbutylacetonicum N1-4_28HMT_29 GCA_000340885.1
580 Clostridium puniceum GCA_002006345.1
BQ
828 Staphylococcus epidermidis ATCC 12228 GCA_000007645.1
759 Staphylococcus capitis subsp. capitis GCA_001028645.1
745 Staphylococcus warneri SG1 GCA_000332735.1
BR
862 Lactobacillus gasseri ATCC 33323 = JCM 1131 GCA_000014425.1
808 Lactobacillus hominis DSM 23910 = CRBIP 24.179 GCA_000296835.1
547 Lactobacillus amylovorus GCA_000191545.1
BS
542 Lactobacillus gasseri ATCC 33323 = JCM 1131 GCA_000014425.1
499 Lactobacillus hominis DSM 23910 = CRBIP 24.179 GCA_000296835.1
369 Lactobacillus iners DSM 13335 GCA_000160875.1
BT
582 Clostridium beijerinckii GCA_000833105.2
548 Clostridium saccharoperbutylacetonicum N1-4_28HMT_29 GCA_000340885.1
543 Clostridium saccharobutylicum DSM 13864 GCA_000473995.1
BU
778 Deinococcus radiodurans R1 GCA_000008565.1
467 Deinococcus deserti VCD115 GCA_000020685.1
463 Deinococcus soli Cha et al. 2016 GCA_001007995.1
BV
429 Streptococcus mutans UA159 GCA_000007465.2
337 Streptococcus ratti FA-1 = DSM 20564 GCA_000286075.1
303 Streptococcus ferus DSM 20646 GCA_000372425.1
BW
741 Bifidobacterium adolescentis ATCC 15703 GCA_000010425.1
635 Bifidobacterium dentium JCM 1195 = DSM 20436 GCA_001042595.1
625 Bifidobacterium angulatum DSM 20098 = JCM 7096 GCA_001025155.1
BX
814 Clostridium beijerinckii GCA_000833105.2
802 Clostridium saccharoperbutylacetonicum N1-4_28HMT_29 GCA_000340885.1

772 Clostridium butyricum GCA_001456065.2
BY
728 Deinococcus radiodurans R1 GCA_000008565.1
466 Deinococcus gobiensis I-0 GCA_000252445.1
443 Deinococcus deserti VCD115 GCA_000020685.1
BZ
483 Bifidobacterium adolescentis ATCC 15703 GCA_000010425.1
397 Bifidobacterium angulatum DSM 20098 = JCM 7096 GCA_001025155.1
392 Bifidobacterium gallicum DSM 20093 = LMG 11596 GCA_000741205.1
C0
756 Lactobacillus gasseri ATCC 33323 = JCM 1131 GCA_000014425.1
697 Lactobacillus hominis DSM 23910 = CRBIP 24.179 GCA_000296835.1
482 Lactobacillus iners DSM 13335 GCA_000160875.1
C1
629 Streptococcus mutans UA159 GCA_000007465.2
512 Streptococcus ratti FA-1 = DSM 20564 GCA_000286075.1
467 Streptococcus criceti HS-6 GCA_000187975.3
467 Streptococcus macacae NCTC 11558 GCA_000187995.3
C2
716 Bifidobacterium adolescentis ATCC 15703 GCA_000010425.1
543 Bifidobacterium angulatum DSM 20098 = JCM 7096 GCA_001025155.1
534 Bifidobacterium dentium JCM 1195 = DSM 20436 GCA_001042595.1
C3
769 Staphylococcus epidermidis ATCC 12228 GCA_000007645.1
735 Staphylococcus capitis subsp. capitis GCA_001028645.1
683 Staphylococcus lugdunensis HKU09-01 GCA_000025085.1
C4
652 Clostridium beijerinckii GCA_000833105.2
628 Clostridium saccharoperbutylacetonicum N1-4_28HMT_29 GCA_000340885.1
613 Clostridium puniceum GCA_002006345.1
C5
586 Rhodobacter sphaeroides 2.4.1 GCA_000012905.2
552 Rhodobacter sphaeroides ATCC 17025 GCA_000016405.1
455 Gemmobacter aquatilis GCA_900110025.1
C6
502 Rhodobacter sphaeroides 2.4.1 GCA_000012905.2
419 Rhodobacter sphaeroides ATCC 17025 GCA_000016405.1
356 Gemmobacter megaterium GCA_900156815.1
C7
766 Bifidobacterium adolescentis ATCC 15703 GCA_000010425.1
642 Bifidobacterium dentium JCM 1195 = DSM 20436 GCA_001042595.1
615 Bifidobacterium angulatum DSM 20098 = JCM 7096 GCA_001025155.1
C8
592 Streptococcus mutans UA159 GCA_000007465.2
467 Clostridium beijerinckii GCA_000833105.2
464 Streptococcus gordonii str. Challis substr. CH1 GCA_000017005.1
C9
432 Clostridium beijerinckii GCA_000833105.2
421 Clostridium saccharoperbutylacetonicum N1-4_28HMT_29 GCA_000340885.1
403 Clostridium puniceum GCA_002006345.1
Ca
698 Clostridium beijerinckii GCA_000833105.2
647 Clostridium saccharobutylicum DSM 13864 GCA_000473995.1
621 Clostridium puniceum GCA_002006345.1
621 Clostridium saccharoperbutylacetonicum N1-4_28HMT_29 GCA_000340885.1
Cb
810 Streptococcus mutans UA159 GCA_000007465.2
602 Streptococcus ratti FA-1 = DSM 20564 GCA_000286075.1
543 Streptococcus equinus GCA_000964315.1
Cc
631 Clostridium beijerinckii GCA_000833105.2
550 Clostridium butyricum GCA_001456065.2
550 Clostridium saccharoperbutylacetonicum N1-4_28HMT_29 GCA_000340885.1
542 Clostridium chromiireducens GCA_002029255.1
542 Clostridium saccharobutylicum DSM 13864 GCA_000473995.1
Cd
745 Streptococcus mutans UA159 GCA_000007465.2
529 Streptococcus macacae NCTC 11558 GCA_000187995.3
525 Streptococcus ratti FA-1 = DSM 20564 GCA_000286075.1
Ce
794 Bacillus anthracis str. Ames GCA_000007845.1
794 Bacillus anthracis str. Sterne GCA_000008165.1
794 _5BBacillus thuringiensis_5D serovar konkukian str. 97-27 GCA_000008505.1
779 Bacillus cereus ATCC 14579 GCA_000007825.1
767 Bacillus thuringiensis YBT-1518 GCA_000497525.2
Cf

723 Deinococcus radiodurans R1 GCA_000008565.1
413 Deinococcus deserti VCD115 GCA_000020685.1
390 Deinococcus gobiensis I-0 GCA_000252445.1
Cg
725 Clostridium beijerinckii GCA_000833105.2
699 Clostridium saccharoperbutylacetonicum N1-4_28HMT_29 GCA_000340885.1
679 Clostridium saccharobutylicum DSM 13864 GCA_000473995.1
Ch
808 Streptococcus mutans UA159 GCA_000007465.2
574 Streptococcus gordonii str. Challis substr. CH1 GCA_000017005.1
546 Streptococcus ratti FA-1 = DSM 20564 GCA_000286075.1
Ci
767 Enterococcus faecalis V583 GCA_000007785.1
726 Streptomyces cinnamoneus GCA_001885705.1
654 Enterococcus asini ATCC 700915 GCA_000407365.1
Cj
677 Deinococcus radiodurans R1 GCA_000008565.1
439 Deinococcus gobiensis I-0 GCA_000252445.1
430 Deinococcus murrayi DSM 11303 GCA_000482805.1
Ck
820 Clostridium beijerinckii GCA_000833105.2
795 Clostridium saccharoperbutylacetonicum N1-4_28HMT_29 GCA_000340885.1
761 Clostridium saccharobutylicum DSM 13864 GCA_000473995.1
Cl
485 Rhodobacter sphaeroides 2.4.1 GCA_000012905.2
428 Rhodobacter sphaeroides ATCC 17025 GCA_000016405.1
406 Defluviimonas alba GCA_001620265.1
Cm
613 Staphylococcus epidermidis ATCC 12228 GCA_000007645.1
598 Staphylococcus capitis subsp. capitis GCA_001028645.1
563 Megasphaera cerevisiae DSM 20462 GCA_001045675.1
563 Staphylococcus warneri SG1 GCA_000332735.1
Cn
681 Rhodobacter sphaeroides 2.4.1 GCA_000012905.2
561 Rhodobacter sphaeroides ATCC 17025 GCA_000016405.1
522 Gemmobacter megaterium GCA_900156815.1
Co
742 Escherichia coli str. K-12 substr. MG1655 GCA_000005845.2
722 Escherichia coli O157_3AH7 str. Sakai GCA_000008865.1
722 Shigella flexneri 2a str. 301 GCA_000006925.2
716 Escherichia coli O104_3AH4 str. 2011C-3493 GCA_000299455.1
Cp
588 Rhodobacter sphaeroides 2.4.1 GCA_000012905.2
480 Rhodobacter sphaeroides ATCC 17025 GCA_000016405.1
467 Gemmobacter aquatilis GCA_900110025.1
Cq
345 Lactobacillus gasseri ATCC 33323 = JCM 1131 GCA_000014425.1
331 Lactobacillus hominis DSM 23910 = CRBIP 24.179 GCA_000296835.1
240 Lactobacillus iners DSM 13335 GCA_000160875.1
Cr
814 Streptococcus mutans UA159 GCA_000007465.2
646 Streptococcus ratti FA-1 = DSM 20564 GCA_000286075.1
612 Streptococcus ferus DSM 20646 GCA_000372425.1
Cs
574 Bifidobacterium adolescentis ATCC 15703 GCA_000010425.1
471 Bifidobacterium dentium JCM 1195 = DSM 20436 GCA_001042595.1
437 Bifidobacterium angulatum DSM 20098 = JCM 7096 GCA_001025155.1
Ct
620 Streptococcus mutans UA159 GCA_000007465.2
465 Streptococcus ratti FA-1 = DSM 20564 GCA_000286075.1
407 Streptococcus sanguinis SK36 GCA_000014205.1
Cu
712 _5BBacillus thuringiensis_5D serovar konkukian str. 97-27 GCA_000008505.1
711 Bacillus anthracis str. Ames GCA_000007845.1
711 Bacillus anthracis str. Sterne GCA_000008165.1
703 Bacillus cereus ATCC 14579 GCA_000007825.1
Cv
548 Clostridium beijerinckii GCA_000833105.2
515 Clostridium saccharoperbutylacetonicum N1-4_28HMT_29 GCA_000340885.1
492 Clostridium saccharobutylicum DSM 13864 GCA_000473995.1
Cw
941 Clostridium beijerinckii GCA_000833105.2
898 Clostridium saccharoperbutylacetonicum N1-4_28HMT_29 GCA_000340885.1
878 Clostridium puniceum GCA_002006345.1
Cx
665 Clostridium beijerinckii GCA_000833105.2

637 Clostridium saccharoperbutylacetonicum N1-4_28HMT_29 GCA_000340885.1
587 Clostridium puniceum GCA_002006345.1
Cy
800 Deinococcus radiodurans R1 GCA_000008565.1
508 Deinococcus deserti VCD115 GCA_000020685.1
495 Deinococcus soli Cha et al. 2016 GCA_001007995.1
Cz
474 Clostridium beijerinckii GCA_000833105.2
473 Clostridium saccharoperbutylacetonicum N1-4_28HMT_29 GCA_000340885.1
459 Clostridium saccharobutylicum DSM 13864 GCA_000473995.1
CA
698 Streptococcus mutans UA159 GCA_000007465.2
505 Streptococcus ratti FA-1 = DSM 20564 GCA_000286075.1
472 Streptococcus gordonii str. Challis substr. CH1 GCA_000017005.1
CB
518 Bifidobacterium adolescentis ATCC 15703 GCA_000010425.1
472 Bifidobacterium dentium JCM 1195 = DSM 20436 GCA_001042595.1
457 Bifidobacterium coryneforme GCA_000737865.1
CC
674 Bifidobacterium adolescentis ATCC 15703 GCA_000010425.1
543 Bifidobacterium dentium JCM 1195 = DSM 20436 GCA_001042595.1
530 Bifidobacterium angulatum DSM 20098 = JCM 7096 GCA_001025155.1
CD
594 Bifidobacterium adolescentis ATCC 15703 GCA_000010425.1
563 Bifidobacterium dentium JCM 1195 = DSM 20436 GCA_001042595.1
510 Bifidobacterium stellenboschense GCA_000741785.1
CE
884 Streptococcus mutans UA159 GCA_000007465.2
710 Streptococcus ratti FA-1 = DSM 20564 GCA_000286075.1
658 Streptococcus gordonii str. Challis substr. CH1 GCA_000017005.1
658 Streptococcus henryi DSM 19005 GCA_000376985.1
CF
640 Bacillus cereus ATCC 14579 GCA_000007825.1
630 Bacillus thuringiensis YBT-1518 GCA_000497525.2
628 Bacillus anthracis str. Ames GCA_000007845.1
628 Bacillus anthracis str. Sterne GCA_000008165.1
628 _5BBacillus thuringiensis_5D serovar konkukian str. 97-27 GCA_000008505.1
CG
641 Clostridium beijerinckii GCA_000833105.2
604 Clostridium saccharoperbutylacetonicum N1-4_28HMT_29 GCA_000340885.1
592 Clostridium saccharobutylicum DSM 13864 GCA_000473995.1
CH
685 Staphylococcus epidermidis ATCC 12228 GCA_000007645.1
642 Staphylococcus capitis subsp. capitis GCA_001028645.1
594 Staphylococcus haemolyticus JCSC1435 GCA_000009865.1
CI
657 Lactobacillus gasseri ATCC 33323 = JCM 1131 GCA_000014425.1
641 Lactobacillus hominis DSM 23910 = CRBIP 24.179 GCA_000296835.1
461 Lactobacillus iners DSM 13335 GCA_000160875.1
CJ
784 Clostridium beijerinckii GCA_000833105.2
737 Clostridium saccharoperbutylacetonicum N1-4_28HMT_29 GCA_000340885.1
705 Clostridium saccharobutylicum DSM 13864 GCA_000473995.1
CK
639 Clostridium beijerinckii GCA_000833105.2
605 Clostridium saccharoperbutylacetonicum N1-4_28HMT_29 GCA_000340885.1
598 Clostridium saccharobutylicum DSM 13864 GCA_000473995.1
CL
421 Bacillus anthracis str. Ames GCA_000007845.1
421 Bacillus anthracis str. Sterne GCA_000008165.1
421 _5BBacillus thuringiensis_5D serovar konkukian str. 97-27 GCA_000008505.1
413 Bacillus pseudomycoides DSM 12442 GCA_000161455.1
406 Bacillus cereus ATCC 14579 GCA_000007825.1
406 Bacillus thuringiensis YBT-1518 GCA_000497525.2
CM
798 Staphylococcus epidermidis ATCC 12228 GCA_000007645.1
749 Staphylococcus capitis subsp. capitis GCA_001028645.1
726 Staphylococcus hominis subsp. hominis C80 GCA_000183685.1
CN
741 Staphylococcus epidermidis ATCC 12228 GCA_000007645.1
729 Staphylococcus capitis subsp. capitis GCA_001028645.1
675 Staphylococcus aureus subsp. aureus NCTC 8325 GCA_000013425.1
CO
405 Clostridium beijerinckii GCA_000833105.2
404 Clostridium saccharoperbutylacetonicum N1-4_28HMT_29 GCA_000340885.1
360 Clostridium puniceum GCA_002006345.1

CP
611 Clostridium puniceum GCA_002006345.1
605 Clostridium beijerinckii GCA_000833105.2
603 Clostridium saccharoperbutylacetonicum N1-4_28HMT_29 GCA_000340885.1
CQ
604 Deinococcus radiodurans R1 GCA_000008565.1
351 Deinococcus deserti VCD115 GCA_000020685.1
307 Deinococcus marmoris DSM 12784 GCA_000701405.1
307 Deinococcus soli Cha et al. 2016 GCA_001007995.1
CR
684 Deinococcus radiodurans R1 GCA_000008565.1
373 Deinococcus deserti VCD115 GCA_000020685.1
370 Deinococcus gobiensis I-0 GCA_000252445.1
CS
529 Staphylococcus epidermidis ATCC 12228 GCA_000007645.1
521 Staphylococcus capitis subsp. capitis GCA_001028645.1
485 Staphylococcus aureus subsp. aureus NCTC 8325 GCA_000013425.1
CT
502 Staphylococcus epidermidis ATCC 12228 GCA_000007645.1
459 Staphylococcus capitis subsp. capitis GCA_001028645.1
457 Staphylococcus aureus subsp. aureus NCTC 8325 GCA_000013425.1
CU
669 Deinococcus radiodurans R1 GCA_000008565.1
388 Deinococcus soli Cha et al. 2016 GCA_001007995.1
380 Deinococcus gobiensis I-0 GCA_000252445.1
CV
733 Clostridium beijerinckii GCA_000833105.2
689 Clostridium saccharoperbutylacetonicum N1-4_28HMT_29 GCA_000340885.1
664 Clostridium puniceum GCA_002006345.1
CW
627 Rhodobacter sphaeroides 2.4.1 GCA_000012905.2
530 Rhodobacter sphaeroides ATCC 17025 GCA_000016405.1
460 Gemmobacter aquatilis GCA_900110025.1
CX
436 Lactobacillus gasseri ATCC 33323 = JCM 1131 GCA_000014425.1
422 Lactobacillus hominis DSM 23910 = CRBIP 24.179 GCA_000296835.1
323 Lactobacillus jensenii GCA_001936235.1
323 Lactobacillus psittaci DSM 15354 GCA_000425905.1
CY
739 Clostridium beijerinckii GCA_000833105.2
709 Clostridium saccharoperbutylacetonicum N1-4_28HMT_29 GCA_000340885.1
670 Clostridium saccharobutylicum DSM 13864 GCA_000473995.1
CZ
770 Deinococcus radiodurans R1 GCA_000008565.1
473 Deinococcus gobiensis I-0 GCA_000252445.1
450 Deinococcus deserti VCD115 GCA_000020685.1
D0
709 Shigella flexneri 2a str. 301 GCA_000006925.2
698 Escherichia coli UMN026 GCA_000026325.2
698 Escherichia coli str. K-12 substr. MG1655 GCA_000005845.2
693 Escherichia coli O157_3AH7 str. Sakai GCA_000008865.1
D1
802 Deinococcus radiodurans R1 GCA_000008565.1
591 Deinococcus deserti VCD115 GCA_000020685.1
556 Deinococcus puniceus GCA_001644565.1
D2
698 Enterococcus faecalis V583 GCA_000007785.1
651 Streptomyces cinnamoneus GCA_001885705.1
603 Enterococcus faecium DO GCA_000174395.2
D3
542 Lactobacillus gasseri ATCC 33323 = JCM 1131 GCA_000014425.1
485 Lactobacillus hominis DSM 23910 = CRBIP 24.179 GCA_000296835.1
328 Lactobacillus iners DSM 13335 GCA_000160875.1
D4
585 Rhodobacter sphaeroides 2.4.1 GCA_000012905.2
514 Rhodobacter sphaeroides ATCC 17025 GCA_000016405.1
469 Gemmobacter aquatilis GCA_900110025.1
D5
620 Clostridium beijerinckii GCA_000833105.2
597 Clostridium saccharobutylicum DSM 13864 GCA_000473995.1
586 Clostridium saccharoperbutylacetonicum N1-4_28HMT_29 GCA_000340885.1
D6
471 Rhodobacter sphaeroides 2.4.1 GCA_000012905.2
441 Rhodobacter sphaeroides ATCC 17025 GCA_000016405.1
363 Pseudorhodobacter psychrotolerans GCA_001294535.1
D7

635 Clostridium beijerinckii GCA_000833105.2
600 Clostridium saccharoperbutylacetonicum N1-4_28HMT_29 GCA_000340885.1
580 Clostridium saccharobutylicum DSM 13864 GCA_000473995.1
D8
681 Deinococcus radiodurans R1 GCA_000008565.1
441 Deinococcus deserti VCD115 GCA_000020685.1
389 Deinococcus gobiensis I-0 GCA_000252445.1
D9
627 Staphylococcus epidermidis ATCC 12228 GCA_000007645.1
578 Staphylococcus capitis subsp. capitis GCA_001028645.1
546 Staphylococcus lugdunensis HKU09-01 GCA_000025085.1
Da
825 Staphylococcus epidermidis ATCC 12228 GCA_000007645.1
754 Staphylococcus capitis subsp. capitis GCA_001028645.1
679 Staphylococcus lugdunensis HKU09-01 GCA_000025085.1
Db
661 Staphylococcus epidermidis ATCC 12228 GCA_000007645.1
623 Staphylococcus capitis subsp. capitis GCA_001028645.1
565 Staphylococcus warneri SG1 GCA_000332735.1
Dc
782 Deinococcus radiodurans R1 GCA_000008565.1
463 Deinococcus deserti VCD115 GCA_000020685.1
447 Deinococcus gobiensis I-0 GCA_000252445.1
Dd
697 Clostridium beijerinckii GCA_000833105.2
657 Clostridium saccharoperbutylacetonicum N1-4_28HMT_29 GCA_000340885.1
622 Clostridium saccharobutylicum DSM 13864 GCA_000473995.1
De
591 Deinococcus radiodurans R1 GCA_000008565.1
375 Deinococcus puniceus GCA_001644565.1
362 Deinococcus deserti VCD115 GCA_000020685.1
Df
705 Bacillus anthracis str. Ames GCA_000007845.1
705 Bacillus anthracis str. Sterne GCA_000008165.1
705 _5BBacillus thuringiensis_5D serovar konkukian str. 97-27 GCA_000008505.1
690 Bacillus cereus ATCC 14579 GCA_000007825.1
672 Bacillus thuringiensis YBT-1518 GCA_000497525.2
Dg
787 Staphylococcus epidermidis ATCC 12228 GCA_000007645.1
775 Staphylococcus capitis subsp. capitis GCA_001028645.1
706 Staphylococcus warneri SG1 GCA_000332735.1
Dh
412 Rhodobacter sphaeroides 2.4.1 GCA_000012905.2
397 Rhodobacter sphaeroides ATCC 17025 GCA_000016405.1
328 Gemmobacter megaterium GCA_900156815.1
Di
823 Bifidobacterium adolescentis ATCC 15703 GCA_000010425.1
728 Bifidobacterium dentium JCM 1195 = DSM 20436 GCA_001042595.1
665 Bifidobacterium angulatum DSM 20098 = JCM 7096 GCA_001025155.1
Dj
744 Lactobacillus gasseri ATCC 33323 = JCM 1131 GCA_000014425.1
662 Lactobacillus hominis DSM 23910 = CRBIP 24.179 GCA_000296835.1
498 Lactobacillus iners DSM 13335 GCA_000160875.1
Dk
715 Rhodobacter sphaeroides 2.4.1 GCA_000012905.2
645 Rhodobacter sphaeroides ATCC 17025 GCA_000016405.1
546 Thioclava indica GCA_000714545.1
Dl
848 Clostridium saccharoperbutylacetonicum N1-4_28HMT_29 GCA_000340885.1
814 Clostridium beijerinckii GCA_000833105.2
779 Clostridium puniceum GCA_002006345.1
Dm
626 Clostridium beijerinckii GCA_000833105.2
571 Clostridium saccharoperbutylacetonicum N1-4_28HMT_29 GCA_000340885.1
566 Clostridium saccharobutylicum DSM 13864 GCA_000473995.1
Dn
662 Rhodobacter sphaeroides 2.4.1 GCA_000012905.2
609 Rhodobacter sphaeroides ATCC 17025 GCA_000016405.1
506 Pseudorhodobacter ferrugineus DSM 5888 GCA_000420745.1
506 Pseudorhodobacter wandonensis GCA_001202035.1
Do
764 Escherichia coli O157_3AH7 str. Sakai GCA_000008865.1
764 Escherichia coli str. K-12 substr. MG1655 GCA_000005845.2
764 Shigella flexneri 2a str. 301 GCA_000006925.2
761 Escherichia coli O83_3AH1 str. NRG 857C GCA_000183345.1
759 Escherichia coli UMN026 GCA_000026325.2

Dp
754 Lactobacillus gasseri ATCC 33323 = JCM 1131 GCA_000014425.1
706 Lactobacillus hominis DSM 23910 = CRBIP 24.179 GCA_000296835.1
499 Lactobacillus hamsteri DSM 5661 = JCM 6256 GCA_000615445.1
Dq
687 Lactobacillus gasseri ATCC 33323 = JCM 1131 GCA_000014425.1
639 Lactobacillus hominis DSM 23910 = CRBIP 24.179 GCA_000296835.1
489 Lactobacillus iners DSM 13335 GCA_000160875.1
Dr
642 Rhodobacter sphaeroides 2.4.1 GCA_000012905.2
612 Rhodobacter sphaeroides ATCC 17025 GCA_000016405.1
458 Pseudorhodobacter psychrotolerans GCA_001294535.1
Ds
772 Rhodobacter sphaeroides 2.4.1 GCA_000012905.2
735 Rhodobacter sphaeroides ATCC 17025 GCA_000016405.1
609 Gemmobacter aquatilis GCA_900110025.1
Dt
632 Enterococcus faecalis V583 GCA_000007785.1
585 Streptomyces cinnamoneus GCA_001885705.1
559 Enterococcus thailandicus GCA_001652875.1
Du
621 Escherichia coli O104_3AH4 str. 2011C-3493 GCA_000299455.1
621 Escherichia coli str. K-12 substr. MG1655 GCA_000005845.2
611 Shigella flexneri 2a str. 301 GCA_000006925.2
600 Escherichia coli IAI39 GCA_000026345.1
600 Escherichia coli O157_3AH7 str. Sakai GCA_000008865.1
Dv
812 Rhodobacter sphaeroides 2.4.1 GCA_000012905.2
734 Rhodobacter sphaeroides ATCC 17025 GCA_000016405.1
613 Gemmobacter megaterium GCA_900156815.1
Dw
605 Bifidobacterium adolescentis ATCC 15703 GCA_000010425.1
479 Bifidobacterium angulatum DSM 20098 = JCM 7096 GCA_001025155.1
475 Bifidobacterium callitrichos DSM 23973 GCA_000741175.1
Dx
730 Lactobacillus gasseri ATCC 33323 = JCM 1131 GCA_000014425.1
689 Lactobacillus hominis DSM 23910 = CRBIP 24.179 GCA_000296835.1
568 Lactobacillus iners DSM 13335 GCA_000160875.1
Dy
736 Bifidobacterium adolescentis ATCC 15703 GCA_000010425.1
643 Bifidobacterium angulatum DSM 20098 = JCM 7096 GCA_001025155.1
603 Bifidobacterium dentium JCM 1195 = DSM 20436 GCA_001042595.1
Dz
649 Staphylococcus epidermidis ATCC 12228 GCA_000007645.1
597 Staphylococcus capitis subsp. capitis GCA_001028645.1
595 Staphylococcus warneri SG1 GCA_000332735.1
DA
758 Deinococcus radiodurans R1 GCA_000008565.1
463 Deinococcus deserti VCD115 GCA_000020685.1
430 Deinococcus gobiensis I-0 GCA_000252445.1
DB
694 Staphylococcus capitis subsp. capitis GCA_001028645.1
694 Staphylococcus epidermidis ATCC 12228 GCA_000007645.1
637 Staphylococcus haemolyticus JCSC1435 GCA_000009865.1
626 Staphylococcus aureus subsp. aureus NCTC 8325 GCA_000013425.1
DC
975 Lactobacillus gasseri ATCC 33323 = JCM 1131 GCA_000014425.1
895 Lactobacillus hominis DSM 23910 = CRBIP 24.179 GCA_000296835.1
737 Lactobacillus hamsteri DSM 5661 = JCM 6256 GCA_000615445.1
DD
744 Bifidobacterium adolescentis ATCC 15703 GCA_000010425.1
603 Bifidobacterium angulatum DSM 20098 = JCM 7096 GCA_001025155.1
562 Bifidobacterium callitrichos DSM 23973 GCA_000741175.1
DE
644 Bifidobacterium adolescentis ATCC 15703 GCA_000010425.1
507 Bifidobacterium dentium JCM 1195 = DSM 20436 GCA_001042595.1
492 Bifidobacterium angulatum DSM 20098 = JCM 7096 GCA_001025155.1
DF
594 Bifidobacterium adolescentis ATCC 15703 GCA_000010425.1
494 Bifidobacterium angulatum DSM 20098 = JCM 7096 GCA_001025155.1
469 Bifidobacterium dentium JCM 1195 = DSM 20436 GCA_001042595.1
DG
496 Staphylococcus epidermidis ATCC 12228 GCA_000007645.1
455 Staphylococcus hominis subsp. hominis C80 GCA_000183685.1
441 Staphylococcus lugdunensis HKU09-01 GCA_000025085.1
DH

608 Rhodobacter sphaeroides 2.4.1 GCA_000012905.2
574 Rhodobacter sphaeroides ATCC 17025 GCA_000016405.1
501 Gemmobacter aquatilis GCA_900110025.1
DI
674 Rhodobacter sphaeroides 2.4.1 GCA_000012905.2
593 Rhodobacter sphaeroides ATCC 17025 GCA_000016405.1
452 Pseudorhodobacter psychrotolerans GCA_001294535.1
DJ
687 Lactobacillus gasseri ATCC 33323 = JCM 1131 GCA_000014425.1
670 Lactobacillus hominis DSM 23910 = CRBIP 24.179 GCA_000296835.1
470 Lactobacillus iners DSM 13335 GCA_000160875.1
DK
745 Enterococcus faecalis V583 GCA_000007785.1
698 Streptomyces cinnamoneus GCA_001885705.1
637 Enterococcus rivorum GCA_001742285.1
DL
526 Staphylococcus epidermidis ATCC 12228 GCA_000007645.1
510 Staphylococcus capitis subsp. capitis GCA_001028645.1
494 Staphylococcus haemolyticus JCSC1435 GCA_000009865.1
DM
590 Bifidobacterium adolescentis ATCC 15703 GCA_000010425.1
475 Bifidobacterium callitrichos DSM 23973 GCA_000741175.1
470 Bifidobacterium dentium JCM 1195 = DSM 20436 GCA_001042595.1
DN
795 Escherichia coli O157_3AH7 str. Sakai GCA_000008865.1
795 Escherichia coli str. K-12 substr. MG1655 GCA_000005845.2
787 Escherichia coli O83_3AH1 str. NRG 857C GCA_000183345.1
787 Escherichia coli UMN026 GCA_000026325.2
786 Escherichia coli IAI39 GCA_000026345.1
DO
614 Escherichia coli str. K-12 substr. MG1655 GCA_000005845.2
608 Escherichia coli O157_3AH7 str. Sakai GCA_000008865.1
596 Escherichia coli UMN026 GCA_000026325.2
DP
652 Clostridium beijerinckii GCA_000833105.2
652 Clostridium saccharoperbutylacetonicum N1-4_28HMT_29 GCA_000340885.1
631 Clostridium puniceum GCA_002006345.1
628 Clostridium saccharobutylicum DSM 13864 GCA_000473995.1
DQ
639 Enterococcus faecalis V583 GCA_000007785.1
593 Streptomyces cinnamoneus GCA_001885705.1
553 Enterococcus faecium DO GCA_000174395.2
DR
654 Rhodobacter sphaeroides 2.4.1 GCA_000012905.2
592 Rhodobacter sphaeroides ATCC 17025 GCA_000016405.1
491 Defluviimonas alba GCA_001620265.1
DS
809 Escherichia coli O104_3AH4 str. 2011C-3493 GCA_000299455.1
809 Escherichia coli str. K-12 substr. MG1655 GCA_000005845.2
809 Shigella flexneri 2a str. 301 GCA_000006925.2
782 Escherichia coli O157_3AH7 str. Sakai GCA_000008865.1
770 Escherichia coli O83_3AH1 str. NRG 857C GCA_000183345.1
DT
769 Deinococcus radiodurans R1 GCA_000008565.1
489 Deinococcus puniceus GCA_001644565.1
485 Deinococcus hopiensis KR-140 GCA_900176165.1
DU
534 Bacillus anthracis str. Ames GCA_000007845.1
534 Bacillus anthracis str. Sterne GCA_000008165.1
534 _5BBacillus thuringiensis_5D serovar konkukian str. 97-27 GCA_000008505.1
521 Bacillus pseudomycoides DSM 12442 GCA_000161455.1
518 Bacillus cereus ATCC 14579 GCA_000007825.1
518 Bacillus thuringiensis YBT-1518 GCA_000497525.2
DV
554 Clostridium beijerinckii GCA_000833105.2
506 Clostridium saccharobutylicum DSM 13864 GCA_000473995.1
503 Clostridium butyricum GCA_001456065.2
503 Clostridium saccharoperbutylacetonicum N1-4_28HMT_29 GCA_000340885.1
DW
680 Staphylococcus epidermidis ATCC 12228 GCA_000007645.1
650 Staphylococcus capitis subsp. capitis GCA_001028645.1
584 Staphylococcus haemolyticus JCSC1435 GCA_000009865.1
DX
744 Streptococcus mutans UA159 GCA_000007465.2
564 Streptococcus ratti FA-1 = DSM 20564 GCA_000286075.1
519 Streptococcus macacae NCTC 11558 GCA_000187995.3

DY
390 Clostridium beijerinckii GCA_000833105.2
351 Clostridium saccharoperbutylacetonicum N1-4_28HMT_29 GCA_000340885.1
350 Clostridium butyricum GCA_001456065.2
DZ
790 Bacillus anthracis str. Ames GCA_000007845.1
790 Bacillus anthracis str. Sterne GCA_000008165.1
790 _5BBacillus thuringiensis_5D serovar konkukian str. 97-27 GCA_000008505.1
788 Bacillus pseudomycoides DSM 12442 GCA_000161455.1
775 Bacillus cereus ATCC 14579 GCA_000007825.1
E0
371 Bifidobacterium adolescentis ATCC 15703 GCA_000010425.1
313 Bifidobacterium angulatum DSM 20098 = JCM 7096 GCA_001025155.1
307 Bifidobacterium dentium JCM 1195 = DSM 20436 GCA_001042595.1
E1
790 Bifidobacterium adolescentis ATCC 15703 GCA_000010425.1
656 Bifidobacterium dentium JCM 1195 = DSM 20436 GCA_001042595.1
630 Bifidobacterium angulatum DSM 20098 = JCM 7096 GCA_001025155.1
E2
836 Enterococcus faecalis V583 GCA_000007785.1
804 Streptomyces cinnamoneus GCA_001885705.1
719 Enterococcus rivorum GCA_001742285.1
E3
702 Staphylococcus epidermidis ATCC 12228 GCA_000007645.1
673 Staphylococcus capitis subsp. capitis GCA_001028645.1
633 Staphylococcus haemolyticus JCSC1435 GCA_000009865.1
E4
628 Deinococcus radiodurans R1 GCA_000008565.1
405 Deinococcus deserti VCD115 GCA_000020685.1
383 Deinococcus gobiensis I-0 GCA_000252445.1
E5
855 Streptococcus mutans UA159 GCA_000007465.2
633 Streptococcus ratti FA-1 = DSM 20564 GCA_000286075.1
603 Streptococcus gordonii str. Challis substr. CH1 GCA_000017005.1
E6
726 Deinococcus radiodurans R1 GCA_000008565.1
400 Deinococcus hopiensis KR-140 GCA_900176165.1
392 Deinococcus deserti VCD115 GCA_000020685.1
E7
907 Clostridium beijerinckii GCA_000833105.2
883 Clostridium saccharoperbutylacetonicum N1-4_28HMT_29 GCA_000340885.1
869 Clostridium saccharobutylicum DSM 13864 GCA_000473995.1
E8
311 Clostridium beijerinckii GCA_000833105.2
296 Clostridium saccharobutylicum DSM 13864 GCA_000473995.1
287 Clostridium butyricum GCA_001456065.2
E9
684 Streptococcus mutans UA159 GCA_000007465.2
482 Streptococcus ratti FA-1 = DSM 20564 GCA_000286075.1
445 Streptococcus macacae NCTC 11558 GCA_000187995.3
Ea
532 Bifidobacterium adolescentis ATCC 15703 GCA_000010425.1
528 Bifidobacterium dentium JCM 1195 = DSM 20436 GCA_001042595.1
473 Bifidobacterium angulatum DSM 20098 = JCM 7096 GCA_001025155.1
Eb
725 Clostridium beijerinckii GCA_000833105.2
689 Clostridium saccharoperbutylacetonicum N1-4_28HMT_29 GCA_000340885.1
670 Clostridium puniceum GCA_002006345.1
Ec
505 Escherichia coli O157_3AH7 str. Sakai GCA_000008865.1
505 Escherichia coli UMN026 GCA_000026325.2
505 Escherichia coli str. K-12 substr. MG1655 GCA_000005845.2
486 Escherichia coli IAI39 GCA_000026345.1
486 Escherichia coli O83_3AH1 str. NRG 857C GCA_000183345.1
486 Shigella dysenteriae Sd197 GCA_000012005.1
486 Shigella flexneri 2a str. 301 GCA_000006925.2
465 Escherichia coli O104_3AH4 str. 2011C-3493 GCA_000299455.1
Ed
581 Clostridium beijerinckii GCA_000833105.2
556 Clostridium puniceum GCA_002006345.1
550 Clostridium saccharoperbutylacetonicum N1-4_28HMT_29 GCA_000340885.1
Ee
757 Deinococcus radiodurans R1 GCA_000008565.1
531 Deinococcus gobiensis I-0 GCA_000252445.1
527 Deinococcus hopiensis KR-140 GCA_900176165.1
Ef

828 Lactobacillus gasseri ATCC 33323 = JCM 1131 GCA_000014425.1
757 Lactobacillus hominis DSM 23910 = CRBIP 24.179 GCA_000296835.1
514 Lactobacillus psittaci DSM 15354 GCA_000425905.1
Eg
671 Staphylococcus epidermidis ATCC 12228 GCA_000007645.1
639 Staphylococcus capitis subsp. capitis GCA_001028645.1
582 Megasphaera cerevisiae DSM 20462 GCA_001045675.1
582 Staphylococcus warneri SG1 GCA_000332735.1
Eh
858 Staphylococcus epidermidis ATCC 12228 GCA_000007645.1
801 Staphylococcus capitis subsp. capitis GCA_001028645.1
756 Staphylococcus lugdunensis HKU09-01 GCA_000025085.1
Ei
465 Bifidobacterium adolescentis ATCC 15703 GCA_000010425.1
334 Bifidobacterium dentium JCM 1195 = DSM 20436 GCA_001042595.1
316 Bifidobacterium angulatum DSM 20098 = JCM 7096 GCA_001025155.1
Ej
653 Deinococcus radiodurans R1 GCA_000008565.1
427 Deinococcus deserti VCD115 GCA_000020685.1
404 Deinococcus frigens DSM 12807 GCA_000701425.1
404 Deinococcus soli Cha et al. 2016 GCA_001007995.1
Ek
660 Bacillus anthracis str. Ames GCA_000007845.1
660 Bacillus anthracis str. Sterne GCA_000008165.1
660 _5BBacillus thuringiensis_5D serovar konkukian str. 97-27 GCA_000008505.1
651 Bacillus cereus ATCC 14579 GCA_000007825.1
639 Bacillus pseudomycoides DSM 12442 GCA_000161455.1
El
478 Clostridium beijerinckii GCA_000833105.2
475 Clostridium saccharobutylicum DSM 13864 GCA_000473995.1
471 Clostridium saccharoperbutylacetonicum N1-4_28HMT_29 GCA_000340885.1
Em
518 Lactobacillus gasseri ATCC 33323 = JCM 1131 GCA_000014425.1
467 Lactobacillus hominis DSM 23910 = CRBIP 24.179 GCA_000296835.1
317 Lactobacillus iners DSM 13335 GCA_000160875.1
En
805 Streptococcus mutans UA159 GCA_000007465.2
586 Streptococcus gordonii str. Challis substr. CH1 GCA_000017005.1
576 Streptococcus ratti FA-1 = DSM 20564 GCA_000286075.1
Eo
367 Deinococcus radiodurans R1 GCA_000008565.1
259 Deinococcus gobiensis I-0 GCA_000252445.1
211 Deinococcus deserti VCD115 GCA_000020685.1
Ep
896 Bacillus anthracis str. Ames GCA_000007845.1
896 Bacillus anthracis str. Sterne GCA_000008165.1
890 Bacillus cereus ATCC 14579 GCA_000007825.1
890 _5BBacillus thuringiensis_5D serovar konkukian str. 97-27 GCA_000008505.1
881 Bacillus pseudomycoides DSM 12442 GCA_000161455.1
Eq
661 Rhodobacter sphaeroides 2.4.1 GCA_000012905.2
585 Rhodobacter sphaeroides ATCC 17025 GCA_000016405.1
503 Gemmobacter aquatilis GCA_900110025.1
Er
919 Rhodobacter sphaeroides 2.4.1 GCA_000012905.2
855 Rhodobacter sphaeroides ATCC 17025 GCA_000016405.1
711 Gemmobacter aquatilis GCA_900110025.1
Es
683 Clostridium beijerinckii GCA_000833105.2
673 Clostridium saccharoperbutylacetonicum N1-4_28HMT_29 GCA_000340885.1
631 Clostridium saccharobutylicum DSM 13864 GCA_000473995.1
Et
874 Streptococcus mutans UA159 GCA_000007465.2
655 Streptococcus ratti FA-1 = DSM 20564 GCA_000286075.1
611 Streptococcus gordonii str. Challis substr. CH1 GCA_000017005.1
Eu
804 Clostridium beijerinckii GCA_000833105.2
768 Clostridium saccharoperbutylacetonicum N1-4_28HMT_29 GCA_000340885.1
742 Clostridium puniceum GCA_002006345.1
Ev
679 Lactobacillus gasseri ATCC 33323 = JCM 1131 GCA_000014425.1
615 Lactobacillus hominis DSM 23910 = CRBIP 24.179 GCA_000296835.1
451 Lactobacillus psittaci DSM 15354 GCA_000425905.1
Ew
727 Streptococcus mutans UA159 GCA_000007465.2
569 Streptococcus gordonii str. Challis substr. CH1 GCA_000017005.1

560 Streptococcus ratti FA-1 = DSM 20564 GCA_000286075.1
Ex
537 Rhodobacter sphaeroides 2.4.1 GCA_000012905.2
503 Rhodobacter sphaeroides ATCC 17025 GCA_000016405.1
391 Pseudorhodobacter ferrugineus DSM 5888 GCA_000420745.1
Ey
605 Rhodobacter sphaeroides 2.4.1 GCA_000012905.2
540 Rhodobacter sphaeroides ATCC 17025 GCA_000016405.1
437 Thioclava nitratireducens GCA_001940525.2
Ez
514 Streptococcus mutans UA159 GCA_000007465.2
362 Streptococcus gordonii str. Challis substr. CH1 GCA_000017005.1
338 Streptococcus iniae GCA_000831485.1
EA
901 Streptococcus mutans UA159 GCA_000007465.2
654 Streptococcus ratti FA-1 = DSM 20564 GCA_000286075.1
600 Streptococcus gordonii str. Challis substr. CH1 GCA_000017005.1
EB
719 Rhodobacter sphaeroides 2.4.1 GCA_000012905.2
615 Rhodobacter sphaeroides ATCC 17025 GCA_000016405.1
521 Pseudorhodobacter ferrugineus DSM 5888 GCA_000420745.1
EC
680 Staphylococcus epidermidis ATCC 12228 GCA_000007645.1
613 Staphylococcus capitis subsp. capitis GCA_001028645.1
595 Staphylococcus warneri SG1 GCA_000332735.1
ED
700 Streptococcus mutans UA159 GCA_000007465.2
468 Streptococcus ratti FA-1 = DSM 20564 GCA_000286075.1
446 Streptococcus gordonii str. Challis substr. CH1 GCA_000017005.1
EE
946 Staphylococcus epidermidis ATCC 12228 GCA_000007645.1
922 Staphylococcus capitis subsp. capitis GCA_001028645.1
857 Staphylococcus haemolyticus JCSC1435 GCA_000009865.1
EF
700 Lactobacillus gasseri ATCC 33323 = JCM 1131 GCA_000014425.1
668 Lactobacillus hominis DSM 23910 = CRBIP 24.179 GCA_000296835.1
525 Lactobacillus psittaci DSM 15354 GCA_000425905.1
EG
812 Rhodobacter sphaeroides 2.4.1 GCA_000012905.2
723 Rhodobacter sphaeroides ATCC 17025 GCA_000016405.1
603 Gemmobacter megaterium GCA_900156815.1
EH
655 Streptococcus mutans UA159 GCA_000007465.2
448 Streptococcus ratti FA-1 = DSM 20564 GCA_000286075.1
426 Streptococcus gordonii str. Challis substr. CH1 GCA_000017005.1
EI
568 Staphylococcus epidermidis ATCC 12228 GCA_000007645.1
542 Staphylococcus capitis subsp. capitis GCA_001028645.1
518 Staphylococcus simulans GCA_001559115.1
EJ
715 Clostridium beijerinckii GCA_000833105.2
707 Clostridium saccharoperbutylacetonicum N1-4_28HMT_29 GCA_000340885.1
696 Clostridium butyricum GCA_001456065.2
EK
719 Bifidobacterium adolescentis ATCC 15703 GCA_000010425.1
544 Bifidobacterium angulatum DSM 20098 = JCM 7096 GCA_001025155.1
537 Bifidobacterium dentium JCM 1195 = DSM 20436 GCA_001042595.1
EL
610 Lactobacillus gasseri ATCC 33323 = JCM 1131 GCA_000014425.1
540 Lactobacillus hominis DSM 23910 = CRBIP 24.179 GCA_000296835.1
376 Lactobacillus iners DSM 13335 GCA_000160875.1
EM
473 Enterococcus faecalis V583 GCA_000007785.1
438 Enterococcus haemoperoxidus ATCC BAA-382 GCA_000407165.1
435 Enterococcus rivorum GCA_001742285.1
EN
724 Staphylococcus epidermidis ATCC 12228 GCA_000007645.1
688 Staphylococcus lugdunensis HKU09-01 GCA_000025085.1
677 Staphylococcus capitis subsp. capitis GCA_001028645.1
EO
821 Bacillus pseudomycoides DSM 12442 GCA_000161455.1
819 Bacillus anthracis str. Ames GCA_000007845.1
819 Bacillus anthracis str. Sterne GCA_000008165.1
819 Bacillus cereus ATCC 14579 GCA_000007825.1
819 _5BBacillus thuringiensis_5D serovar konkukian str. 97-27 GCA_000008505.1
797 Bacillus mycoides GCA_000832605.1

EP
879 Streptococcus mutans UA159 GCA_000007465.2
659 Streptococcus ratti FA-1 = DSM 20564 GCA_000286075.1
609 Streptococcus gordonii str. Challis substr. CH1 GCA_000017005.1
EQ
662 Lactobacillus gasseri ATCC 33323 = JCM 1131 GCA_000014425.1
634 Lactobacillus hominis DSM 23910 = CRBIP 24.179 GCA_000296835.1
503 Lactobacillus psittaci DSM 15354 GCA_000425905.1
ER
564 Staphylococcus epidermidis ATCC 12228 GCA_000007645.1
509 Staphylococcus capitis subsp. capitis GCA_001028645.1
459 Staphylococcus simulans GCA_001559115.1
ES
608 Clostridium beijerinckii GCA_000833105.2
584 Clostridium saccharoperbutylacetonicum N1-4_28HMT_29 GCA_000340885.1
583 Clostridium saccharobutylicum DSM 13864 GCA_000473995.1
ET
504 Staphylococcus epidermidis ATCC 12228 GCA_000007645.1
498 Staphylococcus capitis subsp. capitis GCA_001028645.1
467 Staphylococcus aureus subsp. aureus NCTC 8325 GCA_000013425.1
467 Staphylococcus haemolyticus JCSC1435 GCA_000009865.1
467 Staphylococcus hominis subsp. hominis C80 GCA_000183685.1
EU
634 Bifidobacterium adolescentis ATCC 15703 GCA_000010425.1
503 Bifidobacterium angulatum DSM 20098 = JCM 7096 GCA_001025155.1
496 Bifidobacterium dentium JCM 1195 = DSM 20436 GCA_001042595.1
EV
561 Staphylococcus epidermidis ATCC 12228 GCA_000007645.1
507 Staphylococcus aureus subsp. aureus NCTC 8325 GCA_000013425.1
506 Staphylococcus capitis subsp. capitis GCA_001028645.1
EW
734 Rhodobacter sphaeroides 2.4.1 GCA_000012905.2
651 Rhodobacter sphaeroides ATCC 17025 GCA_000016405.1
547 Gemmobacter aquatilis GCA_900110025.1
EX
519 Streptococcus mutans UA159 GCA_000007465.2
412 Streptococcus ratti FA-1 = DSM 20564 GCA_000286075.1
367 Streptococcus criceti HS-6 GCA_000187975.3
EY
729 Bifidobacterium adolescentis ATCC 15703 GCA_000010425.1
631 Bifidobacterium dentium JCM 1195 = DSM 20436 GCA_001042595.1
563 Bifidobacterium angulatum DSM 20098 = JCM 7096 GCA_001025155.1
EZ
609 Bifidobacterium adolescentis ATCC 15703 GCA_000010425.1
531 Bifidobacterium dentium JCM 1195 = DSM 20436 GCA_001042595.1
529 Bifidobacterium angulatum DSM 20098 = JCM 7096 GCA_001025155.1
F0
834 Lactobacillus gasseri ATCC 33323 = JCM 1131 GCA_000014425.1
726 Lactobacillus hominis DSM 23910 = CRBIP 24.179 GCA_000296835.1
540 Lactobacillus iners DSM 13335 GCA_000160875.1
F1
216 Bifidobacterium adolescentis ATCC 15703 GCA_000010425.1
215 Bifidobacterium dentium JCM 1195 = DSM 20436 GCA_001042595.1
194 Bifidobacterium breve DSM 20213 = JCM 1192 GCA_001025175.1
194 Bifidobacterium lemurum GCA_001895165.1
F2
495 Clostridium beijerinckii GCA_000833105.2
455 Clostridium saccharoperbutylacetonicum N1-4_28HMT_29 GCA_000340885.1
417 Clostridium butyricum GCA_001456065.2
F3
506 Clostridium beijerinckii GCA_000833105.2
462 Clostridium saccharoperbutylacetonicum N1-4_28HMT_29 GCA_000340885.1
460 Clostridium puniceum GCA_002006345.1
F4
602 Clostridium beijerinckii GCA_000833105.2
593 Clostridium saccharobutylicum DSM 13864 GCA_000473995.1
591 Clostridium saccharoperbutylacetonicum N1-4_28HMT_29 GCA_000340885.1
F5
278 Escherichia coli IAI39 GCA_000026345.1
278 Escherichia coli O104_3AH4 str. 2011C-3493 GCA_000299455.1
278 Escherichia coli O157_3AH7 str. Sakai GCA_000008865.1
278 Escherichia coli str. K-12 substr. MG1655 GCA_000005845.2
278 Shigella dysenteriae Sd197 GCA_000012005.1
278 Shigella flexneri 2a str. 301 GCA_000006925.2
270 Escherichia coli O83_3AH1 str. NRG 857C GCA_000183345.1
270 Escherichia coli UMN026 GCA_000026325.2

246 Tumebacillus flagellatus GCA_000714935.1
F6
721 Staphylococcus epidermidis ATCC 12228 GCA_000007645.1
679 Staphylococcus capitis subsp. capitis GCA_001028645.1
626 Staphylococcus aureus subsp. aureus NCTC 8325 GCA_000013425.1
F7
807 Streptococcus mutans UA159 GCA_000007465.2
577 Streptococcus gordonii str. Challis substr. CH1 GCA_000017005.1
560 Streptococcus equinus GCA_000964315.1
F8
691 Rhodobacter sphaeroides 2.4.1 GCA_000012905.2
620 Rhodobacter sphaeroides ATCC 17025 GCA_000016405.1
515 Pseudorhodobacter ferrugineus DSM 5888 GCA_000420745.1
F9
365 Bifidobacterium adolescentis ATCC 15703 GCA_000010425.1
286 Bifidobacterium angulatum DSM 20098 = JCM 7096 GCA_001025155.1
269 Bifidobacterium dentium JCM 1195 = DSM 20436 GCA_001042595.1
Fa
308 Rhodobacter sphaeroides 2.4.1 GCA_000012905.2
294 Rhodobacter sphaeroides ATCC 17025 GCA_000016405.1
241 Gemmobacter aquatilis GCA_900110025.1
Fb
722 Staphylococcus epidermidis ATCC 12228 GCA_000007645.1
664 Staphylococcus capitis subsp. capitis GCA_001028645.1
601 Megasphaera cerevisiae DSM 20462 GCA_001045675.1
601 Staphylococcus warneri SG1 GCA_000332735.1
Fc
868 Streptococcus mutans UA159 GCA_000007465.2
640 Streptococcus ratti FA-1 = DSM 20564 GCA_000286075.1
625 Streptococcus gordonii str. Challis substr. CH1 GCA_000017005.1
Fd
512 Rhodobacter sphaeroides 2.4.1 GCA_000012905.2
451 Rhodobacter sphaeroides ATCC 17025 GCA_000016405.1
386 Pseudorhodobacter ferrugineus DSM 5888 GCA_000420745.1
Fe
716 Rhodobacter sphaeroides 2.4.1 GCA_000012905.2
594 Rhodobacter sphaeroides ATCC 17025 GCA_000016405.1
520 Gemmobacter aquatilis GCA_900110025.1
Ff
423 Deinococcus radiodurans R1 GCA_000008565.1
262 Deinococcus proteolyticus MRP GCA_000190555.1
245 Deinococcus frigens DSM 12807 GCA_000701425.1
Fg
597 Rhodobacter sphaeroides 2.4.1 GCA_000012905.2
505 Rhodobacter sphaeroides ATCC 17025 GCA_000016405.1
406 Gemmobacter aquatilis GCA_900110025.1
Fh
713 Staphylococcus epidermidis ATCC 12228 GCA_000007645.1
684 Staphylococcus capitis subsp. capitis GCA_001028645.1
623 Staphylococcus simulans GCA_001559115.1
Fi
752 Streptococcus mutans UA159 GCA_000007465.2
536 Streptococcus ratti FA-1 = DSM 20564 GCA_000286075.1
529 Streptococcus equinus GCA_000964315.1
529 Streptococcus henryi DSM 19005 GCA_000376985.1
Fj
438 Clostridium saccharoperbutylacetonicum N1-4_28HMT_29 GCA_000340885.1
425 Clostridium beijerinckii GCA_000833105.2
404 Clostridium saccharobutylicum DSM 13864 GCA_000473995.1
Fk
433 Deinococcus radiodurans R1 GCA_000008565.1
298 Deinococcus marmoris DSM 12784 GCA_000701405.1
295 Deinococcus soli Cha et al. 2016 GCA_001007995.1
Fl
980 Bacillus anthracis str. Ames GCA_000007845.1
980 Bacillus anthracis str. Sterne GCA_000008165.1
980 _5BBacillus thuringiensis_5D serovar konkukian str. 97-27 GCA_000008505.1
971 Bacillus cereus ATCC 14579 GCA_000007825.1
956 Bacillus thuringiensis YBT-1518 GCA_000497525.2
Fm
886 Bifidobacterium adolescentis ATCC 15703 GCA_000010425.1
757 Bifidobacterium dentium JCM 1195 = DSM 20436 GCA_001042595.1
730 Bifidobacterium angulatum DSM 20098 = JCM 7096 GCA_001025155.1
Fn
840 Lactobacillus gasseri ATCC 33323 = JCM 1131 GCA_000014425.1
740 Lactobacillus hominis DSM 23910 = CRBIP 24.179 GCA_000296835.1

553 Lactobacillus gallinarum GCA_001314245.2
Fo
397 Deinococcus radiodurans R1 GCA_000008565.1
273 Deinococcus gobiensis I-0 GCA_000252445.1
265 Deinococcus soli Cha et al. 2016 GCA_001007995.1
Fp
620 Clostridium beijerinckii GCA_000833105.2
607 Clostridium saccharobutylicum DSM 13864 GCA_000473995.1
602 Clostridium saccharoperbutylacetonicum N1-4_28HMT_29 GCA_000340885.1
Fq
645 Clostridium beijerinckii GCA_000833105.2
633 Clostridium saccharoperbutylacetonicum N1-4_28HMT_29 GCA_000340885.1
604 Clostridium puniceum GCA_002006345.1
Fr
794 Lactobacillus gasseri ATCC 33323 = JCM 1131 GCA_000014425.1
720 Lactobacillus hominis DSM 23910 = CRBIP 24.179 GCA_000296835.1
515 Lactobacillus iners DSM 13335 GCA_000160875.1
Fs
374 Streptococcus mutans UA159 GCA_000007465.2
267 Streptococcus iniae GCA_000831485.1
267 Streptococcus uberis 0140J GCA_000009545.1
266 Streptococcus dysgalactiae subsp. equisimilis AC-2713 GCA_000317855.1
Ft
663 Clostridium beijerinckii GCA_000833105.2
654 Clostridium saccharoperbutylacetonicum N1-4_28HMT_29 GCA_000340885.1
645 Clostridium saccharobutylicum DSM 13864 GCA_000473995.1
Fu
695 Staphylococcus epidermidis ATCC 12228 GCA_000007645.1
654 Staphylococcus capitis subsp. capitis GCA_001028645.1
613 Staphylococcus lugdunensis HKU09-01 GCA_000025085.1
Fv
835 Lactobacillus gasseri ATCC 33323 = JCM 1131 GCA_000014425.1
807 Lactobacillus hominis DSM 23910 = CRBIP 24.179 GCA_000296835.1
632 Lactobacillus jensenii GCA_001936235.1
Fw
588 Bacillus anthracis str. Ames GCA_000007845.1
588 Bacillus anthracis str. Sterne GCA_000008165.1
588 _5BBacillus thuringiensis_5D serovar konkukian str. 97-27 GCA_000008505.1
578 Bacillus cereus ATCC 14579 GCA_000007825.1
578 Bacillus thuringiensis YBT-1518 GCA_000497525.2
577 Bacillus pseudomycoides DSM 12442 GCA_000161455.1
Fx
736 Staphylococcus epidermidis ATCC 12228 GCA_000007645.1
682 Staphylococcus capitis subsp. capitis GCA_001028645.1
665 Staphylococcus haemolyticus JCSC1435 GCA_000009865.1
Fy
746 Bifidobacterium adolescentis ATCC 15703 GCA_000010425.1
608 Bifidobacterium angulatum DSM 20098 = JCM 7096 GCA_001025155.1
589 Bifidobacterium dentium JCM 1195 = DSM 20436 GCA_001042595.1
Fz
475 Clostridium beijerinckii GCA_000833105.2
445 Clostridium butyricum GCA_001456065.2
441 Clostridium neonatale GCA_001458595.1
FA
709 Clostridium beijerinckii GCA_000833105.2
667 Clostridium puniceum GCA_002006345.1
667 Clostridium saccharoperbutylacetonicum N1-4_28HMT_29 GCA_000340885.1
634 Clostridium saccharobutylicum DSM 13864 GCA_000473995.1
FB
531 Escherichia coli str. K-12 substr. MG1655 GCA_000005845.2
530 Escherichia coli IAI39 GCA_000026345.1
530 Escherichia coli O157_3AH7 str. Sakai GCA_000008865.1
522 Kosakonia cowanii GCA_001975225.1
FC
638 Rhodobacter sphaeroides 2.4.1 GCA_000012905.2
586 Rhodobacter sphaeroides ATCC 17025 GCA_000016405.1
489 Pseudorhodobacter psychrotolerans GCA_001294535.1
FD
685 Staphylococcus epidermidis ATCC 12228 GCA_000007645.1
656 Staphylococcus capitis subsp. capitis GCA_001028645.1
639 Staphylococcus haemolyticus JCSC1435 GCA_000009865.1
FE
838 Rhodobacter sphaeroides 2.4.1 GCA_000012905.2
727 Rhodobacter sphaeroides ATCC 17025 GCA_000016405.1
606 Gemmobacter megaterium GCA_900156815.1
FF

938 Bacillus anthracis str. Ames GCA_000007845.1
938 Bacillus anthracis str. Sterne GCA_000008165.1
938 _5BBacillus thuringiensis_5D serovar konkukian str. 97-27 GCA_000008505.1
930 Bacillus cereus ATCC 14579 GCA_000007825.1
925 Bacillus pseudomycoides DSM 12442 GCA_000161455.1
FG
542 Bacillus anthracis str. Ames GCA_000007845.1
542 Bacillus anthracis str. Sterne GCA_000008165.1
542 Bacillus cereus ATCC 14579 GCA_000007825.1
542 Bacillus thuringiensis YBT-1518 GCA_000497525.2
542 _5BBacillus thuringiensis_5D serovar konkukian str. 97-27 GCA_000008505.1
527 Bacillus pseudomycoides DSM 12442 GCA_000161455.1
523 Bacillus mycoides GCA_000832605.1
FH
687 Deinococcus radiodurans R1 GCA_000008565.1
397 Deinococcus puniceus GCA_001644565.1
395 Deinococcus deserti VCD115 GCA_000020685.1
FI
493 Rhodobacter sphaeroides 2.4.1 GCA_000012905.2
420 Rhodobacter sphaeroides ATCC 17025 GCA_000016405.1
408 Defluviimonas alba GCA_001620265.1
FJ
630 Lactobacillus gasseri ATCC 33323 = JCM 1131 GCA_000014425.1
556 Lactobacillus hominis DSM 23910 = CRBIP 24.179 GCA_000296835.1
440 Lactobacillus iners DSM 13335 GCA_000160875.1
FK
962 Streptococcus mutans UA159 GCA_000007465.2
720 Streptococcus ratti FA-1 = DSM 20564 GCA_000286075.1
689 Streptococcus gallolyticus subsp. gallolyticus DSM 16831 GCA_002000985.1
FL
742 Lactobacillus gasseri ATCC 33323 = JCM 1131 GCA_000014425.1
655 Lactobacillus hominis DSM 23910 = CRBIP 24.179 GCA_000296835.1
512 Lactobacillus jensenii GCA_001936235.1
FM
819 Staphylococcus epidermidis ATCC 12228 GCA_000007645.1
786 Staphylococcus capitis subsp. capitis GCA_001028645.1
718 Staphylococcus warneri SG1 GCA_000332735.1
FN
661 Deinococcus radiodurans R1 GCA_000008565.1
352 Deinococcus gobiensis I-0 GCA_000252445.1
344 Deinococcus deserti VCD115 GCA_000020685.1
FO
485 Rhodobacter sphaeroides 2.4.1 GCA_000012905.2
468 Rhodobacter sphaeroides ATCC 17025 GCA_000016405.1
406 Defluviimonas alba GCA_001620265.1
FP
736 Staphylococcus epidermidis ATCC 12228 GCA_000007645.1
667 Staphylococcus capitis subsp. capitis GCA_001028645.1
626 Staphylococcus simulans GCA_001559115.1
FQ
661 Rhodobacter sphaeroides 2.4.1 GCA_000012905.2
607 Rhodobacter sphaeroides ATCC 17025 GCA_000016405.1
524 Gemmobacter aquatilis GCA_900110025.1
FR
678 Deinococcus radiodurans R1 GCA_000008565.1
421 Deinococcus gobiensis I-0 GCA_000252445.1
408 Deinococcus deserti VCD115 GCA_000020685.1
FS
894 Lactobacillus gasseri ATCC 33323 = JCM 1131 GCA_000014425.1
829 Lactobacillus hominis DSM 23910 = CRBIP 24.179 GCA_000296835.1
656 Lactobacillus jensenii GCA_001936235.1
FT
323 Rhodobacter sphaeroides 2.4.1 GCA_000012905.2
299 Rhodobacter sphaeroides ATCC 17025 GCA_000016405.1
234 Rhodobacter capsulatus SB 1003 GCA_000021865.1
FU
460 Clostridium beijerinckii GCA_000833105.2
437 Clostridium saccharoperbutylacetonicum N1-4_28HMT_29 GCA_000340885.1
412 Clostridium butyricum GCA_001456065.2
FV
629 Escherichia coli O157_3AH7 str. Sakai GCA_000008865.1
629 Escherichia coli UMN026 GCA_000026325.2
629 Escherichia coli str. K-12 substr. MG1655 GCA_000005845.2
628 Escherichia coli O83_3AH1 str. NRG 857C GCA_000183345.1
624 Escherichia coli IAI39 GCA_000026345.1
FW

426 Bifidobacterium adolescentis ATCC 15703 GCA_000010425.1
295 Bifidobacterium callitrichos DSM 23973 GCA_000741175.1
293 Bifidobacterium angulatum DSM 20098 = JCM 7096 GCA_001025155.1
FX
674 Streptococcus mutans UA159 GCA_000007465.2
478 Streptococcus ratti FA-1 = DSM 20564 GCA_000286075.1
397 Streptococcus gordonii str. Challis substr. CH1 GCA_000017005.1
FY
765 Clostridium beijerinckii GCA_000833105.2
737 Clostridium saccharoperbutylacetonicum N1-4_28HMT_29 GCA_000340885.1
708 Clostridium saccharobutylicum DSM 13864 GCA_000473995.1
FZ
822 Streptococcus mutans UA159 GCA_000007465.2
640 Streptococcus equinus GCA_000964315.1
637 Streptococcus gallolyticus subsp. gallolyticus DSM 16831 GCA_002000985.1
G0
800 Enterococcus faecalis V583 GCA_000007785.1
753 Streptomyces cinnamoneus GCA_001885705.1
712 Enterococcus rivorum GCA_001742285.1
G1
715 Staphylococcus epidermidis ATCC 12228 GCA_000007645.1
704 Staphylococcus capitis subsp. capitis GCA_001028645.1
672 Staphylococcus warneri SG1 GCA_000332735.1
G2
661 Lactobacillus gasseri ATCC 33323 = JCM 1131 GCA_000014425.1
656 Lactobacillus hominis DSM 23910 = CRBIP 24.179 GCA_000296835.1
463 Lactobacillus iners DSM 13335 GCA_000160875.1
G3
611 Clostridium beijerinckii GCA_000833105.2
598 Clostridium saccharoperbutylacetonicum N1-4_28HMT_29 GCA_000340885.1
585 Clostridium puniceum GCA_002006345.1
G4
724 Clostridium beijerinckii GCA_000833105.2
669 Clostridium saccharoperbutylacetonicum N1-4_28HMT_29 GCA_000340885.1
657 Clostridium puniceum GCA_002006345.1
G5
443 Rhodobacter sphaeroides 2.4.1 GCA_000012905.2
360 Rhodobacter sphaeroides ATCC 17025 GCA_000016405.1
349 Gemmobacter aquatilis GCA_900110025.1
G6
678 Clostridium beijerinckii GCA_000833105.2
619 Clostridium puniceum GCA_002006345.1
615 Clostridium saccharoperbutylacetonicum N1-4_28HMT_29 GCA_000340885.1
G7
780 Staphylococcus epidermidis ATCC 12228 GCA_000007645.1
765 Staphylococcus capitis subsp. capitis GCA_001028645.1
706 Staphylococcus hominis subsp. hominis C80 GCA_000183685.1
G8
727 Staphylococcus epidermidis ATCC 12228 GCA_000007645.1
712 Staphylococcus capitis subsp. capitis GCA_001028645.1
641 Staphylococcus warneri SG1 GCA_000332735.1
G9
762 Lactobacillus gasseri ATCC 33323 = JCM 1131 GCA_000014425.1
676 Lactobacillus hominis DSM 23910 = CRBIP 24.179 GCA_000296835.1
487 Lactobacillus helveticus GCA_001308285.1
Ga
674 Bifidobacterium adolescentis ATCC 15703 GCA_000010425.1
520 Bifidobacterium dentium JCM 1195 = DSM 20436 GCA_001042595.1
484 Bifidobacterium longum NCC2705 GCA_000007525.1
Gb
683 Lactobacillus gasseri ATCC 33323 = JCM 1131 GCA_000014425.1
606 Lactobacillus hominis DSM 23910 = CRBIP 24.179 GCA_000296835.1
462 Lactobacillus psittaci DSM 15354 GCA_000425905.1
Gc
719 Rhodobacter sphaeroides 2.4.1 GCA_000012905.2
633 Rhodobacter sphaeroides ATCC 17025 GCA_000016405.1
567 Gemmobacter aquatilis GCA_900110025.1
Gd
853 Clostridium beijerinckii GCA_000833105.2
843 Clostridium saccharoperbutylacetonicum N1-4_28HMT_29 GCA_000340885.1
781 Clostridium puniceum GCA_002006345.1
Ge
689 Lactobacillus gasseri ATCC 33323 = JCM 1131 GCA_000014425.1
646 Lactobacillus hominis DSM 23910 = CRBIP 24.179 GCA_000296835.1
508 Lactobacillus iners DSM 13335 GCA_000160875.1
Gf

714 Rhodobacter sphaeroides 2.4.1 GCA_000012905.2
655 Rhodobacter sphaeroides ATCC 17025 GCA_000016405.1
560 Gemmobacter aquatilis GCA_900110025.1
Gg
819 Lactobacillus gasseri ATCC 33323 = JCM 1131 GCA_000014425.1
743 Lactobacillus hominis DSM 23910 = CRBIP 24.179 GCA_000296835.1
566 Lactobacillus psittaci DSM 15354 GCA_000425905.1
Gh
589 Lactobacillus gasseri ATCC 33323 = JCM 1131 GCA_000014425.1
540 Lactobacillus hominis DSM 23910 = CRBIP 24.179 GCA_000296835.1
431 Lactobacillus iners DSM 13335 GCA_000160875.1
Gi
677 Lactobacillus gasseri ATCC 33323 = JCM 1131 GCA_000014425.1
586 Lactobacillus hominis DSM 23910 = CRBIP 24.179 GCA_000296835.1
453 Lactobacillus hamsteri DSM 5661 = JCM 6256 GCA_000615445.1
Gj
652 Bacillus thuringiensis YBT-1518 GCA_000497525.2
638 Bacillus anthracis str. Ames GCA_000007845.1
638 Bacillus anthracis str. Sterne GCA_000008165.1
638 Bacillus cereus ATCC 14579 GCA_000007825.1
638 _5BBacillus thuringiensis_5D serovar konkukian str. 97-27 GCA_000008505.1
611 Bacillus pseudomycoides DSM 12442 GCA_000161455.1
Gk
630 Streptococcus mutans UA159 GCA_000007465.2
476 Streptococcus ratti FA-1 = DSM 20564 GCA_000286075.1
451 Streptococcus gordonii str. Challis substr. CH1 GCA_000017005.1
451 Streptococcus macacae NCTC 11558 GCA_000187995.3
Gl
422 Lactobacillus gasseri ATCC 33323 = JCM 1131 GCA_000014425.1
407 Lactobacillus hominis DSM 23910 = CRBIP 24.179 GCA_000296835.1
317 Lactobacillus iners DSM 13335 GCA_000160875.1
Gm
325 Staphylococcus epidermidis ATCC 12228 GCA_000007645.1
277 Staphylococcus capitis subsp. capitis GCA_001028645.1
269 Staphylococcus hominis subsp. hominis C80 GCA_000183685.1
269 Staphylococcus lugdunensis HKU09-01 GCA_000025085.1
Gn
343 Deinococcus radiodurans R1 GCA_000008565.1
189 Deinococcus gobiensis I-0 GCA_000252445.1
187 Deinococcus deserti VCD115 GCA_000020685.1
Go
596 Staphylococcus epidermidis ATCC 12228 GCA_000007645.1
552 Staphylococcus capitis subsp. capitis GCA_001028645.1
509 Staphylococcus pettenkoferi GCA_002208805.1
Gp
709 Clostridium beijerinckii GCA_000833105.2
667 Clostridium saccharoperbutylacetonicum N1-4_28HMT_29 GCA_000340885.1
652 Clostridium puniceum GCA_002006345.1
Gq
672 Streptococcus mutans UA159 GCA_000007465.2
427 Streptococcus ratti FA-1 = DSM 20564 GCA_000286075.1
406 Streptococcus gordonii str. Challis substr. CH1 GCA_000017005.1
Gr
491 Deinococcus radiodurans R1 GCA_000008565.1
306 Deinococcus deserti VCD115 GCA_000020685.1
296 Deinococcus puniceus GCA_001644565.1
Gs
755 Lactobacillus gasseri ATCC 33323 = JCM 1131 GCA_000014425.1
687 Lactobacillus hominis DSM 23910 = CRBIP 24.179 GCA_000296835.1
557 Lactobacillus iners DSM 13335 GCA_000160875.1
Gt
875 Enterococcus faecalis V583 GCA_000007785.1
828 Streptomyces cinnamoneus GCA_001885705.1
737 Enterococcus rivorum GCA_001742285.1
Gu
723 Deinococcus radiodurans R1 GCA_000008565.1
454 Deinococcus gobiensis I-0 GCA_000252445.1
429 Deinococcus deserti VCD115 GCA_000020685.1
Gv
804 Escherichia coli O157_3AH7 str. Sakai GCA_000008865.1
804 Escherichia coli str. K-12 substr. MG1655 GCA_000005845.2
804 Shigella flexneri 2a str. 301 GCA_000006925.2
797 Escherichia coli O83_3AH1 str. NRG 857C GCA_000183345.1
794 Escherichia coli UMN026 GCA_000026325.2
Gw
628 Rhodobacter sphaeroides 2.4.1 GCA_000012905.2

552 Rhodobacter sphaeroides ATCC 17025 GCA_000016405.1
532 Gemmobacter aquatilis GCA_900110025.1
Gx
766 Rhodobacter sphaeroides 2.4.1 GCA_000012905.2
677 Rhodobacter sphaeroides ATCC 17025 GCA_000016405.1
659 Gemmobacter megaterium GCA_900156815.1
Gy
552 Clostridium beijerinckii GCA_000833105.2
528 Clostridium saccharoperbutylacetonicum N1-4_28HMT_29 GCA_000340885.1
501 Clostridium puniceum GCA_002006345.1
Gz
926 Streptococcus mutans UA159 GCA_000007465.2
674 Streptococcus gordonii str. Challis substr. CH1 GCA_000017005.1
666 Streptococcus ratti FA-1 = DSM 20564 GCA_000286075.1
GA
751 Rhodobacter sphaeroides 2.4.1 GCA_000012905.2
695 Rhodobacter sphaeroides ATCC 17025 GCA_000016405.1
581 Gemmobacter aquatilis GCA_900110025.1
GB
914 Enterococcus faecalis V583 GCA_000007785.1
867 Streptomyces cinnamoneus GCA_001885705.1
770 Enterococcus rivorum GCA_001742285.1
GC
923 Staphylococcus epidermidis ATCC 12228 GCA_000007645.1
875 Staphylococcus capitis subsp. capitis GCA_001028645.1
809 Staphylococcus hominis subsp. hominis C80 GCA_000183685.1
GD
734 Staphylococcus epidermidis ATCC 12228 GCA_000007645.1
668 Staphylococcus capitis subsp. capitis GCA_001028645.1
645 Staphylococcus aureus subsp. aureus NCTC 8325 GCA_000013425.1
GE
856 Deinococcus radiodurans R1 GCA_000008565.1
566 Deinococcus deserti VCD115 GCA_000020685.1
553 Deinococcus soli Cha et al. 2016 GCA_001007995.1
GF
504 Lactobacillus gasseri ATCC 33323 = JCM 1131 GCA_000014425.1
417 Lactobacillus hominis DSM 23910 = CRBIP 24.179 GCA_000296835.1
318 Lactobacillus kalixensis DSM 16043 GCA_001434335.1
GG
633 Streptococcus mutans UA159 GCA_000007465.2
440 Streptococcus ratti FA-1 = DSM 20564 GCA_000286075.1
378 Streptococcus gordonii str. Challis substr. CH1 GCA_000017005.1
GH
786 Rhodobacter sphaeroides 2.4.1 GCA_000012905.2
737 Rhodobacter sphaeroides ATCC 17025 GCA_000016405.1
561 Gemmobacter aquatilis GCA_900110025.1
561 Pseudorhodobacter ferrugineus DSM 5888 GCA_000420745.1
GI
738 Lactobacillus gasseri ATCC 33323 = JCM 1131 GCA_000014425.1
672 Lactobacillus hominis DSM 23910 = CRBIP 24.179 GCA_000296835.1
528 Lactobacillus iners DSM 13335 GCA_000160875.1
GJ
834 Staphylococcus epidermidis ATCC 12228 GCA_000007645.1
807 Staphylococcus capitis subsp. capitis GCA_001028645.1
763 Staphylococcus hominis subsp. hominis C80 GCA_000183685.1
GK
658 Rhodobacter sphaeroides 2.4.1 GCA_000012905.2
582 Rhodobacter sphaeroides ATCC 17025 GCA_000016405.1
534 Gemmobacter megaterium GCA_900156815.1
GL
459 Rhodobacter sphaeroides 2.4.1 GCA_000012905.2
401 Rhodobacter sphaeroides ATCC 17025 GCA_000016405.1
332 Gemmobacter aquatilis GCA_900110025.1
GM
915 Escherichia coli str. K-12 substr. MG1655 GCA_000005845.2
911 Escherichia coli IAI39 GCA_000026345.1
911 Escherichia coli O157_3AH7 str. Sakai GCA_000008865.1
908 Escherichia coli UMN026 GCA_000026325.2
GN
713 Deinococcus radiodurans R1 GCA_000008565.1
500 Deinococcus hopiensis KR-140 GCA_900176165.1
474 Deinococcus deserti VCD115 GCA_000020685.1
GO
522 Enterococcus faecalis V583 GCA_000007785.1
475 Streptomyces cinnamoneus GCA_001885705.1
446 Enterococcus faecium DO GCA_000174395.2

GP
732 Deinococcus radiodurans R1 GCA_000008565.1
451 Deinococcus soli Cha et al. 2016 GCA_001007995.1
444 Deinococcus deserti VCD115 GCA_000020685.1
GQ
526 Streptococcus mutans UA159 GCA_000007465.2
406 Streptococcus sobrinus DSM 20742 = ATCC 33478 GCA_000686605.1
397 Streptococcus ratti FA-1 = DSM 20564 GCA_000286075.1
GR
354 Bifidobacterium adolescentis ATCC 15703 GCA_000010425.1
290 Bifidobacterium dentium JCM 1195 = DSM 20436 GCA_001042595.1
241 Bifidobacterium thermophilum GCA_000741495.1
GS
504 Clostridium beijerinckii GCA_000833105.2
471 Clostridium saccharoperbutylacetonicum N1-4_28HMT_29 GCA_000340885.1
439 Clostridium butyricum GCA_001456065.2
GT
773 Rhodobacter sphaeroides 2.4.1 GCA_000012905.2
733 Rhodobacter sphaeroides ATCC 17025 GCA_000016405.1
635 Gemmobacter aquatilis GCA_900110025.1
GU
844 Streptococcus mutans UA159 GCA_000007465.2
643 Streptococcus gordonii str. Challis substr. CH1 GCA_000017005.1
623 Streptococcus ratti FA-1 = DSM 20564 GCA_000286075.1
GV
372 Streptococcus mutans UA159 GCA_000007465.2
304 Streptococcus ratti FA-1 = DSM 20564 GCA_000286075.1
251 Streptococcus macacae NCTC 11558 GCA_000187995.3
GW
739 Staphylococcus epidermidis ATCC 12228 GCA_000007645.1
682 Staphylococcus capitis subsp. capitis GCA_001028645.1
641 Staphylococcus haemolyticus JCSC1435 GCA_000009865.1
641 Staphylococcus warneri SG1 GCA_000332735.1
GX
676 Enterococcus faecalis V583 GCA_000007785.1
629 Streptomyces cinnamoneus GCA_001885705.1
620 Enterococcus asini ATCC 700915 GCA_000407365.1
GY
677 Bifidobacterium adolescentis ATCC 15703 GCA_000010425.1
549 Bifidobacterium dentium JCM 1195 = DSM 20436 GCA_001042595.1
543 Bifidobacterium breve DSM 20213 = JCM 1192 GCA_001025175.1
GZ
644 Staphylococcus epidermidis ATCC 12228 GCA_000007645.1
635 Staphylococcus capitis subsp. capitis GCA_001028645.1
589 Staphylococcus aureus subsp. aureus NCTC 8325 GCA_000013425.1
H0
633 Lactobacillus gasseri ATCC 33323 = JCM 1131 GCA_000014425.1
575 Lactobacillus hominis DSM 23910 = CRBIP 24.179 GCA_000296835.1
428 Lactobacillus iners DSM 13335 GCA_000160875.1
H1
852 Rhodobacter sphaeroides 2.4.1 GCA_000012905.2
775 Rhodobacter sphaeroides ATCC 17025 GCA_000016405.1
616 Pseudorhodobacter ferrugineus DSM 5888 GCA_000420745.1
H2
733 Clostridium beijerinckii GCA_000833105.2
733 Clostridium saccharoperbutylacetonicum N1-4_28HMT_29 GCA_000340885.1
702 Clostridium saccharobutylicum DSM 13864 GCA_000473995.1
674 Clostridium puniceum GCA_002006345.1
H3
588 Deinococcus radiodurans R1 GCA_000008565.1
391 Deinococcus deserti VCD115 GCA_000020685.1
385 Deinococcus hopiensis KR-140 GCA_900176165.1
H4
790 Bacillus thuringiensis YBT-1518 GCA_000497525.2
777 Bacillus anthracis str. Ames GCA_000007845.1
777 Bacillus anthracis str. Sterne GCA_000008165.1
777 _5BBacillus thuringiensis_5D serovar konkukian str. 97-27 GCA_000008505.1
776 Bacillus cereus ATCC 14579 GCA_000007825.1
H5
717 Streptococcus mutans UA159 GCA_000007465.2
428 Streptococcus ratti FA-1 = DSM 20564 GCA_000286075.1
415 Streptococcus equinus GCA_000964315.1
H6
545 Clostridium saccharoperbutylacetonicum N1-4_28HMT_29 GCA_000340885.1
538 Clostridium beijerinckii GCA_000833105.2
516 Clostridium saccharobutylicum DSM 13864 GCA_000473995.1

H7
665 Lactobacillus gasseri ATCC 33323 = JCM 1131 GCA_000014425.1
636 Lactobacillus hominis DSM 23910 = CRBIP 24.179 GCA_000296835.1
485 Lactobacillus iners DSM 13335 GCA_000160875.1
H8
408 Clostridium beijerinckii GCA_000833105.2
404 Clostridium saccharobutylicum DSM 13864 GCA_000473995.1
393 Clostridium saccharoperbutylacetonicum N1-4_28HMT_29 GCA_000340885.1
H9
798 Staphylococcus epidermidis ATCC 12228 GCA_000007645.1
749 Staphylococcus capitis subsp. capitis GCA_001028645.1
717 Staphylococcus haemolyticus JCSC1435 GCA_000009865.1
Ha
781 Streptococcus mutans UA159 GCA_000007465.2
579 Streptococcus ratti FA-1 = DSM 20564 GCA_000286075.1
563 Streptococcus gordonii str. Challis substr. CH1 GCA_000017005.1
Hb
774 Lactobacillus gasseri ATCC 33323 = JCM 1131 GCA_000014425.1
717 Lactobacillus hominis DSM 23910 = CRBIP 24.179 GCA_000296835.1
555 Lactobacillus iners DSM 13335 GCA_000160875.1
Hc
561 Streptococcus mutans UA159 GCA_000007465.2
495 Streptococcus gordonii str. Challis substr. CH1 GCA_000017005.1
461 Streptococcus salivarius GCA_000785515.1
461 Streptococcus thermophilus JIM 8232 GCA_000253395.1
Hd
688 Rhodobacter sphaeroides 2.4.1 GCA_000012905.2
595 Rhodobacter sphaeroides ATCC 17025 GCA_000016405.1
510 Defluviimonas alba GCA_001620265.1
He
747 Bacillus anthracis str. Ames GCA_000007845.1
747 Bacillus anthracis str. Sterne GCA_000008165.1
747 _5BBacillus thuringiensis_5D serovar konkukian str. 97-27 GCA_000008505.1
732 Bacillus pseudomycoides DSM 12442 GCA_000161455.1
722 Bacillus cereus ATCC 14579 GCA_000007825.1
722 Bacillus thuringiensis YBT-1518 GCA_000497525.2
Hf
749 Streptococcus mutans UA159 GCA_000007465.2
583 Streptococcus ratti FA-1 = DSM 20564 GCA_000286075.1
571 Streptococcus gordonii str. Challis substr. CH1 GCA_000017005.1
Hg
619 Streptococcus mutans UA159 GCA_000007465.2
398 Streptococcus halotolerans GCA_001598035.1
392 Streptococcus sobrinus DSM 20742 = ATCC 33478 GCA_000686605.1
Hh
716 Bifidobacterium adolescentis ATCC 15703 GCA_000010425.1
614 Bifidobacterium angulatum DSM 20098 = JCM 7096 GCA_001025155.1
589 Bifidobacterium stellenboschense GCA_000741785.1
Hi
821 Enterococcus faecalis V583 GCA_000007785.1
774 Streptomyces cinnamoneus GCA_001885705.1
733 Enterococcus phoeniculicola ATCC BAA-412 GCA_000407505.1
Hj
466 Enterococcus faecalis V583 GCA_000007785.1
434 Streptomyces cinnamoneus GCA_001885705.1
386 Enterococcus faecium DO GCA_000174395.2
Hk
502 Rhodobacter sphaeroides 2.4.1 GCA_000012905.2
476 Rhodobacter sphaeroides ATCC 17025 GCA_000016405.1
437 Thioclava dalianensis GCA_000715505.1
437 Thioclava indica GCA_000714545.1
Hl
786 _5BBacillus thuringiensis_5D serovar konkukian str. 97-27 GCA_000008505.1
785 Bacillus anthracis str. Ames GCA_000007845.1
785 Bacillus anthracis str. Sterne GCA_000008165.1
769 Bacillus pseudomycoides DSM 12442 GCA_000161455.1
Hm
698 Lactobacillus gasseri ATCC 33323 = JCM 1131 GCA_000014425.1
633 Lactobacillus hominis DSM 23910 = CRBIP 24.179 GCA_000296835.1
524 Lactobacillus iners DSM 13335 GCA_000160875.1
Hn
673 Clostridium beijerinckii GCA_000833105.2
635 Clostridium saccharoperbutylacetonicum N1-4_28HMT_29 GCA_000340885.1
609 Clostridium puniceum GCA_002006345.1
Ho
882 Deinococcus radiodurans R1 GCA_000008565.1

609 Deinococcus deserti VCD115 GCA_000020685.1
591 Deinococcus gobiensis I-0 GCA_000252445.1
Hp
697 Clostridium beijerinckii GCA_000833105.2
676 Clostridium saccharoperbutylacetonicum N1-4_28HMT_29 GCA_000340885.1
661 Clostridium saccharobutylicum DSM 13864 GCA_000473995.1
Hq
621 Staphylococcus epidermidis ATCC 12228 GCA_000007645.1
568 Staphylococcus hominis subsp. hominis C80 GCA_000183685.1
567 Staphylococcus capitis subsp. capitis GCA_001028645.1
Hr
731 Escherichia coli IAI39 GCA_000026345.1
724 Escherichia coli O157_3AH7 str. Sakai GCA_000008865.1
724 Escherichia coli str. K-12 substr. MG1655 GCA_000005845.2
722 Shigella dysenteriae Sd197 GCA_000012005.1
Hs
806 Rhodobacter sphaeroides 2.4.1 GCA_000012905.2
711 Rhodobacter sphaeroides ATCC 17025 GCA_000016405.1
574 Pseudorhodobacter wandonensis GCA_001202035.1
Ht
601 Streptococcus mutans UA159 GCA_000007465.2
459 Streptococcus ratti FA-1 = DSM 20564 GCA_000286075.1
412 Streptococcus equinus GCA_000964315.1
Hu
777 Lactobacillus gasseri ATCC 33323 = JCM 1131 GCA_000014425.1
706 Lactobacillus hominis DSM 23910 = CRBIP 24.179 GCA_000296835.1
557 Lactobacillus iners DSM 13335 GCA_000160875.1
Hv
620 Lactobacillus gasseri ATCC 33323 = JCM 1131 GCA_000014425.1
592 Lactobacillus hominis DSM 23910 = CRBIP 24.179 GCA_000296835.1
437 Lactobacillus iners DSM 13335 GCA_000160875.1
Hw
716 Escherichia coli str. K-12 substr. MG1655 GCA_000005845.2
704 Shigella flexneri 2a str. 301 GCA_000006925.2
698 Escherichia coli O104_3AH4 str. 2011C-3493 GCA_000299455.1
Hx
825 Escherichia coli O104_3AH4 str. 2011C-3493 GCA_000299455.1
825 Escherichia coli str. K-12 substr. MG1655 GCA_000005845.2
791 Shigella flexneri 2a str. 301 GCA_000006925.2
784 Escherichia coli O157_3AH7 str. Sakai GCA_000008865.1
Hy
571 Staphylococcus epidermidis ATCC 12228 GCA_000007645.1
526 Staphylococcus capitis subsp. capitis GCA_001028645.1
506 Staphylococcus lugdunensis HKU09-01 GCA_000025085.1
Hz
468 Enterococcus faecalis V583 GCA_000007785.1
422 Streptomyces cinnamoneus GCA_001885705.1
374 Enterococcus rivorum GCA_001742285.1
HA
674 Deinococcus radiodurans R1 GCA_000008565.1
461 Deinococcus gobiensis I-0 GCA_000252445.1
455 Deinococcus deserti VCD115 GCA_000020685.1
HB
897 Clostridium beijerinckii GCA_000833105.2
841 Clostridium saccharoperbutylacetonicum N1-4_28HMT_29 GCA_000340885.1
804 Clostridium saccharobutylicum DSM 13864 GCA_000473995.1
HC
431 Enterococcus faecalis V583 GCA_000007785.1
428 Enterococcus rivorum GCA_001742285.1
414 Enterococcus canis NBRC 100695 GCA_001544375.1
414 Enterococcus dispar ATCC 51266 GCA_000406945.1
HD
592 Deinococcus radiodurans R1 GCA_000008565.1
359 Deinococcus deserti VCD115 GCA_000020685.1
345 Deinococcus gobiensis I-0 GCA_000252445.1
HE
897 Bifidobacterium adolescentis ATCC 15703 GCA_000010425.1
782 Bifidobacterium dentium JCM 1195 = DSM 20436 GCA_001042595.1
737 Bifidobacterium stellenboschense GCA_000741785.1
HF
720 Bacillus anthracis str. Ames GCA_000007845.1
720 Bacillus anthracis str. Sterne GCA_000008165.1
720 Bacillus cereus ATCC 14579 GCA_000007825.1
720 _5BBacillus thuringiensis_5D serovar konkukian str. 97-27 GCA_000008505.1
706 Bacillus pseudomycoides DSM 12442 GCA_000161455.1
701 Bacillus thuringiensis YBT-1518 GCA_000497525.2

HG
555 Deinococcus radiodurans R1 GCA_000008565.1
309 Deinococcus deserti VCD115 GCA_000020685.1
305 Deinococcus gobiensis I-0 GCA_000252445.1
HH
611 Deinococcus radiodurans R1 GCA_000008565.1
366 Deinococcus deserti VCD115 GCA_000020685.1
329 Deinococcus frigens DSM 12807 GCA_000701425.1
HI
636 Lactobacillus gasseri ATCC 33323 = JCM 1131 GCA_000014425.1
573 Lactobacillus hominis DSM 23910 = CRBIP 24.179 GCA_000296835.1
352 Lactobacillus iners DSM 13335 GCA_000160875.1
HJ
675 Rhodobacter sphaeroides 2.4.1 GCA_000012905.2
634 Rhodobacter sphaeroides ATCC 17025 GCA_000016405.1
478 Gemmobacter aquatilis GCA_900110025.1
HK
858 Clostridium beijerinckii GCA_000833105.2
846 Clostridium saccharoperbutylacetonicum N1-4_28HMT_29 GCA_000340885.1
835 Clostridium saccharobutylicum DSM 13864 GCA_000473995.1
HL
834 Lactobacillus gasseri ATCC 33323 = JCM 1131 GCA_000014425.1
781 Lactobacillus hominis DSM 23910 = CRBIP 24.179 GCA_000296835.1
573 Lactobacillus psittaci DSM 15354 GCA_000425905.1
HM
484 Lactobacillus gasseri ATCC 33323 = JCM 1131 GCA_000014425.1
424 Lactobacillus hominis DSM 23910 = CRBIP 24.179 GCA_000296835.1
313 Lactobacillus iners DSM 13335 GCA_000160875.1
HN
586 Streptococcus mutans UA159 GCA_000007465.2
460 Streptococcus ratti FA-1 = DSM 20564 GCA_000286075.1
417 Streptococcus gordonii str. Challis substr. CH1 GCA_000017005.1
HO
465 Clostridium beijerinckii GCA_000833105.2
450 Clostridium saccharoperbutylacetonicum N1-4_28HMT_29 GCA_000340885.1
447 Clostridium saccharobutylicum DSM 13864 GCA_000473995.1
HP
864 Lactobacillus gasseri ATCC 33323 = JCM 1131 GCA_000014425.1
784 Lactobacillus hominis DSM 23910 = CRBIP 24.179 GCA_000296835.1
607 Lactobacillus iners DSM 13335 GCA_000160875.1
HQ
588 Clostridium beijerinckii GCA_000833105.2
559 Clostridium saccharoperbutylacetonicum N1-4_28HMT_29 GCA_000340885.1
554 Clostridium saccharobutylicum DSM 13864 GCA_000473995.1
HR
875 Streptococcus mutans UA159 GCA_000007465.2
689 Streptococcus ratti FA-1 = DSM 20564 GCA_000286075.1
640 Streptococcus equi subsp. zooepidemicus GCA_000026605.1
640 Streptococcus sobrinus DSM 20742 = ATCC 33478 GCA_000686605.1
HS
624 Deinococcus radiodurans R1 GCA_000008565.1
376 Deinococcus deserti VCD115 GCA_000020685.1
335 Deinococcus hopiensis KR-140 GCA_900176165.1
HT
756 Staphylococcus epidermidis ATCC 12228 GCA_000007645.1
718 Staphylococcus capitis subsp. capitis GCA_001028645.1
645 Staphylococcus haemolyticus JCSC1435 GCA_000009865.1
HU
545 Clostridium beijerinckii GCA_000833105.2
540 Clostridium saccharoperbutylacetonicum N1-4_28HMT_29 GCA_000340885.1
523 Clostridium puniceum GCA_002006345.1
HV
693 Clostridium beijerinckii GCA_000833105.2
671 Clostridium saccharoperbutylacetonicum N1-4_28HMT_29 GCA_000340885.1
655 Clostridium puniceum GCA_002006345.1
HW
794 Rhodobacter sphaeroides 2.4.1 GCA_000012905.2
734 Rhodobacter sphaeroides ATCC 17025 GCA_000016405.1
646 Gemmobacter aquatilis GCA_900110025.1
HX
582 Clostridium beijerinckii GCA_000833105.2
553 Clostridium saccharoperbutylacetonicum N1-4_28HMT_29 GCA_000340885.1
534 Clostridium puniceum GCA_002006345.1
HY
841 Bacillus thuringiensis YBT-1518 GCA_000497525.2
824 Bacillus cereus ATCC 14579 GCA_000007825.1

822 Bacillus anthracis str. Ames GCA_000007845.1
822 Bacillus anthracis str. Sterne GCA_000008165.1
822 _5BBacillus thuringiensis_5D serovar konkukian str. 97-27 GCA_000008505.1
HZ
671 Staphylococcus lugdunensis HKU09-01 GCA_000025085.1
665 Staphylococcus epidermidis ATCC 12228 GCA_000007645.1
631 Staphylococcus haemolyticus JCSC1435 GCA_000009865.1
631 Staphylococcus hominis subsp. hominis C80 GCA_000183685.1
I0
807 Streptococcus mutans UA159 GCA_000007465.2
569 Streptococcus ratti FA-1 = DSM 20564 GCA_000286075.1
520 Streptococcus equinus GCA_000964315.1
520 Streptococcus gordonii str. Challis substr. CH1 GCA_000017005.1
I1
766 Bifidobacterium adolescentis ATCC 15703 GCA_000010425.1
610 Bifidobacterium dentium JCM 1195 = DSM 20436 GCA_001042595.1
575 Bifidobacterium callitrichos DSM 23973 GCA_000741175.1
I2
990 Staphylococcus epidermidis ATCC 12228 GCA_000007645.1
955 Staphylococcus capitis subsp. capitis GCA_001028645.1
864 Staphylococcus warneri SG1 GCA_000332735.1
I3
732 Deinococcus radiodurans R1 GCA_000008565.1
438 Deinococcus gobiensis I-0 GCA_000252445.1
425 Deinococcus deserti VCD115 GCA_000020685.1
I4
890 Clostridium beijerinckii GCA_000833105.2
867 Clostridium saccharoperbutylacetonicum N1-4_28HMT_29 GCA_000340885.1
846 Clostridium saccharobutylicum DSM 13864 GCA_000473995.1
I5
729 Bacillus anthracis str. Ames GCA_000007845.1
729 Bacillus anthracis str. Sterne GCA_000008165.1
729 Bacillus cereus ATCC 14579 GCA_000007825.1
729 _5BBacillus thuringiensis_5D serovar konkukian str. 97-27 GCA_000008505.1
718 Bacillus pseudomycoides DSM 12442 GCA_000161455.1
716 Bacillus thuringiensis YBT-1518 GCA_000497525.2
I6
758 Clostridium beijerinckii GCA_000833105.2
742 Clostridium saccharoperbutylacetonicum N1-4_28HMT_29 GCA_000340885.1
735 Clostridium puniceum GCA_002006345.1
I7
892 Lactobacillus gasseri ATCC 33323 = JCM 1131 GCA_000014425.1
829 Lactobacillus hominis DSM 23910 = CRBIP 24.179 GCA_000296835.1
616 Lactobacillus iners DSM 13335 GCA_000160875.1
I8
666 Rhodobacter sphaeroides 2.4.1 GCA_000012905.2
597 Rhodobacter sphaeroides ATCC 17025 GCA_000016405.1
464 Pseudorhodobacter psychrotolerans GCA_001294535.1
I9
539 Lactobacillus gasseri ATCC 33323 = JCM 1131 GCA_000014425.1
494 Lactobacillus hominis DSM 23910 = CRBIP 24.179 GCA_000296835.1
404 Lactobacillus iners DSM 13335 GCA_000160875.1
Ia
786 Staphylococcus epidermidis ATCC 12228 GCA_000007645.1
745 Staphylococcus capitis subsp. capitis GCA_001028645.1
699 Staphylococcus warneri SG1 GCA_000332735.1
Ib
443 Bifidobacterium adolescentis ATCC 15703 GCA_000010425.1
297 Bifidobacterium dentium JCM 1195 = DSM 20436 GCA_001042595.1
296 Bifidobacterium pseudolongum PV8-2 GCA_000800475.2
Ic
668 Clostridium beijerinckii GCA_000833105.2
618 Clostridium saccharoperbutylacetonicum N1-4_28HMT_29 GCA_000340885.1
603 Clostridium puniceum GCA_002006345.1
Id
651 Escherichia coli O157_3AH7 str. Sakai GCA_000008865.1
651 Escherichia coli str. K-12 substr. MG1655 GCA_000005845.2
648 Escherichia coli UMN026 GCA_000026325.2
620 Escherichia coli O83_3AH1 str. NRG 857C GCA_000183345.1
Ie
592 Streptococcus mutans UA159 GCA_000007465.2
455 Streptococcus ratti FA-1 = DSM 20564 GCA_000286075.1
428 Streptococcus equinus GCA_000964315.1
If
630 Deinococcus radiodurans R1 GCA_000008565.1
415 Deinococcus deserti VCD115 GCA_000020685.1

392 Deinococcus proteolyticus MRP GCA_000190555.1
Ig
744 Streptococcus mutans UA159 GCA_000007465.2
528 Streptococcus gordonii str. Challis substr. CH1 GCA_000017005.1
527 Streptococcus ratti FA-1 = DSM 20564 GCA_000286075.1
Ih
725 Escherichia coli O104_3AH4 str. 2011C-3493 GCA_000299455.1
725 Escherichia coli str. K-12 substr. MG1655 GCA_000005845.2
714 Escherichia coli O157_3AH7 str. Sakai GCA_000008865.1
711 Escherichia coli O83_3AH1 str. NRG 857C GCA_000183345.1
Ii
749 Rhodobacter sphaeroides 2.4.1 GCA_000012905.2
673 Rhodobacter sphaeroides ATCC 17025 GCA_000016405.1
548 Gemmobacter aquatilis GCA_900110025.1
Ij
602 Escherichia coli str. K-12 substr. MG1655 GCA_000005845.2
595 Escherichia coli O104_3AH4 str. 2011C-3493 GCA_000299455.1
587 Shigella flexneri 2a str. 301 GCA_000006925.2
Ik
534 Rhodobacter sphaeroides 2.4.1 GCA_000012905.2
462 Rhodobacter sphaeroides ATCC 17025 GCA_000016405.1
395 Pseudorhodobacter ferrugineus DSM 5888 GCA_000420745.1
Il
399 Rhodobacter sphaeroides 2.4.1 GCA_000012905.2
361 Rhodobacter sphaeroides ATCC 17025 GCA_000016405.1
320 Gemmobacter aquatilis GCA_900110025.1
Im
588 Lactobacillus gasseri ATCC 33323 = JCM 1131 GCA_000014425.1
535 Lactobacillus hominis DSM 23910 = CRBIP 24.179 GCA_000296835.1
425 Lactobacillus iners DSM 13335 GCA_000160875.1
In
873 Escherichia coli str. K-12 substr. MG1655 GCA_000005845.2
872 Escherichia coli O157_3AH7 str. Sakai GCA_000008865.1
862 Escherichia coli O83_3AH1 str. NRG 857C GCA_000183345.1
Io
753 Lactobacillus gasseri ATCC 33323 = JCM 1131 GCA_000014425.1
753 Lactobacillus hominis DSM 23910 = CRBIP 24.179 GCA_000296835.1
577 Lactobacillus iners DSM 13335 GCA_000160875.1
573 Lactobacillus hamsteri DSM 5661 = JCM 6256 GCA_000615445.1
Ip
593 Rhodobacter sphaeroides 2.4.1 GCA_000012905.2
534 Rhodobacter sphaeroides ATCC 17025 GCA_000016405.1
434 Pseudorhodobacter ferrugineus DSM 5888 GCA_000420745.1
Iq
436 Enterococcus faecalis V583 GCA_000007785.1
389 Streptomyces cinnamoneus GCA_001885705.1
347 Enterococcus faecium DO GCA_000174395.2
Ir
599 Rhodobacter sphaeroides 2.4.1 GCA_000012905.2
528 Rhodobacter sphaeroides ATCC 17025 GCA_000016405.1
460 Pseudorhodobacter ferrugineus DSM 5888 GCA_000420745.1
Is
511 Escherichia coli O104_3AH4 str. 2011C-3493 GCA_000299455.1
511 Escherichia coli str. K-12 substr. MG1655 GCA_000005845.2
505 Escherichia coli IAI39 GCA_000026345.1
505 Escherichia coli UMN026 GCA_000026325.2
504 Shigella flexneri 2a str. 301 GCA_000006925.2
It
403 Clostridium beijerinckii GCA_000833105.2
378 Clostridium saccharobutylicum DSM 13864 GCA_000473995.1
378 Clostridium saccharoperbutylacetonicum N1-4_28HMT_29 GCA_000340885.1
374 Clostridium puniceum GCA_002006345.1
Iu
861 Lactobacillus gasseri ATCC 33323 = JCM 1131 GCA_000014425.1
792 Lactobacillus hominis DSM 23910 = CRBIP 24.179 GCA_000296835.1
577 Lactobacillus iners DSM 13335 GCA_000160875.1
Iv
781 Streptococcus mutans UA159 GCA_000007465.2
576 Streptococcus ratti FA-1 = DSM 20564 GCA_000286075.1
533 Streptococcus gordonii str. Challis substr. CH1 GCA_000017005.1
Iw
729 Bifidobacterium adolescentis ATCC 15703 GCA_000010425.1
647 Bifidobacterium dentium JCM 1195 = DSM 20436 GCA_001042595.1
617 Bifidobacterium angulatum DSM 20098 = JCM 7096 GCA_001025155.1
Ix
651 Staphylococcus epidermidis ATCC 12228 GCA_000007645.1

593 Staphylococcus capitis subsp. capitis GCA_001028645.1
564 Staphylococcus warneri SG1 GCA_000332735.1
Iy
801 Clostridium saccharoperbutylacetonicum N1-4_28HMT_29 GCA_000340885.1
795 Clostridium beijerinckii GCA_000833105.2
780 Clostridium saccharobutylicum DSM 13864 GCA_000473995.1
Iz
753 Clostridium beijerinckii GCA_000833105.2
734 Clostridium saccharoperbutylacetonicum N1-4_28HMT_29 GCA_000340885.1
715 Clostridium puniceum GCA_002006345.1
IA
673 Staphylococcus epidermidis ATCC 12228 GCA_000007645.1
640 Staphylococcus capitis subsp. capitis GCA_001028645.1
630 Staphylococcus lugdunensis HKU09-01 GCA_000025085.1
IB
382 Enterococcus faecalis V583 GCA_000007785.1
382 Streptomyces cinnamoneus GCA_001885705.1
367 Enterococcus canis NBRC 100695 GCA_001544375.1
367 Enterococcus hirae ATCC 9790 GCA_000271405.2
367 Enterococcus rivorum GCA_001742285.1
367 Enterococcus saccharolyticus subsp. saccharolyticus ATCC 43076 GCA_000407285.1
366 Enterococcus casseliflavus EC20 GCA_000157355.2
366 Enterococcus dispar ATCC 51266 GCA_000406945.1
366 Enterococcus gilvus ATCC BAA-350 GCA_000407545.1
366 Enterococcus malodoratus ATCC 43197 GCA_000407185.1
366 Enterococcus pallens ATCC BAA-351 GCA_000407485.1
IC
770 Rhodobacter sphaeroides 2.4.1 GCA_000012905.2
700 Rhodobacter sphaeroides ATCC 17025 GCA_000016405.1
577 Pseudorhodobacter ferrugineus DSM 5888 GCA_000420745.1
ID
529 Clostridium beijerinckii GCA_000833105.2
525 Clostridium saccharoperbutylacetonicum N1-4_28HMT_29 GCA_000340885.1
489 Clostridium puniceum GCA_002006345.1
IE
533 Clostridium beijerinckii GCA_000833105.2
502 Clostridium saccharobutylicum DSM 13864 GCA_000473995.1
500 Clostridium saccharoperbutylacetonicum N1-4_28HMT_29 GCA_000340885.1
IF
466 Lactobacillus gasseri ATCC 33323 = JCM 1131 GCA_000014425.1
442 Lactobacillus hominis DSM 23910 = CRBIP 24.179 GCA_000296835.1
340 Lactobacillus iners DSM 13335 GCA_000160875.1
IG
657 Enterococcus faecalis V583 GCA_000007785.1
611 Streptomyces cinnamoneus GCA_001885705.1
543 Enterococcus thailandicus GCA_001652875.1
IH
699 Escherichia coli O157_3AH7 str. Sakai GCA_000008865.1
699 Escherichia coli str. K-12 substr. MG1655 GCA_000005845.2
689 Escherichia coli O83_3AH1 str. NRG 857C GCA_000183345.1
688 Escherichia coli UMN026 GCA_000026325.2
II
434 Streptococcus mutans UA159 GCA_000007465.2
273 Streptococcus macacae NCTC 11558 GCA_000187995.3
272 Streptococcus ratti FA-1 = DSM 20564 GCA_000286075.1
IJ
869 Streptococcus mutans UA159 GCA_000007465.2
602 Streptococcus gordonii str. Challis substr. CH1 GCA_000017005.1
596 Streptococcus ratti FA-1 = DSM 20564 GCA_000286075.1
IK
560 Staphylococcus epidermidis ATCC 12228 GCA_000007645.1
551 Staphylococcus capitis subsp. capitis GCA_001028645.1
520 Staphylococcus simulans GCA_001559115.1
IL
830 Deinococcus radiodurans R1 GCA_000008565.1
469 Deinococcus gobiensis I-0 GCA_000252445.1
439 Deinococcus deserti VCD115 GCA_000020685.1
IM
839 Bacillus cereus ATCC 14579 GCA_000007825.1
834 Bacillus thuringiensis YBT-1518 GCA_000497525.2
833 Bacillus anthracis str. Ames GCA_000007845.1
833 Bacillus anthracis str. Sterne GCA_000008165.1
833 _5BBacillus thuringiensis_5D serovar konkukian str. 97-27 GCA_000008505.1
IN
728 Clostridium beijerinckii GCA_000833105.2
716 Clostridium saccharoperbutylacetonicum N1-4_28HMT_29 GCA_000340885.1

682 Clostridium butyricum GCA_001456065.2
IO
672 Rhodobacter sphaeroides 2.4.1 GCA_000012905.2
628 Rhodobacter sphaeroides ATCC 17025 GCA_000016405.1
528 Pseudorhodobacter ferrugineus DSM 5888 GCA_000420745.1
528 Pseudorhodobacter wandonensis GCA_001202035.1
IP
949 Staphylococcus epidermidis ATCC 12228 GCA_000007645.1
903 Staphylococcus capitis subsp. capitis GCA_001028645.1
846 Staphylococcus haemolyticus JCSC1435 GCA_000009865.1
IQ
830 Streptococcus mutans UA159 GCA_000007465.2
628 Streptococcus ratti FA-1 = DSM 20564 GCA_000286075.1
613 Streptococcus ferus DSM 20646 GCA_000372425.1
IR
821 Clostridium beijerinckii GCA_000833105.2
800 Clostridium saccharobutylicum DSM 13864 GCA_000473995.1
796 Clostridium saccharoperbutylacetonicum N1-4_28HMT_29 GCA_000340885.1
IS
672 Clostridium beijerinckii GCA_000833105.2
642 Clostridium saccharobutylicum DSM 13864 GCA_000473995.1
633 Clostridium saccharoperbutylacetonicum N1-4_28HMT_29 GCA_000340885.1
IT
410 Rhodobacter sphaeroides 2.4.1 GCA_000012905.2
374 Rhodobacter sphaeroides ATCC 17025 GCA_000016405.1
293 Gemmobacter aquatilis GCA_900110025.1
293 Pseudorhodobacter psychrotolerans GCA_001294535.1
IU
607 Lactobacillus gasseri ATCC 33323 = JCM 1131 GCA_000014425.1
541 Lactobacillus hominis DSM 23910 = CRBIP 24.179 GCA_000296835.1
396 Lactobacillus iners DSM 13335 GCA_000160875.1
IV
765 Enterococcus faecalis V583 GCA_000007785.1
724 Streptomyces cinnamoneus GCA_001885705.1
704 Enterococcus rivorum GCA_001742285.1
IW
724 Bacillus thuringiensis YBT-1518 GCA_000497525.2
713 Bacillus anthracis str. Ames GCA_000007845.1
713 Bacillus anthracis str. Sterne GCA_000008165.1
713 _5BBacillus thuringiensis_5D serovar konkukian str. 97-27 GCA_000008505.1
707 Bacillus cereus ATCC 14579 GCA_000007825.1
IX
701 Deinococcus radiodurans R1 GCA_000008565.1
397 Deinococcus deserti VCD115 GCA_000020685.1
372 Deinococcus hopiensis KR-140 GCA_900176165.1
IY
548 Lactobacillus gasseri ATCC 33323 = JCM 1131 GCA_000014425.1
525 Lactobacillus hominis DSM 23910 = CRBIP 24.179 GCA_000296835.1
451 Lactobacillus iners DSM 13335 GCA_000160875.1
IZ
135 Rhodobacter sphaeroides 2.4.1 GCA_000012905.2
119 Rhodobacter sphaeroides ATCC 17025 GCA_000016405.1
85 Pseudorhodobacter psychrotolerans GCA_001294535.1
85 Pseudorhodobacter wandonensis GCA_001202035.1
J0
830 _5BBacillus thuringiensis_5D serovar konkukian str. 97-27 GCA_000008505.1
829 Bacillus anthracis str. Ames GCA_000007845.1
829 Bacillus anthracis str. Sterne GCA_000008165.1
821 Bacillus cereus ATCC 14579 GCA_000007825.1
J1
802 Streptococcus mutans UA159 GCA_000007465.2
573 Streptococcus ratti FA-1 = DSM 20564 GCA_000286075.1
561 Streptococcus gordonii str. Challis substr. CH1 GCA_000017005.1
J2
779 Deinococcus radiodurans R1 GCA_000008565.1
527 Deinococcus deserti VCD115 GCA_000020685.1
479 Deinococcus gobiensis I-0 GCA_000252445.1
J3
772 Clostridium beijerinckii GCA_000833105.2
735 Clostridium saccharoperbutylacetonicum N1-4_28HMT_29 GCA_000340885.1
726 Clostridium saccharobutylicum DSM 13864 GCA_000473995.1
J4
818 Lactobacillus gasseri ATCC 33323 = JCM 1131 GCA_000014425.1
737 Lactobacillus hominis DSM 23910 = CRBIP 24.179 GCA_000296835.1
544 Lactobacillus psittaci DSM 15354 GCA_000425905.1
J5

614 Staphylococcus epidermidis ATCC 12228 GCA_000007645.1
564 Staphylococcus capitis subsp. capitis GCA_001028645.1
534 Staphylococcus lugdunensis HKU09-01 GCA_000025085.1
J6
699 Clostridium beijerinckii GCA_000833105.2
655 Clostridium saccharoperbutylacetonicum N1-4_28HMT_29 GCA_000340885.1
640 Clostridium puniceum GCA_002006345.1
J7
525 Deinococcus radiodurans R1 GCA_000008565.1
281 Deinococcus hopiensis KR-140 GCA_900176165.1
277 Deinococcus gobiensis I-0 GCA_000252445.1
J8
683 Lactobacillus gasseri ATCC 33323 = JCM 1131 GCA_000014425.1
635 Lactobacillus hominis DSM 23910 = CRBIP 24.179 GCA_000296835.1
487 Lactobacillus iners DSM 13335 GCA_000160875.1
J9
519 Streptococcus mutans UA159 GCA_000007465.2
355 Streptococcus gordonii str. Challis substr. CH1 GCA_000017005.1
353 Streptococcus iniae GCA_000831485.1
Ja
596 Clostridium beijerinckii GCA_000833105.2
562 Clostridium saccharoperbutylacetonicum N1-4_28HMT_29 GCA_000340885.1
530 Clostridium puniceum GCA_002006345.1
Jb
871 Streptococcus mutans UA159 GCA_000007465.2
654 Streptococcus ratti FA-1 = DSM 20564 GCA_000286075.1
653 Streptococcus gordonii str. Challis substr. CH1 GCA_000017005.1
Jc
776 Deinococcus radiodurans R1 GCA_000008565.1
478 Deinococcus deserti VCD115 GCA_000020685.1
416 Deinococcus soli Cha et al. 2016 GCA_001007995.1
Jd
330 Rhodobacter sphaeroides 2.4.1 GCA_000012905.2
309 Rhodobacter sphaeroides ATCC 17025 GCA_000016405.1
227 Gemmobacter aquatilis GCA_900110025.1
Je
582 Streptococcus mutans UA159 GCA_000007465.2
431 Streptococcus ratti FA-1 = DSM 20564 GCA_000286075.1
372 Streptococcus sobrinus DSM 20742 = ATCC 33478 GCA_000686605.1
Jf
673 Deinococcus radiodurans R1 GCA_000008565.1
415 Deinococcus deserti VCD115 GCA_000020685.1
398 Deinococcus gobiensis I-0 GCA_000252445.1
Jg
698 Clostridium beijerinckii GCA_000833105.2
698 Clostridium saccharoperbutylacetonicum N1-4_28HMT_29 GCA_000340885.1
678 Clostridium puniceum GCA_002006345.1
675 Clostridium saccharobutylicum DSM 13864 GCA_000473995.1
Jh
639 Staphylococcus epidermidis ATCC 12228 GCA_000007645.1
612 Staphylococcus capitis subsp. capitis GCA_001028645.1
587 Staphylococcus aureus subsp. aureus NCTC 8325 GCA_000013425.1
Ji
542 Lactobacillus gasseri ATCC 33323 = JCM 1131 GCA_000014425.1
471 Lactobacillus hominis DSM 23910 = CRBIP 24.179 GCA_000296835.1
312 Lactobacillus iners DSM 13335 GCA_000160875.1
Jj
428 Bacillus thuringiensis YBT-1518 GCA_000497525.2
412 Bacillus anthracis str. Ames GCA_000007845.1
412 Bacillus anthracis str. Sterne GCA_000008165.1
412 Bacillus cereus ATCC 14579 GCA_000007825.1
412 _5BBacillus thuringiensis_5D serovar konkukian str. 97-27 GCA_000008505.1
403 Bacillus pseudomycoides DSM 12442 GCA_000161455.1
Jk
800 Lactobacillus gasseri ATCC 33323 = JCM 1131 GCA_000014425.1
772 Lactobacillus hominis DSM 23910 = CRBIP 24.179 GCA_000296835.1
647 Lactobacillus iners DSM 13335 GCA_000160875.1
Jl
631 Bacillus anthracis str. Ames GCA_000007845.1
631 Bacillus anthracis str. Sterne GCA_000008165.1
631 _5BBacillus thuringiensis_5D serovar konkukian str. 97-27 GCA_000008505.1
611 Bacillus cereus ATCC 14579 GCA_000007825.1
611 Bacillus thuringiensis YBT-1518 GCA_000497525.2
590 Bacillus pseudomycoides DSM 12442 GCA_000161455.1
Jm
653 Bifidobacterium adolescentis ATCC 15703 GCA_000010425.1

516 Bifidobacterium dentium JCM 1195 = DSM 20436 GCA_001042595.1
504 Bifidobacterium angulatum DSM 20098 = JCM 7096 GCA_001025155.1
Jn
720 Clostridium beijerinckii GCA_000833105.2
702 Clostridium saccharoperbutylacetonicum N1-4_28HMT_29 GCA_000340885.1
691 Clostridium saccharobutylicum DSM 13864 GCA_000473995.1
Jo
578 Rhodobacter sphaeroides 2.4.1 GCA_000012905.2
490 Rhodobacter sphaeroides ATCC 17025 GCA_000016405.1
445 Defluviimonas alba GCA_001620265.1
Jp
694 Enterococcus faecalis V583 GCA_000007785.1
648 Streptomyces cinnamoneus GCA_001885705.1
643 Enterococcus faecium DO GCA_000174395.2
Jq
534 Clostridium beijerinckii GCA_000833105.2
483 Clostridium saccharoperbutylacetonicum N1-4_28HMT_29 GCA_000340885.1
461 Clostridium puniceum GCA_002006345.1
Jr
809 Bacillus anthracis str. Ames GCA_000007845.1
809 Bacillus anthracis str. Sterne GCA_000008165.1
809 Bacillus cereus ATCC 14579 GCA_000007825.1
809 _5BBacillus thuringiensis_5D serovar konkukian str. 97-27 GCA_000008505.1
794 Bacillus pseudomycoides DSM 12442 GCA_000161455.1
794 Bacillus thuringiensis YBT-1518 GCA_000497525.2
756 Bacillus mycoides GCA_000832605.1
Js
590 Lactobacillus gasseri ATCC 33323 = JCM 1131 GCA_000014425.1
573 Lactobacillus hominis DSM 23910 = CRBIP 24.179 GCA_000296835.1
448 Lactobacillus psittaci DSM 15354 GCA_000425905.1
Jt
626 Bifidobacterium adolescentis ATCC 15703 GCA_000010425.1
494 Bifidobacterium dentium JCM 1195 = DSM 20436 GCA_001042595.1
473 Bifidobacterium callitrichos DSM 23973 GCA_000741175.1
Ju
556 Rhodobacter sphaeroides 2.4.1 GCA_000012905.2
513 Rhodobacter sphaeroides ATCC 17025 GCA_000016405.1
401 Pseudorhodobacter ferrugineus DSM 5888 GCA_000420745.1
Jv
476 Bacillus thuringiensis YBT-1518 GCA_000497525.2
461 Bacillus anthracis str. Ames GCA_000007845.1
461 Bacillus anthracis str. Sterne GCA_000008165.1
461 Bacillus cereus ATCC 14579 GCA_000007825.1
461 _5BBacillus thuringiensis_5D serovar konkukian str. 97-27 GCA_000008505.1
431 Bacillus pseudomycoides DSM 12442 GCA_000161455.1
Jw
788 Streptococcus mutans UA159 GCA_000007465.2
570 Streptococcus ratti FA-1 = DSM 20564 GCA_000286075.1
555 Streptococcus equinus GCA_000964315.1
Jx
583 Bacillus thuringiensis YBT-1518 GCA_000497525.2
570 Bacillus anthracis str. Ames GCA_000007845.1
570 Bacillus anthracis str. Sterne GCA_000008165.1
570 Bacillus cereus ATCC 14579 GCA_000007825.1
570 Bacillus pseudomycoides DSM 12442 GCA_000161455.1
570 _5BBacillus thuringiensis_5D serovar konkukian str. 97-27 GCA_000008505.1
523 Bacillus cytotoxicus NVH 391-98 GCA_000017425.1
Jy
827 Enterococcus faecalis V583 GCA_000007785.1
784 Streptomyces cinnamoneus GCA_001885705.1
744 Enterococcus rivorum GCA_001742285.1
Jz
690 Bifidobacterium adolescentis ATCC 15703 GCA_000010425.1
572 Bifidobacterium dentium JCM 1195 = DSM 20436 GCA_001042595.1
535 Bifidobacterium angulatum DSM 20098 = JCM 7096 GCA_001025155.1
JA
699 Clostridium beijerinckii GCA_000833105.2
669 Clostridium saccharoperbutylacetonicum N1-4_28HMT_29 GCA_000340885.1
644 Clostridium puniceum GCA_002006345.1
JB
588 Clostridium beijerinckii GCA_000833105.2
569 Clostridium saccharobutylicum DSM 13864 GCA_000473995.1
569 Clostridium saccharoperbutylacetonicum N1-4_28HMT_29 GCA_000340885.1
543 Clostridium puniceum GCA_002006345.1
JC
372 Clostridium beijerinckii GCA_000833105.2

338 Clostridium saccharoperbutylacetonicum N1-4_28HMT_29 GCA_000340885.1
336 Clostridium saccharobutylicum DSM 13864 GCA_000473995.1
JD
647 Streptococcus mutans UA159 GCA_000007465.2
450 Streptococcus ratti FA-1 = DSM 20564 GCA_000286075.1
440 Streptococcus gordonii str. Challis substr. CH1 GCA_000017005.1
JE
672 Deinococcus radiodurans R1 GCA_000008565.1
425 Deinococcus deserti VCD115 GCA_000020685.1
379 Deinococcus hopiensis KR-140 GCA_900176165.1
JF
701 Bacillus anthracis str. Ames GCA_000007845.1
701 Bacillus anthracis str. Sterne GCA_000008165.1
701 _5BBacillus thuringiensis_5D serovar konkukian str. 97-27 GCA_000008505.1
691 Bacillus cereus ATCC 14579 GCA_000007825.1
691 Bacillus thuringiensis YBT-1518 GCA_000497525.2
643 Bacillus pseudomycoides DSM 12442 GCA_000161455.1
JG
806 Deinococcus radiodurans R1 GCA_000008565.1
520 Deinococcus deserti VCD115 GCA_000020685.1
501 Deinococcus hopiensis KR-140 GCA_900176165.1
JH
541 Lactobacillus gasseri ATCC 33323 = JCM 1131 GCA_000014425.1
541 Lactobacillus hominis DSM 23910 = CRBIP 24.179 GCA_000296835.1
433 Lactobacillus iners DSM 13335 GCA_000160875.1
354 Lactobacillus farraginis DSM 18382 = JCM 14108 GCA_000583655.1
JI
502 Staphylococcus epidermidis ATCC 12228 GCA_000007645.1
493 Staphylococcus capitis subsp. capitis GCA_001028645.1
457 Staphylococcus aureus subsp. aureus NCTC 8325 GCA_000013425.1
JJ
672 Clostridium saccharoperbutylacetonicum N1-4_28HMT_29 GCA_000340885.1
671 Clostridium beijerinckii GCA_000833105.2
665 Clostridium saccharobutylicum DSM 13864 GCA_000473995.1
JK
798 Lactobacillus gasseri ATCC 33323 = JCM 1131 GCA_000014425.1
766 Lactobacillus hominis DSM 23910 = CRBIP 24.179 GCA_000296835.1
595 Lactobacillus iners DSM 13335 GCA_000160875.1
JL
524 Rhodobacter sphaeroides 2.4.1 GCA_000012905.2
466 Rhodobacter sphaeroides ATCC 17025 GCA_000016405.1
381 Defluviimonas alba GCA_001620265.1
JM
355 Rhodobacter sphaeroides 2.4.1 GCA_000012905.2
295 Rhodobacter sphaeroides ATCC 17025 GCA_000016405.1
274 Gemmobacter megaterium GCA_900156815.1
JN
663 Clostridium beijerinckii GCA_000833105.2
638 Clostridium saccharoperbutylacetonicum N1-4_28HMT_29 GCA_000340885.1
630 Clostridium puniceum GCA_002006345.1
JO
611 Enterococcus faecalis V583 GCA_000007785.1
564 Streptomyces cinnamoneus GCA_001885705.1
537 Enterococcus rivorum GCA_001742285.1
JP
805 Rhodobacter sphaeroides 2.4.1 GCA_000012905.2
714 Rhodobacter sphaeroides ATCC 17025 GCA_000016405.1
623 Pseudorhodobacter psychrotolerans GCA_001294535.1
JQ
530 Clostridium beijerinckii GCA_000833105.2
485 Clostridium saccharoperbutylacetonicum N1-4_28HMT_29 GCA_000340885.1
478 Clostridium puniceum GCA_002006345.1
JR
872 Lactobacillus gasseri ATCC 33323 = JCM 1131 GCA_000014425.1
796 Lactobacillus hominis DSM 23910 = CRBIP 24.179 GCA_000296835.1
598 Lactobacillus iners DSM 13335 GCA_000160875.1
JS
549 Clostridium beijerinckii GCA_000833105.2
519 Clostridium butyricum GCA_001456065.2
519 Clostridium puniceum GCA_002006345.1
519 Clostridium saccharoperbutylacetonicum N1-4_28HMT_29 GCA_000340885.1
492 Clostridium saccharobutylicum DSM 13864 GCA_000473995.1
JT
535 Enterococcus faecalis V583 GCA_000007785.1
488 Streptomyces cinnamoneus GCA_001885705.1
472 Enterococcus asini ATCC 700915 GCA_000407365.1

JU
530 Bacillus anthracis str. Ames GCA_000007845.1
530 Bacillus anthracis str. Sterne GCA_000008165.1
530 _5BBacillus thuringiensis_5D serovar konkukian str. 97-27 GCA_000008505.1
509 Bacillus cereus ATCC 14579 GCA_000007825.1
504 Bacillus pseudomycoides DSM 12442 GCA_000161455.1
JV
771 Clostridium beijerinckii GCA_000833105.2
707 Clostridium saccharoperbutylacetonicum N1-4_28HMT_29 GCA_000340885.1
679 Clostridium saccharobutylicum DSM 13864 GCA_000473995.1
JW
640 Clostridium beijerinckii GCA_000833105.2
626 Clostridium saccharoperbutylacetonicum N1-4_28HMT_29 GCA_000340885.1
601 Clostridium puniceum GCA_002006345.1
JX
805 Lactobacillus gasseri ATCC 33323 = JCM 1131 GCA_000014425.1
777 Lactobacillus hominis DSM 23910 = CRBIP 24.179 GCA_000296835.1
600 Lactobacillus iners DSM 13335 GCA_000160875.1
JY
539 Deinococcus radiodurans R1 GCA_000008565.1
308 Deinococcus deserti VCD115 GCA_000020685.1
280 Deinococcus soli Cha et al. 2016 GCA_001007995.1
JZ
589 Rhodobacter sphaeroides 2.4.1 GCA_000012905.2
497 Rhodobacter sphaeroides ATCC 17025 GCA_000016405.1
435 Gemmobacter aquatilis GCA_900110025.1
K0
888 Bacillus anthracis str. Ames GCA_000007845.1
888 Bacillus anthracis str. Sterne GCA_000008165.1
888 _5BBacillus thuringiensis_5D serovar konkukian str. 97-27 GCA_000008505.1
877 Bacillus cereus ATCC 14579 GCA_000007825.1
863 Bacillus thuringiensis YBT-1518 GCA_000497525.2
K1
359 Clostridium beijerinckii GCA_000833105.2
359 Clostridium saccharoperbutylacetonicum N1-4_28HMT_29 GCA_000340885.1
352 Clostridium saccharobutylicum DSM 13864 GCA_000473995.1
302 Clostridium butyricum GCA_001456065.2
K2
675 Bifidobacterium adolescentis ATCC 15703 GCA_000010425.1
527 Bifidobacterium dentium JCM 1195 = DSM 20436 GCA_001042595.1
507 Bifidobacterium breve DSM 20213 = JCM 1192 GCA_001025175.1
507 Bifidobacterium reuteri DSM 23975 GCA_000741695.1
K3
794 Lactobacillus gasseri ATCC 33323 = JCM 1131 GCA_000014425.1
736 Lactobacillus hominis DSM 23910 = CRBIP 24.179 GCA_000296835.1
499 Lactobacillus iners DSM 13335 GCA_000160875.1
K4
717 Clostridium beijerinckii GCA_000833105.2
696 Clostridium saccharoperbutylacetonicum N1-4_28HMT_29 GCA_000340885.1
646 Clostridium saccharobutylicum DSM 13864 GCA_000473995.1
K5
419 Rhodobacter sphaeroides 2.4.1 GCA_000012905.2
376 Rhodobacter sphaeroides ATCC 17025 GCA_000016405.1
364 Gemmobacter megaterium GCA_900156815.1
K6
472 Bacillus anthracis str. Ames GCA_000007845.1
472 Bacillus anthracis str. Sterne GCA_000008165.1
472 Bacillus pseudomycoides DSM 12442 GCA_000161455.1
472 _5BBacillus thuringiensis_5D serovar konkukian str. 97-27 GCA_000008505.1
456 Bacillus cereus ATCC 14579 GCA_000007825.1
456 Bacillus mycoides GCA_000832605.1
442 Bacillus cytotoxicus NVH 391-98 GCA_000017425.1
K7
922 Rhodobacter sphaeroides 2.4.1 GCA_000012905.2
872 Rhodobacter sphaeroides ATCC 17025 GCA_000016405.1
713 Pseudorhodobacter psychrotolerans GCA_001294535.1
K8
460 Staphylococcus epidermidis ATCC 12228 GCA_000007645.1
457 Staphylococcus capitis subsp. capitis GCA_001028645.1
418 Staphylococcus warneri SG1 GCA_000332735.1
K9
830 Clostridium beijerinckii GCA_000833105.2
796 Clostridium saccharoperbutylacetonicum N1-4_28HMT_29 GCA_000340885.1
758 Clostridium puniceum GCA_002006345.1
Ka
793 Deinococcus radiodurans R1 GCA_000008565.1

517 Deinococcus deserti VCD115 GCA_000020685.1
487 Deinococcus murrayi DSM 11303 GCA_000482805.1
Kb
760 Rhodobacter sphaeroides 2.4.1 GCA_000012905.2
713 Rhodobacter sphaeroides ATCC 17025 GCA_000016405.1
616 Gemmobacter aquatilis GCA_900110025.1
Kc
741 Deinococcus radiodurans R1 GCA_000008565.1
523 Deinococcus soli Cha et al. 2016 GCA_001007995.1
505 Deinococcus deserti VCD115 GCA_000020685.1
Kd
648 Streptococcus mutans UA159 GCA_000007465.2
448 Streptococcus ratti FA-1 = DSM 20564 GCA_000286075.1
446 Streptococcus equinus GCA_000964315.1
Ke
722 Streptococcus mutans UA159 GCA_000007465.2
497 Streptococcus equinus GCA_000964315.1
497 Streptococcus ratti FA-1 = DSM 20564 GCA_000286075.1
496 Streptococcus gallolyticus subsp. gallolyticus DSM 16831 GCA_002000985.1
496 Streptococcus gordonii str. Challis substr. CH1 GCA_000017005.1
Kf
537 Staphylococcus epidermidis ATCC 12228 GCA_000007645.1
490 Staphylococcus capitis subsp. capitis GCA_001028645.1
469 Staphylococcus haemolyticus JCSC1435 GCA_000009865.1
Kg
809 Streptococcus mutans UA159 GCA_000007465.2
612 Streptococcus ratti FA-1 = DSM 20564 GCA_000286075.1
569 Streptococcus gordonii str. Challis substr. CH1 GCA_000017005.1
Kh
715 Deinococcus radiodurans R1 GCA_000008565.1
492 Deinococcus soli Cha et al. 2016 GCA_001007995.1
483 Deinococcus deserti VCD115 GCA_000020685.1
Ki
809 Enterococcus faecalis V583 GCA_000007785.1
768 Streptomyces cinnamoneus GCA_001885705.1
708 Enterococcus saccharolyticus subsp. saccharolyticus ATCC 43076 GCA_000407285.1
Kj
577 Deinococcus radiodurans R1 GCA_000008565.1
377 Deinococcus deserti VCD115 GCA_000020685.1
368 Deinococcus proteolyticus MRP GCA_000190555.1
Kk
467 Bifidobacterium adolescentis ATCC 15703 GCA_000010425.1
403 Bifidobacterium dentium JCM 1195 = DSM 20436 GCA_001042595.1
383 Bifidobacterium bifidum PRL2010 GCA_000165905.1
Kl
693 Bacillus thuringiensis YBT-1518 GCA_000497525.2
685 Bacillus anthracis str. Ames GCA_000007845.1
685 Bacillus anthracis str. Sterne GCA_000008165.1
685 _5BBacillus thuringiensis_5D serovar konkukian str. 97-27 GCA_000008505.1
679 Bacillus cereus ATCC 14579 GCA_000007825.1
Km
635 Lactobacillus gasseri ATCC 33323 = JCM 1131 GCA_000014425.1
605 Lactobacillus hominis DSM 23910 = CRBIP 24.179 GCA_000296835.1
450 Lactobacillus iners DSM 13335 GCA_000160875.1
Kn
592 Deinococcus radiodurans R1 GCA_000008565.1
347 Deinococcus deserti VCD115 GCA_000020685.1
321 Deinococcus gobiensis I-0 GCA_000252445.1
Ko
868 Rhodobacter sphaeroides 2.4.1 GCA_000012905.2
773 Rhodobacter sphaeroides ATCC 17025 GCA_000016405.1
633 Gemmobacter megaterium GCA_900156815.1
Kp
826 Clostridium beijerinckii GCA_000833105.2
796 Clostridium saccharoperbutylacetonicum N1-4_28HMT_29 GCA_000340885.1
758 Clostridium saccharobutylicum DSM 13864 GCA_000473995.1
Kq
642 Bifidobacterium adolescentis ATCC 15703 GCA_000010425.1
544 Bifidobacterium dentium JCM 1195 = DSM 20436 GCA_001042595.1
490 Bifidobacterium stellenboschense GCA_000741785.1
Kr
575 Lactobacillus gasseri ATCC 33323 = JCM 1131 GCA_000014425.1
476 Lactobacillus hominis DSM 23910 = CRBIP 24.179 GCA_000296835.1
322 Lactobacillus iners DSM 13335 GCA_000160875.1
Ks
742 Streptococcus mutans UA159 GCA_000007465.2

555 Streptococcus ratti FA-1 = DSM 20564 GCA_000286075.1
524 Streptococcus ferus DSM 20646 GCA_000372425.1
Kt
735 Lactobacillus gasseri ATCC 33323 = JCM 1131 GCA_000014425.1
703 Lactobacillus hominis DSM 23910 = CRBIP 24.179 GCA_000296835.1
577 Lactobacillus psittaci DSM 15354 GCA_000425905.1
Ku
702 Bacillus anthracis str. Ames GCA_000007845.1
702 Bacillus anthracis str. Sterne GCA_000008165.1
702 _5BBacillus thuringiensis_5D serovar konkukian str. 97-27 GCA_000008505.1
689 Bacillus cereus ATCC 14579 GCA_000007825.1
689 Bacillus thuringiensis YBT-1518 GCA_000497525.2
676 Bacillus pseudomycoides DSM 12442 GCA_000161455.1
Kv
760 Staphylococcus capitis subsp. capitis GCA_001028645.1
760 Staphylococcus epidermidis ATCC 12228 GCA_000007645.1
677 Staphylococcus aureus subsp. aureus NCTC 8325 GCA_000013425.1
667 Staphylococcus haemolyticus JCSC1435 GCA_000009865.1
Kw
509 Clostridium beijerinckii GCA_000833105.2
490 Clostridium butyricum GCA_001456065.2
481 Clostridium saccharobutylicum DSM 13864 GCA_000473995.1
Kx
645 Clostridium beijerinckii GCA_000833105.2
609 Clostridium saccharoperbutylacetonicum N1-4_28HMT_29 GCA_000340885.1
582 Clostridium puniceum GCA_002006345.1
Ky
805 Clostridium beijerinckii GCA_000833105.2
764 Clostridium saccharoperbutylacetonicum N1-4_28HMT_29 GCA_000340885.1
744 Clostridium puniceum GCA_002006345.1
Kz
638 Lactobacillus gasseri ATCC 33323 = JCM 1131 GCA_000014425.1
560 Lactobacillus hominis DSM 23910 = CRBIP 24.179 GCA_000296835.1
481 Lactobacillus iners DSM 13335 GCA_000160875.1
KA
753 Clostridium beijerinckii GCA_000833105.2
693 Clostridium saccharoperbutylacetonicum N1-4_28HMT_29 GCA_000340885.1
663 Clostridium saccharobutylicum DSM 13864 GCA_000473995.1
KB
361 Staphylococcus epidermidis ATCC 12228 GCA_000007645.1
356 Staphylococcus aureus subsp. aureus NCTC 8325 GCA_000013425.1
352 Staphylococcus capitis subsp. capitis GCA_001028645.1
KC
547 Clostridium beijerinckii GCA_000833105.2
532 Clostridium butyricum GCA_001456065.2
528 Clostridium puniceum GCA_002006345.1
528 Clostridium saccharoperbutylacetonicum N1-4_28HMT_29 GCA_000340885.1
KD
907 Enterococcus faecalis V583 GCA_000007785.1
861 Streptomyces cinnamoneus GCA_001885705.1
799 Enterococcus rivorum GCA_001742285.1
KE
768 Rhodobacter sphaeroides 2.4.1 GCA_000012905.2
675 Rhodobacter sphaeroides ATCC 17025 GCA_000016405.1
595 Defluviimonas alba GCA_001620265.1
KF
736 Staphylococcus epidermidis ATCC 12228 GCA_000007645.1
694 Staphylococcus capitis subsp. capitis GCA_001028645.1
641 Staphylococcus warneri SG1 GCA_000332735.1
KG
509 Enterococcus faecalis V583 GCA_000007785.1
490 Enterococcus saccharolyticus subsp. saccharolyticus ATCC 43076 GCA_000407285.1
477 Enterococcus sulfureus ATCC 49903 GCA_000407605.1
KH
843 Deinococcus radiodurans R1 GCA_000008565.1
507 Deinococcus gobiensis I-0 GCA_000252445.1
485 Deinococcus deserti VCD115 GCA_000020685.1
KI
692 Escherichia coli O157_3AH7 str. Sakai GCA_000008865.1
692 Escherichia coli str. K-12 substr. MG1655 GCA_000005845.2
691 Escherichia coli UMN026 GCA_000026325.2
677 Shigella flexneri 2a str. 301 GCA_000006925.2
KJ
638 Rhodobacter sphaeroides 2.4.1 GCA_000012905.2
580 Rhodobacter sphaeroides ATCC 17025 GCA_000016405.1
490 Defluviimonas alba GCA_001620265.1

KK
689 Clostridium beijerinckii GCA_000833105.2
621 Clostridium chromiireducens GCA_002029255.1
611 Clostridium saccharoperbutylacetonicum N1-4_28HMT_29 GCA_000340885.1
KL
269 Lactobacillus gasseri ATCC 33323 = JCM 1131 GCA_000014425.1
240 Lactobacillus hominis DSM 23910 = CRBIP 24.179 GCA_000296835.1
166 Lactobacillus acetotolerans GCA_001042405.1
KM
700 Deinococcus radiodurans R1 GCA_000008565.1
464 Deinococcus deserti VCD115 GCA_000020685.1
425 Deinococcus soli Cha et al. 2016 GCA_001007995.1
KN
801 Streptococcus mutans UA159 GCA_000007465.2
661 Streptococcus ratti FA-1 = DSM 20564 GCA_000286075.1
616 Streptococcus criceti HS-6 GCA_000187975.3
616 Streptococcus gordonii str. Challis substr. CH1 GCA_000017005.1
KO
584 Enterococcus faecalis V583 GCA_000007785.1
555 Streptomyces cinnamoneus GCA_001885705.1
523 Enterococcus rivorum GCA_001742285.1
KP
828 Enterococcus faecalis V583 GCA_000007785.1
781 Streptomyces cinnamoneus GCA_001885705.1
768 Enterococcus faecium DO GCA_000174395.2
KQ
409 Streptococcus mutans UA159 GCA_000007465.2
266 Streptococcus macacae NCTC 11558 GCA_000187995.3
255 Streptococcus ratti FA-1 = DSM 20564 GCA_000286075.1
KR
751 _5BBacillus thuringiensis_5D serovar konkukian str. 97-27 GCA_000008505.1
749 Bacillus anthracis str. Ames GCA_000007845.1
749 Bacillus anthracis str. Sterne GCA_000008165.1
737 Bacillus cereus ATCC 14579 GCA_000007825.1
KS
465 Lactobacillus gasseri ATCC 33323 = JCM 1131 GCA_000014425.1
407 Lactobacillus hominis DSM 23910 = CRBIP 24.179 GCA_000296835.1
350 Lactobacillus iners DSM 13335 GCA_000160875.1
KT
445 Rhodobacter sphaeroides 2.4.1 GCA_000012905.2
371 Rhodobacter sphaeroides ATCC 17025 GCA_000016405.1
317 Gemmobacter megaterium GCA_900156815.1
KU
427 Rhodobacter sphaeroides 2.4.1 GCA_000012905.2
408 Rhodobacter sphaeroides ATCC 17025 GCA_000016405.1
317 Gemmobacter nectariphilus DSM 15620 GCA_000429765.1
317 Pseudorhodobacter psychrotolerans GCA_001294535.1
KV
638 Lactobacillus gasseri ATCC 33323 = JCM 1131 GCA_000014425.1
587 Lactobacillus hominis DSM 23910 = CRBIP 24.179 GCA_000296835.1
433 Lactobacillus iners DSM 13335 GCA_000160875.1
KW
596 Streptococcus mutans UA159 GCA_000007465.2
447 Streptococcus ratti FA-1 = DSM 20564 GCA_000286075.1
402 Streptococcus ferus DSM 20646 GCA_000372425.1
KX
647 Clostridium beijerinckii GCA_000833105.2
632 Clostridium saccharoperbutylacetonicum N1-4_28HMT_29 GCA_000340885.1
619 Clostridium saccharobutylicum DSM 13864 GCA_000473995.1
KY
948 Streptococcus mutans UA159 GCA_000007465.2
620 Streptococcus gordonii str. Challis substr. CH1 GCA_000017005.1
614 Streptococcus ratti FA-1 = DSM 20564 GCA_000286075.1
KZ
813 Deinococcus radiodurans R1 GCA_000008565.1
472 Deinococcus deserti VCD115 GCA_000020685.1
456 Deinococcus gobiensis I-0 GCA_000252445.1
L0
981 Lactobacillus gasseri ATCC 33323 = JCM 1131 GCA_000014425.1
923 Lactobacillus hominis DSM 23910 = CRBIP 24.179 GCA_000296835.1
728 Lactobacillus iners DSM 13335 GCA_000160875.1
L1
512 Lactobacillus gasseri ATCC 33323 = JCM 1131 GCA_000014425.1
434 Lactobacillus hominis DSM 23910 = CRBIP 24.179 GCA_000296835.1
367 Lactobacillus iners DSM 13335 GCA_000160875.1
L2

660 Rhodobacter sphaeroides 2.4.1 GCA_000012905.2
571 Rhodobacter sphaeroides ATCC 17025 GCA_000016405.1
461 Gemmobacter aquatilis GCA_900110025.1
L3
479 Rhodobacter sphaeroides 2.4.1 GCA_000012905.2
468 Rhodobacter sphaeroides ATCC 17025 GCA_000016405.1
360 Pseudorhodobacter wandonensis GCA_001202035.1
L4
835 Staphylococcus epidermidis ATCC 12228 GCA_000007645.1
833 Staphylococcus capitis subsp. capitis GCA_001028645.1
775 Staphylococcus aureus subsp. aureus NCTC 8325 GCA_000013425.1
L5
614 Bacillus anthracis str. Ames GCA_000007845.1
614 Bacillus anthracis str. Sterne GCA_000008165.1
614 Bacillus cereus ATCC 14579 GCA_000007825.1
614 _5BBacillus thuringiensis_5D serovar konkukian str. 97-27 GCA_000008505.1
608 Bacillus thuringiensis YBT-1518 GCA_000497525.2
607 Bacillus mycoides GCA_000832605.1
L6
656 Deinococcus radiodurans R1 GCA_000008565.1
439 Deinococcus puniceus GCA_001644565.1
428 Deinococcus hopiensis KR-140 GCA_900176165.1
L7
804 Enterococcus faecalis V583 GCA_000007785.1
757 Streptomyces cinnamoneus GCA_001885705.1
714 Enterococcus asini ATCC 700915 GCA_000407365.1
714 Enterococcus malodoratus ATCC 43197 GCA_000407185.1
L8
513 Rhodobacter sphaeroides 2.4.1 GCA_000012905.2
495 Rhodobacter sphaeroides ATCC 17025 GCA_000016405.1
449 Gemmobacter aquatilis GCA_900110025.1
L9
365 Enterococcus faecalis V583 GCA_000007785.1
318 Streptomyces cinnamoneus GCA_001885705.1
305 Enterococcus dispar ATCC 51266 GCA_000406945.1
305 Enterococcus phoeniculicola ATCC BAA-412 GCA_000407505.1
La
558 Escherichia coli str. K-12 substr. MG1655 GCA_000005845.2
547 Escherichia coli IAI39 GCA_000026345.1
547 Escherichia coli O104_3AH4 str. 2011C-3493 GCA_000299455.1
547 Shigella flexneri 2a str. 301 GCA_000006925.2
546 Escherichia coli UMN026 GCA_000026325.2
Lb
576 Streptococcus mutans UA159 GCA_000007465.2
434 Streptococcus sobrinus DSM 20742 = ATCC 33478 GCA_000686605.1
433 Streptococcus ratti FA-1 = DSM 20564 GCA_000286075.1
Lc
835 Enterococcus faecalis V583 GCA_000007785.1
789 Streptomyces cinnamoneus GCA_001885705.1
700 Enterococcus rivorum GCA_001742285.1
Ld
664 Bifidobacterium adolescentis ATCC 15703 GCA_000010425.1
525 Bifidobacterium dentium JCM 1195 = DSM 20436 GCA_001042595.1
496 Bifidobacterium angulatum DSM 20098 = JCM 7096 GCA_001025155.1
Le
690 Enterococcus faecalis V583 GCA_000007785.1
643 Streptomyces cinnamoneus GCA_001885705.1
641 Enterococcus rivorum GCA_001742285.1
Lf
632 Lactobacillus gasseri ATCC 33323 = JCM 1131 GCA_000014425.1
605 Lactobacillus hominis DSM 23910 = CRBIP 24.179 GCA_000296835.1
451 Lactobacillus psittaci DSM 15354 GCA_000425905.1
Lg
658 Streptococcus mutans UA159 GCA_000007465.2
481 Streptococcus ratti FA-1 = DSM 20564 GCA_000286075.1
481 Streptococcus sobrinus DSM 20742 = ATCC 33478 GCA_000686605.1
467 Streptococcus gordonii str. Challis substr. CH1 GCA_000017005.1
Lh
765 Staphylococcus epidermidis ATCC 12228 GCA_000007645.1
713 Staphylococcus capitis subsp. capitis GCA_001028645.1
695 Staphylococcus hominis subsp. hominis C80 GCA_000183685.1
Li
758 Staphylococcus epidermidis ATCC 12228 GCA_000007645.1
727 Staphylococcus capitis subsp. capitis GCA_001028645.1
642 Staphylococcus lugdunensis HKU09-01 GCA_000025085.1
Lj

548 Rhodobacter sphaeroides 2.4.1 GCA_000012905.2
483 Rhodobacter sphaeroides ATCC 17025 GCA_000016405.1
374 Pseudorhodobacter ferrugineus DSM 5888 GCA_000420745.1
374 Pseudorhodobacter psychrotolerans GCA_001294535.1
Lk
863 Rhodobacter sphaeroides 2.4.1 GCA_000012905.2
774 Rhodobacter sphaeroides ATCC 17025 GCA_000016405.1
669 Gemmobacter megaterium GCA_900156815.1
Ll
737 Clostridium beijerinckii GCA_000833105.2
720 Clostridium saccharoperbutylacetonicum N1-4_28HMT_29 GCA_000340885.1
675 Clostridium saccharobutylicum DSM 13864 GCA_000473995.1
Lm
506 Bacillus cereus ATCC 14579 GCA_000007825.1
506 Bacillus mycoides GCA_000832605.1
506 Bacillus thuringiensis YBT-1518 GCA_000497525.2
504 Bacillus anthracis str. Ames GCA_000007845.1
504 Bacillus anthracis str. Sterne GCA_000008165.1
504 _5BBacillus thuringiensis_5D serovar konkukian str. 97-27 GCA_000008505.1
486 Bacillus pseudomycoides DSM 12442 GCA_000161455.1
Ln
462 Bifidobacterium adolescentis ATCC 15703 GCA_000010425.1
385 Bifidobacterium angulatum DSM 20098 = JCM 7096 GCA_001025155.1
324 Bifidobacterium callitrichos DSM 23973 GCA_000741175.1
Lo
681 Rhodobacter sphaeroides 2.4.1 GCA_000012905.2
606 Rhodobacter sphaeroides ATCC 17025 GCA_000016405.1
547 Defluviimonas alba GCA_001620265.1
Lp
779 Streptococcus mutans UA159 GCA_000007465.2
587 Streptococcus ratti FA-1 = DSM 20564 GCA_000286075.1
547 Streptococcus gordonii str. Challis substr. CH1 GCA_000017005.1
Lq
491 Lactobacillus gasseri ATCC 33323 = JCM 1131 GCA_000014425.1
431 Lactobacillus hominis DSM 23910 = CRBIP 24.179 GCA_000296835.1
330 Lactobacillus iners DSM 13335 GCA_000160875.1
Lr
796 Enterococcus faecalis V583 GCA_000007785.1
749 Streptomyces cinnamoneus GCA_001885705.1
667 Enterococcus mundtii QU 25 GCA_000504125.1
Ls
506 Staphylococcus epidermidis ATCC 12228 GCA_000007645.1
482 Staphylococcus capitis subsp. capitis GCA_001028645.1
445 Staphylococcus haemolyticus JCSC1435 GCA_000009865.1
445 Staphylococcus hominis subsp. hominis C80 GCA_000183685.1
Lt
753 Rhodobacter sphaeroides 2.4.1 GCA_000012905.2
679 Rhodobacter sphaeroides ATCC 17025 GCA_000016405.1
597 Defluviimonas alba GCA_001620265.1
Lu
836 Streptococcus mutans UA159 GCA_000007465.2
575 Streptococcus ratti FA-1 = DSM 20564 GCA_000286075.1
562 Streptococcus sobrinus DSM 20742 = ATCC 33478 GCA_000686605.1
Lv
709 Streptococcus mutans UA159 GCA_000007465.2
549 Streptococcus ratti FA-1 = DSM 20564 GCA_000286075.1
473 Streptococcus sobrinus DSM 20742 = ATCC 33478 GCA_000686605.1
Lw
844 Bacillus anthracis str. Ames GCA_000007845.1
844 Bacillus anthracis str. Sterne GCA_000008165.1
844 Bacillus cereus ATCC 14579 GCA_000007825.1
844 Bacillus pseudomycoides DSM 12442 GCA_000161455.1
844 _5BBacillus thuringiensis_5D serovar konkukian str. 97-27 GCA_000008505.1
833 Bacillus mycoides GCA_000832605.1
821 Bacillus thuringiensis YBT-1518 GCA_000497525.2
Lx
861 Staphylococcus epidermidis ATCC 12228 GCA_000007645.1
801 Staphylococcus capitis subsp. capitis GCA_001028645.1
751 Staphylococcus haemolyticus JCSC1435 GCA_000009865.1
Ly
401 Clostridium beijerinckii GCA_000833105.2
365 Clostridium puniceum GCA_002006345.1
365 Clostridium saccharoperbutylacetonicum N1-4_28HMT_29 GCA_000340885.1
358 Clostridium butyricum GCA_001456065.2
Lz
855 Streptococcus mutans UA159 GCA_000007465.2

630 Streptococcus ratti FA-1 = DSM 20564 GCA_000286075.1
575 Streptococcus merionis DSM 19192 GCA_000380085.1
LA
593 Rhodobacter sphaeroides 2.4.1 GCA_000012905.2
572 Rhodobacter sphaeroides ATCC 17025 GCA_000016405.1
470 Gemmobacter aquatilis GCA_900110025.1
LB
500 Bifidobacterium adolescentis ATCC 15703 GCA_000010425.1
460 Bifidobacterium dentium JCM 1195 = DSM 20436 GCA_001042595.1
423 Bifidobacterium callitrichos DSM 23973 GCA_000741175.1
LC
676 Clostridium beijerinckii GCA_000833105.2
635 Clostridium saccharoperbutylacetonicum N1-4_28HMT_29 GCA_000340885.1
610 Clostridium butyricum GCA_001456065.2
LD
661 Bacillus anthracis str. Ames GCA_000007845.1
661 Bacillus anthracis str. Sterne GCA_000008165.1
661 Bacillus cereus ATCC 14579 GCA_000007825.1
661 Bacillus pseudomycoides DSM 12442 GCA_000161455.1
661 Bacillus thuringiensis YBT-1518 GCA_000497525.2
661 _5BBacillus thuringiensis_5D serovar konkukian str. 97-27 GCA_000008505.1
645 Bacillus mycoides GCA_000832605.1
624 Bacillus cytotoxicus NVH 391-98 GCA_000017425.1
LE
768 Rhodobacter sphaeroides 2.4.1 GCA_000012905.2
654 Rhodobacter sphaeroides ATCC 17025 GCA_000016405.1
593 Defluviimonas alba GCA_001620265.1
LF
540 Lactobacillus gasseri ATCC 33323 = JCM 1131 GCA_000014425.1
498 Lactobacillus hominis DSM 23910 = CRBIP 24.179 GCA_000296835.1
397 Lactobacillus jensenii GCA_001936235.1
LG
764 Streptococcus mutans UA159 GCA_000007465.2
552 Streptococcus ratti FA-1 = DSM 20564 GCA_000286075.1
525 Streptococcus gordonii str. Challis substr. CH1 GCA_000017005.1
LH
618 Clostridium beijerinckii GCA_000833105.2
610 Clostridium saccharoperbutylacetonicum N1-4_28HMT_29 GCA_000340885.1
591 Clostridium saccharobutylicum DSM 13864 GCA_000473995.1
LI
733 Clostridium beijerinckii GCA_000833105.2
730 Clostridium saccharoperbutylacetonicum N1-4_28HMT_29 GCA_000340885.1
702 Clostridium saccharobutylicum DSM 13864 GCA_000473995.1
LJ
540 Escherichia coli O104_3AH4 str. 2011C-3493 GCA_000299455.1
525 Escherichia coli str. K-12 substr. MG1655 GCA_000005845.2
525 Shigella flexneri 2a str. 301 GCA_000006925.2
510 Escherichia coli IAI39 GCA_000026345.1
LK
620 Streptococcus mutans UA159 GCA_000007465.2
369 Streptococcus ratti FA-1 = DSM 20564 GCA_000286075.1
355 Streptococcus equinus GCA_000964315.1
355 Streptococcus gallolyticus subsp. gallolyticus DSM 16831 GCA_002000985.1
LL
707 Escherichia coli str. K-12 substr. MG1655 GCA_000005845.2
691 Escherichia coli O157_3AH7 str. Sakai GCA_000008865.1
691 Escherichia coli UMN026 GCA_000026325.2
691 Shigella flexneri 2a str. 301 GCA_000006925.2
690 Escherichia coli O104_3AH4 str. 2011C-3493 GCA_000299455.1
LM
692 Rhodobacter sphaeroides 2.4.1 GCA_000012905.2
647 Rhodobacter sphaeroides ATCC 17025 GCA_000016405.1
512 Gemmobacter megaterium GCA_900156815.1
LN
884 Enterococcus faecalis V583 GCA_000007785.1
838 Streptomyces cinnamoneus GCA_001885705.1
759 Enterococcus asini ATCC 700915 GCA_000407365.1
LO
283 Staphylococcus capitis subsp. capitis GCA_001028645.1
283 Staphylococcus epidermidis ATCC 12228 GCA_000007645.1
241 Staphylococcus warneri SG1 GCA_000332735.1
240 Staphylococcus aureus subsp. aureus NCTC 8325 GCA_000013425.1
LP
439 Clostridium puniceum GCA_002006345.1
439 Clostridium saccharoperbutylacetonicum N1-4_28HMT_29 GCA_000340885.1
434 Clostridium beijerinckii GCA_000833105.2

430 Clostridium saccharobutylicum DSM 13864 GCA_000473995.1
LQ
538 Clostridium saccharoperbutylacetonicum N1-4_28HMT_29 GCA_000340885.1
533 Clostridium beijerinckii GCA_000833105.2
495 Clostridium saccharobutylicum DSM 13864 GCA_000473995.1
LR
776 Deinococcus radiodurans R1 GCA_000008565.1
491 Deinococcus deserti VCD115 GCA_000020685.1
464 Deinococcus gobiensis I-0 GCA_000252445.1
LS
479 Clostridium beijerinckii GCA_000833105.2
470 Clostridium saccharoperbutylacetonicum N1-4_28HMT_29 GCA_000340885.1
463 Clostridium saccharobutylicum DSM 13864 GCA_000473995.1
LT
488 Deinococcus radiodurans R1 GCA_000008565.1
272 Deinococcus deserti VCD115 GCA_000020685.1
266 Deinococcus marmoris DSM 12784 GCA_000701405.1
266 Deinococcus swuensis GCA_000800395.1
LU
454 Lactobacillus gasseri ATCC 33323 = JCM 1131 GCA_000014425.1
439 Lactobacillus hominis DSM 23910 = CRBIP 24.179 GCA_000296835.1
319 Lactobacillus iners DSM 13335 GCA_000160875.1
LV
609 Rhodobacter sphaeroides 2.4.1 GCA_000012905.2
547 Rhodobacter sphaeroides ATCC 17025 GCA_000016405.1
458 Gemmobacter aquatilis GCA_900110025.1
LW
799 Enterococcus faecalis V583 GCA_000007785.1
752 Streptomyces cinnamoneus GCA_001885705.1
701 Enterococcus rivorum GCA_001742285.1
LX
765 Enterococcus faecalis V583 GCA_000007785.1
718 Streptomyces cinnamoneus GCA_001885705.1
698 Enterococcus rivorum GCA_001742285.1
LY
521 Lactobacillus gasseri ATCC 33323 = JCM 1131 GCA_000014425.1
488 Lactobacillus hominis DSM 23910 = CRBIP 24.179 GCA_000296835.1
388 Lactobacillus iners DSM 13335 GCA_000160875.1
LZ
786 Escherichia coli str. K-12 substr. MG1655 GCA_000005845.2
784 Escherichia coli O157_3AH7 str. Sakai GCA_000008865.1
779 Escherichia coli UMN026 GCA_000026325.2
M0
946 Bifidobacterium adolescentis ATCC 15703 GCA_000010425.1
761 Bifidobacterium angulatum DSM 20098 = JCM 7096 GCA_001025155.1
738 Bifidobacterium dentium JCM 1195 = DSM 20436 GCA_001042595.1
M1
620 Bifidobacterium adolescentis ATCC 15703 GCA_000010425.1
543 Bifidobacterium breve DSM 20213 = JCM 1192 GCA_001025175.1
535 Bifidobacterium dentium JCM 1195 = DSM 20436 GCA_001042595.1
M2
800 Staphylococcus epidermidis ATCC 12228 GCA_000007645.1
759 Staphylococcus capitis subsp. capitis GCA_001028645.1
684 Staphylococcus hominis subsp. hominis C80 GCA_000183685.1
M3
754 Streptococcus mutans UA159 GCA_000007465.2
646 Streptococcus ratti FA-1 = DSM 20564 GCA_000286075.1
630 Streptococcus gordonii str. Challis substr. CH1 GCA_000017005.1
M4
761 Bifidobacterium adolescentis ATCC 15703 GCA_000010425.1
651 Bifidobacterium dentium JCM 1195 = DSM 20436 GCA_001042595.1
613 Bifidobacterium angulatum DSM 20098 = JCM 7096 GCA_001025155.1
M5
686 Staphylococcus epidermidis ATCC 12228 GCA_000007645.1
657 Staphylococcus capitis subsp. capitis GCA_001028645.1
593 Staphylococcus aureus subsp. aureus NCTC 8325 GCA_000013425.1
M6
811 Streptococcus mutans UA159 GCA_000007465.2
610 Streptococcus ratti FA-1 = DSM 20564 GCA_000286075.1
546 Streptococcus macacae NCTC 11558 GCA_000187995.3
M7
638 Enterococcus faecalis V583 GCA_000007785.1
591 Streptomyces cinnamoneus GCA_001885705.1
538 Enterococcus rivorum GCA_001742285.1
M8
271 Rhodobacter sphaeroides 2.4.1 GCA_000012905.2

260 Rhodobacter sphaeroides ATCC 17025 GCA_000016405.1
209 Gemmobacter aquatilis GCA_900110025.1
M9
652 Enterococcus faecalis V583 GCA_000007785.1
605 Streptomyces cinnamoneus GCA_001885705.1
564 Enterococcus faecium DO GCA_000174395.2
Ma
717 Clostridium beijerinckii GCA_000833105.2
688 Clostridium saccharoperbutylacetonicum N1-4_28HMT_29 GCA_000340885.1
673 Clostridium saccharobutylicum DSM 13864 GCA_000473995.1
Mb
546 Rhodobacter sphaeroides 2.4.1 GCA_000012905.2
502 Rhodobacter sphaeroides ATCC 17025 GCA_000016405.1
429 Pseudorhodobacter ferrugineus DSM 5888 GCA_000420745.1
429 Pseudorhodobacter psychrotolerans GCA_001294535.1
429 Pseudorhodobacter wandonensis GCA_001202035.1
Mc
619 Rhodobacter sphaeroides 2.4.1 GCA_000012905.2
550 Rhodobacter sphaeroides ATCC 17025 GCA_000016405.1
454 Gemmobacter aquatilis GCA_900110025.1
Md
443 Streptococcus mutans UA159 GCA_000007465.2
367 Streptococcus anginosus C238 GCA_000463505.1
361 Streptococcus merionis DSM 19192 GCA_000380085.1
Me
543 Bifidobacterium adolescentis ATCC 15703 GCA_000010425.1
420 Bifidobacterium dentium JCM 1195 = DSM 20436 GCA_001042595.1
419 Bifidobacterium angulatum DSM 20098 = JCM 7096 GCA_001025155.1
Mf
507 Staphylococcus epidermidis ATCC 12228 GCA_000007645.1
486 Staphylococcus haemolyticus JCSC1435 GCA_000009865.1
481 Staphylococcus capitis subsp. capitis GCA_001028645.1
Mg
517 Bifidobacterium adolescentis ATCC 15703 GCA_000010425.1
364 Bifidobacterium dentium JCM 1195 = DSM 20436 GCA_001042595.1
353 Bifidobacterium angulatum DSM 20098 = JCM 7096 GCA_001025155.1
Mh
783 Enterococcus faecalis V583 GCA_000007785.1
783 Streptomyces cinnamoneus GCA_001885705.1
736 Enterococcus asini ATCC 700915 GCA_000407365.1
723 Enterococcus italicus DSM 15952 GCA_000185365.1
Mi
757 Escherichia coli O157_3AH7 str. Sakai GCA_000008865.1
757 Escherichia coli str. K-12 substr. MG1655 GCA_000005845.2
748 Escherichia coli IAI39 GCA_000026345.1
737 Escherichia coli UMN026 GCA_000026325.2
Mj
732 Clostridium beijerinckii GCA_000833105.2
698 Clostridium saccharoperbutylacetonicum N1-4_28HMT_29 GCA_000340885.1
664 Clostridium puniceum GCA_002006345.1
Mk
668 Enterococcus faecalis V583 GCA_000007785.1
622 Streptomyces cinnamoneus GCA_001885705.1
565 Enterococcus faecium DO GCA_000174395.2
Ml
645 Bacillus anthracis str. Ames GCA_000007845.1
645 Bacillus anthracis str. Sterne GCA_000008165.1
645 Bacillus cereus ATCC 14579 GCA_000007825.1
645 Bacillus mycoides GCA_000832605.1
645 Bacillus pseudomycoides DSM 12442 GCA_000161455.1
645 _5BBacillus thuringiensis_5D serovar konkukian str. 97-27 GCA_000008505.1
632 Bacillus thuringiensis YBT-1518 GCA_000497525.2
558 Bacillus cytotoxicus NVH 391-98 GCA_000017425.1
Mm
735 Streptococcus mutans UA159 GCA_000007465.2
498 Streptococcus ratti FA-1 = DSM 20564 GCA_000286075.1
460 Streptococcus ferus DSM 20646 GCA_000372425.1
Mn
845 Enterococcus faecalis V583 GCA_000007785.1
804 Streptomyces cinnamoneus GCA_001885705.1
749 Enterococcus rivorum GCA_001742285.1
Mo
819 Deinococcus radiodurans R1 GCA_000008565.1
513 Deinococcus marmoris DSM 12784 GCA_000701405.1
502 Deinococcus deserti VCD115 GCA_000020685.1
