## Additional File 5 for "Full-length 16S rRNA gene amplicon analysis of human gut microbiota using MinION™ nanopore sequencing confers species-level resolution"

Alignment search results for V3-V4 amplicon sequencing of the mock community.
The top three hits for 3,000 query sequences are shown. The values represent the bit score of alignment.

1
198 Escherichia coli IAI39 GCA_000026345.1
198 Escherichia coli O104_3AH4 str. 2011C-3493 GCA_000299455.1
198 Escherichia coli O157_3AH7 str. Sakai GCA_000008865.1
198 Escherichia coli O83_3AH1 str. NRG 857C GCA_000183345.1
198 Escherichia coli UMN026 GCA_000026325.2
198 Escherichia coli str. K-12 substr. MG1655 GCA_000005845.2
198 Shigella dysenteriae Sd197 GCA_000012005.1
198 Shigella flexneri 2a str. 301 GCA_000006925.2
198 Tumebacillus flagellatus GCA_000714935.1
183 Erwinia iniecta GCA_001267535.1
183 Rosenbergiella nectarea GCA_900111105.1
177 Cronobacter sakazakii GCA_000982825.1
177 Erwinia toletana DAPP-PG 735 GCA_000336255.1
2
244 Enterococcus faecalis V583 GCA_000007785.1
244 Streptomyces cinnamoneus GCA_001885705.1
207 Enterococcus canis NBRC 100695 GCA_001544375.1
207 Enterococcus casseliflavus EC20 GCA_000157355.2
207 Enterococcus dispar ATCC 51266 GCA_000406945.1
207 Enterococcus faecium DO GCA_000174395.2
207 Enterococcus hirae ATCC 9790 GCA_000271405.2
207 Enterococcus mundtii QU 25 GCA_000504125.1
207 Enterococcus rivorum GCA_001742285.1
207 Enterococcus saccharolyticus subsp. saccharolyticus ATCC 43076 GCA_000407285.1
202 Enterococcus haemoperoxidus ATCC BAA-382 GCA_000407165.1
202 Enterococcus phoeniculicola ATCC BAA-412 GCA_000407505.1
202 Enterococcus thailandicus GCA_001652875.1
3
343 Enterococcus faecalis V583 GCA_000007785.1
343 Streptomyces cinnamoneus GCA_001885705.1
328 Enterococcus dispar ATCC 51266 GCA_000406945.1
325 Enterococcus canis NBRC 100695 GCA_001544375.1
325 Enterococcus faecium DO GCA_000174395.2
325 Enterococcus hirae ATCC 9790 GCA_000271405.2
325 Enterococcus mundtii QU 25 GCA_000504125.1
325 Enterococcus rivorum GCA_001742285.1
4
196 Bacillus anthracis str. Ames GCA_000007845.1
196 Bacillus anthracis str. Sterne GCA_000008165.1
196 Bacillus cereus ATCC 14579 GCA_000007825.1
196 Bacillus mycoides GCA_000832605.1
196 Bacillus pseudomycoides DSM 12442 GCA_000161455.1
196 Bacillus thuringiensis YBT-1518 GCA_000497525.2
196 _5BBacillus thuringiensis_5D serovar konkukian str. 97-27 GCA_000008505.1
166 Massilibacterium senegalense GCA_001375675.1
153 Bacillus flexus GCA_002024265.1
153 Bacillus megaterium NBRC 15308 = ATCC 14581 GCA_000832985.1
5
247 Enterococcus faecalis V583 GCA_000007785.1
247 Streptomyces cinnamoneus GCA_001885705.1
216 Enterococcus asini ATCC 700915 GCA_000407365.1
216 Enterococcus canis NBRC 100695 GCA_001544375.1
216 Enterococcus cecorum GCA_001318405.1
216 Enterococcus columbae DSM 7374 = ATCC 51263 GCA_000406925.1
216 Enterococcus dispar ATCC 51266 GCA_000406945.1
216 Enterococcus faecium DO GCA_000174395.2
216 Enterococcus haemoperoxidus ATCC BAA-382 GCA_000407165.1
216 Enterococcus hirae ATCC 9790 GCA_000271405.2
216 Enterococcus massiliensis GCA_001050095.1
216 Enterococcus mundtii QU 25 GCA_000504125.1
216 Enterococcus phoeniculicola ATCC BAA-412 GCA_000407505.1
216 Enterococcus rivorum GCA_001742285.1
216 Enterococcus thailandicus GCA_001652875.1
216 Melissococcus plutonius S1 GCA_000747585.1
211 Enterococcus aquimarinus GCA_001885765.1
211 Enterococcus gilvus ATCC BAA-350 GCA_000407545.1
211 Enterococcus hermanniensis GCA_001885945.1
211 Enterococcus italicus DSM 15952 GCA_000185365.1
211 Enterococcus malodoratus ATCC 43197 GCA_000407185.1
211 Enterococcus pallens ATCC BAA-351 GCA_000407485.1
211 Enterococcus pseudoavium NBRC 100491 GCA_001544295.1

211 Enterococcus sulfureus ATCC 49903 GCA_000407605.1
211 Vagococcus fluvialis bH819 GCA_900163795.1
211 Vagococcus lutrae LBD1 GCA_000498295.1
211 Vagococcus teuberi GCA_001870205.1
6
304 Bacillus thuringiensis YBT-1518 GCA_000497525.2
303 Bacillus anthracis str. Ames GCA_000007845.1
303 Bacillus anthracis str. Sterne GCA_000008165.1
303 Bacillus cereus ATCC 14579 GCA_000007825.1
303 Bacillus mycoides GCA_000832605.1
303 Bacillus pseudomycoides DSM 12442 GCA_000161455.1
303 _5BBacillus thuringiensis_5D serovar konkukian str. 97-27 GCA_000008505.1
259 Bacillus marisflavi GCA_001274775.1
7
218 Rhodobacter sphaeroides 2.4.1 GCA_000012905.2
204 Rhodobacter sphaeroides ATCC 17025 GCA_000016405.1
170 Gemmobacter aquatilis GCA_900110025.1
8
177 Deinococcus radiodurans R1 GCA_000008565.1
105 Deinococcus gobiensis I-0 GCA_000252445.1
94 Deinococcus proteolyticus MRP GCA_000190555.1
9
228 Bacillus mycoides GCA_000832605.1
214 Staphylococcus equorum GCA_001432245.1
206 Staphylococcus arlettae CVD059 GCA_000295715.1
206 Staphylococcus cohnii subsp. cohnii GCA_000972575.1
206 Staphylococcus epidermidis ATCC 12228 GCA_000007645.1
206 Staphylococcus gallinarum GCA_000875895.1
206 Staphylococcus saprophyticus GCA_001074355.1
206 Staphylococcus saprophyticus subsp. saprophyticus ATCC 15305 GCA_000010125.1
206 Staphylococcus succinus GCA_001902315.1
206 Staphylococcus xylosus GCA_000706685.1
a
165 Deinococcus radiodurans R1 GCA_000008565.1
92 Deinococcus hopiensis KR-140 GCA_900176165.1
90 Deinococcus gobiensis I-0 GCA_000252445.1
90 Deinococcus marmoris DSM 12784 GCA_000701405.1
90 Deinococcus swuensis GCA_000800395.1
b
282 Deinococcus radiodurans R1 GCA_000008565.1
157 Deinococcus gobiensis I-0 GCA_000252445.1
155 Deinococcus marmoris DSM 12784 GCA_000701405.1
155 Deinococcus swuensis GCA_000800395.1
c
223 Streptococcus mutans UA159 GCA_000007465.2
122 Streptococcus ratti FA-1 = DSM 20564 GCA_000286075.1
120 Streptococcus macacae NCTC 11558 GCA_000187995.3
120 Streptococcus marimammalium DSM 18627 GCA_000380045.1
120 Streptococcus sobrinus DSM 20742 = ATCC 33478 GCA_000686605.1
d
159 Erwinia iniecta GCA_001267535.1
159 Escherichia coli IAI39 GCA_000026345.1
159 Escherichia coli O104_3AH4 str. 2011C-3493 GCA_000299455.1
159 Escherichia coli O157_3AH7 str. Sakai GCA_000008865.1
159 Escherichia coli UMN026 GCA_000026325.2
159 Escherichia coli str. K-12 substr. MG1655 GCA_000005845.2
159 Shigella flexneri 2a str. 301 GCA_000006925.2
159 Tumebacillus flagellatus GCA_000714935.1
158 Escherichia coli O83_3AH1 str. NRG 857C GCA_000183345.1
150 Shigella dysenteriae Sd197 GCA_000012005.1
e
279 Streptococcus mutans UA159 GCA_000007465.2
203 Streptococcus ratti FA-1 = DSM 20564 GCA_000286075.1
194 Streptococcus gordonii str. Challis substr. CH1 GCA_000017005.1
f
187 Clostridium beijerinckii GCA_000833105.2
187 Clostridium saccharoperbutylacetonicum N1-4_28HMT_29 GCA_000340885.1
183 Clostridium butyricum GCA_001456065.2
181 Clostridium puniceum GCA_002006345.1
g
226 Clostridium beijerinckii GCA_000833105.2
226 Clostridium puniceum GCA_002006345.1
226 Clostridium saccharoperbutylacetonicum N1-4_28HMT_29 GCA_000340885.1
217 Clostridium saccharobutylicum DSM 13864 GCA_000473995.1
192 Clostridium chromiireducens GCA_002029255.1
h

171 Bacillus thuringiensis YBT-1518 GCA_000497525.2
165 Bacillus anthracis str. Ames GCA_000007845.1
165 Bacillus anthracis str. Sterne GCA_000008165.1
165 Bacillus cereus ATCC 14579 GCA_000007825.1
165 Bacillus mycoides GCA_000832605.1
165 Bacillus pseudomycoides DSM 12442 GCA_000161455.1
165 _5BBacillus thuringiensis_5D serovar konkukian str. 97-27 GCA_000008505.1
144 Bacillus manliponensis GCA_000712595.1
i
139 Lactobacillus acetotolerans GCA_001042405.1
139 Lactobacillus amylovorus GCA_000191545.1
139 Lactobacillus crispatus ST1 GCA_000091765.1
139 Lactobacillus gasseri ATCC 33323 = JCM 1131 GCA_000014425.1
139 Lactobacillus helveticus GCA_001308285.1
139 Lactobacillus hominis DSM 23910 = CRBIP 24.179 GCA_000296835.1
139 Lactobacillus kalixensis DSM 16043 GCA_001434335.1
139 Lactobacillus kullabergensis GCA_000967195.1
139 Lactobacillus psittaci DSM 15354 GCA_000425905.1
130 Lactobacillus apis GCA_000970735.1
130 Lactobacillus mellis GCA_000967245.1
130 Pediococcus cellicola GCA_001438655.1
130 Pediococcus ethanolidurans GCA_001437405.1
123 Lactobacillus acidophilus NCFM GCA_000011985.1
123 Lactobacillus bombicola GCA_900112665.1
123 Lactobacillus camelliae DSM 22697 = JCM 13995 GCA_001311195.1
123 Lactobacillus farraginis DSM 18382 = JCM 14108 GCA_000583655.1
123 Lactobacillus gigeriorum DSM 23908 = CRBIP 24.85 GCA_001436575.1
123 Lactobacillus hamsteri DSM 5661 = JCM 6256 GCA_000615445.1
123 Lactobacillus intestinalis DSM 6629 GCA_001435325.1
123 Lactobacillus jensenii GCA_001936235.1
123 Lactobacillus nasuensis JCM 17158 GCA_001434705.1
123 Lactobacillus pantheris DSM 15945 = JCM 12539 = NBRC 106106 GCA_001311175.1
123 Lactobacillus paracasei ATCC 334 GCA_000014525.1
123 Lactobacillus paracasei subsp. paracasei JCM 8130 GCA_000829035.1
123 Lactobacillus pasteurii DSM 23907 = CRBIP 24.76 GCA_000297025.1
123 Lactobacillus rhamnosus GG GCA_000026505.1
123 Lactobacillus thailandensis DSM 22698 = JCM 13996 GCA_001312865.1
123 Pediococcus acidilactici GCA_001767275.1
123 Pediococcus pentosaceus ATCC 25745 GCA_000014505.1
123 Pediococcus stilesii GCA_001437075.1
j
280 Clostridium beijerinckii GCA_000833105.2
280 Clostridium puniceum GCA_002006345.1
280 Clostridium saccharoperbutylacetonicum N1-4_28HMT_29 GCA_000340885.1
266 Clostridium saccharobutylicum DSM 13864 GCA_000473995.1
229 Clostridium chromiireducens GCA_002029255.1
229 Clostridium taeniosporum GCA_001735765.1
k
257 Deinococcus radiodurans R1 GCA_000008565.1
207 Deinococcus gobiensis I-0 GCA_000252445.1
204 Deinococcus puniceus GCA_001644565.1
l
174 Erwinia iniecta GCA_001267535.1
174 Escherichia coli IAI39 GCA_000026345.1
174 Escherichia coli O104_3AH4 str. 2011C-3493 GCA_000299455.1
174 Escherichia coli O157_3AH7 str. Sakai GCA_000008865.1
174 Escherichia coli O83_3AH1 str. NRG 857C GCA_000183345.1
174 Escherichia coli UMN026 GCA_000026325.2
174 Escherichia coli str. K-12 substr. MG1655 GCA_000005845.2
174 Shigella dysenteriae Sd197 GCA_000012005.1
174 Shigella flexneri 2a str. 301 GCA_000006925.2
174 Tumebacillus flagellatus GCA_000714935.1
171 Enterobacter hormaechei subsp. steigerwaltii GCA_001729725.1
171 Erwinia toletana DAPP-PG 735 GCA_000336255.1
171 Klebsiella oxytoca GCA_001022195.1
171 Kosakonia cowanii GCA_001975225.1
171 Pantoea agglomerans GCA_001709315.1
171 Pantoea ananatis LMG 20103 GCA_000025405.2
171 Pantoea rwandensis GCA_000759475.1
171 Pantoea septica GCA_002095575.1
171 Pluralibacter gergoviae GCA_000757785.1
171 Pseudescherichia vulneris NBRC 102420 GCA_000759795.1
171 Shimwellia blattae DSM 4481 = NBRC 105725 GCA_000262305.1
170 Kosakonia sacchari SP1 GCA_000300455.4
m
311 Bacillus anthracis str. Ames GCA_000007845.1

311 Bacillus anthracis str. Sterne GCA_000008165.1
311 Bacillus cereus ATCC 14579 GCA_000007825.1
311 Bacillus mycoides GCA_000832605.1
311 Bacillus pseudomycoides DSM 12442 GCA_000161455.1
311 Bacillus thuringiensis YBT-1518 GCA_000497525.2
311 _5BBacillus thuringiensis_5D serovar konkukian str. 97-27 GCA_000008505.1
277 Bacillus manliponensis GCA_000712595.1
256 Bacillus cytotoxicus NVH 391-98 GCA_000017425.1
n
139 Bifidobacterium breve DSM 20213 = JCM 1192 GCA_001025175.1
138 Bifidobacterium adolescentis ATCC 15703 GCA_000010425.1
137 Bifidobacterium longum NCC2705 GCA_000007525.1
o
148 Escherichia coli IAI39 GCA_000026345.1
148 Escherichia coli O104_3AH4 str. 2011C-3493 GCA_000299455.1
148 Escherichia coli O157_3AH7 str. Sakai GCA_000008865.1
148 Escherichia coli O83_3AH1 str. NRG 857C GCA_000183345.1
148 Escherichia coli UMN026 GCA_000026325.2
148 Escherichia coli str. K-12 substr. MG1655 GCA_000005845.2
148 Shigella dysenteriae Sd197 GCA_000012005.1
148 Shigella flexneri 2a str. 301 GCA_000006925.2
148 Tumebacillus flagellatus GCA_000714935.1
134 Pantoea agglomerans GCA_001709315.1
133 Cronobacter sakazakii GCA_000982825.1
133 Enterobacter cloacae subsp. cloacae ATCC 13047 GCA_000025565.1
133 Enterobacter hormaechei subsp. steigerwaltii GCA_001729725.1
133 Enterobacter kobei GCA_900185885.1
133 Erwinia amylovora CFBP1430 GCA_000091565.1
133 Erwinia billingiae Eb661 GCA_000196615.1
133 Erwinia gerundensis GCA_001517405.1
133 Erwinia iniecta GCA_001267535.1
133 Erwinia toletana DAPP-PG 735 GCA_000336255.1
133 Klebsiella oxytoca GCA_001022195.1
133 Kosakonia cowanii GCA_001975225.1
133 Kosakonia sacchari SP1 GCA_000300455.4
133 Pantoea alhagi GCA_002101395.1
133 Pantoea ananatis LMG 20103 GCA_000025405.2
133 Pantoea dispersa EGD-AAK13 GCA_000465555.2
133 Pantoea rwandensis GCA_000759475.1
133 Pantoea septica GCA_002095575.1
133 Pluralibacter gergoviae GCA_000757785.1
133 Pseudescherichia vulneris NBRC 102420 GCA_000759795.1
133 Salmonella enterica subsp. enterica serovar Typhi str. CT18 GCA_000195995.1
133 Salmonella enterica subsp. enterica serovar Typhimurium str. LT2 GCA_000006945.2
133 Shimwellia blattae DSM 4481 = NBRC 105725 GCA_000262305.1
133 Trabulsiella odontotermitis GCA_001297765.1
p
226 Clostridium beijerinckii GCA_000833105.2
226 Clostridium saccharobutylicum DSM 13864 GCA_000473995.1
226 Clostridium saccharoperbutylacetonicum N1-4_28HMT_29 GCA_000340885.1
219 Clostridium puniceum GCA_002006345.1
208 Clostridium chromiireducens GCA_002029255.1
q
277 Deinococcus radiodurans R1 GCA_000008565.1
211 Deinococcus deserti VCD115 GCA_000020685.1
208 Deinococcus gobiensis I-0 GCA_000252445.1
r
127 Bifidobacterium adolescentis ATCC 15703 GCA_000010425.1
127 Bifidobacterium dentium JCM 1195 = DSM 20436 GCA_001042595.1
112 Bifidobacterium tsurumiense GCA_000741765.1
94 Bifidobacterium thermophilum GCA_000741495.1
s
204 Clostridium beijerinckii GCA_000833105.2
204 Clostridium puniceum GCA_002006345.1
204 Clostridium saccharobutylicum DSM 13864 GCA_000473995.1
204 Clostridium saccharoperbutylacetonicum N1-4_28HMT_29 GCA_000340885.1
185 Clostridium chromiireducens GCA_002029255.1
184 Clostridium neonatale GCA_001458595.1
t
257 Rhodobacter sphaeroides 2.4.1 GCA_000012905.2
246 Rhodobacter sphaeroides ATCC 17025 GCA_000016405.1
227 Pseudorhodobacter ferrugineus DSM 5888 GCA_000420745.1
227 Pseudorhodobacter wandonensis GCA_001202035.1
u
232 Deinococcus radiodurans R1 GCA_000008565.1
165 Deinococcus gobiensis I-0 GCA_000252445.1

150 Deinococcus proteolyticus MRP GCA_000190555.1
v
184 Rhodobacter sphaeroides 2.4.1 GCA_000012905.2
170 Rhodobacter sphaeroides ATCC 17025 GCA_000016405.1
134 Pseudorhodobacter ferrugineus DSM 5888 GCA_000420745.1
134 Pseudorhodobacter psychrotolerans GCA_001294535.1
134 Pseudorhodobacter wandonensis GCA_001202035.1
w
246 Deinococcus radiodurans R1 GCA_000008565.1
190 Deinococcus gobiensis I-0 GCA_000252445.1
149 Deinococcus deserti VCD115 GCA_000020685.1
x
155 Clostridium beijerinckii GCA_000833105.2
155 Clostridium puniceum GCA_002006345.1
155 Clostridium saccharobutylicum DSM 13864 GCA_000473995.1
155 Clostridium saccharoperbutylacetonicum N1-4_28HMT_29 GCA_000340885.1
144 Clostridium chromiireducens GCA_002029255.1
144 Clostridium taeniosporum GCA_001735765.1
138 Clostridium butyricum GCA_001456065.2
y
291 Deinococcus radiodurans R1 GCA_000008565.1
188 Deinococcus gobiensis I-0 GCA_000252445.1
184 Deinococcus marmoris DSM 12784 GCA_000701405.1
184 Deinococcus swuensis GCA_000800395.1
z
233 Deinococcus radiodurans R1 GCA_000008565.1
164 Deinococcus marmoris DSM 12784 GCA_000701405.1
164 Deinococcus swuensis GCA_000800395.1
147 Deinococcus gobiensis I-0 GCA_000252445.1
A
176 Escherichia coli IAI39 GCA_000026345.1
176 Escherichia coli O104_3AH4 str. 2011C-3493 GCA_000299455.1
176 Escherichia coli O157_3AH7 str. Sakai GCA_000008865.1
176 Escherichia coli O83_3AH1 str. NRG 857C GCA_000183345.1
176 Escherichia coli UMN026 GCA_000026325.2
176 Escherichia coli str. K-12 substr. MG1655 GCA_000005845.2
176 Shigella dysenteriae Sd197 GCA_000012005.1
176 Shigella flexneri 2a str. 301 GCA_000006925.2
176 Tumebacillus flagellatus GCA_000714935.1
161 Erwinia iniecta GCA_001267535.1
155 Cronobacter sakazakii GCA_000982825.1
155 Erwinia toletana DAPP-PG 735 GCA_000336255.1
B
307 Bacillus thuringiensis YBT-1518 GCA_000497525.2
290 Bacillus anthracis str. Ames GCA_000007845.1
290 Bacillus anthracis str. Sterne GCA_000008165.1
290 Bacillus cereus ATCC 14579 GCA_000007825.1
290 Bacillus mycoides GCA_000832605.1
290 Bacillus pseudomycoides DSM 12442 GCA_000161455.1
290 _5BBacillus thuringiensis_5D serovar konkukian str. 97-27 GCA_000008505.1
213 Bacillus manliponensis GCA_000712595.1
C
213 Lactobacillus gasseri ATCC 33323 = JCM 1131 GCA_000014425.1
198 Lactobacillus hominis DSM 23910 = CRBIP 24.179 GCA_000296835.1
191 Lactobacillus amylovorus GCA_000191545.1
191 Lactobacillus crispatus ST1 GCA_000091765.1
191 Lactobacillus kalixensis DSM 16043 GCA_001434335.1
191 Lactobacillus psittaci DSM 15354 GCA_000425905.1
D
287 Rhodobacter sphaeroides 2.4.1 GCA_000012905.2
280 Rhodobacter sphaeroides ATCC 17025 GCA_000016405.1
236 Gemmobacter aquatilis GCA_900110025.1
E
207 Escherichia coli IAI39 GCA_000026345.1
207 Escherichia coli O104_3AH4 str. 2011C-3493 GCA_000299455.1
207 Escherichia coli O157_3AH7 str. Sakai GCA_000008865.1
207 Escherichia coli O83_3AH1 str. NRG 857C GCA_000183345.1
207 Escherichia coli UMN026 GCA_000026325.2
207 Escherichia coli str. K-12 substr. MG1655 GCA_000005845.2
207 Shigella dysenteriae Sd197 GCA_000012005.1
207 Shigella flexneri 2a str. 301 GCA_000006925.2
207 Tumebacillus flagellatus GCA_000714935.1
185 Erwinia iniecta GCA_001267535.1
173 Enterobacter hormaechei subsp. steigerwaltii GCA_001729725.1
173 Erwinia toletana DAPP-PG 735 GCA_000336255.1
173 Klebsiella oxytoca GCA_001022195.1

173 Klebsiella pneumoniae subsp. pneumoniae HS11286 GCA_000240185.2
173 Kosakonia cowanii GCA_001975225.1
173 Pantoea alhagi GCA_002101395.1
173 Pluralibacter gergoviae GCA_000757785.1
173 Pseudescherichia vulneris NBRC 102420 GCA_000759795.1
F
154 Megasphaera cerevisiae DSM 20462 GCA_001045675.1
154 Staphylococcus aureus subsp. aureus NCTC 8325 GCA_000013425.1
154 Staphylococcus capitis subsp. capitis GCA_001028645.1
154 Staphylococcus epidermidis ATCC 12228 GCA_000007645.1
154 Staphylococcus haemolyticus JCSC1435 GCA_000009865.1
154 Staphylococcus hominis subsp. hominis C80 GCA_000183685.1
154 Staphylococcus lugdunensis HKU09-01 GCA_000025085.1
154 Staphylococcus lutrae GCA_002101335.1
154 Staphylococcus simiae CCM 7213 GCA_000235645.2
154 Staphylococcus warneri SG1 GCA_000332735.1
151 Staphylococcus arlettae CVD059 GCA_000295715.1
151 Staphylococcus cohnii subsp. cohnii GCA_000972575.1
151 Staphylococcus gallinarum GCA_000875895.1
151 Staphylococcus lentus F1142 GCA_000286395.1
151 Staphylococcus saprophyticus GCA_001074355.1
151 Staphylococcus saprophyticus subsp. saprophyticus ATCC 15305 GCA_000010125.1
151 Staphylococcus sciuri GCA_002209165.1
151 Staphylococcus succinus GCA_001902315.1
151 Staphylococcus xylosus GCA_000706685.1
147 Staphylococcus auricularis GCA_001500315.1
G
167 Streptococcus mutans UA159 GCA_000007465.2
138 Streptococcus gordonii str. Challis substr. CH1 GCA_000017005.1
135 Streptococcus gallolyticus subsp. gallolyticus DSM 16831 GCA_002000985.1
H
186 Streptococcus mutans UA159 GCA_000007465.2
112 Streptococcus ratti FA-1 = DSM 20564 GCA_000286075.1
110 Streptococcus gordonii str. Challis substr. CH1 GCA_000017005.1
I
279 Clostridium saccharobutylicum DSM 13864 GCA_000473995.1
276 Clostridium butyricum GCA_001456065.2
275 Clostridium chromiireducens GCA_002029255.1
J
281 Clostridium saccharobutylicum DSM 13864 GCA_000473995.1
279 Clostridium beijerinckii GCA_000833105.2
279 Clostridium saccharoperbutylacetonicum N1-4_28HMT_29 GCA_000340885.1
267 Clostridium chromiireducens GCA_002029255.1
K
142 Streptococcus mutans UA159 GCA_000007465.2
137 Streptococcus massiliensis DSM 18628 GCA_000380065.1
130 Streptococcus ferus DSM 20646 GCA_000372425.1
L
184 Clostridium beijerinckii GCA_000833105.2
184 Clostridium butyricum GCA_001456065.2
184 Clostridium chromiireducens GCA_002029255.1
184 Clostridium puniceum GCA_002006345.1
184 Clostridium saccharobutylicum DSM 13864 GCA_000473995.1
184 Clostridium saccharoperbutylacetonicum N1-4_28HMT_29 GCA_000340885.1
178 Clostridium neonatale GCA_001458595.1
157 Clostridium fallax GCA_900129365.1
M
232 Escherichia coli IAI39 GCA_000026345.1
232 Escherichia coli O104_3AH4 str. 2011C-3493 GCA_000299455.1
232 Escherichia coli O157_3AH7 str. Sakai GCA_000008865.1
232 Escherichia coli O83_3AH1 str. NRG 857C GCA_000183345.1
232 Escherichia coli UMN026 GCA_000026325.2
232 Escherichia coli str. K-12 substr. MG1655 GCA_000005845.2
232 Shigella flexneri 2a str. 301 GCA_000006925.2
232 Tumebacillus flagellatus GCA_000714935.1
223 Shigella dysenteriae Sd197 GCA_000012005.1
199 Erwinia iniecta GCA_001267535.1
N
148 Rhodobacter sphaeroides 2.4.1 GCA_000012905.2
143 Rhodobacter sphaeroides ATCC 17025 GCA_000016405.1
133 Pseudorhodobacter ferrugineus DSM 5888 GCA_000420745.1
133 Pseudorhodobacter wandonensis GCA_001202035.1
O
160 Streptococcus mutans UA159 GCA_000007465.2
124 Streptococcus gordonii str. Challis substr. CH1 GCA_000017005.1
120 Streptococcus agalactiae 2603V_2FR GCA_000007265.1

120 Streptococcus cristatus AS 1.3089 GCA_000385925.1
120 Streptococcus dysgalactiae subsp. equisimilis AC-2713 GCA_000317855.1
120 Streptococcus ictaluri 707-05 GCA_000188015.3
P
193 Bifidobacterium tsurumiense GCA_000741765.1
191 Bifidobacterium gallicum DSM 20093 = LMG 11596 GCA_000741205.1
189 Bifidobacterium longum NCC2705 GCA_000007525.1
189 Bifidobacterium scardovii JCM 12489 = DSM 13734 GCA_001042635.1
189 Bifidobacterium thermophilum GCA_000741495.1
Q
236 Clostridium beijerinckii GCA_000833105.2
236 Clostridium puniceum GCA_002006345.1
236 Clostridium saccharoperbutylacetonicum N1-4_28HMT_29 GCA_000340885.1
222 Clostridium saccharobutylicum DSM 13864 GCA_000473995.1
218 Clostridium taeniosporum GCA_001735765.1
R
227 Rhodobacter sphaeroides 2.4.1 GCA_000012905.2
191 Defluviimonas alba GCA_001620265.1
175 Rhodobacter sphaeroides ATCC 17025 GCA_000016405.1
S
113 Rhodobacter sphaeroides 2.4.1 GCA_000012905.2
107 Rhodobacter sphaeroides ATCC 17025 GCA_000016405.1
93 Pseudorhodobacter ferrugineus DSM 5888 GCA_000420745.1
93 Pseudorhodobacter wandonensis GCA_001202035.1
T
109 Erwinia iniecta GCA_001267535.1
109 Escherichia coli IAI39 GCA_000026345.1
109 Escherichia coli O104_3AH4 str. 2011C-3493 GCA_000299455.1
109 Escherichia coli O157_3AH7 str. Sakai GCA_000008865.1
109 Escherichia coli O83_3AH1 str. NRG 857C GCA_000183345.1
109 Escherichia coli UMN026 GCA_000026325.2
109 Escherichia coli str. K-12 substr. MG1655 GCA_000005845.2
109 Shigella flexneri 2a str. 301 GCA_000006925.2
109 Tumebacillus flagellatus GCA_000714935.1
96 Shigella dysenteriae Sd197 GCA_000012005.1
94 Cronobacter sakazakii GCA_000982825.1
94 Dickeya zeae Ech586 GCA_000025065.1
94 Erwinia amylovora CFBP1430 GCA_000091565.1
94 Erwinia toletana DAPP-PG 735 GCA_000336255.1
94 Kosakonia cowanii GCA_001975225.1
94 Pantoea ananatis LMG 20103 GCA_000025405.2
94 Pantoea dispersa EGD-AAK13 GCA_000465555.2
94 Plautia stali symbiont GCA_000180175.2
94 Pluralibacter gergoviae GCA_000757785.1
94 Shimwellia blattae DSM 4481 = NBRC 105725 GCA_000262305.1
U
285 Streptococcus mutans UA159 GCA_000007465.2
202 Streptococcus marimammalium DSM 18627 GCA_000380045.1
194 Streptococcus ferus DSM 20646 GCA_000372425.1
V
213 Rhodobacter sphaeroides 2.4.1 GCA_000012905.2
189 Rhodobacter sphaeroides ATCC 17025 GCA_000016405.1
169 Defluviimonas alba GCA_001620265.1
W
269 Bacillus thuringiensis YBT-1518 GCA_000497525.2
256 Bacillus mycoides GCA_000832605.1
247 Bacillus anthracis str. Ames GCA_000007845.1
247 Bacillus anthracis str. Sterne GCA_000008165.1
247 Bacillus cereus ATCC 14579 GCA_000007825.1
247 Bacillus pseudomycoides DSM 12442 GCA_000161455.1
247 _5BBacillus thuringiensis_5D serovar konkukian str. 97-27 GCA_000008505.1
X
125 Deinococcus radiodurans R1 GCA_000008565.1
85 Deinococcus misasensis DSM 22328 GCA_000745915.1
83 Deinococcus marmoris DSM 12784 GCA_000701405.1
83 Deinococcus swuensis GCA_000800395.1
Y
147 Deinococcus radiodurans R1 GCA_000008565.1
92 Deinococcus frigens DSM 12807 GCA_000701425.1
92 Deinococcus marmoris DSM 12784 GCA_000701405.1
92 Deinococcus swuensis GCA_000800395.1
88 Deinococcus hopiensis KR-140 GCA_900176165.1
88 Deinococcus puniceus GCA_001644565.1
88 Deinococcus soli Cha et al. 2016 GCA_001007995.1
Z
286 Streptococcus mutans UA159 GCA_000007465.2

202 Streptococcus ferus DSM 20646 GCA_000372425.1
201 Streptococcus gordonii str. Challis substr. CH1 GCA_000017005.1
10
268 Clostridium beijerinckii GCA_000833105.2
268 Clostridium saccharoperbutylacetonicum N1-4_28HMT_29 GCA_000340885.1
254 Clostridium saccharobutylicum DSM 13864 GCA_000473995.1
251 Clostridium puniceum GCA_002006345.1
11
177 Escherichia coli IAI39 GCA_000026345.1
177 Escherichia coli O104_3AH4 str. 2011C-3493 GCA_000299455.1
177 Escherichia coli O157_3AH7 str. Sakai GCA_000008865.1
177 Escherichia coli O83_3AH1 str. NRG 857C GCA_000183345.1
177 Escherichia coli UMN026 GCA_000026325.2
177 Escherichia coli str. K-12 substr. MG1655 GCA_000005845.2
177 Shigella dysenteriae Sd197 GCA_000012005.1
177 Shigella flexneri 2a str. 301 GCA_000006925.2
177 Tumebacillus flagellatus GCA_000714935.1
169 Erwinia iniecta GCA_001267535.1
158 Rosenbergiella nectarea GCA_900111105.1
12
131 Megasphaera cerevisiae DSM 20462 GCA_001045675.1
131 Staphylococcus arlettae CVD059 GCA_000295715.1
131 Staphylococcus aureus subsp. aureus NCTC 8325 GCA_000013425.1
131 Staphylococcus capitis subsp. capitis GCA_001028645.1
131 Staphylococcus cohnii subsp. cohnii GCA_000972575.1
131 Staphylococcus epidermidis ATCC 12228 GCA_000007645.1
131 Staphylococcus equorum GCA_001432245.1
131 Staphylococcus gallinarum GCA_000875895.1
131 Staphylococcus haemolyticus JCSC1435 GCA_000009865.1
131 Staphylococcus hominis subsp. hominis C80 GCA_000183685.1
131 Staphylococcus lentus F1142 GCA_000286395.1
131 Staphylococcus lugdunensis HKU09-01 GCA_000025085.1
131 Staphylococcus lutrae GCA_002101335.1
131 Staphylococcus saprophyticus GCA_001074355.1
131 Staphylococcus saprophyticus subsp. saprophyticus ATCC 15305 GCA_000010125.1
131 Staphylococcus sciuri GCA_002209165.1
131 Staphylococcus simiae CCM 7213 GCA_000235645.2
131 Staphylococcus succinus GCA_001902315.1
131 Staphylococcus warneri SG1 GCA_000332735.1
131 Staphylococcus xylosus GCA_000706685.1
117 Staphylococcus condimenti GCA_001618885.1
117 Staphylococcus microti GCA_000934465.1
117 Staphylococcus pettenkoferi GCA_002208805.1
117 Staphylococcus pseudintermedius HKU10-03 GCA_000185885.1
117 Staphylococcus simulans GCA_001559115.1
116 Staphylococcus auricularis GCA_001500315.1
13
223 Clostridium beijerinckii GCA_000833105.2
223 Clostridium puniceum GCA_002006345.1
223 Clostridium saccharoperbutylacetonicum N1-4_28HMT_29 GCA_000340885.1
214 Clostridium saccharobutylicum DSM 13864 GCA_000473995.1
204 Clostridium butyricum GCA_001456065.2
14
224 Streptococcus mutans UA159 GCA_000007465.2
140 Streptococcus ferus DSM 20646 GCA_000372425.1
139 Streptococcus gordonii str. Challis substr. CH1 GCA_000017005.1
15
299 Lactobacillus gasseri ATCC 33323 = JCM 1131 GCA_000014425.1
284 Lactobacillus hominis DSM 23910 = CRBIP 24.179 GCA_000296835.1
228 Lactobacillus acidophilus NCFM GCA_000011985.1
16
133 Rhodobacter sphaeroides 2.4.1 GCA_000012905.2
119 Rhodobacter sphaeroides ATCC 17025 GCA_000016405.1
106 Pseudorhodobacter ferrugineus DSM 5888 GCA_000420745.1
106 Pseudorhodobacter wandonensis GCA_001202035.1
17
179 Deinococcus radiodurans R1 GCA_000008565.1
87 Deinococcus deserti VCD115 GCA_000020685.1
87 Deinococcus frigens DSM 12807 GCA_000701425.1
87 Deinococcus geothermalis DSM 11300 GCA_000196275.1
87 Deinococcus gobiensis I-0 GCA_000252445.1
87 Deinococcus hopiensis KR-140 GCA_900176165.1
87 Deinococcus puniceus GCA_001644565.1
86 Deinococcus marmoris DSM 12784 GCA_000701405.1
86 Deinococcus misasensis DSM 22328 GCA_000745915.1
86 Deinococcus swuensis GCA_000800395.1

18
139 Rhodobacter sphaeroides 2.4.1 GCA_000012905.2
123 Rhodobacter sphaeroides ATCC 17025 GCA_000016405.1
108 Defluviimonas alba GCA_001620265.1
19
226 Escherichia coli IAI39 GCA_000026345.1
226 Escherichia coli O104_3AH4 str. 2011C-3493 GCA_000299455.1
226 Escherichia coli O157_3AH7 str. Sakai GCA_000008865.1
226 Escherichia coli O83_3AH1 str. NRG 857C GCA_000183345.1
226 Escherichia coli UMN026 GCA_000026325.2
226 Escherichia coli str. K-12 substr. MG1655 GCA_000005845.2
226 Shigella dysenteriae Sd197 GCA_000012005.1
226 Shigella flexneri 2a str. 301 GCA_000006925.2
226 Tumebacillus flagellatus GCA_000714935.1
222 Erwinia iniecta GCA_001267535.1
211 Erwinia gerundensis GCA_001517405.1
1a
236 Streptococcus mutans UA159 GCA_000007465.2
157 Streptococcus cristatus AS 1.3089 GCA_000385925.1
157 Streptococcus gordonii str. Challis substr. CH1 GCA_000017005.1
157 Streptococcus mitis B6 GCA_000027165.1
157 Streptococcus parasanguinis ATCC 15912 GCA_000164675.2
157 Streptococcus pneumoniae R6 GCA_000007045.1
141 Streptococcus sobrinus DSM 20742 = ATCC 33478 GCA_000686605.1
1b
188 Rhodobacter sphaeroides 2.4.1 GCA_000012905.2
150 Rhodobacter sphaeroides ATCC 17025 GCA_000016405.1
146 Rhodobacter capsulatus SB 1003 GCA_000021865.1
1c
100 Rhodobacter sphaeroides 2.4.1 GCA_000012905.2
97 Gemmobacter megaterium GCA_900156815.1
81 Pseudorhodobacter ferrugineus DSM 5888 GCA_000420745.1
81 Pseudorhodobacter wandonensis GCA_001202035.1
81 Rhodobacter sphaeroides ATCC 17025 GCA_000016405.1
1d
235 Clostridium beijerinckii GCA_000833105.2
235 Clostridium saccharobutylicum DSM 13864 GCA_000473995.1
235 Clostridium saccharoperbutylacetonicum N1-4_28HMT_29 GCA_000340885.1
229 Clostridium puniceum GCA_002006345.1
203 Clostridium chromiireducens GCA_002029255.1
1e
184 Clostridium saccharobutylicum DSM 13864 GCA_000473995.1
182 Clostridium beijerinckii GCA_000833105.2
182 Clostridium saccharoperbutylacetonicum N1-4_28HMT_29 GCA_000340885.1
175 Clostridium puniceum GCA_002006345.1
1f
64 Deinococcus radiodurans R1 GCA_000008565.1
41 Deinococcus gobiensis I-0 GCA_000252445.1
1g
218 Streptococcus mutans UA159 GCA_000007465.2
156 Streptococcus merionis DSM 19192 GCA_000380085.1
155 Streptococcus equinus GCA_000964315.1
155 Streptococcus gallolyticus subsp. gallolyticus DSM 16831 GCA_002000985.1
1h
260 Lactobacillus gasseri ATCC 33323 = JCM 1131 GCA_000014425.1
244 Lactobacillus hominis DSM 23910 = CRBIP 24.179 GCA_000296835.1
201 Lactobacillus crispatus ST1 GCA_000091765.1
1i
277 Enterococcus faecalis V583 GCA_000007785.1
277 Streptomyces cinnamoneus GCA_001885705.1
236 Enterococcus asini ATCC 700915 GCA_000407365.1
236 Enterococcus canis NBRC 100695 GCA_001544375.1
236 Enterococcus dispar ATCC 51266 GCA_000406945.1
236 Enterococcus faecium DO GCA_000174395.2
236 Enterococcus haemoperoxidus ATCC BAA-382 GCA_000407165.1
236 Enterococcus hirae ATCC 9790 GCA_000271405.2
236 Enterococcus mundtii QU 25 GCA_000504125.1
236 Enterococcus phoeniculicola ATCC BAA-412 GCA_000407505.1
236 Enterococcus rivorum GCA_001742285.1
236 Enterococcus thailandicus GCA_001652875.1
233 Melissococcus plutonius S1 GCA_000747585.1
1j
176 Bacillus thuringiensis YBT-1518 GCA_000497525.2
170 Bacillus anthracis str. Ames GCA_000007845.1
170 Bacillus anthracis str. Sterne GCA_000008165.1
170 Bacillus cereus ATCC 14579 GCA_000007825.1

170 Bacillus mycoides GCA_000832605.1
170 Bacillus pseudomycoides DSM 12442 GCA_000161455.1
170 _5BBacillus thuringiensis_5D serovar konkukian str. 97-27 GCA_000008505.1
165 Staphylococcus lentus F1142 GCA_000286395.1
165 Staphylococcus lutrae GCA_002101335.1
165 Staphylococcus sciuri GCA_002209165.1
1k
251 Staphylococcus epidermidis ATCC 12228 GCA_000007645.1
251 Staphylococcus haemolyticus JCSC1435 GCA_000009865.1
251 Staphylococcus hominis subsp. hominis C80 GCA_000183685.1
242 Staphylococcus cohnii subsp. cohnii GCA_000972575.1
240 Staphylococcus lugdunensis HKU09-01 GCA_000025085.1
1l
174 Clostridium beijerinckii GCA_000833105.2
174 Clostridium saccharobutylicum DSM 13864 GCA_000473995.1
174 Clostridium saccharoperbutylacetonicum N1-4_28HMT_29 GCA_000340885.1
170 Clostridium puniceum GCA_002006345.1
154 Clostridium chromiireducens GCA_002029255.1
1m
161 Lactobacillus gasseri ATCC 33323 = JCM 1131 GCA_000014425.1
148 Lactobacillus hominis DSM 23910 = CRBIP 24.179 GCA_000296835.1
97 Lactobacillus acidophilus NCFM GCA_000011985.1
97 Lactobacillus crispatus ST1 GCA_000091765.1
97 Lactobacillus gallinarum GCA_001314245.2
97 Lactobacillus hamsteri DSM 5661 = JCM 6256 GCA_000615445.1
1n
373 Lactobacillus gasseri ATCC 33323 = JCM 1131 GCA_000014425.1
358 Lactobacillus hominis DSM 23910 = CRBIP 24.179 GCA_000296835.1
300 Lactobacillus crispatus ST1 GCA_000091765.1
1o
241 Escherichia coli IAI39 GCA_000026345.1
241 Escherichia coli O104_3AH4 str. 2011C-3493 GCA_000299455.1
241 Escherichia coli O157_3AH7 str. Sakai GCA_000008865.1
241 Escherichia coli O83_3AH1 str. NRG 857C GCA_000183345.1
241 Escherichia coli UMN026 GCA_000026325.2
241 Escherichia coli str. K-12 substr. MG1655 GCA_000005845.2
241 Shigella dysenteriae Sd197 GCA_000012005.1
241 Shigella flexneri 2a str. 301 GCA_000006925.2
241 Tumebacillus flagellatus GCA_000714935.1
220 Erwinia iniecta GCA_001267535.1
212 Rosenbergiella nectarea GCA_900111105.1
1p
206 Clostridium butyricum GCA_001456065.2
205 Clostridium beijerinckii GCA_000833105.2
205 Clostridium saccharobutylicum DSM 13864 GCA_000473995.1
205 Clostridium saccharoperbutylacetonicum N1-4_28HMT_29 GCA_000340885.1
190 Clostridium chromiireducens GCA_002029255.1
190 Clostridium puniceum GCA_002006345.1
1q
163 Streptococcus mutans UA159 GCA_000007465.2
69 Enterococcus asini ATCC 700915 GCA_000407365.1
69 Enterococcus canis NBRC 100695 GCA_001544375.1
69 Enterococcus casseliflavus EC20 GCA_000157355.2
69 Enterococcus dispar ATCC 51266 GCA_000406945.1
69 Enterococcus faecalis V583 GCA_000007785.1
69 Enterococcus faecium DO GCA_000174395.2
69 Enterococcus gilvus ATCC BAA-350 GCA_000407545.1
69 Enterococcus hermanniensis GCA_001885945.1
69 Enterococcus hirae ATCC 9790 GCA_000271405.2
69 Enterococcus italicus DSM 15952 GCA_000185365.1
69 Enterococcus malodoratus ATCC 43197 GCA_000407185.1
69 Enterococcus massiliensis GCA_001050095.1
69 Enterococcus mundtii QU 25 GCA_000504125.1
69 Enterococcus pallens ATCC BAA-351 GCA_000407485.1
69 Enterococcus pseudoavium NBRC 100491 GCA_001544295.1
69 Enterococcus rivorum GCA_001742285.1
69 Enterococcus saccharolyticus subsp. saccharolyticus ATCC 43076 GCA_000407285.1
69 Melissococcus plutonius S1 GCA_000747585.1
69 Streptomyces cinnamoneus GCA_001885705.1
66 Streptococcus macacae NCTC 11558 GCA_000187995.3
1r
314 Staphylococcus capitis subsp. capitis GCA_001028645.1
314 Staphylococcus epidermidis ATCC 12228 GCA_000007645.1
314 Staphylococcus haemolyticus JCSC1435 GCA_000009865.1
303 Staphylococcus hominis subsp. hominis C80 GCA_000183685.1
296 Staphylococcus lugdunensis HKU09-01 GCA_000025085.1

1s
246 Clostridium butyricum GCA_001456065.2
240 Clostridium beijerinckii GCA_000833105.2
240 Clostridium saccharoperbutylacetonicum N1-4_28HMT_29 GCA_000340885.1
236 Clostridium puniceum GCA_002006345.1
1t
265 Bifidobacterium adolescentis ATCC 15703 GCA_000010425.1
257 Bifidobacterium tsurumiense GCA_000741765.1
256 Bifidobacterium choerinum GCA_000741135.1
256 Bifidobacterium pseudolongum PV8-2 GCA_000800475.2
1u
221 Clostridium beijerinckii GCA_000833105.2
221 Clostridium puniceum GCA_002006345.1
221 Clostridium saccharobutylicum DSM 13864 GCA_000473995.1
221 Clostridium saccharoperbutylacetonicum N1-4_28HMT_29 GCA_000340885.1
210 Clostridium chromiireducens GCA_002029255.1
210 Clostridium taeniosporum GCA_001735765.1
199 Clostridium botulinum B str. Eklund 17B _28NRP_29 GCA_000020165.1
1v
261 Escherichia coli IAI39 GCA_000026345.1
261 Escherichia coli O104_3AH4 str. 2011C-3493 GCA_000299455.1
261 Escherichia coli O157_3AH7 str. Sakai GCA_000008865.1
261 Escherichia coli O83_3AH1 str. NRG 857C GCA_000183345.1
261 Escherichia coli UMN026 GCA_000026325.2
261 Escherichia coli str. K-12 substr. MG1655 GCA_000005845.2
261 Shigella flexneri 2a str. 301 GCA_000006925.2
261 Tumebacillus flagellatus GCA_000714935.1
252 Shigella dysenteriae Sd197 GCA_000012005.1
242 Erwinia iniecta GCA_001267535.1
1w
207 Deinococcus radiodurans R1 GCA_000008565.1
123 Deinococcus gobiensis I-0 GCA_000252445.1
120 Deinococcus deserti VCD115 GCA_000020685.1
1x
208 Bifidobacterium adolescentis ATCC 15703 GCA_000010425.1
193 Bifidobacterium dentium JCM 1195 = DSM 20436 GCA_001042595.1
183 Bifidobacterium gallicum DSM 20093 = LMG 11596 GCA_000741205.1
1y
284 Clostridium puniceum GCA_002006345.1
280 Clostridium beijerinckii GCA_000833105.2
280 Clostridium saccharobutylicum DSM 13864 GCA_000473995.1
280 Clostridium saccharoperbutylacetonicum N1-4_28HMT_29 GCA_000340885.1
265 Clostridium chromiireducens GCA_002029255.1
1z
298 Staphylococcus epidermidis ATCC 12228 GCA_000007645.1
293 Megasphaera cerevisiae DSM 20462 GCA_001045675.1
293 Staphylococcus capitis subsp. capitis GCA_001028645.1
293 Staphylococcus lugdunensis HKU09-01 GCA_000025085.1
293 Staphylococcus warneri SG1 GCA_000332735.1
291 Staphylococcus aureus subsp. aureus NCTC 8325 GCA_000013425.1
291 Staphylococcus haemolyticus JCSC1435 GCA_000009865.1
291 Staphylococcus hominis subsp. hominis C80 GCA_000183685.1
291 Staphylococcus simiae CCM 7213 GCA_000235645.2
1A
124 Clostridium butyricum GCA_001456065.2
118 Clostridium chromiireducens GCA_002029255.1
118 Clostridium saccharobutylicum DSM 13864 GCA_000473995.1
116 Clostridium beijerinckii GCA_000833105.2
116 Clostridium puniceum GCA_002006345.1
116 Clostridium saccharoperbutylacetonicum N1-4_28HMT_29 GCA_000340885.1
1B
95 Staphylococcus pseudintermedius HKU10-03 GCA_000185885.1
93 Megasphaera cerevisiae DSM 20462 GCA_001045675.1
93 Staphylococcus capitis subsp. capitis GCA_001028645.1
93 Staphylococcus cohnii subsp. cohnii GCA_000972575.1
93 Staphylococcus epidermidis ATCC 12228 GCA_000007645.1
93 Staphylococcus warneri SG1 GCA_000332735.1
78 Nosocomiicoccus massiliensis GCA_000438455.1
78 Staphylococcus aureus subsp. aureus NCTC 8325 GCA_000013425.1
78 Staphylococcus condimenti GCA_001618885.1
78 Staphylococcus haemolyticus JCSC1435 GCA_000009865.1
78 Staphylococcus hominis subsp. hominis C80 GCA_000183685.1
78 Staphylococcus hyicus GCA_000816085.1
78 Staphylococcus lugdunensis HKU09-01 GCA_000025085.1
78 Staphylococcus lutrae GCA_002101335.1
78 Staphylococcus microti GCA_000934465.1

78 Staphylococcus pettenkoferi GCA_002208805.1
78 Staphylococcus saprophyticus GCA_001074355.1
78 Staphylococcus saprophyticus subsp. saprophyticus ATCC 15305 GCA_000010125.1
78 Staphylococcus sciuri GCA_002209165.1
78 Staphylococcus simiae CCM 7213 GCA_000235645.2
78 Staphylococcus simulans GCA_001559115.1
78 Staphylococcus xylosus GCA_000706685.1
1C
267 Lactobacillus gasseri ATCC 33323 = JCM 1131 GCA_000014425.1
255 Lactobacillus hominis DSM 23910 = CRBIP 24.179 GCA_000296835.1
216 Lactobacillus acidophilus NCFM GCA_000011985.1
216 Lactobacillus gallinarum GCA_001314245.2
1D
290 Clostridium beijerinckii GCA_000833105.2
290 Clostridium puniceum GCA_002006345.1
290 Clostridium saccharoperbutylacetonicum N1-4_28HMT_29 GCA_000340885.1
282 Clostridium saccharobutylicum DSM 13864 GCA_000473995.1
263 Clostridium chromiireducens GCA_002029255.1
1E
166 Rhodobacter sphaeroides 2.4.1 GCA_000012905.2
161 Rhodobacter sphaeroides ATCC 17025 GCA_000016405.1
149 Gemmobacter megaterium GCA_900156815.1
1F
184 Clostridium beijerinckii GCA_000833105.2
184 Clostridium puniceum GCA_002006345.1
184 Clostridium saccharoperbutylacetonicum N1-4_28HMT_29 GCA_000340885.1
175 Clostridium chromiireducens GCA_002029255.1
175 Clostridium saccharobutylicum DSM 13864 GCA_000473995.1
169 Clostridium butyricum GCA_001456065.2
1G
295 Bacillus anthracis str. Ames GCA_000007845.1
295 Bacillus anthracis str. Sterne GCA_000008165.1
295 Bacillus cereus ATCC 14579 GCA_000007825.1
295 Bacillus mycoides GCA_000832605.1
295 Bacillus pseudomycoides DSM 12442 GCA_000161455.1
295 Bacillus thuringiensis YBT-1518 GCA_000497525.2
295 _5BBacillus thuringiensis_5D serovar konkukian str. 97-27 GCA_000008505.1
231 Bacillus manliponensis GCA_000712595.1
231 Staphylococcus condimenti GCA_001618885.1
229 Massilibacterium senegalense GCA_001375675.1
1H
213 Streptococcus mutans UA159 GCA_000007465.2
149 Streptococcus macacae NCTC 11558 GCA_000187995.3
138 Streptococcus sobrinus DSM 20742 = ATCC 33478 GCA_000686605.1
1I
241 Bacillus anthracis str. Ames GCA_000007845.1
241 Bacillus anthracis str. Sterne GCA_000008165.1
241 Bacillus cereus ATCC 14579 GCA_000007825.1
241 Bacillus mycoides GCA_000832605.1
241 Bacillus pseudomycoides DSM 12442 GCA_000161455.1
241 Bacillus thuringiensis YBT-1518 GCA_000497525.2
241 _5BBacillus thuringiensis_5D serovar konkukian str. 97-27 GCA_000008505.1
209 Bacillus manliponensis GCA_000712595.1
200 Bacillus cytotoxicus NVH 391-98 GCA_000017425.1
1J
162 Rhodobacter sphaeroides 2.4.1 GCA_000012905.2
129 Pseudorhodobacter psychrotolerans GCA_001294535.1
119 Gemmobacter aquatilis GCA_900110025.1
119 Rhodobacter sphaeroides ATCC 17025 GCA_000016405.1
1K
231 Clostridium saccharobutylicum DSM 13864 GCA_000473995.1
229 Clostridium beijerinckii GCA_000833105.2
229 Clostridium puniceum GCA_002006345.1
229 Clostridium saccharoperbutylacetonicum N1-4_28HMT_29 GCA_000340885.1
224 Clostridium chromiireducens GCA_002029255.1
1L
296 Streptococcus mutans UA159 GCA_000007465.2
183 Streptococcus gordonii str. Challis substr. CH1 GCA_000017005.1
179 Streptococcus cristatus AS 1.3089 GCA_000385925.1
1M
191 Streptococcus mutans UA159 GCA_000007465.2
132 Streptococcus marimammalium DSM 18627 GCA_000380045.1
126 Streptococcus salivarius GCA_000785515.1
126 Streptococcus thermophilus JIM 8232 GCA_000253395.1
1N
216 Escherichia coli IAI39 GCA_000026345.1

216 Escherichia coli O104_3AH4 str. 2011C-3493 GCA_000299455.1
216 Escherichia coli O157_3AH7 str. Sakai GCA_000008865.1
216 Escherichia coli O83_3AH1 str. NRG 857C GCA_000183345.1
216 Escherichia coli UMN026 GCA_000026325.2
216 Escherichia coli str. K-12 substr. MG1655 GCA_000005845.2
216 Shigella dysenteriae Sd197 GCA_000012005.1
216 Shigella flexneri 2a str. 301 GCA_000006925.2
216 Tumebacillus flagellatus GCA_000714935.1
189 Erwinia iniecta GCA_001267535.1
172 Enterobacter hormaechei subsp. steigerwaltii GCA_001729725.1
172 Erwinia toletana DAPP-PG 735 GCA_000336255.1
172 Klebsiella oxytoca GCA_001022195.1
172 Kosakonia cowanii GCA_001975225.1
172 Kosakonia sacchari SP1 GCA_000300455.4
172 Pluralibacter gergoviae GCA_000757785.1
172 Pseudescherichia vulneris NBRC 102420 GCA_000759795.1
1O
101 Deinococcus radiodurans R1 GCA_000008565.1
1P
267 Bifidobacterium adolescentis ATCC 15703 GCA_000010425.1
224 Bifidobacterium breve DSM 20213 = JCM 1192 GCA_001025175.1
214 Bifidobacterium callitrichos DSM 23973 GCA_000741175.1
1Q
218 Bacillus anthracis str. Ames GCA_000007845.1
218 Bacillus anthracis str. Sterne GCA_000008165.1
218 Bacillus cereus ATCC 14579 GCA_000007825.1
218 Bacillus mycoides GCA_000832605.1
218 Bacillus pseudomycoides DSM 12442 GCA_000161455.1
218 Bacillus thuringiensis YBT-1518 GCA_000497525.2
218 _5BBacillus thuringiensis_5D serovar konkukian str. 97-27 GCA_000008505.1
199 Bacillus manliponensis GCA_000712595.1
164 Bacillus horneckiae GCA_001636335.1
164 Bacillus massiliogorillae GCA_000380245.2
164 Bacillus weihaiensis GCA_001889165.1
1R
240 Escherichia coli IAI39 GCA_000026345.1
240 Escherichia coli O104_3AH4 str. 2011C-3493 GCA_000299455.1
240 Escherichia coli O157_3AH7 str. Sakai GCA_000008865.1
240 Escherichia coli O83_3AH1 str. NRG 857C GCA_000183345.1
240 Escherichia coli UMN026 GCA_000026325.2
240 Escherichia coli str. K-12 substr. MG1655 GCA_000005845.2
240 Shigella flexneri 2a str. 301 GCA_000006925.2
240 Tumebacillus flagellatus GCA_000714935.1
231 Shigella dysenteriae Sd197 GCA_000012005.1
194 Erwinia iniecta GCA_001267535.1
1S
209 Bacillus thuringiensis YBT-1518 GCA_000497525.2
203 Bacillus anthracis str. Ames GCA_000007845.1
203 Bacillus anthracis str. Sterne GCA_000008165.1
203 Bacillus cereus ATCC 14579 GCA_000007825.1
203 Bacillus mycoides GCA_000832605.1
203 Bacillus pseudomycoides DSM 12442 GCA_000161455.1
203 _5BBacillus thuringiensis_5D serovar konkukian str. 97-27 GCA_000008505.1
171 Bacillus cytotoxicus NVH 391-98 GCA_000017425.1
1T
136 Rhodobacter sphaeroides 2.4.1 GCA_000012905.2
106 Gemmobacter aquatilis GCA_900110025.1
106 Pseudorhodobacter psychrotolerans GCA_001294535.1
97 Haematobacter massiliensis GCA_000740795.1
1U
199 Enterococcus faecalis V583 GCA_000007785.1
199 Streptomyces cinnamoneus GCA_001885705.1
187 Bavariicoccus seileri DSM 19936 GCA_000421665.1
187 Isobaculum melis GCA_900111355.1
184 Enterococcus hirae ATCC 9790 GCA_000271405.2
1V
156 Rhodobacter sphaeroides ATCC 17025 GCA_000016405.1
155 Rhodobacter sphaeroides 2.4.1 GCA_000012905.2
150 Haematobacter massiliensis GCA_000740795.1
1W
191 Deinococcus radiodurans R1 GCA_000008565.1
154 Deinococcus gobiensis I-0 GCA_000252445.1
118 Deinococcus puniceus GCA_001644565.1
1X
230 Clostridium neonatale GCA_001458595.1
225 Clostridium beijerinckii GCA_000833105.2

225 Clostridium puniceum GCA_002006345.1
225 Clostridium saccharobutylicum DSM 13864 GCA_000473995.1
225 Clostridium saccharoperbutylacetonicum N1-4_28HMT_29 GCA_000340885.1
214 Clostridium butyricum GCA_001456065.2
1Y
312 Deinococcus radiodurans R1 GCA_000008565.1
171 Deinococcus gobiensis I-0 GCA_000252445.1
167 Deinococcus deserti VCD115 GCA_000020685.1
1Z
321 Enterococcus faecalis V583 GCA_000007785.1
321 Streptomyces cinnamoneus GCA_001885705.1
293 Enterococcus canis NBRC 100695 GCA_001544375.1
293 Enterococcus casseliflavus EC20 GCA_000157355.2
293 Enterococcus dispar ATCC 51266 GCA_000406945.1
293 Enterococcus faecium DO GCA_000174395.2
293 Enterococcus haemoperoxidus ATCC BAA-382 GCA_000407165.1
293 Enterococcus hirae ATCC 9790 GCA_000271405.2
293 Enterococcus mundtii QU 25 GCA_000504125.1
293 Enterococcus phoeniculicola ATCC BAA-412 GCA_000407505.1
293 Enterococcus rivorum GCA_001742285.1
293 Enterococcus saccharolyticus subsp. saccharolyticus ATCC 43076 GCA_000407285.1
293 Enterococcus thailandicus GCA_001652875.1
284 Enterococcus massiliensis GCA_001050095.1
20
188 Deinococcus radiodurans R1 GCA_000008565.1
123 Deinococcus gobiensis I-0 GCA_000252445.1
117 Deinococcus marmoris DSM 12784 GCA_000701405.1
117 Deinococcus swuensis GCA_000800395.1
21
218 Bifidobacterium adolescentis ATCC 15703 GCA_000010425.1
178 Bifidobacterium callitrichos DSM 23973 GCA_000741175.1
178 Bifidobacterium thermophilum GCA_000741495.1
175 Bifidobacterium breve DSM 20213 = JCM 1192 GCA_001025175.1
22
174 Clostridium beijerinckii GCA_000833105.2
174 Clostridium chromiireducens GCA_002029255.1
174 Clostridium puniceum GCA_002006345.1
174 Clostridium saccharobutylicum DSM 13864 GCA_000473995.1
174 Clostridium saccharoperbutylacetonicum N1-4_28HMT_29 GCA_000340885.1
167 Clostridium butyricum GCA_001456065.2
154 Clostridium taeniosporum GCA_001735765.1
23
96 Rhodobacter capsulatus SB 1003 GCA_000021865.1
87 Rhodobacter sphaeroides 2.4.1 GCA_000012905.2
86 Gemmobacter megaterium GCA_900156815.1
86 Gemmobacter nectariphilus DSM 15620 GCA_000429765.1
86 Haematobacter massiliensis GCA_000740795.1
86 Paracoccus yeei GCA_002073635.1
24
164 Bacillus anthracis str. Ames GCA_000007845.1
164 Bacillus anthracis str. Sterne GCA_000008165.1
164 Bacillus cereus ATCC 14579 GCA_000007825.1
164 Bacillus mycoides GCA_000832605.1
164 Bacillus pseudomycoides DSM 12442 GCA_000161455.1
164 Bacillus thuringiensis YBT-1518 GCA_000497525.2
164 _5BBacillus thuringiensis_5D serovar konkukian str. 97-27 GCA_000008505.1
137 Bacillus manliponensis GCA_000712595.1
133 Bacillus horneckiae GCA_001636335.1
133 Bacillus solani GCA_001420595.1
25
256 Rhodobacter sphaeroides 2.4.1 GCA_000012905.2
212 Rhodobacter sphaeroides ATCC 17025 GCA_000016405.1
196 Pseudorhodobacter psychrotolerans GCA_001294535.1
26
181 Bifidobacterium adolescentis ATCC 15703 GCA_000010425.1
177 Bifidobacterium callitrichos DSM 23973 GCA_000741175.1
166 Bifidobacterium longum NCC2705 GCA_000007525.1
166 Bifidobacterium thermophilum GCA_000741495.1
27
333 Bacillus thuringiensis YBT-1518 GCA_000497525.2
318 Bacillus anthracis str. Ames GCA_000007845.1
318 Bacillus anthracis str. Sterne GCA_000008165.1
318 Bacillus cereus ATCC 14579 GCA_000007825.1
318 Bacillus mycoides GCA_000832605.1
318 Bacillus pseudomycoides DSM 12442 GCA_000161455.1
318 _5BBacillus thuringiensis_5D serovar konkukian str. 97-27 GCA_000008505.1

279 Bacillus marisflavi GCA_001274775.1
28
197 Bifidobacterium adolescentis ATCC 15703 GCA_000010425.1
168 Bifidobacterium dentium JCM 1195 = DSM 20436 GCA_001042595.1
163 Bifidobacterium callitrichos DSM 23973 GCA_000741175.1
29
247 Bifidobacterium adolescentis ATCC 15703 GCA_000010425.1
220 Bifidobacterium angulatum DSM 20098 = JCM 7096 GCA_001025155.1
220 Bifidobacterium asteroides PRL2011 GCA_000304215.1
220 Bifidobacterium callitrichos DSM 23973 GCA_000741175.1
216 Bifidobacterium coryneforme GCA_000737865.1
2a
229 Clostridium saccharobutylicum DSM 13864 GCA_000473995.1
227 Clostridium butyricum GCA_001456065.2
224 Clostridium beijerinckii GCA_000833105.2
224 Clostridium saccharoperbutylacetonicum N1-4_28HMT_29 GCA_000340885.1
2b
172 Rhodobacter sphaeroides 2.4.1 GCA_000012905.2
172 Rhodobacter sphaeroides ATCC 17025 GCA_000016405.1
152 Pseudorhodobacter psychrotolerans GCA_001294535.1
144 Pseudorhodobacter ferrugineus DSM 5888 GCA_000420745.1
144 Pseudorhodobacter wandonensis GCA_001202035.1
2c
235 Clostridium beijerinckii GCA_000833105.2
235 Clostridium puniceum GCA_002006345.1
235 Clostridium saccharoperbutylacetonicum N1-4_28HMT_29 GCA_000340885.1
226 Clostridium saccharobutylicum DSM 13864 GCA_000473995.1
207 Clostridium chromiireducens GCA_002029255.1
2d
255 Bacillus anthracis str. Ames GCA_000007845.1
255 Bacillus anthracis str. Sterne GCA_000008165.1
255 Bacillus cereus ATCC 14579 GCA_000007825.1
255 Bacillus mycoides GCA_000832605.1
255 Bacillus pseudomycoides DSM 12442 GCA_000161455.1
255 Bacillus thuringiensis YBT-1518 GCA_000497525.2
255 _5BBacillus thuringiensis_5D serovar konkukian str. 97-27 GCA_000008505.1
197 Bacillus manliponensis GCA_000712595.1
185 Bacillus aquimaris TF-12 GCA_001648555.1
2e
180 Clostridium beijerinckii GCA_000833105.2
180 Clostridium chromiireducens GCA_002029255.1
180 Clostridium puniceum GCA_002006345.1
180 Clostridium saccharobutylicum DSM 13864 GCA_000473995.1
180 Clostridium saccharoperbutylacetonicum N1-4_28HMT_29 GCA_000340885.1
149 Clostridium butyricum GCA_001456065.2
146 Clostridium neonatale GCA_001458595.1
2f
248 Deinococcus radiodurans R1 GCA_000008565.1
165 Deinococcus gobiensis I-0 GCA_000252445.1
158 Deinococcus deserti VCD115 GCA_000020685.1
2g
302 Deinococcus radiodurans R1 GCA_000008565.1
195 Deinococcus gobiensis I-0 GCA_000252445.1
171 Deinococcus proteolyticus MRP GCA_000190555.1
2h
244 Escherichia coli IAI39 GCA_000026345.1
244 Escherichia coli O104_3AH4 str. 2011C-3493 GCA_000299455.1
244 Escherichia coli O157_3AH7 str. Sakai GCA_000008865.1
244 Escherichia coli O83_3AH1 str. NRG 857C GCA_000183345.1
244 Escherichia coli UMN026 GCA_000026325.2
244 Escherichia coli str. K-12 substr. MG1655 GCA_000005845.2
244 Shigella flexneri 2a str. 301 GCA_000006925.2
244 Tumebacillus flagellatus GCA_000714935.1
235 Shigella dysenteriae Sd197 GCA_000012005.1
220 Erwinia iniecta GCA_001267535.1
2i
264 Clostridium beijerinckii GCA_000833105.2
264 Clostridium saccharobutylicum DSM 13864 GCA_000473995.1
264 Clostridium saccharoperbutylacetonicum N1-4_28HMT_29 GCA_000340885.1
260 Clostridium puniceum GCA_002006345.1
249 Clostridium chromiireducens GCA_002029255.1
2j
270 Clostridium beijerinckii GCA_000833105.2
270 Clostridium saccharoperbutylacetonicum N1-4_28HMT_29 GCA_000340885.1
264 Clostridium puniceum GCA_002006345.1
262 Clostridium saccharobutylicum DSM 13864 GCA_000473995.1

2k
333 Lactobacillus gasseri ATCC 33323 = JCM 1131 GCA_000014425.1
320 Lactobacillus hominis DSM 23910 = CRBIP 24.179 GCA_000296835.1
240 Lactobacillus crispatus ST1 GCA_000091765.1
2l
227 Bacillus anthracis str. Ames GCA_000007845.1
227 Bacillus anthracis str. Sterne GCA_000008165.1
227 Bacillus mycoides GCA_000832605.1
227 Bacillus pseudomycoides DSM 12442 GCA_000161455.1
227 Bacillus thuringiensis YBT-1518 GCA_000497525.2
227 _5BBacillus thuringiensis_5D serovar konkukian str. 97-27 GCA_000008505.1
226 Bacillus cereus ATCC 14579 GCA_000007825.1
169 Bacillus halmapalus GCA_002019665.1
169 Bacillus vietnamensis NBRC 101237 GCA_001591825.1
2m
139 Bifidobacterium adolescentis ATCC 15703 GCA_000010425.1
129 Bifidobacterium thermophilum GCA_000741495.1
129 Bifidobacterium thermophilum RBL67 GCA_000347695.1
120 Bifidobacterium dentium JCM 1195 = DSM 20436 GCA_001042595.1
120 Bifidobacterium tsurumiense GCA_000741765.1
2n
169 Clostridium beijerinckii GCA_000833105.2
169 Clostridium chromiireducens GCA_002029255.1
169 Clostridium puniceum GCA_002006345.1
169 Clostridium saccharobutylicum DSM 13864 GCA_000473995.1
169 Clostridium saccharoperbutylacetonicum N1-4_28HMT_29 GCA_000340885.1
159 Clostridium butyricum GCA_001456065.2
133 Clostridium neonatale GCA_001458595.1
2o
196 Bifidobacterium adolescentis ATCC 15703 GCA_000010425.1
192 Bifidobacterium thermophilum GCA_000741495.1
181 Bifidobacterium dentium JCM 1195 = DSM 20436 GCA_001042595.1
2p
78 Clostridium beijerinckii GCA_000833105.2
78 Clostridium puniceum GCA_002006345.1
78 Clostridium saccharobutylicum DSM 13864 GCA_000473995.1
78 Clostridium saccharoperbutylacetonicum N1-4_28HMT_29 GCA_000340885.1
77 Clostridium taeniosporum GCA_001735765.1
74 Clostridium neonatale GCA_001458595.1
2q
68 Clostridium beijerinckii GCA_000833105.2
68 Clostridium neonatale GCA_001458595.1
68 Clostridium puniceum GCA_002006345.1
68 Clostridium saccharobutylicum DSM 13864 GCA_000473995.1
68 Clostridium saccharoperbutylacetonicum N1-4_28HMT_29 GCA_000340885.1
53 Clostridium butyricum GCA_001456065.2
53 Clostridium chromiireducens GCA_002029255.1
46 Clostridium intestinale URNW GCA_000469625.2
46 Desnuesiella massiliensis GCA_001403615.1
2r
229 Deinococcus radiodurans R1 GCA_000008565.1
158 Deinococcus marmoris DSM 12784 GCA_000701405.1
158 Deinococcus swuensis GCA_000800395.1
140 Deinococcus soli Cha et al. 2016 GCA_001007995.1
2s
186 Bifidobacterium adolescentis ATCC 15703 GCA_000010425.1
181 Bifidobacterium longum NCC2705 GCA_000007525.1
180 Bifidobacterium breve DSM 20213 = JCM 1192 GCA_001025175.1
2t
286 Bacillus mycoides GCA_000832605.1
285 Bacillus anthracis str. Ames GCA_000007845.1
285 Bacillus anthracis str. Sterne GCA_000008165.1
285 Bacillus cereus ATCC 14579 GCA_000007825.1
285 Bacillus pseudomycoides DSM 12442 GCA_000161455.1
285 Bacillus thuringiensis YBT-1518 GCA_000497525.2
285 _5BBacillus thuringiensis_5D serovar konkukian str. 97-27 GCA_000008505.1
250 Bacillus halmapalus GCA_002019665.1
2u
208 Staphylococcus condimenti GCA_001618885.1
207 Staphylococcus aureus subsp. aureus NCTC 8325 GCA_000013425.1
207 Staphylococcus capitis subsp. capitis GCA_001028645.1
207 Staphylococcus epidermidis ATCC 12228 GCA_000007645.1
207 Staphylococcus haemolyticus JCSC1435 GCA_000009865.1
207 Staphylococcus hominis subsp. hominis C80 GCA_000183685.1
207 Staphylococcus lugdunensis HKU09-01 GCA_000025085.1
207 Staphylococcus simiae CCM 7213 GCA_000235645.2

205 Megasphaera cerevisiae DSM 20462 GCA_001045675.1
205 Staphylococcus warneri SG1 GCA_000332735.1
2v
98 Deinococcus radiodurans R1 GCA_000008565.1
83 Deinococcus hopiensis KR-140 GCA_900176165.1
71 Deinococcus deserti VCD115 GCA_000020685.1
71 Deinococcus gobiensis I-0 GCA_000252445.1
71 Deinococcus soli Cha et al. 2016 GCA_001007995.1
2w
317 Bacillus anthracis str. Ames GCA_000007845.1
317 Bacillus anthracis str. Sterne GCA_000008165.1
317 Bacillus cereus ATCC 14579 GCA_000007825.1
317 Bacillus mycoides GCA_000832605.1
317 Bacillus pseudomycoides DSM 12442 GCA_000161455.1
317 Bacillus thuringiensis YBT-1518 GCA_000497525.2
317 _5BBacillus thuringiensis_5D serovar konkukian str. 97-27 GCA_000008505.1
256 Bacillus manliponensis GCA_000712595.1
240 Bacillus cytotoxicus NVH 391-98 GCA_000017425.1
2x
137 Cronobacter sakazakii GCA_000982825.1
137 Escherichia coli IAI39 GCA_000026345.1
137 Escherichia coli O104_3AH4 str. 2011C-3493 GCA_000299455.1
137 Escherichia coli O157_3AH7 str. Sakai GCA_000008865.1
137 Escherichia coli O83_3AH1 str. NRG 857C GCA_000183345.1
137 Escherichia coli UMN026 GCA_000026325.2
137 Escherichia coli str. K-12 substr. MG1655 GCA_000005845.2
137 Shigella dysenteriae Sd197 GCA_000012005.1
137 Shigella flexneri 2a str. 301 GCA_000006925.2
137 Shimwellia blattae DSM 4481 = NBRC 105725 GCA_000262305.1
137 Tumebacillus flagellatus GCA_000714935.1
127 Obesumbacterium proteus GCA_001586165.1
127 Rahnella aquatilis HX2 GCA_000255535.1
127 Rouxiella chamberiensis GCA_000951135.1
127 Serratia fonticola GCA_001006005.1
127 Serratia liquefaciens ATCC 27592 GCA_000422085.1
127 Serratia marcescens subsp. marcescens Db11 GCA_000513215.1
127 Serratia rubidaea GCA_001572725.1
127 Serratia symbiotica GCA_000821185.1
127 Yersinia enterocolitica subsp. enterocolitica 8081 GCA_000009345.1
127 Yersinia pestis CO92 GCA_000009065.1
127 Yersinia ruckeri GCA_000964565.1
122 Erwinia gerundensis GCA_001517405.1
122 Erwinia iniecta GCA_001267535.1
122 Erwinia toletana DAPP-PG 735 GCA_000336255.1
122 Pantoea agglomerans GCA_001709315.1
122 Pantoea ananatis LMG 20103 GCA_000025405.2
122 Pantoea dispersa EGD-AAK13 GCA_000465555.2
122 Pectobacterium carotovorum subsp. carotovorum PC1 GCA_000023605.1
122 Plautia stali symbiont GCA_000180175.2
2y
119 Deinococcus radiodurans R1 GCA_000008565.1
78 Deinococcus deserti VCD115 GCA_000020685.1
78 Deinococcus hopiensis KR-140 GCA_900176165.1
72 Deinococcus soli Cha et al. 2016 GCA_001007995.1
2z
295 Enterococcus faecalis V583 GCA_000007785.1
295 Streptomyces cinnamoneus GCA_001885705.1
257 Enterococcus dispar ATCC 51266 GCA_000406945.1
257 Enterococcus hirae ATCC 9790 GCA_000271405.2
254 Enterococcus canis NBRC 100695 GCA_001544375.1
254 Enterococcus casseliflavus EC20 GCA_000157355.2
254 Enterococcus faecium DO GCA_000174395.2
254 Enterococcus haemoperoxidus ATCC BAA-382 GCA_000407165.1
254 Enterococcus mundtii QU 25 GCA_000504125.1
254 Enterococcus phoeniculicola ATCC BAA-412 GCA_000407505.1
254 Enterococcus rivorum GCA_001742285.1
254 Enterococcus saccharolyticus subsp. saccharolyticus ATCC 43076 GCA_000407285.1
254 Enterococcus thailandicus GCA_001652875.1
2A
232 Clostridium beijerinckii GCA_000833105.2
232 Clostridium puniceum GCA_002006345.1
232 Clostridium saccharoperbutylacetonicum N1-4_28HMT_29 GCA_000340885.1
224 Clostridium saccharobutylicum DSM 13864 GCA_000473995.1
214 Clostridium butyricum GCA_001456065.2
2B
144 Megasphaera cerevisiae DSM 20462 GCA_001045675.1

144 Staphylococcus aureus subsp. aureus NCTC 8325 GCA_000013425.1
144 Staphylococcus capitis subsp. capitis GCA_001028645.1
144 Staphylococcus epidermidis ATCC 12228 GCA_000007645.1
144 Staphylococcus haemolyticus JCSC1435 GCA_000009865.1
144 Staphylococcus hominis subsp. hominis C80 GCA_000183685.1
144 Staphylococcus lugdunensis HKU09-01 GCA_000025085.1
144 Staphylococcus simiae CCM 7213 GCA_000235645.2
144 Staphylococcus warneri SG1 GCA_000332735.1
139 Staphylococcus hyicus GCA_000816085.1
139 Staphylococcus lutrae GCA_002101335.1
132 Staphylococcus condimenti GCA_001618885.1
132 Staphylococcus simulans GCA_001559115.1
2C
161 Deinococcus radiodurans R1 GCA_000008565.1
114 Deinococcus gobiensis I-0 GCA_000252445.1
99 Deinococcus deserti VCD115 GCA_000020685.1
2D
172 Clostridium saccharobutylicum DSM 13864 GCA_000473995.1
170 Clostridium beijerinckii GCA_000833105.2
170 Clostridium saccharoperbutylacetonicum N1-4_28HMT_29 GCA_000340885.1
163 Clostridium puniceum GCA_002006345.1
2E
221 Bacillus thuringiensis YBT-1518 GCA_000497525.2
206 Bacillus anthracis str. Ames GCA_000007845.1
206 Bacillus anthracis str. Sterne GCA_000008165.1
206 Bacillus cereus ATCC 14579 GCA_000007825.1
206 Bacillus mycoides GCA_000832605.1
206 Bacillus pseudomycoides DSM 12442 GCA_000161455.1
206 _5BBacillus thuringiensis_5D serovar konkukian str. 97-27 GCA_000008505.1
171 Bacillus manliponensis GCA_000712595.1
2F
222 Deinococcus radiodurans R1 GCA_000008565.1
133 Deinococcus gobiensis I-0 GCA_000252445.1
115 Deinococcus deserti VCD115 GCA_000020685.1
2G
204 Escherichia coli IAI39 GCA_000026345.1
204 Escherichia coli O104_3AH4 str. 2011C-3493 GCA_000299455.1
204 Escherichia coli O157_3AH7 str. Sakai GCA_000008865.1
204 Escherichia coli O83_3AH1 str. NRG 857C GCA_000183345.1
204 Escherichia coli UMN026 GCA_000026325.2
204 Escherichia coli str. K-12 substr. MG1655 GCA_000005845.2
204 Shigella dysenteriae Sd197 GCA_000012005.1
204 Tumebacillus flagellatus GCA_000714935.1
191 Shigella flexneri 2a str. 301 GCA_000006925.2
182 Erwinia iniecta GCA_001267535.1
2H
152 Rhodobacter sphaeroides 2.4.1 GCA_000012905.2
137 Defluviimonas alba GCA_001620265.1
122 Haematobacter massiliensis GCA_000740795.1
2I
128 Deinococcus radiodurans R1 GCA_000008565.1
88 Deinococcus deserti VCD115 GCA_000020685.1
84 Deinococcus hopiensis KR-140 GCA_900176165.1
2J
200 Deinococcus radiodurans R1 GCA_000008565.1
115 Deinococcus puniceus GCA_001644565.1
115 Deinococcus soli Cha et al. 2016 GCA_001007995.1
112 Deinococcus deserti VCD115 GCA_000020685.1
112 Deinococcus gobiensis I-0 GCA_000252445.1
112 Deinococcus hopiensis KR-140 GCA_900176165.1
2K
232 Streptococcus mutans UA159 GCA_000007465.2
173 Streptococcus equinus GCA_000964315.1
173 Streptococcus gallolyticus subsp. gallolyticus DSM 16831 GCA_002000985.1
158 Streptococcus orisratti DSM 15617 GCA_000380105.1
158 Streptococcus ratti FA-1 = DSM 20564 GCA_000286075.1
2L
279 Lactobacillus gasseri ATCC 33323 = JCM 1131 GCA_000014425.1
247 Lactobacillus hominis DSM 23910 = CRBIP 24.179 GCA_000296835.1
163 Lactobacillus iners DSM 13335 GCA_000160875.1
2M
238 Escherichia coli IAI39 GCA_000026345.1
238 Escherichia coli O104_3AH4 str. 2011C-3493 GCA_000299455.1
238 Escherichia coli O157_3AH7 str. Sakai GCA_000008865.1
238 Escherichia coli O83_3AH1 str. NRG 857C GCA_000183345.1
238 Escherichia coli UMN026 GCA_000026325.2

238 Escherichia coli str. K-12 substr. MG1655 GCA_000005845.2
238 Shigella flexneri 2a str. 301 GCA_000006925.2
238 Tumebacillus flagellatus GCA_000714935.1
223 Shigella dysenteriae Sd197 GCA_000012005.1
190 Erwinia iniecta GCA_001267535.1
2N
176 Clostridium chromiireducens GCA_002029255.1
174 Clostridium beijerinckii GCA_000833105.2
174 Clostridium butyricum GCA_001456065.2
174 Clostridium puniceum GCA_002006345.1
174 Clostridium saccharoperbutylacetonicum N1-4_28HMT_29 GCA_000340885.1
165 Clostridium saccharobutylicum DSM 13864 GCA_000473995.1
2O
230 Bifidobacterium adolescentis ATCC 15703 GCA_000010425.1
218 Bifidobacterium choerinum GCA_000741135.1
218 Bifidobacterium pseudolongum PV8-2 GCA_000800475.2
217 Bifidobacterium angulatum DSM 20098 = JCM 7096 GCA_001025155.1
2P
200 Lactobacillus gasseri ATCC 33323 = JCM 1131 GCA_000014425.1
200 Lactobacillus hominis DSM 23910 = CRBIP 24.179 GCA_000296835.1
190 Lactobacillus psittaci DSM 15354 GCA_000425905.1
187 Lactobacillus amylolyticus GCA_002075105.1
2Q
160 Bifidobacterium adolescentis ATCC 15703 GCA_000010425.1
147 Bifidobacterium thermophilum GCA_000741495.1
122 Bifidobacterium tsurumiense GCA_000741765.1
2R
263 Clostridium beijerinckii GCA_000833105.2
263 Clostridium saccharoperbutylacetonicum N1-4_28HMT_29 GCA_000340885.1
248 Clostridium saccharobutylicum DSM 13864 GCA_000473995.1
246 Clostridium puniceum GCA_002006345.1
2S
277 Bacillus thuringiensis YBT-1518 GCA_000497525.2
274 Bacillus anthracis str. Ames GCA_000007845.1
274 Bacillus anthracis str. Sterne GCA_000008165.1
274 Bacillus cereus ATCC 14579 GCA_000007825.1
274 Bacillus mycoides GCA_000832605.1
274 Bacillus pseudomycoides DSM 12442 GCA_000161455.1
274 _5BBacillus thuringiensis_5D serovar konkukian str. 97-27 GCA_000008505.1
220 Bacillus manliponensis GCA_000712595.1
2T
203 Escherichia coli IAI39 GCA_000026345.1
203 Escherichia coli O104_3AH4 str. 2011C-3493 GCA_000299455.1
203 Escherichia coli O157_3AH7 str. Sakai GCA_000008865.1
203 Escherichia coli O83_3AH1 str. NRG 857C GCA_000183345.1
203 Escherichia coli UMN026 GCA_000026325.2
203 Escherichia coli str. K-12 substr. MG1655 GCA_000005845.2
203 Shigella dysenteriae Sd197 GCA_000012005.1
203 Shigella flexneri 2a str. 301 GCA_000006925.2
203 Tumebacillus flagellatus GCA_000714935.1
183 Erwinia iniecta GCA_001267535.1
169 Brenneria goodwinii GCA_001049335.1
169 Enterobacter hormaechei subsp. steigerwaltii GCA_001729725.1
169 Erwinia toletana DAPP-PG 735 GCA_000336255.1
169 Klebsiella oxytoca GCA_001022195.1
169 Kosakonia cowanii GCA_001975225.1
169 Kosakonia sacchari SP1 GCA_000300455.4
169 Pantoea agglomerans GCA_001709315.1
169 Pantoea alhagi GCA_002101395.1
169 Pantoea rwandensis GCA_000759475.1
169 Pantoea septica GCA_002095575.1
169 Plesiomonas shigelloides GCA_900087055.1
169 Pseudescherichia vulneris NBRC 102420 GCA_000759795.1
169 Shimwellia blattae DSM 4481 = NBRC 105725 GCA_000262305.1
2U
236 Deinococcus radiodurans R1 GCA_000008565.1
155 Deinococcus puniceus GCA_001644565.1
147 Deinococcus deserti VCD115 GCA_000020685.1
147 Deinococcus geothermalis DSM 11300 GCA_000196275.1
2V
253 Bacillus anthracis str. Ames GCA_000007845.1
253 Bacillus anthracis str. Sterne GCA_000008165.1
253 Bacillus cereus ATCC 14579 GCA_000007825.1
253 Bacillus mycoides GCA_000832605.1
253 Bacillus pseudomycoides DSM 12442 GCA_000161455.1
253 Bacillus thuringiensis YBT-1518 GCA_000497525.2

253 _5BBacillus thuringiensis_5D serovar konkukian str. 97-27 GCA_000008505.1
203 Bacillus infantis NRRL B-14911 GCA_000473245.1
194 Bacillus coahuilensis m4-4 GCA_000171615.1
194 Bacillus vietnamensis NBRC 101237 GCA_001591825.1
2W
258 Clostridium beijerinckii GCA_000833105.2
258 Clostridium saccharoperbutylacetonicum N1-4_28HMT_29 GCA_000340885.1
254 Clostridium butyricum GCA_001456065.2
254 Clostridium puniceum GCA_002006345.1
254 Clostridium saccharobutylicum DSM 13864 GCA_000473995.1
245 Clostridium chromiireducens GCA_002029255.1
2X
130 Rhodobacter sphaeroides 2.4.1 GCA_000012905.2
130 Rhodobacter sphaeroides ATCC 17025 GCA_000016405.1
124 Clostridium beijerinckii GCA_000833105.2
124 Clostridium chromiireducens GCA_002029255.1
124 Clostridium puniceum GCA_002006345.1
124 Clostridium saccharobutylicum DSM 13864 GCA_000473995.1
124 Clostridium saccharoperbutylacetonicum N1-4_28HMT_29 GCA_000340885.1
114 Clostridium butyricum GCA_001456065.2
2Y
187 Clostridium beijerinckii GCA_000833105.2
187 Clostridium puniceum GCA_002006345.1
187 Clostridium saccharoperbutylacetonicum N1-4_28HMT_29 GCA_000340885.1
178 Clostridium saccharobutylicum DSM 13864 GCA_000473995.1
175 Clostridium butyricum GCA_001456065.2
175 Clostridium taeniosporum GCA_001735765.1
2Z
225 Clostridium beijerinckii GCA_000833105.2
225 Clostridium puniceum GCA_002006345.1
225 Clostridium saccharobutylicum DSM 13864 GCA_000473995.1
225 Clostridium saccharoperbutylacetonicum N1-4_28HMT_29 GCA_000340885.1
217 Clostridium chromiireducens GCA_002029255.1
207 Clostridium butyricum GCA_001456065.2
30
297 Enterococcus faecalis V583 GCA_000007785.1
297 Streptomyces cinnamoneus GCA_001885705.1
266 Enterococcus canis NBRC 100695 GCA_001544375.1
266 Enterococcus casseliflavus EC20 GCA_000157355.2
266 Enterococcus dispar ATCC 51266 GCA_000406945.1
266 Enterococcus faecium DO GCA_000174395.2
266 Enterococcus gilvus ATCC BAA-350 GCA_000407545.1
266 Enterococcus hermanniensis GCA_001885945.1
266 Enterococcus hirae ATCC 9790 GCA_000271405.2
266 Enterococcus malodoratus ATCC 43197 GCA_000407185.1
266 Enterococcus mundtii QU 25 GCA_000504125.1
266 Enterococcus pallens ATCC BAA-351 GCA_000407485.1
266 Enterococcus pseudoavium NBRC 100491 GCA_001544295.1
266 Enterococcus rivorum GCA_001742285.1
266 Enterococcus saccharolyticus subsp. saccharolyticus ATCC 43076 GCA_000407285.1
261 Enterococcus haemoperoxidus ATCC BAA-382 GCA_000407165.1
261 Enterococcus phoeniculicola ATCC BAA-412 GCA_000407505.1
261 Enterococcus thailandicus GCA_001652875.1
31
315 Lactobacillus gasseri ATCC 33323 = JCM 1131 GCA_000014425.1
300 Lactobacillus hominis DSM 23910 = CRBIP 24.179 GCA_000296835.1
243 Lactobacillus psittaci DSM 15354 GCA_000425905.1
32
260 Deinococcus radiodurans R1 GCA_000008565.1
198 Deinococcus gobiensis I-0 GCA_000252445.1
196 Deinococcus deserti VCD115 GCA_000020685.1
33
212 Staphylococcus capitis subsp. capitis GCA_001028645.1
212 Staphylococcus epidermidis ATCC 12228 GCA_000007645.1
212 Staphylococcus haemolyticus JCSC1435 GCA_000009865.1
201 Staphylococcus hominis subsp. hominis C80 GCA_000183685.1
201 Staphylococcus lugdunensis HKU09-01 GCA_000025085.1
195 Staphylococcus condimenti GCA_001618885.1
195 Staphylococcus simulans GCA_001559115.1
34
170 Erwinia iniecta GCA_001267535.1
170 Escherichia coli IAI39 GCA_000026345.1
170 Escherichia coli O104_3AH4 str. 2011C-3493 GCA_000299455.1
170 Escherichia coli O157_3AH7 str. Sakai GCA_000008865.1
170 Escherichia coli UMN026 GCA_000026325.2
170 Escherichia coli str. K-12 substr. MG1655 GCA_000005845.2

170 Shigella flexneri 2a str. 301 GCA_000006925.2
170 Tumebacillus flagellatus GCA_000714935.1
169 Escherichia coli O83_3AH1 str. NRG 857C GCA_000183345.1
156 Shigella dysenteriae Sd197 GCA_000012005.1
35
75 Deinococcus radiodurans R1 GCA_000008565.1
57 Deinococcus gobiensis I-0 GCA_000252445.1
42 Deinococcus deserti VCD115 GCA_000020685.1
42 Deinococcus puniceus GCA_001644565.1
36
201 Bifidobacterium longum NCC2705 GCA_000007525.1
199 Bifidobacterium adolescentis ATCC 15703 GCA_000010425.1
199 Bifidobacterium breve DSM 20213 = JCM 1192 GCA_001025175.1
192 Bifidobacterium angulatum DSM 20098 = JCM 7096 GCA_001025155.1
37
251 Escherichia coli IAI39 GCA_000026345.1
251 Escherichia coli O104_3AH4 str. 2011C-3493 GCA_000299455.1
251 Escherichia coli O157_3AH7 str. Sakai GCA_000008865.1
251 Escherichia coli O83_3AH1 str. NRG 857C GCA_000183345.1
251 Escherichia coli UMN026 GCA_000026325.2
251 Escherichia coli str. K-12 substr. MG1655 GCA_000005845.2
251 Shigella flexneri 2a str. 301 GCA_000006925.2
251 Tumebacillus flagellatus GCA_000714935.1
241 Shigella dysenteriae Sd197 GCA_000012005.1
228 Erwinia iniecta GCA_001267535.1
38
178 Clostridium beijerinckii GCA_000833105.2
178 Clostridium saccharobutylicum DSM 13864 GCA_000473995.1
178 Clostridium saccharoperbutylacetonicum N1-4_28HMT_29 GCA_000340885.1
172 Clostridium puniceum GCA_002006345.1
142 Clostridium chromiireducens GCA_002029255.1
39
196 Bacillus thuringiensis YBT-1518 GCA_000497525.2
173 Bacillus anthracis str. Ames GCA_000007845.1
173 Bacillus anthracis str. Sterne GCA_000008165.1
173 Bacillus cereus ATCC 14579 GCA_000007825.1
173 Bacillus mycoides GCA_000832605.1
173 Bacillus pseudomycoides DSM 12442 GCA_000161455.1
173 _5BBacillus thuringiensis_5D serovar konkukian str. 97-27 GCA_000008505.1
147 Bacillus horneckiae GCA_001636335.1
147 Bacillus solani GCA_001420595.1
3a
158 Lactobacillus gasseri ATCC 33323 = JCM 1131 GCA_000014425.1
158 Lactobacillus hominis DSM 23910 = CRBIP 24.179 GCA_000296835.1
98 Lactobacillus amylophilus DSM 20533 = JCM 1125 GCA_001936335.1
98 Lactobacillus iners DSM 13335 GCA_000160875.1
98 Lactobacillus siliginis GCA_001437435.1
91 Lactobacillus antri DSM 16041 GCA_000160835.1
91 Lactobacillus brevis ATCC 367 GCA_000014465.1
91 Lactobacillus coleohominis 101-4-CHN GCA_000161935.1
91 Lactobacillus curieae GCA_000785105.2
91 Lactobacillus fabifermentans DSM 21115 GCA_000498955.2
91 Lactobacillus fermentum IFO 3956 GCA_000010145.1
91 Lactobacillus frumenti DSM 13145 GCA_001436045.1
91 Lactobacillus gastricus DSM 16045 GCA_001434365.1
91 Lactobacillus herbarum GCA_001039045.1
91 Lactobacillus ingluviei str. Autruche 4 GCA_000312405.1
91 Lactobacillus koreensis GCA_001050435.1
91 Lactobacillus mucosae LM1 GCA_000248095.3
91 Lactobacillus odoratitofui DSM 19909 = JCM 15043 GCA_001434895.1
91 Lactobacillus ozensis DSM 23829 = JCM 17196 GCA_001435995.1
91 Lactobacillus paucivorans GCA_001437125.1
91 Lactobacillus plantarum WCFS1 GCA_000203855.3
91 Lactobacillus pontis DSM 8475 GCA_001435345.1
91 Lactobacillus rennini DSM 20253 GCA_001436505.1
91 Lactobacillus reuteri DSM 20016 GCA_000016825.1
91 Lactobacillus secaliphilus GCA_001437055.1
91 Lactobacillus senioris DSM 24302 = JCM 17472 GCA_001436555.1
91 Lactobacillus senmaizukei DSM 21775 = NBRC 103853 GCA_001592085.1
91 Lactobacillus silagei JCM 19001 GCA_002217945.1
91 Lactobacillus similis DSM 23365 = JCM 2765 GCA_001311075.1
91 Lactobacillus spicheri DSM 15429 GCA_001435095.1
91 Lactobacillus vaginalis DSM 5837 = ATCC 49540 GCA_000159435.1
91 Lactobacillus xiangfangensis GCA_001438845.1
91 Lactobacillus zymae GCA_900183405.1
91 Pediococcus acidilactici GCA_001767275.1

91 Pediococcus argentinicus GCA_001437605.1
91 Pediococcus claussenii ATCC BAA-344 GCA_000237995.2
91 Pediococcus pentosaceus ATCC 25745 GCA_000014505.1
91 Pediococcus stilesii GCA_001437075.1
3b
223 Bacillus anthracis str. Ames GCA_000007845.1
223 Bacillus anthracis str. Sterne GCA_000008165.1
223 Bacillus cereus ATCC 14579 GCA_000007825.1
223 Bacillus mycoides GCA_000832605.1
223 Bacillus pseudomycoides DSM 12442 GCA_000161455.1
223 Bacillus thuringiensis YBT-1518 GCA_000497525.2
223 _5BBacillus thuringiensis_5D serovar konkukian str. 97-27 GCA_000008505.1
178 Bacillus manliponensis GCA_000712595.1
172 Bacillus hemicellulosilyticus JCM 9152 GCA_000513115.1
3c
246 Streptococcus mutans UA159 GCA_000007465.2
176 Streptococcus macacae NCTC 11558 GCA_000187995.3
175 Streptococcus ratti FA-1 = DSM 20564 GCA_000286075.1
3d
262 Bifidobacterium adolescentis ATCC 15703 GCA_000010425.1
246 Bifidobacterium dentium JCM 1195 = DSM 20436 GCA_001042595.1
233 Bifidobacterium thermophilum GCA_000741495.1
233 Bifidobacterium thermophilum RBL67 GCA_000347695.1
3e
205 Bifidobacterium dentium JCM 1195 = DSM 20436 GCA_001042595.1
204 Bifidobacterium adolescentis ATCC 15703 GCA_000010425.1
199 Bifidobacterium angulatum DSM 20098 = JCM 7096 GCA_001025155.1
3f
238 Bifidobacterium adolescentis ATCC 15703 GCA_000010425.1
206 Bifidobacterium thermophilum GCA_000741495.1
197 Bifidobacterium breve DSM 20213 = JCM 1192 GCA_001025175.1
3g
190 Streptococcus mutans UA159 GCA_000007465.2
102 Oceanobacillus caeni GCA_001298135.1
93 Streptococcus ferus DSM 20646 GCA_000372425.1
3h
232 Bifidobacterium adolescentis ATCC 15703 GCA_000010425.1
213 Bifidobacterium thermophilum GCA_000741495.1
197 Bifidobacterium asteroides PRL2011 GCA_000304215.1
3i
117 Bacillus anthracis str. Ames GCA_000007845.1
117 Bacillus anthracis str. Sterne GCA_000008165.1
117 Bacillus cereus ATCC 14579 GCA_000007825.1
117 Bacillus manliponensis GCA_000712595.1
117 Bacillus mycoides GCA_000832605.1
117 Bacillus pseudomycoides DSM 12442 GCA_000161455.1
117 Bacillus thuringiensis YBT-1518 GCA_000497525.2
117 _5BBacillus thuringiensis_5D serovar konkukian str. 97-27 GCA_000008505.1
114 Facklamia hominis CCUG 36813 GCA_000301035.1
112 Globicatella sulfidifaciens DSM 15739 GCA_900167405.1
3j
149 Clostridium beijerinckii GCA_000833105.2
149 Clostridium butyricum GCA_001456065.2
149 Clostridium chromiireducens GCA_002029255.1
149 Clostridium puniceum GCA_002006345.1
149 Clostridium saccharobutylicum DSM 13864 GCA_000473995.1
149 Clostridium saccharoperbutylacetonicum N1-4_28HMT_29 GCA_000340885.1
147 Clostridium ventriculi GCA_001404895.1
142 Clostridium cavendishii DSM 21758 GCA_900141845.1
3k
266 Clostridium beijerinckii GCA_000833105.2
266 Clostridium saccharoperbutylacetonicum N1-4_28HMT_29 GCA_000340885.1
252 Clostridium saccharobutylicum DSM 13864 GCA_000473995.1
251 Clostridium puniceum GCA_002006345.1
3l
124 Clostridium beijerinckii GCA_000833105.2
124 Clostridium puniceum GCA_002006345.1
124 Clostridium saccharobutylicum DSM 13864 GCA_000473995.1
124 Clostridium saccharoperbutylacetonicum N1-4_28HMT_29 GCA_000340885.1
116 Clostridium chromiireducens GCA_002029255.1
107 Clostridium butyricum GCA_001456065.2
3m
252 Escherichia coli IAI39 GCA_000026345.1
252 Escherichia coli O104_3AH4 str. 2011C-3493 GCA_000299455.1
252 Escherichia coli O157_3AH7 str. Sakai GCA_000008865.1
252 Escherichia coli O83_3AH1 str. NRG 857C GCA_000183345.1

252 Escherichia coli UMN026 GCA_000026325.2
252 Escherichia coli str. K-12 substr. MG1655 GCA_000005845.2
252 Shigella flexneri 2a str. 301 GCA_000006925.2
252 Tumebacillus flagellatus GCA_000714935.1
243 Shigella dysenteriae Sd197 GCA_000012005.1
220 Erwinia iniecta GCA_001267535.1
3n
224 Enterococcus faecalis V583 GCA_000007785.1
224 Streptomyces cinnamoneus GCA_001885705.1
200 Enterococcus canis NBRC 100695 GCA_001544375.1
200 Enterococcus dispar ATCC 51266 GCA_000406945.1
200 Enterococcus faecium DO GCA_000174395.2
200 Enterococcus haemoperoxidus ATCC BAA-382 GCA_000407165.1
200 Enterococcus hirae ATCC 9790 GCA_000271405.2
200 Enterococcus mundtii QU 25 GCA_000504125.1
200 Enterococcus phoeniculicola ATCC BAA-412 GCA_000407505.1
200 Enterococcus rivorum GCA_001742285.1
200 Enterococcus thailandicus GCA_001652875.1
191 Enterococcus massiliensis GCA_001050095.1
3o
118 Rhodobacter capsulatus SB 1003 GCA_000021865.1
118 Rhodobacter sphaeroides 2.4.1 GCA_000012905.2
103 Gemmobacter aquatilis GCA_900110025.1
103 Gemmobacter nectariphilus DSM 15620 GCA_000429765.1
103 Haematobacter massiliensis GCA_000740795.1
103 Pseudorhodobacter psychrotolerans GCA_001294535.1
100 Gemmobacter megaterium GCA_900156815.1
100 Paracoccus yeei GCA_002073635.1
100 Rhodobacter sphaeroides ATCC 17025 GCA_000016405.1
3p
259 Bacillus anthracis str. Ames GCA_000007845.1
259 Bacillus anthracis str. Sterne GCA_000008165.1
259 Bacillus cereus ATCC 14579 GCA_000007825.1
259 Bacillus mycoides GCA_000832605.1
259 Bacillus pseudomycoides DSM 12442 GCA_000161455.1
259 Bacillus thuringiensis YBT-1518 GCA_000497525.2
259 _5BBacillus thuringiensis_5D serovar konkukian str. 97-27 GCA_000008505.1
231 Bacillus manliponensis GCA_000712595.1
230 Bacillus aquimaris TF-12 GCA_001648555.1
230 Bacillus coahuilensis m4-4 GCA_000171615.1
230 Bacillus marisflavi GCA_001274775.1
230 Bacillus simplex GCA_001578185.1
230 Bacillus testis GCA_001243895.1
230 Bacillus vietnamensis NBRC 101237 GCA_001591825.1
3q
264 Rhodobacter sphaeroides 2.4.1 GCA_000012905.2
243 Rhodobacter sphaeroides ATCC 17025 GCA_000016405.1
225 Pseudorhodobacter ferrugineus DSM 5888 GCA_000420745.1
225 Pseudorhodobacter psychrotolerans GCA_001294535.1
225 Pseudorhodobacter wandonensis GCA_001202035.1
3r
206 Escherichia coli IAI39 GCA_000026345.1
206 Escherichia coli O104_3AH4 str. 2011C-3493 GCA_000299455.1
206 Escherichia coli O157_3AH7 str. Sakai GCA_000008865.1
206 Escherichia coli O83_3AH1 str. NRG 857C GCA_000183345.1
206 Escherichia coli UMN026 GCA_000026325.2
206 Escherichia coli str. K-12 substr. MG1655 GCA_000005845.2
206 Shigella flexneri 2a str. 301 GCA_000006925.2
206 Tumebacillus flagellatus GCA_000714935.1
197 Shigella dysenteriae Sd197 GCA_000012005.1
191 Erwinia iniecta GCA_001267535.1
3s
196 Clostridium beijerinckii GCA_000833105.2
196 Clostridium puniceum GCA_002006345.1
196 Clostridium saccharoperbutylacetonicum N1-4_28HMT_29 GCA_000340885.1
187 Clostridium saccharobutylicum DSM 13864 GCA_000473995.1
161 Clostridium taeniosporum GCA_001735765.1
3t
144 Erwinia iniecta GCA_001267535.1
144 Escherichia coli IAI39 GCA_000026345.1
144 Escherichia coli O104_3AH4 str. 2011C-3493 GCA_000299455.1
144 Escherichia coli O157_3AH7 str. Sakai GCA_000008865.1
144 Escherichia coli UMN026 GCA_000026325.2
144 Escherichia coli str. K-12 substr. MG1655 GCA_000005845.2
144 Shigella dysenteriae Sd197 GCA_000012005.1
144 Shigella flexneri 2a str. 301 GCA_000006925.2

144 Tumebacillus flagellatus GCA_000714935.1
143 Escherichia coli O83_3AH1 str. NRG 857C GCA_000183345.1
141 Rosenbergiella nectarea GCA_900111105.1
3u
331 Clostridium beijerinckii GCA_000833105.2
331 Clostridium puniceum GCA_002006345.1
331 Clostridium saccharoperbutylacetonicum N1-4_28HMT_29 GCA_000340885.1
322 Clostridium saccharobutylicum DSM 13864 GCA_000473995.1
307 Clostridium chromiireducens GCA_002029255.1
3v
265 Bifidobacterium adolescentis ATCC 15703 GCA_000010425.1
246 Bifidobacterium dentium JCM 1195 = DSM 20436 GCA_001042595.1
246 Bifidobacterium tsurumiense GCA_000741765.1
241 Bifidobacterium longum NCC2705 GCA_000007525.1
3w
161 Staphylococcus hominis subsp. hominis C80 GCA_000183685.1
156 Staphylococcus arlettae CVD059 GCA_000295715.1
156 Staphylococcus microti GCA_000934465.1
153 Bacillus thuringiensis YBT-1518 GCA_000497525.2
3x
285 Escherichia coli IAI39 GCA_000026345.1
285 Escherichia coli O104_3AH4 str. 2011C-3493 GCA_000299455.1
285 Escherichia coli O157_3AH7 str. Sakai GCA_000008865.1
285 Escherichia coli O83_3AH1 str. NRG 857C GCA_000183345.1
285 Escherichia coli UMN026 GCA_000026325.2
285 Escherichia coli str. K-12 substr. MG1655 GCA_000005845.2
285 Shigella dysenteriae Sd197 GCA_000012005.1
285 Shigella flexneri 2a str. 301 GCA_000006925.2
285 Tumebacillus flagellatus GCA_000714935.1
264 Erwinia iniecta GCA_001267535.1
244 Cronobacter sakazakii GCA_000982825.1
244 Erwinia toletana DAPP-PG 735 GCA_000336255.1
3y
379 Lactobacillus gasseri ATCC 33323 = JCM 1131 GCA_000014425.1
365 Lactobacillus hominis DSM 23910 = CRBIP 24.179 GCA_000296835.1
282 Lactobacillus psittaci DSM 15354 GCA_000425905.1
3z
217 Deinococcus radiodurans R1 GCA_000008565.1
138 Deinococcus gobiensis I-0 GCA_000252445.1
129 Deinococcus marmoris DSM 12784 GCA_000701405.1
129 Deinococcus swuensis GCA_000800395.1
3A
238 Deinococcus radiodurans R1 GCA_000008565.1
172 Deinococcus puniceus GCA_001644565.1
165 Deinococcus gobiensis I-0 GCA_000252445.1
3B
259 Enterococcus faecalis V583 GCA_000007785.1
259 Streptomyces cinnamoneus GCA_001885705.1
232 Enterococcus hirae ATCC 9790 GCA_000271405.2
231 Enterococcus asini ATCC 700915 GCA_000407365.1
231 Enterococcus canis NBRC 100695 GCA_001544375.1
231 Enterococcus dispar ATCC 51266 GCA_000406945.1
231 Enterococcus faecium DO GCA_000174395.2
231 Enterococcus haemoperoxidus ATCC BAA-382 GCA_000407165.1
231 Enterococcus massiliensis GCA_001050095.1
231 Enterococcus mundtii QU 25 GCA_000504125.1
231 Enterococcus phoeniculicola ATCC BAA-412 GCA_000407505.1
231 Enterococcus rivorum GCA_001742285.1
231 Enterococcus thailandicus GCA_001652875.1
3C
218 Clostridium beijerinckii GCA_000833105.2
218 Clostridium puniceum GCA_002006345.1
218 Clostridium saccharoperbutylacetonicum N1-4_28HMT_29 GCA_000340885.1
209 Clostridium saccharobutylicum DSM 13864 GCA_000473995.1
194 Clostridium chromiireducens GCA_002029255.1
3D
157 Escherichia coli IAI39 GCA_000026345.1
157 Escherichia coli O104_3AH4 str. 2011C-3493 GCA_000299455.1
157 Escherichia coli O157_3AH7 str. Sakai GCA_000008865.1
157 Escherichia coli O83_3AH1 str. NRG 857C GCA_000183345.1
157 Escherichia coli UMN026 GCA_000026325.2
157 Escherichia coli str. K-12 substr. MG1655 GCA_000005845.2
157 Shigella dysenteriae Sd197 GCA_000012005.1
157 Shigella flexneri 2a str. 301 GCA_000006925.2
157 Tumebacillus flagellatus GCA_000714935.1
139 Erwinia iniecta GCA_001267535.1

136 Rosenbergiella nectarea GCA_900111105.1
3E
165 Clostridium beijerinckii GCA_000833105.2
165 Clostridium puniceum GCA_002006345.1
165 Clostridium saccharoperbutylacetonicum N1-4_28HMT_29 GCA_000340885.1
156 Clostridium chromiireducens GCA_002029255.1
156 Clostridium saccharobutylicum DSM 13864 GCA_000473995.1
156 Clostridium taeniosporum GCA_001735765.1
151 Bacillus anthracis str. Ames GCA_000007845.1
151 Bacillus anthracis str. Sterne GCA_000008165.1
151 Bacillus cereus ATCC 14579 GCA_000007825.1
151 Bacillus mycoides GCA_000832605.1
151 Bacillus pseudomycoides DSM 12442 GCA_000161455.1
151 Bacillus thuringiensis YBT-1518 GCA_000497525.2
151 _5BBacillus thuringiensis_5D serovar konkukian str. 97-27 GCA_000008505.1
3F
211 Deinococcus radiodurans R1 GCA_000008565.1
143 Deinococcus gobiensis I-0 GCA_000252445.1
124 Deinococcus marmoris DSM 12784 GCA_000701405.1
124 Deinococcus swuensis GCA_000800395.1
3G
307 Bifidobacterium adolescentis ATCC 15703 GCA_000010425.1
281 Bifidobacterium callitrichos DSM 23973 GCA_000741175.1
280 Bifidobacterium coryneforme GCA_000737865.1
3H
264 Streptococcus mutans UA159 GCA_000007465.2
175 Streptococcus ferus DSM 20646 GCA_000372425.1
170 Streptococcus macacae NCTC 11558 GCA_000187995.3
3I
232 Deinococcus radiodurans R1 GCA_000008565.1
137 Deinococcus deserti VCD115 GCA_000020685.1
134 Deinococcus marmoris DSM 12784 GCA_000701405.1
134 Deinococcus swuensis GCA_000800395.1
3J
145 Rhodobacter sphaeroides 2.4.1 GCA_000012905.2
124 Litorimicrobium taeanense GCA_900110775.1
123 Thioclava dalianensis GCA_000715505.1
123 Thioclava indica GCA_000714545.1
3K
242 Lactobacillus gasseri ATCC 33323 = JCM 1131 GCA_000014425.1
226 Lactobacillus hominis DSM 23910 = CRBIP 24.179 GCA_000296835.1
184 Lactobacillus acidophilus NCFM GCA_000011985.1
184 Lactobacillus crispatus ST1 GCA_000091765.1
184 Lactobacillus hamsteri DSM 5661 = JCM 6256 GCA_000615445.1
184 Lactobacillus helveticus GCA_001308285.1
3L
290 Clostridium beijerinckii GCA_000833105.2
290 Clostridium saccharoperbutylacetonicum N1-4_28HMT_29 GCA_000340885.1
286 Clostridium puniceum GCA_002006345.1
282 Clostridium saccharobutylicum DSM 13864 GCA_000473995.1
3M
351 Bacillus anthracis str. Ames GCA_000007845.1
351 Bacillus anthracis str. Sterne GCA_000008165.1
351 Bacillus cereus ATCC 14579 GCA_000007825.1
351 Bacillus mycoides GCA_000832605.1
351 Bacillus pseudomycoides DSM 12442 GCA_000161455.1
351 Bacillus thuringiensis YBT-1518 GCA_000497525.2
351 _5BBacillus thuringiensis_5D serovar konkukian str. 97-27 GCA_000008505.1
266 Bacillus marisflavi GCA_001274775.1
260 Bacillus coahuilensis m4-4 GCA_000171615.1
3N
184 Rhodobacter sphaeroides 2.4.1 GCA_000012905.2
178 Pseudorhodobacter ferrugineus DSM 5888 GCA_000420745.1
178 Pseudorhodobacter wandonensis GCA_001202035.1
178 Rhodobacter sphaeroides ATCC 17025 GCA_000016405.1
167 Pseudorhodobacter psychrotolerans GCA_001294535.1
3O
129 Escherichia coli IAI39 GCA_000026345.1
129 Escherichia coli O104_3AH4 str. 2011C-3493 GCA_000299455.1
129 Escherichia coli O157_3AH7 str. Sakai GCA_000008865.1
129 Escherichia coli O83_3AH1 str. NRG 857C GCA_000183345.1
129 Escherichia coli UMN026 GCA_000026325.2
129 Escherichia coli str. K-12 substr. MG1655 GCA_000005845.2
129 Shigella dysenteriae Sd197 GCA_000012005.1
129 Shigella flexneri 2a str. 301 GCA_000006925.2
129 Tumebacillus flagellatus GCA_000714935.1

115 Erwinia iniecta GCA_001267535.1
115 Rosenbergiella nectarea GCA_900111105.1
113 Cedecea neteri GCA_000757825.1
113 Edwardsiella anguillarum ET080813 GCA_000264765.2
3P
213 Clostridium butyricum GCA_001456065.2
208 Clostridium beijerinckii GCA_000833105.2
208 Clostridium puniceum GCA_002006345.1
208 Clostridium saccharoperbutylacetonicum N1-4_28HMT_29 GCA_000340885.1
200 Clostridium chromiireducens GCA_002029255.1
200 Clostridium saccharobutylicum DSM 13864 GCA_000473995.1
3Q
223 Escherichia coli IAI39 GCA_000026345.1
223 Escherichia coli O104_3AH4 str. 2011C-3493 GCA_000299455.1
223 Escherichia coli O157_3AH7 str. Sakai GCA_000008865.1
223 Escherichia coli O83_3AH1 str. NRG 857C GCA_000183345.1
223 Escherichia coli UMN026 GCA_000026325.2
223 Escherichia coli str. K-12 substr. MG1655 GCA_000005845.2
223 Shigella flexneri 2a str. 301 GCA_000006925.2
223 Tumebacillus flagellatus GCA_000714935.1
214 Shigella dysenteriae Sd197 GCA_000012005.1
191 Erwinia iniecta GCA_001267535.1
3R
211 Rhodobacter sphaeroides 2.4.1 GCA_000012905.2
184 Rhodobacter sphaeroides ATCC 17025 GCA_000016405.1
166 Gemmobacter megaterium GCA_900156815.1
3S
185 Bacillus thuringiensis YBT-1518 GCA_000497525.2
171 Bacillus anthracis str. Ames GCA_000007845.1
171 Bacillus anthracis str. Sterne GCA_000008165.1
171 Bacillus cereus ATCC 14579 GCA_000007825.1
171 Bacillus mycoides GCA_000832605.1
171 Bacillus pseudomycoides DSM 12442 GCA_000161455.1
171 _5BBacillus thuringiensis_5D serovar konkukian str. 97-27 GCA_000008505.1
148 Bacillus horneckiae GCA_001636335.1
148 Bacillus solani GCA_001420595.1
3T
167 Streptococcus mutans UA159 GCA_000007465.2
124 Streptococcus ferus DSM 20646 GCA_000372425.1
110 Streptococcus massiliensis DSM 18628 GCA_000380065.1
3U
226 Bacillus anthracis str. Ames GCA_000007845.1
226 Bacillus anthracis str. Sterne GCA_000008165.1
226 Bacillus cereus ATCC 14579 GCA_000007825.1
226 Bacillus mycoides GCA_000832605.1
226 Bacillus pseudomycoides DSM 12442 GCA_000161455.1
226 Bacillus thuringiensis YBT-1518 GCA_000497525.2
226 _5BBacillus thuringiensis_5D serovar konkukian str. 97-27 GCA_000008505.1
213 Staphylococcus equorum GCA_001432245.1
202 Jeotgalicoccus saudimassiliensis GCA_000756715.2
202 Megasphaera cerevisiae DSM 20462 GCA_001045675.1
202 Staphylococcus aureus subsp. aureus NCTC 8325 GCA_000013425.1
202 Staphylococcus epidermidis ATCC 12228 GCA_000007645.1
202 Staphylococcus haemolyticus JCSC1435 GCA_000009865.1
202 Staphylococcus hominis subsp. hominis C80 GCA_000183685.1
202 Staphylococcus lugdunensis HKU09-01 GCA_000025085.1
202 Staphylococcus simiae CCM 7213 GCA_000235645.2
202 Staphylococcus warneri SG1 GCA_000332735.1
3V
247 Clostridium butyricum GCA_001456065.2
244 Clostridium beijerinckii GCA_000833105.2
244 Clostridium puniceum GCA_002006345.1
244 Clostridium saccharoperbutylacetonicum N1-4_28HMT_29 GCA_000340885.1
235 Clostridium saccharobutylicum DSM 13864 GCA_000473995.1
3W
122 Escherichia coli IAI39 GCA_000026345.1
122 Escherichia coli O104_3AH4 str. 2011C-3493 GCA_000299455.1
122 Escherichia coli O157_3AH7 str. Sakai GCA_000008865.1
122 Escherichia coli O83_3AH1 str. NRG 857C GCA_000183345.1
122 Escherichia coli UMN026 GCA_000026325.2
122 Escherichia coli str. K-12 substr. MG1655 GCA_000005845.2
122 Shigella flexneri 2a str. 301 GCA_000006925.2
122 Tumebacillus flagellatus GCA_000714935.1
120 Bacillus humi GCA_001439915.1
118 Erwinia iniecta GCA_001267535.1
118 Serratia marcescens subsp. marcescens Db11 GCA_000513215.1

3X
306 Escherichia coli IAI39 GCA_000026345.1
306 Escherichia coli O104_3AH4 str. 2011C-3493 GCA_000299455.1
306 Escherichia coli O157_3AH7 str. Sakai GCA_000008865.1
306 Escherichia coli O83_3AH1 str. NRG 857C GCA_000183345.1
306 Escherichia coli UMN026 GCA_000026325.2
306 Escherichia coli str. K-12 substr. MG1655 GCA_000005845.2
306 Shigella flexneri 2a str. 301 GCA_000006925.2
306 Tumebacillus flagellatus GCA_000714935.1
291 Shigella dysenteriae Sd197 GCA_000012005.1
262 Erwinia iniecta GCA_001267535.1
3Y
266 Deinococcus radiodurans R1 GCA_000008565.1
171 Deinococcus gobiensis I-0 GCA_000252445.1
157 Deinococcus deserti VCD115 GCA_000020685.1
3Z
249 Rhodobacter sphaeroides 2.4.1 GCA_000012905.2
240 Pseudorhodobacter psychrotolerans GCA_001294535.1
232 Gemmobacter megaterium GCA_900156815.1
40
225 Streptococcus mutans UA159 GCA_000007465.2
151 Streptococcus macacae NCTC 11558 GCA_000187995.3
143 Streptococcus equinus GCA_000964315.1
143 Streptococcus gallolyticus subsp. gallolyticus DSM 16831 GCA_002000985.1
143 Streptococcus orisratti DSM 15617 GCA_000380105.1
143 Streptococcus ratti FA-1 = DSM 20564 GCA_000286075.1
41
243 Bifidobacterium adolescentis ATCC 15703 GCA_000010425.1
230 Bifidobacterium callitrichos DSM 23973 GCA_000741175.1
229 Bifidobacterium angulatum DSM 20098 = JCM 7096 GCA_001025155.1
42
315 Streptococcus mutans UA159 GCA_000007465.2
191 Streptococcus ferus DSM 20646 GCA_000372425.1
186 Streptococcus massiliensis DSM 18628 GCA_000380065.1
43
263 Bacillus anthracis str. Ames GCA_000007845.1
263 Bacillus anthracis str. Sterne GCA_000008165.1
263 Bacillus cereus ATCC 14579 GCA_000007825.1
263 Bacillus mycoides GCA_000832605.1
263 Bacillus pseudomycoides DSM 12442 GCA_000161455.1
263 Bacillus thuringiensis YBT-1518 GCA_000497525.2
263 _5BBacillus thuringiensis_5D serovar konkukian str. 97-27 GCA_000008505.1
205 Bacillus cytotoxicus NVH 391-98 GCA_000017425.1
200 Bacillus vietnamensis NBRC 101237 GCA_001591825.1
44
178 Escherichia coli IAI39 GCA_000026345.1
178 Escherichia coli O104_3AH4 str. 2011C-3493 GCA_000299455.1
178 Escherichia coli O157_3AH7 str. Sakai GCA_000008865.1
178 Escherichia coli O83_3AH1 str. NRG 857C GCA_000183345.1
178 Escherichia coli UMN026 GCA_000026325.2
178 Escherichia coli str. K-12 substr. MG1655 GCA_000005845.2
178 Shigella dysenteriae Sd197 GCA_000012005.1
178 Shigella flexneri 2a str. 301 GCA_000006925.2
178 Tumebacillus flagellatus GCA_000714935.1
162 Erwinia iniecta GCA_001267535.1
158 Pectobacterium carotovorum subsp. carotovorum PC1 GCA_000023605.1
158 Plesiomonas shigelloides GCA_900087055.1
45
233 Enterococcus faecalis V583 GCA_000007785.1
233 Streptomyces cinnamoneus GCA_001885705.1
205 Enterococcus faecium DO GCA_000174395.2
199 Enterococcus canis NBRC 100695 GCA_001544375.1
199 Enterococcus casseliflavus EC20 GCA_000157355.2
199 Enterococcus dispar ATCC 51266 GCA_000406945.1
199 Enterococcus haemoperoxidus ATCC BAA-382 GCA_000407165.1
199 Enterococcus hirae ATCC 9790 GCA_000271405.2
199 Enterococcus massiliensis GCA_001050095.1
199 Enterococcus mundtii QU 25 GCA_000504125.1
199 Enterococcus phoeniculicola ATCC BAA-412 GCA_000407505.1
199 Enterococcus rivorum GCA_001742285.1
199 Enterococcus saccharolyticus subsp. saccharolyticus ATCC 43076 GCA_000407285.1
199 Enterococcus thailandicus GCA_001652875.1
46
210 Streptococcus mutans UA159 GCA_000007465.2
135 Streptococcus ferus DSM 20646 GCA_000372425.1
132 Streptococcus gordonii str. Challis substr. CH1 GCA_000017005.1

47
142 Clostridium beijerinckii GCA_000833105.2
142 Clostridium saccharobutylicum DSM 13864 GCA_000473995.1
142 Clostridium saccharoperbutylacetonicum N1-4_28HMT_29 GCA_000340885.1
132 Clostridium intestinale URNW GCA_000469625.2
117 Clostridium botulinum B str. Eklund 17B _28NRP_29 GCA_000020165.1
117 Clostridium butyricum GCA_001456065.2
117 Clostridium chromiireducens GCA_002029255.1
117 Clostridium puniceum GCA_002006345.1
117 Clostridium taeniosporum GCA_001735765.1
117 Desnuesiella massiliensis GCA_001403615.1
48
267 Streptococcus mutans UA159 GCA_000007465.2
174 Streptococcus equinus GCA_000964315.1
174 Streptococcus gallolyticus subsp. gallolyticus DSM 16831 GCA_002000985.1
173 Streptococcus gordonii str. Challis substr. CH1 GCA_000017005.1
49
240 Clostridium beijerinckii GCA_000833105.2
240 Clostridium saccharoperbutylacetonicum N1-4_28HMT_29 GCA_000340885.1
231 Clostridium saccharobutylicum DSM 13864 GCA_000473995.1
225 Clostridium puniceum GCA_002006345.1
4a
369 Enterococcus faecalis V583 GCA_000007785.1
369 Streptomyces cinnamoneus GCA_001885705.1
343 Enterococcus asini ATCC 700915 GCA_000407365.1
343 Enterococcus canis NBRC 100695 GCA_001544375.1
343 Enterococcus dispar ATCC 51266 GCA_000406945.1
343 Enterococcus faecium DO GCA_000174395.2
343 Enterococcus hirae ATCC 9790 GCA_000271405.2
343 Enterococcus mundtii QU 25 GCA_000504125.1
343 Enterococcus rivorum GCA_001742285.1
338 Enterococcus haemoperoxidus ATCC BAA-382 GCA_000407165.1
338 Enterococcus phoeniculicola ATCC BAA-412 GCA_000407505.1
338 Enterococcus thailandicus GCA_001652875.1
4b
292 Clostridium beijerinckii GCA_000833105.2
292 Clostridium saccharoperbutylacetonicum N1-4_28HMT_29 GCA_000340885.1
286 Clostridium puniceum GCA_002006345.1
284 Clostridium saccharobutylicum DSM 13864 GCA_000473995.1
4c
219 Escherichia coli IAI39 GCA_000026345.1
219 Escherichia coli O104_3AH4 str. 2011C-3493 GCA_000299455.1
219 Escherichia coli O157_3AH7 str. Sakai GCA_000008865.1
219 Escherichia coli O83_3AH1 str. NRG 857C GCA_000183345.1
219 Escherichia coli UMN026 GCA_000026325.2
219 Escherichia coli str. K-12 substr. MG1655 GCA_000005845.2
219 Shigella dysenteriae Sd197 GCA_000012005.1
219 Shigella flexneri 2a str. 301 GCA_000006925.2
219 Tumebacillus flagellatus GCA_000714935.1
196 Erwinia iniecta GCA_001267535.1
196 Erwinia toletana DAPP-PG 735 GCA_000336255.1
192 Cronobacter sakazakii GCA_000982825.1
192 Pantoea agglomerans GCA_001709315.1
192 Pantoea ananatis LMG 20103 GCA_000025405.2
192 Shimwellia blattae DSM 4481 = NBRC 105725 GCA_000262305.1
4d
203 Escherichia coli IAI39 GCA_000026345.1
203 Escherichia coli O104_3AH4 str. 2011C-3493 GCA_000299455.1
203 Escherichia coli O157_3AH7 str. Sakai GCA_000008865.1
203 Escherichia coli O83_3AH1 str. NRG 857C GCA_000183345.1
203 Escherichia coli UMN026 GCA_000026325.2
203 Escherichia coli str. K-12 substr. MG1655 GCA_000005845.2
203 Shigella dysenteriae Sd197 GCA_000012005.1
203 Shigella flexneri 2a str. 301 GCA_000006925.2
203 Tumebacillus flagellatus GCA_000714935.1
170 Erwinia iniecta GCA_001267535.1
167 Cronobacter sakazakii GCA_000982825.1
167 Enterobacter cloacae subsp. cloacae ATCC 13047 GCA_000025565.1
167 Enterobacter hormaechei subsp. steigerwaltii GCA_001729725.1
167 Enterobacter kobei GCA_900185885.1
167 Erwinia amylovora CFBP1430 GCA_000091565.1
167 Erwinia billingiae Eb661 GCA_000196615.1
167 Erwinia gerundensis GCA_001517405.1
167 Erwinia persicina NBRC 102418 GCA_001571305.1
167 Erwinia toletana DAPP-PG 735 GCA_000336255.1
167 Klebsiella oxytoca GCA_001022195.1

167 Kosakonia cowanii GCA_001975225.1
167 Kosakonia sacchari SP1 GCA_000300455.4
167 Pantoea agglomerans GCA_001709315.1
167 Pantoea alhagi GCA_002101395.1
167 Pantoea ananatis LMG 20103 GCA_000025405.2
167 Pantoea dispersa EGD-AAK13 GCA_000465555.2
167 Pantoea rwandensis GCA_000759475.1
167 Pantoea septica GCA_002095575.1
167 Pluralibacter gergoviae GCA_000757785.1
167 Pseudescherichia vulneris NBRC 102420 GCA_000759795.1
167 Salmonella enterica subsp. enterica serovar Typhi str. CT18 GCA_000195995.1
167 Shimwellia blattae DSM 4481 = NBRC 105725 GCA_000262305.1
167 Trabulsiella odontotermitis GCA_001297765.1
4e
179 Clostridium beijerinckii GCA_000833105.2
179 Clostridium saccharobutylicum DSM 13864 GCA_000473995.1
179 Clostridium saccharoperbutylacetonicum N1-4_28HMT_29 GCA_000340885.1
175 Clostridium butyricum GCA_001456065.2
166 Clostridium puniceum GCA_002006345.1
4f
203 Deinococcus radiodurans R1 GCA_000008565.1
139 Deinococcus gobiensis I-0 GCA_000252445.1
133 Deinococcus deserti VCD115 GCA_000020685.1
133 Deinococcus hopiensis KR-140 GCA_900176165.1
133 Deinococcus marmoris DSM 12784 GCA_000701405.1
133 Deinococcus swuensis GCA_000800395.1
4g
215 Bacillus anthracis str. Ames GCA_000007845.1
215 Bacillus anthracis str. Sterne GCA_000008165.1
215 Bacillus cereus ATCC 14579 GCA_000007825.1
215 Bacillus mycoides GCA_000832605.1
215 Bacillus pseudomycoides DSM 12442 GCA_000161455.1
215 Bacillus thuringiensis YBT-1518 GCA_000497525.2
215 _5BBacillus thuringiensis_5D serovar konkukian str. 97-27 GCA_000008505.1
201 Bacillus solani GCA_001420595.1
200 Bacillus horneckiae GCA_001636335.1
4h
175 Rhodobacter sphaeroides 2.4.1 GCA_000012905.2
152 Rhodobacter sphaeroides ATCC 17025 GCA_000016405.1
133 Gemmobacter megaterium GCA_900156815.1
133 Pseudorhodobacter ferrugineus DSM 5888 GCA_000420745.1
133 Pseudorhodobacter wandonensis GCA_001202035.1
4i
142 Staphylococcus succinus GCA_001902315.1
130 Megasphaera cerevisiae DSM 20462 GCA_001045675.1
130 Staphylococcus aureus subsp. aureus NCTC 8325 GCA_000013425.1
130 Staphylococcus capitis subsp. capitis GCA_001028645.1
130 Staphylococcus cohnii subsp. cohnii GCA_000972575.1
130 Staphylococcus condimenti GCA_001618885.1
130 Staphylococcus epidermidis ATCC 12228 GCA_000007645.1
130 Staphylococcus haemolyticus JCSC1435 GCA_000009865.1
130 Staphylococcus hominis subsp. hominis C80 GCA_000183685.1
130 Staphylococcus sciuri GCA_002209165.1
130 Staphylococcus simiae CCM 7213 GCA_000235645.2
130 Staphylococcus simulans GCA_001559115.1
130 Staphylococcus warneri SG1 GCA_000332735.1
125 Staphylococcus hyicus GCA_000816085.1
125 Staphylococcus lutrae GCA_002101335.1
125 Staphylococcus saprophyticus GCA_001074355.1
4j
257 Escherichia coli IAI39 GCA_000026345.1
257 Escherichia coli O104_3AH4 str. 2011C-3493 GCA_000299455.1
257 Escherichia coli O157_3AH7 str. Sakai GCA_000008865.1
257 Escherichia coli O83_3AH1 str. NRG 857C GCA_000183345.1
257 Escherichia coli UMN026 GCA_000026325.2
257 Escherichia coli str. K-12 substr. MG1655 GCA_000005845.2
257 Shigella dysenteriae Sd197 GCA_000012005.1
257 Shigella flexneri 2a str. 301 GCA_000006925.2
257 Tumebacillus flagellatus GCA_000714935.1
231 Erwinia iniecta GCA_001267535.1
231 Erwinia toletana DAPP-PG 735 GCA_000336255.1
231 Pantoea agglomerans GCA_001709315.1
231 Pantoea ananatis LMG 20103 GCA_000025405.2
231 Shimwellia blattae DSM 4481 = NBRC 105725 GCA_000262305.1
216 Cronobacter sakazakii GCA_000982825.1
216 Enterobacter hormaechei subsp. steigerwaltii GCA_001729725.1

216 Erwinia billingiae Eb661 GCA_000196615.1
216 Erwinia gerundensis GCA_001517405.1
216 Klebsiella oxytoca GCA_001022195.1
216 Kosakonia cowanii GCA_001975225.1
216 Kosakonia sacchari SP1 GCA_000300455.4
216 Pantoea alhagi GCA_002101395.1
216 Pantoea dispersa EGD-AAK13 GCA_000465555.2
216 Pantoea rwandensis GCA_000759475.1
216 Pantoea septica GCA_002095575.1
216 Plautia stali symbiont GCA_000180175.2
216 Pluralibacter gergoviae GCA_000757785.1
216 Pseudescherichia vulneris NBRC 102420 GCA_000759795.1
4k
225 Clostridium beijerinckii GCA_000833105.2
225 Clostridium puniceum GCA_002006345.1
225 Clostridium saccharoperbutylacetonicum N1-4_28HMT_29 GCA_000340885.1
216 Clostridium butyricum GCA_001456065.2
216 Clostridium chromiireducens GCA_002029255.1
216 Clostridium saccharobutylicum DSM 13864 GCA_000473995.1
214 Clostridium taeniosporum GCA_001735765.1
4l
167 Clostridium taeniosporum GCA_001735765.1
163 Clostridium beijerinckii GCA_000833105.2
163 Clostridium puniceum GCA_002006345.1
163 Clostridium saccharoperbutylacetonicum N1-4_28HMT_29 GCA_000340885.1
158 Clostridium botulinum B str. Eklund 17B _28NRP_29 GCA_000020165.1
4m
170 Escherichia coli IAI39 GCA_000026345.1
170 Escherichia coli O104_3AH4 str. 2011C-3493 GCA_000299455.1
170 Escherichia coli O157_3AH7 str. Sakai GCA_000008865.1
170 Escherichia coli O83_3AH1 str. NRG 857C GCA_000183345.1
170 Escherichia coli UMN026 GCA_000026325.2
170 Escherichia coli str. K-12 substr. MG1655 GCA_000005845.2
170 Shigella dysenteriae Sd197 GCA_000012005.1
170 Shigella flexneri 2a str. 301 GCA_000006925.2
170 Tumebacillus flagellatus GCA_000714935.1
130 Erwinia iniecta GCA_001267535.1
111 Cedecea neteri GCA_000757825.1
111 Rosenbergiella nectarea GCA_900111105.1
4n
131 Staphylococcus aureus subsp. aureus NCTC 8325 GCA_000013425.1
131 Staphylococcus capitis subsp. capitis GCA_001028645.1
131 Staphylococcus epidermidis ATCC 12228 GCA_000007645.1
131 Staphylococcus haemolyticus JCSC1435 GCA_000009865.1
131 Staphylococcus hominis subsp. hominis C80 GCA_000183685.1
131 Staphylococcus lugdunensis HKU09-01 GCA_000025085.1
131 Staphylococcus simiae CCM 7213 GCA_000235645.2
128 Staphylococcus condimenti GCA_001618885.1
128 Staphylococcus simulans GCA_001559115.1
118 Megasphaera cerevisiae DSM 20462 GCA_001045675.1
118 Staphylococcus warneri SG1 GCA_000332735.1
4o
295 Staphylococcus capitis subsp. capitis GCA_001028645.1
295 Staphylococcus epidermidis ATCC 12228 GCA_000007645.1
292 Staphylococcus haemolyticus JCSC1435 GCA_000009865.1
282 Staphylococcus cohnii subsp. cohnii GCA_000972575.1
4p
263 Deinococcus radiodurans R1 GCA_000008565.1
193 Deinococcus gobiensis I-0 GCA_000252445.1
179 Deinococcus puniceus GCA_001644565.1
4q
179 Clostridium beijerinckii GCA_000833105.2
179 Clostridium puniceum GCA_002006345.1
179 Clostridium saccharoperbutylacetonicum N1-4_28HMT_29 GCA_000340885.1
171 Clostridium butyricum GCA_001456065.2
163 Clostridium saccharobutylicum DSM 13864 GCA_000473995.1
4r
121 Rhodobacter sphaeroides 2.4.1 GCA_000012905.2
92 Rhodobacter sphaeroides ATCC 17025 GCA_000016405.1
80 Gemmobacter megaterium GCA_900156815.1
80 Pseudorhodobacter ferrugineus DSM 5888 GCA_000420745.1
80 Pseudorhodobacter wandonensis GCA_001202035.1
4s
196 Clostridium beijerinckii GCA_000833105.2
196 Clostridium puniceum GCA_002006345.1
196 Clostridium saccharoperbutylacetonicum N1-4_28HMT_29 GCA_000340885.1

188 Clostridium saccharobutylicum DSM 13864 GCA_000473995.1
178 Clostridium butyricum GCA_001456065.2
4t
287 Clostridium saccharobutylicum DSM 13864 GCA_000473995.1
272 Clostridium beijerinckii GCA_000833105.2
272 Clostridium saccharoperbutylacetonicum N1-4_28HMT_29 GCA_000340885.1
266 Clostridium chromiireducens GCA_002029255.1
266 Clostridium puniceum GCA_002006345.1
4u
208 Deinococcus radiodurans R1 GCA_000008565.1
141 Deinococcus marmoris DSM 12784 GCA_000701405.1
141 Deinococcus swuensis GCA_000800395.1
134 Deinococcus gobiensis I-0 GCA_000252445.1
4v
163 Staphylococcus aureus subsp. aureus NCTC 8325 GCA_000013425.1
163 Staphylococcus capitis subsp. capitis GCA_001028645.1
163 Staphylococcus cohnii subsp. cohnii GCA_000972575.1
163 Staphylococcus condimenti GCA_001618885.1
163 Staphylococcus epidermidis ATCC 12228 GCA_000007645.1
163 Staphylococcus haemolyticus JCSC1435 GCA_000009865.1
163 Staphylococcus hominis subsp. hominis C80 GCA_000183685.1
163 Staphylococcus simiae CCM 7213 GCA_000235645.2
163 Staphylococcus simulans GCA_001559115.1
158 Staphylococcus succinus GCA_001902315.1
153 Staphylococcus microti GCA_000934465.1
153 Staphylococcus saprophyticus GCA_001074355.1
4w
245 Clostridium beijerinckii GCA_000833105.2
245 Clostridium puniceum GCA_002006345.1
245 Clostridium saccharobutylicum DSM 13864 GCA_000473995.1
245 Clostridium saccharoperbutylacetonicum N1-4_28HMT_29 GCA_000340885.1
227 Clostridium chromiireducens GCA_002029255.1
225 Clostridium butyricum GCA_001456065.2
4x
206 Clostridium beijerinckii GCA_000833105.2
206 Clostridium saccharoperbutylacetonicum N1-4_28HMT_29 GCA_000340885.1
202 Clostridium puniceum GCA_002006345.1
197 Clostridium saccharobutylicum DSM 13864 GCA_000473995.1
4y
130 Staphylococcus aureus subsp. aureus NCTC 8325 GCA_000013425.1
130 Staphylococcus capitis subsp. capitis GCA_001028645.1
130 Staphylococcus epidermidis ATCC 12228 GCA_000007645.1
130 Staphylococcus haemolyticus JCSC1435 GCA_000009865.1
130 Staphylococcus hominis subsp. hominis C80 GCA_000183685.1
130 Staphylococcus lugdunensis HKU09-01 GCA_000025085.1
130 Staphylococcus simiae CCM 7213 GCA_000235645.2
127 Megasphaera cerevisiae DSM 20462 GCA_001045675.1
127 Staphylococcus arlettae CVD059 GCA_000295715.1
127 Staphylococcus cohnii subsp. cohnii GCA_000972575.1
127 Staphylococcus gallinarum GCA_000875895.1
127 Staphylococcus lutrae GCA_002101335.1
127 Staphylococcus saprophyticus GCA_001074355.1
127 Staphylococcus saprophyticus subsp. saprophyticus ATCC 15305 GCA_000010125.1
127 Staphylococcus succinus GCA_001902315.1
127 Staphylococcus warneri SG1 GCA_000332735.1
127 Staphylococcus xylosus GCA_000706685.1
119 Salinicoccus alkaliphilus DSM 16010 GCA_900142805.1
119 Salinicoccus carnicancri Crm GCA_000330705.1
119 Salinicoccus halodurans GCA_001005905.1
119 Salinicoccus qingdaonensis GCA_900101075.1
119 Staphylococcus lentus F1142 GCA_000286395.1
119 Staphylococcus sciuri GCA_002209165.1
119 Staphylococcus vitulinus F1028 GCA_000286335.1
4z
173 Lactobacillus gasseri ATCC 33323 = JCM 1131 GCA_000014425.1
173 Lactobacillus hominis DSM 23910 = CRBIP 24.179 GCA_000296835.1
124 Lactobacillus psittaci DSM 15354 GCA_000425905.1
123 Lactobacillus iners DSM 13335 GCA_000160875.1
4A
253 Rhodobacter sphaeroides 2.4.1 GCA_000012905.2
247 Rhodobacter sphaeroides ATCC 17025 GCA_000016405.1
189 Gemmobacter aquatilis GCA_900110025.1
189 Rhodobacter capsulatus SB 1003 GCA_000021865.1
4B
284 Escherichia coli IAI39 GCA_000026345.1
284 Escherichia coli O104_3AH4 str. 2011C-3493 GCA_000299455.1

284 Escherichia coli O157_3AH7 str. Sakai GCA_000008865.1
284 Escherichia coli O83_3AH1 str. NRG 857C GCA_000183345.1
284 Escherichia coli UMN026 GCA_000026325.2
284 Escherichia coli str. K-12 substr. MG1655 GCA_000005845.2
284 Shigella flexneri 2a str. 301 GCA_000006925.2
284 Tumebacillus flagellatus GCA_000714935.1
275 Shigella dysenteriae Sd197 GCA_000012005.1
263 Erwinia iniecta GCA_001267535.1
4C
237 Clostridium saccharobutylicum DSM 13864 GCA_000473995.1
235 Clostridium beijerinckii GCA_000833105.2
235 Clostridium saccharoperbutylacetonicum N1-4_28HMT_29 GCA_000340885.1
219 Clostridium butyricum GCA_001456065.2
219 Clostridium puniceum GCA_002006345.1
4D
299 Clostridium beijerinckii GCA_000833105.2
299 Clostridium puniceum GCA_002006345.1
299 Clostridium saccharoperbutylacetonicum N1-4_28HMT_29 GCA_000340885.1
290 Clostridium saccharobutylicum DSM 13864 GCA_000473995.1
278 Clostridium taeniosporum GCA_001735765.1
4E
230 Clostridium beijerinckii GCA_000833105.2
230 Clostridium puniceum GCA_002006345.1
230 Clostridium saccharoperbutylacetonicum N1-4_28HMT_29 GCA_000340885.1
222 Clostridium saccharobutylicum DSM 13864 GCA_000473995.1
218 Clostridium butyricum GCA_001456065.2
4F
188 Deinococcus radiodurans R1 GCA_000008565.1
122 Deinococcus gobiensis I-0 GCA_000252445.1
121 Deinococcus deserti VCD115 GCA_000020685.1
4G
205 Clostridium beijerinckii GCA_000833105.2
205 Clostridium saccharoperbutylacetonicum N1-4_28HMT_29 GCA_000340885.1
198 Clostridium puniceum GCA_002006345.1
191 Clostridium saccharobutylicum DSM 13864 GCA_000473995.1
4H
242 Bacillus anthracis str. Ames GCA_000007845.1
242 Bacillus anthracis str. Sterne GCA_000008165.1
242 Bacillus cereus ATCC 14579 GCA_000007825.1
242 Bacillus mycoides GCA_000832605.1
242 Bacillus pseudomycoides DSM 12442 GCA_000161455.1
242 Bacillus thuringiensis YBT-1518 GCA_000497525.2
242 _5BBacillus thuringiensis_5D serovar konkukian str. 97-27 GCA_000008505.1
200 Bacillus manliponensis GCA_000712595.1
199 Bacillus tuaregi GCA_900104575.1
4I
241 Lactobacillus gasseri ATCC 33323 = JCM 1131 GCA_000014425.1
226 Lactobacillus hominis DSM 23910 = CRBIP 24.179 GCA_000296835.1
172 Lactobacillus psittaci DSM 15354 GCA_000425905.1
4J
309 Streptococcus mutans UA159 GCA_000007465.2
159 Streptococcus macacae NCTC 11558 GCA_000187995.3
154 Streptococcus massiliensis DSM 18628 GCA_000380065.1
4K
229 Enterococcus faecalis V583 GCA_000007785.1
229 Streptomyces cinnamoneus GCA_001885705.1
194 Enterococcus canis NBRC 100695 GCA_001544375.1
194 Enterococcus casseliflavus EC20 GCA_000157355.2
194 Enterococcus dispar ATCC 51266 GCA_000406945.1
194 Enterococcus faecium DO GCA_000174395.2
194 Enterococcus haemoperoxidus ATCC BAA-382 GCA_000407165.1
194 Enterococcus hirae ATCC 9790 GCA_000271405.2
194 Enterococcus mundtii QU 25 GCA_000504125.1
194 Enterococcus phoeniculicola ATCC BAA-412 GCA_000407505.1
194 Enterococcus rivorum GCA_001742285.1
194 Enterococcus saccharolyticus subsp. saccharolyticus ATCC 43076 GCA_000407285.1
194 Enterococcus thailandicus GCA_001652875.1
192 Enterococcus asini ATCC 700915 GCA_000407365.1
4L
271 Escherichia coli IAI39 GCA_000026345.1
271 Escherichia coli O104_3AH4 str. 2011C-3493 GCA_000299455.1
271 Escherichia coli O157_3AH7 str. Sakai GCA_000008865.1
271 Escherichia coli O83_3AH1 str. NRG 857C GCA_000183345.1
271 Escherichia coli UMN026 GCA_000026325.2
271 Escherichia coli str. K-12 substr. MG1655 GCA_000005845.2
271 Tumebacillus flagellatus GCA_000714935.1

267 Shigella dysenteriae Sd197 GCA_000012005.1
263 Shigella flexneri 2a str. 301 GCA_000006925.2
4M
220 Bacillus thuringiensis YBT-1518 GCA_000497525.2
191 Bacillus anthracis str. Ames GCA_000007845.1
191 Bacillus anthracis str. Sterne GCA_000008165.1
191 Bacillus cereus ATCC 14579 GCA_000007825.1
191 Bacillus mycoides GCA_000832605.1
191 Bacillus pseudomycoides DSM 12442 GCA_000161455.1
191 _5BBacillus thuringiensis_5D serovar konkukian str. 97-27 GCA_000008505.1
178 Bacillus acidicola GCA_001636425.1
178 Bacillus shackletonii GCA_001420715.1
4N
248 Deinococcus radiodurans R1 GCA_000008565.1
150 Deinococcus gobiensis I-0 GCA_000252445.1
145 Deinococcus marmoris DSM 12784 GCA_000701405.1
145 Deinococcus swuensis GCA_000800395.1
4O
264 Bacillus anthracis str. Ames GCA_000007845.1
264 Bacillus anthracis str. Sterne GCA_000008165.1
264 Bacillus mycoides GCA_000832605.1
264 Bacillus pseudomycoides DSM 12442 GCA_000161455.1
264 Bacillus thuringiensis YBT-1518 GCA_000497525.2
264 _5BBacillus thuringiensis_5D serovar konkukian str. 97-27 GCA_000008505.1
263 Bacillus cereus ATCC 14579 GCA_000007825.1
230 Bacillus manliponensis GCA_000712595.1
4P
254 Escherichia coli IAI39 GCA_000026345.1
254 Escherichia coli O104_3AH4 str. 2011C-3493 GCA_000299455.1
254 Escherichia coli O157_3AH7 str. Sakai GCA_000008865.1
254 Escherichia coli O83_3AH1 str. NRG 857C GCA_000183345.1
254 Escherichia coli UMN026 GCA_000026325.2
254 Escherichia coli str. K-12 substr. MG1655 GCA_000005845.2
254 Shigella dysenteriae Sd197 GCA_000012005.1
254 Shigella flexneri 2a str. 301 GCA_000006925.2
254 Tumebacillus flagellatus GCA_000714935.1
222 Enterobacter hormaechei subsp. steigerwaltii GCA_001729725.1
222 Erwinia iniecta GCA_001267535.1
222 Erwinia toletana DAPP-PG 735 GCA_000336255.1
222 Klebsiella oxytoca GCA_001022195.1
222 Kosakonia cowanii GCA_001975225.1
222 Pluralibacter gergoviae GCA_000757785.1
222 Pseudescherichia vulneris NBRC 102420 GCA_000759795.1
218 Cronobacter sakazakii GCA_000982825.1
218 Enterobacter cloacae subsp. cloacae ATCC 13047 GCA_000025565.1
218 Enterobacter kobei GCA_900185885.1
218 Kosakonia sacchari SP1 GCA_000300455.4
218 Salmonella enterica subsp. enterica serovar Typhi str. CT18 GCA_000195995.1
4Q
309 Streptococcus mutans UA159 GCA_000007465.2
203 Streptococcus sobrinus DSM 20742 = ATCC 33478 GCA_000686605.1
190 Streptococcus gordonii str. Challis substr. CH1 GCA_000017005.1
4R
237 Clostridium beijerinckii GCA_000833105.2
237 Clostridium puniceum GCA_002006345.1
237 Clostridium saccharoperbutylacetonicum N1-4_28HMT_29 GCA_000340885.1
228 Clostridium chromiireducens GCA_002029255.1
228 Clostridium saccharobutylicum DSM 13864 GCA_000473995.1
222 Clostridium butyricum GCA_001456065.2
4S
226 Bacillus anthracis str. Ames GCA_000007845.1
226 Bacillus anthracis str. Sterne GCA_000008165.1
226 Bacillus cereus ATCC 14579 GCA_000007825.1
226 Bacillus mycoides GCA_000832605.1
226 Bacillus pseudomycoides DSM 12442 GCA_000161455.1
226 Bacillus thuringiensis YBT-1518 GCA_000497525.2
226 _5BBacillus thuringiensis_5D serovar konkukian str. 97-27 GCA_000008505.1
192 Bacillus flexus GCA_002024265.1
192 Bacillus megaterium NBRC 15308 = ATCC 14581 GCA_000832985.1
186 Bacillus marisflavi GCA_001274775.1
4T
183 Enterococcus faecalis V583 GCA_000007785.1
183 Streptomyces cinnamoneus GCA_001885705.1
174 Lactobacillus ginsenosidimutans GCA_001050475.1
174 Lactobacillus versmoldensis DSM 14857 = KCTC 3814 GCA_001434295.1
173 Catellicoccus marimammalium M35_2F04_2F3 GCA_000313915.1

4U
287 Lactobacillus gasseri ATCC 33323 = JCM 1131 GCA_000014425.1
274 Lactobacillus hominis DSM 23910 = CRBIP 24.179 GCA_000296835.1
213 Lactobacillus crispatus ST1 GCA_000091765.1
213 Lactobacillus helveticus GCA_001308285.1
4V
157 Clostridium beijerinckii GCA_000833105.2
157 Clostridium puniceum GCA_002006345.1
157 Clostridium saccharoperbutylacetonicum N1-4_28HMT_29 GCA_000340885.1
146 Clostridium saccharobutylicum DSM 13864 GCA_000473995.1
146 Clostridium taeniosporum GCA_001735765.1
137 Clostridium intestinale URNW GCA_000469625.2
137 Desnuesiella massiliensis GCA_001403615.1
4W
269 Clostridium saccharobutylicum DSM 13864 GCA_000473995.1
267 Clostridium beijerinckii GCA_000833105.2
267 Clostridium puniceum GCA_002006345.1
267 Clostridium saccharoperbutylacetonicum N1-4_28HMT_29 GCA_000340885.1
254 Clostridium chromiireducens GCA_002029255.1
4X
272 Bacillus anthracis str. Ames GCA_000007845.1
272 Bacillus anthracis str. Sterne GCA_000008165.1
272 Bacillus cereus ATCC 14579 GCA_000007825.1
272 Bacillus mycoides GCA_000832605.1
272 Bacillus pseudomycoides DSM 12442 GCA_000161455.1
272 Bacillus thuringiensis YBT-1518 GCA_000497525.2
272 _5BBacillus thuringiensis_5D serovar konkukian str. 97-27 GCA_000008505.1
213 Bacillus manliponensis GCA_000712595.1
211 Staphylococcus condimenti GCA_001618885.1
4Y
214 Streptococcus mutans UA159 GCA_000007465.2
158 Streptococcus ferus DSM 20646 GCA_000372425.1
150 Streptococcus macacae NCTC 11558 GCA_000187995.3
4Z
132 Clostridium beijerinckii GCA_000833105.2
132 Clostridium butyricum GCA_001456065.2
132 Clostridium puniceum GCA_002006345.1
132 Clostridium saccharoperbutylacetonicum N1-4_28HMT_29 GCA_000340885.1
123 Clostridium chromiireducens GCA_002029255.1
123 Clostridium saccharobutylicum DSM 13864 GCA_000473995.1
118 Clostridium uliginosum GCA_900112485.1
50
243 Clostridium saccharobutylicum DSM 13864 GCA_000473995.1
241 Clostridium beijerinckii GCA_000833105.2
241 Clostridium puniceum GCA_002006345.1
241 Clostridium saccharoperbutylacetonicum N1-4_28HMT_29 GCA_000340885.1
233 Clostridium butyricum GCA_001456065.2
51
161 Escherichia coli IAI39 GCA_000026345.1
161 Escherichia coli O104_3AH4 str. 2011C-3493 GCA_000299455.1
161 Escherichia coli O157_3AH7 str. Sakai GCA_000008865.1
161 Escherichia coli O83_3AH1 str. NRG 857C GCA_000183345.1
161 Escherichia coli UMN026 GCA_000026325.2
161 Escherichia coli str. K-12 substr. MG1655 GCA_000005845.2
161 Shigella dysenteriae Sd197 GCA_000012005.1
161 Shigella flexneri 2a str. 301 GCA_000006925.2
161 Tumebacillus flagellatus GCA_000714935.1
144 Enterobacter cloacae subsp. cloacae ATCC 13047 GCA_000025565.1
142 Cronobacter sakazakii GCA_000982825.1
142 Enterobacter hormaechei subsp. steigerwaltii GCA_001729725.1
142 Enterobacter kobei GCA_900185885.1
142 Erwinia iniecta GCA_001267535.1
142 Erwinia toletana DAPP-PG 735 GCA_000336255.1
142 Klebsiella oxytoca GCA_001022195.1
142 Kosakonia cowanii GCA_001975225.1
142 Kosakonia sacchari SP1 GCA_000300455.4
142 Pluralibacter gergoviae GCA_000757785.1
142 Pseudescherichia vulneris NBRC 102420 GCA_000759795.1
142 Salmonella enterica subsp. enterica serovar Typhi str. CT18 GCA_000195995.1
52
103 Buttiauxella ferragutiae ATCC 51602 GCA_001654915.1
103 Cedecea neteri GCA_000757825.1
103 Citrobacter freundii CFNIH1 GCA_000648515.1
103 Cronobacter sakazakii GCA_000982825.1
103 Edwardsiella anguillarum ET080813 GCA_000264765.2
103 Enterobacter cloacae subsp. cloacae ATCC 13047 GCA_000025565.1

103 Enterobacter hormaechei subsp. steigerwaltii GCA_001729725.1
103 Enterobacter kobei GCA_900185885.1
103 Erwinia amylovora CFBP1430 GCA_000091565.1
103 Erwinia billingiae Eb661 GCA_000196615.1
103 Erwinia gerundensis GCA_001517405.1
103 Erwinia iniecta GCA_001267535.1
103 Erwinia persicina NBRC 102418 GCA_001571305.1
103 Erwinia toletana DAPP-PG 735 GCA_000336255.1
103 Escherichia coli IAI39 GCA_000026345.1
103 Escherichia coli O104_3AH4 str. 2011C-3493 GCA_000299455.1
103 Escherichia coli O157_3AH7 str. Sakai GCA_000008865.1
103 Escherichia coli O83_3AH1 str. NRG 857C GCA_000183345.1
103 Escherichia coli UMN026 GCA_000026325.2
103 Escherichia coli str. K-12 substr. MG1655 GCA_000005845.2
103 Klebsiella aerogenes KCTC 2190 GCA_000215745.1
103 Klebsiella oxytoca GCA_001022195.1
103 Kluyvera ascorbata ATCC 33433 GCA_000735365.1
103 Kosakonia cowanii GCA_001975225.1
103 Kosakonia sacchari SP1 GCA_000300455.4
103 Mangrovibacter phragmitis GCA_001655675.1
103 Obesumbacterium proteus GCA_001586165.1
103 Pantoea agglomerans GCA_001709315.1
103 Pantoea alhagi GCA_002101395.1
103 Pantoea ananatis LMG 20103 GCA_000025405.2
103 Pantoea dispersa EGD-AAK13 GCA_000465555.2
103 Pantoea rwandensis GCA_000759475.1
103 Pantoea septica GCA_002095575.1
103 Pluralibacter gergoviae GCA_000757785.1
103 Pseudescherichia vulneris NBRC 102420 GCA_000759795.1
103 Rosenbergiella nectarea GCA_900111105.1
103 Salmonella enterica subsp. enterica serovar Typhi str. CT18 GCA_000195995.1
103 Shigella dysenteriae Sd197 GCA_000012005.1
103 Shigella flexneri 2a str. 301 GCA_000006925.2
103 Shimwellia blattae DSM 4481 = NBRC 105725 GCA_000262305.1
103 Tatumella citrea GCA_002163585.1
103 Tatumella saanichensis GCA_000439375.1
103 Trabulsiella odontotermitis GCA_001297765.1
103 Tumebacillus flagellatus GCA_000714935.1
103 Yokenella regensburgei ATCC 49455 GCA_000735455.1
103 _5BEnterobacter_5D lignolyticus SCF1 GCA_000164865.1
96 Klebsiella pneumoniae subsp. pneumoniae HS11286 GCA_000240185.2
96 Plesiomonas shigelloides GCA_900087055.1
96 Serratia liquefaciens ATCC 27592 GCA_000422085.1
96 Serratia marcescens subsp. marcescens Db11 GCA_000513215.1
96 Serratia rubidaea GCA_001572725.1
96 Serratia symbiotica GCA_000821185.1
96 Yersinia enterocolitica subsp. enterocolitica 8081 GCA_000009345.1
96 Yersinia pestis CO92 GCA_000009065.1
96 Yersinia ruckeri GCA_000964565.1
91 Salmonella enterica subsp. enterica serovar Typhimurium str. LT2 GCA_000006945.2
53
203 Deinococcus radiodurans R1 GCA_000008565.1
137 Deinococcus gobiensis I-0 GCA_000252445.1
136 Deinococcus frigens DSM 12807 GCA_000701425.1
54
149 Escherichia coli IAI39 GCA_000026345.1
149 Escherichia coli O104_3AH4 str. 2011C-3493 GCA_000299455.1
149 Escherichia coli O157_3AH7 str. Sakai GCA_000008865.1
149 Escherichia coli UMN026 GCA_000026325.2
149 Escherichia coli str. K-12 substr. MG1655 GCA_000005845.2
149 Shigella flexneri 2a str. 301 GCA_000006925.2
149 Tumebacillus flagellatus GCA_000714935.1
134 Escherichia coli O83_3AH1 str. NRG 857C GCA_000183345.1
134 Shigella dysenteriae Sd197 GCA_000012005.1
128 Erwinia iniecta GCA_001267535.1
55
255 Rhodobacter sphaeroides 2.4.1 GCA_000012905.2
234 Rhodobacter sphaeroides ATCC 17025 GCA_000016405.1
226 Pseudorhodobacter ferrugineus DSM 5888 GCA_000420745.1
226 Pseudorhodobacter wandonensis GCA_001202035.1
56
126 Bacillus aquimaris TF-12 GCA_001648555.1
126 Bacillus coahuilensis m4-4 GCA_000171615.1
126 Bacillus eiseniae GCA_001636325.1
126 Bacillus infantis NRRL B-14911 GCA_000473245.1
126 Bacillus marisflavi GCA_001274775.1

126 Bacillus vietnamensis NBRC 101237 GCA_001591825.1
126 Planococcus antarcticus DSM 14505 GCA_001687565.2
126 Planococcus kocurii GCA_001465835.2
126 Planococcus maritimus GCA_001999905.1
126 Planococcus rifietoensis GCA_001465795.2
122 Bacillus anthracis str. Ames GCA_000007845.1
122 Bacillus anthracis str. Sterne GCA_000008165.1
122 Bacillus cereus ATCC 14579 GCA_000007825.1
122 Bacillus mycoides GCA_000832605.1
122 Bacillus pseudomycoides DSM 12442 GCA_000161455.1
122 Bacillus thuringiensis YBT-1518 GCA_000497525.2
122 Staphylococcus equorum GCA_001432245.1
122 _5BBacillus thuringiensis_5D serovar konkukian str. 97-27 GCA_000008505.1
117 Anaerobacillus macyae GCA_001039475.1
117 Bacillus acidicola GCA_001636425.1
117 Bacillus akibai JCM 9157 GCA_000513135.1
117 Bacillus alkalinitrilicus GCA_002019605.1
117 Bacillus badius GCA_001630115.1
117 Bacillus circulans NBRC 13626 GCA_001591585.1
117 Bacillus flexus GCA_002024265.1
117 Bacillus gottheilii GCA_001636345.1
117 Bacillus koreensis GCA_001274935.1
117 Bacillus krulwichiae GCA_002109385.1
117 Bacillus megaterium NBRC 15308 = ATCC 14581 GCA_000832985.1
117 Bacillus pseudofirmus OF4 GCA_000005825.2
117 Bacillus shackletonii GCA_001420715.1
117 Bacillus sporothermodurans GCA_001587375.1
117 Bacillus tuaregi GCA_900104575.1
117 Domibacillus antri GCA_001936625.1
117 Quasibacillus thermotolerans GCA_000812025.2
117 Virgibacillus halodenitrificans GCA_001878675.1
57
287 Lactobacillus gasseri ATCC 33323 = JCM 1131 GCA_000014425.1
287 Lactobacillus hominis DSM 23910 = CRBIP 24.179 GCA_000296835.1
218 Lactobacillus iners DSM 13335 GCA_000160875.1
218 Lactobacillus psittaci DSM 15354 GCA_000425905.1
203 Lactobacillus crispatus ST1 GCA_000091765.1
58
242 Enterococcus faecalis V583 GCA_000007785.1
242 Streptomyces cinnamoneus GCA_001885705.1
211 Enterococcus dispar ATCC 51266 GCA_000406945.1
208 Enterococcus canis NBRC 100695 GCA_001544375.1
208 Enterococcus faecium DO GCA_000174395.2
208 Enterococcus haemoperoxidus ATCC BAA-382 GCA_000407165.1
208 Enterococcus hirae ATCC 9790 GCA_000271405.2
208 Enterococcus massiliensis GCA_001050095.1
208 Enterococcus mundtii QU 25 GCA_000504125.1
208 Enterococcus phoeniculicola ATCC BAA-412 GCA_000407505.1
208 Enterococcus rivorum GCA_001742285.1
208 Enterococcus thailandicus GCA_001652875.1
59
309 Streptococcus mutans UA159 GCA_000007465.2
210 Streptococcus equinus GCA_000964315.1
210 Streptococcus gallolyticus subsp. gallolyticus DSM 16831 GCA_002000985.1
206 Streptococcus cristatus AS 1.3089 GCA_000385925.1
5a
266 Deinococcus radiodurans R1 GCA_000008565.1
181 Deinococcus gobiensis I-0 GCA_000252445.1
162 Deinococcus puniceus GCA_001644565.1
5b
157 Clostridium beijerinckii GCA_000833105.2
157 Clostridium puniceum GCA_002006345.1
157 Clostridium saccharobutylicum DSM 13864 GCA_000473995.1
157 Clostridium saccharoperbutylacetonicum N1-4_28HMT_29 GCA_000340885.1
154 Clostridium neonatale GCA_001458595.1
153 Clostridium butyricum GCA_001456065.2
153 Clostridium chromiireducens GCA_002029255.1
5c
318 Bacillus thuringiensis YBT-1518 GCA_000497525.2
317 Bacillus anthracis str. Ames GCA_000007845.1
317 Bacillus anthracis str. Sterne GCA_000008165.1
317 Bacillus cereus ATCC 14579 GCA_000007825.1
317 Bacillus mycoides GCA_000832605.1
317 Bacillus pseudomycoides DSM 12442 GCA_000161455.1
317 _5BBacillus thuringiensis_5D serovar konkukian str. 97-27 GCA_000008505.1
264 Bacillus manliponensis GCA_000712595.1

264 Bacillus marisflavi GCA_001274775.1
5d
255 Clostridium saccharobutylicum DSM 13864 GCA_000473995.1
253 Clostridium beijerinckii GCA_000833105.2
253 Clostridium saccharoperbutylacetonicum N1-4_28HMT_29 GCA_000340885.1
246 Clostridium puniceum GCA_002006345.1
5e
237 Bifidobacterium adolescentis ATCC 15703 GCA_000010425.1
221 Bifidobacterium gallicum DSM 20093 = LMG 11596 GCA_000741205.1
215 Bifidobacterium dentium JCM 1195 = DSM 20436 GCA_001042595.1
5f
201 Escherichia coli IAI39 GCA_000026345.1
201 Escherichia coli O104_3AH4 str. 2011C-3493 GCA_000299455.1
201 Escherichia coli O157_3AH7 str. Sakai GCA_000008865.1
201 Escherichia coli O83_3AH1 str. NRG 857C GCA_000183345.1
201 Escherichia coli UMN026 GCA_000026325.2
201 Escherichia coli str. K-12 substr. MG1655 GCA_000005845.2
201 Shigella flexneri 2a str. 301 GCA_000006925.2
201 Tumebacillus flagellatus GCA_000714935.1
185 Shigella dysenteriae Sd197 GCA_000012005.1
172 Erwinia iniecta GCA_001267535.1
5g
203 Clostridium beijerinckii GCA_000833105.2
203 Clostridium chromiireducens GCA_002029255.1
203 Clostridium puniceum GCA_002006345.1
203 Clostridium saccharobutylicum DSM 13864 GCA_000473995.1
203 Clostridium saccharoperbutylacetonicum N1-4_28HMT_29 GCA_000340885.1
183 Clostridium butyricum GCA_001456065.2
165 Clostridium neonatale GCA_001458595.1
5h
228 Erwinia iniecta GCA_001267535.1
227 Escherichia coli IAI39 GCA_000026345.1
227 Escherichia coli O104_3AH4 str. 2011C-3493 GCA_000299455.1
227 Escherichia coli O157_3AH7 str. Sakai GCA_000008865.1
227 Escherichia coli O83_3AH1 str. NRG 857C GCA_000183345.1
227 Escherichia coli UMN026 GCA_000026325.2
227 Escherichia coli str. K-12 substr. MG1655 GCA_000005845.2
227 Shigella flexneri 2a str. 301 GCA_000006925.2
227 Tumebacillus flagellatus GCA_000714935.1
218 Shigella dysenteriae Sd197 GCA_000012005.1
5i
317 Clostridium chromiireducens GCA_002029255.1
316 Clostridium saccharobutylicum DSM 13864 GCA_000473995.1
314 Clostridium beijerinckii GCA_000833105.2
314 Clostridium puniceum GCA_002006345.1
314 Clostridium saccharoperbutylacetonicum N1-4_28HMT_29 GCA_000340885.1
5j
217 Rhodobacter sphaeroides 2.4.1 GCA_000012905.2
212 Rhodobacter sphaeroides ATCC 17025 GCA_000016405.1
198 Pseudorhodobacter ferrugineus DSM 5888 GCA_000420745.1
198 Pseudorhodobacter wandonensis GCA_001202035.1
5k
277 Lactobacillus gasseri ATCC 33323 = JCM 1131 GCA_000014425.1
252 Lactobacillus hominis DSM 23910 = CRBIP 24.179 GCA_000296835.1
203 Lactobacillus amylophilus DSM 20533 = JCM 1125 GCA_001936335.1
5l
147 Streptococcus mutans UA159 GCA_000007465.2
118 Streptococcus equinus GCA_000964315.1
118 Streptococcus gallolyticus subsp. gallolyticus DSM 16831 GCA_002000985.1
118 Streptococcus orisratti DSM 15617 GCA_000380105.1
118 Streptococcus ratti FA-1 = DSM 20564 GCA_000286075.1
105 Streptococcus agalactiae 2603V_2FR GCA_000007265.1
105 Streptococcus cristatus AS 1.3089 GCA_000385925.1
105 Streptococcus dysgalactiae subsp. equisimilis AC-2713 GCA_000317855.1
105 Streptococcus gordonii str. Challis substr. CH1 GCA_000017005.1
105 Streptococcus ictaluri 707-05 GCA_000188015.3
105 Streptococcus mitis B6 GCA_000027165.1
105 Streptococcus pneumoniae R6 GCA_000007045.1
5m
293 Escherichia coli IAI39 GCA_000026345.1
293 Escherichia coli O104_3AH4 str. 2011C-3493 GCA_000299455.1
293 Escherichia coli O157_3AH7 str. Sakai GCA_000008865.1
293 Escherichia coli O83_3AH1 str. NRG 857C GCA_000183345.1
293 Escherichia coli UMN026 GCA_000026325.2
293 Escherichia coli str. K-12 substr. MG1655 GCA_000005845.2
293 Shigella flexneri 2a str. 301 GCA_000006925.2

293 Tumebacillus flagellatus GCA_000714935.1
284 Shigella dysenteriae Sd197 GCA_000012005.1
276 Erwinia iniecta GCA_001267535.1
5n
187 Deinococcus radiodurans R1 GCA_000008565.1
148 Deinococcus gobiensis I-0 GCA_000252445.1
104 Deinococcus deserti VCD115 GCA_000020685.1
5o
307 Lactobacillus gasseri ATCC 33323 = JCM 1131 GCA_000014425.1
276 Lactobacillus hominis DSM 23910 = CRBIP 24.179 GCA_000296835.1
205 Lactobacillus iners DSM 13335 GCA_000160875.1
5p
240 Staphylococcus epidermidis ATCC 12228 GCA_000007645.1
235 Staphylococcus haemolyticus JCSC1435 GCA_000009865.1
224 Staphylococcus hominis subsp. hominis C80 GCA_000183685.1
224 Staphylococcus lugdunensis HKU09-01 GCA_000025085.1
5q
236 Lactobacillus gasseri ATCC 33323 = JCM 1131 GCA_000014425.1
222 Lactobacillus hominis DSM 23910 = CRBIP 24.179 GCA_000296835.1
184 Lactobacillus amylovorus GCA_000191545.1
184 Lactobacillus crispatus ST1 GCA_000091765.1
5r
187 Bifidobacterium adolescentis ATCC 15703 GCA_000010425.1
172 Bifidobacterium breve DSM 20213 = JCM 1192 GCA_001025175.1
172 Bifidobacterium longum NCC2705 GCA_000007525.1
172 Bifidobacterium saguini DSM 23967 GCA_000741715.1
171 Bifidobacterium tsurumiense GCA_000741765.1
5s
280 Escherichia coli IAI39 GCA_000026345.1
280 Escherichia coli O104_3AH4 str. 2011C-3493 GCA_000299455.1
280 Escherichia coli O157_3AH7 str. Sakai GCA_000008865.1
280 Escherichia coli UMN026 GCA_000026325.2
280 Escherichia coli str. K-12 substr. MG1655 GCA_000005845.2
280 Shigella flexneri 2a str. 301 GCA_000006925.2
280 Tumebacillus flagellatus GCA_000714935.1
279 Escherichia coli O83_3AH1 str. NRG 857C GCA_000183345.1
271 Shigella dysenteriae Sd197 GCA_000012005.1
5t
280 Enterococcus faecalis V583 GCA_000007785.1
280 Streptomyces cinnamoneus GCA_001885705.1
253 Enterococcus dispar ATCC 51266 GCA_000406945.1
250 Enterococcus canis NBRC 100695 GCA_001544375.1
250 Enterococcus faecium DO GCA_000174395.2
250 Enterococcus hirae ATCC 9790 GCA_000271405.2
250 Enterococcus massiliensis GCA_001050095.1
250 Enterococcus mundtii QU 25 GCA_000504125.1
250 Enterococcus rivorum GCA_001742285.1
5u
150 Morganella morganii subsp. morganii KT GCA_000286435.2
148 Serratia rubidaea GCA_001572725.1
147 Klebsiella pneumoniae subsp. pneumoniae HS11286 GCA_000240185.2
5v
175 Erwinia iniecta GCA_001267535.1
175 Erwinia toletana DAPP-PG 735 GCA_000336255.1
175 Escherichia coli IAI39 GCA_000026345.1
175 Escherichia coli O104_3AH4 str. 2011C-3493 GCA_000299455.1
175 Escherichia coli O157_3AH7 str. Sakai GCA_000008865.1
175 Escherichia coli O83_3AH1 str. NRG 857C GCA_000183345.1
175 Escherichia coli UMN026 GCA_000026325.2
175 Escherichia coli str. K-12 substr. MG1655 GCA_000005845.2
175 Shigella dysenteriae Sd197 GCA_000012005.1
175 Shigella flexneri 2a str. 301 GCA_000006925.2
175 Tumebacillus flagellatus GCA_000714935.1
171 Cronobacter sakazakii GCA_000982825.1
167 Pantoea agglomerans GCA_001709315.1
167 Pantoea ananatis LMG 20103 GCA_000025405.2
167 Shimwellia blattae DSM 4481 = NBRC 105725 GCA_000262305.1
5w
96 Enterococcus faecalis V583 GCA_000007785.1
96 Streptomyces cinnamoneus GCA_001885705.1
89 Macrococcus canis GCA_002119805.1
89 Salinicoccus carnicancri Crm GCA_000330705.1
89 Staphylococcus arlettae CVD059 GCA_000295715.1
89 Staphylococcus saprophyticus GCA_001074355.1
89 Staphylococcus vitulinus F1028 GCA_000286335.1
83 Enterococcus hirae ATCC 9790 GCA_000271405.2

83 Vagococcus penaei GCA_001998885.1
5x
236 Rhodobacter sphaeroides 2.4.1 GCA_000012905.2
219 Pseudorhodobacter psychrotolerans GCA_001294535.1
210 Rhodobacter sphaeroides ATCC 17025 GCA_000016405.1
5y
96 Escherichia coli IAI39 GCA_000026345.1
96 Escherichia coli O104_3AH4 str. 2011C-3493 GCA_000299455.1
96 Escherichia coli O157_3AH7 str. Sakai GCA_000008865.1
96 Escherichia coli O83_3AH1 str. NRG 857C GCA_000183345.1
96 Escherichia coli UMN026 GCA_000026325.2
96 Escherichia coli str. K-12 substr. MG1655 GCA_000005845.2
96 Shigella flexneri 2a str. 301 GCA_000006925.2
96 Tumebacillus flagellatus GCA_000714935.1
81 Erwinia iniecta GCA_001267535.1
81 Erwinia toletana DAPP-PG 735 GCA_000336255.1
81 Kosakonia cowanii GCA_001975225.1
81 Pluralibacter gergoviae GCA_000757785.1
81 Shigella dysenteriae Sd197 GCA_000012005.1
81 Sodalis praecaptivus GCA_000517425.1
77 Cronobacter sakazakii GCA_000982825.1
77 Kosakonia sacchari SP1 GCA_000300455.4
5z
261 Staphylococcus capitis subsp. capitis GCA_001028645.1
261 Staphylococcus epidermidis ATCC 12228 GCA_000007645.1
261 Staphylococcus haemolyticus JCSC1435 GCA_000009865.1
244 Staphylococcus lutrae GCA_002101335.1
243 Megasphaera cerevisiae DSM 20462 GCA_001045675.1
243 Staphylococcus aureus subsp. aureus NCTC 8325 GCA_000013425.1
243 Staphylococcus hominis subsp. hominis C80 GCA_000183685.1
243 Staphylococcus lugdunensis HKU09-01 GCA_000025085.1
243 Staphylococcus simiae CCM 7213 GCA_000235645.2
243 Staphylococcus warneri SG1 GCA_000332735.1
5A
245 Bacillus cereus ATCC 14579 GCA_000007825.1
233 Bacillus anthracis str. Ames GCA_000007845.1
233 Bacillus anthracis str. Sterne GCA_000008165.1
233 Bacillus mycoides GCA_000832605.1
233 Bacillus pseudomycoides DSM 12442 GCA_000161455.1
233 Bacillus thuringiensis YBT-1518 GCA_000497525.2
233 _5BBacillus thuringiensis_5D serovar konkukian str. 97-27 GCA_000008505.1
177 Bacillus horneckiae GCA_001636335.1
177 Bacillus solani GCA_001420595.1
5B
213 Clostridium beijerinckii GCA_000833105.2
213 Clostridium puniceum GCA_002006345.1
213 Clostridium saccharobutylicum DSM 13864 GCA_000473995.1
213 Clostridium saccharoperbutylacetonicum N1-4_28HMT_29 GCA_000340885.1
197 Clostridium chromiireducens GCA_002029255.1
188 Clostridium butyricum GCA_001456065.2
5C
269 Enterococcus faecalis V583 GCA_000007785.1
269 Streptomyces cinnamoneus GCA_001885705.1
252 Enterococcus asini ATCC 700915 GCA_000407365.1
252 Enterococcus canis NBRC 100695 GCA_001544375.1
252 Enterococcus casseliflavus EC20 GCA_000157355.2
252 Enterococcus dispar ATCC 51266 GCA_000406945.1
252 Enterococcus faecium DO GCA_000174395.2
252 Enterococcus haemoperoxidus ATCC BAA-382 GCA_000407165.1
252 Enterococcus hirae ATCC 9790 GCA_000271405.2
252 Enterococcus mundtii QU 25 GCA_000504125.1
252 Enterococcus phoeniculicola ATCC BAA-412 GCA_000407505.1
252 Enterococcus rivorum GCA_001742285.1
252 Enterococcus saccharolyticus subsp. saccharolyticus ATCC 43076 GCA_000407285.1
252 Enterococcus thailandicus GCA_001652875.1
242 Enterococcus gilvus ATCC BAA-350 GCA_000407545.1
242 Enterococcus hermanniensis GCA_001885945.1
242 Enterococcus italicus DSM 15952 GCA_000185365.1
242 Enterococcus malodoratus ATCC 43197 GCA_000407185.1
242 Enterococcus pallens ATCC BAA-351 GCA_000407485.1
242 Enterococcus pseudoavium NBRC 100491 GCA_001544295.1
242 Enterococcus sulfureus ATCC 49903 GCA_000407605.1
5D
300 Clostridium beijerinckii GCA_000833105.2
300 Clostridium saccharoperbutylacetonicum N1-4_28HMT_29 GCA_000340885.1
291 Clostridium saccharobutylicum DSM 13864 GCA_000473995.1

286 Clostridium puniceum GCA_002006345.1
5E
325 Clostridium beijerinckii GCA_000833105.2
325 Clostridium puniceum GCA_002006345.1
325 Clostridium saccharoperbutylacetonicum N1-4_28HMT_29 GCA_000340885.1
310 Clostridium saccharobutylicum DSM 13864 GCA_000473995.1
288 Clostridium chromiireducens GCA_002029255.1
5F
187 Clostridium beijerinckii GCA_000833105.2
187 Clostridium saccharobutylicum DSM 13864 GCA_000473995.1
187 Clostridium saccharoperbutylacetonicum N1-4_28HMT_29 GCA_000340885.1
180 Clostridium puniceum GCA_002006345.1
165 Clostridium chromiireducens GCA_002029255.1
165 Clostridium taeniosporum GCA_001735765.1
5G
205 Bifidobacterium adolescentis ATCC 15703 GCA_000010425.1
173 Bifidobacterium tsurumiense GCA_000741765.1
167 Bifidobacterium thermophilum GCA_000741495.1
5H
171 Rhodobacter sphaeroides 2.4.1 GCA_000012905.2
126 Thioclava dalianensis GCA_000715505.1
126 Thioclava indica GCA_000714545.1
124 Pseudaminobacter salicylatoxidans KCT001 GCA_000304395.1
5I
245 Deinococcus radiodurans R1 GCA_000008565.1
122 Deinococcus marmoris DSM 12784 GCA_000701405.1
122 Deinococcus swuensis GCA_000800395.1
110 Deinococcus puniceus GCA_001644565.1
5J
97 Clostridium neonatale GCA_001458595.1
96 Clostridium butyricum GCA_001456065.2
95 Clostridium beijerinckii GCA_000833105.2
95 Clostridium puniceum GCA_002006345.1
95 Clostridium saccharobutylicum DSM 13864 GCA_000473995.1
95 Clostridium saccharoperbutylacetonicum N1-4_28HMT_29 GCA_000340885.1
5K
261 Escherichia coli IAI39 GCA_000026345.1
261 Escherichia coli O104_3AH4 str. 2011C-3493 GCA_000299455.1
261 Escherichia coli O157_3AH7 str. Sakai GCA_000008865.1
261 Escherichia coli O83_3AH1 str. NRG 857C GCA_000183345.1
261 Escherichia coli UMN026 GCA_000026325.2
261 Escherichia coli str. K-12 substr. MG1655 GCA_000005845.2
261 Shigella dysenteriae Sd197 GCA_000012005.1
261 Shigella flexneri 2a str. 301 GCA_000006925.2
261 Tumebacillus flagellatus GCA_000714935.1
238 Enterobacter hormaechei subsp. steigerwaltii GCA_001729725.1
238 Erwinia iniecta GCA_001267535.1
238 Erwinia toletana DAPP-PG 735 GCA_000336255.1
238 Klebsiella oxytoca GCA_001022195.1
238 Kosakonia cowanii GCA_001975225.1
238 Pseudescherichia vulneris NBRC 102420 GCA_000759795.1
237 Kosakonia sacchari SP1 GCA_000300455.4
5L
184 Staphylococcus epidermidis ATCC 12228 GCA_000007645.1
184 Staphylococcus haemolyticus JCSC1435 GCA_000009865.1
179 Staphylococcus capitis subsp. capitis GCA_001028645.1
170 Staphylococcus pettenkoferi GCA_002208805.1
5M
305 Streptococcus mutans UA159 GCA_000007465.2
211 Streptococcus ferus DSM 20646 GCA_000372425.1
206 Streptococcus gordonii str. Challis substr. CH1 GCA_000017005.1
5N
166 Deinococcus radiodurans R1 GCA_000008565.1
110 Deinococcus marmoris DSM 12784 GCA_000701405.1
110 Deinococcus swuensis GCA_000800395.1
105 Deinococcus hopiensis KR-140 GCA_900176165.1
5O
212 Deinococcus radiodurans R1 GCA_000008565.1
150 Deinococcus gobiensis I-0 GCA_000252445.1
133 Deinococcus marmoris DSM 12784 GCA_000701405.1
133 Deinococcus swuensis GCA_000800395.1
5P
205 Clostridium beijerinckii GCA_000833105.2
205 Clostridium puniceum GCA_002006345.1
205 Clostridium saccharoperbutylacetonicum N1-4_28HMT_29 GCA_000340885.1
196 Clostridium saccharobutylicum DSM 13864 GCA_000473995.1

187 Clostridium neonatale GCA_001458595.1
5Q
227 Clostridium beijerinckii GCA_000833105.2
227 Clostridium saccharoperbutylacetonicum N1-4_28HMT_29 GCA_000340885.1
212 Clostridium puniceum GCA_002006345.1
212 Clostridium saccharobutylicum DSM 13864 GCA_000473995.1
185 Clostridium butyricum GCA_001456065.2
5R
154 Clostridium chromiireducens GCA_002029255.1
154 Clostridium saccharobutylicum DSM 13864 GCA_000473995.1
153 Clostridium beijerinckii GCA_000833105.2
153 Clostridium puniceum GCA_002006345.1
153 Clostridium saccharoperbutylacetonicum N1-4_28HMT_29 GCA_000340885.1
138 Clostridium butyricum GCA_001456065.2
5S
139 Clostridium neonatale GCA_001458595.1
135 Clostridium beijerinckii GCA_000833105.2
135 Clostridium puniceum GCA_002006345.1
135 Clostridium saccharobutylicum DSM 13864 GCA_000473995.1
135 Clostridium saccharoperbutylacetonicum N1-4_28HMT_29 GCA_000340885.1
133 Clostridium intestinale URNW GCA_000469625.2
133 Clostridium taeniosporum GCA_001735765.1
5T
319 Clostridium beijerinckii GCA_000833105.2
319 Clostridium saccharobutylicum DSM 13864 GCA_000473995.1
319 Clostridium saccharoperbutylacetonicum N1-4_28HMT_29 GCA_000340885.1
313 Clostridium puniceum GCA_002006345.1
295 Clostridium chromiireducens GCA_002029255.1
5U
264 Deinococcus radiodurans R1 GCA_000008565.1
193 Deinococcus gobiensis I-0 GCA_000252445.1
173 Deinococcus puniceus GCA_001644565.1
5V
267 Escherichia coli IAI39 GCA_000026345.1
267 Escherichia coli O104_3AH4 str. 2011C-3493 GCA_000299455.1
267 Escherichia coli O157_3AH7 str. Sakai GCA_000008865.1
267 Escherichia coli O83_3AH1 str. NRG 857C GCA_000183345.1
267 Escherichia coli UMN026 GCA_000026325.2
267 Escherichia coli str. K-12 substr. MG1655 GCA_000005845.2
267 Shigella flexneri 2a str. 301 GCA_000006925.2
267 Tumebacillus flagellatus GCA_000714935.1
258 Shigella dysenteriae Sd197 GCA_000012005.1
244 Erwinia iniecta GCA_001267535.1
5W
334 Lactobacillus gasseri ATCC 33323 = JCM 1131 GCA_000014425.1
314 Lactobacillus hominis DSM 23910 = CRBIP 24.179 GCA_000296835.1
229 Lactobacillus psittaci DSM 15354 GCA_000425905.1
5X
227 Rhodobacter sphaeroides 2.4.1 GCA_000012905.2
192 Defluviimonas alba GCA_001620265.1
178 Rhodobacter sphaeroides ATCC 17025 GCA_000016405.1
5Y
261 Escherichia coli IAI39 GCA_000026345.1
261 Escherichia coli O104_3AH4 str. 2011C-3493 GCA_000299455.1
261 Escherichia coli O157_3AH7 str. Sakai GCA_000008865.1
261 Escherichia coli O83_3AH1 str. NRG 857C GCA_000183345.1
261 Escherichia coli UMN026 GCA_000026325.2
261 Escherichia coli str. K-12 substr. MG1655 GCA_000005845.2
261 Shigella dysenteriae Sd197 GCA_000012005.1
261 Tumebacillus flagellatus GCA_000714935.1
247 Shigella flexneri 2a str. 301 GCA_000006925.2
233 Cronobacter sakazakii GCA_000982825.1
233 Erwinia iniecta GCA_001267535.1
233 Erwinia toletana DAPP-PG 735 GCA_000336255.1
5Z
268 Streptococcus mutans UA159 GCA_000007465.2
187 Streptococcus ferus DSM 20646 GCA_000372425.1
170 Streptococcus macacae NCTC 11558 GCA_000187995.3
60
242 Deinococcus radiodurans R1 GCA_000008565.1
203 Deinococcus gobiensis I-0 GCA_000252445.1
197 Deinococcus puniceus GCA_001644565.1
61
235 Bifidobacterium adolescentis ATCC 15703 GCA_000010425.1
201 Bifidobacterium asteroides PRL2011 GCA_000304215.1
201 Bifidobacterium thermophilum GCA_000741495.1

200 Bifidobacterium callitrichos DSM 23973 GCA_000741175.1
62
193 Bacillus anthracis str. Ames GCA_000007845.1
193 Bacillus anthracis str. Sterne GCA_000008165.1
193 Bacillus cereus ATCC 14579 GCA_000007825.1
193 Bacillus mycoides GCA_000832605.1
193 Bacillus pseudomycoides DSM 12442 GCA_000161455.1
193 Bacillus thuringiensis YBT-1518 GCA_000497525.2
193 _5BBacillus thuringiensis_5D serovar konkukian str. 97-27 GCA_000008505.1
168 Bacillus cytotoxicus NVH 391-98 GCA_000017425.1
162 Bacillus horneckiae GCA_001636335.1
162 Bacillus solani GCA_001420595.1
63
205 Clostridium beijerinckii GCA_000833105.2
205 Clostridium puniceum GCA_002006345.1
205 Clostridium saccharobutylicum DSM 13864 GCA_000473995.1
205 Clostridium saccharoperbutylacetonicum N1-4_28HMT_29 GCA_000340885.1
190 Clostridium chromiireducens GCA_002029255.1
188 Clostridium butyricum GCA_001456065.2
64
212 Rhodobacter sphaeroides 2.4.1 GCA_000012905.2
183 Rhodobacter sphaeroides ATCC 17025 GCA_000016405.1
152 Defluviimonas alba GCA_001620265.1
152 Gemmobacter megaterium GCA_900156815.1
152 Pseudorhodobacter ferrugineus DSM 5888 GCA_000420745.1
152 Pseudorhodobacter psychrotolerans GCA_001294535.1
152 Pseudorhodobacter wandonensis GCA_001202035.1
65
258 Streptococcus mutans UA159 GCA_000007465.2
163 Streptococcus ferus DSM 20646 GCA_000372425.1
152 Streptococcus sobrinus DSM 20742 = ATCC 33478 GCA_000686605.1
66
244 Enterococcus faecalis V583 GCA_000007785.1
244 Streptomyces cinnamoneus GCA_001885705.1
220 Enterococcus canis NBRC 100695 GCA_001544375.1
220 Enterococcus dispar ATCC 51266 GCA_000406945.1
220 Enterococcus faecium DO GCA_000174395.2
220 Enterococcus gilvus ATCC BAA-350 GCA_000407545.1
220 Enterococcus haemoperoxidus ATCC BAA-382 GCA_000407165.1
220 Enterococcus hermanniensis GCA_001885945.1
220 Enterococcus hirae ATCC 9790 GCA_000271405.2
220 Enterococcus malodoratus ATCC 43197 GCA_000407185.1
220 Enterococcus massiliensis GCA_001050095.1
220 Enterococcus mundtii QU 25 GCA_000504125.1
220 Enterococcus pallens ATCC BAA-351 GCA_000407485.1
220 Enterococcus phoeniculicola ATCC BAA-412 GCA_000407505.1
220 Enterococcus pseudoavium NBRC 100491 GCA_001544295.1
220 Enterococcus rivorum GCA_001742285.1
220 Enterococcus thailandicus GCA_001652875.1
213 Enterococcus asini ATCC 700915 GCA_000407365.1
67
209 Enterococcus faecalis V583 GCA_000007785.1
209 Streptomyces cinnamoneus GCA_001885705.1
174 Enterococcus canis NBRC 100695 GCA_001544375.1
174 Enterococcus casseliflavus EC20 GCA_000157355.2
174 Enterococcus dispar ATCC 51266 GCA_000406945.1
174 Enterococcus faecium DO GCA_000174395.2
174 Enterococcus hirae ATCC 9790 GCA_000271405.2
174 Enterococcus mundtii QU 25 GCA_000504125.1
174 Enterococcus rivorum GCA_001742285.1
174 Enterococcus saccharolyticus subsp. saccharolyticus ATCC 43076 GCA_000407285.1
163 Enterococcus gilvus ATCC BAA-350 GCA_000407545.1
163 Enterococcus hermanniensis GCA_001885945.1
163 Enterococcus malodoratus ATCC 43197 GCA_000407185.1
163 Enterococcus pallens ATCC BAA-351 GCA_000407485.1
163 Enterococcus pseudoavium NBRC 100491 GCA_001544295.1
68
244 Bifidobacterium adolescentis ATCC 15703 GCA_000010425.1
220 Bifidobacterium thermophilum GCA_000741495.1
215 Bifidobacterium breve DSM 20213 = JCM 1192 GCA_001025175.1
69
255 Escherichia coli IAI39 GCA_000026345.1
255 Escherichia coli O104_3AH4 str. 2011C-3493 GCA_000299455.1
255 Escherichia coli O157_3AH7 str. Sakai GCA_000008865.1
255 Escherichia coli O83_3AH1 str. NRG 857C GCA_000183345.1
255 Escherichia coli UMN026 GCA_000026325.2

255 Escherichia coli str. K-12 substr. MG1655 GCA_000005845.2
255 Shigella flexneri 2a str. 301 GCA_000006925.2
255 Tumebacillus flagellatus GCA_000714935.1
246 Shigella dysenteriae Sd197 GCA_000012005.1
231 Erwinia iniecta GCA_001267535.1
6a
201 Bacillus thuringiensis YBT-1518 GCA_000497525.2
185 Bacillus anthracis str. Ames GCA_000007845.1
185 Bacillus anthracis str. Sterne GCA_000008165.1
185 Bacillus cereus ATCC 14579 GCA_000007825.1
185 Bacillus mycoides GCA_000832605.1
185 Bacillus pseudomycoides DSM 12442 GCA_000161455.1
185 _5BBacillus thuringiensis_5D serovar konkukian str. 97-27 GCA_000008505.1
181 Bacillus horneckiae GCA_001636335.1
181 Bacillus marisflavi GCA_001274775.1
181 Bacillus solani GCA_001420595.1
6b
244 Escherichia coli IAI39 GCA_000026345.1
244 Escherichia coli O104_3AH4 str. 2011C-3493 GCA_000299455.1
244 Escherichia coli O157_3AH7 str. Sakai GCA_000008865.1
244 Escherichia coli O83_3AH1 str. NRG 857C GCA_000183345.1
244 Escherichia coli UMN026 GCA_000026325.2
244 Escherichia coli str. K-12 substr. MG1655 GCA_000005845.2
244 Shigella dysenteriae Sd197 GCA_000012005.1
244 Shigella flexneri 2a str. 301 GCA_000006925.2
244 Tumebacillus flagellatus GCA_000714935.1
238 Erwinia iniecta GCA_001267535.1
203 Rosenbergiella nectarea GCA_900111105.1
6c
147 Deinococcus radiodurans R1 GCA_000008565.1
81 Deinococcus marmoris DSM 12784 GCA_000701405.1
81 Deinococcus swuensis GCA_000800395.1
76 Deinococcus gobiensis I-0 GCA_000252445.1
76 Deinococcus puniceus GCA_001644565.1
6d
165 Enterococcus faecalis V583 GCA_000007785.1
165 Streptomyces cinnamoneus GCA_001885705.1
155 Enterococcus canis NBRC 100695 GCA_001544375.1
155 Enterococcus casseliflavus EC20 GCA_000157355.2
155 Enterococcus dispar ATCC 51266 GCA_000406945.1
155 Enterococcus faecium DO GCA_000174395.2
155 Enterococcus haemoperoxidus ATCC BAA-382 GCA_000407165.1
155 Enterococcus hirae ATCC 9790 GCA_000271405.2
155 Enterococcus massiliensis GCA_001050095.1
155 Enterococcus mundtii QU 25 GCA_000504125.1
155 Enterococcus phoeniculicola ATCC BAA-412 GCA_000407505.1
155 Enterococcus rivorum GCA_001742285.1
155 Enterococcus saccharolyticus subsp. saccharolyticus ATCC 43076 GCA_000407285.1
155 Enterococcus thailandicus GCA_001652875.1
145 Enterococcus aquimarinus GCA_001885765.1
145 Enterococcus gilvus ATCC BAA-350 GCA_000407545.1
145 Enterococcus hermanniensis GCA_001885945.1
145 Enterococcus malodoratus ATCC 43197 GCA_000407185.1
145 Enterococcus pallens ATCC BAA-351 GCA_000407485.1
145 Enterococcus pseudoavium NBRC 100491 GCA_001544295.1
6e
180 Bifidobacterium adolescentis ATCC 15703 GCA_000010425.1
179 Bifidobacterium longum NCC2705 GCA_000007525.1
177 Bifidobacterium breve DSM 20213 = JCM 1192 GCA_001025175.1
6f
142 Escherichia coli IAI39 GCA_000026345.1
142 Escherichia coli O104_3AH4 str. 2011C-3493 GCA_000299455.1
142 Escherichia coli O157_3AH7 str. Sakai GCA_000008865.1
142 Escherichia coli O83_3AH1 str. NRG 857C GCA_000183345.1
142 Escherichia coli UMN026 GCA_000026325.2
142 Escherichia coli str. K-12 substr. MG1655 GCA_000005845.2
142 Shigella dysenteriae Sd197 GCA_000012005.1
142 Shigella flexneri 2a str. 301 GCA_000006925.2
142 Tumebacillus flagellatus GCA_000714935.1
128 Pantoea agglomerans GCA_001709315.1
127 Brenneria goodwinii GCA_001049335.1
127 Enterobacter hormaechei subsp. steigerwaltii GCA_001729725.1
127 Erwinia billingiae Eb661 GCA_000196615.1
127 Erwinia iniecta GCA_001267535.1
127 Erwinia toletana DAPP-PG 735 GCA_000336255.1
127 Klebsiella oxytoca GCA_001022195.1

127 Kosakonia cowanii GCA_001975225.1
127 Pantoea alhagi GCA_002101395.1
127 Pantoea ananatis LMG 20103 GCA_000025405.2
127 Pantoea rwandensis GCA_000759475.1
127 Pantoea septica GCA_002095575.1
127 Pluralibacter gergoviae GCA_000757785.1
127 Pseudescherichia vulneris NBRC 102420 GCA_000759795.1
127 Shimwellia blattae DSM 4481 = NBRC 105725 GCA_000262305.1
6g
160 Bifidobacterium adolescentis ATCC 15703 GCA_000010425.1
149 Bifidobacterium animalis subsp. lactis DSM 10140 GCA_000022965.1
145 Bifidobacterium dentium JCM 1195 = DSM 20436 GCA_001042595.1
145 Bifidobacterium tsurumiense GCA_000741765.1
6h
206 Deinococcus radiodurans R1 GCA_000008565.1
141 Deinococcus gobiensis I-0 GCA_000252445.1
120 Deinococcus deserti VCD115 GCA_000020685.1
6i
213 Escherichia coli IAI39 GCA_000026345.1
213 Escherichia coli O104_3AH4 str. 2011C-3493 GCA_000299455.1
213 Escherichia coli O157_3AH7 str. Sakai GCA_000008865.1
213 Escherichia coli O83_3AH1 str. NRG 857C GCA_000183345.1
213 Escherichia coli UMN026 GCA_000026325.2
213 Escherichia coli str. K-12 substr. MG1655 GCA_000005845.2
213 Shigella dysenteriae Sd197 GCA_000012005.1
213 Shigella flexneri 2a str. 301 GCA_000006925.2
213 Tumebacillus flagellatus GCA_000714935.1
197 Erwinia iniecta GCA_001267535.1
197 Erwinia toletana DAPP-PG 735 GCA_000336255.1
197 Pantoea ananatis LMG 20103 GCA_000025405.2
197 Shimwellia blattae DSM 4481 = NBRC 105725 GCA_000262305.1
193 Cronobacter sakazakii GCA_000982825.1
193 Pantoea dispersa EGD-AAK13 GCA_000465555.2
6j
277 Deinococcus radiodurans R1 GCA_000008565.1
181 Deinococcus gobiensis I-0 GCA_000252445.1
153 Deinococcus puniceus GCA_001644565.1
6k
214 _5BBacillus thuringiensis_5D serovar konkukian str. 97-27 GCA_000008505.1
204 Bacillus anthracis str. Ames GCA_000007845.1
204 Bacillus anthracis str. Sterne GCA_000008165.1
204 Bacillus cereus ATCC 14579 GCA_000007825.1
204 Bacillus mycoides GCA_000832605.1
204 Bacillus pseudomycoides DSM 12442 GCA_000161455.1
204 Bacillus thuringiensis YBT-1518 GCA_000497525.2
168 Bacillus marisflavi GCA_001274775.1
6l
216 Staphylococcus epidermidis ATCC 12228 GCA_000007645.1
216 Staphylococcus haemolyticus JCSC1435 GCA_000009865.1
202 Staphylococcus capitis subsp. capitis GCA_001028645.1
200 Staphylococcus aureus subsp. aureus NCTC 8325 GCA_000013425.1
200 Staphylococcus condimenti GCA_001618885.1
200 Staphylococcus hominis subsp. hominis C80 GCA_000183685.1
200 Staphylococcus simiae CCM 7213 GCA_000235645.2
200 Staphylococcus simulans GCA_001559115.1
6m
199 Bacillus anthracis str. Ames GCA_000007845.1
199 Bacillus anthracis str. Sterne GCA_000008165.1
199 Bacillus cereus ATCC 14579 GCA_000007825.1
199 Bacillus mycoides GCA_000832605.1
199 Bacillus pseudomycoides DSM 12442 GCA_000161455.1
199 Bacillus thuringiensis YBT-1518 GCA_000497525.2
199 _5BBacillus thuringiensis_5D serovar konkukian str. 97-27 GCA_000008505.1
181 Staphylococcus lentus F1142 GCA_000286395.1
181 Staphylococcus sciuri GCA_002209165.1
178 Bacillus manliponensis GCA_000712595.1
6n
217 Clostridium beijerinckii GCA_000833105.2
217 Clostridium saccharobutylicum DSM 13864 GCA_000473995.1
217 Clostridium saccharoperbutylacetonicum N1-4_28HMT_29 GCA_000340885.1
210 Clostridium puniceum GCA_002006345.1
185 Clostridium neonatale GCA_001458595.1
6o
148 Clostridium butyricum GCA_001456065.2
147 Clostridium beijerinckii GCA_000833105.2
147 Clostridium botulinum B str. Eklund 17B _28NRP_29 GCA_000020165.1

147 Clostridium chromiireducens GCA_002029255.1
147 Clostridium puniceum GCA_002006345.1
147 Clostridium saccharobutylicum DSM 13864 GCA_000473995.1
147 Clostridium saccharoperbutylacetonicum N1-4_28HMT_29 GCA_000340885.1
147 Clostridium taeniosporum GCA_001735765.1
145 Clostridium neonatale GCA_001458595.1
6p
197 Clostridium beijerinckii GCA_000833105.2
197 Clostridium saccharobutylicum DSM 13864 GCA_000473995.1
197 Clostridium saccharoperbutylacetonicum N1-4_28HMT_29 GCA_000340885.1
182 Clostridium puniceum GCA_002006345.1
166 Clostridium chromiireducens GCA_002029255.1
6q
287 Bifidobacterium adolescentis ATCC 15703 GCA_000010425.1
253 Bifidobacterium dentium JCM 1195 = DSM 20436 GCA_001042595.1
243 Bifidobacterium tsurumiense GCA_000741765.1
6r
189 Clostridium puniceum GCA_002006345.1
176 Clostridium beijerinckii GCA_000833105.2
176 Clostridium saccharoperbutylacetonicum N1-4_28HMT_29 GCA_000340885.1
173 Clostridium taeniosporum GCA_001735765.1
6s
248 Clostridium beijerinckii GCA_000833105.2
248 Clostridium saccharobutylicum DSM 13864 GCA_000473995.1
248 Clostridium saccharoperbutylacetonicum N1-4_28HMT_29 GCA_000340885.1
239 Clostridium butyricum GCA_001456065.2
233 Clostridium puniceum GCA_002006345.1
6t
156 Streptococcus mutans UA159 GCA_000007465.2
122 Streptococcus macacae NCTC 11558 GCA_000187995.3
120 Streptococcus ferus DSM 20646 GCA_000372425.1
6u
220 Erwinia iniecta GCA_001267535.1
220 Escherichia coli IAI39 GCA_000026345.1
220 Escherichia coli O104_3AH4 str. 2011C-3493 GCA_000299455.1
220 Escherichia coli O157_3AH7 str. Sakai GCA_000008865.1
220 Escherichia coli UMN026 GCA_000026325.2
220 Escherichia coli str. K-12 substr. MG1655 GCA_000005845.2
220 Shigella flexneri 2a str. 301 GCA_000006925.2
220 Tumebacillus flagellatus GCA_000714935.1
219 Escherichia coli O83_3AH1 str. NRG 857C GCA_000183345.1
219 Xenorhabdus bovienii SS-2004 GCA_000027225.1
217 Cronobacter sakazakii GCA_000982825.1
217 Erwinia toletana DAPP-PG 735 GCA_000336255.1
217 Kosakonia cowanii GCA_001975225.1
217 Kosakonia sacchari SP1 GCA_000300455.4
6v
197 Enterococcus faecalis V583 GCA_000007785.1
197 Streptomyces cinnamoneus GCA_001885705.1
164 Enterococcus hirae ATCC 9790 GCA_000271405.2
153 Enterococcus dispar ATCC 51266 GCA_000406945.1
6w
337 Bacillus anthracis str. Ames GCA_000007845.1
337 Bacillus anthracis str. Sterne GCA_000008165.1
337 Bacillus cereus ATCC 14579 GCA_000007825.1
337 Bacillus mycoides GCA_000832605.1
337 Bacillus pseudomycoides DSM 12442 GCA_000161455.1
337 Bacillus thuringiensis YBT-1518 GCA_000497525.2
337 _5BBacillus thuringiensis_5D serovar konkukian str. 97-27 GCA_000008505.1
281 Bacillus manliponensis GCA_000712595.1
264 Massilibacterium senegalense GCA_001375675.1
6x
253 Bacillus anthracis str. Ames GCA_000007845.1
253 Bacillus anthracis str. Sterne GCA_000008165.1
253 Bacillus cereus ATCC 14579 GCA_000007825.1
253 Bacillus mycoides GCA_000832605.1
253 Bacillus pseudomycoides DSM 12442 GCA_000161455.1
253 Bacillus thuringiensis YBT-1518 GCA_000497525.2
253 _5BBacillus thuringiensis_5D serovar konkukian str. 97-27 GCA_000008505.1
209 Bacillus manliponensis GCA_000712595.1
193 Bacillus cytotoxicus NVH 391-98 GCA_000017425.1
6y
240 Lactobacillus gasseri ATCC 33323 = JCM 1131 GCA_000014425.1
232 Lactobacillus hominis DSM 23910 = CRBIP 24.179 GCA_000296835.1
152 Lactobacillus amylophilus DSM 20533 = JCM 1125 GCA_001936335.1
6z

151 Serratia symbiotica GCA_000821185.1
146 Escherichia coli IAI39 GCA_000026345.1
146 Escherichia coli O104_3AH4 str. 2011C-3493 GCA_000299455.1
146 Escherichia coli O157_3AH7 str. Sakai GCA_000008865.1
146 Escherichia coli O83_3AH1 str. NRG 857C GCA_000183345.1
146 Escherichia coli UMN026 GCA_000026325.2
146 Escherichia coli str. K-12 substr. MG1655 GCA_000005845.2
146 Shigella dysenteriae Sd197 GCA_000012005.1
146 Shigella flexneri 2a str. 301 GCA_000006925.2
146 Tumebacillus flagellatus GCA_000714935.1
144 Bacillus humi GCA_001439915.1
6A
278 Deinococcus radiodurans R1 GCA_000008565.1
219 Deinococcus gobiensis I-0 GCA_000252445.1
187 Deinococcus marmoris DSM 12784 GCA_000701405.1
187 Deinococcus swuensis GCA_000800395.1
6B
220 Rhodobacter sphaeroides 2.4.1 GCA_000012905.2
204 Rhodobacter capsulatus SB 1003 GCA_000021865.1
203 Haematobacter massiliensis GCA_000740795.1
6C
138 Bifidobacterium adolescentis ATCC 15703 GCA_000010425.1
129 Bifidobacterium asteroides PRL2011 GCA_000304215.1
126 Bifidobacterium callitrichos DSM 23973 GCA_000741175.1
126 Bifidobacterium coryneforme GCA_000737865.1
6D
185 Rhodobacter sphaeroides 2.4.1 GCA_000012905.2
185 Rhodobacter sphaeroides ATCC 17025 GCA_000016405.1
182 Pseudorhodobacter ferrugineus DSM 5888 GCA_000420745.1
182 Pseudorhodobacter psychrotolerans GCA_001294535.1
182 Pseudorhodobacter wandonensis GCA_001202035.1
167 Pseudorhodobacter aquimaris GCA_001202025.1
6E
182 Clostridium beijerinckii GCA_000833105.2
182 Clostridium puniceum GCA_002006345.1
182 Clostridium saccharoperbutylacetonicum N1-4_28HMT_29 GCA_000340885.1
180 Clostridium chromiireducens GCA_002029255.1
180 Clostridium saccharobutylicum DSM 13864 GCA_000473995.1
178 Clostridium butyricum GCA_001456065.2
6F
281 Deinococcus radiodurans R1 GCA_000008565.1
205 Deinococcus gobiensis I-0 GCA_000252445.1
196 Deinococcus puniceus GCA_001644565.1
6G
156 Enterococcus faecalis V583 GCA_000007785.1
156 Streptomyces cinnamoneus GCA_001885705.1
146 Enterococcus canis NBRC 100695 GCA_001544375.1
146 Enterococcus faecium DO GCA_000174395.2
146 Enterococcus haemoperoxidus ATCC BAA-382 GCA_000407165.1
146 Enterococcus hirae ATCC 9790 GCA_000271405.2
146 Enterococcus mundtii QU 25 GCA_000504125.1
146 Enterococcus phoeniculicola ATCC BAA-412 GCA_000407505.1
146 Enterococcus rivorum GCA_001742285.1
146 Enterococcus thailandicus GCA_001652875.1
145 Enterococcus casseliflavus EC20 GCA_000157355.2
145 Enterococcus dispar ATCC 51266 GCA_000406945.1
145 Enterococcus saccharolyticus subsp. saccharolyticus ATCC 43076 GCA_000407285.1
6H
298 Clostridium beijerinckii GCA_000833105.2
298 Clostridium puniceum GCA_002006345.1
298 Clostridium saccharoperbutylacetonicum N1-4_28HMT_29 GCA_000340885.1
290 Clostridium butyricum GCA_001456065.2
283 Clostridium saccharobutylicum DSM 13864 GCA_000473995.1
6I
197 Escherichia coli IAI39 GCA_000026345.1
197 Escherichia coli O104_3AH4 str. 2011C-3493 GCA_000299455.1
197 Escherichia coli O157_3AH7 str. Sakai GCA_000008865.1
197 Escherichia coli O83_3AH1 str. NRG 857C GCA_000183345.1
197 Escherichia coli UMN026 GCA_000026325.2
197 Escherichia coli str. K-12 substr. MG1655 GCA_000005845.2
197 Shigella flexneri 2a str. 301 GCA_000006925.2
197 Tumebacillus flagellatus GCA_000714935.1
188 Shigella dysenteriae Sd197 GCA_000012005.1
172 Erwinia iniecta GCA_001267535.1
6J
304 Bacillus anthracis str. Ames GCA_000007845.1

304 Bacillus anthracis str. Sterne GCA_000008165.1
304 Bacillus cereus ATCC 14579 GCA_000007825.1
304 Bacillus mycoides GCA_000832605.1
304 Bacillus pseudomycoides DSM 12442 GCA_000161455.1
304 Bacillus thuringiensis YBT-1518 GCA_000497525.2
304 _5BBacillus thuringiensis_5D serovar konkukian str. 97-27 GCA_000008505.1
225 Bacillus manliponensis GCA_000712595.1
223 Bacillus akibai JCM 9157 GCA_000513135.1
223 Bacillus cytotoxicus NVH 391-98 GCA_000017425.1
223 Bacillus marisflavi GCA_001274775.1
223 Bacillus pseudofirmus OF4 GCA_000005825.2
6K
157 Escherichia coli IAI39 GCA_000026345.1
157 Escherichia coli O104_3AH4 str. 2011C-3493 GCA_000299455.1
157 Escherichia coli O157_3AH7 str. Sakai GCA_000008865.1
157 Escherichia coli O83_3AH1 str. NRG 857C GCA_000183345.1
157 Escherichia coli UMN026 GCA_000026325.2
157 Escherichia coli str. K-12 substr. MG1655 GCA_000005845.2
157 Shigella dysenteriae Sd197 GCA_000012005.1
157 Shigella flexneri 2a str. 301 GCA_000006925.2
157 Tumebacillus flagellatus GCA_000714935.1
150 Cedecea neteri GCA_000757825.1
150 Erwinia iniecta GCA_001267535.1
150 Rosenbergiella nectarea GCA_900111105.1
135 Kosakonia cowanii GCA_001975225.1
6L
216 Lactobacillus gasseri ATCC 33323 = JCM 1131 GCA_000014425.1
193 Lactobacillus hominis DSM 23910 = CRBIP 24.179 GCA_000296835.1
160 Lactobacillus psittaci DSM 15354 GCA_000425905.1
6M
216 Clostridium beijerinckii GCA_000833105.2
216 Clostridium puniceum GCA_002006345.1
216 Clostridium saccharobutylicum DSM 13864 GCA_000473995.1
216 Clostridium saccharoperbutylacetonicum N1-4_28HMT_29 GCA_000340885.1
212 Clostridium chromiireducens GCA_002029255.1
195 Clostridium botulinum B str. Eklund 17B _28NRP_29 GCA_000020165.1
195 Clostridium taeniosporum GCA_001735765.1
6N
261 Deinococcus radiodurans R1 GCA_000008565.1
212 Deinococcus gobiensis I-0 GCA_000252445.1
185 Deinococcus deserti VCD115 GCA_000020685.1
6O
277 Rhodobacter sphaeroides 2.4.1 GCA_000012905.2
236 Thioclava dalianensis GCA_000715505.1
236 Thioclava indica GCA_000714545.1
230 Defluviimonas alba GCA_001620265.1
6P
194 Rhodobacter sphaeroides 2.4.1 GCA_000012905.2
153 Rhodobacter sphaeroides ATCC 17025 GCA_000016405.1
146 Gemmobacter aquatilis GCA_900110025.1
146 Gemmobacter nectariphilus DSM 15620 GCA_000429765.1
146 Haematobacter massiliensis GCA_000740795.1
146 Pseudorhodobacter psychrotolerans GCA_001294535.1
146 Rhodobacter capsulatus SB 1003 GCA_000021865.1
6Q
192 Bacillus anthracis str. Ames GCA_000007845.1
192 Bacillus anthracis str. Sterne GCA_000008165.1
192 Bacillus cereus ATCC 14579 GCA_000007825.1
192 Bacillus mycoides GCA_000832605.1
192 Bacillus pseudomycoides DSM 12442 GCA_000161455.1
192 Bacillus thuringiensis YBT-1518 GCA_000497525.2
192 _5BBacillus thuringiensis_5D serovar konkukian str. 97-27 GCA_000008505.1
157 Bacillus manliponensis GCA_000712595.1
153 Bacillus cytotoxicus NVH 391-98 GCA_000017425.1
153 Bacillus humi GCA_001439915.1
153 Bacillus sinesaloumensis GCA_900156865.1
153 Bacillus timonensis GCA_000285535.1
6R
194 Escherichia coli IAI39 GCA_000026345.1
194 Escherichia coli O104_3AH4 str. 2011C-3493 GCA_000299455.1
194 Escherichia coli O157_3AH7 str. Sakai GCA_000008865.1
194 Escherichia coli O83_3AH1 str. NRG 857C GCA_000183345.1
194 Escherichia coli UMN026 GCA_000026325.2
194 Escherichia coli str. K-12 substr. MG1655 GCA_000005845.2
194 Shigella dysenteriae Sd197 GCA_000012005.1
194 Shigella flexneri 2a str. 301 GCA_000006925.2

194 Tumebacillus flagellatus GCA_000714935.1
159 Erwinia iniecta GCA_001267535.1
153 Erwinia toletana DAPP-PG 735 GCA_000336255.1
6S
163 Bacillus anthracis str. Ames GCA_000007845.1
163 Bacillus anthracis str. Sterne GCA_000008165.1
163 Bacillus cereus ATCC 14579 GCA_000007825.1
163 Bacillus mycoides GCA_000832605.1
163 Bacillus pseudomycoides DSM 12442 GCA_000161455.1
163 Bacillus thuringiensis YBT-1518 GCA_000497525.2
163 _5BBacillus thuringiensis_5D serovar konkukian str. 97-27 GCA_000008505.1
145 Bacillus cytotoxicus NVH 391-98 GCA_000017425.1
143 Bacillus manliponensis GCA_000712595.1
6T
117 Clostridium beijerinckii GCA_000833105.2
117 Clostridium puniceum GCA_002006345.1
117 Clostridium saccharoperbutylacetonicum N1-4_28HMT_29 GCA_000340885.1
117 Clostridium taeniosporum GCA_001735765.1
105 Clostridium butyricum GCA_001456065.2
102 Clostridium chromiireducens GCA_002029255.1
102 Clostridium saccharobutylicum DSM 13864 GCA_000473995.1
6U
307 Deinococcus radiodurans R1 GCA_000008565.1
188 Deinococcus gobiensis I-0 GCA_000252445.1
178 Deinococcus geothermalis DSM 11300 GCA_000196275.1
6V
255 Escherichia coli IAI39 GCA_000026345.1
255 Escherichia coli O104_3AH4 str. 2011C-3493 GCA_000299455.1
255 Escherichia coli O157_3AH7 str. Sakai GCA_000008865.1
255 Escherichia coli O83_3AH1 str. NRG 857C GCA_000183345.1
255 Escherichia coli UMN026 GCA_000026325.2
255 Escherichia coli str. K-12 substr. MG1655 GCA_000005845.2
255 Shigella dysenteriae Sd197 GCA_000012005.1
255 Shigella flexneri 2a str. 301 GCA_000006925.2
255 Tumebacillus flagellatus GCA_000714935.1
235 Erwinia iniecta GCA_001267535.1
205 Cedecea neteri GCA_000757825.1
205 Rosenbergiella nectarea GCA_900111105.1
6W
228 Bifidobacterium callitrichos DSM 23973 GCA_000741175.1
223 Bifidobacterium adolescentis ATCC 15703 GCA_000010425.1
223 Bifidobacterium angulatum DSM 20098 = JCM 7096 GCA_001025155.1
223 Bifidobacterium stellenboschense GCA_000741785.1
212 Bifidobacterium dentium JCM 1195 = DSM 20436 GCA_001042595.1
212 Bifidobacterium saguini DSM 23967 GCA_000741715.1
212 Bifidobacterium thermophilum GCA_000741495.1
212 Bifidobacterium tsurumiense GCA_000741765.1
6X
172 Bifidobacterium adolescentis ATCC 15703 GCA_000010425.1
139 Bifidobacterium dentium JCM 1195 = DSM 20436 GCA_001042595.1
133 Bifidobacterium asteroides PRL2011 GCA_000304215.1
6Y
203 Rhodobacter sphaeroides 2.4.1 GCA_000012905.2
203 Rhodobacter sphaeroides ATCC 17025 GCA_000016405.1
183 Gemmobacter megaterium GCA_900156815.1
180 Pseudorhodobacter ferrugineus DSM 5888 GCA_000420745.1
180 Pseudorhodobacter wandonensis GCA_001202035.1
6Z
228 Clostridium beijerinckii GCA_000833105.2
228 Clostridium puniceum GCA_002006345.1
228 Clostridium saccharobutylicum DSM 13864 GCA_000473995.1
228 Clostridium saccharoperbutylacetonicum N1-4_28HMT_29 GCA_000340885.1
217 Clostridium chromiireducens GCA_002029255.1
211 Clostridium taeniosporum GCA_001735765.1
70
278 Enterococcus faecalis V583 GCA_000007785.1
278 Streptomyces cinnamoneus GCA_001885705.1
243 Enterococcus asini ATCC 700915 GCA_000407365.1
243 Enterococcus canis NBRC 100695 GCA_001544375.1
243 Enterococcus dispar ATCC 51266 GCA_000406945.1
243 Enterococcus faecium DO GCA_000174395.2
243 Enterococcus haemoperoxidus ATCC BAA-382 GCA_000407165.1
243 Enterococcus hirae ATCC 9790 GCA_000271405.2
243 Enterococcus mundtii QU 25 GCA_000504125.1
243 Enterococcus phoeniculicola ATCC BAA-412 GCA_000407505.1
243 Enterococcus rivorum GCA_001742285.1

243 Enterococcus thailandicus GCA_001652875.1
234 Enterococcus casseliflavus EC20 GCA_000157355.2
234 Enterococcus massiliensis GCA_001050095.1
234 Enterococcus saccharolyticus subsp. saccharolyticus ATCC 43076 GCA_000407285.1
71
248 Deinococcus radiodurans R1 GCA_000008565.1
186 Deinococcus gobiensis I-0 GCA_000252445.1
178 Deinococcus deserti VCD115 GCA_000020685.1
72
276 Streptococcus mutans UA159 GCA_000007465.2
152 Streptococcus gordonii str. Challis substr. CH1 GCA_000017005.1
151 Streptococcus ferus DSM 20646 GCA_000372425.1
73
242 Rhodobacter sphaeroides 2.4.1 GCA_000012905.2
213 Pseudorhodobacter psychrotolerans GCA_001294535.1
210 Rhodobacter sphaeroides ATCC 17025 GCA_000016405.1
74
157 Staphylococcus saprophyticus GCA_001074355.1
156 Staphylococcus capitis subsp. capitis GCA_001028645.1
156 Staphylococcus epidermidis ATCC 12228 GCA_000007645.1
156 Staphylococcus haemolyticus JCSC1435 GCA_000009865.1
152 Staphylococcus cohnii subsp. cohnii GCA_000972575.1
75
159 Staphylococcus aureus subsp. aureus NCTC 8325 GCA_000013425.1
159 Staphylococcus capitis subsp. capitis GCA_001028645.1
159 Staphylococcus epidermidis ATCC 12228 GCA_000007645.1
159 Staphylococcus haemolyticus JCSC1435 GCA_000009865.1
159 Staphylococcus hominis subsp. hominis C80 GCA_000183685.1
159 Staphylococcus lugdunensis HKU09-01 GCA_000025085.1
159 Staphylococcus simiae CCM 7213 GCA_000235645.2
155 Staphylococcus arlettae CVD059 GCA_000295715.1
155 Staphylococcus cohnii subsp. cohnii GCA_000972575.1
155 Staphylococcus equorum GCA_001432245.1
155 Staphylococcus gallinarum GCA_000875895.1
155 Staphylococcus saprophyticus GCA_001074355.1
155 Staphylococcus saprophyticus subsp. saprophyticus ATCC 15305 GCA_000010125.1
155 Staphylococcus succinus GCA_001902315.1
155 Staphylococcus xylosus GCA_000706685.1
150 Staphylococcus condimenti GCA_001618885.1
150 Staphylococcus simulans GCA_001559115.1
76
246 Enterococcus faecalis V583 GCA_000007785.1
246 Streptomyces cinnamoneus GCA_001885705.1
225 Melissococcus plutonius S1 GCA_000747585.1
223 Enterococcus asini ATCC 700915 GCA_000407365.1
223 Enterococcus canis NBRC 100695 GCA_001544375.1
223 Enterococcus dispar ATCC 51266 GCA_000406945.1
223 Enterococcus faecium DO GCA_000174395.2
223 Enterococcus gilvus ATCC BAA-350 GCA_000407545.1
223 Enterococcus hermanniensis GCA_001885945.1
223 Enterococcus hirae ATCC 9790 GCA_000271405.2
223 Enterococcus italicus DSM 15952 GCA_000185365.1
223 Enterococcus malodoratus ATCC 43197 GCA_000407185.1
223 Enterococcus massiliensis GCA_001050095.1
223 Enterococcus mundtii QU 25 GCA_000504125.1
223 Enterococcus pallens ATCC BAA-351 GCA_000407485.1
223 Enterococcus pseudoavium NBRC 100491 GCA_001544295.1
223 Enterococcus rivorum GCA_001742285.1
77
257 Streptococcus mutans UA159 GCA_000007465.2
178 Streptococcus ferus DSM 20646 GCA_000372425.1
167 Streptococcus macacae NCTC 11558 GCA_000187995.3
78
111 Staphylococcus capitis subsp. capitis GCA_001028645.1
111 Staphylococcus epidermidis ATCC 12228 GCA_000007645.1
111 Staphylococcus pettenkoferi GCA_002208805.1
108 Staphylococcus cohnii subsp. cohnii GCA_000972575.1
108 Staphylococcus lugdunensis HKU09-01 GCA_000025085.1
108 Staphylococcus saprophyticus subsp. saprophyticus ATCC 15305 GCA_000010125.1
108 Staphylococcus xylosus GCA_000706685.1
100 Staphylococcus haemolyticus JCSC1435 GCA_000009865.1
79
230 Deinococcus radiodurans R1 GCA_000008565.1
152 Deinococcus proteolyticus MRP GCA_000190555.1
144 Deinococcus deserti VCD115 GCA_000020685.1
144 Deinococcus hopiensis KR-140 GCA_900176165.1

144 Deinococcus puniceus GCA_001644565.1
144 Deinococcus soli Cha et al. 2016 GCA_001007995.1
7a
252 Megasphaera cerevisiae DSM 20462 GCA_001045675.1
252 Staphylococcus cohnii subsp. cohnii GCA_000972575.1
252 Staphylococcus epidermidis ATCC 12228 GCA_000007645.1
252 Staphylococcus warneri SG1 GCA_000332735.1
247 Staphylococcus capitis subsp. capitis GCA_001028645.1
239 Staphylococcus lugdunensis HKU09-01 GCA_000025085.1
7b
209 Lactobacillus gasseri ATCC 33323 = JCM 1131 GCA_000014425.1
193 Lactobacillus hominis DSM 23910 = CRBIP 24.179 GCA_000296835.1
149 Lactobacillus psittaci DSM 15354 GCA_000425905.1
7c
76 Streptococcus mutans UA159 GCA_000007465.2
55 Streptococcus gordonii str. Challis substr. CH1 GCA_000017005.1
46 Mesoplasma photuris ATCC 49581 GCA_000702725.1
46 Sporosarcina newyorkensis 2681 GCA_000220335.1
46 Sporosarcina psychrophila GCA_001590685.1
46 Sporosarcina ureae GCA_002082015.1
46 Streptococcus cristatus AS 1.3089 GCA_000385925.1
46 Streptococcus mitis B6 GCA_000027165.1
46 Streptococcus parasanguinis ATCC 15912 GCA_000164675.2
46 Streptococcus phocae subsp. salmonis GCA_000772915.1
46 Streptococcus pneumoniae R6 GCA_000007045.1
7d
218 Clostridium butyricum GCA_001456065.2
207 Clostridium saccharobutylicum DSM 13864 GCA_000473995.1
204 Clostridium chromiireducens GCA_002029255.1
204 Clostridium taeniosporum GCA_001735765.1
7e
256 Lactobacillus gasseri ATCC 33323 = JCM 1131 GCA_000014425.1
252 Lactobacillus hominis DSM 23910 = CRBIP 24.179 GCA_000296835.1
204 Lactobacillus acidophilus NCFM GCA_000011985.1
204 Lactobacillus crispatus ST1 GCA_000091765.1
204 Lactobacillus hamsteri DSM 5661 = JCM 6256 GCA_000615445.1
7f
159 Staphylococcus cohnii subsp. cohnii GCA_000972575.1
159 Staphylococcus epidermidis ATCC 12228 GCA_000007645.1
159 Staphylococcus lugdunensis HKU09-01 GCA_000025085.1
159 Staphylococcus saprophyticus subsp. saprophyticus ATCC 15305 GCA_000010125.1
159 Staphylococcus xylosus GCA_000706685.1
153 Staphylococcus capitis subsp. capitis GCA_001028645.1
150 Staphylococcus pettenkoferi GCA_002208805.1
7g
250 Staphylococcus capitis subsp. capitis GCA_001028645.1
250 Staphylococcus epidermidis ATCC 12228 GCA_000007645.1
250 Staphylococcus haemolyticus JCSC1435 GCA_000009865.1
250 Staphylococcus hominis subsp. hominis C80 GCA_000183685.1
250 Staphylococcus lugdunensis HKU09-01 GCA_000025085.1
236 Staphylococcus microti GCA_000934465.1
235 Megasphaera cerevisiae DSM 20462 GCA_001045675.1
235 Staphylococcus aureus subsp. aureus NCTC 8325 GCA_000013425.1
235 Staphylococcus lutrae GCA_002101335.1
235 Staphylococcus simiae CCM 7213 GCA_000235645.2
235 Staphylococcus warneri SG1 GCA_000332735.1
7h
242 Clostridium saccharobutylicum DSM 13864 GCA_000473995.1
231 Clostridium beijerinckii GCA_000833105.2
231 Clostridium puniceum GCA_002006345.1
231 Clostridium saccharoperbutylacetonicum N1-4_28HMT_29 GCA_000340885.1
226 Clostridium butyricum GCA_001456065.2
7i
291 Bacillus anthracis str. Ames GCA_000007845.1
291 Bacillus anthracis str. Sterne GCA_000008165.1
291 Bacillus cereus ATCC 14579 GCA_000007825.1
291 Bacillus mycoides GCA_000832605.1
291 Bacillus pseudomycoides DSM 12442 GCA_000161455.1
291 Bacillus thuringiensis YBT-1518 GCA_000497525.2
291 _5BBacillus thuringiensis_5D serovar konkukian str. 97-27 GCA_000008505.1
237 Bacillus horneckiae GCA_001636335.1
237 Bacillus solani GCA_001420595.1
233 Bacillus coahuilensis m4-4 GCA_000171615.1
233 Bacillus marisflavi GCA_001274775.1
7j
229 Rhodobacter sphaeroides 2.4.1 GCA_000012905.2

208 Pseudorhodobacter psychrotolerans GCA_001294535.1
201 Pseudorhodobacter ferrugineus DSM 5888 GCA_000420745.1
201 Pseudorhodobacter wandonensis GCA_001202035.1
7k
160 Escherichia coli IAI39 GCA_000026345.1
160 Escherichia coli O104_3AH4 str. 2011C-3493 GCA_000299455.1
160 Escherichia coli O157_3AH7 str. Sakai GCA_000008865.1
160 Escherichia coli O83_3AH1 str. NRG 857C GCA_000183345.1
160 Escherichia coli UMN026 GCA_000026325.2
160 Escherichia coli str. K-12 substr. MG1655 GCA_000005845.2
160 Shigella flexneri 2a str. 301 GCA_000006925.2
160 Tumebacillus flagellatus GCA_000714935.1
153 Erwinia iniecta GCA_001267535.1
145 Shigella dysenteriae Sd197 GCA_000012005.1
7l
187 Rhodobacter sphaeroides 2.4.1 GCA_000012905.2
170 Rhodobacter sphaeroides ATCC 17025 GCA_000016405.1
161 Gemmobacter megaterium GCA_900156815.1
7m
162 Deinococcus radiodurans R1 GCA_000008565.1
150 Deinococcus soli Cha et al. 2016 GCA_001007995.1
139 Deinococcus hopiensis KR-140 GCA_900176165.1
7n
209 Clostridium butyricum GCA_001456065.2
208 Clostridium beijerinckii GCA_000833105.2
208 Clostridium chromiireducens GCA_002029255.1
208 Clostridium puniceum GCA_002006345.1
208 Clostridium saccharobutylicum DSM 13864 GCA_000473995.1
208 Clostridium saccharoperbutylacetonicum N1-4_28HMT_29 GCA_000340885.1
192 Clostridium neonatale GCA_001458595.1
7o
93 Staphylococcus capitis subsp. capitis GCA_001028645.1
93 Staphylococcus epidermidis ATCC 12228 GCA_000007645.1
93 Staphylococcus haemolyticus JCSC1435 GCA_000009865.1
82 Staphylococcus cohnii subsp. cohnii GCA_000972575.1
82 Staphylococcus hominis subsp. hominis C80 GCA_000183685.1
82 Staphylococcus simulans GCA_001559115.1
77 Staphylococcus microti GCA_000934465.1
7p
312 Deinococcus radiodurans R1 GCA_000008565.1
196 Deinococcus gobiensis I-0 GCA_000252445.1
194 Deinococcus deserti VCD115 GCA_000020685.1
7q
251 Bacillus anthracis str. Ames GCA_000007845.1
251 Bacillus anthracis str. Sterne GCA_000008165.1
251 Bacillus cereus ATCC 14579 GCA_000007825.1
251 Bacillus mycoides GCA_000832605.1
251 Bacillus pseudomycoides DSM 12442 GCA_000161455.1
251 Bacillus thuringiensis YBT-1518 GCA_000497525.2
251 _5BBacillus thuringiensis_5D serovar konkukian str. 97-27 GCA_000008505.1
205 Bacillus manliponensis GCA_000712595.1
197 Bacillus cytotoxicus NVH 391-98 GCA_000017425.1
7r
166 Clostridium beijerinckii GCA_000833105.2
166 Clostridium puniceum GCA_002006345.1
166 Clostridium saccharoperbutylacetonicum N1-4_28HMT_29 GCA_000340885.1
166 Clostridium taeniosporum GCA_001735765.1
162 Clostridium butyricum GCA_001456065.2
157 Clostridium botulinum B str. Eklund 17B _28NRP_29 GCA_000020165.1
157 Clostridium chromiireducens GCA_002029255.1
157 Clostridium saccharobutylicum DSM 13864 GCA_000473995.1
7s
180 Deinococcus radiodurans R1 GCA_000008565.1
120 Deinococcus gobiensis I-0 GCA_000252445.1
113 Deinococcus deserti VCD115 GCA_000020685.1
7t
151 Lactobacillus gasseri ATCC 33323 = JCM 1131 GCA_000014425.1
151 Lactobacillus hominis DSM 23910 = CRBIP 24.179 GCA_000296835.1
122 Lactobacillus iners DSM 13335 GCA_000160875.1
111 Lactobacillus farraginis DSM 18382 = JCM 14108 GCA_000583655.1
7u
143 Enterococcus faecalis V583 GCA_000007785.1
143 Streptomyces cinnamoneus GCA_001885705.1
133 Enterococcus asini ATCC 700915 GCA_000407365.1
133 Enterococcus canis NBRC 100695 GCA_001544375.1
133 Enterococcus casseliflavus EC20 GCA_000157355.2

133 Enterococcus dispar ATCC 51266 GCA_000406945.1
133 Enterococcus faecium DO GCA_000174395.2
133 Enterococcus gilvus ATCC BAA-350 GCA_000407545.1
133 Enterococcus hermanniensis GCA_001885945.1
133 Enterococcus hirae ATCC 9790 GCA_000271405.2
133 Enterococcus italicus DSM 15952 GCA_000185365.1
133 Enterococcus malodoratus ATCC 43197 GCA_000407185.1
133 Enterococcus massiliensis GCA_001050095.1
133 Enterococcus mundtii QU 25 GCA_000504125.1
133 Enterococcus pallens ATCC BAA-351 GCA_000407485.1
133 Enterococcus pseudoavium NBRC 100491 GCA_001544295.1
133 Enterococcus rivorum GCA_001742285.1
133 Enterococcus saccharolyticus subsp. saccharolyticus ATCC 43076 GCA_000407285.1
133 Melissococcus plutonius S1 GCA_000747585.1
128 Listeria aquatica FSL S10-1188 GCA_000525795.1
128 Listeria floridensis FSL S10-1187 GCA_000525875.1
7v
227 Escherichia coli IAI39 GCA_000026345.1
227 Escherichia coli O104_3AH4 str. 2011C-3493 GCA_000299455.1
227 Escherichia coli O157_3AH7 str. Sakai GCA_000008865.1
227 Escherichia coli O83_3AH1 str. NRG 857C GCA_000183345.1
227 Escherichia coli UMN026 GCA_000026325.2
227 Escherichia coli str. K-12 substr. MG1655 GCA_000005845.2
227 Shigella dysenteriae Sd197 GCA_000012005.1
227 Shigella flexneri 2a str. 301 GCA_000006925.2
227 Tumebacillus flagellatus GCA_000714935.1
216 Erwinia iniecta GCA_001267535.1
200 Cedecea neteri GCA_000757825.1
200 Cronobacter sakazakii GCA_000982825.1
200 Enterobacter cloacae subsp. cloacae ATCC 13047 GCA_000025565.1
200 Enterobacter hormaechei subsp. steigerwaltii GCA_001729725.1
200 Enterobacter kobei GCA_900185885.1
200 Erwinia persicina NBRC 102418 GCA_001571305.1
200 Erwinia toletana DAPP-PG 735 GCA_000336255.1
200 Klebsiella oxytoca GCA_001022195.1
200 Kosakonia cowanii GCA_001975225.1
200 Pseudescherichia vulneris NBRC 102420 GCA_000759795.1
200 Rosenbergiella nectarea GCA_900111105.1
200 Salmonella enterica subsp. enterica serovar Typhi str. CT18 GCA_000195995.1
200 Salmonella enterica subsp. enterica serovar Typhimurium str. LT2 GCA_000006945.2
7w
224 Deinococcus radiodurans R1 GCA_000008565.1
141 Deinococcus gobiensis I-0 GCA_000252445.1
118 Deinococcus puniceus GCA_001644565.1
7x
222 Deinococcus radiodurans R1 GCA_000008565.1
135 Deinococcus gobiensis I-0 GCA_000252445.1
120 Deinococcus deserti VCD115 GCA_000020685.1
120 Deinococcus geothermalis DSM 11300 GCA_000196275.1
120 Deinococcus hopiensis KR-140 GCA_900176165.1
7y
202 Enterococcus faecalis V583 GCA_000007785.1
202 Streptomyces cinnamoneus GCA_001885705.1
178 Enterococcus aquimarinus GCA_001885765.1
178 Enterococcus canis NBRC 100695 GCA_001544375.1
178 Enterococcus casseliflavus EC20 GCA_000157355.2
178 Enterococcus dispar ATCC 51266 GCA_000406945.1
178 Enterococcus faecium DO GCA_000174395.2
178 Enterococcus gilvus ATCC BAA-350 GCA_000407545.1
178 Enterococcus haemoperoxidus ATCC BAA-382 GCA_000407165.1
178 Enterococcus hermanniensis GCA_001885945.1
178 Enterococcus hirae ATCC 9790 GCA_000271405.2
178 Enterococcus malodoratus ATCC 43197 GCA_000407185.1
178 Enterococcus massiliensis GCA_001050095.1
178 Enterococcus mundtii QU 25 GCA_000504125.1
178 Enterococcus pallens ATCC BAA-351 GCA_000407485.1
178 Enterococcus phoeniculicola ATCC BAA-412 GCA_000407505.1
178 Enterococcus pseudoavium NBRC 100491 GCA_001544295.1
178 Enterococcus rivorum GCA_001742285.1
178 Enterococcus saccharolyticus subsp. saccharolyticus ATCC 43076 GCA_000407285.1
178 Enterococcus thailandicus GCA_001652875.1
171 Enterococcus asini ATCC 700915 GCA_000407365.1
7z
202 Escherichia coli IAI39 GCA_000026345.1
202 Escherichia coli O104_3AH4 str. 2011C-3493 GCA_000299455.1
202 Escherichia coli O157_3AH7 str. Sakai GCA_000008865.1

202 Escherichia coli O83_3AH1 str. NRG 857C GCA_000183345.1
202 Escherichia coli UMN026 GCA_000026325.2
202 Escherichia coli str. K-12 substr. MG1655 GCA_000005845.2
202 Shigella dysenteriae Sd197 GCA_000012005.1
202 Shigella flexneri 2a str. 301 GCA_000006925.2
202 Tumebacillus flagellatus GCA_000714935.1
195 Erwinia iniecta GCA_001267535.1
190 Pantoea agglomerans GCA_001709315.1
7A
237 Escherichia coli IAI39 GCA_000026345.1
237 Escherichia coli O104_3AH4 str. 2011C-3493 GCA_000299455.1
237 Escherichia coli O157_3AH7 str. Sakai GCA_000008865.1
237 Escherichia coli O83_3AH1 str. NRG 857C GCA_000183345.1
237 Escherichia coli UMN026 GCA_000026325.2
237 Escherichia coli str. K-12 substr. MG1655 GCA_000005845.2
237 Shigella flexneri 2a str. 301 GCA_000006925.2
237 Tumebacillus flagellatus GCA_000714935.1
233 Erwinia iniecta GCA_001267535.1
233 Erwinia toletana DAPP-PG 735 GCA_000336255.1
231 Pantoea ananatis LMG 20103 GCA_000025405.2
231 Shimwellia blattae DSM 4481 = NBRC 105725 GCA_000262305.1
7B
250 Escherichia coli IAI39 GCA_000026345.1
250 Escherichia coli O104_3AH4 str. 2011C-3493 GCA_000299455.1
250 Escherichia coli O157_3AH7 str. Sakai GCA_000008865.1
250 Escherichia coli O83_3AH1 str. NRG 857C GCA_000183345.1
250 Escherichia coli UMN026 GCA_000026325.2
250 Escherichia coli str. K-12 substr. MG1655 GCA_000005845.2
250 Shigella dysenteriae Sd197 GCA_000012005.1
250 Shigella flexneri 2a str. 301 GCA_000006925.2
250 Tumebacillus flagellatus GCA_000714935.1
213 Erwinia iniecta GCA_001267535.1
201 Enterobacter hormaechei subsp. steigerwaltii GCA_001729725.1
201 Erwinia toletana DAPP-PG 735 GCA_000336255.1
201 Klebsiella oxytoca GCA_001022195.1
201 Kosakonia cowanii GCA_001975225.1
201 Kosakonia sacchari SP1 GCA_000300455.4
201 Pluralibacter gergoviae GCA_000757785.1
201 Pseudescherichia vulneris NBRC 102420 GCA_000759795.1
7C
303 Deinococcus radiodurans R1 GCA_000008565.1
206 Deinococcus puniceus GCA_001644565.1
194 Deinococcus marmoris DSM 12784 GCA_000701405.1
194 Deinococcus swuensis GCA_000800395.1
7D
142 Deinococcus radiodurans R1 GCA_000008565.1
106 Deinococcus gobiensis I-0 GCA_000252445.1
82 Deinococcus marmoris DSM 12784 GCA_000701405.1
82 Deinococcus swuensis GCA_000800395.1
7E
191 Bacillus anthracis str. Ames GCA_000007845.1
191 Bacillus anthracis str. Sterne GCA_000008165.1
191 Bacillus cereus ATCC 14579 GCA_000007825.1
191 Bacillus mycoides GCA_000832605.1
191 Bacillus pseudomycoides DSM 12442 GCA_000161455.1
191 Bacillus thuringiensis YBT-1518 GCA_000497525.2
191 _5BBacillus thuringiensis_5D serovar konkukian str. 97-27 GCA_000008505.1
166 Bacillus manliponensis GCA_000712595.1
158 Bacillus halmapalus GCA_002019665.1
7F
178 Bacillus thuringiensis YBT-1518 GCA_000497525.2
172 Bacillus anthracis str. Ames GCA_000007845.1
172 Bacillus anthracis str. Sterne GCA_000008165.1
172 Bacillus cereus ATCC 14579 GCA_000007825.1
172 Bacillus mycoides GCA_000832605.1
172 Bacillus pseudomycoides DSM 12442 GCA_000161455.1
172 _5BBacillus thuringiensis_5D serovar konkukian str. 97-27 GCA_000008505.1
140 Bacillus hemicellulosilyticus JCM 9152 GCA_000513115.1
7G
87 Erwinia iniecta GCA_001267535.1
87 Erwinia toletana DAPP-PG 735 GCA_000336255.1
87 Escherichia coli IAI39 GCA_000026345.1
87 Escherichia coli O104_3AH4 str. 2011C-3493 GCA_000299455.1
87 Escherichia coli O157_3AH7 str. Sakai GCA_000008865.1
87 Escherichia coli O83_3AH1 str. NRG 857C GCA_000183345.1
87 Escherichia coli UMN026 GCA_000026325.2

87 Escherichia coli str. K-12 substr. MG1655 GCA_000005845.2
87 Pantoea agglomerans GCA_001709315.1
87 Pantoea ananatis LMG 20103 GCA_000025405.2
87 Shigella dysenteriae Sd197 GCA_000012005.1
87 Shigella flexneri 2a str. 301 GCA_000006925.2
87 Shimwellia blattae DSM 4481 = NBRC 105725 GCA_000262305.1
87 Tumebacillus flagellatus GCA_000714935.1
83 Cronobacter sakazakii GCA_000982825.1
83 Erwinia gerundensis GCA_001517405.1
83 Erwinia oleae GCA_000770305.1
83 Erwinia teleogrylli GCA_001484765.1
83 Pantoea dispersa EGD-AAK13 GCA_000465555.2
83 Plautia stali symbiont GCA_000180175.2
83 Plesiomonas shigelloides GCA_900087055.1
83 Proteus mirabilis HI4320 GCA_000069965.1
83 Sodalis praecaptivus GCA_000517425.1
83 Streptomyces gilvigriseus GCA_001879105.1
80 Buchnera aphidicola str. Bp _28Baizongia pistaciae_29 GCA_000007725.1
80 Buchnera aphidicola str. Sg _28Schizaphis graminum_29 GCA_000007365.1
7H
196 Escherichia coli IAI39 GCA_000026345.1
196 Escherichia coli O104_3AH4 str. 2011C-3493 GCA_000299455.1
196 Escherichia coli O157_3AH7 str. Sakai GCA_000008865.1
196 Escherichia coli O83_3AH1 str. NRG 857C GCA_000183345.1
196 Escherichia coli UMN026 GCA_000026325.2
196 Escherichia coli str. K-12 substr. MG1655 GCA_000005845.2
196 Shigella dysenteriae Sd197 GCA_000012005.1
196 Shigella flexneri 2a str. 301 GCA_000006925.2
196 Tumebacillus flagellatus GCA_000714935.1
160 Erwinia iniecta GCA_001267535.1
140 Cedecea neteri GCA_000757825.1
140 Rosenbergiella nectarea GCA_900111105.1
7I
164 Cronobacter sakazakii GCA_000982825.1
164 Erwinia iniecta GCA_001267535.1
164 Erwinia toletana DAPP-PG 735 GCA_000336255.1
164 Escherichia coli IAI39 GCA_000026345.1
164 Escherichia coli O104_3AH4 str. 2011C-3493 GCA_000299455.1
164 Escherichia coli O157_3AH7 str. Sakai GCA_000008865.1
164 Escherichia coli O83_3AH1 str. NRG 857C GCA_000183345.1
164 Escherichia coli UMN026 GCA_000026325.2
164 Escherichia coli str. K-12 substr. MG1655 GCA_000005845.2
164 Pantoea ananatis LMG 20103 GCA_000025405.2
164 Pantoea dispersa EGD-AAK13 GCA_000465555.2
164 Shigella flexneri 2a str. 301 GCA_000006925.2
164 Shimwellia blattae DSM 4481 = NBRC 105725 GCA_000262305.1
164 Tumebacillus flagellatus GCA_000714935.1
156 Kosakonia sacchari SP1 GCA_000300455.4
156 Plautia stali symbiont GCA_000180175.2
155 Erwinia gerundensis GCA_001517405.1
155 Pantoea agglomerans GCA_001709315.1
155 Shigella dysenteriae Sd197 GCA_000012005.1
7J
261 Bacillus mycoides GCA_000832605.1
260 Bacillus anthracis str. Ames GCA_000007845.1
260 Bacillus anthracis str. Sterne GCA_000008165.1
260 Bacillus cereus ATCC 14579 GCA_000007825.1
260 Bacillus pseudomycoides DSM 12442 GCA_000161455.1
260 Bacillus thuringiensis YBT-1518 GCA_000497525.2
260 _5BBacillus thuringiensis_5D serovar konkukian str. 97-27 GCA_000008505.1
198 Bacillus horneckiae GCA_001636335.1
198 Bacillus okuhidensis GCA_001274915.1
198 Bacillus solani GCA_001420595.1
7K
237 Clostridium beijerinckii GCA_000833105.2
237 Clostridium puniceum GCA_002006345.1
237 Clostridium saccharoperbutylacetonicum N1-4_28HMT_29 GCA_000340885.1
230 Clostridium taeniosporum GCA_001735765.1
228 Clostridium saccharobutylicum DSM 13864 GCA_000473995.1
7L
115 Bifidobacterium lemurum GCA_001895165.1
114 Bifidobacterium dentium JCM 1195 = DSM 20436 GCA_001042595.1
114 Bifidobacterium thermophilum GCA_000741495.1
114 Bifidobacterium tsurumiense GCA_000741765.1
112 Bifidobacterium gallicum DSM 20093 = LMG 11596 GCA_000741205.1
7M

329 Streptococcus mutans UA159 GCA_000007465.2
232 Streptococcus ratti FA-1 = DSM 20564 GCA_000286075.1
222 Streptococcus gordonii str. Challis substr. CH1 GCA_000017005.1
222 Streptococcus salivarius GCA_000785515.1
222 Streptococcus thermophilus JIM 8232 GCA_000253395.1
7N
162 Erwinia iniecta GCA_001267535.1
162 Escherichia coli IAI39 GCA_000026345.1
162 Escherichia coli O104_3AH4 str. 2011C-3493 GCA_000299455.1
162 Escherichia coli O157_3AH7 str. Sakai GCA_000008865.1
162 Escherichia coli O83_3AH1 str. NRG 857C GCA_000183345.1
162 Escherichia coli UMN026 GCA_000026325.2
162 Escherichia coli str. K-12 substr. MG1655 GCA_000005845.2
162 Shigella flexneri 2a str. 301 GCA_000006925.2
162 Tumebacillus flagellatus GCA_000714935.1
150 Serratia marcescens subsp. marcescens Db11 GCA_000513215.1
150 Serratia rubidaea GCA_001572725.1
147 Shigella dysenteriae Sd197 GCA_000012005.1
7O
286 Bacillus thuringiensis YBT-1518 GCA_000497525.2
269 Bacillus anthracis str. Ames GCA_000007845.1
269 Bacillus anthracis str. Sterne GCA_000008165.1
269 Bacillus cereus ATCC 14579 GCA_000007825.1
269 Bacillus mycoides GCA_000832605.1
269 Bacillus pseudomycoides DSM 12442 GCA_000161455.1
269 _5BBacillus thuringiensis_5D serovar konkukian str. 97-27 GCA_000008505.1
211 Massilibacterium senegalense GCA_001375675.1
7P
144 Staphylococcus pettenkoferi GCA_002208805.1
133 Megasphaera cerevisiae DSM 20462 GCA_001045675.1
133 Staphylococcus epidermidis ATCC 12228 GCA_000007645.1
133 Staphylococcus lugdunensis HKU09-01 GCA_000025085.1
133 Staphylococcus pseudintermedius HKU10-03 GCA_000185885.1
133 Staphylococcus warneri SG1 GCA_000332735.1
132 Staphylococcus condimenti GCA_001618885.1
132 Staphylococcus simulans GCA_001559115.1
7Q
128 Hathewaya proteolytica DSM 3090 GCA_900142225.1
126 Clostridium beijerinckii GCA_000833105.2
126 Clostridium saccharobutylicum DSM 13864 GCA_000473995.1
126 Clostridium saccharoperbutylacetonicum N1-4_28HMT_29 GCA_000340885.1
120 Clostridium puniceum GCA_002006345.1
7R
174 Rhodobacter sphaeroides 2.4.1 GCA_000012905.2
149 Pseudorhodobacter ferrugineus DSM 5888 GCA_000420745.1
149 Pseudorhodobacter wandonensis GCA_001202035.1
149 Thioclava dalianensis GCA_000715505.1
149 Thioclava indica GCA_000714545.1
143 Gemmobacter megaterium GCA_900156815.1
143 Paracoccus yeei GCA_002073635.1
143 Pseudorhodobacter psychrotolerans GCA_001294535.1
7S
188 Rhodobacter sphaeroides 2.4.1 GCA_000012905.2
188 Rhodobacter sphaeroides ATCC 17025 GCA_000016405.1
178 Gemmobacter megaterium GCA_900156815.1
164 Pseudorhodobacter psychrotolerans GCA_001294535.1
7T
127 Staphylococcus capitis subsp. capitis GCA_001028645.1
127 Staphylococcus cohnii subsp. cohnii GCA_000972575.1
127 Staphylococcus epidermidis ATCC 12228 GCA_000007645.1
127 Staphylococcus lugdunensis HKU09-01 GCA_000025085.1
127 Staphylococcus saprophyticus subsp. saprophyticus ATCC 15305 GCA_000010125.1
127 Staphylococcus xylosus GCA_000706685.1
126 Staphylococcus pseudintermedius HKU10-03 GCA_000185885.1
122 Staphylococcus microti GCA_000934465.1
122 Staphylococcus saprophyticus GCA_001074355.1
7U
271 Streptococcus mutans UA159 GCA_000007465.2
204 Streptococcus gallolyticus subsp. gallolyticus DSM 16831 GCA_002000985.1
203 Streptococcus equinus GCA_000964315.1
203 Streptococcus ratti FA-1 = DSM 20564 GCA_000286075.1
7V
214 Rhodobacter sphaeroides 2.4.1 GCA_000012905.2
208 Rhodobacter sphaeroides ATCC 17025 GCA_000016405.1
184 Pseudorhodobacter ferrugineus DSM 5888 GCA_000420745.1
184 Pseudorhodobacter psychrotolerans GCA_001294535.1

184 Pseudorhodobacter wandonensis GCA_001202035.1
7W
172 Clostridium beijerinckii GCA_000833105.2
172 Clostridium puniceum GCA_002006345.1
172 Clostridium saccharoperbutylacetonicum N1-4_28HMT_29 GCA_000340885.1
168 Clostridium saccharobutylicum DSM 13864 GCA_000473995.1
159 Clostridium butyricum GCA_001456065.2
7X
203 Enterococcus faecalis V583 GCA_000007785.1
203 Streptomyces cinnamoneus GCA_001885705.1
183 Melissococcus plutonius S1 GCA_000747585.1
181 Enterococcus asini ATCC 700915 GCA_000407365.1
181 Enterococcus canis NBRC 100695 GCA_001544375.1
181 Enterococcus casseliflavus EC20 GCA_000157355.2
181 Enterococcus dispar ATCC 51266 GCA_000406945.1
181 Enterococcus faecium DO GCA_000174395.2
181 Enterococcus gilvus ATCC BAA-350 GCA_000407545.1
181 Enterococcus hermanniensis GCA_001885945.1
181 Enterococcus hirae ATCC 9790 GCA_000271405.2
181 Enterococcus italicus DSM 15952 GCA_000185365.1
181 Enterococcus malodoratus ATCC 43197 GCA_000407185.1
181 Enterococcus mundtii QU 25 GCA_000504125.1
181 Enterococcus pallens ATCC BAA-351 GCA_000407485.1
181 Enterococcus pseudoavium NBRC 100491 GCA_001544295.1
181 Enterococcus rivorum GCA_001742285.1
181 Enterococcus saccharolyticus subsp. saccharolyticus ATCC 43076 GCA_000407285.1
7Y
244 Escherichia coli IAI39 GCA_000026345.1
244 Escherichia coli O104_3AH4 str. 2011C-3493 GCA_000299455.1
244 Escherichia coli O157_3AH7 str. Sakai GCA_000008865.1
244 Escherichia coli O83_3AH1 str. NRG 857C GCA_000183345.1
244 Escherichia coli UMN026 GCA_000026325.2
244 Escherichia coli str. K-12 substr. MG1655 GCA_000005845.2
244 Shigella flexneri 2a str. 301 GCA_000006925.2
244 Tumebacillus flagellatus GCA_000714935.1
235 Shigella dysenteriae Sd197 GCA_000012005.1
208 Erwinia iniecta GCA_001267535.1
7Z
269 Clostridium beijerinckii GCA_000833105.2
269 Clostridium saccharoperbutylacetonicum N1-4_28HMT_29 GCA_000340885.1
262 Clostridium puniceum GCA_002006345.1
254 Clostridium saccharobutylicum DSM 13864 GCA_000473995.1
80
93 Rhodobacter sphaeroides ATCC 17025 GCA_000016405.1
79 Rhodobacter sphaeroides 2.4.1 GCA_000012905.2
65 Rhodobacter capsulatus SB 1003 GCA_000021865.1
81
207 Clostridium beijerinckii GCA_000833105.2
207 Clostridium puniceum GCA_002006345.1
207 Clostridium saccharobutylicum DSM 13864 GCA_000473995.1
207 Clostridium saccharoperbutylacetonicum N1-4_28HMT_29 GCA_000340885.1
205 Clostridium butyricum GCA_001456065.2
199 Clostridium chromiireducens GCA_002029255.1
82
292 Clostridium beijerinckii GCA_000833105.2
292 Clostridium saccharoperbutylacetonicum N1-4_28HMT_29 GCA_000340885.1
288 Clostridium puniceum GCA_002006345.1
284 Clostridium saccharobutylicum DSM 13864 GCA_000473995.1
83
163 Pseudorhodobacter psychrotolerans GCA_001294535.1
163 Rhodobacter sphaeroides 2.4.1 GCA_000012905.2
163 Rhodobacter sphaeroides ATCC 17025 GCA_000016405.1
155 Pseudorhodobacter ferrugineus DSM 5888 GCA_000420745.1
155 Pseudorhodobacter wandonensis GCA_001202035.1
150 Pseudorhodobacter aquimaris GCA_001202025.1
84
237 Clostridium puniceum GCA_002006345.1
233 Clostridium beijerinckii GCA_000833105.2
233 Clostridium saccharobutylicum DSM 13864 GCA_000473995.1
233 Clostridium saccharoperbutylacetonicum N1-4_28HMT_29 GCA_000340885.1
219 Clostridium chromiireducens GCA_002029255.1
85
144 Staphylococcus capitis subsp. capitis GCA_001028645.1
144 Staphylococcus epidermidis ATCC 12228 GCA_000007645.1
144 Staphylococcus haemolyticus JCSC1435 GCA_000009865.1
137 Staphylococcus pettenkoferi GCA_002208805.1

128 Staphylococcus aureus subsp. aureus NCTC 8325 GCA_000013425.1
128 Staphylococcus hominis subsp. hominis C80 GCA_000183685.1
128 Staphylococcus lugdunensis HKU09-01 GCA_000025085.1
128 Staphylococcus simiae CCM 7213 GCA_000235645.2
86
302 Escherichia coli IAI39 GCA_000026345.1
302 Escherichia coli O104_3AH4 str. 2011C-3493 GCA_000299455.1
302 Escherichia coli O157_3AH7 str. Sakai GCA_000008865.1
302 Escherichia coli O83_3AH1 str. NRG 857C GCA_000183345.1
302 Escherichia coli UMN026 GCA_000026325.2
302 Escherichia coli str. K-12 substr. MG1655 GCA_000005845.2
302 Shigella dysenteriae Sd197 GCA_000012005.1
302 Shigella flexneri 2a str. 301 GCA_000006925.2
302 Tumebacillus flagellatus GCA_000714935.1
285 Erwinia iniecta GCA_001267535.1
266 Rosenbergiella nectarea GCA_900111105.1
87
248 Deinococcus radiodurans R1 GCA_000008565.1
185 Deinococcus gobiensis I-0 GCA_000252445.1
184 Deinococcus frigens DSM 12807 GCA_000701425.1
184 Deinococcus marmoris DSM 12784 GCA_000701405.1
184 Deinococcus swuensis GCA_000800395.1
88
278 Bacillus anthracis str. Ames GCA_000007845.1
278 Bacillus anthracis str. Sterne GCA_000008165.1
278 Bacillus cereus ATCC 14579 GCA_000007825.1
278 Bacillus mycoides GCA_000832605.1
278 Bacillus pseudomycoides DSM 12442 GCA_000161455.1
278 Bacillus thuringiensis YBT-1518 GCA_000497525.2
278 _5BBacillus thuringiensis_5D serovar konkukian str. 97-27 GCA_000008505.1
216 Bacillus manliponensis GCA_000712595.1
206 Bacillus marisflavi GCA_001274775.1
89
188 Rhodobacter sphaeroides 2.4.1 GCA_000012905.2
179 Rhodobacter sphaeroides ATCC 17025 GCA_000016405.1
138 Pseudorhodobacter ferrugineus DSM 5888 GCA_000420745.1
138 Pseudorhodobacter psychrotolerans GCA_001294535.1
138 Pseudorhodobacter wandonensis GCA_001202035.1
8a
248 Escherichia coli IAI39 GCA_000026345.1
248 Escherichia coli O104_3AH4 str. 2011C-3493 GCA_000299455.1
248 Escherichia coli O157_3AH7 str. Sakai GCA_000008865.1
248 Escherichia coli O83_3AH1 str. NRG 857C GCA_000183345.1
248 Escherichia coli UMN026 GCA_000026325.2
248 Escherichia coli str. K-12 substr. MG1655 GCA_000005845.2
248 Shigella flexneri 2a str. 301 GCA_000006925.2
248 Tumebacillus flagellatus GCA_000714935.1
239 Shigella dysenteriae Sd197 GCA_000012005.1
238 Erwinia iniecta GCA_001267535.1
8b
194 Lactobacillus gasseri ATCC 33323 = JCM 1131 GCA_000014425.1
186 Lactobacillus hominis DSM 23910 = CRBIP 24.179 GCA_000296835.1
138 Lactobacillus amylovorus GCA_000191545.1
138 Lactobacillus crispatus ST1 GCA_000091765.1
138 Lactobacillus delbrueckii subsp. bulgaricus ATCC 11842 = JCM 1002 GCA_000056065.1
138 Lactobacillus kalixensis DSM 16043 GCA_001434335.1
8c
270 Deinococcus radiodurans R1 GCA_000008565.1
162 Deinococcus puniceus GCA_001644565.1
155 Deinococcus gobiensis I-0 GCA_000252445.1
8d
157 Bacillus thuringiensis YBT-1518 GCA_000497525.2
127 Bacillus psychrosaccharolyticus ATCC 23296 GCA_000305495.2
125 Bacillus anthracis str. Ames GCA_000007845.1
125 Bacillus anthracis str. Sterne GCA_000008165.1
125 Bacillus cereus ATCC 14579 GCA_000007825.1
125 Bacillus mycoides GCA_000832605.1
125 Bacillus pseudomycoides DSM 12442 GCA_000161455.1
125 _5BBacillus thuringiensis_5D serovar konkukian str. 97-27 GCA_000008505.1
8e
211 Deinococcus radiodurans R1 GCA_000008565.1
152 Deinococcus deserti VCD115 GCA_000020685.1
143 Deinococcus gobiensis I-0 GCA_000252445.1
8f
158 Streptococcus mutans UA159 GCA_000007465.2
117 Streptococcus equinus GCA_000964315.1

117 Streptococcus gallolyticus subsp. gallolyticus DSM 16831 GCA_002000985.1
114 Streptococcus cristatus AS 1.3089 GCA_000385925.1
114 Streptococcus gordonii str. Challis substr. CH1 GCA_000017005.1
114 Streptococcus mitis B6 GCA_000027165.1
114 Streptococcus pneumoniae R6 GCA_000007045.1
8g
261 Bacillus anthracis str. Ames GCA_000007845.1
261 Bacillus anthracis str. Sterne GCA_000008165.1
261 Bacillus cereus ATCC 14579 GCA_000007825.1
261 Bacillus mycoides GCA_000832605.1
261 Bacillus pseudomycoides DSM 12442 GCA_000161455.1
261 Bacillus thuringiensis YBT-1518 GCA_000497525.2
261 _5BBacillus thuringiensis_5D serovar konkukian str. 97-27 GCA_000008505.1
202 Bacillus marisflavi GCA_001274775.1
200 Bacillus manliponensis GCA_000712595.1
8h
94 Bacillus cereus ATCC 14579 GCA_000007825.1
94 Bacillus thuringiensis YBT-1518 GCA_000497525.2
79 Bacillus anthracis str. Ames GCA_000007845.1
79 Bacillus anthracis str. Sterne GCA_000008165.1
79 Bacillus mycoides GCA_000832605.1
79 Bacillus pseudomycoides DSM 12442 GCA_000161455.1
79 _5BBacillus thuringiensis_5D serovar konkukian str. 97-27 GCA_000008505.1
68 Bacillus manliponensis GCA_000712595.1
68 Paenirhodobacter enshiensis GCA_000740785.1
8i
148 Rhodobacter sphaeroides 2.4.1 GCA_000012905.2
118 Rhodobacter sphaeroides ATCC 17025 GCA_000016405.1
110 Defluviimonas alba GCA_001620265.1
8j
220 Clostridium saccharobutylicum DSM 13864 GCA_000473995.1
216 Clostridium chromiireducens GCA_002029255.1
209 Clostridium butyricum GCA_001456065.2
8k
226 Enterococcus faecalis V583 GCA_000007785.1
226 Streptomyces cinnamoneus GCA_001885705.1
203 Enterococcus dispar ATCC 51266 GCA_000406945.1
203 Enterococcus hirae ATCC 9790 GCA_000271405.2
200 Enterococcus asini ATCC 700915 GCA_000407365.1
200 Enterococcus canis NBRC 100695 GCA_001544375.1
200 Enterococcus casseliflavus EC20 GCA_000157355.2
200 Enterococcus faecium DO GCA_000174395.2
200 Enterococcus haemoperoxidus ATCC BAA-382 GCA_000407165.1
200 Enterococcus massiliensis GCA_001050095.1
200 Enterococcus mundtii QU 25 GCA_000504125.1
200 Enterococcus phoeniculicola ATCC BAA-412 GCA_000407505.1
200 Enterococcus rivorum GCA_001742285.1
200 Enterococcus saccharolyticus subsp. saccharolyticus ATCC 43076 GCA_000407285.1
200 Enterococcus thailandicus GCA_001652875.1
8l
269 Clostridium beijerinckii GCA_000833105.2
269 Clostridium puniceum GCA_002006345.1
269 Clostridium saccharoperbutylacetonicum N1-4_28HMT_29 GCA_000340885.1
261 Clostridium saccharobutylicum DSM 13864 GCA_000473995.1
236 Clostridium chromiireducens GCA_002029255.1
8m
266 Bacillus anthracis str. Ames GCA_000007845.1
266 Bacillus anthracis str. Sterne GCA_000008165.1
266 Bacillus cereus ATCC 14579 GCA_000007825.1
266 Bacillus mycoides GCA_000832605.1
266 Bacillus pseudomycoides DSM 12442 GCA_000161455.1
266 Bacillus thuringiensis YBT-1518 GCA_000497525.2
266 _5BBacillus thuringiensis_5D serovar konkukian str. 97-27 GCA_000008505.1
251 Bacillus cytotoxicus NVH 391-98 GCA_000017425.1
247 Bacillus manliponensis GCA_000712595.1
8n
302 Deinococcus radiodurans R1 GCA_000008565.1
190 Deinococcus geothermalis DSM 11300 GCA_000196275.1
190 Deinococcus gobiensis I-0 GCA_000252445.1
186 Deinococcus proteolyticus MRP GCA_000190555.1
8o
238 Escherichia coli IAI39 GCA_000026345.1
238 Escherichia coli O104_3AH4 str. 2011C-3493 GCA_000299455.1
238 Escherichia coli O157_3AH7 str. Sakai GCA_000008865.1
238 Escherichia coli UMN026 GCA_000026325.2
238 Escherichia coli str. K-12 substr. MG1655 GCA_000005845.2

238 Shigella dysenteriae Sd197 GCA_000012005.1
238 Shigella flexneri 2a str. 301 GCA_000006925.2
238 Tumebacillus flagellatus GCA_000714935.1
237 Escherichia coli O83_3AH1 str. NRG 857C GCA_000183345.1
192 Erwinia iniecta GCA_001267535.1
192 Rosenbergiella nectarea GCA_900111105.1
8p
243 Clostridium saccharobutylicum DSM 13864 GCA_000473995.1
241 Clostridium beijerinckii GCA_000833105.2
241 Clostridium saccharoperbutylacetonicum N1-4_28HMT_29 GCA_000340885.1
226 Clostridium puniceum GCA_002006345.1
8q
253 Deinococcus radiodurans R1 GCA_000008565.1
171 Deinococcus deserti VCD115 GCA_000020685.1
161 Deinococcus puniceus GCA_001644565.1
8r
245 Bifidobacterium adolescentis ATCC 15703 GCA_000010425.1
243 Bifidobacterium angulatum DSM 20098 = JCM 7096 GCA_001025155.1
236 Bifidobacterium callitrichos DSM 23973 GCA_000741175.1
8s
194 Deinococcus radiodurans R1 GCA_000008565.1
126 Deinococcus puniceus GCA_001644565.1
125 Deinococcus gobiensis I-0 GCA_000252445.1
8t
212 Rhodobacter sphaeroides 2.4.1 GCA_000012905.2
176 Gemmobacter aquatilis GCA_900110025.1
176 Rhodobacter capsulatus SB 1003 GCA_000021865.1
175 Haematobacter massiliensis GCA_000740795.1
8u
161 Streptococcus mutans UA159 GCA_000007465.2
123 Streptococcus equinus GCA_000964315.1
123 Streptococcus gallolyticus subsp. gallolyticus DSM 16831 GCA_002000985.1
123 Streptococcus orisratti DSM 15617 GCA_000380105.1
123 Streptococcus ratti FA-1 = DSM 20564 GCA_000286075.1
122 Streptococcus macacae NCTC 11558 GCA_000187995.3
8v
297 Escherichia coli IAI39 GCA_000026345.1
297 Escherichia coli O104_3AH4 str. 2011C-3493 GCA_000299455.1
297 Escherichia coli O157_3AH7 str. Sakai GCA_000008865.1
297 Escherichia coli O83_3AH1 str. NRG 857C GCA_000183345.1
297 Escherichia coli UMN026 GCA_000026325.2
297 Escherichia coli str. K-12 substr. MG1655 GCA_000005845.2
297 Shigella flexneri 2a str. 301 GCA_000006925.2
297 Tumebacillus flagellatus GCA_000714935.1
293 Erwinia iniecta GCA_001267535.1
288 Shigella dysenteriae Sd197 GCA_000012005.1
8w
163 Staphylococcus arlettae CVD059 GCA_000295715.1
163 Staphylococcus saprophyticus GCA_001074355.1
160 Bacillus anthracis str. Ames GCA_000007845.1
160 Bacillus anthracis str. Sterne GCA_000008165.1
160 Bacillus cereus ATCC 14579 GCA_000007825.1
160 Bacillus mycoides GCA_000832605.1
160 Bacillus pseudomycoides DSM 12442 GCA_000161455.1
160 Bacillus thuringiensis YBT-1518 GCA_000497525.2
160 _5BBacillus thuringiensis_5D serovar konkukian str. 97-27 GCA_000008505.1
159 Staphylococcus lutrae GCA_002101335.1
8x
256 Bifidobacterium adolescentis ATCC 15703 GCA_000010425.1
241 Bifidobacterium callitrichos DSM 23973 GCA_000741175.1
232 Bifidobacterium coryneforme GCA_000737865.1
8y
198 Bifidobacterium adolescentis ATCC 15703 GCA_000010425.1
184 Bifidobacterium angulatum DSM 20098 = JCM 7096 GCA_001025155.1
184 Bifidobacterium dentium JCM 1195 = DSM 20436 GCA_001042595.1
183 Bifidobacterium tsurumiense GCA_000741765.1
8z
219 Staphylococcus capitis subsp. capitis GCA_001028645.1
219 Staphylococcus epidermidis ATCC 12228 GCA_000007645.1
219 Staphylococcus pettenkoferi GCA_002208805.1
208 Staphylococcus cohnii subsp. cohnii GCA_000972575.1
208 Staphylococcus lugdunensis HKU09-01 GCA_000025085.1
208 Staphylococcus saprophyticus subsp. saprophyticus ATCC 15305 GCA_000010125.1
203 Staphylococcus haemolyticus JCSC1435 GCA_000009865.1
8A
280 Enterococcus faecalis V583 GCA_000007785.1

280 Streptomyces cinnamoneus GCA_001885705.1
263 Melissococcus plutonius S1 GCA_000747585.1
262 Enterococcus asini ATCC 700915 GCA_000407365.1
262 Enterococcus canis NBRC 100695 GCA_001544375.1
262 Enterococcus dispar ATCC 51266 GCA_000406945.1
262 Enterococcus faecium DO GCA_000174395.2
262 Enterococcus gilvus ATCC BAA-350 GCA_000407545.1
262 Enterococcus haemoperoxidus ATCC BAA-382 GCA_000407165.1
262 Enterococcus hermanniensis GCA_001885945.1
262 Enterococcus hirae ATCC 9790 GCA_000271405.2
262 Enterococcus italicus DSM 15952 GCA_000185365.1
262 Enterococcus malodoratus ATCC 43197 GCA_000407185.1
262 Enterococcus mundtii QU 25 GCA_000504125.1
262 Enterococcus pallens ATCC BAA-351 GCA_000407485.1
262 Enterococcus phoeniculicola ATCC BAA-412 GCA_000407505.1
262 Enterococcus pseudoavium NBRC 100491 GCA_001544295.1
262 Enterococcus rivorum GCA_001742285.1
262 Enterococcus sulfureus ATCC 49903 GCA_000407605.1
262 Enterococcus thailandicus GCA_001652875.1
8B
220 Clostridium beijerinckii GCA_000833105.2
220 Clostridium puniceum GCA_002006345.1
220 Clostridium saccharobutylicum DSM 13864 GCA_000473995.1
220 Clostridium saccharoperbutylacetonicum N1-4_28HMT_29 GCA_000340885.1
209 Clostridium chromiireducens GCA_002029255.1
194 Clostridium butyricum GCA_001456065.2
8C
227 Bifidobacterium adolescentis ATCC 15703 GCA_000010425.1
215 Bifidobacterium callitrichos DSM 23973 GCA_000741175.1
212 Bifidobacterium coryneforme GCA_000737865.1
8D
66 Lactobacillus gasseri ATCC 33323 = JCM 1131 GCA_000014425.1
66 Lactobacillus hominis DSM 23910 = CRBIP 24.179 GCA_000296835.1
59 Lactobacillus acidophilus NCFM GCA_000011985.1
59 Lactobacillus amylovorus GCA_000191545.1
59 Lactobacillus crispatus ST1 GCA_000091765.1
59 Lactobacillus hamsteri DSM 5661 = JCM 6256 GCA_000615445.1
59 Lactobacillus jensenii GCA_001936235.1
59 Lactobacillus psittaci DSM 15354 GCA_000425905.1
56 Lactobacillus kalixensis DSM 16043 GCA_001434335.1
56 Streptococcus macacae NCTC 11558 GCA_000187995.3
56 Streptococcus mutans UA159 GCA_000007465.2
8E
179 Deinococcus radiodurans R1 GCA_000008565.1
127 Bacillus cecembensis GCA_001439635.1
127 Escherichia coli IAI39 GCA_000026345.1
127 Escherichia coli O104_3AH4 str. 2011C-3493 GCA_000299455.1
127 Escherichia coli O157_3AH7 str. Sakai GCA_000008865.1
127 Escherichia coli O83_3AH1 str. NRG 857C GCA_000183345.1
127 Escherichia coli UMN026 GCA_000026325.2
127 Escherichia coli str. K-12 substr. MG1655 GCA_000005845.2
127 Shigella dysenteriae Sd197 GCA_000012005.1
127 Shigella flexneri 2a str. 301 GCA_000006925.2
127 Tumebacillus flagellatus GCA_000714935.1
117 Deinococcus geothermalis DSM 11300 GCA_000196275.1
8F
274 Escherichia coli IAI39 GCA_000026345.1
274 Escherichia coli O104_3AH4 str. 2011C-3493 GCA_000299455.1
274 Escherichia coli O157_3AH7 str. Sakai GCA_000008865.1
274 Escherichia coli UMN026 GCA_000026325.2
274 Escherichia coli str. K-12 substr. MG1655 GCA_000005845.2
274 Shigella flexneri 2a str. 301 GCA_000006925.2
274 Tumebacillus flagellatus GCA_000714935.1
273 Escherichia coli O83_3AH1 str. NRG 857C GCA_000183345.1
265 Shigella dysenteriae Sd197 GCA_000012005.1
8G
175 Escherichia coli IAI39 GCA_000026345.1
175 Escherichia coli O104_3AH4 str. 2011C-3493 GCA_000299455.1
175 Escherichia coli O157_3AH7 str. Sakai GCA_000008865.1
175 Escherichia coli O83_3AH1 str. NRG 857C GCA_000183345.1
175 Escherichia coli UMN026 GCA_000026325.2
175 Escherichia coli str. K-12 substr. MG1655 GCA_000005845.2
175 Shigella dysenteriae Sd197 GCA_000012005.1
175 Tumebacillus flagellatus GCA_000714935.1
166 Shigella flexneri 2a str. 301 GCA_000006925.2
151 Erwinia iniecta GCA_001267535.1

8H
96 Bifidobacterium adolescentis ATCC 15703 GCA_000010425.1
84 Bifidobacterium dentium JCM 1195 = DSM 20436 GCA_001042595.1
84 Bifidobacterium thermophilum GCA_000741495.1
84 Bifidobacterium tsurumiense GCA_000741765.1
82 Bifidobacterium angulatum DSM 20098 = JCM 7096 GCA_001025155.1
82 Bifidobacterium callitrichos DSM 23973 GCA_000741175.1
82 Nesterenkonia alba DSM 19423 GCA_000421745.1
82 Pseudoglutamicibacter albus DNF00011 GCA_000758985.1
8I
265 Bifidobacterium adolescentis ATCC 15703 GCA_000010425.1
245 Bifidobacterium choerinum GCA_000741135.1
245 Bifidobacterium pseudolongum PV8-2 GCA_000800475.2
244 Bifidobacterium angulatum DSM 20098 = JCM 7096 GCA_001025155.1
8J
239 Deinococcus radiodurans R1 GCA_000008565.1
129 Deinococcus marmoris DSM 12784 GCA_000701405.1
129 Deinococcus swuensis GCA_000800395.1
121 Deinococcus puniceus GCA_001644565.1
8K
237 Enterococcus faecalis V583 GCA_000007785.1
237 Streptomyces cinnamoneus GCA_001885705.1
225 Enterococcus hirae ATCC 9790 GCA_000271405.2
211 Enterococcus canis NBRC 100695 GCA_001544375.1
211 Enterococcus faecium DO GCA_000174395.2
211 Enterococcus haemoperoxidus ATCC BAA-382 GCA_000407165.1
211 Enterococcus mundtii QU 25 GCA_000504125.1
211 Enterococcus phoeniculicola ATCC BAA-412 GCA_000407505.1
211 Enterococcus rivorum GCA_001742285.1
211 Enterococcus thailandicus GCA_001652875.1
8L
196 Escherichia coli IAI39 GCA_000026345.1
196 Escherichia coli O104_3AH4 str. 2011C-3493 GCA_000299455.1
196 Escherichia coli O157_3AH7 str. Sakai GCA_000008865.1
196 Escherichia coli O83_3AH1 str. NRG 857C GCA_000183345.1
196 Escherichia coli UMN026 GCA_000026325.2
196 Escherichia coli str. K-12 substr. MG1655 GCA_000005845.2
196 Shigella dysenteriae Sd197 GCA_000012005.1
196 Shigella flexneri 2a str. 301 GCA_000006925.2
196 Tumebacillus flagellatus GCA_000714935.1
192 Cronobacter sakazakii GCA_000982825.1
192 Erwinia iniecta GCA_001267535.1
192 Erwinia toletana DAPP-PG 735 GCA_000336255.1
192 Rosenbergiella nectarea GCA_900111105.1
190 Erwinia gerundensis GCA_001517405.1
190 Pantoea agglomerans GCA_001709315.1
190 Pantoea ananatis LMG 20103 GCA_000025405.2
190 Pantoea dispersa EGD-AAK13 GCA_000465555.2
190 Shimwellia blattae DSM 4481 = NBRC 105725 GCA_000262305.1
8M
170 Megasphaera cerevisiae DSM 20462 GCA_001045675.1
170 Staphylococcus aureus subsp. aureus NCTC 8325 GCA_000013425.1
170 Staphylococcus capitis subsp. capitis GCA_001028645.1
170 Staphylococcus condimenti GCA_001618885.1
170 Staphylococcus epidermidis ATCC 12228 GCA_000007645.1
170 Staphylococcus haemolyticus JCSC1435 GCA_000009865.1
170 Staphylococcus hominis subsp. hominis C80 GCA_000183685.1
170 Staphylococcus lugdunensis HKU09-01 GCA_000025085.1
170 Staphylococcus simiae CCM 7213 GCA_000235645.2
170 Staphylococcus simulans GCA_001559115.1
170 Staphylococcus warneri SG1 GCA_000332735.1
166 Staphylococcus arlettae CVD059 GCA_000295715.1
166 Staphylococcus cohnii subsp. cohnii GCA_000972575.1
166 Staphylococcus equorum GCA_001432245.1
166 Staphylococcus gallinarum GCA_000875895.1
166 Staphylococcus saprophyticus GCA_001074355.1
166 Staphylococcus saprophyticus subsp. saprophyticus ATCC 15305 GCA_000010125.1
166 Staphylococcus succinus GCA_001902315.1
166 Staphylococcus xylosus GCA_000706685.1
162 Staphylococcus hyicus GCA_000816085.1
162 Staphylococcus lutrae GCA_002101335.1
162 Staphylococcus pseudintermedius HKU10-03 GCA_000185885.1
8N
211 Deinococcus radiodurans R1 GCA_000008565.1
138 Deinococcus deserti VCD115 GCA_000020685.1
138 Deinococcus gobiensis I-0 GCA_000252445.1

119 Deinococcus puniceus GCA_001644565.1
8O
184 Megasphaera cerevisiae DSM 20462 GCA_001045675.1
184 Staphylococcus capitis subsp. capitis GCA_001028645.1
184 Staphylococcus condimenti GCA_001618885.1
184 Staphylococcus epidermidis ATCC 12228 GCA_000007645.1
184 Staphylococcus simulans GCA_001559115.1
184 Staphylococcus warneri SG1 GCA_000332735.1
180 Staphylococcus pseudintermedius HKU10-03 GCA_000185885.1
178 Staphylococcus aureus subsp. aureus NCTC 8325 GCA_000013425.1
178 Staphylococcus simiae CCM 7213 GCA_000235645.2
8P
249 Escherichia coli IAI39 GCA_000026345.1
249 Escherichia coli O104_3AH4 str. 2011C-3493 GCA_000299455.1
249 Escherichia coli O157_3AH7 str. Sakai GCA_000008865.1
249 Escherichia coli O83_3AH1 str. NRG 857C GCA_000183345.1
249 Escherichia coli UMN026 GCA_000026325.2
249 Escherichia coli str. K-12 substr. MG1655 GCA_000005845.2
249 Shigella flexneri 2a str. 301 GCA_000006925.2
249 Tumebacillus flagellatus GCA_000714935.1
234 Shigella dysenteriae Sd197 GCA_000012005.1
215 Erwinia iniecta GCA_001267535.1
8Q
310 Enterococcus faecalis V583 GCA_000007785.1
310 Streptomyces cinnamoneus GCA_001885705.1
269 Enterococcus canis NBRC 100695 GCA_001544375.1
269 Enterococcus dispar ATCC 51266 GCA_000406945.1
269 Enterococcus faecium DO GCA_000174395.2
269 Enterococcus haemoperoxidus ATCC BAA-382 GCA_000407165.1
269 Enterococcus hirae ATCC 9790 GCA_000271405.2
269 Enterococcus massiliensis GCA_001050095.1
269 Enterococcus mundtii QU 25 GCA_000504125.1
269 Enterococcus phoeniculicola ATCC BAA-412 GCA_000407505.1
269 Enterococcus rivorum GCA_001742285.1
269 Enterococcus thailandicus GCA_001652875.1
259 Enterococcus aquimarinus GCA_001885765.1
259 Enterococcus asini ATCC 700915 GCA_000407365.1
259 Enterococcus cecorum GCA_001318405.1
259 Enterococcus columbae DSM 7374 = ATCC 51263 GCA_000406925.1
259 Enterococcus gilvus ATCC BAA-350 GCA_000407545.1
259 Enterococcus hermanniensis GCA_001885945.1
259 Enterococcus malodoratus ATCC 43197 GCA_000407185.1
259 Enterococcus pallens ATCC BAA-351 GCA_000407485.1
259 Enterococcus pseudoavium NBRC 100491 GCA_001544295.1
8R
243 Bacillus thuringiensis YBT-1518 GCA_000497525.2
229 Bacillus anthracis str. Ames GCA_000007845.1
229 Bacillus anthracis str. Sterne GCA_000008165.1
229 Bacillus cereus ATCC 14579 GCA_000007825.1
229 Bacillus mycoides GCA_000832605.1
229 Bacillus pseudomycoides DSM 12442 GCA_000161455.1
229 _5BBacillus thuringiensis_5D serovar konkukian str. 97-27 GCA_000008505.1
164 Bacillus horneckiae GCA_001636335.1
164 Bacillus solani GCA_001420595.1
8S
111 Deinococcus radiodurans R1 GCA_000008565.1
64 Deinococcus gobiensis I-0 GCA_000252445.1
62 Deinococcus deserti VCD115 GCA_000020685.1
8T
235 Deinococcus radiodurans R1 GCA_000008565.1
178 Deinococcus gobiensis I-0 GCA_000252445.1
137 Deinococcus marmoris DSM 12784 GCA_000701405.1
137 Deinococcus swuensis GCA_000800395.1
8U
285 Bacillus anthracis str. Ames GCA_000007845.1
285 Bacillus anthracis str. Sterne GCA_000008165.1
285 Bacillus cereus ATCC 14579 GCA_000007825.1
285 Bacillus mycoides GCA_000832605.1
285 Bacillus pseudomycoides DSM 12442 GCA_000161455.1
285 Bacillus thuringiensis YBT-1518 GCA_000497525.2
285 _5BBacillus thuringiensis_5D serovar konkukian str. 97-27 GCA_000008505.1
221 Bacillus coahuilensis m4-4 GCA_000171615.1
219 Staphylococcus saprophyticus GCA_001074355.1
8V
251 Clostridium beijerinckii GCA_000833105.2
251 Clostridium saccharobutylicum DSM 13864 GCA_000473995.1

251 Clostridium saccharoperbutylacetonicum N1-4_28HMT_29 GCA_000340885.1
234 Clostridium puniceum GCA_002006345.1
229 Clostridium butyricum GCA_001456065.2
8W
218 Megasphaera cerevisiae DSM 20462 GCA_001045675.1
218 Staphylococcus aureus subsp. aureus NCTC 8325 GCA_000013425.1
218 Staphylococcus epidermidis ATCC 12228 GCA_000007645.1
218 Staphylococcus haemolyticus JCSC1435 GCA_000009865.1
218 Staphylococcus hominis subsp. hominis C80 GCA_000183685.1
218 Staphylococcus simiae CCM 7213 GCA_000235645.2
218 Staphylococcus warneri SG1 GCA_000332735.1
214 Staphylococcus cohnii subsp. cohnii GCA_000972575.1
213 Staphylococcus capitis subsp. capitis GCA_001028645.1
213 Staphylococcus condimenti GCA_001618885.1
213 Staphylococcus lutrae GCA_002101335.1
213 Staphylococcus simulans GCA_001559115.1
8X
222 Staphylococcus epidermidis ATCC 12228 GCA_000007645.1
219 Staphylococcus capitis subsp. capitis GCA_001028645.1
219 Staphylococcus haemolyticus JCSC1435 GCA_000009865.1
207 Staphylococcus cohnii subsp. cohnii GCA_000972575.1
207 Staphylococcus saprophyticus GCA_001074355.1
8Y
292 Lactobacillus gasseri ATCC 33323 = JCM 1131 GCA_000014425.1
276 Lactobacillus hominis DSM 23910 = CRBIP 24.179 GCA_000296835.1
216 Lactobacillus iners DSM 13335 GCA_000160875.1
8Z
278 Clostridium beijerinckii GCA_000833105.2
278 Clostridium saccharobutylicum DSM 13864 GCA_000473995.1
278 Clostridium saccharoperbutylacetonicum N1-4_28HMT_29 GCA_000340885.1
272 Clostridium puniceum GCA_002006345.1
246 Clostridium chromiireducens GCA_002029255.1
90
134 Streptococcus mutans UA159 GCA_000007465.2
98 Streptococcus cristatus AS 1.3089 GCA_000385925.1
98 Streptococcus gordonii str. Challis substr. CH1 GCA_000017005.1
98 Streptococcus mitis B6 GCA_000027165.1
98 Streptococcus parasanguinis ATCC 15912 GCA_000164675.2
98 Streptococcus pneumoniae R6 GCA_000007045.1
90 Tetragenococcus halophilus NBRC 12172 GCA_000283615.1
91
129 Clostridium beijerinckii GCA_000833105.2
129 Clostridium puniceum GCA_002006345.1
129 Clostridium saccharoperbutylacetonicum N1-4_28HMT_29 GCA_000340885.1
115 Clostridium saccharobutylicum DSM 13864 GCA_000473995.1
100 Clostridium amylolyticum GCA_900142075.1
100 Clostridium intestinale URNW GCA_000469625.2
100 Clostridium polynesiense GCA_000820705.1
92
180 Clostridium saccharobutylicum DSM 13864 GCA_000473995.1
176 Clostridium beijerinckii GCA_000833105.2
176 Clostridium puniceum GCA_002006345.1
176 Clostridium saccharoperbutylacetonicum N1-4_28HMT_29 GCA_000340885.1
165 Clostridium chromiireducens GCA_002029255.1
93
275 Bacillus thuringiensis YBT-1518 GCA_000497525.2
260 Bacillus anthracis str. Ames GCA_000007845.1
260 Bacillus anthracis str. Sterne GCA_000008165.1
260 Bacillus cereus ATCC 14579 GCA_000007825.1
260 Bacillus mycoides GCA_000832605.1
260 Bacillus pseudomycoides DSM 12442 GCA_000161455.1
260 _5BBacillus thuringiensis_5D serovar konkukian str. 97-27 GCA_000008505.1
225 Bacillus manliponensis GCA_000712595.1
94
227 Staphylococcus epidermidis ATCC 12228 GCA_000007645.1
224 Staphylococcus haemolyticus JCSC1435 GCA_000009865.1
219 Staphylococcus capitis subsp. capitis GCA_001028645.1
95
301 Enterococcus faecalis V583 GCA_000007785.1
301 Streptomyces cinnamoneus GCA_001885705.1
273 Enterococcus canis NBRC 100695 GCA_001544375.1
273 Enterococcus dispar ATCC 51266 GCA_000406945.1
273 Enterococcus faecium DO GCA_000174395.2
273 Enterococcus gilvus ATCC BAA-350 GCA_000407545.1
273 Enterococcus haemoperoxidus ATCC BAA-382 GCA_000407165.1
273 Enterococcus hermanniensis GCA_001885945.1

273 Enterococcus hirae ATCC 9790 GCA_000271405.2
273 Enterococcus malodoratus ATCC 43197 GCA_000407185.1
273 Enterococcus mundtii QU 25 GCA_000504125.1
273 Enterococcus pallens ATCC BAA-351 GCA_000407485.1
273 Enterococcus phoeniculicola ATCC BAA-412 GCA_000407505.1
273 Enterococcus pseudoavium NBRC 100491 GCA_001544295.1
273 Enterococcus rivorum GCA_001742285.1
273 Enterococcus thailandicus GCA_001652875.1
264 Enterococcus massiliensis GCA_001050095.1
96
199 Streptococcus mutans UA159 GCA_000007465.2
136 Streptococcus ratti FA-1 = DSM 20564 GCA_000286075.1
132 Streptococcus gordonii str. Challis substr. CH1 GCA_000017005.1
132 Streptococcus macacae NCTC 11558 GCA_000187995.3
97
242 Streptococcus mutans UA159 GCA_000007465.2
163 Streptococcus macacae NCTC 11558 GCA_000187995.3
157 Streptococcus ratti FA-1 = DSM 20564 GCA_000286075.1
98
237 Clostridium beijerinckii GCA_000833105.2
237 Clostridium saccharoperbutylacetonicum N1-4_28HMT_29 GCA_000340885.1
229 Clostridium saccharobutylicum DSM 13864 GCA_000473995.1
223 Clostridium puniceum GCA_002006345.1
99
212 Deinococcus radiodurans R1 GCA_000008565.1
134 Deinococcus proteolyticus MRP GCA_000190555.1
132 Deinococcus deserti VCD115 GCA_000020685.1
132 Deinococcus gobiensis I-0 GCA_000252445.1
132 Deinococcus soli Cha et al. 2016 GCA_001007995.1
9a
238 Enterococcus faecalis V583 GCA_000007785.1
238 Streptomyces cinnamoneus GCA_001885705.1
224 Enterococcus dispar ATCC 51266 GCA_000406945.1
221 Enterococcus asini ATCC 700915 GCA_000407365.1
221 Enterococcus canis NBRC 100695 GCA_001544375.1
221 Enterococcus cecorum GCA_001318405.1
221 Enterococcus columbae DSM 7374 = ATCC 51263 GCA_000406925.1
221 Enterococcus faecium DO GCA_000174395.2
221 Enterococcus haemoperoxidus ATCC BAA-382 GCA_000407165.1
221 Enterococcus hirae ATCC 9790 GCA_000271405.2
221 Enterococcus massiliensis GCA_001050095.1
221 Enterococcus mundtii QU 25 GCA_000504125.1
221 Enterococcus phoeniculicola ATCC BAA-412 GCA_000407505.1
221 Enterococcus rivorum GCA_001742285.1
221 Enterococcus thailandicus GCA_001652875.1
9b
193 Staphylococcus simulans GCA_001559115.1
179 Staphylococcus aureus subsp. aureus NCTC 8325 GCA_000013425.1
179 Staphylococcus capitis subsp. capitis GCA_001028645.1
179 Staphylococcus condimenti GCA_001618885.1
179 Staphylococcus epidermidis ATCC 12228 GCA_000007645.1
179 Staphylococcus haemolyticus JCSC1435 GCA_000009865.1
179 Staphylococcus hominis subsp. hominis C80 GCA_000183685.1
179 Staphylococcus simiae CCM 7213 GCA_000235645.2
172 Staphylococcus lugdunensis HKU09-01 GCA_000025085.1
9c
196 Rhodobacter sphaeroides 2.4.1 GCA_000012905.2
182 Rhodobacter sphaeroides ATCC 17025 GCA_000016405.1
172 Pseudorhodobacter psychrotolerans GCA_001294535.1
9d
254 Lactobacillus gasseri ATCC 33323 = JCM 1131 GCA_000014425.1
238 Lactobacillus hominis DSM 23910 = CRBIP 24.179 GCA_000296835.1
201 Lactobacillus iners DSM 13335 GCA_000160875.1
9e
146 Rhodobacter sphaeroides 2.4.1 GCA_000012905.2
146 Rhodobacter sphaeroides ATCC 17025 GCA_000016405.1
122 Gemmobacter megaterium GCA_900156815.1
122 Pseudorhodobacter ferrugineus DSM 5888 GCA_000420745.1
122 Pseudorhodobacter wandonensis GCA_001202035.1
122 Roseivivax lentus GCA_900156805.1
122 Thioclava dalianensis GCA_000715505.1
122 Thioclava indica GCA_000714545.1
116 Rhodobacter capsulatus SB 1003 GCA_000021865.1
9f
294 Rhodobacter sphaeroides 2.4.1 GCA_000012905.2
264 Pseudorhodobacter psychrotolerans GCA_001294535.1

256 Rhodobacter sphaeroides ATCC 17025 GCA_000016405.1
9g
273 Deinococcus radiodurans R1 GCA_000008565.1
171 Deinococcus gobiensis I-0 GCA_000252445.1
159 Deinococcus proteolyticus MRP GCA_000190555.1
9h
258 Bacillus anthracis str. Ames GCA_000007845.1
258 Bacillus anthracis str. Sterne GCA_000008165.1
258 Bacillus cereus ATCC 14579 GCA_000007825.1
258 Bacillus mycoides GCA_000832605.1
258 Bacillus pseudomycoides DSM 12442 GCA_000161455.1
258 Bacillus thuringiensis YBT-1518 GCA_000497525.2
258 _5BBacillus thuringiensis_5D serovar konkukian str. 97-27 GCA_000008505.1
198 Bacillus manliponensis GCA_000712595.1
189 Bacillus horneckiae GCA_001636335.1
9i
219 Clostridium beijerinckii GCA_000833105.2
219 Clostridium puniceum GCA_002006345.1
219 Clostridium saccharobutylicum DSM 13864 GCA_000473995.1
219 Clostridium saccharoperbutylacetonicum N1-4_28HMT_29 GCA_000340885.1
201 Clostridium butyricum GCA_001456065.2
197 Clostridium chromiireducens GCA_002029255.1
9j
165 Deinococcus radiodurans R1 GCA_000008565.1
95 Deinococcus gobiensis I-0 GCA_000252445.1
93 Deinococcus deserti VCD115 GCA_000020685.1
9k
215 Bacillus anthracis str. Ames GCA_000007845.1
215 Bacillus anthracis str. Sterne GCA_000008165.1
215 Bacillus cereus ATCC 14579 GCA_000007825.1
215 Bacillus mycoides GCA_000832605.1
215 Bacillus pseudomycoides DSM 12442 GCA_000161455.1
215 Bacillus thuringiensis YBT-1518 GCA_000497525.2
215 _5BBacillus thuringiensis_5D serovar konkukian str. 97-27 GCA_000008505.1
193 Bacillus halmapalus GCA_002019665.1
187 Bacillus horikoshii GCA_002157855.1
9l
187 Bifidobacterium adolescentis ATCC 15703 GCA_000010425.1
172 Bifidobacterium thermophilum GCA_000741495.1
172 Bifidobacterium tsurumiense GCA_000741765.1
164 Bifidobacterium dentium JCM 1195 = DSM 20436 GCA_001042595.1
9m
169 Deinococcus radiodurans R1 GCA_000008565.1
106 Deinococcus puniceus GCA_001644565.1
103 Deinococcus deserti VCD115 GCA_000020685.1
9n
225 Deinococcus radiodurans R1 GCA_000008565.1
143 Deinococcus gobiensis I-0 GCA_000252445.1
116 Deinococcus deserti VCD115 GCA_000020685.1
9o
337 Staphylococcus epidermidis ATCC 12228 GCA_000007645.1
327 Staphylococcus haemolyticus JCSC1435 GCA_000009865.1
327 Staphylococcus lugdunensis HKU09-01 GCA_000025085.1
318 Staphylococcus capitis subsp. capitis GCA_001028645.1
9p
249 Escherichia coli IAI39 GCA_000026345.1
249 Escherichia coli O104_3AH4 str. 2011C-3493 GCA_000299455.1
249 Escherichia coli O157_3AH7 str. Sakai GCA_000008865.1
249 Escherichia coli O83_3AH1 str. NRG 857C GCA_000183345.1
249 Escherichia coli UMN026 GCA_000026325.2
249 Escherichia coli str. K-12 substr. MG1655 GCA_000005845.2
249 Shigella dysenteriae Sd197 GCA_000012005.1
249 Tumebacillus flagellatus GCA_000714935.1
236 Shigella flexneri 2a str. 301 GCA_000006925.2
226 Cronobacter sakazakii GCA_000982825.1
226 Erwinia iniecta GCA_001267535.1
226 Erwinia toletana DAPP-PG 735 GCA_000336255.1
226 Kosakonia cowanii GCA_001975225.1
9q
195 Clostridium beijerinckii GCA_000833105.2
195 Clostridium saccharobutylicum DSM 13864 GCA_000473995.1
195 Clostridium saccharoperbutylacetonicum N1-4_28HMT_29 GCA_000340885.1
188 Clostridium puniceum GCA_002006345.1
173 Clostridium chromiireducens GCA_002029255.1
9r
209 Escherichia coli UMN026 GCA_000026325.2

209 Shigella flexneri 2a str. 301 GCA_000006925.2
205 Escherichia coli IAI39 GCA_000026345.1
205 Escherichia coli O104_3AH4 str. 2011C-3493 GCA_000299455.1
205 Escherichia coli O157_3AH7 str. Sakai GCA_000008865.1
205 Escherichia coli O83_3AH1 str. NRG 857C GCA_000183345.1
205 Escherichia coli str. K-12 substr. MG1655 GCA_000005845.2
205 Shigella dysenteriae Sd197 GCA_000012005.1
205 Tumebacillus flagellatus GCA_000714935.1
186 Erwinia iniecta GCA_001267535.1
9s
272 Deinococcus radiodurans R1 GCA_000008565.1
171 Deinococcus gobiensis I-0 GCA_000252445.1
157 Deinococcus puniceus GCA_001644565.1
9t
206 Bifidobacterium adolescentis ATCC 15703 GCA_000010425.1
195 Bifidobacterium asteroides PRL2011 GCA_000304215.1
176 Bifidobacterium callitrichos DSM 23973 GCA_000741175.1
9u
207 Deinococcus radiodurans R1 GCA_000008565.1
131 Deinococcus marmoris DSM 12784 GCA_000701405.1
131 Deinococcus swuensis GCA_000800395.1
116 Deinococcus frigens DSM 12807 GCA_000701425.1
9v
279 Bacillus anthracis str. Ames GCA_000007845.1
279 Bacillus anthracis str. Sterne GCA_000008165.1
279 Bacillus cereus ATCC 14579 GCA_000007825.1
279 Bacillus mycoides GCA_000832605.1
279 Bacillus pseudomycoides DSM 12442 GCA_000161455.1
279 Bacillus thuringiensis YBT-1518 GCA_000497525.2
279 _5BBacillus thuringiensis_5D serovar konkukian str. 97-27 GCA_000008505.1
220 Bacillus manliponensis GCA_000712595.1
207 Bacillus marisflavi GCA_001274775.1
9w
219 Escherichia coli IAI39 GCA_000026345.1
219 Escherichia coli O104_3AH4 str. 2011C-3493 GCA_000299455.1
219 Escherichia coli O157_3AH7 str. Sakai GCA_000008865.1
219 Escherichia coli O83_3AH1 str. NRG 857C GCA_000183345.1
219 Escherichia coli UMN026 GCA_000026325.2
219 Escherichia coli str. K-12 substr. MG1655 GCA_000005845.2
219 Shigella dysenteriae Sd197 GCA_000012005.1
219 Shigella flexneri 2a str. 301 GCA_000006925.2
219 Tumebacillus flagellatus GCA_000714935.1
179 Erwinia iniecta GCA_001267535.1
179 Rosenbergiella nectarea GCA_900111105.1
177 Cedecea neteri GCA_000757825.1
177 Edwardsiella anguillarum ET080813 GCA_000264765.2
9x
265 Clostridium beijerinckii GCA_000833105.2
265 Clostridium puniceum GCA_002006345.1
265 Clostridium saccharoperbutylacetonicum N1-4_28HMT_29 GCA_000340885.1
257 Clostridium saccharobutylicum DSM 13864 GCA_000473995.1
239 Clostridium chromiireducens GCA_002029255.1
239 Clostridium taeniosporum GCA_001735765.1
9y
237 Staphylococcus aureus subsp. aureus NCTC 8325 GCA_000013425.1
237 Staphylococcus epidermidis ATCC 12228 GCA_000007645.1
237 Staphylococcus haemolyticus JCSC1435 GCA_000009865.1
237 Staphylococcus hominis subsp. hominis C80 GCA_000183685.1
237 Staphylococcus lugdunensis HKU09-01 GCA_000025085.1
237 Staphylococcus simiae CCM 7213 GCA_000235645.2
232 Staphylococcus condimenti GCA_001618885.1
232 Staphylococcus microti GCA_000934465.1
232 Staphylococcus simulans GCA_001559115.1
231 Staphylococcus capitis subsp. capitis GCA_001028645.1
9z
181 Deinococcus radiodurans R1 GCA_000008565.1
114 Deinococcus gobiensis I-0 GCA_000252445.1
96 Deinococcus deserti VCD115 GCA_000020685.1
9A
263 Streptococcus mutans UA159 GCA_000007465.2
181 Streptococcus macacae NCTC 11558 GCA_000187995.3
176 Streptococcus ferus DSM 20646 GCA_000372425.1
9B
202 Deinococcus radiodurans R1 GCA_000008565.1
139 Deinococcus marmoris DSM 12784 GCA_000701405.1
139 Deinococcus swuensis GCA_000800395.1

135 Deinococcus deserti VCD115 GCA_000020685.1
9C
194 Staphylococcus epidermidis ATCC 12228 GCA_000007645.1
194 Staphylococcus haemolyticus JCSC1435 GCA_000009865.1
180 Staphylococcus capitis subsp. capitis GCA_001028645.1
180 Staphylococcus cohnii subsp. cohnii GCA_000972575.1
178 Megasphaera cerevisiae DSM 20462 GCA_001045675.1
178 Staphylococcus aureus subsp. aureus NCTC 8325 GCA_000013425.1
178 Staphylococcus condimenti GCA_001618885.1
178 Staphylococcus hominis subsp. hominis C80 GCA_000183685.1
178 Staphylococcus simiae CCM 7213 GCA_000235645.2
178 Staphylococcus simulans GCA_001559115.1
178 Staphylococcus warneri SG1 GCA_000332735.1
9D
210 Lactobacillus gasseri ATCC 33323 = JCM 1131 GCA_000014425.1
202 Lactobacillus hominis DSM 23910 = CRBIP 24.179 GCA_000296835.1
171 Lactobacillus antri DSM 16041 GCA_000160835.1
171 Lactobacillus frumenti DSM 13145 GCA_001436045.1
171 Lactobacillus reuteri DSM 20016 GCA_000016825.1
171 Lactobacillus secaliphilus GCA_001437055.1
171 Lactobacillus vaginalis DSM 5837 = ATCC 49540 GCA_000159435.1
9E
167 Escherichia coli IAI39 GCA_000026345.1
167 Escherichia coli O104_3AH4 str. 2011C-3493 GCA_000299455.1
167 Escherichia coli O157_3AH7 str. Sakai GCA_000008865.1
167 Escherichia coli O83_3AH1 str. NRG 857C GCA_000183345.1
167 Escherichia coli UMN026 GCA_000026325.2
167 Escherichia coli str. K-12 substr. MG1655 GCA_000005845.2
167 Shigella dysenteriae Sd197 GCA_000012005.1
167 Shigella flexneri 2a str. 301 GCA_000006925.2
167 Tumebacillus flagellatus GCA_000714935.1
149 Erwinia iniecta GCA_001267535.1
146 Erwinia toletana DAPP-PG 735 GCA_000336255.1
9F
213 Bifidobacterium tsurumiense GCA_000741765.1
211 Bifidobacterium adolescentis ATCC 15703 GCA_000010425.1
198 Bifidobacterium angulatum DSM 20098 = JCM 7096 GCA_001025155.1
9G
323 Clostridium beijerinckii GCA_000833105.2
323 Clostridium puniceum GCA_002006345.1
323 Clostridium saccharoperbutylacetonicum N1-4_28HMT_29 GCA_000340885.1
314 Clostridium saccharobutylicum DSM 13864 GCA_000473995.1
310 Clostridium chromiireducens GCA_002029255.1
9H
198 Clostridium beijerinckii GCA_000833105.2
198 Clostridium puniceum GCA_002006345.1
198 Clostridium saccharoperbutylacetonicum N1-4_28HMT_29 GCA_000340885.1
183 Clostridium saccharobutylicum DSM 13864 GCA_000473995.1
168 Clostridium butyricum GCA_001456065.2
168 Clostridium chromiireducens GCA_002029255.1
9I
193 Clostridium beijerinckii GCA_000833105.2
193 Clostridium puniceum GCA_002006345.1
193 Clostridium saccharoperbutylacetonicum N1-4_28HMT_29 GCA_000340885.1
189 Clostridium butyricum GCA_001456065.2
177 Clostridium chromiireducens GCA_002029255.1
177 Clostridium saccharobutylicum DSM 13864 GCA_000473995.1
9J
273 Enterococcus faecalis V583 GCA_000007785.1
273 Streptomyces cinnamoneus GCA_001885705.1
258 Enterococcus asini ATCC 700915 GCA_000407365.1
258 Enterococcus canis NBRC 100695 GCA_001544375.1
258 Enterococcus dispar ATCC 51266 GCA_000406945.1
258 Enterococcus faecium DO GCA_000174395.2
258 Enterococcus haemoperoxidus ATCC BAA-382 GCA_000407165.1
258 Enterococcus hirae ATCC 9790 GCA_000271405.2
258 Enterococcus massiliensis GCA_001050095.1
258 Enterococcus mundtii QU 25 GCA_000504125.1
258 Enterococcus phoeniculicola ATCC BAA-412 GCA_000407505.1
258 Enterococcus rivorum GCA_001742285.1
258 Enterococcus thailandicus GCA_001652875.1
252 Isobaculum melis GCA_900111355.1
9K
259 Streptococcus mutans UA159 GCA_000007465.2
180 Streptococcus ferus DSM 20646 GCA_000372425.1
169 Streptococcus macacae NCTC 11558 GCA_000187995.3

9L
158 Rhodobacter sphaeroides 2.4.1 GCA_000012905.2
143 Rhodobacter sphaeroides ATCC 17025 GCA_000016405.1
117 Rhodobacter capsulatus SB 1003 GCA_000021865.1
9M
202 Staphylococcus epidermidis ATCC 12228 GCA_000007645.1
190 Staphylococcus capitis subsp. capitis GCA_001028645.1
190 Staphylococcus lugdunensis HKU09-01 GCA_000025085.1
176 Staphylococcus cohnii subsp. cohnii GCA_000972575.1
176 Staphylococcus saprophyticus subsp. saprophyticus ATCC 15305 GCA_000010125.1
9N
284 Lactobacillus gasseri ATCC 33323 = JCM 1131 GCA_000014425.1
254 Lactobacillus hominis DSM 23910 = CRBIP 24.179 GCA_000296835.1
179 Lactobacillus psittaci DSM 15354 GCA_000425905.1
9O
177 Rhodobacter sphaeroides ATCC 17025 GCA_000016405.1
173 Rhodobacter sphaeroides 2.4.1 GCA_000012905.2
144 Pseudorhodobacter aquimaris GCA_001202025.1
9P
251 Staphylococcus capitis subsp. capitis GCA_001028645.1
251 Staphylococcus epidermidis ATCC 12228 GCA_000007645.1
240 Staphylococcus haemolyticus JCSC1435 GCA_000009865.1
232 Megasphaera cerevisiae DSM 20462 GCA_001045675.1
232 Staphylococcus lugdunensis HKU09-01 GCA_000025085.1
232 Staphylococcus warneri SG1 GCA_000332735.1
9Q
307 Escherichia coli IAI39 GCA_000026345.1
307 Escherichia coli O104_3AH4 str. 2011C-3493 GCA_000299455.1
307 Escherichia coli O157_3AH7 str. Sakai GCA_000008865.1
307 Escherichia coli O83_3AH1 str. NRG 857C GCA_000183345.1
307 Escherichia coli UMN026 GCA_000026325.2
307 Escherichia coli str. K-12 substr. MG1655 GCA_000005845.2
307 Shigella flexneri 2a str. 301 GCA_000006925.2
307 Tumebacillus flagellatus GCA_000714935.1
292 Shigella dysenteriae Sd197 GCA_000012005.1
261 Erwinia iniecta GCA_001267535.1
9R
162 Bacillus anthracis str. Ames GCA_000007845.1
162 Bacillus anthracis str. Sterne GCA_000008165.1
162 Bacillus cereus ATCC 14579 GCA_000007825.1
162 Bacillus manliponensis GCA_000712595.1
162 Bacillus mycoides GCA_000832605.1
162 Bacillus pseudomycoides DSM 12442 GCA_000161455.1
162 Bacillus thuringiensis YBT-1518 GCA_000497525.2
162 _5BBacillus thuringiensis_5D serovar konkukian str. 97-27 GCA_000008505.1
156 Planococcus antarcticus DSM 14505 GCA_001687565.2
156 Planococcus kocurii GCA_001465835.2
154 Bacillus cihuensis GCA_000504145.1
9S
155 Bacillus thuringiensis YBT-1518 GCA_000497525.2
140 Bacillus anthracis str. Ames GCA_000007845.1
140 Bacillus anthracis str. Sterne GCA_000008165.1
140 Bacillus mycoides GCA_000832605.1
140 Bacillus pseudomycoides DSM 12442 GCA_000161455.1
140 _5BBacillus thuringiensis_5D serovar konkukian str. 97-27 GCA_000008505.1
139 Bacillus cereus ATCC 14579 GCA_000007825.1
9T
189 Rhodobacter sphaeroides 2.4.1 GCA_000012905.2
168 Rhodobacter sphaeroides ATCC 17025 GCA_000016405.1
152 Defluviimonas alba GCA_001620265.1
9U
160 Deinococcus radiodurans R1 GCA_000008565.1
104 Deinococcus deserti VCD115 GCA_000020685.1
104 Deinococcus misasensis DSM 22328 GCA_000745915.1
104 Deinococcus puniceus GCA_001644565.1
102 Deinococcus gobiensis I-0 GCA_000252445.1
9V
161 Rhodobacter sphaeroides 2.4.1 GCA_000012905.2
146 Rhodobacter sphaeroides ATCC 17025 GCA_000016405.1
119 Gemmobacter megaterium GCA_900156815.1
119 Pseudorhodobacter ferrugineus DSM 5888 GCA_000420745.1
119 Pseudorhodobacter wandonensis GCA_001202035.1
9W
227 Deinococcus radiodurans R1 GCA_000008565.1
164 Deinococcus gobiensis I-0 GCA_000252445.1
162 Deinococcus deserti VCD115 GCA_000020685.1

9X
181 Escherichia coli IAI39 GCA_000026345.1
181 Escherichia coli O104_3AH4 str. 2011C-3493 GCA_000299455.1
181 Escherichia coli O157_3AH7 str. Sakai GCA_000008865.1
181 Escherichia coli O83_3AH1 str. NRG 857C GCA_000183345.1
181 Escherichia coli UMN026 GCA_000026325.2
181 Escherichia coli str. K-12 substr. MG1655 GCA_000005845.2
181 Shigella dysenteriae Sd197 GCA_000012005.1
181 Tumebacillus flagellatus GCA_000714935.1
168 Shigella flexneri 2a str. 301 GCA_000006925.2
155 Erwinia iniecta GCA_001267535.1
9Y
171 Lactobacillus gasseri ATCC 33323 = JCM 1131 GCA_000014425.1
158 Lactobacillus hominis DSM 23910 = CRBIP 24.179 GCA_000296835.1
116 Lactobacillus amylovorus GCA_000191545.1
116 Lactobacillus crispatus ST1 GCA_000091765.1
116 Lactobacillus kalixensis DSM 16043 GCA_001434335.1
116 Lactobacillus psittaci DSM 15354 GCA_000425905.1
9Z
237 Clostridium beijerinckii GCA_000833105.2
237 Clostridium saccharobutylicum DSM 13864 GCA_000473995.1
237 Clostridium saccharoperbutylacetonicum N1-4_28HMT_29 GCA_000340885.1
231 Clostridium puniceum GCA_002006345.1
212 Clostridium chromiireducens GCA_002029255.1
a0
229 Escherichia coli IAI39 GCA_000026345.1
229 Escherichia coli O104_3AH4 str. 2011C-3493 GCA_000299455.1
229 Escherichia coli O157_3AH7 str. Sakai GCA_000008865.1
229 Escherichia coli O83_3AH1 str. NRG 857C GCA_000183345.1
229 Escherichia coli UMN026 GCA_000026325.2
229 Escherichia coli str. K-12 substr. MG1655 GCA_000005845.2
229 Shigella flexneri 2a str. 301 GCA_000006925.2
229 Tumebacillus flagellatus GCA_000714935.1
220 Shigella dysenteriae Sd197 GCA_000012005.1
205 Erwinia iniecta GCA_001267535.1
a1
263 Streptococcus mutans UA159 GCA_000007465.2
190 Streptococcus sobrinus DSM 20742 = ATCC 33478 GCA_000686605.1
185 Streptococcus ratti FA-1 = DSM 20564 GCA_000286075.1
a2
250 Escherichia coli IAI39 GCA_000026345.1
250 Escherichia coli O104_3AH4 str. 2011C-3493 GCA_000299455.1
250 Escherichia coli O157_3AH7 str. Sakai GCA_000008865.1
250 Escherichia coli O83_3AH1 str. NRG 857C GCA_000183345.1
250 Escherichia coli UMN026 GCA_000026325.2
250 Escherichia coli str. K-12 substr. MG1655 GCA_000005845.2
250 Shigella dysenteriae Sd197 GCA_000012005.1
250 Shigella flexneri 2a str. 301 GCA_000006925.2
250 Tumebacillus flagellatus GCA_000714935.1
203 Erwinia iniecta GCA_001267535.1
197 Cronobacter sakazakii GCA_000982825.1
197 Erwinia toletana DAPP-PG 735 GCA_000336255.1
a3
286 Deinococcus radiodurans R1 GCA_000008565.1
219 Deinococcus gobiensis I-0 GCA_000252445.1
191 Deinococcus puniceus GCA_001644565.1
a4
316 Clostridium beijerinckii GCA_000833105.2
316 Clostridium saccharoperbutylacetonicum N1-4_28HMT_29 GCA_000340885.1
308 Clostridium butyricum GCA_001456065.2
307 Clostridium saccharobutylicum DSM 13864 GCA_000473995.1
a5
252 Clostridium beijerinckii GCA_000833105.2
252 Clostridium saccharoperbutylacetonicum N1-4_28HMT_29 GCA_000340885.1
248 Clostridium saccharobutylicum DSM 13864 GCA_000473995.1
245 Clostridium puniceum GCA_002006345.1
a6
144 Clostridium beijerinckii GCA_000833105.2
144 Clostridium puniceum GCA_002006345.1
144 Clostridium saccharoperbutylacetonicum N1-4_28HMT_29 GCA_000340885.1
142 Clostridium chromiireducens GCA_002029255.1
142 Clostridium saccharobutylicum DSM 13864 GCA_000473995.1
141 Clostridium taeniosporum GCA_001735765.1
a7
256 Streptococcus mutans UA159 GCA_000007465.2
165 Streptococcus equinus GCA_000964315.1

165 Streptococcus gallolyticus subsp. gallolyticus DSM 16831 GCA_002000985.1
165 Streptococcus gordonii str. Challis substr. CH1 GCA_000017005.1
165 Streptococcus orisratti DSM 15617 GCA_000380105.1
165 Streptococcus ratti FA-1 = DSM 20564 GCA_000286075.1
156 Streptococcus cristatus AS 1.3089 GCA_000385925.1
156 Streptococcus macacae NCTC 11558 GCA_000187995.3
156 Streptococcus mitis B6 GCA_000027165.1
156 Streptococcus pneumoniae R6 GCA_000007045.1
a8
236 Escherichia coli IAI39 GCA_000026345.1
236 Escherichia coli O104_3AH4 str. 2011C-3493 GCA_000299455.1
236 Escherichia coli O157_3AH7 str. Sakai GCA_000008865.1
236 Escherichia coli O83_3AH1 str. NRG 857C GCA_000183345.1
236 Escherichia coli UMN026 GCA_000026325.2
236 Escherichia coli str. K-12 substr. MG1655 GCA_000005845.2
236 Shigella flexneri 2a str. 301 GCA_000006925.2
236 Tumebacillus flagellatus GCA_000714935.1
229 Erwinia iniecta GCA_001267535.1
227 Shigella dysenteriae Sd197 GCA_000012005.1
a9
169 Clostridium neonatale GCA_001458595.1
167 Clostridium butyricum GCA_001456065.2
161 Clostridium saccharobutylicum DSM 13864 GCA_000473995.1
aa
271 Bacillus thuringiensis YBT-1518 GCA_000497525.2
254 Bacillus anthracis str. Ames GCA_000007845.1
254 Bacillus anthracis str. Sterne GCA_000008165.1
254 Bacillus cereus ATCC 14579 GCA_000007825.1
254 Bacillus mycoides GCA_000832605.1
254 Bacillus pseudomycoides DSM 12442 GCA_000161455.1
254 _5BBacillus thuringiensis_5D serovar konkukian str. 97-27 GCA_000008505.1
184 Massilibacterium senegalense GCA_001375675.1
ab
153 Rhodobacter sphaeroides 2.4.1 GCA_000012905.2
150 Pseudorhodobacter psychrotolerans GCA_001294535.1
140 Gemmobacter aquatilis GCA_900110025.1
140 Gemmobacter megaterium GCA_900156815.1
140 Rhodobacter sphaeroides ATCC 17025 GCA_000016405.1
ac
264 Bacillus anthracis str. Ames GCA_000007845.1
264 Bacillus anthracis str. Sterne GCA_000008165.1
264 Bacillus cereus ATCC 14579 GCA_000007825.1
264 Bacillus mycoides GCA_000832605.1
264 Bacillus pseudomycoides DSM 12442 GCA_000161455.1
264 Bacillus thuringiensis YBT-1518 GCA_000497525.2
264 _5BBacillus thuringiensis_5D serovar konkukian str. 97-27 GCA_000008505.1
206 Bacillus manliponensis GCA_000712595.1
196 Bacillus marisflavi GCA_001274775.1
ad
247 Escherichia coli IAI39 GCA_000026345.1
247 Escherichia coli O104_3AH4 str. 2011C-3493 GCA_000299455.1
247 Escherichia coli O157_3AH7 str. Sakai GCA_000008865.1
247 Escherichia coli O83_3AH1 str. NRG 857C GCA_000183345.1
247 Escherichia coli UMN026 GCA_000026325.2
247 Escherichia coli str. K-12 substr. MG1655 GCA_000005845.2
247 Shigella dysenteriae Sd197 GCA_000012005.1
247 Shigella flexneri 2a str. 301 GCA_000006925.2
247 Tumebacillus flagellatus GCA_000714935.1
224 Enterobacter hormaechei subsp. steigerwaltii GCA_001729725.1
224 Erwinia iniecta GCA_001267535.1
224 Erwinia toletana DAPP-PG 735 GCA_000336255.1
224 Klebsiella oxytoca GCA_001022195.1
224 Kosakonia cowanii GCA_001975225.1
224 Kosakonia sacchari SP1 GCA_000300455.4
224 Pantoea alhagi GCA_002101395.1
224 Pseudescherichia vulneris NBRC 102420 GCA_000759795.1
220 Cronobacter sakazakii GCA_000982825.1
220 Enterobacter cloacae subsp. cloacae ATCC 13047 GCA_000025565.1
220 Enterobacter kobei GCA_900185885.1
220 Salmonella enterica subsp. enterica serovar Typhi str. CT18 GCA_000195995.1
220 Salmonella enterica subsp. enterica serovar Typhimurium str. LT2 GCA_000006945.2
220 Trabulsiella odontotermitis GCA_001297765.1
ae
282 Clostridium beijerinckii GCA_000833105.2
274 Clostridium chromiireducens GCA_002029255.1
274 Clostridium puniceum GCA_002006345.1

274 Clostridium saccharobutylicum DSM 13864 GCA_000473995.1
274 Clostridium saccharoperbutylacetonicum N1-4_28HMT_29 GCA_000340885.1
265 Clostridium butyricum GCA_001456065.2
af
186 Bacillus thuringiensis YBT-1518 GCA_000497525.2
180 Bacillus anthracis str. Ames GCA_000007845.1
180 Bacillus anthracis str. Sterne GCA_000008165.1
180 Bacillus cereus ATCC 14579 GCA_000007825.1
180 Bacillus mycoides GCA_000832605.1
180 Bacillus pseudomycoides DSM 12442 GCA_000161455.1
180 _5BBacillus thuringiensis_5D serovar konkukian str. 97-27 GCA_000008505.1
150 Staphylococcus equorum GCA_001432245.1
ag
322 Staphylococcus epidermidis ATCC 12228 GCA_000007645.1
317 Staphylococcus capitis subsp. capitis GCA_001028645.1
313 Megasphaera cerevisiae DSM 20462 GCA_001045675.1
313 Staphylococcus warneri SG1 GCA_000332735.1
ah
291 Deinococcus radiodurans R1 GCA_000008565.1
178 Deinococcus gobiensis I-0 GCA_000252445.1
170 Deinococcus marmoris DSM 12784 GCA_000701405.1
170 Deinococcus swuensis GCA_000800395.1
ai
186 Deinococcus radiodurans R1 GCA_000008565.1
94 Deinococcus marmoris DSM 12784 GCA_000701405.1
94 Deinococcus swuensis GCA_000800395.1
80 Deinococcus gobiensis I-0 GCA_000252445.1
80 Deinococcus puniceus GCA_001644565.1
aj
240 Lactobacillus gasseri ATCC 33323 = JCM 1131 GCA_000014425.1
222 Lactobacillus hominis DSM 23910 = CRBIP 24.179 GCA_000296835.1
196 Lactobacillus jensenii GCA_001936235.1
ak
207 Streptococcus mutans UA159 GCA_000007465.2
126 Streptococcus ferus DSM 20646 GCA_000372425.1
115 Streptococcus marimammalium DSM 18627 GCA_000380045.1
al
225 Clostridium beijerinckii GCA_000833105.2
225 Clostridium puniceum GCA_002006345.1
225 Clostridium saccharobutylicum DSM 13864 GCA_000473995.1
225 Clostridium saccharoperbutylacetonicum N1-4_28HMT_29 GCA_000340885.1
207 Clostridium chromiireducens GCA_002029255.1
201 Clostridium taeniosporum GCA_001735765.1
am
240 Staphylococcus capitis subsp. capitis GCA_001028645.1
240 Staphylococcus epidermidis ATCC 12228 GCA_000007645.1
240 Staphylococcus haemolyticus JCSC1435 GCA_000009865.1
231 Staphylococcus cohnii subsp. cohnii GCA_000972575.1
231 Staphylococcus saprophyticus subsp. saprophyticus ATCC 15305 GCA_000010125.1
229 Staphylococcus hominis subsp. hominis C80 GCA_000183685.1
229 Staphylococcus lugdunensis HKU09-01 GCA_000025085.1
an
219 Deinococcus radiodurans R1 GCA_000008565.1
120 Deinococcus gobiensis I-0 GCA_000252445.1
114 Deinococcus frigens DSM 12807 GCA_000701425.1
114 Deinococcus marmoris DSM 12784 GCA_000701405.1
114 Deinococcus swuensis GCA_000800395.1
ao
181 Clostridium beijerinckii GCA_000833105.2
181 Clostridium puniceum GCA_002006345.1
181 Clostridium saccharobutylicum DSM 13864 GCA_000473995.1
181 Clostridium saccharoperbutylacetonicum N1-4_28HMT_29 GCA_000340885.1
170 Clostridium chromiireducens GCA_002029255.1
159 Clostridium botulinum B str. Eklund 17B _28NRP_29 GCA_000020165.1
159 Clostridium taeniosporum GCA_001735765.1
ap
215 Streptococcus mutans UA159 GCA_000007465.2
144 Streptococcus sobrinus DSM 20742 = ATCC 33478 GCA_000686605.1
139 Streptococcus gordonii str. Challis substr. CH1 GCA_000017005.1
aq
149 Enterococcus faecalis V583 GCA_000007785.1
149 Streptomyces cinnamoneus GCA_001885705.1
113 Enterococcus asini ATCC 700915 GCA_000407365.1
113 Enterococcus canis NBRC 100695 GCA_001544375.1
113 Enterococcus casseliflavus EC20 GCA_000157355.2
113 Enterococcus dispar ATCC 51266 GCA_000406945.1

113 Enterococcus faecium DO GCA_000174395.2
113 Enterococcus haemoperoxidus ATCC BAA-382 GCA_000407165.1
113 Enterococcus hirae ATCC 9790 GCA_000271405.2
113 Enterococcus massiliensis GCA_001050095.1
113 Enterococcus mundtii QU 25 GCA_000504125.1
113 Enterococcus phoeniculicola ATCC BAA-412 GCA_000407505.1
113 Enterococcus rivorum GCA_001742285.1
113 Enterococcus saccharolyticus subsp. saccharolyticus ATCC 43076 GCA_000407285.1
113 Enterococcus thailandicus GCA_001652875.1
113 Isobaculum melis GCA_900111355.1
113 Vagococcus penaei GCA_001998885.1
112 Carnobacterium divergens DSM 20623 GCA_000744255.1
112 Granulicatella balaenopterae GCA_900111135.1
ar
230 Deinococcus radiodurans R1 GCA_000008565.1
136 Deinococcus gobiensis I-0 GCA_000252445.1
122 Deinococcus hopiensis KR-140 GCA_900176165.1
122 Deinococcus puniceus GCA_001644565.1
as
142 Megasphaera cerevisiae DSM 20462 GCA_001045675.1
142 Staphylococcus aureus subsp. aureus NCTC 8325 GCA_000013425.1
142 Staphylococcus epidermidis ATCC 12228 GCA_000007645.1
142 Staphylococcus haemolyticus JCSC1435 GCA_000009865.1
142 Staphylococcus hominis subsp. hominis C80 GCA_000183685.1
142 Staphylococcus lugdunensis HKU09-01 GCA_000025085.1
142 Staphylococcus simiae CCM 7213 GCA_000235645.2
142 Staphylococcus warneri SG1 GCA_000332735.1
137 Staphylococcus lutrae GCA_002101335.1
134 Staphylococcus condimenti GCA_001618885.1
134 Staphylococcus simulans GCA_001559115.1
at
229 Deinococcus radiodurans R1 GCA_000008565.1
168 Deinococcus hopiensis KR-140 GCA_900176165.1
166 Deinococcus deserti VCD115 GCA_000020685.1
au
67 Clostridium botulinum A str. ATCC 3502 GCA_000063585.1
67 Clostridium botulinum A str. Hall GCA_000017045.1
67 Clostridium butyricum GCA_001456065.2
52 Clostridium beijerinckii GCA_000833105.2
52 Clostridium chromiireducens GCA_002029255.1
52 Clostridium cylindrosporum DSM 605 GCA_001047375.1
52 Clostridium fallax GCA_900129365.1
52 Clostridium neonatale GCA_001458595.1
52 Clostridium puniceum GCA_002006345.1
52 Clostridium saccharobutylicum DSM 13864 GCA_000473995.1
52 Clostridium saccharoperbutylacetonicum N1-4_28HMT_29 GCA_000340885.1
52 Clostridium tepidum GCA_002008345.1
52 Fervidicella metallireducens AeB GCA_000601455.1
44 Clostridium cavendishii DSM 21758 GCA_900141845.1
44 Clostridium tepidiprofundi DSM 19306 GCA_001594005.1
av
244 Clostridium beijerinckii GCA_000833105.2
244 Clostridium saccharobutylicum DSM 13864 GCA_000473995.1
244 Clostridium saccharoperbutylacetonicum N1-4_28HMT_29 GCA_000340885.1
240 Clostridium puniceum GCA_002006345.1
225 Clostridium chromiireducens GCA_002029255.1
aw
167 Streptococcus mutans UA159 GCA_000007465.2
105 Streptococcus macacae NCTC 11558 GCA_000187995.3
90 Clostridium beijerinckii GCA_000833105.2
90 Clostridium puniceum GCA_002006345.1
90 Clostridium saccharobutylicum DSM 13864 GCA_000473995.1
90 Clostridium saccharoperbutylacetonicum N1-4_28HMT_29 GCA_000340885.1
ax
207 Bifidobacterium adolescentis ATCC 15703 GCA_000010425.1
193 Bifidobacterium callitrichos DSM 23973 GCA_000741175.1
178 Bifidobacterium lemurum GCA_001895165.1
178 Bifidobacterium longum NCC2705 GCA_000007525.1
178 Bifidobacterium minimum GCA_000741645.1
178 Bifidobacterium mongoliense DSM 21395 GCA_000741285.1
178 Bifidobacterium subtile GCA_000741775.1
ay
193 Deinococcus radiodurans R1 GCA_000008565.1
162 Deinococcus gobiensis I-0 GCA_000252445.1
161 Deinococcus deserti VCD115 GCA_000020685.1
az

270 Streptococcus mutans UA159 GCA_000007465.2
180 Streptococcus gordonii str. Challis substr. CH1 GCA_000017005.1
175 Streptococcus salivarius GCA_000785515.1
175 Streptococcus thermophilus JIM 8232 GCA_000253395.1
aA
213 Bacillus anthracis str. Ames GCA_000007845.1
213 Bacillus anthracis str. Sterne GCA_000008165.1
213 Bacillus cereus ATCC 14579 GCA_000007825.1
213 Bacillus mycoides GCA_000832605.1
213 Bacillus pseudomycoides DSM 12442 GCA_000161455.1
213 Bacillus thuringiensis YBT-1518 GCA_000497525.2
213 _5BBacillus thuringiensis_5D serovar konkukian str. 97-27 GCA_000008505.1
165 Massilibacterium senegalense GCA_001375675.1
155 Oceanobacillus caeni GCA_001298135.1
aB
238 Streptococcus mutans UA159 GCA_000007465.2
107 Streptococcus sobrinus DSM 20742 = ATCC 33478 GCA_000686605.1
105 Streptococcus massiliensis DSM 18628 GCA_000380065.1
aC
297 Deinococcus radiodurans R1 GCA_000008565.1
194 Deinococcus gobiensis I-0 GCA_000252445.1
191 Deinococcus deserti VCD115 GCA_000020685.1
aD
324 Bacillus anthracis str. Ames GCA_000007845.1
324 Bacillus anthracis str. Sterne GCA_000008165.1
324 Bacillus cereus ATCC 14579 GCA_000007825.1
324 Bacillus mycoides GCA_000832605.1
324 Bacillus pseudomycoides DSM 12442 GCA_000161455.1
324 Bacillus thuringiensis YBT-1518 GCA_000497525.2
324 _5BBacillus thuringiensis_5D serovar konkukian str. 97-27 GCA_000008505.1
268 Bacillus manliponensis GCA_000712595.1
242 Bacillus marisflavi GCA_001274775.1
aE
189 Bacillus anthracis str. Ames GCA_000007845.1
189 Bacillus anthracis str. Sterne GCA_000008165.1
189 Bacillus cereus ATCC 14579 GCA_000007825.1
189 Bacillus mycoides GCA_000832605.1
189 Bacillus pseudomycoides DSM 12442 GCA_000161455.1
189 Bacillus thuringiensis YBT-1518 GCA_000497525.2
189 _5BBacillus thuringiensis_5D serovar konkukian str. 97-27 GCA_000008505.1
151 Bacillus cohnii NBRC 15565 GCA_001591425.1
149 Bacillus cytotoxicus NVH 391-98 GCA_000017425.1
149 Bacillus halmapalus GCA_002019665.1
149 Bacillus horikoshii GCA_002157855.1
aF
270 Bifidobacterium adolescentis ATCC 15703 GCA_000010425.1
261 Bifidobacterium dentium JCM 1195 = DSM 20436 GCA_001042595.1
252 Bifidobacterium callitrichos DSM 23973 GCA_000741175.1
aG
267 Escherichia coli IAI39 GCA_000026345.1
267 Escherichia coli O104_3AH4 str. 2011C-3493 GCA_000299455.1
267 Escherichia coli O157_3AH7 str. Sakai GCA_000008865.1
267 Escherichia coli O83_3AH1 str. NRG 857C GCA_000183345.1
267 Escherichia coli UMN026 GCA_000026325.2
267 Escherichia coli str. K-12 substr. MG1655 GCA_000005845.2
267 Shigella dysenteriae Sd197 GCA_000012005.1
267 Tumebacillus flagellatus GCA_000714935.1
247 Shigella flexneri 2a str. 301 GCA_000006925.2
229 Erwinia iniecta GCA_001267535.1
229 Erwinia toletana DAPP-PG 735 GCA_000336255.1
229 Kosakonia cowanii GCA_001975225.1
229 Pluralibacter gergoviae GCA_000757785.1
aH
184 Deinococcus radiodurans R1 GCA_000008565.1
114 Deinococcus puniceus GCA_001644565.1
109 Deinococcus geothermalis DSM 11300 GCA_000196275.1
aI
181 Rhodobacter sphaeroides 2.4.1 GCA_000012905.2
168 Gemmobacter megaterium GCA_900156815.1
166 Gemmobacter aquatilis GCA_900110025.1
aJ
267 Clostridium beijerinckii GCA_000833105.2
267 Clostridium saccharoperbutylacetonicum N1-4_28HMT_29 GCA_000340885.1
259 Clostridium saccharobutylicum DSM 13864 GCA_000473995.1
252 Clostridium puniceum GCA_002006345.1
aK

227 Lactobacillus gasseri ATCC 33323 = JCM 1131 GCA_000014425.1
223 Lactobacillus hominis DSM 23910 = CRBIP 24.179 GCA_000296835.1
171 Lactobacillus psittaci DSM 15354 GCA_000425905.1
aL
320 Staphylococcus epidermidis ATCC 12228 GCA_000007645.1
320 Staphylococcus haemolyticus JCSC1435 GCA_000009865.1
320 Staphylococcus hominis subsp. hominis C80 GCA_000183685.1
320 Staphylococcus lugdunensis HKU09-01 GCA_000025085.1
314 Staphylococcus capitis subsp. capitis GCA_001028645.1
311 Staphylococcus aureus subsp. aureus NCTC 8325 GCA_000013425.1
311 Staphylococcus simiae CCM 7213 GCA_000235645.2
aM
113 Enterococcus faecalis V583 GCA_000007785.1
113 Streptomyces cinnamoneus GCA_001885705.1
113 Vagococcus penaei GCA_001998885.1
112 Bacillus coahuilensis m4-4 GCA_000171615.1
112 Bacillus marisflavi GCA_001274775.1
108 Bacillus anthracis str. Ames GCA_000007845.1
108 Bacillus anthracis str. Sterne GCA_000008165.1
108 Bacillus cereus ATCC 14579 GCA_000007825.1
108 Bacillus mycoides GCA_000832605.1
108 Bacillus pseudomycoides DSM 12442 GCA_000161455.1
108 Bacillus thuringiensis YBT-1518 GCA_000497525.2
108 _5BBacillus thuringiensis_5D serovar konkukian str. 97-27 GCA_000008505.1
aN
189 Deinococcus radiodurans R1 GCA_000008565.1
94 Deinococcus gobiensis I-0 GCA_000252445.1
76 Deinococcus puniceus GCA_001644565.1
aO
228 Enterococcus faecalis V583 GCA_000007785.1
228 Streptomyces cinnamoneus GCA_001885705.1
203 Enterococcus canis NBRC 100695 GCA_001544375.1
203 Enterococcus casseliflavus EC20 GCA_000157355.2
203 Enterococcus dispar ATCC 51266 GCA_000406945.1
203 Enterococcus faecium DO GCA_000174395.2
203 Enterococcus gilvus ATCC BAA-350 GCA_000407545.1
203 Enterococcus hermanniensis GCA_001885945.1
203 Enterococcus hirae ATCC 9790 GCA_000271405.2
203 Enterococcus malodoratus ATCC 43197 GCA_000407185.1
203 Enterococcus massiliensis GCA_001050095.1
203 Enterococcus mundtii QU 25 GCA_000504125.1
203 Enterococcus pallens ATCC BAA-351 GCA_000407485.1
203 Enterococcus pseudoavium NBRC 100491 GCA_001544295.1
203 Enterococcus rivorum GCA_001742285.1
203 Enterococcus saccharolyticus subsp. saccharolyticus ATCC 43076 GCA_000407285.1
191 Enterococcus asini ATCC 700915 GCA_000407365.1
aP
244 Streptococcus mutans UA159 GCA_000007465.2
169 Streptococcus marimammalium DSM 18627 GCA_000380045.1
156 Streptococcus macacae NCTC 11558 GCA_000187995.3
aQ
170 Bacillus anthracis str. Ames GCA_000007845.1
170 Bacillus anthracis str. Sterne GCA_000008165.1
170 Bacillus cereus ATCC 14579 GCA_000007825.1
170 Bacillus mycoides GCA_000832605.1
170 Bacillus pseudomycoides DSM 12442 GCA_000161455.1
170 Bacillus thuringiensis YBT-1518 GCA_000497525.2
170 _5BBacillus thuringiensis_5D serovar konkukian str. 97-27 GCA_000008505.1
146 Staphylococcus equorum GCA_001432245.1
129 Bacillus aquimaris TF-12 GCA_001648555.1
129 Bacillus manliponensis GCA_000712595.1
129 Bacillus marisflavi GCA_001274775.1
129 Bacillus solani GCA_001420595.1
aR
88 Staphylococcus simulans GCA_001559115.1
86 Megasphaera cerevisiae DSM 20462 GCA_001045675.1
86 Staphylococcus capitis subsp. capitis GCA_001028645.1
86 Staphylococcus cohnii subsp. cohnii GCA_000972575.1
86 Staphylococcus epidermidis ATCC 12228 GCA_000007645.1
86 Staphylococcus warneri SG1 GCA_000332735.1
76 Alkalibacterium pelagium GCA_900109325.1
76 Marinilactibacillus piezotolerans GCA_900169305.1
aS
168 Clostridium saccharobutylicum DSM 13864 GCA_000473995.1
166 Clostridium beijerinckii GCA_000833105.2
166 Clostridium puniceum GCA_002006345.1

166 Clostridium saccharoperbutylacetonicum N1-4_28HMT_29 GCA_000340885.1
153 Clostridium chromiireducens GCA_002029255.1
aT
126 Deinococcus radiodurans R1 GCA_000008565.1
88 Deinococcus marmoris DSM 12784 GCA_000701405.1
88 Deinococcus swuensis GCA_000800395.1
75 Deinococcus proteolyticus MRP GCA_000190555.1
aU
266 Escherichia coli IAI39 GCA_000026345.1
266 Escherichia coli O104_3AH4 str. 2011C-3493 GCA_000299455.1
266 Escherichia coli O157_3AH7 str. Sakai GCA_000008865.1
266 Escherichia coli O83_3AH1 str. NRG 857C GCA_000183345.1
266 Escherichia coli UMN026 GCA_000026325.2
266 Escherichia coli str. K-12 substr. MG1655 GCA_000005845.2
266 Shigella dysenteriae Sd197 GCA_000012005.1
266 Shigella flexneri 2a str. 301 GCA_000006925.2
266 Tumebacillus flagellatus GCA_000714935.1
237 Erwinia iniecta GCA_001267535.1
222 Rosenbergiella nectarea GCA_900111105.1
aV
245 Clostridium taeniosporum GCA_001735765.1
244 Clostridium chromiireducens GCA_002029255.1
244 Clostridium saccharobutylicum DSM 13864 GCA_000473995.1
242 Clostridium beijerinckii GCA_000833105.2
242 Clostridium puniceum GCA_002006345.1
242 Clostridium saccharoperbutylacetonicum N1-4_28HMT_29 GCA_000340885.1
aW
115 Megasphaera cerevisiae DSM 20462 GCA_001045675.1
115 Staphylococcus capitis subsp. capitis GCA_001028645.1
115 Staphylococcus cohnii subsp. cohnii GCA_000972575.1
115 Staphylococcus epidermidis ATCC 12228 GCA_000007645.1
115 Staphylococcus warneri SG1 GCA_000332735.1
114 Staphylococcus condimenti GCA_001618885.1
114 Staphylococcus simulans GCA_001559115.1
112 Staphylococcus pseudintermedius HKU10-03 GCA_000185885.1
aX
242 Deinococcus radiodurans R1 GCA_000008565.1
205 Deinococcus gobiensis I-0 GCA_000252445.1
192 Deinococcus puniceus GCA_001644565.1
aY
152 Bifidobacterium adolescentis ATCC 15703 GCA_000010425.1
137 Bifidobacterium tsurumiense GCA_000741765.1
133 Bifidobacterium choerinum GCA_000741135.1
133 Bifidobacterium pseudolongum PV8-2 GCA_000800475.2
aZ
64 Clostridium beijerinckii GCA_000833105.2
64 Clostridium chromiireducens GCA_002029255.1
64 Clostridium puniceum GCA_002006345.1
64 Clostridium saccharobutylicum DSM 13864 GCA_000473995.1
64 Clostridium saccharoperbutylacetonicum N1-4_28HMT_29 GCA_000340885.1
62 Clostridium botulinum B str. Eklund 17B _28NRP_29 GCA_000020165.1
62 Clostridium fallax GCA_900129365.1
62 Clostridium gasigenes GCA_900104115.1
62 Clostridium taeniosporum GCA_001735765.1
62 Clostridium uliginosum GCA_900112485.1
61 Clostridium algidicarnis GCA_000703125.1
61 Clostridium amylolyticum GCA_900142075.1
61 Clostridium intestinale URNW GCA_000469625.2
61 Clostridium polynesiense GCA_000820705.1
b0
172 Bacillus anthracis str. Ames GCA_000007845.1
172 Bacillus anthracis str. Sterne GCA_000008165.1
172 Bacillus cereus ATCC 14579 GCA_000007825.1
172 Bacillus mycoides GCA_000832605.1
172 Bacillus pseudomycoides DSM 12442 GCA_000161455.1
172 Bacillus thuringiensis YBT-1518 GCA_000497525.2
172 _5BBacillus thuringiensis_5D serovar konkukian str. 97-27 GCA_000008505.1
161 Bacillus marisflavi GCA_001274775.1
158 Bacillus horneckiae GCA_001636335.1
b1
307 Escherichia coli IAI39 GCA_000026345.1
307 Escherichia coli O104_3AH4 str. 2011C-3493 GCA_000299455.1
307 Escherichia coli O157_3AH7 str. Sakai GCA_000008865.1
307 Escherichia coli UMN026 GCA_000026325.2
307 Escherichia coli str. K-12 substr. MG1655 GCA_000005845.2
307 Tumebacillus flagellatus GCA_000714935.1

306 Escherichia coli O83_3AH1 str. NRG 857C GCA_000183345.1
300 Shigella flexneri 2a str. 301 GCA_000006925.2
b2
266 Streptococcus mutans UA159 GCA_000007465.2
161 Streptococcus sobrinus DSM 20742 = ATCC 33478 GCA_000686605.1
155 Streptococcus massiliensis DSM 18628 GCA_000380065.1
b3
245 Streptococcus mutans UA159 GCA_000007465.2
137 Streptococcus macacae NCTC 11558 GCA_000187995.3
133 Streptococcus ferus DSM 20646 GCA_000372425.1
b4
262 Clostridium saccharobutylicum DSM 13864 GCA_000473995.1
257 Clostridium beijerinckii GCA_000833105.2
257 Clostridium saccharoperbutylacetonicum N1-4_28HMT_29 GCA_000340885.1
255 Clostridium butyricum GCA_001456065.2
b5
225 Escherichia coli IAI39 GCA_000026345.1
225 Escherichia coli O104_3AH4 str. 2011C-3493 GCA_000299455.1
225 Escherichia coli O157_3AH7 str. Sakai GCA_000008865.1
225 Escherichia coli O83_3AH1 str. NRG 857C GCA_000183345.1
225 Escherichia coli UMN026 GCA_000026325.2
225 Escherichia coli str. K-12 substr. MG1655 GCA_000005845.2
225 Shigella dysenteriae Sd197 GCA_000012005.1
225 Shigella flexneri 2a str. 301 GCA_000006925.2
225 Tumebacillus flagellatus GCA_000714935.1
199 Cronobacter sakazakii GCA_000982825.1
199 Enterobacter cloacae subsp. cloacae ATCC 13047 GCA_000025565.1
199 Enterobacter hormaechei subsp. steigerwaltii GCA_001729725.1
199 Enterobacter kobei GCA_900185885.1
199 Erwinia amylovora CFBP1430 GCA_000091565.1
199 Erwinia gerundensis GCA_001517405.1
199 Erwinia iniecta GCA_001267535.1
199 Erwinia persicina NBRC 102418 GCA_001571305.1
199 Erwinia toletana DAPP-PG 735 GCA_000336255.1
199 Klebsiella oxytoca GCA_001022195.1
199 Kosakonia cowanii GCA_001975225.1
199 Pantoea agglomerans GCA_001709315.1
199 Pantoea alhagi GCA_002101395.1
199 Pantoea dispersa EGD-AAK13 GCA_000465555.2
199 Pantoea rwandensis GCA_000759475.1
199 Pantoea septica GCA_002095575.1
199 Pluralibacter gergoviae GCA_000757785.1
199 Pseudescherichia vulneris NBRC 102420 GCA_000759795.1
199 Salmonella enterica subsp. enterica serovar Typhi str. CT18 GCA_000195995.1
199 Salmonella enterica subsp. enterica serovar Typhimurium str. LT2 GCA_000006945.2
199 Shimwellia blattae DSM 4481 = NBRC 105725 GCA_000262305.1
199 Trabulsiella odontotermitis GCA_001297765.1
198 Erwinia billingiae Eb661 GCA_000196615.1
b6
171 Clostridium beijerinckii GCA_000833105.2
171 Clostridium puniceum GCA_002006345.1
171 Clostridium saccharobutylicum DSM 13864 GCA_000473995.1
171 Clostridium saccharoperbutylacetonicum N1-4_28HMT_29 GCA_000340885.1
167 Clostridium chromiireducens GCA_002029255.1
166 Clostridium fallax GCA_900129365.1
b7
194 Bacillus anthracis str. Ames GCA_000007845.1
194 Bacillus anthracis str. Sterne GCA_000008165.1
194 Bacillus cereus ATCC 14579 GCA_000007825.1
194 Bacillus mycoides GCA_000832605.1
194 Bacillus pseudomycoides DSM 12442 GCA_000161455.1
194 Bacillus thuringiensis YBT-1518 GCA_000497525.2
194 _5BBacillus thuringiensis_5D serovar konkukian str. 97-27 GCA_000008505.1
140 Vagococcus penaei GCA_001998885.1
139 Viridibacillus arvi GCA_001274945.1
b8
287 Clostridium beijerinckii GCA_000833105.2
287 Clostridium puniceum GCA_002006345.1
287 Clostridium saccharoperbutylacetonicum N1-4_28HMT_29 GCA_000340885.1
278 Clostridium saccharobutylicum DSM 13864 GCA_000473995.1
274 Clostridium butyricum GCA_001456065.2
274 Clostridium chromiireducens GCA_002029255.1
b9
224 Lactobacillus gasseri ATCC 33323 = JCM 1131 GCA_000014425.1
209 Lactobacillus hominis DSM 23910 = CRBIP 24.179 GCA_000296835.1
166 Lactobacillus antri DSM 16041 GCA_000160835.1

166 Lactobacillus frumenti DSM 13145 GCA_001436045.1
166 Lactobacillus pontis DSM 8475 GCA_001435345.1
166 Lactobacillus reuteri DSM 20016 GCA_000016825.1
166 Lactobacillus secaliphilus GCA_001437055.1
166 Lactobacillus vaginalis DSM 5837 = ATCC 49540 GCA_000159435.1
ba
129 Enterococcus faecalis V583 GCA_000007785.1
129 Streptomyces cinnamoneus GCA_001885705.1
126 Isobaculum melis GCA_900111355.1
121 Catellicoccus marimammalium M35_2F04_2F3 GCA_000313915.1
bb
243 Staphylococcus condimenti GCA_001618885.1
243 Staphylococcus simulans GCA_001559115.1
230 Staphylococcus microti GCA_000934465.1
228 Staphylococcus aureus subsp. aureus NCTC 8325 GCA_000013425.1
228 Staphylococcus capitis subsp. capitis GCA_001028645.1
228 Staphylococcus epidermidis ATCC 12228 GCA_000007645.1
228 Staphylococcus haemolyticus JCSC1435 GCA_000009865.1
228 Staphylococcus hominis subsp. hominis C80 GCA_000183685.1
228 Staphylococcus lugdunensis HKU09-01 GCA_000025085.1
228 Staphylococcus pettenkoferi GCA_002208805.1
228 Staphylococcus simiae CCM 7213 GCA_000235645.2
bc
151 Bifidobacterium adolescentis ATCC 15703 GCA_000010425.1
146 Bifidobacterium tsurumiense GCA_000741765.1
136 Bifidobacterium breve DSM 20213 = JCM 1192 GCA_001025175.1
136 Bifidobacterium longum NCC2705 GCA_000007525.1
136 Bifidobacterium saguini DSM 23967 GCA_000741715.1
bd
242 Clostridium beijerinckii GCA_000833105.2
242 Clostridium puniceum GCA_002006345.1
242 Clostridium saccharoperbutylacetonicum N1-4_28HMT_29 GCA_000340885.1
231 Clostridium butyricum GCA_001456065.2
228 Clostridium neonatale GCA_001458595.1
be
202 Lactobacillus gasseri ATCC 33323 = JCM 1131 GCA_000014425.1
172 Lactobacillus hominis DSM 23910 = CRBIP 24.179 GCA_000296835.1
124 Lactobacillus antri DSM 16041 GCA_000160835.1
124 Lactobacillus frumenti DSM 13145 GCA_001436045.1
124 Lactobacillus mucosae LM1 GCA_000248095.3
124 Lactobacillus pontis DSM 8475 GCA_001435345.1
124 Lactobacillus reuteri DSM 20016 GCA_000016825.1
124 Lactobacillus secaliphilus GCA_001437055.1
124 Lactobacillus vaginalis DSM 5837 = ATCC 49540 GCA_000159435.1
bf
253 Streptococcus mutans UA159 GCA_000007465.2
168 Streptococcus sobrinus DSM 20742 = ATCC 33478 GCA_000686605.1
155 Streptococcus macacae NCTC 11558 GCA_000187995.3
bg
248 Clostridium saccharobutylicum DSM 13864 GCA_000473995.1
246 Clostridium beijerinckii GCA_000833105.2
246 Clostridium puniceum GCA_002006345.1
246 Clostridium saccharoperbutylacetonicum N1-4_28HMT_29 GCA_000340885.1
232 Clostridium chromiireducens GCA_002029255.1
bh
236 Clostridium beijerinckii GCA_000833105.2
236 Clostridium puniceum GCA_002006345.1
236 Clostridium saccharoperbutylacetonicum N1-4_28HMT_29 GCA_000340885.1
228 Clostridium saccharobutylicum DSM 13864 GCA_000473995.1
224 Clostridium chromiireducens GCA_002029255.1
bi
212 Streptococcus mutans UA159 GCA_000007465.2
143 Streptococcus cristatus AS 1.3089 GCA_000385925.1
143 Streptococcus gordonii str. Challis substr. CH1 GCA_000017005.1
143 Streptococcus mitis B6 GCA_000027165.1
143 Streptococcus pneumoniae R6 GCA_000007045.1
137 Streptococcus parasanguinis ATCC 15912 GCA_000164675.2
bj
150 Enterococcus faecalis V583 GCA_000007785.1
150 Streptomyces cinnamoneus GCA_001885705.1
137 Enterococcus hirae ATCC 9790 GCA_000271405.2
135 Melissococcus plutonius S1 GCA_000747585.1
bk
257 Bacillus anthracis str. Ames GCA_000007845.1
257 Bacillus anthracis str. Sterne GCA_000008165.1
257 Bacillus cereus ATCC 14579 GCA_000007825.1

257 Bacillus mycoides GCA_000832605.1
257 Bacillus pseudomycoides DSM 12442 GCA_000161455.1
257 Bacillus thuringiensis YBT-1518 GCA_000497525.2
257 _5BBacillus thuringiensis_5D serovar konkukian str. 97-27 GCA_000008505.1
211 Bacillus manliponensis GCA_000712595.1
190 Bacillus cytotoxicus NVH 391-98 GCA_000017425.1
bl
290 Staphylococcus epidermidis ATCC 12228 GCA_000007645.1
286 Staphylococcus lugdunensis HKU09-01 GCA_000025085.1
284 Staphylococcus capitis subsp. capitis GCA_001028645.1
bm
182 Deinococcus radiodurans R1 GCA_000008565.1
124 Deinococcus deserti VCD115 GCA_000020685.1
116 Deinococcus gobiensis I-0 GCA_000252445.1
bn
207 Bacillus anthracis str. Ames GCA_000007845.1
207 Bacillus anthracis str. Sterne GCA_000008165.1
207 Bacillus cereus ATCC 14579 GCA_000007825.1
207 Bacillus mycoides GCA_000832605.1
207 Bacillus pseudomycoides DSM 12442 GCA_000161455.1
207 Bacillus thuringiensis YBT-1518 GCA_000497525.2
207 _5BBacillus thuringiensis_5D serovar konkukian str. 97-27 GCA_000008505.1
176 Bacillus manliponensis GCA_000712595.1
171 Staphylococcus equorum GCA_001432245.1
bo
271 Deinococcus radiodurans R1 GCA_000008565.1
182 Deinococcus deserti VCD115 GCA_000020685.1
179 Deinococcus gobiensis I-0 GCA_000252445.1
bp
269 Deinococcus radiodurans R1 GCA_000008565.1
203 Deinococcus gobiensis I-0 GCA_000252445.1
186 Deinococcus deserti VCD115 GCA_000020685.1
bq
233 Clostridium beijerinckii GCA_000833105.2
233 Clostridium puniceum GCA_002006345.1
233 Clostridium saccharobutylicum DSM 13864 GCA_000473995.1
233 Clostridium saccharoperbutylacetonicum N1-4_28HMT_29 GCA_000340885.1
226 Clostridium butyricum GCA_001456065.2
218 Clostridium chromiireducens GCA_002029255.1
br
101 Deinococcus radiodurans R1 GCA_000008565.1
48 Deinococcus deserti VCD115 GCA_000020685.1
45 Deinococcus gobiensis I-0 GCA_000252445.1
bs
223 Clostridium beijerinckii GCA_000833105.2
223 Clostridium puniceum GCA_002006345.1
223 Clostridium saccharoperbutylacetonicum N1-4_28HMT_29 GCA_000340885.1
215 Clostridium saccharobutylicum DSM 13864 GCA_000473995.1
184 Clostridium chromiireducens GCA_002029255.1
bt
210 Rhodobacter sphaeroides 2.4.1 GCA_000012905.2
210 Rhodobacter sphaeroides ATCC 17025 GCA_000016405.1
191 Pseudorhodobacter ferrugineus DSM 5888 GCA_000420745.1
191 Pseudorhodobacter wandonensis GCA_001202035.1
177 Pseudorhodobacter psychrotolerans GCA_001294535.1
bu
143 Enterococcus faecalis V583 GCA_000007785.1
143 Streptomyces cinnamoneus GCA_001885705.1
113 Enterococcus asini ATCC 700915 GCA_000407365.1
113 Enterococcus canis NBRC 100695 GCA_001544375.1
113 Enterococcus casseliflavus EC20 GCA_000157355.2
113 Enterococcus dispar ATCC 51266 GCA_000406945.1
113 Enterococcus faecium DO GCA_000174395.2
113 Enterococcus gilvus ATCC BAA-350 GCA_000407545.1
113 Enterococcus haemoperoxidus ATCC BAA-382 GCA_000407165.1
113 Enterococcus hermanniensis GCA_001885945.1
113 Enterococcus hirae ATCC 9790 GCA_000271405.2
113 Enterococcus italicus DSM 15952 GCA_000185365.1
113 Enterococcus malodoratus ATCC 43197 GCA_000407185.1
113 Enterococcus mundtii QU 25 GCA_000504125.1
113 Enterococcus pallens ATCC BAA-351 GCA_000407485.1
113 Enterococcus phoeniculicola ATCC BAA-412 GCA_000407505.1
113 Enterococcus pseudoavium NBRC 100491 GCA_001544295.1
113 Enterococcus rivorum GCA_001742285.1
113 Enterococcus saccharolyticus subsp. saccharolyticus ATCC 43076 GCA_000407285.1
113 Enterococcus sulfureus ATCC 49903 GCA_000407605.1

113 Enterococcus thailandicus GCA_001652875.1
113 Granulicatella adiacens ATCC 49175 GCA_000160675.1
113 Melissococcus plutonius S1 GCA_000747585.1
104 Enterococcus aquimarinus GCA_001885765.1
104 Enterococcus cecorum GCA_001318405.1
104 Enterococcus massiliensis GCA_001050095.1
104 Isobaculum melis GCA_900111355.1
104 Macrococcus canis GCA_002119805.1
bv
192 Bifidobacterium adolescentis ATCC 15703 GCA_000010425.1
167 Bifidobacterium dentium JCM 1195 = DSM 20436 GCA_001042595.1
167 Bifidobacterium tsurumiense GCA_000741765.1
162 Bifidobacterium longum NCC2705 GCA_000007525.1
162 Bifidobacterium saguini DSM 23967 GCA_000741715.1
bw
239 Staphylococcus epidermidis ATCC 12228 GCA_000007645.1
236 Megasphaera cerevisiae DSM 20462 GCA_001045675.1
236 Staphylococcus lugdunensis HKU09-01 GCA_000025085.1
236 Staphylococcus pseudintermedius HKU10-03 GCA_000185885.1
236 Staphylococcus warneri SG1 GCA_000332735.1
228 Staphylococcus haemolyticus JCSC1435 GCA_000009865.1
bx
264 Vagococcus penaei GCA_001998885.1
261 Enterococcus asini ATCC 700915 GCA_000407365.1
261 Enterococcus canis NBRC 100695 GCA_001544375.1
261 Enterococcus cecorum GCA_001318405.1
261 Enterococcus columbae DSM 7374 = ATCC 51263 GCA_000406925.1
261 Enterococcus dispar ATCC 51266 GCA_000406945.1
261 Enterococcus faecalis V583 GCA_000007785.1
261 Enterococcus faecium DO GCA_000174395.2
261 Enterococcus haemoperoxidus ATCC BAA-382 GCA_000407165.1
261 Enterococcus hirae ATCC 9790 GCA_000271405.2
261 Enterococcus massiliensis GCA_001050095.1
261 Enterococcus mundtii QU 25 GCA_000504125.1
261 Enterococcus phoeniculicola ATCC BAA-412 GCA_000407505.1
261 Enterococcus rivorum GCA_001742285.1
261 Enterococcus thailandicus GCA_001652875.1
261 Streptomyces cinnamoneus GCA_001885705.1
253 Melissococcus plutonius S1 GCA_000747585.1
by
237 Deinococcus radiodurans R1 GCA_000008565.1
165 Deinococcus gobiensis I-0 GCA_000252445.1
157 Deinococcus deserti VCD115 GCA_000020685.1
bz
205 Bifidobacterium adolescentis ATCC 15703 GCA_000010425.1
199 Bifidobacterium tsurumiense GCA_000741765.1
197 Bifidobacterium dentium JCM 1195 = DSM 20436 GCA_001042595.1
bA
202 Rhodobacter sphaeroides 2.4.1 GCA_000012905.2
187 Rhodobacter sphaeroides ATCC 17025 GCA_000016405.1
175 Gemmobacter megaterium GCA_900156815.1
bB
243 Enterococcus faecalis V583 GCA_000007785.1
243 Streptomyces cinnamoneus GCA_001885705.1
225 Enterococcus asini ATCC 700915 GCA_000407365.1
225 Enterococcus canis NBRC 100695 GCA_001544375.1
225 Enterococcus casseliflavus EC20 GCA_000157355.2
225 Enterococcus dispar ATCC 51266 GCA_000406945.1
225 Enterococcus faecium DO GCA_000174395.2
225 Enterococcus hirae ATCC 9790 GCA_000271405.2
225 Enterococcus massiliensis GCA_001050095.1
225 Enterococcus mundtii QU 25 GCA_000504125.1
225 Enterococcus rivorum GCA_001742285.1
225 Enterococcus saccharolyticus subsp. saccharolyticus ATCC 43076 GCA_000407285.1
217 Enterococcus haemoperoxidus ATCC BAA-382 GCA_000407165.1
217 Enterococcus phoeniculicola ATCC BAA-412 GCA_000407505.1
217 Enterococcus thailandicus GCA_001652875.1
bC
159 Rhodobacter sphaeroides 2.4.1 GCA_000012905.2
140 Defluviimonas alba GCA_001620265.1
125 Haematobacter massiliensis GCA_000740795.1
bD
190 Deinococcus radiodurans R1 GCA_000008565.1
159 Deinococcus deserti VCD115 GCA_000020685.1
157 Deinococcus puniceus GCA_001644565.1
bE

236 Clostridium beijerinckii GCA_000833105.2
236 Clostridium puniceum GCA_002006345.1
236 Clostridium saccharoperbutylacetonicum N1-4_28HMT_29 GCA_000340885.1
228 Clostridium saccharobutylicum DSM 13864 GCA_000473995.1
213 Clostridium chromiireducens GCA_002029255.1
bF
297 Escherichia coli IAI39 GCA_000026345.1
297 Escherichia coli O104_3AH4 str. 2011C-3493 GCA_000299455.1
297 Escherichia coli O157_3AH7 str. Sakai GCA_000008865.1
297 Escherichia coli O83_3AH1 str. NRG 857C GCA_000183345.1
297 Escherichia coli UMN026 GCA_000026325.2
297 Escherichia coli str. K-12 substr. MG1655 GCA_000005845.2
297 Shigella dysenteriae Sd197 GCA_000012005.1
297 Shigella flexneri 2a str. 301 GCA_000006925.2
297 Tumebacillus flagellatus GCA_000714935.1
278 Erwinia iniecta GCA_001267535.1
252 Cronobacter sakazakii GCA_000982825.1
252 Erwinia toletana DAPP-PG 735 GCA_000336255.1
252 Pantoea agglomerans GCA_001709315.1
252 Pantoea ananatis LMG 20103 GCA_000025405.2
252 Pantoea dispersa EGD-AAK13 GCA_000465555.2
252 Shimwellia blattae DSM 4481 = NBRC 105725 GCA_000262305.1
bG
182 Escherichia coli IAI39 GCA_000026345.1
182 Escherichia coli O104_3AH4 str. 2011C-3493 GCA_000299455.1
182 Escherichia coli O157_3AH7 str. Sakai GCA_000008865.1
182 Escherichia coli O83_3AH1 str. NRG 857C GCA_000183345.1
182 Escherichia coli UMN026 GCA_000026325.2
182 Escherichia coli str. K-12 substr. MG1655 GCA_000005845.2
182 Shigella flexneri 2a str. 301 GCA_000006925.2
182 Tumebacillus flagellatus GCA_000714935.1
168 Shigella dysenteriae Sd197 GCA_000012005.1
154 Cronobacter sakazakii GCA_000982825.1
154 Erwinia amylovora CFBP1430 GCA_000091565.1
154 Erwinia iniecta GCA_001267535.1
154 Erwinia toletana DAPP-PG 735 GCA_000336255.1
154 Kosakonia cowanii GCA_001975225.1
154 Kosakonia sacchari SP1 GCA_000300455.4
154 Pantoea ananatis LMG 20103 GCA_000025405.2
154 Pantoea dispersa EGD-AAK13 GCA_000465555.2
154 Plautia stali symbiont GCA_000180175.2
154 Pluralibacter gergoviae GCA_000757785.1
154 Shimwellia blattae DSM 4481 = NBRC 105725 GCA_000262305.1
bH
242 Bacillus anthracis str. Ames GCA_000007845.1
242 Bacillus anthracis str. Sterne GCA_000008165.1
242 Bacillus cereus ATCC 14579 GCA_000007825.1
242 Bacillus mycoides GCA_000832605.1
242 Bacillus pseudomycoides DSM 12442 GCA_000161455.1
242 Bacillus thuringiensis YBT-1518 GCA_000497525.2
242 _5BBacillus thuringiensis_5D serovar konkukian str. 97-27 GCA_000008505.1
188 Massilibacterium senegalense GCA_001375675.1
185 Staphylococcus pettenkoferi GCA_002208805.1
bI
251 Streptococcus mutans UA159 GCA_000007465.2
196 Streptococcus ferus DSM 20646 GCA_000372425.1
183 Streptococcus macacae NCTC 11558 GCA_000187995.3
bJ
225 Clostridium beijerinckii GCA_000833105.2
225 Clostridium saccharoperbutylacetonicum N1-4_28HMT_29 GCA_000340885.1
221 Clostridium puniceum GCA_002006345.1
221 Clostridium saccharobutylicum DSM 13864 GCA_000473995.1
206 Clostridium chromiireducens GCA_002029255.1
bK
86 Rhodobacter sphaeroides 2.4.1 GCA_000012905.2
71 Defluviimonas alba GCA_001620265.1
71 Haematobacter massiliensis GCA_000740795.1
70 Albimonas donghaensis GCA_900106695.1
70 Thioclava dalianensis GCA_000715505.1
70 Thioclava indica GCA_000714545.1
bL
242 Deinococcus radiodurans R1 GCA_000008565.1
167 Deinococcus deserti VCD115 GCA_000020685.1
153 Deinococcus gobiensis I-0 GCA_000252445.1
bM
243 Deinococcus radiodurans R1 GCA_000008565.1

180 Deinococcus puniceus GCA_001644565.1
176 Deinococcus gobiensis I-0 GCA_000252445.1
bN
251 Lactobacillus gasseri ATCC 33323 = JCM 1131 GCA_000014425.1
230 Lactobacillus hominis DSM 23910 = CRBIP 24.179 GCA_000296835.1
161 Lactobacillus acidophilus NCFM GCA_000011985.1
161 Lactobacillus crispatus ST1 GCA_000091765.1
161 Lactobacillus gallinarum GCA_001314245.2
161 Lactobacillus hamsteri DSM 5661 = JCM 6256 GCA_000615445.1
161 Lactobacillus psittaci DSM 15354 GCA_000425905.1
bO
185 Rhodobacter sphaeroides 2.4.1 GCA_000012905.2
170 Rhodobacter sphaeroides ATCC 17025 GCA_000016405.1
165 Pseudorhodobacter ferrugineus DSM 5888 GCA_000420745.1
165 Pseudorhodobacter wandonensis GCA_001202035.1
bP
239 Lactobacillus gasseri ATCC 33323 = JCM 1131 GCA_000014425.1
223 Lactobacillus hominis DSM 23910 = CRBIP 24.179 GCA_000296835.1
195 Lactobacillus amylovorus GCA_000191545.1
195 Lactobacillus crispatus ST1 GCA_000091765.1
195 Lactobacillus kalixensis DSM 16043 GCA_001434335.1
195 Lactobacillus psittaci DSM 15354 GCA_000425905.1
bQ
181 Escherichia coli IAI39 GCA_000026345.1
181 Escherichia coli O104_3AH4 str. 2011C-3493 GCA_000299455.1
181 Escherichia coli O157_3AH7 str. Sakai GCA_000008865.1
181 Escherichia coli O83_3AH1 str. NRG 857C GCA_000183345.1
181 Escherichia coli UMN026 GCA_000026325.2
181 Escherichia coli str. K-12 substr. MG1655 GCA_000005845.2
181 Shigella dysenteriae Sd197 GCA_000012005.1
181 Shigella flexneri 2a str. 301 GCA_000006925.2
181 Tumebacillus flagellatus GCA_000714935.1
167 Erwinia iniecta GCA_001267535.1
167 Rosenbergiella nectarea GCA_900111105.1
165 Cedecea neteri GCA_000757825.1
bR
224 Clostridium neonatale GCA_001458595.1
219 Clostridium beijerinckii GCA_000833105.2
219 Clostridium puniceum GCA_002006345.1
219 Clostridium saccharobutylicum DSM 13864 GCA_000473995.1
219 Clostridium saccharoperbutylacetonicum N1-4_28HMT_29 GCA_000340885.1
212 Clostridium butyricum GCA_001456065.2
bS
284 Lactobacillus gasseri ATCC 33323 = JCM 1131 GCA_000014425.1
260 Lactobacillus hominis DSM 23910 = CRBIP 24.179 GCA_000296835.1
193 Lactobacillus psittaci DSM 15354 GCA_000425905.1
bT
197 Escherichia coli IAI39 GCA_000026345.1
197 Escherichia coli O104_3AH4 str. 2011C-3493 GCA_000299455.1
197 Escherichia coli O157_3AH7 str. Sakai GCA_000008865.1
197 Escherichia coli UMN026 GCA_000026325.2
197 Escherichia coli str. K-12 substr. MG1655 GCA_000005845.2
197 Shigella flexneri 2a str. 301 GCA_000006925.2
197 Tumebacillus flagellatus GCA_000714935.1
196 Escherichia coli O83_3AH1 str. NRG 857C GCA_000183345.1
186 Brenneria goodwinii GCA_001049335.1
186 Erwinia iniecta GCA_001267535.1
186 Erwinia toletana DAPP-PG 735 GCA_000336255.1
186 Kosakonia cowanii GCA_001975225.1
186 Kosakonia sacchari SP1 GCA_000300455.4
186 Shimwellia blattae DSM 4481 = NBRC 105725 GCA_000262305.1
bU
196 Bifidobacterium adolescentis ATCC 15703 GCA_000010425.1
193 Bifidobacterium dentium JCM 1195 = DSM 20436 GCA_001042595.1
166 Bifidobacterium callitrichos DSM 23973 GCA_000741175.1
bV
191 Bacillus anthracis str. Ames GCA_000007845.1
191 Bacillus anthracis str. Sterne GCA_000008165.1
191 Bacillus cereus ATCC 14579 GCA_000007825.1
191 Bacillus mycoides GCA_000832605.1
191 Bacillus pseudomycoides DSM 12442 GCA_000161455.1
191 Bacillus thuringiensis YBT-1518 GCA_000497525.2
191 _5BBacillus thuringiensis_5D serovar konkukian str. 97-27 GCA_000008505.1
130 Bacillus flexus GCA_002024265.1
130 Bacillus megaterium NBRC 15308 = ATCC 14581 GCA_000832985.1
130 Thalassospira mesophila GCA_002115755.1

124 Bacillus marisflavi GCA_001274775.1
bW
295 Deinococcus radiodurans R1 GCA_000008565.1
224 Deinococcus gobiensis I-0 GCA_000252445.1
196 Deinococcus deserti VCD115 GCA_000020685.1
bX
192 Bifidobacterium adolescentis ATCC 15703 GCA_000010425.1
169 Bifidobacterium asteroides PRL2011 GCA_000304215.1
168 Bifidobacterium dentium JCM 1195 = DSM 20436 GCA_001042595.1
bY
81 Bacillus anthracis str. Ames GCA_000007845.1
81 Bacillus anthracis str. Sterne GCA_000008165.1
81 Bacillus cereus ATCC 14579 GCA_000007825.1
81 Bacillus flexus GCA_002024265.1
81 Bacillus megaterium NBRC 15308 = ATCC 14581 GCA_000832985.1
81 Bacillus mycoides GCA_000832605.1
81 Bacillus pseudomycoides DSM 12442 GCA_000161455.1
81 Bacillus thuringiensis YBT-1518 GCA_000497525.2
81 _5BBacillus thuringiensis_5D serovar konkukian str. 97-27 GCA_000008505.1
66 Bhargavaea cecembensis DSE10 GCA_000348905.1
66 Bhargavaea cecembensis GCA_000826125.2
66 Edaphobacillus lindanitolerans GCA_900156305.1
66 Listeria floridensis FSL S10-1187 GCA_000525875.1
66 Planococcus antarcticus DSM 14505 GCA_001687565.2
66 Planococcus kocurii GCA_001465835.2
66 Planococcus maritimus GCA_001999905.1
66 Planococcus rifietoensis GCA_001465795.2
65 Anaerobacillus macyae GCA_001039475.1
65 Anoxybacillus amylolyticus GCA_001634285.1
65 Anoxybacillus gonensis GCA_001187595.1
65 Bacillus abyssalis GCA_002019595.1
65 Bacillus acidicola GCA_001636425.1
65 Bacillus akibai JCM 9157 GCA_000513135.1
65 Bacillus alkalinitrilicus GCA_002019605.1
65 Bacillus aquimaris TF-12 GCA_001648555.1
65 Bacillus badius GCA_001630115.1
65 Bacillus cihuensis GCA_000504145.1
65 Bacillus circulans NBRC 13626 GCA_001591585.1
65 Bacillus coahuilensis m4-4 GCA_000171615.1
65 Bacillus cytotoxicus NVH 391-98 GCA_000017425.1
65 Bacillus eiseniae GCA_001636325.1
65 Bacillus fastidiosus NBRC 101226 GCA_001591625.1
65 Bacillus gottheilii GCA_001636345.1
65 Bacillus halosaccharovorans GCA_002019635.1
65 Bacillus humi GCA_001439915.1
65 Bacillus infantis NRRL B-14911 GCA_000473245.1
65 Bacillus koreensis GCA_001274935.1
65 Bacillus krulwichiae GCA_002109385.1
65 Bacillus ligniniphilus GCA_000334155.1
65 Bacillus litoralis GCA_001654695.1
65 Bacillus manliponensis GCA_000712595.1
65 Bacillus marisflavi GCA_001274775.1
65 Bacillus methanolicus MGA3 GCA_000724485.1
65 Bacillus niacini NBRC 15566 GCA_001591505.1
65 Bacillus pseudofirmus OF4 GCA_000005825.2
65 Bacillus salsus GCA_900104555.1
65 Bacillus shackletonii GCA_001420715.1
65 Bacillus simplex GCA_001578185.1
65 Bacillus sinesaloumensis GCA_900156865.1
65 Bacillus smithii GCA_001050115.1
65 Bacillus sporothermodurans GCA_001587375.1
65 Bacillus testis GCA_001243895.1
65 Bacillus timonensis GCA_000285535.1
65 Bacillus tuaregi GCA_900104575.1
65 Bacillus vietnamensis NBRC 101237 GCA_001591825.1
65 Bacillus weihaiensis GCA_001889165.1
65 Carnobacterium maltaromaticum LMA28 GCA_000317975.2
65 Domibacillus antri GCA_001936625.1
65 Gemella asaccharolytica GCA_001553005.1
65 Granulicatella balaenopterae GCA_900111135.1
65 Halobacillus halophilus DSM 2266 GCA_000284515.1
65 Halobacillus hunanensis GCA_900166655.1
65 Halobacillus mangrovi GCA_002097535.1
65 Halobacillus massiliensis GCA_900166625.1
65 Listeria monocytogenes EGD-e GCA_000196035.1
65 Massilibacterium senegalense GCA_001375675.1

65 Quasibacillus thermotolerans GCA_000812025.2
65 Staphylococcus equorum GCA_001432245.1
65 Thalassobacillus devorans MSP14 GCA_000496835.1
65 Virgibacillus halodenitrificans GCA_001878675.1
bZ
244 Clostridium beijerinckii GCA_000833105.2
244 Clostridium puniceum GCA_002006345.1
244 Clostridium saccharoperbutylacetonicum N1-4_28HMT_29 GCA_000340885.1
236 Clostridium saccharobutylicum DSM 13864 GCA_000473995.1
203 Clostridium chromiireducens GCA_002029255.1
c0
244 Escherichia coli IAI39 GCA_000026345.1
244 Escherichia coli O104_3AH4 str. 2011C-3493 GCA_000299455.1
244 Escherichia coli O157_3AH7 str. Sakai GCA_000008865.1
244 Escherichia coli O83_3AH1 str. NRG 857C GCA_000183345.1
244 Escherichia coli UMN026 GCA_000026325.2
244 Escherichia coli str. K-12 substr. MG1655 GCA_000005845.2
244 Shigella flexneri 2a str. 301 GCA_000006925.2
244 Tumebacillus flagellatus GCA_000714935.1
235 Shigella dysenteriae Sd197 GCA_000012005.1
223 Erwinia iniecta GCA_001267535.1
c1
249 Deinococcus radiodurans R1 GCA_000008565.1
173 Deinococcus deserti VCD115 GCA_000020685.1
167 Deinococcus gobiensis I-0 GCA_000252445.1
c2
159 Deinococcus radiodurans R1 GCA_000008565.1
102 Deinococcus deserti VCD115 GCA_000020685.1
102 Deinococcus gobiensis I-0 GCA_000252445.1
95 Deinococcus maricopensis DSM 21211 GCA_000186385.1
c3
230 Bifidobacterium adolescentis ATCC 15703 GCA_000010425.1
219 Bifidobacterium dentium JCM 1195 = DSM 20436 GCA_001042595.1
209 Bifidobacterium callitrichos DSM 23973 GCA_000741175.1
c4
173 Enterococcus faecalis V583 GCA_000007785.1
173 Streptomyces cinnamoneus GCA_001885705.1
161 Enterococcus casseliflavus EC20 GCA_000157355.2
161 Enterococcus saccharolyticus subsp. saccharolyticus ATCC 43076 GCA_000407285.1
155 Enterococcus asini ATCC 700915 GCA_000407365.1
155 Enterococcus canis NBRC 100695 GCA_001544375.1
155 Enterococcus dispar ATCC 51266 GCA_000406945.1
155 Enterococcus faecium DO GCA_000174395.2
155 Enterococcus gilvus ATCC BAA-350 GCA_000407545.1
155 Enterococcus haemoperoxidus ATCC BAA-382 GCA_000407165.1
155 Enterococcus hermanniensis GCA_001885945.1
155 Enterococcus hirae ATCC 9790 GCA_000271405.2
155 Enterococcus italicus DSM 15952 GCA_000185365.1
155 Enterococcus malodoratus ATCC 43197 GCA_000407185.1
155 Enterococcus massiliensis GCA_001050095.1
155 Enterococcus mundtii QU 25 GCA_000504125.1
155 Enterococcus pallens ATCC BAA-351 GCA_000407485.1
155 Enterococcus phoeniculicola ATCC BAA-412 GCA_000407505.1
155 Enterococcus pseudoavium NBRC 100491 GCA_001544295.1
155 Enterococcus rivorum GCA_001742285.1
155 Enterococcus sulfureus ATCC 49903 GCA_000407605.1
155 Enterococcus thailandicus GCA_001652875.1
155 Vagococcus fluvialis bH819 GCA_900163795.1
155 Vagococcus lutrae LBD1 GCA_000498295.1
155 Vagococcus penaei GCA_001998885.1
c5
254 Clostridium beijerinckii GCA_000833105.2
254 Clostridium saccharoperbutylacetonicum N1-4_28HMT_29 GCA_000340885.1
247 Clostridium puniceum GCA_002006345.1
246 Clostridium saccharobutylicum DSM 13864 GCA_000473995.1
c6
121 Bifidobacterium adolescentis ATCC 15703 GCA_000010425.1
121 Bifidobacterium thermophilum GCA_000741495.1
112 Bifidobacterium asteroides PRL2011 GCA_000304215.1
106 Bifidobacterium thermophilum RBL67 GCA_000347695.1
c7
98 Staphylococcus condimenti GCA_001618885.1
98 Staphylococcus simulans GCA_001559115.1
97 Megasphaera cerevisiae DSM 20462 GCA_001045675.1
97 Staphylococcus aureus subsp. aureus NCTC 8325 GCA_000013425.1
97 Staphylococcus cohnii subsp. cohnii GCA_000972575.1

97 Staphylococcus epidermidis ATCC 12228 GCA_000007645.1
97 Staphylococcus haemolyticus JCSC1435 GCA_000009865.1
97 Staphylococcus hominis subsp. hominis C80 GCA_000183685.1
97 Staphylococcus lutrae GCA_002101335.1
97 Staphylococcus saprophyticus GCA_001074355.1
97 Staphylococcus sciuri GCA_002209165.1
97 Staphylococcus simiae CCM 7213 GCA_000235645.2
97 Staphylococcus warneri SG1 GCA_000332735.1
92 Staphylococcus capitis subsp. capitis GCA_001028645.1
c8
215 Streptococcus mutans UA159 GCA_000007465.2
117 Streptococcus marimammalium DSM 18627 GCA_000380045.1
103 Streptococcus salivarius GCA_000785515.1
103 Streptococcus thermophilus JIM 8232 GCA_000253395.1
c9
195 Bifidobacterium adolescentis ATCC 15703 GCA_000010425.1
163 Bifidobacterium thermophilum GCA_000741495.1
161 Bifidobacterium callitrichos DSM 23973 GCA_000741175.1
ca
268 Enterococcus faecalis V583 GCA_000007785.1
268 Streptomyces cinnamoneus GCA_001885705.1
237 Enterococcus hirae ATCC 9790 GCA_000271405.2
234 Enterococcus canis NBRC 100695 GCA_001544375.1
234 Enterococcus casseliflavus EC20 GCA_000157355.2
234 Enterococcus dispar ATCC 51266 GCA_000406945.1
234 Enterococcus faecium DO GCA_000174395.2
234 Enterococcus haemoperoxidus ATCC BAA-382 GCA_000407165.1
234 Enterococcus mundtii QU 25 GCA_000504125.1
234 Enterococcus phoeniculicola ATCC BAA-412 GCA_000407505.1
234 Enterococcus rivorum GCA_001742285.1
234 Enterococcus saccharolyticus subsp. saccharolyticus ATCC 43076 GCA_000407285.1
234 Enterococcus thailandicus GCA_001652875.1
cb
226 Bifidobacterium adolescentis ATCC 15703 GCA_000010425.1
212 Bifidobacterium breve DSM 20213 = JCM 1192 GCA_001025175.1
204 Bifidobacterium longum NCC2705 GCA_000007525.1
cc
285 Deinococcus radiodurans R1 GCA_000008565.1
219 Deinococcus gobiensis I-0 GCA_000252445.1
207 Deinococcus deserti VCD115 GCA_000020685.1
cd
302 Bifidobacterium adolescentis ATCC 15703 GCA_000010425.1
263 Bifidobacterium asteroides PRL2011 GCA_000304215.1
249 Bifidobacterium tsurumiense GCA_000741765.1
ce
181 Lactobacillus gasseri ATCC 33323 = JCM 1131 GCA_000014425.1
168 Lactobacillus hominis DSM 23910 = CRBIP 24.179 GCA_000296835.1
140 Lactobacillus gastricus DSM 16045 GCA_001434365.1
140 Lactobacillus jensenii GCA_001936235.1
140 Lactobacillus psittaci DSM 15354 GCA_000425905.1
cf
179 Klebsiella pneumoniae subsp. pneumoniae HS11286 GCA_000240185.2
178 Cronobacter sakazakii GCA_000982825.1
178 Erwinia iniecta GCA_001267535.1
178 Erwinia toletana DAPP-PG 735 GCA_000336255.1
178 Escherichia coli IAI39 GCA_000026345.1
178 Escherichia coli O104_3AH4 str. 2011C-3493 GCA_000299455.1
178 Escherichia coli O157_3AH7 str. Sakai GCA_000008865.1
178 Escherichia coli O83_3AH1 str. NRG 857C GCA_000183345.1
178 Escherichia coli UMN026 GCA_000026325.2
178 Escherichia coli str. K-12 substr. MG1655 GCA_000005845.2
178 Shigella dysenteriae Sd197 GCA_000012005.1
178 Shigella flexneri 2a str. 301 GCA_000006925.2
178 Tumebacillus flagellatus GCA_000714935.1
176 Erwinia billingiae Eb661 GCA_000196615.1
176 Erwinia gerundensis GCA_001517405.1
176 Pantoea agglomerans GCA_001709315.1
176 Pantoea ananatis LMG 20103 GCA_000025405.2
176 Pantoea dispersa EGD-AAK13 GCA_000465555.2
176 Shimwellia blattae DSM 4481 = NBRC 105725 GCA_000262305.1
cg
168 Clostridium beijerinckii GCA_000833105.2
168 Clostridium saccharobutylicum DSM 13864 GCA_000473995.1
168 Clostridium saccharoperbutylacetonicum N1-4_28HMT_29 GCA_000340885.1
161 Clostridium puniceum GCA_002006345.1
151 Clostridium amylolyticum GCA_900142075.1

151 Clostridium intestinale URNW GCA_000469625.2
151 Clostridium polynesiense GCA_000820705.1
ch
293 Deinococcus radiodurans R1 GCA_000008565.1
194 Deinococcus deserti VCD115 GCA_000020685.1
178 Deinococcus puniceus GCA_001644565.1
ci
261 Bacillus thuringiensis YBT-1518 GCA_000497525.2
253 Bacillus anthracis str. Ames GCA_000007845.1
253 Bacillus anthracis str. Sterne GCA_000008165.1
253 Bacillus cereus ATCC 14579 GCA_000007825.1
253 Bacillus mycoides GCA_000832605.1
253 Bacillus pseudomycoides DSM 12442 GCA_000161455.1
253 _5BBacillus thuringiensis_5D serovar konkukian str. 97-27 GCA_000008505.1
196 Bacillus cytotoxicus NVH 391-98 GCA_000017425.1
cj
225 Enterococcus faecalis V583 GCA_000007785.1
225 Streptomyces cinnamoneus GCA_001885705.1
212 Enterococcus hirae ATCC 9790 GCA_000271405.2
209 Enterococcus dispar ATCC 51266 GCA_000406945.1
ck
155 Deinococcus radiodurans R1 GCA_000008565.1
95 Deinococcus frigens DSM 12807 GCA_000701425.1
93 Deinococcus puniceus GCA_001644565.1
cl
180 Bifidobacterium adolescentis ATCC 15703 GCA_000010425.1
166 Bifidobacterium thermophilum GCA_000741495.1
152 Bifidobacterium breve DSM 20213 = JCM 1192 GCA_001025175.1
cm
232 Deinococcus radiodurans R1 GCA_000008565.1
159 Deinococcus deserti VCD115 GCA_000020685.1
158 Deinococcus puniceus GCA_001644565.1
cn
81 Bifidobacterium asteroides PRL2011 GCA_000304215.1
81 Bifidobacterium callitrichos DSM 23973 GCA_000741175.1
81 Bifidobacterium coryneforme GCA_000737865.1
79 Bifidobacterium adolescentis ATCC 15703 GCA_000010425.1
79 Deinococcus radiodurans R1 GCA_000008565.1
76 Bifidobacterium minimum GCA_000741645.1
76 Bifidobacterium mongoliense DSM 21395 GCA_000741285.1
co
204 Clostridium beijerinckii GCA_000833105.2
204 Clostridium saccharobutylicum DSM 13864 GCA_000473995.1
204 Clostridium saccharoperbutylacetonicum N1-4_28HMT_29 GCA_000340885.1
189 Clostridium puniceum GCA_002006345.1
184 Clostridium butyricum GCA_001456065.2
cp
240 Bifidobacterium adolescentis ATCC 15703 GCA_000010425.1
221 Bifidobacterium tsurumiense GCA_000741765.1
220 Bifidobacterium callitrichos DSM 23973 GCA_000741175.1
220 Bifidobacterium gallicum DSM 20093 = LMG 11596 GCA_000741205.1
cq
141 Escherichia coli IAI39 GCA_000026345.1
141 Escherichia coli O104_3AH4 str. 2011C-3493 GCA_000299455.1
141 Escherichia coli O157_3AH7 str. Sakai GCA_000008865.1
141 Escherichia coli O83_3AH1 str. NRG 857C GCA_000183345.1
141 Escherichia coli UMN026 GCA_000026325.2
141 Escherichia coli str. K-12 substr. MG1655 GCA_000005845.2
141 Shigella dysenteriae Sd197 GCA_000012005.1
141 Shigella flexneri 2a str. 301 GCA_000006925.2
141 Tumebacillus flagellatus GCA_000714935.1
103 Pantoea agglomerans GCA_001709315.1
102 Cronobacter sakazakii GCA_000982825.1
102 Enterobacter cloacae subsp. cloacae ATCC 13047 GCA_000025565.1
102 Enterobacter hormaechei subsp. steigerwaltii GCA_001729725.1
102 Enterobacter kobei GCA_900185885.1
102 Erwinia amylovora CFBP1430 GCA_000091565.1
102 Erwinia iniecta GCA_001267535.1
102 Erwinia toletana DAPP-PG 735 GCA_000336255.1
102 Klebsiella oxytoca GCA_001022195.1
102 Kosakonia cowanii GCA_001975225.1
102 Kosakonia sacchari SP1 GCA_000300455.4
102 Pantoea alhagi GCA_002101395.1
102 Pantoea ananatis LMG 20103 GCA_000025405.2
102 Pantoea dispersa EGD-AAK13 GCA_000465555.2
102 Pantoea rwandensis GCA_000759475.1

102 Pantoea septica GCA_002095575.1
102 Photorhabdus temperata subsp. thracensis GCA_001010285.1
102 Proteus mirabilis HI4320 GCA_000069965.1
102 Pseudescherichia vulneris NBRC 102420 GCA_000759795.1
102 Salmonella enterica subsp. enterica serovar Typhi str. CT18 GCA_000195995.1
102 Salmonella enterica subsp. enterica serovar Typhimurium str. LT2 GCA_000006945.2
102 Shimwellia blattae DSM 4481 = NBRC 105725 GCA_000262305.1
102 Trabulsiella odontotermitis GCA_001297765.1
102 Xenorhabdus bovienii SS-2004 GCA_000027225.1
102 Xenorhabdus cabanillasii JM26 GCA_000531755.1
102 Xenorhabdus doucetiae GCA_000968195.1
102 Xenorhabdus hominickii GCA_001721185.1
102 Xenorhabdus innexi GCA_900155355.1
102 Xenorhabdus japonica GCA_900115195.1
102 Xenorhabdus koppenhoeferi GCA_900116635.1
102 Xenorhabdus nematophila AN6_2F1 GCA_000953355.1
cr
302 Lactobacillus gasseri ATCC 33323 = JCM 1131 GCA_000014425.1
287 Lactobacillus hominis DSM 23910 = CRBIP 24.179 GCA_000296835.1
219 Lactobacillus psittaci DSM 15354 GCA_000425905.1
cs
213 Escherichia coli IAI39 GCA_000026345.1
213 Escherichia coli O104_3AH4 str. 2011C-3493 GCA_000299455.1
213 Escherichia coli O157_3AH7 str. Sakai GCA_000008865.1
213 Escherichia coli O83_3AH1 str. NRG 857C GCA_000183345.1
213 Escherichia coli UMN026 GCA_000026325.2
213 Escherichia coli str. K-12 substr. MG1655 GCA_000005845.2
213 Shigella dysenteriae Sd197 GCA_000012005.1
213 Shigella flexneri 2a str. 301 GCA_000006925.2
213 Tumebacillus flagellatus GCA_000714935.1
179 Erwinia iniecta GCA_001267535.1
179 Rosenbergiella nectarea GCA_900111105.1
179 Serratia marcescens subsp. marcescens Db11 GCA_000513215.1
177 Cedecea neteri GCA_000757825.1
ct
254 Clostridium saccharobutylicum DSM 13864 GCA_000473995.1
252 Clostridium beijerinckii GCA_000833105.2
252 Clostridium puniceum GCA_002006345.1
252 Clostridium saccharoperbutylacetonicum N1-4_28HMT_29 GCA_000340885.1
243 Clostridium chromiireducens GCA_002029255.1
cu
150 Escherichia coli IAI39 GCA_000026345.1
150 Escherichia coli O104_3AH4 str. 2011C-3493 GCA_000299455.1
150 Escherichia coli O157_3AH7 str. Sakai GCA_000008865.1
150 Escherichia coli O83_3AH1 str. NRG 857C GCA_000183345.1
150 Escherichia coli UMN026 GCA_000026325.2
150 Escherichia coli str. K-12 substr. MG1655 GCA_000005845.2
150 Shigella flexneri 2a str. 301 GCA_000006925.2
150 Tumebacillus flagellatus GCA_000714935.1
146 Erwinia iniecta GCA_001267535.1
141 Shigella dysenteriae Sd197 GCA_000012005.1
cv
131 Gemmobacter megaterium GCA_900156815.1
131 Rhodobacter sphaeroides 2.4.1 GCA_000012905.2
122 Pseudorhodobacter ferrugineus DSM 5888 GCA_000420745.1
122 Pseudorhodobacter wandonensis GCA_001202035.1
113 Gemmobacter aquatilis GCA_900110025.1
113 Pseudorhodobacter psychrotolerans GCA_001294535.1
cw
208 Clostridium saccharobutylicum DSM 13864 GCA_000473995.1
193 Clostridium beijerinckii GCA_000833105.2
193 Clostridium chromiireducens GCA_002029255.1
193 Clostridium puniceum GCA_002006345.1
193 Clostridium saccharoperbutylacetonicum N1-4_28HMT_29 GCA_000340885.1
184 Clostridium butyricum GCA_001456065.2
cx
169 Escherichia coli IAI39 GCA_000026345.1
169 Escherichia coli O104_3AH4 str. 2011C-3493 GCA_000299455.1
169 Escherichia coli O157_3AH7 str. Sakai GCA_000008865.1
169 Escherichia coli O83_3AH1 str. NRG 857C GCA_000183345.1
169 Escherichia coli UMN026 GCA_000026325.2
169 Escherichia coli str. K-12 substr. MG1655 GCA_000005845.2
169 Shigella dysenteriae Sd197 GCA_000012005.1
169 Shigella flexneri 2a str. 301 GCA_000006925.2
169 Tumebacillus flagellatus GCA_000714935.1
143 Enterobacter hormaechei subsp. steigerwaltii GCA_001729725.1

143 Erwinia iniecta GCA_001267535.1
143 Erwinia toletana DAPP-PG 735 GCA_000336255.1
143 Klebsiella oxytoca GCA_001022195.1
143 Kosakonia cowanii GCA_001975225.1
143 Kosakonia sacchari SP1 GCA_000300455.4
143 Pantoea alhagi GCA_002101395.1
143 Pseudescherichia vulneris NBRC 102420 GCA_000759795.1
137 Pluralibacter gergoviae GCA_000757785.1
cy
283 Clostridium beijerinckii GCA_000833105.2
283 Clostridium saccharobutylicum DSM 13864 GCA_000473995.1
283 Clostridium saccharoperbutylacetonicum N1-4_28HMT_29 GCA_000340885.1
277 Clostridium puniceum GCA_002006345.1
240 Clostridium chromiireducens GCA_002029255.1
cz
205 Clostridium beijerinckii GCA_000833105.2
205 Clostridium saccharobutylicum DSM 13864 GCA_000473995.1
205 Clostridium saccharoperbutylacetonicum N1-4_28HMT_29 GCA_000340885.1
199 Clostridium puniceum GCA_002006345.1
194 Clostridium butyricum GCA_001456065.2
cA
248 Rhodobacter sphaeroides 2.4.1 GCA_000012905.2
211 Pseudorhodobacter psychrotolerans GCA_001294535.1
209 Gemmobacter megaterium GCA_900156815.1
209 Rhodobacter sphaeroides ATCC 17025 GCA_000016405.1
cB
231 Rhodobacter sphaeroides 2.4.1 GCA_000012905.2
196 Rhodobacter sphaeroides ATCC 17025 GCA_000016405.1
188 Thioclava dalianensis GCA_000715505.1
188 Thioclava indica GCA_000714545.1
cC
115 Deinococcus radiodurans R1 GCA_000008565.1
75 Deinococcus gobiensis I-0 GCA_000252445.1
59 Deinococcus puniceus GCA_001644565.1
cD
273 Staphylococcus capitis subsp. capitis GCA_001028645.1
273 Staphylococcus epidermidis ATCC 12228 GCA_000007645.1
273 Staphylococcus haemolyticus JCSC1435 GCA_000009865.1
262 Staphylococcus hominis subsp. hominis C80 GCA_000183685.1
262 Staphylococcus lugdunensis HKU09-01 GCA_000025085.1
254 Staphylococcus simulans GCA_001559115.1
cE
276 Clostridium beijerinckii GCA_000833105.2
276 Clostridium saccharobutylicum DSM 13864 GCA_000473995.1
276 Clostridium saccharoperbutylacetonicum N1-4_28HMT_29 GCA_000340885.1
269 Clostridium puniceum GCA_002006345.1
254 Clostridium chromiireducens GCA_002029255.1
cF
303 Escherichia coli IAI39 GCA_000026345.1
303 Escherichia coli O104_3AH4 str. 2011C-3493 GCA_000299455.1
303 Escherichia coli O157_3AH7 str. Sakai GCA_000008865.1
303 Escherichia coli UMN026 GCA_000026325.2
303 Escherichia coli str. K-12 substr. MG1655 GCA_000005845.2
303 Shigella flexneri 2a str. 301 GCA_000006925.2
303 Tumebacillus flagellatus GCA_000714935.1
302 Escherichia coli O83_3AH1 str. NRG 857C GCA_000183345.1
294 Shigella dysenteriae Sd197 GCA_000012005.1
cG
195 Deinococcus radiodurans R1 GCA_000008565.1
101 Deinococcus gobiensis I-0 GCA_000252445.1
93 Deinococcus marmoris DSM 12784 GCA_000701405.1
93 Deinococcus swuensis GCA_000800395.1
cH
211 Rhodobacter sphaeroides 2.4.1 GCA_000012905.2
189 Pseudorhodobacter psychrotolerans GCA_001294535.1
183 Rhodobacter sphaeroides ATCC 17025 GCA_000016405.1
cI
228 Clostridium beijerinckii GCA_000833105.2
228 Clostridium puniceum GCA_002006345.1
228 Clostridium saccharoperbutylacetonicum N1-4_28HMT_29 GCA_000340885.1
213 Clostridium saccharobutylicum DSM 13864 GCA_000473995.1
190 Clostridium chromiireducens GCA_002029255.1
cJ
163 Rhodobacter sphaeroides 2.4.1 GCA_000012905.2
136 Defluviimonas alba GCA_001620265.1
131 Haematobacter massiliensis GCA_000740795.1

131 Pseudoruegeria lutimaris GCA_900099935.1
131 Tateyamaria omphalii GCA_001969365.1
cK
114 Serratia symbiotica GCA_000821185.1
110 Pantoea agglomerans GCA_001709315.1
110 Yersinia ruckeri GCA_000964565.1
109 Cronobacter sakazakii GCA_000982825.1
109 Erwinia gerundensis GCA_001517405.1
109 Erwinia iniecta GCA_001267535.1
109 Erwinia toletana DAPP-PG 735 GCA_000336255.1
109 Escherichia coli IAI39 GCA_000026345.1
109 Escherichia coli O104_3AH4 str. 2011C-3493 GCA_000299455.1
109 Escherichia coli O157_3AH7 str. Sakai GCA_000008865.1
109 Escherichia coli O83_3AH1 str. NRG 857C GCA_000183345.1
109 Escherichia coli UMN026 GCA_000026325.2
109 Escherichia coli str. K-12 substr. MG1655 GCA_000005845.2
109 Obesumbacterium proteus GCA_001586165.1
109 Pantoea ananatis LMG 20103 GCA_000025405.2
109 Pantoea dispersa EGD-AAK13 GCA_000465555.2
109 Plesiomonas shigelloides GCA_900087055.1
109 Rosenbergiella nectarea GCA_900111105.1
109 Serratia fonticola GCA_001006005.1
109 Serratia liquefaciens ATCC 27592 GCA_000422085.1
109 Serratia marcescens subsp. marcescens Db11 GCA_000513215.1
109 Serratia rubidaea GCA_001572725.1
109 Shigella dysenteriae Sd197 GCA_000012005.1
109 Shigella flexneri 2a str. 301 GCA_000006925.2
109 Shimwellia blattae DSM 4481 = NBRC 105725 GCA_000262305.1
109 Tumebacillus flagellatus GCA_000714935.1
109 Yersinia enterocolitica subsp. enterocolitica 8081 GCA_000009345.1
109 Yersinia pestis CO92 GCA_000009065.1
cL
169 Clostridium beijerinckii GCA_000833105.2
169 Clostridium butyricum GCA_001456065.2
169 Clostridium puniceum GCA_002006345.1
169 Clostridium saccharoperbutylacetonicum N1-4_28HMT_29 GCA_000340885.1
161 Clostridium chromiireducens GCA_002029255.1
161 Clostridium saccharobutylicum DSM 13864 GCA_000473995.1
143 Clostridium taeniosporum GCA_001735765.1
cM
327 Streptococcus mutans UA159 GCA_000007465.2
212 Streptococcus ferus DSM 20646 GCA_000372425.1
208 Streptococcus ratti FA-1 = DSM 20564 GCA_000286075.1
cN
315 Escherichia coli IAI39 GCA_000026345.1
315 Escherichia coli O104_3AH4 str. 2011C-3493 GCA_000299455.1
315 Escherichia coli O157_3AH7 str. Sakai GCA_000008865.1
315 Escherichia coli O83_3AH1 str. NRG 857C GCA_000183345.1
315 Escherichia coli UMN026 GCA_000026325.2
315 Escherichia coli str. K-12 substr. MG1655 GCA_000005845.2
315 Shigella dysenteriae Sd197 GCA_000012005.1
315 Shigella flexneri 2a str. 301 GCA_000006925.2
315 Tumebacillus flagellatus GCA_000714935.1
275 Erwinia iniecta GCA_001267535.1
252 Rosenbergiella nectarea GCA_900111105.1
cO
176 Clostridium beijerinckii GCA_000833105.2
176 Clostridium puniceum GCA_002006345.1
176 Clostridium saccharoperbutylacetonicum N1-4_28HMT_29 GCA_000340885.1
167 Clostridium saccharobutylicum DSM 13864 GCA_000473995.1
151 Clostridium neonatale GCA_001458595.1
cP
161 Clostridium beijerinckii GCA_000833105.2
161 Clostridium puniceum GCA_002006345.1
161 Clostridium saccharobutylicum DSM 13864 GCA_000473995.1
161 Clostridium saccharoperbutylacetonicum N1-4_28HMT_29 GCA_000340885.1
153 Clostridium chromiireducens GCA_002029255.1
153 Clostridium taeniosporum GCA_001735765.1
143 Clostridium butyricum GCA_001456065.2
cQ
192 Clostridium beijerinckii GCA_000833105.2
192 Clostridium chromiireducens GCA_002029255.1
192 Clostridium puniceum GCA_002006345.1
192 Clostridium saccharobutylicum DSM 13864 GCA_000473995.1
192 Clostridium saccharoperbutylacetonicum N1-4_28HMT_29 GCA_000340885.1
187 Clostridium butyricum GCA_001456065.2

181 Clostridium taeniosporum GCA_001735765.1
cR
164 Rhodobacter sphaeroides 2.4.1 GCA_000012905.2
142 Rhodobacter capsulatus SB 1003 GCA_000021865.1
142 Rhodobacter sphaeroides ATCC 17025 GCA_000016405.1
135 Haematobacter massiliensis GCA_000740795.1
cS
204 Streptococcus mutans UA159 GCA_000007465.2
142 Streptococcus cristatus AS 1.3089 GCA_000385925.1
142 Streptococcus gordonii str. Challis substr. CH1 GCA_000017005.1
142 Streptococcus mitis B6 GCA_000027165.1
142 Streptococcus parasanguinis ATCC 15912 GCA_000164675.2
142 Streptococcus pneumoniae R6 GCA_000007045.1
128 Streptococcus marimammalium DSM 18627 GCA_000380045.1
cT
239 Escherichia coli IAI39 GCA_000026345.1
239 Escherichia coli O104_3AH4 str. 2011C-3493 GCA_000299455.1
239 Escherichia coli O157_3AH7 str. Sakai GCA_000008865.1
239 Escherichia coli O83_3AH1 str. NRG 857C GCA_000183345.1
239 Escherichia coli UMN026 GCA_000026325.2
239 Escherichia coli str. K-12 substr. MG1655 GCA_000005845.2
239 Shigella flexneri 2a str. 301 GCA_000006925.2
239 Tumebacillus flagellatus GCA_000714935.1
230 Shigella dysenteriae Sd197 GCA_000012005.1
209 Erwinia iniecta GCA_001267535.1
209 Plautia stali symbiont GCA_000180175.2
cU
244 Bifidobacterium adolescentis ATCC 15703 GCA_000010425.1
213 Bifidobacterium dentium JCM 1195 = DSM 20436 GCA_001042595.1
211 Bifidobacterium thermophilum GCA_000741495.1
cV
298 Staphylococcus epidermidis ATCC 12228 GCA_000007645.1
287 Staphylococcus haemolyticus JCSC1435 GCA_000009865.1
282 Staphylococcus capitis subsp. capitis GCA_001028645.1
cW
222 Enterococcus faecalis V583 GCA_000007785.1
222 Streptomyces cinnamoneus GCA_001885705.1
207 Enterococcus faecium DO GCA_000174395.2
197 Enterococcus canis NBRC 100695 GCA_001544375.1
197 Enterococcus casseliflavus EC20 GCA_000157355.2
197 Enterococcus dispar ATCC 51266 GCA_000406945.1
197 Enterococcus gilvus ATCC BAA-350 GCA_000407545.1
197 Enterococcus haemoperoxidus ATCC BAA-382 GCA_000407165.1
197 Enterococcus hermanniensis GCA_001885945.1
197 Enterococcus hirae ATCC 9790 GCA_000271405.2
197 Enterococcus malodoratus ATCC 43197 GCA_000407185.1
197 Enterococcus mundtii QU 25 GCA_000504125.1
197 Enterococcus pallens ATCC BAA-351 GCA_000407485.1
197 Enterococcus phoeniculicola ATCC BAA-412 GCA_000407505.1
197 Enterococcus pseudoavium NBRC 100491 GCA_001544295.1
197 Enterococcus rivorum GCA_001742285.1
197 Enterococcus saccharolyticus subsp. saccharolyticus ATCC 43076 GCA_000407285.1
197 Enterococcus thailandicus GCA_001652875.1
cX
133 Erwinia iniecta GCA_001267535.1
133 Escherichia coli IAI39 GCA_000026345.1
133 Escherichia coli O104_3AH4 str. 2011C-3493 GCA_000299455.1
133 Escherichia coli O157_3AH7 str. Sakai GCA_000008865.1
133 Escherichia coli O83_3AH1 str. NRG 857C GCA_000183345.1
133 Escherichia coli UMN026 GCA_000026325.2
133 Escherichia coli str. K-12 substr. MG1655 GCA_000005845.2
133 Shigella dysenteriae Sd197 GCA_000012005.1
133 Shigella flexneri 2a str. 301 GCA_000006925.2
133 Tumebacillus flagellatus GCA_000714935.1
117 Rosenbergiella nectarea GCA_900111105.1
115 Cedecea neteri GCA_000757825.1
cY
193 Lactobacillus gasseri ATCC 33323 = JCM 1131 GCA_000014425.1
173 Lactobacillus hominis DSM 23910 = CRBIP 24.179 GCA_000296835.1
158 Lactobacillus acidophilus NCFM GCA_000011985.1
158 Lactobacillus gallinarum GCA_001314245.2
cZ
247 Clostridium beijerinckii GCA_000833105.2
247 Clostridium puniceum GCA_002006345.1
247 Clostridium saccharobutylicum DSM 13864 GCA_000473995.1
247 Clostridium saccharoperbutylacetonicum N1-4_28HMT_29 GCA_000340885.1

228 Clostridium chromiireducens GCA_002029255.1
214 Clostridium amylolyticum GCA_900142075.1
214 Clostridium butyricum GCA_001456065.2
214 Clostridium polynesiense GCA_000820705.1
d0
212 Rhodobacter sphaeroides 2.4.1 GCA_000012905.2
175 Pseudorhodobacter ferrugineus DSM 5888 GCA_000420745.1
175 Pseudorhodobacter wandonensis GCA_001202035.1
163 Thioclava dalianensis GCA_000715505.1
163 Thioclava indica GCA_000714545.1
d1
293 Rhodobacter sphaeroides 2.4.1 GCA_000012905.2
264 Rhodobacter sphaeroides ATCC 17025 GCA_000016405.1
247 Pseudorhodobacter ferrugineus DSM 5888 GCA_000420745.1
247 Pseudorhodobacter wandonensis GCA_001202035.1
d2
184 Escherichia coli IAI39 GCA_000026345.1
184 Escherichia coli O104_3AH4 str. 2011C-3493 GCA_000299455.1
184 Escherichia coli O157_3AH7 str. Sakai GCA_000008865.1
184 Escherichia coli O83_3AH1 str. NRG 857C GCA_000183345.1
184 Escherichia coli UMN026 GCA_000026325.2
184 Escherichia coli str. K-12 substr. MG1655 GCA_000005845.2
184 Shigella dysenteriae Sd197 GCA_000012005.1
184 Shigella flexneri 2a str. 301 GCA_000006925.2
184 Tumebacillus flagellatus GCA_000714935.1
171 Kosakonia cowanii GCA_001975225.1
170 Enterobacter hormaechei subsp. steigerwaltii GCA_001729725.1
170 Erwinia iniecta GCA_001267535.1
170 Erwinia toletana DAPP-PG 735 GCA_000336255.1
170 Klebsiella oxytoca GCA_001022195.1
170 Kosakonia sacchari SP1 GCA_000300455.4
170 Pluralibacter gergoviae GCA_000757785.1
170 Pseudescherichia vulneris NBRC 102420 GCA_000759795.1
d3
287 Bifidobacterium adolescentis ATCC 15703 GCA_000010425.1
246 Bifidobacterium callitrichos DSM 23973 GCA_000741175.1
220 Bifidobacterium breve DSM 20213 = JCM 1192 GCA_001025175.1
d4
199 Enterococcus faecalis V583 GCA_000007785.1
199 Streptomyces cinnamoneus GCA_001885705.1
180 Enterococcus aquimarinus GCA_001885765.1
180 Enterococcus canis NBRC 100695 GCA_001544375.1
180 Enterococcus casseliflavus EC20 GCA_000157355.2
180 Enterococcus dispar ATCC 51266 GCA_000406945.1
180 Enterococcus faecium DO GCA_000174395.2
180 Enterococcus gilvus ATCC BAA-350 GCA_000407545.1
180 Enterococcus haemoperoxidus ATCC BAA-382 GCA_000407165.1
180 Enterococcus hermanniensis GCA_001885945.1
180 Enterococcus hirae ATCC 9790 GCA_000271405.2
180 Enterococcus malodoratus ATCC 43197 GCA_000407185.1
180 Enterococcus massiliensis GCA_001050095.1
180 Enterococcus mundtii QU 25 GCA_000504125.1
180 Enterococcus pallens ATCC BAA-351 GCA_000407485.1
180 Enterococcus phoeniculicola ATCC BAA-412 GCA_000407505.1
180 Enterococcus pseudoavium NBRC 100491 GCA_001544295.1
180 Enterococcus rivorum GCA_001742285.1
180 Enterococcus saccharolyticus subsp. saccharolyticus ATCC 43076 GCA_000407285.1
180 Enterococcus thailandicus GCA_001652875.1
166 Enterococcus asini ATCC 700915 GCA_000407365.1
166 Enterococcus cecorum GCA_001318405.1
166 Enterococcus columbae DSM 7374 = ATCC 51263 GCA_000406925.1
166 Enterococcus italicus DSM 15952 GCA_000185365.1
166 Enterococcus sulfureus ATCC 49903 GCA_000407605.1
166 Melissococcus plutonius S1 GCA_000747585.1
166 Vagococcus fluvialis bH819 GCA_900163795.1
166 Vagococcus lutrae LBD1 GCA_000498295.1
166 Vagococcus penaei GCA_001998885.1
166 Vagococcus teuberi GCA_001870205.1
d5
99 Deinococcus radiodurans R1 GCA_000008565.1
93 Clostridium beijerinckii GCA_000833105.2
93 Clostridium puniceum GCA_002006345.1
93 Clostridium saccharobutylicum DSM 13864 GCA_000473995.1
93 Clostridium saccharoperbutylacetonicum N1-4_28HMT_29 GCA_000340885.1
82 Clostridium botulinum B str. Eklund 17B _28NRP_29 GCA_000020165.1
82 Clostridium butyricum GCA_001456065.2

82 Clostridium chromiireducens GCA_002029255.1
82 Clostridium taeniosporum GCA_001735765.1
82 Clostridium uliginosum GCA_900112485.1
d6
211 Clostridium beijerinckii GCA_000833105.2
211 Clostridium saccharoperbutylacetonicum N1-4_28HMT_29 GCA_000340885.1
205 Clostridium puniceum GCA_002006345.1
196 Clostridium saccharobutylicum DSM 13864 GCA_000473995.1
d7
260 Escherichia coli IAI39 GCA_000026345.1
260 Escherichia coli O104_3AH4 str. 2011C-3493 GCA_000299455.1
260 Escherichia coli O157_3AH7 str. Sakai GCA_000008865.1
260 Escherichia coli O83_3AH1 str. NRG 857C GCA_000183345.1
260 Escherichia coli UMN026 GCA_000026325.2
260 Escherichia coli str. K-12 substr. MG1655 GCA_000005845.2
260 Shigella flexneri 2a str. 301 GCA_000006925.2
260 Tumebacillus flagellatus GCA_000714935.1
251 Shigella dysenteriae Sd197 GCA_000012005.1
221 Erwinia iniecta GCA_001267535.1
d8
231 Deinococcus radiodurans R1 GCA_000008565.1
163 Deinococcus marmoris DSM 12784 GCA_000701405.1
163 Deinococcus swuensis GCA_000800395.1
149 Deinococcus gobiensis I-0 GCA_000252445.1
d9
62 Lactobacillus acidophilus NCFM GCA_000011985.1
62 Lactobacillus amylophilus DSM 20533 = JCM 1125 GCA_001936335.1
62 Lactobacillus bombicola GCA_900112665.1
62 Lactobacillus florum DSM 22689 = JCM 16035 GCA_001436645.1
62 Lactobacillus gallinarum GCA_001314245.2
62 Lactobacillus gigeriorum DSM 23908 = CRBIP 24.85 GCA_001436575.1
62 Lactobacillus helveticus GCA_001308285.1
62 Lactobacillus jensenii GCA_001936235.1
62 Lactobacillus lindneri GCA_001702135.1
62 Lactobacillus pasteurii DSM 23907 = CRBIP 24.76 GCA_000297025.1
62 Lactobacillus sanfranciscensis TMW 1.1304 GCA_000225325.1
62 Lactobacillus vini DSM 20605 GCA_000255495.2
60 Deinococcus radiodurans R1 GCA_000008565.1
56 Lactobacillus acetotolerans GCA_001042405.1
56 Lactobacillus amylolyticus GCA_002075105.1
56 Lactobacillus amylovorus GCA_000191545.1
56 Lactobacillus apis GCA_000970735.1
56 Lactobacillus crispatus ST1 GCA_000091765.1
56 Lactobacillus delbrueckii subsp. bulgaricus ATCC 11842 = JCM 1002 GCA_000056065.1
56 Lactobacillus gasseri ATCC 33323 = JCM 1131 GCA_000014425.1
56 Lactobacillus hamsteri DSM 5661 = JCM 6256 GCA_000615445.1
56 Lactobacillus harbinensis DSM 16991 GCA_000425885.1
56 Lactobacillus hominis DSM 23910 = CRBIP 24.179 GCA_000296835.1
56 Lactobacillus iners DSM 13335 GCA_000160875.1
56 Lactobacillus intestinalis DSM 6629 GCA_001435325.1
56 Lactobacillus kalixensis DSM 16043 GCA_001434335.1
56 Lactobacillus kefiranofaciens ZW3 GCA_000214785.1
56 Lactobacillus kullabergensis GCA_000967195.1
56 Lactobacillus perolens DSM 12744 GCA_001435585.1
56 Lactobacillus psittaci DSM 15354 GCA_000425905.1
56 Lactobacillus shenzhenensis LY-73 GCA_000469325.1
da
224 Deinococcus radiodurans R1 GCA_000008565.1
174 Deinococcus gobiensis I-0 GCA_000252445.1
160 Deinococcus puniceus GCA_001644565.1
db
128 Enterococcus canis NBRC 100695 GCA_001544375.1
128 Enterococcus faecalis V583 GCA_000007785.1
128 Enterococcus faecium DO GCA_000174395.2
128 Enterococcus haemoperoxidus ATCC BAA-382 GCA_000407165.1
128 Enterococcus hirae ATCC 9790 GCA_000271405.2
128 Enterococcus mundtii QU 25 GCA_000504125.1
128 Enterococcus phoeniculicola ATCC BAA-412 GCA_000407505.1
128 Enterococcus rivorum GCA_001742285.1
128 Enterococcus thailandicus GCA_001652875.1
128 Streptomyces cinnamoneus GCA_001885705.1
127 Enterococcus dispar ATCC 51266 GCA_000406945.1
125 Enterococcus asini ATCC 700915 GCA_000407365.1
dc
220 Deinococcus radiodurans R1 GCA_000008565.1
151 Deinococcus puniceus GCA_001644565.1

145 Deinococcus gobiensis I-0 GCA_000252445.1
dd
228 Deinococcus radiodurans R1 GCA_000008565.1
169 Deinococcus puniceus GCA_001644565.1
167 Deinococcus deserti VCD115 GCA_000020685.1
de
178 Escherichia coli IAI39 GCA_000026345.1
178 Escherichia coli O104_3AH4 str. 2011C-3493 GCA_000299455.1
178 Escherichia coli O157_3AH7 str. Sakai GCA_000008865.1
178 Escherichia coli O83_3AH1 str. NRG 857C GCA_000183345.1
178 Escherichia coli UMN026 GCA_000026325.2
178 Escherichia coli str. K-12 substr. MG1655 GCA_000005845.2
178 Shigella flexneri 2a str. 301 GCA_000006925.2
178 Tumebacillus flagellatus GCA_000714935.1
169 Shigella dysenteriae Sd197 GCA_000012005.1
142 Bacillus humi GCA_001439915.1
df
293 Deinococcus radiodurans R1 GCA_000008565.1
206 Deinococcus gobiensis I-0 GCA_000252445.1
185 Deinococcus deserti VCD115 GCA_000020685.1
185 Deinococcus geothermalis DSM 11300 GCA_000196275.1
dg
288 Clostridium beijerinckii GCA_000833105.2
288 Clostridium puniceum GCA_002006345.1
288 Clostridium saccharobutylicum DSM 13864 GCA_000473995.1
288 Clostridium saccharoperbutylacetonicum N1-4_28HMT_29 GCA_000340885.1
272 Clostridium chromiireducens GCA_002029255.1
250 Clostridium butyricum GCA_001456065.2
dh
108 Deinococcus radiodurans R1 GCA_000008565.1
64 Deinococcus proteolyticus MRP GCA_000190555.1
54 Deinococcus marmoris DSM 12784 GCA_000701405.1
54 Deinococcus swuensis GCA_000800395.1
di
236 Deinococcus radiodurans R1 GCA_000008565.1
156 Deinococcus marmoris DSM 12784 GCA_000701405.1
156 Deinococcus swuensis GCA_000800395.1
155 Deinococcus puniceus GCA_001644565.1
dj
214 Clostridium beijerinckii GCA_000833105.2
214 Clostridium puniceum GCA_002006345.1
214 Clostridium saccharoperbutylacetonicum N1-4_28HMT_29 GCA_000340885.1
206 Clostridium saccharobutylicum DSM 13864 GCA_000473995.1
196 Clostridium taeniosporum GCA_001735765.1
dk
219 Streptococcus mutans UA159 GCA_000007465.2
165 Streptococcus ratti FA-1 = DSM 20564 GCA_000286075.1
152 Streptococcus ferus DSM 20646 GCA_000372425.1
dl
205 Bifidobacterium adolescentis ATCC 15703 GCA_000010425.1
191 Bifidobacterium dentium JCM 1195 = DSM 20436 GCA_001042595.1
176 Bifidobacterium callitrichos DSM 23973 GCA_000741175.1
dm
147 Clostridium beijerinckii GCA_000833105.2
147 Clostridium puniceum GCA_002006345.1
147 Clostridium saccharoperbutylacetonicum N1-4_28HMT_29 GCA_000340885.1
133 Clostridium saccharobutylicum DSM 13864 GCA_000473995.1
132 Clostridium taeniosporum GCA_001735765.1
dn
166 Rhodobacter sphaeroides ATCC 17025 GCA_000016405.1
151 Rhodobacter sphaeroides 2.4.1 GCA_000012905.2
138 Gemmobacter megaterium GCA_900156815.1
138 Pseudorhodobacter ferrugineus DSM 5888 GCA_000420745.1
138 Pseudorhodobacter wandonensis GCA_001202035.1
do
256 Enterococcus faecalis V583 GCA_000007785.1
256 Streptomyces cinnamoneus GCA_001885705.1
223 Enterococcus canis NBRC 100695 GCA_001544375.1
223 Enterococcus dispar ATCC 51266 GCA_000406945.1
223 Enterococcus faecium DO GCA_000174395.2
223 Enterococcus haemoperoxidus ATCC BAA-382 GCA_000407165.1
223 Enterococcus hirae ATCC 9790 GCA_000271405.2
223 Enterococcus mundtii QU 25 GCA_000504125.1
223 Enterococcus phoeniculicola ATCC BAA-412 GCA_000407505.1
223 Enterococcus rivorum GCA_001742285.1
223 Enterococcus thailandicus GCA_001652875.1

208 Enterococcus asini ATCC 700915 GCA_000407365.1
208 Enterococcus gilvus ATCC BAA-350 GCA_000407545.1
208 Enterococcus hermanniensis GCA_001885945.1
208 Enterococcus malodoratus ATCC 43197 GCA_000407185.1
208 Enterococcus massiliensis GCA_001050095.1
208 Enterococcus pallens ATCC BAA-351 GCA_000407485.1
208 Enterococcus pseudoavium NBRC 100491 GCA_001544295.1
208 Listeria floridensis FSL S10-1187 GCA_000525875.1
dp
190 Enterococcus faecalis V583 GCA_000007785.1
190 Streptomyces cinnamoneus GCA_001885705.1
165 Enterococcus dispar ATCC 51266 GCA_000406945.1
162 Enterococcus canis NBRC 100695 GCA_001544375.1
162 Enterococcus casseliflavus EC20 GCA_000157355.2
162 Enterococcus faecium DO GCA_000174395.2
162 Enterococcus hirae ATCC 9790 GCA_000271405.2
162 Enterococcus massiliensis GCA_001050095.1
162 Enterococcus mundtii QU 25 GCA_000504125.1
162 Enterococcus rivorum GCA_001742285.1
162 Enterococcus saccharolyticus subsp. saccharolyticus ATCC 43076 GCA_000407285.1
dq
237 Bacillus anthracis str. Ames GCA_000007845.1
237 Bacillus anthracis str. Sterne GCA_000008165.1
237 Bacillus cereus ATCC 14579 GCA_000007825.1
237 Bacillus mycoides GCA_000832605.1
237 Bacillus pseudomycoides DSM 12442 GCA_000161455.1
237 Bacillus thuringiensis YBT-1518 GCA_000497525.2
237 _5BBacillus thuringiensis_5D serovar konkukian str. 97-27 GCA_000008505.1
174 Bacillus halmapalus GCA_002019665.1
169 Bacillus aquimaris TF-12 GCA_001648555.1
169 Bacillus manliponensis GCA_000712595.1
dr
229 Rhodobacter sphaeroides 2.4.1 GCA_000012905.2
206 Pseudorhodobacter ferrugineus DSM 5888 GCA_000420745.1
206 Pseudorhodobacter wandonensis GCA_001202035.1
200 Gemmobacter megaterium GCA_900156815.1
200 Rhodobacter sphaeroides ATCC 17025 GCA_000016405.1
ds
124 Clostridium beijerinckii GCA_000833105.2
124 Clostridium saccharoperbutylacetonicum N1-4_28HMT_29 GCA_000340885.1
111 Clostridium puniceum GCA_002006345.1
105 Clostridium saccharobutylicum DSM 13864 GCA_000473995.1
dt
245 Escherichia coli IAI39 GCA_000026345.1
245 Escherichia coli O104_3AH4 str. 2011C-3493 GCA_000299455.1
245 Escherichia coli O157_3AH7 str. Sakai GCA_000008865.1
245 Escherichia coli O83_3AH1 str. NRG 857C GCA_000183345.1
245 Escherichia coli UMN026 GCA_000026325.2
245 Escherichia coli str. K-12 substr. MG1655 GCA_000005845.2
245 Shigella flexneri 2a str. 301 GCA_000006925.2
245 Tumebacillus flagellatus GCA_000714935.1
236 Shigella dysenteriae Sd197 GCA_000012005.1
220 Erwinia iniecta GCA_001267535.1
du
163 Enterococcus faecalis V583 GCA_000007785.1
163 Streptomyces cinnamoneus GCA_001885705.1
148 Enterococcus canis NBRC 100695 GCA_001544375.1
148 Enterococcus dispar ATCC 51266 GCA_000406945.1
148 Enterococcus faecium DO GCA_000174395.2
148 Enterococcus hirae ATCC 9790 GCA_000271405.2
148 Enterococcus massiliensis GCA_001050095.1
148 Enterococcus mundtii QU 25 GCA_000504125.1
148 Enterococcus rivorum GCA_001742285.1
143 Enterococcus haemoperoxidus ATCC BAA-382 GCA_000407165.1
143 Enterococcus phoeniculicola ATCC BAA-412 GCA_000407505.1
143 Enterococcus thailandicus GCA_001652875.1
143 Isobaculum melis GCA_900111355.1
dv
190 Clostridium beijerinckii GCA_000833105.2
190 Clostridium puniceum GCA_002006345.1
190 Clostridium saccharoperbutylacetonicum N1-4_28HMT_29 GCA_000340885.1
175 Clostridium saccharobutylicum DSM 13864 GCA_000473995.1
163 Clostridium butyricum GCA_001456065.2
dw
205 Escherichia coli IAI39 GCA_000026345.1
205 Escherichia coli O104_3AH4 str. 2011C-3493 GCA_000299455.1

205 Escherichia coli O157_3AH7 str. Sakai GCA_000008865.1
205 Escherichia coli O83_3AH1 str. NRG 857C GCA_000183345.1
205 Escherichia coli UMN026 GCA_000026325.2
205 Escherichia coli str. K-12 substr. MG1655 GCA_000005845.2
205 Shigella dysenteriae Sd197 GCA_000012005.1
205 Shigella flexneri 2a str. 301 GCA_000006925.2
205 Tumebacillus flagellatus GCA_000714935.1
201 Erwinia iniecta GCA_001267535.1
198 Enterobacter hormaechei subsp. steigerwaltii GCA_001729725.1
198 Erwinia toletana DAPP-PG 735 GCA_000336255.1
198 Klebsiella oxytoca GCA_001022195.1
198 Kosakonia cowanii GCA_001975225.1
198 Kosakonia sacchari SP1 GCA_000300455.4
198 Pseudescherichia vulneris NBRC 102420 GCA_000759795.1
dx
259 Clostridium beijerinckii GCA_000833105.2
259 Clostridium puniceum GCA_002006345.1
259 Clostridium saccharoperbutylacetonicum N1-4_28HMT_29 GCA_000340885.1
251 Clostridium saccharobutylicum DSM 13864 GCA_000473995.1
236 Clostridium chromiireducens GCA_002029255.1
dy
221 Escherichia coli IAI39 GCA_000026345.1
221 Escherichia coli O104_3AH4 str. 2011C-3493 GCA_000299455.1
221 Escherichia coli O157_3AH7 str. Sakai GCA_000008865.1
221 Escherichia coli O83_3AH1 str. NRG 857C GCA_000183345.1
221 Escherichia coli UMN026 GCA_000026325.2
221 Escherichia coli str. K-12 substr. MG1655 GCA_000005845.2
221 Shigella dysenteriae Sd197 GCA_000012005.1
221 Shigella flexneri 2a str. 301 GCA_000006925.2
221 Tumebacillus flagellatus GCA_000714935.1
207 Erwinia iniecta GCA_001267535.1
187 Cedecea neteri GCA_000757825.1
187 Enterobacter hormaechei subsp. steigerwaltii GCA_001729725.1
187 Erwinia toletana DAPP-PG 735 GCA_000336255.1
187 Klebsiella oxytoca GCA_001022195.1
187 Kosakonia cowanii GCA_001975225.1
187 Kosakonia sacchari SP1 GCA_000300455.4
187 Pseudescherichia vulneris NBRC 102420 GCA_000759795.1
187 Rosenbergiella nectarea GCA_900111105.1
dz
155 Bacillus anthracis str. Ames GCA_000007845.1
155 Bacillus anthracis str. Sterne GCA_000008165.1
155 Bacillus cereus ATCC 14579 GCA_000007825.1
155 Bacillus manliponensis GCA_000712595.1
155 Bacillus mycoides GCA_000832605.1
155 Bacillus pseudomycoides DSM 12442 GCA_000161455.1
155 Bacillus thuringiensis YBT-1518 GCA_000497525.2
155 _5BBacillus thuringiensis_5D serovar konkukian str. 97-27 GCA_000008505.1
148 Bacillus halmapalus GCA_002019665.1
141 Bacillus aquimaris TF-12 GCA_001648555.1
141 Bacillus marisflavi GCA_001274775.1
dA
193 Escherichia coli IAI39 GCA_000026345.1
193 Escherichia coli O104_3AH4 str. 2011C-3493 GCA_000299455.1
193 Escherichia coli O157_3AH7 str. Sakai GCA_000008865.1
193 Escherichia coli UMN026 GCA_000026325.2
193 Escherichia coli str. K-12 substr. MG1655 GCA_000005845.2
193 Shigella dysenteriae Sd197 GCA_000012005.1
193 Shigella flexneri 2a str. 301 GCA_000006925.2
193 Tumebacillus flagellatus GCA_000714935.1
192 Escherichia coli O83_3AH1 str. NRG 857C GCA_000183345.1
167 Erwinia iniecta GCA_001267535.1
dB
134 Clostridium beijerinckii GCA_000833105.2
134 Clostridium puniceum GCA_002006345.1
134 Clostridium saccharobutylicum DSM 13864 GCA_000473995.1
134 Clostridium saccharoperbutylacetonicum N1-4_28HMT_29 GCA_000340885.1
115 Clostridium amylolyticum GCA_900142075.1
115 Clostridium disporicum GCA_001405015.1
115 Clostridium intestinale URNW GCA_000469625.2
115 Clostridium polynesiense GCA_000820705.1
114 Provencibacterium massiliense GCA_900169495.1
dC
210 Bifidobacterium adolescentis ATCC 15703 GCA_000010425.1
164 Bifidobacterium callitrichos DSM 23973 GCA_000741175.1
154 Bifidobacterium gallicum DSM 20093 = LMG 11596 GCA_000741205.1

dD
224 Bacillus anthracis str. Ames GCA_000007845.1
224 Bacillus anthracis str. Sterne GCA_000008165.1
224 Bacillus cereus ATCC 14579 GCA_000007825.1
224 Bacillus mycoides GCA_000832605.1
224 Bacillus pseudomycoides DSM 12442 GCA_000161455.1
224 Bacillus thuringiensis YBT-1518 GCA_000497525.2
224 _5BBacillus thuringiensis_5D serovar konkukian str. 97-27 GCA_000008505.1
216 Bacillus manliponensis GCA_000712595.1
192 Bacillus cytotoxicus NVH 391-98 GCA_000017425.1
dE
209 Clostridium beijerinckii GCA_000833105.2
209 Clostridium saccharobutylicum DSM 13864 GCA_000473995.1
209 Clostridium saccharoperbutylacetonicum N1-4_28HMT_29 GCA_000340885.1
202 Clostridium puniceum GCA_002006345.1
191 Clostridium chromiireducens GCA_002029255.1
dF
271 Clostridium butyricum GCA_001456065.2
271 Clostridium saccharobutylicum DSM 13864 GCA_000473995.1
268 Clostridium beijerinckii GCA_000833105.2
268 Clostridium puniceum GCA_002006345.1
268 Clostridium saccharoperbutylacetonicum N1-4_28HMT_29 GCA_000340885.1
255 Clostridium chromiireducens GCA_002029255.1
dG
225 Enterococcus dispar ATCC 51266 GCA_000406945.1
223 Enterococcus canis NBRC 100695 GCA_001544375.1
223 Enterococcus faecalis V583 GCA_000007785.1
223 Enterococcus faecium DO GCA_000174395.2
223 Enterococcus haemoperoxidus ATCC BAA-382 GCA_000407165.1
223 Enterococcus hirae ATCC 9790 GCA_000271405.2
223 Enterococcus mundtii QU 25 GCA_000504125.1
223 Enterococcus phoeniculicola ATCC BAA-412 GCA_000407505.1
223 Enterococcus rivorum GCA_001742285.1
223 Enterococcus thailandicus GCA_001652875.1
223 Streptomyces cinnamoneus GCA_001885705.1
222 Enterococcus asini ATCC 700915 GCA_000407365.1
222 Enterococcus casseliflavus EC20 GCA_000157355.2
222 Enterococcus massiliensis GCA_001050095.1
222 Enterococcus saccharolyticus subsp. saccharolyticus ATCC 43076 GCA_000407285.1
dH
191 Staphylococcus pettenkoferi GCA_002208805.1
175 Staphylococcus epidermidis ATCC 12228 GCA_000007645.1
172 Staphylococcus cohnii subsp. cohnii GCA_000972575.1
172 Staphylococcus haemolyticus JCSC1435 GCA_000009865.1
172 Staphylococcus saprophyticus subsp. saprophyticus ATCC 15305 GCA_000010125.1
dI
271 Deinococcus radiodurans R1 GCA_000008565.1
210 Deinococcus gobiensis I-0 GCA_000252445.1
182 Deinococcus puniceus GCA_001644565.1
dJ
141 Streptococcus mutans UA159 GCA_000007465.2
131 Lactobacillus selangorensis GCA_001437205.1
127 Lactobacillus backii GCA_001663675.1
127 Lactobacillus concavus DSM 17758 GCA_001435835.1
127 Lactobacillus dextrinicus DSM 20335 GCA_001436095.1
127 Lactobacillus harbinensis DSM 16991 GCA_000425885.1
127 Lactobacillus perolens DSM 12744 GCA_001435585.1
127 Lactobacillus shenzhenensis LY-73 GCA_000469325.1
dK
197 Clostridium beijerinckii GCA_000833105.2
197 Clostridium saccharobutylicum DSM 13864 GCA_000473995.1
197 Clostridium saccharoperbutylacetonicum N1-4_28HMT_29 GCA_000340885.1
193 Clostridium puniceum GCA_002006345.1
185 Clostridium butyricum GCA_001456065.2
dL
184 Escherichia coli IAI39 GCA_000026345.1
184 Escherichia coli O104_3AH4 str. 2011C-3493 GCA_000299455.1
184 Escherichia coli O157_3AH7 str. Sakai GCA_000008865.1
184 Escherichia coli O83_3AH1 str. NRG 857C GCA_000183345.1
184 Escherichia coli UMN026 GCA_000026325.2
184 Escherichia coli str. K-12 substr. MG1655 GCA_000005845.2
184 Shigella dysenteriae Sd197 GCA_000012005.1
184 Shigella flexneri 2a str. 301 GCA_000006925.2
184 Tumebacillus flagellatus GCA_000714935.1
140 Cedecea neteri GCA_000757825.1
138 Erwinia iniecta GCA_001267535.1

dM
157 Escherichia coli IAI39 GCA_000026345.1
157 Escherichia coli O104_3AH4 str. 2011C-3493 GCA_000299455.1
157 Escherichia coli O157_3AH7 str. Sakai GCA_000008865.1
157 Escherichia coli O83_3AH1 str. NRG 857C GCA_000183345.1
157 Escherichia coli UMN026 GCA_000026325.2
157 Escherichia coli str. K-12 substr. MG1655 GCA_000005845.2
157 Shigella dysenteriae Sd197 GCA_000012005.1
157 Shigella flexneri 2a str. 301 GCA_000006925.2
157 Tumebacillus flagellatus GCA_000714935.1
133 Klebsiella pneumoniae subsp. pneumoniae HS11286 GCA_000240185.2
125 Dickeya zeae Ech586 GCA_000025065.1
dN
260 Bacillus anthracis str. Ames GCA_000007845.1
260 Bacillus anthracis str. Sterne GCA_000008165.1
260 Bacillus cereus ATCC 14579 GCA_000007825.1
260 Bacillus mycoides GCA_000832605.1
260 Bacillus pseudomycoides DSM 12442 GCA_000161455.1
260 Bacillus thuringiensis YBT-1518 GCA_000497525.2
260 _5BBacillus thuringiensis_5D serovar konkukian str. 97-27 GCA_000008505.1
236 Bacillus manliponensis GCA_000712595.1
227 Bacillus weihaiensis GCA_001889165.1
dO
102 Clostridium beijerinckii GCA_000833105.2
102 Clostridium intestinale URNW GCA_000469625.2
102 Clostridium neonatale GCA_001458595.1
102 Clostridium puniceum GCA_002006345.1
102 Clostridium saccharobutylicum DSM 13864 GCA_000473995.1
102 Clostridium saccharoperbutylacetonicum N1-4_28HMT_29 GCA_000340885.1
102 Desnuesiella massiliensis GCA_001403615.1
91 Clostridium baratii str. Sullivan GCA_000789395.1
91 Clostridium botulinum B str. Eklund 17B _28NRP_29 GCA_000020165.1
91 Clostridium butyricum GCA_001456065.2
91 Clostridium chromiireducens GCA_002029255.1
91 Clostridium taeniosporum GCA_001735765.1
90 Clostridium bornimense GCA_000577895.1
dP
125 Bifidobacterium adolescentis ATCC 15703 GCA_000010425.1
123 Bifidobacterium asteroides PRL2011 GCA_000304215.1
121 Bifidobacterium tsurumiense GCA_000741765.1
dQ
201 Staphylococcus aureus subsp. aureus NCTC 8325 GCA_000013425.1
201 Staphylococcus cohnii subsp. cohnii GCA_000972575.1
201 Staphylococcus epidermidis ATCC 12228 GCA_000007645.1
201 Staphylococcus haemolyticus JCSC1435 GCA_000009865.1
201 Staphylococcus hominis subsp. hominis C80 GCA_000183685.1
201 Staphylococcus simiae CCM 7213 GCA_000235645.2
197 Staphylococcus succinus GCA_001902315.1
196 Staphylococcus lugdunensis HKU09-01 GCA_000025085.1
dR
126 Rhodobacter sphaeroides 2.4.1 GCA_000012905.2
126 Rhodobacter sphaeroides ATCC 17025 GCA_000016405.1
109 Gemmobacter aquatilis GCA_900110025.1
105 Defluviimonas alba GCA_001620265.1
dS
234 Bacillus anthracis str. Ames GCA_000007845.1
234 Bacillus anthracis str. Sterne GCA_000008165.1
234 Bacillus cereus ATCC 14579 GCA_000007825.1
234 Bacillus mycoides GCA_000832605.1
234 Bacillus pseudomycoides DSM 12442 GCA_000161455.1
234 Bacillus thuringiensis YBT-1518 GCA_000497525.2
234 _5BBacillus thuringiensis_5D serovar konkukian str. 97-27 GCA_000008505.1
196 Bacillus marisflavi GCA_001274775.1
194 Bacillus manliponensis GCA_000712595.1
dT
233 Escherichia coli IAI39 GCA_000026345.1
233 Escherichia coli O104_3AH4 str. 2011C-3493 GCA_000299455.1
233 Escherichia coli O157_3AH7 str. Sakai GCA_000008865.1
233 Escherichia coli O83_3AH1 str. NRG 857C GCA_000183345.1
233 Escherichia coli UMN026 GCA_000026325.2
233 Escherichia coli str. K-12 substr. MG1655 GCA_000005845.2
233 Shigella flexneri 2a str. 301 GCA_000006925.2
233 Tumebacillus flagellatus GCA_000714935.1
224 Shigella dysenteriae Sd197 GCA_000012005.1
191 Erwinia iniecta GCA_001267535.1
dU

255 Escherichia coli IAI39 GCA_000026345.1
255 Escherichia coli O104_3AH4 str. 2011C-3493 GCA_000299455.1
255 Escherichia coli O157_3AH7 str. Sakai GCA_000008865.1
255 Escherichia coli O83_3AH1 str. NRG 857C GCA_000183345.1
255 Escherichia coli UMN026 GCA_000026325.2
255 Escherichia coli str. K-12 substr. MG1655 GCA_000005845.2
255 Shigella flexneri 2a str. 301 GCA_000006925.2
255 Tumebacillus flagellatus GCA_000714935.1
246 Shigella dysenteriae Sd197 GCA_000012005.1
219 Erwinia iniecta GCA_001267535.1
dV
221 Clostridium beijerinckii GCA_000833105.2
221 Clostridium chromiireducens GCA_002029255.1
221 Clostridium puniceum GCA_002006345.1
221 Clostridium saccharobutylicum DSM 13864 GCA_000473995.1
221 Clostridium saccharoperbutylacetonicum N1-4_28HMT_29 GCA_000340885.1
214 Clostridium neonatale GCA_001458595.1
212 Clostridium butyricum GCA_001456065.2
dW
87 Clostridium beijerinckii GCA_000833105.2
87 Clostridium chromiireducens GCA_002029255.1
87 Clostridium puniceum GCA_002006345.1
87 Clostridium saccharobutylicum DSM 13864 GCA_000473995.1
87 Clostridium saccharoperbutylacetonicum N1-4_28HMT_29 GCA_000340885.1
74 Clostridium butyricum GCA_001456065.2
72 Clostridium botulinum B str. Eklund 17B _28NRP_29 GCA_000020165.1
72 Clostridium taeniosporum GCA_001735765.1
72 Clostridium uliginosum GCA_900112485.1
72 Clostridium ventriculi GCA_001404895.1
dX
231 Streptococcus mutans UA159 GCA_000007465.2
181 Streptococcus gordonii str. Challis substr. CH1 GCA_000017005.1
177 Streptococcus cristatus AS 1.3089 GCA_000385925.1
177 Streptococcus mitis B6 GCA_000027165.1
177 Streptococcus parasanguinis ATCC 15912 GCA_000164675.2
177 Streptococcus pneumoniae R6 GCA_000007045.1
dY
236 Deinococcus radiodurans R1 GCA_000008565.1
183 Deinococcus gobiensis I-0 GCA_000252445.1
143 Deinococcus marmoris DSM 12784 GCA_000701405.1
143 Deinococcus swuensis GCA_000800395.1
dZ
230 Staphylococcus epidermidis ATCC 12228 GCA_000007645.1
230 Staphylococcus haemolyticus JCSC1435 GCA_000009865.1
225 Staphylococcus simulans GCA_001559115.1
224 Staphylococcus capitis subsp. capitis GCA_001028645.1
e0
235 Escherichia coli IAI39 GCA_000026345.1
235 Escherichia coli O104_3AH4 str. 2011C-3493 GCA_000299455.1
235 Escherichia coli O157_3AH7 str. Sakai GCA_000008865.1
235 Escherichia coli O83_3AH1 str. NRG 857C GCA_000183345.1
235 Escherichia coli UMN026 GCA_000026325.2
235 Escherichia coli str. K-12 substr. MG1655 GCA_000005845.2
235 Shigella flexneri 2a str. 301 GCA_000006925.2
235 Tumebacillus flagellatus GCA_000714935.1
231 Erwinia iniecta GCA_001267535.1
231 Shigella dysenteriae Sd197 GCA_000012005.1
225 Cronobacter sakazakii GCA_000982825.1
225 Erwinia toletana DAPP-PG 735 GCA_000336255.1
e1
165 Clostridium beijerinckii GCA_000833105.2
165 Clostridium saccharobutylicum DSM 13864 GCA_000473995.1
165 Clostridium saccharoperbutylacetonicum N1-4_28HMT_29 GCA_000340885.1
158 Clostridium puniceum GCA_002006345.1
141 Clostridium neonatale GCA_001458595.1
e2
173 Rhodobacter sphaeroides 2.4.1 GCA_000012905.2
171 Pseudorhodobacter psychrotolerans GCA_001294535.1
150 Gemmobacter aquatilis GCA_900110025.1
e3
211 Cronobacter sakazakii GCA_000982825.1
211 Enterobacter cloacae subsp. cloacae ATCC 13047 GCA_000025565.1
211 Enterobacter hormaechei subsp. steigerwaltii GCA_001729725.1
211 Enterobacter kobei GCA_900185885.1
211 Erwinia iniecta GCA_001267535.1
211 Erwinia toletana DAPP-PG 735 GCA_000336255.1

211 Klebsiella oxytoca GCA_001022195.1
211 Pseudescherichia vulneris NBRC 102420 GCA_000759795.1
211 Salmonella enterica subsp. enterica serovar Typhi str. CT18 GCA_000195995.1
210 Escherichia coli IAI39 GCA_000026345.1
210 Escherichia coli O104_3AH4 str. 2011C-3493 GCA_000299455.1
210 Escherichia coli O157_3AH7 str. Sakai GCA_000008865.1
210 Escherichia coli O83_3AH1 str. NRG 857C GCA_000183345.1
210 Escherichia coli UMN026 GCA_000026325.2
210 Escherichia coli str. K-12 substr. MG1655 GCA_000005845.2
210 Kosakonia cowanii GCA_001975225.1
210 Kosakonia sacchari SP1 GCA_000300455.4
210 Shigella dysenteriae Sd197 GCA_000012005.1
210 Shigella flexneri 2a str. 301 GCA_000006925.2
210 Tumebacillus flagellatus GCA_000714935.1
208 Salmonella enterica subsp. enterica serovar Typhimurium str. LT2 GCA_000006945.2
e4
168 Streptococcus mutans UA159 GCA_000007465.2
139 Streptococcus sobrinus DSM 20742 = ATCC 33478 GCA_000686605.1
126 Streptococcus macacae NCTC 11558 GCA_000187995.3
e5
99 Deinococcus radiodurans R1 GCA_000008565.1
54 Deinococcus gobiensis I-0 GCA_000252445.1
45 Deinococcus deserti VCD115 GCA_000020685.1
45 Deinococcus marmoris DSM 12784 GCA_000701405.1
45 Deinococcus swuensis GCA_000800395.1
e6
143 Enterococcus faecalis V583 GCA_000007785.1
143 Streptomyces cinnamoneus GCA_001885705.1
120 Enterococcus asini ATCC 700915 GCA_000407365.1
120 Enterococcus canis NBRC 100695 GCA_001544375.1
120 Enterococcus casseliflavus EC20 GCA_000157355.2
120 Enterococcus cecorum GCA_001318405.1
120 Enterococcus columbae DSM 7374 = ATCC 51263 GCA_000406925.1
120 Enterococcus dispar ATCC 51266 GCA_000406945.1
120 Enterococcus faecium DO GCA_000174395.2
120 Enterococcus haemoperoxidus ATCC BAA-382 GCA_000407165.1
120 Enterococcus hirae ATCC 9790 GCA_000271405.2
120 Enterococcus massiliensis GCA_001050095.1
120 Enterococcus mundtii QU 25 GCA_000504125.1
120 Enterococcus phoeniculicola ATCC BAA-412 GCA_000407505.1
120 Enterococcus rivorum GCA_001742285.1
120 Enterococcus saccharolyticus subsp. saccharolyticus ATCC 43076 GCA_000407285.1
120 Enterococcus thailandicus GCA_001652875.1
115 Granulicatella balaenopterae GCA_900111135.1
e7
168 Clostridium beijerinckii GCA_000833105.2
168 Clostridium saccharobutylicum DSM 13864 GCA_000473995.1
168 Clostridium saccharoperbutylacetonicum N1-4_28HMT_29 GCA_000340885.1
149 Clostridium butyricum GCA_001456065.2
143 Clostridium puniceum GCA_002006345.1
e8
215 Streptococcus mutans UA159 GCA_000007465.2
153 Streptococcus ferus DSM 20646 GCA_000372425.1
153 Streptococcus sobrinus DSM 20742 = ATCC 33478 GCA_000686605.1
148 Streptococcus macacae NCTC 11558 GCA_000187995.3
e9
276 Deinococcus radiodurans R1 GCA_000008565.1
205 Deinococcus puniceus GCA_001644565.1
183 Deinococcus gobiensis I-0 GCA_000252445.1
ea
250 Clostridium beijerinckii GCA_000833105.2
250 Clostridium saccharobutylicum DSM 13864 GCA_000473995.1
250 Clostridium saccharoperbutylacetonicum N1-4_28HMT_29 GCA_000340885.1
243 Clostridium puniceum GCA_002006345.1
237 Clostridium butyricum GCA_001456065.2
eb
207 Deinococcus radiodurans R1 GCA_000008565.1
175 Deinococcus deserti VCD115 GCA_000020685.1
165 Deinococcus soli Cha et al. 2016 GCA_001007995.1
ec
232 Escherichia coli IAI39 GCA_000026345.1
232 Escherichia coli O104_3AH4 str. 2011C-3493 GCA_000299455.1
232 Escherichia coli O157_3AH7 str. Sakai GCA_000008865.1
232 Escherichia coli O83_3AH1 str. NRG 857C GCA_000183345.1
232 Escherichia coli UMN026 GCA_000026325.2
232 Escherichia coli str. K-12 substr. MG1655 GCA_000005845.2

232 Shigella dysenteriae Sd197 GCA_000012005.1
232 Shigella flexneri 2a str. 301 GCA_000006925.2
232 Tumebacillus flagellatus GCA_000714935.1
203 Erwinia iniecta GCA_001267535.1
198 Pantoea agglomerans GCA_001709315.1
ed
144 Clostridium beijerinckii GCA_000833105.2
144 Clostridium saccharobutylicum DSM 13864 GCA_000473995.1
144 Clostridium saccharoperbutylacetonicum N1-4_28HMT_29 GCA_000340885.1
119 Clostridium chromiireducens GCA_002029255.1
119 Clostridium puniceum GCA_002006345.1
98 Clostridium botulinum B str. Eklund 17B _28NRP_29 GCA_000020165.1
98 Clostridium butyricum GCA_001456065.2
98 Clostridium taeniosporum GCA_001735765.1
98 Clostridium thermobutyricum DSM 4928 GCA_002050515.1
ee
291 Clostridium beijerinckii GCA_000833105.2
291 Clostridium puniceum GCA_002006345.1
291 Clostridium saccharoperbutylacetonicum N1-4_28HMT_29 GCA_000340885.1
287 Clostridium saccharobutylicum DSM 13864 GCA_000473995.1
276 Clostridium chromiireducens GCA_002029255.1
ef
244 Bifidobacterium adolescentis ATCC 15703 GCA_000010425.1
223 Bifidobacterium angulatum DSM 20098 = JCM 7096 GCA_001025155.1
210 Bifidobacterium choerinum GCA_000741135.1
210 Bifidobacterium pseudolongum PV8-2 GCA_000800475.2
eg
138 Rhodobacter sphaeroides 2.4.1 GCA_000012905.2
120 Rhodobacter sphaeroides ATCC 17025 GCA_000016405.1
110 Gemmobacter aquatilis GCA_900110025.1
110 Pseudorhodobacter psychrotolerans GCA_001294535.1
eh
284 Clostridium butyricum GCA_001456065.2
281 Clostridium beijerinckii GCA_000833105.2
281 Clostridium saccharobutylicum DSM 13864 GCA_000473995.1
281 Clostridium saccharoperbutylacetonicum N1-4_28HMT_29 GCA_000340885.1
274 Clostridium puniceum GCA_002006345.1
ei
188 Clostridium saccharobutylicum DSM 13864 GCA_000473995.1
187 Clostridium beijerinckii GCA_000833105.2
187 Clostridium puniceum GCA_002006345.1
187 Clostridium saccharoperbutylacetonicum N1-4_28HMT_29 GCA_000340885.1
165 Clostridium chromiireducens GCA_002029255.1
165 Clostridium taeniosporum GCA_001735765.1
ej
185 Staphylococcus capitis subsp. capitis GCA_001028645.1
185 Staphylococcus epidermidis ATCC 12228 GCA_000007645.1
185 Staphylococcus haemolyticus JCSC1435 GCA_000009865.1
180 Staphylococcus condimenti GCA_001618885.1
180 Staphylococcus simulans GCA_001559115.1
174 Staphylococcus hominis subsp. hominis C80 GCA_000183685.1
ek
223 Clostridium butyricum GCA_001456065.2
219 Clostridium neonatale GCA_001458595.1
214 Clostridium beijerinckii GCA_000833105.2
214 Clostridium puniceum GCA_002006345.1
214 Clostridium saccharobutylicum DSM 13864 GCA_000473995.1
214 Clostridium saccharoperbutylacetonicum N1-4_28HMT_29 GCA_000340885.1
el
256 Clostridium beijerinckii GCA_000833105.2
256 Clostridium puniceum GCA_002006345.1
256 Clostridium saccharobutylicum DSM 13864 GCA_000473995.1
256 Clostridium saccharoperbutylacetonicum N1-4_28HMT_29 GCA_000340885.1
246 Clostridium butyricum GCA_001456065.2
245 Clostridium chromiireducens GCA_002029255.1
em
199 Rhodobacter sphaeroides 2.4.1 GCA_000012905.2
174 Gemmobacter megaterium GCA_900156815.1
174 Pseudorhodobacter psychrotolerans GCA_001294535.1
171 Haematobacter massiliensis GCA_000740795.1
171 Pseudorhodobacter ferrugineus DSM 5888 GCA_000420745.1
171 Pseudorhodobacter wandonensis GCA_001202035.1
en
127 Clostridium beijerinckii GCA_000833105.2
127 Clostridium puniceum GCA_002006345.1
127 Clostridium saccharobutylicum DSM 13864 GCA_000473995.1

127 Clostridium saccharoperbutylacetonicum N1-4_28HMT_29 GCA_000340885.1
119 Clostridium neonatale GCA_001458595.1
116 Clostridium botulinum B str. Eklund 17B _28NRP_29 GCA_000020165.1
116 Clostridium butyricum GCA_001456065.2
116 Clostridium chromiireducens GCA_002029255.1
116 Clostridium taeniosporum GCA_001735765.1
eo
295 Streptococcus mutans UA159 GCA_000007465.2
181 Streptococcus macacae NCTC 11558 GCA_000187995.3
177 Streptococcus marimammalium DSM 18627 GCA_000380045.1
ep
82 Celeribacter baekdonensis GCA_900102315.1
82 Celeribacter marinus GCA_001308265.1
82 Halocynthiibacter arcticus GCA_000812665.2
82 Oceanicola granulosus HTCC2516 GCA_000153305.1
82 Rhodobacter sphaeroides 2.4.1 GCA_000012905.2
81 Loktanella fryxellensis GCA_900110065.1
81 Loktanella salsilacus GCA_900114485.1
81 Roseovarius lutimaris GCA_900115165.1
81 Sulfitobacter pseudonitzschiae GCA_002222635.1
81 Thioclava dalianensis GCA_000715505.1
81 Thioclava indica GCA_000714545.1
69 Ahrensia kielensis DSM 5890 GCA_000374465.1
69 Aquamicrobium aerolatum DSM 21857 GCA_900113935.1
69 Celeribacter ethanolicus GCA_001550095.1
69 Celeribacter halophilus GCA_900114135.1
69 Celeribacter neptunius GCA_900113955.1
69 Defluviimonas alba GCA_001620265.1
69 Haematobacter massiliensis GCA_000740795.1
69 Loktanella koreensis GCA_900109295.1
69 Octadecabacter arcticus 238 GCA_000155735.2
69 Planktomarina temperata RCA23 GCA_000738435.1
69 Planktotalea frisia GCA_001890925.1
69 Pseudaminobacter salicylatoxidans KCT001 GCA_000304395.1
69 Pseudoruegeria sabulilitoris GCA_001558155.1
69 Thalassobius gelatinovorus GCA_001458355.1
eq
240 Bifidobacterium adolescentis ATCC 15703 GCA_000010425.1
205 Bifidobacterium dentium JCM 1195 = DSM 20436 GCA_001042595.1
203 Bifidobacterium thermophilum GCA_000741495.1
er
329 Bacillus anthracis str. Ames GCA_000007845.1
329 Bacillus anthracis str. Sterne GCA_000008165.1
329 Bacillus cereus ATCC 14579 GCA_000007825.1
329 Bacillus mycoides GCA_000832605.1
329 Bacillus pseudomycoides DSM 12442 GCA_000161455.1
329 Bacillus thuringiensis YBT-1518 GCA_000497525.2
329 _5BBacillus thuringiensis_5D serovar konkukian str. 97-27 GCA_000008505.1
256 Bacillus manliponensis GCA_000712595.1
230 Bacillus marisflavi GCA_001274775.1
es
285 Deinococcus radiodurans R1 GCA_000008565.1
182 Deinococcus marmoris DSM 12784 GCA_000701405.1
182 Deinococcus swuensis GCA_000800395.1
162 Deinococcus frigens DSM 12807 GCA_000701425.1
et
238 Escherichia coli IAI39 GCA_000026345.1
238 Escherichia coli O104_3AH4 str. 2011C-3493 GCA_000299455.1
238 Escherichia coli O157_3AH7 str. Sakai GCA_000008865.1
238 Escherichia coli O83_3AH1 str. NRG 857C GCA_000183345.1
238 Escherichia coli UMN026 GCA_000026325.2
238 Escherichia coli str. K-12 substr. MG1655 GCA_000005845.2
238 Shigella flexneri 2a str. 301 GCA_000006925.2
238 Tumebacillus flagellatus GCA_000714935.1
229 Shigella dysenteriae Sd197 GCA_000012005.1
221 Erwinia iniecta GCA_001267535.1
eu
245 Lactobacillus gasseri ATCC 33323 = JCM 1131 GCA_000014425.1
245 Lactobacillus hominis DSM 23910 = CRBIP 24.179 GCA_000296835.1
204 Lactobacillus iners DSM 13335 GCA_000160875.1
196 Lactobacillus amylovorus GCA_000191545.1
196 Lactobacillus crispatus ST1 GCA_000091765.1
196 Lactobacillus helveticus GCA_001308285.1
196 Lactobacillus kalixensis DSM 16043 GCA_001434335.1
196 Lactobacillus psittaci DSM 15354 GCA_000425905.1
196 Pediococcus pentosaceus ATCC 25745 GCA_000014505.1

196 Pediococcus stilesii GCA_001437075.1
ev
213 Clostridium beijerinckii GCA_000833105.2
213 Clostridium butyricum GCA_001456065.2
213 Clostridium puniceum GCA_002006345.1
213 Clostridium saccharoperbutylacetonicum N1-4_28HMT_29 GCA_000340885.1
205 Clostridium chromiireducens GCA_002029255.1
205 Clostridium saccharobutylicum DSM 13864 GCA_000473995.1
194 Clostridium taeniosporum GCA_001735765.1
ew
352 Staphylococcus epidermidis ATCC 12228 GCA_000007645.1
347 Staphylococcus capitis subsp. capitis GCA_001028645.1
347 Staphylococcus lugdunensis HKU09-01 GCA_000025085.1
337 Staphylococcus haemolyticus JCSC1435 GCA_000009865.1
337 Staphylococcus hominis subsp. hominis C80 GCA_000183685.1
ex
191 Escherichia coli IAI39 GCA_000026345.1
191 Escherichia coli O104_3AH4 str. 2011C-3493 GCA_000299455.1
191 Escherichia coli O157_3AH7 str. Sakai GCA_000008865.1
191 Escherichia coli O83_3AH1 str. NRG 857C GCA_000183345.1
191 Escherichia coli UMN026 GCA_000026325.2
191 Escherichia coli str. K-12 substr. MG1655 GCA_000005845.2
191 Shigella dysenteriae Sd197 GCA_000012005.1
191 Shigella flexneri 2a str. 301 GCA_000006925.2
191 Tumebacillus flagellatus GCA_000714935.1
177 Erwinia iniecta GCA_001267535.1
175 _5BEnterobacter_5D lignolyticus SCF1 GCA_000164865.1
ey
219 Deinococcus radiodurans R1 GCA_000008565.1
173 Deinococcus puniceus GCA_001644565.1
171 Deinococcus deserti VCD115 GCA_000020685.1
ez
182 Rhodobacter sphaeroides 2.4.1 GCA_000012905.2
164 Rhodobacter sphaeroides ATCC 17025 GCA_000016405.1
155 Gemmobacter megaterium GCA_900156815.1
eA
138 Deinococcus radiodurans R1 GCA_000008565.1
109 Deinococcus puniceus GCA_001644565.1
108 Deinococcus deserti VCD115 GCA_000020685.1
eB
225 Streptococcus mutans UA159 GCA_000007465.2
150 Streptococcus equinus GCA_000964315.1
150 Streptococcus gallolyticus subsp. gallolyticus DSM 16831 GCA_002000985.1
147 Streptococcus ratti FA-1 = DSM 20564 GCA_000286075.1
eC
294 Clostridium beijerinckii GCA_000833105.2
294 Clostridium puniceum GCA_002006345.1
294 Clostridium saccharoperbutylacetonicum N1-4_28HMT_29 GCA_000340885.1
286 Clostridium saccharobutylicum DSM 13864 GCA_000473995.1
275 Clostridium chromiireducens GCA_002029255.1
eD
232 Bacillus anthracis str. Ames GCA_000007845.1
232 Bacillus anthracis str. Sterne GCA_000008165.1
232 Bacillus cereus ATCC 14579 GCA_000007825.1
232 Bacillus mycoides GCA_000832605.1
232 Bacillus pseudomycoides DSM 12442 GCA_000161455.1
232 Bacillus thuringiensis YBT-1518 GCA_000497525.2
232 _5BBacillus thuringiensis_5D serovar konkukian str. 97-27 GCA_000008505.1
192 Bacillus manliponensis GCA_000712595.1
190 Bacillus eiseniae GCA_001636325.1
eE
201 Clostridium beijerinckii GCA_000833105.2
201 Clostridium saccharoperbutylacetonicum N1-4_28HMT_29 GCA_000340885.1
194 Clostridium puniceum GCA_002006345.1
192 Clostridium saccharobutylicum DSM 13864 GCA_000473995.1
eF
292 Clostridium beijerinckii GCA_000833105.2
292 Clostridium puniceum GCA_002006345.1
292 Clostridium saccharoperbutylacetonicum N1-4_28HMT_29 GCA_000340885.1
287 Clostridium saccharobutylicum DSM 13864 GCA_000473995.1
272 Clostridium chromiireducens GCA_002029255.1
eG
252 Deinococcus radiodurans R1 GCA_000008565.1
171 Deinococcus puniceus GCA_001644565.1
169 Deinococcus gobiensis I-0 GCA_000252445.1
eH

135 Clostridium beijerinckii GCA_000833105.2
135 Clostridium puniceum GCA_002006345.1
135 Clostridium saccharobutylicum DSM 13864 GCA_000473995.1
135 Clostridium saccharoperbutylacetonicum N1-4_28HMT_29 GCA_000340885.1
127 Clostridium chromiireducens GCA_002029255.1
112 Clostridium butyricum GCA_001456065.2
112 Clostridium fallax GCA_900129365.1
eI
195 Bacillus anthracis str. Ames GCA_000007845.1
195 Bacillus anthracis str. Sterne GCA_000008165.1
195 Bacillus mycoides GCA_000832605.1
195 Bacillus pseudomycoides DSM 12442 GCA_000161455.1
195 Bacillus thuringiensis YBT-1518 GCA_000497525.2
195 _5BBacillus thuringiensis_5D serovar konkukian str. 97-27 GCA_000008505.1
194 Bacillus cereus ATCC 14579 GCA_000007825.1
153 Bacillus marisflavi GCA_001274775.1
eJ
218 Clostridium beijerinckii GCA_000833105.2
218 Clostridium chromiireducens GCA_002029255.1
218 Clostridium puniceum GCA_002006345.1
218 Clostridium saccharobutylicum DSM 13864 GCA_000473995.1
218 Clostridium saccharoperbutylacetonicum N1-4_28HMT_29 GCA_000340885.1
206 Clostridium butyricum GCA_001456065.2
179 Clostridium neonatale GCA_001458595.1
eK
219 Staphylococcus epidermidis ATCC 12228 GCA_000007645.1
212 Staphylococcus capitis subsp. capitis GCA_001028645.1
209 Staphylococcus haemolyticus JCSC1435 GCA_000009865.1
eL
263 Lactobacillus gasseri ATCC 33323 = JCM 1131 GCA_000014425.1
259 Lactobacillus hominis DSM 23910 = CRBIP 24.179 GCA_000296835.1
169 Lactobacillus amylophilus DSM 20533 = JCM 1125 GCA_001936335.1
eM
194 Streptococcus mutans UA159 GCA_000007465.2
138 Streptococcus sobrinus DSM 20742 = ATCC 33478 GCA_000686605.1
137 Streptococcus marimammalium DSM 18627 GCA_000380045.1
eN
263 Lactobacillus gasseri ATCC 33323 = JCM 1131 GCA_000014425.1
254 Lactobacillus hominis DSM 23910 = CRBIP 24.179 GCA_000296835.1
189 Pediococcus pentosaceus ATCC 25745 GCA_000014505.1
189 Pediococcus stilesii GCA_001437075.1
eO
220 Bacillus anthracis str. Ames GCA_000007845.1
220 Bacillus anthracis str. Sterne GCA_000008165.1
220 Bacillus cereus ATCC 14579 GCA_000007825.1
220 Bacillus mycoides GCA_000832605.1
220 Bacillus pseudomycoides DSM 12442 GCA_000161455.1
220 Bacillus thuringiensis YBT-1518 GCA_000497525.2
220 _5BBacillus thuringiensis_5D serovar konkukian str. 97-27 GCA_000008505.1
173 Bacillus cytotoxicus NVH 391-98 GCA_000017425.1
172 Bacillus manliponensis GCA_000712595.1
172 Staphylococcus equorum GCA_001432245.1
eP
266 Enterococcus faecalis V583 GCA_000007785.1
266 Streptomyces cinnamoneus GCA_001885705.1
232 Melissococcus plutonius S1 GCA_000747585.1
230 Enterococcus aquimarinus GCA_001885765.1
230 Enterococcus asini ATCC 700915 GCA_000407365.1
230 Enterococcus canis NBRC 100695 GCA_001544375.1
230 Enterococcus dispar ATCC 51266 GCA_000406945.1
230 Enterococcus faecium DO GCA_000174395.2
230 Enterococcus gilvus ATCC BAA-350 GCA_000407545.1
230 Enterococcus haemoperoxidus ATCC BAA-382 GCA_000407165.1
230 Enterococcus hermanniensis GCA_001885945.1
230 Enterococcus hirae ATCC 9790 GCA_000271405.2
230 Enterococcus italicus DSM 15952 GCA_000185365.1
230 Enterococcus malodoratus ATCC 43197 GCA_000407185.1
230 Enterococcus massiliensis GCA_001050095.1
230 Enterococcus mundtii QU 25 GCA_000504125.1
230 Enterococcus pallens ATCC BAA-351 GCA_000407485.1
230 Enterococcus phoeniculicola ATCC BAA-412 GCA_000407505.1
230 Enterococcus pseudoavium NBRC 100491 GCA_001544295.1
230 Enterococcus rivorum GCA_001742285.1
230 Enterococcus sulfureus ATCC 49903 GCA_000407605.1
230 Enterococcus thailandicus GCA_001652875.1
eQ

247 Deinococcus radiodurans R1 GCA_000008565.1
176 Deinococcus gobiensis I-0 GCA_000252445.1
171 Deinococcus deserti VCD115 GCA_000020685.1
eR
233 Lactobacillus gasseri ATCC 33323 = JCM 1131 GCA_000014425.1
233 Lactobacillus hominis DSM 23910 = CRBIP 24.179 GCA_000296835.1
194 Lactobacillus iners DSM 13335 GCA_000160875.1
180 Lactobacillus psittaci DSM 15354 GCA_000425905.1
eS
177 Clostridium beijerinckii GCA_000833105.2
177 Clostridium puniceum GCA_002006345.1
177 Clostridium saccharoperbutylacetonicum N1-4_28HMT_29 GCA_000340885.1
169 Clostridium butyricum GCA_001456065.2
168 Clostridium saccharobutylicum DSM 13864 GCA_000473995.1
eT
216 Rhodobacter sphaeroides 2.4.1 GCA_000012905.2
181 Rhodobacter sphaeroides ATCC 17025 GCA_000016405.1
170 Gemmobacter megaterium GCA_900156815.1
eU
227 Staphylococcus capitis subsp. capitis GCA_001028645.1
227 Staphylococcus epidermidis ATCC 12228 GCA_000007645.1
227 Staphylococcus haemolyticus JCSC1435 GCA_000009865.1
227 Staphylococcus hominis subsp. hominis C80 GCA_000183685.1
227 Staphylococcus lugdunensis HKU09-01 GCA_000025085.1
227 Staphylococcus microti GCA_000934465.1
218 Staphylococcus aureus subsp. aureus NCTC 8325 GCA_000013425.1
218 Staphylococcus condimenti GCA_001618885.1
218 Staphylococcus simiae CCM 7213 GCA_000235645.2
218 Staphylococcus simulans GCA_001559115.1
214 Staphylococcus arlettae CVD059 GCA_000295715.1
214 Staphylococcus cohnii subsp. cohnii GCA_000972575.1
214 Staphylococcus saprophyticus subsp. saprophyticus ATCC 15305 GCA_000010125.1
eV
239 Staphylococcus cohnii subsp. cohnii GCA_000972575.1
239 Staphylococcus saprophyticus GCA_001074355.1
236 Staphylococcus aureus subsp. aureus NCTC 8325 GCA_000013425.1
236 Staphylococcus condimenti GCA_001618885.1
236 Staphylococcus epidermidis ATCC 12228 GCA_000007645.1
236 Staphylococcus haemolyticus JCSC1435 GCA_000009865.1
236 Staphylococcus hominis subsp. hominis C80 GCA_000183685.1
236 Staphylococcus simiae CCM 7213 GCA_000235645.2
236 Staphylococcus simulans GCA_001559115.1
231 Staphylococcus capitis subsp. capitis GCA_001028645.1
eW
245 Escherichia coli IAI39 GCA_000026345.1
245 Escherichia coli O104_3AH4 str. 2011C-3493 GCA_000299455.1
245 Escherichia coli O157_3AH7 str. Sakai GCA_000008865.1
245 Escherichia coli O83_3AH1 str. NRG 857C GCA_000183345.1
245 Escherichia coli UMN026 GCA_000026325.2
245 Escherichia coli str. K-12 substr. MG1655 GCA_000005845.2
245 Shigella flexneri 2a str. 301 GCA_000006925.2
245 Tumebacillus flagellatus GCA_000714935.1
236 Shigella dysenteriae Sd197 GCA_000012005.1
230 Erwinia iniecta GCA_001267535.1
eX
203 Bacillus anthracis str. Ames GCA_000007845.1
203 Bacillus anthracis str. Sterne GCA_000008165.1
203 Bacillus cereus ATCC 14579 GCA_000007825.1
203 Bacillus mycoides GCA_000832605.1
203 Bacillus pseudomycoides DSM 12442 GCA_000161455.1
203 Bacillus thuringiensis YBT-1518 GCA_000497525.2
203 _5BBacillus thuringiensis_5D serovar konkukian str. 97-27 GCA_000008505.1
183 Bacillus manliponensis GCA_000712595.1
177 Bacillus aquimaris TF-12 GCA_001648555.1
eY
223 Clostridium beijerinckii GCA_000833105.2
223 Clostridium puniceum GCA_002006345.1
223 Clostridium saccharobutylicum DSM 13864 GCA_000473995.1
223 Clostridium saccharoperbutylacetonicum N1-4_28HMT_29 GCA_000340885.1
208 Clostridium chromiireducens GCA_002029255.1
199 Clostridium butyricum GCA_001456065.2
eZ
218 Deinococcus radiodurans R1 GCA_000008565.1
122 Deinococcus marmoris DSM 12784 GCA_000701405.1
122 Deinococcus swuensis GCA_000800395.1
116 Deinococcus deserti VCD115 GCA_000020685.1

116 Deinococcus gobiensis I-0 GCA_000252445.1
f0
211 Lactobacillus gasseri ATCC 33323 = JCM 1131 GCA_000014425.1
190 Lactobacillus hominis DSM 23910 = CRBIP 24.179 GCA_000296835.1
173 Lactobacillus helveticus GCA_001308285.1
f1
210 Escherichia coli IAI39 GCA_000026345.1
210 Escherichia coli O104_3AH4 str. 2011C-3493 GCA_000299455.1
210 Escherichia coli O157_3AH7 str. Sakai GCA_000008865.1
210 Escherichia coli O83_3AH1 str. NRG 857C GCA_000183345.1
210 Escherichia coli UMN026 GCA_000026325.2
210 Escherichia coli str. K-12 substr. MG1655 GCA_000005845.2
210 Shigella dysenteriae Sd197 GCA_000012005.1
210 Shigella flexneri 2a str. 301 GCA_000006925.2
210 Tumebacillus flagellatus GCA_000714935.1
186 Erwinia iniecta GCA_001267535.1
180 Cronobacter sakazakii GCA_000982825.1
180 Erwinia gerundensis GCA_001517405.1
180 Erwinia toletana DAPP-PG 735 GCA_000336255.1
180 Pantoea agglomerans GCA_001709315.1
180 Pantoea ananatis LMG 20103 GCA_000025405.2
180 Pantoea dispersa EGD-AAK13 GCA_000465555.2
180 Shimwellia blattae DSM 4481 = NBRC 105725 GCA_000262305.1
f2
133 Rhodobacter sphaeroides 2.4.1 GCA_000012905.2
133 Rhodobacter sphaeroides ATCC 17025 GCA_000016405.1
126 Gemmobacter aquatilis GCA_900110025.1
119 Rhodobacter capsulatus SB 1003 GCA_000021865.1
f3
137 Streptococcus mutans UA159 GCA_000007465.2
117 Streptococcus salivarius GCA_000785515.1
117 Streptococcus thermophilus JIM 8232 GCA_000253395.1
115 Streptococcus macacae NCTC 11558 GCA_000187995.3
f4
144 Escherichia coli IAI39 GCA_000026345.1
144 Escherichia coli O104_3AH4 str. 2011C-3493 GCA_000299455.1
144 Escherichia coli O157_3AH7 str. Sakai GCA_000008865.1
144 Escherichia coli O83_3AH1 str. NRG 857C GCA_000183345.1
144 Escherichia coli UMN026 GCA_000026325.2
144 Escherichia coli str. K-12 substr. MG1655 GCA_000005845.2
144 Shigella dysenteriae Sd197 GCA_000012005.1
144 Shigella flexneri 2a str. 301 GCA_000006925.2
144 Tumebacillus flagellatus GCA_000714935.1
131 Buttiauxella ferragutiae ATCC 51602 GCA_001654915.1
131 Cedecea neteri GCA_000757825.1
131 Citrobacter freundii CFNIH1 GCA_000648515.1
131 Kluyvera ascorbata ATCC 33433 GCA_000735365.1
131 Serratia marcescens subsp. marcescens Db11 GCA_000513215.1
131 _5BEnterobacter_5D lignolyticus SCF1 GCA_000164865.1
130 Klebsiella aerogenes KCTC 2190 GCA_000215745.1
130 Yokenella regensburgei ATCC 49455 GCA_000735455.1
f5
175 Deinococcus radiodurans R1 GCA_000008565.1
131 Deinococcus deserti VCD115 GCA_000020685.1
131 Deinococcus gobiensis I-0 GCA_000252445.1
108 Deinococcus marmoris DSM 12784 GCA_000701405.1
108 Deinococcus soli Cha et al. 2016 GCA_001007995.1
108 Deinococcus swuensis GCA_000800395.1
f6
227 Lactobacillus gasseri ATCC 33323 = JCM 1131 GCA_000014425.1
227 Lactobacillus hominis DSM 23910 = CRBIP 24.179 GCA_000296835.1
182 Lactobacillus psittaci DSM 15354 GCA_000425905.1
167 Lactobacillus amylovorus GCA_000191545.1
167 Lactobacillus crispatus ST1 GCA_000091765.1
f7
249 Streptococcus mutans UA159 GCA_000007465.2
182 Streptococcus iniae GCA_000831485.1
171 Streptococcus massiliensis DSM 18628 GCA_000380065.1
f8
351 Clostridium beijerinckii GCA_000833105.2
351 Clostridium saccharoperbutylacetonicum N1-4_28HMT_29 GCA_000340885.1
347 Clostridium puniceum GCA_002006345.1
343 Clostridium saccharobutylicum DSM 13864 GCA_000473995.1
f9
150 Clostridium butyricum GCA_001456065.2
135 Clostridium beijerinckii GCA_000833105.2

135 Clostridium chromiireducens GCA_002029255.1
135 Clostridium puniceum GCA_002006345.1
135 Clostridium saccharobutylicum DSM 13864 GCA_000473995.1
135 Clostridium saccharoperbutylacetonicum N1-4_28HMT_29 GCA_000340885.1
131 Clostridium fallax GCA_900129365.1
131 Clostridium gasigenes GCA_900104115.1
fa
183 Escherichia coli IAI39 GCA_000026345.1
183 Escherichia coli O104_3AH4 str. 2011C-3493 GCA_000299455.1
183 Escherichia coli O157_3AH7 str. Sakai GCA_000008865.1
183 Escherichia coli O83_3AH1 str. NRG 857C GCA_000183345.1
183 Escherichia coli UMN026 GCA_000026325.2
183 Escherichia coli str. K-12 substr. MG1655 GCA_000005845.2
183 Shigella dysenteriae Sd197 GCA_000012005.1
183 Shigella flexneri 2a str. 301 GCA_000006925.2
183 Tumebacillus flagellatus GCA_000714935.1
163 Erwinia iniecta GCA_001267535.1
149 Cedecea neteri GCA_000757825.1
149 Rosenbergiella nectarea GCA_900111105.1
149 Serratia marcescens subsp. marcescens Db11 GCA_000513215.1
fb
228 Streptococcus mutans UA159 GCA_000007465.2
137 Streptococcus sobrinus DSM 20742 = ATCC 33478 GCA_000686605.1
131 Streptococcus marimammalium DSM 18627 GCA_000380045.1
fc
175 Rhodobacter sphaeroides 2.4.1 GCA_000012905.2
143 Rhodobacter sphaeroides ATCC 17025 GCA_000016405.1
131 Defluviimonas alba GCA_001620265.1
fd
168 Bacillus anthracis str. Ames GCA_000007845.1
168 Bacillus anthracis str. Sterne GCA_000008165.1
168 Bacillus cereus ATCC 14579 GCA_000007825.1
168 Bacillus mycoides GCA_000832605.1
168 Bacillus pseudomycoides DSM 12442 GCA_000161455.1
168 Bacillus thuringiensis YBT-1518 GCA_000497525.2
168 _5BBacillus thuringiensis_5D serovar konkukian str. 97-27 GCA_000008505.1
147 Bacillus manliponensis GCA_000712595.1
141 Bacillus halmapalus GCA_002019665.1
fe
291 Bacillus anthracis str. Ames GCA_000007845.1
291 Bacillus anthracis str. Sterne GCA_000008165.1
291 Bacillus cereus ATCC 14579 GCA_000007825.1
291 Bacillus mycoides GCA_000832605.1
291 Bacillus pseudomycoides DSM 12442 GCA_000161455.1
291 Bacillus thuringiensis YBT-1518 GCA_000497525.2
291 _5BBacillus thuringiensis_5D serovar konkukian str. 97-27 GCA_000008505.1
218 Bacillus manliponensis GCA_000712595.1
213 Bacillus aquimaris TF-12 GCA_001648555.1
ff
171 Clostridium puniceum GCA_002006345.1
160 Clostridium chromiireducens GCA_002029255.1
159 Clostridium neonatale GCA_001458595.1
fg
218 Staphylococcus capitis subsp. capitis GCA_001028645.1
218 Staphylococcus epidermidis ATCC 12228 GCA_000007645.1
215 Staphylococcus haemolyticus JCSC1435 GCA_000009865.1
210 Staphylococcus condimenti GCA_001618885.1
210 Staphylococcus simulans GCA_001559115.1
fh
232 Staphylococcus capitis subsp. capitis GCA_001028645.1
232 Staphylococcus epidermidis ATCC 12228 GCA_000007645.1
229 Megasphaera cerevisiae DSM 20462 GCA_001045675.1
229 Staphylococcus lugdunensis HKU09-01 GCA_000025085.1
229 Staphylococcus warneri SG1 GCA_000332735.1
225 Staphylococcus cohnii subsp. cohnii GCA_000972575.1
225 Staphylococcus equorum GCA_001432245.1
225 Staphylococcus saprophyticus subsp. saprophyticus ATCC 15305 GCA_000010125.1
225 Staphylococcus xylosus GCA_000706685.1
fi
153 Deinococcus radiodurans R1 GCA_000008565.1
110 Deinococcus deserti VCD115 GCA_000020685.1
96 Deinococcus gobiensis I-0 GCA_000252445.1
fj
83 Rhodobacter sphaeroides 2.4.1 GCA_000012905.2
83 Rhodobacter sphaeroides ATCC 17025 GCA_000016405.1
82 Deinococcus radiodurans R1 GCA_000008565.1

70 Rhodobacter capsulatus SB 1003 GCA_000021865.1
fk
222 Deinococcus radiodurans R1 GCA_000008565.1
144 Deinococcus gobiensis I-0 GCA_000252445.1
125 Deinococcus puniceus GCA_001644565.1
fl
270 Lactobacillus gasseri ATCC 33323 = JCM 1131 GCA_000014425.1
237 Lactobacillus hominis DSM 23910 = CRBIP 24.179 GCA_000296835.1
200 Lactobacillus psittaci DSM 15354 GCA_000425905.1
fm
123 _5BEnterobacter_5D lignolyticus SCF1 GCA_000164865.1
114 Buttiauxella ferragutiae ATCC 51602 GCA_001654915.1
114 Cedecea neteri GCA_000757825.1
114 Citrobacter freundii CFNIH1 GCA_000648515.1
114 Klebsiella aerogenes KCTC 2190 GCA_000215745.1
114 Kluyvera ascorbata ATCC 33433 GCA_000735365.1
114 Serratia marcescens subsp. marcescens Db11 GCA_000513215.1
114 Yokenella regensburgei ATCC 49455 GCA_000735455.1
102 Tatumella saanichensis GCA_000439375.1
fn
141 Deinococcus radiodurans R1 GCA_000008565.1
70 Deinococcus deserti VCD115 GCA_000020685.1
60 Deinococcus marmoris DSM 12784 GCA_000701405.1
60 Deinococcus swuensis GCA_000800395.1
fo
219 Rhodobacter sphaeroides 2.4.1 GCA_000012905.2
185 Gemmobacter megaterium GCA_900156815.1
176 Pseudorhodobacter ferrugineus DSM 5888 GCA_000420745.1
176 Pseudorhodobacter wandonensis GCA_001202035.1
fp
278 Bacillus anthracis str. Ames GCA_000007845.1
278 Bacillus anthracis str. Sterne GCA_000008165.1
278 Bacillus cereus ATCC 14579 GCA_000007825.1
278 Bacillus mycoides GCA_000832605.1
278 Bacillus pseudomycoides DSM 12442 GCA_000161455.1
278 Bacillus thuringiensis YBT-1518 GCA_000497525.2
278 _5BBacillus thuringiensis_5D serovar konkukian str. 97-27 GCA_000008505.1
249 Bacillus manliponensis GCA_000712595.1
242 Bacillus cytotoxicus NVH 391-98 GCA_000017425.1
fq
162 Escherichia coli IAI39 GCA_000026345.1
162 Escherichia coli O104_3AH4 str. 2011C-3493 GCA_000299455.1
162 Escherichia coli O157_3AH7 str. Sakai GCA_000008865.1
162 Escherichia coli O83_3AH1 str. NRG 857C GCA_000183345.1
162 Escherichia coli UMN026 GCA_000026325.2
162 Escherichia coli str. K-12 substr. MG1655 GCA_000005845.2
162 Shigella dysenteriae Sd197 GCA_000012005.1
162 Shigella flexneri 2a str. 301 GCA_000006925.2
162 Tumebacillus flagellatus GCA_000714935.1
143 Cronobacter sakazakii GCA_000982825.1
143 Erwinia iniecta GCA_001267535.1
143 Erwinia toletana DAPP-PG 735 GCA_000336255.1
142 Xenorhabdus bovienii SS-2004 GCA_000027225.1
fr
147 Rhodobacter sphaeroides 2.4.1 GCA_000012905.2
147 Rhodobacter sphaeroides ATCC 17025 GCA_000016405.1
141 Gemmobacter megaterium GCA_900156815.1
141 Pseudorhodobacter ferrugineus DSM 5888 GCA_000420745.1
141 Pseudorhodobacter wandonensis GCA_001202035.1
137 Defluviimonas alba GCA_001620265.1
fs
190 Deinococcus radiodurans R1 GCA_000008565.1
127 Deinococcus gobiensis I-0 GCA_000252445.1
110 Deinococcus frigens DSM 12807 GCA_000701425.1
110 Deinococcus marmoris DSM 12784 GCA_000701405.1
110 Deinococcus swuensis GCA_000800395.1
ft
331 Bifidobacterium adolescentis ATCC 15703 GCA_000010425.1
302 Bifidobacterium tsurumiense GCA_000741765.1
284 Bifidobacterium dentium JCM 1195 = DSM 20436 GCA_001042595.1
fu
200 Escherichia coli IAI39 GCA_000026345.1
200 Escherichia coli O104_3AH4 str. 2011C-3493 GCA_000299455.1
200 Escherichia coli O157_3AH7 str. Sakai GCA_000008865.1
200 Escherichia coli O83_3AH1 str. NRG 857C GCA_000183345.1
200 Escherichia coli UMN026 GCA_000026325.2

200 Escherichia coli str. K-12 substr. MG1655 GCA_000005845.2
200 Shigella dysenteriae Sd197 GCA_000012005.1
200 Shigella flexneri 2a str. 301 GCA_000006925.2
200 Tumebacillus flagellatus GCA_000714935.1
187 Erwinia iniecta GCA_001267535.1
162 Erwinia toletana DAPP-PG 735 GCA_000336255.1
162 Pantoea agglomerans GCA_001709315.1
162 Pantoea ananatis LMG 20103 GCA_000025405.2
162 Shimwellia blattae DSM 4481 = NBRC 105725 GCA_000262305.1
fv
260 Escherichia coli IAI39 GCA_000026345.1
260 Escherichia coli O104_3AH4 str. 2011C-3493 GCA_000299455.1
260 Escherichia coli O157_3AH7 str. Sakai GCA_000008865.1
260 Escherichia coli O83_3AH1 str. NRG 857C GCA_000183345.1
260 Escherichia coli UMN026 GCA_000026325.2
260 Escherichia coli str. K-12 substr. MG1655 GCA_000005845.2
260 Shigella dysenteriae Sd197 GCA_000012005.1
260 Shigella flexneri 2a str. 301 GCA_000006925.2
260 Tumebacillus flagellatus GCA_000714935.1
243 Erwinia iniecta GCA_001267535.1
229 Pantoea agglomerans GCA_001709315.1
fw
184 Deinococcus radiodurans R1 GCA_000008565.1
115 Deinococcus deserti VCD115 GCA_000020685.1
115 Deinococcus gobiensis I-0 GCA_000252445.1
100 Deinococcus soli Cha et al. 2016 GCA_001007995.1
fx
307 Bacillus anthracis str. Ames GCA_000007845.1
307 Bacillus anthracis str. Sterne GCA_000008165.1
307 Bacillus cereus ATCC 14579 GCA_000007825.1
307 Bacillus mycoides GCA_000832605.1
307 Bacillus pseudomycoides DSM 12442 GCA_000161455.1
307 Bacillus thuringiensis YBT-1518 GCA_000497525.2
307 _5BBacillus thuringiensis_5D serovar konkukian str. 97-27 GCA_000008505.1
251 Bacillus horneckiae GCA_001636335.1
251 Bacillus solani GCA_001420595.1
250 Bacillus manliponensis GCA_000712595.1
fy
142 Rhodobacter sphaeroides 2.4.1 GCA_000012905.2
133 Haematobacter massiliensis GCA_000740795.1
115 Gemmobacter aquatilis GCA_900110025.1
115 Gemmobacter nectariphilus DSM 15620 GCA_000429765.1
115 Pseudorhodobacter psychrotolerans GCA_001294535.1
115 Rhodobacter capsulatus SB 1003 GCA_000021865.1
fz
300 Enterococcus faecalis V583 GCA_000007785.1
300 Streptomyces cinnamoneus GCA_001885705.1
272 Enterococcus asini ATCC 700915 GCA_000407365.1
272 Enterococcus canis NBRC 100695 GCA_001544375.1
272 Enterococcus dispar ATCC 51266 GCA_000406945.1
272 Enterococcus faecium DO GCA_000174395.2
272 Enterococcus gilvus ATCC BAA-350 GCA_000407545.1
272 Enterococcus hermanniensis GCA_001885945.1
272 Enterococcus hirae ATCC 9790 GCA_000271405.2
272 Enterococcus malodoratus ATCC 43197 GCA_000407185.1
272 Enterococcus mundtii QU 25 GCA_000504125.1
272 Enterococcus pallens ATCC BAA-351 GCA_000407485.1
272 Enterococcus pseudoavium NBRC 100491 GCA_001544295.1
272 Enterococcus rivorum GCA_001742285.1
267 Enterococcus haemoperoxidus ATCC BAA-382 GCA_000407165.1
267 Enterococcus italicus DSM 15952 GCA_000185365.1
267 Enterococcus phoeniculicola ATCC BAA-412 GCA_000407505.1
267 Enterococcus thailandicus GCA_001652875.1
fA
279 Clostridium beijerinckii GCA_000833105.2
279 Clostridium puniceum GCA_002006345.1
279 Clostridium saccharoperbutylacetonicum N1-4_28HMT_29 GCA_000340885.1
265 Clostridium saccharobutylicum DSM 13864 GCA_000473995.1
255 Clostridium butyricum GCA_001456065.2
fB
230 Deinococcus radiodurans R1 GCA_000008565.1
150 Deinococcus puniceus GCA_001644565.1
140 Deinococcus gobiensis I-0 GCA_000252445.1
fC
219 Clostridium beijerinckii GCA_000833105.2
219 Clostridium puniceum GCA_002006345.1

219 Clostridium saccharobutylicum DSM 13864 GCA_000473995.1
219 Clostridium saccharoperbutylacetonicum N1-4_28HMT_29 GCA_000340885.1
201 Clostridium neonatale GCA_001458595.1
199 Clostridium chromiireducens GCA_002029255.1
fD
223 Escherichia coli IAI39 GCA_000026345.1
223 Escherichia coli O104_3AH4 str. 2011C-3493 GCA_000299455.1
223 Escherichia coli O157_3AH7 str. Sakai GCA_000008865.1
223 Escherichia coli O83_3AH1 str. NRG 857C GCA_000183345.1
223 Escherichia coli UMN026 GCA_000026325.2
223 Escherichia coli str. K-12 substr. MG1655 GCA_000005845.2
223 Shigella flexneri 2a str. 301 GCA_000006925.2
223 Tumebacillus flagellatus GCA_000714935.1
217 Erwinia iniecta GCA_001267535.1
214 Shigella dysenteriae Sd197 GCA_000012005.1
fE
197 Staphylococcus condimenti GCA_001618885.1
197 Staphylococcus microti GCA_000934465.1
197 Staphylococcus simulans GCA_001559115.1
193 Staphylococcus pettenkoferi GCA_002208805.1
186 Staphylococcus aureus subsp. aureus NCTC 8325 GCA_000013425.1
186 Staphylococcus capitis subsp. capitis GCA_001028645.1
186 Staphylococcus epidermidis ATCC 12228 GCA_000007645.1
186 Staphylococcus haemolyticus JCSC1435 GCA_000009865.1
186 Staphylococcus hominis subsp. hominis C80 GCA_000183685.1
186 Staphylococcus lugdunensis HKU09-01 GCA_000025085.1
186 Staphylococcus simiae CCM 7213 GCA_000235645.2
fF
130 Streptococcus mutans UA159 GCA_000007465.2
112 Streptococcus sobrinus DSM 20742 = ATCC 33478 GCA_000686605.1
109 Streptococcus dysgalactiae subsp. equisimilis AC-2713 GCA_000317855.1
fG
163 Gemmobacter aquatilis GCA_900110025.1
142 Pseudorhodobacter psychrotolerans GCA_001294535.1
142 Rhodobacter sphaeroides 2.4.1 GCA_000012905.2
140 Gemmobacter megaterium GCA_900156815.1
fH
206 Enterococcus faecalis V583 GCA_000007785.1
206 Streptomyces cinnamoneus GCA_001885705.1
165 Enterococcus asini ATCC 700915 GCA_000407365.1
165 Enterococcus casseliflavus EC20 GCA_000157355.2
165 Enterococcus dispar ATCC 51266 GCA_000406945.1
165 Enterococcus massiliensis GCA_001050095.1
165 Enterococcus saccharolyticus subsp. saccharolyticus ATCC 43076 GCA_000407285.1
164 Enterococcus canis NBRC 100695 GCA_001544375.1
164 Enterococcus faecium DO GCA_000174395.2
164 Enterococcus haemoperoxidus ATCC BAA-382 GCA_000407165.1
164 Enterococcus hirae ATCC 9790 GCA_000271405.2
164 Enterococcus mundtii QU 25 GCA_000504125.1
164 Enterococcus phoeniculicola ATCC BAA-412 GCA_000407505.1
164 Enterococcus rivorum GCA_001742285.1
164 Enterococcus thailandicus GCA_001652875.1
164 Vagococcus penaei GCA_001998885.1
fI
233 Lactobacillus gasseri ATCC 33323 = JCM 1131 GCA_000014425.1
217 Lactobacillus hominis DSM 23910 = CRBIP 24.179 GCA_000296835.1
173 Lactobacillus psittaci DSM 15354 GCA_000425905.1
fJ
186 Lactobacillus gasseri ATCC 33323 = JCM 1131 GCA_000014425.1
186 Lactobacillus hominis DSM 23910 = CRBIP 24.179 GCA_000296835.1
151 Lactobacillus acidophilus NCFM GCA_000011985.1
151 Lactobacillus crispatus ST1 GCA_000091765.1
151 Lactobacillus delbrueckii subsp. bulgaricus ATCC 11842 = JCM 1002 GCA_000056065.1
151 Lactobacillus helveticus GCA_001308285.1
151 Lactobacillus kalixensis DSM 16043 GCA_001434335.1
147 Lactobacillus pasteurii DSM 23907 = CRBIP 24.76 GCA_000297025.1
fK
343 Lactobacillus gasseri ATCC 33323 = JCM 1131 GCA_000014425.1
330 Lactobacillus hominis DSM 23910 = CRBIP 24.179 GCA_000296835.1
246 Lactobacillus psittaci DSM 15354 GCA_000425905.1
fL
271 Lactobacillus gasseri ATCC 33323 = JCM 1131 GCA_000014425.1
258 Lactobacillus hominis DSM 23910 = CRBIP 24.179 GCA_000296835.1
191 Lactobacillus acidophilus NCFM GCA_000011985.1
191 Lactobacillus gallinarum GCA_001314245.2
fM

192 Rhodobacter sphaeroides 2.4.1 GCA_000012905.2
188 Defluviimonas alba GCA_001620265.1
187 Gemmobacter aquatilis GCA_900110025.1
187 Pseudorhodobacter psychrotolerans GCA_001294535.1
fN
315 Lactobacillus gasseri ATCC 33323 = JCM 1131 GCA_000014425.1
284 Lactobacillus hominis DSM 23910 = CRBIP 24.179 GCA_000296835.1
191 Lactobacillus iners DSM 13335 GCA_000160875.1
fO
230 Rhodobacter sphaeroides 2.4.1 GCA_000012905.2
217 Gemmobacter megaterium GCA_900156815.1
208 Gemmobacter aquatilis GCA_900110025.1
fP
172 Staphylococcus capitis subsp. capitis GCA_001028645.1
172 Staphylococcus epidermidis ATCC 12228 GCA_000007645.1
172 Staphylococcus haemolyticus JCSC1435 GCA_000009865.1
161 Staphylococcus hominis subsp. hominis C80 GCA_000183685.1
146 Staphylococcus aureus subsp. aureus NCTC 8325 GCA_000013425.1
146 Staphylococcus hyicus GCA_000816085.1
146 Staphylococcus lugdunensis HKU09-01 GCA_000025085.1
146 Staphylococcus simiae CCM 7213 GCA_000235645.2
fQ
207 Escherichia coli IAI39 GCA_000026345.1
207 Escherichia coli O104_3AH4 str. 2011C-3493 GCA_000299455.1
207 Escherichia coli O157_3AH7 str. Sakai GCA_000008865.1
207 Escherichia coli O83_3AH1 str. NRG 857C GCA_000183345.1
207 Escherichia coli UMN026 GCA_000026325.2
207 Escherichia coli str. K-12 substr. MG1655 GCA_000005845.2
207 Shigella flexneri 2a str. 301 GCA_000006925.2
207 Tumebacillus flagellatus GCA_000714935.1
193 Shigella dysenteriae Sd197 GCA_000012005.1
177 Erwinia iniecta GCA_001267535.1
fR
185 Rhodobacter sphaeroides 2.4.1 GCA_000012905.2
185 Rhodobacter sphaeroides ATCC 17025 GCA_000016405.1
163 Pseudorhodobacter ferrugineus DSM 5888 GCA_000420745.1
163 Pseudorhodobacter wandonensis GCA_001202035.1
158 Pseudorhodobacter psychrotolerans GCA_001294535.1
fS
236 Lactobacillus gasseri ATCC 33323 = JCM 1131 GCA_000014425.1
228 Lactobacillus hominis DSM 23910 = CRBIP 24.179 GCA_000296835.1
189 Lactobacillus acidophilus NCFM GCA_000011985.1
fT
207 Rhodobacter sphaeroides 2.4.1 GCA_000012905.2
178 Rhodobacter sphaeroides ATCC 17025 GCA_000016405.1
176 Gemmobacter aquatilis GCA_900110025.1
fU
217 Staphylococcus epidermidis ATCC 12228 GCA_000007645.1
217 Staphylococcus haemolyticus JCSC1435 GCA_000009865.1
212 Staphylococcus capitis subsp. capitis GCA_001028645.1
207 Staphylococcus hominis subsp. hominis C80 GCA_000183685.1
fV
220 Escherichia coli IAI39 GCA_000026345.1
220 Escherichia coli O104_3AH4 str. 2011C-3493 GCA_000299455.1
220 Escherichia coli O157_3AH7 str. Sakai GCA_000008865.1
220 Escherichia coli O83_3AH1 str. NRG 857C GCA_000183345.1
220 Escherichia coli UMN026 GCA_000026325.2
220 Escherichia coli str. K-12 substr. MG1655 GCA_000005845.2
220 Shigella flexneri 2a str. 301 GCA_000006925.2
220 Tumebacillus flagellatus GCA_000714935.1
206 Shigella dysenteriae Sd197 GCA_000012005.1
202 Kosakonia cowanii GCA_001975225.1
fW
212 Streptococcus mutans UA159 GCA_000007465.2
169 Streptococcus ratti FA-1 = DSM 20564 GCA_000286075.1
152 Streptococcus equinus GCA_000964315.1
152 Streptococcus gallolyticus subsp. gallolyticus DSM 16831 GCA_002000985.1
152 Streptococcus orisratti DSM 15617 GCA_000380105.1
fX
154 Megasphaera cerevisiae DSM 20462 GCA_001045675.1
154 Staphylococcus aureus subsp. aureus NCTC 8325 GCA_000013425.1
154 Staphylococcus cohnii subsp. cohnii GCA_000972575.1
154 Staphylococcus condimenti GCA_001618885.1
154 Staphylococcus epidermidis ATCC 12228 GCA_000007645.1
154 Staphylococcus haemolyticus JCSC1435 GCA_000009865.1
154 Staphylococcus hominis subsp. hominis C80 GCA_000183685.1

154 Staphylococcus hyicus GCA_000816085.1
154 Staphylococcus lugdunensis HKU09-01 GCA_000025085.1
154 Staphylococcus lutrae GCA_002101335.1
154 Staphylococcus microti GCA_000934465.1
154 Staphylococcus pseudintermedius HKU10-03 GCA_000185885.1
154 Staphylococcus saprophyticus GCA_001074355.1
154 Staphylococcus simiae CCM 7213 GCA_000235645.2
154 Staphylococcus simulans GCA_001559115.1
154 Staphylococcus warneri SG1 GCA_000332735.1
148 Staphylococcus arlettae CVD059 GCA_000295715.1
148 Staphylococcus equorum GCA_001432245.1
148 Staphylococcus gallinarum GCA_000875895.1
148 Staphylococcus saprophyticus subsp. saprophyticus ATCC 15305 GCA_000010125.1
148 Staphylococcus succinus GCA_001902315.1
148 Staphylococcus xylosus GCA_000706685.1
142 Staphylococcus capitis subsp. capitis GCA_001028645.1
fY
258 Staphylococcus epidermidis ATCC 12228 GCA_000007645.1
254 Staphylococcus lugdunensis HKU09-01 GCA_000025085.1
252 Staphylococcus capitis subsp. capitis GCA_001028645.1
fZ
256 Lactobacillus gasseri ATCC 33323 = JCM 1131 GCA_000014425.1
228 Lactobacillus hominis DSM 23910 = CRBIP 24.179 GCA_000296835.1
168 Lactobacillus acidophilus NCFM GCA_000011985.1
168 Lactobacillus crispatus ST1 GCA_000091765.1
g0
202 Enterococcus faecalis V583 GCA_000007785.1
202 Streptomyces cinnamoneus GCA_001885705.1
197 Enterococcus faecium DO GCA_000174395.2
192 Enterococcus asini ATCC 700915 GCA_000407365.1
192 Enterococcus canis NBRC 100695 GCA_001544375.1
192 Enterococcus casseliflavus EC20 GCA_000157355.2
192 Enterococcus dispar ATCC 51266 GCA_000406945.1
192 Enterococcus gilvus ATCC BAA-350 GCA_000407545.1
192 Enterococcus haemoperoxidus ATCC BAA-382 GCA_000407165.1
192 Enterococcus hermanniensis GCA_001885945.1
192 Enterococcus hirae ATCC 9790 GCA_000271405.2
192 Enterococcus malodoratus ATCC 43197 GCA_000407185.1
192 Enterococcus mundtii QU 25 GCA_000504125.1
192 Enterococcus pallens ATCC BAA-351 GCA_000407485.1
192 Enterococcus phoeniculicola ATCC BAA-412 GCA_000407505.1
192 Enterococcus pseudoavium NBRC 100491 GCA_001544295.1
192 Enterococcus rivorum GCA_001742285.1
192 Enterococcus saccharolyticus subsp. saccharolyticus ATCC 43076 GCA_000407285.1
192 Enterococcus thailandicus GCA_001652875.1
g1
124 Clostridium neonatale GCA_001458595.1
115 Clostridium beijerinckii GCA_000833105.2
115 Clostridium puniceum GCA_002006345.1
115 Clostridium saccharobutylicum DSM 13864 GCA_000473995.1
115 Clostridium saccharoperbutylacetonicum N1-4_28HMT_29 GCA_000340885.1
99 Clostridium amylolyticum GCA_900142075.1
99 Clostridium polynesiense GCA_000820705.1
g2
202 Bifidobacterium adolescentis ATCC 15703 GCA_000010425.1
180 Bifidobacterium callitrichos DSM 23973 GCA_000741175.1
168 Bifidobacterium thermophilum GCA_000741495.1
168 Bifidobacterium tsurumiense GCA_000741765.1
g3
217 Clostridium beijerinckii GCA_000833105.2
217 Clostridium puniceum GCA_002006345.1
217 Clostridium saccharoperbutylacetonicum N1-4_28HMT_29 GCA_000340885.1
213 Clostridium chromiireducens GCA_002029255.1
213 Clostridium saccharobutylicum DSM 13864 GCA_000473995.1
204 Clostridium butyricum GCA_001456065.2
g4
238 Staphylococcus lutrae GCA_002101335.1
237 Megasphaera cerevisiae DSM 20462 GCA_001045675.1
237 Staphylococcus aureus subsp. aureus NCTC 8325 GCA_000013425.1
237 Staphylococcus epidermidis ATCC 12228 GCA_000007645.1
237 Staphylococcus haemolyticus JCSC1435 GCA_000009865.1
237 Staphylococcus hominis subsp. hominis C80 GCA_000183685.1
237 Staphylococcus simiae CCM 7213 GCA_000235645.2
237 Staphylococcus warneri SG1 GCA_000332735.1
232 Staphylococcus capitis subsp. capitis GCA_001028645.1
g5

142 Escherichia coli IAI39 GCA_000026345.1
142 Escherichia coli O104_3AH4 str. 2011C-3493 GCA_000299455.1
142 Escherichia coli O157_3AH7 str. Sakai GCA_000008865.1
142 Escherichia coli UMN026 GCA_000026325.2
142 Escherichia coli str. K-12 substr. MG1655 GCA_000005845.2
142 Tumebacillus flagellatus GCA_000714935.1
134 Shigella flexneri 2a str. 301 GCA_000006925.2
132 Shigella dysenteriae Sd197 GCA_000012005.1
g6
178 Streptococcus mutans UA159 GCA_000007465.2
130 Streptococcus gordonii str. Challis substr. CH1 GCA_000017005.1
127 Streptococcus iniae GCA_000831485.1
g7
169 Deinococcus radiodurans R1 GCA_000008565.1
128 Deinococcus puniceus GCA_001644565.1
122 Deinococcus gobiensis I-0 GCA_000252445.1
g8
213 Lactobacillus gasseri ATCC 33323 = JCM 1131 GCA_000014425.1
195 Lactobacillus hominis DSM 23910 = CRBIP 24.179 GCA_000296835.1
180 Lactobacillus iners DSM 13335 GCA_000160875.1
g9
267 Bacillus anthracis str. Ames GCA_000007845.1
267 Bacillus anthracis str. Sterne GCA_000008165.1
267 Bacillus cereus ATCC 14579 GCA_000007825.1
267 Bacillus mycoides GCA_000832605.1
267 Bacillus pseudomycoides DSM 12442 GCA_000161455.1
267 Bacillus thuringiensis YBT-1518 GCA_000497525.2
267 _5BBacillus thuringiensis_5D serovar konkukian str. 97-27 GCA_000008505.1
214 Bacillus acidicola GCA_001636425.1
214 Bacillus shackletonii GCA_001420715.1
211 Bacillus cytotoxicus NVH 391-98 GCA_000017425.1
ga
139 Streptococcus mutans UA159 GCA_000007465.2
114 Streptococcus equinus GCA_000964315.1
114 Streptococcus gallolyticus subsp. gallolyticus DSM 16831 GCA_002000985.1
112 Streptococcus marimammalium DSM 18627 GCA_000380045.1
gb
172 Clostridium puniceum GCA_002006345.1
168 Clostridium beijerinckii GCA_000833105.2
168 Clostridium saccharobutylicum DSM 13864 GCA_000473995.1
168 Clostridium saccharoperbutylacetonicum N1-4_28HMT_29 GCA_000340885.1
145 Clostridium chromiireducens GCA_002029255.1
gc
63 Streptococcus mutans UA159 GCA_000007465.2
56 Enterococcus faecium DO GCA_000174395.2
48 Streptococcus ferus DSM 20646 GCA_000372425.1
48 Streptococcus macacae NCTC 11558 GCA_000187995.3
48 Streptococcus marimammalium DSM 18627 GCA_000380045.1
48 Streptococcus sobrinus DSM 20742 = ATCC 33478 GCA_000686605.1
gd
102 Bifidobacterium asteroides PRL2011 GCA_000304215.1
101 Bifidobacterium adolescentis ATCC 15703 GCA_000010425.1
92 Bifidobacterium bifidum PRL2010 GCA_000165905.1
ge
82 Rhodobacter sphaeroides 2.4.1 GCA_000012905.2
79 Defluviimonas alba GCA_001620265.1
74 Haematobacter massiliensis GCA_000740795.1
gf
258 Deinococcus radiodurans R1 GCA_000008565.1
177 Deinococcus gobiensis I-0 GCA_000252445.1
172 Deinococcus deserti VCD115 GCA_000020685.1
gg
157 Deinococcus radiodurans R1 GCA_000008565.1
124 Deinococcus gobiensis I-0 GCA_000252445.1
118 Deinococcus deserti VCD115 GCA_000020685.1
118 Deinococcus proteolyticus MRP GCA_000190555.1
gh
178 Bifidobacterium adolescentis ATCC 15703 GCA_000010425.1
173 Bifidobacterium angulatum DSM 20098 = JCM 7096 GCA_001025155.1
173 Bifidobacterium choerinum GCA_000741135.1
173 Bifidobacterium pseudolongum PV8-2 GCA_000800475.2
171 Bifidobacterium thermophilum GCA_000741495.1
gi
176 Deinococcus radiodurans R1 GCA_000008565.1
133 Deinococcus marmoris DSM 12784 GCA_000701405.1
133 Deinococcus puniceus GCA_001644565.1

133 Deinococcus swuensis GCA_000800395.1
131 Deinococcus gobiensis I-0 GCA_000252445.1
gj
134 Escherichia coli IAI39 GCA_000026345.1
134 Escherichia coli O104_3AH4 str. 2011C-3493 GCA_000299455.1
134 Escherichia coli O157_3AH7 str. Sakai GCA_000008865.1
134 Escherichia coli O83_3AH1 str. NRG 857C GCA_000183345.1
134 Escherichia coli UMN026 GCA_000026325.2
134 Escherichia coli str. K-12 substr. MG1655 GCA_000005845.2
134 Shigella flexneri 2a str. 301 GCA_000006925.2
134 Tumebacillus flagellatus GCA_000714935.1
125 Shigella dysenteriae Sd197 GCA_000012005.1
105 Bacillus cecembensis GCA_001439635.1
gk
215 Clostridium beijerinckii GCA_000833105.2
215 Clostridium puniceum GCA_002006345.1
215 Clostridium saccharoperbutylacetonicum N1-4_28HMT_29 GCA_000340885.1
213 Clostridium butyricum GCA_001456065.2
207 Clostridium saccharobutylicum DSM 13864 GCA_000473995.1
gl
289 Streptococcus mutans UA159 GCA_000007465.2
179 Streptococcus gordonii str. Challis substr. CH1 GCA_000017005.1
174 Streptococcus salivarius GCA_000785515.1
174 Streptococcus thermophilus JIM 8232 GCA_000253395.1
gm
218 Staphylococcus epidermidis ATCC 12228 GCA_000007645.1
203 Staphylococcus capitis subsp. capitis GCA_001028645.1
200 Staphylococcus haemolyticus JCSC1435 GCA_000009865.1
200 Staphylococcus lugdunensis HKU09-01 GCA_000025085.1
gn
230 Bacillus anthracis str. Ames GCA_000007845.1
230 Bacillus anthracis str. Sterne GCA_000008165.1
230 Bacillus cereus ATCC 14579 GCA_000007825.1
230 Bacillus mycoides GCA_000832605.1
230 Bacillus pseudomycoides DSM 12442 GCA_000161455.1
230 Bacillus thuringiensis YBT-1518 GCA_000497525.2
230 _5BBacillus thuringiensis_5D serovar konkukian str. 97-27 GCA_000008505.1
178 Bacillus horneckiae GCA_001636335.1
178 Bacillus solani GCA_001420595.1
177 Bacillus manliponensis GCA_000712595.1
go
137 Rhodobacter sphaeroides 2.4.1 GCA_000012905.2
122 Rhodobacter sphaeroides ATCC 17025 GCA_000016405.1
96 Aliiroseovarius sediminilitoris GCA_900109955.1
96 Maribius pelagius GCA_900110115.1
96 Oceanicola granulosus HTCC2516 GCA_000153305.1
gp
257 Escherichia coli IAI39 GCA_000026345.1
257 Escherichia coli O104_3AH4 str. 2011C-3493 GCA_000299455.1
257 Escherichia coli O157_3AH7 str. Sakai GCA_000008865.1
257 Escherichia coli O83_3AH1 str. NRG 857C GCA_000183345.1
257 Escherichia coli UMN026 GCA_000026325.2
257 Escherichia coli str. K-12 substr. MG1655 GCA_000005845.2
257 Shigella dysenteriae Sd197 GCA_000012005.1
257 Tumebacillus flagellatus GCA_000714935.1
249 Shigella flexneri 2a str. 301 GCA_000006925.2
223 Erwinia iniecta GCA_001267535.1
gq
252 Clostridium beijerinckii GCA_000833105.2
252 Clostridium saccharobutylicum DSM 13864 GCA_000473995.1
252 Clostridium saccharoperbutylacetonicum N1-4_28HMT_29 GCA_000340885.1
246 Clostridium puniceum GCA_002006345.1
231 Clostridium chromiireducens GCA_002029255.1
gr
250 Clostridium puniceum GCA_002006345.1
244 Clostridium chromiireducens GCA_002029255.1
240 Clostridium beijerinckii GCA_000833105.2
240 Clostridium saccharobutylicum DSM 13864 GCA_000473995.1
240 Clostridium saccharoperbutylacetonicum N1-4_28HMT_29 GCA_000340885.1
gs
217 Staphylococcus epidermidis ATCC 12228 GCA_000007645.1
215 Staphylococcus haemolyticus JCSC1435 GCA_000009865.1
209 Staphylococcus capitis subsp. capitis GCA_001028645.1
gt
233 Bacillus thuringiensis YBT-1518 GCA_000497525.2
218 Bacillus anthracis str. Ames GCA_000007845.1

218 Bacillus anthracis str. Sterne GCA_000008165.1
218 Bacillus cereus ATCC 14579 GCA_000007825.1
218 Bacillus mycoides GCA_000832605.1
218 Bacillus pseudomycoides DSM 12442 GCA_000161455.1
218 _5BBacillus thuringiensis_5D serovar konkukian str. 97-27 GCA_000008505.1
205 Bacillus marisflavi GCA_001274775.1
gu
215 Rhodobacter sphaeroides 2.4.1 GCA_000012905.2
203 Pseudorhodobacter ferrugineus DSM 5888 GCA_000420745.1
203 Pseudorhodobacter wandonensis GCA_001202035.1
202 Gemmobacter megaterium GCA_900156815.1
202 Pseudorhodobacter psychrotolerans GCA_001294535.1
gv
259 Deinococcus radiodurans R1 GCA_000008565.1
194 Deinococcus marmoris DSM 12784 GCA_000701405.1
194 Deinococcus swuensis GCA_000800395.1
185 Deinococcus frigens DSM 12807 GCA_000701425.1
gw
333 Enterococcus faecalis V583 GCA_000007785.1
333 Streptomyces cinnamoneus GCA_001885705.1
305 Enterococcus canis NBRC 100695 GCA_001544375.1
305 Enterococcus dispar ATCC 51266 GCA_000406945.1
305 Enterococcus faecium DO GCA_000174395.2
305 Enterococcus gilvus ATCC BAA-350 GCA_000407545.1
305 Enterococcus hermanniensis GCA_001885945.1
305 Enterococcus hirae ATCC 9790 GCA_000271405.2
305 Enterococcus malodoratus ATCC 43197 GCA_000407185.1
305 Enterococcus mundtii QU 25 GCA_000504125.1
305 Enterococcus pallens ATCC BAA-351 GCA_000407485.1
305 Enterococcus pseudoavium NBRC 100491 GCA_001544295.1
305 Enterococcus rivorum GCA_001742285.1
303 Enterococcus asini ATCC 700915 GCA_000407365.1
gx
338 Deinococcus radiodurans R1 GCA_000008565.1
228 Deinococcus gobiensis I-0 GCA_000252445.1
227 Deinococcus puniceus GCA_001644565.1
gy
263 Bacillus anthracis str. Ames GCA_000007845.1
263 Bacillus anthracis str. Sterne GCA_000008165.1
263 Bacillus cereus ATCC 14579 GCA_000007825.1
263 Bacillus mycoides GCA_000832605.1
263 Bacillus pseudomycoides DSM 12442 GCA_000161455.1
263 Bacillus thuringiensis YBT-1518 GCA_000497525.2
263 _5BBacillus thuringiensis_5D serovar konkukian str. 97-27 GCA_000008505.1
242 Bacillus manliponensis GCA_000712595.1
240 Planococcus antarcticus DSM 14505 GCA_001687565.2
gz
162 Escherichia coli IAI39 GCA_000026345.1
162 Escherichia coli O104_3AH4 str. 2011C-3493 GCA_000299455.1
162 Escherichia coli O157_3AH7 str. Sakai GCA_000008865.1
162 Escherichia coli UMN026 GCA_000026325.2
162 Escherichia coli str. K-12 substr. MG1655 GCA_000005845.2
162 Shigella dysenteriae Sd197 GCA_000012005.1
162 Shigella flexneri 2a str. 301 GCA_000006925.2
162 Tumebacillus flagellatus GCA_000714935.1
147 Escherichia coli O83_3AH1 str. NRG 857C GCA_000183345.1
142 Cronobacter sakazakii GCA_000982825.1
142 Erwinia iniecta GCA_001267535.1
142 Erwinia toletana DAPP-PG 735 GCA_000336255.1
142 Rosenbergiella nectarea GCA_900111105.1
gA
184 Bifidobacterium adolescentis ATCC 15703 GCA_000010425.1
147 Bifidobacterium asteroides PRL2011 GCA_000304215.1
139 Bifidobacterium bifidum PRL2010 GCA_000165905.1
139 Bifidobacterium breve DSM 20213 = JCM 1192 GCA_001025175.1
gB
256 Rhodobacter sphaeroides 2.4.1 GCA_000012905.2
230 Rhodobacter sphaeroides ATCC 17025 GCA_000016405.1
203 Pseudorhodobacter psychrotolerans GCA_001294535.1
gC
170 Escherichia coli IAI39 GCA_000026345.1
170 Escherichia coli O104_3AH4 str. 2011C-3493 GCA_000299455.1
170 Escherichia coli O157_3AH7 str. Sakai GCA_000008865.1
170 Escherichia coli O83_3AH1 str. NRG 857C GCA_000183345.1
170 Escherichia coli UMN026 GCA_000026325.2
170 Escherichia coli str. K-12 substr. MG1655 GCA_000005845.2

170 Shigella dysenteriae Sd197 GCA_000012005.1
170 Shigella flexneri 2a str. 301 GCA_000006925.2
170 Tumebacillus flagellatus GCA_000714935.1
147 Cronobacter sakazakii GCA_000982825.1
147 Enterobacter cloacae subsp. cloacae ATCC 13047 GCA_000025565.1
147 Enterobacter hormaechei subsp. steigerwaltii GCA_001729725.1
147 Enterobacter kobei GCA_900185885.1
147 Erwinia amylovora CFBP1430 GCA_000091565.1
147 Erwinia billingiae Eb661 GCA_000196615.1
147 Erwinia gerundensis GCA_001517405.1
147 Erwinia iniecta GCA_001267535.1
147 Erwinia toletana DAPP-PG 735 GCA_000336255.1
147 Klebsiella oxytoca GCA_001022195.1
147 Kosakonia cowanii GCA_001975225.1
147 Kosakonia sacchari SP1 GCA_000300455.4
147 Pantoea agglomerans GCA_001709315.1
147 Pantoea alhagi GCA_002101395.1
147 Pantoea ananatis LMG 20103 GCA_000025405.2
147 Pantoea dispersa EGD-AAK13 GCA_000465555.2
147 Pantoea rwandensis GCA_000759475.1
147 Pantoea septica GCA_002095575.1
147 Pluralibacter gergoviae GCA_000757785.1
147 Pseudescherichia vulneris NBRC 102420 GCA_000759795.1
147 Salmonella enterica subsp. enterica serovar Typhi str. CT18 GCA_000195995.1
147 Salmonella enterica subsp. enterica serovar Typhimurium str. LT2 GCA_000006945.2
147 Shimwellia blattae DSM 4481 = NBRC 105725 GCA_000262305.1
147 Trabulsiella odontotermitis GCA_001297765.1
135 Bacillus cecembensis GCA_001439635.1
gD
219 Staphylococcus simulans GCA_001559115.1
212 Megasphaera cerevisiae DSM 20462 GCA_001045675.1
212 Staphylococcus aureus subsp. aureus NCTC 8325 GCA_000013425.1
212 Staphylococcus epidermidis ATCC 12228 GCA_000007645.1
212 Staphylococcus haemolyticus JCSC1435 GCA_000009865.1
212 Staphylococcus hominis subsp. hominis C80 GCA_000183685.1
212 Staphylococcus simiae CCM 7213 GCA_000235645.2
212 Staphylococcus warneri SG1 GCA_000332735.1
207 Staphylococcus capitis subsp. capitis GCA_001028645.1
gE
281 Streptococcus mutans UA159 GCA_000007465.2
185 Streptococcus sobrinus DSM 20742 = ATCC 33478 GCA_000686605.1
183 Streptococcus gordonii str. Challis substr. CH1 GCA_000017005.1
gF
289 Enterococcus faecalis V583 GCA_000007785.1
289 Streptomyces cinnamoneus GCA_001885705.1
256 Enterococcus asini ATCC 700915 GCA_000407365.1
256 Enterococcus canis NBRC 100695 GCA_001544375.1
256 Enterococcus casseliflavus EC20 GCA_000157355.2
256 Enterococcus dispar ATCC 51266 GCA_000406945.1
256 Enterococcus faecium DO GCA_000174395.2
256 Enterococcus hirae ATCC 9790 GCA_000271405.2
256 Enterococcus mundtii QU 25 GCA_000504125.1
256 Enterococcus rivorum GCA_001742285.1
256 Enterococcus saccharolyticus subsp. saccharolyticus ATCC 43076 GCA_000407285.1
251 Enterococcus haemoperoxidus ATCC BAA-382 GCA_000407165.1
251 Enterococcus phoeniculicola ATCC BAA-412 GCA_000407505.1
251 Enterococcus thailandicus GCA_001652875.1
gG
185 Escherichia coli IAI39 GCA_000026345.1
185 Escherichia coli O104_3AH4 str. 2011C-3493 GCA_000299455.1
185 Escherichia coli O157_3AH7 str. Sakai GCA_000008865.1
185 Escherichia coli O83_3AH1 str. NRG 857C GCA_000183345.1
185 Escherichia coli UMN026 GCA_000026325.2
185 Escherichia coli str. K-12 substr. MG1655 GCA_000005845.2
185 Tumebacillus flagellatus GCA_000714935.1
175 Shigella dysenteriae Sd197 GCA_000012005.1
175 Shigella flexneri 2a str. 301 GCA_000006925.2
152 Photorhabdus temperata subsp. thracensis GCA_001010285.1
gH
286 Enterococcus faecalis V583 GCA_000007785.1
286 Streptomyces cinnamoneus GCA_001885705.1
251 Enterococcus hirae ATCC 9790 GCA_000271405.2
248 Enterococcus asini ATCC 700915 GCA_000407365.1
248 Enterococcus canis NBRC 100695 GCA_001544375.1
248 Enterococcus dispar ATCC 51266 GCA_000406945.1
248 Enterococcus faecium DO GCA_000174395.2

248 Enterococcus haemoperoxidus ATCC BAA-382 GCA_000407165.1
248 Enterococcus mundtii QU 25 GCA_000504125.1
248 Enterococcus phoeniculicola ATCC BAA-412 GCA_000407505.1
248 Enterococcus rivorum GCA_001742285.1
248 Enterococcus thailandicus GCA_001652875.1
gI
340 Bacillus anthracis str. Ames GCA_000007845.1
340 Bacillus anthracis str. Sterne GCA_000008165.1
340 Bacillus cereus ATCC 14579 GCA_000007825.1
340 Bacillus mycoides GCA_000832605.1
340 Bacillus pseudomycoides DSM 12442 GCA_000161455.1
340 Bacillus thuringiensis YBT-1518 GCA_000497525.2
340 _5BBacillus thuringiensis_5D serovar konkukian str. 97-27 GCA_000008505.1
288 Bacillus coahuilensis m4-4 GCA_000171615.1
284 Bacillus marisflavi GCA_001274775.1
gJ
122 Deinococcus radiodurans R1 GCA_000008565.1
106 Deinococcus puniceus GCA_001644565.1
94 Deinococcus gobiensis I-0 GCA_000252445.1
gK
255 Rhodobacter sphaeroides 2.4.1 GCA_000012905.2
228 Rhodobacter sphaeroides ATCC 17025 GCA_000016405.1
226 Rhodobacter capsulatus SB 1003 GCA_000021865.1
gL
194 Clostridium beijerinckii GCA_000833105.2
194 Clostridium puniceum GCA_002006345.1
194 Clostridium saccharoperbutylacetonicum N1-4_28HMT_29 GCA_000340885.1
190 Clostridium saccharobutylicum DSM 13864 GCA_000473995.1
171 Clostridium taeniosporum GCA_001735765.1
gM
180 Bacillus anthracis str. Ames GCA_000007845.1
180 Bacillus anthracis str. Sterne GCA_000008165.1
180 Bacillus cereus ATCC 14579 GCA_000007825.1
180 Bacillus mycoides GCA_000832605.1
180 Bacillus pseudomycoides DSM 12442 GCA_000161455.1
180 Bacillus thuringiensis YBT-1518 GCA_000497525.2
180 _5BBacillus thuringiensis_5D serovar konkukian str. 97-27 GCA_000008505.1
162 Staphylococcus lentus F1142 GCA_000286395.1
162 Staphylococcus sciuri GCA_002209165.1
157 Staphylococcus vitulinus F1028 GCA_000286335.1
gN
182 Bacillus anthracis str. Ames GCA_000007845.1
182 Bacillus anthracis str. Sterne GCA_000008165.1
182 Bacillus cereus ATCC 14579 GCA_000007825.1
182 Bacillus mycoides GCA_000832605.1
182 Bacillus pseudomycoides DSM 12442 GCA_000161455.1
182 Bacillus thuringiensis YBT-1518 GCA_000497525.2
182 _5BBacillus thuringiensis_5D serovar konkukian str. 97-27 GCA_000008505.1
162 Viridibacillus arvi GCA_001274945.1
155 Bacillus aquimaris TF-12 GCA_001648555.1
155 Bacillus marisflavi GCA_001274775.1
gO
307 Deinococcus radiodurans R1 GCA_000008565.1
240 Deinococcus gobiensis I-0 GCA_000252445.1
213 Deinococcus puniceus GCA_001644565.1
gP
206 Enterococcus faecalis V583 GCA_000007785.1
206 Streptomyces cinnamoneus GCA_001885705.1
183 Enterococcus casseliflavus EC20 GCA_000157355.2
183 Enterococcus saccharolyticus subsp. saccharolyticus ATCC 43076 GCA_000407285.1
178 Catellicoccus marimammalium M35_2F04_2F3 GCA_000313915.1
gQ
261 Bacillus anthracis str. Ames GCA_000007845.1
261 Bacillus anthracis str. Sterne GCA_000008165.1
261 Bacillus cereus ATCC 14579 GCA_000007825.1
261 Bacillus mycoides GCA_000832605.1
261 Bacillus pseudomycoides DSM 12442 GCA_000161455.1
261 Bacillus thuringiensis YBT-1518 GCA_000497525.2
261 _5BBacillus thuringiensis_5D serovar konkukian str. 97-27 GCA_000008505.1
226 Bacillus manliponensis GCA_000712595.1
205 Bacillus cytotoxicus NVH 391-98 GCA_000017425.1
gR
266 Bifidobacterium adolescentis ATCC 15703 GCA_000010425.1
234 Bifidobacterium dentium JCM 1195 = DSM 20436 GCA_001042595.1
233 Bifidobacterium thermophilum GCA_000741495.1
233 Bifidobacterium thermophilum RBL67 GCA_000347695.1

gS
242 Deinococcus radiodurans R1 GCA_000008565.1
191 Deinococcus geothermalis DSM 11300 GCA_000196275.1
185 Deinococcus deserti VCD115 GCA_000020685.1
gT
298 Deinococcus radiodurans R1 GCA_000008565.1
212 Deinococcus gobiensis I-0 GCA_000252445.1
178 Deinococcus geothermalis DSM 11300 GCA_000196275.1
gU
135 Pantoea agglomerans GCA_001709315.1
134 Cronobacter sakazakii GCA_000982825.1
134 Enterobacter cloacae subsp. cloacae ATCC 13047 GCA_000025565.1
134 Enterobacter hormaechei subsp. steigerwaltii GCA_001729725.1
134 Enterobacter kobei GCA_900185885.1
134 Erwinia amylovora CFBP1430 GCA_000091565.1
134 Erwinia gerundensis GCA_001517405.1
134 Erwinia iniecta GCA_001267535.1
134 Erwinia toletana DAPP-PG 735 GCA_000336255.1
134 Escherichia coli IAI39 GCA_000026345.1
134 Escherichia coli O104_3AH4 str. 2011C-3493 GCA_000299455.1
134 Escherichia coli O157_3AH7 str. Sakai GCA_000008865.1
134 Escherichia coli O83_3AH1 str. NRG 857C GCA_000183345.1
134 Escherichia coli UMN026 GCA_000026325.2
134 Escherichia coli str. K-12 substr. MG1655 GCA_000005845.2
134 Klebsiella oxytoca GCA_001022195.1
134 Kosakonia cowanii GCA_001975225.1
134 Kosakonia sacchari SP1 GCA_000300455.4
134 Pantoea alhagi GCA_002101395.1
134 Pantoea ananatis LMG 20103 GCA_000025405.2
134 Pantoea dispersa EGD-AAK13 GCA_000465555.2
134 Pantoea rwandensis GCA_000759475.1
134 Pantoea septica GCA_002095575.1
134 Plautia stali symbiont GCA_000180175.2
134 Pluralibacter gergoviae GCA_000757785.1
134 Pseudescherichia vulneris NBRC 102420 GCA_000759795.1
134 Salmonella enterica subsp. enterica serovar Typhi str. CT18 GCA_000195995.1
134 Shigella dysenteriae Sd197 GCA_000012005.1
134 Shigella flexneri 2a str. 301 GCA_000006925.2
134 Shimwellia blattae DSM 4481 = NBRC 105725 GCA_000262305.1
134 Trabulsiella odontotermitis GCA_001297765.1
134 Tumebacillus flagellatus GCA_000714935.1
123 Salmonella enterica subsp. enterica serovar Typhimurium str. LT2 GCA_000006945.2
123 _5BEnterobacter_5D lignolyticus SCF1 GCA_000164865.1
gV
174 Deinococcus radiodurans R1 GCA_000008565.1
120 Deinococcus gobiensis I-0 GCA_000252445.1
114 Deinococcus frigens DSM 12807 GCA_000701425.1
114 Deinococcus marmoris DSM 12784 GCA_000701405.1
114 Deinococcus swuensis GCA_000800395.1
gW
243 Clostridium beijerinckii GCA_000833105.2
243 Clostridium saccharoperbutylacetonicum N1-4_28HMT_29 GCA_000340885.1
239 Clostridium puniceum GCA_002006345.1
235 Clostridium saccharobutylicum DSM 13864 GCA_000473995.1
gX
231 Rhodobacter sphaeroides 2.4.1 GCA_000012905.2
223 Pseudorhodobacter ferrugineus DSM 5888 GCA_000420745.1
223 Pseudorhodobacter wandonensis GCA_001202035.1
218 Pseudorhodobacter psychrotolerans GCA_001294535.1
gY
138 Streptococcus mutans UA159 GCA_000007465.2
87 Streptococcus salivarius GCA_000785515.1
87 Streptococcus thermophilus JIM 8232 GCA_000253395.1
85 Streptococcus gordonii str. Challis substr. CH1 GCA_000017005.1
gZ
282 Bacillus anthracis str. Ames GCA_000007845.1
282 Bacillus anthracis str. Sterne GCA_000008165.1
282 Bacillus cereus ATCC 14579 GCA_000007825.1
282 Bacillus mycoides GCA_000832605.1
282 Bacillus pseudomycoides DSM 12442 GCA_000161455.1
282 Bacillus thuringiensis YBT-1518 GCA_000497525.2
282 _5BBacillus thuringiensis_5D serovar konkukian str. 97-27 GCA_000008505.1
220 Staphylococcus saprophyticus GCA_001074355.1
215 Bacillus coahuilensis m4-4 GCA_000171615.1
h0
203 Streptococcus mutans UA159 GCA_000007465.2

117 Streptococcus gordonii str. Challis substr. CH1 GCA_000017005.1
113 Streptococcus cristatus AS 1.3089 GCA_000385925.1
113 Streptococcus parasanguinis ATCC 15912 GCA_000164675.2
h1
293 Clostridium beijerinckii GCA_000833105.2
293 Clostridium saccharoperbutylacetonicum N1-4_28HMT_29 GCA_000340885.1
289 Clostridium puniceum GCA_002006345.1
289 Clostridium saccharobutylicum DSM 13864 GCA_000473995.1
285 Clostridium chromiireducens GCA_002029255.1
h2
177 Clostridium beijerinckii GCA_000833105.2
177 Clostridium puniceum GCA_002006345.1
177 Clostridium saccharoperbutylacetonicum N1-4_28HMT_29 GCA_000340885.1
177 Clostridium taeniosporum GCA_001735765.1
172 Clostridium butyricum GCA_001456065.2
168 Clostridium botulinum B str. Eklund 17B _28NRP_29 GCA_000020165.1
168 Clostridium chromiireducens GCA_002029255.1
168 Clostridium saccharobutylicum DSM 13864 GCA_000473995.1
h3
215 Clostridium beijerinckii GCA_000833105.2
215 Clostridium puniceum GCA_002006345.1
215 Clostridium saccharobutylicum DSM 13864 GCA_000473995.1
215 Clostridium saccharoperbutylacetonicum N1-4_28HMT_29 GCA_000340885.1
200 Clostridium chromiireducens GCA_002029255.1
195 Clostridium butyricum GCA_001456065.2
h4
289 Deinococcus radiodurans R1 GCA_000008565.1
212 Deinococcus deserti VCD115 GCA_000020685.1
194 Deinococcus soli Cha et al. 2016 GCA_001007995.1
h5
151 Bifidobacterium adolescentis ATCC 15703 GCA_000010425.1
124 Bifidobacterium asteroides PRL2011 GCA_000304215.1
121 Bifidobacterium callitrichos DSM 23973 GCA_000741175.1
121 Bifidobacterium coryneforme GCA_000737865.1
121 Bifidobacterium minimum GCA_000741645.1
121 Bifidobacterium mongoliense DSM 21395 GCA_000741285.1
h6
293 Deinococcus radiodurans R1 GCA_000008565.1
204 Deinococcus puniceus GCA_001644565.1
182 Deinococcus gobiensis I-0 GCA_000252445.1
h7
253 Lactobacillus gasseri ATCC 33323 = JCM 1131 GCA_000014425.1
239 Lactobacillus hominis DSM 23910 = CRBIP 24.179 GCA_000296835.1
180 Lactobacillus acidophilus NCFM GCA_000011985.1
180 Lactobacillus crispatus ST1 GCA_000091765.1
180 Lactobacillus gallinarum GCA_001314245.2
180 Lactobacillus hamsteri DSM 5661 = JCM 6256 GCA_000615445.1
h8
333 Deinococcus radiodurans R1 GCA_000008565.1
232 Deinococcus gobiensis I-0 GCA_000252445.1
224 Deinococcus puniceus GCA_001644565.1
h9
187 Lactobacillus gasseri ATCC 33323 = JCM 1131 GCA_000014425.1
172 Lactobacillus hominis DSM 23910 = CRBIP 24.179 GCA_000296835.1
132 Lactobacillus acidophilus NCFM GCA_000011985.1
132 Lactobacillus gallinarum GCA_001314245.2
ha
134 Clostridium beijerinckii GCA_000833105.2
134 Clostridium puniceum GCA_002006345.1
134 Clostridium saccharoperbutylacetonicum N1-4_28HMT_29 GCA_000340885.1
125 Clostridium intestinale URNW GCA_000469625.2
125 Clostridium saccharobutylicum DSM 13864 GCA_000473995.1
116 Clostridium neonatale GCA_001458595.1
116 Desnuesiella massiliensis GCA_001403615.1
hb
232 Deinococcus radiodurans R1 GCA_000008565.1
160 Deinococcus gobiensis I-0 GCA_000252445.1
150 Deinococcus puniceus GCA_001644565.1
hc
253 Bacillus anthracis str. Ames GCA_000007845.1
253 Bacillus anthracis str. Sterne GCA_000008165.1
253 Bacillus cereus ATCC 14579 GCA_000007825.1
253 Bacillus mycoides GCA_000832605.1
253 Bacillus pseudomycoides DSM 12442 GCA_000161455.1
253 Bacillus thuringiensis YBT-1518 GCA_000497525.2
253 _5BBacillus thuringiensis_5D serovar konkukian str. 97-27 GCA_000008505.1

199 Massilibacterium senegalense GCA_001375675.1
194 Bacillus solimangrovi GCA_001742425.1
hd
107 Deinococcus radiodurans R1 GCA_000008565.1
82 Defluviimonas alba GCA_001620265.1
82 Pseudorhodobacter psychrotolerans GCA_001294535.1
80 Rhodobacter capsulatus SB 1003 GCA_000021865.1
he
229 Rhodobacter sphaeroides 2.4.1 GCA_000012905.2
177 Rhodobacter sphaeroides ATCC 17025 GCA_000016405.1
156 Pseudorhodobacter psychrotolerans GCA_001294535.1
hf
236 Staphylococcus capitis subsp. capitis GCA_001028645.1
236 Staphylococcus epidermidis ATCC 12228 GCA_000007645.1
231 Megasphaera cerevisiae DSM 20462 GCA_001045675.1
231 Staphylococcus warneri SG1 GCA_000332735.1
224 Staphylococcus lugdunensis HKU09-01 GCA_000025085.1
hg
220 Deinococcus radiodurans R1 GCA_000008565.1
153 Deinococcus gobiensis I-0 GCA_000252445.1
150 Deinococcus deserti VCD115 GCA_000020685.1
hh
237 Escherichia coli IAI39 GCA_000026345.1
237 Escherichia coli O104_3AH4 str. 2011C-3493 GCA_000299455.1
237 Escherichia coli O157_3AH7 str. Sakai GCA_000008865.1
237 Escherichia coli O83_3AH1 str. NRG 857C GCA_000183345.1
237 Escherichia coli UMN026 GCA_000026325.2
237 Escherichia coli str. K-12 substr. MG1655 GCA_000005845.2
237 Shigella flexneri 2a str. 301 GCA_000006925.2
237 Tumebacillus flagellatus GCA_000714935.1
222 Shigella dysenteriae Sd197 GCA_000012005.1
216 Erwinia iniecta GCA_001267535.1
hi
167 Bifidobacterium adolescentis ATCC 15703 GCA_000010425.1
167 Bifidobacterium breve DSM 20213 = JCM 1192 GCA_001025175.1
165 Bifidobacterium gallicum DSM 20093 = LMG 11596 GCA_000741205.1
159 Bifidobacterium longum NCC2705 GCA_000007525.1
hj
179 Lactobacillus gasseri ATCC 33323 = JCM 1131 GCA_000014425.1
158 Lactobacillus hominis DSM 23910 = CRBIP 24.179 GCA_000296835.1
125 Lactobacillus acidophilus NCFM GCA_000011985.1
hk
208 Clostridium beijerinckii GCA_000833105.2
208 Clostridium saccharoperbutylacetonicum N1-4_28HMT_29 GCA_000340885.1
201 Clostridium puniceum GCA_002006345.1
194 Clostridium saccharobutylicum DSM 13864 GCA_000473995.1
hl
183 Streptococcus mutans UA159 GCA_000007465.2
123 Streptococcus massiliensis DSM 18628 GCA_000380065.1
116 Lactobacillus acidophilus NCFM GCA_000011985.1
116 Lactobacillus amylovorus GCA_000191545.1
116 Lactobacillus crispatus ST1 GCA_000091765.1
116 Lactobacillus delbrueckii subsp. bulgaricus ATCC 11842 = JCM 1002 GCA_000056065.1
116 Lactobacillus hamsteri DSM 5661 = JCM 6256 GCA_000615445.1
116 Lactobacillus helveticus GCA_001308285.1
116 Lactobacillus kalixensis DSM 16043 GCA_001434335.1
hm
255 Clostridium beijerinckii GCA_000833105.2
255 Clostridium puniceum GCA_002006345.1
255 Clostridium saccharobutylicum DSM 13864 GCA_000473995.1
255 Clostridium saccharoperbutylacetonicum N1-4_28HMT_29 GCA_000340885.1
237 Clostridium chromiireducens GCA_002029255.1
226 Clostridium butyricum GCA_001456065.2
hn
209 Deinococcus radiodurans R1 GCA_000008565.1
144 Deinococcus deserti VCD115 GCA_000020685.1
142 Deinococcus puniceus GCA_001644565.1
ho
196 Deinococcus radiodurans R1 GCA_000008565.1
122 Deinococcus marmoris DSM 12784 GCA_000701405.1
122 Deinococcus swuensis GCA_000800395.1
114 Deinococcus gobiensis I-0 GCA_000252445.1
hp
335 Streptococcus mutans UA159 GCA_000007465.2
201 Streptococcus gordonii str. Challis substr. CH1 GCA_000017005.1
194 Streptococcus marimammalium DSM 18627 GCA_000380045.1

hq
256 Streptococcus mutans UA159 GCA_000007465.2
181 Streptococcus ratti FA-1 = DSM 20564 GCA_000286075.1
179 Streptococcus ferus DSM 20646 GCA_000372425.1
179 Streptococcus macacae NCTC 11558 GCA_000187995.3
179 Streptococcus sobrinus DSM 20742 = ATCC 33478 GCA_000686605.1
hr
158 Bifidobacterium choerinum GCA_000741135.1
158 Bifidobacterium pseudolongum PV8-2 GCA_000800475.2
155 Bifidobacterium thermophilum GCA_000741495.1
145 Bifidobacterium adolescentis ATCC 15703 GCA_000010425.1
145 Bifidobacterium breve DSM 20213 = JCM 1192 GCA_001025175.1
hs
289 Enterococcus faecalis V583 GCA_000007785.1
289 Streptomyces cinnamoneus GCA_001885705.1
266 Enterococcus canis NBRC 100695 GCA_001544375.1
266 Enterococcus casseliflavus EC20 GCA_000157355.2
266 Enterococcus dispar ATCC 51266 GCA_000406945.1
266 Enterococcus faecium DO GCA_000174395.2
266 Enterococcus hirae ATCC 9790 GCA_000271405.2
266 Enterococcus mundtii QU 25 GCA_000504125.1
266 Enterococcus rivorum GCA_001742285.1
266 Enterococcus saccharolyticus subsp. saccharolyticus ATCC 43076 GCA_000407285.1
261 Enterococcus gilvus ATCC BAA-350 GCA_000407545.1
261 Enterococcus haemoperoxidus ATCC BAA-382 GCA_000407165.1
261 Enterococcus hermanniensis GCA_001885945.1
261 Enterococcus malodoratus ATCC 43197 GCA_000407185.1
261 Enterococcus pallens ATCC BAA-351 GCA_000407485.1
261 Enterococcus phoeniculicola ATCC BAA-412 GCA_000407505.1
261 Enterococcus pseudoavium NBRC 100491 GCA_001544295.1
261 Enterococcus thailandicus GCA_001652875.1
ht
177 Deinococcus radiodurans R1 GCA_000008565.1
112 Deinococcus gobiensis I-0 GCA_000252445.1
93 Deinococcus deserti VCD115 GCA_000020685.1
hu
114 Clostridium saccharobutylicum DSM 13864 GCA_000473995.1
110 Clostridium butyricum GCA_001456065.2
105 Clostridium beijerinckii GCA_000833105.2
105 Clostridium saccharoperbutylacetonicum N1-4_28HMT_29 GCA_000340885.1
hv
183 Bacillus thuringiensis YBT-1518 GCA_000497525.2
177 Bacillus anthracis str. Ames GCA_000007845.1
177 Bacillus anthracis str. Sterne GCA_000008165.1
177 Bacillus cereus ATCC 14579 GCA_000007825.1
177 Bacillus mycoides GCA_000832605.1
177 Bacillus pseudomycoides DSM 12442 GCA_000161455.1
177 _5BBacillus thuringiensis_5D serovar konkukian str. 97-27 GCA_000008505.1
134 Bacillus eiseniae GCA_001636325.1
hw
218 Rhodobacter sphaeroides 2.4.1 GCA_000012905.2
190 Rhodobacter sphaeroides ATCC 17025 GCA_000016405.1
189 Pseudorhodobacter psychrotolerans GCA_001294535.1
hx
141 Deinococcus radiodurans R1 GCA_000008565.1
76 Deinococcus gobiensis I-0 GCA_000252445.1
75 Deinococcus puniceus GCA_001644565.1
hy
149 Clostridium beijerinckii GCA_000833105.2
149 Clostridium saccharobutylicum DSM 13864 GCA_000473995.1
149 Clostridium saccharoperbutylacetonicum N1-4_28HMT_29 GCA_000340885.1
143 Clostridium puniceum GCA_002006345.1
132 Clostridium chromiireducens GCA_002029255.1
132 Clostridium taeniosporum GCA_001735765.1
hz
58 Deinococcus radiodurans R1 GCA_000008565.1
45 Deinococcus geothermalis DSM 11300 GCA_000196275.1
45 Deinococcus puniceus GCA_001644565.1
hA
183 Staphylococcus epidermidis ATCC 12228 GCA_000007645.1
183 Staphylococcus haemolyticus JCSC1435 GCA_000009865.1
178 Staphylococcus capitis subsp. capitis GCA_001028645.1
172 Staphylococcus cohnii subsp. cohnii GCA_000972575.1
172 Staphylococcus hominis subsp. hominis C80 GCA_000183685.1
hB
238 Clostridium beijerinckii GCA_000833105.2

238 Clostridium saccharoperbutylacetonicum N1-4_28HMT_29 GCA_000340885.1
234 Clostridium puniceum GCA_002006345.1
234 Clostridium saccharobutylicum DSM 13864 GCA_000473995.1
219 Clostridium taeniosporum GCA_001735765.1
hC
253 Bacillus anthracis str. Ames GCA_000007845.1
253 Bacillus anthracis str. Sterne GCA_000008165.1
253 Bacillus cereus ATCC 14579 GCA_000007825.1
253 Bacillus mycoides GCA_000832605.1
253 Bacillus pseudomycoides DSM 12442 GCA_000161455.1
253 Bacillus thuringiensis YBT-1518 GCA_000497525.2
253 _5BBacillus thuringiensis_5D serovar konkukian str. 97-27 GCA_000008505.1
172 Bacillus cytotoxicus NVH 391-98 GCA_000017425.1
172 Bacillus manliponensis GCA_000712595.1
171 Viridibacillus arvi GCA_001274945.1
hD
109 Clostridium beijerinckii GCA_000833105.2
109 Clostridium butyricum GCA_001456065.2
109 Clostridium chromiireducens GCA_002029255.1
109 Clostridium puniceum GCA_002006345.1
109 Clostridium saccharobutylicum DSM 13864 GCA_000473995.1
109 Clostridium saccharoperbutylacetonicum N1-4_28HMT_29 GCA_000340885.1
100 Clostridium neonatale GCA_001458595.1
98 Clostridium thermobutyricum DSM 4928 GCA_002050515.1
hE
213 Deinococcus radiodurans R1 GCA_000008565.1
124 Deinococcus marmoris DSM 12784 GCA_000701405.1
124 Deinococcus swuensis GCA_000800395.1
120 Deinococcus gobiensis I-0 GCA_000252445.1
hF
298 Escherichia coli IAI39 GCA_000026345.1
298 Escherichia coli O104_3AH4 str. 2011C-3493 GCA_000299455.1
298 Escherichia coli O157_3AH7 str. Sakai GCA_000008865.1
298 Escherichia coli O83_3AH1 str. NRG 857C GCA_000183345.1
298 Escherichia coli UMN026 GCA_000026325.2
298 Escherichia coli str. K-12 substr. MG1655 GCA_000005845.2
298 Shigella flexneri 2a str. 301 GCA_000006925.2
298 Tumebacillus flagellatus GCA_000714935.1
289 Shigella dysenteriae Sd197 GCA_000012005.1
281 Erwinia iniecta GCA_001267535.1
hG
178 Rhodobacter sphaeroides 2.4.1 GCA_000012905.2
167 Rhodobacter sphaeroides ATCC 17025 GCA_000016405.1
157 Haematobacter massiliensis GCA_000740795.1
hH
305 Escherichia coli IAI39 GCA_000026345.1
305 Escherichia coli O104_3AH4 str. 2011C-3493 GCA_000299455.1
305 Escherichia coli O157_3AH7 str. Sakai GCA_000008865.1
305 Escherichia coli O83_3AH1 str. NRG 857C GCA_000183345.1
305 Escherichia coli UMN026 GCA_000026325.2
305 Escherichia coli str. K-12 substr. MG1655 GCA_000005845.2
305 Shigella dysenteriae Sd197 GCA_000012005.1
305 Tumebacillus flagellatus GCA_000714935.1
297 Shigella flexneri 2a str. 301 GCA_000006925.2
266 Erwinia iniecta GCA_001267535.1
hI
203 Lactobacillus gasseri ATCC 33323 = JCM 1131 GCA_000014425.1
187 Lactobacillus hominis DSM 23910 = CRBIP 24.179 GCA_000296835.1
116 Lactobacillus psittaci DSM 15354 GCA_000425905.1
hJ
262 Lactobacillus gasseri ATCC 33323 = JCM 1131 GCA_000014425.1
241 Lactobacillus hominis DSM 23910 = CRBIP 24.179 GCA_000296835.1
173 Lactobacillus iners DSM 13335 GCA_000160875.1
hK
168 Escherichia coli IAI39 GCA_000026345.1
168 Escherichia coli O104_3AH4 str. 2011C-3493 GCA_000299455.1
168 Escherichia coli O157_3AH7 str. Sakai GCA_000008865.1
168 Escherichia coli O83_3AH1 str. NRG 857C GCA_000183345.1
168 Escherichia coli UMN026 GCA_000026325.2
168 Escherichia coli str. K-12 substr. MG1655 GCA_000005845.2
168 Shigella dysenteriae Sd197 GCA_000012005.1
168 Shigella flexneri 2a str. 301 GCA_000006925.2
168 Tumebacillus flagellatus GCA_000714935.1
135 Cronobacter sakazakii GCA_000982825.1
135 Enterobacter cloacae subsp. cloacae ATCC 13047 GCA_000025565.1
135 Enterobacter hormaechei subsp. steigerwaltii GCA_001729725.1

135 Enterobacter kobei GCA_900185885.1
135 Erwinia iniecta GCA_001267535.1
135 Erwinia toletana DAPP-PG 735 GCA_000336255.1
135 Klebsiella oxytoca GCA_001022195.1
135 Kosakonia cowanii GCA_001975225.1
135 Kosakonia sacchari SP1 GCA_000300455.4
135 Pantoea alhagi GCA_002101395.1
135 Pseudescherichia vulneris NBRC 102420 GCA_000759795.1
135 Salmonella enterica subsp. enterica serovar Typhi str. CT18 GCA_000195995.1
135 Trabulsiella odontotermitis GCA_001297765.1
132 Pluralibacter gergoviae GCA_000757785.1
hL
174 Erwinia iniecta GCA_001267535.1
174 Escherichia coli IAI39 GCA_000026345.1
174 Escherichia coli O104_3AH4 str. 2011C-3493 GCA_000299455.1
174 Escherichia coli O157_3AH7 str. Sakai GCA_000008865.1
174 Escherichia coli O83_3AH1 str. NRG 857C GCA_000183345.1
174 Escherichia coli UMN026 GCA_000026325.2
174 Escherichia coli str. K-12 substr. MG1655 GCA_000005845.2
174 Shigella dysenteriae Sd197 GCA_000012005.1
174 Shigella flexneri 2a str. 301 GCA_000006925.2
174 Tumebacillus flagellatus GCA_000714935.1
160 Enterobacter hormaechei subsp. steigerwaltii GCA_001729725.1
160 Erwinia toletana DAPP-PG 735 GCA_000336255.1
160 Klebsiella oxytoca GCA_001022195.1
160 Kosakonia cowanii GCA_001975225.1
160 Pantoea alhagi GCA_002101395.1
160 Pseudescherichia vulneris NBRC 102420 GCA_000759795.1
157 Pluralibacter gergoviae GCA_000757785.1
hM
247 Streptococcus mutans UA159 GCA_000007465.2
189 Streptococcus ferus DSM 20646 GCA_000372425.1
187 Streptococcus marimammalium DSM 18627 GCA_000380045.1
hN
260 Bacillus anthracis str. Ames GCA_000007845.1
260 Bacillus anthracis str. Sterne GCA_000008165.1
260 Bacillus cereus ATCC 14579 GCA_000007825.1
260 Bacillus mycoides GCA_000832605.1
260 Bacillus pseudomycoides DSM 12442 GCA_000161455.1
260 Bacillus thuringiensis YBT-1518 GCA_000497525.2
260 _5BBacillus thuringiensis_5D serovar konkukian str. 97-27 GCA_000008505.1
221 Bacillus coahuilensis m4-4 GCA_000171615.1
220 Bacillus solani GCA_001420595.1
hO
153 Rhodobacter sphaeroides 2.4.1 GCA_000012905.2
139 Pseudorhodobacter psychrotolerans GCA_001294535.1
139 Rhodobacter sphaeroides ATCC 17025 GCA_000016405.1
125 Defluviimonas alba GCA_001620265.1
hP
237 Clostridium beijerinckii GCA_000833105.2
237 Clostridium saccharoperbutylacetonicum N1-4_28HMT_29 GCA_000340885.1
230 Clostridium puniceum GCA_002006345.1
228 Clostridium saccharobutylicum DSM 13864 GCA_000473995.1
hQ
162 Clostridium beijerinckii GCA_000833105.2
162 Clostridium puniceum GCA_002006345.1
162 Clostridium saccharoperbutylacetonicum N1-4_28HMT_29 GCA_000340885.1
151 Clostridium butyricum GCA_001456065.2
147 Clostridium saccharobutylicum DSM 13864 GCA_000473995.1
hR
177 Rhodobacter sphaeroides 2.4.1 GCA_000012905.2
150 Rhodobacter sphaeroides ATCC 17025 GCA_000016405.1
137 Thioclava dalianensis GCA_000715505.1
137 Thioclava indica GCA_000714545.1
hS
84 Pseudorhodobacter aquimaris GCA_001202025.1
84 Pseudorhodobacter ferrugineus DSM 5888 GCA_000420745.1
84 Pseudorhodobacter wandonensis GCA_001202035.1
73 Pseudorhodobacter psychrotolerans GCA_001294535.1
70 Rhodobacter sphaeroides 2.4.1 GCA_000012905.2
70 Rhodobacter sphaeroides ATCC 17025 GCA_000016405.1
hT
259 Rhodobacter sphaeroides 2.4.1 GCA_000012905.2
219 Rhodobacter sphaeroides ATCC 17025 GCA_000016405.1
215 Pseudorhodobacter psychrotolerans GCA_001294535.1
hU

153 Streptococcus mutans UA159 GCA_000007465.2
91 Streptococcus ferus DSM 20646 GCA_000372425.1
87 Lactobacillus suebicus DSM 5007 = KCTC 3549 GCA_001434475.1
87 Lactobacillus vaccinostercus DSM 20634 GCA_001436295.1
hV
263 Deinococcus radiodurans R1 GCA_000008565.1
155 Deinococcus puniceus GCA_001644565.1
154 Deinococcus gobiensis I-0 GCA_000252445.1
hW
284 Clostridium beijerinckii GCA_000833105.2
284 Clostridium saccharoperbutylacetonicum N1-4_28HMT_29 GCA_000340885.1
275 Clostridium saccharobutylicum DSM 13864 GCA_000473995.1
269 Clostridium puniceum GCA_002006345.1
hX
176 Enterococcus faecalis V583 GCA_000007785.1
176 Streptomyces cinnamoneus GCA_001885705.1
149 Enterococcus aquimarinus GCA_001885765.1
149 Enterococcus asini ATCC 700915 GCA_000407365.1
149 Enterococcus canis NBRC 100695 GCA_001544375.1
149 Enterococcus casseliflavus EC20 GCA_000157355.2
149 Enterococcus cecorum GCA_001318405.1
149 Enterococcus columbae DSM 7374 = ATCC 51263 GCA_000406925.1
149 Enterococcus dispar ATCC 51266 GCA_000406945.1
149 Enterococcus faecium DO GCA_000174395.2
149 Enterococcus gilvus ATCC BAA-350 GCA_000407545.1
149 Enterococcus haemoperoxidus ATCC BAA-382 GCA_000407165.1
149 Enterococcus hermanniensis GCA_001885945.1
149 Enterococcus hirae ATCC 9790 GCA_000271405.2
149 Enterococcus italicus DSM 15952 GCA_000185365.1
149 Enterococcus malodoratus ATCC 43197 GCA_000407185.1
149 Enterococcus massiliensis GCA_001050095.1
149 Enterococcus mundtii QU 25 GCA_000504125.1
149 Enterococcus pallens ATCC BAA-351 GCA_000407485.1
149 Enterococcus phoeniculicola ATCC BAA-412 GCA_000407505.1
149 Enterococcus pseudoavium NBRC 100491 GCA_001544295.1
149 Enterococcus rivorum GCA_001742285.1
149 Enterococcus saccharolyticus subsp. saccharolyticus ATCC 43076 GCA_000407285.1
149 Enterococcus sulfureus ATCC 49903 GCA_000407605.1
149 Enterococcus thailandicus GCA_001652875.1
149 Melissococcus plutonius S1 GCA_000747585.1
149 Vagococcus fluvialis bH819 GCA_900163795.1
149 Vagococcus lutrae LBD1 GCA_000498295.1
149 Vagococcus penaei GCA_001998885.1
149 Vagococcus teuberi GCA_001870205.1
141 Catellicoccus marimammalium M35_2F04_2F3 GCA_000313915.1
hY
194 Escherichia coli IAI39 GCA_000026345.1
194 Escherichia coli O104_3AH4 str. 2011C-3493 GCA_000299455.1
194 Escherichia coli O157_3AH7 str. Sakai GCA_000008865.1
194 Escherichia coli O83_3AH1 str. NRG 857C GCA_000183345.1
194 Escherichia coli UMN026 GCA_000026325.2
194 Escherichia coli str. K-12 substr. MG1655 GCA_000005845.2
194 Shigella flexneri 2a str. 301 GCA_000006925.2
194 Tumebacillus flagellatus GCA_000714935.1
185 Shigella dysenteriae Sd197 GCA_000012005.1
175 Erwinia iniecta GCA_001267535.1
hZ
255 Enterococcus faecalis V583 GCA_000007785.1
255 Streptomyces cinnamoneus GCA_001885705.1
224 Enterococcus dispar ATCC 51266 GCA_000406945.1
222 Enterococcus canis NBRC 100695 GCA_001544375.1
222 Enterococcus faecium DO GCA_000174395.2
222 Enterococcus haemoperoxidus ATCC BAA-382 GCA_000407165.1
222 Enterococcus hirae ATCC 9790 GCA_000271405.2
222 Enterococcus mundtii QU 25 GCA_000504125.1
222 Enterococcus phoeniculicola ATCC BAA-412 GCA_000407505.1
222 Enterococcus rivorum GCA_001742285.1
222 Enterococcus thailandicus GCA_001652875.1
i0
276 Lactobacillus gasseri ATCC 33323 = JCM 1131 GCA_000014425.1
271 Lactobacillus hominis DSM 23910 = CRBIP 24.179 GCA_000296835.1
211 Lactobacillus acidophilus NCFM GCA_000011985.1
211 Lactobacillus gallinarum GCA_001314245.2
i1
193 Deinococcus radiodurans R1 GCA_000008565.1
145 Deinococcus gobiensis I-0 GCA_000252445.1

135 Deinococcus deserti VCD115 GCA_000020685.1
i2
172 Deinococcus radiodurans R1 GCA_000008565.1
137 Deinococcus gobiensis I-0 GCA_000252445.1
134 Deinococcus marmoris DSM 12784 GCA_000701405.1
134 Deinococcus swuensis GCA_000800395.1
i3
199 Deinococcus radiodurans R1 GCA_000008565.1
144 Deinococcus deserti VCD115 GCA_000020685.1
137 Deinococcus gobiensis I-0 GCA_000252445.1
137 Deinococcus soli Cha et al. 2016 GCA_001007995.1
i4
253 Clostridium beijerinckii GCA_000833105.2
253 Clostridium saccharobutylicum DSM 13864 GCA_000473995.1
253 Clostridium saccharoperbutylacetonicum N1-4_28HMT_29 GCA_000340885.1
238 Clostridium puniceum GCA_002006345.1
232 Clostridium butyricum GCA_001456065.2
i5
161 Megasphaera cerevisiae DSM 20462 GCA_001045675.1
161 Staphylococcus aureus subsp. aureus NCTC 8325 GCA_000013425.1
161 Staphylococcus capitis subsp. capitis GCA_001028645.1
161 Staphylococcus cohnii subsp. cohnii GCA_000972575.1
161 Staphylococcus condimenti GCA_001618885.1
161 Staphylococcus epidermidis ATCC 12228 GCA_000007645.1
161 Staphylococcus haemolyticus JCSC1435 GCA_000009865.1
161 Staphylococcus hominis subsp. hominis C80 GCA_000183685.1
161 Staphylococcus hyicus GCA_000816085.1
161 Staphylococcus lutrae GCA_002101335.1
161 Staphylococcus microti GCA_000934465.1
161 Staphylococcus pseudintermedius HKU10-03 GCA_000185885.1
161 Staphylococcus saprophyticus GCA_001074355.1
161 Staphylococcus sciuri GCA_002209165.1
161 Staphylococcus simiae CCM 7213 GCA_000235645.2
161 Staphylococcus simulans GCA_001559115.1
161 Staphylococcus warneri SG1 GCA_000332735.1
146 Nosocomiicoccus massiliensis GCA_000438455.1
146 Staphylococcus arlettae CVD059 GCA_000295715.1
146 Staphylococcus gallinarum GCA_000875895.1
146 Staphylococcus lentus F1142 GCA_000286395.1
146 Staphylococcus lugdunensis HKU09-01 GCA_000025085.1
146 Staphylococcus pettenkoferi GCA_002208805.1
146 Staphylococcus saprophyticus subsp. saprophyticus ATCC 15305 GCA_000010125.1
146 Staphylococcus succinus GCA_001902315.1
146 Staphylococcus xylosus GCA_000706685.1
137 Facklamia miroungae GCA_900100775.1
i6
254 Deinococcus radiodurans R1 GCA_000008565.1
216 Deinococcus gobiensis I-0 GCA_000252445.1
176 Deinococcus marmoris DSM 12784 GCA_000701405.1
176 Deinococcus swuensis GCA_000800395.1
i7
146 _5BEnterobacter_5D lignolyticus SCF1 GCA_000164865.1
145 Cronobacter sakazakii GCA_000982825.1
145 Erwinia iniecta GCA_001267535.1
145 Erwinia toletana DAPP-PG 735 GCA_000336255.1
145 Escherichia coli IAI39 GCA_000026345.1
145 Escherichia coli O104_3AH4 str. 2011C-3493 GCA_000299455.1
145 Escherichia coli O157_3AH7 str. Sakai GCA_000008865.1
145 Escherichia coli O83_3AH1 str. NRG 857C GCA_000183345.1
145 Escherichia coli UMN026 GCA_000026325.2
145 Escherichia coli str. K-12 substr. MG1655 GCA_000005845.2
145 Kosakonia cowanii GCA_001975225.1
145 Kosakonia sacchari SP1 GCA_000300455.4
145 Mangrovibacter phragmitis GCA_001655675.1
145 Pluralibacter gergoviae GCA_000757785.1
145 Serratia marcescens subsp. marcescens Db11 GCA_000513215.1
145 Shigella flexneri 2a str. 301 GCA_000006925.2
145 Tumebacillus flagellatus GCA_000714935.1
136 Aeromonas eucrenophila GCA_000819865.1
136 Aeromonas fluvialis GCA_000819885.1
136 Aeromonas hydrophila subsp. hydrophila ATCC 7966 GCA_000014805.1
136 Aeromonas salmonicida subsp. salmonicida A449 GCA_000196395.1
136 Aeromonas schubertii GCA_001447335.1
136 Aeromonas simiae GCA_000820125.1
136 Buttiauxella ferragutiae ATCC 51602 GCA_001654915.1
136 Cedecea neteri GCA_000757825.1

136 Citrobacter freundii CFNIH1 GCA_000648515.1
136 Enterobacter cloacae subsp. cloacae ATCC 13047 GCA_000025565.1
136 Enterobacter hormaechei subsp. steigerwaltii GCA_001729725.1
136 Enterobacter kobei GCA_900185885.1
136 Erwinia persicina NBRC 102418 GCA_001571305.1
136 Klebsiella aerogenes KCTC 2190 GCA_000215745.1
136 Klebsiella oxytoca GCA_001022195.1
136 Klebsiella pneumoniae subsp. pneumoniae HS11286 GCA_000240185.2
136 Kluyvera ascorbata ATCC 33433 GCA_000735365.1
136 Kluyvera cryocrescens NBRC 102467 GCA_001571285.1
136 Pseudescherichia vulneris NBRC 102420 GCA_000759795.1
136 Rosenbergiella nectarea GCA_900111105.1
136 Salmonella enterica subsp. enterica serovar Typhi str. CT18 GCA_000195995.1
136 Shigella dysenteriae Sd197 GCA_000012005.1
136 Yokenella regensburgei ATCC 49455 GCA_000735455.1
i8
211 Clostridium beijerinckii GCA_000833105.2
211 Clostridium puniceum GCA_002006345.1
211 Clostridium saccharoperbutylacetonicum N1-4_28HMT_29 GCA_000340885.1
203 Clostridium saccharobutylicum DSM 13864 GCA_000473995.1
196 Clostridium taeniosporum GCA_001735765.1
i9
152 Bacillus anthracis str. Ames GCA_000007845.1
152 Bacillus anthracis str. Sterne GCA_000008165.1
152 Bacillus cereus ATCC 14579 GCA_000007825.1
152 Bacillus mycoides GCA_000832605.1
152 Bacillus pseudomycoides DSM 12442 GCA_000161455.1
152 Bacillus thuringiensis YBT-1518 GCA_000497525.2
152 _5BBacillus thuringiensis_5D serovar konkukian str. 97-27 GCA_000008505.1
126 Catellicoccus marimammalium M35_2F04_2F3 GCA_000313915.1
126 Enterococcus asini ATCC 700915 GCA_000407365.1
126 Enterococcus canis NBRC 100695 GCA_001544375.1
126 Enterococcus cecorum GCA_001318405.1
126 Enterococcus columbae DSM 7374 = ATCC 51263 GCA_000406925.1
126 Enterococcus dispar ATCC 51266 GCA_000406945.1
126 Enterococcus faecalis V583 GCA_000007785.1
126 Enterococcus faecium DO GCA_000174395.2
126 Enterococcus haemoperoxidus ATCC BAA-382 GCA_000407165.1
126 Enterococcus hirae ATCC 9790 GCA_000271405.2
126 Enterococcus massiliensis GCA_001050095.1
126 Enterococcus mundtii QU 25 GCA_000504125.1
126 Enterococcus phoeniculicola ATCC BAA-412 GCA_000407505.1
126 Enterococcus rivorum GCA_001742285.1
126 Enterococcus thailandicus GCA_001652875.1
126 Streptomyces cinnamoneus GCA_001885705.1
126 Vagococcus penaei GCA_001998885.1
125 Fictibacillus phosphorivorans GCA_001629705.2
ia
216 Deinococcus radiodurans R1 GCA_000008565.1
149 Deinococcus deserti VCD115 GCA_000020685.1
147 Deinococcus puniceus GCA_001644565.1
ib
235 Deinococcus radiodurans R1 GCA_000008565.1
190 Deinococcus puniceus GCA_001644565.1
179 Deinococcus marmoris DSM 12784 GCA_000701405.1
179 Deinococcus swuensis GCA_000800395.1
ic
245 Rhodobacter sphaeroides 2.4.1 GCA_000012905.2
222 Pseudorhodobacter ferrugineus DSM 5888 GCA_000420745.1
222 Pseudorhodobacter wandonensis GCA_001202035.1
220 Pseudorhodobacter psychrotolerans GCA_001294535.1
id
294 Clostridium beijerinckii GCA_000833105.2
294 Clostridium puniceum GCA_002006345.1
294 Clostridium saccharobutylicum DSM 13864 GCA_000473995.1
294 Clostridium saccharoperbutylacetonicum N1-4_28HMT_29 GCA_000340885.1
279 Clostridium chromiireducens GCA_002029255.1
270 Clostridium botulinum B str. Eklund 17B _28NRP_29 GCA_000020165.1
270 Clostridium butyricum GCA_001456065.2
270 Clostridium taeniosporum GCA_001735765.1
ie
236 Deinococcus radiodurans R1 GCA_000008565.1
162 Deinococcus gobiensis I-0 GCA_000252445.1
155 Deinococcus marmoris DSM 12784 GCA_000701405.1
155 Deinococcus swuensis GCA_000800395.1
if

293 Bacillus anthracis str. Ames GCA_000007845.1
293 Bacillus anthracis str. Sterne GCA_000008165.1
293 Bacillus cereus ATCC 14579 GCA_000007825.1
293 Bacillus mycoides GCA_000832605.1
293 Bacillus pseudomycoides DSM 12442 GCA_000161455.1
293 Bacillus thuringiensis YBT-1518 GCA_000497525.2
293 _5BBacillus thuringiensis_5D serovar konkukian str. 97-27 GCA_000008505.1
260 Bacillus manliponensis GCA_000712595.1
237 Bacillus cytotoxicus NVH 391-98 GCA_000017425.1
ig
183 Streptococcus mutans UA159 GCA_000007465.2
129 Streptococcus ferus DSM 20646 GCA_000372425.1
129 Streptococcus macacae NCTC 11558 GCA_000187995.3
127 Streptococcus iniae GCA_000831485.1
ih
235 Escherichia coli IAI39 GCA_000026345.1
235 Escherichia coli O104_3AH4 str. 2011C-3493 GCA_000299455.1
235 Escherichia coli O157_3AH7 str. Sakai GCA_000008865.1
235 Escherichia coli O83_3AH1 str. NRG 857C GCA_000183345.1
235 Escherichia coli UMN026 GCA_000026325.2
235 Escherichia coli str. K-12 substr. MG1655 GCA_000005845.2
235 Shigella dysenteriae Sd197 GCA_000012005.1
235 Shigella flexneri 2a str. 301 GCA_000006925.2
235 Tumebacillus flagellatus GCA_000714935.1
216 Erwinia iniecta GCA_001267535.1
193 Cronobacter sakazakii GCA_000982825.1
193 Erwinia toletana DAPP-PG 735 GCA_000336255.1
ii
209 Bacillus anthracis str. Ames GCA_000007845.1
209 Bacillus anthracis str. Sterne GCA_000008165.1
209 Bacillus cereus ATCC 14579 GCA_000007825.1
209 Bacillus mycoides GCA_000832605.1
209 Bacillus pseudomycoides DSM 12442 GCA_000161455.1
209 Bacillus thuringiensis YBT-1518 GCA_000497525.2
209 _5BBacillus thuringiensis_5D serovar konkukian str. 97-27 GCA_000008505.1
183 Bacillus horneckiae GCA_001636335.1
183 Bacillus solani GCA_001420595.1
159 Bacillus aquimaris TF-12 GCA_001648555.1
ij
215 Lactobacillus gasseri ATCC 33323 = JCM 1131 GCA_000014425.1
215 Lactobacillus hominis DSM 23910 = CRBIP 24.179 GCA_000296835.1
194 Lactobacillus acidophilus NCFM GCA_000011985.1
194 Lactobacillus gallinarum GCA_001314245.2
187 Lactobacillus pasteurii DSM 23907 = CRBIP 24.76 GCA_000297025.1
ik
228 Escherichia coli IAI39 GCA_000026345.1
228 Escherichia coli O104_3AH4 str. 2011C-3493 GCA_000299455.1
228 Escherichia coli O157_3AH7 str. Sakai GCA_000008865.1
228 Escherichia coli O83_3AH1 str. NRG 857C GCA_000183345.1
228 Escherichia coli UMN026 GCA_000026325.2
228 Escherichia coli str. K-12 substr. MG1655 GCA_000005845.2
228 Shigella flexneri 2a str. 301 GCA_000006925.2
228 Tumebacillus flagellatus GCA_000714935.1
221 Erwinia iniecta GCA_001267535.1
213 Shigella dysenteriae Sd197 GCA_000012005.1
il
97 Clostridium beijerinckii GCA_000833105.2
97 Clostridium puniceum GCA_002006345.1
97 Clostridium saccharobutylicum DSM 13864 GCA_000473995.1
97 Clostridium saccharoperbutylacetonicum N1-4_28HMT_29 GCA_000340885.1
89 Clostridium botulinum B str. Eklund 17B _28NRP_29 GCA_000020165.1
89 Clostridium chromiireducens GCA_002029255.1
89 Clostridium taeniosporum GCA_001735765.1
89 Clostridium uliginosum GCA_900112485.1
82 Clostridium butyricum GCA_001456065.2
im
203 Escherichia coli IAI39 GCA_000026345.1
203 Escherichia coli O104_3AH4 str. 2011C-3493 GCA_000299455.1
203 Escherichia coli O157_3AH7 str. Sakai GCA_000008865.1
203 Escherichia coli O83_3AH1 str. NRG 857C GCA_000183345.1
203 Escherichia coli UMN026 GCA_000026325.2
203 Escherichia coli str. K-12 substr. MG1655 GCA_000005845.2
203 Shigella flexneri 2a str. 301 GCA_000006925.2
203 Tumebacillus flagellatus GCA_000714935.1
188 Shigella dysenteriae Sd197 GCA_000012005.1
164 Erwinia iniecta GCA_001267535.1

in
127 Rhodobacter sphaeroides 2.4.1 GCA_000012905.2
127 Rhodobacter sphaeroides ATCC 17025 GCA_000016405.1
112 Rhodobacter capsulatus SB 1003 GCA_000021865.1
99 Actibacterium atlanticum GCA_000671395.1
99 Actibacterium mucosum KCTC 23349 GCA_000647975.1
99 Aestuariivita atlantica GCA_001205715.1
99 Aestuariivita boseongensis GCA_001262635.1
99 Ahrensia kielensis DSM 5890 GCA_000374465.1
99 Aliiroseovarius crassostreae GCA_001307765.1
99 Aliiroseovarius sediminilitoris GCA_900109955.1
99 Antarctobacter heliothermus GCA_900188425.1
99 Aquamicrobium aerolatum DSM 21857 GCA_900113935.1
99 Aquamicrobium defluvii GCA_000585625.1
99 Aquimixticola soesokkakensis GCA_900172375.1
99 Celeribacter indicus GCA_000819565.1
99 Celeribacter neptunius GCA_900113955.1
99 Citreicella marina GCA_900100085.1
99 Citreicella thiooxidans GCA_900102075.1
99 Citreimonas salinaria GCA_900107235.1
99 Cribrihabitans marinus GCA_900109035.1
99 Defluviimonas alba GCA_001620265.1
99 Defluviimonas indica GCA_900106675.1
99 Dinoroseobacter shibae DFL 12 = DSM 16493 GCA_000018145.1
99 Donghicola eburneus GCA_900115865.1
99 Epibacterium ulvae GCA_900102795.1
99 Gemmobacter aquatilis GCA_900110025.1
99 Gemmobacter megaterium GCA_900156815.1
99 Gemmobacter nectariphilus DSM 15620 GCA_000429765.1
99 Haematobacter massiliensis GCA_000740795.1
99 Hoeflea olei GCA_001703635.1
99 Hoeflea phototrophica DFL-43 GCA_000154705.2
99 Jannaschia aquimarina GCA_000877395.1
99 Jannaschia donghaensis GCA_001403795.1
99 Jannaschia faecimaris GCA_900107415.1
99 Jannaschia helgolandensis GCA_900109285.1
99 Jannaschia pohangensis GCA_900113875.1
99 Jannaschia rubra GCA_001403735.1
99 Jannaschia seosinensis GCA_001408515.1
99 Ketogulonicigenium robustum GCA_002117445.1
99 Ketogulonicigenium vulgare WSH-001 GCA_000223375.1
99 Leisingera daeponensis DSM 23529 GCA_000473145.1
99 Leisingera methylohalidivorans DSM 14336 GCA_000511355.1
99 Loktanella atrilutea GCA_900128995.1
99 Loktanella cinnabarina LL-001 GCA_000466965.1
99 Loktanella fryxellensis GCA_900110065.1
99 Loktanella hongkongensis DSM 17492 GCA_000600975.2
99 Loktanella koreensis GCA_900109295.1
99 Loktanella pyoseonensis GCA_900102015.1
99 Loktanella salsilacus GCA_900114485.1
99 Loktanella soesokkakensis GCA_900172345.1
99 Lutimaribacter saemankumensis GCA_900100005.1
99 Mameliella alba GCA_900101505.1
99 Maribius pelagius GCA_900110115.1
99 Marinovum algicola DG 898 GCA_001046955.1
99 Maritimibacter alkaliphilus HTCC2654 GCA_000152805.1
99 Marivita cryptomonadis GCA_002115725.1
99 Marivita geojedonensis GCA_002115805.1
99 Marivita hallyeonensis GCA_900129875.1
99 Nautella italica GCA_001258055.1
99 Nioella nitratireducens GCA_001879715.1
99 Nioella sediminis GCA_001879695.1
99 Nitratireductor aquibiodomus NL21 = JCM 21793 GCA_000615975.1
99 Nitratireductor basaltis GCA_000733725.1
99 Nitratireductor indicus C115 GCA_000300515.1
99 Nitratireductor pacificus pht-3B GCA_000300335.1
99 Oceanicella actignis GCA_900143155.1
99 Oceanicola granulosus HTCC2516 GCA_000153305.1
99 Oceaniovalibus guishaninsula JLT2003 GCA_000299575.1
99 Octadecabacter arcticus 238 GCA_000155735.2
99 Paenirhodobacter enshiensis GCA_000740785.1
99 Paracoccus alcaliphilus GCA_900110285.1
99 Paracoccus alkenifer GCA_900108405.1
99 Paracoccus aminophilus JCM 7686 GCA_000444995.1
99 Paracoccus chinensis GCA_900102885.1
99 Paracoccus contaminans GCA_002105555.1

99 Paracoccus denitrificans PD1222 GCA_000203895.1
99 Paracoccus halophilus GCA_000763905.1
99 Paracoccus isoporae GCA_900101865.1
99 Paracoccus saliphilus GCA_900156835.1
99 Paracoccus sanguinis GCA_900106665.1
99 Paracoccus sediminis GCA_900188295.1
99 Paracoccus solventivorans GCA_900142875.1
99 Paracoccus tibetensis GCA_900102505.1
99 Paracoccus versutus GCA_000763885.1
99 Paracoccus yeei GCA_002073635.1
99 Phaeobacter gallaeciensis DSM 26640 GCA_000511385.1
99 Phaeobacter gallaeciensis GCA_001678945.1
99 Phaeobacter porticola GCA_001888185.1
99 Planktotalea frisia GCA_001890925.1
99 Poseidonocella pacifica GCA_900111875.1
99 Poseidonocella sedimentorum GCA_900116005.1
99 Pseudaminobacter salicylatoxidans KCT001 GCA_000304395.1
99 Pseudooceanicola atlanticus GCA_000768315.1
99 Pseudooceanicola batsensis HTCC2597 GCA_000152725.1
99 Pseudooceanicola nanhaiensis DSM 18065 GCA_000688295.1
99 Pseudooceanicola nitratireducens GCA_900112545.1
99 Pseudooctadecabacter jejudonensis GCA_900172275.1
99 Pseudorhodobacter aquimaris GCA_001202025.1
99 Pseudorhodobacter ferrugineus DSM 5888 GCA_000420745.1
99 Pseudorhodobacter psychrotolerans GCA_001294535.1
99 Pseudorhodobacter wandonensis GCA_001202035.1
99 Pseudoruegeria aquimaris GCA_900172235.1
99 Pseudoruegeria lutimaris GCA_900099935.1
99 Pseudoruegeria marinistellae GCA_001509585.1
99 Pseudoruegeria sabulilitoris GCA_001558155.1
99 Puniceibacterium sediminis GCA_900188035.1
99 Rhodobacter aestuarii GCA_900156655.1
99 Rhodovulum sulfidophilum DSM 1374 GCA_001633165.1
99 Rhodovulum sulfidophilum GCA_001941715.1
99 Roseibacterium elongatum DSM 19469 GCA_000590925.1
99 Roseicitreum antarcticum GCA_900107025.1
99 Roseisalinus antarcticus GCA_900172355.1
99 Roseivivax halodurans JCM 10272 GCA_000521785.1
99 Roseivivax isoporae LMG 25204 GCA_000521865.1
99 Roseivivax jejudonensis GCA_900172265.1
99 Roseivivax lentus GCA_900156805.1
99 Roseovarius halotolerans GCA_900172255.1
99 Roseovarius lutimaris GCA_900115165.1
99 Roseovarius mucosus GCA_002080415.1
99 Roseovarius pacificus GCA_900142665.1
99 Ruegeria atlantica GCA_001458195.1
99 Ruegeria halocynthiae GCA_000743705.1
99 Ruegeria marina GCA_900101475.1
99 Ruegeria meonggei GCA_900172215.1
99 Ruegeria mobilis F1926 GCA_000376545.2
99 Ruegeria pomeroyi DSS-3 GCA_000011965.2
99 Sagittula stellata E-37 GCA_000169415.1
99 Salinihabitans flavidus GCA_900110425.1
99 Shimia marina GCA_001458175.1
99 Silicibacter lacuscaerulensis ITI-1157 GCA_000161775.1
99 Sulfitobacter pseudonitzschiae GCA_002222635.1
99 Tateyamaria omphalii GCA_001969365.1
99 Thalassobius aestuarii GCA_900114635.1
99 Thalassobius gelatinovorus GCA_001458355.1
99 Thalassobius mediterraneus GCA_001458435.1
99 Thioclava atlantica GCA_000737065.1
99 Thioclava dalianensis GCA_000715505.1
99 Thioclava indica GCA_000714545.1
99 Thioclava marina GCA_002020135.1
99 Thioclava nitratireducens GCA_001940525.2
99 Tropicibacter litoreus R37 GCA_900172225.1
99 Tropicibacter multivorans GCA_001458415.1
99 Tropicibacter naphthalenivorans GCA_900176475.1
99 Tropicimonas isoalkanivorans GCA_900112335.1
99 Tropicimonas sediminicola GCA_900188335.1
99 Wenxinia marina DSM 24838 GCA_000379485.1
99 Wenxinia saemankumensis GCA_900141735.1
io
224 Deinococcus radiodurans R1 GCA_000008565.1
144 Deinococcus gobiensis I-0 GCA_000252445.1
132 Deinococcus deserti VCD115 GCA_000020685.1

ip
283 Escherichia coli IAI39 GCA_000026345.1
283 Escherichia coli O104_3AH4 str. 2011C-3493 GCA_000299455.1
283 Escherichia coli O157_3AH7 str. Sakai GCA_000008865.1
283 Escherichia coli UMN026 GCA_000026325.2
283 Escherichia coli str. K-12 substr. MG1655 GCA_000005845.2
283 Shigella flexneri 2a str. 301 GCA_000006925.2
283 Tumebacillus flagellatus GCA_000714935.1
282 Escherichia coli O83_3AH1 str. NRG 857C GCA_000183345.1
270 Erwinia iniecta GCA_001267535.1
iq
319 Bifidobacterium adolescentis ATCC 15703 GCA_000010425.1
275 Bifidobacterium dentium JCM 1195 = DSM 20436 GCA_001042595.1
273 Bifidobacterium thermophilum GCA_000741495.1
ir
155 Deinococcus radiodurans R1 GCA_000008565.1
89 Deinococcus marmoris DSM 12784 GCA_000701405.1
89 Deinococcus swuensis GCA_000800395.1
83 Deinococcus gobiensis I-0 GCA_000252445.1
is
154 Streptococcus mutans UA159 GCA_000007465.2
87 Streptococcus equinus GCA_000964315.1
87 Streptococcus gallolyticus subsp. gallolyticus DSM 16831 GCA_002000985.1
87 Streptococcus massiliensis DSM 18628 GCA_000380065.1
87 Streptococcus orisratti DSM 15617 GCA_000380105.1
87 Streptococcus ratti FA-1 = DSM 20564 GCA_000286075.1
85 Streptococcus salivarius GCA_000785515.1
85 Streptococcus thermophilus JIM 8232 GCA_000253395.1
it
179 Megasphaera cerevisiae DSM 20462 GCA_001045675.1
179 Staphylococcus aureus subsp. aureus NCTC 8325 GCA_000013425.1
179 Staphylococcus epidermidis ATCC 12228 GCA_000007645.1
179 Staphylococcus haemolyticus JCSC1435 GCA_000009865.1
179 Staphylococcus hominis subsp. hominis C80 GCA_000183685.1
179 Staphylococcus simiae CCM 7213 GCA_000235645.2
179 Staphylococcus warneri SG1 GCA_000332735.1
174 Staphylococcus capitis subsp. capitis GCA_001028645.1
174 Staphylococcus lutrae GCA_002101335.1
173 Staphylococcus cohnii subsp. cohnii GCA_000972575.1
173 Staphylococcus sciuri GCA_002209165.1
iu
118 Bacillus bataviensis LMG 21833 GCA_000307875.1
118 Lactococcus lactis subsp. lactis Il1403 GCA_000006865.1
118 Lactococcus piscium MKFS47 GCA_000981525.1
118 Streptococcus mutans UA159 GCA_000007465.2
118 Streptococcus sobrinus DSM 20742 = ATCC 33478 GCA_000686605.1
117 Bacillus azotoformans LMG 9581 GCA_000307855.1
103 Streptococcus agalactiae 2603V_2FR GCA_000007265.1
103 Streptococcus cristatus AS 1.3089 GCA_000385925.1
103 Streptococcus dysgalactiae subsp. equisimilis AC-2713 GCA_000317855.1
103 Streptococcus gordonii str. Challis substr. CH1 GCA_000017005.1
103 Streptococcus ictaluri 707-05 GCA_000188015.3
103 Streptococcus mitis B6 GCA_000027165.1
103 Streptococcus pneumoniae R6 GCA_000007045.1
iv
192 Bacillus thuringiensis YBT-1518 GCA_000497525.2
179 Bacillus coahuilensis m4-4 GCA_000171615.1
170 Bacillus anthracis str. Ames GCA_000007845.1
170 Bacillus anthracis str. Sterne GCA_000008165.1
170 Bacillus cereus ATCC 14579 GCA_000007825.1
170 Bacillus mycoides GCA_000832605.1
170 Bacillus pseudomycoides DSM 12442 GCA_000161455.1
170 _5BBacillus thuringiensis_5D serovar konkukian str. 97-27 GCA_000008505.1
iw
242 Clostridium beijerinckii GCA_000833105.2
242 Clostridium saccharoperbutylacetonicum N1-4_28HMT_29 GCA_000340885.1
238 Clostridium puniceum GCA_002006345.1
227 Clostridium saccharobutylicum DSM 13864 GCA_000473995.1
ix
239 Staphylococcus capitis subsp. capitis GCA_001028645.1
239 Staphylococcus epidermidis ATCC 12228 GCA_000007645.1
239 Staphylococcus haemolyticus JCSC1435 GCA_000009865.1
239 Staphylococcus hominis subsp. hominis C80 GCA_000183685.1
239 Staphylococcus simulans GCA_001559115.1
233 Staphylococcus microti GCA_000934465.1
224 Staphylococcus lugdunensis HKU09-01 GCA_000025085.1

iy
113 Alloscardovia macacae GCA_002127695.1
113 Bifidobacterium callitrichos DSM 23973 GCA_000741175.1
113 Bifidobacterium coryneforme GCA_000737865.1
113 Bifidobacterium subtile GCA_000741775.1
112 Bifidobacterium adolescentis ATCC 15703 GCA_000010425.1
112 Bifidobacterium angulatum DSM 20098 = JCM 7096 GCA_001025155.1
111 Alloscardovia criceti DSM 17774 GCA_000376885.1
iz
108 Bifidobacterium animalis subsp. lactis DSM 10140 GCA_000022965.1
106 Bifidobacterium adolescentis ATCC 15703 GCA_000010425.1
106 Bifidobacterium dentium JCM 1195 = DSM 20436 GCA_001042595.1
106 Bifidobacterium tsurumiense GCA_000741765.1
90 Bifidobacterium asteroides PRL2011 GCA_000304215.1
iA
313 Bacillus anthracis str. Ames GCA_000007845.1
313 Bacillus anthracis str. Sterne GCA_000008165.1
313 Bacillus mycoides GCA_000832605.1
313 Bacillus pseudomycoides DSM 12442 GCA_000161455.1
313 Bacillus thuringiensis YBT-1518 GCA_000497525.2
313 _5BBacillus thuringiensis_5D serovar konkukian str. 97-27 GCA_000008505.1
312 Bacillus cereus ATCC 14579 GCA_000007825.1
268 Bacillus manliponensis GCA_000712595.1
iB
236 Escherichia coli IAI39 GCA_000026345.1
236 Escherichia coli O104_3AH4 str. 2011C-3493 GCA_000299455.1
236 Escherichia coli O157_3AH7 str. Sakai GCA_000008865.1
236 Escherichia coli O83_3AH1 str. NRG 857C GCA_000183345.1
236 Escherichia coli UMN026 GCA_000026325.2
236 Escherichia coli str. K-12 substr. MG1655 GCA_000005845.2
236 Shigella flexneri 2a str. 301 GCA_000006925.2
236 Tumebacillus flagellatus GCA_000714935.1
228 Erwinia iniecta GCA_001267535.1
227 Shigella dysenteriae Sd197 GCA_000012005.1
iC
219 Bacillus anthracis str. Ames GCA_000007845.1
219 Bacillus anthracis str. Sterne GCA_000008165.1
219 Bacillus cereus ATCC 14579 GCA_000007825.1
219 Bacillus mycoides GCA_000832605.1
219 Bacillus pseudomycoides DSM 12442 GCA_000161455.1
219 Bacillus thuringiensis YBT-1518 GCA_000497525.2
219 _5BBacillus thuringiensis_5D serovar konkukian str. 97-27 GCA_000008505.1
184 Isobaculum melis GCA_900111355.1
174 Bacillus horneckiae GCA_001636335.1
174 Bacillus solani GCA_001420595.1
iD
267 Streptococcus mutans UA159 GCA_000007465.2
188 Streptococcus ferus DSM 20646 GCA_000372425.1
185 Streptococcus ratti FA-1 = DSM 20564 GCA_000286075.1
iE
232 Escherichia coli IAI39 GCA_000026345.1
232 Escherichia coli O104_3AH4 str. 2011C-3493 GCA_000299455.1
232 Escherichia coli O157_3AH7 str. Sakai GCA_000008865.1
232 Escherichia coli O83_3AH1 str. NRG 857C GCA_000183345.1
232 Escherichia coli UMN026 GCA_000026325.2
232 Escherichia coli str. K-12 substr. MG1655 GCA_000005845.2
232 Shigella dysenteriae Sd197 GCA_000012005.1
232 Shigella flexneri 2a str. 301 GCA_000006925.2
232 Tumebacillus flagellatus GCA_000714935.1
226 Erwinia iniecta GCA_001267535.1
220 Erwinia toletana DAPP-PG 735 GCA_000336255.1
iF
249 Clostridium beijerinckii GCA_000833105.2
249 Clostridium chromiireducens GCA_002029255.1
249 Clostridium puniceum GCA_002006345.1
249 Clostridium saccharobutylicum DSM 13864 GCA_000473995.1
249 Clostridium saccharoperbutylacetonicum N1-4_28HMT_29 GCA_000340885.1
222 Clostridium botulinum B str. Eklund 17B _28NRP_29 GCA_000020165.1
222 Clostridium taeniosporum GCA_001735765.1
217 Clostridium fallax GCA_900129365.1
iG
196 Staphylococcus condimenti GCA_001618885.1
196 Staphylococcus simulans GCA_001559115.1
190 Staphylococcus aureus subsp. aureus NCTC 8325 GCA_000013425.1
190 Staphylococcus simiae CCM 7213 GCA_000235645.2
181 Staphylococcus epidermidis ATCC 12228 GCA_000007645.1

181 Staphylococcus haemolyticus JCSC1435 GCA_000009865.1
181 Staphylococcus hominis subsp. hominis C80 GCA_000183685.1
181 Staphylococcus lugdunensis HKU09-01 GCA_000025085.1
iH
287 Deinococcus radiodurans R1 GCA_000008565.1
225 Deinococcus deserti VCD115 GCA_000020685.1
219 Deinococcus gobiensis I-0 GCA_000252445.1
iI
280 Bacillus anthracis str. Ames GCA_000007845.1
280 Bacillus anthracis str. Sterne GCA_000008165.1
280 Bacillus cereus ATCC 14579 GCA_000007825.1
280 Bacillus mycoides GCA_000832605.1
280 Bacillus pseudomycoides DSM 12442 GCA_000161455.1
280 Bacillus thuringiensis YBT-1518 GCA_000497525.2
280 _5BBacillus thuringiensis_5D serovar konkukian str. 97-27 GCA_000008505.1
245 Bacillus marisflavi GCA_001274775.1
233 Bacillus aquimaris TF-12 GCA_001648555.1
233 Bacillus coahuilensis m4-4 GCA_000171615.1
iJ
289 Escherichia coli IAI39 GCA_000026345.1
289 Escherichia coli O104_3AH4 str. 2011C-3493 GCA_000299455.1
289 Escherichia coli O157_3AH7 str. Sakai GCA_000008865.1
289 Escherichia coli O83_3AH1 str. NRG 857C GCA_000183345.1
289 Escherichia coli UMN026 GCA_000026325.2
289 Escherichia coli str. K-12 substr. MG1655 GCA_000005845.2
289 Shigella dysenteriae Sd197 GCA_000012005.1
289 Shigella flexneri 2a str. 301 GCA_000006925.2
289 Tumebacillus flagellatus GCA_000714935.1
252 Erwinia iniecta GCA_001267535.1
232 Cronobacter sakazakii GCA_000982825.1
232 Erwinia toletana DAPP-PG 735 GCA_000336255.1
iK
203 Cronobacter sakazakii GCA_000982825.1
203 Edwardsiella anguillarum ET080813 GCA_000264765.2
203 Enterobacter cloacae subsp. cloacae ATCC 13047 GCA_000025565.1
203 Enterobacter hormaechei subsp. steigerwaltii GCA_001729725.1
203 Enterobacter kobei GCA_900185885.1
203 Erwinia amylovora CFBP1430 GCA_000091565.1
203 Erwinia billingiae Eb661 GCA_000196615.1
203 Erwinia gerundensis GCA_001517405.1
203 Erwinia iniecta GCA_001267535.1
203 Erwinia persicina NBRC 102418 GCA_001571305.1
203 Erwinia toletana DAPP-PG 735 GCA_000336255.1
203 Escherichia coli IAI39 GCA_000026345.1
203 Escherichia coli O104_3AH4 str. 2011C-3493 GCA_000299455.1
203 Escherichia coli O157_3AH7 str. Sakai GCA_000008865.1
203 Escherichia coli O83_3AH1 str. NRG 857C GCA_000183345.1
203 Escherichia coli UMN026 GCA_000026325.2
203 Escherichia coli str. K-12 substr. MG1655 GCA_000005845.2
203 Izhakiella capsodis GCA_900115045.1
203 Klebsiella oxytoca GCA_001022195.1
203 Kosakonia cowanii GCA_001975225.1
203 Kosakonia sacchari SP1 GCA_000300455.4
203 Mangrovibacter phragmitis GCA_001655675.1
203 Pantoea agglomerans GCA_001709315.1
203 Pantoea alhagi GCA_002101395.1
203 Pantoea ananatis LMG 20103 GCA_000025405.2
203 Pantoea dispersa EGD-AAK13 GCA_000465555.2
203 Pantoea rwandensis GCA_000759475.1
203 Pantoea septica GCA_002095575.1
203 Pluralibacter gergoviae GCA_000757785.1
203 Pseudescherichia vulneris NBRC 102420 GCA_000759795.1
203 Rosenbergiella nectarea GCA_900111105.1
203 Salmonella enterica subsp. enterica serovar Typhi str. CT18 GCA_000195995.1
203 Shigella dysenteriae Sd197 GCA_000012005.1
203 Shigella flexneri 2a str. 301 GCA_000006925.2
203 Shimwellia blattae DSM 4481 = NBRC 105725 GCA_000262305.1
203 Tatumella citrea GCA_002163585.1
203 Trabulsiella odontotermitis GCA_001297765.1
203 Tumebacillus flagellatus GCA_000714935.1
202 Buttiauxella ferragutiae ATCC 51602 GCA_001654915.1
202 Cedecea neteri GCA_000757825.1
202 Citrobacter freundii CFNIH1 GCA_000648515.1
202 Klebsiella aerogenes KCTC 2190 GCA_000215745.1
202 Kluyvera ascorbata ATCC 33433 GCA_000735365.1
202 Nissabacter archeti GCA_900130115.1

202 Obesumbacterium proteus GCA_001586165.1
202 Tatumella saanichensis GCA_000439375.1
202 Yokenella regensburgei ATCC 49455 GCA_000735455.1
202 _5BEnterobacter_5D lignolyticus SCF1 GCA_000164865.1
195 Brenneria goodwinii GCA_001049335.1
195 Lonsdalea quercina subsp. quercina GCA_000688655.1
iL
193 Clostridium beijerinckii GCA_000833105.2
193 Clostridium puniceum GCA_002006345.1
193 Clostridium saccharoperbutylacetonicum N1-4_28HMT_29 GCA_000340885.1
184 Clostridium saccharobutylicum DSM 13864 GCA_000473995.1
172 Clostridium neonatale GCA_001458595.1
iM
316 Streptococcus mutans UA159 GCA_000007465.2
195 Streptococcus marimammalium DSM 18627 GCA_000380045.1
182 Streptococcus ferus DSM 20646 GCA_000372425.1
182 Streptococcus gordonii str. Challis substr. CH1 GCA_000017005.1
iN
215 Clostridium beijerinckii GCA_000833105.2
215 Clostridium chromiireducens GCA_002029255.1
215 Clostridium fallax GCA_900129365.1
215 Clostridium puniceum GCA_002006345.1
215 Clostridium saccharobutylicum DSM 13864 GCA_000473995.1
215 Clostridium saccharoperbutylacetonicum N1-4_28HMT_29 GCA_000340885.1
211 Clostridium butyricum GCA_001456065.2
201 Clostridium gasigenes GCA_900104115.1
iO
196 Enterobacter hormaechei subsp. steigerwaltii GCA_001729725.1
196 Erwinia billingiae Eb661 GCA_000196615.1
196 Erwinia iniecta GCA_001267535.1
196 Erwinia toletana DAPP-PG 735 GCA_000336255.1
196 Escherichia coli IAI39 GCA_000026345.1
196 Escherichia coli O104_3AH4 str. 2011C-3493 GCA_000299455.1
196 Escherichia coli O157_3AH7 str. Sakai GCA_000008865.1
196 Escherichia coli O83_3AH1 str. NRG 857C GCA_000183345.1
196 Escherichia coli UMN026 GCA_000026325.2
196 Escherichia coli str. K-12 substr. MG1655 GCA_000005845.2
196 Klebsiella oxytoca GCA_001022195.1
196 Klebsiella pneumoniae subsp. pneumoniae HS11286 GCA_000240185.2
196 Kosakonia cowanii GCA_001975225.1
196 Kosakonia sacchari SP1 GCA_000300455.4
196 Pantoea agglomerans GCA_001709315.1
196 Pantoea alhagi GCA_002101395.1
196 Pantoea ananatis LMG 20103 GCA_000025405.2
196 Pantoea rwandensis GCA_000759475.1
196 Pantoea septica GCA_002095575.1
196 Pseudescherichia vulneris NBRC 102420 GCA_000759795.1
196 Shigella dysenteriae Sd197 GCA_000012005.1
196 Shigella flexneri 2a str. 301 GCA_000006925.2
196 Shimwellia blattae DSM 4481 = NBRC 105725 GCA_000262305.1
196 Tumebacillus flagellatus GCA_000714935.1
194 Xenorhabdus cabanillasii JM26 GCA_000531755.1
194 Xenorhabdus doucetiae GCA_000968195.1
194 Xenorhabdus hominickii GCA_001721185.1
194 Xenorhabdus innexi GCA_900155355.1
194 Xenorhabdus japonica GCA_900115195.1
194 Xenorhabdus koppenhoeferi GCA_900116635.1
194 Xenorhabdus nematophila AN6_2F1 GCA_000953355.1
190 Pluralibacter gergoviae GCA_000757785.1
iP
218 Bacillus thuringiensis YBT-1518 GCA_000497525.2
203 Bacillus anthracis str. Ames GCA_000007845.1
203 Bacillus anthracis str. Sterne GCA_000008165.1
203 Bacillus cereus ATCC 14579 GCA_000007825.1
203 Bacillus mycoides GCA_000832605.1
203 Bacillus pseudomycoides DSM 12442 GCA_000161455.1
203 _5BBacillus thuringiensis_5D serovar konkukian str. 97-27 GCA_000008505.1
170 Massilibacterium senegalense GCA_001375675.1
iQ
297 Enterococcus faecalis V583 GCA_000007785.1
297 Streptomyces cinnamoneus GCA_001885705.1
264 Enterococcus asini ATCC 700915 GCA_000407365.1
264 Enterococcus canis NBRC 100695 GCA_001544375.1
264 Enterococcus dispar ATCC 51266 GCA_000406945.1
264 Enterococcus faecium DO GCA_000174395.2
264 Enterococcus gilvus ATCC BAA-350 GCA_000407545.1

264 Enterococcus haemoperoxidus ATCC BAA-382 GCA_000407165.1
264 Enterococcus hermanniensis GCA_001885945.1
264 Enterococcus hirae ATCC 9790 GCA_000271405.2
264 Enterococcus italicus DSM 15952 GCA_000185365.1
264 Enterococcus malodoratus ATCC 43197 GCA_000407185.1
264 Enterococcus mundtii QU 25 GCA_000504125.1
264 Enterococcus pallens ATCC BAA-351 GCA_000407485.1
264 Enterococcus phoeniculicola ATCC BAA-412 GCA_000407505.1
264 Enterococcus pseudoavium NBRC 100491 GCA_001544295.1
264 Enterococcus rivorum GCA_001742285.1
264 Enterococcus sulfureus ATCC 49903 GCA_000407605.1
264 Enterococcus thailandicus GCA_001652875.1
264 Vagococcus lutrae LBD1 GCA_000498295.1
260 Melissococcus plutonius S1 GCA_000747585.1
iR
174 Escherichia coli IAI39 GCA_000026345.1
174 Escherichia coli O104_3AH4 str. 2011C-3493 GCA_000299455.1
174 Escherichia coli O157_3AH7 str. Sakai GCA_000008865.1
174 Escherichia coli O83_3AH1 str. NRG 857C GCA_000183345.1
174 Escherichia coli UMN026 GCA_000026325.2
174 Escherichia coli str. K-12 substr. MG1655 GCA_000005845.2
174 Shigella dysenteriae Sd197 GCA_000012005.1
174 Shigella flexneri 2a str. 301 GCA_000006925.2
174 Tumebacillus flagellatus GCA_000714935.1
167 Kosakonia cowanii GCA_001975225.1
166 Enterobacter hormaechei subsp. steigerwaltii GCA_001729725.1
166 Erwinia billingiae Eb661 GCA_000196615.1
166 Erwinia iniecta GCA_001267535.1
166 Erwinia toletana DAPP-PG 735 GCA_000336255.1
166 Klebsiella oxytoca GCA_001022195.1
166 Klebsiella pneumoniae subsp. pneumoniae HS11286 GCA_000240185.2
166 Kosakonia sacchari SP1 GCA_000300455.4
166 Pantoea agglomerans GCA_001709315.1
166 Pantoea alhagi GCA_002101395.1
166 Pantoea ananatis LMG 20103 GCA_000025405.2
166 Pantoea rwandensis GCA_000759475.1
166 Pantoea septica GCA_002095575.1
166 Pluralibacter gergoviae GCA_000757785.1
166 Pseudescherichia vulneris NBRC 102420 GCA_000759795.1
166 Shimwellia blattae DSM 4481 = NBRC 105725 GCA_000262305.1
iS
275 Deinococcus radiodurans R1 GCA_000008565.1
161 Deinococcus gobiensis I-0 GCA_000252445.1
142 Deinococcus deserti VCD115 GCA_000020685.1
iT
249 Lactobacillus gasseri ATCC 33323 = JCM 1131 GCA_000014425.1
234 Lactobacillus hominis DSM 23910 = CRBIP 24.179 GCA_000296835.1
210 Lactobacillus acidophilus NCFM GCA_000011985.1
iU
225 Deinococcus radiodurans R1 GCA_000008565.1
166 Deinococcus gobiensis I-0 GCA_000252445.1
159 Deinococcus puniceus GCA_001644565.1
iV
220 Clostridium beijerinckii GCA_000833105.2
220 Clostridium saccharoperbutylacetonicum N1-4_28HMT_29 GCA_000340885.1
213 Clostridium puniceum GCA_002006345.1
206 Clostridium saccharobutylicum DSM 13864 GCA_000473995.1
iW
238 Escherichia coli IAI39 GCA_000026345.1
238 Escherichia coli O104_3AH4 str. 2011C-3493 GCA_000299455.1
238 Escherichia coli O157_3AH7 str. Sakai GCA_000008865.1
238 Escherichia coli O83_3AH1 str. NRG 857C GCA_000183345.1
238 Escherichia coli UMN026 GCA_000026325.2
238 Escherichia coli str. K-12 substr. MG1655 GCA_000005845.2
238 Shigella flexneri 2a str. 301 GCA_000006925.2
238 Tumebacillus flagellatus GCA_000714935.1
232 Erwinia iniecta GCA_001267535.1
229 Shigella dysenteriae Sd197 GCA_000012005.1
iX
200 Streptococcus mutans UA159 GCA_000007465.2
124 Streptococcus sobrinus DSM 20742 = ATCC 33478 GCA_000686605.1
119 Streptococcus ferus DSM 20646 GCA_000372425.1
iY
205 Bacillus thuringiensis YBT-1518 GCA_000497525.2
201 Bacillus anthracis str. Ames GCA_000007845.1
201 Bacillus anthracis str. Sterne GCA_000008165.1

201 Bacillus cereus ATCC 14579 GCA_000007825.1
201 Bacillus mycoides GCA_000832605.1
201 Bacillus pseudomycoides DSM 12442 GCA_000161455.1
201 _5BBacillus thuringiensis_5D serovar konkukian str. 97-27 GCA_000008505.1
196 Bacillus manliponensis GCA_000712595.1
iZ
236 Clostridium beijerinckii GCA_000833105.2
236 Clostridium puniceum GCA_002006345.1
236 Clostridium saccharobutylicum DSM 13864 GCA_000473995.1
236 Clostridium saccharoperbutylacetonicum N1-4_28HMT_29 GCA_000340885.1
232 Clostridium chromiireducens GCA_002029255.1
223 Clostridium taeniosporum GCA_001735765.1
j0
173 Clostridium beijerinckii GCA_000833105.2
173 Clostridium saccharobutylicum DSM 13864 GCA_000473995.1
173 Clostridium saccharoperbutylacetonicum N1-4_28HMT_29 GCA_000340885.1
159 Clostridium puniceum GCA_002006345.1
158 Hathewaya proteolytica DSM 3090 GCA_900142225.1
j1
124 Clostridium beijerinckii GCA_000833105.2
124 Clostridium puniceum GCA_002006345.1
124 Clostridium saccharobutylicum DSM 13864 GCA_000473995.1
124 Clostridium saccharoperbutylacetonicum N1-4_28HMT_29 GCA_000340885.1
120 Clostridium intestinale URNW GCA_000469625.2
109 Clostridium chromiireducens GCA_002029255.1
j2
180 Bacillus thuringiensis YBT-1518 GCA_000497525.2
163 Bacillus anthracis str. Ames GCA_000007845.1
163 Bacillus anthracis str. Sterne GCA_000008165.1
163 Bacillus cereus ATCC 14579 GCA_000007825.1
163 Bacillus mycoides GCA_000832605.1
163 Bacillus pseudomycoides DSM 12442 GCA_000161455.1
163 _5BBacillus thuringiensis_5D serovar konkukian str. 97-27 GCA_000008505.1
117 Massilibacterium senegalense GCA_001375675.1
j3
141 Bifidobacterium adolescentis ATCC 15703 GCA_000010425.1
141 Bifidobacterium dentium JCM 1195 = DSM 20436 GCA_001042595.1
133 Bifidobacterium angulatum DSM 20098 = JCM 7096 GCA_001025155.1
126 Bifidobacterium thermophilum GCA_000741495.1
j4
239 Staphylococcus aureus subsp. aureus NCTC 8325 GCA_000013425.1
239 Staphylococcus simiae CCM 7213 GCA_000235645.2
237 Staphylococcus capitis subsp. capitis GCA_001028645.1
237 Staphylococcus epidermidis ATCC 12228 GCA_000007645.1
237 Staphylococcus lugdunensis HKU09-01 GCA_000025085.1
234 Staphylococcus xylosus GCA_000706685.1
j5
235 Deinococcus radiodurans R1 GCA_000008565.1
188 Deinococcus marmoris DSM 12784 GCA_000701405.1
188 Deinococcus swuensis GCA_000800395.1
180 Deinococcus frigens DSM 12807 GCA_000701425.1
j6
196 Clostridium beijerinckii GCA_000833105.2
196 Clostridium puniceum GCA_002006345.1
196 Clostridium saccharobutylicum DSM 13864 GCA_000473995.1
196 Clostridium saccharoperbutylacetonicum N1-4_28HMT_29 GCA_000340885.1
188 Clostridium botulinum B str. Eklund 17B _28NRP_29 GCA_000020165.1
188 Clostridium butyricum GCA_001456065.2
188 Clostridium chromiireducens GCA_002029255.1
188 Clostridium taeniosporum GCA_001735765.1
186 Clostridium uliginosum GCA_900112485.1
j7
255 Bifidobacterium adolescentis ATCC 15703 GCA_000010425.1
229 Bifidobacterium asteroides PRL2011 GCA_000304215.1
204 Bifidobacterium callitrichos DSM 23973 GCA_000741175.1
204 Bifidobacterium dentium JCM 1195 = DSM 20436 GCA_001042595.1
j8
167 Escherichia coli IAI39 GCA_000026345.1
167 Escherichia coli O104_3AH4 str. 2011C-3493 GCA_000299455.1
167 Escherichia coli O157_3AH7 str. Sakai GCA_000008865.1
167 Escherichia coli O83_3AH1 str. NRG 857C GCA_000183345.1
167 Escherichia coli UMN026 GCA_000026325.2
167 Escherichia coli str. K-12 substr. MG1655 GCA_000005845.2
167 Shigella flexneri 2a str. 301 GCA_000006925.2
167 Tumebacillus flagellatus GCA_000714935.1
158 Shigella dysenteriae Sd197 GCA_000012005.1

153 Erwinia iniecta GCA_001267535.1
j9
113 Sagittula stellata E-37 GCA_000169415.1
109 Haematobacter massiliensis GCA_000740795.1
107 Gemmobacter aquatilis GCA_900110025.1
107 Gemmobacter megaterium GCA_900156815.1
107 Gemmobacter nectariphilus DSM 15620 GCA_000429765.1
107 Paracoccus yeei GCA_002073635.1
107 Rhodobacter capsulatus SB 1003 GCA_000021865.1
ja
170 Staphylococcus epidermidis ATCC 12228 GCA_000007645.1
159 Staphylococcus capitis subsp. capitis GCA_001028645.1
156 Staphylococcus lugdunensis HKU09-01 GCA_000025085.1
jb
204 Rhodobacter sphaeroides 2.4.1 GCA_000012905.2
177 Rhodobacter sphaeroides ATCC 17025 GCA_000016405.1
174 Pseudorhodobacter ferrugineus DSM 5888 GCA_000420745.1
174 Pseudorhodobacter wandonensis GCA_001202035.1
jc
313 _5BBacillus thuringiensis_5D serovar konkukian str. 97-27 GCA_000008505.1
294 Bacillus anthracis str. Ames GCA_000007845.1
294 Bacillus anthracis str. Sterne GCA_000008165.1
294 Bacillus cereus ATCC 14579 GCA_000007825.1
294 Bacillus mycoides GCA_000832605.1
294 Bacillus pseudomycoides DSM 12442 GCA_000161455.1
294 Bacillus thuringiensis YBT-1518 GCA_000497525.2
254 Bacillus manliponensis GCA_000712595.1
jd
100 Deinococcus radiodurans R1 GCA_000008565.1
92 Staphylococcus epidermidis ATCC 12228 GCA_000007645.1
92 Staphylococcus haemolyticus JCSC1435 GCA_000009865.1
83 Deinococcus gobiensis I-0 GCA_000252445.1
je
294 Deinococcus radiodurans R1 GCA_000008565.1
204 Deinococcus gobiensis I-0 GCA_000252445.1
197 Deinococcus puniceus GCA_001644565.1
jf
176 Deinococcus radiodurans R1 GCA_000008565.1
115 Deinococcus marmoris DSM 12784 GCA_000701405.1
115 Deinococcus swuensis GCA_000800395.1
109 Deinococcus proteolyticus MRP GCA_000190555.1
jg
293 Staphylococcus epidermidis ATCC 12228 GCA_000007645.1
293 Staphylococcus haemolyticus JCSC1435 GCA_000009865.1
282 Staphylococcus hominis subsp. hominis C80 GCA_000183685.1
279 Staphylococcus capitis subsp. capitis GCA_001028645.1
jh
225 Clostridium beijerinckii GCA_000833105.2
225 Clostridium saccharoperbutylacetonicum N1-4_28HMT_29 GCA_000340885.1
218 Clostridium puniceum GCA_002006345.1
216 Clostridium saccharobutylicum DSM 13864 GCA_000473995.1
ji
105 Anaerobacillus macyae GCA_001039475.1
105 Anoxybacillus amylolyticus GCA_001634285.1
105 Anoxybacillus gonensis GCA_001187595.1
105 Bacillus coagulans DSM 1 = ATCC 7050 GCA_000832905.1
105 Bacillus hemicellulosilyticus JCM 9152 GCA_000513115.1
105 Enterococcus asini ATCC 700915 GCA_000407365.1
105 Enterococcus canis NBRC 100695 GCA_001544375.1
105 Enterococcus dispar ATCC 51266 GCA_000406945.1
105 Enterococcus faecalis V583 GCA_000007785.1
105 Enterococcus faecium DO GCA_000174395.2
105 Enterococcus gilvus ATCC BAA-350 GCA_000407545.1
105 Enterococcus hermanniensis GCA_001885945.1
105 Enterococcus hirae ATCC 9790 GCA_000271405.2
105 Enterococcus italicus DSM 15952 GCA_000185365.1
105 Enterococcus malodoratus ATCC 43197 GCA_000407185.1
105 Enterococcus massiliensis GCA_001050095.1
105 Enterococcus mundtii QU 25 GCA_000504125.1
105 Enterococcus pallens ATCC BAA-351 GCA_000407485.1
105 Enterococcus pseudoavium NBRC 100491 GCA_001544295.1
105 Enterococcus rivorum GCA_001742285.1
105 Listeria aquatica FSL S10-1188 GCA_000525795.1
105 Listeria fleischmannii subsp. coloradonensis GCA_000252625.2
105 Listeria floridensis FSL S10-1187 GCA_000525875.1
105 Melissococcus plutonius S1 GCA_000747585.1

105 Sporolactobacillus laevolacticus DSM 442 GCA_000497245.1
105 Sporolactobacillus nakayamae GCA_900113325.1
105 Streptomyces cinnamoneus GCA_001885705.1
103 Bacillus mycoides GCA_000832605.1
102 Bacillus anthracis str. Ames GCA_000007845.1
102 Bacillus anthracis str. Sterne GCA_000008165.1
102 Bacillus cereus ATCC 14579 GCA_000007825.1
102 Bacillus pseudomycoides DSM 12442 GCA_000161455.1
102 Bacillus thuringiensis YBT-1518 GCA_000497525.2
102 _5BBacillus thuringiensis_5D serovar konkukian str. 97-27 GCA_000008505.1
jj
286 Lactobacillus gasseri ATCC 33323 = JCM 1131 GCA_000014425.1
268 Lactobacillus hominis DSM 23910 = CRBIP 24.179 GCA_000296835.1
195 Lactobacillus acidophilus NCFM GCA_000011985.1
195 Lactobacillus gallinarum GCA_001314245.2
jk
202 Escherichia coli IAI39 GCA_000026345.1
202 Escherichia coli O104_3AH4 str. 2011C-3493 GCA_000299455.1
202 Escherichia coli O157_3AH7 str. Sakai GCA_000008865.1
202 Escherichia coli UMN026 GCA_000026325.2
202 Escherichia coli str. K-12 substr. MG1655 GCA_000005845.2
202 Shigella dysenteriae Sd197 GCA_000012005.1
202 Shigella flexneri 2a str. 301 GCA_000006925.2
202 Tumebacillus flagellatus GCA_000714935.1
201 Escherichia coli O83_3AH1 str. NRG 857C GCA_000183345.1
181 Erwinia iniecta GCA_001267535.1
jl
119 Enterococcus faecalis V583 GCA_000007785.1
119 Streptomyces cinnamoneus GCA_001885705.1
117 Enterococcus hirae ATCC 9790 GCA_000271405.2
113 Hymenobacter terrenus GCA_000972495.1
jm
238 Bifidobacterium adolescentis ATCC 15703 GCA_000010425.1
223 Bifidobacterium dentium JCM 1195 = DSM 20436 GCA_001042595.1
200 Bifidobacterium tsurumiense GCA_000741765.1
jn
223 Streptococcus mutans UA159 GCA_000007465.2
165 Streptococcus equinus GCA_000964315.1
165 Streptococcus gallolyticus subsp. gallolyticus DSM 16831 GCA_002000985.1
161 Streptococcus ratti FA-1 = DSM 20564 GCA_000286075.1
jo
146 Clostridium beijerinckii GCA_000833105.2
146 Clostridium puniceum GCA_002006345.1
146 Clostridium saccharobutylicum DSM 13864 GCA_000473995.1
146 Clostridium saccharoperbutylacetonicum N1-4_28HMT_29 GCA_000340885.1
138 Clostridium chromiireducens GCA_002029255.1
125 Clostridium butyricum GCA_001456065.2
jp
240 Deinococcus radiodurans R1 GCA_000008565.1
153 Deinococcus gobiensis I-0 GCA_000252445.1
121 Deinococcus deserti VCD115 GCA_000020685.1
121 Deinococcus soli Cha et al. 2016 GCA_001007995.1
jq
304 Lactobacillus gasseri ATCC 33323 = JCM 1131 GCA_000014425.1
295 Lactobacillus hominis DSM 23910 = CRBIP 24.179 GCA_000296835.1
226 Lactobacillus psittaci DSM 15354 GCA_000425905.1
jr
264 Lactobacillus gasseri ATCC 33323 = JCM 1131 GCA_000014425.1
251 Lactobacillus hominis DSM 23910 = CRBIP 24.179 GCA_000296835.1
218 Lactobacillus iners DSM 13335 GCA_000160875.1
js
168 Bacillus thuringiensis YBT-1518 GCA_000497525.2
160 Bacillus anthracis str. Ames GCA_000007845.1
160 Bacillus anthracis str. Sterne GCA_000008165.1
160 Bacillus cereus ATCC 14579 GCA_000007825.1
160 Bacillus mycoides GCA_000832605.1
160 Bacillus pseudomycoides DSM 12442 GCA_000161455.1
160 _5BBacillus thuringiensis_5D serovar konkukian str. 97-27 GCA_000008505.1
118 Bacillus hemicellulosilyticus JCM 9152 GCA_000513115.1
118 Bacillus licheniformis DSM 13 = ATCC 14580 GCA_000011645.1
118 Oceanobacillus iheyensis HTE831 GCA_000011245.1
jt
167 Streptococcus mutans UA159 GCA_000007465.2
119 Streptococcus sobrinus DSM 20742 = ATCC 33478 GCA_000686605.1
118 Streptococcus macacae NCTC 11558 GCA_000187995.3
ju

239 Streptococcus mutans UA159 GCA_000007465.2
222 Enterococcus faecalis V583 GCA_000007785.1
222 Streptomyces cinnamoneus GCA_001885705.1
189 Enterococcus asini ATCC 700915 GCA_000407365.1
189 Enterococcus canis NBRC 100695 GCA_001544375.1
189 Enterococcus casseliflavus EC20 GCA_000157355.2
189 Enterococcus cecorum GCA_001318405.1
189 Enterococcus columbae DSM 7374 = ATCC 51263 GCA_000406925.1
189 Enterococcus dispar ATCC 51266 GCA_000406945.1
189 Enterococcus faecium DO GCA_000174395.2
189 Enterococcus gilvus ATCC BAA-350 GCA_000407545.1
189 Enterococcus haemoperoxidus ATCC BAA-382 GCA_000407165.1
189 Enterococcus hermanniensis GCA_001885945.1
189 Enterococcus hirae ATCC 9790 GCA_000271405.2
189 Enterococcus italicus DSM 15952 GCA_000185365.1
189 Enterococcus malodoratus ATCC 43197 GCA_000407185.1
189 Enterococcus massiliensis GCA_001050095.1
189 Enterococcus mundtii QU 25 GCA_000504125.1
189 Enterococcus pallens ATCC BAA-351 GCA_000407485.1
189 Enterococcus phoeniculicola ATCC BAA-412 GCA_000407505.1
189 Enterococcus pseudoavium NBRC 100491 GCA_001544295.1
189 Enterococcus rivorum GCA_001742285.1
189 Enterococcus saccharolyticus subsp. saccharolyticus ATCC 43076 GCA_000407285.1
189 Enterococcus sulfureus ATCC 49903 GCA_000407605.1
189 Enterococcus thailandicus GCA_001652875.1
189 Vagococcus fluvialis bH819 GCA_900163795.1
189 Vagococcus lutrae LBD1 GCA_000498295.1
189 Vagococcus penaei GCA_001998885.1
jv
215 Clostridium butyricum GCA_001456065.2
204 Clostridium beijerinckii GCA_000833105.2
204 Clostridium puniceum GCA_002006345.1
204 Clostridium saccharobutylicum DSM 13864 GCA_000473995.1
204 Clostridium saccharoperbutylacetonicum N1-4_28HMT_29 GCA_000340885.1
200 Clostridium chromiireducens GCA_002029255.1
jw
288 Staphylococcus capitis subsp. capitis GCA_001028645.1
288 Staphylococcus epidermidis ATCC 12228 GCA_000007645.1
285 Staphylococcus haemolyticus JCSC1435 GCA_000009865.1
284 Staphylococcus cohnii subsp. cohnii GCA_000972575.1
jx
256 Rhodobacter sphaeroides 2.4.1 GCA_000012905.2
226 Rhodobacter sphaeroides ATCC 17025 GCA_000016405.1
194 Defluviimonas alba GCA_001620265.1
jy
174 Escherichia coli IAI39 GCA_000026345.1
174 Escherichia coli O104_3AH4 str. 2011C-3493 GCA_000299455.1
174 Escherichia coli O157_3AH7 str. Sakai GCA_000008865.1
174 Escherichia coli UMN026 GCA_000026325.2
174 Escherichia coli str. K-12 substr. MG1655 GCA_000005845.2
174 Shigella dysenteriae Sd197 GCA_000012005.1
174 Shigella flexneri 2a str. 301 GCA_000006925.2
174 Tumebacillus flagellatus GCA_000714935.1
173 Escherichia coli O83_3AH1 str. NRG 857C GCA_000183345.1
159 Cronobacter sakazakii GCA_000982825.1
159 Erwinia gerundensis GCA_001517405.1
159 Erwinia iniecta GCA_001267535.1
159 Erwinia toletana DAPP-PG 735 GCA_000336255.1
159 Pantoea agglomerans GCA_001709315.1
159 Pantoea ananatis LMG 20103 GCA_000025405.2
159 Pantoea dispersa EGD-AAK13 GCA_000465555.2
159 Shimwellia blattae DSM 4481 = NBRC 105725 GCA_000262305.1
jz
89 Bacillus anthracis str. Ames GCA_000007845.1
89 Bacillus anthracis str. Sterne GCA_000008165.1
89 Bacillus cereus ATCC 14579 GCA_000007825.1
89 Bacillus mycoides GCA_000832605.1
89 Bacillus pseudomycoides DSM 12442 GCA_000161455.1
89 Bacillus thuringiensis YBT-1518 GCA_000497525.2
89 _5BBacillus thuringiensis_5D serovar konkukian str. 97-27 GCA_000008505.1
85 Pelosinus fermentans JBW45 GCA_000271665.2
79 Clostridium beijerinckii GCA_000833105.2
79 Clostridium puniceum GCA_002006345.1
79 Clostridium saccharoperbutylacetonicum N1-4_28HMT_29 GCA_000340885.1
79 Granulicatella balaenopterae GCA_900111135.1
79 Listeria aquatica FSL S10-1188 GCA_000525795.1

79 Listeria floridensis FSL S10-1187 GCA_000525875.1
jA
265 Lactobacillus gasseri ATCC 33323 = JCM 1131 GCA_000014425.1
255 Lactobacillus hominis DSM 23910 = CRBIP 24.179 GCA_000296835.1
201 Lactobacillus psittaci DSM 15354 GCA_000425905.1
jB
239 Rhodobacter sphaeroides 2.4.1 GCA_000012905.2
200 Rhodobacter capsulatus SB 1003 GCA_000021865.1
198 Gemmobacter megaterium GCA_900156815.1
jC
263 Deinococcus radiodurans R1 GCA_000008565.1
177 Deinococcus gobiensis I-0 GCA_000252445.1
169 Deinococcus puniceus GCA_001644565.1
jD
199 Rhodobacter sphaeroides ATCC 17025 GCA_000016405.1
181 Rhodobacter aestuarii GCA_900156655.1
176 Rhodobacter sphaeroides 2.4.1 GCA_000012905.2
176 Rhodobacter vinaykumarii GCA_900156695.1
jE
215 Rhodobacter sphaeroides 2.4.1 GCA_000012905.2
172 Rhodobacter sphaeroides ATCC 17025 GCA_000016405.1
165 Thioclava dalianensis GCA_000715505.1
165 Thioclava indica GCA_000714545.1
jF
257 Bifidobacterium adolescentis ATCC 15703 GCA_000010425.1
201 Bifidobacterium breve DSM 20213 = JCM 1192 GCA_001025175.1
198 Bifidobacterium thermophilum GCA_000741495.1
jG
166 Escherichia coli IAI39 GCA_000026345.1
166 Escherichia coli O104_3AH4 str. 2011C-3493 GCA_000299455.1
166 Escherichia coli O157_3AH7 str. Sakai GCA_000008865.1
166 Escherichia coli O83_3AH1 str. NRG 857C GCA_000183345.1
166 Escherichia coli UMN026 GCA_000026325.2
166 Escherichia coli str. K-12 substr. MG1655 GCA_000005845.2
166 Shigella dysenteriae Sd197 GCA_000012005.1
166 Shigella flexneri 2a str. 301 GCA_000006925.2
166 Tumebacillus flagellatus GCA_000714935.1
154 _5BEnterobacter_5D lignolyticus SCF1 GCA_000164865.1
151 Buttiauxella ferragutiae ATCC 51602 GCA_001654915.1
151 Cedecea neteri GCA_000757825.1
151 Citrobacter freundii CFNIH1 GCA_000648515.1
151 Cronobacter sakazakii GCA_000982825.1
151 Enterobacter cloacae subsp. cloacae ATCC 13047 GCA_000025565.1
151 Enterobacter hormaechei subsp. steigerwaltii GCA_001729725.1
151 Enterobacter kobei GCA_900185885.1
151 Erwinia iniecta GCA_001267535.1
151 Erwinia persicina NBRC 102418 GCA_001571305.1
151 Erwinia toletana DAPP-PG 735 GCA_000336255.1
151 Klebsiella oxytoca GCA_001022195.1
151 Klebsiella pneumoniae subsp. pneumoniae HS11286 GCA_000240185.2
151 Kluyvera ascorbata ATCC 33433 GCA_000735365.1
151 Kosakonia cowanii GCA_001975225.1
151 Pseudescherichia vulneris NBRC 102420 GCA_000759795.1
151 Rosenbergiella nectarea GCA_900111105.1
151 Salmonella enterica subsp. enterica serovar Typhi str. CT18 GCA_000195995.1
151 Yokenella regensburgei ATCC 49455 GCA_000735455.1
jH
193 Bifidobacterium adolescentis ATCC 15703 GCA_000010425.1
170 Bifidobacterium callitrichos DSM 23973 GCA_000741175.1
163 Bifidobacterium angulatum DSM 20098 = JCM 7096 GCA_001025155.1
jI
266 Bacillus anthracis str. Ames GCA_000007845.1
266 Bacillus anthracis str. Sterne GCA_000008165.1
266 Bacillus cereus ATCC 14579 GCA_000007825.1
266 Bacillus mycoides GCA_000832605.1
266 Bacillus pseudomycoides DSM 12442 GCA_000161455.1
266 Bacillus thuringiensis YBT-1518 GCA_000497525.2
266 _5BBacillus thuringiensis_5D serovar konkukian str. 97-27 GCA_000008505.1
214 Bacillus manliponensis GCA_000712595.1
209 Bacillus horneckiae GCA_001636335.1
209 Bacillus solani GCA_001420595.1
jJ
226 Staphylococcus capitis subsp. capitis GCA_001028645.1
226 Staphylococcus cohnii subsp. cohnii GCA_000972575.1
226 Staphylococcus epidermidis ATCC 12228 GCA_000007645.1
226 Staphylococcus lugdunensis HKU09-01 GCA_000025085.1

226 Staphylococcus saprophyticus subsp. saprophyticus ATCC 15305 GCA_000010125.1
219 Staphylococcus arlettae CVD059 GCA_000295715.1
219 Staphylococcus haemolyticus JCSC1435 GCA_000009865.1
219 Staphylococcus hominis subsp. hominis C80 GCA_000183685.1
217 Staphylococcus equorum GCA_001432245.1
217 Staphylococcus xylosus GCA_000706685.1
jK
193 Bifidobacterium adolescentis ATCC 15703 GCA_000010425.1
179 Bifidobacterium breve DSM 20213 = JCM 1192 GCA_001025175.1
179 Bifidobacterium longum NCC2705 GCA_000007525.1
174 Bifidobacterium saguini DSM 23967 GCA_000741715.1
jL
222 Deinococcus radiodurans R1 GCA_000008565.1
145 Deinococcus marmoris DSM 12784 GCA_000701405.1
145 Deinococcus swuensis GCA_000800395.1
139 Deinococcus geothermalis DSM 11300 GCA_000196275.1
jM
325 Deinococcus radiodurans R1 GCA_000008565.1
236 Deinococcus gobiensis I-0 GCA_000252445.1
236 Deinococcus puniceus GCA_001644565.1
216 Deinococcus deserti VCD115 GCA_000020685.1
jN
256 Bifidobacterium dentium JCM 1195 = DSM 20436 GCA_001042595.1
241 Bifidobacterium tsurumiense GCA_000741765.1
235 Bifidobacterium adolescentis ATCC 15703 GCA_000010425.1
jO
246 Deinococcus radiodurans R1 GCA_000008565.1
180 Deinococcus gobiensis I-0 GCA_000252445.1
163 Deinococcus deserti VCD115 GCA_000020685.1
jP
302 Streptococcus mutans UA159 GCA_000007465.2
178 Streptococcus macacae NCTC 11558 GCA_000187995.3
175 Streptococcus ferus DSM 20646 GCA_000372425.1
jQ
243 Deinococcus radiodurans R1 GCA_000008565.1
211 Deinococcus puniceus GCA_001644565.1
188 Deinococcus hopiensis KR-140 GCA_900176165.1
jR
237 Clostridium beijerinckii GCA_000833105.2
237 Clostridium saccharobutylicum DSM 13864 GCA_000473995.1
237 Clostridium saccharoperbutylacetonicum N1-4_28HMT_29 GCA_000340885.1
233 Clostridium butyricum GCA_001456065.2
230 Clostridium puniceum GCA_002006345.1
jS
164 Rhodobacter sphaeroides 2.4.1 GCA_000012905.2
133 Rhodobacter sphaeroides ATCC 17025 GCA_000016405.1
124 Thioclava dalianensis GCA_000715505.1
124 Thioclava indica GCA_000714545.1
jT
155 Bacillus anthracis str. Ames GCA_000007845.1
155 Bacillus anthracis str. Sterne GCA_000008165.1
155 Bacillus cereus ATCC 14579 GCA_000007825.1
155 Bacillus mycoides GCA_000832605.1
155 Bacillus pseudomycoides DSM 12442 GCA_000161455.1
155 Bacillus thuringiensis YBT-1518 GCA_000497525.2
155 _5BBacillus thuringiensis_5D serovar konkukian str. 97-27 GCA_000008505.1
147 Bacillus flexus GCA_002024265.1
147 Bacillus megaterium NBRC 15308 = ATCC 14581 GCA_000832985.1
133 Bacillus coagulans DSM 1 = ATCC 7050 GCA_000832905.1
133 Planococcus antarcticus DSM 14505 GCA_001687565.2
133 Planococcus kocurii GCA_001465835.2
jU
190 Streptococcus mutans UA159 GCA_000007465.2
158 Streptococcus sobrinus DSM 20742 = ATCC 33478 GCA_000686605.1
138 Streptococcus macacae NCTC 11558 GCA_000187995.3
jV
274 Lactobacillus gasseri ATCC 33323 = JCM 1131 GCA_000014425.1
255 Lactobacillus hominis DSM 23910 = CRBIP 24.179 GCA_000296835.1
198 Lactobacillus amylovorus GCA_000191545.1
198 Lactobacillus crispatus ST1 GCA_000091765.1
198 Lactobacillus kalixensis DSM 16043 GCA_001434335.1
jW
99 Deinococcus radiodurans R1 GCA_000008565.1
77 Deinococcus soli Cha et al. 2016 GCA_001007995.1
74 Deinococcus puniceus GCA_001644565.1
jX

294 Staphylococcus aureus subsp. aureus NCTC 8325 GCA_000013425.1
294 Staphylococcus capitis subsp. capitis GCA_001028645.1
294 Staphylococcus epidermidis ATCC 12228 GCA_000007645.1
294 Staphylococcus haemolyticus JCSC1435 GCA_000009865.1
294 Staphylococcus hominis subsp. hominis C80 GCA_000183685.1
294 Staphylococcus simiae CCM 7213 GCA_000235645.2
285 Staphylococcus condimenti GCA_001618885.1
285 Staphylococcus simulans GCA_001559115.1
282 Staphylococcus lugdunensis HKU09-01 GCA_000025085.1
jY
273 Escherichia coli IAI39 GCA_000026345.1
273 Escherichia coli O104_3AH4 str. 2011C-3493 GCA_000299455.1
273 Escherichia coli O157_3AH7 str. Sakai GCA_000008865.1
273 Escherichia coli O83_3AH1 str. NRG 857C GCA_000183345.1
273 Escherichia coli UMN026 GCA_000026325.2
273 Escherichia coli str. K-12 substr. MG1655 GCA_000005845.2
273 Shigella dysenteriae Sd197 GCA_000012005.1
273 Tumebacillus flagellatus GCA_000714935.1
265 Shigella flexneri 2a str. 301 GCA_000006925.2
247 Erwinia iniecta GCA_001267535.1
jZ
260 Escherichia coli IAI39 GCA_000026345.1
260 Escherichia coli O104_3AH4 str. 2011C-3493 GCA_000299455.1
260 Escherichia coli O157_3AH7 str. Sakai GCA_000008865.1
260 Escherichia coli UMN026 GCA_000026325.2
260 Escherichia coli str. K-12 substr. MG1655 GCA_000005845.2
260 Shigella flexneri 2a str. 301 GCA_000006925.2
260 Tumebacillus flagellatus GCA_000714935.1
259 Escherichia coli O83_3AH1 str. NRG 857C GCA_000183345.1
246 Shigella dysenteriae Sd197 GCA_000012005.1
k0
201 Bifidobacterium adolescentis ATCC 15703 GCA_000010425.1
165 Bifidobacterium choerinum GCA_000741135.1
165 Bifidobacterium pseudolongum PV8-2 GCA_000800475.2
163 Bifidobacterium breve DSM 20213 = JCM 1192 GCA_001025175.1
163 Bifidobacterium longum NCC2705 GCA_000007525.1
163 Bifidobacterium saguini DSM 23967 GCA_000741715.1
k1
182 Enterococcus faecalis V583 GCA_000007785.1
182 Streptomyces cinnamoneus GCA_001885705.1
178 Vagococcus penaei GCA_001998885.1
169 Enterococcus dispar ATCC 51266 GCA_000406945.1
k2
149 Clostridium beijerinckii GCA_000833105.2
149 Clostridium puniceum GCA_002006345.1
149 Clostridium saccharoperbutylacetonicum N1-4_28HMT_29 GCA_000340885.1
134 Clostridium saccharobutylicum DSM 13864 GCA_000473995.1
132 Clostridium butyricum GCA_001456065.2
132 Clostridium neonatale GCA_001458595.1
132 Clostridium taeniosporum GCA_001735765.1
k3
176 Escherichia coli IAI39 GCA_000026345.1
176 Escherichia coli O104_3AH4 str. 2011C-3493 GCA_000299455.1
176 Escherichia coli O157_3AH7 str. Sakai GCA_000008865.1
176 Escherichia coli O83_3AH1 str. NRG 857C GCA_000183345.1
176 Escherichia coli UMN026 GCA_000026325.2
176 Escherichia coli str. K-12 substr. MG1655 GCA_000005845.2
176 Shigella dysenteriae Sd197 GCA_000012005.1
176 Shigella flexneri 2a str. 301 GCA_000006925.2
176 Tumebacillus flagellatus GCA_000714935.1
158 Kosakonia cowanii GCA_001975225.1
157 Enterobacter hormaechei subsp. steigerwaltii GCA_001729725.1
157 Erwinia iniecta GCA_001267535.1
157 Erwinia toletana DAPP-PG 735 GCA_000336255.1
157 Klebsiella oxytoca GCA_001022195.1
157 Kosakonia sacchari SP1 GCA_000300455.4
157 Pluralibacter gergoviae GCA_000757785.1
157 Pseudescherichia vulneris NBRC 102420 GCA_000759795.1
k4
148 Bifidobacterium adolescentis ATCC 15703 GCA_000010425.1
103 Bifidobacterium asteroides PRL2011 GCA_000304215.1
95 Bifidobacterium animalis subsp. lactis DSM 10140 GCA_000022965.1
95 Bifidobacterium breve DSM 20213 = JCM 1192 GCA_001025175.1
95 Bifidobacterium callitrichos DSM 23973 GCA_000741175.1
95 Bifidobacterium choerinum GCA_000741135.1
95 Bifidobacterium dentium JCM 1195 = DSM 20436 GCA_001042595.1

95 Bifidobacterium longum NCC2705 GCA_000007525.1
95 Bifidobacterium pseudolongum PV8-2 GCA_000800475.2
95 Bifidobacterium thermophilum GCA_000741495.1
95 Bifidobacterium thermophilum RBL67 GCA_000347695.1
95 Bifidobacterium tsurumiense GCA_000741765.1
k5
276 Clostridium beijerinckii GCA_000833105.2
276 Clostridium saccharobutylicum DSM 13864 GCA_000473995.1
276 Clostridium saccharoperbutylacetonicum N1-4_28HMT_29 GCA_000340885.1
269 Clostridium puniceum GCA_002006345.1
254 Clostridium chromiireducens GCA_002029255.1
k6
115 Enterococcus canis NBRC 100695 GCA_001544375.1
115 Enterococcus casseliflavus EC20 GCA_000157355.2
115 Enterococcus dispar ATCC 51266 GCA_000406945.1
115 Enterococcus faecalis V583 GCA_000007785.1
115 Enterococcus faecium DO GCA_000174395.2
115 Enterococcus haemoperoxidus ATCC BAA-382 GCA_000407165.1
115 Enterococcus hirae ATCC 9790 GCA_000271405.2
115 Enterococcus massiliensis GCA_001050095.1
115 Enterococcus mundtii QU 25 GCA_000504125.1
115 Enterococcus phoeniculicola ATCC BAA-412 GCA_000407505.1
115 Enterococcus rivorum GCA_001742285.1
115 Enterococcus saccharolyticus subsp. saccharolyticus ATCC 43076 GCA_000407285.1
115 Enterococcus thailandicus GCA_001652875.1
115 Isobaculum melis GCA_900111355.1
115 Streptomyces cinnamoneus GCA_001885705.1
106 Lactobacillus kimchicus JCM 15530 GCA_001433995.1
106 Lactobacillus mixtipabuli GCA_002217925.1
106 Lactobacillus nasuensis JCM 17158 GCA_001434705.1
106 Lactobacillus odoratitofui DSM 19909 = JCM 15043 GCA_001434895.1
106 Lactobacillus silagei JCM 19001 GCA_002217945.1
106 Lactobacillus similis DSM 23365 = JCM 2765 GCA_001311075.1
105 Enterococcus asini ATCC 700915 GCA_000407365.1
105 Enterococcus cecorum GCA_001318405.1
105 Enterococcus columbae DSM 7374 = ATCC 51263 GCA_000406925.1
k7
173 Deinococcus radiodurans R1 GCA_000008565.1
127 Deinococcus puniceus GCA_001644565.1
122 Deinococcus proteolyticus MRP GCA_000190555.1
k8
157 Deinococcus radiodurans R1 GCA_000008565.1
131 Deinococcus puniceus GCA_001644565.1
118 Deinococcus gobiensis I-0 GCA_000252445.1
k9
223 Clostridium beijerinckii GCA_000833105.2
223 Clostridium puniceum GCA_002006345.1
223 Clostridium saccharoperbutylacetonicum N1-4_28HMT_29 GCA_000340885.1
214 Clostridium saccharobutylicum DSM 13864 GCA_000473995.1
192 Clostridium taeniosporum GCA_001735765.1
ka
258 Streptococcus mutans UA159 GCA_000007465.2
169 Streptococcus ratti FA-1 = DSM 20564 GCA_000286075.1
167 Streptococcus gordonii str. Challis substr. CH1 GCA_000017005.1
kb
148 Rhodobacter sphaeroides 2.4.1 GCA_000012905.2
99 Rhodobacter sphaeroides ATCC 17025 GCA_000016405.1
81 Haematobacter massiliensis GCA_000740795.1
81 Pseudorhodobacter psychrotolerans GCA_001294535.1
kc
200 Streptococcus mutans UA159 GCA_000007465.2
150 Streptococcus marimammalium DSM 18627 GCA_000380045.1
143 Streptococcus macacae NCTC 11558 GCA_000187995.3
kd
156 Rhodobacter sphaeroides 2.4.1 GCA_000012905.2
136 Gemmobacter megaterium GCA_900156815.1
127 Pseudorhodobacter ferrugineus DSM 5888 GCA_000420745.1
127 Pseudorhodobacter wandonensis GCA_001202035.1
ke
241 Deinococcus radiodurans R1 GCA_000008565.1
182 Deinococcus gobiensis I-0 GCA_000252445.1
170 Deinococcus hopiensis KR-140 GCA_900176165.1
170 Deinococcus puniceus GCA_001644565.1
kf
204 Staphylococcus capitis subsp. capitis GCA_001028645.1
204 Staphylococcus epidermidis ATCC 12228 GCA_000007645.1

188 Megasphaera cerevisiae DSM 20462 GCA_001045675.1
188 Staphylococcus haemolyticus JCSC1435 GCA_000009865.1
188 Staphylococcus hyicus GCA_000816085.1
188 Staphylococcus warneri SG1 GCA_000332735.1
173 Staphylococcus aureus subsp. aureus NCTC 8325 GCA_000013425.1
173 Staphylococcus hominis subsp. hominis C80 GCA_000183685.1
173 Staphylococcus lugdunensis HKU09-01 GCA_000025085.1
173 Staphylococcus lutrae GCA_002101335.1
173 Staphylococcus simiae CCM 7213 GCA_000235645.2
kg
253 Escherichia coli IAI39 GCA_000026345.1
253 Escherichia coli O104_3AH4 str. 2011C-3493 GCA_000299455.1
253 Escherichia coli O157_3AH7 str. Sakai GCA_000008865.1
253 Escherichia coli O83_3AH1 str. NRG 857C GCA_000183345.1
253 Escherichia coli UMN026 GCA_000026325.2
253 Escherichia coli str. K-12 substr. MG1655 GCA_000005845.2
253 Shigella dysenteriae Sd197 GCA_000012005.1
253 Shigella flexneri 2a str. 301 GCA_000006925.2
253 Tumebacillus flagellatus GCA_000714935.1
214 Erwinia iniecta GCA_001267535.1
205 Cronobacter sakazakii GCA_000982825.1
205 Erwinia toletana DAPP-PG 735 GCA_000336255.1
kh
329 Escherichia coli IAI39 GCA_000026345.1
329 Escherichia coli O104_3AH4 str. 2011C-3493 GCA_000299455.1
329 Escherichia coli O157_3AH7 str. Sakai GCA_000008865.1
329 Escherichia coli UMN026 GCA_000026325.2
329 Escherichia coli str. K-12 substr. MG1655 GCA_000005845.2
329 Shigella flexneri 2a str. 301 GCA_000006925.2
329 Tumebacillus flagellatus GCA_000714935.1
328 Escherichia coli O83_3AH1 str. NRG 857C GCA_000183345.1
323 Erwinia iniecta GCA_001267535.1
ki
182 Clostridium beijerinckii GCA_000833105.2
182 Clostridium puniceum GCA_002006345.1
182 Clostridium saccharoperbutylacetonicum N1-4_28HMT_29 GCA_000340885.1
174 Clostridium saccharobutylicum DSM 13864 GCA_000473995.1
155 Clostridium chromiireducens GCA_002029255.1
kj
206 Rhodobacter sphaeroides 2.4.1 GCA_000012905.2
188 Rhodobacter sphaeroides ATCC 17025 GCA_000016405.1
183 Pseudorhodobacter psychrotolerans GCA_001294535.1
kk
183 Bacillus thuringiensis YBT-1518 GCA_000497525.2
168 Bacillus anthracis str. Ames GCA_000007845.1
168 Bacillus anthracis str. Sterne GCA_000008165.1
168 Bacillus mycoides GCA_000832605.1
168 Bacillus pseudomycoides DSM 12442 GCA_000161455.1
168 _5BBacillus thuringiensis_5D serovar konkukian str. 97-27 GCA_000008505.1
167 Bacillus cereus ATCC 14579 GCA_000007825.1
kl
162 Rhodobacter sphaeroides 2.4.1 GCA_000012905.2
151 Rhodobacter sphaeroides ATCC 17025 GCA_000016405.1
124 Pseudorhodobacter ferrugineus DSM 5888 GCA_000420745.1
124 Pseudorhodobacter wandonensis GCA_001202035.1
km
190 Escherichia coli IAI39 GCA_000026345.1
190 Escherichia coli O104_3AH4 str. 2011C-3493 GCA_000299455.1
190 Escherichia coli O157_3AH7 str. Sakai GCA_000008865.1
190 Escherichia coli O83_3AH1 str. NRG 857C GCA_000183345.1
190 Escherichia coli UMN026 GCA_000026325.2
190 Escherichia coli str. K-12 substr. MG1655 GCA_000005845.2
190 Shigella dysenteriae Sd197 GCA_000012005.1
190 Shigella flexneri 2a str. 301 GCA_000006925.2
190 Tumebacillus flagellatus GCA_000714935.1
164 Buttiauxella ferragutiae ATCC 51602 GCA_001654915.1
164 Cedecea neteri GCA_000757825.1
164 Citrobacter freundii CFNIH1 GCA_000648515.1
164 Cronobacter sakazakii GCA_000982825.1
164 Enterobacter cloacae subsp. cloacae ATCC 13047 GCA_000025565.1
164 Enterobacter hormaechei subsp. steigerwaltii GCA_001729725.1
164 Enterobacter kobei GCA_900185885.1
164 Erwinia iniecta GCA_001267535.1
164 Erwinia persicina NBRC 102418 GCA_001571305.1
164 Erwinia toletana DAPP-PG 735 GCA_000336255.1
164 Klebsiella oxytoca GCA_001022195.1

164 Klebsiella pneumoniae subsp. pneumoniae HS11286 GCA_000240185.2
164 Kluyvera ascorbata ATCC 33433 GCA_000735365.1
164 Kluyvera cryocrescens NBRC 102467 GCA_001571285.1
164 Kosakonia cowanii GCA_001975225.1
164 Kosakonia sacchari SP1 GCA_000300455.4
164 Mangrovibacter phragmitis GCA_001655675.1
164 Pseudescherichia vulneris NBRC 102420 GCA_000759795.1
164 Rosenbergiella nectarea GCA_900111105.1
164 Salmonella enterica subsp. enterica serovar Typhi str. CT18 GCA_000195995.1
164 Yokenella regensburgei ATCC 49455 GCA_000735455.1
164 _5BEnterobacter_5D lignolyticus SCF1 GCA_000164865.1
161 Salmonella enterica subsp. enterica serovar Typhimurium str. LT2 GCA_000006945.2
kn
228 Deinococcus radiodurans R1 GCA_000008565.1
183 Deinococcus deserti VCD115 GCA_000020685.1
157 Deinococcus soli Cha et al. 2016 GCA_001007995.1
ko
213 Escherichia coli IAI39 GCA_000026345.1
213 Escherichia coli O104_3AH4 str. 2011C-3493 GCA_000299455.1
213 Escherichia coli O157_3AH7 str. Sakai GCA_000008865.1
213 Escherichia coli O83_3AH1 str. NRG 857C GCA_000183345.1
213 Escherichia coli UMN026 GCA_000026325.2
213 Escherichia coli str. K-12 substr. MG1655 GCA_000005845.2
213 Shigella flexneri 2a str. 301 GCA_000006925.2
213 Tumebacillus flagellatus GCA_000714935.1
209 Erwinia iniecta GCA_001267535.1
198 Shigella dysenteriae Sd197 GCA_000012005.1
kp
252 Clostridium saccharobutylicum DSM 13864 GCA_000473995.1
248 Clostridium beijerinckii GCA_000833105.2
248 Clostridium chromiireducens GCA_002029255.1
248 Clostridium puniceum GCA_002006345.1
248 Clostridium saccharoperbutylacetonicum N1-4_28HMT_29 GCA_000340885.1
244 Clostridium butyricum GCA_001456065.2
kq
251 Clostridium beijerinckii GCA_000833105.2
251 Clostridium saccharobutylicum DSM 13864 GCA_000473995.1
251 Clostridium saccharoperbutylacetonicum N1-4_28HMT_29 GCA_000340885.1
249 Clostridium butyricum GCA_001456065.2
237 Clostridium ventriculi GCA_001404895.1
kr
290 Bacillus anthracis str. Ames GCA_000007845.1
290 Bacillus anthracis str. Sterne GCA_000008165.1
290 Bacillus cereus ATCC 14579 GCA_000007825.1
290 Bacillus mycoides GCA_000832605.1
290 Bacillus pseudomycoides DSM 12442 GCA_000161455.1
290 Bacillus thuringiensis YBT-1518 GCA_000497525.2
290 _5BBacillus thuringiensis_5D serovar konkukian str. 97-27 GCA_000008505.1
254 Bacillus manliponensis GCA_000712595.1
230 Bacillus cytotoxicus NVH 391-98 GCA_000017425.1
ks
272 Clostridium beijerinckii GCA_000833105.2
272 Clostridium puniceum GCA_002006345.1
272 Clostridium saccharoperbutylacetonicum N1-4_28HMT_29 GCA_000340885.1
263 Clostridium saccharobutylicum DSM 13864 GCA_000473995.1
238 Clostridium butyricum GCA_001456065.2
kt
195 Deinococcus radiodurans R1 GCA_000008565.1
134 Deinococcus hopiensis KR-140 GCA_900176165.1
132 Deinococcus puniceus GCA_001644565.1
ku
238 Escherichia coli IAI39 GCA_000026345.1
238 Escherichia coli O104_3AH4 str. 2011C-3493 GCA_000299455.1
238 Escherichia coli O157_3AH7 str. Sakai GCA_000008865.1
238 Escherichia coli O83_3AH1 str. NRG 857C GCA_000183345.1
238 Escherichia coli UMN026 GCA_000026325.2
238 Escherichia coli str. K-12 substr. MG1655 GCA_000005845.2
238 Shigella dysenteriae Sd197 GCA_000012005.1
238 Tumebacillus flagellatus GCA_000714935.1
230 Shigella flexneri 2a str. 301 GCA_000006925.2
215 Cronobacter sakazakii GCA_000982825.1
215 Enterobacter cloacae subsp. cloacae ATCC 13047 GCA_000025565.1
215 Enterobacter hormaechei subsp. steigerwaltii GCA_001729725.1
215 Enterobacter kobei GCA_900185885.1
215 Erwinia amylovora CFBP1430 GCA_000091565.1
215 Erwinia billingiae Eb661 GCA_000196615.1

215 Erwinia gerundensis GCA_001517405.1
215 Erwinia iniecta GCA_001267535.1
215 Erwinia toletana DAPP-PG 735 GCA_000336255.1
215 Klebsiella oxytoca GCA_001022195.1
215 Kosakonia cowanii GCA_001975225.1
215 Pantoea agglomerans GCA_001709315.1
215 Pantoea alhagi GCA_002101395.1
215 Pantoea ananatis LMG 20103 GCA_000025405.2
215 Pantoea dispersa EGD-AAK13 GCA_000465555.2
215 Pantoea rwandensis GCA_000759475.1
215 Pantoea septica GCA_002095575.1
215 Pluralibacter gergoviae GCA_000757785.1
215 Pseudescherichia vulneris NBRC 102420 GCA_000759795.1
215 Salmonella enterica subsp. enterica serovar Typhi str. CT18 GCA_000195995.1
215 Salmonella enterica subsp. enterica serovar Typhimurium str. LT2 GCA_000006945.2
215 Shimwellia blattae DSM 4481 = NBRC 105725 GCA_000262305.1
215 Trabulsiella odontotermitis GCA_001297765.1
kv
289 Bifidobacterium adolescentis ATCC 15703 GCA_000010425.1
268 Bifidobacterium longum NCC2705 GCA_000007525.1
266 Bifidobacterium breve DSM 20213 = JCM 1192 GCA_001025175.1
kw
213 Escherichia coli IAI39 GCA_000026345.1
213 Escherichia coli O104_3AH4 str. 2011C-3493 GCA_000299455.1
213 Escherichia coli O157_3AH7 str. Sakai GCA_000008865.1
213 Escherichia coli O83_3AH1 str. NRG 857C GCA_000183345.1
213 Escherichia coli UMN026 GCA_000026325.2
213 Escherichia coli str. K-12 substr. MG1655 GCA_000005845.2
213 Shigella dysenteriae Sd197 GCA_000012005.1
213 Shigella flexneri 2a str. 301 GCA_000006925.2
213 Tumebacillus flagellatus GCA_000714935.1
207 Erwinia iniecta GCA_001267535.1
186 Rosenbergiella nectarea GCA_900111105.1
kx
178 Escherichia coli IAI39 GCA_000026345.1
178 Escherichia coli O104_3AH4 str. 2011C-3493 GCA_000299455.1
178 Escherichia coli O157_3AH7 str. Sakai GCA_000008865.1
178 Escherichia coli O83_3AH1 str. NRG 857C GCA_000183345.1
178 Escherichia coli UMN026 GCA_000026325.2
178 Escherichia coli str. K-12 substr. MG1655 GCA_000005845.2
178 Shigella dysenteriae Sd197 GCA_000012005.1
178 Shigella flexneri 2a str. 301 GCA_000006925.2
178 Tumebacillus flagellatus GCA_000714935.1
172 Erwinia iniecta GCA_001267535.1
172 Erwinia toletana DAPP-PG 735 GCA_000336255.1
172 Pantoea agglomerans GCA_001709315.1
172 Pantoea ananatis LMG 20103 GCA_000025405.2
172 Shimwellia blattae DSM 4481 = NBRC 105725 GCA_000262305.1
168 Cronobacter sakazakii GCA_000982825.1
168 Pantoea dispersa EGD-AAK13 GCA_000465555.2
168 Plautia stali symbiont GCA_000180175.2
ky
224 Bifidobacterium adolescentis ATCC 15703 GCA_000010425.1
199 Bifidobacterium dentium JCM 1195 = DSM 20436 GCA_001042595.1
194 Bifidobacterium asteroides PRL2011 GCA_000304215.1
kz
225 Deinococcus radiodurans R1 GCA_000008565.1
140 Deinococcus deserti VCD115 GCA_000020685.1
124 Deinococcus marmoris DSM 12784 GCA_000701405.1
124 Deinococcus swuensis GCA_000800395.1
kA
298 Deinococcus radiodurans R1 GCA_000008565.1
209 Deinococcus gobiensis I-0 GCA_000252445.1
207 Deinococcus deserti VCD115 GCA_000020685.1
kB
147 Clostridium beijerinckii GCA_000833105.2
147 Clostridium puniceum GCA_002006345.1
147 Clostridium saccharobutylicum DSM 13864 GCA_000473995.1
147 Clostridium saccharoperbutylacetonicum N1-4_28HMT_29 GCA_000340885.1
136 Clostridium chromiireducens GCA_002029255.1
121 Clostridium butyricum GCA_001456065.2
kC
232 Rhodobacter sphaeroides 2.4.1 GCA_000012905.2
228 Pseudorhodobacter psychrotolerans GCA_001294535.1
211 Gemmobacter aquatilis GCA_900110025.1
kD

227 Escherichia coli IAI39 GCA_000026345.1
227 Escherichia coli O104_3AH4 str. 2011C-3493 GCA_000299455.1
227 Escherichia coli O157_3AH7 str. Sakai GCA_000008865.1
227 Escherichia coli O83_3AH1 str. NRG 857C GCA_000183345.1
227 Escherichia coli UMN026 GCA_000026325.2
227 Escherichia coli str. K-12 substr. MG1655 GCA_000005845.2
227 Shigella dysenteriae Sd197 GCA_000012005.1
227 Shigella flexneri 2a str. 301 GCA_000006925.2
227 Tumebacillus flagellatus GCA_000714935.1
212 Erwinia iniecta GCA_001267535.1
204 Klebsiella pneumoniae subsp. pneumoniae HS11286 GCA_000240185.2
kE
106 Clostridium beijerinckii GCA_000833105.2
106 Clostridium chromiireducens GCA_002029255.1
106 Clostridium puniceum GCA_002006345.1
106 Clostridium saccharobutylicum DSM 13864 GCA_000473995.1
106 Clostridium saccharoperbutylacetonicum N1-4_28HMT_29 GCA_000340885.1
97 Clostridium butyricum GCA_001456065.2
82 Clostridium neonatale GCA_001458595.1
kF
212 Clostridium beijerinckii GCA_000833105.2
212 Clostridium puniceum GCA_002006345.1
212 Clostridium saccharoperbutylacetonicum N1-4_28HMT_29 GCA_000340885.1
198 Clostridium saccharobutylicum DSM 13864 GCA_000473995.1
186 Clostridium taeniosporum GCA_001735765.1
kG
132 Bifidobacterium adolescentis ATCC 15703 GCA_000010425.1
107 Bifidobacterium angulatum DSM 20098 = JCM 7096 GCA_001025155.1
103 Bifidobacterium breve DSM 20213 = JCM 1192 GCA_001025175.1
103 Bifidobacterium longum NCC2705 GCA_000007525.1
103 Bifidobacterium saguini DSM 23967 GCA_000741715.1
kH
173 Bacillus anthracis str. Ames GCA_000007845.1
173 Bacillus anthracis str. Sterne GCA_000008165.1
173 Bacillus cereus ATCC 14579 GCA_000007825.1
173 Bacillus mycoides GCA_000832605.1
173 Bacillus pseudomycoides DSM 12442 GCA_000161455.1
173 Bacillus thuringiensis YBT-1518 GCA_000497525.2
173 _5BBacillus thuringiensis_5D serovar konkukian str. 97-27 GCA_000008505.1
132 Bacillus horneckiae GCA_001636335.1
132 Bacillus solani GCA_001420595.1
131 Bacillus halmapalus GCA_002019665.1
kI
208 Streptococcus mutans UA159 GCA_000007465.2
163 Streptococcus ferus DSM 20646 GCA_000372425.1
160 Streptococcus macacae NCTC 11558 GCA_000187995.3
kJ
168 Enterococcus faecalis V583 GCA_000007785.1
168 Streptomyces cinnamoneus GCA_001885705.1
145 Viridibacillus arvi GCA_001274945.1
143 Listeria floridensis FSL S10-1187 GCA_000525875.1
kK
286 Rhodobacter sphaeroides 2.4.1 GCA_000012905.2
274 Rhodobacter sphaeroides ATCC 17025 GCA_000016405.1
267 Pseudorhodobacter psychrotolerans GCA_001294535.1
kL
270 Clostridium beijerinckii GCA_000833105.2
270 Clostridium saccharoperbutylacetonicum N1-4_28HMT_29 GCA_000340885.1
266 Clostridium puniceum GCA_002006345.1
262 Clostridium saccharobutylicum DSM 13864 GCA_000473995.1
kM
153 Deinococcus radiodurans R1 GCA_000008565.1
96 Deinococcus gobiensis I-0 GCA_000252445.1
94 Deinococcus deserti VCD115 GCA_000020685.1
94 Deinococcus soli Cha et al. 2016 GCA_001007995.1
kN
241 Staphylococcus epidermidis ATCC 12228 GCA_000007645.1
241 Staphylococcus haemolyticus JCSC1435 GCA_000009865.1
233 Staphylococcus capitis subsp. capitis GCA_001028645.1
230 Staphylococcus hominis subsp. hominis C80 GCA_000183685.1
kO
308 Lactobacillus gasseri ATCC 33323 = JCM 1131 GCA_000014425.1
290 Lactobacillus hominis DSM 23910 = CRBIP 24.179 GCA_000296835.1
227 Lactobacillus acidophilus NCFM GCA_000011985.1
227 Lactobacillus crispatus ST1 GCA_000091765.1
kP

142 Clostridium amylolyticum GCA_900142075.1
142 Clostridium intestinale URNW GCA_000469625.2
142 Clostridium polynesiense GCA_000820705.1
127 Clostridium beijerinckii GCA_000833105.2
127 Clostridium saccharobutylicum DSM 13864 GCA_000473995.1
127 Clostridium saccharoperbutylacetonicum N1-4_28HMT_29 GCA_000340885.1
127 Desnuesiella massiliensis GCA_001403615.1
120 Clostridium algidicarnis GCA_000703125.1
120 Clostridium botulinum B str. Eklund 17B _28NRP_29 GCA_000020165.1
120 Clostridium fallax GCA_900129365.1
120 Clostridium gasigenes GCA_900104115.1
120 Clostridium neonatale GCA_001458595.1
120 Clostridium puniceum GCA_002006345.1
120 Clostridium taeniosporum GCA_001735765.1
120 Clostridium uliginosum GCA_900112485.1
kQ
127 Staphylococcus epidermidis ATCC 12228 GCA_000007645.1
124 Staphylococcus haemolyticus JCSC1435 GCA_000009865.1
123 Staphylococcus cohnii subsp. cohnii GCA_000972575.1
kR
209 Clostridium beijerinckii GCA_000833105.2
209 Clostridium saccharoperbutylacetonicum N1-4_28HMT_29 GCA_000340885.1
195 Clostridium saccharobutylicum DSM 13864 GCA_000473995.1
194 Clostridium puniceum GCA_002006345.1
kS
198 Bifidobacterium adolescentis ATCC 15703 GCA_000010425.1
196 Bifidobacterium gallicum DSM 20093 = LMG 11596 GCA_000741205.1
190 Bifidobacterium dentium JCM 1195 = DSM 20436 GCA_001042595.1
kT
161 Enterococcus faecalis V583 GCA_000007785.1
161 Streptomyces cinnamoneus GCA_001885705.1
146 Enterococcus asini ATCC 700915 GCA_000407365.1
146 Enterococcus canis NBRC 100695 GCA_001544375.1
146 Enterococcus casseliflavus EC20 GCA_000157355.2
146 Enterococcus dispar ATCC 51266 GCA_000406945.1
146 Enterococcus faecium DO GCA_000174395.2
146 Enterococcus hirae ATCC 9790 GCA_000271405.2
146 Enterococcus massiliensis GCA_001050095.1
146 Enterococcus mundtii QU 25 GCA_000504125.1
146 Enterococcus rivorum GCA_001742285.1
146 Enterococcus saccharolyticus subsp. saccharolyticus ATCC 43076 GCA_000407285.1
135 Enterococcus gilvus ATCC BAA-350 GCA_000407545.1
135 Enterococcus hermanniensis GCA_001885945.1
135 Enterococcus malodoratus ATCC 43197 GCA_000407185.1
135 Enterococcus pallens ATCC BAA-351 GCA_000407485.1
135 Enterococcus pseudoavium NBRC 100491 GCA_001544295.1
135 Melissococcus plutonius S1 GCA_000747585.1
kU
295 Clostridium beijerinckii GCA_000833105.2
295 Clostridium saccharoperbutylacetonicum N1-4_28HMT_29 GCA_000340885.1
287 Clostridium saccharobutylicum DSM 13864 GCA_000473995.1
280 Clostridium puniceum GCA_002006345.1
kV
290 Clostridium saccharobutylicum DSM 13864 GCA_000473995.1
288 Clostridium butyricum GCA_001456065.2
279 Clostridium beijerinckii GCA_000833105.2
279 Clostridium puniceum GCA_002006345.1
279 Clostridium saccharoperbutylacetonicum N1-4_28HMT_29 GCA_000340885.1
kW
162 Deinococcus radiodurans R1 GCA_000008565.1
146 Deinococcus deserti VCD115 GCA_000020685.1
146 Deinococcus hopiensis KR-140 GCA_900176165.1
144 Deinococcus puniceus GCA_001644565.1
kX
293 Clostridium puniceum GCA_002006345.1
289 Clostridium beijerinckii GCA_000833105.2
289 Clostridium saccharobutylicum DSM 13864 GCA_000473995.1
289 Clostridium saccharoperbutylacetonicum N1-4_28HMT_29 GCA_000340885.1
274 Clostridium chromiireducens GCA_002029255.1
kY
251 Lactobacillus gasseri ATCC 33323 = JCM 1131 GCA_000014425.1
242 Lactobacillus hominis DSM 23910 = CRBIP 24.179 GCA_000296835.1
176 Lactobacillus psittaci DSM 15354 GCA_000425905.1
kZ
209 Clostridium beijerinckii GCA_000833105.2
209 Clostridium saccharobutylicum DSM 13864 GCA_000473995.1

209 Clostridium saccharoperbutylacetonicum N1-4_28HMT_29 GCA_000340885.1
205 Clostridium puniceum GCA_002006345.1
187 Clostridium chromiireducens GCA_002029255.1
l0
189 Clostridium taeniosporum GCA_001735765.1
181 Clostridium beijerinckii GCA_000833105.2
181 Clostridium butyricum GCA_001456065.2
181 Clostridium saccharobutylicum DSM 13864 GCA_000473995.1
181 Clostridium saccharoperbutylacetonicum N1-4_28HMT_29 GCA_000340885.1
177 Clostridium puniceum GCA_002006345.1
l1
241 Clostridium beijerinckii GCA_000833105.2
241 Clostridium puniceum GCA_002006345.1
241 Clostridium saccharoperbutylacetonicum N1-4_28HMT_29 GCA_000340885.1
239 Clostridium butyricum GCA_001456065.2
233 Clostridium chromiireducens GCA_002029255.1
233 Clostridium saccharobutylicum DSM 13864 GCA_000473995.1
l2
180 Clostridium beijerinckii GCA_000833105.2
180 Clostridium saccharobutylicum DSM 13864 GCA_000473995.1
180 Clostridium saccharoperbutylacetonicum N1-4_28HMT_29 GCA_000340885.1
174 Clostridium puniceum GCA_002006345.1
163 Clostridium chromiireducens GCA_002029255.1
l3
161 Bifidobacterium adolescentis ATCC 15703 GCA_000010425.1
148 Bifidobacterium thermophilum GCA_000741495.1
146 Bifidobacterium subtile GCA_000741775.1
l4
204 Streptococcus mutans UA159 GCA_000007465.2
143 Streptococcus ferus DSM 20646 GCA_000372425.1
136 Streptococcus ratti FA-1 = DSM 20564 GCA_000286075.1
l5
261 Lactobacillus gasseri ATCC 33323 = JCM 1131 GCA_000014425.1
241 Lactobacillus hominis DSM 23910 = CRBIP 24.179 GCA_000296835.1
169 Lactobacillus psittaci DSM 15354 GCA_000425905.1
l6
136 Rhodobacter capsulatus SB 1003 GCA_000021865.1
130 Rhodobacter sphaeroides 2.4.1 GCA_000012905.2
120 Gemmobacter aquatilis GCA_900110025.1
120 Gemmobacter nectariphilus DSM 15620 GCA_000429765.1
120 Haematobacter massiliensis GCA_000740795.1
l7
285 Lactobacillus gasseri ATCC 33323 = JCM 1131 GCA_000014425.1
285 Lactobacillus hominis DSM 23910 = CRBIP 24.179 GCA_000296835.1
225 Lactobacillus crispatus ST1 GCA_000091765.1
221 Lactobacillus iners DSM 13335 GCA_000160875.1
l8
187 Escherichia coli IAI39 GCA_000026345.1
187 Escherichia coli O104_3AH4 str. 2011C-3493 GCA_000299455.1
187 Escherichia coli O157_3AH7 str. Sakai GCA_000008865.1
187 Escherichia coli O83_3AH1 str. NRG 857C GCA_000183345.1
187 Escherichia coli UMN026 GCA_000026325.2
187 Escherichia coli str. K-12 substr. MG1655 GCA_000005845.2
187 Shigella dysenteriae Sd197 GCA_000012005.1
187 Shigella flexneri 2a str. 301 GCA_000006925.2
187 Tumebacillus flagellatus GCA_000714935.1
171 Erwinia iniecta GCA_001267535.1
169 Pantoea agglomerans GCA_001709315.1
l9
258 Bacillus anthracis str. Ames GCA_000007845.1
258 Bacillus anthracis str. Sterne GCA_000008165.1
258 Bacillus cereus ATCC 14579 GCA_000007825.1
258 Bacillus mycoides GCA_000832605.1
258 Bacillus pseudomycoides DSM 12442 GCA_000161455.1
258 Bacillus thuringiensis YBT-1518 GCA_000497525.2
258 _5BBacillus thuringiensis_5D serovar konkukian str. 97-27 GCA_000008505.1
218 Bacillus infantis NRRL B-14911 GCA_000473245.1
211 Bacillus vietnamensis NBRC 101237 GCA_001591825.1
la
241 Enterococcus faecalis V583 GCA_000007785.1
241 Streptomyces cinnamoneus GCA_001885705.1
217 Enterococcus canis NBRC 100695 GCA_001544375.1
217 Enterococcus dispar ATCC 51266 GCA_000406945.1
217 Enterococcus faecium DO GCA_000174395.2
217 Enterococcus gilvus ATCC BAA-350 GCA_000407545.1
217 Enterococcus haemoperoxidus ATCC BAA-382 GCA_000407165.1

217 Enterococcus hermanniensis GCA_001885945.1
217 Enterococcus hirae ATCC 9790 GCA_000271405.2
217 Enterococcus malodoratus ATCC 43197 GCA_000407185.1
217 Enterococcus mundtii QU 25 GCA_000504125.1
217 Enterococcus pallens ATCC BAA-351 GCA_000407485.1
217 Enterococcus phoeniculicola ATCC BAA-412 GCA_000407505.1
217 Enterococcus pseudoavium NBRC 100491 GCA_001544295.1
217 Enterococcus rivorum GCA_001742285.1
217 Enterococcus thailandicus GCA_001652875.1
207 Carnobacterium divergens DSM 20623 GCA_000744255.1
lb
184 Erwinia iniecta GCA_001267535.1
184 Escherichia coli IAI39 GCA_000026345.1
184 Escherichia coli O104_3AH4 str. 2011C-3493 GCA_000299455.1
184 Escherichia coli O157_3AH7 str. Sakai GCA_000008865.1
184 Escherichia coli O83_3AH1 str. NRG 857C GCA_000183345.1
184 Escherichia coli UMN026 GCA_000026325.2
184 Escherichia coli str. K-12 substr. MG1655 GCA_000005845.2
184 Shigella dysenteriae Sd197 GCA_000012005.1
184 Shigella flexneri 2a str. 301 GCA_000006925.2
184 Tumebacillus flagellatus GCA_000714935.1
169 Cronobacter sakazakii GCA_000982825.1
169 Enterobacter cloacae subsp. cloacae ATCC 13047 GCA_000025565.1
169 Enterobacter hormaechei subsp. steigerwaltii GCA_001729725.1
169 Enterobacter kobei GCA_900185885.1
169 Erwinia amylovora CFBP1430 GCA_000091565.1
169 Erwinia toletana DAPP-PG 735 GCA_000336255.1
169 Klebsiella oxytoca GCA_001022195.1
169 Kosakonia cowanii GCA_001975225.1
169 Kosakonia sacchari SP1 GCA_000300455.4
169 Pantoea agglomerans GCA_001709315.1
169 Pantoea alhagi GCA_002101395.1
169 Pantoea ananatis LMG 20103 GCA_000025405.2
169 Pantoea dispersa EGD-AAK13 GCA_000465555.2
169 Pantoea rwandensis GCA_000759475.1
169 Pantoea septica GCA_002095575.1
169 Pseudescherichia vulneris NBRC 102420 GCA_000759795.1
169 Salmonella enterica subsp. enterica serovar Typhi str. CT18 GCA_000195995.1
169 Shimwellia blattae DSM 4481 = NBRC 105725 GCA_000262305.1
169 Trabulsiella odontotermitis GCA_001297765.1
169 Xenorhabdus bovienii SS-2004 GCA_000027225.1
169 Xenorhabdus doucetiae GCA_000968195.1
169 Xenorhabdus hominickii GCA_001721185.1
169 Xenorhabdus innexi GCA_900155355.1
169 Xenorhabdus japonica GCA_900115195.1
169 Xenorhabdus koppenhoeferi GCA_900116635.1
169 Xenorhabdus nematophila AN6_2F1 GCA_000953355.1
168 Proteus mirabilis HI4320 GCA_000069965.1
168 Xenorhabdus cabanillasii JM26 GCA_000531755.1
lc
109 Staphylococcus epidermidis ATCC 12228 GCA_000007645.1
109 Staphylococcus pettenkoferi GCA_002208805.1
106 Staphylococcus haemolyticus JCSC1435 GCA_000009865.1
105 Staphylococcus cohnii subsp. cohnii GCA_000972575.1
ld
122 Bacillus thuringiensis YBT-1518 GCA_000497525.2
108 Bacillus anthracis str. Ames GCA_000007845.1
108 Bacillus anthracis str. Sterne GCA_000008165.1
108 Bacillus cereus ATCC 14579 GCA_000007825.1
108 Bacillus mycoides GCA_000832605.1
108 Bacillus pseudomycoides DSM 12442 GCA_000161455.1
108 _5BBacillus thuringiensis_5D serovar konkukian str. 97-27 GCA_000008505.1
93 Bacillus dakarensis GCA_900156875.1
93 Bacillus tuaregi GCA_900104575.1
93 Isobaculum melis GCA_900111355.1
le
139 Streptococcus mutans UA159 GCA_000007465.2
84 Streptococcus sobrinus DSM 20742 = ATCC 33478 GCA_000686605.1
81 Streptococcus ferus DSM 20646 GCA_000372425.1
lf
232 Clostridium beijerinckii GCA_000833105.2
232 Clostridium puniceum GCA_002006345.1
232 Clostridium saccharobutylicum DSM 13864 GCA_000473995.1
232 Clostridium saccharoperbutylacetonicum N1-4_28HMT_29 GCA_000340885.1
220 Clostridium neonatale GCA_001458595.1
201 Clostridium chromiireducens GCA_002029255.1

lg
175 Clostridium beijerinckii GCA_000833105.2
175 Clostridium puniceum GCA_002006345.1
175 Clostridium saccharoperbutylacetonicum N1-4_28HMT_29 GCA_000340885.1
166 Clostridium saccharobutylicum DSM 13864 GCA_000473995.1
155 Clostridium chromiireducens GCA_002029255.1
lh
122 Streptococcus mutans UA159 GCA_000007465.2
95 Bacillus azotoformans LMG 9581 GCA_000307855.1
91 Bacillus bataviensis LMG 21833 GCA_000307875.1
91 Lactococcus lactis subsp. lactis Il1403 GCA_000006865.1
91 Vagococcus penaei GCA_001998885.1
li
166 Rhodobacter sphaeroides 2.4.1 GCA_000012905.2
160 Rhodobacter capsulatus SB 1003 GCA_000021865.1
160 Rhodobacter sphaeroides ATCC 17025 GCA_000016405.1
146 Paracoccus sediminis GCA_900188295.1
lj
187 Clostridium beijerinckii GCA_000833105.2
187 Clostridium puniceum GCA_002006345.1
187 Clostridium saccharoperbutylacetonicum N1-4_28HMT_29 GCA_000340885.1
173 Clostridium saccharobutylicum DSM 13864 GCA_000473995.1
156 Clostridium taeniosporum GCA_001735765.1
lk
272 Enterococcus faecalis V583 GCA_000007785.1
272 Streptomyces cinnamoneus GCA_001885705.1
245 Enterococcus dispar ATCC 51266 GCA_000406945.1
242 Enterococcus canis NBRC 100695 GCA_001544375.1
242 Enterococcus casseliflavus EC20 GCA_000157355.2
242 Enterococcus faecium DO GCA_000174395.2
242 Enterococcus hirae ATCC 9790 GCA_000271405.2
242 Enterococcus mundtii QU 25 GCA_000504125.1
242 Enterococcus rivorum GCA_001742285.1
242 Enterococcus saccharolyticus subsp. saccharolyticus ATCC 43076 GCA_000407285.1
ll
213 Clostridium fallax GCA_900129365.1
212 Clostridium butyricum GCA_001456065.2
209 Clostridium beijerinckii GCA_000833105.2
209 Clostridium neonatale GCA_001458595.1
209 Clostridium puniceum GCA_002006345.1
209 Clostridium saccharobutylicum DSM 13864 GCA_000473995.1
209 Clostridium saccharoperbutylacetonicum N1-4_28HMT_29 GCA_000340885.1
lm
107 Clostridium beijerinckii GCA_000833105.2
107 Clostridium saccharobutylicum DSM 13864 GCA_000473995.1
107 Clostridium saccharoperbutylacetonicum N1-4_28HMT_29 GCA_000340885.1
100 Clostridium neonatale GCA_001458595.1
100 Clostridium puniceum GCA_002006345.1
85 Clostridium butyricum GCA_001456065.2
85 Clostridium chromiireducens GCA_002029255.1
ln
148 Clostridium beijerinckii GCA_000833105.2
148 Clostridium butyricum GCA_001456065.2
148 Clostridium chromiireducens GCA_002029255.1
148 Clostridium puniceum GCA_002006345.1
148 Clostridium saccharobutylicum DSM 13864 GCA_000473995.1
148 Clostridium saccharoperbutylacetonicum N1-4_28HMT_29 GCA_000340885.1
139 Clostridium cavendishii DSM 21758 GCA_900141845.1
122 Clostridium botulinum B str. Eklund 17B _28NRP_29 GCA_000020165.1
122 Clostridium taeniosporum GCA_001735765.1
122 Clostridium uliginosum GCA_900112485.1
lo
230 Bifidobacterium adolescentis ATCC 15703 GCA_000010425.1
217 Bifidobacterium thermophilum GCA_000741495.1
209 Bifidobacterium angulatum DSM 20098 = JCM 7096 GCA_001025155.1
lp
229 Clostridium beijerinckii GCA_000833105.2
229 Clostridium saccharobutylicum DSM 13864 GCA_000473995.1
229 Clostridium saccharoperbutylacetonicum N1-4_28HMT_29 GCA_000340885.1
219 Clostridium puniceum GCA_002006345.1
207 Clostridium chromiireducens GCA_002029255.1
lq
296 Staphylococcus capitis subsp. capitis GCA_001028645.1
296 Staphylococcus epidermidis ATCC 12228 GCA_000007645.1
296 Staphylococcus haemolyticus JCSC1435 GCA_000009865.1
282 Staphylococcus succinus GCA_001902315.1

281 Staphylococcus aureus subsp. aureus NCTC 8325 GCA_000013425.1
281 Staphylococcus condimenti GCA_001618885.1
281 Staphylococcus hominis subsp. hominis C80 GCA_000183685.1
281 Staphylococcus lugdunensis HKU09-01 GCA_000025085.1
281 Staphylococcus simiae CCM 7213 GCA_000235645.2
281 Staphylococcus simulans GCA_001559115.1
lr
204 Clostridium beijerinckii GCA_000833105.2
204 Clostridium puniceum GCA_002006345.1
204 Clostridium saccharobutylicum DSM 13864 GCA_000473995.1
204 Clostridium saccharoperbutylacetonicum N1-4_28HMT_29 GCA_000340885.1
188 Clostridium chromiireducens GCA_002029255.1
181 Clostridium butyricum GCA_001456065.2
ls
216 Escherichia coli IAI39 GCA_000026345.1
216 Escherichia coli O104_3AH4 str. 2011C-3493 GCA_000299455.1
216 Escherichia coli O157_3AH7 str. Sakai GCA_000008865.1
216 Escherichia coli O83_3AH1 str. NRG 857C GCA_000183345.1
216 Escherichia coli UMN026 GCA_000026325.2
216 Escherichia coli str. K-12 substr. MG1655 GCA_000005845.2
216 Shigella flexneri 2a str. 301 GCA_000006925.2
216 Tumebacillus flagellatus GCA_000714935.1
203 Erwinia iniecta GCA_001267535.1
201 Shigella dysenteriae Sd197 GCA_000012005.1
lt
315 Bacillus thuringiensis YBT-1518 GCA_000497525.2
307 Bacillus anthracis str. Ames GCA_000007845.1
307 Bacillus anthracis str. Sterne GCA_000008165.1
307 Bacillus cereus ATCC 14579 GCA_000007825.1
307 Bacillus mycoides GCA_000832605.1
307 Bacillus pseudomycoides DSM 12442 GCA_000161455.1
307 _5BBacillus thuringiensis_5D serovar konkukian str. 97-27 GCA_000008505.1
223 Bacillus vietnamensis NBRC 101237 GCA_001591825.1
lu
188 Megasphaera cerevisiae DSM 20462 GCA_001045675.1
188 Staphylococcus aureus subsp. aureus NCTC 8325 GCA_000013425.1
188 Staphylococcus auricularis GCA_001500315.1
188 Staphylococcus capitis subsp. capitis GCA_001028645.1
188 Staphylococcus epidermidis ATCC 12228 GCA_000007645.1
188 Staphylococcus haemolyticus JCSC1435 GCA_000009865.1
188 Staphylococcus hominis subsp. hominis C80 GCA_000183685.1
188 Staphylococcus lugdunensis HKU09-01 GCA_000025085.1
188 Staphylococcus lutrae GCA_002101335.1
188 Staphylococcus simiae CCM 7213 GCA_000235645.2
188 Staphylococcus warneri SG1 GCA_000332735.1
183 Staphylococcus condimenti GCA_001618885.1
183 Staphylococcus microti GCA_000934465.1
183 Staphylococcus pseudintermedius HKU10-03 GCA_000185885.1
183 Staphylococcus simulans GCA_001559115.1
180 Staphylococcus cohnii subsp. cohnii GCA_000972575.1
180 Staphylococcus gallinarum GCA_000875895.1
180 Staphylococcus saprophyticus subsp. saprophyticus ATCC 15305 GCA_000010125.1
180 Staphylococcus sciuri GCA_002209165.1
180 Staphylococcus succinus GCA_001902315.1
180 Staphylococcus xylosus GCA_000706685.1
lv
203 Deinococcus radiodurans R1 GCA_000008565.1
132 Deinococcus gobiensis I-0 GCA_000252445.1
124 Deinococcus proteolyticus MRP GCA_000190555.1
lw
203 Staphylococcus capitis subsp. capitis GCA_001028645.1
203 Staphylococcus epidermidis ATCC 12228 GCA_000007645.1
203 Staphylococcus haemolyticus JCSC1435 GCA_000009865.1
193 Staphylococcus pettenkoferi GCA_002208805.1
192 Staphylococcus arlettae CVD059 GCA_000295715.1
192 Staphylococcus cohnii subsp. cohnii GCA_000972575.1
192 Staphylococcus hominis subsp. hominis C80 GCA_000183685.1
192 Staphylococcus lugdunensis HKU09-01 GCA_000025085.1
192 Staphylococcus saprophyticus subsp. saprophyticus ATCC 15305 GCA_000010125.1
lx
248 Bacillus anthracis str. Ames GCA_000007845.1
248 Bacillus anthracis str. Sterne GCA_000008165.1
248 Bacillus cereus ATCC 14579 GCA_000007825.1
248 Bacillus mycoides GCA_000832605.1
248 Bacillus pseudomycoides DSM 12442 GCA_000161455.1
248 Bacillus thuringiensis YBT-1518 GCA_000497525.2

248 _5BBacillus thuringiensis_5D serovar konkukian str. 97-27 GCA_000008505.1
198 Oceanobacillus caeni GCA_001298135.1
192 Bacillus manliponensis GCA_000712595.1
ly
253 Rhodobacter sphaeroides 2.4.1 GCA_000012905.2
228 Defluviimonas alba GCA_001620265.1
226 Pseudorhodobacter psychrotolerans GCA_001294535.1
lz
185 Clostridium saccharobutylicum DSM 13864 GCA_000473995.1
183 Clostridium beijerinckii GCA_000833105.2
183 Clostridium saccharoperbutylacetonicum N1-4_28HMT_29 GCA_000340885.1
179 Clostridium puniceum GCA_002006345.1
lA
229 Clostridium beijerinckii GCA_000833105.2
229 Clostridium puniceum GCA_002006345.1
229 Clostridium saccharobutylicum DSM 13864 GCA_000473995.1
229 Clostridium saccharoperbutylacetonicum N1-4_28HMT_29 GCA_000340885.1
218 Clostridium chromiireducens GCA_002029255.1
209 Clostridium botulinum B str. Eklund 17B _28NRP_29 GCA_000020165.1
209 Clostridium taeniosporum GCA_001735765.1
lB
204 Clostridium beijerinckii GCA_000833105.2
204 Clostridium saccharobutylicum DSM 13864 GCA_000473995.1
204 Clostridium saccharoperbutylacetonicum N1-4_28HMT_29 GCA_000340885.1
190 Clostridium puniceum GCA_002006345.1
182 Clostridium chromiireducens GCA_002029255.1
lC
184 Clostridium beijerinckii GCA_000833105.2
184 Clostridium saccharobutylicum DSM 13864 GCA_000473995.1
184 Clostridium saccharoperbutylacetonicum N1-4_28HMT_29 GCA_000340885.1
178 Clostridium puniceum GCA_002006345.1
177 Clostridium butyricum GCA_001456065.2
lD
188 Clostridium beijerinckii GCA_000833105.2
188 Clostridium chromiireducens GCA_002029255.1
188 Clostridium puniceum GCA_002006345.1
188 Clostridium saccharobutylicum DSM 13864 GCA_000473995.1
188 Clostridium saccharoperbutylacetonicum N1-4_28HMT_29 GCA_000340885.1
183 Clostridium butyricum GCA_001456065.2
165 Clostridium taeniosporum GCA_001735765.1
lE
183 Lactobacillus gasseri ATCC 33323 = JCM 1131 GCA_000014425.1
170 Lactobacillus hominis DSM 23910 = CRBIP 24.179 GCA_000296835.1
163 Lactobacillus iners DSM 13335 GCA_000160875.1
lF
232 Bacillus thuringiensis YBT-1518 GCA_000497525.2
226 Bacillus anthracis str. Ames GCA_000007845.1
226 Bacillus anthracis str. Sterne GCA_000008165.1
226 Bacillus cereus ATCC 14579 GCA_000007825.1
226 Bacillus mycoides GCA_000832605.1
226 Bacillus pseudomycoides DSM 12442 GCA_000161455.1
226 _5BBacillus thuringiensis_5D serovar konkukian str. 97-27 GCA_000008505.1
220 Bacillus manliponensis GCA_000712595.1
lG
210 Clostridium beijerinckii GCA_000833105.2
210 Clostridium saccharobutylicum DSM 13864 GCA_000473995.1
210 Clostridium saccharoperbutylacetonicum N1-4_28HMT_29 GCA_000340885.1
204 Clostridium puniceum GCA_002006345.1
193 Clostridium chromiireducens GCA_002029255.1
lH
163 Rhodobacter sphaeroides 2.4.1 GCA_000012905.2
121 Haematobacter massiliensis GCA_000740795.1
121 Maribius pelagius GCA_900110115.1
121 Xuhuaishuia manganoxidans GCA_001981245.1
120 Rhodobacter capsulatus SB 1003 GCA_000021865.1
lI
121 Escherichia coli IAI39 GCA_000026345.1
121 Escherichia coli O104_3AH4 str. 2011C-3493 GCA_000299455.1
121 Escherichia coli O157_3AH7 str. Sakai GCA_000008865.1
121 Escherichia coli O83_3AH1 str. NRG 857C GCA_000183345.1
121 Escherichia coli UMN026 GCA_000026325.2
121 Escherichia coli str. K-12 substr. MG1655 GCA_000005845.2
121 Shigella flexneri 2a str. 301 GCA_000006925.2
121 Tumebacillus flagellatus GCA_000714935.1
120 Brenneria goodwinii GCA_001049335.1
117 Cronobacter sakazakii GCA_000982825.1

117 Erwinia amylovora CFBP1430 GCA_000091565.1
117 Erwinia iniecta GCA_001267535.1
117 Erwinia toletana DAPP-PG 735 GCA_000336255.1
117 Kosakonia cowanii GCA_001975225.1
117 Pantoea dispersa EGD-AAK13 GCA_000465555.2
117 Pluralibacter gergoviae GCA_000757785.1
117 Shimwellia blattae DSM 4481 = NBRC 105725 GCA_000262305.1
117 _5BEnterobacter_5D lignolyticus SCF1 GCA_000164865.1
lJ
225 Lactobacillus gasseri ATCC 33323 = JCM 1131 GCA_000014425.1
215 Lactobacillus hominis DSM 23910 = CRBIP 24.179 GCA_000296835.1
157 Lactobacillus iners DSM 13335 GCA_000160875.1
lK
156 Enterococcus asini ATCC 700915 GCA_000407365.1
156 Enterococcus canis NBRC 100695 GCA_001544375.1
156 Enterococcus dispar ATCC 51266 GCA_000406945.1
156 Enterococcus faecalis V583 GCA_000007785.1
156 Enterococcus faecium DO GCA_000174395.2
156 Enterococcus haemoperoxidus ATCC BAA-382 GCA_000407165.1
156 Enterococcus hirae ATCC 9790 GCA_000271405.2
156 Enterococcus mundtii QU 25 GCA_000504125.1
156 Enterococcus phoeniculicola ATCC BAA-412 GCA_000407505.1
156 Enterococcus rivorum GCA_001742285.1
156 Enterococcus thailandicus GCA_001652875.1
156 Melissococcus plutonius S1 GCA_000747585.1
156 Streptomyces cinnamoneus GCA_001885705.1
142 Enterococcus gilvus ATCC BAA-350 GCA_000407545.1
142 Enterococcus hermanniensis GCA_001885945.1
142 Enterococcus italicus DSM 15952 GCA_000185365.1
142 Enterococcus malodoratus ATCC 43197 GCA_000407185.1
142 Enterococcus pallens ATCC BAA-351 GCA_000407485.1
142 Enterococcus pseudoavium NBRC 100491 GCA_001544295.1
142 Enterococcus sulfureus ATCC 49903 GCA_000407605.1
142 Vagococcus lutrae LBD1 GCA_000498295.1
141 Enterococcus casseliflavus EC20 GCA_000157355.2
141 Enterococcus saccharolyticus subsp. saccharolyticus ATCC 43076 GCA_000407285.1
lL
205 Deinococcus radiodurans R1 GCA_000008565.1
118 Deinococcus marmoris DSM 12784 GCA_000701405.1
118 Deinococcus swuensis GCA_000800395.1
110 Deinococcus frigens DSM 12807 GCA_000701425.1
lM
231 Bacillus thuringiensis YBT-1518 GCA_000497525.2
223 Bacillus anthracis str. Ames GCA_000007845.1
223 Bacillus anthracis str. Sterne GCA_000008165.1
223 Bacillus mycoides GCA_000832605.1
223 Bacillus pseudomycoides DSM 12442 GCA_000161455.1
223 _5BBacillus thuringiensis_5D serovar konkukian str. 97-27 GCA_000008505.1
208 Bacillus cereus ATCC 14579 GCA_000007825.1
lN
297 Staphylococcus epidermidis ATCC 12228 GCA_000007645.1
292 Staphylococcus capitis subsp. capitis GCA_001028645.1
286 Staphylococcus haemolyticus JCSC1435 GCA_000009865.1
lO
194 Rhodobacter sphaeroides 2.4.1 GCA_000012905.2
178 Pseudorhodobacter ferrugineus DSM 5888 GCA_000420745.1
178 Pseudorhodobacter wandonensis GCA_001202035.1
175 Gemmobacter aquatilis GCA_900110025.1
175 Gemmobacter megaterium GCA_900156815.1
lP
153 Deinococcus radiodurans R1 GCA_000008565.1
77 Deinococcus deserti VCD115 GCA_000020685.1
75 Deinococcus soli Cha et al. 2016 GCA_001007995.1
lQ
140 Enterobacter cloacae subsp. cloacae ATCC 13047 GCA_000025565.1
125 Buttiauxella ferragutiae ATCC 51602 GCA_001654915.1
125 Cedecea neteri GCA_000757825.1
125 Citrobacter freundii CFNIH1 GCA_000648515.1
125 Cronobacter sakazakii GCA_000982825.1
125 Edwardsiella anguillarum ET080813 GCA_000264765.2
125 Enterobacter hormaechei subsp. steigerwaltii GCA_001729725.1
125 Enterobacter kobei GCA_900185885.1
125 Erwinia amylovora CFBP1430 GCA_000091565.1
125 Erwinia billingiae Eb661 GCA_000196615.1
125 Erwinia gerundensis GCA_001517405.1
125 Erwinia iniecta GCA_001267535.1

125 Erwinia persicina NBRC 102418 GCA_001571305.1
125 Erwinia toletana DAPP-PG 735 GCA_000336255.1
125 Escherichia coli IAI39 GCA_000026345.1
125 Escherichia coli O104_3AH4 str. 2011C-3493 GCA_000299455.1
125 Escherichia coli O157_3AH7 str. Sakai GCA_000008865.1
125 Escherichia coli O83_3AH1 str. NRG 857C GCA_000183345.1
125 Escherichia coli UMN026 GCA_000026325.2
125 Escherichia coli str. K-12 substr. MG1655 GCA_000005845.2
125 Klebsiella aerogenes KCTC 2190 GCA_000215745.1
125 Klebsiella oxytoca GCA_001022195.1
125 Kluyvera ascorbata ATCC 33433 GCA_000735365.1
125 Kosakonia cowanii GCA_001975225.1
125 Kosakonia sacchari SP1 GCA_000300455.4
125 Mangrovibacter phragmitis GCA_001655675.1
125 Obesumbacterium proteus GCA_001586165.1
125 Pantoea agglomerans GCA_001709315.1
125 Pantoea alhagi GCA_002101395.1
125 Pantoea ananatis LMG 20103 GCA_000025405.2
125 Pantoea dispersa EGD-AAK13 GCA_000465555.2
125 Pantoea rwandensis GCA_000759475.1
125 Pantoea septica GCA_002095575.1
125 Plautia stali symbiont GCA_000180175.2
125 Pluralibacter gergoviae GCA_000757785.1
125 Pseudescherichia vulneris NBRC 102420 GCA_000759795.1
125 Rosenbergiella nectarea GCA_900111105.1
125 Salmonella enterica subsp. enterica serovar Typhi str. CT18 GCA_000195995.1
125 Shigella dysenteriae Sd197 GCA_000012005.1
125 Shigella flexneri 2a str. 301 GCA_000006925.2
125 Shimwellia blattae DSM 4481 = NBRC 105725 GCA_000262305.1
125 Tatumella citrea GCA_002163585.1
125 Tatumella saanichensis GCA_000439375.1
125 Trabulsiella odontotermitis GCA_001297765.1
125 Tumebacillus flagellatus GCA_000714935.1
125 Yokenella regensburgei ATCC 49455 GCA_000735455.1
125 _5BEnterobacter_5D lignolyticus SCF1 GCA_000164865.1
121 Morganella morganii subsp. morganii KT GCA_000286435.2
121 Morganella psychrotolerans GCA_001676155.1
121 Proteus mirabilis HI4320 GCA_000069965.1
121 Providencia alcalifaciens Dmel2 GCA_000314875.2
121 Providencia sneebia DSM 19967 GCA_000314895.2
121 Providencia stuartii MRSN 2154 GCA_000259175.1
121 Xenorhabdus bovienii SS-2004 GCA_000027225.1
121 Xenorhabdus doucetiae GCA_000968195.1
121 Xenorhabdus hominickii GCA_001721185.1
121 Xenorhabdus innexi GCA_900155355.1
121 Xenorhabdus japonica GCA_900115195.1
121 Xenorhabdus koppenhoeferi GCA_900116635.1
121 Xenorhabdus nematophila AN6_2F1 GCA_000953355.1
lR
279 Escherichia coli IAI39 GCA_000026345.1
279 Escherichia coli O104_3AH4 str. 2011C-3493 GCA_000299455.1
279 Escherichia coli O157_3AH7 str. Sakai GCA_000008865.1
279 Escherichia coli O83_3AH1 str. NRG 857C GCA_000183345.1
279 Escherichia coli UMN026 GCA_000026325.2
279 Escherichia coli str. K-12 substr. MG1655 GCA_000005845.2
279 Shigella dysenteriae Sd197 GCA_000012005.1
279 Shigella flexneri 2a str. 301 GCA_000006925.2
279 Tumebacillus flagellatus GCA_000714935.1
256 Erwinia iniecta GCA_001267535.1
256 Erwinia toletana DAPP-PG 735 GCA_000336255.1
256 Kosakonia cowanii GCA_001975225.1
255 Pluralibacter gergoviae GCA_000757785.1
lS
230 Clostridium beijerinckii GCA_000833105.2
230 Clostridium puniceum GCA_002006345.1
230 Clostridium saccharoperbutylacetonicum N1-4_28HMT_29 GCA_000340885.1
226 Clostridium saccharobutylicum DSM 13864 GCA_000473995.1
218 Clostridium chromiireducens GCA_002029255.1
lT
88 Escherichia coli IAI39 GCA_000026345.1
88 Escherichia coli O104_3AH4 str. 2011C-3493 GCA_000299455.1
88 Escherichia coli O157_3AH7 str. Sakai GCA_000008865.1
88 Escherichia coli O83_3AH1 str. NRG 857C GCA_000183345.1
88 Escherichia coli UMN026 GCA_000026325.2
88 Escherichia coli str. K-12 substr. MG1655 GCA_000005845.2
88 Shigella dysenteriae Sd197 GCA_000012005.1

88 Shigella flexneri 2a str. 301 GCA_000006925.2
88 Tumebacillus flagellatus GCA_000714935.1
77 Vibrio coralliilyticus GCA_000763535.2
66 Buchnera aphidicola str. G002 _28Myzus persicae_29 GCA_000521565.1
66 Buchnera aphidicola str. Ua _28Uroleucon ambrosiae_29 GCA_000225465.1
lU
280 Bacillus thuringiensis YBT-1518 GCA_000497525.2
272 Bacillus anthracis str. Ames GCA_000007845.1
272 Bacillus anthracis str. Sterne GCA_000008165.1
272 Bacillus cereus ATCC 14579 GCA_000007825.1
272 Bacillus mycoides GCA_000832605.1
272 Bacillus pseudomycoides DSM 12442 GCA_000161455.1
272 _5BBacillus thuringiensis_5D serovar konkukian str. 97-27 GCA_000008505.1
216 Bacillus manliponensis GCA_000712595.1
lV
294 Lactobacillus gasseri ATCC 33323 = JCM 1131 GCA_000014425.1
285 Lactobacillus hominis DSM 23910 = CRBIP 24.179 GCA_000296835.1
214 Lactobacillus iners DSM 13335 GCA_000160875.1
lW
229 Bacillus anthracis str. Ames GCA_000007845.1
229 Bacillus anthracis str. Sterne GCA_000008165.1
229 Bacillus cereus ATCC 14579 GCA_000007825.1
229 Bacillus mycoides GCA_000832605.1
229 Bacillus pseudomycoides DSM 12442 GCA_000161455.1
229 Bacillus thuringiensis YBT-1518 GCA_000497525.2
229 _5BBacillus thuringiensis_5D serovar konkukian str. 97-27 GCA_000008505.1
177 Bacillus manliponensis GCA_000712595.1
170 Bacillus infantis NRRL B-14911 GCA_000473245.1
170 Planococcus antarcticus DSM 14505 GCA_001687565.2
170 Planococcus kocurii GCA_001465835.2
170 Planococcus maritimus GCA_001999905.1
170 Planococcus rifietoensis GCA_001465795.2
lX
300 Deinococcus radiodurans R1 GCA_000008565.1
177 Deinococcus marmoris DSM 12784 GCA_000701405.1
177 Deinococcus swuensis GCA_000800395.1
169 Deinococcus gobiensis I-0 GCA_000252445.1
lY
256 Bifidobacterium adolescentis ATCC 15703 GCA_000010425.1
227 Bifidobacterium asteroides PRL2011 GCA_000304215.1
221 Bifidobacterium callitrichos DSM 23973 GCA_000741175.1
lZ
227 Rhodobacter sphaeroides ATCC 17025 GCA_000016405.1
212 Pseudorhodobacter ferrugineus DSM 5888 GCA_000420745.1
212 Pseudorhodobacter wandonensis GCA_001202035.1
212 Rhodobacter sphaeroides 2.4.1 GCA_000012905.2
198 Pseudorhodobacter psychrotolerans GCA_001294535.1
m0
217 Clostridium saccharobutylicum DSM 13864 GCA_000473995.1
215 Clostridium beijerinckii GCA_000833105.2
215 Clostridium saccharoperbutylacetonicum N1-4_28HMT_29 GCA_000340885.1
213 Clostridium chromiireducens GCA_002029255.1
m1
183 Deinococcus radiodurans R1 GCA_000008565.1
89 Deinococcus puniceus GCA_001644565.1
81 Deinococcus deserti VCD115 GCA_000020685.1
81 Deinococcus gobiensis I-0 GCA_000252445.1
m2
239 Enterococcus faecalis V583 GCA_000007785.1
239 Streptomyces cinnamoneus GCA_001885705.1
215 Carnobacterium divergens DSM 20623 GCA_000744255.1
205 Enterococcus asini ATCC 700915 GCA_000407365.1
205 Enterococcus canis NBRC 100695 GCA_001544375.1
205 Enterococcus dispar ATCC 51266 GCA_000406945.1
205 Enterococcus faecium DO GCA_000174395.2
205 Enterococcus gilvus ATCC BAA-350 GCA_000407545.1
205 Enterococcus haemoperoxidus ATCC BAA-382 GCA_000407165.1
205 Enterococcus hermanniensis GCA_001885945.1
205 Enterococcus hirae ATCC 9790 GCA_000271405.2
205 Enterococcus italicus DSM 15952 GCA_000185365.1
205 Enterococcus malodoratus ATCC 43197 GCA_000407185.1
205 Enterococcus mundtii QU 25 GCA_000504125.1
205 Enterococcus pallens ATCC BAA-351 GCA_000407485.1
205 Enterococcus phoeniculicola ATCC BAA-412 GCA_000407505.1
205 Enterococcus pseudoavium NBRC 100491 GCA_001544295.1
205 Enterococcus rivorum GCA_001742285.1

205 Enterococcus sulfureus ATCC 49903 GCA_000407605.1
205 Enterococcus thailandicus GCA_001652875.1
205 Melissococcus plutonius S1 GCA_000747585.1
m3
186 Streptococcus mutans UA159 GCA_000007465.2
131 Streptococcus iniae GCA_000831485.1
125 Streptococcus ferus DSM 20646 GCA_000372425.1
m4
223 Deinococcus radiodurans R1 GCA_000008565.1
166 Deinococcus gobiensis I-0 GCA_000252445.1
163 Deinococcus puniceus GCA_001644565.1
m5
272 Streptococcus mutans UA159 GCA_000007465.2
183 Streptococcus sobrinus DSM 20742 = ATCC 33478 GCA_000686605.1
180 Streptococcus macacae NCTC 11558 GCA_000187995.3
m6
224 Clostridium beijerinckii GCA_000833105.2
224 Clostridium puniceum GCA_002006345.1
224 Clostridium saccharoperbutylacetonicum N1-4_28HMT_29 GCA_000340885.1
220 Clostridium saccharobutylicum DSM 13864 GCA_000473995.1
215 Clostridium butyricum GCA_001456065.2
m7
190 Clostridium beijerinckii GCA_000833105.2
190 Clostridium saccharoperbutylacetonicum N1-4_28HMT_29 GCA_000340885.1
182 Clostridium saccharobutylicum DSM 13864 GCA_000473995.1
175 Clostridium puniceum GCA_002006345.1
m8
228 Rhodobacter sphaeroides 2.4.1 GCA_000012905.2
214 Pseudorhodobacter ferrugineus DSM 5888 GCA_000420745.1
214 Pseudorhodobacter wandonensis GCA_001202035.1
212 Pseudorhodobacter psychrotolerans GCA_001294535.1
m9
257 Clostridium beijerinckii GCA_000833105.2
257 Clostridium saccharobutylicum DSM 13864 GCA_000473995.1
257 Clostridium saccharoperbutylacetonicum N1-4_28HMT_29 GCA_000340885.1
250 Clostridium puniceum GCA_002006345.1
229 Clostridium butyricum GCA_001456065.2
ma
157 Erwinia iniecta GCA_001267535.1
157 Escherichia coli IAI39 GCA_000026345.1
157 Escherichia coli O104_3AH4 str. 2011C-3493 GCA_000299455.1
157 Escherichia coli O157_3AH7 str. Sakai GCA_000008865.1
157 Escherichia coli O83_3AH1 str. NRG 857C GCA_000183345.1
157 Escherichia coli UMN026 GCA_000026325.2
157 Escherichia coli str. K-12 substr. MG1655 GCA_000005845.2
157 Shigella flexneri 2a str. 301 GCA_000006925.2
157 Tumebacillus flagellatus GCA_000714935.1
151 Erwinia toletana DAPP-PG 735 GCA_000336255.1
147 Cronobacter sakazakii GCA_000982825.1
147 Pantoea ananatis LMG 20103 GCA_000025405.2
147 Shimwellia blattae DSM 4481 = NBRC 105725 GCA_000262305.1
mb
242 Streptococcus mutans UA159 GCA_000007465.2
168 Streptococcus ferus DSM 20646 GCA_000372425.1
168 Streptococcus massiliensis DSM 18628 GCA_000380065.1
161 Streptococcus sobrinus DSM 20742 = ATCC 33478 GCA_000686605.1
mc
188 Clostridium beijerinckii GCA_000833105.2
188 Clostridium saccharobutylicum DSM 13864 GCA_000473995.1
188 Clostridium saccharoperbutylacetonicum N1-4_28HMT_29 GCA_000340885.1
179 Clostridium algidicarnis GCA_000703125.1
174 Clostridium cylindrosporum DSM 605 GCA_001047375.1
md
293 Lactobacillus gasseri ATCC 33323 = JCM 1131 GCA_000014425.1
285 Lactobacillus hominis DSM 23910 = CRBIP 24.179 GCA_000296835.1
220 Lactobacillus iners DSM 13335 GCA_000160875.1
me
228 Clostridium beijerinckii GCA_000833105.2
228 Clostridium saccharobutylicum DSM 13864 GCA_000473995.1
228 Clostridium saccharoperbutylacetonicum N1-4_28HMT_29 GCA_000340885.1
222 Clostridium puniceum GCA_002006345.1
207 Clostridium chromiireducens GCA_002029255.1
mf
150 Bifidobacterium adolescentis ATCC 15703 GCA_000010425.1
130 Bifidobacterium choerinum GCA_000741135.1
130 Bifidobacterium pseudolongum PV8-2 GCA_000800475.2

126 Bifidobacterium longum NCC2705 GCA_000007525.1
mg
120 Clostridium beijerinckii GCA_000833105.2
120 Clostridium puniceum GCA_002006345.1
120 Clostridium saccharoperbutylacetonicum N1-4_28HMT_29 GCA_000340885.1
111 Clostridium saccharobutylicum DSM 13864 GCA_000473995.1
109 Clostridium butyricum GCA_001456065.2
109 Clostridium taeniosporum GCA_001735765.1
mh
305 Escherichia coli IAI39 GCA_000026345.1
305 Escherichia coli O104_3AH4 str. 2011C-3493 GCA_000299455.1
305 Escherichia coli O157_3AH7 str. Sakai GCA_000008865.1
305 Escherichia coli UMN026 GCA_000026325.2
305 Escherichia coli str. K-12 substr. MG1655 GCA_000005845.2
305 Shigella dysenteriae Sd197 GCA_000012005.1
305 Shigella flexneri 2a str. 301 GCA_000006925.2
305 Tumebacillus flagellatus GCA_000714935.1
301 Escherichia coli O83_3AH1 str. NRG 857C GCA_000183345.1
277 Erwinia iniecta GCA_001267535.1
mi
147 Deinococcus radiodurans R1 GCA_000008565.1
92 Deinococcus gobiensis I-0 GCA_000252445.1
89 Deinococcus soli Cha et al. 2016 GCA_001007995.1
mj
211 Deinococcus radiodurans R1 GCA_000008565.1
160 Deinococcus gobiensis I-0 GCA_000252445.1
155 Deinococcus marmoris DSM 12784 GCA_000701405.1
155 Deinococcus swuensis GCA_000800395.1
mk
111 Clostridium beijerinckii GCA_000833105.2
111 Clostridium chromiireducens GCA_002029255.1
111 Clostridium puniceum GCA_002006345.1
111 Clostridium saccharobutylicum DSM 13864 GCA_000473995.1
111 Clostridium saccharoperbutylacetonicum N1-4_28HMT_29 GCA_000340885.1
101 Clostridium butyricum GCA_001456065.2
91 Clostridium cylindrosporum DSM 605 GCA_001047375.1
91 Clostridium taeniosporum GCA_001735765.1
91 Fervidicella metallireducens AeB GCA_000601455.1
ml
262 Bacillus anthracis str. Ames GCA_000007845.1
262 Bacillus anthracis str. Sterne GCA_000008165.1
262 Bacillus cereus ATCC 14579 GCA_000007825.1
262 Bacillus mycoides GCA_000832605.1
262 Bacillus pseudomycoides DSM 12442 GCA_000161455.1
262 Bacillus thuringiensis YBT-1518 GCA_000497525.2
262 _5BBacillus thuringiensis_5D serovar konkukian str. 97-27 GCA_000008505.1
194 Bacillus cytotoxicus NVH 391-98 GCA_000017425.1
191 Bacillus tuaregi GCA_900104575.1
mm
188 Rhodobacter sphaeroides 2.4.1 GCA_000012905.2
179 Rhodobacter sphaeroides ATCC 17025 GCA_000016405.1
169 Gemmobacter nectariphilus DSM 15620 GCA_000429765.1
169 Haematobacter massiliensis GCA_000740795.1
169 Pseudorhodobacter psychrotolerans GCA_001294535.1
169 Rhodobacter capsulatus SB 1003 GCA_000021865.1
mn
187 Rhodobacter sphaeroides 2.4.1 GCA_000012905.2
168 Gemmobacter megaterium GCA_900156815.1
159 Gemmobacter aquatilis GCA_900110025.1
mo
254 Staphylococcus aureus subsp. aureus NCTC 8325 GCA_000013425.1
254 Staphylococcus capitis subsp. capitis GCA_001028645.1
254 Staphylococcus epidermidis ATCC 12228 GCA_000007645.1
254 Staphylococcus haemolyticus JCSC1435 GCA_000009865.1
254 Staphylococcus hominis subsp. hominis C80 GCA_000183685.1
254 Staphylococcus lugdunensis HKU09-01 GCA_000025085.1
254 Staphylococcus simiae CCM 7213 GCA_000235645.2
246 Staphylococcus condimenti GCA_001618885.1
246 Staphylococcus simulans GCA_001559115.1
244 Megasphaera cerevisiae DSM 20462 GCA_001045675.1
244 Staphylococcus warneri SG1 GCA_000332735.1
mp
225 Rhodobacter sphaeroides 2.4.1 GCA_000012905.2
203 Gemmobacter megaterium GCA_900156815.1
198 Rhodobacter sphaeroides ATCC 17025 GCA_000016405.1
mq

273 Streptococcus mutans UA159 GCA_000007465.2
166 Streptococcus ferus DSM 20646 GCA_000372425.1
164 Streptococcus sobrinus DSM 20742 = ATCC 33478 GCA_000686605.1
mr
260 Lactobacillus gasseri ATCC 33323 = JCM 1131 GCA_000014425.1
222 Lactobacillus hominis DSM 23910 = CRBIP 24.179 GCA_000296835.1
173 Lactobacillus amylophilus DSM 20533 = JCM 1125 GCA_001936335.1
173 Lactobacillus jensenii GCA_001936235.1
ms
215 Streptococcus mutans UA159 GCA_000007465.2
182 Streptococcus sobrinus DSM 20742 = ATCC 33478 GCA_000686605.1
169 Streptococcus gordonii str. Challis substr. CH1 GCA_000017005.1
mt
234 Bacillus thuringiensis YBT-1518 GCA_000497525.2
228 Bacillus anthracis str. Ames GCA_000007845.1
228 Bacillus anthracis str. Sterne GCA_000008165.1
228 Bacillus cereus ATCC 14579 GCA_000007825.1
228 Bacillus mycoides GCA_000832605.1
228 Bacillus pseudomycoides DSM 12442 GCA_000161455.1
228 _5BBacillus thuringiensis_5D serovar konkukian str. 97-27 GCA_000008505.1
198 Bacillus manliponensis GCA_000712595.1
mu
312 Clostridium beijerinckii GCA_000833105.2
312 Clostridium saccharobutylicum DSM 13864 GCA_000473995.1
312 Clostridium saccharoperbutylacetonicum N1-4_28HMT_29 GCA_000340885.1
305 Clostridium puniceum GCA_002006345.1
290 Clostridium chromiireducens GCA_002029255.1
mv
198 Escherichia coli IAI39 GCA_000026345.1
198 Escherichia coli O104_3AH4 str. 2011C-3493 GCA_000299455.1
198 Escherichia coli O157_3AH7 str. Sakai GCA_000008865.1
198 Escherichia coli O83_3AH1 str. NRG 857C GCA_000183345.1
198 Escherichia coli UMN026 GCA_000026325.2
198 Escherichia coli str. K-12 substr. MG1655 GCA_000005845.2
198 Shigella dysenteriae Sd197 GCA_000012005.1
198 Tumebacillus flagellatus GCA_000714935.1
190 Shigella flexneri 2a str. 301 GCA_000006925.2
175 Cronobacter sakazakii GCA_000982825.1
175 Erwinia amylovora CFBP1430 GCA_000091565.1
175 Erwinia iniecta GCA_001267535.1
175 Erwinia toletana DAPP-PG 735 GCA_000336255.1
175 Kosakonia cowanii GCA_001975225.1
175 Pantoea ananatis LMG 20103 GCA_000025405.2
175 Pantoea dispersa EGD-AAK13 GCA_000465555.2
175 Photorhabdus temperata subsp. thracensis GCA_001010285.1
175 Pluralibacter gergoviae GCA_000757785.1
175 Shimwellia blattae DSM 4481 = NBRC 105725 GCA_000262305.1
175 Xenorhabdus bovienii SS-2004 GCA_000027225.1
175 Xenorhabdus cabanillasii JM26 GCA_000531755.1
175 Xenorhabdus doucetiae GCA_000968195.1
175 Xenorhabdus hominickii GCA_001721185.1
175 Xenorhabdus innexi GCA_900155355.1
175 Xenorhabdus japonica GCA_900115195.1
175 Xenorhabdus koppenhoeferi GCA_900116635.1
175 Xenorhabdus nematophila AN6_2F1 GCA_000953355.1
mw
111 Deinococcus radiodurans R1 GCA_000008565.1
64 Deinococcus hopiensis KR-140 GCA_900176165.1
55 Deinococcus misasensis DSM 22328 GCA_000745915.1
mx
247 Clostridium beijerinckii GCA_000833105.2
247 Clostridium saccharobutylicum DSM 13864 GCA_000473995.1
247 Clostridium saccharoperbutylacetonicum N1-4_28HMT_29 GCA_000340885.1
241 Clostridium puniceum GCA_002006345.1
216 Clostridium chromiireducens GCA_002029255.1
my
244 Bifidobacterium adolescentis ATCC 15703 GCA_000010425.1
232 Bifidobacterium longum NCC2705 GCA_000007525.1
232 Bifidobacterium saguini DSM 23967 GCA_000741715.1
231 Bifidobacterium breve DSM 20213 = JCM 1192 GCA_001025175.1
mz
295 Enterococcus faecalis V583 GCA_000007785.1
295 Streptomyces cinnamoneus GCA_001885705.1
277 Enterococcus asini ATCC 700915 GCA_000407365.1
277 Enterococcus canis NBRC 100695 GCA_001544375.1
277 Enterococcus casseliflavus EC20 GCA_000157355.2

277 Enterococcus dispar ATCC 51266 GCA_000406945.1
277 Enterococcus faecium DO GCA_000174395.2
277 Enterococcus haemoperoxidus ATCC BAA-382 GCA_000407165.1
277 Enterococcus hirae ATCC 9790 GCA_000271405.2
277 Enterococcus massiliensis GCA_001050095.1
277 Enterococcus mundtii QU 25 GCA_000504125.1
277 Enterococcus phoeniculicola ATCC BAA-412 GCA_000407505.1
277 Enterococcus rivorum GCA_001742285.1
277 Enterococcus saccharolyticus subsp. saccharolyticus ATCC 43076 GCA_000407285.1
277 Enterococcus thailandicus GCA_001652875.1
268 Enterococcus cecorum GCA_001318405.1
mA
210 Clostridium beijerinckii GCA_000833105.2
210 Clostridium puniceum GCA_002006345.1
210 Clostridium saccharobutylicum DSM 13864 GCA_000473995.1
210 Clostridium saccharoperbutylacetonicum N1-4_28HMT_29 GCA_000340885.1
209 Clostridium butyricum GCA_001456065.2
202 Clostridium chromiireducens GCA_002029255.1
mB
301 Escherichia coli IAI39 GCA_000026345.1
301 Escherichia coli O104_3AH4 str. 2011C-3493 GCA_000299455.1
301 Escherichia coli O157_3AH7 str. Sakai GCA_000008865.1
301 Escherichia coli O83_3AH1 str. NRG 857C GCA_000183345.1
301 Escherichia coli UMN026 GCA_000026325.2
301 Escherichia coli str. K-12 substr. MG1655 GCA_000005845.2
301 Shigella flexneri 2a str. 301 GCA_000006925.2
301 Tumebacillus flagellatus GCA_000714935.1
292 Shigella dysenteriae Sd197 GCA_000012005.1
281 Erwinia iniecta GCA_001267535.1
mC
204 Deinococcus radiodurans R1 GCA_000008565.1
114 Deinococcus gobiensis I-0 GCA_000252445.1
107 Deinococcus proteolyticus MRP GCA_000190555.1
mD
138 Deinococcus radiodurans R1 GCA_000008565.1
110 Deinococcus marmoris DSM 12784 GCA_000701405.1
110 Deinococcus swuensis GCA_000800395.1
109 Deinococcus gobiensis I-0 GCA_000252445.1
mE
176 Deinococcus radiodurans R1 GCA_000008565.1
109 Deinococcus deserti VCD115 GCA_000020685.1
107 Deinococcus puniceus GCA_001644565.1
mF
146 Deinococcus radiodurans R1 GCA_000008565.1
122 Deinococcus puniceus GCA_001644565.1
105 Deinococcus deserti VCD115 GCA_000020685.1
mG
254 Deinococcus radiodurans R1 GCA_000008565.1
174 Deinococcus gobiensis I-0 GCA_000252445.1
168 Deinococcus frigens DSM 12807 GCA_000701425.1
168 Deinococcus marmoris DSM 12784 GCA_000701405.1
168 Deinococcus swuensis GCA_000800395.1
mH
312 Lactobacillus gasseri ATCC 33323 = JCM 1131 GCA_000014425.1
302 Lactobacillus hominis DSM 23910 = CRBIP 24.179 GCA_000296835.1
214 Lactobacillus iners DSM 13335 GCA_000160875.1
mI
239 Deinococcus radiodurans R1 GCA_000008565.1
178 Deinococcus deserti VCD115 GCA_000020685.1
175 Deinococcus phoenicis GCA_000599865.1
mJ
212 Escherichia coli IAI39 GCA_000026345.1
212 Escherichia coli O104_3AH4 str. 2011C-3493 GCA_000299455.1
212 Escherichia coli O157_3AH7 str. Sakai GCA_000008865.1
212 Escherichia coli O83_3AH1 str. NRG 857C GCA_000183345.1
212 Escherichia coli UMN026 GCA_000026325.2
212 Escherichia coli str. K-12 substr. MG1655 GCA_000005845.2
212 Shigella flexneri 2a str. 301 GCA_000006925.2
212 Tumebacillus flagellatus GCA_000714935.1
197 Shigella dysenteriae Sd197 GCA_000012005.1
173 Erwinia iniecta GCA_001267535.1
173 Erwinia toletana DAPP-PG 735 GCA_000336255.1
173 Kosakonia cowanii GCA_001975225.1
173 Pantoea ananatis LMG 20103 GCA_000025405.2
173 Shimwellia blattae DSM 4481 = NBRC 105725 GCA_000262305.1
mK

279 Vagococcus lutrae LBD1 GCA_000498295.1
274 Enterococcus hirae ATCC 9790 GCA_000271405.2
271 Enterococcus asini ATCC 700915 GCA_000407365.1
271 Enterococcus canis NBRC 100695 GCA_001544375.1
271 Enterococcus dispar ATCC 51266 GCA_000406945.1
271 Enterococcus faecalis V583 GCA_000007785.1
271 Enterococcus faecium DO GCA_000174395.2
271 Enterococcus gilvus ATCC BAA-350 GCA_000407545.1
271 Enterococcus hermanniensis GCA_001885945.1
271 Enterococcus malodoratus ATCC 43197 GCA_000407185.1
271 Enterococcus mundtii QU 25 GCA_000504125.1
271 Enterococcus pallens ATCC BAA-351 GCA_000407485.1
271 Enterococcus pseudoavium NBRC 100491 GCA_001544295.1
271 Enterococcus rivorum GCA_001742285.1
271 Streptomyces cinnamoneus GCA_001885705.1
mL
273 Lactobacillus gasseri ATCC 33323 = JCM 1131 GCA_000014425.1
265 Lactobacillus hominis DSM 23910 = CRBIP 24.179 GCA_000296835.1
241 Lactobacillus iners DSM 13335 GCA_000160875.1
mM
235 Rhodobacter sphaeroides 2.4.1 GCA_000012905.2
213 Pseudorhodobacter psychrotolerans GCA_001294535.1
199 Gemmobacter aquatilis GCA_900110025.1
mN
109 Erwinia iniecta GCA_001267535.1
109 Escherichia coli IAI39 GCA_000026345.1
109 Escherichia coli O104_3AH4 str. 2011C-3493 GCA_000299455.1
109 Escherichia coli O157_3AH7 str. Sakai GCA_000008865.1
109 Escherichia coli O83_3AH1 str. NRG 857C GCA_000183345.1
109 Escherichia coli UMN026 GCA_000026325.2
109 Escherichia coli str. K-12 substr. MG1655 GCA_000005845.2
109 Shigella dysenteriae Sd197 GCA_000012005.1
109 Shigella flexneri 2a str. 301 GCA_000006925.2
109 Tumebacillus flagellatus GCA_000714935.1
98 Xenorhabdus bovienii SS-2004 GCA_000027225.1
94 Rosenbergiella nectarea GCA_900111105.1
mO
226 Streptococcus mutans UA159 GCA_000007465.2
148 Streptococcus ferus DSM 20646 GCA_000372425.1
148 Streptococcus macacae NCTC 11558 GCA_000187995.3
148 Streptococcus sobrinus DSM 20742 = ATCC 33478 GCA_000686605.1
133 Streptococcus massiliensis DSM 18628 GCA_000380065.1
mP
180 Clostridium beijerinckii GCA_000833105.2
180 Clostridium chromiireducens GCA_002029255.1
180 Clostridium puniceum GCA_002006345.1
180 Clostridium saccharobutylicum DSM 13864 GCA_000473995.1
180 Clostridium saccharoperbutylacetonicum N1-4_28HMT_29 GCA_000340885.1
163 Clostridium butyricum GCA_001456065.2
138 Clostridium taeniosporum GCA_001735765.1
mQ
148 Erwinia iniecta GCA_001267535.1
148 Escherichia coli IAI39 GCA_000026345.1
148 Escherichia coli O104_3AH4 str. 2011C-3493 GCA_000299455.1
148 Escherichia coli O157_3AH7 str. Sakai GCA_000008865.1
148 Escherichia coli O83_3AH1 str. NRG 857C GCA_000183345.1
148 Escherichia coli UMN026 GCA_000026325.2
148 Escherichia coli str. K-12 substr. MG1655 GCA_000005845.2
148 Shigella dysenteriae Sd197 GCA_000012005.1
148 Shigella flexneri 2a str. 301 GCA_000006925.2
148 Tumebacillus flagellatus GCA_000714935.1
143 Rosenbergiella nectarea GCA_900111105.1
131 Bacillus humi GCA_001439915.1
mR
194 Deinococcus radiodurans R1 GCA_000008565.1
101 Deinococcus deserti VCD115 GCA_000020685.1
95 Deinococcus puniceus GCA_001644565.1
mS
192 Clostridium beijerinckii GCA_000833105.2
192 Clostridium saccharobutylicum DSM 13864 GCA_000473995.1
192 Clostridium saccharoperbutylacetonicum N1-4_28HMT_29 GCA_000340885.1
185 Clostridium puniceum GCA_002006345.1
174 Clostridium butyricum GCA_001456065.2
174 Clostridium chromiireducens GCA_002029255.1
mT
251 Clostridium beijerinckii GCA_000833105.2

251 Clostridium puniceum GCA_002006345.1
251 Clostridium saccharoperbutylacetonicum N1-4_28HMT_29 GCA_000340885.1
246 Clostridium neonatale GCA_001458595.1
242 Clostridium saccharobutylicum DSM 13864 GCA_000473995.1
mU
207 Deinococcus radiodurans R1 GCA_000008565.1
138 Deinococcus gobiensis I-0 GCA_000252445.1
116 Deinococcus puniceus GCA_001644565.1
mV
193 Rhodobacter sphaeroides ATCC 17025 GCA_000016405.1
178 Pseudorhodobacter psychrotolerans GCA_001294535.1
178 Rhodobacter sphaeroides 2.4.1 GCA_000012905.2
176 Gemmobacter aquatilis GCA_900110025.1
mW
214 Clostridium beijerinckii GCA_000833105.2
214 Clostridium puniceum GCA_002006345.1
214 Clostridium saccharoperbutylacetonicum N1-4_28HMT_29 GCA_000340885.1
210 Clostridium butyricum GCA_001456065.2
205 Clostridium chromiireducens GCA_002029255.1
205 Clostridium saccharobutylicum DSM 13864 GCA_000473995.1
mX
225 Escherichia coli IAI39 GCA_000026345.1
225 Escherichia coli O104_3AH4 str. 2011C-3493 GCA_000299455.1
225 Escherichia coli O157_3AH7 str. Sakai GCA_000008865.1
225 Escherichia coli O83_3AH1 str. NRG 857C GCA_000183345.1
225 Escherichia coli UMN026 GCA_000026325.2
225 Escherichia coli str. K-12 substr. MG1655 GCA_000005845.2
225 Shigella flexneri 2a str. 301 GCA_000006925.2
225 Tumebacillus flagellatus GCA_000714935.1
216 Shigella dysenteriae Sd197 GCA_000012005.1
208 Erwinia iniecta GCA_001267535.1
mY
233 Deinococcus radiodurans R1 GCA_000008565.1
181 Deinococcus gobiensis I-0 GCA_000252445.1
159 Deinococcus deserti VCD115 GCA_000020685.1
mZ
285 Staphylococcus capitis subsp. capitis GCA_001028645.1
285 Staphylococcus epidermidis ATCC 12228 GCA_000007645.1
285 Staphylococcus haemolyticus JCSC1435 GCA_000009865.1
285 Staphylococcus hominis subsp. hominis C80 GCA_000183685.1
278 Staphylococcus lugdunensis HKU09-01 GCA_000025085.1
270 Megasphaera cerevisiae DSM 20462 GCA_001045675.1
270 Staphylococcus aureus subsp. aureus NCTC 8325 GCA_000013425.1
270 Staphylococcus simiae CCM 7213 GCA_000235645.2
270 Staphylococcus warneri SG1 GCA_000332735.1
n0
188 Clostridium beijerinckii GCA_000833105.2
188 Clostridium puniceum GCA_002006345.1
188 Clostridium saccharobutylicum DSM 13864 GCA_000473995.1
188 Clostridium saccharoperbutylacetonicum N1-4_28HMT_29 GCA_000340885.1
174 Clostridium chromiireducens GCA_002029255.1
170 Clostridium amylolyticum GCA_900142075.1
170 Clostridium polynesiense GCA_000820705.1
n1
205 Rhodobacter sphaeroides 2.4.1 GCA_000012905.2
173 Rhodobacter sphaeroides ATCC 17025 GCA_000016405.1
168 Haematobacter massiliensis GCA_000740795.1
n2
226 Escherichia coli IAI39 GCA_000026345.1
226 Escherichia coli O104_3AH4 str. 2011C-3493 GCA_000299455.1
226 Escherichia coli O157_3AH7 str. Sakai GCA_000008865.1
226 Escherichia coli O83_3AH1 str. NRG 857C GCA_000183345.1
226 Escherichia coli UMN026 GCA_000026325.2
226 Escherichia coli str. K-12 substr. MG1655 GCA_000005845.2
226 Shigella flexneri 2a str. 301 GCA_000006925.2
226 Tumebacillus flagellatus GCA_000714935.1
217 Shigella dysenteriae Sd197 GCA_000012005.1
186 Erwinia iniecta GCA_001267535.1
n3
108 Clostridium butyricum GCA_001456065.2
107 Clostridium beijerinckii GCA_000833105.2
107 Clostridium saccharoperbutylacetonicum N1-4_28HMT_29 GCA_000340885.1
94 Clostridium puniceum GCA_002006345.1
n4
254 Deinococcus radiodurans R1 GCA_000008565.1
188 Deinococcus deserti VCD115 GCA_000020685.1

172 Deinococcus soli Cha et al. 2016 GCA_001007995.1
n5
202 Clostridium beijerinckii GCA_000833105.2
202 Clostridium chromiireducens GCA_002029255.1
202 Clostridium puniceum GCA_002006345.1
202 Clostridium saccharobutylicum DSM 13864 GCA_000473995.1
202 Clostridium saccharoperbutylacetonicum N1-4_28HMT_29 GCA_000340885.1
194 Clostridium butyricum GCA_001456065.2
182 Clostridium cavendishii DSM 21758 GCA_900141845.1
n6
223 Enterococcus faecalis V583 GCA_000007785.1
223 Streptomyces cinnamoneus GCA_001885705.1
220 Enterococcus haemoperoxidus ATCC BAA-382 GCA_000407165.1
214 Enterococcus hirae ATCC 9790 GCA_000271405.2
n7
176 Streptococcus mutans UA159 GCA_000007465.2
114 Streptococcus salivarius GCA_000785515.1
114 Streptococcus thermophilus JIM 8232 GCA_000253395.1
103 Streptococcus equinus GCA_000964315.1
103 Streptococcus gallolyticus subsp. gallolyticus DSM 16831 GCA_002000985.1
103 Streptococcus orisratti DSM 15617 GCA_000380105.1
103 Streptococcus ratti FA-1 = DSM 20564 GCA_000286075.1
n8
256 Deinococcus radiodurans R1 GCA_000008565.1
185 Deinococcus gobiensis I-0 GCA_000252445.1
179 Deinococcus puniceus GCA_001644565.1
n9
128 Deinococcus radiodurans R1 GCA_000008565.1
77 Deinococcus gobiensis I-0 GCA_000252445.1
75 Deinococcus deserti VCD115 GCA_000020685.1
na
213 Clostridium beijerinckii GCA_000833105.2
213 Clostridium saccharobutylicum DSM 13864 GCA_000473995.1
213 Clostridium saccharoperbutylacetonicum N1-4_28HMT_29 GCA_000340885.1
198 Clostridium chromiireducens GCA_002029255.1
198 Clostridium puniceum GCA_002006345.1
190 Clostridium butyricum GCA_001456065.2
nb
255 Escherichia coli IAI39 GCA_000026345.1
255 Escherichia coli O104_3AH4 str. 2011C-3493 GCA_000299455.1
255 Escherichia coli O157_3AH7 str. Sakai GCA_000008865.1
255 Escherichia coli O83_3AH1 str. NRG 857C GCA_000183345.1
255 Escherichia coli UMN026 GCA_000026325.2
255 Escherichia coli str. K-12 substr. MG1655 GCA_000005845.2
255 Shigella dysenteriae Sd197 GCA_000012005.1
255 Shigella flexneri 2a str. 301 GCA_000006925.2
255 Tumebacillus flagellatus GCA_000714935.1
221 Erwinia iniecta GCA_001267535.1
212 Cronobacter sakazakii GCA_000982825.1
212 Erwinia toletana DAPP-PG 735 GCA_000336255.1
nc
335 Clostridium beijerinckii GCA_000833105.2
335 Clostridium puniceum GCA_002006345.1
335 Clostridium saccharoperbutylacetonicum N1-4_28HMT_29 GCA_000340885.1
327 Clostridium saccharobutylicum DSM 13864 GCA_000473995.1
312 Clostridium chromiireducens GCA_002029255.1
nd
161 Bifidobacterium adolescentis ATCC 15703 GCA_000010425.1
149 Bifidobacterium asteroides PRL2011 GCA_000304215.1
145 Bifidobacterium choerinum GCA_000741135.1
145 Bifidobacterium stellenboschense GCA_000741785.1
ne
260 Streptococcus mutans UA159 GCA_000007465.2
220 Streptococcus ferus DSM 20646 GCA_000372425.1
212 Streptococcus macacae NCTC 11558 GCA_000187995.3
nf
235 Clostridium beijerinckii GCA_000833105.2
235 Clostridium puniceum GCA_002006345.1
235 Clostridium saccharoperbutylacetonicum N1-4_28HMT_29 GCA_000340885.1
226 Clostridium saccharobutylicum DSM 13864 GCA_000473995.1
219 Clostridium butyricum GCA_001456065.2
ng
192 Staphylococcus cohnii subsp. cohnii GCA_000972575.1
192 Staphylococcus gallinarum GCA_000875895.1
192 Staphylococcus saprophyticus subsp. saprophyticus ATCC 15305 GCA_000010125.1
192 Staphylococcus sciuri GCA_002209165.1

192 Staphylococcus succinus GCA_001902315.1
192 Staphylococcus xylosus GCA_000706685.1
186 Megasphaera cerevisiae DSM 20462 GCA_001045675.1
186 Staphylococcus arlettae CVD059 GCA_000295715.1
186 Staphylococcus aureus subsp. aureus NCTC 8325 GCA_000013425.1
186 Staphylococcus epidermidis ATCC 12228 GCA_000007645.1
186 Staphylococcus haemolyticus JCSC1435 GCA_000009865.1
186 Staphylococcus hominis subsp. hominis C80 GCA_000183685.1
186 Staphylococcus hyicus GCA_000816085.1
186 Staphylococcus lentus F1142 GCA_000286395.1
186 Staphylococcus lugdunensis HKU09-01 GCA_000025085.1
186 Staphylococcus lutrae GCA_002101335.1
186 Staphylococcus saprophyticus GCA_001074355.1
186 Staphylococcus simiae CCM 7213 GCA_000235645.2
186 Staphylococcus warneri SG1 GCA_000332735.1
174 Staphylococcus capitis subsp. capitis GCA_001028645.1
nh
222 Clostridium beijerinckii GCA_000833105.2
222 Clostridium saccharobutylicum DSM 13864 GCA_000473995.1
222 Clostridium saccharoperbutylacetonicum N1-4_28HMT_29 GCA_000340885.1
216 Clostridium puniceum GCA_002006345.1
205 Clostridium chromiireducens GCA_002029255.1
ni
266 Enterococcus faecalis V583 GCA_000007785.1
266 Streptomyces cinnamoneus GCA_001885705.1
240 Enterococcus asini ATCC 700915 GCA_000407365.1
240 Enterococcus canis NBRC 100695 GCA_001544375.1
240 Enterococcus casseliflavus EC20 GCA_000157355.2
240 Enterococcus dispar ATCC 51266 GCA_000406945.1
240 Enterococcus faecium DO GCA_000174395.2
240 Enterococcus hirae ATCC 9790 GCA_000271405.2
240 Enterococcus mundtii QU 25 GCA_000504125.1
240 Enterococcus rivorum GCA_001742285.1
240 Enterococcus saccharolyticus subsp. saccharolyticus ATCC 43076 GCA_000407285.1
235 Enterococcus gilvus ATCC BAA-350 GCA_000407545.1
235 Enterococcus haemoperoxidus ATCC BAA-382 GCA_000407165.1
235 Enterococcus hermanniensis GCA_001885945.1
235 Enterococcus italicus DSM 15952 GCA_000185365.1
235 Enterococcus malodoratus ATCC 43197 GCA_000407185.1
235 Enterococcus pallens ATCC BAA-351 GCA_000407485.1
235 Enterococcus phoeniculicola ATCC BAA-412 GCA_000407505.1
235 Enterococcus pseudoavium NBRC 100491 GCA_001544295.1
235 Enterococcus thailandicus GCA_001652875.1
nj
261 Deinococcus radiodurans R1 GCA_000008565.1
171 Deinococcus deserti VCD115 GCA_000020685.1
167 Deinococcus puniceus GCA_001644565.1
nk
184 Rhodobacter sphaeroides 2.4.1 GCA_000012905.2
184 Rhodobacter sphaeroides ATCC 17025 GCA_000016405.1
176 Gemmobacter aquatilis GCA_900110025.1
176 Gemmobacter nectariphilus DSM 15620 GCA_000429765.1
176 Haematobacter massiliensis GCA_000740795.1
176 Pseudorhodobacter psychrotolerans GCA_001294535.1
176 Rhodobacter capsulatus SB 1003 GCA_000021865.1
169 Gemmobacter megaterium GCA_900156815.1
nl
77 Escherichia coli IAI39 GCA_000026345.1
77 Escherichia coli O104_3AH4 str. 2011C-3493 GCA_000299455.1
77 Escherichia coli O157_3AH7 str. Sakai GCA_000008865.1
77 Escherichia coli UMN026 GCA_000026325.2
77 Escherichia coli str. K-12 substr. MG1655 GCA_000005845.2
77 Shigella dysenteriae Sd197 GCA_000012005.1
77 Shigella flexneri 2a str. 301 GCA_000006925.2
77 Tumebacillus flagellatus GCA_000714935.1
76 Escherichia coli O83_3AH1 str. NRG 857C GCA_000183345.1
73 Oceanisphaera profunda GCA_002157895.1
nm
240 Deinococcus radiodurans R1 GCA_000008565.1
166 Deinococcus gobiensis I-0 GCA_000252445.1
161 Deinococcus deserti VCD115 GCA_000020685.1
nn
257 Enterococcus faecalis V583 GCA_000007785.1
257 Streptomyces cinnamoneus GCA_001885705.1
232 Enterococcus hirae ATCC 9790 GCA_000271405.2
221 Enterococcus canis NBRC 100695 GCA_001544375.1

221 Enterococcus casseliflavus EC20 GCA_000157355.2
221 Enterococcus dispar ATCC 51266 GCA_000406945.1
221 Enterococcus faecium DO GCA_000174395.2
221 Enterococcus haemoperoxidus ATCC BAA-382 GCA_000407165.1
221 Enterococcus mundtii QU 25 GCA_000504125.1
221 Enterococcus phoeniculicola ATCC BAA-412 GCA_000407505.1
221 Enterococcus rivorum GCA_001742285.1
221 Enterococcus saccharolyticus subsp. saccharolyticus ATCC 43076 GCA_000407285.1
221 Enterococcus thailandicus GCA_001652875.1
no
200 Escherichia coli IAI39 GCA_000026345.1
200 Escherichia coli O104_3AH4 str. 2011C-3493 GCA_000299455.1
200 Escherichia coli O157_3AH7 str. Sakai GCA_000008865.1
200 Escherichia coli O83_3AH1 str. NRG 857C GCA_000183345.1
200 Escherichia coli UMN026 GCA_000026325.2
200 Escherichia coli str. K-12 substr. MG1655 GCA_000005845.2
200 Shigella dysenteriae Sd197 GCA_000012005.1
200 Shigella flexneri 2a str. 301 GCA_000006925.2
200 Tumebacillus flagellatus GCA_000714935.1
185 Erwinia iniecta GCA_001267535.1
181 Plesiomonas shigelloides GCA_900087055.1
np
258 Enterococcus faecalis V583 GCA_000007785.1
258 Streptomyces cinnamoneus GCA_001885705.1
238 Enterococcus canis NBRC 100695 GCA_001544375.1
238 Enterococcus dispar ATCC 51266 GCA_000406945.1
238 Enterococcus faecium DO GCA_000174395.2
238 Enterococcus haemoperoxidus ATCC BAA-382 GCA_000407165.1
238 Enterococcus hirae ATCC 9790 GCA_000271405.2
238 Enterococcus massiliensis GCA_001050095.1
238 Enterococcus mundtii QU 25 GCA_000504125.1
238 Enterococcus phoeniculicola ATCC BAA-412 GCA_000407505.1
238 Enterococcus rivorum GCA_001742285.1
238 Enterococcus thailandicus GCA_001652875.1
226 Enterococcus asini ATCC 700915 GCA_000407365.1
nq
283 Streptococcus mutans UA159 GCA_000007465.2
196 Streptococcus macacae NCTC 11558 GCA_000187995.3
193 Streptococcus sobrinus DSM 20742 = ATCC 33478 GCA_000686605.1
nr
169 Clostridium beijerinckii GCA_000833105.2
169 Clostridium chromiireducens GCA_002029255.1
169 Clostridium puniceum GCA_002006345.1
169 Clostridium saccharobutylicum DSM 13864 GCA_000473995.1
169 Clostridium saccharoperbutylacetonicum N1-4_28HMT_29 GCA_000340885.1
164 Clostridium butyricum GCA_001456065.2
140 Clostridium neonatale GCA_001458595.1
ns
248 Deinococcus radiodurans R1 GCA_000008565.1
178 Deinococcus gobiensis I-0 GCA_000252445.1
161 Deinococcus deserti VCD115 GCA_000020685.1
nt
229 Rhodobacter sphaeroides 2.4.1 GCA_000012905.2
203 Rhodobacter sphaeroides ATCC 17025 GCA_000016405.1
163 Thioclava dalianensis GCA_000715505.1
163 Thioclava indica GCA_000714545.1
nu
242 Clostridium beijerinckii GCA_000833105.2
242 Clostridium saccharobutylicum DSM 13864 GCA_000473995.1
242 Clostridium saccharoperbutylacetonicum N1-4_28HMT_29 GCA_000340885.1
227 Clostridium puniceum GCA_002006345.1
215 Clostridium botulinum B str. Eklund 17B _28NRP_29 GCA_000020165.1
215 Clostridium chromiireducens GCA_002029255.1
215 Clostridium taeniosporum GCA_001735765.1
nv
193 Streptococcus mutans UA159 GCA_000007465.2
100 Streptococcus macacae NCTC 11558 GCA_000187995.3
99 Streptococcus ferus DSM 20646 GCA_000372425.1
99 Streptococcus sobrinus DSM 20742 = ATCC 33478 GCA_000686605.1
nw
208 Rhodobacter sphaeroides 2.4.1 GCA_000012905.2
204 Rhodobacter sphaeroides ATCC 17025 GCA_000016405.1
182 Rhodobacter capsulatus SB 1003 GCA_000021865.1
nx
230 Deinococcus radiodurans R1 GCA_000008565.1
159 Deinococcus deserti VCD115 GCA_000020685.1

151 Deinococcus gobiensis I-0 GCA_000252445.1
ny
309 Bacillus thuringiensis YBT-1518 GCA_000497525.2
301 Bacillus anthracis str. Ames GCA_000007845.1
301 Bacillus anthracis str. Sterne GCA_000008165.1
301 Bacillus cereus ATCC 14579 GCA_000007825.1
301 Bacillus mycoides GCA_000832605.1
301 Bacillus pseudomycoides DSM 12442 GCA_000161455.1
301 _5BBacillus thuringiensis_5D serovar konkukian str. 97-27 GCA_000008505.1
225 Bacillus manliponensis GCA_000712595.1
nz
207 Bacillus anthracis str. Ames GCA_000007845.1
207 Bacillus anthracis str. Sterne GCA_000008165.1
207 Bacillus cereus ATCC 14579 GCA_000007825.1
207 Bacillus mycoides GCA_000832605.1
207 Bacillus pseudomycoides DSM 12442 GCA_000161455.1
207 Bacillus thuringiensis YBT-1518 GCA_000497525.2
207 _5BBacillus thuringiensis_5D serovar konkukian str. 97-27 GCA_000008505.1
184 Bacillus manliponensis GCA_000712595.1
171 Bacillus flexus GCA_002024265.1
171 Bacillus megaterium NBRC 15308 = ATCC 14581 GCA_000832985.1
nA
160 Rhodobacter sphaeroides 2.4.1 GCA_000012905.2
130 Gemmobacter megaterium GCA_900156815.1
130 Pseudorhodobacter ferrugineus DSM 5888 GCA_000420745.1
130 Pseudorhodobacter wandonensis GCA_001202035.1
130 Rhodobacter sphaeroides ATCC 17025 GCA_000016405.1
115 Defluviimonas alba GCA_001620265.1
115 Gemmobacter aquatilis GCA_900110025.1
115 Pseudorhodobacter psychrotolerans GCA_001294535.1
nB
203 Rhodobacter sphaeroides 2.4.1 GCA_000012905.2
197 Rhodobacter sphaeroides ATCC 17025 GCA_000016405.1
174 Pseudorhodobacter psychrotolerans GCA_001294535.1
nC
173 Escherichia coli IAI39 GCA_000026345.1
173 Escherichia coli O104_3AH4 str. 2011C-3493 GCA_000299455.1
173 Escherichia coli O157_3AH7 str. Sakai GCA_000008865.1
173 Escherichia coli O83_3AH1 str. NRG 857C GCA_000183345.1
173 Escherichia coli UMN026 GCA_000026325.2
173 Escherichia coli str. K-12 substr. MG1655 GCA_000005845.2
173 Shigella dysenteriae Sd197 GCA_000012005.1
173 Shigella flexneri 2a str. 301 GCA_000006925.2
173 Tumebacillus flagellatus GCA_000714935.1
169 Erwinia iniecta GCA_001267535.1
147 Rosenbergiella nectarea GCA_900111105.1
nD
165 Clostridium beijerinckii GCA_000833105.2
165 Clostridium puniceum GCA_002006345.1
165 Clostridium saccharobutylicum DSM 13864 GCA_000473995.1
165 Clostridium saccharoperbutylacetonicum N1-4_28HMT_29 GCA_000340885.1
151 Clostridium chromiireducens GCA_002029255.1
142 Clostridium butyricum GCA_001456065.2
nE
258 Clostridium beijerinckii GCA_000833105.2
258 Clostridium saccharobutylicum DSM 13864 GCA_000473995.1
258 Clostridium saccharoperbutylacetonicum N1-4_28HMT_29 GCA_000340885.1
252 Clostridium puniceum GCA_002006345.1
222 Clostridium amylolyticum GCA_900142075.1
222 Clostridium polynesiense GCA_000820705.1
nF
233 Bifidobacterium adolescentis ATCC 15703 GCA_000010425.1
217 Bifidobacterium breve DSM 20213 = JCM 1192 GCA_001025175.1
216 Bifidobacterium callitrichos DSM 23973 GCA_000741175.1
nG
152 Rhodobacter sphaeroides ATCC 17025 GCA_000016405.1
138 Rhodobacter sphaeroides 2.4.1 GCA_000012905.2
123 Pseudorhodobacter ferrugineus DSM 5888 GCA_000420745.1
123 Pseudorhodobacter wandonensis GCA_001202035.1
nH
251 Bifidobacterium adolescentis ATCC 15703 GCA_000010425.1
224 Bifidobacterium dentium JCM 1195 = DSM 20436 GCA_001042595.1
224 Bifidobacterium tsurumiense GCA_000741765.1
221 Bifidobacterium thermophilum GCA_000741495.1
nI
217 Clostridium beijerinckii GCA_000833105.2

217 Clostridium chromiireducens GCA_002029255.1
217 Clostridium puniceum GCA_002006345.1
217 Clostridium saccharobutylicum DSM 13864 GCA_000473995.1
217 Clostridium saccharoperbutylacetonicum N1-4_28HMT_29 GCA_000340885.1
209 Clostridium taeniosporum GCA_001735765.1
207 Clostridium butyricum GCA_001456065.2
nJ
187 Lactobacillus gasseri ATCC 33323 = JCM 1131 GCA_000014425.1
160 Lactobacillus hominis DSM 23910 = CRBIP 24.179 GCA_000296835.1
126 Lactobacillus antri DSM 16041 GCA_000160835.1
126 Lactobacillus fermentum IFO 3956 GCA_000010145.1
126 Lactobacillus frumenti DSM 13145 GCA_001436045.1
126 Lactobacillus mucosae LM1 GCA_000248095.3
126 Lactobacillus reuteri DSM 20016 GCA_000016825.1
126 Lactobacillus secaliphilus GCA_001437055.1
126 Lactobacillus vaginalis DSM 5837 = ATCC 49540 GCA_000159435.1
nK
233 Rhodobacter sphaeroides 2.4.1 GCA_000012905.2
203 Rhodobacter sphaeroides ATCC 17025 GCA_000016405.1
198 Pseudorhodobacter psychrotolerans GCA_001294535.1
nL
310 Bifidobacterium adolescentis ATCC 15703 GCA_000010425.1
257 Bifidobacterium breve DSM 20213 = JCM 1192 GCA_001025175.1
255 Bifidobacterium tsurumiense GCA_000741765.1
nM
205 Escherichia coli IAI39 GCA_000026345.1
205 Escherichia coli O104_3AH4 str. 2011C-3493 GCA_000299455.1
205 Escherichia coli O157_3AH7 str. Sakai GCA_000008865.1
205 Escherichia coli O83_3AH1 str. NRG 857C GCA_000183345.1
205 Escherichia coli UMN026 GCA_000026325.2
205 Escherichia coli str. K-12 substr. MG1655 GCA_000005845.2
205 Shigella flexneri 2a str. 301 GCA_000006925.2
205 Tumebacillus flagellatus GCA_000714935.1
196 Shigella dysenteriae Sd197 GCA_000012005.1
153 Cronobacter sakazakii GCA_000982825.1
153 Erwinia iniecta GCA_001267535.1
nN
308 Bacillus thuringiensis YBT-1518 GCA_000497525.2
304 Bacillus anthracis str. Ames GCA_000007845.1
304 Bacillus anthracis str. Sterne GCA_000008165.1
304 Bacillus cereus ATCC 14579 GCA_000007825.1
304 Bacillus mycoides GCA_000832605.1
304 Bacillus pseudomycoides DSM 12442 GCA_000161455.1
304 _5BBacillus thuringiensis_5D serovar konkukian str. 97-27 GCA_000008505.1
245 Bacillus cytotoxicus NVH 391-98 GCA_000017425.1
nO
309 Deinococcus radiodurans R1 GCA_000008565.1
208 Deinococcus marmoris DSM 12784 GCA_000701405.1
208 Deinococcus swuensis GCA_000800395.1
206 Deinococcus puniceus GCA_001644565.1
nP
137 Bacillus halmapalus GCA_002019665.1
116 Bacillus aquimaris TF-12 GCA_001648555.1
116 Bacillus pseudofirmus OF4 GCA_000005825.2
115 Bacillus vietnamensis NBRC 101237 GCA_001591825.1
nQ
256 Bacillus anthracis str. Ames GCA_000007845.1
256 Bacillus anthracis str. Sterne GCA_000008165.1
256 Bacillus cereus ATCC 14579 GCA_000007825.1
256 Bacillus mycoides GCA_000832605.1
256 Bacillus pseudomycoides DSM 12442 GCA_000161455.1
256 Bacillus thuringiensis YBT-1518 GCA_000497525.2
256 _5BBacillus thuringiensis_5D serovar konkukian str. 97-27 GCA_000008505.1
228 Bacillus flexus GCA_002024265.1
228 Bacillus megaterium NBRC 15308 = ATCC 14581 GCA_000832985.1
215 Bacillus manliponensis GCA_000712595.1
nR
235 Bacillus anthracis str. Ames GCA_000007845.1
235 Bacillus anthracis str. Sterne GCA_000008165.1
235 Bacillus cereus ATCC 14579 GCA_000007825.1
235 Bacillus mycoides GCA_000832605.1
235 Bacillus pseudomycoides DSM 12442 GCA_000161455.1
235 Bacillus thuringiensis YBT-1518 GCA_000497525.2
235 _5BBacillus thuringiensis_5D serovar konkukian str. 97-27 GCA_000008505.1
197 Bacillus horneckiae GCA_001636335.1
197 Bacillus solani GCA_001420595.1

195 Bacillus manliponensis GCA_000712595.1
nS
205 Escherichia coli IAI39 GCA_000026345.1
205 Escherichia coli O104_3AH4 str. 2011C-3493 GCA_000299455.1
205 Escherichia coli O157_3AH7 str. Sakai GCA_000008865.1
205 Escherichia coli O83_3AH1 str. NRG 857C GCA_000183345.1
205 Escherichia coli UMN026 GCA_000026325.2
205 Escherichia coli str. K-12 substr. MG1655 GCA_000005845.2
205 Shigella dysenteriae Sd197 GCA_000012005.1
205 Tumebacillus flagellatus GCA_000714935.1
197 Shigella flexneri 2a str. 301 GCA_000006925.2
170 Cronobacter sakazakii GCA_000982825.1
170 Enterobacter cloacae subsp. cloacae ATCC 13047 GCA_000025565.1
170 Enterobacter hormaechei subsp. steigerwaltii GCA_001729725.1
170 Enterobacter kobei GCA_900185885.1
170 Erwinia iniecta GCA_001267535.1
170 Erwinia toletana DAPP-PG 735 GCA_000336255.1
170 Klebsiella oxytoca GCA_001022195.1
170 Kosakonia cowanii GCA_001975225.1
170 Pantoea alhagi GCA_002101395.1
170 Pluralibacter gergoviae GCA_000757785.1
170 Pseudescherichia vulneris NBRC 102420 GCA_000759795.1
170 Salmonella enterica subsp. enterica serovar Typhi str. CT18 GCA_000195995.1
170 Salmonella enterica subsp. enterica serovar Typhimurium str. LT2 GCA_000006945.2
170 Trabulsiella odontotermitis GCA_001297765.1
nT
182 Deinococcus radiodurans R1 GCA_000008565.1
147 Deinococcus deserti VCD115 GCA_000020685.1
147 Deinococcus gobiensis I-0 GCA_000252445.1
141 Deinococcus puniceus GCA_001644565.1
nU
179 Streptococcus mutans UA159 GCA_000007465.2
110 Streptococcus equinus GCA_000964315.1
110 Streptococcus gallolyticus subsp. gallolyticus DSM 16831 GCA_002000985.1
110 Streptococcus orisratti DSM 15617 GCA_000380105.1
110 Streptococcus ratti FA-1 = DSM 20564 GCA_000286075.1
106 Streptococcus gordonii str. Challis substr. CH1 GCA_000017005.1
106 Streptococcus phocae subsp. salmonis GCA_000772915.1
nV
174 Clostridium beijerinckii GCA_000833105.2
174 Clostridium puniceum GCA_002006345.1
174 Clostridium saccharoperbutylacetonicum N1-4_28HMT_29 GCA_000340885.1
160 Clostridium saccharobutylicum DSM 13864 GCA_000473995.1
153 Clostridium butyricum GCA_001456065.2
nW
166 Deinococcus radiodurans R1 GCA_000008565.1
126 Deinococcus geothermalis DSM 11300 GCA_000196275.1
118 Deinococcus proteolyticus MRP GCA_000190555.1
nX
241 Streptococcus mutans UA159 GCA_000007465.2
211 Streptococcus sobrinus DSM 20742 = ATCC 33478 GCA_000686605.1
194 Streptococcus gordonii str. Challis substr. CH1 GCA_000017005.1
nY
272 Streptococcus mutans UA159 GCA_000007465.2
202 Streptococcus marimammalium DSM 18627 GCA_000380045.1
200 Streptococcus macacae NCTC 11558 GCA_000187995.3
nZ
257 Escherichia coli IAI39 GCA_000026345.1
257 Escherichia coli O104_3AH4 str. 2011C-3493 GCA_000299455.1
257 Escherichia coli O157_3AH7 str. Sakai GCA_000008865.1
257 Escherichia coli O83_3AH1 str. NRG 857C GCA_000183345.1
257 Escherichia coli UMN026 GCA_000026325.2
257 Escherichia coli str. K-12 substr. MG1655 GCA_000005845.2
257 Shigella dysenteriae Sd197 GCA_000012005.1
257 Shigella flexneri 2a str. 301 GCA_000006925.2
257 Tumebacillus flagellatus GCA_000714935.1
242 Erwinia iniecta GCA_001267535.1
239 Rosenbergiella nectarea GCA_900111105.1
o0
204 Lactobacillus gasseri ATCC 33323 = JCM 1131 GCA_000014425.1
172 Lactobacillus hominis DSM 23910 = CRBIP 24.179 GCA_000296835.1
126 Lactobacillus iners DSM 13335 GCA_000160875.1
o1
225 Deinococcus radiodurans R1 GCA_000008565.1
149 Deinococcus proteolyticus MRP GCA_000190555.1
148 Deinococcus geothermalis DSM 11300 GCA_000196275.1

o2
262 Clostridium beijerinckii GCA_000833105.2
262 Clostridium saccharoperbutylacetonicum N1-4_28HMT_29 GCA_000340885.1
256 Clostridium puniceum GCA_002006345.1
254 Clostridium saccharobutylicum DSM 13864 GCA_000473995.1
o3
163 Clostridium beijerinckii GCA_000833105.2
163 Clostridium chromiireducens GCA_002029255.1
163 Clostridium puniceum GCA_002006345.1
163 Clostridium saccharobutylicum DSM 13864 GCA_000473995.1
163 Clostridium saccharoperbutylacetonicum N1-4_28HMT_29 GCA_000340885.1
163 Clostridium taeniosporum GCA_001735765.1
152 Clostridium botulinum B str. Eklund 17B _28NRP_29 GCA_000020165.1
152 Clostridium uliginosum GCA_900112485.1
139 Clostridium butyricum GCA_001456065.2
o4
187 Bifidobacterium adolescentis ATCC 15703 GCA_000010425.1
172 Bifidobacterium thermophilum GCA_000741495.1
168 Bifidobacterium dentium JCM 1195 = DSM 20436 GCA_001042595.1
168 Bifidobacterium tsurumiense GCA_000741765.1
o5
169 Deinococcus radiodurans R1 GCA_000008565.1
137 Deinococcus deserti VCD115 GCA_000020685.1
132 Deinococcus proteolyticus MRP GCA_000190555.1
o6
178 Rhodobacter sphaeroides 2.4.1 GCA_000012905.2
146 Rhodobacter sphaeroides ATCC 17025 GCA_000016405.1
132 Haematobacter massiliensis GCA_000740795.1
o7
158 Enterococcus canis NBRC 100695 GCA_001544375.1
158 Enterococcus faecalis V583 GCA_000007785.1
158 Enterococcus faecium DO GCA_000174395.2
158 Enterococcus haemoperoxidus ATCC BAA-382 GCA_000407165.1
158 Enterococcus hirae ATCC 9790 GCA_000271405.2
158 Enterococcus mundtii QU 25 GCA_000504125.1
158 Enterococcus phoeniculicola ATCC BAA-412 GCA_000407505.1
158 Enterococcus rivorum GCA_001742285.1
158 Enterococcus thailandicus GCA_001652875.1
158 Streptomyces cinnamoneus GCA_001885705.1
157 Enterococcus asini ATCC 700915 GCA_000407365.1
157 Enterococcus casseliflavus EC20 GCA_000157355.2
157 Enterococcus dispar ATCC 51266 GCA_000406945.1
157 Enterococcus saccharolyticus subsp. saccharolyticus ATCC 43076 GCA_000407285.1
153 Melissococcus plutonius S1 GCA_000747585.1
o8
248 Bacillus thuringiensis YBT-1518 GCA_000497525.2
247 Bacillus anthracis str. Ames GCA_000007845.1
247 Bacillus anthracis str. Sterne GCA_000008165.1
247 Bacillus cereus ATCC 14579 GCA_000007825.1
247 Bacillus mycoides GCA_000832605.1
247 Bacillus pseudomycoides DSM 12442 GCA_000161455.1
247 _5BBacillus thuringiensis_5D serovar konkukian str. 97-27 GCA_000008505.1
197 Bacillus lentus NBRC 16444 GCA_001591545.1
o9
217 Enterococcus faecalis V583 GCA_000007785.1
217 Streptomyces cinnamoneus GCA_001885705.1
180 Enterococcus canis NBRC 100695 GCA_001544375.1
180 Enterococcus dispar ATCC 51266 GCA_000406945.1
180 Enterococcus faecium DO GCA_000174395.2
180 Enterococcus gilvus ATCC BAA-350 GCA_000407545.1
180 Enterococcus hermanniensis GCA_001885945.1
180 Enterococcus hirae ATCC 9790 GCA_000271405.2
180 Enterococcus malodoratus ATCC 43197 GCA_000407185.1
180 Enterococcus mundtii QU 25 GCA_000504125.1
180 Enterococcus pallens ATCC BAA-351 GCA_000407485.1
180 Enterococcus pseudoavium NBRC 100491 GCA_001544295.1
180 Enterococcus rivorum GCA_001742285.1
175 Enterococcus haemoperoxidus ATCC BAA-382 GCA_000407165.1
175 Enterococcus phoeniculicola ATCC BAA-412 GCA_000407505.1
175 Enterococcus thailandicus GCA_001652875.1
oa
197 Clostridium beijerinckii GCA_000833105.2
197 Clostridium saccharobutylicum DSM 13864 GCA_000473995.1
197 Clostridium saccharoperbutylacetonicum N1-4_28HMT_29 GCA_000340885.1
190 Clostridium puniceum GCA_002006345.1
171 Clostridium chromiireducens GCA_002029255.1

ob
220 Bacillus thuringiensis YBT-1518 GCA_000497525.2
206 Bacillus anthracis str. Ames GCA_000007845.1
206 Bacillus anthracis str. Sterne GCA_000008165.1
206 Bacillus mycoides GCA_000832605.1
206 Bacillus pseudomycoides DSM 12442 GCA_000161455.1
206 _5BBacillus thuringiensis_5D serovar konkukian str. 97-27 GCA_000008505.1
205 Bacillus cereus ATCC 14579 GCA_000007825.1
oc
204 Bifidobacterium adolescentis ATCC 15703 GCA_000010425.1
177 Bifidobacterium breve DSM 20213 = JCM 1192 GCA_001025175.1
176 Bifidobacterium asteroides PRL2011 GCA_000304215.1
176 Bifidobacterium tsurumiense GCA_000741765.1
od
297 Enterococcus faecalis V583 GCA_000007785.1
297 Streptomyces cinnamoneus GCA_001885705.1
261 Enterococcus canis NBRC 100695 GCA_001544375.1
261 Enterococcus casseliflavus EC20 GCA_000157355.2
261 Enterococcus dispar ATCC 51266 GCA_000406945.1
261 Enterococcus faecium DO GCA_000174395.2
261 Enterococcus haemoperoxidus ATCC BAA-382 GCA_000407165.1
261 Enterococcus hirae ATCC 9790 GCA_000271405.2
261 Enterococcus mundtii QU 25 GCA_000504125.1
261 Enterococcus phoeniculicola ATCC BAA-412 GCA_000407505.1
261 Enterococcus rivorum GCA_001742285.1
261 Enterococcus saccharolyticus subsp. saccharolyticus ATCC 43076 GCA_000407285.1
261 Enterococcus thailandicus GCA_001652875.1
256 Enterococcus gilvus ATCC BAA-350 GCA_000407545.1
256 Enterococcus hermanniensis GCA_001885945.1
256 Enterococcus malodoratus ATCC 43197 GCA_000407185.1
256 Enterococcus pallens ATCC BAA-351 GCA_000407485.1
256 Enterococcus pseudoavium NBRC 100491 GCA_001544295.1
oe
102 Clostridium beijerinckii GCA_000833105.2
102 Clostridium butyricum GCA_001456065.2
102 Clostridium chromiireducens GCA_002029255.1
102 Clostridium puniceum GCA_002006345.1
102 Clostridium saccharobutylicum DSM 13864 GCA_000473995.1
102 Clostridium saccharoperbutylacetonicum N1-4_28HMT_29 GCA_000340885.1
94 Clostridium neonatale GCA_001458595.1
87 Clostridium thermobutyricum DSM 4928 GCA_002050515.1
of
192 Bacillus anthracis str. Ames GCA_000007845.1
192 Bacillus anthracis str. Sterne GCA_000008165.1
192 Bacillus cereus ATCC 14579 GCA_000007825.1
192 Bacillus mycoides GCA_000832605.1
192 Bacillus pseudomycoides DSM 12442 GCA_000161455.1
192 Bacillus thuringiensis YBT-1518 GCA_000497525.2
192 _5BBacillus thuringiensis_5D serovar konkukian str. 97-27 GCA_000008505.1
145 Salinicoccus halodurans GCA_001005905.1
145 Salinicoccus qingdaonensis GCA_900101075.1
144 Bacillus manliponensis GCA_000712595.1
og
248 Streptococcus mutans UA159 GCA_000007465.2
160 Streptococcus sobrinus DSM 20742 = ATCC 33478 GCA_000686605.1
159 Streptococcus iniae GCA_000831485.1
oh
258 Lactobacillus gasseri ATCC 33323 = JCM 1131 GCA_000014425.1
258 Lactobacillus hominis DSM 23910 = CRBIP 24.179 GCA_000296835.1
182 Lactobacillus iners DSM 13335 GCA_000160875.1
182 Lactobacillus psittaci DSM 15354 GCA_000425905.1
181 Lactobacillus amylovorus GCA_000191545.1
181 Lactobacillus crispatus ST1 GCA_000091765.1
oi
239 Clostridium beijerinckii GCA_000833105.2
239 Clostridium puniceum GCA_002006345.1
239 Clostridium saccharoperbutylacetonicum N1-4_28HMT_29 GCA_000340885.1
229 Clostridium butyricum GCA_001456065.2
224 Clostridium taeniosporum GCA_001735765.1
oj
304 Deinococcus radiodurans R1 GCA_000008565.1
236 Deinococcus gobiensis I-0 GCA_000252445.1
186 Deinococcus deserti VCD115 GCA_000020685.1
186 Deinococcus puniceus GCA_001644565.1
ok
254 Rhodobacter sphaeroides 2.4.1 GCA_000012905.2

240 Pseudorhodobacter psychrotolerans GCA_001294535.1
231 Haematobacter massiliensis GCA_000740795.1
ol
185 Rhodobacter sphaeroides 2.4.1 GCA_000012905.2
143 Thioclava dalianensis GCA_000715505.1
143 Thioclava indica GCA_000714545.1
142 Gemmobacter megaterium GCA_900156815.1
142 Pseudorhodobacter ferrugineus DSM 5888 GCA_000420745.1
142 Pseudorhodobacter wandonensis GCA_001202035.1
142 Rhodobacter sphaeroides ATCC 17025 GCA_000016405.1
om
178 Haematobacter massiliensis GCA_000740795.1
170 Defluviimonas alba GCA_001620265.1
170 Rhodobacter capsulatus SB 1003 GCA_000021865.1
166 Rhodobacter sphaeroides 2.4.1 GCA_000012905.2
on
120 Pediococcus ethanolidurans GCA_001437405.1
117 Enterococcus faecalis V583 GCA_000007785.1
117 Streptomyces cinnamoneus GCA_001885705.1
114 Pediococcus cellicola GCA_001438655.1
oo
213 Clostridium beijerinckii GCA_000833105.2
213 Clostridium puniceum GCA_002006345.1
213 Clostridium saccharobutylicum DSM 13864 GCA_000473995.1
213 Clostridium saccharoperbutylacetonicum N1-4_28HMT_29 GCA_000340885.1
202 Clostridium chromiireducens GCA_002029255.1
198 Clostridium butyricum GCA_001456065.2
op
202 Rhodobacter sphaeroides 2.4.1 GCA_000012905.2
174 Rhodobacter sphaeroides ATCC 17025 GCA_000016405.1
172 Pseudorhodobacter psychrotolerans GCA_001294535.1
oq
129 Escherichia coli IAI39 GCA_000026345.1
129 Escherichia coli O104_3AH4 str. 2011C-3493 GCA_000299455.1
129 Escherichia coli O157_3AH7 str. Sakai GCA_000008865.1
129 Escherichia coli O83_3AH1 str. NRG 857C GCA_000183345.1
129 Escherichia coli UMN026 GCA_000026325.2
129 Escherichia coli str. K-12 substr. MG1655 GCA_000005845.2
129 Shigella dysenteriae Sd197 GCA_000012005.1
129 Shigella flexneri 2a str. 301 GCA_000006925.2
129 Tumebacillus flagellatus GCA_000714935.1
109 Buttiauxella ferragutiae ATCC 51602 GCA_001654915.1
109 Cedecea neteri GCA_000757825.1
109 Citrobacter freundii CFNIH1 GCA_000648515.1
109 Cronobacter sakazakii GCA_000982825.1
109 Enterobacter cloacae subsp. cloacae ATCC 13047 GCA_000025565.1
109 Enterobacter hormaechei subsp. steigerwaltii GCA_001729725.1
109 Enterobacter kobei GCA_900185885.1
109 Erwinia iniecta GCA_001267535.1
109 Erwinia persicina NBRC 102418 GCA_001571305.1
109 Erwinia toletana DAPP-PG 735 GCA_000336255.1
109 Klebsiella aerogenes KCTC 2190 GCA_000215745.1
109 Klebsiella oxytoca GCA_001022195.1
109 Kluyvera ascorbata ATCC 33433 GCA_000735365.1
109 Kosakonia cowanii GCA_001975225.1
109 Pluralibacter gergoviae GCA_000757785.1
109 Pseudescherichia vulneris NBRC 102420 GCA_000759795.1
109 Rosenbergiella nectarea GCA_900111105.1
109 Salmonella enterica subsp. enterica serovar Typhi str. CT18 GCA_000195995.1
109 Salmonella enterica subsp. enterica serovar Typhimurium str. LT2 GCA_000006945.2
109 Yokenella regensburgei ATCC 49455 GCA_000735455.1
109 _5BEnterobacter_5D lignolyticus SCF1 GCA_000164865.1
108 Kosakonia sacchari SP1 GCA_000300455.4
108 Mangrovibacter phragmitis GCA_001655675.1
or
220 Lactobacillus gasseri ATCC 33323 = JCM 1131 GCA_000014425.1
204 Lactobacillus hominis DSM 23910 = CRBIP 24.179 GCA_000296835.1
145 Lactobacillus iners DSM 13335 GCA_000160875.1
os
136 Lactobacillus gasseri ATCC 33323 = JCM 1131 GCA_000014425.1
101 Lactobacillus hominis DSM 23910 = CRBIP 24.179 GCA_000296835.1
92 Salinicoccus carnicancri Crm GCA_000330705.1
ot
192 Rhodobacter sphaeroides 2.4.1 GCA_000012905.2
176 Pseudorhodobacter ferrugineus DSM 5888 GCA_000420745.1
176 Pseudorhodobacter wandonensis GCA_001202035.1

170 Aquamicrobium aerolatum DSM 21857 GCA_900113935.1
170 Gemmobacter aquatilis GCA_900110025.1
170 Hoeflea olei GCA_001703635.1
170 Hoeflea phototrophica DFL-43 GCA_000154705.2
170 Nitratireductor basaltis GCA_000733725.1
170 Pseudaminobacter salicylatoxidans KCT001 GCA_000304395.1
170 Pseudorhodobacter psychrotolerans GCA_001294535.1
170 Thioclava dalianensis GCA_000715505.1
170 Thioclava indica GCA_000714545.1
ou
275 Deinococcus radiodurans R1 GCA_000008565.1
183 Deinococcus puniceus GCA_001644565.1
182 Deinococcus deserti VCD115 GCA_000020685.1
ov
146 Deinococcus radiodurans R1 GCA_000008565.1
109 Deinococcus geothermalis DSM 11300 GCA_000196275.1
109 Deinococcus proteolyticus MRP GCA_000190555.1
98 Deinococcus misasensis DSM 22328 GCA_000745915.1
ow
108 Streptococcus mutans UA159 GCA_000007465.2
94 Streptococcus ferus DSM 20646 GCA_000372425.1
82 Aerococcus urinae GCA_001543175.1
82 Streptococcus caballi DSM 19004 GCA_000379985.1
82 Streptococcus criceti HS-6 GCA_000187975.3
82 Streptococcus equinus GCA_000964315.1
82 Streptococcus gallolyticus subsp. gallolyticus DSM 16831 GCA_002000985.1
82 Streptococcus henryi DSM 19005 GCA_000376985.1
82 Streptococcus himalayensis GCA_001708305.1
82 Streptococcus marimammalium DSM 18627 GCA_000380045.1
82 Streptococcus marmotae GCA_001623565.1
82 Streptococcus merionis DSM 19192 GCA_000380085.1
82 Streptococcus minor DSM 17118 GCA_000377005.1
82 Streptococcus ovis DSM 16829 GCA_000380125.1
82 Streptococcus plurextorum DSM 22810 GCA_000423745.1
82 Streptococcus porci DSM 23759 GCA_000423765.1
82 Streptococcus salivarius GCA_000785515.1
82 Streptococcus sanguinis SK36 GCA_000014205.1
82 Streptococcus suis BM407 GCA_000026745.1
82 Streptococcus thermophilus JIM 8232 GCA_000253395.1
82 Streptococcus varani GCA_001375655.1
ox
245 Deinococcus radiodurans R1 GCA_000008565.1
188 Deinococcus puniceus GCA_001644565.1
175 Deinococcus gobiensis I-0 GCA_000252445.1
oy
171 Clostridium beijerinckii GCA_000833105.2
171 Clostridium saccharoperbutylacetonicum N1-4_28HMT_29 GCA_000340885.1
164 Clostridium puniceum GCA_002006345.1
156 Clostridium saccharobutylicum DSM 13864 GCA_000473995.1
oz
198 Deinococcus radiodurans R1 GCA_000008565.1
143 Deinococcus gobiensis I-0 GCA_000252445.1
111 Deinococcus frigens DSM 12807 GCA_000701425.1
111 Deinococcus marmoris DSM 12784 GCA_000701405.1
111 Deinococcus swuensis GCA_000800395.1
oA
201 Bacillus thuringiensis YBT-1518 GCA_000497525.2
185 Bacillus anthracis str. Ames GCA_000007845.1
185 Bacillus anthracis str. Sterne GCA_000008165.1
185 Bacillus cereus ATCC 14579 GCA_000007825.1
185 Bacillus mycoides GCA_000832605.1
185 Bacillus pseudomycoides DSM 12442 GCA_000161455.1
185 _5BBacillus thuringiensis_5D serovar konkukian str. 97-27 GCA_000008505.1
149 Bacillus flexus GCA_002024265.1
149 Bacillus megaterium NBRC 15308 = ATCC 14581 GCA_000832985.1
149 Massilibacterium senegalense GCA_001375675.1
oB
260 Staphylococcus epidermidis ATCC 12228 GCA_000007645.1
260 Staphylococcus haemolyticus JCSC1435 GCA_000009865.1
254 Staphylococcus capitis subsp. capitis GCA_001028645.1
245 Megasphaera cerevisiae DSM 20462 GCA_001045675.1
245 Staphylococcus aureus subsp. aureus NCTC 8325 GCA_000013425.1
245 Staphylococcus hominis subsp. hominis C80 GCA_000183685.1
245 Staphylococcus lugdunensis HKU09-01 GCA_000025085.1
245 Staphylococcus simiae CCM 7213 GCA_000235645.2
245 Staphylococcus warneri SG1 GCA_000332735.1

oC
292 Clostridium beijerinckii GCA_000833105.2
292 Clostridium saccharoperbutylacetonicum N1-4_28HMT_29 GCA_000340885.1
285 Clostridium puniceum GCA_002006345.1
284 Clostridium saccharobutylicum DSM 13864 GCA_000473995.1
oD
253 Streptococcus mutans UA159 GCA_000007465.2
152 Streptococcus marimammalium DSM 18627 GCA_000380045.1
148 Streptococcus macacae NCTC 11558 GCA_000187995.3
148 Streptococcus sobrinus DSM 20742 = ATCC 33478 GCA_000686605.1
oE
269 Streptococcus mutans UA159 GCA_000007465.2
184 Streptococcus gordonii str. Challis substr. CH1 GCA_000017005.1
179 Streptococcus ferus DSM 20646 GCA_000372425.1
oF
301 Streptococcus mutans UA159 GCA_000007465.2
201 Streptococcus sobrinus DSM 20742 = ATCC 33478 GCA_000686605.1
198 Streptococcus cristatus AS 1.3089 GCA_000385925.1
198 Streptococcus gordonii str. Challis substr. CH1 GCA_000017005.1
198 Streptococcus mitis B6 GCA_000027165.1
198 Streptococcus pneumoniae R6 GCA_000007045.1
oG
197 Rhodobacter sphaeroides 2.4.1 GCA_000012905.2
168 Rhodobacter sphaeroides ATCC 17025 GCA_000016405.1
165 Pseudorhodobacter psychrotolerans GCA_001294535.1
oH
305 Escherichia coli IAI39 GCA_000026345.1
305 Escherichia coli O104_3AH4 str. 2011C-3493 GCA_000299455.1
305 Escherichia coli O157_3AH7 str. Sakai GCA_000008865.1
305 Escherichia coli O83_3AH1 str. NRG 857C GCA_000183345.1
305 Escherichia coli UMN026 GCA_000026325.2
305 Escherichia coli str. K-12 substr. MG1655 GCA_000005845.2
305 Shigella dysenteriae Sd197 GCA_000012005.1
305 Shigella flexneri 2a str. 301 GCA_000006925.2
305 Tumebacillus flagellatus GCA_000714935.1
265 Erwinia iniecta GCA_001267535.1
231 Cronobacter sakazakii GCA_000982825.1
231 Enterobacter cloacae subsp. cloacae ATCC 13047 GCA_000025565.1
231 Enterobacter hormaechei subsp. steigerwaltii GCA_001729725.1
231 Enterobacter kobei GCA_900185885.1
231 Erwinia toletana DAPP-PG 735 GCA_000336255.1
231 Klebsiella oxytoca GCA_001022195.1
231 Kosakonia cowanii GCA_001975225.1
231 Kosakonia sacchari SP1 GCA_000300455.4
231 Pseudescherichia vulneris NBRC 102420 GCA_000759795.1
231 Salmonella enterica subsp. enterica serovar Typhi str. CT18 GCA_000195995.1
oI
262 Escherichia coli IAI39 GCA_000026345.1
262 Escherichia coli O104_3AH4 str. 2011C-3493 GCA_000299455.1
262 Escherichia coli O157_3AH7 str. Sakai GCA_000008865.1
262 Escherichia coli UMN026 GCA_000026325.2
262 Escherichia coli str. K-12 substr. MG1655 GCA_000005845.2
262 Shigella flexneri 2a str. 301 GCA_000006925.2
262 Tumebacillus flagellatus GCA_000714935.1
261 Escherichia coli O83_3AH1 str. NRG 857C GCA_000183345.1
253 Shigella dysenteriae Sd197 GCA_000012005.1
oJ
219 Rhodobacter sphaeroides 2.4.1 GCA_000012905.2
207 Pseudorhodobacter ferrugineus DSM 5888 GCA_000420745.1
207 Pseudorhodobacter wandonensis GCA_001202035.1
201 Gemmobacter megaterium GCA_900156815.1
oK
200 Escherichia coli IAI39 GCA_000026345.1
200 Escherichia coli O104_3AH4 str. 2011C-3493 GCA_000299455.1
200 Escherichia coli O157_3AH7 str. Sakai GCA_000008865.1
200 Escherichia coli O83_3AH1 str. NRG 857C GCA_000183345.1
200 Escherichia coli UMN026 GCA_000026325.2
200 Escherichia coli str. K-12 substr. MG1655 GCA_000005845.2
200 Shigella flexneri 2a str. 301 GCA_000006925.2
200 Tumebacillus flagellatus GCA_000714935.1
187 Shigella dysenteriae Sd197 GCA_000012005.1
179 Erwinia iniecta GCA_001267535.1
oL
143 Cronobacter sakazakii GCA_000982825.1
143 Erwinia iniecta GCA_001267535.1
143 Erwinia toletana DAPP-PG 735 GCA_000336255.1

143 Escherichia coli IAI39 GCA_000026345.1
143 Escherichia coli O104_3AH4 str. 2011C-3493 GCA_000299455.1
143 Escherichia coli O157_3AH7 str. Sakai GCA_000008865.1
143 Escherichia coli O83_3AH1 str. NRG 857C GCA_000183345.1
143 Escherichia coli UMN026 GCA_000026325.2
143 Escherichia coli str. K-12 substr. MG1655 GCA_000005845.2
143 Rosenbergiella nectarea GCA_900111105.1
143 Shigella dysenteriae Sd197 GCA_000012005.1
143 Shigella flexneri 2a str. 301 GCA_000006925.2
143 Tumebacillus flagellatus GCA_000714935.1
128 Buttiauxella ferragutiae ATCC 51602 GCA_001654915.1
128 Cedecea neteri GCA_000757825.1
128 Citrobacter freundii CFNIH1 GCA_000648515.1
128 Enterobacter cloacae subsp. cloacae ATCC 13047 GCA_000025565.1
128 Enterobacter hormaechei subsp. steigerwaltii GCA_001729725.1
128 Enterobacter kobei GCA_900185885.1
128 Erwinia persicina NBRC 102418 GCA_001571305.1
128 Klebsiella oxytoca GCA_001022195.1
128 Kluyvera ascorbata ATCC 33433 GCA_000735365.1
128 Kosakonia cowanii GCA_001975225.1
128 Kosakonia sacchari SP1 GCA_000300455.4
128 Mangrovibacter phragmitis GCA_001655675.1
128 Pseudescherichia vulneris NBRC 102420 GCA_000759795.1
128 Salmonella enterica subsp. enterica serovar Typhi str. CT18 GCA_000195995.1
128 Salmonella enterica subsp. enterica serovar Typhimurium str. LT2 GCA_000006945.2
128 Yokenella regensburgei ATCC 49455 GCA_000735455.1
128 _5BEnterobacter_5D lignolyticus SCF1 GCA_000164865.1
125 Klebsiella aerogenes KCTC 2190 GCA_000215745.1
125 Pluralibacter gergoviae GCA_000757785.1
125 Serratia marcescens subsp. marcescens Db11 GCA_000513215.1
oM
198 Escherichia coli IAI39 GCA_000026345.1
198 Escherichia coli O104_3AH4 str. 2011C-3493 GCA_000299455.1
198 Escherichia coli O157_3AH7 str. Sakai GCA_000008865.1
198 Escherichia coli O83_3AH1 str. NRG 857C GCA_000183345.1
198 Escherichia coli UMN026 GCA_000026325.2
198 Escherichia coli str. K-12 substr. MG1655 GCA_000005845.2
198 Shigella dysenteriae Sd197 GCA_000012005.1
198 Shigella flexneri 2a str. 301 GCA_000006925.2
198 Tumebacillus flagellatus GCA_000714935.1
155 Enterobacter hormaechei subsp. steigerwaltii GCA_001729725.1
155 Erwinia billingiae Eb661 GCA_000196615.1
155 Erwinia iniecta GCA_001267535.1
155 Erwinia toletana DAPP-PG 735 GCA_000336255.1
155 Klebsiella oxytoca GCA_001022195.1
155 Kosakonia cowanii GCA_001975225.1
155 Pantoea agglomerans GCA_001709315.1
155 Pantoea alhagi GCA_002101395.1
155 Pantoea ananatis LMG 20103 GCA_000025405.2
155 Pantoea rwandensis GCA_000759475.1
155 Pantoea septica GCA_002095575.1
155 Pluralibacter gergoviae GCA_000757785.1
155 Pseudescherichia vulneris NBRC 102420 GCA_000759795.1
155 Shimwellia blattae DSM 4481 = NBRC 105725 GCA_000262305.1
151 Kosakonia sacchari SP1 GCA_000300455.4
oN
196 Bifidobacterium adolescentis ATCC 15703 GCA_000010425.1
180 Bifidobacterium dentium JCM 1195 = DSM 20436 GCA_001042595.1
177 Bifidobacterium angulatum DSM 20098 = JCM 7096 GCA_001025155.1
oO
236 Escherichia coli IAI39 GCA_000026345.1
236 Escherichia coli O104_3AH4 str. 2011C-3493 GCA_000299455.1
236 Escherichia coli O157_3AH7 str. Sakai GCA_000008865.1
236 Escherichia coli O83_3AH1 str. NRG 857C GCA_000183345.1
236 Escherichia coli UMN026 GCA_000026325.2
236 Escherichia coli str. K-12 substr. MG1655 GCA_000005845.2
236 Shigella dysenteriae Sd197 GCA_000012005.1
236 Shigella flexneri 2a str. 301 GCA_000006925.2
236 Tumebacillus flagellatus GCA_000714935.1
213 Erwinia iniecta GCA_001267535.1
193 Erwinia toletana DAPP-PG 735 GCA_000336255.1
oP
279 Rhodobacter sphaeroides 2.4.1 GCA_000012905.2
255 Rhodobacter sphaeroides ATCC 17025 GCA_000016405.1
247 Pseudorhodobacter psychrotolerans GCA_001294535.1
oQ

178 Rhodobacter sphaeroides 2.4.1 GCA_000012905.2
153 Rhodobacter sphaeroides ATCC 17025 GCA_000016405.1
138 Pseudorhodobacter ferrugineus DSM 5888 GCA_000420745.1
138 Pseudorhodobacter wandonensis GCA_001202035.1
oR
216 Staphylococcus epidermidis ATCC 12228 GCA_000007645.1
216 Staphylococcus haemolyticus JCSC1435 GCA_000009865.1
210 Staphylococcus capitis subsp. capitis GCA_001028645.1
205 Staphylococcus hominis subsp. hominis C80 GCA_000183685.1
205 Staphylococcus lugdunensis HKU09-01 GCA_000025085.1
oS
214 Streptococcus mutans UA159 GCA_000007465.2
136 Streptococcus marimammalium DSM 18627 GCA_000380045.1
118 Streptococcus equinus GCA_000964315.1
118 Streptococcus gallolyticus subsp. gallolyticus DSM 16831 GCA_002000985.1
oT
191 Clostridium beijerinckii GCA_000833105.2
191 Clostridium puniceum GCA_002006345.1
191 Clostridium saccharoperbutylacetonicum N1-4_28HMT_29 GCA_000340885.1
187 Clostridium neonatale GCA_001458595.1
182 Clostridium saccharobutylicum DSM 13864 GCA_000473995.1
oU
214 Lactobacillus gasseri ATCC 33323 = JCM 1131 GCA_000014425.1
206 Lactobacillus hominis DSM 23910 = CRBIP 24.179 GCA_000296835.1
164 Lactobacillus hamsteri DSM 5661 = JCM 6256 GCA_000615445.1
oV
140 Bifidobacterium asteroides PRL2011 GCA_000304215.1
137 Bifidobacterium adolescentis ATCC 15703 GCA_000010425.1
137 Bifidobacterium angulatum DSM 20098 = JCM 7096 GCA_001025155.1
137 Bifidobacterium choerinum GCA_000741135.1
137 Bifidobacterium gallicum DSM 20093 = LMG 11596 GCA_000741205.1
137 Bifidobacterium pseudolongum PV8-2 GCA_000800475.2
137 Bifidobacterium reuteri DSM 23975 GCA_000741695.1
137 Bifidobacterium thermophilum GCA_000741495.1
137 Bifidobacterium tsurumiense GCA_000741765.1
136 Bifidobacterium callitrichos DSM 23973 GCA_000741175.1
oW
191 Escherichia coli UMN026 GCA_000026325.2
191 Shigella flexneri 2a str. 301 GCA_000006925.2
176 Escherichia coli IAI39 GCA_000026345.1
176 Escherichia coli O104_3AH4 str. 2011C-3493 GCA_000299455.1
176 Escherichia coli O157_3AH7 str. Sakai GCA_000008865.1
176 Escherichia coli O83_3AH1 str. NRG 857C GCA_000183345.1
176 Escherichia coli str. K-12 substr. MG1655 GCA_000005845.2
176 Shigella dysenteriae Sd197 GCA_000012005.1
176 Tumebacillus flagellatus GCA_000714935.1
163 Pantoea agglomerans GCA_001709315.1
163 Pantoea ananatis LMG 20103 GCA_000025405.2
163 Shimwellia blattae DSM 4481 = NBRC 105725 GCA_000262305.1
oX
154 Enterococcus faecalis V583 GCA_000007785.1
154 Streptomyces cinnamoneus GCA_001885705.1
132 Streptococcus phocae subsp. salmonis GCA_000772915.1
131 Catellicoccus marimammalium M35_2F04_2F3 GCA_000313915.1
131 Enterococcus aquimarinus GCA_001885765.1
131 Enterococcus asini ATCC 700915 GCA_000407365.1
131 Enterococcus canis NBRC 100695 GCA_001544375.1
131 Enterococcus dispar ATCC 51266 GCA_000406945.1
131 Enterococcus faecium DO GCA_000174395.2
131 Enterococcus gilvus ATCC BAA-350 GCA_000407545.1
131 Enterococcus haemoperoxidus ATCC BAA-382 GCA_000407165.1
131 Enterococcus hermanniensis GCA_001885945.1
131 Enterococcus hirae ATCC 9790 GCA_000271405.2
131 Enterococcus italicus DSM 15952 GCA_000185365.1
131 Enterococcus malodoratus ATCC 43197 GCA_000407185.1
131 Enterococcus massiliensis GCA_001050095.1
131 Enterococcus mundtii QU 25 GCA_000504125.1
131 Enterococcus pallens ATCC BAA-351 GCA_000407485.1
131 Enterococcus phoeniculicola ATCC BAA-412 GCA_000407505.1
131 Enterococcus pseudoavium NBRC 100491 GCA_001544295.1
131 Enterococcus rivorum GCA_001742285.1
131 Enterococcus sulfureus ATCC 49903 GCA_000407605.1
131 Enterococcus thailandicus GCA_001652875.1
131 Melissococcus plutonius S1 GCA_000747585.1
131 Vagococcus fluvialis bH819 GCA_900163795.1
131 Vagococcus lutrae LBD1 GCA_000498295.1

131 Vagococcus teuberi GCA_001870205.1
oY
231 Bifidobacterium adolescentis ATCC 15703 GCA_000010425.1
220 Bifidobacterium tsurumiense GCA_000741765.1
216 Bifidobacterium thermophilum GCA_000741495.1
oZ
279 Lactobacillus gasseri ATCC 33323 = JCM 1131 GCA_000014425.1
274 Lactobacillus hominis DSM 23910 = CRBIP 24.179 GCA_000296835.1
215 Lactobacillus amylovorus GCA_000191545.1
215 Lactobacillus crispatus ST1 GCA_000091765.1
p0
157 Lactobacillus hominis DSM 23910 = CRBIP 24.179 GCA_000296835.1
155 Lactobacillus gasseri ATCC 33323 = JCM 1131 GCA_000014425.1
127 Lactobacillus crispatus ST1 GCA_000091765.1
127 Lactobacillus kefiranofaciens ZW3 GCA_000214785.1
127 Lactobacillus psittaci DSM 15354 GCA_000425905.1
p1
245 Lactobacillus gasseri ATCC 33323 = JCM 1131 GCA_000014425.1
229 Lactobacillus hominis DSM 23910 = CRBIP 24.179 GCA_000296835.1
166 Lactobacillus crispatus ST1 GCA_000091765.1
p2
242 Clostridium beijerinckii GCA_000833105.2
242 Clostridium puniceum GCA_002006345.1
242 Clostridium saccharoperbutylacetonicum N1-4_28HMT_29 GCA_000340885.1
238 Clostridium saccharobutylicum DSM 13864 GCA_000473995.1
227 Clostridium chromiireducens GCA_002029255.1
p3
218 Staphylococcus arlettae CVD059 GCA_000295715.1
218 Staphylococcus cohnii subsp. cohnii GCA_000972575.1
218 Staphylococcus saprophyticus subsp. saprophyticus ATCC 15305 GCA_000010125.1
211 Staphylococcus epidermidis ATCC 12228 GCA_000007645.1
211 Staphylococcus pettenkoferi GCA_002208805.1
209 Staphylococcus succinus GCA_001902315.1
209 Staphylococcus xylosus GCA_000706685.1
p4
225 Streptococcus mutans UA159 GCA_000007465.2
139 Streptococcus macacae NCTC 11558 GCA_000187995.3
123 Streptococcus sobrinus DSM 20742 = ATCC 33478 GCA_000686605.1
p5
267 Deinococcus radiodurans R1 GCA_000008565.1
200 Deinococcus gobiensis I-0 GCA_000252445.1
200 Deinococcus puniceus GCA_001644565.1
187 Deinococcus deserti VCD115 GCA_000020685.1
p6
255 Staphylococcus capitis subsp. capitis GCA_001028645.1
255 Staphylococcus epidermidis ATCC 12228 GCA_000007645.1
244 Staphylococcus haemolyticus JCSC1435 GCA_000009865.1
238 Staphylococcus cohnii subsp. cohnii GCA_000972575.1
p7
211 Rhodobacter sphaeroides 2.4.1 GCA_000012905.2
203 Pseudorhodobacter psychrotolerans GCA_001294535.1
194 Pseudorhodobacter ferrugineus DSM 5888 GCA_000420745.1
194 Pseudorhodobacter wandonensis GCA_001202035.1
p8
252 Enterococcus faecalis V583 GCA_000007785.1
252 Streptomyces cinnamoneus GCA_001885705.1
230 Carnobacterium divergens DSM 20623 GCA_000744255.1
220 Enterococcus asini ATCC 700915 GCA_000407365.1
220 Enterococcus canis NBRC 100695 GCA_001544375.1
220 Enterococcus dispar ATCC 51266 GCA_000406945.1
220 Enterococcus faecium DO GCA_000174395.2
220 Enterococcus gilvus ATCC BAA-350 GCA_000407545.1
220 Enterococcus haemoperoxidus ATCC BAA-382 GCA_000407165.1
220 Enterococcus hermanniensis GCA_001885945.1
220 Enterococcus hirae ATCC 9790 GCA_000271405.2
220 Enterococcus italicus DSM 15952 GCA_000185365.1
220 Enterococcus malodoratus ATCC 43197 GCA_000407185.1
220 Enterococcus mundtii QU 25 GCA_000504125.1
220 Enterococcus pallens ATCC BAA-351 GCA_000407485.1
220 Enterococcus phoeniculicola ATCC BAA-412 GCA_000407505.1
220 Enterococcus pseudoavium NBRC 100491 GCA_001544295.1
220 Enterococcus rivorum GCA_001742285.1
220 Enterococcus sulfureus ATCC 49903 GCA_000407605.1
220 Enterococcus thailandicus GCA_001652875.1
220 Granulicatella adiacens ATCC 49175 GCA_000160675.1
p9

133 Lactobacillus gasseri ATCC 33323 = JCM 1131 GCA_000014425.1
103 Lactobacillus hominis DSM 23910 = CRBIP 24.179 GCA_000296835.1
81 Lactobacillus alimentarius DSM 20249 GCA_001434745.1
pa
199 Clostridium beijerinckii GCA_000833105.2
199 Clostridium butyricum GCA_001456065.2
199 Clostridium chromiireducens GCA_002029255.1
199 Clostridium puniceum GCA_002006345.1
199 Clostridium saccharobutylicum DSM 13864 GCA_000473995.1
199 Clostridium saccharoperbutylacetonicum N1-4_28HMT_29 GCA_000340885.1
171 Clostridium taeniosporum GCA_001735765.1
167 Clostridium cavendishii DSM 21758 GCA_900141845.1
167 Clostridium thermobutyricum DSM 4928 GCA_002050515.1
pb
190 Deinococcus radiodurans R1 GCA_000008565.1
141 Deinococcus gobiensis I-0 GCA_000252445.1
129 Deinococcus frigens DSM 12807 GCA_000701425.1
pc
169 Deinococcus radiodurans R1 GCA_000008565.1
137 Deinococcus deserti VCD115 GCA_000020685.1
121 Deinococcus hopiensis KR-140 GCA_900176165.1
pd
193 Rhodobacter sphaeroides 2.4.1 GCA_000012905.2
152 Rhodobacter sphaeroides ATCC 17025 GCA_000016405.1
152 Thioclava dalianensis GCA_000715505.1
152 Thioclava indica GCA_000714545.1
150 Defluviimonas alba GCA_001620265.1
pe
201 Clostridium beijerinckii GCA_000833105.2
201 Clostridium puniceum GCA_002006345.1
201 Clostridium saccharoperbutylacetonicum N1-4_28HMT_29 GCA_000340885.1
186 Clostridium saccharobutylicum DSM 13864 GCA_000473995.1
180 Clostridium butyricum GCA_001456065.2
pf
211 Lactobacillus gasseri ATCC 33323 = JCM 1131 GCA_000014425.1
211 Lactobacillus hominis DSM 23910 = CRBIP 24.179 GCA_000296835.1
173 Lactobacillus amylophilus DSM 20533 = JCM 1125 GCA_001936335.1
173 Lactobacillus iners DSM 13335 GCA_000160875.1
162 Lactobacillus delbrueckii subsp. bulgaricus ATCC 11842 = JCM 1002 GCA_000056065.1
162 Lactobacillus pasteurii DSM 23907 = CRBIP 24.76 GCA_000297025.1
pg
223 Rhodobacter sphaeroides 2.4.1 GCA_000012905.2
179 Rhodobacter sphaeroides ATCC 17025 GCA_000016405.1
178 Defluviimonas alba GCA_001620265.1
ph
227 Clostridium beijerinckii GCA_000833105.2
227 Clostridium puniceum GCA_002006345.1
227 Clostridium saccharobutylicum DSM 13864 GCA_000473995.1
227 Clostridium saccharoperbutylacetonicum N1-4_28HMT_29 GCA_000340885.1
214 Clostridium butyricum GCA_001456065.2
212 Clostridium neonatale GCA_001458595.1
pi
185 Clostridium beijerinckii GCA_000833105.2
185 Clostridium puniceum GCA_002006345.1
185 Clostridium saccharoperbutylacetonicum N1-4_28HMT_29 GCA_000340885.1
177 Clostridium saccharobutylicum DSM 13864 GCA_000473995.1
166 Clostridium chromiireducens GCA_002029255.1
pj
326 Escherichia coli IAI39 GCA_000026345.1
326 Escherichia coli O104_3AH4 str. 2011C-3493 GCA_000299455.1
326 Escherichia coli O157_3AH7 str. Sakai GCA_000008865.1
326 Escherichia coli O83_3AH1 str. NRG 857C GCA_000183345.1
326 Escherichia coli UMN026 GCA_000026325.2
326 Escherichia coli str. K-12 substr. MG1655 GCA_000005845.2
326 Shigella flexneri 2a str. 301 GCA_000006925.2
326 Tumebacillus flagellatus GCA_000714935.1
317 Shigella dysenteriae Sd197 GCA_000012005.1
303 Erwinia iniecta GCA_001267535.1
pk
226 Streptococcus mutans UA159 GCA_000007465.2
165 Streptococcus sobrinus DSM 20742 = ATCC 33478 GCA_000686605.1
162 Streptococcus cristatus AS 1.3089 GCA_000385925.1
162 Streptococcus parasanguinis ATCC 15912 GCA_000164675.2
pl
236 Deinococcus radiodurans R1 GCA_000008565.1
184 Deinococcus geothermalis DSM 11300 GCA_000196275.1

176 Deinococcus puniceus GCA_001644565.1
pm
217 Streptococcus mutans UA159 GCA_000007465.2
155 Streptococcus ratti FA-1 = DSM 20564 GCA_000286075.1
150 Streptococcus cristatus AS 1.3089 GCA_000385925.1
150 Streptococcus gordonii str. Challis substr. CH1 GCA_000017005.1
150 Streptococcus parasanguinis ATCC 15912 GCA_000164675.2
pn
196 Clostridium beijerinckii GCA_000833105.2
196 Clostridium saccharoperbutylacetonicum N1-4_28HMT_29 GCA_000340885.1
189 Clostridium puniceum GCA_002006345.1
187 Clostridium saccharobutylicum DSM 13864 GCA_000473995.1
po
186 Clostridium beijerinckii GCA_000833105.2
186 Clostridium saccharobutylicum DSM 13864 GCA_000473995.1
186 Clostridium saccharoperbutylacetonicum N1-4_28HMT_29 GCA_000340885.1
171 Clostridium puniceum GCA_002006345.1
165 Clostridium cylindrosporum DSM 605 GCA_001047375.1
pp
301 Streptococcus mutans UA159 GCA_000007465.2
216 Streptococcus macacae NCTC 11558 GCA_000187995.3
210 Streptococcus sobrinus DSM 20742 = ATCC 33478 GCA_000686605.1
pq
160 Escherichia coli IAI39 GCA_000026345.1
160 Escherichia coli O104_3AH4 str. 2011C-3493 GCA_000299455.1
160 Escherichia coli O157_3AH7 str. Sakai GCA_000008865.1
160 Escherichia coli O83_3AH1 str. NRG 857C GCA_000183345.1
160 Escherichia coli UMN026 GCA_000026325.2
160 Escherichia coli str. K-12 substr. MG1655 GCA_000005845.2
160 Shigella dysenteriae Sd197 GCA_000012005.1
160 Shigella flexneri 2a str. 301 GCA_000006925.2
160 Tumebacillus flagellatus GCA_000714935.1
132 Bacillus humi GCA_001439915.1
132 Erwinia iniecta GCA_001267535.1
132 Serratia marcescens subsp. marcescens Db11 GCA_000513215.1
132 Tolumonas auensis DSM 9187 GCA_000023065.1
128 Rosenbergiella nectarea GCA_900111105.1
pr
142 Bifidobacterium adolescentis ATCC 15703 GCA_000010425.1
138 Bifidobacterium tsurumiense GCA_000741765.1
124 Bifidobacterium callitrichos DSM 23973 GCA_000741175.1
ps
241 Clostridium saccharobutylicum DSM 13864 GCA_000473995.1
236 Clostridium beijerinckii GCA_000833105.2
236 Clostridium saccharoperbutylacetonicum N1-4_28HMT_29 GCA_000340885.1
234 Clostridium butyricum GCA_001456065.2
234 Clostridium chromiireducens GCA_002029255.1
pt
113 Clostridium beijerinckii GCA_000833105.2
113 Clostridium chromiireducens GCA_002029255.1
113 Clostridium puniceum GCA_002006345.1
113 Clostridium saccharobutylicum DSM 13864 GCA_000473995.1
113 Clostridium saccharoperbutylacetonicum N1-4_28HMT_29 GCA_000340885.1
97 Clostridium sulfidigenes GCA_000732635.1
93 Clostridium butyricum GCA_001456065.2
pu
238 Streptococcus mutans UA159 GCA_000007465.2
176 Streptococcus ferus DSM 20646 GCA_000372425.1
166 Streptococcus macacae NCTC 11558 GCA_000187995.3
pv
211 Deinococcus radiodurans R1 GCA_000008565.1
140 Deinococcus puniceus GCA_001644565.1
135 Deinococcus gobiensis I-0 GCA_000252445.1
pw
193 Bifidobacterium adolescentis ATCC 15703 GCA_000010425.1
174 Bifidobacterium choerinum GCA_000741135.1
174 Bifidobacterium pseudolongum PV8-2 GCA_000800475.2
173 Bifidobacterium breve DSM 20213 = JCM 1192 GCA_001025175.1
173 Bifidobacterium longum NCC2705 GCA_000007525.1
173 Bifidobacterium reuteri DSM 23975 GCA_000741695.1
173 Bifidobacterium saguini DSM 23967 GCA_000741715.1
px
299 Escherichia coli IAI39 GCA_000026345.1
299 Escherichia coli O104_3AH4 str. 2011C-3493 GCA_000299455.1
299 Escherichia coli O157_3AH7 str. Sakai GCA_000008865.1
299 Escherichia coli O83_3AH1 str. NRG 857C GCA_000183345.1

299 Escherichia coli UMN026 GCA_000026325.2
299 Escherichia coli str. K-12 substr. MG1655 GCA_000005845.2
299 Shigella dysenteriae Sd197 GCA_000012005.1
299 Tumebacillus flagellatus GCA_000714935.1
293 Shigella flexneri 2a str. 301 GCA_000006925.2
261 Erwinia iniecta GCA_001267535.1
py
224 Streptococcus mutans UA159 GCA_000007465.2
127 Streptococcus agalactiae 2603V_2FR GCA_000007265.1
127 Streptococcus cristatus AS 1.3089 GCA_000385925.1
127 Streptococcus dysgalactiae subsp. equisimilis AC-2713 GCA_000317855.1
127 Streptococcus gordonii str. Challis substr. CH1 GCA_000017005.1
127 Streptococcus ictaluri 707-05 GCA_000188015.3
127 Streptococcus mitis B6 GCA_000027165.1
127 Streptococcus pneumoniae R6 GCA_000007045.1
127 Streptococcus sobrinus DSM 20742 = ATCC 33478 GCA_000686605.1
121 Streptococcus parasanguinis ATCC 15912 GCA_000164675.2
pz
289 Lactobacillus gasseri ATCC 33323 = JCM 1131 GCA_000014425.1
279 Lactobacillus hominis DSM 23910 = CRBIP 24.179 GCA_000296835.1
186 Lactobacillus crispatus ST1 GCA_000091765.1
186 Lactobacillus helveticus GCA_001308285.1
pA
164 Bifidobacterium adolescentis ATCC 15703 GCA_000010425.1
161 Bifidobacterium callitrichos DSM 23973 GCA_000741175.1
161 Bifidobacterium coryneforme GCA_000737865.1
150 Bifidobacterium tsurumiense GCA_000741765.1
pB
186 Staphylococcus capitis subsp. capitis GCA_001028645.1
186 Staphylococcus epidermidis ATCC 12228 GCA_000007645.1
175 Staphylococcus cohnii subsp. cohnii GCA_000972575.1
171 Staphylococcus haemolyticus JCSC1435 GCA_000009865.1
pC
226 Clostridium beijerinckii GCA_000833105.2
226 Clostridium puniceum GCA_002006345.1
226 Clostridium saccharoperbutylacetonicum N1-4_28HMT_29 GCA_000340885.1
217 Clostridium saccharobutylicum DSM 13864 GCA_000473995.1
213 Clostridium chromiireducens GCA_002029255.1
pD
205 Clostridium beijerinckii GCA_000833105.2
205 Clostridium saccharobutylicum DSM 13864 GCA_000473995.1
205 Clostridium saccharoperbutylacetonicum N1-4_28HMT_29 GCA_000340885.1
201 Clostridium puniceum GCA_002006345.1
194 Clostridium butyricum GCA_001456065.2
pE
147 Rhodobacter sphaeroides 2.4.1 GCA_000012905.2
132 Rhodobacter sphaeroides ATCC 17025 GCA_000016405.1
116 Pseudorhodobacter ferrugineus DSM 5888 GCA_000420745.1
116 Pseudorhodobacter wandonensis GCA_001202035.1
pF
156 Rhodobacter sphaeroides 2.4.1 GCA_000012905.2
128 Pseudorhodobacter psychrotolerans GCA_001294535.1
125 Rhodobacter sphaeroides ATCC 17025 GCA_000016405.1
pG
234 Escherichia coli IAI39 GCA_000026345.1
234 Escherichia coli O104_3AH4 str. 2011C-3493 GCA_000299455.1
234 Escherichia coli O157_3AH7 str. Sakai GCA_000008865.1
234 Escherichia coli O83_3AH1 str. NRG 857C GCA_000183345.1
234 Escherichia coli UMN026 GCA_000026325.2
234 Escherichia coli str. K-12 substr. MG1655 GCA_000005845.2
234 Shigella dysenteriae Sd197 GCA_000012005.1
234 Shigella flexneri 2a str. 301 GCA_000006925.2
234 Tumebacillus flagellatus GCA_000714935.1
212 Erwinia iniecta GCA_001267535.1
206 Erwinia toletana DAPP-PG 735 GCA_000336255.1
206 Pantoea agglomerans GCA_001709315.1
206 Pantoea ananatis LMG 20103 GCA_000025405.2
206 Shimwellia blattae DSM 4481 = NBRC 105725 GCA_000262305.1
pH
191 Streptococcus mutans UA159 GCA_000007465.2
96 Streptococcus massiliensis DSM 18628 GCA_000380065.1
94 Streptococcus cristatus AS 1.3089 GCA_000385925.1
94 Streptococcus ferus DSM 20646 GCA_000372425.1
94 Streptococcus gordonii str. Challis substr. CH1 GCA_000017005.1
94 Streptococcus mitis B6 GCA_000027165.1
94 Streptococcus parasanguinis ATCC 15912 GCA_000164675.2

94 Streptococcus pneumoniae R6 GCA_000007045.1
pI
295 Lactobacillus gasseri ATCC 33323 = JCM 1131 GCA_000014425.1
263 Lactobacillus hominis DSM 23910 = CRBIP 24.179 GCA_000296835.1
221 Lactobacillus psittaci DSM 15354 GCA_000425905.1
pJ
250 Clostridium beijerinckii GCA_000833105.2
250 Clostridium puniceum GCA_002006345.1
250 Clostridium saccharoperbutylacetonicum N1-4_28HMT_29 GCA_000340885.1
241 Clostridium chromiireducens GCA_002029255.1
241 Clostridium saccharobutylicum DSM 13864 GCA_000473995.1
228 Clostridium taeniosporum GCA_001735765.1
pK
327 Bacillus anthracis str. Ames GCA_000007845.1
327 Bacillus anthracis str. Sterne GCA_000008165.1
327 Bacillus cereus ATCC 14579 GCA_000007825.1
327 Bacillus mycoides GCA_000832605.1
327 Bacillus pseudomycoides DSM 12442 GCA_000161455.1
327 Bacillus thuringiensis YBT-1518 GCA_000497525.2
327 _5BBacillus thuringiensis_5D serovar konkukian str. 97-27 GCA_000008505.1
269 Bacillus marisflavi GCA_001274775.1
268 Bacillus manliponensis GCA_000712595.1
pL
310 Streptococcus mutans UA159 GCA_000007465.2
201 Streptococcus equinus GCA_000964315.1
201 Streptococcus gallolyticus subsp. gallolyticus DSM 16831 GCA_002000985.1
197 Streptococcus gordonii str. Challis substr. CH1 GCA_000017005.1
pM
197 Lactobacillus gasseri ATCC 33323 = JCM 1131 GCA_000014425.1
182 Lactobacillus hominis DSM 23910 = CRBIP 24.179 GCA_000296835.1
153 Lactobacillus acetotolerans GCA_001042405.1
153 Lactobacillus amylovorus GCA_000191545.1
153 Lactobacillus crispatus ST1 GCA_000091765.1
153 Lactobacillus helveticus GCA_001308285.1
153 Lactobacillus kalixensis DSM 16043 GCA_001434335.1
153 Lactobacillus kullabergensis GCA_000967195.1
153 Lactobacillus psittaci DSM 15354 GCA_000425905.1
pN
111 Lactobacillus gasseri ATCC 33323 = JCM 1131 GCA_000014425.1
94 Lactobacillus hominis DSM 23910 = CRBIP 24.179 GCA_000296835.1
79 Lactobacillus acidophilus NCFM GCA_000011985.1
79 Lactobacillus amylolyticus GCA_002075105.1
79 Lactobacillus amylovorus GCA_000191545.1
79 Lactobacillus crispatus ST1 GCA_000091765.1
79 Lactobacillus hamsteri DSM 5661 = JCM 6256 GCA_000615445.1
79 Lactobacillus jensenii GCA_001936235.1
79 Lactobacillus kefiranofaciens ZW3 GCA_000214785.1
79 Lactobacillus psittaci DSM 15354 GCA_000425905.1
pO
217 Clostridium beijerinckii GCA_000833105.2
217 Clostridium puniceum GCA_002006345.1
217 Clostridium saccharobutylicum DSM 13864 GCA_000473995.1
217 Clostridium saccharoperbutylacetonicum N1-4_28HMT_29 GCA_000340885.1
203 Clostridium chromiireducens GCA_002029255.1
202 Clostridium butyricum GCA_001456065.2
pP
161 Deinococcus radiodurans R1 GCA_000008565.1
109 Deinococcus gobiensis I-0 GCA_000252445.1
108 Deinococcus puniceus GCA_001644565.1
pQ
244 Staphylococcus aureus subsp. aureus NCTC 8325 GCA_000013425.1
244 Staphylococcus epidermidis ATCC 12228 GCA_000007645.1
244 Staphylococcus haemolyticus JCSC1435 GCA_000009865.1
244 Staphylococcus hominis subsp. hominis C80 GCA_000183685.1
244 Staphylococcus lugdunensis HKU09-01 GCA_000025085.1
244 Staphylococcus simiae CCM 7213 GCA_000235645.2
240 Staphylococcus arlettae CVD059 GCA_000295715.1
240 Staphylococcus cohnii subsp. cohnii GCA_000972575.1
240 Staphylococcus equorum GCA_001432245.1
240 Staphylococcus gallinarum GCA_000875895.1
240 Staphylococcus saprophyticus subsp. saprophyticus ATCC 15305 GCA_000010125.1
240 Staphylococcus succinus GCA_001902315.1
240 Staphylococcus xylosus GCA_000706685.1
238 Staphylococcus capitis subsp. capitis GCA_001028645.1
pR
236 Streptococcus mutans UA159 GCA_000007465.2

160 Streptococcus sobrinus DSM 20742 = ATCC 33478 GCA_000686605.1
146 Streptococcus ferus DSM 20646 GCA_000372425.1
146 Streptococcus iniae GCA_000831485.1
146 Streptococcus macacae NCTC 11558 GCA_000187995.3
pS
212 Deinococcus radiodurans R1 GCA_000008565.1
147 Deinococcus gobiensis I-0 GCA_000252445.1
138 Deinococcus deserti VCD115 GCA_000020685.1
pT
236 Clostridium beijerinckii GCA_000833105.2
236 Clostridium saccharoperbutylacetonicum N1-4_28HMT_29 GCA_000340885.1
228 Clostridium saccharobutylicum DSM 13864 GCA_000473995.1
222 Clostridium puniceum GCA_002006345.1
pU
264 Escherichia coli IAI39 GCA_000026345.1
264 Escherichia coli O104_3AH4 str. 2011C-3493 GCA_000299455.1
264 Escherichia coli O157_3AH7 str. Sakai GCA_000008865.1
264 Escherichia coli O83_3AH1 str. NRG 857C GCA_000183345.1
264 Escherichia coli UMN026 GCA_000026325.2
264 Escherichia coli str. K-12 substr. MG1655 GCA_000005845.2
264 Shigella flexneri 2a str. 301 GCA_000006925.2
264 Tumebacillus flagellatus GCA_000714935.1
258 Erwinia iniecta GCA_001267535.1
255 Shigella dysenteriae Sd197 GCA_000012005.1
pV
255 Bacillus thuringiensis YBT-1518 GCA_000497525.2
234 Bacillus anthracis str. Ames GCA_000007845.1
234 Bacillus anthracis str. Sterne GCA_000008165.1
234 Bacillus cereus ATCC 14579 GCA_000007825.1
234 Bacillus mycoides GCA_000832605.1
234 Bacillus pseudomycoides DSM 12442 GCA_000161455.1
234 _5BBacillus thuringiensis_5D serovar konkukian str. 97-27 GCA_000008505.1
219 Bacillus flexus GCA_002024265.1
219 Bacillus megaterium NBRC 15308 = ATCC 14581 GCA_000832985.1
pW
142 Clostridium beijerinckii GCA_000833105.2
142 Clostridium puniceum GCA_002006345.1
142 Clostridium saccharoperbutylacetonicum N1-4_28HMT_29 GCA_000340885.1
138 Clostridium chromiireducens GCA_002029255.1
138 Clostridium saccharobutylicum DSM 13864 GCA_000473995.1
133 Clostridium butyricum GCA_001456065.2
pX
247 Deinococcus radiodurans R1 GCA_000008565.1
188 Deinococcus deserti VCD115 GCA_000020685.1
175 Deinococcus soli Cha et al. 2016 GCA_001007995.1
pY
216 Escherichia coli IAI39 GCA_000026345.1
216 Escherichia coli O104_3AH4 str. 2011C-3493 GCA_000299455.1
216 Escherichia coli O157_3AH7 str. Sakai GCA_000008865.1
216 Escherichia coli O83_3AH1 str. NRG 857C GCA_000183345.1
216 Escherichia coli UMN026 GCA_000026325.2
216 Escherichia coli str. K-12 substr. MG1655 GCA_000005845.2
216 Shigella dysenteriae Sd197 GCA_000012005.1
216 Shigella flexneri 2a str. 301 GCA_000006925.2
216 Tumebacillus flagellatus GCA_000714935.1
210 Erwinia iniecta GCA_001267535.1
190 Cedecea neteri GCA_000757825.1
190 Rosenbergiella nectarea GCA_900111105.1
pZ
116 Bifidobacterium adolescentis ATCC 15703 GCA_000010425.1
101 Bifidobacterium thermophilum GCA_000741495.1
101 Bifidobacterium thermophilum RBL67 GCA_000347695.1
100 Bifidobacterium callitrichos DSM 23973 GCA_000741175.1
100 Bifidobacterium coryneforme GCA_000737865.1
q0
72 Enterococcus canis NBRC 100695 GCA_001544375.1
72 Enterococcus casseliflavus EC20 GCA_000157355.2
72 Enterococcus dispar ATCC 51266 GCA_000406945.1
72 Enterococcus faecalis V583 GCA_000007785.1
72 Enterococcus faecium DO GCA_000174395.2
72 Enterococcus gilvus ATCC BAA-350 GCA_000407545.1
72 Enterococcus haemoperoxidus ATCC BAA-382 GCA_000407165.1
72 Enterococcus hermanniensis GCA_001885945.1
72 Enterococcus hirae ATCC 9790 GCA_000271405.2
72 Enterococcus malodoratus ATCC 43197 GCA_000407185.1
72 Enterococcus mundtii QU 25 GCA_000504125.1

72 Enterococcus pallens ATCC BAA-351 GCA_000407485.1
72 Enterococcus phoeniculicola ATCC BAA-412 GCA_000407505.1
72 Enterococcus pseudoavium NBRC 100491 GCA_001544295.1
72 Enterococcus rivorum GCA_001742285.1
72 Enterococcus saccharolyticus subsp. saccharolyticus ATCC 43076 GCA_000407285.1
72 Enterococcus thailandicus GCA_001652875.1
72 Listeria weihenstephanensis FSL R9-0317 GCA_000525955.1
72 Streptomyces cinnamoneus GCA_001885705.1
59 Anaerobacillus arseniciselenatis GCA_001865995.1
59 Bacillus glycinifermentans GCA_900093775.1
59 Bacillus indicus GCA_000709935.2
59 Bacillus licheniformis DSM 13 = ATCC 14580 GCA_000011645.1
57 Streptococcus equinus GCA_000964315.1
57 Streptococcus gallolyticus subsp. gallolyticus DSM 16831 GCA_002000985.1
57 Streptococcus orisratti DSM 15617 GCA_000380105.1
57 Streptococcus ratti FA-1 = DSM 20564 GCA_000286075.1
q1
221 Clostridium beijerinckii GCA_000833105.2
221 Clostridium saccharoperbutylacetonicum N1-4_28HMT_29 GCA_000340885.1
212 Clostridium saccharobutylicum DSM 13864 GCA_000473995.1
207 Clostridium butyricum GCA_001456065.2
207 Clostridium puniceum GCA_002006345.1
q2
203 Deinococcus radiodurans R1 GCA_000008565.1
146 Deinococcus deserti VCD115 GCA_000020685.1
143 Deinococcus gobiensis I-0 GCA_000252445.1
q3
292 Escherichia coli IAI39 GCA_000026345.1
292 Escherichia coli O104_3AH4 str. 2011C-3493 GCA_000299455.1
292 Escherichia coli O157_3AH7 str. Sakai GCA_000008865.1
292 Escherichia coli O83_3AH1 str. NRG 857C GCA_000183345.1
292 Escherichia coli UMN026 GCA_000026325.2
292 Escherichia coli str. K-12 substr. MG1655 GCA_000005845.2
292 Shigella dysenteriae Sd197 GCA_000012005.1
292 Shigella flexneri 2a str. 301 GCA_000006925.2
292 Tumebacillus flagellatus GCA_000714935.1
272 Erwinia billingiae Eb661 GCA_000196615.1
266 Erwinia iniecta GCA_001267535.1
266 Erwinia toletana DAPP-PG 735 GCA_000336255.1
q4
209 Deinococcus radiodurans R1 GCA_000008565.1
147 Deinococcus gobiensis I-0 GCA_000252445.1
132 Deinococcus puniceus GCA_001644565.1
q5
221 Enterococcus hirae ATCC 9790 GCA_000271405.2
218 Enterococcus faecalis V583 GCA_000007785.1
218 Streptomyces cinnamoneus GCA_001885705.1
203 Enterococcus asini ATCC 700915 GCA_000407365.1
203 Enterococcus canis NBRC 100695 GCA_001544375.1
203 Enterococcus casseliflavus EC20 GCA_000157355.2
203 Enterococcus dispar ATCC 51266 GCA_000406945.1
203 Enterococcus faecium DO GCA_000174395.2
203 Enterococcus haemoperoxidus ATCC BAA-382 GCA_000407165.1
203 Enterococcus massiliensis GCA_001050095.1
203 Enterococcus mundtii QU 25 GCA_000504125.1
203 Enterococcus phoeniculicola ATCC BAA-412 GCA_000407505.1
203 Enterococcus rivorum GCA_001742285.1
203 Enterococcus saccharolyticus subsp. saccharolyticus ATCC 43076 GCA_000407285.1
203 Enterococcus thailandicus GCA_001652875.1
q6
255 Rhodobacter sphaeroides 2.4.1 GCA_000012905.2
237 Rhodobacter sphaeroides ATCC 17025 GCA_000016405.1
223 Gemmobacter megaterium GCA_900156815.1
223 Pseudorhodobacter psychrotolerans GCA_001294535.1
q7
190 Staphylococcus cohnii subsp. cohnii GCA_000972575.1
188 Staphylococcus capitis subsp. capitis GCA_001028645.1
188 Staphylococcus epidermidis ATCC 12228 GCA_000007645.1
185 Staphylococcus haemolyticus JCSC1435 GCA_000009865.1
q8
234 Bifidobacterium adolescentis ATCC 15703 GCA_000010425.1
225 Bifidobacterium dentium JCM 1195 = DSM 20436 GCA_001042595.1
225 Bifidobacterium tsurumiense GCA_000741765.1
216 Bifidobacterium thermophilum GCA_000741495.1
216 Bifidobacterium thermophilum RBL67 GCA_000347695.1
q9

248 Deinococcus radiodurans R1 GCA_000008565.1
169 Deinococcus gobiensis I-0 GCA_000252445.1
148 Deinococcus deserti VCD115 GCA_000020685.1
qa
169 Rhodobacter sphaeroides 2.4.1 GCA_000012905.2
149 Rhodobacter sphaeroides ATCC 17025 GCA_000016405.1
136 Rhodobacter capsulatus SB 1003 GCA_000021865.1
qb
280 Enterococcus faecalis V583 GCA_000007785.1
280 Streptomyces cinnamoneus GCA_001885705.1
233 Enterococcus canis NBRC 100695 GCA_001544375.1
233 Enterococcus casseliflavus EC20 GCA_000157355.2
233 Enterococcus dispar ATCC 51266 GCA_000406945.1
233 Enterococcus faecium DO GCA_000174395.2
233 Enterococcus hirae ATCC 9790 GCA_000271405.2
233 Enterococcus mundtii QU 25 GCA_000504125.1
233 Enterococcus rivorum GCA_001742285.1
233 Enterococcus saccharolyticus subsp. saccharolyticus ATCC 43076 GCA_000407285.1
223 Enterococcus asini ATCC 700915 GCA_000407365.1
qc
239 Deinococcus radiodurans R1 GCA_000008565.1
159 Deinococcus gobiensis I-0 GCA_000252445.1
157 Deinococcus puniceus GCA_001644565.1
qd
175 Rhodobacter sphaeroides 2.4.1 GCA_000012905.2
154 Defluviimonas alba GCA_001620265.1
149 Haematobacter massiliensis GCA_000740795.1
149 Pseudoruegeria lutimaris GCA_900099935.1
qe
276 Deinococcus radiodurans R1 GCA_000008565.1
183 Deinococcus gobiensis I-0 GCA_000252445.1
175 Deinococcus marmoris DSM 12784 GCA_000701405.1
175 Deinococcus swuensis GCA_000800395.1
qf
210 Escherichia coli IAI39 GCA_000026345.1
210 Escherichia coli O104_3AH4 str. 2011C-3493 GCA_000299455.1
210 Escherichia coli O157_3AH7 str. Sakai GCA_000008865.1
210 Escherichia coli O83_3AH1 str. NRG 857C GCA_000183345.1
210 Escherichia coli UMN026 GCA_000026325.2
210 Escherichia coli str. K-12 substr. MG1655 GCA_000005845.2
210 Shigella dysenteriae Sd197 GCA_000012005.1
210 Shigella flexneri 2a str. 301 GCA_000006925.2
210 Tumebacillus flagellatus GCA_000714935.1
197 Lonsdalea quercina subsp. quercina GCA_000688655.1
195 Erwinia iniecta GCA_001267535.1
195 Erwinia toletana DAPP-PG 735 GCA_000336255.1
195 Pantoea agglomerans GCA_001709315.1
195 Pantoea ananatis LMG 20103 GCA_000025405.2
195 Shimwellia blattae DSM 4481 = NBRC 105725 GCA_000262305.1
qg
137 Escherichia coli IAI39 GCA_000026345.1
137 Escherichia coli O104_3AH4 str. 2011C-3493 GCA_000299455.1
137 Escherichia coli O157_3AH7 str. Sakai GCA_000008865.1
137 Escherichia coli O83_3AH1 str. NRG 857C GCA_000183345.1
137 Escherichia coli UMN026 GCA_000026325.2
137 Escherichia coli str. K-12 substr. MG1655 GCA_000005845.2
137 Shigella flexneri 2a str. 301 GCA_000006925.2
137 Tumebacillus flagellatus GCA_000714935.1
128 Shigella dysenteriae Sd197 GCA_000012005.1
104 Erwinia iniecta GCA_001267535.1
qh
252 Streptococcus mutans UA159 GCA_000007465.2
137 Streptococcus iniae GCA_000831485.1
137 Streptococcus macacae NCTC 11558 GCA_000187995.3
129 Streptococcus sobrinus DSM 20742 = ATCC 33478 GCA_000686605.1
qi
222 Clostridium beijerinckii GCA_000833105.2
222 Clostridium chromiireducens GCA_002029255.1
222 Clostridium puniceum GCA_002006345.1
222 Clostridium saccharobutylicum DSM 13864 GCA_000473995.1
222 Clostridium saccharoperbutylacetonicum N1-4_28HMT_29 GCA_000340885.1
222 Clostridium taeniosporum GCA_001735765.1
217 Clostridium butyricum GCA_001456065.2
211 Clostridium botulinum B str. Eklund 17B _28NRP_29 GCA_000020165.1
211 Clostridium uliginosum GCA_900112485.1
qj

209 Clostridium beijerinckii GCA_000833105.2
209 Clostridium puniceum GCA_002006345.1
209 Clostridium saccharoperbutylacetonicum N1-4_28HMT_29 GCA_000340885.1
201 Clostridium saccharobutylicum DSM 13864 GCA_000473995.1
186 Clostridium chromiireducens GCA_002029255.1
qk
168 Staphylococcus capitis subsp. capitis GCA_001028645.1
168 Staphylococcus epidermidis ATCC 12228 GCA_000007645.1
164 Staphylococcus cohnii subsp. cohnii GCA_000972575.1
163 Megasphaera cerevisiae DSM 20462 GCA_001045675.1
163 Staphylococcus warneri SG1 GCA_000332735.1
ql
236 Bifidobacterium adolescentis ATCC 15703 GCA_000010425.1
221 Bifidobacterium thermophilum GCA_000741495.1
214 Bifidobacterium thermophilum RBL67 GCA_000347695.1
qm
218 Rhodobacter sphaeroides 2.4.1 GCA_000012905.2
179 Rhodobacter sphaeroides ATCC 17025 GCA_000016405.1
177 Defluviimonas alba GCA_001620265.1
qn
252 Clostridium beijerinckii GCA_000833105.2
252 Clostridium saccharoperbutylacetonicum N1-4_28HMT_29 GCA_000340885.1
248 Clostridium saccharobutylicum DSM 13864 GCA_000473995.1
246 Clostridium puniceum GCA_002006345.1
qo
220 Clostridium beijerinckii GCA_000833105.2
220 Clostridium saccharoperbutylacetonicum N1-4_28HMT_29 GCA_000340885.1
216 Clostridium puniceum GCA_002006345.1
201 Clostridium saccharobutylicum DSM 13864 GCA_000473995.1
qp
157 Streptococcus mutans UA159 GCA_000007465.2
77 Streptococcus sobrinus DSM 20742 = ATCC 33478 GCA_000686605.1
71 Streptococcus ratti FA-1 = DSM 20564 GCA_000286075.1
qq
222 Enterococcus faecalis V583 GCA_000007785.1
222 Streptomyces cinnamoneus GCA_001885705.1
207 Enterococcus canis NBRC 100695 GCA_001544375.1
207 Enterococcus faecium DO GCA_000174395.2
207 Enterococcus hirae ATCC 9790 GCA_000271405.2
207 Enterococcus mundtii QU 25 GCA_000504125.1
207 Enterococcus rivorum GCA_001742285.1
206 Enterococcus dispar ATCC 51266 GCA_000406945.1
206 Enterococcus massiliensis GCA_001050095.1
qr
171 Staphylococcus aureus subsp. aureus NCTC 8325 GCA_000013425.1
171 Staphylococcus capitis subsp. capitis GCA_001028645.1
171 Staphylococcus epidermidis ATCC 12228 GCA_000007645.1
171 Staphylococcus haemolyticus JCSC1435 GCA_000009865.1
171 Staphylococcus hominis subsp. hominis C80 GCA_000183685.1
171 Staphylococcus lugdunensis HKU09-01 GCA_000025085.1
171 Staphylococcus simiae CCM 7213 GCA_000235645.2
161 Megasphaera cerevisiae DSM 20462 GCA_001045675.1
161 Staphylococcus warneri SG1 GCA_000332735.1
158 Staphylococcus arlettae CVD059 GCA_000295715.1
158 Staphylococcus cohnii subsp. cohnii GCA_000972575.1
158 Staphylococcus equorum GCA_001432245.1
158 Staphylococcus gallinarum GCA_000875895.1
158 Staphylococcus saprophyticus subsp. saprophyticus ATCC 15305 GCA_000010125.1
158 Staphylococcus succinus GCA_001902315.1
158 Staphylococcus xylosus GCA_000706685.1
qs
200 Escherichia coli IAI39 GCA_000026345.1
200 Escherichia coli O104_3AH4 str. 2011C-3493 GCA_000299455.1
200 Escherichia coli O157_3AH7 str. Sakai GCA_000008865.1
200 Escherichia coli UMN026 GCA_000026325.2
200 Escherichia coli str. K-12 substr. MG1655 GCA_000005845.2
200 Shigella dysenteriae Sd197 GCA_000012005.1
200 Shigella flexneri 2a str. 301 GCA_000006925.2
200 Tumebacillus flagellatus GCA_000714935.1
199 Escherichia coli O83_3AH1 str. NRG 857C GCA_000183345.1
176 Enterobacter hormaechei subsp. steigerwaltii GCA_001729725.1
176 Erwinia billingiae Eb661 GCA_000196615.1
176 Erwinia iniecta GCA_001267535.1
176 Erwinia toletana DAPP-PG 735 GCA_000336255.1
176 Klebsiella oxytoca GCA_001022195.1
176 Kosakonia cowanii GCA_001975225.1

176 Pantoea agglomerans GCA_001709315.1
176 Pantoea alhagi GCA_002101395.1
176 Pantoea ananatis LMG 20103 GCA_000025405.2
176 Pantoea rwandensis GCA_000759475.1
176 Pantoea septica GCA_002095575.1
176 Pluralibacter gergoviae GCA_000757785.1
176 Pseudescherichia vulneris NBRC 102420 GCA_000759795.1
176 Shimwellia blattae DSM 4481 = NBRC 105725 GCA_000262305.1
qt
245 Deinococcus radiodurans R1 GCA_000008565.1
168 Deinococcus puniceus GCA_001644565.1
167 Deinococcus gobiensis I-0 GCA_000252445.1
qu
171 Clostridium beijerinckii GCA_000833105.2
171 Clostridium saccharobutylicum DSM 13864 GCA_000473995.1
171 Clostridium saccharoperbutylacetonicum N1-4_28HMT_29 GCA_000340885.1
167 Clostridium puniceum GCA_002006345.1
164 Clostridium taeniosporum GCA_001735765.1
qv
259 Bacillus thuringiensis YBT-1518 GCA_000497525.2
247 Bacillus anthracis str. Ames GCA_000007845.1
247 Bacillus anthracis str. Sterne GCA_000008165.1
247 Bacillus cereus ATCC 14579 GCA_000007825.1
247 Bacillus mycoides GCA_000832605.1
247 Bacillus pseudomycoides DSM 12442 GCA_000161455.1
247 _5BBacillus thuringiensis_5D serovar konkukian str. 97-27 GCA_000008505.1
201 Bacillus manliponensis GCA_000712595.1
qw
214 Escherichia coli IAI39 GCA_000026345.1
214 Escherichia coli O104_3AH4 str. 2011C-3493 GCA_000299455.1
214 Escherichia coli O157_3AH7 str. Sakai GCA_000008865.1
214 Escherichia coli O83_3AH1 str. NRG 857C GCA_000183345.1
214 Escherichia coli UMN026 GCA_000026325.2
214 Escherichia coli str. K-12 substr. MG1655 GCA_000005845.2
214 Shigella dysenteriae Sd197 GCA_000012005.1
214 Shigella flexneri 2a str. 301 GCA_000006925.2
214 Tumebacillus flagellatus GCA_000714935.1
184 Erwinia iniecta GCA_001267535.1
184 Rosenbergiella nectarea GCA_900111105.1
175 Cedecea neteri GCA_000757825.1
qx
106 Escherichia coli IAI39 GCA_000026345.1
106 Escherichia coli O104_3AH4 str. 2011C-3493 GCA_000299455.1
106 Escherichia coli O157_3AH7 str. Sakai GCA_000008865.1
106 Escherichia coli O83_3AH1 str. NRG 857C GCA_000183345.1
106 Escherichia coli UMN026 GCA_000026325.2
106 Escherichia coli str. K-12 substr. MG1655 GCA_000005845.2
106 Shigella dysenteriae Sd197 GCA_000012005.1
106 Shigella flexneri 2a str. 301 GCA_000006925.2
106 Tumebacillus flagellatus GCA_000714935.1
77 Cronobacter sakazakii GCA_000982825.1
77 Erwinia iniecta GCA_001267535.1
77 Erwinia toletana DAPP-PG 735 GCA_000336255.1
77 Rosenbergiella nectarea GCA_900111105.1
74 Pantoea agglomerans GCA_001709315.1
qy
340 Escherichia coli IAI39 GCA_000026345.1
340 Escherichia coli O104_3AH4 str. 2011C-3493 GCA_000299455.1
340 Escherichia coli O157_3AH7 str. Sakai GCA_000008865.1
340 Escherichia coli O83_3AH1 str. NRG 857C GCA_000183345.1
340 Escherichia coli UMN026 GCA_000026325.2
340 Escherichia coli str. K-12 substr. MG1655 GCA_000005845.2
340 Shigella flexneri 2a str. 301 GCA_000006925.2
340 Tumebacillus flagellatus GCA_000714935.1
331 Shigella dysenteriae Sd197 GCA_000012005.1
330 Erwinia iniecta GCA_001267535.1
qz
91 Roseovarius aestuarii GCA_900172285.1
91 Sedimentitalea nanhaiensis DSM 24252 GCA_000473225.1
91 Sulfitobacter guttiformis KCTC 32187 GCA_000622425.1
89 Aliiroseovarius crassostreae GCA_001307765.1
89 Aliiroseovarius sediminilitoris GCA_900109955.1
89 Antarctobacter heliothermus GCA_900188425.1
89 Aquimixticola soesokkakensis GCA_900172375.1
89 Citreicella marina GCA_900100085.1
89 Citreicella thiooxidans GCA_900102075.1

89 Citreimonas salinaria GCA_900107235.1
89 Donghicola eburneus GCA_900115865.1
89 Gemmobacter megaterium GCA_900156815.1
89 Ketogulonicigenium robustum GCA_002117445.1
89 Loktanella cinnabarina LL-001 GCA_000466965.1
89 Loktanella hongkongensis DSM 17492 GCA_000600975.2
89 Loktanella pyoseonensis GCA_900102015.1
89 Loktanella soesokkakensis GCA_900172345.1
89 Maribius pelagius GCA_900110115.1
89 Maritimibacter alkaliphilus HTCC2654 GCA_000152805.1
89 Marivita cryptomonadis GCA_002115725.1
89 Marivita hallyeonensis GCA_900129875.1
89 Nioella sediminis GCA_001879695.1
89 Oceanicola granulosus HTCC2516 GCA_000153305.1
89 Pelagibaca abyssi GCA_001975705.1
89 Poseidonocella pacifica GCA_900111875.1
89 Pseudooceanicola atlanticus GCA_000768315.1
89 Pseudooceanicola batsensis HTCC2597 GCA_000152725.1
89 Pseudooceanicola marinus GCA_900172385.1
89 Pseudooceanicola nanhaiensis DSM 18065 GCA_000688295.1
89 Pseudooceanicola nitratireducens GCA_900112545.1
89 Pseudorhodobacter aquimaris GCA_001202025.1
89 Pseudorhodobacter ferrugineus DSM 5888 GCA_000420745.1
89 Pseudorhodobacter wandonensis GCA_001202035.1
89 Puniceibacterium sediminis GCA_900188035.1
89 Rhodobacter sphaeroides 2.4.1 GCA_000012905.2
89 Rhodobacter sphaeroides ATCC 17025 GCA_000016405.1
89 Roseivivax isoporae LMG 25204 GCA_000521865.1
89 Roseivivax jejudonensis GCA_900172265.1
89 Roseivivax lentus GCA_900156805.1
89 Roseovarius albus GCA_900172335.1
89 Roseovarius atlanticus GCA_001441615.1
89 Roseovarius azorensis GCA_900109455.1
89 Roseovarius lutimaris GCA_900115165.1
89 Roseovarius mucosus GCA_002080415.1
89 Salinihabitans flavidus GCA_900110425.1
89 Sulfitobacter pseudonitzschiae GCA_002222635.1
89 Thalassobius maritimus GCA_900129685.1
89 Thioclava dalianensis GCA_000715505.1
89 Thioclava indica GCA_000714545.1
86 Roseivivax halotolerans GCA_900115815.1
86 Roseovarius nanhaiticus GCA_900156535.1
86 Roseovarius tolerans GCA_900109855.1
qA
354 Streptococcus mutans UA159 GCA_000007465.2
242 Streptococcus equinus GCA_000964315.1
242 Streptococcus gallolyticus subsp. gallolyticus DSM 16831 GCA_002000985.1
242 Streptococcus orisratti DSM 15617 GCA_000380105.1
242 Streptococcus ratti FA-1 = DSM 20564 GCA_000286075.1
241 Streptococcus ferus DSM 20646 GCA_000372425.1
qB
284 Lactobacillus gasseri ATCC 33323 = JCM 1131 GCA_000014425.1
268 Lactobacillus hominis DSM 23910 = CRBIP 24.179 GCA_000296835.1
221 Lactobacillus crispatus ST1 GCA_000091765.1
221 Lactobacillus kefiranofaciens ZW3 GCA_000214785.1
qC
237 Deinococcus radiodurans R1 GCA_000008565.1
176 Deinococcus gobiensis I-0 GCA_000252445.1
157 Deinococcus puniceus GCA_001644565.1
qD
246 Bacillus thuringiensis YBT-1518 GCA_000497525.2
240 Bacillus anthracis str. Ames GCA_000007845.1
240 Bacillus anthracis str. Sterne GCA_000008165.1
240 Bacillus cereus ATCC 14579 GCA_000007825.1
240 Bacillus mycoides GCA_000832605.1
240 Bacillus pseudomycoides DSM 12442 GCA_000161455.1
240 _5BBacillus thuringiensis_5D serovar konkukian str. 97-27 GCA_000008505.1
202 Bacillus megaterium NBRC 15308 = ATCC 14581 GCA_000832985.1
qE
318 Bacillus anthracis str. Ames GCA_000007845.1
318 Bacillus anthracis str. Sterne GCA_000008165.1
318 Bacillus cereus ATCC 14579 GCA_000007825.1
318 Bacillus mycoides GCA_000832605.1
318 Bacillus pseudomycoides DSM 12442 GCA_000161455.1
318 Bacillus thuringiensis YBT-1518 GCA_000497525.2
318 _5BBacillus thuringiensis_5D serovar konkukian str. 97-27 GCA_000008505.1

272 Bacillus manliponensis GCA_000712595.1
263 Bacillus cytotoxicus NVH 391-98 GCA_000017425.1
qF
331 Escherichia coli IAI39 GCA_000026345.1
331 Escherichia coli O104_3AH4 str. 2011C-3493 GCA_000299455.1
331 Escherichia coli O157_3AH7 str. Sakai GCA_000008865.1
331 Escherichia coli O83_3AH1 str. NRG 857C GCA_000183345.1
331 Escherichia coli UMN026 GCA_000026325.2
331 Escherichia coli str. K-12 substr. MG1655 GCA_000005845.2
331 Shigella flexneri 2a str. 301 GCA_000006925.2
331 Tumebacillus flagellatus GCA_000714935.1
322 Shigella dysenteriae Sd197 GCA_000012005.1
299 Erwinia iniecta GCA_001267535.1
qG
270 Deinococcus radiodurans R1 GCA_000008565.1
192 Deinococcus gobiensis I-0 GCA_000252445.1
182 Deinococcus marmoris DSM 12784 GCA_000701405.1
182 Deinococcus swuensis GCA_000800395.1
qH
294 Escherichia coli IAI39 GCA_000026345.1
294 Escherichia coli O104_3AH4 str. 2011C-3493 GCA_000299455.1
294 Escherichia coli O157_3AH7 str. Sakai GCA_000008865.1
294 Escherichia coli UMN026 GCA_000026325.2
294 Escherichia coli str. K-12 substr. MG1655 GCA_000005845.2
294 Shigella flexneri 2a str. 301 GCA_000006925.2
294 Tumebacillus flagellatus GCA_000714935.1
290 Escherichia coli O83_3AH1 str. NRG 857C GCA_000183345.1
285 Shigella dysenteriae Sd197 GCA_000012005.1
qI
164 Bifidobacterium longum NCC2705 GCA_000007525.1
164 Bifidobacterium saguini DSM 23967 GCA_000741715.1
163 Bifidobacterium adolescentis ATCC 15703 GCA_000010425.1
162 Bifidobacterium breve DSM 20213 = JCM 1192 GCA_001025175.1
qJ
172 Clostridium beijerinckii GCA_000833105.2
172 Clostridium saccharobutylicum DSM 13864 GCA_000473995.1
172 Clostridium saccharoperbutylacetonicum N1-4_28HMT_29 GCA_000340885.1
158 Clostridium butyricum GCA_001456065.2
157 Clostridium chromiireducens GCA_002029255.1
157 Clostridium puniceum GCA_002006345.1
qK
133 Escherichia coli IAI39 GCA_000026345.1
133 Escherichia coli O104_3AH4 str. 2011C-3493 GCA_000299455.1
133 Escherichia coli O157_3AH7 str. Sakai GCA_000008865.1
133 Escherichia coli O83_3AH1 str. NRG 857C GCA_000183345.1
133 Escherichia coli UMN026 GCA_000026325.2
133 Escherichia coli str. K-12 substr. MG1655 GCA_000005845.2
133 Shigella dysenteriae Sd197 GCA_000012005.1
133 Shigella flexneri 2a str. 301 GCA_000006925.2
133 Tumebacillus flagellatus GCA_000714935.1
113 Cronobacter sakazakii GCA_000982825.1
113 Erwinia iniecta GCA_001267535.1
113 Kosakonia cowanii GCA_001975225.1
113 Pantoea ananatis LMG 20103 GCA_000025405.2
113 Pantoea dispersa EGD-AAK13 GCA_000465555.2
113 Shimwellia blattae DSM 4481 = NBRC 105725 GCA_000262305.1
111 Xenorhabdus cabanillasii JM26 GCA_000531755.1
111 Xenorhabdus japonica GCA_900115195.1
111 Xenorhabdus koppenhoeferi GCA_900116635.1
qL
217 Rhodobacter sphaeroides 2.4.1 GCA_000012905.2
189 Rhodobacter sphaeroides ATCC 17025 GCA_000016405.1
183 Pseudorhodobacter psychrotolerans GCA_001294535.1
qM
249 Bacillus anthracis str. Ames GCA_000007845.1
249 Bacillus anthracis str. Sterne GCA_000008165.1
249 Bacillus cereus ATCC 14579 GCA_000007825.1
249 Bacillus mycoides GCA_000832605.1
249 Bacillus pseudomycoides DSM 12442 GCA_000161455.1
249 Bacillus thuringiensis YBT-1518 GCA_000497525.2
249 _5BBacillus thuringiensis_5D serovar konkukian str. 97-27 GCA_000008505.1
218 Bacillus dakarensis GCA_900156875.1
213 Bacillus acidicola GCA_001636425.1
213 Bacillus korlensis NBRC 107688 GCA_001591645.1
213 Bacillus okuhidensis GCA_001274915.1
213 Bacillus shackletonii GCA_001420715.1

213 Bacillus sporothermodurans GCA_001587375.1
qN
275 Clostridium beijerinckii GCA_000833105.2
275 Clostridium saccharobutylicum DSM 13864 GCA_000473995.1
275 Clostridium saccharoperbutylacetonicum N1-4_28HMT_29 GCA_000340885.1
268 Clostridium puniceum GCA_002006345.1
257 Clostridium butyricum GCA_001456065.2
257 Clostridium chromiireducens GCA_002029255.1
qO
242 Rhodobacter sphaeroides 2.4.1 GCA_000012905.2
203 Rhodobacter capsulatus SB 1003 GCA_000021865.1
200 Haematobacter massiliensis GCA_000740795.1
qP
196 Clostridium beijerinckii GCA_000833105.2
196 Clostridium saccharobutylicum DSM 13864 GCA_000473995.1
196 Clostridium saccharoperbutylacetonicum N1-4_28HMT_29 GCA_000340885.1
181 Clostridium chromiireducens GCA_002029255.1
181 Clostridium puniceum GCA_002006345.1
168 Clostridium amylolyticum GCA_900142075.1
168 Clostridium polynesiense GCA_000820705.1
qQ
271 Rhodobacter sphaeroides 2.4.1 GCA_000012905.2
242 Rhodobacter sphaeroides ATCC 17025 GCA_000016405.1
241 Pseudorhodobacter ferrugineus DSM 5888 GCA_000420745.1
241 Pseudorhodobacter wandonensis GCA_001202035.1
241 Thioclava dalianensis GCA_000715505.1
241 Thioclava indica GCA_000714545.1
qR
203 Clostridium beijerinckii GCA_000833105.2
203 Clostridium puniceum GCA_002006345.1
203 Clostridium saccharobutylicum DSM 13864 GCA_000473995.1
203 Clostridium saccharoperbutylacetonicum N1-4_28HMT_29 GCA_000340885.1
191 Clostridium chromiireducens GCA_002029255.1
182 Clostridium butyricum GCA_001456065.2
qS
287 Escherichia coli IAI39 GCA_000026345.1
287 Escherichia coli O104_3AH4 str. 2011C-3493 GCA_000299455.1
287 Escherichia coli O157_3AH7 str. Sakai GCA_000008865.1
287 Escherichia coli O83_3AH1 str. NRG 857C GCA_000183345.1
287 Escherichia coli UMN026 GCA_000026325.2
287 Escherichia coli str. K-12 substr. MG1655 GCA_000005845.2
287 Shigella flexneri 2a str. 301 GCA_000006925.2
287 Tumebacillus flagellatus GCA_000714935.1
279 Erwinia iniecta GCA_001267535.1
278 Shigella dysenteriae Sd197 GCA_000012005.1
qT
228 Bacillus anthracis str. Ames GCA_000007845.1
228 Bacillus anthracis str. Sterne GCA_000008165.1
228 Bacillus cereus ATCC 14579 GCA_000007825.1
228 Bacillus mycoides GCA_000832605.1
228 Bacillus pseudomycoides DSM 12442 GCA_000161455.1
228 Bacillus thuringiensis YBT-1518 GCA_000497525.2
228 _5BBacillus thuringiensis_5D serovar konkukian str. 97-27 GCA_000008505.1
194 Bacillus manliponensis GCA_000712595.1
188 Massilibacterium senegalense GCA_001375675.1
qU
171 Gemmobacter aquatilis GCA_900110025.1
158 Pseudorhodobacter psychrotolerans GCA_001294535.1
158 Rhodobacter sphaeroides 2.4.1 GCA_000012905.2
156 Gemmobacter megaterium GCA_900156815.1
156 Gemmobacter nectariphilus DSM 15620 GCA_000429765.1
156 Haematobacter massiliensis GCA_000740795.1
156 Rhodobacter capsulatus SB 1003 GCA_000021865.1
qV
196 Bacillus anthracis str. Ames GCA_000007845.1
196 Bacillus anthracis str. Sterne GCA_000008165.1
196 Bacillus cereus ATCC 14579 GCA_000007825.1
196 Bacillus mycoides GCA_000832605.1
196 Bacillus pseudomycoides DSM 12442 GCA_000161455.1
196 Bacillus thuringiensis YBT-1518 GCA_000497525.2
196 _5BBacillus thuringiensis_5D serovar konkukian str. 97-27 GCA_000008505.1
160 Bacillus manliponensis GCA_000712595.1
153 Bacillus coahuilensis m4-4 GCA_000171615.1
153 Bacillus vietnamensis NBRC 101237 GCA_001591825.1
qW
266 Staphylococcus capitis subsp. capitis GCA_001028645.1

266 Staphylococcus epidermidis ATCC 12228 GCA_000007645.1
263 Staphylococcus haemolyticus JCSC1435 GCA_000009865.1
258 Megasphaera cerevisiae DSM 20462 GCA_001045675.1
258 Staphylococcus warneri SG1 GCA_000332735.1
qX
258 Erwinia iniecta GCA_001267535.1
258 Escherichia coli IAI39 GCA_000026345.1
258 Escherichia coli O104_3AH4 str. 2011C-3493 GCA_000299455.1
258 Escherichia coli O157_3AH7 str. Sakai GCA_000008865.1
258 Escherichia coli UMN026 GCA_000026325.2
258 Escherichia coli str. K-12 substr. MG1655 GCA_000005845.2
258 Shigella flexneri 2a str. 301 GCA_000006925.2
258 Tumebacillus flagellatus GCA_000714935.1
257 Escherichia coli O83_3AH1 str. NRG 857C GCA_000183345.1
249 Shigella dysenteriae Sd197 GCA_000012005.1
qY
206 Clostridium beijerinckii GCA_000833105.2
206 Clostridium puniceum GCA_002006345.1
206 Clostridium saccharoperbutylacetonicum N1-4_28HMT_29 GCA_000340885.1
205 Clostridium taeniosporum GCA_001735765.1
187 Clostridium butyricum GCA_001456065.2
187 Clostridium chromiireducens GCA_002029255.1
187 Clostridium saccharobutylicum DSM 13864 GCA_000473995.1
qZ
329 Deinococcus radiodurans R1 GCA_000008565.1
200 Deinococcus gobiensis I-0 GCA_000252445.1
194 Deinococcus puniceus GCA_001644565.1
r0
271 Streptococcus mutans UA159 GCA_000007465.2
197 Streptococcus equinus GCA_000964315.1
197 Streptococcus gallolyticus subsp. gallolyticus DSM 16831 GCA_002000985.1
197 Streptococcus ratti FA-1 = DSM 20564 GCA_000286075.1
188 Streptococcus orisratti DSM 15617 GCA_000380105.1
r1
240 Bacillus anthracis str. Ames GCA_000007845.1
240 Bacillus anthracis str. Sterne GCA_000008165.1
240 Bacillus cereus ATCC 14579 GCA_000007825.1
240 Bacillus mycoides GCA_000832605.1
240 Bacillus pseudomycoides DSM 12442 GCA_000161455.1
240 Bacillus thuringiensis YBT-1518 GCA_000497525.2
240 _5BBacillus thuringiensis_5D serovar konkukian str. 97-27 GCA_000008505.1
210 Bacillus marisflavi GCA_001274775.1
201 Viridibacillus arvi GCA_001274945.1
r2
183 Enterococcus faecalis V583 GCA_000007785.1
183 Streptomyces cinnamoneus GCA_001885705.1
144 Enterococcus asini ATCC 700915 GCA_000407365.1
144 Enterococcus canis NBRC 100695 GCA_001544375.1
144 Enterococcus casseliflavus EC20 GCA_000157355.2
144 Enterococcus dispar ATCC 51266 GCA_000406945.1
144 Enterococcus faecium DO GCA_000174395.2
144 Enterococcus haemoperoxidus ATCC BAA-382 GCA_000407165.1
144 Enterococcus hirae ATCC 9790 GCA_000271405.2
144 Enterococcus massiliensis GCA_001050095.1
144 Enterococcus mundtii QU 25 GCA_000504125.1
144 Enterococcus phoeniculicola ATCC BAA-412 GCA_000407505.1
144 Enterococcus rivorum GCA_001742285.1
144 Enterococcus saccharolyticus subsp. saccharolyticus ATCC 43076 GCA_000407285.1
144 Enterococcus thailandicus GCA_001652875.1
139 Enterococcus cecorum GCA_001318405.1
139 Vagococcus penaei GCA_001998885.1
r3
200 Clostridium beijerinckii GCA_000833105.2
200 Clostridium puniceum GCA_002006345.1
200 Clostridium saccharoperbutylacetonicum N1-4_28HMT_29 GCA_000340885.1
186 Clostridium taeniosporum GCA_001735765.1
185 Clostridium saccharobutylicum DSM 13864 GCA_000473995.1
r4
109 Bifidobacterium adolescentis ATCC 15703 GCA_000010425.1
109 Bifidobacterium bohemicum DSM 22767 GCA_000741525.1
109 Bifidobacterium breve DSM 20213 = JCM 1192 GCA_001025175.1
109 Bifidobacterium choerinum GCA_000741135.1
109 Bifidobacterium longum NCC2705 GCA_000007525.1
109 Bifidobacterium pseudolongum PV8-2 GCA_000800475.2
100 Bifidobacterium saguini DSM 23967 GCA_000741715.1
94 Bifidobacterium angulatum DSM 20098 = JCM 7096 GCA_001025155.1

94 Bifidobacterium biavatii DSM 23969 GCA_000741165.1
94 Bifidobacterium callitrichos DSM 23973 GCA_000741175.1
94 Bifidobacterium coryneforme GCA_000737865.1
94 Bifidobacterium dentium JCM 1195 = DSM 20436 GCA_001042595.1
94 Bifidobacterium gallicum DSM 20093 = LMG 11596 GCA_000741205.1
94 Bifidobacterium gallinarum GCA_000741215.1
94 Bifidobacterium scardovii JCM 12489 = DSM 13734 GCA_001042635.1
94 Bifidobacterium tsurumiense GCA_000741765.1
r5
238 Escherichia coli IAI39 GCA_000026345.1
238 Escherichia coli O104_3AH4 str. 2011C-3493 GCA_000299455.1
238 Escherichia coli O157_3AH7 str. Sakai GCA_000008865.1
238 Escherichia coli O83_3AH1 str. NRG 857C GCA_000183345.1
238 Escherichia coli UMN026 GCA_000026325.2
238 Escherichia coli str. K-12 substr. MG1655 GCA_000005845.2
238 Shigella dysenteriae Sd197 GCA_000012005.1
238 Shigella flexneri 2a str. 301 GCA_000006925.2
238 Tumebacillus flagellatus GCA_000714935.1
208 Cronobacter sakazakii GCA_000982825.1
208 Erwinia iniecta GCA_001267535.1
208 Erwinia toletana DAPP-PG 735 GCA_000336255.1
206 Erwinia billingiae Eb661 GCA_000196615.1
206 Erwinia gerundensis GCA_001517405.1
206 Pantoea agglomerans GCA_001709315.1
206 Pantoea ananatis LMG 20103 GCA_000025405.2
206 Pantoea dispersa EGD-AAK13 GCA_000465555.2
206 Shimwellia blattae DSM 4481 = NBRC 105725 GCA_000262305.1
r6
218 Clostridium beijerinckii GCA_000833105.2
218 Clostridium puniceum GCA_002006345.1
218 Clostridium saccharobutylicum DSM 13864 GCA_000473995.1
218 Clostridium saccharoperbutylacetonicum N1-4_28HMT_29 GCA_000340885.1
203 Clostridium butyricum GCA_001456065.2
203 Clostridium chromiireducens GCA_002029255.1
201 Clostridium neonatale GCA_001458595.1
r7
129 Enterococcus faecalis V583 GCA_000007785.1
129 Streptomyces cinnamoneus GCA_001885705.1
110 Enterococcus canis NBRC 100695 GCA_001544375.1
110 Enterococcus faecium DO GCA_000174395.2
110 Enterococcus haemoperoxidus ATCC BAA-382 GCA_000407165.1
110 Enterococcus hirae ATCC 9790 GCA_000271405.2
110 Enterococcus mundtii QU 25 GCA_000504125.1
110 Enterococcus phoeniculicola ATCC BAA-412 GCA_000407505.1
110 Enterococcus rivorum GCA_001742285.1
110 Enterococcus thailandicus GCA_001652875.1
110 Isobaculum melis GCA_900111355.1
109 Abiotrophia defectiva ATCC 49176 GCA_000160075.2
109 Enterococcus asini ATCC 700915 GCA_000407365.1
109 Enterococcus casseliflavus EC20 GCA_000157355.2
109 Enterococcus cecorum GCA_001318405.1
109 Enterococcus dispar ATCC 51266 GCA_000406945.1
109 Enterococcus massiliensis GCA_001050095.1
109 Enterococcus saccharolyticus subsp. saccharolyticus ATCC 43076 GCA_000407285.1
r8
191 Staphylococcus condimenti GCA_001618885.1
191 Staphylococcus simulans GCA_001559115.1
190 Staphylococcus aureus subsp. aureus NCTC 8325 GCA_000013425.1
190 Staphylococcus capitis subsp. capitis GCA_001028645.1
190 Staphylococcus cohnii subsp. cohnii GCA_000972575.1
190 Staphylococcus epidermidis ATCC 12228 GCA_000007645.1
190 Staphylococcus haemolyticus JCSC1435 GCA_000009865.1
190 Staphylococcus hominis subsp. hominis C80 GCA_000183685.1
190 Staphylococcus saprophyticus GCA_001074355.1
190 Staphylococcus simiae CCM 7213 GCA_000235645.2
176 Staphylococcus microti GCA_000934465.1
r9
131 Rhodobacter sphaeroides 2.4.1 GCA_000012905.2
109 Defluviimonas alba GCA_001620265.1
106 Gemmobacter aquatilis GCA_900110025.1
106 Pseudorhodobacter psychrotolerans GCA_001294535.1
106 Rhodobacter sphaeroides ATCC 17025 GCA_000016405.1
ra
182 Bifidobacterium adolescentis ATCC 15703 GCA_000010425.1
158 Bifidobacterium asteroides PRL2011 GCA_000304215.1
158 Bifidobacterium callitrichos DSM 23973 GCA_000741175.1

158 Bifidobacterium coryneforme GCA_000737865.1
149 Bifidobacterium dentium JCM 1195 = DSM 20436 GCA_001042595.1
rb
205 Clostridium beijerinckii GCA_000833105.2
205 Clostridium puniceum GCA_002006345.1
205 Clostridium saccharoperbutylacetonicum N1-4_28HMT_29 GCA_000340885.1
198 Clostridium butyricum GCA_001456065.2
191 Clostridium taeniosporum GCA_001735765.1
rc
179 Bifidobacterium adolescentis ATCC 15703 GCA_000010425.1
157 Bifidobacterium callitrichos DSM 23973 GCA_000741175.1
157 Bifidobacterium coryneforme GCA_000737865.1
157 Bifidobacterium minimum GCA_000741645.1
157 Bifidobacterium mongoliense DSM 21395 GCA_000741285.1
155 Bifidobacterium thermophilum GCA_000741495.1
rd
88 Rhodobacter sphaeroides 2.4.1 GCA_000012905.2
74 Pseudorhodobacter psychrotolerans GCA_001294535.1
74 Rhodobacter sphaeroides ATCC 17025 GCA_000016405.1
60 Pseudorhodobacter ferrugineus DSM 5888 GCA_000420745.1
60 Pseudorhodobacter wandonensis GCA_001202035.1
re
216 Bacillus cereus ATCC 14579 GCA_000007825.1
205 Bacillus anthracis str. Ames GCA_000007845.1
205 Bacillus anthracis str. Sterne GCA_000008165.1
205 Bacillus mycoides GCA_000832605.1
205 Bacillus pseudomycoides DSM 12442 GCA_000161455.1
205 Bacillus thuringiensis YBT-1518 GCA_000497525.2
205 _5BBacillus thuringiensis_5D serovar konkukian str. 97-27 GCA_000008505.1
170 Viridibacillus arvi GCA_001274945.1
rf
186 Streptococcus mutans UA159 GCA_000007465.2
117 Streptococcus iniae GCA_000831485.1
109 Streptococcus macacae NCTC 11558 GCA_000187995.3
rg
132 Escherichia coli IAI39 GCA_000026345.1
132 Escherichia coli O104_3AH4 str. 2011C-3493 GCA_000299455.1
132 Escherichia coli O157_3AH7 str. Sakai GCA_000008865.1
132 Escherichia coli O83_3AH1 str. NRG 857C GCA_000183345.1
132 Escherichia coli UMN026 GCA_000026325.2
132 Escherichia coli str. K-12 substr. MG1655 GCA_000005845.2
132 Shigella dysenteriae Sd197 GCA_000012005.1
132 Shigella flexneri 2a str. 301 GCA_000006925.2
132 Tumebacillus flagellatus GCA_000714935.1
128 Abiotrophia defectiva ATCC 49176 GCA_000160075.2
128 Enterococcus asini ATCC 700915 GCA_000407365.1
128 Enterococcus canis NBRC 100695 GCA_001544375.1
128 Enterococcus cecorum GCA_001318405.1
128 Enterococcus columbae DSM 7374 = ATCC 51263 GCA_000406925.1
128 Enterococcus dispar ATCC 51266 GCA_000406945.1
128 Enterococcus faecalis V583 GCA_000007785.1
128 Enterococcus faecium DO GCA_000174395.2
128 Enterococcus haemoperoxidus ATCC BAA-382 GCA_000407165.1
128 Enterococcus hirae ATCC 9790 GCA_000271405.2
128 Enterococcus massiliensis GCA_001050095.1
128 Enterococcus mundtii QU 25 GCA_000504125.1
128 Enterococcus phoeniculicola ATCC BAA-412 GCA_000407505.1
128 Enterococcus rivorum GCA_001742285.1
128 Enterococcus thailandicus GCA_001652875.1
128 Isobaculum melis GCA_900111355.1
128 Melissococcus plutonius S1 GCA_000747585.1
128 Streptomyces cinnamoneus GCA_001885705.1
128 Vagococcus penaei GCA_001998885.1
124 Photorhabdus temperata subsp. thracensis GCA_001010285.1
rh
200 Clostridium beijerinckii GCA_000833105.2
200 Clostridium saccharobutylicum DSM 13864 GCA_000473995.1
200 Clostridium saccharoperbutylacetonicum N1-4_28HMT_29 GCA_000340885.1
186 Clostridium puniceum GCA_002006345.1
179 Clostridium butyricum GCA_001456065.2
ri
151 Bifidobacterium adolescentis ATCC 15703 GCA_000010425.1
147 Bifidobacterium tsurumiense GCA_000741765.1
136 Bifidobacterium longum NCC2705 GCA_000007525.1
rj
240 Clostridium beijerinckii GCA_000833105.2

240 Clostridium puniceum GCA_002006345.1
240 Clostridium saccharobutylicum DSM 13864 GCA_000473995.1
240 Clostridium saccharoperbutylacetonicum N1-4_28HMT_29 GCA_000340885.1
229 Clostridium chromiireducens GCA_002029255.1
226 Clostridium butyricum GCA_001456065.2
rk
183 Lactobacillus gasseri ATCC 33323 = JCM 1131 GCA_000014425.1
171 Streptococcus mutans UA159 GCA_000007465.2
159 Lactobacillus hominis DSM 23910 = CRBIP 24.179 GCA_000296835.1
rl
222 Staphylococcus epidermidis ATCC 12228 GCA_000007645.1
217 Staphylococcus lugdunensis HKU09-01 GCA_000025085.1
214 Staphylococcus capitis subsp. capitis GCA_001028645.1
rm
269 Streptococcus mutans UA159 GCA_000007465.2
170 Streptococcus marimammalium DSM 18627 GCA_000380045.1
155 Streptococcus macacae NCTC 11558 GCA_000187995.3
rn
203 Clostridium beijerinckii GCA_000833105.2
203 Clostridium puniceum GCA_002006345.1
203 Clostridium saccharoperbutylacetonicum N1-4_28HMT_29 GCA_000340885.1
199 Clostridium saccharobutylicum DSM 13864 GCA_000473995.1
183 Clostridium chromiireducens GCA_002029255.1
ro
284 Clostridium beijerinckii GCA_000833105.2
284 Clostridium puniceum GCA_002006345.1
284 Clostridium saccharoperbutylacetonicum N1-4_28HMT_29 GCA_000340885.1
275 Clostridium saccharobutylicum DSM 13864 GCA_000473995.1
265 Clostridium butyricum GCA_001456065.2
rp
216 Deinococcus radiodurans R1 GCA_000008565.1
154 Deinococcus gobiensis I-0 GCA_000252445.1
110 Deinococcus puniceus GCA_001644565.1
rq
153 Rhodobacter sphaeroides 2.4.1 GCA_000012905.2
134 Rhodobacter sphaeroides ATCC 17025 GCA_000016405.1
128 Defluviimonas alba GCA_001620265.1
rr
177 Clostridium beijerinckii GCA_000833105.2
177 Clostridium puniceum GCA_002006345.1
177 Clostridium saccharobutylicum DSM 13864 GCA_000473995.1
177 Clostridium saccharoperbutylacetonicum N1-4_28HMT_29 GCA_000340885.1
164 Clostridium neonatale GCA_001458595.1
151 Clostridium chromiireducens GCA_002029255.1
rs
252 Escherichia coli IAI39 GCA_000026345.1
252 Escherichia coli O104_3AH4 str. 2011C-3493 GCA_000299455.1
252 Escherichia coli O157_3AH7 str. Sakai GCA_000008865.1
252 Escherichia coli O83_3AH1 str. NRG 857C GCA_000183345.1
252 Escherichia coli UMN026 GCA_000026325.2
252 Escherichia coli str. K-12 substr. MG1655 GCA_000005845.2
252 Tumebacillus flagellatus GCA_000714935.1
248 Shigella dysenteriae Sd197 GCA_000012005.1
246 Shigella flexneri 2a str. 301 GCA_000006925.2
rt
234 Lactobacillus gasseri ATCC 33323 = JCM 1131 GCA_000014425.1
217 Lactobacillus hominis DSM 23910 = CRBIP 24.179 GCA_000296835.1
176 Lactobacillus iners DSM 13335 GCA_000160875.1
ru
205 Deinococcus radiodurans R1 GCA_000008565.1
108 Deinococcus puniceus GCA_001644565.1
99 Deinococcus gobiensis I-0 GCA_000252445.1
rv
174 Clostridium saccharobutylicum DSM 13864 GCA_000473995.1
170 Clostridium beijerinckii GCA_000833105.2
170 Clostridium botulinum B str. Eklund 17B _28NRP_29 GCA_000020165.1
170 Clostridium chromiireducens GCA_002029255.1
170 Clostridium puniceum GCA_002006345.1
170 Clostridium saccharoperbutylacetonicum N1-4_28HMT_29 GCA_000340885.1
170 Clostridium taeniosporum GCA_001735765.1
155 Clostridium uliginosum GCA_900112485.1
rw
155 Staphylococcus pettenkoferi GCA_002208805.1
155 Staphylococcus simulans GCA_001559115.1
144 Massilibacterium senegalense GCA_001375675.1
141 Staphylococcus condimenti GCA_001618885.1

141 Staphylococcus hyicus GCA_000816085.1
141 Staphylococcus lentus F1142 GCA_000286395.1
141 Staphylococcus lutrae GCA_002101335.1
141 Staphylococcus pseudintermedius HKU10-03 GCA_000185885.1
141 Staphylococcus sciuri GCA_002209165.1
rx
231 Staphylococcus arlettae CVD059 GCA_000295715.1
231 Staphylococcus capitis subsp. capitis GCA_001028645.1
231 Staphylococcus cohnii subsp. cohnii GCA_000972575.1
231 Staphylococcus epidermidis ATCC 12228 GCA_000007645.1
231 Staphylococcus haemolyticus JCSC1435 GCA_000009865.1
231 Staphylococcus hominis subsp. hominis C80 GCA_000183685.1
231 Staphylococcus lugdunensis HKU09-01 GCA_000025085.1
231 Staphylococcus saprophyticus subsp. saprophyticus ATCC 15305 GCA_000010125.1
216 Staphylococcus condimenti GCA_001618885.1
216 Staphylococcus simulans GCA_001559115.1
215 Staphylococcus aureus subsp. aureus NCTC 8325 GCA_000013425.1
215 Staphylococcus gallinarum GCA_000875895.1
215 Staphylococcus simiae CCM 7213 GCA_000235645.2
215 Staphylococcus succinus GCA_001902315.1
215 Staphylococcus xylosus GCA_000706685.1
ry
295 Staphylococcus epidermidis ATCC 12228 GCA_000007645.1
284 Staphylococcus aureus subsp. aureus NCTC 8325 GCA_000013425.1
284 Staphylococcus haemolyticus JCSC1435 GCA_000009865.1
284 Staphylococcus hominis subsp. hominis C80 GCA_000183685.1
284 Staphylococcus simiae CCM 7213 GCA_000235645.2
281 Staphylococcus cohnii subsp. cohnii GCA_000972575.1
rz
189 Bifidobacterium adolescentis ATCC 15703 GCA_000010425.1
178 Bifidobacterium asteroides PRL2011 GCA_000304215.1
166 Bifidobacterium bifidum PRL2010 GCA_000165905.1
rA
205 Megasphaera cerevisiae DSM 20462 GCA_001045675.1
205 Staphylococcus aureus subsp. aureus NCTC 8325 GCA_000013425.1
205 Staphylococcus capitis subsp. capitis GCA_001028645.1
205 Staphylococcus epidermidis ATCC 12228 GCA_000007645.1
205 Staphylococcus haemolyticus JCSC1435 GCA_000009865.1
205 Staphylococcus hominis subsp. hominis C80 GCA_000183685.1
205 Staphylococcus lugdunensis HKU09-01 GCA_000025085.1
205 Staphylococcus simiae CCM 7213 GCA_000235645.2
205 Staphylococcus warneri SG1 GCA_000332735.1
200 Staphylococcus lutrae GCA_002101335.1
198 Staphylococcus arlettae CVD059 GCA_000295715.1
198 Staphylococcus cohnii subsp. cohnii GCA_000972575.1
198 Staphylococcus gallinarum GCA_000875895.1
198 Staphylococcus saprophyticus GCA_001074355.1
198 Staphylococcus saprophyticus subsp. saprophyticus ATCC 15305 GCA_000010125.1
198 Staphylococcus succinus GCA_001902315.1
198 Staphylococcus xylosus GCA_000706685.1
rB
164 Bifidobacterium adolescentis ATCC 15703 GCA_000010425.1
142 Bifidobacterium angulatum DSM 20098 = JCM 7096 GCA_001025155.1
136 Bifidobacterium breve DSM 20213 = JCM 1192 GCA_001025175.1
136 Bifidobacterium longum NCC2705 GCA_000007525.1
rC
257 Bacillus thuringiensis YBT-1518 GCA_000497525.2
251 Bacillus anthracis str. Ames GCA_000007845.1
251 Bacillus anthracis str. Sterne GCA_000008165.1
251 Bacillus cereus ATCC 14579 GCA_000007825.1
251 Bacillus mycoides GCA_000832605.1
251 Bacillus pseudomycoides DSM 12442 GCA_000161455.1
251 _5BBacillus thuringiensis_5D serovar konkukian str. 97-27 GCA_000008505.1
181 Bacillus manliponensis GCA_000712595.1
rD
244 Deinococcus radiodurans R1 GCA_000008565.1
160 Deinococcus puniceus GCA_001644565.1
140 Deinococcus deserti VCD115 GCA_000020685.1
rE
171 Clostridium beijerinckii GCA_000833105.2
171 Clostridium puniceum GCA_002006345.1
171 Clostridium saccharoperbutylacetonicum N1-4_28HMT_29 GCA_000340885.1
156 Clostridium saccharobutylicum DSM 13864 GCA_000473995.1
153 Clostridium butyricum GCA_001456065.2
153 Clostridium taeniosporum GCA_001735765.1
rF

210 Deinococcus radiodurans R1 GCA_000008565.1
118 Deinococcus puniceus GCA_001644565.1
107 Deinococcus deserti VCD115 GCA_000020685.1
rG
272 Clostridium beijerinckii GCA_000833105.2
272 Clostridium saccharoperbutylacetonicum N1-4_28HMT_29 GCA_000340885.1
268 Clostridium puniceum GCA_002006345.1
264 Clostridium saccharobutylicum DSM 13864 GCA_000473995.1
rH
230 Staphylococcus epidermidis ATCC 12228 GCA_000007645.1
223 Staphylococcus capitis subsp. capitis GCA_001028645.1
208 Staphylococcus haemolyticus JCSC1435 GCA_000009865.1
rI
282 Bacillus anthracis str. Ames GCA_000007845.1
282 Bacillus anthracis str. Sterne GCA_000008165.1
282 Bacillus cereus ATCC 14579 GCA_000007825.1
282 Bacillus mycoides GCA_000832605.1
282 Bacillus pseudomycoides DSM 12442 GCA_000161455.1
282 Bacillus thuringiensis YBT-1518 GCA_000497525.2
282 _5BBacillus thuringiensis_5D serovar konkukian str. 97-27 GCA_000008505.1
205 Bacillus horneckiae GCA_001636335.1
205 Bacillus solani GCA_001420595.1
199 Bacillus manliponensis GCA_000712595.1
rJ
308 Streptococcus mutans UA159 GCA_000007465.2
221 Streptococcus sobrinus DSM 20742 = ATCC 33478 GCA_000686605.1
219 Streptococcus ratti FA-1 = DSM 20564 GCA_000286075.1
rK
220 Deinococcus radiodurans R1 GCA_000008565.1
159 Deinococcus gobiensis I-0 GCA_000252445.1
159 Deinococcus proteolyticus MRP GCA_000190555.1
147 Deinococcus puniceus GCA_001644565.1
rL
264 Streptococcus mutans UA159 GCA_000007465.2
202 Streptococcus ferus DSM 20646 GCA_000372425.1
193 Streptococcus macacae NCTC 11558 GCA_000187995.3
rM
261 Deinococcus radiodurans R1 GCA_000008565.1
188 Deinococcus gobiensis I-0 GCA_000252445.1
186 Deinococcus deserti VCD115 GCA_000020685.1
rN
204 Clostridium beijerinckii GCA_000833105.2
204 Clostridium puniceum GCA_002006345.1
204 Clostridium saccharobutylicum DSM 13864 GCA_000473995.1
204 Clostridium saccharoperbutylacetonicum N1-4_28HMT_29 GCA_000340885.1
197 Clostridium chromiireducens GCA_002029255.1
192 Clostridium butyricum GCA_001456065.2
rO
178 Clostridium beijerinckii GCA_000833105.2
178 Clostridium saccharobutylicum DSM 13864 GCA_000473995.1
178 Clostridium saccharoperbutylacetonicum N1-4_28HMT_29 GCA_000340885.1
175 Clostridium butyricum GCA_001456065.2
171 Clostridium puniceum GCA_002006345.1
rP
273 Escherichia coli IAI39 GCA_000026345.1
273 Escherichia coli O104_3AH4 str. 2011C-3493 GCA_000299455.1
273 Escherichia coli O157_3AH7 str. Sakai GCA_000008865.1
273 Escherichia coli O83_3AH1 str. NRG 857C GCA_000183345.1
273 Escherichia coli UMN026 GCA_000026325.2
273 Escherichia coli str. K-12 substr. MG1655 GCA_000005845.2
273 Shigella dysenteriae Sd197 GCA_000012005.1
273 Shigella flexneri 2a str. 301 GCA_000006925.2
273 Tumebacillus flagellatus GCA_000714935.1
244 Erwinia iniecta GCA_001267535.1
230 Pantoea agglomerans GCA_001709315.1
rQ
263 Staphylococcus capitis subsp. capitis GCA_001028645.1
263 Staphylococcus epidermidis ATCC 12228 GCA_000007645.1
263 Staphylococcus lugdunensis HKU09-01 GCA_000025085.1
256 Staphylococcus haemolyticus JCSC1435 GCA_000009865.1
256 Staphylococcus hominis subsp. hominis C80 GCA_000183685.1
251 Staphylococcus cohnii subsp. cohnii GCA_000972575.1
251 Staphylococcus pettenkoferi GCA_002208805.1
251 Staphylococcus saprophyticus subsp. saprophyticus ATCC 15305 GCA_000010125.1
rR
257 Deinococcus radiodurans R1 GCA_000008565.1

186 Deinococcus gobiensis I-0 GCA_000252445.1
165 Deinococcus puniceus GCA_001644565.1
rS
303 Bacillus thuringiensis YBT-1518 GCA_000497525.2
291 Bacillus anthracis str. Ames GCA_000007845.1
291 Bacillus anthracis str. Sterne GCA_000008165.1
291 Bacillus cereus ATCC 14579 GCA_000007825.1
291 Bacillus mycoides GCA_000832605.1
291 Bacillus pseudomycoides DSM 12442 GCA_000161455.1
291 _5BBacillus thuringiensis_5D serovar konkukian str. 97-27 GCA_000008505.1
262 Bacillus aquimaris TF-12 GCA_001648555.1
rT
196 Enterococcus faecalis V583 GCA_000007785.1
196 Streptomyces cinnamoneus GCA_001885705.1
173 Enterococcus hirae ATCC 9790 GCA_000271405.2
165 Enterococcus canis NBRC 100695 GCA_001544375.1
165 Enterococcus casseliflavus EC20 GCA_000157355.2
165 Enterococcus dispar ATCC 51266 GCA_000406945.1
165 Enterococcus faecium DO GCA_000174395.2
165 Enterococcus haemoperoxidus ATCC BAA-382 GCA_000407165.1
165 Enterococcus mundtii QU 25 GCA_000504125.1
165 Enterococcus phoeniculicola ATCC BAA-412 GCA_000407505.1
165 Enterococcus rivorum GCA_001742285.1
165 Enterococcus saccharolyticus subsp. saccharolyticus ATCC 43076 GCA_000407285.1
165 Enterococcus thailandicus GCA_001652875.1
rU
220 Lactobacillus gasseri ATCC 33323 = JCM 1131 GCA_000014425.1
195 Lactobacillus hominis DSM 23910 = CRBIP 24.179 GCA_000296835.1
132 Lactobacillus acidophilus NCFM GCA_000011985.1
132 Lactobacillus crispatus ST1 GCA_000091765.1
132 Lactobacillus gallinarum GCA_001314245.2
rV
174 Deinococcus radiodurans R1 GCA_000008565.1
92 Deinococcus gobiensis I-0 GCA_000252445.1
89 Deinococcus puniceus GCA_001644565.1
rW
216 Bifidobacterium adolescentis ATCC 15703 GCA_000010425.1
189 Bifidobacterium tsurumiense GCA_000741765.1
187 Bifidobacterium thermophilum GCA_000741495.1
rX
317 Escherichia coli IAI39 GCA_000026345.1
317 Escherichia coli O104_3AH4 str. 2011C-3493 GCA_000299455.1
317 Escherichia coli O157_3AH7 str. Sakai GCA_000008865.1
317 Escherichia coli UMN026 GCA_000026325.2
317 Escherichia coli str. K-12 substr. MG1655 GCA_000005845.2
317 Shigella dysenteriae Sd197 GCA_000012005.1
317 Shigella flexneri 2a str. 301 GCA_000006925.2
317 Tumebacillus flagellatus GCA_000714935.1
302 Escherichia coli O83_3AH1 str. NRG 857C GCA_000183345.1
281 Erwinia iniecta GCA_001267535.1
rY
245 Deinococcus radiodurans R1 GCA_000008565.1
139 Deinococcus gobiensis I-0 GCA_000252445.1
122 Deinococcus marmoris DSM 12784 GCA_000701405.1
122 Deinococcus swuensis GCA_000800395.1
rZ
121 Streptococcus equinus GCA_000964315.1
121 Streptococcus gallolyticus subsp. gallolyticus DSM 16831 GCA_002000985.1
121 Streptococcus mutans UA159 GCA_000007465.2
114 Streptococcus cristatus AS 1.3089 GCA_000385925.1
114 Streptococcus gordonii str. Challis substr. CH1 GCA_000017005.1
114 Streptococcus mitis B6 GCA_000027165.1
114 Streptococcus pneumoniae R6 GCA_000007045.1
114 Streptococcus salivarius GCA_000785515.1
114 Streptococcus sanguinis SK36 GCA_000014205.1
114 Streptococcus thermophilus JIM 8232 GCA_000253395.1
112 Streptococcus orisratti DSM 15617 GCA_000380105.1
112 Streptococcus ratti FA-1 = DSM 20564 GCA_000286075.1
s0
183 Clostridium beijerinckii GCA_000833105.2
183 Clostridium puniceum GCA_002006345.1
183 Clostridium saccharobutylicum DSM 13864 GCA_000473995.1
183 Clostridium saccharoperbutylacetonicum N1-4_28HMT_29 GCA_000340885.1
168 Clostridium chromiireducens GCA_002029255.1
163 Clostridium butyricum GCA_001456065.2
s1

103 Clostridium beijerinckii GCA_000833105.2
103 Clostridium saccharobutylicum DSM 13864 GCA_000473995.1
103 Clostridium saccharoperbutylacetonicum N1-4_28HMT_29 GCA_000340885.1
99 Clostridium chromiireducens GCA_002029255.1
99 Clostridium puniceum GCA_002006345.1
98 Clostridium taeniosporum GCA_001735765.1
s2
197 Bifidobacterium adolescentis ATCC 15703 GCA_000010425.1
179 Bifidobacterium callitrichos DSM 23973 GCA_000741175.1
169 Bifidobacterium tsurumiense GCA_000741765.1
s3
178 Deinococcus radiodurans R1 GCA_000008565.1
95 Deinococcus gobiensis I-0 GCA_000252445.1
89 Deinococcus puniceus GCA_001644565.1
s4
158 Clostridium beijerinckii GCA_000833105.2
158 Clostridium chromiireducens GCA_002029255.1
158 Clostridium puniceum GCA_002006345.1
158 Clostridium saccharobutylicum DSM 13864 GCA_000473995.1
158 Clostridium saccharoperbutylacetonicum N1-4_28HMT_29 GCA_000340885.1
155 Clostridium botulinum B str. Eklund 17B _28NRP_29 GCA_000020165.1
155 Clostridium taeniosporum GCA_001735765.1
155 Clostridium uliginosum GCA_900112485.1
150 Clostridium fallax GCA_900129365.1
150 Clostridium gasigenes GCA_900104115.1
s5
225 Streptococcus mutans UA159 GCA_000007465.2
133 Streptococcus macacae NCTC 11558 GCA_000187995.3
125 Streptococcus marimammalium DSM 18627 GCA_000380045.1
s6
137 Clostridium beijerinckii GCA_000833105.2
137 Clostridium puniceum GCA_002006345.1
137 Clostridium saccharobutylicum DSM 13864 GCA_000473995.1
137 Clostridium saccharoperbutylacetonicum N1-4_28HMT_29 GCA_000340885.1
123 Clostridium neonatale GCA_001458595.1
122 Clostridium butyricum GCA_001456065.2
122 Clostridium chromiireducens GCA_002029255.1
s7
232 Escherichia coli IAI39 GCA_000026345.1
232 Escherichia coli O104_3AH4 str. 2011C-3493 GCA_000299455.1
232 Escherichia coli O157_3AH7 str. Sakai GCA_000008865.1
232 Escherichia coli O83_3AH1 str. NRG 857C GCA_000183345.1
232 Escherichia coli UMN026 GCA_000026325.2
232 Escherichia coli str. K-12 substr. MG1655 GCA_000005845.2
232 Shigella dysenteriae Sd197 GCA_000012005.1
232 Shigella flexneri 2a str. 301 GCA_000006925.2
232 Tumebacillus flagellatus GCA_000714935.1
212 Erwinia iniecta GCA_001267535.1
202 Rosenbergiella nectarea GCA_900111105.1
s8
232 Rhodobacter sphaeroides 2.4.1 GCA_000012905.2
209 Pseudorhodobacter psychrotolerans GCA_001294535.1
205 Rhodobacter sphaeroides ATCC 17025 GCA_000016405.1
s9
249 Clostridium beijerinckii GCA_000833105.2
249 Clostridium saccharoperbutylacetonicum N1-4_28HMT_29 GCA_000340885.1
243 Clostridium puniceum GCA_002006345.1
234 Clostridium saccharobutylicum DSM 13864 GCA_000473995.1
sa
197 Bacillus anthracis str. Ames GCA_000007845.1
197 Bacillus anthracis str. Sterne GCA_000008165.1
197 Bacillus cereus ATCC 14579 GCA_000007825.1
197 Bacillus manliponensis GCA_000712595.1
197 Bacillus mycoides GCA_000832605.1
197 Bacillus pseudomycoides DSM 12442 GCA_000161455.1
197 Bacillus thuringiensis YBT-1518 GCA_000497525.2
197 _5BBacillus thuringiensis_5D serovar konkukian str. 97-27 GCA_000008505.1
192 Bacillus cytotoxicus NVH 391-98 GCA_000017425.1
182 Anaerobacillus alkalilacustris GCA_001866005.1
182 Anaerobacillus arseniciselenatis GCA_001865995.1
182 Listeria floridensis FSL S10-1187 GCA_000525875.1
sb
255 Streptococcus mutans UA159 GCA_000007465.2
183 Streptococcus macacae NCTC 11558 GCA_000187995.3
179 Streptococcus ratti FA-1 = DSM 20564 GCA_000286075.1
sc

241 Deinococcus radiodurans R1 GCA_000008565.1
145 Deinococcus marmoris DSM 12784 GCA_000701405.1
145 Deinococcus swuensis GCA_000800395.1
144 Deinococcus puniceus GCA_001644565.1
sd
176 Rhodobacter sphaeroides 2.4.1 GCA_000012905.2
160 Rhodobacter sphaeroides ATCC 17025 GCA_000016405.1
145 Pseudorhodobacter ferrugineus DSM 5888 GCA_000420745.1
145 Pseudorhodobacter wandonensis GCA_001202035.1
se
181 Lactobacillus gasseri ATCC 33323 = JCM 1131 GCA_000014425.1
162 Lactobacillus hominis DSM 23910 = CRBIP 24.179 GCA_000296835.1
106 Lactobacillus acetotolerans GCA_001042405.1
106 Lactobacillus amylovorus GCA_000191545.1
106 Lactobacillus crispatus ST1 GCA_000091765.1
106 Lactobacillus delbrueckii subsp. bulgaricus ATCC 11842 = JCM 1002 GCA_000056065.1
106 Lactobacillus helveticus GCA_001308285.1
106 Lactobacillus kalixensis DSM 16043 GCA_001434335.1
106 Lactobacillus psittaci DSM 15354 GCA_000425905.1
sf
238 Lactobacillus gasseri ATCC 33323 = JCM 1131 GCA_000014425.1
210 Lactobacillus hominis DSM 23910 = CRBIP 24.179 GCA_000296835.1
174 Lactobacillus acidophilus NCFM GCA_000011985.1
174 Lactobacillus crispatus ST1 GCA_000091765.1
sg
286 Escherichia coli IAI39 GCA_000026345.1
286 Escherichia coli O104_3AH4 str. 2011C-3493 GCA_000299455.1
286 Escherichia coli O157_3AH7 str. Sakai GCA_000008865.1
286 Escherichia coli O83_3AH1 str. NRG 857C GCA_000183345.1
286 Escherichia coli UMN026 GCA_000026325.2
286 Escherichia coli str. K-12 substr. MG1655 GCA_000005845.2
286 Shigella flexneri 2a str. 301 GCA_000006925.2
286 Tumebacillus flagellatus GCA_000714935.1
271 Erwinia iniecta GCA_001267535.1
270 Shigella dysenteriae Sd197 GCA_000012005.1
sh
96 Bifidobacterium adolescentis ATCC 15703 GCA_000010425.1
91 Bifidobacterium breve DSM 20213 = JCM 1192 GCA_001025175.1
77 Bifidobacterium choerinum GCA_000741135.1
77 Bifidobacterium pseudolongum PV8-2 GCA_000800475.2
si
267 Clostridium saccharobutylicum DSM 13864 GCA_000473995.1
265 Clostridium beijerinckii GCA_000833105.2
265 Clostridium puniceum GCA_002006345.1
265 Clostridium saccharoperbutylacetonicum N1-4_28HMT_29 GCA_000340885.1
256 Clostridium chromiireducens GCA_002029255.1
sj
207 Rhodobacter sphaeroides 2.4.1 GCA_000012905.2
169 Rhodobacter sphaeroides ATCC 17025 GCA_000016405.1
151 Gemmobacter megaterium GCA_900156815.1
sk
226 Clostridium beijerinckii GCA_000833105.2
226 Clostridium puniceum GCA_002006345.1
226 Clostridium saccharoperbutylacetonicum N1-4_28HMT_29 GCA_000340885.1
217 Clostridium saccharobutylicum DSM 13864 GCA_000473995.1
198 Clostridium neonatale GCA_001458595.1
sl
281 Enterococcus faecalis V583 GCA_000007785.1
281 Streptomyces cinnamoneus GCA_001885705.1
253 Enterococcus canis NBRC 100695 GCA_001544375.1
253 Enterococcus dispar ATCC 51266 GCA_000406945.1
253 Enterococcus faecium DO GCA_000174395.2
253 Enterococcus haemoperoxidus ATCC BAA-382 GCA_000407165.1
253 Enterococcus hirae ATCC 9790 GCA_000271405.2
253 Enterococcus massiliensis GCA_001050095.1
253 Enterococcus mundtii QU 25 GCA_000504125.1
253 Enterococcus phoeniculicola ATCC BAA-412 GCA_000407505.1
253 Enterococcus rivorum GCA_001742285.1
253 Enterococcus thailandicus GCA_001652875.1
246 Enterococcus asini ATCC 700915 GCA_000407365.1
sm
151 Staphylococcus lentus F1142 GCA_000286395.1
151 Staphylococcus sciuri GCA_002209165.1
151 Staphylococcus vitulinus F1028 GCA_000286335.1
150 Bacillus anthracis str. Ames GCA_000007845.1
150 Bacillus anthracis str. Sterne GCA_000008165.1

150 Bacillus cereus ATCC 14579 GCA_000007825.1
150 Bacillus mycoides GCA_000832605.1
150 Bacillus pseudomycoides DSM 12442 GCA_000161455.1
150 Bacillus thuringiensis YBT-1518 GCA_000497525.2
150 _5BBacillus thuringiensis_5D serovar konkukian str. 97-27 GCA_000008505.1
147 Staphylococcus arlettae CVD059 GCA_000295715.1
147 Staphylococcus cohnii subsp. cohnii GCA_000972575.1
147 Staphylococcus equorum GCA_001432245.1
147 Staphylococcus gallinarum GCA_000875895.1
147 Staphylococcus pettenkoferi GCA_002208805.1
147 Staphylococcus saprophyticus GCA_001074355.1
147 Staphylococcus saprophyticus subsp. saprophyticus ATCC 15305 GCA_000010125.1
147 Staphylococcus succinus GCA_001902315.1
147 Staphylococcus xylosus GCA_000706685.1
sn
216 Enterococcus faecalis V583 GCA_000007785.1
216 Streptomyces cinnamoneus GCA_001885705.1
176 Enterococcus canis NBRC 100695 GCA_001544375.1
176 Enterococcus casseliflavus EC20 GCA_000157355.2
176 Enterococcus dispar ATCC 51266 GCA_000406945.1
176 Enterococcus faecium DO GCA_000174395.2
176 Enterococcus hirae ATCC 9790 GCA_000271405.2
176 Enterococcus massiliensis GCA_001050095.1
176 Enterococcus mundtii QU 25 GCA_000504125.1
176 Enterococcus rivorum GCA_001742285.1
176 Enterococcus saccharolyticus subsp. saccharolyticus ATCC 43076 GCA_000407285.1
171 Enterococcus gilvus ATCC BAA-350 GCA_000407545.1
171 Enterococcus hermanniensis GCA_001885945.1
171 Enterococcus malodoratus ATCC 43197 GCA_000407185.1
171 Enterococcus pallens ATCC BAA-351 GCA_000407485.1
171 Enterococcus pseudoavium NBRC 100491 GCA_001544295.1
so
227 Bifidobacterium adolescentis ATCC 15703 GCA_000010425.1
192 Bifidobacterium asteroides PRL2011 GCA_000304215.1
185 Bifidobacterium callitrichos DSM 23973 GCA_000741175.1
sp
241 Rhodobacter sphaeroides 2.4.1 GCA_000012905.2
216 Rhodobacter sphaeroides ATCC 17025 GCA_000016405.1
180 Rhodobacter capsulatus SB 1003 GCA_000021865.1
sq
319 Streptococcus mutans UA159 GCA_000007465.2
183 Streptococcus massiliensis DSM 18628 GCA_000380065.1
181 Streptococcus equinus GCA_000964315.1
181 Streptococcus gallolyticus subsp. gallolyticus DSM 16831 GCA_002000985.1
sr
228 Clostridium chromiireducens GCA_002029255.1
228 Clostridium saccharobutylicum DSM 13864 GCA_000473995.1
226 Clostridium beijerinckii GCA_000833105.2
226 Clostridium puniceum GCA_002006345.1
226 Clostridium saccharoperbutylacetonicum N1-4_28HMT_29 GCA_000340885.1
225 Clostridium botulinum B str. Eklund 17B _28NRP_29 GCA_000020165.1
225 Clostridium taeniosporum GCA_001735765.1
ss
255 Deinococcus radiodurans R1 GCA_000008565.1
139 Deinococcus puniceus GCA_001644565.1
117 Deinococcus hopiensis KR-140 GCA_900176165.1
st
311 Staphylococcus epidermidis ATCC 12228 GCA_000007645.1
308 Staphylococcus haemolyticus JCSC1435 GCA_000009865.1
302 Staphylococcus lugdunensis HKU09-01 GCA_000025085.1
su
151 Clostridium beijerinckii GCA_000833105.2
151 Clostridium puniceum GCA_002006345.1
151 Clostridium saccharoperbutylacetonicum N1-4_28HMT_29 GCA_000340885.1
147 Clostridium butyricum GCA_001456065.2
147 Clostridium taeniosporum GCA_001735765.1
142 Clostridium baratii str. Sullivan GCA_000789395.1
142 Clostridium saccharobutylicum DSM 13864 GCA_000473995.1
sv
234 Lactobacillus gasseri ATCC 33323 = JCM 1131 GCA_000014425.1
207 Lactobacillus hominis DSM 23910 = CRBIP 24.179 GCA_000296835.1
173 Lactobacillus antri DSM 16041 GCA_000160835.1
173 Lactobacillus brevis ATCC 367 GCA_000014465.1
173 Lactobacillus frumenti DSM 13145 GCA_001436045.1
173 Lactobacillus ingluviei str. Autruche 4 GCA_000312405.1
173 Lactobacillus koreensis GCA_001050435.1

173 Lactobacillus paucivorans GCA_001437125.1
173 Lactobacillus reuteri DSM 20016 GCA_000016825.1
173 Lactobacillus secaliphilus GCA_001437055.1
173 Lactobacillus senmaizukei DSM 21775 = NBRC 103853 GCA_001592085.1
173 Lactobacillus spicheri DSM 15429 GCA_001435095.1
173 Lactobacillus vaginalis DSM 5837 = ATCC 49540 GCA_000159435.1
173 Lactobacillus zymae GCA_900183405.1
sw
223 Streptococcus mutans UA159 GCA_000007465.2
116 Streptococcus marimammalium DSM 18627 GCA_000380045.1
112 Streptococcus sobrinus DSM 20742 = ATCC 33478 GCA_000686605.1
sx
214 Rhodobacter sphaeroides 2.4.1 GCA_000012905.2
211 Rhodobacter sphaeroides ATCC 17025 GCA_000016405.1
195 Pseudorhodobacter psychrotolerans GCA_001294535.1
sy
248 Clostridium beijerinckii GCA_000833105.2
248 Clostridium saccharoperbutylacetonicum N1-4_28HMT_29 GCA_000340885.1
240 Clostridium saccharobutylicum DSM 13864 GCA_000473995.1
234 Clostridium puniceum GCA_002006345.1
sz
299 Deinococcus radiodurans R1 GCA_000008565.1
230 Deinococcus gobiensis I-0 GCA_000252445.1
182 Deinococcus deserti VCD115 GCA_000020685.1
sA
280 Lactobacillus gasseri ATCC 33323 = JCM 1131 GCA_000014425.1
259 Lactobacillus hominis DSM 23910 = CRBIP 24.179 GCA_000296835.1
203 Lactobacillus psittaci DSM 15354 GCA_000425905.1
sB
215 Staphylococcus capitis subsp. capitis GCA_001028645.1
215 Staphylococcus epidermidis ATCC 12228 GCA_000007645.1
212 Staphylococcus haemolyticus JCSC1435 GCA_000009865.1
206 Megasphaera cerevisiae DSM 20462 GCA_001045675.1
206 Staphylococcus lugdunensis HKU09-01 GCA_000025085.1
206 Staphylococcus warneri SG1 GCA_000332735.1
sC
166 Deinococcus radiodurans R1 GCA_000008565.1
111 Deinococcus geothermalis DSM 11300 GCA_000196275.1
98 Deinococcus marmoris DSM 12784 GCA_000701405.1
98 Deinococcus swuensis GCA_000800395.1
sD
245 Lactobacillus gasseri ATCC 33323 = JCM 1131 GCA_000014425.1
238 Lactobacillus hominis DSM 23910 = CRBIP 24.179 GCA_000296835.1
208 Lactobacillus crispatus ST1 GCA_000091765.1
sE
274 Escherichia coli IAI39 GCA_000026345.1
274 Escherichia coli O104_3AH4 str. 2011C-3493 GCA_000299455.1
274 Escherichia coli O157_3AH7 str. Sakai GCA_000008865.1
274 Escherichia coli O83_3AH1 str. NRG 857C GCA_000183345.1
274 Escherichia coli UMN026 GCA_000026325.2
274 Escherichia coli str. K-12 substr. MG1655 GCA_000005845.2
274 Shigella dysenteriae Sd197 GCA_000012005.1
274 Shigella flexneri 2a str. 301 GCA_000006925.2
274 Tumebacillus flagellatus GCA_000714935.1
238 Erwinia iniecta GCA_001267535.1
213 Rosenbergiella nectarea GCA_900111105.1
sF
219 Bacillus anthracis str. Ames GCA_000007845.1
219 Bacillus anthracis str. Sterne GCA_000008165.1
219 Bacillus cereus ATCC 14579 GCA_000007825.1
219 Bacillus mycoides GCA_000832605.1
219 Bacillus pseudomycoides DSM 12442 GCA_000161455.1
219 Bacillus thuringiensis YBT-1518 GCA_000497525.2
219 _5BBacillus thuringiensis_5D serovar konkukian str. 97-27 GCA_000008505.1
180 Bacillus acidicola GCA_001636425.1
180 Bacillus shackletonii GCA_001420715.1
178 Bacillus cohnii NBRC 15565 GCA_001591425.1
sG
221 Clostridium beijerinckii GCA_000833105.2
221 Clostridium puniceum GCA_002006345.1
221 Clostridium saccharoperbutylacetonicum N1-4_28HMT_29 GCA_000340885.1
212 Clostridium saccharobutylicum DSM 13864 GCA_000473995.1
202 Clostridium butyricum GCA_001456065.2
sH
229 Rhodobacter sphaeroides 2.4.1 GCA_000012905.2
211 Rhodobacter sphaeroides ATCC 17025 GCA_000016405.1

210 Defluviimonas alba GCA_001620265.1
sI
179 Bacillus thuringiensis YBT-1518 GCA_000497525.2
164 Bacillus anthracis str. Ames GCA_000007845.1
164 Bacillus anthracis str. Sterne GCA_000008165.1
164 Bacillus cereus ATCC 14579 GCA_000007825.1
164 Bacillus mycoides GCA_000832605.1
164 Bacillus pseudomycoides DSM 12442 GCA_000161455.1
164 _5BBacillus thuringiensis_5D serovar konkukian str. 97-27 GCA_000008505.1
133 Bacillus flexus GCA_002024265.1
133 Bacillus megaterium NBRC 15308 = ATCC 14581 GCA_000832985.1
133 Thalassospira mesophila GCA_002115755.1
sJ
202 Deinococcus radiodurans R1 GCA_000008565.1
183 Deinococcus puniceus GCA_001644565.1
173 Deinococcus gobiensis I-0 GCA_000252445.1
sK
246 Deinococcus radiodurans R1 GCA_000008565.1
184 Deinococcus puniceus GCA_001644565.1
171 Deinococcus gobiensis I-0 GCA_000252445.1
sL
266 Staphylococcus lutrae GCA_002101335.1
265 Staphylococcus equorum GCA_001432245.1
264 Bacillus anthracis str. Ames GCA_000007845.1
264 Bacillus anthracis str. Sterne GCA_000008165.1
264 Bacillus mycoides GCA_000832605.1
264 Bacillus pseudomycoides DSM 12442 GCA_000161455.1
264 Bacillus thuringiensis YBT-1518 GCA_000497525.2
264 _5BBacillus thuringiensis_5D serovar konkukian str. 97-27 GCA_000008505.1
sM
135 Streptococcus mutans UA159 GCA_000007465.2
110 Streptococcus cristatus AS 1.3089 GCA_000385925.1
110 Streptococcus parasanguinis ATCC 15912 GCA_000164675.2
99 Streptococcus gordonii str. Challis substr. CH1 GCA_000017005.1
99 Streptococcus mitis B6 GCA_000027165.1
99 Streptococcus pneumoniae R6 GCA_000007045.1
sN
244 Deinococcus radiodurans R1 GCA_000008565.1
166 Deinococcus puniceus GCA_001644565.1
155 Deinococcus soli Cha et al. 2016 GCA_001007995.1
sO
195 Lactobacillus gasseri ATCC 33323 = JCM 1131 GCA_000014425.1
186 Lactobacillus hominis DSM 23910 = CRBIP 24.179 GCA_000296835.1
144 Lactobacillus iners DSM 13335 GCA_000160875.1
sP
123 Escherichia coli IAI39 GCA_000026345.1
123 Escherichia coli O104_3AH4 str. 2011C-3493 GCA_000299455.1
123 Escherichia coli O157_3AH7 str. Sakai GCA_000008865.1
123 Escherichia coli O83_3AH1 str. NRG 857C GCA_000183345.1
123 Escherichia coli UMN026 GCA_000026325.2
123 Escherichia coli str. K-12 substr. MG1655 GCA_000005845.2
123 Shigella dysenteriae Sd197 GCA_000012005.1
123 Shigella flexneri 2a str. 301 GCA_000006925.2
123 Tumebacillus flagellatus GCA_000714935.1
102 Pantoea agglomerans GCA_001709315.1
101 Cronobacter sakazakii GCA_000982825.1
101 Enterobacter cloacae subsp. cloacae ATCC 13047 GCA_000025565.1
101 Enterobacter hormaechei subsp. steigerwaltii GCA_001729725.1
101 Enterobacter kobei GCA_900185885.1
101 Erwinia amylovora CFBP1430 GCA_000091565.1
101 Erwinia iniecta GCA_001267535.1
101 Erwinia persicina NBRC 102418 GCA_001571305.1
101 Erwinia toletana DAPP-PG 735 GCA_000336255.1
101 Klebsiella oxytoca GCA_001022195.1
101 Klebsiella pneumoniae subsp. pneumoniae HS11286 GCA_000240185.2
101 Kosakonia cowanii GCA_001975225.1
101 Kosakonia sacchari SP1 GCA_000300455.4
101 Pantoea alhagi GCA_002101395.1
101 Pantoea ananatis LMG 20103 GCA_000025405.2
101 Pantoea dispersa EGD-AAK13 GCA_000465555.2
101 Pantoea rwandensis GCA_000759475.1
101 Pantoea septica GCA_002095575.1
101 Plautia stali symbiont GCA_000180175.2
101 Proteus mirabilis HI4320 GCA_000069965.1
101 Pseudescherichia vulneris NBRC 102420 GCA_000759795.1
101 Salmonella enterica subsp. enterica serovar Typhi str. CT18 GCA_000195995.1

101 Salmonella enterica subsp. enterica serovar Typhimurium str. LT2 GCA_000006945.2
101 Shimwellia blattae DSM 4481 = NBRC 105725 GCA_000262305.1
101 Trabulsiella odontotermitis GCA_001297765.1
101 Xenorhabdus bovienii SS-2004 GCA_000027225.1
101 Xenorhabdus cabanillasii JM26 GCA_000531755.1
101 Xenorhabdus doucetiae GCA_000968195.1
101 Xenorhabdus hominickii GCA_001721185.1
101 Xenorhabdus innexi GCA_900155355.1
101 Xenorhabdus japonica GCA_900115195.1
101 Xenorhabdus koppenhoeferi GCA_900116635.1
101 Xenorhabdus nematophila AN6_2F1 GCA_000953355.1
sQ
273 Enterococcus faecalis V583 GCA_000007785.1
273 Streptomyces cinnamoneus GCA_001885705.1
243 Enterococcus dispar ATCC 51266 GCA_000406945.1
240 Enterococcus asini ATCC 700915 GCA_000407365.1
240 Enterococcus canis NBRC 100695 GCA_001544375.1
240 Enterococcus casseliflavus EC20 GCA_000157355.2
240 Enterococcus faecium DO GCA_000174395.2
240 Enterococcus hirae ATCC 9790 GCA_000271405.2
240 Enterococcus mundtii QU 25 GCA_000504125.1
240 Enterococcus rivorum GCA_001742285.1
240 Enterococcus saccharolyticus subsp. saccharolyticus ATCC 43076 GCA_000407285.1
sR
177 Bacillus thuringiensis YBT-1518 GCA_000497525.2
157 Bacillus mycoides GCA_000832605.1
156 Bacillus anthracis str. Ames GCA_000007845.1
156 Bacillus anthracis str. Sterne GCA_000008165.1
156 Bacillus cereus ATCC 14579 GCA_000007825.1
156 Bacillus pseudomycoides DSM 12442 GCA_000161455.1
156 _5BBacillus thuringiensis_5D serovar konkukian str. 97-27 GCA_000008505.1
sS
234 Deinococcus radiodurans R1 GCA_000008565.1
131 Deinococcus marmoris DSM 12784 GCA_000701405.1
131 Deinococcus swuensis GCA_000800395.1
123 Deinococcus proteolyticus MRP GCA_000190555.1
sT
122 Rhodobacter sphaeroides 2.4.1 GCA_000012905.2
104 Rhodobacter sphaeroides ATCC 17025 GCA_000016405.1
86 Roseivivax halotolerans GCA_900115815.1
sU
140 Bacillus mycoides GCA_000832605.1
130 Staphylococcus vitulinus F1028 GCA_000286335.1
125 Bacillus anthracis str. Ames GCA_000007845.1
125 Bacillus anthracis str. Sterne GCA_000008165.1
125 Bacillus cereus ATCC 14579 GCA_000007825.1
125 Bacillus pseudomycoides DSM 12442 GCA_000161455.1
125 Bacillus solimangrovi GCA_001742425.1
125 Bacillus thuringiensis YBT-1518 GCA_000497525.2
125 _5BBacillus thuringiensis_5D serovar konkukian str. 97-27 GCA_000008505.1
sV
219 Rhodobacter sphaeroides 2.4.1 GCA_000012905.2
219 Rhodobacter sphaeroides ATCC 17025 GCA_000016405.1
209 Pseudorhodobacter ferrugineus DSM 5888 GCA_000420745.1
209 Pseudorhodobacter wandonensis GCA_001202035.1
194 Gemmobacter megaterium GCA_900156815.1
194 Pseudorhodobacter psychrotolerans GCA_001294535.1
sW
297 Deinococcus radiodurans R1 GCA_000008565.1
226 Deinococcus deserti VCD115 GCA_000020685.1
219 Deinococcus soli Cha et al. 2016 GCA_001007995.1
sX
189 Staphylococcus epidermidis ATCC 12228 GCA_000007645.1
186 Megasphaera cerevisiae DSM 20462 GCA_001045675.1
186 Staphylococcus warneri SG1 GCA_000332735.1
184 Staphylococcus capitis subsp. capitis GCA_001028645.1
sY
109 Clostridium beijerinckii GCA_000833105.2
109 Clostridium puniceum GCA_002006345.1
109 Clostridium saccharobutylicum DSM 13864 GCA_000473995.1
109 Clostridium saccharoperbutylacetonicum N1-4_28HMT_29 GCA_000340885.1
91 Clostridium butyricum GCA_001456065.2
91 Clostridium chromiireducens GCA_002029255.1
78 Clostridium taeniosporum GCA_001735765.1
sZ
352 Bacillus anthracis str. Ames GCA_000007845.1

352 Bacillus anthracis str. Sterne GCA_000008165.1
352 Bacillus cereus ATCC 14579 GCA_000007825.1
352 Bacillus mycoides GCA_000832605.1
352 Bacillus pseudomycoides DSM 12442 GCA_000161455.1
352 Bacillus thuringiensis YBT-1518 GCA_000497525.2
352 _5BBacillus thuringiensis_5D serovar konkukian str. 97-27 GCA_000008505.1
276 Bacillus acidicola GCA_001636425.1
276 Bacillus shackletonii GCA_001420715.1
274 Bacillus koreensis GCA_001274935.1
t0
143 Lactobacillus gasseri ATCC 33323 = JCM 1131 GCA_000014425.1
129 Lactobacillus hominis DSM 23910 = CRBIP 24.179 GCA_000296835.1
77 Lactobacillus collinoides DSM 20515 = JCM 1123 GCA_001435975.1
77 Lactobacillus coryniformis subsp. coryniformis KCTC 3167 = DSM 20001 GCA_001433765.1
77 Lactobacillus paracollinoides GCA_001702175.1
77 Lactobacillus rhamnosus GG GCA_000026505.1
t1
256 Deinococcus radiodurans R1 GCA_000008565.1
160 Deinococcus gobiensis I-0 GCA_000252445.1
153 Deinococcus deserti VCD115 GCA_000020685.1
t2
199 Deinococcus radiodurans R1 GCA_000008565.1
123 Deinococcus puniceus GCA_001644565.1
118 Deinococcus marmoris DSM 12784 GCA_000701405.1
118 Deinococcus swuensis GCA_000800395.1
t3
130 Clostridium cellulovorans 743B GCA_000145275.1
125 Clostridium beijerinckii GCA_000833105.2
125 Clostridium chromiireducens GCA_002029255.1
125 Clostridium puniceum GCA_002006345.1
125 Clostridium saccharobutylicum DSM 13864 GCA_000473995.1
125 Clostridium saccharoperbutylacetonicum N1-4_28HMT_29 GCA_000340885.1
123 Clostridium akagii DSM 12554 GCA_000686725.1
123 Clostridium amylolyticum GCA_900142075.1
123 Clostridium arbusti SL206 GCA_000246895.2
123 Clostridium butyricum GCA_001456065.2
123 Clostridium celatum DSM 1785 GCA_000320405.1
123 Clostridium chauvoei JF4335 GCA_900168365.1
123 Clostridium fallax GCA_900129365.1
123 Clostridium gasigenes GCA_900104115.1
123 Clostridium neonatale GCA_001458595.1
123 Clostridium pasteurianum BC1 GCA_000389635.1
123 Clostridium pasteurianum DSM 525 = ATCC 6013 GCA_000807255.1
123 Clostridium pasteurianum GCA_001705235.1
123 Clostridium perfringens ATCC 13124 GCA_000013285.1
123 Clostridium polynesiense GCA_000820705.1
123 Clostridium sartagoforme AAU1 GCA_000401215.1
123 Clostridium ventriculi GCA_001404895.1
t4
119 Deinococcus radiodurans R1 GCA_000008565.1
90 Deinococcus hopiensis KR-140 GCA_900176165.1
90 Deinococcus puniceus GCA_001644565.1
79 Deinococcus marmoris DSM 12784 GCA_000701405.1
79 Deinococcus misasensis DSM 22328 GCA_000745915.1
79 Deinococcus swuensis GCA_000800395.1
t5
188 Staphylococcus capitis subsp. capitis GCA_001028645.1
188 Staphylococcus epidermidis ATCC 12228 GCA_000007645.1
188 Staphylococcus haemolyticus JCSC1435 GCA_000009865.1
178 Staphylococcus condimenti GCA_001618885.1
178 Staphylococcus simulans GCA_001559115.1
172 Staphylococcus hominis subsp. hominis C80 GCA_000183685.1
t6
222 Clostridium beijerinckii GCA_000833105.2
222 Clostridium saccharoperbutylacetonicum N1-4_28HMT_29 GCA_000340885.1
215 Clostridium puniceum GCA_002006345.1
207 Clostridium saccharobutylicum DSM 13864 GCA_000473995.1
t7
233 Escherichia coli IAI39 GCA_000026345.1
233 Escherichia coli O104_3AH4 str. 2011C-3493 GCA_000299455.1
233 Escherichia coli O157_3AH7 str. Sakai GCA_000008865.1
233 Escherichia coli O83_3AH1 str. NRG 857C GCA_000183345.1
233 Escherichia coli UMN026 GCA_000026325.2
233 Escherichia coli str. K-12 substr. MG1655 GCA_000005845.2
233 Shigella flexneri 2a str. 301 GCA_000006925.2
233 Tumebacillus flagellatus GCA_000714935.1

224 Shigella dysenteriae Sd197 GCA_000012005.1
211 Erwinia iniecta GCA_001267535.1
t8
244 Clostridium beijerinckii GCA_000833105.2
244 Clostridium saccharoperbutylacetonicum N1-4_28HMT_29 GCA_000340885.1
240 Clostridium saccharobutylicum DSM 13864 GCA_000473995.1
229 Clostridium puniceum GCA_002006345.1
t9
222 Bifidobacterium adolescentis ATCC 15703 GCA_000010425.1
214 Bifidobacterium asteroides PRL2011 GCA_000304215.1
203 Bifidobacterium dentium JCM 1195 = DSM 20436 GCA_001042595.1
ta
232 Erwinia iniecta GCA_001267535.1
232 Escherichia coli IAI39 GCA_000026345.1
232 Escherichia coli O104_3AH4 str. 2011C-3493 GCA_000299455.1
232 Escherichia coli O157_3AH7 str. Sakai GCA_000008865.1
232 Escherichia coli O83_3AH1 str. NRG 857C GCA_000183345.1
232 Escherichia coli UMN026 GCA_000026325.2
232 Escherichia coli str. K-12 substr. MG1655 GCA_000005845.2
232 Shigella flexneri 2a str. 301 GCA_000006925.2
232 Tumebacillus flagellatus GCA_000714935.1
217 Shigella dysenteriae Sd197 GCA_000012005.1
197 Cronobacter sakazakii GCA_000982825.1
197 Erwinia toletana DAPP-PG 735 GCA_000336255.1
197 Kosakonia cowanii GCA_001975225.1
197 Kosakonia sacchari SP1 GCA_000300455.4
197 Rosenbergiella nectarea GCA_900111105.1
tb
240 Deinococcus radiodurans R1 GCA_000008565.1
177 Deinococcus deserti VCD115 GCA_000020685.1
165 Deinococcus soli Cha et al. 2016 GCA_001007995.1
tc
227 Staphylococcus capitis subsp. capitis GCA_001028645.1
227 Staphylococcus epidermidis ATCC 12228 GCA_000007645.1
227 Staphylococcus haemolyticus JCSC1435 GCA_000009865.1
216 Staphylococcus cohnii subsp. cohnii GCA_000972575.1
216 Staphylococcus hominis subsp. hominis C80 GCA_000183685.1
207 Staphylococcus aureus subsp. aureus NCTC 8325 GCA_000013425.1
207 Staphylococcus simiae CCM 7213 GCA_000235645.2
207 Staphylococcus succinus GCA_001902315.1
td
193 Clostridium beijerinckii GCA_000833105.2
193 Clostridium puniceum GCA_002006345.1
193 Clostridium saccharobutylicum DSM 13864 GCA_000473995.1
193 Clostridium saccharoperbutylacetonicum N1-4_28HMT_29 GCA_000340885.1
180 Clostridium butyricum GCA_001456065.2
174 Clostridium chromiireducens GCA_002029255.1
te
226 Clostridium beijerinckii GCA_000833105.2
226 Clostridium puniceum GCA_002006345.1
226 Clostridium saccharobutylicum DSM 13864 GCA_000473995.1
226 Clostridium saccharoperbutylacetonicum N1-4_28HMT_29 GCA_000340885.1
222 Clostridium chromiireducens GCA_002029255.1
209 Clostridium botulinum B str. Eklund 17B _28NRP_29 GCA_000020165.1
209 Clostridium taeniosporum GCA_001735765.1
tf
242 Enterococcus faecalis V583 GCA_000007785.1
242 Streptomyces cinnamoneus GCA_001885705.1
231 Enterococcus hirae ATCC 9790 GCA_000271405.2
230 Enterococcus dispar ATCC 51266 GCA_000406945.1
tg
105 Bacillus anthracis str. Ames GCA_000007845.1
105 Bacillus anthracis str. Sterne GCA_000008165.1
105 Bacillus cereus ATCC 14579 GCA_000007825.1
105 Bacillus manliponensis GCA_000712595.1
105 Bacillus mycoides GCA_000832605.1
105 Bacillus pseudomycoides DSM 12442 GCA_000161455.1
105 Bacillus thuringiensis YBT-1518 GCA_000497525.2
105 Paenirhodobacter enshiensis GCA_000740785.1
105 _5BBacillus thuringiensis_5D serovar konkukian str. 97-27 GCA_000008505.1
96 Escherichia coli IAI39 GCA_000026345.1
96 Escherichia coli O104_3AH4 str. 2011C-3493 GCA_000299455.1
96 Escherichia coli O157_3AH7 str. Sakai GCA_000008865.1
96 Escherichia coli O83_3AH1 str. NRG 857C GCA_000183345.1
96 Escherichia coli UMN026 GCA_000026325.2
96 Escherichia coli str. K-12 substr. MG1655 GCA_000005845.2

96 Shigella dysenteriae Sd197 GCA_000012005.1
96 Shigella flexneri 2a str. 301 GCA_000006925.2
96 Tumebacillus flagellatus GCA_000714935.1
95 Bacillus halmapalus GCA_002019665.1
th
250 Rhodobacter sphaeroides 2.4.1 GCA_000012905.2
220 Rhodobacter sphaeroides ATCC 17025 GCA_000016405.1
196 Gemmobacter megaterium GCA_900156815.1
ti
257 Bifidobacterium adolescentis ATCC 15703 GCA_000010425.1
241 Bifidobacterium tsurumiense GCA_000741765.1
233 Bifidobacterium angulatum DSM 20098 = JCM 7096 GCA_001025155.1
tj
229 Clostridium beijerinckii GCA_000833105.2
229 Clostridium puniceum GCA_002006345.1
229 Clostridium saccharobutylicum DSM 13864 GCA_000473995.1
229 Clostridium saccharoperbutylacetonicum N1-4_28HMT_29 GCA_000340885.1
215 Clostridium chromiireducens GCA_002029255.1
215 Clostridium neonatale GCA_001458595.1
203 Clostridium butyricum GCA_001456065.2
tk
220 Streptococcus mutans UA159 GCA_000007465.2
174 Streptococcus ratti FA-1 = DSM 20564 GCA_000286075.1
173 Streptococcus gordonii str. Challis substr. CH1 GCA_000017005.1
tl
212 Enterococcus faecalis V583 GCA_000007785.1
212 Streptomyces cinnamoneus GCA_001885705.1
170 Enterococcus canis NBRC 100695 GCA_001544375.1
170 Enterococcus faecium DO GCA_000174395.2
170 Enterococcus haemoperoxidus ATCC BAA-382 GCA_000407165.1
170 Enterococcus hirae ATCC 9790 GCA_000271405.2
170 Enterococcus mundtii QU 25 GCA_000504125.1
170 Enterococcus phoeniculicola ATCC BAA-412 GCA_000407505.1
170 Enterococcus rivorum GCA_001742285.1
170 Enterococcus thailandicus GCA_001652875.1
169 Enterococcus casseliflavus EC20 GCA_000157355.2
169 Enterococcus dispar ATCC 51266 GCA_000406945.1
169 Enterococcus gilvus ATCC BAA-350 GCA_000407545.1
169 Enterococcus hermanniensis GCA_001885945.1
169 Enterococcus malodoratus ATCC 43197 GCA_000407185.1
169 Enterococcus pallens ATCC BAA-351 GCA_000407485.1
169 Enterococcus pseudoavium NBRC 100491 GCA_001544295.1
169 Enterococcus saccharolyticus subsp. saccharolyticus ATCC 43076 GCA_000407285.1
169 Lactobacillus sharpeae JCM 1186 = DSM 20505 GCA_001436225.1
tm
198 Rhodobacter sphaeroides 2.4.1 GCA_000012905.2
186 Pseudorhodobacter psychrotolerans GCA_001294535.1
186 Rhodobacter sphaeroides ATCC 17025 GCA_000016405.1
176 Gemmobacter aquatilis GCA_900110025.1
176 Gemmobacter nectariphilus DSM 15620 GCA_000429765.1
176 Haematobacter massiliensis GCA_000740795.1
176 Rhodobacter capsulatus SB 1003 GCA_000021865.1
tn
236 Staphylococcus capitis subsp. capitis GCA_001028645.1
236 Staphylococcus epidermidis ATCC 12228 GCA_000007645.1
236 Staphylococcus haemolyticus JCSC1435 GCA_000009865.1
236 Staphylococcus hominis subsp. hominis C80 GCA_000183685.1
236 Staphylococcus lugdunensis HKU09-01 GCA_000025085.1
227 Staphylococcus aureus subsp. aureus NCTC 8325 GCA_000013425.1
227 Staphylococcus condimenti GCA_001618885.1
227 Staphylococcus microti GCA_000934465.1
227 Staphylococcus simiae CCM 7213 GCA_000235645.2
227 Staphylococcus simulans GCA_001559115.1
224 Staphylococcus arlettae CVD059 GCA_000295715.1
224 Staphylococcus cohnii subsp. cohnii GCA_000972575.1
224 Staphylococcus saprophyticus subsp. saprophyticus ATCC 15305 GCA_000010125.1
to
171 Escherichia coli IAI39 GCA_000026345.1
171 Escherichia coli O104_3AH4 str. 2011C-3493 GCA_000299455.1
171 Escherichia coli O157_3AH7 str. Sakai GCA_000008865.1
171 Escherichia coli O83_3AH1 str. NRG 857C GCA_000183345.1
171 Escherichia coli UMN026 GCA_000026325.2
171 Escherichia coli str. K-12 substr. MG1655 GCA_000005845.2
171 Shigella flexneri 2a str. 301 GCA_000006925.2
171 Tumebacillus flagellatus GCA_000714935.1
156 Shigella dysenteriae Sd197 GCA_000012005.1

138 Erwinia iniecta GCA_001267535.1
tp
266 Deinococcus radiodurans R1 GCA_000008565.1
164 Deinococcus puniceus GCA_001644565.1
162 Deinococcus gobiensis I-0 GCA_000252445.1
tq
208 Escherichia coli IAI39 GCA_000026345.1
208 Escherichia coli O104_3AH4 str. 2011C-3493 GCA_000299455.1
208 Escherichia coli O157_3AH7 str. Sakai GCA_000008865.1
208 Escherichia coli O83_3AH1 str. NRG 857C GCA_000183345.1
208 Escherichia coli UMN026 GCA_000026325.2
208 Escherichia coli str. K-12 substr. MG1655 GCA_000005845.2
208 Shigella dysenteriae Sd197 GCA_000012005.1
208 Shigella flexneri 2a str. 301 GCA_000006925.2
208 Tumebacillus flagellatus GCA_000714935.1
168 Erwinia iniecta GCA_001267535.1
161 Enterobacter cloacae subsp. cloacae ATCC 13047 GCA_000025565.1
tr
185 Streptococcus mutans UA159 GCA_000007465.2
132 Streptococcus macacae NCTC 11558 GCA_000187995.3
129 Streptococcus ratti FA-1 = DSM 20564 GCA_000286075.1
ts
250 Bacillus thuringiensis YBT-1518 GCA_000497525.2
249 Bacillus anthracis str. Ames GCA_000007845.1
249 Bacillus anthracis str. Sterne GCA_000008165.1
249 Bacillus cereus ATCC 14579 GCA_000007825.1
249 Bacillus mycoides GCA_000832605.1
249 Bacillus pseudomycoides DSM 12442 GCA_000161455.1
249 _5BBacillus thuringiensis_5D serovar konkukian str. 97-27 GCA_000008505.1
170 Bacillus cytotoxicus NVH 391-98 GCA_000017425.1
tt
288 Bifidobacterium adolescentis ATCC 15703 GCA_000010425.1
259 Bifidobacterium thermophilum GCA_000741495.1
257 Bifidobacterium callitrichos DSM 23973 GCA_000741175.1
tu
175 Streptococcus mutans UA159 GCA_000007465.2
114 Streptococcus equinus GCA_000964315.1
114 Streptococcus gallolyticus subsp. gallolyticus DSM 16831 GCA_002000985.1
114 Streptococcus orisratti DSM 15617 GCA_000380105.1
114 Streptococcus ratti FA-1 = DSM 20564 GCA_000286075.1
104 Streptococcus massiliensis DSM 18628 GCA_000380065.1
tv
203 Escherichia coli IAI39 GCA_000026345.1
203 Escherichia coli O104_3AH4 str. 2011C-3493 GCA_000299455.1
203 Escherichia coli O157_3AH7 str. Sakai GCA_000008865.1
203 Escherichia coli UMN026 GCA_000026325.2
203 Escherichia coli str. K-12 substr. MG1655 GCA_000005845.2
203 Shigella flexneri 2a str. 301 GCA_000006925.2
203 Tumebacillus flagellatus GCA_000714935.1
202 Escherichia coli O83_3AH1 str. NRG 857C GCA_000183345.1
194 Shigella dysenteriae Sd197 GCA_000012005.1
tw
314 Deinococcus radiodurans R1 GCA_000008565.1
212 Deinococcus deserti VCD115 GCA_000020685.1
208 Deinococcus gobiensis I-0 GCA_000252445.1
tx
173 Lactobacillus gasseri ATCC 33323 = JCM 1131 GCA_000014425.1
173 Lactobacillus hominis DSM 23910 = CRBIP 24.179 GCA_000296835.1
146 Lactobacillus iners DSM 13335 GCA_000160875.1
141 Lactobacillus psittaci DSM 15354 GCA_000425905.1
ty
202 Deinococcus radiodurans R1 GCA_000008565.1
119 Deinococcus gobiensis I-0 GCA_000252445.1
117 Deinococcus deserti VCD115 GCA_000020685.1
tz
191 Streptococcus mutans UA159 GCA_000007465.2
107 Streptococcus ferus DSM 20646 GCA_000372425.1
105 Streptococcus marimammalium DSM 18627 GCA_000380045.1
tA
216 Deinococcus radiodurans R1 GCA_000008565.1
121 Deinococcus puniceus GCA_001644565.1
117 Deinococcus deserti VCD115 GCA_000020685.1
117 Deinococcus marmoris DSM 12784 GCA_000701405.1
117 Deinococcus swuensis GCA_000800395.1
tB
313 Escherichia coli IAI39 GCA_000026345.1

313 Escherichia coli O104_3AH4 str. 2011C-3493 GCA_000299455.1
313 Escherichia coli O157_3AH7 str. Sakai GCA_000008865.1
313 Escherichia coli O83_3AH1 str. NRG 857C GCA_000183345.1
313 Escherichia coli UMN026 GCA_000026325.2
313 Escherichia coli str. K-12 substr. MG1655 GCA_000005845.2
313 Shigella flexneri 2a str. 301 GCA_000006925.2
313 Tumebacillus flagellatus GCA_000714935.1
307 Erwinia iniecta GCA_001267535.1
304 Shigella dysenteriae Sd197 GCA_000012005.1
tC
185 Lactobacillus gasseri ATCC 33323 = JCM 1131 GCA_000014425.1
170 Lactobacillus hominis DSM 23910 = CRBIP 24.179 GCA_000296835.1
141 Lactobacillus amylophilus DSM 20533 = JCM 1125 GCA_001936335.1
tD
239 Staphylococcus epidermidis ATCC 12228 GCA_000007645.1
236 Staphylococcus cohnii subsp. cohnii GCA_000972575.1
236 Staphylococcus haemolyticus JCSC1435 GCA_000009865.1
236 Staphylococcus lugdunensis HKU09-01 GCA_000025085.1
231 Megasphaera cerevisiae DSM 20462 GCA_001045675.1
231 Staphylococcus pettenkoferi GCA_002208805.1
231 Staphylococcus warneri SG1 GCA_000332735.1
tE
218 Deinococcus radiodurans R1 GCA_000008565.1
164 Deinococcus gobiensis I-0 GCA_000252445.1
141 Deinococcus deserti VCD115 GCA_000020685.1
tF
275 Lactobacillus gasseri ATCC 33323 = JCM 1131 GCA_000014425.1
256 Lactobacillus hominis DSM 23910 = CRBIP 24.179 GCA_000296835.1
210 Lactobacillus iners DSM 13335 GCA_000160875.1
tG
287 Clostridium beijerinckii GCA_000833105.2
287 Clostridium saccharobutylicum DSM 13864 GCA_000473995.1
287 Clostridium saccharoperbutylacetonicum N1-4_28HMT_29 GCA_000340885.1
281 Clostridium puniceum GCA_002006345.1
266 Clostridium chromiireducens GCA_002029255.1
tH
301 Bacillus anthracis str. Ames GCA_000007845.1
301 Bacillus anthracis str. Sterne GCA_000008165.1
301 Bacillus cereus ATCC 14579 GCA_000007825.1
301 Bacillus mycoides GCA_000832605.1
301 Bacillus pseudomycoides DSM 12442 GCA_000161455.1
301 Bacillus thuringiensis YBT-1518 GCA_000497525.2
301 _5BBacillus thuringiensis_5D serovar konkukian str. 97-27 GCA_000008505.1
217 Bacillus korlensis NBRC 107688 GCA_001591645.1
216 Bacillus acidicola GCA_001636425.1
216 Bacillus shackletonii GCA_001420715.1
tI
212 Clostridium beijerinckii GCA_000833105.2
212 Clostridium puniceum GCA_002006345.1
212 Clostridium saccharoperbutylacetonicum N1-4_28HMT_29 GCA_000340885.1
199 Clostridium saccharobutylicum DSM 13864 GCA_000473995.1
193 Clostridium butyricum GCA_001456065.2
tJ
94 Clostridium beijerinckii GCA_000833105.2
94 Clostridium puniceum GCA_002006345.1
94 Clostridium saccharobutylicum DSM 13864 GCA_000473995.1
94 Clostridium saccharoperbutylacetonicum N1-4_28HMT_29 GCA_000340885.1
93 Clostridium neonatale GCA_001458595.1
80 Clostridium butyricum GCA_001456065.2
80 Clostridium chromiireducens GCA_002029255.1
tK
193 Escherichia coli IAI39 GCA_000026345.1
193 Escherichia coli O104_3AH4 str. 2011C-3493 GCA_000299455.1
193 Escherichia coli O157_3AH7 str. Sakai GCA_000008865.1
193 Escherichia coli O83_3AH1 str. NRG 857C GCA_000183345.1
193 Escherichia coli UMN026 GCA_000026325.2
193 Escherichia coli str. K-12 substr. MG1655 GCA_000005845.2
193 Shigella dysenteriae Sd197 GCA_000012005.1
193 Shigella flexneri 2a str. 301 GCA_000006925.2
193 Tumebacillus flagellatus GCA_000714935.1
174 Erwinia iniecta GCA_001267535.1
159 Cronobacter sakazakii GCA_000982825.1
tL
275 Rhodobacter sphaeroides 2.4.1 GCA_000012905.2
251 Rhodobacter sphaeroides ATCC 17025 GCA_000016405.1
233 Pseudorhodobacter psychrotolerans GCA_001294535.1

tM
216 Deinococcus radiodurans R1 GCA_000008565.1
146 Deinococcus deserti VCD115 GCA_000020685.1
139 Deinococcus gobiensis I-0 GCA_000252445.1
tN
313 Deinococcus radiodurans R1 GCA_000008565.1
227 Deinococcus gobiensis I-0 GCA_000252445.1
201 Deinococcus puniceus GCA_001644565.1
tO
261 Enterococcus faecalis V583 GCA_000007785.1
261 Streptomyces cinnamoneus GCA_001885705.1
228 Enterococcus asini ATCC 700915 GCA_000407365.1
228 Enterococcus canis NBRC 100695 GCA_001544375.1
228 Enterococcus casseliflavus EC20 GCA_000157355.2
228 Enterococcus dispar ATCC 51266 GCA_000406945.1
228 Enterococcus faecium DO GCA_000174395.2
228 Enterococcus hirae ATCC 9790 GCA_000271405.2
228 Enterococcus mundtii QU 25 GCA_000504125.1
228 Enterococcus rivorum GCA_001742285.1
228 Enterococcus saccharolyticus subsp. saccharolyticus ATCC 43076 GCA_000407285.1
223 Enterococcus haemoperoxidus ATCC BAA-382 GCA_000407165.1
223 Enterococcus phoeniculicola ATCC BAA-412 GCA_000407505.1
223 Enterococcus thailandicus GCA_001652875.1
tP
196 Staphylococcus capitis subsp. capitis GCA_001028645.1
196 Staphylococcus epidermidis ATCC 12228 GCA_000007645.1
196 Staphylococcus pettenkoferi GCA_002208805.1
192 Staphylococcus cohnii subsp. cohnii GCA_000972575.1
192 Staphylococcus lugdunensis HKU09-01 GCA_000025085.1
192 Staphylococcus saprophyticus subsp. saprophyticus ATCC 15305 GCA_000010125.1
185 Staphylococcus haemolyticus JCSC1435 GCA_000009865.1
tQ
236 Lactobacillus gasseri ATCC 33323 = JCM 1131 GCA_000014425.1
236 Lactobacillus hominis DSM 23910 = CRBIP 24.179 GCA_000296835.1
175 Lactobacillus acidophilus NCFM GCA_000011985.1
175 Lactobacillus gallinarum GCA_001314245.2
174 Lactobacillus crispatus ST1 GCA_000091765.1
174 Lactobacillus hamsteri DSM 5661 = JCM 6256 GCA_000615445.1
174 Lactobacillus kefiranofaciens ZW3 GCA_000214785.1
tR
179 Deinococcus radiodurans R1 GCA_000008565.1
134 Deinococcus gobiensis I-0 GCA_000252445.1
132 Deinococcus deserti VCD115 GCA_000020685.1
tS
181 Escherichia coli IAI39 GCA_000026345.1
181 Escherichia coli O104_3AH4 str. 2011C-3493 GCA_000299455.1
181 Escherichia coli O157_3AH7 str. Sakai GCA_000008865.1
181 Escherichia coli O83_3AH1 str. NRG 857C GCA_000183345.1
181 Escherichia coli UMN026 GCA_000026325.2
181 Escherichia coli str. K-12 substr. MG1655 GCA_000005845.2
181 Shigella dysenteriae Sd197 GCA_000012005.1
181 Shigella flexneri 2a str. 301 GCA_000006925.2
181 Tumebacillus flagellatus GCA_000714935.1
170 Erwinia iniecta GCA_001267535.1
160 Xenorhabdus bovienii SS-2004 GCA_000027225.1
tT
151 Bacillus thuringiensis YBT-1518 GCA_000497525.2
148 Lactobacillus harbinensis DSM 16991 GCA_000425885.1
148 Lactobacillus shenzhenensis LY-73 GCA_000469325.1
147 Abiotrophia defectiva ATCC 49176 GCA_000160075.2
tU
254 Streptococcus mutans UA159 GCA_000007465.2
196 Streptococcus macacae NCTC 11558 GCA_000187995.3
193 Streptococcus ratti FA-1 = DSM 20564 GCA_000286075.1
tV
152 Staphylococcus cohnii subsp. cohnii GCA_000972575.1
148 Staphylococcus equorum GCA_001432245.1
141 Staphylococcus saprophyticus subsp. saprophyticus ATCC 15305 GCA_000010125.1
141 Staphylococcus xylosus GCA_000706685.1
tW
94 Staphylococcus pettenkoferi GCA_002208805.1
91 Megasphaera cerevisiae DSM 20462 GCA_001045675.1
91 Staphylococcus arlettae CVD059 GCA_000295715.1
91 Staphylococcus aureus subsp. aureus NCTC 8325 GCA_000013425.1
91 Staphylococcus capitis subsp. capitis GCA_001028645.1
91 Staphylococcus cohnii subsp. cohnii GCA_000972575.1

91 Staphylococcus epidermidis ATCC 12228 GCA_000007645.1
91 Staphylococcus equorum GCA_001432245.1
91 Staphylococcus gallinarum GCA_000875895.1
91 Staphylococcus haemolyticus JCSC1435 GCA_000009865.1
91 Staphylococcus hominis subsp. hominis C80 GCA_000183685.1
91 Staphylococcus lugdunensis HKU09-01 GCA_000025085.1
91 Staphylococcus saprophyticus GCA_001074355.1
91 Staphylococcus saprophyticus subsp. saprophyticus ATCC 15305 GCA_000010125.1
91 Staphylococcus simiae CCM 7213 GCA_000235645.2
91 Staphylococcus succinus GCA_001902315.1
91 Staphylococcus warneri SG1 GCA_000332735.1
91 Staphylococcus xylosus GCA_000706685.1
86 Jeotgalicoccus halophilus GCA_900101065.1
86 Jeotgalicoccus marinus DSM 19772 GCA_000425825.1
86 Jeotgalicoccus saudimassiliensis GCA_000756715.2
86 Macrococcus canis GCA_002119805.1
86 Nosocomiicoccus massiliensis GCA_000438455.1
86 Salinicoccus alkaliphilus DSM 16010 GCA_900142805.1
86 Salinicoccus carnicancri Crm GCA_000330705.1
86 Salinicoccus halodurans GCA_001005905.1
86 Salinicoccus qingdaonensis GCA_900101075.1
86 Staphylococcus hyicus GCA_000816085.1
86 Staphylococcus lentus F1142 GCA_000286395.1
86 Staphylococcus lutrae GCA_002101335.1
86 Staphylococcus sciuri GCA_002209165.1
86 Staphylococcus vitulinus F1028 GCA_000286335.1
tX
232 Escherichia coli IAI39 GCA_000026345.1
232 Escherichia coli O104_3AH4 str. 2011C-3493 GCA_000299455.1
232 Escherichia coli O157_3AH7 str. Sakai GCA_000008865.1
232 Escherichia coli O83_3AH1 str. NRG 857C GCA_000183345.1
232 Escherichia coli UMN026 GCA_000026325.2
232 Escherichia coli str. K-12 substr. MG1655 GCA_000005845.2
232 Shigella dysenteriae Sd197 GCA_000012005.1
232 Shigella flexneri 2a str. 301 GCA_000006925.2
232 Tumebacillus flagellatus GCA_000714935.1
206 Erwinia iniecta GCA_001267535.1
206 Rosenbergiella nectarea GCA_900111105.1
197 Cedecea neteri GCA_000757825.1
197 Cronobacter sakazakii GCA_000982825.1
197 Edwardsiella anguillarum ET080813 GCA_000264765.2
197 Erwinia gerundensis GCA_001517405.1
197 Erwinia toletana DAPP-PG 735 GCA_000336255.1
197 Pantoea agglomerans GCA_001709315.1
197 Pantoea dispersa EGD-AAK13 GCA_000465555.2
197 Shimwellia blattae DSM 4481 = NBRC 105725 GCA_000262305.1
tY
306 Streptococcus mutans UA159 GCA_000007465.2
188 Streptococcus marimammalium DSM 18627 GCA_000380045.1
183 Streptococcus iniae GCA_000831485.1
tZ
247 Escherichia coli IAI39 GCA_000026345.1
247 Escherichia coli O104_3AH4 str. 2011C-3493 GCA_000299455.1
247 Escherichia coli O157_3AH7 str. Sakai GCA_000008865.1
247 Escherichia coli O83_3AH1 str. NRG 857C GCA_000183345.1
247 Escherichia coli UMN026 GCA_000026325.2
247 Escherichia coli str. K-12 substr. MG1655 GCA_000005845.2
247 Shigella flexneri 2a str. 301 GCA_000006925.2
247 Tumebacillus flagellatus GCA_000714935.1
238 Shigella dysenteriae Sd197 GCA_000012005.1
232 Erwinia iniecta GCA_001267535.1
u0
240 Deinococcus radiodurans R1 GCA_000008565.1
170 Deinococcus gobiensis I-0 GCA_000252445.1
154 Deinococcus deserti VCD115 GCA_000020685.1
u1
221 Clostridium beijerinckii GCA_000833105.2
221 Clostridium puniceum GCA_002006345.1
221 Clostridium saccharobutylicum DSM 13864 GCA_000473995.1
221 Clostridium saccharoperbutylacetonicum N1-4_28HMT_29 GCA_000340885.1
202 Clostridium chromiireducens GCA_002029255.1
183 Clostridium amylolyticum GCA_900142075.1
183 Clostridium polynesiense GCA_000820705.1
u2
283 Streptococcus mutans UA159 GCA_000007465.2
186 Streptococcus macacae NCTC 11558 GCA_000187995.3

181 Streptococcus ferus DSM 20646 GCA_000372425.1
u3
240 Streptococcus mutans UA159 GCA_000007465.2
149 Streptococcus macacae NCTC 11558 GCA_000187995.3
146 Streptococcus ratti FA-1 = DSM 20564 GCA_000286075.1
u4
269 Bacillus anthracis str. Ames GCA_000007845.1
269 Bacillus anthracis str. Sterne GCA_000008165.1
269 Bacillus cereus ATCC 14579 GCA_000007825.1
269 Bacillus mycoides GCA_000832605.1
269 Bacillus pseudomycoides DSM 12442 GCA_000161455.1
269 Bacillus thuringiensis YBT-1518 GCA_000497525.2
269 _5BBacillus thuringiensis_5D serovar konkukian str. 97-27 GCA_000008505.1
237 Bacillus manliponensis GCA_000712595.1
218 Bacillus aquimaris TF-12 GCA_001648555.1
u5
277 Bacillus anthracis str. Ames GCA_000007845.1
277 Bacillus anthracis str. Sterne GCA_000008165.1
277 Bacillus cereus ATCC 14579 GCA_000007825.1
277 Bacillus mycoides GCA_000832605.1
277 Bacillus pseudomycoides DSM 12442 GCA_000161455.1
277 Bacillus thuringiensis YBT-1518 GCA_000497525.2
277 _5BBacillus thuringiensis_5D serovar konkukian str. 97-27 GCA_000008505.1
210 Bacillus manliponensis GCA_000712595.1
196 Bacillus coahuilensis m4-4 GCA_000171615.1
196 Bacillus vietnamensis NBRC 101237 GCA_001591825.1
u6
266 Staphylococcus epidermidis ATCC 12228 GCA_000007645.1
263 Staphylococcus haemolyticus JCSC1435 GCA_000009865.1
262 Staphylococcus lugdunensis HKU09-01 GCA_000025085.1
u7
249 Clostridium saccharobutylicum DSM 13864 GCA_000473995.1
245 Clostridium beijerinckii GCA_000833105.2
245 Clostridium saccharoperbutylacetonicum N1-4_28HMT_29 GCA_000340885.1
241 Clostridium chromiireducens GCA_002029255.1
241 Clostridium puniceum GCA_002006345.1
u8
202 Staphylococcus cohnii subsp. cohnii GCA_000972575.1
202 Staphylococcus saprophyticus subsp. saprophyticus ATCC 15305 GCA_000010125.1
196 Staphylococcus capitis subsp. capitis GCA_001028645.1
196 Staphylococcus epidermidis ATCC 12228 GCA_000007645.1
196 Staphylococcus lugdunensis HKU09-01 GCA_000025085.1
193 Staphylococcus xylosus GCA_000706685.1
u9
244 Deinococcus radiodurans R1 GCA_000008565.1
137 Deinococcus proteolyticus MRP GCA_000190555.1
137 Deinococcus puniceus GCA_001644565.1
131 Deinococcus soli Cha et al. 2016 GCA_001007995.1
ua
259 Escherichia coli IAI39 GCA_000026345.1
259 Escherichia coli O104_3AH4 str. 2011C-3493 GCA_000299455.1
259 Escherichia coli O157_3AH7 str. Sakai GCA_000008865.1
259 Escherichia coli O83_3AH1 str. NRG 857C GCA_000183345.1
259 Escherichia coli UMN026 GCA_000026325.2
259 Escherichia coli str. K-12 substr. MG1655 GCA_000005845.2
259 Shigella dysenteriae Sd197 GCA_000012005.1
259 Shigella flexneri 2a str. 301 GCA_000006925.2
259 Tumebacillus flagellatus GCA_000714935.1
248 Erwinia iniecta GCA_001267535.1
243 Pantoea agglomerans GCA_001709315.1
ub
283 Lactobacillus gasseri ATCC 33323 = JCM 1131 GCA_000014425.1
246 Lactobacillus hominis DSM 23910 = CRBIP 24.179 GCA_000296835.1
175 Lactobacillus acidophilus NCFM GCA_000011985.1
175 Lactobacillus gallinarum GCA_001314245.2
uc
205 Clostridium beijerinckii GCA_000833105.2
205 Clostridium puniceum GCA_002006345.1
205 Clostridium saccharobutylicum DSM 13864 GCA_000473995.1
205 Clostridium saccharoperbutylacetonicum N1-4_28HMT_29 GCA_000340885.1
201 Clostridium butyricum GCA_001456065.2
201 Clostridium chromiireducens GCA_002029255.1
188 Clostridium taeniosporum GCA_001735765.1
ud
204 Clostridium beijerinckii GCA_000833105.2
204 Clostridium puniceum GCA_002006345.1

204 Clostridium saccharoperbutylacetonicum N1-4_28HMT_29 GCA_000340885.1
189 Clostridium saccharobutylicum DSM 13864 GCA_000473995.1
187 Clostridium butyricum GCA_001456065.2
ue
316 Lactobacillus gasseri ATCC 33323 = JCM 1131 GCA_000014425.1
311 Lactobacillus hominis DSM 23910 = CRBIP 24.179 GCA_000296835.1
241 Lactobacillus jensenii GCA_001936235.1
uf
219 Escherichia coli IAI39 GCA_000026345.1
219 Escherichia coli O104_3AH4 str. 2011C-3493 GCA_000299455.1
219 Escherichia coli O157_3AH7 str. Sakai GCA_000008865.1
219 Escherichia coli O83_3AH1 str. NRG 857C GCA_000183345.1
219 Escherichia coli UMN026 GCA_000026325.2
219 Escherichia coli str. K-12 substr. MG1655 GCA_000005845.2
219 Shigella dysenteriae Sd197 GCA_000012005.1
219 Shigella flexneri 2a str. 301 GCA_000006925.2
219 Tumebacillus flagellatus GCA_000714935.1
185 Erwinia iniecta GCA_001267535.1
157 Rosenbergiella nectarea GCA_900111105.1
ug
217 Escherichia coli IAI39 GCA_000026345.1
217 Escherichia coli O104_3AH4 str. 2011C-3493 GCA_000299455.1
217 Escherichia coli O157_3AH7 str. Sakai GCA_000008865.1
217 Escherichia coli O83_3AH1 str. NRG 857C GCA_000183345.1
217 Escherichia coli UMN026 GCA_000026325.2
217 Escherichia coli str. K-12 substr. MG1655 GCA_000005845.2
217 Shigella dysenteriae Sd197 GCA_000012005.1
217 Shigella flexneri 2a str. 301 GCA_000006925.2
217 Tumebacillus flagellatus GCA_000714935.1
203 Erwinia iniecta GCA_001267535.1
185 Pragia fontium GCA_001026985.1
uh
219 Bacillus anthracis str. Ames GCA_000007845.1
219 Bacillus anthracis str. Sterne GCA_000008165.1
219 Bacillus cereus ATCC 14579 GCA_000007825.1
219 Bacillus mycoides GCA_000832605.1
219 Bacillus pseudomycoides DSM 12442 GCA_000161455.1
219 Bacillus thuringiensis YBT-1518 GCA_000497525.2
219 _5BBacillus thuringiensis_5D serovar konkukian str. 97-27 GCA_000008505.1
188 Bacillus marisflavi GCA_001274775.1
182 Bacillus manliponensis GCA_000712595.1
ui
243 Streptococcus mutans UA159 GCA_000007465.2
177 Streptococcus gordonii str. Challis substr. CH1 GCA_000017005.1
173 Streptococcus cristatus AS 1.3089 GCA_000385925.1
173 Streptococcus mitis B6 GCA_000027165.1
173 Streptococcus pneumoniae R6 GCA_000007045.1
uj
136 Salinicoccus carnicancri Crm GCA_000330705.1
126 Salinicoccus alkaliphilus DSM 16010 GCA_900142805.1
126 Salinicoccus halodurans GCA_001005905.1
126 Salinicoccus qingdaonensis GCA_900101075.1
124 Macrococcus canis GCA_002119805.1
124 Staphylococcus lentus F1142 GCA_000286395.1
124 Staphylococcus lutrae GCA_002101335.1
124 Staphylococcus sciuri GCA_002209165.1
uk
290 Deinococcus radiodurans R1 GCA_000008565.1
197 Deinococcus gobiensis I-0 GCA_000252445.1
181 Deinococcus puniceus GCA_001644565.1
ul
200 Bacillus anthracis str. Ames GCA_000007845.1
200 Bacillus anthracis str. Sterne GCA_000008165.1
200 Bacillus mycoides GCA_000832605.1
200 Bacillus pseudomycoides DSM 12442 GCA_000161455.1
200 Bacillus thuringiensis YBT-1518 GCA_000497525.2
200 _5BBacillus thuringiensis_5D serovar konkukian str. 97-27 GCA_000008505.1
199 Bacillus cereus ATCC 14579 GCA_000007825.1
158 Bacillus marisflavi GCA_001274775.1
um
186 Megasphaera cerevisiae DSM 20462 GCA_001045675.1
186 Staphylococcus arlettae CVD059 GCA_000295715.1
186 Staphylococcus aureus subsp. aureus NCTC 8325 GCA_000013425.1
186 Staphylococcus capitis subsp. capitis GCA_001028645.1
186 Staphylococcus cohnii subsp. cohnii GCA_000972575.1
186 Staphylococcus epidermidis ATCC 12228 GCA_000007645.1

186 Staphylococcus gallinarum GCA_000875895.1
186 Staphylococcus haemolyticus JCSC1435 GCA_000009865.1
186 Staphylococcus hominis subsp. hominis C80 GCA_000183685.1
186 Staphylococcus lugdunensis HKU09-01 GCA_000025085.1
186 Staphylococcus saprophyticus GCA_001074355.1
186 Staphylococcus saprophyticus subsp. saprophyticus ATCC 15305 GCA_000010125.1
186 Staphylococcus simiae CCM 7213 GCA_000235645.2
186 Staphylococcus succinus GCA_001902315.1
186 Staphylococcus warneri SG1 GCA_000332735.1
186 Staphylococcus xylosus GCA_000706685.1
181 Staphylococcus condimenti GCA_001618885.1
181 Staphylococcus simulans GCA_001559115.1
177 Staphylococcus lutrae GCA_002101335.1
un
247 Rhodobacter sphaeroides 2.4.1 GCA_000012905.2
233 Defluviimonas alba GCA_001620265.1
229 Rhodobacter sphaeroides ATCC 17025 GCA_000016405.1
uo
124 Staphylococcus epidermidis ATCC 12228 GCA_000007645.1
122 Staphylococcus capitis subsp. capitis GCA_001028645.1
122 Staphylococcus haemolyticus JCSC1435 GCA_000009865.1
122 Staphylococcus hominis subsp. hominis C80 GCA_000183685.1
122 Staphylococcus lugdunensis HKU09-01 GCA_000025085.1
122 Staphylococcus microti GCA_000934465.1
122 Staphylococcus pseudintermedius HKU10-03 GCA_000185885.1
122 Staphylococcus simulans GCA_001559115.1
109 Staphylococcus arlettae CVD059 GCA_000295715.1
109 Staphylococcus cohnii subsp. cohnii GCA_000972575.1
109 Staphylococcus saprophyticus subsp. saprophyticus ATCC 15305 GCA_000010125.1
up
306 Bacillus anthracis str. Ames GCA_000007845.1
306 Bacillus anthracis str. Sterne GCA_000008165.1
306 Bacillus cereus ATCC 14579 GCA_000007825.1
306 Bacillus mycoides GCA_000832605.1
306 Bacillus pseudomycoides DSM 12442 GCA_000161455.1
306 Bacillus thuringiensis YBT-1518 GCA_000497525.2
306 _5BBacillus thuringiensis_5D serovar konkukian str. 97-27 GCA_000008505.1
270 Bacillus cytotoxicus NVH 391-98 GCA_000017425.1
268 Bacillus manliponensis GCA_000712595.1
uq
257 Clostridium beijerinckii GCA_000833105.2
257 Clostridium saccharoperbutylacetonicum N1-4_28HMT_29 GCA_000340885.1
250 Clostridium puniceum GCA_002006345.1
248 Clostridium saccharobutylicum DSM 13864 GCA_000473995.1
ur
205 Deinococcus radiodurans R1 GCA_000008565.1
147 Deinococcus deserti VCD115 GCA_000020685.1
138 Deinococcus proteolyticus MRP GCA_000190555.1
us
313 Lactobacillus gasseri ATCC 33323 = JCM 1131 GCA_000014425.1
279 Lactobacillus hominis DSM 23910 = CRBIP 24.179 GCA_000296835.1
246 Lactobacillus psittaci DSM 15354 GCA_000425905.1
ut
182 Streptococcus mutans UA159 GCA_000007465.2
124 Streptococcus iniae GCA_000831485.1
114 Bacillus bataviensis LMG 21833 GCA_000307875.1
114 Lactococcus lactis subsp. lactis Il1403 GCA_000006865.1
114 Lactococcus piscium MKFS47 GCA_000981525.1
114 Streptococcus agalactiae 2603V_2FR GCA_000007265.1
114 Streptococcus cristatus AS 1.3089 GCA_000385925.1
114 Streptococcus dysgalactiae subsp. equisimilis AC-2713 GCA_000317855.1
114 Streptococcus gordonii str. Challis substr. CH1 GCA_000017005.1
114 Streptococcus ictaluri 707-05 GCA_000188015.3
114 Streptococcus mitis B6 GCA_000027165.1
114 Streptococcus parasanguinis ATCC 15912 GCA_000164675.2
114 Streptococcus phocae subsp. salmonis GCA_000772915.1
114 Streptococcus pneumoniae R6 GCA_000007045.1
uu
195 Streptococcus mutans UA159 GCA_000007465.2
136 Streptococcus dysgalactiae subsp. equisimilis AC-2713 GCA_000317855.1
131 Streptococcus iniae GCA_000831485.1
uv
317 Bacillus cereus ATCC 14579 GCA_000007825.1
308 Bacillus anthracis str. Ames GCA_000007845.1
308 Bacillus anthracis str. Sterne GCA_000008165.1
308 Bacillus mycoides GCA_000832605.1

308 Bacillus pseudomycoides DSM 12442 GCA_000161455.1
308 Bacillus thuringiensis YBT-1518 GCA_000497525.2
308 _5BBacillus thuringiensis_5D serovar konkukian str. 97-27 GCA_000008505.1
235 Bacillus manliponensis GCA_000712595.1
uw
178 Deinococcus radiodurans R1 GCA_000008565.1
137 Deinococcus gobiensis I-0 GCA_000252445.1
116 Deinococcus puniceus GCA_001644565.1
ux
245 Escherichia coli IAI39 GCA_000026345.1
245 Escherichia coli O104_3AH4 str. 2011C-3493 GCA_000299455.1
245 Escherichia coli O157_3AH7 str. Sakai GCA_000008865.1
245 Escherichia coli O83_3AH1 str. NRG 857C GCA_000183345.1
245 Escherichia coli UMN026 GCA_000026325.2
245 Escherichia coli str. K-12 substr. MG1655 GCA_000005845.2
245 Shigella dysenteriae Sd197 GCA_000012005.1
245 Shigella flexneri 2a str. 301 GCA_000006925.2
245 Tumebacillus flagellatus GCA_000714935.1
206 Erwinia iniecta GCA_001267535.1
176 Rosenbergiella nectarea GCA_900111105.1
uy
195 Deinococcus radiodurans R1 GCA_000008565.1
138 Deinococcus gobiensis I-0 GCA_000252445.1
126 Deinococcus marmoris DSM 12784 GCA_000701405.1
126 Deinococcus swuensis GCA_000800395.1
uz
243 Clostridium beijerinckii GCA_000833105.2
243 Clostridium puniceum GCA_002006345.1
243 Clostridium saccharoperbutylacetonicum N1-4_28HMT_29 GCA_000340885.1
228 Clostridium saccharobutylicum DSM 13864 GCA_000473995.1
213 Clostridium chromiireducens GCA_002029255.1
uA
125 Streptococcus mutans UA159 GCA_000007465.2
116 Streptococcus ferus DSM 20646 GCA_000372425.1
97 Streptococcus marimammalium DSM 18627 GCA_000380045.1
97 Streptococcus sobrinus DSM 20742 = ATCC 33478 GCA_000686605.1
uB
197 Clostridium beijerinckii GCA_000833105.2
197 Clostridium saccharobutylicum DSM 13864 GCA_000473995.1
197 Clostridium saccharoperbutylacetonicum N1-4_28HMT_29 GCA_000340885.1
190 Clostridium puniceum GCA_002006345.1
175 Clostridium chromiireducens GCA_002029255.1
uC
146 Escherichia coli IAI39 GCA_000026345.1
146 Escherichia coli O104_3AH4 str. 2011C-3493 GCA_000299455.1
146 Escherichia coli O157_3AH7 str. Sakai GCA_000008865.1
146 Escherichia coli O83_3AH1 str. NRG 857C GCA_000183345.1
146 Escherichia coli UMN026 GCA_000026325.2
146 Escherichia coli str. K-12 substr. MG1655 GCA_000005845.2
146 Shigella dysenteriae Sd197 GCA_000012005.1
146 Shigella flexneri 2a str. 301 GCA_000006925.2
146 Tumebacillus flagellatus GCA_000714935.1
142 Erwinia iniecta GCA_001267535.1
127 Cronobacter sakazakii GCA_000982825.1
127 Enterobacter cloacae subsp. cloacae ATCC 13047 GCA_000025565.1
127 Enterobacter hormaechei subsp. steigerwaltii GCA_001729725.1
127 Enterobacter kobei GCA_900185885.1
127 Erwinia persicina NBRC 102418 GCA_001571305.1
127 Erwinia toletana DAPP-PG 735 GCA_000336255.1
127 Klebsiella oxytoca GCA_001022195.1
127 Kosakonia cowanii GCA_001975225.1
127 Pseudescherichia vulneris NBRC 102420 GCA_000759795.1
127 Salmonella enterica subsp. enterica serovar Typhi str. CT18 GCA_000195995.1
uD
194 Staphylococcus epidermidis ATCC 12228 GCA_000007645.1
188 Staphylococcus capitis subsp. capitis GCA_001028645.1
179 Megasphaera cerevisiae DSM 20462 GCA_001045675.1
179 Staphylococcus haemolyticus JCSC1435 GCA_000009865.1
179 Staphylococcus lugdunensis HKU09-01 GCA_000025085.1
179 Staphylococcus pettenkoferi GCA_002208805.1
179 Staphylococcus warneri SG1 GCA_000332735.1
uE
275 Streptococcus mutans UA159 GCA_000007465.2
160 Streptococcus massiliensis DSM 18628 GCA_000380065.1
157 Streptococcus gordonii str. Challis substr. CH1 GCA_000017005.1
uF

269 Staphylococcus epidermidis ATCC 12228 GCA_000007645.1
269 Staphylococcus haemolyticus JCSC1435 GCA_000009865.1
260 Staphylococcus cohnii subsp. cohnii GCA_000972575.1
258 Staphylococcus hominis subsp. hominis C80 GCA_000183685.1
uG
226 Granulicatella elegans ATCC 700633 GCA_000162475.2
221 Melissococcus plutonius S1 GCA_000747585.1
218 Abiotrophia defectiva ATCC 49176 GCA_000160075.2
uH
239 Deinococcus radiodurans R1 GCA_000008565.1
184 Deinococcus gobiensis I-0 GCA_000252445.1
167 Deinococcus soli Cha et al. 2016 GCA_001007995.1
uI
308 Enterococcus faecalis V583 GCA_000007785.1
308 Streptomyces cinnamoneus GCA_001885705.1
292 Enterococcus asini ATCC 700915 GCA_000407365.1
292 Enterococcus dispar ATCC 51266 GCA_000406945.1
291 Enterococcus canis NBRC 100695 GCA_001544375.1
291 Enterococcus faecium DO GCA_000174395.2
291 Enterococcus haemoperoxidus ATCC BAA-382 GCA_000407165.1
291 Enterococcus hirae ATCC 9790 GCA_000271405.2
291 Enterococcus mundtii QU 25 GCA_000504125.1
291 Enterococcus phoeniculicola ATCC BAA-412 GCA_000407505.1
291 Enterococcus rivorum GCA_001742285.1
291 Enterococcus thailandicus GCA_001652875.1
uJ
191 Rhodobacter sphaeroides 2.4.1 GCA_000012905.2
134 Gemmobacter megaterium GCA_900156815.1
126 Rhodobacter capsulatus SB 1003 GCA_000021865.1
uK
132 Bifidobacterium choerinum GCA_000741135.1
132 Bifidobacterium pseudolongum PV8-2 GCA_000800475.2
128 Bifidobacterium thermophilum GCA_000741495.1
124 Bifidobacterium adolescentis ATCC 15703 GCA_000010425.1
uL
268 Bacillus anthracis str. Ames GCA_000007845.1
268 Bacillus anthracis str. Sterne GCA_000008165.1
268 Bacillus cereus ATCC 14579 GCA_000007825.1
268 Bacillus mycoides GCA_000832605.1
268 Bacillus pseudomycoides DSM 12442 GCA_000161455.1
268 Bacillus thuringiensis YBT-1518 GCA_000497525.2
268 _5BBacillus thuringiensis_5D serovar konkukian str. 97-27 GCA_000008505.1
220 Bacillus flexus GCA_002024265.1
220 Bacillus megaterium NBRC 15308 = ATCC 14581 GCA_000832985.1
219 Bacillus tuaregi GCA_900104575.1
uM
246 Staphylococcus aureus subsp. aureus NCTC 8325 GCA_000013425.1
246 Staphylococcus cohnii subsp. cohnii GCA_000972575.1
246 Staphylococcus epidermidis ATCC 12228 GCA_000007645.1
246 Staphylococcus haemolyticus JCSC1435 GCA_000009865.1
246 Staphylococcus hominis subsp. hominis C80 GCA_000183685.1
246 Staphylococcus saprophyticus GCA_001074355.1
246 Staphylococcus simiae CCM 7213 GCA_000235645.2
242 Megasphaera cerevisiae DSM 20462 GCA_001045675.1
242 Staphylococcus warneri SG1 GCA_000332735.1
241 Staphylococcus capitis subsp. capitis GCA_001028645.1
uN
50 Clostridium kluyveri DSM 555 GCA_000016505.1
50 Clostridium oryzae GCA_002029235.1
46 Tepidibacter formicigenes DSM 15518 GCA_900142235.1
46 Tepidibacter thalassicus DSM 15285 GCA_900129915.1
45 Clostridium acetobutylicum ATCC 824 GCA_000008765.1
45 Clostridium baratii str. Sullivan GCA_000789395.1
45 Clostridium beijerinckii GCA_000833105.2
45 Clostridium botulinum B str. Eklund 17B _28NRP_29 GCA_000020165.1
45 Clostridium butyricum GCA_001456065.2
45 Clostridium chromiireducens GCA_002029255.1
45 Clostridium felsineum DSM 794 GCA_002006355.1
45 Clostridium puniceum GCA_002006345.1
45 Clostridium saccharobutylicum DSM 13864 GCA_000473995.1
45 Clostridium saccharoperbutylacetonicum N1-4_28HMT_29 GCA_000340885.1
45 Clostridium taeniosporum GCA_001735765.1
45 Clostridium uliginosum GCA_900112485.1
uO
267 Bifidobacterium adolescentis ATCC 15703 GCA_000010425.1
235 Bifidobacterium breve DSM 20213 = JCM 1192 GCA_001025175.1

225 Bifidobacterium callitrichos DSM 23973 GCA_000741175.1
uP
91 Deinococcus radiodurans R1 GCA_000008565.1
75 Deinococcus geothermalis DSM 11300 GCA_000196275.1
64 Deinococcus deserti VCD115 GCA_000020685.1
64 Deinococcus proteolyticus MRP GCA_000190555.1
uQ
141 Streptococcus mutans UA159 GCA_000007465.2
83 Streptococcus ferus DSM 20646 GCA_000372425.1
77 Streptococcus macacae NCTC 11558 GCA_000187995.3
uR
232 Bacillus thuringiensis YBT-1518 GCA_000497525.2
231 Bacillus anthracis str. Ames GCA_000007845.1
231 Bacillus anthracis str. Sterne GCA_000008165.1
231 Bacillus cereus ATCC 14579 GCA_000007825.1
231 Bacillus mycoides GCA_000832605.1
231 Bacillus pseudomycoides DSM 12442 GCA_000161455.1
231 _5BBacillus thuringiensis_5D serovar konkukian str. 97-27 GCA_000008505.1
199 Bacillus horneckiae GCA_001636335.1
199 Bacillus solani GCA_001420595.1
uS
275 Rhodobacter sphaeroides 2.4.1 GCA_000012905.2
245 Rhodobacter sphaeroides ATCC 17025 GCA_000016405.1
227 Gemmobacter megaterium GCA_900156815.1
uT
172 Deinococcus radiodurans R1 GCA_000008565.1
136 Deinococcus marmoris DSM 12784 GCA_000701405.1
136 Deinococcus swuensis GCA_000800395.1
127 Deinococcus soli Cha et al. 2016 GCA_001007995.1
uU
134 Clostridium beijerinckii GCA_000833105.2
134 Clostridium puniceum GCA_002006345.1
134 Clostridium saccharobutylicum DSM 13864 GCA_000473995.1
134 Clostridium saccharoperbutylacetonicum N1-4_28HMT_29 GCA_000340885.1
119 Clostridium chromiireducens GCA_002029255.1
112 Clostridium neonatale GCA_001458595.1
uV
243 Deinococcus radiodurans R1 GCA_000008565.1
179 Deinococcus deserti VCD115 GCA_000020685.1
176 Deinococcus gobiensis I-0 GCA_000252445.1
uW
202 Clostridium beijerinckii GCA_000833105.2
202 Clostridium saccharobutylicum DSM 13864 GCA_000473995.1
202 Clostridium saccharoperbutylacetonicum N1-4_28HMT_29 GCA_000340885.1
195 Clostridium puniceum GCA_002006345.1
180 Clostridium botulinum B str. Eklund 17B _28NRP_29 GCA_000020165.1
180 Clostridium chromiireducens GCA_002029255.1
180 Clostridium taeniosporum GCA_001735765.1
uX
181 Enterococcus faecalis V583 GCA_000007785.1
181 Streptomyces cinnamoneus GCA_001885705.1
151 Enterococcus canis NBRC 100695 GCA_001544375.1
151 Enterococcus casseliflavus EC20 GCA_000157355.2
151 Enterococcus dispar ATCC 51266 GCA_000406945.1
151 Enterococcus faecium DO GCA_000174395.2
151 Enterococcus gilvus ATCC BAA-350 GCA_000407545.1
151 Enterococcus haemoperoxidus ATCC BAA-382 GCA_000407165.1
151 Enterococcus hermanniensis GCA_001885945.1
151 Enterococcus hirae ATCC 9790 GCA_000271405.2
151 Enterococcus malodoratus ATCC 43197 GCA_000407185.1
151 Enterococcus mundtii QU 25 GCA_000504125.1
151 Enterococcus pallens ATCC BAA-351 GCA_000407485.1
151 Enterococcus phoeniculicola ATCC BAA-412 GCA_000407505.1
151 Enterococcus pseudoavium NBRC 100491 GCA_001544295.1
151 Enterococcus rivorum GCA_001742285.1
151 Enterococcus saccharolyticus subsp. saccharolyticus ATCC 43076 GCA_000407285.1
151 Enterococcus thailandicus GCA_001652875.1
149 Catellicoccus marimammalium M35_2F04_2F3 GCA_000313915.1
149 Enterococcus asini ATCC 700915 GCA_000407365.1
149 Lactobacillus curvatus GCA_002224425.1
149 Lactobacillus kimchicus JCM 15530 GCA_001433995.1
149 Lactobacillus mixtipabuli GCA_002217925.1
149 Lactobacillus odoratitofui DSM 19909 = JCM 15043 GCA_001434895.1
149 Lactobacillus sakei GCA_002224565.1
149 Lactobacillus silagei JCM 19001 GCA_002217945.1
149 Lactobacillus similis DSM 23365 = JCM 2765 GCA_001311075.1

uY
231 Clostridium beijerinckii GCA_000833105.2
231 Clostridium saccharoperbutylacetonicum N1-4_28HMT_29 GCA_000340885.1
224 Clostridium puniceum GCA_002006345.1
222 Clostridium saccharobutylicum DSM 13864 GCA_000473995.1
uZ
236 Bifidobacterium adolescentis ATCC 15703 GCA_000010425.1
222 Bifidobacterium tsurumiense GCA_000741765.1
217 Bifidobacterium angulatum DSM 20098 = JCM 7096 GCA_001025155.1
217 Bifidobacterium callitrichos DSM 23973 GCA_000741175.1
v0
254 Rhodobacter sphaeroides 2.4.1 GCA_000012905.2
213 Rhodobacter sphaeroides ATCC 17025 GCA_000016405.1
212 Defluviimonas alba GCA_001620265.1
v1
82 Deinococcus radiodurans R1 GCA_000008565.1
63 Deinococcus puniceus GCA_001644565.1
62 Deinococcus hopiensis KR-140 GCA_900176165.1
v2
85 Deinococcus radiodurans R1 GCA_000008565.1
78 Deinococcus gobiensis I-0 GCA_000252445.1
69 Deinococcus marmoris DSM 12784 GCA_000701405.1
69 Deinococcus swuensis GCA_000800395.1
v3
227 Escherichia coli IAI39 GCA_000026345.1
227 Escherichia coli O104_3AH4 str. 2011C-3493 GCA_000299455.1
227 Escherichia coli O157_3AH7 str. Sakai GCA_000008865.1
227 Escherichia coli O83_3AH1 str. NRG 857C GCA_000183345.1
227 Escherichia coli UMN026 GCA_000026325.2
227 Escherichia coli str. K-12 substr. MG1655 GCA_000005845.2
227 Shigella dysenteriae Sd197 GCA_000012005.1
227 Shigella flexneri 2a str. 301 GCA_000006925.2
227 Tumebacillus flagellatus GCA_000714935.1
187 Erwinia iniecta GCA_001267535.1
178 Cronobacter sakazakii GCA_000982825.1
178 Erwinia toletana DAPP-PG 735 GCA_000336255.1
v4
226 Escherichia coli IAI39 GCA_000026345.1
226 Escherichia coli O104_3AH4 str. 2011C-3493 GCA_000299455.1
226 Escherichia coli O157_3AH7 str. Sakai GCA_000008865.1
226 Escherichia coli O83_3AH1 str. NRG 857C GCA_000183345.1
226 Escherichia coli UMN026 GCA_000026325.2
226 Escherichia coli str. K-12 substr. MG1655 GCA_000005845.2
226 Shigella dysenteriae Sd197 GCA_000012005.1
226 Shigella flexneri 2a str. 301 GCA_000006925.2
226 Tumebacillus flagellatus GCA_000714935.1
221 Cedecea neteri GCA_000757825.1
210 Serratia marcescens subsp. marcescens Db11 GCA_000513215.1
v5
244 Bifidobacterium adolescentis ATCC 15703 GCA_000010425.1
232 Bifidobacterium asteroides PRL2011 GCA_000304215.1
229 Bifidobacterium gallicum DSM 20093 = LMG 11596 GCA_000741205.1
v6
259 Deinococcus radiodurans R1 GCA_000008565.1
183 Deinococcus gobiensis I-0 GCA_000252445.1
177 Deinococcus deserti VCD115 GCA_000020685.1
v7
143 Erwinia iniecta GCA_001267535.1
143 Erwinia toletana DAPP-PG 735 GCA_000336255.1
143 Escherichia coli IAI39 GCA_000026345.1
143 Escherichia coli O104_3AH4 str. 2011C-3493 GCA_000299455.1
143 Escherichia coli O157_3AH7 str. Sakai GCA_000008865.1
143 Escherichia coli O83_3AH1 str. NRG 857C GCA_000183345.1
143 Escherichia coli UMN026 GCA_000026325.2
143 Escherichia coli str. K-12 substr. MG1655 GCA_000005845.2
143 Kosakonia cowanii GCA_001975225.1
143 Pantoea ananatis LMG 20103 GCA_000025405.2
143 Pluralibacter gergoviae GCA_000757785.1
143 Shigella flexneri 2a str. 301 GCA_000006925.2
143 Shimwellia blattae DSM 4481 = NBRC 105725 GCA_000262305.1
143 Tumebacillus flagellatus GCA_000714935.1
142 Kosakonia sacchari SP1 GCA_000300455.4
128 Atlantibacter hermannii NBRC 105704 GCA_000248015.2
128 Cronobacter sakazakii GCA_000982825.1
128 Enterobacter hormaechei subsp. steigerwaltii GCA_001729725.1
128 Erwinia amylovora CFBP1430 GCA_000091565.1

128 Erwinia billingiae Eb661 GCA_000196615.1
128 Klebsiella oxytoca GCA_001022195.1
128 Pantoea agglomerans GCA_001709315.1
128 Pantoea alhagi GCA_002101395.1
128 Pantoea dispersa EGD-AAK13 GCA_000465555.2
128 Pantoea rwandensis GCA_000759475.1
128 Pantoea septica GCA_002095575.1
128 Pseudescherichia vulneris NBRC 102420 GCA_000759795.1
128 Shigella dysenteriae Sd197 GCA_000012005.1
v8
140 Deinococcus radiodurans R1 GCA_000008565.1
70 Deinococcus deserti VCD115 GCA_000020685.1
70 Deinococcus gobiensis I-0 GCA_000252445.1
55 Deinococcus frigens DSM 12807 GCA_000701425.1
55 Deinococcus hopiensis KR-140 GCA_900176165.1
55 Deinococcus marmoris DSM 12784 GCA_000701405.1
55 Deinococcus puniceus GCA_001644565.1
55 Deinococcus swuensis GCA_000800395.1
v9
254 Bacillus thuringiensis YBT-1518 GCA_000497525.2
253 Bacillus anthracis str. Ames GCA_000007845.1
253 Bacillus anthracis str. Sterne GCA_000008165.1
253 Bacillus cereus ATCC 14579 GCA_000007825.1
253 Bacillus mycoides GCA_000832605.1
253 Bacillus pseudomycoides DSM 12442 GCA_000161455.1
253 _5BBacillus thuringiensis_5D serovar konkukian str. 97-27 GCA_000008505.1
186 Bacillus manliponensis GCA_000712595.1
va
76 Streptococcus mutans UA159 GCA_000007465.2
67 Streptococcus macacae NCTC 11558 GCA_000187995.3
66 Streptococcus sobrinus DSM 20742 = ATCC 33478 GCA_000686605.1
vb
205 Deinococcus radiodurans R1 GCA_000008565.1
129 Deinococcus gobiensis I-0 GCA_000252445.1
116 Deinococcus puniceus GCA_001644565.1
vc
278 Clostridium beijerinckii GCA_000833105.2
278 Clostridium puniceum GCA_002006345.1
278 Clostridium saccharoperbutylacetonicum N1-4_28HMT_29 GCA_000340885.1
269 Clostridium saccharobutylicum DSM 13864 GCA_000473995.1
264 Clostridium butyricum GCA_001456065.2
vd
285 Deinococcus radiodurans R1 GCA_000008565.1
230 Deinococcus gobiensis I-0 GCA_000252445.1
198 Deinococcus puniceus GCA_001644565.1
ve
231 Deinococcus radiodurans R1 GCA_000008565.1
156 Deinococcus gobiensis I-0 GCA_000252445.1
141 Deinococcus deserti VCD115 GCA_000020685.1
141 Deinococcus soli Cha et al. 2016 GCA_001007995.1
vf
142 Clostridium beijerinckii GCA_000833105.2
142 Clostridium puniceum GCA_002006345.1
142 Clostridium saccharoperbutylacetonicum N1-4_28HMT_29 GCA_000340885.1
138 Clostridium saccharobutylicum DSM 13864 GCA_000473995.1
121 Clostridium chromiireducens GCA_002029255.1
vg
182 Bacillus anthracis str. Ames GCA_000007845.1
182 Bacillus anthracis str. Sterne GCA_000008165.1
182 Bacillus cereus ATCC 14579 GCA_000007825.1
182 Bacillus mycoides GCA_000832605.1
182 Bacillus pseudomycoides DSM 12442 GCA_000161455.1
182 Bacillus thuringiensis YBT-1518 GCA_000497525.2
182 _5BBacillus thuringiensis_5D serovar konkukian str. 97-27 GCA_000008505.1
164 Staphylococcus arlettae CVD059 GCA_000295715.1
159 Megasphaera cerevisiae DSM 20462 GCA_001045675.1
159 Staphylococcus aureus subsp. aureus NCTC 8325 GCA_000013425.1
159 Staphylococcus capitis subsp. capitis GCA_001028645.1
159 Staphylococcus cohnii subsp. cohnii GCA_000972575.1
159 Staphylococcus condimenti GCA_001618885.1
159 Staphylococcus epidermidis ATCC 12228 GCA_000007645.1
159 Staphylococcus haemolyticus JCSC1435 GCA_000009865.1
159 Staphylococcus hominis subsp. hominis C80 GCA_000183685.1
159 Staphylococcus lugdunensis HKU09-01 GCA_000025085.1
159 Staphylococcus saprophyticus subsp. saprophyticus ATCC 15305 GCA_000010125.1
159 Staphylococcus simiae CCM 7213 GCA_000235645.2

159 Staphylococcus simulans GCA_001559115.1
159 Staphylococcus succinus GCA_001902315.1
159 Staphylococcus warneri SG1 GCA_000332735.1
159 Staphylococcus xylosus GCA_000706685.1
vh
218 Rhodobacter sphaeroides 2.4.1 GCA_000012905.2
172 Pseudorhodobacter psychrotolerans GCA_001294535.1
165 Rhodobacter sphaeroides ATCC 17025 GCA_000016405.1
vi
146 Bifidobacterium asteroides PRL2011 GCA_000304215.1
137 Bifidobacterium adolescentis ATCC 15703 GCA_000010425.1
130 Bifidobacterium callitrichos DSM 23973 GCA_000741175.1
vj
318 Lactobacillus gasseri ATCC 33323 = JCM 1131 GCA_000014425.1
316 Lactobacillus hominis DSM 23910 = CRBIP 24.179 GCA_000296835.1
237 Lactobacillus psittaci DSM 15354 GCA_000425905.1
vk
246 Megasphaera cerevisiae DSM 20462 GCA_001045675.1
246 Staphylococcus warneri SG1 GCA_000332735.1
242 Staphylococcus xylosus GCA_000706685.1
231 Staphylococcus capitis subsp. capitis GCA_001028645.1
231 Staphylococcus epidermidis ATCC 12228 GCA_000007645.1
231 Staphylococcus lugdunensis HKU09-01 GCA_000025085.1
vl
211 Bifidobacterium adolescentis ATCC 15703 GCA_000010425.1
206 Bifidobacterium dentium JCM 1195 = DSM 20436 GCA_001042595.1
192 Bifidobacterium breve DSM 20213 = JCM 1192 GCA_001025175.1
vm
307 Enterococcus faecalis V583 GCA_000007785.1
307 Streptomyces cinnamoneus GCA_001885705.1
284 Melissococcus plutonius S1 GCA_000747585.1
282 Enterococcus aquimarinus GCA_001885765.1
282 Enterococcus asini ATCC 700915 GCA_000407365.1
282 Enterococcus canis NBRC 100695 GCA_001544375.1
282 Enterococcus dispar ATCC 51266 GCA_000406945.1
282 Enterococcus faecium DO GCA_000174395.2
282 Enterococcus gilvus ATCC BAA-350 GCA_000407545.1
282 Enterococcus haemoperoxidus ATCC BAA-382 GCA_000407165.1
282 Enterococcus hermanniensis GCA_001885945.1
282 Enterococcus hirae ATCC 9790 GCA_000271405.2
282 Enterococcus italicus DSM 15952 GCA_000185365.1
282 Enterococcus malodoratus ATCC 43197 GCA_000407185.1
282 Enterococcus massiliensis GCA_001050095.1
282 Enterococcus mundtii QU 25 GCA_000504125.1
282 Enterococcus pallens ATCC BAA-351 GCA_000407485.1
282 Enterococcus phoeniculicola ATCC BAA-412 GCA_000407505.1
282 Enterococcus pseudoavium NBRC 100491 GCA_001544295.1
282 Enterococcus rivorum GCA_001742285.1
282 Enterococcus sulfureus ATCC 49903 GCA_000407605.1
282 Enterococcus thailandicus GCA_001652875.1
vn
256 Escherichia coli IAI39 GCA_000026345.1
256 Escherichia coli O104_3AH4 str. 2011C-3493 GCA_000299455.1
256 Escherichia coli O157_3AH7 str. Sakai GCA_000008865.1
256 Escherichia coli O83_3AH1 str. NRG 857C GCA_000183345.1
256 Escherichia coli UMN026 GCA_000026325.2
256 Escherichia coli str. K-12 substr. MG1655 GCA_000005845.2
256 Shigella flexneri 2a str. 301 GCA_000006925.2
256 Tumebacillus flagellatus GCA_000714935.1
241 Shigella dysenteriae Sd197 GCA_000012005.1
213 Erwinia iniecta GCA_001267535.1
vo
123 Rhodobacter sphaeroides 2.4.1 GCA_000012905.2
112 Defluviimonas alba GCA_001620265.1
102 Gemmobacter aquatilis GCA_900110025.1
102 Gemmobacter megaterium GCA_900156815.1
102 Pseudorhodobacter psychrotolerans GCA_001294535.1
vp
212 Rhodobacter sphaeroides 2.4.1 GCA_000012905.2
207 Pseudorhodobacter ferrugineus DSM 5888 GCA_000420745.1
207 Pseudorhodobacter wandonensis GCA_001202035.1
207 Rhodobacter sphaeroides ATCC 17025 GCA_000016405.1
193 Gemmobacter megaterium GCA_900156815.1
vq
220 Streptococcus mutans UA159 GCA_000007465.2
153 Streptococcus sobrinus DSM 20742 = ATCC 33478 GCA_000686605.1

134 Streptococcus gordonii str. Challis substr. CH1 GCA_000017005.1
vr
264 Rhodobacter sphaeroides 2.4.1 GCA_000012905.2
226 Gemmobacter megaterium GCA_900156815.1
218 Defluviimonas alba GCA_001620265.1
vs
198 Clostridium beijerinckii GCA_000833105.2
198 Clostridium puniceum GCA_002006345.1
198 Clostridium saccharoperbutylacetonicum N1-4_28HMT_29 GCA_000340885.1
183 Clostridium saccharobutylicum DSM 13864 GCA_000473995.1
180 Clostridium butyricum GCA_001456065.2
vt
111 Lactobacillus gasseri ATCC 33323 = JCM 1131 GCA_000014425.1
111 Lactobacillus hominis DSM 23910 = CRBIP 24.179 GCA_000296835.1
96 Lactobacillus acidophilus NCFM GCA_000011985.1
96 Lactobacillus amylovorus GCA_000191545.1
96 Lactobacillus crispatus ST1 GCA_000091765.1
96 Lactobacillus delbrueckii subsp. bulgaricus ATCC 11842 = JCM 1002 GCA_000056065.1
96 Lactobacillus gallinarum GCA_001314245.2
96 Lactobacillus hamsteri DSM 5661 = JCM 6256 GCA_000615445.1
96 Lactobacillus kalixensis DSM 16043 GCA_001434335.1
94 Bifidobacterium adolescentis ATCC 15703 GCA_000010425.1
vu
266 Deinococcus radiodurans R1 GCA_000008565.1
190 Deinococcus gobiensis I-0 GCA_000252445.1
182 Deinococcus puniceus GCA_001644565.1
vv
155 Rhodobacter sphaeroides 2.4.1 GCA_000012905.2
141 Rhodobacter sphaeroides ATCC 17025 GCA_000016405.1
126 Aquimixticola soesokkakensis GCA_900172375.1
126 Donghicola eburneus GCA_900115865.1
126 Gemmobacter megaterium GCA_900156815.1
126 Jannaschia donghaensis GCA_001403795.1
126 Loktanella atrilutea GCA_900128995.1
126 Loktanella fryxellensis GCA_900110065.1
126 Loktanella salsilacus GCA_900114485.1
126 Oceanicola granulosus HTCC2516 GCA_000153305.1
126 Pseudooceanicola atlanticus GCA_000768315.1
126 Pseudooceanicola batsensis HTCC2597 GCA_000152725.1
126 Pseudooceanicola nitratireducens GCA_900112545.1
126 Pseudorhodobacter ferrugineus DSM 5888 GCA_000420745.1
126 Pseudorhodobacter wandonensis GCA_001202035.1
126 Roseivivax lentus GCA_900156805.1
126 Roseovarius lutimaris GCA_900115165.1
126 Roseovarius mucosus GCA_002080415.1
126 Salinihabitans flavidus GCA_900110425.1
126 Sulfitobacter pseudonitzschiae GCA_002222635.1
126 Thioclava dalianensis GCA_000715505.1
126 Thioclava indica GCA_000714545.1
vw
179 Rhodobacter sphaeroides 2.4.1 GCA_000012905.2
179 Rhodobacter sphaeroides ATCC 17025 GCA_000016405.1
168 Pseudorhodobacter psychrotolerans GCA_001294535.1
163 Pseudorhodobacter ferrugineus DSM 5888 GCA_000420745.1
163 Pseudorhodobacter wandonensis GCA_001202035.1
vx
208 Deinococcus radiodurans R1 GCA_000008565.1
148 Deinococcus deserti VCD115 GCA_000020685.1
138 Deinococcus hopiensis KR-140 GCA_900176165.1
138 Deinococcus puniceus GCA_001644565.1
vy
167 Lactobacillus gasseri ATCC 33323 = JCM 1131 GCA_000014425.1
157 Lactobacillus hominis DSM 23910 = CRBIP 24.179 GCA_000296835.1
142 Lactobacillus iners DSM 13335 GCA_000160875.1
vz
256 Streptococcus mutans UA159 GCA_000007465.2
136 Streptococcus ratti FA-1 = DSM 20564 GCA_000286075.1
128 Streptococcus ferus DSM 20646 GCA_000372425.1
vA
231 Escherichia coli IAI39 GCA_000026345.1
231 Escherichia coli O104_3AH4 str. 2011C-3493 GCA_000299455.1
231 Escherichia coli O157_3AH7 str. Sakai GCA_000008865.1
231 Escherichia coli O83_3AH1 str. NRG 857C GCA_000183345.1
231 Escherichia coli UMN026 GCA_000026325.2
231 Escherichia coli str. K-12 substr. MG1655 GCA_000005845.2
231 Shigella dysenteriae Sd197 GCA_000012005.1

231 Shigella flexneri 2a str. 301 GCA_000006925.2
231 Tumebacillus flagellatus GCA_000714935.1
217 Erwinia iniecta GCA_001267535.1
192 Cronobacter sakazakii GCA_000982825.1
192 Kosakonia cowanii GCA_001975225.1
192 Pantoea ananatis LMG 20103 GCA_000025405.2
192 Pantoea dispersa EGD-AAK13 GCA_000465555.2
192 Shimwellia blattae DSM 4481 = NBRC 105725 GCA_000262305.1
vB
214 Escherichia coli IAI39 GCA_000026345.1
214 Escherichia coli O104_3AH4 str. 2011C-3493 GCA_000299455.1
214 Escherichia coli O157_3AH7 str. Sakai GCA_000008865.1
214 Escherichia coli O83_3AH1 str. NRG 857C GCA_000183345.1
214 Escherichia coli UMN026 GCA_000026325.2
214 Escherichia coli str. K-12 substr. MG1655 GCA_000005845.2
214 Shigella dysenteriae Sd197 GCA_000012005.1
214 Shigella flexneri 2a str. 301 GCA_000006925.2
214 Tumebacillus flagellatus GCA_000714935.1
200 Enterobacter hormaechei subsp. steigerwaltii GCA_001729725.1
200 Erwinia iniecta GCA_001267535.1
200 Erwinia toletana DAPP-PG 735 GCA_000336255.1
200 Klebsiella oxytoca GCA_001022195.1
200 Kosakonia cowanii GCA_001975225.1
200 Kosakonia sacchari SP1 GCA_000300455.4
200 Pantoea agglomerans GCA_001709315.1
200 Pantoea alhagi GCA_002101395.1
200 Pantoea ananatis LMG 20103 GCA_000025405.2
200 Pantoea rwandensis GCA_000759475.1
200 Pantoea septica GCA_002095575.1
200 Pluralibacter gergoviae GCA_000757785.1
200 Pseudescherichia vulneris NBRC 102420 GCA_000759795.1
200 Shimwellia blattae DSM 4481 = NBRC 105725 GCA_000262305.1
197 Enterobacter cloacae subsp. cloacae ATCC 13047 GCA_000025565.1
vC
239 Deinococcus radiodurans R1 GCA_000008565.1
146 Deinococcus deserti VCD115 GCA_000020685.1
140 Deinococcus gobiensis I-0 GCA_000252445.1
vD
268 Clostridium beijerinckii GCA_000833105.2
268 Clostridium saccharoperbutylacetonicum N1-4_28HMT_29 GCA_000340885.1
262 Clostridium puniceum GCA_002006345.1
255 Clostridium saccharobutylicum DSM 13864 GCA_000473995.1
vE
239 Clostridium beijerinckii GCA_000833105.2
239 Clostridium puniceum GCA_002006345.1
239 Clostridium saccharobutylicum DSM 13864 GCA_000473995.1
239 Clostridium saccharoperbutylacetonicum N1-4_28HMT_29 GCA_000340885.1
217 Clostridium chromiireducens GCA_002029255.1
202 Clostridium butyricum GCA_001456065.2
vF
307 Enterococcus faecalis V583 GCA_000007785.1
307 Streptomyces cinnamoneus GCA_001885705.1
277 Enterococcus hirae ATCC 9790 GCA_000271405.2
274 Enterococcus dispar ATCC 51266 GCA_000406945.1
vG
231 Escherichia coli IAI39 GCA_000026345.1
231 Escherichia coli O104_3AH4 str. 2011C-3493 GCA_000299455.1
231 Escherichia coli O157_3AH7 str. Sakai GCA_000008865.1
231 Escherichia coli O83_3AH1 str. NRG 857C GCA_000183345.1
231 Escherichia coli UMN026 GCA_000026325.2
231 Escherichia coli str. K-12 substr. MG1655 GCA_000005845.2
231 Shigella dysenteriae Sd197 GCA_000012005.1
231 Shigella flexneri 2a str. 301 GCA_000006925.2
231 Tumebacillus flagellatus GCA_000714935.1
209 Enterobacter hormaechei subsp. steigerwaltii GCA_001729725.1
209 Erwinia iniecta GCA_001267535.1
209 Erwinia toletana DAPP-PG 735 GCA_000336255.1
209 Klebsiella oxytoca GCA_001022195.1
209 Kosakonia cowanii GCA_001975225.1
209 Kosakonia sacchari SP1 GCA_000300455.4
209 Pluralibacter gergoviae GCA_000757785.1
209 Pseudescherichia vulneris NBRC 102420 GCA_000759795.1
205 Cronobacter sakazakii GCA_000982825.1
205 Enterobacter cloacae subsp. cloacae ATCC 13047 GCA_000025565.1
205 Enterobacter kobei GCA_900185885.1
205 Salmonella enterica subsp. enterica serovar Typhi str. CT18 GCA_000195995.1

205 Salmonella enterica subsp. enterica serovar Typhimurium str. LT2 GCA_000006945.2
vH
190 Lactobacillus gasseri ATCC 33323 = JCM 1131 GCA_000014425.1
182 Lactobacillus hominis DSM 23910 = CRBIP 24.179 GCA_000296835.1
144 Lactobacillus hamsteri DSM 5661 = JCM 6256 GCA_000615445.1
vI
191 Rhodobacter sphaeroides 2.4.1 GCA_000012905.2
162 Gemmobacter megaterium GCA_900156815.1
162 Pseudorhodobacter ferrugineus DSM 5888 GCA_000420745.1
162 Pseudorhodobacter wandonensis GCA_001202035.1
159 Rhodobacter sphaeroides ATCC 17025 GCA_000016405.1
vJ
243 Enterococcus faecalis V583 GCA_000007785.1
243 Streptomyces cinnamoneus GCA_001885705.1
205 Enterococcus canis NBRC 100695 GCA_001544375.1
205 Enterococcus faecium DO GCA_000174395.2
205 Enterococcus hirae ATCC 9790 GCA_000271405.2
205 Enterococcus mundtii QU 25 GCA_000504125.1
205 Enterococcus rivorum GCA_001742285.1
204 Enterococcus asini ATCC 700915 GCA_000407365.1
204 Enterococcus casseliflavus EC20 GCA_000157355.2
204 Enterococcus dispar ATCC 51266 GCA_000406945.1
204 Enterococcus massiliensis GCA_001050095.1
204 Enterococcus saccharolyticus subsp. saccharolyticus ATCC 43076 GCA_000407285.1
vK
265 Clostridium beijerinckii GCA_000833105.2
265 Clostridium puniceum GCA_002006345.1
265 Clostridium saccharoperbutylacetonicum N1-4_28HMT_29 GCA_000340885.1
257 Clostridium saccharobutylicum DSM 13864 GCA_000473995.1
246 Clostridium chromiireducens GCA_002029255.1
vL
251 Escherichia coli IAI39 GCA_000026345.1
251 Escherichia coli O104_3AH4 str. 2011C-3493 GCA_000299455.1
251 Escherichia coli O157_3AH7 str. Sakai GCA_000008865.1
251 Escherichia coli O83_3AH1 str. NRG 857C GCA_000183345.1
251 Escherichia coli UMN026 GCA_000026325.2
251 Escherichia coli str. K-12 substr. MG1655 GCA_000005845.2
251 Shigella dysenteriae Sd197 GCA_000012005.1
251 Shigella flexneri 2a str. 301 GCA_000006925.2
251 Tumebacillus flagellatus GCA_000714935.1
201 Erwinia iniecta GCA_001267535.1
187 Erwinia teleogrylli GCA_001484765.1
vM
114 Lactobacillus gasseri ATCC 33323 = JCM 1131 GCA_000014425.1
114 Lactobacillus hominis DSM 23910 = CRBIP 24.179 GCA_000296835.1
99 Lactobacillus amylovorus GCA_000191545.1
99 Lactobacillus crispatus ST1 GCA_000091765.1
99 Lactobacillus psittaci DSM 15354 GCA_000425905.1
96 Lactobacillus kalixensis DSM 16043 GCA_001434335.1
vN
196 Staphylococcus condimenti GCA_001618885.1
196 Staphylococcus simulans GCA_001559115.1
195 Staphylococcus epidermidis ATCC 12228 GCA_000007645.1
195 Staphylococcus haemolyticus JCSC1435 GCA_000009865.1
189 Staphylococcus capitis subsp. capitis GCA_001028645.1
vO
287 Clostridium beijerinckii GCA_000833105.2
287 Clostridium saccharobutylicum DSM 13864 GCA_000473995.1
287 Clostridium saccharoperbutylacetonicum N1-4_28HMT_29 GCA_000340885.1
283 Clostridium puniceum GCA_002006345.1
268 Clostridium chromiireducens GCA_002029255.1
vP
237 Clostridium beijerinckii GCA_000833105.2
237 Clostridium saccharobutylicum DSM 13864 GCA_000473995.1
237 Clostridium saccharoperbutylacetonicum N1-4_28HMT_29 GCA_000340885.1
230 Clostridium puniceum GCA_002006345.1
225 Clostridium butyricum GCA_001456065.2
vQ
244 Clostridium beijerinckii GCA_000833105.2
244 Clostridium puniceum GCA_002006345.1
244 Clostridium saccharoperbutylacetonicum N1-4_28HMT_29 GCA_000340885.1
236 Clostridium saccharobutylicum DSM 13864 GCA_000473995.1
226 Clostridium butyricum GCA_001456065.2
vR
130 Escherichia coli IAI39 GCA_000026345.1
130 Escherichia coli O104_3AH4 str. 2011C-3493 GCA_000299455.1

130 Escherichia coli O157_3AH7 str. Sakai GCA_000008865.1
130 Escherichia coli O83_3AH1 str. NRG 857C GCA_000183345.1
130 Escherichia coli UMN026 GCA_000026325.2
130 Escherichia coli str. K-12 substr. MG1655 GCA_000005845.2
130 Shigella dysenteriae Sd197 GCA_000012005.1
130 Shigella flexneri 2a str. 301 GCA_000006925.2
130 Tumebacillus flagellatus GCA_000714935.1
127 Enterobacter cloacae subsp. cloacae ATCC 13047 GCA_000025565.1
126 Cronobacter sakazakii GCA_000982825.1
126 Enterobacter hormaechei subsp. steigerwaltii GCA_001729725.1
126 Enterobacter kobei GCA_900185885.1
126 Erwinia iniecta GCA_001267535.1
126 Erwinia toletana DAPP-PG 735 GCA_000336255.1
126 Klebsiella oxytoca GCA_001022195.1
126 Kosakonia cowanii GCA_001975225.1
126 Kosakonia sacchari SP1 GCA_000300455.4
126 Pseudescherichia vulneris NBRC 102420 GCA_000759795.1
126 Salmonella enterica subsp. enterica serovar Typhi str. CT18 GCA_000195995.1
126 Salmonella enterica subsp. enterica serovar Typhimurium str. LT2 GCA_000006945.2
vS
271 Bacillus anthracis str. Ames GCA_000007845.1
271 Bacillus anthracis str. Sterne GCA_000008165.1
271 Bacillus cereus ATCC 14579 GCA_000007825.1
271 Bacillus mycoides GCA_000832605.1
271 Bacillus pseudomycoides DSM 12442 GCA_000161455.1
271 Bacillus thuringiensis YBT-1518 GCA_000497525.2
271 _5BBacillus thuringiensis_5D serovar konkukian str. 97-27 GCA_000008505.1
254 Bacillus manliponensis GCA_000712595.1
246 Bacillus cytotoxicus NVH 391-98 GCA_000017425.1
vT
260 Deinococcus radiodurans R1 GCA_000008565.1
150 Deinococcus puniceus GCA_001644565.1
147 Deinococcus gobiensis I-0 GCA_000252445.1
vU
221 Bacillus thuringiensis YBT-1518 GCA_000497525.2
198 Bacillus anthracis str. Ames GCA_000007845.1
198 Bacillus anthracis str. Sterne GCA_000008165.1
198 Bacillus cereus ATCC 14579 GCA_000007825.1
198 Bacillus mycoides GCA_000832605.1
198 Bacillus pseudomycoides DSM 12442 GCA_000161455.1
198 _5BBacillus thuringiensis_5D serovar konkukian str. 97-27 GCA_000008505.1
190 Oceanobacillus iheyensis HTE831 GCA_000011245.1
vV
183 Staphylococcus capitis subsp. capitis GCA_001028645.1
183 Staphylococcus epidermidis ATCC 12228 GCA_000007645.1
183 Staphylococcus lugdunensis HKU09-01 GCA_000025085.1
170 Staphylococcus cohnii subsp. cohnii GCA_000972575.1
170 Staphylococcus saprophyticus subsp. saprophyticus ATCC 15305 GCA_000010125.1
168 Staphylococcus haemolyticus JCSC1435 GCA_000009865.1
168 Staphylococcus hominis subsp. hominis C80 GCA_000183685.1
168 Staphylococcus microti GCA_000934465.1
vW
197 Rhodobacter sphaeroides 2.4.1 GCA_000012905.2
147 Rhodobacter sphaeroides ATCC 17025 GCA_000016405.1
140 Gemmobacter megaterium GCA_900156815.1
vX
147 Bacillus anthracis str. Ames GCA_000007845.1
147 Bacillus anthracis str. Sterne GCA_000008165.1
147 Bacillus cereus ATCC 14579 GCA_000007825.1
147 Bacillus mycoides GCA_000832605.1
147 Bacillus pseudomycoides DSM 12442 GCA_000161455.1
147 Bacillus thuringiensis YBT-1518 GCA_000497525.2
147 _5BBacillus thuringiensis_5D serovar konkukian str. 97-27 GCA_000008505.1
129 Bacillus cytotoxicus NVH 391-98 GCA_000017425.1
113 Bacillus manliponensis GCA_000712595.1
vY
193 Clostridium beijerinckii GCA_000833105.2
193 Clostridium puniceum GCA_002006345.1
193 Clostridium saccharobutylicum DSM 13864 GCA_000473995.1
193 Clostridium saccharoperbutylacetonicum N1-4_28HMT_29 GCA_000340885.1
190 Clostridium neonatale GCA_001458595.1
182 Clostridium butyricum GCA_001456065.2
182 Clostridium chromiireducens GCA_002029255.1
vZ
331 Bacillus cereus ATCC 14579 GCA_000007825.1
316 Bacillus anthracis str. Ames GCA_000007845.1

316 Bacillus anthracis str. Sterne GCA_000008165.1
316 Bacillus mycoides GCA_000832605.1
316 Bacillus pseudomycoides DSM 12442 GCA_000161455.1
316 Bacillus thuringiensis YBT-1518 GCA_000497525.2
316 _5BBacillus thuringiensis_5D serovar konkukian str. 97-27 GCA_000008505.1
274 Bacillus cytotoxicus NVH 391-98 GCA_000017425.1
w0
223 Bifidobacterium adolescentis ATCC 15703 GCA_000010425.1
219 Bifidobacterium dentium JCM 1195 = DSM 20436 GCA_001042595.1
219 Bifidobacterium tsurumiense GCA_000741765.1
218 Bifidobacterium breve DSM 20213 = JCM 1192 GCA_001025175.1
218 Bifidobacterium gallicum DSM 20093 = LMG 11596 GCA_000741205.1
w1
229 Clostridium puniceum GCA_002006345.1
225 Clostridium beijerinckii GCA_000833105.2
225 Clostridium saccharobutylicum DSM 13864 GCA_000473995.1
225 Clostridium saccharoperbutylacetonicum N1-4_28HMT_29 GCA_000340885.1
210 Clostridium chromiireducens GCA_002029255.1
w2
294 Streptococcus mutans UA159 GCA_000007465.2
204 Streptococcus salivarius GCA_000785515.1
204 Streptococcus thermophilus JIM 8232 GCA_000253395.1
203 Streptococcus ratti FA-1 = DSM 20564 GCA_000286075.1
w3
254 Escherichia coli IAI39 GCA_000026345.1
254 Escherichia coli O104_3AH4 str. 2011C-3493 GCA_000299455.1
254 Escherichia coli O157_3AH7 str. Sakai GCA_000008865.1
254 Escherichia coli O83_3AH1 str. NRG 857C GCA_000183345.1
254 Escherichia coli UMN026 GCA_000026325.2
254 Escherichia coli str. K-12 substr. MG1655 GCA_000005845.2
254 Shigella dysenteriae Sd197 GCA_000012005.1
254 Shigella flexneri 2a str. 301 GCA_000006925.2
254 Tumebacillus flagellatus GCA_000714935.1
248 Erwinia iniecta GCA_001267535.1
236 Cedecea neteri GCA_000757825.1
236 Rosenbergiella nectarea GCA_900111105.1
w4
186 Escherichia coli IAI39 GCA_000026345.1
186 Escherichia coli O104_3AH4 str. 2011C-3493 GCA_000299455.1
186 Escherichia coli O157_3AH7 str. Sakai GCA_000008865.1
186 Escherichia coli O83_3AH1 str. NRG 857C GCA_000183345.1
186 Escherichia coli UMN026 GCA_000026325.2
186 Escherichia coli str. K-12 substr. MG1655 GCA_000005845.2
186 Shigella flexneri 2a str. 301 GCA_000006925.2
186 Tumebacillus flagellatus GCA_000714935.1
180 Erwinia iniecta GCA_001267535.1
180 Erwinia toletana DAPP-PG 735 GCA_000336255.1
180 Pantoea ananatis LMG 20103 GCA_000025405.2
180 Shimwellia blattae DSM 4481 = NBRC 105725 GCA_000262305.1
176 Cronobacter sakazakii GCA_000982825.1
176 Pantoea dispersa EGD-AAK13 GCA_000465555.2
176 Plautia stali symbiont GCA_000180175.2
w5
315 Deinococcus radiodurans R1 GCA_000008565.1
221 Deinococcus gobiensis I-0 GCA_000252445.1
212 Deinococcus marmoris DSM 12784 GCA_000701405.1
212 Deinococcus swuensis GCA_000800395.1
w6
322 Rhodobacter sphaeroides 2.4.1 GCA_000012905.2
267 Rhodobacter sphaeroides ATCC 17025 GCA_000016405.1
264 Pseudorhodobacter psychrotolerans GCA_001294535.1
w7
240 Streptococcus mutans UA159 GCA_000007465.2
125 Streptococcus massiliensis DSM 18628 GCA_000380065.1
119 Streptococcus ferus DSM 20646 GCA_000372425.1
w8
246 Clostridium saccharobutylicum DSM 13864 GCA_000473995.1
235 Clostridium beijerinckii GCA_000833105.2
235 Clostridium puniceum GCA_002006345.1
235 Clostridium saccharoperbutylacetonicum N1-4_28HMT_29 GCA_000340885.1
231 Clostridium chromiireducens GCA_002029255.1
w9
238 Streptococcus mutans UA159 GCA_000007465.2
146 Streptococcus sobrinus DSM 20742 = ATCC 33478 GCA_000686605.1
134 Streptococcus gordonii str. Challis substr. CH1 GCA_000017005.1
wa

98 Rhodobacter sphaeroides 2.4.1 GCA_000012905.2
89 Escherichia coli IAI39 GCA_000026345.1
89 Escherichia coli O104_3AH4 str. 2011C-3493 GCA_000299455.1
89 Escherichia coli O157_3AH7 str. Sakai GCA_000008865.1
89 Escherichia coli O83_3AH1 str. NRG 857C GCA_000183345.1
89 Escherichia coli UMN026 GCA_000026325.2
89 Escherichia coli str. K-12 substr. MG1655 GCA_000005845.2
89 Shigella dysenteriae Sd197 GCA_000012005.1
89 Shigella flexneri 2a str. 301 GCA_000006925.2
89 Tumebacillus flagellatus GCA_000714935.1
85 Xenorhabdus bovienii SS-2004 GCA_000027225.1
wb
266 Deinococcus radiodurans R1 GCA_000008565.1
134 Deinococcus deserti VCD115 GCA_000020685.1
134 Deinococcus proteolyticus MRP GCA_000190555.1
131 Deinococcus marmoris DSM 12784 GCA_000701405.1
131 Deinococcus swuensis GCA_000800395.1
wc
137 Escherichia coli IAI39 GCA_000026345.1
137 Escherichia coli O104_3AH4 str. 2011C-3493 GCA_000299455.1
137 Escherichia coli O157_3AH7 str. Sakai GCA_000008865.1
137 Escherichia coli UMN026 GCA_000026325.2
137 Escherichia coli str. K-12 substr. MG1655 GCA_000005845.2
137 Shigella dysenteriae Sd197 GCA_000012005.1
137 Shigella flexneri 2a str. 301 GCA_000006925.2
137 Tumebacillus flagellatus GCA_000714935.1
122 Escherichia coli O83_3AH1 str. NRG 857C GCA_000183345.1
122 _5BEnterobacter_5D lignolyticus SCF1 GCA_000164865.1
107 Buttiauxella ferragutiae ATCC 51602 GCA_001654915.1
107 Cedecea neteri GCA_000757825.1
107 Citrobacter freundii CFNIH1 GCA_000648515.1
107 Cronobacter sakazakii GCA_000982825.1
107 Edwardsiella anguillarum ET080813 GCA_000264765.2
107 Enterobacter cloacae subsp. cloacae ATCC 13047 GCA_000025565.1
107 Enterobacter hormaechei subsp. steigerwaltii GCA_001729725.1
107 Enterobacter kobei GCA_900185885.1
107 Erwinia amylovora CFBP1430 GCA_000091565.1
107 Erwinia billingiae Eb661 GCA_000196615.1
107 Erwinia gerundensis GCA_001517405.1
107 Erwinia iniecta GCA_001267535.1
107 Erwinia persicina NBRC 102418 GCA_001571305.1
107 Erwinia toletana DAPP-PG 735 GCA_000336255.1
107 Klebsiella aerogenes KCTC 2190 GCA_000215745.1
107 Klebsiella oxytoca GCA_001022195.1
107 Kluyvera ascorbata ATCC 33433 GCA_000735365.1
107 Kosakonia cowanii GCA_001975225.1
107 Kosakonia sacchari SP1 GCA_000300455.4
107 Mangrovibacter phragmitis GCA_001655675.1
107 Morganella morganii subsp. morganii KT GCA_000286435.2
107 Obesumbacterium proteus GCA_001586165.1
107 Pantoea agglomerans GCA_001709315.1
107 Pantoea alhagi GCA_002101395.1
107 Pantoea ananatis LMG 20103 GCA_000025405.2
107 Pantoea dispersa EGD-AAK13 GCA_000465555.2
107 Pantoea rwandensis GCA_000759475.1
107 Pantoea septica GCA_002095575.1
107 Photorhabdus temperata subsp. thracensis GCA_001010285.1
107 Pluralibacter gergoviae GCA_000757785.1
107 Pseudescherichia vulneris NBRC 102420 GCA_000759795.1
107 Rosenbergiella nectarea GCA_900111105.1
107 Salmonella enterica subsp. enterica serovar Typhi str. CT18 GCA_000195995.1
107 Salmonella enterica subsp. enterica serovar Typhimurium str. LT2 GCA_000006945.2
107 Shimwellia blattae DSM 4481 = NBRC 105725 GCA_000262305.1
107 Tatumella citrea GCA_002163585.1
107 Tatumella saanichensis GCA_000439375.1
107 Trabulsiella odontotermitis GCA_001297765.1
107 Xenorhabdus bovienii SS-2004 GCA_000027225.1
107 Xenorhabdus doucetiae GCA_000968195.1
107 Xenorhabdus hominickii GCA_001721185.1
107 Xenorhabdus innexi GCA_900155355.1
107 Xenorhabdus japonica GCA_900115195.1
107 Xenorhabdus koppenhoeferi GCA_900116635.1
107 Xenorhabdus nematophila AN6_2F1 GCA_000953355.1
107 Yokenella regensburgei ATCC 49455 GCA_000735455.1
wd
266 Bifidobacterium adolescentis ATCC 15703 GCA_000010425.1

250 Bifidobacterium asteroides PRL2011 GCA_000304215.1
244 Bifidobacterium dentium JCM 1195 = DSM 20436 GCA_001042595.1
we
170 Rhodobacter sphaeroides 2.4.1 GCA_000012905.2
154 Pseudorhodobacter psychrotolerans GCA_001294535.1
149 Rhodobacter capsulatus SB 1003 GCA_000021865.1
wf
218 Clostridium beijerinckii GCA_000833105.2
218 Clostridium saccharoperbutylacetonicum N1-4_28HMT_29 GCA_000340885.1
211 Clostridium puniceum GCA_002006345.1
203 Clostridium butyricum GCA_001456065.2
203 Clostridium saccharobutylicum DSM 13864 GCA_000473995.1
wg
291 Escherichia coli IAI39 GCA_000026345.1
291 Escherichia coli O104_3AH4 str. 2011C-3493 GCA_000299455.1
291 Escherichia coli O157_3AH7 str. Sakai GCA_000008865.1
291 Escherichia coli O83_3AH1 str. NRG 857C GCA_000183345.1
291 Escherichia coli UMN026 GCA_000026325.2
291 Escherichia coli str. K-12 substr. MG1655 GCA_000005845.2
291 Shigella flexneri 2a str. 301 GCA_000006925.2
291 Tumebacillus flagellatus GCA_000714935.1
282 Shigella dysenteriae Sd197 GCA_000012005.1
268 Erwinia iniecta GCA_001267535.1
wh
154 Lactobacillus gasseri ATCC 33323 = JCM 1131 GCA_000014425.1
154 Lactobacillus hominis DSM 23910 = CRBIP 24.179 GCA_000296835.1
111 Lactobacillus amylovorus GCA_000191545.1
111 Lactobacillus crispatus ST1 GCA_000091765.1
111 Lactobacillus kalixensis DSM 16043 GCA_001434335.1
111 Lactobacillus psittaci DSM 15354 GCA_000425905.1
108 Lactobacillus secaliphilus GCA_001437055.1
108 Lactobacillus zymae GCA_900183405.1
wi
150 Rhodobacter sphaeroides 2.4.1 GCA_000012905.2
116 Defluviimonas alba GCA_001620265.1
115 Haematobacter massiliensis GCA_000740795.1
115 Oceanicola granulosus HTCC2516 GCA_000153305.1
wj
245 Lactobacillus gasseri ATCC 33323 = JCM 1131 GCA_000014425.1
222 Lactobacillus hominis DSM 23910 = CRBIP 24.179 GCA_000296835.1
155 Lactobacillus iners DSM 13335 GCA_000160875.1
wk
218 Clostridium beijerinckii GCA_000833105.2
218 Clostridium puniceum GCA_002006345.1
218 Clostridium saccharobutylicum DSM 13864 GCA_000473995.1
218 Clostridium saccharoperbutylacetonicum N1-4_28HMT_29 GCA_000340885.1
207 Clostridium botulinum B str. Eklund 17B _28NRP_29 GCA_000020165.1
207 Clostridium chromiireducens GCA_002029255.1
207 Clostridium taeniosporum GCA_001735765.1
205 Clostridium uliginosum GCA_900112485.1
wl
126 Deinococcus radiodurans R1 GCA_000008565.1
94 Deinococcus deserti VCD115 GCA_000020685.1
86 Deinococcus gobiensis I-0 GCA_000252445.1
wm
299 Staphylococcus capitis subsp. capitis GCA_001028645.1
299 Staphylococcus epidermidis ATCC 12228 GCA_000007645.1
296 Staphylococcus haemolyticus JCSC1435 GCA_000009865.1
291 Staphylococcus cohnii subsp. cohnii GCA_000972575.1
wn
253 Streptococcus mutans UA159 GCA_000007465.2
158 Streptococcus macacae NCTC 11558 GCA_000187995.3
150 Streptococcus ratti FA-1 = DSM 20564 GCA_000286075.1
wo
226 Bacillus anthracis str. Ames GCA_000007845.1
226 Bacillus anthracis str. Sterne GCA_000008165.1
226 Bacillus cereus ATCC 14579 GCA_000007825.1
226 Bacillus mycoides GCA_000832605.1
226 Bacillus pseudomycoides DSM 12442 GCA_000161455.1
226 Bacillus thuringiensis YBT-1518 GCA_000497525.2
226 _5BBacillus thuringiensis_5D serovar konkukian str. 97-27 GCA_000008505.1
190 Bacillus smithii GCA_001050115.1
186 Bacillus humi GCA_001439915.1
186 Bacillus sinesaloumensis GCA_900156865.1
186 Bacillus timonensis GCA_000285535.1
wp

267 Clostridium beijerinckii GCA_000833105.2
267 Clostridium puniceum GCA_002006345.1
267 Clostridium saccharobutylicum DSM 13864 GCA_000473995.1
267 Clostridium saccharoperbutylacetonicum N1-4_28HMT_29 GCA_000340885.1
249 Clostridium chromiireducens GCA_002029255.1
245 Clostridium butyricum GCA_001456065.2
wq
208 Enterococcus faecalis V583 GCA_000007785.1
208 Streptomyces cinnamoneus GCA_001885705.1
179 Enterococcus canis NBRC 100695 GCA_001544375.1
179 Enterococcus casseliflavus EC20 GCA_000157355.2
179 Enterococcus dispar ATCC 51266 GCA_000406945.1
179 Enterococcus faecium DO GCA_000174395.2
179 Enterococcus haemoperoxidus ATCC BAA-382 GCA_000407165.1
179 Enterococcus hirae ATCC 9790 GCA_000271405.2
179 Enterococcus mundtii QU 25 GCA_000504125.1
179 Enterococcus phoeniculicola ATCC BAA-412 GCA_000407505.1
179 Enterococcus rivorum GCA_001742285.1
179 Enterococcus saccharolyticus subsp. saccharolyticus ATCC 43076 GCA_000407285.1
179 Enterococcus thailandicus GCA_001652875.1
176 Salinicoccus carnicancri Crm GCA_000330705.1
wr
260 Clostridium beijerinckii GCA_000833105.2
260 Clostridium saccharobutylicum DSM 13864 GCA_000473995.1
260 Clostridium saccharoperbutylacetonicum N1-4_28HMT_29 GCA_000340885.1
256 Clostridium puniceum GCA_002006345.1
241 Clostridium chromiireducens GCA_002029255.1
ws
293 Streptococcus mutans UA159 GCA_000007465.2
200 Streptococcus macacae NCTC 11558 GCA_000187995.3
188 Streptococcus iniae GCA_000831485.1
wt
196 Enterococcus faecalis V583 GCA_000007785.1
196 Streptomyces cinnamoneus GCA_001885705.1
164 Enterococcus canis NBRC 100695 GCA_001544375.1
164 Enterococcus faecium DO GCA_000174395.2
164 Enterococcus haemoperoxidus ATCC BAA-382 GCA_000407165.1
164 Enterococcus hirae ATCC 9790 GCA_000271405.2
164 Enterococcus mundtii QU 25 GCA_000504125.1
164 Enterococcus phoeniculicola ATCC BAA-412 GCA_000407505.1
164 Enterococcus rivorum GCA_001742285.1
164 Enterococcus thailandicus GCA_001652875.1
164 Isobaculum melis GCA_900111355.1
163 Enterococcus aquimarinus GCA_001885765.1
163 Enterococcus asini ATCC 700915 GCA_000407365.1
163 Enterococcus dispar ATCC 51266 GCA_000406945.1
163 Enterococcus gilvus ATCC BAA-350 GCA_000407545.1
163 Enterococcus hermanniensis GCA_001885945.1
163 Enterococcus malodoratus ATCC 43197 GCA_000407185.1
163 Enterococcus massiliensis GCA_001050095.1
163 Enterococcus pallens ATCC BAA-351 GCA_000407485.1
163 Enterococcus pseudoavium NBRC 100491 GCA_001544295.1
wu
159 Staphylococcus capitis subsp. capitis GCA_001028645.1
159 Staphylococcus cohnii subsp. cohnii GCA_000972575.1
159 Staphylococcus condimenti GCA_001618885.1
159 Staphylococcus epidermidis ATCC 12228 GCA_000007645.1
159 Staphylococcus haemolyticus JCSC1435 GCA_000009865.1
159 Staphylococcus hominis subsp. hominis C80 GCA_000183685.1
159 Staphylococcus pettenkoferi GCA_002208805.1
159 Staphylococcus simulans GCA_001559115.1
152 Staphylococcus arlettae CVD059 GCA_000295715.1
152 Staphylococcus lugdunensis HKU09-01 GCA_000025085.1
152 Staphylococcus saprophyticus subsp. saprophyticus ATCC 15305 GCA_000010125.1
144 Megasphaera cerevisiae DSM 20462 GCA_001045675.1
144 Staphylococcus aureus subsp. aureus NCTC 8325 GCA_000013425.1
144 Staphylococcus simiae CCM 7213 GCA_000235645.2
144 Staphylococcus succinus GCA_001902315.1
144 Staphylococcus warneri SG1 GCA_000332735.1
wv
236 Bifidobacterium adolescentis ATCC 15703 GCA_000010425.1
224 Bifidobacterium breve DSM 20213 = JCM 1192 GCA_001025175.1
214 Bifidobacterium tsurumiense GCA_000741765.1
ww
308 Clostridium beijerinckii GCA_000833105.2
308 Clostridium puniceum GCA_002006345.1

308 Clostridium saccharoperbutylacetonicum N1-4_28HMT_29 GCA_000340885.1
293 Clostridium saccharobutylicum DSM 13864 GCA_000473995.1
285 Clostridium butyricum GCA_001456065.2
wx
209 Bifidobacterium adolescentis ATCC 15703 GCA_000010425.1
190 Bifidobacterium reuteri DSM 23975 GCA_000741695.1
188 Bifidobacterium gallicum DSM 20093 = LMG 11596 GCA_000741205.1
wy
256 Enterococcus faecalis V583 GCA_000007785.1
256 Streptomyces cinnamoneus GCA_001885705.1
229 Enterococcus canis NBRC 100695 GCA_001544375.1
229 Enterococcus casseliflavus EC20 GCA_000157355.2
229 Enterococcus dispar ATCC 51266 GCA_000406945.1
229 Enterococcus faecium DO GCA_000174395.2
229 Enterococcus haemoperoxidus ATCC BAA-382 GCA_000407165.1
229 Enterococcus hirae ATCC 9790 GCA_000271405.2
229 Enterococcus mundtii QU 25 GCA_000504125.1
229 Enterococcus phoeniculicola ATCC BAA-412 GCA_000407505.1
229 Enterococcus rivorum GCA_001742285.1
229 Enterococcus saccharolyticus subsp. saccharolyticus ATCC 43076 GCA_000407285.1
229 Enterococcus thailandicus GCA_001652875.1
214 Enterococcus asini ATCC 700915 GCA_000407365.1
214 Enterococcus massiliensis GCA_001050095.1
wz
231 Bacillus anthracis str. Ames GCA_000007845.1
231 Bacillus anthracis str. Sterne GCA_000008165.1
231 Bacillus cereus ATCC 14579 GCA_000007825.1
231 Bacillus mycoides GCA_000832605.1
231 Bacillus pseudomycoides DSM 12442 GCA_000161455.1
231 Bacillus thuringiensis YBT-1518 GCA_000497525.2
231 _5BBacillus thuringiensis_5D serovar konkukian str. 97-27 GCA_000008505.1
196 Bacillus coahuilensis m4-4 GCA_000171615.1
190 Bacillus marisflavi GCA_001274775.1
wA
249 Escherichia coli IAI39 GCA_000026345.1
249 Escherichia coli O104_3AH4 str. 2011C-3493 GCA_000299455.1
249 Escherichia coli O157_3AH7 str. Sakai GCA_000008865.1
249 Escherichia coli O83_3AH1 str. NRG 857C GCA_000183345.1
249 Escherichia coli UMN026 GCA_000026325.2
249 Escherichia coli str. K-12 substr. MG1655 GCA_000005845.2
249 Shigella flexneri 2a str. 301 GCA_000006925.2
249 Tumebacillus flagellatus GCA_000714935.1
234 Shigella dysenteriae Sd197 GCA_000012005.1
215 Cronobacter sakazakii GCA_000982825.1
215 Erwinia iniecta GCA_001267535.1
215 Erwinia toletana DAPP-PG 735 GCA_000336255.1
215 Xenorhabdus bovienii SS-2004 GCA_000027225.1
wB
262 Bacillus anthracis str. Ames GCA_000007845.1
262 Bacillus anthracis str. Sterne GCA_000008165.1
262 Bacillus mycoides GCA_000832605.1
262 Bacillus pseudomycoides DSM 12442 GCA_000161455.1
262 Bacillus thuringiensis YBT-1518 GCA_000497525.2
262 _5BBacillus thuringiensis_5D serovar konkukian str. 97-27 GCA_000008505.1
261 Bacillus cereus ATCC 14579 GCA_000007825.1
215 Bacillus cytotoxicus NVH 391-98 GCA_000017425.1
wC
245 Clostridium beijerinckii GCA_000833105.2
245 Clostridium saccharoperbutylacetonicum N1-4_28HMT_29 GCA_000340885.1
231 Clostridium puniceum GCA_002006345.1
230 Clostridium saccharobutylicum DSM 13864 GCA_000473995.1
wD
328 Deinococcus radiodurans R1 GCA_000008565.1
234 Deinococcus gobiensis I-0 GCA_000252445.1
217 Deinococcus puniceus GCA_001644565.1
wE
242 Streptococcus mutans UA159 GCA_000007465.2
178 Streptococcus ferus DSM 20646 GCA_000372425.1
162 Streptococcus massiliensis DSM 18628 GCA_000380065.1
wF
214 Enterococcus faecalis V583 GCA_000007785.1
214 Streptomyces cinnamoneus GCA_001885705.1
200 Enterococcus hirae ATCC 9790 GCA_000271405.2
199 Enterococcus asini ATCC 700915 GCA_000407365.1
199 Enterococcus canis NBRC 100695 GCA_001544375.1
199 Enterococcus casseliflavus EC20 GCA_000157355.2

199 Enterococcus dispar ATCC 51266 GCA_000406945.1
199 Enterococcus faecium DO GCA_000174395.2
199 Enterococcus haemoperoxidus ATCC BAA-382 GCA_000407165.1
199 Enterococcus massiliensis GCA_001050095.1
199 Enterococcus mundtii QU 25 GCA_000504125.1
199 Enterococcus phoeniculicola ATCC BAA-412 GCA_000407505.1
199 Enterococcus rivorum GCA_001742285.1
199 Enterococcus saccharolyticus subsp. saccharolyticus ATCC 43076 GCA_000407285.1
199 Enterococcus thailandicus GCA_001652875.1
199 Vagococcus penaei GCA_001998885.1
wG
175 Clostridium beijerinckii GCA_000833105.2
175 Clostridium saccharoperbutylacetonicum N1-4_28HMT_29 GCA_000340885.1
161 Clostridium puniceum GCA_002006345.1
161 Clostridium saccharobutylicum DSM 13864 GCA_000473995.1
151 Clostridium butyricum GCA_001456065.2
wH
258 Rhodobacter sphaeroides 2.4.1 GCA_000012905.2
242 Rhodobacter sphaeroides ATCC 17025 GCA_000016405.1
236 Gemmobacter megaterium GCA_900156815.1
wI
185 Deinococcus radiodurans R1 GCA_000008565.1
147 Deinococcus deserti VCD115 GCA_000020685.1
137 Deinococcus puniceus GCA_001644565.1
wJ
216 Erwinia iniecta GCA_001267535.1
216 Escherichia coli IAI39 GCA_000026345.1
216 Escherichia coli O104_3AH4 str. 2011C-3493 GCA_000299455.1
216 Escherichia coli O157_3AH7 str. Sakai GCA_000008865.1
216 Escherichia coli O83_3AH1 str. NRG 857C GCA_000183345.1
216 Escherichia coli UMN026 GCA_000026325.2
216 Escherichia coli str. K-12 substr. MG1655 GCA_000005845.2
216 Shigella flexneri 2a str. 301 GCA_000006925.2
216 Tumebacillus flagellatus GCA_000714935.1
207 Shigella dysenteriae Sd197 GCA_000012005.1
199 Cronobacter sakazakii GCA_000982825.1
199 Pantoea ananatis LMG 20103 GCA_000025405.2
199 Pantoea dispersa EGD-AAK13 GCA_000465555.2
199 Shimwellia blattae DSM 4481 = NBRC 105725 GCA_000262305.1
wK
248 Rhodobacter sphaeroides 2.4.1 GCA_000012905.2
244 Gemmobacter aquatilis GCA_900110025.1
233 Pseudorhodobacter psychrotolerans GCA_001294535.1
wL
256 Bifidobacterium adolescentis ATCC 15703 GCA_000010425.1
233 Bifidobacterium dentium JCM 1195 = DSM 20436 GCA_001042595.1
229 Bifidobacterium callitrichos DSM 23973 GCA_000741175.1
wM
171 Clostridium beijerinckii GCA_000833105.2
171 Clostridium puniceum GCA_002006345.1
171 Clostridium saccharobutylicum DSM 13864 GCA_000473995.1
171 Clostridium saccharoperbutylacetonicum N1-4_28HMT_29 GCA_000340885.1
158 Clostridium chromiireducens GCA_002029255.1
153 Clostridium taeniosporum GCA_001735765.1
wN
188 Bacillus thuringiensis YBT-1518 GCA_000497525.2
187 Bacillus anthracis str. Ames GCA_000007845.1
187 Bacillus anthracis str. Sterne GCA_000008165.1
187 Bacillus cereus ATCC 14579 GCA_000007825.1
187 Bacillus mycoides GCA_000832605.1
187 Bacillus pseudomycoides DSM 12442 GCA_000161455.1
187 _5BBacillus thuringiensis_5D serovar konkukian str. 97-27 GCA_000008505.1
158 Bacillus cytotoxicus NVH 391-98 GCA_000017425.1
wO
230 Clostridium beijerinckii GCA_000833105.2
230 Clostridium puniceum GCA_002006345.1
230 Clostridium saccharoperbutylacetonicum N1-4_28HMT_29 GCA_000340885.1
227 Clostridium saccharobutylicum DSM 13864 GCA_000473995.1
212 Clostridium butyricum GCA_001456065.2
212 Clostridium chromiireducens GCA_002029255.1
wP
273 Enterococcus faecalis V583 GCA_000007785.1
273 Streptomyces cinnamoneus GCA_001885705.1
236 Enterococcus dispar ATCC 51266 GCA_000406945.1
236 Enterococcus massiliensis GCA_001050095.1
235 Enterococcus canis NBRC 100695 GCA_001544375.1

235 Enterococcus faecium DO GCA_000174395.2
235 Enterococcus haemoperoxidus ATCC BAA-382 GCA_000407165.1
235 Enterococcus hirae ATCC 9790 GCA_000271405.2
235 Enterococcus mundtii QU 25 GCA_000504125.1
235 Enterococcus phoeniculicola ATCC BAA-412 GCA_000407505.1
235 Enterococcus rivorum GCA_001742285.1
235 Enterococcus thailandicus GCA_001652875.1
wQ
259 Bacillus anthracis str. Ames GCA_000007845.1
259 Bacillus anthracis str. Sterne GCA_000008165.1
259 Bacillus cereus ATCC 14579 GCA_000007825.1
259 Bacillus mycoides GCA_000832605.1
259 Bacillus pseudomycoides DSM 12442 GCA_000161455.1
259 Bacillus thuringiensis YBT-1518 GCA_000497525.2
259 _5BBacillus thuringiensis_5D serovar konkukian str. 97-27 GCA_000008505.1
177 Bacillus manliponensis GCA_000712595.1
167 Bacillus cytotoxicus NVH 391-98 GCA_000017425.1
wR
149 Bacillus anthracis str. Ames GCA_000007845.1
149 Bacillus anthracis str. Sterne GCA_000008165.1
149 Bacillus mycoides GCA_000832605.1
149 Bacillus pseudomycoides DSM 12442 GCA_000161455.1
149 Bacillus thuringiensis YBT-1518 GCA_000497525.2
149 _5BBacillus thuringiensis_5D serovar konkukian str. 97-27 GCA_000008505.1
134 Bacillus cereus ATCC 14579 GCA_000007825.1
126 Bacillus flexus GCA_002024265.1
126 Bacillus megaterium NBRC 15308 = ATCC 14581 GCA_000832985.1
wS
95 Bacillus anthracis str. Ames GCA_000007845.1
95 Bacillus anthracis str. Sterne GCA_000008165.1
95 Bacillus cereus ATCC 14579 GCA_000007825.1
95 Bacillus mycoides GCA_000832605.1
95 Bacillus pseudomycoides DSM 12442 GCA_000161455.1
95 Bacillus thuringiensis YBT-1518 GCA_000497525.2
95 _5BBacillus thuringiensis_5D serovar konkukian str. 97-27 GCA_000008505.1
89 Bacillus akibai JCM 9157 GCA_000513135.1
89 Bacillus alcalophilus ATCC 27647 = CGMCC 1.3604 GCA_000292245.2
89 Bacillus hemicellulosilyticus JCM 9152 GCA_000513115.1
89 Bacillus okuhidensis GCA_001274915.1
89 Bacillus pseudofirmus OF4 GCA_000005825.2
89 Bacillus trypoxylicola GCA_001590785.1
83 Bacillus cohnii NBRC 15565 GCA_001591425.1
wT
177 Staphylococcus capitis subsp. capitis GCA_001028645.1
177 Staphylococcus cohnii subsp. cohnii GCA_000972575.1
177 Staphylococcus epidermidis ATCC 12228 GCA_000007645.1
162 Megasphaera cerevisiae DSM 20462 GCA_001045675.1
162 Staphylococcus haemolyticus JCSC1435 GCA_000009865.1
162 Staphylococcus hominis subsp. hominis C80 GCA_000183685.1
162 Staphylococcus lugdunensis HKU09-01 GCA_000025085.1
162 Staphylococcus saprophyticus subsp. saprophyticus ATCC 15305 GCA_000010125.1
162 Staphylococcus warneri SG1 GCA_000332735.1
150 Staphylococcus condimenti GCA_001618885.1
150 Staphylococcus simulans GCA_001559115.1
wU
238 Rhodobacter sphaeroides 2.4.1 GCA_000012905.2
207 Gemmobacter megaterium GCA_900156815.1
200 Pseudorhodobacter ferrugineus DSM 5888 GCA_000420745.1
200 Pseudorhodobacter wandonensis GCA_001202035.1
wV
202 Clostridium beijerinckii GCA_000833105.2
202 Clostridium puniceum GCA_002006345.1
202 Clostridium saccharoperbutylacetonicum N1-4_28HMT_29 GCA_000340885.1
197 Clostridium butyricum GCA_001456065.2
194 Clostridium saccharobutylicum DSM 13864 GCA_000473995.1
wW
358 Streptococcus mutans UA159 GCA_000007465.2
236 Streptococcus macacae NCTC 11558 GCA_000187995.3
228 Streptococcus ferus DSM 20646 GCA_000372425.1
228 Streptococcus sobrinus DSM 20742 = ATCC 33478 GCA_000686605.1
wX
291 Deinococcus radiodurans R1 GCA_000008565.1
218 Deinococcus gobiensis I-0 GCA_000252445.1
200 Deinococcus puniceus GCA_001644565.1
wY
285 Deinococcus radiodurans R1 GCA_000008565.1

190 Deinococcus gobiensis I-0 GCA_000252445.1
184 Deinococcus hopiensis KR-140 GCA_900176165.1
wZ
201 Bifidobacterium thermophilum GCA_000741495.1
190 Bifidobacterium adolescentis ATCC 15703 GCA_000010425.1
190 Bifidobacterium dentium JCM 1195 = DSM 20436 GCA_001042595.1
190 Bifidobacterium stellenboschense GCA_000741785.1
187 Bifidobacterium angulatum DSM 20098 = JCM 7096 GCA_001025155.1
x0
211 Clostridium saccharobutylicum DSM 13864 GCA_000473995.1
207 Clostridium beijerinckii GCA_000833105.2
207 Clostridium butyricum GCA_001456065.2
207 Clostridium saccharoperbutylacetonicum N1-4_28HMT_29 GCA_000340885.1
203 Clostridium chromiireducens GCA_002029255.1
203 Clostridium puniceum GCA_002006345.1
x1
253 Rhodobacter sphaeroides 2.4.1 GCA_000012905.2
201 Pseudorhodobacter ferrugineus DSM 5888 GCA_000420745.1
201 Pseudorhodobacter wandonensis GCA_001202035.1
201 Rhodobacter sphaeroides ATCC 17025 GCA_000016405.1
188 Thioclava dalianensis GCA_000715505.1
188 Thioclava indica GCA_000714545.1
x2
165 Streptococcus mutans UA159 GCA_000007465.2
141 Tetragenococcus halophilus NBRC 12172 GCA_000283615.1
137 Streptococcus equinus GCA_000964315.1
137 Streptococcus gallolyticus subsp. gallolyticus DSM 16831 GCA_002000985.1
x3
192 Clostridium beijerinckii GCA_000833105.2
192 Clostridium puniceum GCA_002006345.1
192 Clostridium saccharobutylicum DSM 13864 GCA_000473995.1
192 Clostridium saccharoperbutylacetonicum N1-4_28HMT_29 GCA_000340885.1
181 Clostridium chromiireducens GCA_002029255.1
155 Clostridium botulinum B str. Eklund 17B _28NRP_29 GCA_000020165.1
155 Clostridium taeniosporum GCA_001735765.1
x4
185 Lactobacillus gasseri ATCC 33323 = JCM 1131 GCA_000014425.1
165 Lactobacillus hominis DSM 23910 = CRBIP 24.179 GCA_000296835.1
130 Lactobacillus acidophilus NCFM GCA_000011985.1
130 Lactobacillus gallinarum GCA_001314245.2
x5
138 Cronobacter sakazakii GCA_000982825.1
138 Erwinia iniecta GCA_001267535.1
138 Erwinia toletana DAPP-PG 735 GCA_000336255.1
138 Escherichia coli IAI39 GCA_000026345.1
138 Escherichia coli O104_3AH4 str. 2011C-3493 GCA_000299455.1
138 Escherichia coli O157_3AH7 str. Sakai GCA_000008865.1
138 Escherichia coli UMN026 GCA_000026325.2
138 Escherichia coli str. K-12 substr. MG1655 GCA_000005845.2
138 Shigella dysenteriae Sd197 GCA_000012005.1
138 Shigella flexneri 2a str. 301 GCA_000006925.2
138 Tumebacillus flagellatus GCA_000714935.1
137 Pantoea agglomerans GCA_001709315.1
136 Erwinia gerundensis GCA_001517405.1
136 Pantoea ananatis LMG 20103 GCA_000025405.2
136 Pantoea dispersa EGD-AAK13 GCA_000465555.2
136 Plautia stali symbiont GCA_000180175.2
136 Shimwellia blattae DSM 4481 = NBRC 105725 GCA_000262305.1
x6
177 Clostridium chromiireducens GCA_002029255.1
177 Clostridium saccharobutylicum DSM 13864 GCA_000473995.1
175 Clostridium beijerinckii GCA_000833105.2
175 Clostridium puniceum GCA_002006345.1
175 Clostridium saccharoperbutylacetonicum N1-4_28HMT_29 GCA_000340885.1
174 Clostridium butyricum GCA_001456065.2
x7
251 Streptococcus mutans UA159 GCA_000007465.2
145 Streptococcus cristatus AS 1.3089 GCA_000385925.1
145 Streptococcus gordonii str. Challis substr. CH1 GCA_000017005.1
145 Streptococcus mitis B6 GCA_000027165.1
145 Streptococcus pneumoniae R6 GCA_000007045.1
139 Bacillus azotoformans LMG 9581 GCA_000307855.1
139 Bacillus bataviensis LMG 21833 GCA_000307875.1
139 Lactococcus lactis subsp. lactis Il1403 GCA_000006865.1
139 Streptococcus parasanguinis ATCC 15912 GCA_000164675.2
x8

302 Lactobacillus gasseri ATCC 33323 = JCM 1131 GCA_000014425.1
285 Lactobacillus hominis DSM 23910 = CRBIP 24.179 GCA_000296835.1
223 Lactobacillus psittaci DSM 15354 GCA_000425905.1
x9
226 Escherichia coli IAI39 GCA_000026345.1
226 Escherichia coli O104_3AH4 str. 2011C-3493 GCA_000299455.1
226 Escherichia coli O157_3AH7 str. Sakai GCA_000008865.1
226 Escherichia coli O83_3AH1 str. NRG 857C GCA_000183345.1
226 Escherichia coli UMN026 GCA_000026325.2
226 Escherichia coli str. K-12 substr. MG1655 GCA_000005845.2
226 Shigella flexneri 2a str. 301 GCA_000006925.2
226 Tumebacillus flagellatus GCA_000714935.1
216 Shigella dysenteriae Sd197 GCA_000012005.1
201 Cronobacter sakazakii GCA_000982825.1
201 Erwinia iniecta GCA_001267535.1
201 Erwinia toletana DAPP-PG 735 GCA_000336255.1
201 Kosakonia cowanii GCA_001975225.1
201 Pluralibacter gergoviae GCA_000757785.1
xa
235 Lactobacillus gasseri ATCC 33323 = JCM 1131 GCA_000014425.1
219 Lactobacillus hominis DSM 23910 = CRBIP 24.179 GCA_000296835.1
171 Lactobacillus crispatus ST1 GCA_000091765.1
xb
230 Deinococcus radiodurans R1 GCA_000008565.1
152 Deinococcus deserti VCD115 GCA_000020685.1
123 Deinococcus puniceus GCA_001644565.1
xc
174 Escherichia coli IAI39 GCA_000026345.1
174 Escherichia coli O104_3AH4 str. 2011C-3493 GCA_000299455.1
174 Escherichia coli O157_3AH7 str. Sakai GCA_000008865.1
174 Escherichia coli O83_3AH1 str. NRG 857C GCA_000183345.1
174 Escherichia coli UMN026 GCA_000026325.2
174 Escherichia coli str. K-12 substr. MG1655 GCA_000005845.2
174 Shigella flexneri 2a str. 301 GCA_000006925.2
174 Tumebacillus flagellatus GCA_000714935.1
165 Shigella dysenteriae Sd197 GCA_000012005.1
159 Erwinia iniecta GCA_001267535.1
xd
268 Deinococcus radiodurans R1 GCA_000008565.1
161 Deinococcus deserti VCD115 GCA_000020685.1
152 Deinococcus puniceus GCA_001644565.1
xe
292 Deinococcus radiodurans R1 GCA_000008565.1
213 Deinococcus gobiensis I-0 GCA_000252445.1
199 Deinococcus puniceus GCA_001644565.1
xf
153 Deinococcus radiodurans R1 GCA_000008565.1
88 Deinococcus gobiensis I-0 GCA_000252445.1
87 Deinococcus puniceus GCA_001644565.1
xg
135 Streptococcus mutans UA159 GCA_000007465.2
77 Streptococcus ratti FA-1 = DSM 20564 GCA_000286075.1
76 Streptococcus equinus GCA_000964315.1
76 Streptococcus gallolyticus subsp. gallolyticus DSM 16831 GCA_002000985.1
76 Streptococcus orisratti DSM 15617 GCA_000380105.1
xh
226 Clostridium beijerinckii GCA_000833105.2
226 Clostridium saccharobutylicum DSM 13864 GCA_000473995.1
226 Clostridium saccharoperbutylacetonicum N1-4_28HMT_29 GCA_000340885.1
219 Clostridium puniceum GCA_002006345.1
204 Clostridium butyricum GCA_001456065.2
204 Clostridium chromiireducens GCA_002029255.1
xi
244 Streptococcus mutans UA159 GCA_000007465.2
152 Streptococcus gordonii str. Challis substr. CH1 GCA_000017005.1
148 Streptococcus marimammalium DSM 18627 GCA_000380045.1
xj
251 Bacillus anthracis str. Ames GCA_000007845.1
251 Bacillus anthracis str. Sterne GCA_000008165.1
251 Bacillus cereus ATCC 14579 GCA_000007825.1
251 Bacillus mycoides GCA_000832605.1
251 Bacillus pseudomycoides DSM 12442 GCA_000161455.1
251 Bacillus thuringiensis YBT-1518 GCA_000497525.2
251 _5BBacillus thuringiensis_5D serovar konkukian str. 97-27 GCA_000008505.1
203 Bacillus cytotoxicus NVH 391-98 GCA_000017425.1
201 Bacillus manliponensis GCA_000712595.1

xk
248 Lactobacillus gasseri ATCC 33323 = JCM 1131 GCA_000014425.1
241 Lactobacillus hominis DSM 23910 = CRBIP 24.179 GCA_000296835.1
216 Lactobacillus jensenii GCA_001936235.1
xl
134 Clostridium beijerinckii GCA_000833105.2
134 Clostridium puniceum GCA_002006345.1
134 Clostridium saccharoperbutylacetonicum N1-4_28HMT_29 GCA_000340885.1
119 Clostridium saccharobutylicum DSM 13864 GCA_000473995.1
119 Clostridium taeniosporum GCA_001735765.1
114 Clostridium butyricum GCA_001456065.2
xm
269 Escherichia coli IAI39 GCA_000026345.1
269 Escherichia coli O104_3AH4 str. 2011C-3493 GCA_000299455.1
269 Escherichia coli O157_3AH7 str. Sakai GCA_000008865.1
269 Escherichia coli UMN026 GCA_000026325.2
269 Escherichia coli str. K-12 substr. MG1655 GCA_000005845.2
269 Shigella flexneri 2a str. 301 GCA_000006925.2
269 Tumebacillus flagellatus GCA_000714935.1
268 Escherichia coli O83_3AH1 str. NRG 857C GCA_000183345.1
260 Shigella dysenteriae Sd197 GCA_000012005.1
xn
183 Deinococcus radiodurans R1 GCA_000008565.1
129 Deinococcus puniceus GCA_001644565.1
128 Deinococcus deserti VCD115 GCA_000020685.1
xo
258 Clostridium beijerinckii GCA_000833105.2
258 Clostridium saccharoperbutylacetonicum N1-4_28HMT_29 GCA_000340885.1
251 Clostridium puniceum GCA_002006345.1
249 Clostridium saccharobutylicum DSM 13864 GCA_000473995.1
xp
134 Clostridium disporicum GCA_001405015.1
134 Clostridium neonatale GCA_001458595.1
132 Clostridium beijerinckii GCA_000833105.2
132 Clostridium butyricum GCA_001456065.2
132 Clostridium fallax GCA_900129365.1
132 Clostridium puniceum GCA_002006345.1
132 Clostridium saccharobutylicum DSM 13864 GCA_000473995.1
132 Clostridium saccharoperbutylacetonicum N1-4_28HMT_29 GCA_000340885.1
130 Clostridium celatum DSM 1785 GCA_000320405.1
130 Clostridium chauvoei JF4335 GCA_900168365.1
130 Clostridium sartagoforme AAU1 GCA_000401215.1
xq
239 Clostridium beijerinckii GCA_000833105.2
239 Clostridium puniceum GCA_002006345.1
239 Clostridium saccharoperbutylacetonicum N1-4_28HMT_29 GCA_000340885.1
230 Clostridium saccharobutylicum DSM 13864 GCA_000473995.1
225 Clostridium butyricum GCA_001456065.2
xr
203 Streptococcus mutans UA159 GCA_000007465.2
173 Streptococcus marimammalium DSM 18627 GCA_000380045.1
168 Streptococcus gordonii str. Challis substr. CH1 GCA_000017005.1
xs
187 Clostridium beijerinckii GCA_000833105.2
187 Clostridium puniceum GCA_002006345.1
187 Clostridium saccharoperbutylacetonicum N1-4_28HMT_29 GCA_000340885.1
183 Clostridium butyricum GCA_001456065.2
176 Clostridium taeniosporum GCA_001735765.1
xt
229 Clostridium beijerinckii GCA_000833105.2
229 Clostridium saccharoperbutylacetonicum N1-4_28HMT_29 GCA_000340885.1
222 Clostridium puniceum GCA_002006345.1
220 Clostridium saccharobutylicum DSM 13864 GCA_000473995.1
xu
203 Bacillus licheniformis DSM 13 = ATCC 14580 GCA_000011645.1
200 Massilibacterium senegalense GCA_001375675.1
198 Bacillus okuhidensis GCA_001274915.1
xv
250 Lactobacillus gasseri ATCC 33323 = JCM 1131 GCA_000014425.1
231 Lactobacillus hominis DSM 23910 = CRBIP 24.179 GCA_000296835.1
184 Lactobacillus crispatus ST1 GCA_000091765.1
184 Lactobacillus helveticus GCA_001308285.1
xw
160 Bacillus anthracis str. Ames GCA_000007845.1
160 Bacillus anthracis str. Sterne GCA_000008165.1
160 Bacillus cereus ATCC 14579 GCA_000007825.1

160 Bacillus mycoides GCA_000832605.1
160 Bacillus pseudomycoides DSM 12442 GCA_000161455.1
160 Bacillus thuringiensis YBT-1518 GCA_000497525.2
160 _5BBacillus thuringiensis_5D serovar konkukian str. 97-27 GCA_000008505.1
146 Bacillus horneckiae GCA_001636335.1
146 Bacillus solani GCA_001420595.1
136 Bacillus acidicola GCA_001636425.1
136 Bacillus dakarensis GCA_900156875.1
136 Bacillus shackletonii GCA_001420715.1
136 Bacillus sporothermodurans GCA_001587375.1
136 Bacillus tuaregi GCA_900104575.1
xx
120 Clostridium beijerinckii GCA_000833105.2
120 Clostridium puniceum GCA_002006345.1
120 Clostridium saccharobutylicum DSM 13864 GCA_000473995.1
120 Clostridium saccharoperbutylacetonicum N1-4_28HMT_29 GCA_000340885.1
110 Clostridium neonatale GCA_001458595.1
103 Clostridium celatum DSM 1785 GCA_000320405.1
xy
233 Deinococcus radiodurans R1 GCA_000008565.1
120 Deinococcus gobiensis I-0 GCA_000252445.1
94 Deinococcus marmoris DSM 12784 GCA_000701405.1
94 Deinococcus swuensis GCA_000800395.1
xz
275 Streptococcus mutans UA159 GCA_000007465.2
203 Streptococcus gordonii str. Challis substr. CH1 GCA_000017005.1
199 Streptococcus cristatus AS 1.3089 GCA_000385925.1
199 Streptococcus mitis B6 GCA_000027165.1
199 Streptococcus pneumoniae R6 GCA_000007045.1
xA
236 Deinococcus radiodurans R1 GCA_000008565.1
153 Deinococcus gobiensis I-0 GCA_000252445.1
141 Deinococcus marmoris DSM 12784 GCA_000701405.1
141 Deinococcus swuensis GCA_000800395.1
xB
130 Bifidobacterium adolescentis ATCC 15703 GCA_000010425.1
107 Bifidobacterium bohemicum DSM 22767 GCA_000741525.1
107 Bifidobacterium thermophilum GCA_000741495.1
107 Bifidobacterium thermophilum RBL67 GCA_000347695.1
105 Bifidobacterium breve DSM 20213 = JCM 1192 GCA_001025175.1
xC
185 Clostridium beijerinckii GCA_000833105.2
185 Clostridium puniceum GCA_002006345.1
185 Clostridium saccharoperbutylacetonicum N1-4_28HMT_29 GCA_000340885.1
170 Clostridium saccharobutylicum DSM 13864 GCA_000473995.1
136 Clostridium butyricum GCA_001456065.2
136 Clostridium chromiireducens GCA_002029255.1
xD
209 Bifidobacterium adolescentis ATCC 15703 GCA_000010425.1
159 Bifidobacterium asteroides PRL2011 GCA_000304215.1
159 Bifidobacterium callitrichos DSM 23973 GCA_000741175.1
153 Bifidobacterium breve DSM 20213 = JCM 1192 GCA_001025175.1
xE
168 Escherichia coli IAI39 GCA_000026345.1
168 Escherichia coli O104_3AH4 str. 2011C-3493 GCA_000299455.1
168 Escherichia coli O157_3AH7 str. Sakai GCA_000008865.1
168 Escherichia coli O83_3AH1 str. NRG 857C GCA_000183345.1
168 Escherichia coli UMN026 GCA_000026325.2
168 Escherichia coli str. K-12 substr. MG1655 GCA_000005845.2
168 Shigella flexneri 2a str. 301 GCA_000006925.2
168 Tumebacillus flagellatus GCA_000714935.1
162 Erwinia iniecta GCA_001267535.1
156 Cronobacter sakazakii GCA_000982825.1
156 Erwinia toletana DAPP-PG 735 GCA_000336255.1
xF
254 Clostridium saccharobutylicum DSM 13864 GCA_000473995.1
251 Clostridium butyricum GCA_001456065.2
248 Clostridium beijerinckii GCA_000833105.2
248 Clostridium puniceum GCA_002006345.1
248 Clostridium saccharoperbutylacetonicum N1-4_28HMT_29 GCA_000340885.1
xG
221 Deinococcus radiodurans R1 GCA_000008565.1
120 Deinococcus frigens DSM 12807 GCA_000701425.1
119 Deinococcus hopiensis KR-140 GCA_900176165.1
119 Deinococcus puniceus GCA_001644565.1
xH

145 Staphylococcus capitis subsp. capitis GCA_001028645.1
145 Staphylococcus epidermidis ATCC 12228 GCA_000007645.1
142 Staphylococcus haemolyticus JCSC1435 GCA_000009865.1
136 Staphylococcus cohnii subsp. cohnii GCA_000972575.1
136 Staphylococcus lugdunensis HKU09-01 GCA_000025085.1
136 Staphylococcus saprophyticus subsp. saprophyticus ATCC 15305 GCA_000010125.1
136 Staphylococcus xylosus GCA_000706685.1
xI
96 Rhodobacter sphaeroides 2.4.1 GCA_000012905.2
96 Rhodobacter sphaeroides ATCC 17025 GCA_000016405.1
81 Gemmobacter megaterium GCA_900156815.1
81 Pseudorhodobacter ferrugineus DSM 5888 GCA_000420745.1
81 Pseudorhodobacter wandonensis GCA_001202035.1
66 Gemmobacter aquatilis GCA_900110025.1
66 Pseudorhodobacter psychrotolerans GCA_001294535.1
xJ
154 Escherichia coli IAI39 GCA_000026345.1
154 Escherichia coli O104_3AH4 str. 2011C-3493 GCA_000299455.1
154 Escherichia coli O157_3AH7 str. Sakai GCA_000008865.1
154 Escherichia coli O83_3AH1 str. NRG 857C GCA_000183345.1
154 Escherichia coli UMN026 GCA_000026325.2
154 Escherichia coli str. K-12 substr. MG1655 GCA_000005845.2
154 Shigella dysenteriae Sd197 GCA_000012005.1
154 Shigella flexneri 2a str. 301 GCA_000006925.2
154 Tumebacillus flagellatus GCA_000714935.1
143 Plesiomonas shigelloides GCA_900087055.1
139 Buttiauxella ferragutiae ATCC 51602 GCA_001654915.1
139 Cedecea neteri GCA_000757825.1
139 Citrobacter freundii CFNIH1 GCA_000648515.1
139 Klebsiella aerogenes KCTC 2190 GCA_000215745.1
139 Kluyvera ascorbata ATCC 33433 GCA_000735365.1
139 Obesumbacterium proteus GCA_001586165.1
139 Tatumella saanichensis GCA_000439375.1
139 Yokenella regensburgei ATCC 49455 GCA_000735455.1
139 _5BEnterobacter_5D lignolyticus SCF1 GCA_000164865.1
xK
222 Lonsdalea quercina subsp. quercina GCA_000688655.1
219 Escherichia coli IAI39 GCA_000026345.1
219 Escherichia coli O104_3AH4 str. 2011C-3493 GCA_000299455.1
219 Escherichia coli O157_3AH7 str. Sakai GCA_000008865.1
219 Escherichia coli O83_3AH1 str. NRG 857C GCA_000183345.1
219 Escherichia coli UMN026 GCA_000026325.2
219 Escherichia coli str. K-12 substr. MG1655 GCA_000005845.2
219 Shigella dysenteriae Sd197 GCA_000012005.1
219 Shigella flexneri 2a str. 301 GCA_000006925.2
219 Tumebacillus flagellatus GCA_000714935.1
205 Erwinia iniecta GCA_001267535.1
xL
71 Clostridium intestinale URNW GCA_000469625.2
65 Clostridium beijerinckii GCA_000833105.2
65 Clostridium puniceum GCA_002006345.1
65 Clostridium saccharoperbutylacetonicum N1-4_28HMT_29 GCA_000340885.1
64 Clostridium algidicarnis GCA_000703125.1
xM
261 Streptococcus mutans UA159 GCA_000007465.2
176 Streptococcus ratti FA-1 = DSM 20564 GCA_000286075.1
174 Streptococcus macacae NCTC 11558 GCA_000187995.3
xN
216 Escherichia coli IAI39 GCA_000026345.1
216 Escherichia coli O104_3AH4 str. 2011C-3493 GCA_000299455.1
216 Escherichia coli O157_3AH7 str. Sakai GCA_000008865.1
216 Escherichia coli O83_3AH1 str. NRG 857C GCA_000183345.1
216 Escherichia coli UMN026 GCA_000026325.2
216 Escherichia coli str. K-12 substr. MG1655 GCA_000005845.2
216 Shigella dysenteriae Sd197 GCA_000012005.1
216 Shigella flexneri 2a str. 301 GCA_000006925.2
216 Tumebacillus flagellatus GCA_000714935.1
180 Erwinia iniecta GCA_001267535.1
167 Enterobacter cloacae subsp. cloacae ATCC 13047 GCA_000025565.1
xO
239 Escherichia coli IAI39 GCA_000026345.1
239 Escherichia coli O104_3AH4 str. 2011C-3493 GCA_000299455.1
239 Escherichia coli O157_3AH7 str. Sakai GCA_000008865.1
239 Escherichia coli O83_3AH1 str. NRG 857C GCA_000183345.1
239 Escherichia coli UMN026 GCA_000026325.2
239 Escherichia coli str. K-12 substr. MG1655 GCA_000005845.2

239 Shigella flexneri 2a str. 301 GCA_000006925.2
239 Tumebacillus flagellatus GCA_000714935.1
230 Shigella dysenteriae Sd197 GCA_000012005.1
225 Cronobacter sakazakii GCA_000982825.1
225 Erwinia iniecta GCA_001267535.1
225 Erwinia toletana DAPP-PG 735 GCA_000336255.1
xP
224 Deinococcus radiodurans R1 GCA_000008565.1
157 Deinococcus marmoris DSM 12784 GCA_000701405.1
157 Deinococcus swuensis GCA_000800395.1
147 Deinococcus gobiensis I-0 GCA_000252445.1
xQ
137 Rhodobacter sphaeroides 2.4.1 GCA_000012905.2
123 Aliiroseovarius crassostreae GCA_001307765.1
123 Aliiroseovarius sediminilitoris GCA_900109955.1
123 Gemmobacter megaterium GCA_900156815.1
123 Jannaschia donghaensis GCA_001403795.1
123 Jannaschia faecimaris GCA_900107415.1
123 Maribius pelagius GCA_900110115.1
123 Marivita cryptomonadis GCA_002115725.1
123 Oceanicola granulosus HTCC2516 GCA_000153305.1
123 Pseudorhodobacter ferrugineus DSM 5888 GCA_000420745.1
123 Pseudorhodobacter wandonensis GCA_001202035.1
123 Xuhuaishuia manganoxidans GCA_001981245.1
122 Rhodobacter sphaeroides ATCC 17025 GCA_000016405.1
xR
201 Escherichia coli IAI39 GCA_000026345.1
201 Escherichia coli O104_3AH4 str. 2011C-3493 GCA_000299455.1
201 Escherichia coli O157_3AH7 str. Sakai GCA_000008865.1
201 Escherichia coli O83_3AH1 str. NRG 857C GCA_000183345.1
201 Escherichia coli UMN026 GCA_000026325.2
201 Escherichia coli str. K-12 substr. MG1655 GCA_000005845.2
201 Shigella flexneri 2a str. 301 GCA_000006925.2
201 Tumebacillus flagellatus GCA_000714935.1
197 Cronobacter sakazakii GCA_000982825.1
197 Erwinia amylovora CFBP1430 GCA_000091565.1
197 Erwinia iniecta GCA_001267535.1
197 Erwinia toletana DAPP-PG 735 GCA_000336255.1
197 Kosakonia cowanii GCA_001975225.1
197 Pantoea ananatis LMG 20103 GCA_000025405.2
197 Pantoea dispersa EGD-AAK13 GCA_000465555.2
197 Pluralibacter gergoviae GCA_000757785.1
197 Shimwellia blattae DSM 4481 = NBRC 105725 GCA_000262305.1
192 Plautia stali symbiont GCA_000180175.2
192 Shigella dysenteriae Sd197 GCA_000012005.1
xS
228 Clostridium beijerinckii GCA_000833105.2
228 Clostridium puniceum GCA_002006345.1
228 Clostridium saccharoperbutylacetonicum N1-4_28HMT_29 GCA_000340885.1
227 Clostridium taeniosporum GCA_001735765.1
214 Clostridium chromiireducens GCA_002029255.1
214 Clostridium saccharobutylicum DSM 13864 GCA_000473995.1
xT
219 Escherichia coli IAI39 GCA_000026345.1
219 Escherichia coli O104_3AH4 str. 2011C-3493 GCA_000299455.1
219 Escherichia coli O157_3AH7 str. Sakai GCA_000008865.1
219 Escherichia coli O83_3AH1 str. NRG 857C GCA_000183345.1
219 Escherichia coli UMN026 GCA_000026325.2
219 Escherichia coli str. K-12 substr. MG1655 GCA_000005845.2
219 Shigella flexneri 2a str. 301 GCA_000006925.2
219 Tumebacillus flagellatus GCA_000714935.1
210 Shigella dysenteriae Sd197 GCA_000012005.1
185 Erwinia iniecta GCA_001267535.1
xU
232 Streptococcus mutans UA159 GCA_000007465.2
177 Streptococcus cristatus AS 1.3089 GCA_000385925.1
175 Streptococcus gordonii str. Challis substr. CH1 GCA_000017005.1
xV
282 Streptococcus mutans UA159 GCA_000007465.2
208 Streptococcus ferus DSM 20646 GCA_000372425.1
195 Streptococcus sobrinus DSM 20742 = ATCC 33478 GCA_000686605.1
xW
256 Bacillus anthracis str. Ames GCA_000007845.1
256 Bacillus anthracis str. Sterne GCA_000008165.1
256 Bacillus cereus ATCC 14579 GCA_000007825.1
256 Bacillus mycoides GCA_000832605.1

256 Bacillus pseudomycoides DSM 12442 GCA_000161455.1
256 Bacillus thuringiensis YBT-1518 GCA_000497525.2
256 _5BBacillus thuringiensis_5D serovar konkukian str. 97-27 GCA_000008505.1
220 Bacillus manliponensis GCA_000712595.1
200 Anaerobacillus alkalilacustris GCA_001866005.1
200 Bacillus solimangrovi GCA_001742425.1
200 Listeria floridensis FSL S10-1187 GCA_000525875.1
xX
243 Bacillus anthracis str. Ames GCA_000007845.1
243 Bacillus anthracis str. Sterne GCA_000008165.1
243 Bacillus cereus ATCC 14579 GCA_000007825.1
243 Bacillus mycoides GCA_000832605.1
243 Bacillus pseudomycoides DSM 12442 GCA_000161455.1
243 Bacillus thuringiensis YBT-1518 GCA_000497525.2
243 _5BBacillus thuringiensis_5D serovar konkukian str. 97-27 GCA_000008505.1
197 Staphylococcus saprophyticus GCA_001074355.1
194 Bacillus manliponensis GCA_000712595.1
xY
237 Bifidobacterium adolescentis ATCC 15703 GCA_000010425.1
218 Bifidobacterium callitrichos DSM 23973 GCA_000741175.1
217 Bifidobacterium dentium JCM 1195 = DSM 20436 GCA_001042595.1
xZ
256 Clostridium beijerinckii GCA_000833105.2
256 Clostridium puniceum GCA_002006345.1
256 Clostridium saccharoperbutylacetonicum N1-4_28HMT_29 GCA_000340885.1
241 Clostridium saccharobutylicum DSM 13864 GCA_000473995.1
223 Clostridium butyricum GCA_001456065.2
y0
110 Bifidobacterium adolescentis ATCC 15703 GCA_000010425.1
109 Bifidobacterium asteroides PRL2011 GCA_000304215.1
105 Bifidobacterium thermophilum GCA_000741495.1
105 Bifidobacterium thermophilum RBL67 GCA_000347695.1
y1
214 Enterococcus faecalis V583 GCA_000007785.1
214 Streptomyces cinnamoneus GCA_001885705.1
189 Enterococcus asini ATCC 700915 GCA_000407365.1
189 Enterococcus canis NBRC 100695 GCA_001544375.1
189 Enterococcus casseliflavus EC20 GCA_000157355.2
189 Enterococcus dispar ATCC 51266 GCA_000406945.1
189 Enterococcus faecium DO GCA_000174395.2
189 Enterococcus hirae ATCC 9790 GCA_000271405.2
189 Enterococcus massiliensis GCA_001050095.1
189 Enterococcus mundtii QU 25 GCA_000504125.1
189 Enterococcus rivorum GCA_001742285.1
189 Enterococcus saccharolyticus subsp. saccharolyticus ATCC 43076 GCA_000407285.1
189 Melissococcus plutonius S1 GCA_000747585.1
174 Enterococcus cecorum GCA_001318405.1
174 Enterococcus columbae DSM 7374 = ATCC 51263 GCA_000406925.1
174 Enterococcus haemoperoxidus ATCC BAA-382 GCA_000407165.1
174 Enterococcus phoeniculicola ATCC BAA-412 GCA_000407505.1
174 Enterococcus thailandicus GCA_001652875.1
y2
227 Deinococcus radiodurans R1 GCA_000008565.1
154 Deinococcus gobiensis I-0 GCA_000252445.1
136 Deinococcus deserti VCD115 GCA_000020685.1
y3
223 Clostridium beijerinckii GCA_000833105.2
223 Clostridium puniceum GCA_002006345.1
223 Clostridium saccharobutylicum DSM 13864 GCA_000473995.1
223 Clostridium saccharoperbutylacetonicum N1-4_28HMT_29 GCA_000340885.1
216 Clostridium chromiireducens GCA_002029255.1
216 Clostridium taeniosporum GCA_001735765.1
193 Clostridium botulinum B str. Eklund 17B _28NRP_29 GCA_000020165.1
y4
245 Clostridium puniceum GCA_002006345.1
229 Clostridium beijerinckii GCA_000833105.2
229 Clostridium saccharobutylicum DSM 13864 GCA_000473995.1
229 Clostridium saccharoperbutylacetonicum N1-4_28HMT_29 GCA_000340885.1
218 Clostridium chromiireducens GCA_002029255.1
y5
233 Bacillus anthracis str. Ames GCA_000007845.1
233 Bacillus anthracis str. Sterne GCA_000008165.1
233 Bacillus cereus ATCC 14579 GCA_000007825.1
233 Bacillus mycoides GCA_000832605.1
233 Bacillus pseudomycoides DSM 12442 GCA_000161455.1
233 Bacillus thuringiensis YBT-1518 GCA_000497525.2

233 _5BBacillus thuringiensis_5D serovar konkukian str. 97-27 GCA_000008505.1
189 Bacillus manliponensis GCA_000712595.1
186 Bacillus aquimaris TF-12 GCA_001648555.1
y6
228 Clostridium beijerinckii GCA_000833105.2
228 Clostridium saccharobutylicum DSM 13864 GCA_000473995.1
228 Clostridium saccharoperbutylacetonicum N1-4_28HMT_29 GCA_000340885.1
218 Clostridium neonatale GCA_001458595.1
211 Clostridium puniceum GCA_002006345.1
y7
164 Rhodobacter sphaeroides 2.4.1 GCA_000012905.2
140 Pseudorhodobacter psychrotolerans GCA_001294535.1
135 Rhodobacter sphaeroides ATCC 17025 GCA_000016405.1
y8
192 Clostridium beijerinckii GCA_000833105.2
192 Clostridium puniceum GCA_002006345.1
192 Clostridium saccharobutylicum DSM 13864 GCA_000473995.1
192 Clostridium saccharoperbutylacetonicum N1-4_28HMT_29 GCA_000340885.1
181 Clostridium botulinum B str. Eklund 17B _28NRP_29 GCA_000020165.1
181 Clostridium chromiireducens GCA_002029255.1
181 Clostridium taeniosporum GCA_001735765.1
174 Clostridium uliginosum GCA_900112485.1
y9
193 Rhodobacter sphaeroides 2.4.1 GCA_000012905.2
178 Rhodobacter sphaeroides ATCC 17025 GCA_000016405.1
152 Gemmobacter aquatilis GCA_900110025.1
152 Pseudorhodobacter psychrotolerans GCA_001294535.1
ya
267 Bifidobacterium adolescentis ATCC 15703 GCA_000010425.1
225 Bifidobacterium angulatum DSM 20098 = JCM 7096 GCA_001025155.1
222 Bifidobacterium thermophilum GCA_000741495.1
yb
124 Deinococcus radiodurans R1 GCA_000008565.1
92 Deinococcus gobiensis I-0 GCA_000252445.1
85 Deinococcus puniceus GCA_001644565.1
yc
220 Deinococcus radiodurans R1 GCA_000008565.1
145 Deinococcus deserti VCD115 GCA_000020685.1
135 Deinococcus gobiensis I-0 GCA_000252445.1
yd
214 Staphylococcus capitis subsp. capitis GCA_001028645.1
214 Staphylococcus epidermidis ATCC 12228 GCA_000007645.1
214 Staphylococcus haemolyticus JCSC1435 GCA_000009865.1
204 Staphylococcus cohnii subsp. cohnii GCA_000972575.1
204 Staphylococcus gallinarum GCA_000875895.1
204 Staphylococcus saprophyticus subsp. saprophyticus ATCC 15305 GCA_000010125.1
204 Staphylococcus succinus GCA_001902315.1
204 Staphylococcus xylosus GCA_000706685.1
202 Staphylococcus pettenkoferi GCA_002208805.1
ye
244 Bifidobacterium adolescentis ATCC 15703 GCA_000010425.1
205 Bifidobacterium asteroides PRL2011 GCA_000304215.1
205 Bifidobacterium bifidum PRL2010 GCA_000165905.1
202 Bifidobacterium callitrichos DSM 23973 GCA_000741175.1
yf
188 Escherichia coli IAI39 GCA_000026345.1
188 Escherichia coli O104_3AH4 str. 2011C-3493 GCA_000299455.1
188 Escherichia coli O157_3AH7 str. Sakai GCA_000008865.1
188 Escherichia coli O83_3AH1 str. NRG 857C GCA_000183345.1
188 Escherichia coli UMN026 GCA_000026325.2
188 Escherichia coli str. K-12 substr. MG1655 GCA_000005845.2
188 Tumebacillus flagellatus GCA_000714935.1
180 Shigella flexneri 2a str. 301 GCA_000006925.2
178 Shigella dysenteriae Sd197 GCA_000012005.1
yg
137 Lactobacillus gasseri ATCC 33323 = JCM 1131 GCA_000014425.1
122 Lactobacillus hominis DSM 23910 = CRBIP 24.179 GCA_000296835.1
109 Lactobacillus fabifermentans DSM 21115 GCA_000498955.2
109 Lactobacillus herbarum GCA_001039045.1
109 Lactobacillus oryzae JCM 18671 GCA_000740055.1
109 Lactobacillus ozensis DSM 23829 = JCM 17196 GCA_001435995.1
109 Lactobacillus plantarum WCFS1 GCA_000203855.3
yh
206 Staphylococcus epidermidis ATCC 12228 GCA_000007645.1
206 Staphylococcus lugdunensis HKU09-01 GCA_000025085.1
205 Staphylococcus cohnii subsp. cohnii GCA_000972575.1

195 Staphylococcus condimenti GCA_001618885.1
195 Staphylococcus simulans GCA_001559115.1
yi
152 Rhodobacter sphaeroides 2.4.1 GCA_000012905.2
137 Rhodobacter sphaeroides ATCC 17025 GCA_000016405.1
126 Gemmobacter aquatilis GCA_900110025.1
yj
206 Streptococcus mutans UA159 GCA_000007465.2
141 Streptococcus minor DSM 17118 GCA_000377005.1
140 Streptococcus equinus GCA_000964315.1
140 Streptococcus gallolyticus subsp. gallolyticus DSM 16831 GCA_002000985.1
140 Streptococcus marimammalium DSM 18627 GCA_000380045.1
yk
215 Clostridium saccharobutylicum DSM 13864 GCA_000473995.1
213 Clostridium beijerinckii GCA_000833105.2
213 Clostridium puniceum GCA_002006345.1
213 Clostridium saccharoperbutylacetonicum N1-4_28HMT_29 GCA_000340885.1
204 Clostridium chromiireducens GCA_002029255.1
204 Clostridium taeniosporum GCA_001735765.1
yl
249 Enterococcus faecalis V583 GCA_000007785.1
249 Streptomyces cinnamoneus GCA_001885705.1
216 Enterococcus canis NBRC 100695 GCA_001544375.1
216 Enterococcus faecium DO GCA_000174395.2
216 Enterococcus haemoperoxidus ATCC BAA-382 GCA_000407165.1
216 Enterococcus hirae ATCC 9790 GCA_000271405.2
216 Enterococcus mundtii QU 25 GCA_000504125.1
216 Enterococcus phoeniculicola ATCC BAA-412 GCA_000407505.1
216 Enterococcus rivorum GCA_001742285.1
216 Enterococcus thailandicus GCA_001652875.1
215 Enterococcus asini ATCC 700915 GCA_000407365.1
215 Enterococcus dispar ATCC 51266 GCA_000406945.1
ym
128 Pseudorhodobacter ferrugineus DSM 5888 GCA_000420745.1
128 Pseudorhodobacter psychrotolerans GCA_001294535.1
128 Pseudorhodobacter wandonensis GCA_001202035.1
128 Rhodobacter sphaeroides 2.4.1 GCA_000012905.2
128 Rhodobacter sphaeroides ATCC 17025 GCA_000016405.1
115 Sphingomonas koreensis GCA_001922385.1
114 Sphingomonas echinoides ATCC 14820 GCA_000241465.1
114 Sphingomonas hengshuiensis GCA_000935025.1
114 Sphingomonas mali NBRC 15500 GCA_001598415.1
114 Sphingomonas mucosissima GCA_002197665.1
114 Sphingomonas panacis GCA_001717955.1
114 Sphingomonas pruni NBRC 15498 GCA_001598455.1
114 Sphingomonas soli NBRC 100801 GCA_001591025.1
yn
188 Escherichia coli IAI39 GCA_000026345.1
188 Escherichia coli O104_3AH4 str. 2011C-3493 GCA_000299455.1
188 Escherichia coli O157_3AH7 str. Sakai GCA_000008865.1
188 Escherichia coli O83_3AH1 str. NRG 857C GCA_000183345.1
188 Escherichia coli UMN026 GCA_000026325.2
188 Escherichia coli str. K-12 substr. MG1655 GCA_000005845.2
188 Shigella dysenteriae Sd197 GCA_000012005.1
188 Shigella flexneri 2a str. 301 GCA_000006925.2
188 Tumebacillus flagellatus GCA_000714935.1
182 Erwinia iniecta GCA_001267535.1
162 Cedecea neteri GCA_000757825.1
162 Cronobacter sakazakii GCA_000982825.1
162 Enterobacter cloacae subsp. cloacae ATCC 13047 GCA_000025565.1
162 Enterobacter hormaechei subsp. steigerwaltii GCA_001729725.1
162 Enterobacter kobei GCA_900185885.1
162 Erwinia persicina NBRC 102418 GCA_001571305.1
162 Erwinia toletana DAPP-PG 735 GCA_000336255.1
162 Klebsiella oxytoca GCA_001022195.1
162 Kosakonia cowanii GCA_001975225.1
162 Kosakonia sacchari SP1 GCA_000300455.4
162 Pluralibacter gergoviae GCA_000757785.1
162 Pseudescherichia vulneris NBRC 102420 GCA_000759795.1
162 Rosenbergiella nectarea GCA_900111105.1
162 Salmonella enterica subsp. enterica serovar Typhi str. CT18 GCA_000195995.1
162 Salmonella enterica subsp. enterica serovar Typhimurium str. LT2 GCA_000006945.2
yo
258 Clostridium beijerinckii GCA_000833105.2
258 Clostridium puniceum GCA_002006345.1
258 Clostridium saccharoperbutylacetonicum N1-4_28HMT_29 GCA_000340885.1

250 Clostridium saccharobutylicum DSM 13864 GCA_000473995.1
239 Clostridium chromiireducens GCA_002029255.1
yp
201 Clostridium beijerinckii GCA_000833105.2
201 Clostridium saccharobutylicum DSM 13864 GCA_000473995.1
201 Clostridium saccharoperbutylacetonicum N1-4_28HMT_29 GCA_000340885.1
190 Clostridium butyricum GCA_001456065.2
186 Clostridium botulinum B str. Eklund 17B _28NRP_29 GCA_000020165.1
186 Clostridium chromiireducens GCA_002029255.1
186 Clostridium puniceum GCA_002006345.1
186 Clostridium taeniosporum GCA_001735765.1
yq
154 Escherichia coli IAI39 GCA_000026345.1
154 Escherichia coli O104_3AH4 str. 2011C-3493 GCA_000299455.1
154 Escherichia coli O157_3AH7 str. Sakai GCA_000008865.1
154 Escherichia coli UMN026 GCA_000026325.2
154 Escherichia coli str. K-12 substr. MG1655 GCA_000005845.2
154 Shigella dysenteriae Sd197 GCA_000012005.1
154 Shigella flexneri 2a str. 301 GCA_000006925.2
154 Tumebacillus flagellatus GCA_000714935.1
153 Escherichia coli O83_3AH1 str. NRG 857C GCA_000183345.1
142 Erwinia iniecta GCA_001267535.1
yr
212 Clostridium beijerinckii GCA_000833105.2
212 Clostridium puniceum GCA_002006345.1
212 Clostridium saccharobutylicum DSM 13864 GCA_000473995.1
212 Clostridium saccharoperbutylacetonicum N1-4_28HMT_29 GCA_000340885.1
204 Clostridium chromiireducens GCA_002029255.1
190 Clostridium neonatale GCA_001458595.1
ys
241 Escherichia coli IAI39 GCA_000026345.1
241 Escherichia coli O104_3AH4 str. 2011C-3493 GCA_000299455.1
241 Escherichia coli O157_3AH7 str. Sakai GCA_000008865.1
241 Escherichia coli O83_3AH1 str. NRG 857C GCA_000183345.1
241 Escherichia coli UMN026 GCA_000026325.2
241 Escherichia coli str. K-12 substr. MG1655 GCA_000005845.2
241 Shigella dysenteriae Sd197 GCA_000012005.1
241 Shigella flexneri 2a str. 301 GCA_000006925.2
241 Tumebacillus flagellatus GCA_000714935.1
204 Cronobacter sakazakii GCA_000982825.1
204 Enterobacter cloacae subsp. cloacae ATCC 13047 GCA_000025565.1
204 Enterobacter hormaechei subsp. steigerwaltii GCA_001729725.1
204 Enterobacter kobei GCA_900185885.1
204 Erwinia iniecta GCA_001267535.1
204 Erwinia persicina NBRC 102418 GCA_001571305.1
204 Erwinia toletana DAPP-PG 735 GCA_000336255.1
204 Klebsiella oxytoca GCA_001022195.1
204 Kosakonia cowanii GCA_001975225.1
204 Kosakonia sacchari SP1 GCA_000300455.4
204 Pluralibacter gergoviae GCA_000757785.1
204 Pseudescherichia vulneris NBRC 102420 GCA_000759795.1
204 Salmonella enterica subsp. enterica serovar Typhi str. CT18 GCA_000195995.1
201 Salmonella enterica subsp. enterica serovar Typhimurium str. LT2 GCA_000006945.2
yt
220 Clostridium beijerinckii GCA_000833105.2
220 Clostridium puniceum GCA_002006345.1
220 Clostridium saccharobutylicum DSM 13864 GCA_000473995.1
220 Clostridium saccharoperbutylacetonicum N1-4_28HMT_29 GCA_000340885.1
195 Clostridium chromiireducens GCA_002029255.1
185 Clostridium neonatale GCA_001458595.1
yu
238 Bacillus thuringiensis YBT-1518 GCA_000497525.2
222 Bacillus anthracis str. Ames GCA_000007845.1
222 Bacillus anthracis str. Sterne GCA_000008165.1
222 Bacillus cereus ATCC 14579 GCA_000007825.1
222 Bacillus mycoides GCA_000832605.1
222 Bacillus pseudomycoides DSM 12442 GCA_000161455.1
222 _5BBacillus thuringiensis_5D serovar konkukian str. 97-27 GCA_000008505.1
175 Massilibacterium senegalense GCA_001375675.1
yv
305 Deinococcus radiodurans R1 GCA_000008565.1
188 Deinococcus gobiensis I-0 GCA_000252445.1
171 Deinococcus marmoris DSM 12784 GCA_000701405.1
171 Deinococcus swuensis GCA_000800395.1
yw
245 Deinococcus radiodurans R1 GCA_000008565.1

161 Deinococcus puniceus GCA_001644565.1
159 Deinococcus gobiensis I-0 GCA_000252445.1
yx
88 Clostridium beijerinckii GCA_000833105.2
88 Clostridium puniceum GCA_002006345.1
88 Clostridium saccharobutylicum DSM 13864 GCA_000473995.1
88 Clostridium saccharoperbutylacetonicum N1-4_28HMT_29 GCA_000340885.1
77 Clostridium butyricum GCA_001456065.2
77 Clostridium chromiireducens GCA_002029255.1
76 Clostridium taeniosporum GCA_001735765.1
yy
313 Bacillus thuringiensis YBT-1518 GCA_000497525.2
300 Bacillus anthracis str. Ames GCA_000007845.1
300 Bacillus anthracis str. Sterne GCA_000008165.1
300 Bacillus cereus ATCC 14579 GCA_000007825.1
300 Bacillus mycoides GCA_000832605.1
300 Bacillus pseudomycoides DSM 12442 GCA_000161455.1
300 _5BBacillus thuringiensis_5D serovar konkukian str. 97-27 GCA_000008505.1
248 Bacillus cytotoxicus NVH 391-98 GCA_000017425.1
yz
274 Streptococcus mutans UA159 GCA_000007465.2
223 Streptococcus sobrinus DSM 20742 = ATCC 33478 GCA_000686605.1
222 Streptococcus ratti FA-1 = DSM 20564 GCA_000286075.1
yA
225 Bacillus mycoides GCA_000832605.1
220 Bacillus thuringiensis YBT-1518 GCA_000497525.2
214 Bacillus anthracis str. Ames GCA_000007845.1
214 Bacillus anthracis str. Sterne GCA_000008165.1
214 Bacillus cereus ATCC 14579 GCA_000007825.1
214 Bacillus pseudomycoides DSM 12442 GCA_000161455.1
214 _5BBacillus thuringiensis_5D serovar konkukian str. 97-27 GCA_000008505.1
yB
269 Enterococcus faecalis V583 GCA_000007785.1
269 Streptomyces cinnamoneus GCA_001885705.1
236 Enterococcus asini ATCC 700915 GCA_000407365.1
236 Enterococcus canis NBRC 100695 GCA_001544375.1
236 Enterococcus dispar ATCC 51266 GCA_000406945.1
236 Enterococcus faecium DO GCA_000174395.2
236 Enterococcus gilvus ATCC BAA-350 GCA_000407545.1
236 Enterococcus hermanniensis GCA_001885945.1
236 Enterococcus hirae ATCC 9790 GCA_000271405.2
236 Enterococcus italicus DSM 15952 GCA_000185365.1
236 Enterococcus malodoratus ATCC 43197 GCA_000407185.1
236 Enterococcus massiliensis GCA_001050095.1
236 Enterococcus mundtii QU 25 GCA_000504125.1
236 Enterococcus pallens ATCC BAA-351 GCA_000407485.1
236 Enterococcus pseudoavium NBRC 100491 GCA_001544295.1
236 Enterococcus rivorum GCA_001742285.1
230 Melissococcus plutonius S1 GCA_000747585.1
yC
205 Escherichia coli IAI39 GCA_000026345.1
205 Escherichia coli O104_3AH4 str. 2011C-3493 GCA_000299455.1
205 Escherichia coli O157_3AH7 str. Sakai GCA_000008865.1
205 Escherichia coli UMN026 GCA_000026325.2
205 Escherichia coli str. K-12 substr. MG1655 GCA_000005845.2
205 Shigella flexneri 2a str. 301 GCA_000006925.2
205 Tumebacillus flagellatus GCA_000714935.1
196 Shigella dysenteriae Sd197 GCA_000012005.1
189 Escherichia coli O83_3AH1 str. NRG 857C GCA_000183345.1
yD
262 Rhodobacter sphaeroides 2.4.1 GCA_000012905.2
229 Gemmobacter megaterium GCA_900156815.1
229 Rhodobacter sphaeroides ATCC 17025 GCA_000016405.1
218 Pseudorhodobacter psychrotolerans GCA_001294535.1
yE
338 Lactobacillus gasseri ATCC 33323 = JCM 1131 GCA_000014425.1
324 Lactobacillus hominis DSM 23910 = CRBIP 24.179 GCA_000296835.1
238 Lactobacillus amylovorus GCA_000191545.1
238 Lactobacillus crispatus ST1 GCA_000091765.1
yF
251 Clostridium beijerinckii GCA_000833105.2
251 Clostridium puniceum GCA_002006345.1
251 Clostridium saccharobutylicum DSM 13864 GCA_000473995.1
251 Clostridium saccharoperbutylacetonicum N1-4_28HMT_29 GCA_000340885.1
233 Clostridium neonatale GCA_001458595.1
232 Clostridium chromiireducens GCA_002029255.1

yG
185 Bacillus thuringiensis YBT-1518 GCA_000497525.2
177 Bacillus anthracis str. Ames GCA_000007845.1
177 Bacillus anthracis str. Sterne GCA_000008165.1
177 Bacillus cereus ATCC 14579 GCA_000007825.1
177 Bacillus mycoides GCA_000832605.1
177 Bacillus pseudomycoides DSM 12442 GCA_000161455.1
177 _5BBacillus thuringiensis_5D serovar konkukian str. 97-27 GCA_000008505.1
150 Oceanobacillus caeni GCA_001298135.1
yH
296 Bacillus anthracis str. Ames GCA_000007845.1
296 Bacillus anthracis str. Sterne GCA_000008165.1
296 Bacillus cereus ATCC 14579 GCA_000007825.1
296 Bacillus mycoides GCA_000832605.1
296 Bacillus pseudomycoides DSM 12442 GCA_000161455.1
296 Bacillus thuringiensis YBT-1518 GCA_000497525.2
296 _5BBacillus thuringiensis_5D serovar konkukian str. 97-27 GCA_000008505.1
242 Bacillus infantis NRRL B-14911 GCA_000473245.1
226 Bacillus pseudofirmus OF4 GCA_000005825.2
yI
194 Staphylococcus condimenti GCA_001618885.1
194 Staphylococcus simulans GCA_001559115.1
191 Staphylococcus aureus subsp. aureus NCTC 8325 GCA_000013425.1
191 Staphylococcus capitis subsp. capitis GCA_001028645.1
191 Staphylococcus epidermidis ATCC 12228 GCA_000007645.1
191 Staphylococcus haemolyticus JCSC1435 GCA_000009865.1
191 Staphylococcus hominis subsp. hominis C80 GCA_000183685.1
191 Staphylococcus lugdunensis HKU09-01 GCA_000025085.1
191 Staphylococcus simiae CCM 7213 GCA_000235645.2
187 Staphylococcus arlettae CVD059 GCA_000295715.1
187 Staphylococcus cohnii subsp. cohnii GCA_000972575.1
187 Staphylococcus gallinarum GCA_000875895.1
187 Staphylococcus saprophyticus subsp. saprophyticus ATCC 15305 GCA_000010125.1
187 Staphylococcus succinus GCA_001902315.1
187 Staphylococcus xylosus GCA_000706685.1
yJ
225 Enterococcus faecalis V583 GCA_000007785.1
225 Streptomyces cinnamoneus GCA_001885705.1
214 Enterococcus columbae DSM 7374 = ATCC 51263 GCA_000406925.1
207 Enterococcus asini ATCC 700915 GCA_000407365.1
207 Enterococcus canis NBRC 100695 GCA_001544375.1
207 Enterococcus dispar ATCC 51266 GCA_000406945.1
207 Enterococcus faecium DO GCA_000174395.2
207 Enterococcus hirae ATCC 9790 GCA_000271405.2
207 Enterococcus massiliensis GCA_001050095.1
207 Enterococcus mundtii QU 25 GCA_000504125.1
207 Enterococcus rivorum GCA_001742285.1
yK
185 Bifidobacterium adolescentis ATCC 15703 GCA_000010425.1
175 Bifidobacterium dentium JCM 1195 = DSM 20436 GCA_001042595.1
175 Bifidobacterium tsurumiense GCA_000741765.1
170 Bifidobacterium thermophilum GCA_000741495.1
yL
255 Lactobacillus gasseri ATCC 33323 = JCM 1131 GCA_000014425.1
255 Lactobacillus hominis DSM 23910 = CRBIP 24.179 GCA_000296835.1
215 Lactobacillus iners DSM 13335 GCA_000160875.1
215 Lactobacillus psittaci DSM 15354 GCA_000425905.1
206 Lactobacillus harbinensis DSM 16991 GCA_000425885.1
206 Lactobacillus perolens DSM 12744 GCA_001435585.1
206 Lactobacillus shenzhenensis LY-73 GCA_000469325.1
yM
160 Clostridium beijerinckii GCA_000833105.2
160 Clostridium saccharobutylicum DSM 13864 GCA_000473995.1
160 Clostridium saccharoperbutylacetonicum N1-4_28HMT_29 GCA_000340885.1
146 Clostridium chromiireducens GCA_002029255.1
146 Clostridium puniceum GCA_002006345.1
134 Clostridium butyricum GCA_001456065.2
yN
255 Streptococcus mutans UA159 GCA_000007465.2
182 Streptococcus gordonii str. Challis substr. CH1 GCA_000017005.1
178 Streptococcus cristatus AS 1.3089 GCA_000385925.1
178 Streptococcus mitis B6 GCA_000027165.1
178 Streptococcus pneumoniae R6 GCA_000007045.1
yO
171 Deinococcus radiodurans R1 GCA_000008565.1
121 Deinococcus deserti VCD115 GCA_000020685.1

118 Deinococcus gobiensis I-0 GCA_000252445.1
yP
257 Lactobacillus gasseri ATCC 33323 = JCM 1131 GCA_000014425.1
233 Lactobacillus hominis DSM 23910 = CRBIP 24.179 GCA_000296835.1
149 Lactobacillus acidophilus NCFM GCA_000011985.1
149 Lactobacillus crispatus ST1 GCA_000091765.1
149 Lactobacillus hamsteri DSM 5661 = JCM 6256 GCA_000615445.1
149 Lactobacillus psittaci DSM 15354 GCA_000425905.1
yQ
226 Clostridium beijerinckii GCA_000833105.2
226 Clostridium puniceum GCA_002006345.1
226 Clostridium saccharoperbutylacetonicum N1-4_28HMT_29 GCA_000340885.1
218 Clostridium saccharobutylicum DSM 13864 GCA_000473995.1
200 Clostridium chromiireducens GCA_002029255.1
yR
251 Bacillus cytotoxicus NVH 391-98 GCA_000017425.1
247 Bacillus thuringiensis YBT-1518 GCA_000497525.2
246 Bacillus flexus GCA_002024265.1
246 Bacillus megaterium NBRC 15308 = ATCC 14581 GCA_000832985.1
yS
199 Deinococcus radiodurans R1 GCA_000008565.1
134 Deinococcus gobiensis I-0 GCA_000252445.1
104 Deinococcus puniceus GCA_001644565.1
yT
217 Escherichia coli IAI39 GCA_000026345.1
217 Escherichia coli O104_3AH4 str. 2011C-3493 GCA_000299455.1
217 Escherichia coli O157_3AH7 str. Sakai GCA_000008865.1
217 Escherichia coli O83_3AH1 str. NRG 857C GCA_000183345.1
217 Escherichia coli UMN026 GCA_000026325.2
217 Escherichia coli str. K-12 substr. MG1655 GCA_000005845.2
217 Shigella flexneri 2a str. 301 GCA_000006925.2
217 Tumebacillus flagellatus GCA_000714935.1
210 Erwinia iniecta GCA_001267535.1
202 Shigella dysenteriae Sd197 GCA_000012005.1
yU
120 Bifidobacterium adolescentis ATCC 15703 GCA_000010425.1
105 Bifidobacterium asteroides PRL2011 GCA_000304215.1
105 Bifidobacterium callitrichos DSM 23973 GCA_000741175.1
102 Bifidobacterium thermophilum GCA_000741495.1
yV
271 Bacillus anthracis str. Ames GCA_000007845.1
271 Bacillus anthracis str. Sterne GCA_000008165.1
271 Bacillus cereus ATCC 14579 GCA_000007825.1
271 Bacillus mycoides GCA_000832605.1
271 Bacillus pseudomycoides DSM 12442 GCA_000161455.1
271 Bacillus thuringiensis YBT-1518 GCA_000497525.2
271 _5BBacillus thuringiensis_5D serovar konkukian str. 97-27 GCA_000008505.1
199 Bacillus cytotoxicus NVH 391-98 GCA_000017425.1
190 Bacillus manliponensis GCA_000712595.1
yW
194 Bifidobacterium adolescentis ATCC 15703 GCA_000010425.1
190 Bifidobacterium tsurumiense GCA_000741765.1
175 Bifidobacterium dentium JCM 1195 = DSM 20436 GCA_001042595.1
yX
205 Deinococcus radiodurans R1 GCA_000008565.1
125 Deinococcus gobiensis I-0 GCA_000252445.1
108 Deinococcus deserti VCD115 GCA_000020685.1
108 Deinococcus hopiensis KR-140 GCA_900176165.1
108 Deinococcus misasensis DSM 22328 GCA_000745915.1
108 Deinococcus puniceus GCA_001644565.1
yY
234 Bacillus anthracis str. Ames GCA_000007845.1
234 Bacillus anthracis str. Sterne GCA_000008165.1
234 Bacillus cereus ATCC 14579 GCA_000007825.1
234 Bacillus mycoides GCA_000832605.1
234 Bacillus pseudomycoides DSM 12442 GCA_000161455.1
234 Bacillus thuringiensis YBT-1518 GCA_000497525.2
234 _5BBacillus thuringiensis_5D serovar konkukian str. 97-27 GCA_000008505.1
171 Bacillus horneckiae GCA_001636335.1
171 Bacillus solani GCA_001420595.1
167 Bacillus hemicellulosilyticus JCM 9152 GCA_000513115.1
yZ
198 Deinococcus radiodurans R1 GCA_000008565.1
170 Deinococcus deserti VCD115 GCA_000020685.1
170 Deinococcus soli Cha et al. 2016 GCA_001007995.1
165 Deinococcus hopiensis KR-140 GCA_900176165.1

165 Deinococcus puniceus GCA_001644565.1
z0
180 Streptococcus equinus GCA_000964315.1
165 Streptococcus gallolyticus subsp. gallolyticus DSM 16831 GCA_002000985.1
165 Streptococcus mutans UA159 GCA_000007465.2
158 Streptococcus cristatus AS 1.3089 GCA_000385925.1
158 Streptococcus gordonii str. Challis substr. CH1 GCA_000017005.1
158 Streptococcus mitis B6 GCA_000027165.1
158 Streptococcus pneumoniae R6 GCA_000007045.1
z1
334 Staphylococcus aureus subsp. aureus NCTC 8325 GCA_000013425.1
334 Staphylococcus epidermidis ATCC 12228 GCA_000007645.1
334 Staphylococcus haemolyticus JCSC1435 GCA_000009865.1
334 Staphylococcus hominis subsp. hominis C80 GCA_000183685.1
334 Staphylococcus lugdunensis HKU09-01 GCA_000025085.1
334 Staphylococcus simiae CCM 7213 GCA_000235645.2
326 Staphylococcus condimenti GCA_001618885.1
326 Staphylococcus simulans GCA_001559115.1
322 Staphylococcus capitis subsp. capitis GCA_001028645.1
z2
157 Deinococcus radiodurans R1 GCA_000008565.1
102 Deinococcus deserti VCD115 GCA_000020685.1
95 Deinococcus soli Cha et al. 2016 GCA_001007995.1
z3
117 Clostridium butyricum GCA_001456065.2
111 Clostridium beijerinckii GCA_000833105.2
111 Clostridium puniceum GCA_002006345.1
111 Clostridium saccharoperbutylacetonicum N1-4_28HMT_29 GCA_000340885.1
105 Clostridium taeniosporum GCA_001735765.1
z4
248 Streptococcus mutans UA159 GCA_000007465.2
170 Streptococcus macacae NCTC 11558 GCA_000187995.3
164 Streptococcus gordonii str. Challis substr. CH1 GCA_000017005.1
z5
240 Bacillus anthracis str. Ames GCA_000007845.1
240 Bacillus anthracis str. Sterne GCA_000008165.1
240 Bacillus cereus ATCC 14579 GCA_000007825.1
240 Bacillus mycoides GCA_000832605.1
240 Bacillus pseudomycoides DSM 12442 GCA_000161455.1
240 Bacillus thuringiensis YBT-1518 GCA_000497525.2
240 _5BBacillus thuringiensis_5D serovar konkukian str. 97-27 GCA_000008505.1
198 Bacillus hemicellulosilyticus JCM 9152 GCA_000513115.1
193 Bacillus vietnamensis NBRC 101237 GCA_001591825.1
z6
206 Escherichia coli IAI39 GCA_000026345.1
206 Escherichia coli O104_3AH4 str. 2011C-3493 GCA_000299455.1
206 Escherichia coli O157_3AH7 str. Sakai GCA_000008865.1
206 Escherichia coli O83_3AH1 str. NRG 857C GCA_000183345.1
206 Escherichia coli UMN026 GCA_000026325.2
206 Escherichia coli str. K-12 substr. MG1655 GCA_000005845.2
206 Shigella flexneri 2a str. 301 GCA_000006925.2
206 Tumebacillus flagellatus GCA_000714935.1
197 Shigella dysenteriae Sd197 GCA_000012005.1
156 Cronobacter sakazakii GCA_000982825.1
156 Erwinia iniecta GCA_001267535.1
156 Erwinia toletana DAPP-PG 735 GCA_000336255.1
156 Kosakonia cowanii GCA_001975225.1
156 Kosakonia sacchari SP1 GCA_000300455.4
156 Mangrovibacter phragmitis GCA_001655675.1
156 Pluralibacter gergoviae GCA_000757785.1
156 Serratia marcescens subsp. marcescens Db11 GCA_000513215.1
156 _5BEnterobacter_5D lignolyticus SCF1 GCA_000164865.1
z7
178 Klebsiella pneumoniae subsp. pneumoniae HS11286 GCA_000240185.2
173 Escherichia coli IAI39 GCA_000026345.1
173 Escherichia coli O104_3AH4 str. 2011C-3493 GCA_000299455.1
173 Escherichia coli O157_3AH7 str. Sakai GCA_000008865.1
173 Escherichia coli O83_3AH1 str. NRG 857C GCA_000183345.1
173 Escherichia coli UMN026 GCA_000026325.2
173 Escherichia coli str. K-12 substr. MG1655 GCA_000005845.2
173 Shigella dysenteriae Sd197 GCA_000012005.1
173 Shigella flexneri 2a str. 301 GCA_000006925.2
173 Tumebacillus flagellatus GCA_000714935.1
163 Serratia marcescens subsp. marcescens Db11 GCA_000513215.1
z8
230 Lactobacillus hominis DSM 23910 = CRBIP 24.179 GCA_000296835.1

221 Lactobacillus gasseri ATCC 33323 = JCM 1131 GCA_000014425.1
195 Lactobacillus kefiranofaciens ZW3 GCA_000214785.1
z9
283 Clostridium beijerinckii GCA_000833105.2
283 Clostridium puniceum GCA_002006345.1
283 Clostridium saccharoperbutylacetonicum N1-4_28HMT_29 GCA_000340885.1
275 Clostridium saccharobutylicum DSM 13864 GCA_000473995.1
252 Clostridium chromiireducens GCA_002029255.1
za
165 Clostridium beijerinckii GCA_000833105.2
165 Clostridium saccharoperbutylacetonicum N1-4_28HMT_29 GCA_000340885.1
156 Clostridium saccharobutylicum DSM 13864 GCA_000473995.1
150 Clostridium puniceum GCA_002006345.1
150 Clostridium taeniosporum GCA_001735765.1
zb
311 Deinococcus radiodurans R1 GCA_000008565.1
193 Deinococcus marmoris DSM 12784 GCA_000701405.1
193 Deinococcus swuensis GCA_000800395.1
188 Deinococcus gobiensis I-0 GCA_000252445.1
zc
147 Clostridium beijerinckii GCA_000833105.2
147 Clostridium puniceum GCA_002006345.1
147 Clostridium saccharobutylicum DSM 13864 GCA_000473995.1
147 Clostridium saccharoperbutylacetonicum N1-4_28HMT_29 GCA_000340885.1
135 Clostridium chromiireducens GCA_002029255.1
132 Clostridium intestinale URNW GCA_000469625.2
zd
239 Bacillus anthracis str. Ames GCA_000007845.1
239 Bacillus anthracis str. Sterne GCA_000008165.1
239 Bacillus cereus ATCC 14579 GCA_000007825.1
239 Bacillus mycoides GCA_000832605.1
239 Bacillus pseudomycoides DSM 12442 GCA_000161455.1
239 Bacillus thuringiensis YBT-1518 GCA_000497525.2
239 _5BBacillus thuringiensis_5D serovar konkukian str. 97-27 GCA_000008505.1
201 Bacillus coahuilensis m4-4 GCA_000171615.1
195 Bacillus horneckiae GCA_001636335.1
195 Bacillus solani GCA_001420595.1
ze
296 Clostridium beijerinckii GCA_000833105.2
296 Clostridium puniceum GCA_002006345.1
296 Clostridium saccharoperbutylacetonicum N1-4_28HMT_29 GCA_000340885.1
287 Clostridium saccharobutylicum DSM 13864 GCA_000473995.1
272 Clostridium chromiireducens GCA_002029255.1
zf
69 Bifidobacterium adolescentis ATCC 15703 GCA_000010425.1
62 Arthrobacter woluwensis GCA_900105345.1
50 Nocardia mikamii NBRC 108933 GCA_001613505.1
50 Nocardia uniformis NBRC 13702 GCA_001613345.1
50 Nocardia vermiculata NBRC 100427 GCA_001613265.1
50 Nocardia veterana NBRC 100344 GCA_000308855.1
50 Nocardia vinacea NBRC 16497 GCA_000308835.1
50 Saccharomonospora saliphila YIM 90502 GCA_000383795.1
zg
194 Rhodobacter sphaeroides 2.4.1 GCA_000012905.2
177 Celeribacter neptunius GCA_900113955.1
177 Defluviimonas alba GCA_001620265.1
177 Haematobacter massiliensis GCA_000740795.1
177 Loktanella koreensis GCA_900109295.1
177 Oceanicola granulosus HTCC2516 GCA_000153305.1
177 Planktotalea frisia GCA_001890925.1
177 Pseudoruegeria sabulilitoris GCA_001558155.1
174 Octadecabacter arcticus 238 GCA_000155735.2
174 Thalassobius gelatinovorus GCA_001458355.1
zh
193 Deinococcus radiodurans R1 GCA_000008565.1
149 Deinococcus gobiensis I-0 GCA_000252445.1
123 Deinococcus marmoris DSM 12784 GCA_000701405.1
123 Deinococcus swuensis GCA_000800395.1
zi
158 Aerococcus urinaehominis GCA_001543245.1
158 Jeotgalicoccus marinus DSM 19772 GCA_000425825.1
158 Salinicoccus qingdaonensis GCA_900101075.1
156 Lactobacillus gasseri ATCC 33323 = JCM 1131 GCA_000014425.1
153 Jeotgalicoccus halophilus GCA_900101065.1
zj
200 Escherichia coli IAI39 GCA_000026345.1

200 Escherichia coli O104_3AH4 str. 2011C-3493 GCA_000299455.1
200 Escherichia coli O157_3AH7 str. Sakai GCA_000008865.1
200 Escherichia coli O83_3AH1 str. NRG 857C GCA_000183345.1
200 Escherichia coli UMN026 GCA_000026325.2
200 Escherichia coli str. K-12 substr. MG1655 GCA_000005845.2
200 Shigella dysenteriae Sd197 GCA_000012005.1
200 Tumebacillus flagellatus GCA_000714935.1
191 Shigella flexneri 2a str. 301 GCA_000006925.2
176 Erwinia iniecta GCA_001267535.1
zk
122 Deinococcus radiodurans R1 GCA_000008565.1
69 Deinococcus deserti VCD115 GCA_000020685.1
69 Deinococcus gobiensis I-0 GCA_000252445.1
54 Deinococcus geothermalis DSM 11300 GCA_000196275.1
zl
281 Lactobacillus gasseri ATCC 33323 = JCM 1131 GCA_000014425.1
273 Lactobacillus hominis DSM 23910 = CRBIP 24.179 GCA_000296835.1
237 Lactobacillus crispatus ST1 GCA_000091765.1
zm
193 Staphylococcus simulans GCA_001559115.1
189 Staphylococcus saprophyticus GCA_001074355.1
185 Megasphaera cerevisiae DSM 20462 GCA_001045675.1
185 Staphylococcus aureus subsp. aureus NCTC 8325 GCA_000013425.1
185 Staphylococcus cohnii subsp. cohnii GCA_000972575.1
185 Staphylococcus epidermidis ATCC 12228 GCA_000007645.1
185 Staphylococcus haemolyticus JCSC1435 GCA_000009865.1
185 Staphylococcus hominis subsp. hominis C80 GCA_000183685.1
185 Staphylococcus simiae CCM 7213 GCA_000235645.2
185 Staphylococcus warneri SG1 GCA_000332735.1
zn
165 Bifidobacterium dentium JCM 1195 = DSM 20436 GCA_001042595.1
160 Bifidobacterium adolescentis ATCC 15703 GCA_000010425.1
160 Bifidobacterium breve DSM 20213 = JCM 1192 GCA_001025175.1
160 Bifidobacterium longum NCC2705 GCA_000007525.1
160 Bifidobacterium saguini DSM 23967 GCA_000741715.1
157 Bifidobacterium angulatum DSM 20098 = JCM 7096 GCA_001025155.1
157 Bifidobacterium tsurumiense GCA_000741765.1
zo
191 Bacillus anthracis str. Ames GCA_000007845.1
191 Bacillus anthracis str. Sterne GCA_000008165.1
191 Bacillus cereus ATCC 14579 GCA_000007825.1
191 Bacillus mycoides GCA_000832605.1
191 Bacillus pseudomycoides DSM 12442 GCA_000161455.1
191 Bacillus thuringiensis YBT-1518 GCA_000497525.2
191 _5BBacillus thuringiensis_5D serovar konkukian str. 97-27 GCA_000008505.1
171 Bacillus manliponensis GCA_000712595.1
158 Paenirhodobacter enshiensis GCA_000740785.1
zp
302 Bacillus anthracis str. Ames GCA_000007845.1
302 Bacillus anthracis str. Sterne GCA_000008165.1
302 Bacillus cereus ATCC 14579 GCA_000007825.1
302 Bacillus mycoides GCA_000832605.1
302 Bacillus pseudomycoides DSM 12442 GCA_000161455.1
302 Bacillus thuringiensis YBT-1518 GCA_000497525.2
302 _5BBacillus thuringiensis_5D serovar konkukian str. 97-27 GCA_000008505.1
239 Bacillus manliponensis GCA_000712595.1
233 Bacillus marisflavi GCA_001274775.1
zq
276 Escherichia coli IAI39 GCA_000026345.1
276 Escherichia coli O104_3AH4 str. 2011C-3493 GCA_000299455.1
276 Escherichia coli O157_3AH7 str. Sakai GCA_000008865.1
276 Escherichia coli O83_3AH1 str. NRG 857C GCA_000183345.1
276 Escherichia coli UMN026 GCA_000026325.2
276 Escherichia coli str. K-12 substr. MG1655 GCA_000005845.2
276 Shigella flexneri 2a str. 301 GCA_000006925.2
276 Tumebacillus flagellatus GCA_000714935.1
267 Shigella dysenteriae Sd197 GCA_000012005.1
237 Erwinia iniecta GCA_001267535.1
237 Erwinia toletana DAPP-PG 735 GCA_000336255.1
237 Kosakonia cowanii GCA_001975225.1
zr
233 Deinococcus radiodurans R1 GCA_000008565.1
131 Deinococcus marmoris DSM 12784 GCA_000701405.1
131 Deinococcus puniceus GCA_001644565.1
131 Deinococcus swuensis GCA_000800395.1
130 Deinococcus hopiensis KR-140 GCA_900176165.1

zs
225 Cronobacter sakazakii GCA_000982825.1
225 Erwinia iniecta GCA_001267535.1
225 Erwinia toletana DAPP-PG 735 GCA_000336255.1
225 Escherichia coli IAI39 GCA_000026345.1
225 Escherichia coli O104_3AH4 str. 2011C-3493 GCA_000299455.1
225 Escherichia coli O157_3AH7 str. Sakai GCA_000008865.1
225 Escherichia coli O83_3AH1 str. NRG 857C GCA_000183345.1
225 Escherichia coli UMN026 GCA_000026325.2
225 Escherichia coli str. K-12 substr. MG1655 GCA_000005845.2
225 Pantoea ananatis LMG 20103 GCA_000025405.2
225 Pantoea dispersa EGD-AAK13 GCA_000465555.2
225 Shigella flexneri 2a str. 301 GCA_000006925.2
225 Shimwellia blattae DSM 4481 = NBRC 105725 GCA_000262305.1
225 Tumebacillus flagellatus GCA_000714935.1
211 Edwardsiella anguillarum ET080813 GCA_000264765.2
211 Mangrovibacter phragmitis GCA_001655675.1
211 Rosenbergiella nectarea GCA_900111105.1
210 Erwinia amylovora CFBP1430 GCA_000091565.1
210 Kosakonia cowanii GCA_001975225.1
210 Kosakonia sacchari SP1 GCA_000300455.4
210 Pantoea agglomerans GCA_001709315.1
210 Plautia stali symbiont GCA_000180175.2
210 Shigella dysenteriae Sd197 GCA_000012005.1
zt
301 Escherichia coli IAI39 GCA_000026345.1
301 Escherichia coli O104_3AH4 str. 2011C-3493 GCA_000299455.1
301 Escherichia coli O157_3AH7 str. Sakai GCA_000008865.1
301 Escherichia coli UMN026 GCA_000026325.2
301 Escherichia coli str. K-12 substr. MG1655 GCA_000005845.2
301 Shigella dysenteriae Sd197 GCA_000012005.1
301 Shigella flexneri 2a str. 301 GCA_000006925.2
301 Tumebacillus flagellatus GCA_000714935.1
300 Escherichia coli O83_3AH1 str. NRG 857C GCA_000183345.1
295 Erwinia iniecta GCA_001267535.1
zu
186 Bifidobacterium adolescentis ATCC 15703 GCA_000010425.1
175 Bifidobacterium angulatum DSM 20098 = JCM 7096 GCA_001025155.1
174 Bifidobacterium subtile GCA_000741775.1
zv
187 Escherichia coli IAI39 GCA_000026345.1
187 Escherichia coli O104_3AH4 str. 2011C-3493 GCA_000299455.1
187 Escherichia coli O157_3AH7 str. Sakai GCA_000008865.1
187 Escherichia coli O83_3AH1 str. NRG 857C GCA_000183345.1
187 Escherichia coli UMN026 GCA_000026325.2
187 Escherichia coli str. K-12 substr. MG1655 GCA_000005845.2
187 Shigella flexneri 2a str. 301 GCA_000006925.2
187 Tumebacillus flagellatus GCA_000714935.1
178 Shigella dysenteriae Sd197 GCA_000012005.1
177 Serratia marcescens subsp. marcescens Db11 GCA_000513215.1
zw
234 Clostridium saccharobutylicum DSM 13864 GCA_000473995.1
229 Clostridium beijerinckii GCA_000833105.2
229 Clostridium saccharoperbutylacetonicum N1-4_28HMT_29 GCA_000340885.1
225 Clostridium butyricum GCA_001456065.2
225 Clostridium chromiireducens GCA_002029255.1
225 Clostridium puniceum GCA_002006345.1
zx
306 Escherichia coli IAI39 GCA_000026345.1
306 Escherichia coli O104_3AH4 str. 2011C-3493 GCA_000299455.1
306 Escherichia coli O157_3AH7 str. Sakai GCA_000008865.1
306 Escherichia coli O83_3AH1 str. NRG 857C GCA_000183345.1
306 Escherichia coli UMN026 GCA_000026325.2
306 Escherichia coli str. K-12 substr. MG1655 GCA_000005845.2
306 Shigella dysenteriae Sd197 GCA_000012005.1
306 Shigella flexneri 2a str. 301 GCA_000006925.2
306 Tumebacillus flagellatus GCA_000714935.1
275 Erwinia iniecta GCA_001267535.1
268 Rosenbergiella nectarea GCA_900111105.1
zy
337 Staphylococcus epidermidis ATCC 12228 GCA_000007645.1
337 Staphylococcus haemolyticus JCSC1435 GCA_000009865.1
337 Staphylococcus hominis subsp. hominis C80 GCA_000183685.1
332 Staphylococcus lugdunensis HKU09-01 GCA_000025085.1
329 Staphylococcus capitis subsp. capitis GCA_001028645.1
zz

221 Rhodobacter sphaeroides 2.4.1 GCA_000012905.2
194 Gemmobacter megaterium GCA_900156815.1
194 Rhodobacter sphaeroides ATCC 17025 GCA_000016405.1
184 Gemmobacter aquatilis GCA_900110025.1
184 Pseudorhodobacter psychrotolerans GCA_001294535.1
zA
267 Lactobacillus gasseri ATCC 33323 = JCM 1131 GCA_000014425.1
234 Lactobacillus hominis DSM 23910 = CRBIP 24.179 GCA_000296835.1
190 Lactobacillus crispatus ST1 GCA_000091765.1
zB
236 Rhodobacter sphaeroides 2.4.1 GCA_000012905.2
207 Rhodobacter sphaeroides ATCC 17025 GCA_000016405.1
193 Gemmobacter megaterium GCA_900156815.1
zC
236 Rhodobacter sphaeroides 2.4.1 GCA_000012905.2
232 Rhodobacter sphaeroides ATCC 17025 GCA_000016405.1
208 Gemmobacter megaterium GCA_900156815.1
zD
297 Bacillus anthracis str. Ames GCA_000007845.1
297 Bacillus anthracis str. Sterne GCA_000008165.1
297 Bacillus cereus ATCC 14579 GCA_000007825.1
297 Bacillus mycoides GCA_000832605.1
297 Bacillus pseudomycoides DSM 12442 GCA_000161455.1
297 Bacillus thuringiensis YBT-1518 GCA_000497525.2
297 _5BBacillus thuringiensis_5D serovar konkukian str. 97-27 GCA_000008505.1
239 Bacillus manliponensis GCA_000712595.1
229 Bacillus aquimaris TF-12 GCA_001648555.1
zE
282 Deinococcus radiodurans R1 GCA_000008565.1
181 Deinococcus gobiensis I-0 GCA_000252445.1
173 Deinococcus puniceus GCA_001644565.1
zF
238 Clostridium beijerinckii GCA_000833105.2
238 Clostridium puniceum GCA_002006345.1
238 Clostridium saccharoperbutylacetonicum N1-4_28HMT_29 GCA_000340885.1
223 Clostridium saccharobutylicum DSM 13864 GCA_000473995.1
209 Clostridium taeniosporum GCA_001735765.1
zG
172 Rhodobacter sphaeroides 2.4.1 GCA_000012905.2
167 Rhodobacter sphaeroides ATCC 17025 GCA_000016405.1
133 Gemmobacter aquatilis GCA_900110025.1
zH
168 Streptococcus mutans UA159 GCA_000007465.2
106 Streptococcus iniae GCA_000831485.1
97 Streptococcus ferus DSM 20646 GCA_000372425.1
zI
252 Enterococcus faecalis V583 GCA_000007785.1
252 Streptomyces cinnamoneus GCA_001885705.1
221 Enterococcus canis NBRC 100695 GCA_001544375.1
221 Enterococcus dispar ATCC 51266 GCA_000406945.1
221 Enterococcus faecium DO GCA_000174395.2
221 Enterococcus gilvus ATCC BAA-350 GCA_000407545.1
221 Enterococcus hermanniensis GCA_001885945.1
221 Enterococcus hirae ATCC 9790 GCA_000271405.2
221 Enterococcus malodoratus ATCC 43197 GCA_000407185.1
221 Enterococcus massiliensis GCA_001050095.1
221 Enterococcus mundtii QU 25 GCA_000504125.1
221 Enterococcus pallens ATCC BAA-351 GCA_000407485.1
221 Enterococcus pseudoavium NBRC 100491 GCA_001544295.1
221 Enterococcus rivorum GCA_001742285.1
209 Enterococcus aquimarinus GCA_001885765.1
209 Enterococcus haemoperoxidus ATCC BAA-382 GCA_000407165.1
209 Enterococcus phoeniculicola ATCC BAA-412 GCA_000407505.1
209 Enterococcus thailandicus GCA_001652875.1
zJ
242 Clostridium beijerinckii GCA_000833105.2
242 Clostridium puniceum GCA_002006345.1
242 Clostridium saccharobutylicum DSM 13864 GCA_000473995.1
242 Clostridium saccharoperbutylacetonicum N1-4_28HMT_29 GCA_000340885.1
231 Clostridium chromiireducens GCA_002029255.1
226 Clostridium butyricum GCA_001456065.2
zK
216 Bacillus thuringiensis YBT-1518 GCA_000497525.2
203 Bacillus anthracis str. Ames GCA_000007845.1
203 Bacillus anthracis str. Sterne GCA_000008165.1
203 Bacillus cereus ATCC 14579 GCA_000007825.1

203 Bacillus mycoides GCA_000832605.1
203 Bacillus pseudomycoides DSM 12442 GCA_000161455.1
203 _5BBacillus thuringiensis_5D serovar konkukian str. 97-27 GCA_000008505.1
182 Bacillus horneckiae GCA_001636335.1
zL
210 Escherichia coli IAI39 GCA_000026345.1
210 Escherichia coli O104_3AH4 str. 2011C-3493 GCA_000299455.1
210 Escherichia coli O157_3AH7 str. Sakai GCA_000008865.1
210 Escherichia coli O83_3AH1 str. NRG 857C GCA_000183345.1
210 Escherichia coli UMN026 GCA_000026325.2
210 Escherichia coli str. K-12 substr. MG1655 GCA_000005845.2
210 Shigella flexneri 2a str. 301 GCA_000006925.2
210 Tumebacillus flagellatus GCA_000714935.1
201 Shigella dysenteriae Sd197 GCA_000012005.1
196 Erwinia iniecta GCA_001267535.1
zM
157 Clostridium beijerinckii GCA_000833105.2
157 Clostridium puniceum GCA_002006345.1
157 Clostridium saccharoperbutylacetonicum N1-4_28HMT_29 GCA_000340885.1
142 Clostridium saccharobutylicum DSM 13864 GCA_000473995.1
140 Clostridium intestinale URNW GCA_000469625.2
zN
227 Clostridium beijerinckii GCA_000833105.2
227 Clostridium saccharobutylicum DSM 13864 GCA_000473995.1
227 Clostridium saccharoperbutylacetonicum N1-4_28HMT_29 GCA_000340885.1
223 Clostridium puniceum GCA_002006345.1
215 Clostridium chromiireducens GCA_002029255.1
zO
162 Bifidobacterium adolescentis ATCC 15703 GCA_000010425.1
147 Bifidobacterium angulatum DSM 20098 = JCM 7096 GCA_001025155.1
135 Bifidobacterium tsurumiense GCA_000741765.1
zP
205 Deinococcus radiodurans R1 GCA_000008565.1
156 Deinococcus hopiensis KR-140 GCA_900176165.1
156 Deinococcus puniceus GCA_001644565.1
154 Deinococcus gobiensis I-0 GCA_000252445.1
zQ
225 Bifidobacterium adolescentis ATCC 15703 GCA_000010425.1
201 Bifidobacterium gallicum DSM 20093 = LMG 11596 GCA_000741205.1
199 Bifidobacterium asteroides PRL2011 GCA_000304215.1
zR
270 Escherichia coli IAI39 GCA_000026345.1
270 Escherichia coli O104_3AH4 str. 2011C-3493 GCA_000299455.1
270 Escherichia coli O157_3AH7 str. Sakai GCA_000008865.1
270 Escherichia coli O83_3AH1 str. NRG 857C GCA_000183345.1
270 Escherichia coli UMN026 GCA_000026325.2
270 Escherichia coli str. K-12 substr. MG1655 GCA_000005845.2
270 Shigella flexneri 2a str. 301 GCA_000006925.2
270 Tumebacillus flagellatus GCA_000714935.1
255 Shigella dysenteriae Sd197 GCA_000012005.1
233 Erwinia iniecta GCA_001267535.1
zS
212 Deinococcus radiodurans R1 GCA_000008565.1
145 Deinococcus deserti VCD115 GCA_000020685.1
140 Deinococcus geothermalis DSM 11300 GCA_000196275.1
140 Deinococcus puniceus GCA_001644565.1
zT
291 Deinococcus radiodurans R1 GCA_000008565.1
178 Deinococcus puniceus GCA_001644565.1
170 Deinococcus gobiensis I-0 GCA_000252445.1
zU
195 Clostridium beijerinckii GCA_000833105.2
195 Clostridium saccharobutylicum DSM 13864 GCA_000473995.1
195 Clostridium saccharoperbutylacetonicum N1-4_28HMT_29 GCA_000340885.1
189 Clostridium puniceum GCA_002006345.1
174 Clostridium butyricum GCA_001456065.2
174 Clostridium chromiireducens GCA_002029255.1
zV
281 Deinococcus radiodurans R1 GCA_000008565.1
174 Deinococcus gobiensis I-0 GCA_000252445.1
159 Deinococcus deserti VCD115 GCA_000020685.1
zW
223 Escherichia coli IAI39 GCA_000026345.1
223 Escherichia coli O104_3AH4 str. 2011C-3493 GCA_000299455.1
223 Escherichia coli O157_3AH7 str. Sakai GCA_000008865.1
223 Escherichia coli O83_3AH1 str. NRG 857C GCA_000183345.1

223 Escherichia coli UMN026 GCA_000026325.2
223 Escherichia coli str. K-12 substr. MG1655 GCA_000005845.2
223 Shigella dysenteriae Sd197 GCA_000012005.1
223 Shigella flexneri 2a str. 301 GCA_000006925.2
223 Tumebacillus flagellatus GCA_000714935.1
216 Erwinia iniecta GCA_001267535.1
196 Erwinia toletana DAPP-PG 735 GCA_000336255.1
196 Rosenbergiella nectarea GCA_900111105.1
zX
218 Clostridium beijerinckii GCA_000833105.2
218 Clostridium puniceum GCA_002006345.1
218 Clostridium saccharoperbutylacetonicum N1-4_28HMT_29 GCA_000340885.1
211 Clostridium taeniosporum GCA_001735765.1
209 Clostridium saccharobutylicum DSM 13864 GCA_000473995.1
zY
186 Rhodobacter sphaeroides 2.4.1 GCA_000012905.2
183 Rhodobacter sphaeroides ATCC 17025 GCA_000016405.1
147 Pseudorhodobacter psychrotolerans GCA_001294535.1
zZ
309 Deinococcus radiodurans R1 GCA_000008565.1
229 Deinococcus gobiensis I-0 GCA_000252445.1
203 Deinococcus puniceus GCA_001644565.1
A0
164 Clostridium beijerinckii GCA_000833105.2
164 Clostridium chromiireducens GCA_002029255.1
164 Clostridium puniceum GCA_002006345.1
164 Clostridium saccharobutylicum DSM 13864 GCA_000473995.1
164 Clostridium saccharoperbutylacetonicum N1-4_28HMT_29 GCA_000340885.1
149 Clostridium butyricum GCA_001456065.2
125 Clostridium neonatale GCA_001458595.1
A1
108 Clostridium beijerinckii GCA_000833105.2
108 Clostridium botulinum B str. Eklund 17B _28NRP_29 GCA_000020165.1
108 Clostridium chromiireducens GCA_002029255.1
108 Clostridium puniceum GCA_002006345.1
108 Clostridium saccharobutylicum DSM 13864 GCA_000473995.1
108 Clostridium saccharoperbutylacetonicum N1-4_28HMT_29 GCA_000340885.1
108 Clostridium taeniosporum GCA_001735765.1
101 Clostridium uliginosum GCA_900112485.1
93 Clostridium butyricum GCA_001456065.2
A2
148 Deinococcus radiodurans R1 GCA_000008565.1
118 Deinococcus marmoris DSM 12784 GCA_000701405.1
118 Deinococcus swuensis GCA_000800395.1
98 Deinococcus deserti VCD115 GCA_000020685.1
A3
295 Deinococcus radiodurans R1 GCA_000008565.1
222 Deinococcus gobiensis I-0 GCA_000252445.1
201 Deinococcus deserti VCD115 GCA_000020685.1
A4
251 Bifidobacterium adolescentis ATCC 15703 GCA_000010425.1
225 Bifidobacterium dentium JCM 1195 = DSM 20436 GCA_001042595.1
212 Bifidobacterium thermophilum GCA_000741495.1
212 Bifidobacterium thermophilum RBL67 GCA_000347695.1
A5
191 Rhodobacter sphaeroides 2.4.1 GCA_000012905.2
162 Gemmobacter megaterium GCA_900156815.1
158 Rhodobacter sphaeroides ATCC 17025 GCA_000016405.1
A6
322 Enterococcus faecalis V583 GCA_000007785.1
322 Streptomyces cinnamoneus GCA_001885705.1
291 Enterococcus canis NBRC 100695 GCA_001544375.1
291 Enterococcus dispar ATCC 51266 GCA_000406945.1
291 Enterococcus faecium DO GCA_000174395.2
291 Enterococcus gilvus ATCC BAA-350 GCA_000407545.1
291 Enterococcus haemoperoxidus ATCC BAA-382 GCA_000407165.1
291 Enterococcus hermanniensis GCA_001885945.1
291 Enterococcus hirae ATCC 9790 GCA_000271405.2
291 Enterococcus malodoratus ATCC 43197 GCA_000407185.1
291 Enterococcus mundtii QU 25 GCA_000504125.1
291 Enterococcus pallens ATCC BAA-351 GCA_000407485.1
291 Enterococcus phoeniculicola ATCC BAA-412 GCA_000407505.1
291 Enterococcus pseudoavium NBRC 100491 GCA_001544295.1
291 Enterococcus rivorum GCA_001742285.1
291 Enterococcus thailandicus GCA_001652875.1
284 Enterococcus asini ATCC 700915 GCA_000407365.1

A7
250 Deinococcus radiodurans R1 GCA_000008565.1
133 Deinococcus gobiensis I-0 GCA_000252445.1
118 Deinococcus proteolyticus MRP GCA_000190555.1
A8
160 Clostridium botulinum B str. Eklund 17B _28NRP_29 GCA_000020165.1
154 Clostridium beijerinckii GCA_000833105.2
154 Clostridium chromiireducens GCA_002029255.1
154 Clostridium puniceum GCA_002006345.1
154 Clostridium saccharobutylicum DSM 13864 GCA_000473995.1
154 Clostridium saccharoperbutylacetonicum N1-4_28HMT_29 GCA_000340885.1
154 Clostridium taeniosporum GCA_001735765.1
145 Clostridium butyricum GCA_001456065.2
A9
175 Escherichia coli IAI39 GCA_000026345.1
175 Escherichia coli O104_3AH4 str. 2011C-3493 GCA_000299455.1
175 Escherichia coli O157_3AH7 str. Sakai GCA_000008865.1
175 Escherichia coli O83_3AH1 str. NRG 857C GCA_000183345.1
175 Escherichia coli UMN026 GCA_000026325.2
175 Escherichia coli str. K-12 substr. MG1655 GCA_000005845.2
175 Shigella dysenteriae Sd197 GCA_000012005.1
175 Shigella flexneri 2a str. 301 GCA_000006925.2
175 Tumebacillus flagellatus GCA_000714935.1
140 Kosakonia cowanii GCA_001975225.1
139 Brenneria goodwinii GCA_001049335.1
139 Enterobacter hormaechei subsp. steigerwaltii GCA_001729725.1
139 Erwinia billingiae Eb661 GCA_000196615.1
139 Erwinia iniecta GCA_001267535.1
139 Erwinia toletana DAPP-PG 735 GCA_000336255.1
139 Klebsiella oxytoca GCA_001022195.1
139 Pantoea agglomerans GCA_001709315.1
139 Pantoea alhagi GCA_002101395.1
139 Pantoea ananatis LMG 20103 GCA_000025405.2
139 Pantoea rwandensis GCA_000759475.1
139 Pantoea septica GCA_002095575.1
139 Pluralibacter gergoviae GCA_000757785.1
139 Pseudescherichia vulneris NBRC 102420 GCA_000759795.1
139 Shimwellia blattae DSM 4481 = NBRC 105725 GCA_000262305.1
Aa
128 Staphylococcus epidermidis ATCC 12228 GCA_000007645.1
128 Staphylococcus haemolyticus JCSC1435 GCA_000009865.1
117 Staphylococcus cohnii subsp. cohnii GCA_000972575.1
117 Staphylococcus hominis subsp. hominis C80 GCA_000183685.1
117 Staphylococcus microti GCA_000934465.1
114 Staphylococcus capitis subsp. capitis GCA_001028645.1
114 Staphylococcus pettenkoferi GCA_002208805.1
Ab
238 Rhodobacter sphaeroides 2.4.1 GCA_000012905.2
173 Rhodobacter sphaeroides ATCC 17025 GCA_000016405.1
160 Defluviimonas alba GCA_001620265.1
Ac
144 Deinococcus radiodurans R1 GCA_000008565.1
95 Deinococcus proteolyticus MRP GCA_000190555.1
68 Deinococcus deserti VCD115 GCA_000020685.1
68 Deinococcus gobiensis I-0 GCA_000252445.1
68 Deinococcus soli Cha et al. 2016 GCA_001007995.1
Ad
150 Clostridium beijerinckii GCA_000833105.2
150 Clostridium puniceum GCA_002006345.1
150 Clostridium saccharoperbutylacetonicum N1-4_28HMT_29 GCA_000340885.1
142 Clostridium chromiireducens GCA_002029255.1
142 Clostridium saccharobutylicum DSM 13864 GCA_000473995.1
138 Clostridium butyricum GCA_001456065.2
Ae
157 Clostridium beijerinckii GCA_000833105.2
157 Clostridium puniceum GCA_002006345.1
157 Clostridium saccharoperbutylacetonicum N1-4_28HMT_29 GCA_000340885.1
148 Clostridium saccharobutylicum DSM 13864 GCA_000473995.1
142 Clostridium butyricum GCA_001456065.2
Af
150 Bifidobacterium adolescentis ATCC 15703 GCA_000010425.1
133 Bifidobacterium tsurumiense GCA_000741765.1
132 Bifidobacterium asteroides PRL2011 GCA_000304215.1
Ag
360 Lactobacillus gasseri ATCC 33323 = JCM 1131 GCA_000014425.1
333 Lactobacillus hominis DSM 23910 = CRBIP 24.179 GCA_000296835.1

261 Lactobacillus crispatus ST1 GCA_000091765.1
Ah
290 Clostridium beijerinckii GCA_000833105.2
290 Clostridium puniceum GCA_002006345.1
290 Clostridium saccharoperbutylacetonicum N1-4_28HMT_29 GCA_000340885.1
288 Clostridium saccharobutylicum DSM 13864 GCA_000473995.1
273 Clostridium chromiireducens GCA_002029255.1
Ai
235 Deinococcus radiodurans R1 GCA_000008565.1
152 Deinococcus gobiensis I-0 GCA_000252445.1
137 Deinococcus puniceus GCA_001644565.1
Aj
209 Deinococcus radiodurans R1 GCA_000008565.1
155 Deinococcus gobiensis I-0 GCA_000252445.1
154 Deinococcus marmoris DSM 12784 GCA_000701405.1
154 Deinococcus swuensis GCA_000800395.1
Ak
166 Streptococcus mutans UA159 GCA_000007465.2
117 Streptococcus equinus GCA_000964315.1
117 Streptococcus gallolyticus subsp. gallolyticus DSM 16831 GCA_002000985.1
114 Streptococcus cristatus AS 1.3089 GCA_000385925.1
114 Streptococcus gordonii str. Challis substr. CH1 GCA_000017005.1
114 Streptococcus parasanguinis ATCC 15912 GCA_000164675.2
Al
252 Clostridium beijerinckii GCA_000833105.2
252 Clostridium puniceum GCA_002006345.1
252 Clostridium saccharoperbutylacetonicum N1-4_28HMT_29 GCA_000340885.1
243 Clostridium saccharobutylicum DSM 13864 GCA_000473995.1
234 Clostridium butyricum GCA_001456065.2
Am
327 Streptococcus mutans UA159 GCA_000007465.2
213 Streptococcus macacae NCTC 11558 GCA_000187995.3
199 Streptococcus sobrinus DSM 20742 = ATCC 33478 GCA_000686605.1
An
212 Rhodobacter sphaeroides 2.4.1 GCA_000012905.2
185 Rhodobacter sphaeroides ATCC 17025 GCA_000016405.1
184 Gemmobacter megaterium GCA_900156815.1
Ao
141 Deinococcus radiodurans R1 GCA_000008565.1
104 Deinococcus frigens DSM 12807 GCA_000701425.1
104 Deinococcus marmoris DSM 12784 GCA_000701405.1
104 Deinococcus soli Cha et al. 2016 GCA_001007995.1
104 Deinococcus swuensis GCA_000800395.1
101 Deinococcus geothermalis DSM 11300 GCA_000196275.1
101 Deinococcus pimensis DSM 21231 GCA_000519345.1
101 Deinococcus puniceus GCA_001644565.1
Ap
285 Bifidobacterium adolescentis ATCC 15703 GCA_000010425.1
249 Bifidobacterium callitrichos DSM 23973 GCA_000741175.1
243 Bifidobacterium asteroides PRL2011 GCA_000304215.1
Aq
86 Erwinia iniecta GCA_001267535.1
86 Escherichia coli IAI39 GCA_000026345.1
86 Escherichia coli O104_3AH4 str. 2011C-3493 GCA_000299455.1
86 Escherichia coli O157_3AH7 str. Sakai GCA_000008865.1
86 Escherichia coli O83_3AH1 str. NRG 857C GCA_000183345.1
86 Escherichia coli UMN026 GCA_000026325.2
86 Escherichia coli str. K-12 substr. MG1655 GCA_000005845.2
86 Shigella dysenteriae Sd197 GCA_000012005.1
86 Shigella flexneri 2a str. 301 GCA_000006925.2
86 Tumebacillus flagellatus GCA_000714935.1
84 Cronobacter sakazakii GCA_000982825.1
84 Shimwellia blattae DSM 4481 = NBRC 105725 GCA_000262305.1
73 Serratia rubidaea GCA_001572725.1
Ar
245 Bacillus anthracis str. Ames GCA_000007845.1
245 Bacillus anthracis str. Sterne GCA_000008165.1
245 Bacillus cereus ATCC 14579 GCA_000007825.1
245 Bacillus mycoides GCA_000832605.1
245 Bacillus pseudomycoides DSM 12442 GCA_000161455.1
245 Bacillus thuringiensis YBT-1518 GCA_000497525.2
245 _5BBacillus thuringiensis_5D serovar konkukian str. 97-27 GCA_000008505.1
176 Bacillus manliponensis GCA_000712595.1
171 Bacillus cytotoxicus NVH 391-98 GCA_000017425.1
As
264 Staphylococcus aureus subsp. aureus NCTC 8325 GCA_000013425.1

264 Staphylococcus capitis subsp. capitis GCA_001028645.1
264 Staphylococcus epidermidis ATCC 12228 GCA_000007645.1
264 Staphylococcus haemolyticus JCSC1435 GCA_000009865.1
264 Staphylococcus hominis subsp. hominis C80 GCA_000183685.1
264 Staphylococcus simiae CCM 7213 GCA_000235645.2
260 Staphylococcus cohnii subsp. cohnii GCA_000972575.1
255 Staphylococcus equorum GCA_001432245.1
255 Staphylococcus succinus GCA_001902315.1
At
183 Rhodobacter sphaeroides 2.4.1 GCA_000012905.2
168 Rhodobacter sphaeroides ATCC 17025 GCA_000016405.1
158 Gemmobacter megaterium GCA_900156815.1
158 Pseudorhodobacter ferrugineus DSM 5888 GCA_000420745.1
158 Pseudorhodobacter wandonensis GCA_001202035.1
Au
216 Staphylococcus capitis subsp. capitis GCA_001028645.1
216 Staphylococcus epidermidis ATCC 12228 GCA_000007645.1
204 Staphylococcus condimenti GCA_001618885.1
204 Staphylococcus simulans GCA_001559115.1
201 Staphylococcus aureus subsp. aureus NCTC 8325 GCA_000013425.1
201 Staphylococcus haemolyticus JCSC1435 GCA_000009865.1
201 Staphylococcus hominis subsp. hominis C80 GCA_000183685.1
201 Staphylococcus lugdunensis HKU09-01 GCA_000025085.1
201 Staphylococcus simiae CCM 7213 GCA_000235645.2
Av
275 Rhodobacter sphaeroides 2.4.1 GCA_000012905.2
254 Pseudorhodobacter ferrugineus DSM 5888 GCA_000420745.1
254 Pseudorhodobacter psychrotolerans GCA_001294535.1
254 Pseudorhodobacter wandonensis GCA_001202035.1
252 Rhodobacter sphaeroides ATCC 17025 GCA_000016405.1
Aw
190 Escherichia coli IAI39 GCA_000026345.1
190 Escherichia coli O104_3AH4 str. 2011C-3493 GCA_000299455.1
190 Escherichia coli O157_3AH7 str. Sakai GCA_000008865.1
190 Escherichia coli O83_3AH1 str. NRG 857C GCA_000183345.1
190 Escherichia coli UMN026 GCA_000026325.2
190 Escherichia coli str. K-12 substr. MG1655 GCA_000005845.2
190 Shigella dysenteriae Sd197 GCA_000012005.1
190 Shigella flexneri 2a str. 301 GCA_000006925.2
190 Tumebacillus flagellatus GCA_000714935.1
163 Kosakonia sacchari SP1 GCA_000300455.4
163 _5BEnterobacter_5D lignolyticus SCF1 GCA_000164865.1
149 Klebsiella pneumoniae subsp. pneumoniae HS11286 GCA_000240185.2
Ax
277 Bacillus anthracis str. Ames GCA_000007845.1
277 Bacillus anthracis str. Sterne GCA_000008165.1
277 Bacillus cereus ATCC 14579 GCA_000007825.1
277 Bacillus mycoides GCA_000832605.1
277 Bacillus pseudomycoides DSM 12442 GCA_000161455.1
277 Bacillus thuringiensis YBT-1518 GCA_000497525.2
277 _5BBacillus thuringiensis_5D serovar konkukian str. 97-27 GCA_000008505.1
251 Bacillus aquimaris TF-12 GCA_001648555.1
247 Bacillus manliponensis GCA_000712595.1
Ay
344 Staphylococcus epidermidis ATCC 12228 GCA_000007645.1
339 Staphylococcus capitis subsp. capitis GCA_001028645.1
337 Staphylococcus haemolyticus JCSC1435 GCA_000009865.1
Az
190 Clostridium beijerinckii GCA_000833105.2
190 Clostridium puniceum GCA_002006345.1
190 Clostridium saccharoperbutylacetonicum N1-4_28HMT_29 GCA_000340885.1
182 Clostridium chromiireducens GCA_002029255.1
182 Clostridium saccharobutylicum DSM 13864 GCA_000473995.1
181 Clostridium taeniosporum GCA_001735765.1
AA
288 Rhodobacter sphaeroides 2.4.1 GCA_000012905.2
273 Rhodobacter sphaeroides ATCC 17025 GCA_000016405.1
252 Pseudorhodobacter psychrotolerans GCA_001294535.1
AB
187 Escherichia coli IAI39 GCA_000026345.1
187 Escherichia coli O104_3AH4 str. 2011C-3493 GCA_000299455.1
187 Escherichia coli O157_3AH7 str. Sakai GCA_000008865.1
187 Escherichia coli O83_3AH1 str. NRG 857C GCA_000183345.1
187 Escherichia coli UMN026 GCA_000026325.2
187 Escherichia coli str. K-12 substr. MG1655 GCA_000005845.2
187 Shigella dysenteriae Sd197 GCA_000012005.1

187 Shigella flexneri 2a str. 301 GCA_000006925.2
187 Tumebacillus flagellatus GCA_000714935.1
147 Erwinia iniecta GCA_001267535.1
147 Rosenbergiella nectarea GCA_900111105.1
144 Izhakiella capsodis GCA_900115045.1
AC
127 Rhodobacter sphaeroides 2.4.1 GCA_000012905.2
124 Haematobacter massiliensis GCA_000740795.1
119 Clostridium amylolyticum GCA_900142075.1
119 Clostridium beijerinckii GCA_000833105.2
119 Clostridium chauvoei JF4335 GCA_900168365.1
119 Clostridium intestinale URNW GCA_000469625.2
119 Clostridium polynesiense GCA_000820705.1
119 Clostridium puniceum GCA_002006345.1
119 Clostridium saccharobutylicum DSM 13864 GCA_000473995.1
119 Clostridium saccharoperbutylacetonicum N1-4_28HMT_29 GCA_000340885.1
119 Clostridium sartagoforme AAU1 GCA_000401215.1
AD
248 Clostridium beijerinckii GCA_000833105.2
248 Clostridium saccharobutylicum DSM 13864 GCA_000473995.1
248 Clostridium saccharoperbutylacetonicum N1-4_28HMT_29 GCA_000340885.1
242 Clostridium puniceum GCA_002006345.1
231 Clostridium chromiireducens GCA_002029255.1
AE
203 Escherichia coli IAI39 GCA_000026345.1
203 Escherichia coli O104_3AH4 str. 2011C-3493 GCA_000299455.1
203 Escherichia coli O157_3AH7 str. Sakai GCA_000008865.1
203 Escherichia coli O83_3AH1 str. NRG 857C GCA_000183345.1
203 Escherichia coli UMN026 GCA_000026325.2
203 Escherichia coli str. K-12 substr. MG1655 GCA_000005845.2
203 Shigella dysenteriae Sd197 GCA_000012005.1
203 Shigella flexneri 2a str. 301 GCA_000006925.2
203 Tumebacillus flagellatus GCA_000714935.1
165 Erwinia iniecta GCA_001267535.1
162 Enterobacter hormaechei subsp. steigerwaltii GCA_001729725.1
162 Erwinia toletana DAPP-PG 735 GCA_000336255.1
162 Klebsiella oxytoca GCA_001022195.1
162 Kosakonia cowanii GCA_001975225.1
162 Kosakonia sacchari SP1 GCA_000300455.4
162 Pluralibacter gergoviae GCA_000757785.1
162 Pseudescherichia vulneris NBRC 102420 GCA_000759795.1
AF
217 Escherichia coli IAI39 GCA_000026345.1
217 Escherichia coli O104_3AH4 str. 2011C-3493 GCA_000299455.1
217 Escherichia coli O157_3AH7 str. Sakai GCA_000008865.1
217 Escherichia coli O83_3AH1 str. NRG 857C GCA_000183345.1
217 Escherichia coli UMN026 GCA_000026325.2
217 Escherichia coli str. K-12 substr. MG1655 GCA_000005845.2
217 Shigella flexneri 2a str. 301 GCA_000006925.2
217 Tumebacillus flagellatus GCA_000714935.1
202 Shigella dysenteriae Sd197 GCA_000012005.1
194 Erwinia iniecta GCA_001267535.1
AG
209 Bifidobacterium breve DSM 20213 = JCM 1192 GCA_001025175.1
209 Bifidobacterium tsurumiense GCA_000741765.1
208 Bifidobacterium adolescentis ATCC 15703 GCA_000010425.1
207 Bifidobacterium longum NCC2705 GCA_000007525.1
207 Bifidobacterium saguini DSM 23967 GCA_000741715.1
AH
245 Deinococcus radiodurans R1 GCA_000008565.1
191 Deinococcus marmoris DSM 12784 GCA_000701405.1
191 Deinococcus swuensis GCA_000800395.1
160 Deinococcus gobiensis I-0 GCA_000252445.1
AI
194 Escherichia coli IAI39 GCA_000026345.1
194 Escherichia coli O104_3AH4 str. 2011C-3493 GCA_000299455.1
194 Escherichia coli O157_3AH7 str. Sakai GCA_000008865.1
194 Escherichia coli O83_3AH1 str. NRG 857C GCA_000183345.1
194 Escherichia coli UMN026 GCA_000026325.2
194 Escherichia coli str. K-12 substr. MG1655 GCA_000005845.2
194 Shigella flexneri 2a str. 301 GCA_000006925.2
194 Tumebacillus flagellatus GCA_000714935.1
179 Shigella dysenteriae Sd197 GCA_000012005.1
154 Erwinia iniecta GCA_001267535.1
AJ
225 Staphylococcus epidermidis ATCC 12228 GCA_000007645.1

212 Staphylococcus haemolyticus JCSC1435 GCA_000009865.1
204 Staphylococcus capitis subsp. capitis GCA_001028645.1
AK
236 Escherichia coli IAI39 GCA_000026345.1
236 Escherichia coli O104_3AH4 str. 2011C-3493 GCA_000299455.1
236 Escherichia coli O157_3AH7 str. Sakai GCA_000008865.1
236 Escherichia coli O83_3AH1 str. NRG 857C GCA_000183345.1
236 Escherichia coli UMN026 GCA_000026325.2
236 Escherichia coli str. K-12 substr. MG1655 GCA_000005845.2
236 Shigella flexneri 2a str. 301 GCA_000006925.2
236 Tumebacillus flagellatus GCA_000714935.1
227 Shigella dysenteriae Sd197 GCA_000012005.1
214 Erwinia iniecta GCA_001267535.1
AL
225 Enterococcus faecalis V583 GCA_000007785.1
225 Streptomyces cinnamoneus GCA_001885705.1
204 Vagococcus penaei GCA_001998885.1
195 Enterococcus dispar ATCC 51266 GCA_000406945.1
AM
292 Lactobacillus gasseri ATCC 33323 = JCM 1131 GCA_000014425.1
285 Lactobacillus hominis DSM 23910 = CRBIP 24.179 GCA_000296835.1
210 Lactobacillus psittaci DSM 15354 GCA_000425905.1
AN
264 Streptococcus mutans UA159 GCA_000007465.2
145 Streptococcus ratti FA-1 = DSM 20564 GCA_000286075.1
143 Streptococcus massiliensis DSM 18628 GCA_000380065.1
AO
228 Clostridium beijerinckii GCA_000833105.2
228 Clostridium saccharobutylicum DSM 13864 GCA_000473995.1
228 Clostridium saccharoperbutylacetonicum N1-4_28HMT_29 GCA_000340885.1
221 Clostridium puniceum GCA_002006345.1
199 Clostridium chromiireducens GCA_002029255.1
AP
241 Pseudorhodobacter psychrotolerans GCA_001294535.1
238 Rhodobacter sphaeroides 2.4.1 GCA_000012905.2
220 Rhodobacter sphaeroides ATCC 17025 GCA_000016405.1
AQ
202 Bifidobacterium adolescentis ATCC 15703 GCA_000010425.1
167 Bifidobacterium thermophilum GCA_000741495.1
162 Bifidobacterium dentium JCM 1195 = DSM 20436 GCA_001042595.1
AR
302 Deinococcus radiodurans R1 GCA_000008565.1
224 Deinococcus gobiensis I-0 GCA_000252445.1
206 Deinococcus marmoris DSM 12784 GCA_000701405.1
206 Deinococcus swuensis GCA_000800395.1
AS
227 Bifidobacterium adolescentis ATCC 15703 GCA_000010425.1
218 Bifidobacterium dentium JCM 1195 = DSM 20436 GCA_001042595.1
207 Bifidobacterium tsurumiense GCA_000741765.1
AT
232 Lactobacillus gasseri ATCC 33323 = JCM 1131 GCA_000014425.1
219 Lactobacillus hominis DSM 23910 = CRBIP 24.179 GCA_000296835.1
176 Lactobacillus crispatus ST1 GCA_000091765.1
176 Lactobacillus kefiranofaciens ZW3 GCA_000214785.1
AU
263 Streptococcus mutans UA159 GCA_000007465.2
164 Streptococcus ratti FA-1 = DSM 20564 GCA_000286075.1
158 Streptococcus equinus GCA_000964315.1
158 Streptococcus gallolyticus subsp. gallolyticus DSM 16831 GCA_002000985.1
158 Streptococcus gordonii str. Challis substr. CH1 GCA_000017005.1
AV
194 Rhodobacter sphaeroides 2.4.1 GCA_000012905.2
194 Rhodobacter sphaeroides ATCC 17025 GCA_000016405.1
169 Gemmobacter megaterium GCA_900156815.1
169 Pseudorhodobacter ferrugineus DSM 5888 GCA_000420745.1
169 Pseudorhodobacter wandonensis GCA_001202035.1
155 Defluviimonas alba GCA_001620265.1
AW
157 Lactobacillus gasseri ATCC 33323 = JCM 1131 GCA_000014425.1
157 Lactobacillus hominis DSM 23910 = CRBIP 24.179 GCA_000296835.1
119 Lactobacillus amylovorus GCA_000191545.1
119 Lactobacillus crispatus ST1 GCA_000091765.1
115 Lactobacillus kalixensis DSM 16043 GCA_001434335.1
AX
269 Escherichia coli IAI39 GCA_000026345.1
269 Escherichia coli O104_3AH4 str. 2011C-3493 GCA_000299455.1

269 Escherichia coli O157_3AH7 str. Sakai GCA_000008865.1
269 Escherichia coli O83_3AH1 str. NRG 857C GCA_000183345.1
269 Escherichia coli UMN026 GCA_000026325.2
269 Escherichia coli str. K-12 substr. MG1655 GCA_000005845.2
269 Shigella flexneri 2a str. 301 GCA_000006925.2
269 Tumebacillus flagellatus GCA_000714935.1
260 Shigella dysenteriae Sd197 GCA_000012005.1
257 Erwinia iniecta GCA_001267535.1
AY
247 Deinococcus radiodurans R1 GCA_000008565.1
193 Deinococcus gobiensis I-0 GCA_000252445.1
163 Deinococcus deserti VCD115 GCA_000020685.1
AZ
180 Clostridium beijerinckii GCA_000833105.2
180 Clostridium botulinum B str. Eklund 17B _28NRP_29 GCA_000020165.1
180 Clostridium chromiireducens GCA_002029255.1
180 Clostridium puniceum GCA_002006345.1
180 Clostridium saccharobutylicum DSM 13864 GCA_000473995.1
180 Clostridium saccharoperbutylacetonicum N1-4_28HMT_29 GCA_000340885.1
180 Clostridium taeniosporum GCA_001735765.1
180 Clostridium uliginosum GCA_900112485.1
176 Clostridium fallax GCA_900129365.1
176 Clostridium gasigenes GCA_900104115.1
171 Clostridium amylolyticum GCA_900142075.1
171 Clostridium butyricum GCA_001456065.2
171 Clostridium polynesiense GCA_000820705.1
B0
243 Clostridium beijerinckii GCA_000833105.2
243 Clostridium saccharoperbutylacetonicum N1-4_28HMT_29 GCA_000340885.1
236 Clostridium puniceum GCA_002006345.1
235 Clostridium saccharobutylicum DSM 13864 GCA_000473995.1
B1
121 Enterococcus canis NBRC 100695 GCA_001544375.1
121 Enterococcus dispar ATCC 51266 GCA_000406945.1
121 Enterococcus faecalis V583 GCA_000007785.1
121 Enterococcus faecium DO GCA_000174395.2
121 Enterococcus gilvus ATCC BAA-350 GCA_000407545.1
121 Enterococcus hermanniensis GCA_001885945.1
121 Enterococcus hirae ATCC 9790 GCA_000271405.2
121 Enterococcus malodoratus ATCC 43197 GCA_000407185.1
121 Enterococcus massiliensis GCA_001050095.1
121 Enterococcus mundtii QU 25 GCA_000504125.1
121 Enterococcus pallens ATCC BAA-351 GCA_000407485.1
121 Enterococcus pseudoavium NBRC 100491 GCA_001544295.1
121 Enterococcus rivorum GCA_001742285.1
121 Streptomyces cinnamoneus GCA_001885705.1
113 Enterococcus aquimarinus GCA_001885765.1
113 Enterococcus haemoperoxidus ATCC BAA-382 GCA_000407165.1
113 Enterococcus phoeniculicola ATCC BAA-412 GCA_000407505.1
113 Enterococcus thailandicus GCA_001652875.1
108 Aerococcus urinaehominis GCA_001543245.1
B2
137 Deinococcus radiodurans R1 GCA_000008565.1
107 Deinococcus puniceus GCA_001644565.1
86 Deinococcus deserti VCD115 GCA_000020685.1
86 Deinococcus proteolyticus MRP GCA_000190555.1
B3
236 Deinococcus radiodurans R1 GCA_000008565.1
134 Deinococcus geothermalis DSM 11300 GCA_000196275.1
132 Deinococcus gobiensis I-0 GCA_000252445.1
132 Deinococcus puniceus GCA_001644565.1
B4
227 Bacillus thuringiensis YBT-1518 GCA_000497525.2
219 Bacillus anthracis str. Ames GCA_000007845.1
219 Bacillus anthracis str. Sterne GCA_000008165.1
219 Bacillus cereus ATCC 14579 GCA_000007825.1
219 Bacillus mycoides GCA_000832605.1
219 Bacillus pseudomycoides DSM 12442 GCA_000161455.1
219 _5BBacillus thuringiensis_5D serovar konkukian str. 97-27 GCA_000008505.1
192 Bacillus manliponensis GCA_000712595.1
B5
221 Escherichia coli IAI39 GCA_000026345.1
221 Escherichia coli O104_3AH4 str. 2011C-3493 GCA_000299455.1
221 Escherichia coli O157_3AH7 str. Sakai GCA_000008865.1
221 Escherichia coli O83_3AH1 str. NRG 857C GCA_000183345.1
221 Escherichia coli UMN026 GCA_000026325.2

221 Escherichia coli str. K-12 substr. MG1655 GCA_000005845.2
221 Shigella flexneri 2a str. 301 GCA_000006925.2
221 Tumebacillus flagellatus GCA_000714935.1
217 Erwinia iniecta GCA_001267535.1
217 Erwinia toletana DAPP-PG 735 GCA_000336255.1
215 Pantoea ananatis LMG 20103 GCA_000025405.2
215 Shimwellia blattae DSM 4481 = NBRC 105725 GCA_000262305.1
B6
238 Deinococcus radiodurans R1 GCA_000008565.1
177 Deinococcus gobiensis I-0 GCA_000252445.1
168 Deinococcus puniceus GCA_001644565.1
B7
164 Bifidobacterium lemurum GCA_001895165.1
155 Bifidobacterium thermophilum GCA_000741495.1
150 Bifidobacterium dentium JCM 1195 = DSM 20436 GCA_001042595.1
B8
208 Clostridium beijerinckii GCA_000833105.2
208 Clostridium saccharoperbutylacetonicum N1-4_28HMT_29 GCA_000340885.1
201 Clostridium puniceum GCA_002006345.1
193 Clostridium saccharobutylicum DSM 13864 GCA_000473995.1
B9
75 Bifidobacterium actinocoloniiforme DSM 22766 GCA_001263395.1
75 Bifidobacterium angulatum DSM 20098 = JCM 7096 GCA_001025155.1
75 Bifidobacterium asteroides PRL2011 GCA_000304215.1
75 Bifidobacterium bombi DSM 19703 GCA_000737845.1
75 Bifidobacterium callitrichos DSM 23973 GCA_000741175.1
75 Bifidobacterium lemurum GCA_001895165.1
75 Bifidobacterium reuteri DSM 23975 GCA_000741695.1
74 Bifidobacterium adolescentis ATCC 15703 GCA_000010425.1
74 Bifidobacterium dentium JCM 1195 = DSM 20436 GCA_001042595.1
74 Bifidobacterium stellenboschense GCA_000741785.1
74 Bifidobacterium thermophilum GCA_000741495.1
74 Bifidobacterium thermophilum RBL67 GCA_000347695.1
74 Bifidobacterium tsurumiense GCA_000741765.1
72 Bifidobacterium animalis subsp. lactis DSM 10140 GCA_000022965.1
72 Bifidobacterium choerinum GCA_000741135.1
72 Bifidobacterium cuniculi GCA_000741575.1
72 Bifidobacterium gallicum DSM 20093 = LMG 11596 GCA_000741205.1
72 Bifidobacterium magnum GCA_000741255.1
72 Bifidobacterium pseudolongum PV8-2 GCA_000800475.2
72 Scardovia wiggsiae F0424 GCA_000269605.1
Ba
187 Clostridium beijerinckii GCA_000833105.2
187 Clostridium puniceum GCA_002006345.1
187 Clostridium saccharobutylicum DSM 13864 GCA_000473995.1
187 Clostridium saccharoperbutylacetonicum N1-4_28HMT_29 GCA_000340885.1
172 Clostridium chromiireducens GCA_002029255.1
172 Clostridium taeniosporum GCA_001735765.1
165 Clostridium butyricum GCA_001456065.2
Bb
226 Enterococcus faecalis V583 GCA_000007785.1
226 Streptomyces cinnamoneus GCA_001885705.1
194 Enterococcus dispar ATCC 51266 GCA_000406945.1
191 Enterococcus asini ATCC 700915 GCA_000407365.1
191 Enterococcus canis NBRC 100695 GCA_001544375.1
191 Enterococcus faecium DO GCA_000174395.2
191 Enterococcus hirae ATCC 9790 GCA_000271405.2
191 Enterococcus massiliensis GCA_001050095.1
191 Enterococcus mundtii QU 25 GCA_000504125.1
191 Enterococcus rivorum GCA_001742285.1
Bc
148 Staphylococcus capitis subsp. capitis GCA_001028645.1
148 Staphylococcus epidermidis ATCC 12228 GCA_000007645.1
144 Staphylococcus lugdunensis HKU09-01 GCA_000025085.1
137 Staphylococcus haemolyticus JCSC1435 GCA_000009865.1
Bd
317 Clostridium beijerinckii GCA_000833105.2
317 Clostridium puniceum GCA_002006345.1
317 Clostridium saccharobutylicum DSM 13864 GCA_000473995.1
317 Clostridium saccharoperbutylacetonicum N1-4_28HMT_29 GCA_000340885.1
313 Clostridium chromiireducens GCA_002029255.1
296 Clostridium butyricum GCA_001456065.2
Be
234 Deinococcus radiodurans R1 GCA_000008565.1
136 Deinococcus puniceus GCA_001644565.1
118 Deinococcus marmoris DSM 12784 GCA_000701405.1

118 Deinococcus swuensis GCA_000800395.1
Bf
197 Rhodobacter sphaeroides 2.4.1 GCA_000012905.2
161 Rhodobacter sphaeroides ATCC 17025 GCA_000016405.1
146 Pseudorhodobacter ferrugineus DSM 5888 GCA_000420745.1
146 Pseudorhodobacter wandonensis GCA_001202035.1
Bg
195 Streptococcus mutans UA159 GCA_000007465.2
130 Streptococcus merionis DSM 19192 GCA_000380085.1
122 Streptococcus equinus GCA_000964315.1
122 Streptococcus gallolyticus subsp. gallolyticus DSM 16831 GCA_002000985.1
Bh
199 Bacillus anthracis str. Ames GCA_000007845.1
199 Bacillus anthracis str. Sterne GCA_000008165.1
199 Bacillus cereus ATCC 14579 GCA_000007825.1
199 Bacillus mycoides GCA_000832605.1
199 Bacillus pseudomycoides DSM 12442 GCA_000161455.1
199 Bacillus thuringiensis YBT-1518 GCA_000497525.2
199 _5BBacillus thuringiensis_5D serovar konkukian str. 97-27 GCA_000008505.1
158 Staphylococcus equorum GCA_001432245.1
155 Bacillus cytotoxicus NVH 391-98 GCA_000017425.1
Bi
249 Clostridium puniceum GCA_002006345.1
234 Clostridium beijerinckii GCA_000833105.2
234 Clostridium saccharoperbutylacetonicum N1-4_28HMT_29 GCA_000340885.1
225 Clostridium chromiireducens GCA_002029255.1
225 Clostridium saccharobutylicum DSM 13864 GCA_000473995.1
Bj
151 Staphylococcus condimenti GCA_001618885.1
151 Staphylococcus simulans GCA_001559115.1
150 Megasphaera cerevisiae DSM 20462 GCA_001045675.1
150 Staphylococcus aureus subsp. aureus NCTC 8325 GCA_000013425.1
150 Staphylococcus capitis subsp. capitis GCA_001028645.1
150 Staphylococcus epidermidis ATCC 12228 GCA_000007645.1
150 Staphylococcus haemolyticus JCSC1435 GCA_000009865.1
150 Staphylococcus hominis subsp. hominis C80 GCA_000183685.1
150 Staphylococcus lugdunensis HKU09-01 GCA_000025085.1
150 Staphylococcus lutrae GCA_002101335.1
150 Staphylococcus simiae CCM 7213 GCA_000235645.2
150 Staphylococcus warneri SG1 GCA_000332735.1
136 Staphylococcus microti GCA_000934465.1
136 Staphylococcus pseudintermedius HKU10-03 GCA_000185885.1
Bk
235 Staphylococcus capitis subsp. capitis GCA_001028645.1
235 Staphylococcus epidermidis ATCC 12228 GCA_000007645.1
235 Staphylococcus lugdunensis HKU09-01 GCA_000025085.1
231 Staphylococcus cohnii subsp. cohnii GCA_000972575.1
231 Staphylococcus saprophyticus subsp. saprophyticus ATCC 15305 GCA_000010125.1
231 Staphylococcus xylosus GCA_000706685.1
228 Megasphaera cerevisiae DSM 20462 GCA_001045675.1
228 Staphylococcus aureus subsp. aureus NCTC 8325 GCA_000013425.1
228 Staphylococcus haemolyticus JCSC1435 GCA_000009865.1
228 Staphylococcus hominis subsp. hominis C80 GCA_000183685.1
228 Staphylococcus simiae CCM 7213 GCA_000235645.2
228 Staphylococcus warneri SG1 GCA_000332735.1
Bl
170 Deinococcus radiodurans R1 GCA_000008565.1
126 Deinococcus puniceus GCA_001644565.1
122 Deinococcus deserti VCD115 GCA_000020685.1
Bm
159 Deinococcus radiodurans R1 GCA_000008565.1
97 Deinococcus gobiensis I-0 GCA_000252445.1
72 Deinococcus geothermalis DSM 11300 GCA_000196275.1
Bn
233 Streptococcus mutans UA159 GCA_000007465.2
189 Streptococcus equinus GCA_000964315.1
189 Streptococcus gallolyticus subsp. gallolyticus DSM 16831 GCA_002000985.1
184 Streptococcus merionis DSM 19192 GCA_000380085.1
Bo
338 Lactobacillus gasseri ATCC 33323 = JCM 1131 GCA_000014425.1
331 Lactobacillus hominis DSM 23910 = CRBIP 24.179 GCA_000296835.1
239 Lactobacillus psittaci DSM 15354 GCA_000425905.1
Bp
265 Clostridium beijerinckii GCA_000833105.2
265 Clostridium puniceum GCA_002006345.1
265 Clostridium saccharobutylicum DSM 13864 GCA_000473995.1

265 Clostridium saccharoperbutylacetonicum N1-4_28HMT_29 GCA_000340885.1
261 Clostridium chromiireducens GCA_002029255.1
238 Clostridium butyricum GCA_001456065.2
Bq
268 Streptococcus mutans UA159 GCA_000007465.2
191 Streptococcus ratti FA-1 = DSM 20564 GCA_000286075.1
179 Streptococcus equinus GCA_000964315.1
179 Streptococcus gallolyticus subsp. gallolyticus DSM 16831 GCA_002000985.1
179 Streptococcus orisratti DSM 15617 GCA_000380105.1
Br
240 Deinococcus radiodurans R1 GCA_000008565.1
171 Deinococcus gobiensis I-0 GCA_000252445.1
163 Deinococcus deserti VCD115 GCA_000020685.1
Bs
247 Rhodobacter sphaeroides 2.4.1 GCA_000012905.2
214 Rhodobacter sphaeroides ATCC 17025 GCA_000016405.1
189 Gemmobacter megaterium GCA_900156815.1
Bt
308 Streptococcus mutans UA159 GCA_000007465.2
179 Streptococcus ratti FA-1 = DSM 20564 GCA_000286075.1
171 Streptococcus ferus DSM 20646 GCA_000372425.1
Bu
277 Enterococcus faecalis V583 GCA_000007785.1
277 Streptomyces cinnamoneus GCA_001885705.1
239 Enterococcus asini ATCC 700915 GCA_000407365.1
239 Enterococcus canis NBRC 100695 GCA_001544375.1
239 Enterococcus dispar ATCC 51266 GCA_000406945.1
239 Enterococcus faecium DO GCA_000174395.2
239 Enterococcus haemoperoxidus ATCC BAA-382 GCA_000407165.1
239 Enterococcus hirae ATCC 9790 GCA_000271405.2
239 Enterococcus massiliensis GCA_001050095.1
239 Enterococcus mundtii QU 25 GCA_000504125.1
239 Enterococcus phoeniculicola ATCC BAA-412 GCA_000407505.1
239 Enterococcus rivorum GCA_001742285.1
239 Enterococcus thailandicus GCA_001652875.1
228 Melissococcus plutonius S1 GCA_000747585.1
Bv
203 Bifidobacterium adolescentis ATCC 15703 GCA_000010425.1
162 Bifidobacterium breve DSM 20213 = JCM 1192 GCA_001025175.1
149 Bifidobacterium callitrichos DSM 23973 GCA_000741175.1
149 Bifidobacterium coryneforme GCA_000737865.1
Bw
352 Lactobacillus gasseri ATCC 33323 = JCM 1131 GCA_000014425.1
326 Lactobacillus hominis DSM 23910 = CRBIP 24.179 GCA_000296835.1
262 Lactobacillus crispatus ST1 GCA_000091765.1
Bx
104 Clostridium beijerinckii GCA_000833105.2
104 Clostridium botulinum A str. ATCC 3502 GCA_000063585.1
104 Clostridium botulinum A str. Hall GCA_000017045.1
104 Clostridium butyricum GCA_001456065.2
104 Clostridium chromiireducens GCA_002029255.1
104 Clostridium neonatale GCA_001458595.1
104 Clostridium puniceum GCA_002006345.1
104 Clostridium saccharobutylicum DSM 13864 GCA_000473995.1
104 Clostridium saccharoperbutylacetonicum N1-4_28HMT_29 GCA_000340885.1
104 Clostridium sulfidigenes GCA_000732635.1
104 Clostridium tepidum GCA_002008345.1
104 Clostridium tyrobutyricum GCA_001642655.1
99 Clostridium acetireducens DSM 10703 GCA_001758365.1
99 Clostridium kluyveri DSM 555 GCA_000016505.1
99 Clostridium ventriculi GCA_001404895.1
97 Clostridium carboxidivorans P7 GCA_001038625.1
97 Clostridium ihumii AP5 GCA_000612845.1
97 Clostridium senegalense JC122 GCA_000285575.1
By
208 Rhodobacter sphaeroides 2.4.1 GCA_000012905.2
188 Rhodobacter sphaeroides ATCC 17025 GCA_000016405.1
176 Pseudorhodobacter ferrugineus DSM 5888 GCA_000420745.1
176 Pseudorhodobacter wandonensis GCA_001202035.1
Bz
272 Clostridium beijerinckii GCA_000833105.2
272 Clostridium saccharoperbutylacetonicum N1-4_28HMT_29 GCA_000340885.1
266 Clostridium puniceum GCA_002006345.1
263 Clostridium saccharobutylicum DSM 13864 GCA_000473995.1
BA
227 Rhodobacter sphaeroides 2.4.1 GCA_000012905.2

212 Rhodobacter sphaeroides ATCC 17025 GCA_000016405.1
194 Gemmobacter megaterium GCA_900156815.1
BB
247 Escherichia coli IAI39 GCA_000026345.1
247 Escherichia coli O104_3AH4 str. 2011C-3493 GCA_000299455.1
247 Escherichia coli O157_3AH7 str. Sakai GCA_000008865.1
247 Escherichia coli O83_3AH1 str. NRG 857C GCA_000183345.1
247 Escherichia coli UMN026 GCA_000026325.2
247 Escherichia coli str. K-12 substr. MG1655 GCA_000005845.2
247 Shigella flexneri 2a str. 301 GCA_000006925.2
247 Tumebacillus flagellatus GCA_000714935.1
238 Shigella dysenteriae Sd197 GCA_000012005.1
208 Erwinia iniecta GCA_001267535.1
BC
129 Bifidobacterium adolescentis ATCC 15703 GCA_000010425.1
119 Bifidobacterium dentium JCM 1195 = DSM 20436 GCA_001042595.1
119 Bifidobacterium tsurumiense GCA_000741765.1
116 Bifidobacterium choerinum GCA_000741135.1
116 Bifidobacterium pseudolongum PV8-2 GCA_000800475.2
BD
184 Staphylococcus epidermidis ATCC 12228 GCA_000007645.1
177 Staphylococcus capitis subsp. capitis GCA_001028645.1
177 Staphylococcus haemolyticus JCSC1435 GCA_000009865.1
165 Staphylococcus hominis subsp. hominis C80 GCA_000183685.1
BE
186 Rhodobacter sphaeroides ATCC 17025 GCA_000016405.1
177 Rhodobacter sphaeroides 2.4.1 GCA_000012905.2
161 Pseudorhodobacter psychrotolerans GCA_001294535.1
BF
131 Bacillus humi GCA_001439915.1
131 Escherichia coli IAI39 GCA_000026345.1
131 Escherichia coli O104_3AH4 str. 2011C-3493 GCA_000299455.1
131 Escherichia coli O157_3AH7 str. Sakai GCA_000008865.1
131 Escherichia coli O83_3AH1 str. NRG 857C GCA_000183345.1
131 Escherichia coli UMN026 GCA_000026325.2
131 Escherichia coli str. K-12 substr. MG1655 GCA_000005845.2
131 Shigella flexneri 2a str. 301 GCA_000006925.2
131 Tumebacillus flagellatus GCA_000714935.1
118 Shigella dysenteriae Sd197 GCA_000012005.1
117 Erwinia iniecta GCA_001267535.1
117 Serratia marcescens subsp. marcescens Db11 GCA_000513215.1
BG
156 Enterococcus faecalis V583 GCA_000007785.1
156 Streptomyces cinnamoneus GCA_001885705.1
150 Enterococcus columbae DSM 7374 = ATCC 51263 GCA_000406925.1
150 Melissococcus plutonius S1 GCA_000747585.1
149 Enterococcus asini ATCC 700915 GCA_000407365.1
149 Enterococcus canis NBRC 100695 GCA_001544375.1
149 Enterococcus casseliflavus EC20 GCA_000157355.2
149 Enterococcus dispar ATCC 51266 GCA_000406945.1
149 Enterococcus faecium DO GCA_000174395.2
149 Enterococcus gilvus ATCC BAA-350 GCA_000407545.1
149 Enterococcus haemoperoxidus ATCC BAA-382 GCA_000407165.1
149 Enterococcus hermanniensis GCA_001885945.1
149 Enterococcus hirae ATCC 9790 GCA_000271405.2
149 Enterococcus italicus DSM 15952 GCA_000185365.1
149 Enterococcus malodoratus ATCC 43197 GCA_000407185.1
149 Enterococcus mundtii QU 25 GCA_000504125.1
149 Enterococcus pallens ATCC BAA-351 GCA_000407485.1
149 Enterococcus phoeniculicola ATCC BAA-412 GCA_000407505.1
149 Enterococcus pseudoavium NBRC 100491 GCA_001544295.1
149 Enterococcus rivorum GCA_001742285.1
149 Enterococcus saccharolyticus subsp. saccharolyticus ATCC 43076 GCA_000407285.1
149 Enterococcus sulfureus ATCC 49903 GCA_000407605.1
149 Enterococcus thailandicus GCA_001652875.1
149 Vagococcus lutrae LBD1 GCA_000498295.1
BH
198 Streptococcus mutans UA159 GCA_000007465.2
130 Streptococcus equinus GCA_000964315.1
130 Streptococcus gallolyticus subsp. gallolyticus DSM 16831 GCA_002000985.1
128 Streptococcus salivarius GCA_000785515.1
128 Streptococcus thermophilus JIM 8232 GCA_000253395.1
BI
271 Streptococcus mutans UA159 GCA_000007465.2
169 Streptococcus ferus DSM 20646 GCA_000372425.1
165 Streptococcus ratti FA-1 = DSM 20564 GCA_000286075.1

BJ
254 Escherichia coli IAI39 GCA_000026345.1
254 Escherichia coli O104_3AH4 str. 2011C-3493 GCA_000299455.1
254 Escherichia coli O157_3AH7 str. Sakai GCA_000008865.1
254 Escherichia coli O83_3AH1 str. NRG 857C GCA_000183345.1
254 Escherichia coli UMN026 GCA_000026325.2
254 Escherichia coli str. K-12 substr. MG1655 GCA_000005845.2
254 Shigella dysenteriae Sd197 GCA_000012005.1
254 Shigella flexneri 2a str. 301 GCA_000006925.2
254 Tumebacillus flagellatus GCA_000714935.1
247 Erwinia iniecta GCA_001267535.1
243 Rosenbergiella nectarea GCA_900111105.1
BK
244 Escherichia coli IAI39 GCA_000026345.1
244 Escherichia coli O104_3AH4 str. 2011C-3493 GCA_000299455.1
244 Escherichia coli O157_3AH7 str. Sakai GCA_000008865.1
244 Escherichia coli UMN026 GCA_000026325.2
244 Escherichia coli str. K-12 substr. MG1655 GCA_000005845.2
244 Shigella flexneri 2a str. 301 GCA_000006925.2
244 Tumebacillus flagellatus GCA_000714935.1
238 Erwinia iniecta GCA_001267535.1
235 Shigella dysenteriae Sd197 GCA_000012005.1
BL
200 Deinococcus radiodurans R1 GCA_000008565.1
143 Deinococcus puniceus GCA_001644565.1
128 Deinococcus hopiensis KR-140 GCA_900176165.1
BM
334 Streptococcus mutans UA159 GCA_000007465.2
244 Streptococcus ratti FA-1 = DSM 20564 GCA_000286075.1
240 Streptococcus sobrinus DSM 20742 = ATCC 33478 GCA_000686605.1
BN
144 Rhodobacter sphaeroides 2.4.1 GCA_000012905.2
129 Rhodobacter sphaeroides ATCC 17025 GCA_000016405.1
114 Defluviimonas alba GCA_001620265.1
BO
222 Escherichia coli IAI39 GCA_000026345.1
222 Escherichia coli O104_3AH4 str. 2011C-3493 GCA_000299455.1
222 Escherichia coli O157_3AH7 str. Sakai GCA_000008865.1
222 Escherichia coli O83_3AH1 str. NRG 857C GCA_000183345.1
222 Escherichia coli UMN026 GCA_000026325.2
222 Escherichia coli str. K-12 substr. MG1655 GCA_000005845.2
222 Shigella flexneri 2a str. 301 GCA_000006925.2
222 Tumebacillus flagellatus GCA_000714935.1
208 Shigella dysenteriae Sd197 GCA_000012005.1
203 Erwinia iniecta GCA_001267535.1
BP
236 Streptococcus mutans UA159 GCA_000007465.2
163 Streptococcus massiliensis DSM 18628 GCA_000380065.1
160 Streptococcus ferus DSM 20646 GCA_000372425.1
BQ
301 Clostridium beijerinckii GCA_000833105.2
301 Clostridium saccharoperbutylacetonicum N1-4_28HMT_29 GCA_000340885.1
294 Clostridium puniceum GCA_002006345.1
286 Clostridium saccharobutylicum DSM 13864 GCA_000473995.1
BR
224 Clostridium beijerinckii GCA_000833105.2
224 Clostridium saccharoperbutylacetonicum N1-4_28HMT_29 GCA_000340885.1
217 Clostridium puniceum GCA_002006345.1
209 Clostridium saccharobutylicum DSM 13864 GCA_000473995.1
BS
166 Rhodobacter sphaeroides 2.4.1 GCA_000012905.2
145 Defluviimonas alba GCA_001620265.1
131 Aliiroseovarius crassostreae GCA_001307765.1
131 Aliiroseovarius sediminilitoris GCA_900109955.1
131 Ketogulonicigenium robustum GCA_002117445.1
131 Ketogulonicigenium vulgare WSH-001 GCA_000223375.1
131 Maribius pelagius GCA_900110115.1
131 Marivita cryptomonadis GCA_002115725.1
131 Oceanicola granulosus HTCC2516 GCA_000153305.1
131 Pelagicola litorisediminis GCA_900172295.1
131 Roseovarius lutimaris GCA_900115165.1
131 Sagittula stellata E-37 GCA_000169415.1
131 Sedimentitalea nanhaiensis DSM 24252 GCA_000473225.1
131 Sulfitobacter guttiformis KCTC 32187 GCA_000622425.1
131 Sulfitobacter pseudonitzschiae GCA_002222635.1
131 Thioclava dalianensis GCA_000715505.1

131 Thioclava indica GCA_000714545.1
BT
256 Bacillus anthracis str. Ames GCA_000007845.1
256 Bacillus anthracis str. Sterne GCA_000008165.1
256 Bacillus cereus ATCC 14579 GCA_000007825.1
256 Bacillus mycoides GCA_000832605.1
256 Bacillus pseudomycoides DSM 12442 GCA_000161455.1
256 Bacillus thuringiensis YBT-1518 GCA_000497525.2
256 _5BBacillus thuringiensis_5D serovar konkukian str. 97-27 GCA_000008505.1
236 Bacillus manliponensis GCA_000712595.1
210 Bacillus aquimaris TF-12 GCA_001648555.1
BU
122 Bifidobacterium adolescentis ATCC 15703 GCA_000010425.1
122 Bifidobacterium callitrichos DSM 23973 GCA_000741175.1
122 Bifidobacterium coryneforme GCA_000737865.1
111 Bifidobacterium minimum GCA_000741645.1
111 Bifidobacterium mongoliense DSM 21395 GCA_000741285.1
109 Bifidobacterium asteroides PRL2011 GCA_000304215.1
109 Bifidobacterium gallicum DSM 20093 = LMG 11596 GCA_000741205.1
109 Haematomicrobium sanguinis DSM 21259 GCA_000688395.1
BV
146 Rhodobacter sphaeroides 2.4.1 GCA_000012905.2
126 Rhodobacter sphaeroides ATCC 17025 GCA_000016405.1
112 Gemmobacter aquatilis GCA_900110025.1
112 Pseudorhodobacter psychrotolerans GCA_001294535.1
BW
246 Bacillus anthracis str. Ames GCA_000007845.1
246 Bacillus anthracis str. Sterne GCA_000008165.1
246 Bacillus cereus ATCC 14579 GCA_000007825.1
246 Bacillus mycoides GCA_000832605.1
246 Bacillus pseudomycoides DSM 12442 GCA_000161455.1
246 Bacillus thuringiensis YBT-1518 GCA_000497525.2
246 _5BBacillus thuringiensis_5D serovar konkukian str. 97-27 GCA_000008505.1
192 Bacillus megaterium NBRC 15308 = ATCC 14581 GCA_000832985.1
191 Bacillus flexus GCA_002024265.1
BX
160 Streptococcus mutans UA159 GCA_000007465.2
115 Streptococcus sobrinus DSM 20742 = ATCC 33478 GCA_000686605.1
113 Streptococcus macacae NCTC 11558 GCA_000187995.3
BY
224 Clostridium beijerinckii GCA_000833105.2
224 Clostridium saccharoperbutylacetonicum N1-4_28HMT_29 GCA_000340885.1
217 Clostridium puniceum GCA_002006345.1
216 Clostridium butyricum GCA_001456065.2
BZ
207 Deinococcus radiodurans R1 GCA_000008565.1
160 Deinococcus hopiensis KR-140 GCA_900176165.1
157 Deinococcus deserti VCD115 GCA_000020685.1
C0
103 Escherichia coli IAI39 GCA_000026345.1
103 Escherichia coli O104_3AH4 str. 2011C-3493 GCA_000299455.1
103 Escherichia coli O157_3AH7 str. Sakai GCA_000008865.1
103 Escherichia coli O83_3AH1 str. NRG 857C GCA_000183345.1
103 Escherichia coli UMN026 GCA_000026325.2
103 Escherichia coli str. K-12 substr. MG1655 GCA_000005845.2
103 Shigella dysenteriae Sd197 GCA_000012005.1
103 Shigella flexneri 2a str. 301 GCA_000006925.2
103 Tumebacillus flagellatus GCA_000714935.1
90 Bacillus humi GCA_001439915.1
90 Serratia symbiotica GCA_000821185.1
76 Aeromonas eucrenophila GCA_000819865.1
76 Aeromonas fluvialis GCA_000819885.1
76 Aeromonas hydrophila subsp. hydrophila ATCC 7966 GCA_000014805.1
76 Aeromonas salmonicida subsp. salmonicida A449 GCA_000196395.1
76 Aeromonas schubertii GCA_001447335.1
76 Aeromonas simiae GCA_000820125.1
76 Buttiauxella ferragutiae ATCC 51602 GCA_001654915.1
76 Cedecea neteri GCA_000757825.1
76 Citrobacter freundii CFNIH1 GCA_000648515.1
76 Cronobacter sakazakii GCA_000982825.1
76 Edwardsiella anguillarum ET080813 GCA_000264765.2
76 Enterobacter cloacae subsp. cloacae ATCC 13047 GCA_000025565.1
76 Enterobacter hormaechei subsp. steigerwaltii GCA_001729725.1
76 Enterobacter kobei GCA_900185885.1
76 Erwinia amylovora CFBP1430 GCA_000091565.1
76 Erwinia billingiae Eb661 GCA_000196615.1

76 Erwinia gerundensis GCA_001517405.1
76 Erwinia iniecta GCA_001267535.1
76 Erwinia persicina NBRC 102418 GCA_001571305.1
76 Erwinia toletana DAPP-PG 735 GCA_000336255.1
76 Izhakiella capsodis GCA_900115045.1
76 Klebsiella aerogenes KCTC 2190 GCA_000215745.1
76 Klebsiella oxytoca GCA_001022195.1
76 Klebsiella pneumoniae subsp. pneumoniae HS11286 GCA_000240185.2
76 Kluyvera ascorbata ATCC 33433 GCA_000735365.1
76 Kluyvera cryocrescens NBRC 102467 GCA_001571285.1
76 Kosakonia cowanii GCA_001975225.1
76 Kosakonia sacchari SP1 GCA_000300455.4
76 Mangrovibacter phragmitis GCA_001655675.1
76 Nissabacter archeti GCA_900130115.1
76 Obesumbacterium proteus GCA_001586165.1
76 Pantoea agglomerans GCA_001709315.1
76 Pantoea alhagi GCA_002101395.1
76 Pantoea ananatis LMG 20103 GCA_000025405.2
76 Pantoea dispersa EGD-AAK13 GCA_000465555.2
76 Pantoea rwandensis GCA_000759475.1
76 Pantoea septica GCA_002095575.1
76 Plautia stali symbiont GCA_000180175.2
76 Plesiomonas shigelloides GCA_900087055.1
76 Pluralibacter gergoviae GCA_000757785.1
76 Pseudescherichia vulneris NBRC 102420 GCA_000759795.1
76 Rahnella aquatilis HX2 GCA_000255535.1
76 Rosenbergiella nectarea GCA_900111105.1
76 Rouxiella chamberiensis GCA_000951135.1
76 Salmonella enterica subsp. enterica serovar Typhi str. CT18 GCA_000195995.1
76 Serratia liquefaciens ATCC 27592 GCA_000422085.1
76 Serratia marcescens subsp. marcescens Db11 GCA_000513215.1
76 Serratia rubidaea GCA_001572725.1
76 Shimwellia blattae DSM 4481 = NBRC 105725 GCA_000262305.1
76 Siccibacter turicensis LMG 23730 GCA_000463155.2
76 Tatumella citrea GCA_002163585.1
76 Tatumella saanichensis GCA_000439375.1
76 Trabulsiella odontotermitis GCA_001297765.1
76 Yersinia enterocolitica subsp. enterocolitica 8081 GCA_000009345.1
76 Yersinia pestis CO92 GCA_000009065.1
76 Yersinia ruckeri GCA_000964565.1
76 Yokenella regensburgei ATCC 49455 GCA_000735455.1
76 _5BEnterobacter_5D lignolyticus SCF1 GCA_000164865.1
C1
165 Bacillus anthracis str. Ames GCA_000007845.1
165 Bacillus anthracis str. Sterne GCA_000008165.1
165 Bacillus cereus ATCC 14579 GCA_000007825.1
165 Bacillus mycoides GCA_000832605.1
165 Bacillus pseudomycoides DSM 12442 GCA_000161455.1
165 Bacillus thuringiensis YBT-1518 GCA_000497525.2
165 _5BBacillus thuringiensis_5D serovar konkukian str. 97-27 GCA_000008505.1
135 Bacillus coahuilensis m4-4 GCA_000171615.1
135 Bacillus marisflavi GCA_001274775.1
135 Oceanobacillus caeni GCA_001298135.1
134 Bacillus halmapalus GCA_002019665.1
134 Bacillus horikoshii GCA_002157855.1
134 Bacillus horneckiae GCA_001636335.1
134 Enterococcus aquimarinus GCA_001885765.1
134 Enterococcus asini ATCC 700915 GCA_000407365.1
134 Enterococcus canis NBRC 100695 GCA_001544375.1
134 Enterococcus casseliflavus EC20 GCA_000157355.2
134 Enterococcus dispar ATCC 51266 GCA_000406945.1
134 Enterococcus faecalis V583 GCA_000007785.1
134 Enterococcus faecium DO GCA_000174395.2
134 Enterococcus gilvus ATCC BAA-350 GCA_000407545.1
134 Enterococcus haemoperoxidus ATCC BAA-382 GCA_000407165.1
134 Enterococcus hermanniensis GCA_001885945.1
134 Enterococcus hirae ATCC 9790 GCA_000271405.2
134 Enterococcus italicus DSM 15952 GCA_000185365.1
134 Enterococcus malodoratus ATCC 43197 GCA_000407185.1
134 Enterococcus massiliensis GCA_001050095.1
134 Enterococcus mundtii QU 25 GCA_000504125.1
134 Enterococcus pallens ATCC BAA-351 GCA_000407485.1
134 Enterococcus phoeniculicola ATCC BAA-412 GCA_000407505.1
134 Enterococcus pseudoavium NBRC 100491 GCA_001544295.1
134 Enterococcus rivorum GCA_001742285.1
134 Enterococcus saccharolyticus subsp. saccharolyticus ATCC 43076 GCA_000407285.1

134 Enterococcus sulfureus ATCC 49903 GCA_000407605.1
134 Enterococcus thailandicus GCA_001652875.1
134 Melissococcus plutonius S1 GCA_000747585.1
134 Streptomyces cinnamoneus GCA_001885705.1
134 Vagococcus fluvialis bH819 GCA_900163795.1
134 Vagococcus lutrae LBD1 GCA_000498295.1
134 Vagococcus penaei GCA_001998885.1
134 Vagococcus teuberi GCA_001870205.1
C2
210 Bacillus anthracis str. Ames GCA_000007845.1
210 Bacillus anthracis str. Sterne GCA_000008165.1
210 Bacillus cereus ATCC 14579 GCA_000007825.1
210 Bacillus mycoides GCA_000832605.1
210 Bacillus pseudomycoides DSM 12442 GCA_000161455.1
210 Bacillus thuringiensis YBT-1518 GCA_000497525.2
210 _5BBacillus thuringiensis_5D serovar konkukian str. 97-27 GCA_000008505.1
195 Bacillus infantis NRRL B-14911 GCA_000473245.1
193 Bacillus aquimaris TF-12 GCA_001648555.1
193 Bacillus eiseniae GCA_001636325.1
193 Bacillus vietnamensis NBRC 101237 GCA_001591825.1
C3
203 Bacillus anthracis str. Ames GCA_000007845.1
203 Bacillus anthracis str. Sterne GCA_000008165.1
203 Bacillus cereus ATCC 14579 GCA_000007825.1
203 Bacillus mycoides GCA_000832605.1
203 Bacillus pseudomycoides DSM 12442 GCA_000161455.1
203 Bacillus thuringiensis YBT-1518 GCA_000497525.2
203 _5BBacillus thuringiensis_5D serovar konkukian str. 97-27 GCA_000008505.1
166 Bacillus coahuilensis m4-4 GCA_000171615.1
166 Bacillus manliponensis GCA_000712595.1
166 Bacillus marisflavi GCA_001274775.1
159 Enterococcus hirae ATCC 9790 GCA_000271405.2
C4
255 Clostridium beijerinckii GCA_000833105.2
255 Clostridium puniceum GCA_002006345.1
255 Clostridium saccharoperbutylacetonicum N1-4_28HMT_29 GCA_000340885.1
246 Clostridium saccharobutylicum DSM 13864 GCA_000473995.1
235 Clostridium butyricum GCA_001456065.2
C5
161 Clostridium beijerinckii GCA_000833105.2
161 Clostridium puniceum GCA_002006345.1
161 Clostridium saccharobutylicum DSM 13864 GCA_000473995.1
161 Clostridium saccharoperbutylacetonicum N1-4_28HMT_29 GCA_000340885.1
158 Clostridium neonatale GCA_001458595.1
150 Clostridium chromiireducens GCA_002029255.1
150 Clostridium taeniosporum GCA_001735765.1
C6
252 Escherichia coli IAI39 GCA_000026345.1
252 Escherichia coli O104_3AH4 str. 2011C-3493 GCA_000299455.1
252 Escherichia coli O157_3AH7 str. Sakai GCA_000008865.1
252 Escherichia coli O83_3AH1 str. NRG 857C GCA_000183345.1
252 Escherichia coli UMN026 GCA_000026325.2
252 Escherichia coli str. K-12 substr. MG1655 GCA_000005845.2
252 Shigella dysenteriae Sd197 GCA_000012005.1
252 Shigella flexneri 2a str. 301 GCA_000006925.2
252 Tumebacillus flagellatus GCA_000714935.1
214 Cronobacter sakazakii GCA_000982825.1
214 Enterobacter cloacae subsp. cloacae ATCC 13047 GCA_000025565.1
214 Enterobacter hormaechei subsp. steigerwaltii GCA_001729725.1
214 Enterobacter kobei GCA_900185885.1
214 Erwinia amylovora CFBP1430 GCA_000091565.1
214 Erwinia iniecta GCA_001267535.1
214 Erwinia persicina NBRC 102418 GCA_001571305.1
214 Erwinia toletana DAPP-PG 735 GCA_000336255.1
214 Klebsiella oxytoca GCA_001022195.1
214 Kosakonia cowanii GCA_001975225.1
214 Pantoea agglomerans GCA_001709315.1
214 Pantoea alhagi GCA_002101395.1
214 Pantoea ananatis LMG 20103 GCA_000025405.2
214 Pantoea dispersa EGD-AAK13 GCA_000465555.2
214 Pantoea rwandensis GCA_000759475.1
214 Pantoea septica GCA_002095575.1
214 Pseudescherichia vulneris NBRC 102420 GCA_000759795.1
214 Salmonella enterica subsp. enterica serovar Typhi str. CT18 GCA_000195995.1
214 Salmonella enterica subsp. enterica serovar Typhimurium str. LT2 GCA_000006945.2
214 Shimwellia blattae DSM 4481 = NBRC 105725 GCA_000262305.1

214 Trabulsiella odontotermitis GCA_001297765.1
211 Erwinia billingiae Eb661 GCA_000196615.1
211 Erwinia gerundensis GCA_001517405.1
211 Pluralibacter gergoviae GCA_000757785.1
C7
252 Escherichia coli IAI39 GCA_000026345.1
252 Escherichia coli O104_3AH4 str. 2011C-3493 GCA_000299455.1
252 Escherichia coli O157_3AH7 str. Sakai GCA_000008865.1
252 Escherichia coli O83_3AH1 str. NRG 857C GCA_000183345.1
252 Escherichia coli UMN026 GCA_000026325.2
252 Escherichia coli str. K-12 substr. MG1655 GCA_000005845.2
252 Shigella flexneri 2a str. 301 GCA_000006925.2
252 Tumebacillus flagellatus GCA_000714935.1
243 Shigella dysenteriae Sd197 GCA_000012005.1
222 Erwinia iniecta GCA_001267535.1
C8
249 Bifidobacterium adolescentis ATCC 15703 GCA_000010425.1
225 Bifidobacterium dentium JCM 1195 = DSM 20436 GCA_001042595.1
207 Bifidobacterium breve DSM 20213 = JCM 1192 GCA_001025175.1
C9
168 Clostridium puniceum GCA_002006345.1
163 Clostridium taeniosporum GCA_001735765.1
161 Clostridium beijerinckii GCA_000833105.2
161 Clostridium saccharobutylicum DSM 13864 GCA_000473995.1
161 Clostridium saccharoperbutylacetonicum N1-4_28HMT_29 GCA_000340885.1
Ca
222 Deinococcus radiodurans R1 GCA_000008565.1
153 Deinococcus gobiensis I-0 GCA_000252445.1
148 Deinococcus puniceus GCA_001644565.1
Cb
198 Escherichia coli IAI39 GCA_000026345.1
198 Escherichia coli O104_3AH4 str. 2011C-3493 GCA_000299455.1
198 Escherichia coli O157_3AH7 str. Sakai GCA_000008865.1
198 Escherichia coli O83_3AH1 str. NRG 857C GCA_000183345.1
198 Escherichia coli UMN026 GCA_000026325.2
198 Escherichia coli str. K-12 substr. MG1655 GCA_000005845.2
198 Shigella dysenteriae Sd197 GCA_000012005.1
198 Shigella flexneri 2a str. 301 GCA_000006925.2
198 Tumebacillus flagellatus GCA_000714935.1
169 Pantoea agglomerans GCA_001709315.1
168 Cronobacter sakazakii GCA_000982825.1
168 Erwinia iniecta GCA_001267535.1
168 Erwinia toletana DAPP-PG 735 GCA_000336255.1
168 Pantoea ananatis LMG 20103 GCA_000025405.2
168 Pantoea dispersa EGD-AAK13 GCA_000465555.2
168 Plautia stali symbiont GCA_000180175.2
168 Shimwellia blattae DSM 4481 = NBRC 105725 GCA_000262305.1
Cc
153 Streptococcus mutans UA159 GCA_000007465.2
92 Streptococcus massiliensis DSM 18628 GCA_000380065.1
90 Streptococcus ferus DSM 20646 GCA_000372425.1
Cd
295 Bifidobacterium adolescentis ATCC 15703 GCA_000010425.1
263 Bifidobacterium tsurumiense GCA_000741765.1
262 Bifidobacterium thermophilum GCA_000741495.1
Ce
183 Clostridium beijerinckii GCA_000833105.2
183 Clostridium saccharobutylicum DSM 13864 GCA_000473995.1
183 Clostridium saccharoperbutylacetonicum N1-4_28HMT_29 GCA_000340885.1
171 Clostridium butyricum GCA_001456065.2
169 Clostridium chromiireducens GCA_002029255.1
169 Clostridium puniceum GCA_002006345.1
Cf
172 Bacillus thuringiensis YBT-1518 GCA_000497525.2
157 Bacillus anthracis str. Ames GCA_000007845.1
157 Bacillus anthracis str. Sterne GCA_000008165.1
157 Bacillus cereus ATCC 14579 GCA_000007825.1
157 Bacillus mycoides GCA_000832605.1
157 Bacillus pseudomycoides DSM 12442 GCA_000161455.1
157 _5BBacillus thuringiensis_5D serovar konkukian str. 97-27 GCA_000008505.1
139 Lysinibacillus fusiformis GCA_000724775.3
Cg
227 Bifidobacterium adolescentis ATCC 15703 GCA_000010425.1
209 Bifidobacterium callitrichos DSM 23973 GCA_000741175.1
200 Bifidobacterium angulatum DSM 20098 = JCM 7096 GCA_001025155.1
Ch

97 Deinococcus radiodurans R1 GCA_000008565.1
83 Enterococcus hirae ATCC 9790 GCA_000271405.2
80 Enterococcus dispar ATCC 51266 GCA_000406945.1
Ci
139 Clostridium beijerinckii GCA_000833105.2
139 Clostridium puniceum GCA_002006345.1
139 Clostridium saccharobutylicum DSM 13864 GCA_000473995.1
139 Clostridium saccharoperbutylacetonicum N1-4_28HMT_29 GCA_000340885.1
128 Clostridium neonatale GCA_001458595.1
118 Clostridium butyricum GCA_001456065.2
Cj
280 Deinococcus radiodurans R1 GCA_000008565.1
192 Deinococcus gobiensis I-0 GCA_000252445.1
190 Deinococcus deserti VCD115 GCA_000020685.1
Ck
152 Defluviimonas alba GCA_001620265.1
152 Rhodobacter sphaeroides 2.4.1 GCA_000012905.2
147 Pseudorhodobacter ferrugineus DSM 5888 GCA_000420745.1
147 Pseudorhodobacter psychrotolerans GCA_001294535.1
147 Pseudorhodobacter wandonensis GCA_001202035.1
131 Gemmobacter megaterium GCA_900156815.1
131 Rhodobacter sphaeroides ATCC 17025 GCA_000016405.1
Cl
213 Clostridium beijerinckii GCA_000833105.2
213 Clostridium saccharobutylicum DSM 13864 GCA_000473995.1
213 Clostridium saccharoperbutylacetonicum N1-4_28HMT_29 GCA_000340885.1
209 Clostridium puniceum GCA_002006345.1
190 Clostridium chromiireducens GCA_002029255.1
Cm
118 Escherichia coli IAI39 GCA_000026345.1
118 Escherichia coli O104_3AH4 str. 2011C-3493 GCA_000299455.1
118 Escherichia coli O157_3AH7 str. Sakai GCA_000008865.1
118 Escherichia coli O83_3AH1 str. NRG 857C GCA_000183345.1
118 Escherichia coli UMN026 GCA_000026325.2
118 Escherichia coli str. K-12 substr. MG1655 GCA_000005845.2
118 Shigella dysenteriae Sd197 GCA_000012005.1
118 Shigella flexneri 2a str. 301 GCA_000006925.2
118 Tumebacillus flagellatus GCA_000714935.1
104 Erwinia iniecta GCA_001267535.1
91 Cedecea neteri GCA_000757825.1
Cn
259 Staphylococcus capitis subsp. capitis GCA_001028645.1
259 Staphylococcus epidermidis ATCC 12228 GCA_000007645.1
252 Staphylococcus aureus subsp. aureus NCTC 8325 GCA_000013425.1
252 Staphylococcus haemolyticus JCSC1435 GCA_000009865.1
252 Staphylococcus hominis subsp. hominis C80 GCA_000183685.1
252 Staphylococcus simiae CCM 7213 GCA_000235645.2
246 Megasphaera cerevisiae DSM 20462 GCA_001045675.1
246 Staphylococcus warneri SG1 GCA_000332735.1
Co
227 Bacillus anthracis str. Ames GCA_000007845.1
227 Bacillus anthracis str. Sterne GCA_000008165.1
227 Bacillus cereus ATCC 14579 GCA_000007825.1
227 Bacillus mycoides GCA_000832605.1
227 Bacillus pseudomycoides DSM 12442 GCA_000161455.1
227 Bacillus thuringiensis YBT-1518 GCA_000497525.2
227 _5BBacillus thuringiensis_5D serovar konkukian str. 97-27 GCA_000008505.1
166 Massilibacterium senegalense GCA_001375675.1
164 Bacillus coahuilensis m4-4 GCA_000171615.1
Cp
272 Enterococcus faecalis V583 GCA_000007785.1
272 Streptomyces cinnamoneus GCA_001885705.1
250 Enterococcus canis NBRC 100695 GCA_001544375.1
250 Enterococcus dispar ATCC 51266 GCA_000406945.1
250 Enterococcus faecium DO GCA_000174395.2
250 Enterococcus hirae ATCC 9790 GCA_000271405.2
250 Enterococcus mundtii QU 25 GCA_000504125.1
250 Enterococcus rivorum GCA_001742285.1
245 Enterococcus haemoperoxidus ATCC BAA-382 GCA_000407165.1
245 Enterococcus phoeniculicola ATCC BAA-412 GCA_000407505.1
245 Enterococcus thailandicus GCA_001652875.1
Cq
259 Lactobacillus gasseri ATCC 33323 = JCM 1131 GCA_000014425.1
239 Lactobacillus hominis DSM 23910 = CRBIP 24.179 GCA_000296835.1
205 Lactobacillus crispatus ST1 GCA_000091765.1
Cr

185 Deinococcus radiodurans R1 GCA_000008565.1
136 Deinococcus deserti VCD115 GCA_000020685.1
127 Deinococcus gobiensis I-0 GCA_000252445.1
Cs
162 Clostridium beijerinckii GCA_000833105.2
162 Clostridium puniceum GCA_002006345.1
162 Clostridium saccharobutylicum DSM 13864 GCA_000473995.1
162 Clostridium saccharoperbutylacetonicum N1-4_28HMT_29 GCA_000340885.1
157 Clostridium butyricum GCA_001456065.2
147 Clostridium botulinum B str. Eklund 17B _28NRP_29 GCA_000020165.1
147 Clostridium chromiireducens GCA_002029255.1
147 Clostridium taeniosporum GCA_001735765.1
Ct
128 Rhodobacter sphaeroides 2.4.1 GCA_000012905.2
112 Rhodobacter sphaeroides ATCC 17025 GCA_000016405.1
101 Gemmobacter aquatilis GCA_900110025.1
101 Phaeobacter porticola GCA_001888185.1
101 Pseudophaeobacter arcticus DSM 23566 GCA_000473205.1
101 Sedimentitalea nanhaiensis DSM 24252 GCA_000473225.1
101 Sulfitobacter guttiformis KCTC 32187 GCA_000622425.1
101 Sulfitobacter pseudonitzschiae GCA_002222635.1
101 Tropicibacter litoreus R37 GCA_900172225.1
Cu
187 Clostridium beijerinckii GCA_000833105.2
187 Clostridium saccharoperbutylacetonicum N1-4_28HMT_29 GCA_000340885.1
180 Clostridium puniceum GCA_002006345.1
173 Clostridium saccharobutylicum DSM 13864 GCA_000473995.1
Cv
159 Clostridium beijerinckii GCA_000833105.2
159 Clostridium butyricum GCA_001456065.2
159 Clostridium puniceum GCA_002006345.1
159 Clostridium saccharobutylicum DSM 13864 GCA_000473995.1
159 Clostridium saccharoperbutylacetonicum N1-4_28HMT_29 GCA_000340885.1
144 Clostridium chromiireducens GCA_002029255.1
138 Clostridium neonatale GCA_001458595.1
Cw
207 Deinococcus radiodurans R1 GCA_000008565.1
155 Deinococcus puniceus GCA_001644565.1
154 Deinococcus gobiensis I-0 GCA_000252445.1
Cx
265 Streptococcus mutans UA159 GCA_000007465.2
187 Streptococcus gordonii str. Challis substr. CH1 GCA_000017005.1
173 Streptococcus cristatus AS 1.3089 GCA_000385925.1
173 Streptococcus mitis B6 GCA_000027165.1
173 Streptococcus pneumoniae R6 GCA_000007045.1
Cy
241 Rhodobacter sphaeroides 2.4.1 GCA_000012905.2
220 Gemmobacter megaterium GCA_900156815.1
215 Gemmobacter aquatilis GCA_900110025.1
Cz
230 Clostridium butyricum GCA_001456065.2
222 Clostridium saccharobutylicum DSM 13864 GCA_000473995.1
220 Clostridium beijerinckii GCA_000833105.2
220 Clostridium saccharoperbutylacetonicum N1-4_28HMT_29 GCA_000340885.1
CA
173 Lactobacillus gasseri ATCC 33323 = JCM 1131 GCA_000014425.1
151 Lactobacillus hominis DSM 23910 = CRBIP 24.179 GCA_000296835.1
126 Lactobacillus harbinensis DSM 16991 GCA_000425885.1
126 Lactobacillus perolens DSM 12744 GCA_001435585.1
126 Lactobacillus shenzhenensis LY-73 GCA_000469325.1
CB
259 Deinococcus radiodurans R1 GCA_000008565.1
182 Deinococcus deserti VCD115 GCA_000020685.1
174 Deinococcus hopiensis KR-140 GCA_900176165.1
CC
137 Deinococcus radiodurans R1 GCA_000008565.1
94 Deinococcus gobiensis I-0 GCA_000252445.1
82 Deinococcus proteolyticus MRP GCA_000190555.1
CD
162 Streptococcus mutans UA159 GCA_000007465.2
114 Streptococcus ferus DSM 20646 GCA_000372425.1
108 Streptococcus macacae NCTC 11558 GCA_000187995.3
CE
253 Clostridium beijerinckii GCA_000833105.2
253 Clostridium saccharoperbutylacetonicum N1-4_28HMT_29 GCA_000340885.1
246 Clostridium puniceum GCA_002006345.1

244 Clostridium saccharobutylicum DSM 13864 GCA_000473995.1
CF
292 Escherichia coli IAI39 GCA_000026345.1
292 Escherichia coli O104_3AH4 str. 2011C-3493 GCA_000299455.1
292 Escherichia coli O157_3AH7 str. Sakai GCA_000008865.1
292 Escherichia coli O83_3AH1 str. NRG 857C GCA_000183345.1
292 Escherichia coli UMN026 GCA_000026325.2
292 Escherichia coli str. K-12 substr. MG1655 GCA_000005845.2
292 Shigella flexneri 2a str. 301 GCA_000006925.2
292 Tumebacillus flagellatus GCA_000714935.1
283 Shigella dysenteriae Sd197 GCA_000012005.1
260 Erwinia iniecta GCA_001267535.1
CG
312 Enterococcus faecalis V583 GCA_000007785.1
312 Streptomyces cinnamoneus GCA_001885705.1
295 Enterococcus canis NBRC 100695 GCA_001544375.1
295 Enterococcus faecium DO GCA_000174395.2
295 Enterococcus haemoperoxidus ATCC BAA-382 GCA_000407165.1
295 Enterococcus hirae ATCC 9790 GCA_000271405.2
295 Enterococcus mundtii QU 25 GCA_000504125.1
295 Enterococcus phoeniculicola ATCC BAA-412 GCA_000407505.1
295 Enterococcus rivorum GCA_001742285.1
295 Enterococcus thailandicus GCA_001652875.1
294 Enterococcus dispar ATCC 51266 GCA_000406945.1
CH
203 Deinococcus radiodurans R1 GCA_000008565.1
124 Deinococcus puniceus GCA_001644565.1
117 Deinococcus deserti VCD115 GCA_000020685.1
117 Deinococcus gobiensis I-0 GCA_000252445.1
CI
255 Enterococcus faecalis V583 GCA_000007785.1
255 Streptomyces cinnamoneus GCA_001885705.1
225 Enterococcus canis NBRC 100695 GCA_001544375.1
225 Enterococcus faecium DO GCA_000174395.2
225 Enterococcus hirae ATCC 9790 GCA_000271405.2
225 Enterococcus mundtii QU 25 GCA_000504125.1
225 Enterococcus rivorum GCA_001742285.1
224 Enterococcus casseliflavus EC20 GCA_000157355.2
224 Enterococcus dispar ATCC 51266 GCA_000406945.1
224 Enterococcus massiliensis GCA_001050095.1
224 Enterococcus saccharolyticus subsp. saccharolyticus ATCC 43076 GCA_000407285.1
CJ
145 Lactobacillus gasseri ATCC 33323 = JCM 1131 GCA_000014425.1
145 Lactobacillus hominis DSM 23910 = CRBIP 24.179 GCA_000296835.1
114 Lactobacillus iners DSM 13335 GCA_000160875.1
112 Lactobacillus amylovorus GCA_000191545.1
112 Lactobacillus crispatus ST1 GCA_000091765.1
112 Lactobacillus delbrueckii subsp. bulgaricus ATCC 11842 = JCM 1002 GCA_000056065.1
112 Lactobacillus kalixensis DSM 16043 GCA_001434335.1
112 Lactobacillus kefiranofaciens ZW3 GCA_000214785.1
CK
249 Bacillus anthracis str. Ames GCA_000007845.1
249 Bacillus anthracis str. Sterne GCA_000008165.1
249 Bacillus cereus ATCC 14579 GCA_000007825.1
249 Bacillus mycoides GCA_000832605.1
249 Bacillus pseudomycoides DSM 12442 GCA_000161455.1
249 Bacillus thuringiensis YBT-1518 GCA_000497525.2
249 _5BBacillus thuringiensis_5D serovar konkukian str. 97-27 GCA_000008505.1
199 Bacillus manliponensis GCA_000712595.1
197 Viridibacillus arvi GCA_001274945.1
CL
184 Rhodobacter sphaeroides 2.4.1 GCA_000012905.2
169 Rhodobacter sphaeroides ATCC 17025 GCA_000016405.1
168 Gemmobacter megaterium GCA_900156815.1
CM
249 Bacillus anthracis str. Ames GCA_000007845.1
249 Bacillus anthracis str. Sterne GCA_000008165.1
249 Bacillus cereus ATCC 14579 GCA_000007825.1
249 Bacillus mycoides GCA_000832605.1
249 Bacillus pseudomycoides DSM 12442 GCA_000161455.1
249 Bacillus thuringiensis YBT-1518 GCA_000497525.2
249 _5BBacillus thuringiensis_5D serovar konkukian str. 97-27 GCA_000008505.1
216 Bacillus coahuilensis m4-4 GCA_000171615.1
207 Bacillus manliponensis GCA_000712595.1
207 Bacillus marisflavi GCA_001274775.1
CN

222 Deinococcus radiodurans R1 GCA_000008565.1
133 Deinococcus gobiensis I-0 GCA_000252445.1
128 Deinococcus proteolyticus MRP GCA_000190555.1
CO
190 Clostridium beijerinckii GCA_000833105.2
190 Clostridium saccharobutylicum DSM 13864 GCA_000473995.1
190 Clostridium saccharoperbutylacetonicum N1-4_28HMT_29 GCA_000340885.1
183 Clostridium puniceum GCA_002006345.1
172 Clostridium butyricum GCA_001456065.2
172 Clostridium chromiireducens GCA_002029255.1
CP
281 Clostridium beijerinckii GCA_000833105.2
281 Clostridium saccharobutylicum DSM 13864 GCA_000473995.1
281 Clostridium saccharoperbutylacetonicum N1-4_28HMT_29 GCA_000340885.1
267 Clostridium chromiireducens GCA_002029255.1
267 Clostridium puniceum GCA_002006345.1
251 Clostridium butyricum GCA_001456065.2
CQ
205 Clostridium beijerinckii GCA_000833105.2
205 Clostridium puniceum GCA_002006345.1
205 Clostridium saccharoperbutylacetonicum N1-4_28HMT_29 GCA_000340885.1
197 Clostridium butyricum GCA_001456065.2
190 Clostridium saccharobutylicum DSM 13864 GCA_000473995.1
CR
237 Clostridium beijerinckii GCA_000833105.2
237 Clostridium puniceum GCA_002006345.1
237 Clostridium saccharobutylicum DSM 13864 GCA_000473995.1
237 Clostridium saccharoperbutylacetonicum N1-4_28HMT_29 GCA_000340885.1
230 Clostridium butyricum GCA_001456065.2
224 Clostridium taeniosporum GCA_001735765.1
CS
144 Clostridium beijerinckii GCA_000833105.2
144 Clostridium puniceum GCA_002006345.1
144 Clostridium saccharobutylicum DSM 13864 GCA_000473995.1
144 Clostridium saccharoperbutylacetonicum N1-4_28HMT_29 GCA_000340885.1
129 Clostridium chromiireducens GCA_002029255.1
107 Clostridium taeniosporum GCA_001735765.1
CT
236 Clostridium beijerinckii GCA_000833105.2
236 Clostridium saccharobutylicum DSM 13864 GCA_000473995.1
236 Clostridium saccharoperbutylacetonicum N1-4_28HMT_29 GCA_000340885.1
230 Clostridium puniceum GCA_002006345.1
219 Clostridium chromiireducens GCA_002029255.1
CU
248 Deinococcus radiodurans R1 GCA_000008565.1
170 Deinococcus gobiensis I-0 GCA_000252445.1
144 Deinococcus marmoris DSM 12784 GCA_000701405.1
144 Deinococcus proteolyticus MRP GCA_000190555.1
144 Deinococcus swuensis GCA_000800395.1
CV
260 Escherichia coli IAI39 GCA_000026345.1
260 Escherichia coli O104_3AH4 str. 2011C-3493 GCA_000299455.1
260 Escherichia coli O157_3AH7 str. Sakai GCA_000008865.1
260 Escherichia coli O83_3AH1 str. NRG 857C GCA_000183345.1
260 Escherichia coli UMN026 GCA_000026325.2
260 Escherichia coli str. K-12 substr. MG1655 GCA_000005845.2
260 Shigella flexneri 2a str. 301 GCA_000006925.2
260 Tumebacillus flagellatus GCA_000714935.1
245 Shigella dysenteriae Sd197 GCA_000012005.1
234 Erwinia iniecta GCA_001267535.1
CW
282 Deinococcus radiodurans R1 GCA_000008565.1
161 Deinococcus marmoris DSM 12784 GCA_000701405.1
161 Deinococcus swuensis GCA_000800395.1
159 Deinococcus puniceus GCA_001644565.1
CX
196 Escherichia coli IAI39 GCA_000026345.1
196 Escherichia coli O104_3AH4 str. 2011C-3493 GCA_000299455.1
196 Escherichia coli O157_3AH7 str. Sakai GCA_000008865.1
196 Escherichia coli O83_3AH1 str. NRG 857C GCA_000183345.1
196 Escherichia coli UMN026 GCA_000026325.2
196 Escherichia coli str. K-12 substr. MG1655 GCA_000005845.2
196 Shigella dysenteriae Sd197 GCA_000012005.1
196 Shigella flexneri 2a str. 301 GCA_000006925.2
196 Tumebacillus flagellatus GCA_000714935.1
193 Erwinia amylovora CFBP1430 GCA_000091565.1

193 Pantoea dispersa EGD-AAK13 GCA_000465555.2
193 Plautia stali symbiont GCA_000180175.2
193 Shimwellia blattae DSM 4481 = NBRC 105725 GCA_000262305.1
192 Cronobacter sakazakii GCA_000982825.1
192 Erwinia iniecta GCA_001267535.1
192 Erwinia toletana DAPP-PG 735 GCA_000336255.1
192 Kosakonia cowanii GCA_001975225.1
192 Pantoea ananatis LMG 20103 GCA_000025405.2
192 Pluralibacter gergoviae GCA_000757785.1
CY
219 Megasphaera cerevisiae DSM 20462 GCA_001045675.1
219 Staphylococcus capitis subsp. capitis GCA_001028645.1
219 Staphylococcus epidermidis ATCC 12228 GCA_000007645.1
219 Staphylococcus lugdunensis HKU09-01 GCA_000025085.1
219 Staphylococcus warneri SG1 GCA_000332735.1
210 Staphylococcus pseudintermedius HKU10-03 GCA_000185885.1
209 Staphylococcus hyicus GCA_000816085.1
CZ
254 Lactobacillus gasseri ATCC 33323 = JCM 1131 GCA_000014425.1
246 Lactobacillus hominis DSM 23910 = CRBIP 24.179 GCA_000296835.1
197 Lactobacillus acidophilus NCFM GCA_000011985.1
197 Lactobacillus amylovorus GCA_000191545.1
197 Lactobacillus crispatus ST1 GCA_000091765.1
197 Lactobacillus gallinarum GCA_001314245.2
197 Lactobacillus hamsteri DSM 5661 = JCM 6256 GCA_000615445.1
D0
306 Deinococcus radiodurans R1 GCA_000008565.1
214 Deinococcus gobiensis I-0 GCA_000252445.1
188 Deinococcus deserti VCD115 GCA_000020685.1
D1
290 Staphylococcus epidermidis ATCC 12228 GCA_000007645.1
290 Staphylococcus haemolyticus JCSC1435 GCA_000009865.1
285 Staphylococcus capitis subsp. capitis GCA_001028645.1
279 Staphylococcus hominis subsp. hominis C80 GCA_000183685.1
D2
243 Escherichia coli IAI39 GCA_000026345.1
243 Escherichia coli O104_3AH4 str. 2011C-3493 GCA_000299455.1
243 Escherichia coli O157_3AH7 str. Sakai GCA_000008865.1
243 Escherichia coli O83_3AH1 str. NRG 857C GCA_000183345.1
243 Escherichia coli UMN026 GCA_000026325.2
243 Escherichia coli str. K-12 substr. MG1655 GCA_000005845.2
243 Shigella flexneri 2a str. 301 GCA_000006925.2
243 Tumebacillus flagellatus GCA_000714935.1
239 Erwinia iniecta GCA_001267535.1
233 Cronobacter sakazakii GCA_000982825.1
233 Erwinia toletana DAPP-PG 735 GCA_000336255.1
D3
148 Erwinia iniecta GCA_001267535.1
148 Erwinia toletana DAPP-PG 735 GCA_000336255.1
148 Escherichia coli IAI39 GCA_000026345.1
148 Escherichia coli O104_3AH4 str. 2011C-3493 GCA_000299455.1
148 Escherichia coli O157_3AH7 str. Sakai GCA_000008865.1
148 Escherichia coli O83_3AH1 str. NRG 857C GCA_000183345.1
148 Escherichia coli UMN026 GCA_000026325.2
148 Escherichia coli str. K-12 substr. MG1655 GCA_000005845.2
148 Shigella dysenteriae Sd197 GCA_000012005.1
148 Shigella flexneri 2a str. 301 GCA_000006925.2
148 Tumebacillus flagellatus GCA_000714935.1
144 Cronobacter sakazakii GCA_000982825.1
144 Pantoea agglomerans GCA_001709315.1
144 Pantoea ananatis LMG 20103 GCA_000025405.2
144 Shimwellia blattae DSM 4481 = NBRC 105725 GCA_000262305.1
140 Pantoea dispersa EGD-AAK13 GCA_000465555.2
D4
261 Enterococcus faecalis V583 GCA_000007785.1
261 Streptomyces cinnamoneus GCA_001885705.1
256 Vagococcus lutrae LBD1 GCA_000498295.1
247 Enterococcus dispar ATCC 51266 GCA_000406945.1
D5
148 Streptococcus equinus GCA_000964315.1
148 Streptococcus gallolyticus subsp. gallolyticus DSM 16831 GCA_002000985.1
146 Streptococcus mutans UA159 GCA_000007465.2
143 Lactobacillus bifermentans DSM 20003 GCA_001434575.1
D6
236 Deinococcus radiodurans R1 GCA_000008565.1
167 Deinococcus puniceus GCA_001644565.1

165 Deinococcus deserti VCD115 GCA_000020685.1
D7
221 Deinococcus radiodurans R1 GCA_000008565.1
116 Deinococcus proteolyticus MRP GCA_000190555.1
111 Deinococcus gobiensis I-0 GCA_000252445.1
D8
85 Enterobacter cloacae subsp. cloacae ATCC 13047 GCA_000025565.1
85 Enterobacter hormaechei subsp. steigerwaltii GCA_001729725.1
85 Enterobacter kobei GCA_900185885.1
85 Erwinia persicina NBRC 102418 GCA_001571305.1
85 Escherichia coli IAI39 GCA_000026345.1
85 Escherichia coli O104_3AH4 str. 2011C-3493 GCA_000299455.1
85 Escherichia coli O157_3AH7 str. Sakai GCA_000008865.1
85 Escherichia coli O83_3AH1 str. NRG 857C GCA_000183345.1
85 Escherichia coli UMN026 GCA_000026325.2
85 Escherichia coli str. K-12 substr. MG1655 GCA_000005845.2
85 Klebsiella oxytoca GCA_001022195.1
85 Klebsiella pneumoniae subsp. pneumoniae HS11286 GCA_000240185.2
85 Pseudescherichia vulneris NBRC 102420 GCA_000759795.1
85 Salmonella enterica subsp. enterica serovar Typhi str. CT18 GCA_000195995.1
85 Salmonella enterica subsp. enterica serovar Typhimurium str. LT2 GCA_000006945.2
85 Shigella dysenteriae Sd197 GCA_000012005.1
85 Shigella flexneri 2a str. 301 GCA_000006925.2
85 Tumebacillus flagellatus GCA_000714935.1
82 Rosenbergiella nectarea GCA_900111105.1
82 Shewanella amazonensis SB2B GCA_000015245.1
82 Shewanella marina JCM 15074 GCA_000614975.1
82 Tolumonas auensis DSM 9187 GCA_000023065.1
81 Buttiauxella ferragutiae ATCC 51602 GCA_001654915.1
81 Cedecea neteri GCA_000757825.1
81 Citrobacter freundii CFNIH1 GCA_000648515.1
81 Erwinia billingiae Eb661 GCA_000196615.1
81 Erwinia gerundensis GCA_001517405.1
81 Klebsiella aerogenes KCTC 2190 GCA_000215745.1
81 Kluyvera ascorbata ATCC 33433 GCA_000735365.1
81 Kluyvera cryocrescens NBRC 102467 GCA_001571285.1
81 Pantoea agglomerans GCA_001709315.1
81 Pantoea rwandensis GCA_000759475.1
81 Pantoea septica GCA_002095575.1
81 Pragia fontium GCA_001026985.1
81 Shewanella pealeana ATCC 700345 GCA_000018285.1
81 Shewanella piezotolerans WP3 GCA_000014885.1
81 Yokenella regensburgei ATCC 49455 GCA_000735455.1
D9
241 Clostridium beijerinckii GCA_000833105.2
241 Clostridium saccharoperbutylacetonicum N1-4_28HMT_29 GCA_000340885.1
237 Clostridium saccharobutylicum DSM 13864 GCA_000473995.1
234 Clostridium puniceum GCA_002006345.1
Da
189 Enterococcus faecalis V583 GCA_000007785.1
189 Streptomyces cinnamoneus GCA_001885705.1
150 Enterococcus canis NBRC 100695 GCA_001544375.1
150 Enterococcus casseliflavus EC20 GCA_000157355.2
150 Enterococcus dispar ATCC 51266 GCA_000406945.1
150 Enterococcus faecium DO GCA_000174395.2
150 Enterococcus haemoperoxidus ATCC BAA-382 GCA_000407165.1
150 Enterococcus hirae ATCC 9790 GCA_000271405.2
150 Enterococcus mundtii QU 25 GCA_000504125.1
150 Enterococcus phoeniculicola ATCC BAA-412 GCA_000407505.1
150 Enterococcus rivorum GCA_001742285.1
150 Enterococcus saccharolyticus subsp. saccharolyticus ATCC 43076 GCA_000407285.1
150 Enterococcus thailandicus GCA_001652875.1
139 Enterococcus gilvus ATCC BAA-350 GCA_000407545.1
139 Enterococcus hermanniensis GCA_001885945.1
139 Enterococcus malodoratus ATCC 43197 GCA_000407185.1
139 Enterococcus pallens ATCC BAA-351 GCA_000407485.1
139 Enterococcus pseudoavium NBRC 100491 GCA_001544295.1
Db
262 Deinococcus radiodurans R1 GCA_000008565.1
186 Deinococcus gobiensis I-0 GCA_000252445.1
176 Deinococcus puniceus GCA_001644565.1
Dc
270 Deinococcus radiodurans R1 GCA_000008565.1
205 Deinococcus gobiensis I-0 GCA_000252445.1
169 Deinococcus deserti VCD115 GCA_000020685.1
Dd

208 Bifidobacterium adolescentis ATCC 15703 GCA_000010425.1
194 Bifidobacterium thermophilum GCA_000741495.1
193 Bifidobacterium dentium JCM 1195 = DSM 20436 GCA_001042595.1
193 Bifidobacterium tsurumiense GCA_000741765.1
De
152 Bacillus anthracis str. Ames GCA_000007845.1
152 Bacillus anthracis str. Sterne GCA_000008165.1
152 Bacillus cereus ATCC 14579 GCA_000007825.1
152 Bacillus mycoides GCA_000832605.1
152 Bacillus pseudomycoides DSM 12442 GCA_000161455.1
152 Bacillus thuringiensis YBT-1518 GCA_000497525.2
152 _5BBacillus thuringiensis_5D serovar konkukian str. 97-27 GCA_000008505.1
114 Bacillus manliponensis GCA_000712595.1
108 Staphylococcus equorum GCA_001432245.1
Df
77 Enterococcus faecalis V583 GCA_000007785.1
77 Streptomyces cinnamoneus GCA_001885705.1
53 Listeria floridensis FSL S10-1187 GCA_000525875.1
51 Enterococcus faecium DO GCA_000174395.2
Dg
155 Clostridium beijerinckii GCA_000833105.2
155 Clostridium puniceum GCA_002006345.1
155 Clostridium saccharobutylicum DSM 13864 GCA_000473995.1
155 Clostridium saccharoperbutylacetonicum N1-4_28HMT_29 GCA_000340885.1
144 Clostridium botulinum B str. Eklund 17B _28NRP_29 GCA_000020165.1
144 Clostridium chromiireducens GCA_002029255.1
144 Clostridium taeniosporum GCA_001735765.1
144 Clostridium uliginosum GCA_900112485.1
136 Clostridium intestinale URNW GCA_000469625.2
Dh
318 Enterococcus faecalis V583 GCA_000007785.1
318 Streptomyces cinnamoneus GCA_001885705.1
294 Enterococcus asini ATCC 700915 GCA_000407365.1
294 Enterococcus canis NBRC 100695 GCA_001544375.1
294 Enterococcus casseliflavus EC20 GCA_000157355.2
294 Enterococcus dispar ATCC 51266 GCA_000406945.1
294 Enterococcus faecium DO GCA_000174395.2
294 Enterococcus hirae ATCC 9790 GCA_000271405.2
294 Enterococcus massiliensis GCA_001050095.1
294 Enterococcus mundtii QU 25 GCA_000504125.1
294 Enterococcus rivorum GCA_001742285.1
294 Enterococcus saccharolyticus subsp. saccharolyticus ATCC 43076 GCA_000407285.1
289 Enterococcus haemoperoxidus ATCC BAA-382 GCA_000407165.1
289 Enterococcus phoeniculicola ATCC BAA-412 GCA_000407505.1
289 Enterococcus thailandicus GCA_001652875.1
Di
205 Deinococcus radiodurans R1 GCA_000008565.1
117 Deinococcus deserti VCD115 GCA_000020685.1
112 Deinococcus gobiensis I-0 GCA_000252445.1
Dj
304 Bacillus anthracis str. Ames GCA_000007845.1
304 Bacillus anthracis str. Sterne GCA_000008165.1
304 Bacillus cereus ATCC 14579 GCA_000007825.1
304 Bacillus mycoides GCA_000832605.1
304 Bacillus pseudomycoides DSM 12442 GCA_000161455.1
304 Bacillus thuringiensis YBT-1518 GCA_000497525.2
304 _5BBacillus thuringiensis_5D serovar konkukian str. 97-27 GCA_000008505.1
229 Bacillus manliponensis GCA_000712595.1
216 Bacillus cytotoxicus NVH 391-98 GCA_000017425.1
Dk
209 Lactobacillus gasseri ATCC 33323 = JCM 1131 GCA_000014425.1
173 Lactobacillus hominis DSM 23910 = CRBIP 24.179 GCA_000296835.1
125 Lactobacillus iners DSM 13335 GCA_000160875.1
Dl
172 Rhodobacter sphaeroides 2.4.1 GCA_000012905.2
138 Haematobacter massiliensis GCA_000740795.1
134 Pseudorhodobacter psychrotolerans GCA_001294535.1
134 Rhodobacter sphaeroides ATCC 17025 GCA_000016405.1
Dm
231 Escherichia coli IAI39 GCA_000026345.1
231 Escherichia coli O104_3AH4 str. 2011C-3493 GCA_000299455.1
231 Escherichia coli O157_3AH7 str. Sakai GCA_000008865.1
231 Escherichia coli O83_3AH1 str. NRG 857C GCA_000183345.1
231 Escherichia coli UMN026 GCA_000026325.2
231 Escherichia coli str. K-12 substr. MG1655 GCA_000005845.2
231 Shigella flexneri 2a str. 301 GCA_000006925.2

231 Tumebacillus flagellatus GCA_000714935.1
216 Shigella dysenteriae Sd197 GCA_000012005.1
203 Erwinia iniecta GCA_001267535.1
203 Erwinia toletana DAPP-PG 735 GCA_000336255.1
203 Pantoea ananatis LMG 20103 GCA_000025405.2
203 Shimwellia blattae DSM 4481 = NBRC 105725 GCA_000262305.1
Dn
262 Staphylococcus capitis subsp. capitis GCA_001028645.1
262 Staphylococcus epidermidis ATCC 12228 GCA_000007645.1
262 Staphylococcus haemolyticus JCSC1435 GCA_000009865.1
252 Staphylococcus hominis subsp. hominis C80 GCA_000183685.1
252 Staphylococcus lugdunensis HKU09-01 GCA_000025085.1
248 Staphylococcus arlettae CVD059 GCA_000295715.1
248 Staphylococcus cohnii subsp. cohnii GCA_000972575.1
248 Staphylococcus saprophyticus subsp. saprophyticus ATCC 15305 GCA_000010125.1
Do
256 Bacillus thuringiensis YBT-1518 GCA_000497525.2
241 Bacillus anthracis str. Ames GCA_000007845.1
241 Bacillus anthracis str. Sterne GCA_000008165.1
241 Bacillus cereus ATCC 14579 GCA_000007825.1
241 Bacillus mycoides GCA_000832605.1
241 Bacillus pseudomycoides DSM 12442 GCA_000161455.1
241 _5BBacillus thuringiensis_5D serovar konkukian str. 97-27 GCA_000008505.1
164 Massilibacterium senegalense GCA_001375675.1
Dp
112 Rhodobacter sphaeroides 2.4.1 GCA_000012905.2
109 Rhodobacter sphaeroides ATCC 17025 GCA_000016405.1
107 Pseudorhodobacter psychrotolerans GCA_001294535.1
Dq
231 Escherichia coli IAI39 GCA_000026345.1
231 Escherichia coli O104_3AH4 str. 2011C-3493 GCA_000299455.1
231 Escherichia coli O157_3AH7 str. Sakai GCA_000008865.1
231 Escherichia coli O83_3AH1 str. NRG 857C GCA_000183345.1
231 Escherichia coli UMN026 GCA_000026325.2
231 Escherichia coli str. K-12 substr. MG1655 GCA_000005845.2
231 Shigella flexneri 2a str. 301 GCA_000006925.2
231 Tumebacillus flagellatus GCA_000714935.1
225 Erwinia iniecta GCA_001267535.1
222 Shigella dysenteriae Sd197 GCA_000012005.1
Dr
346 Staphylococcus epidermidis ATCC 12228 GCA_000007645.1
335 Staphylococcus lugdunensis HKU09-01 GCA_000025085.1
333 Staphylococcus capitis subsp. capitis GCA_001028645.1
Ds
269 Bacillus thuringiensis YBT-1518 GCA_000497525.2
255 Bacillus anthracis str. Ames GCA_000007845.1
255 Bacillus anthracis str. Sterne GCA_000008165.1
255 Bacillus cereus ATCC 14579 GCA_000007825.1
255 Bacillus mycoides GCA_000832605.1
255 Bacillus pseudomycoides DSM 12442 GCA_000161455.1
255 _5BBacillus thuringiensis_5D serovar konkukian str. 97-27 GCA_000008505.1
202 Bacillus flexus GCA_002024265.1
202 Bacillus megaterium NBRC 15308 = ATCC 14581 GCA_000832985.1
Dt
218 Escherichia coli IAI39 GCA_000026345.1
218 Escherichia coli O104_3AH4 str. 2011C-3493 GCA_000299455.1
218 Escherichia coli O157_3AH7 str. Sakai GCA_000008865.1
218 Escherichia coli O83_3AH1 str. NRG 857C GCA_000183345.1
218 Escherichia coli UMN026 GCA_000026325.2
218 Escherichia coli str. K-12 substr. MG1655 GCA_000005845.2
218 Shigella dysenteriae Sd197 GCA_000012005.1
218 Shigella flexneri 2a str. 301 GCA_000006925.2
218 Tumebacillus flagellatus GCA_000714935.1
192 Erwinia iniecta GCA_001267535.1
186 Erwinia toletana DAPP-PG 735 GCA_000336255.1
Du
198 Deinococcus radiodurans R1 GCA_000008565.1
111 Deinococcus gobiensis I-0 GCA_000252445.1
104 Deinococcus puniceus GCA_001644565.1
Dv
228 Clostridium beijerinckii GCA_000833105.2
228 Clostridium puniceum GCA_002006345.1
228 Clostridium saccharoperbutylacetonicum N1-4_28HMT_29 GCA_000340885.1
219 Clostridium chromiireducens GCA_002029255.1
219 Clostridium saccharobutylicum DSM 13864 GCA_000473995.1
217 Clostridium butyricum GCA_001456065.2

Dw
182 Bacillus anthracis str. Ames GCA_000007845.1
182 Bacillus anthracis str. Sterne GCA_000008165.1
182 Bacillus cereus ATCC 14579 GCA_000007825.1
182 Bacillus mycoides GCA_000832605.1
182 Bacillus pseudomycoides DSM 12442 GCA_000161455.1
182 Bacillus thuringiensis YBT-1518 GCA_000497525.2
182 _5BBacillus thuringiensis_5D serovar konkukian str. 97-27 GCA_000008505.1
158 Bacillus manliponensis GCA_000712595.1
150 Bacillus aquimaris TF-12 GCA_001648555.1
150 Bacillus vietnamensis NBRC 101237 GCA_001591825.1
Dx
181 Streptococcus mutans UA159 GCA_000007465.2
138 Streptococcus ferus DSM 20646 GCA_000372425.1
138 Streptococcus merionis DSM 19192 GCA_000380085.1
133 Streptococcus equinus GCA_000964315.1
133 Streptococcus gallolyticus subsp. gallolyticus DSM 16831 GCA_002000985.1
Dy
183 Deinococcus radiodurans R1 GCA_000008565.1
108 Deinococcus marmoris DSM 12784 GCA_000701405.1
108 Deinococcus swuensis GCA_000800395.1
100 Deinococcus hopiensis KR-140 GCA_900176165.1
Dz
157 Enterococcus asini ATCC 700915 GCA_000407365.1
157 Enterococcus canis NBRC 100695 GCA_001544375.1
157 Enterococcus casseliflavus EC20 GCA_000157355.2
157 Enterococcus dispar ATCC 51266 GCA_000406945.1
157 Enterococcus faecalis V583 GCA_000007785.1
157 Enterococcus faecium DO GCA_000174395.2
157 Enterococcus hirae ATCC 9790 GCA_000271405.2
157 Enterococcus massiliensis GCA_001050095.1
157 Enterococcus mundtii QU 25 GCA_000504125.1
157 Enterococcus rivorum GCA_001742285.1
157 Enterococcus saccharolyticus subsp. saccharolyticus ATCC 43076 GCA_000407285.1
157 Streptomyces cinnamoneus GCA_001885705.1
152 Enterococcus haemoperoxidus ATCC BAA-382 GCA_000407165.1
152 Enterococcus phoeniculicola ATCC BAA-412 GCA_000407505.1
152 Enterococcus thailandicus GCA_001652875.1
152 Isobaculum melis GCA_900111355.1
145 Listeria floridensis FSL S10-1187 GCA_000525875.1
DA
242 Bacillus anthracis str. Ames GCA_000007845.1
242 Bacillus anthracis str. Sterne GCA_000008165.1
242 Bacillus cereus ATCC 14579 GCA_000007825.1
242 Bacillus mycoides GCA_000832605.1
242 Bacillus pseudomycoides DSM 12442 GCA_000161455.1
242 Bacillus thuringiensis YBT-1518 GCA_000497525.2
242 _5BBacillus thuringiensis_5D serovar konkukian str. 97-27 GCA_000008505.1
197 Bacillus marisflavi GCA_001274775.1
192 Bacillus manliponensis GCA_000712595.1
DB
223 Rhodobacter sphaeroides 2.4.1 GCA_000012905.2
191 Gemmobacter aquatilis GCA_900110025.1
191 Rhodobacter capsulatus SB 1003 GCA_000021865.1
191 Rhodobacter sphaeroides ATCC 17025 GCA_000016405.1
188 Pseudorhodobacter psychrotolerans GCA_001294535.1
DC
72 Clostridium beijerinckii GCA_000833105.2
72 Clostridium saccharoperbutylacetonicum N1-4_28HMT_29 GCA_000340885.1
65 Clostridium puniceum GCA_002006345.1
57 Clostridium saccharobutylicum DSM 13864 GCA_000473995.1
DD
161 Deinococcus radiodurans R1 GCA_000008565.1
100 Deinococcus gobiensis I-0 GCA_000252445.1
91 Deinococcus geothermalis DSM 11300 GCA_000196275.1
DE
174 Escherichia coli IAI39 GCA_000026345.1
174 Escherichia coli O104_3AH4 str. 2011C-3493 GCA_000299455.1
174 Escherichia coli O157_3AH7 str. Sakai GCA_000008865.1
174 Escherichia coli O83_3AH1 str. NRG 857C GCA_000183345.1
174 Escherichia coli UMN026 GCA_000026325.2
174 Escherichia coli str. K-12 substr. MG1655 GCA_000005845.2
174 Shigella dysenteriae Sd197 GCA_000012005.1
174 Shigella flexneri 2a str. 301 GCA_000006925.2
174 Tumebacillus flagellatus GCA_000714935.1
159 Erwinia iniecta GCA_001267535.1

155 Pantoea agglomerans GCA_001709315.1
DF
171 Deinococcus radiodurans R1 GCA_000008565.1
136 Deinococcus puniceus GCA_001644565.1
120 Deinococcus proteolyticus MRP GCA_000190555.1
DG
239 Clostridium beijerinckii GCA_000833105.2
239 Clostridium saccharobutylicum DSM 13864 GCA_000473995.1
239 Clostridium saccharoperbutylacetonicum N1-4_28HMT_29 GCA_000340885.1
233 Clostridium puniceum GCA_002006345.1
225 Clostridium butyricum GCA_001456065.2
DH
283 Clostridium beijerinckii GCA_000833105.2
283 Clostridium saccharoperbutylacetonicum N1-4_28HMT_29 GCA_000340885.1
274 Clostridium butyricum GCA_001456065.2
268 Clostridium puniceum GCA_002006345.1
268 Clostridium saccharobutylicum DSM 13864 GCA_000473995.1
DI
243 Deinococcus radiodurans R1 GCA_000008565.1
140 Deinococcus puniceus GCA_001644565.1
136 Deinococcus deserti VCD115 GCA_000020685.1
DJ
143 Lactobacillus psittaci DSM 15354 GCA_000425905.1
138 Lactobacillus gasseri ATCC 33323 = JCM 1131 GCA_000014425.1
138 Lactobacillus hominis DSM 23910 = CRBIP 24.179 GCA_000296835.1
116 Lactobacillus acidophilus NCFM GCA_000011985.1
116 Lactobacillus amylolyticus GCA_002075105.1
116 Lactobacillus amylovorus GCA_000191545.1
116 Lactobacillus crispatus ST1 GCA_000091765.1
116 Lactobacillus hamsteri DSM 5661 = JCM 6256 GCA_000615445.1
116 Lactobacillus jensenii GCA_001936235.1
116 Lactobacillus kalixensis DSM 16043 GCA_001434335.1
116 Lactobacillus kefiranofaciens ZW3 GCA_000214785.1
DK
188 Clostridium butyricum GCA_001456065.2
184 Clostridium beijerinckii GCA_000833105.2
184 Clostridium puniceum GCA_002006345.1
184 Clostridium saccharoperbutylacetonicum N1-4_28HMT_29 GCA_000340885.1
174 Clostridium neonatale GCA_001458595.1
DL
245 Staphylococcus epidermidis ATCC 12228 GCA_000007645.1
242 Staphylococcus haemolyticus JCSC1435 GCA_000009865.1
240 Staphylococcus capitis subsp. capitis GCA_001028645.1
DM
218 Rhodobacter sphaeroides 2.4.1 GCA_000012905.2
185 Rhodobacter sphaeroides ATCC 17025 GCA_000016405.1
158 Thioclava dalianensis GCA_000715505.1
158 Thioclava indica GCA_000714545.1
DN
108 Deinococcus radiodurans R1 GCA_000008565.1
62 Deinococcus marmoris DSM 12784 GCA_000701405.1
62 Deinococcus swuensis GCA_000800395.1
45 Deinococcus apachensis DSM 19763 GCA_000381345.1
45 Deinococcus deserti VCD115 GCA_000020685.1
45 Deinococcus frigens DSM 12807 GCA_000701425.1
45 Deinococcus geothermalis DSM 11300 GCA_000196275.1
45 Deinococcus hopiensis KR-140 GCA_900176165.1
45 Deinococcus murrayi DSM 11303 GCA_000482805.1
45 Deinococcus phoenicis GCA_000599865.1
45 Deinococcus puniceus GCA_001644565.1
45 Deinococcus soli Cha et al. 2016 GCA_001007995.1
DO
263 Rhodobacter sphaeroides 2.4.1 GCA_000012905.2
225 Rhodobacter sphaeroides ATCC 17025 GCA_000016405.1
218 Pseudorhodobacter psychrotolerans GCA_001294535.1
DP
91 Deinococcus radiodurans R1 GCA_000008565.1
76 Deinococcus deserti VCD115 GCA_000020685.1
76 Deinococcus hopiensis KR-140 GCA_900176165.1
76 Deinococcus puniceus GCA_001644565.1
75 Deinococcus proteolyticus MRP GCA_000190555.1
DQ
174 Clostridium beijerinckii GCA_000833105.2
174 Clostridium puniceum GCA_002006345.1
174 Clostridium saccharobutylicum DSM 13864 GCA_000473995.1
174 Clostridium saccharoperbutylacetonicum N1-4_28HMT_29 GCA_000340885.1

170 Clostridium chromiireducens GCA_002029255.1
161 Clostridium butyricum GCA_001456065.2
DR
173 Deinococcus radiodurans R1 GCA_000008565.1
132 Deinococcus gobiensis I-0 GCA_000252445.1
123 Deinococcus marmoris DSM 12784 GCA_000701405.1
123 Deinococcus swuensis GCA_000800395.1
DS
201 Deinococcus radiodurans R1 GCA_000008565.1
164 Deinococcus puniceus GCA_001644565.1
156 Deinococcus deserti VCD115 GCA_000020685.1
156 Deinococcus hopiensis KR-140 GCA_900176165.1
DT
218 Rhodobacter sphaeroides 2.4.1 GCA_000012905.2
215 Rhodobacter sphaeroides ATCC 17025 GCA_000016405.1
192 Gemmobacter megaterium GCA_900156815.1
DU
288 Bacillus anthracis str. Ames GCA_000007845.1
288 Bacillus anthracis str. Sterne GCA_000008165.1
288 Bacillus cereus ATCC 14579 GCA_000007825.1
288 Bacillus mycoides GCA_000832605.1
288 Bacillus pseudomycoides DSM 12442 GCA_000161455.1
288 Bacillus thuringiensis YBT-1518 GCA_000497525.2
288 _5BBacillus thuringiensis_5D serovar konkukian str. 97-27 GCA_000008505.1
228 Bacillus manliponensis GCA_000712595.1
224 Bacillus flexus GCA_002024265.1
224 Bacillus megaterium NBRC 15308 = ATCC 14581 GCA_000832985.1
DV
252 Clostridium beijerinckii GCA_000833105.2
252 Clostridium saccharobutylicum DSM 13864 GCA_000473995.1
252 Clostridium saccharoperbutylacetonicum N1-4_28HMT_29 GCA_000340885.1
237 Clostridium puniceum GCA_002006345.1
226 Clostridium butyricum GCA_001456065.2
226 Clostridium chromiireducens GCA_002029255.1
DW
259 Clostridium beijerinckii GCA_000833105.2
259 Clostridium saccharobutylicum DSM 13864 GCA_000473995.1
259 Clostridium saccharoperbutylacetonicum N1-4_28HMT_29 GCA_000340885.1
253 Clostridium puniceum GCA_002006345.1
238 Clostridium chromiireducens GCA_002029255.1
DX
151 Clostridium saccharobutylicum DSM 13864 GCA_000473995.1
149 Clostridium beijerinckii GCA_000833105.2
149 Clostridium saccharoperbutylacetonicum N1-4_28HMT_29 GCA_000340885.1
142 Clostridium puniceum GCA_002006345.1
DY
240 Deinococcus radiodurans R1 GCA_000008565.1
161 Deinococcus gobiensis I-0 GCA_000252445.1
139 Deinococcus deserti VCD115 GCA_000020685.1
139 Deinococcus proteolyticus MRP GCA_000190555.1
DZ
230 Deinococcus radiodurans R1 GCA_000008565.1
199 Deinococcus gobiensis I-0 GCA_000252445.1
190 Deinococcus deserti VCD115 GCA_000020685.1
E0
196 Enterococcus faecalis V583 GCA_000007785.1
196 Streptomyces cinnamoneus GCA_001885705.1
178 Clostridium beijerinckii GCA_000833105.2
178 Clostridium puniceum GCA_002006345.1
178 Clostridium saccharoperbutylacetonicum N1-4_28HMT_29 GCA_000340885.1
169 Enterococcus aquimarinus GCA_001885765.1
169 Enterococcus asini ATCC 700915 GCA_000407365.1
169 Enterococcus canis NBRC 100695 GCA_001544375.1
169 Enterococcus casseliflavus EC20 GCA_000157355.2
169 Enterococcus cecorum GCA_001318405.1
169 Enterococcus columbae DSM 7374 = ATCC 51263 GCA_000406925.1
169 Enterococcus dispar ATCC 51266 GCA_000406945.1
169 Enterococcus faecium DO GCA_000174395.2
169 Enterococcus gilvus ATCC BAA-350 GCA_000407545.1
169 Enterococcus haemoperoxidus ATCC BAA-382 GCA_000407165.1
169 Enterococcus hermanniensis GCA_001885945.1
169 Enterococcus hirae ATCC 9790 GCA_000271405.2
169 Enterococcus italicus DSM 15952 GCA_000185365.1
169 Enterococcus malodoratus ATCC 43197 GCA_000407185.1
169 Enterococcus massiliensis GCA_001050095.1
169 Enterococcus mundtii QU 25 GCA_000504125.1

169 Enterococcus pallens ATCC BAA-351 GCA_000407485.1
169 Enterococcus phoeniculicola ATCC BAA-412 GCA_000407505.1
169 Enterococcus pseudoavium NBRC 100491 GCA_001544295.1
169 Enterococcus rivorum GCA_001742285.1
169 Enterococcus saccharolyticus subsp. saccharolyticus ATCC 43076 GCA_000407285.1
169 Enterococcus sulfureus ATCC 49903 GCA_000407605.1
169 Enterococcus thailandicus GCA_001652875.1
169 Melissococcus plutonius S1 GCA_000747585.1
169 Vagococcus fluvialis bH819 GCA_900163795.1
169 Vagococcus lutrae LBD1 GCA_000498295.1
169 Vagococcus penaei GCA_001998885.1
169 Vagococcus teuberi GCA_001870205.1
E1
291 Lactobacillus gasseri ATCC 33323 = JCM 1131 GCA_000014425.1
279 Lactobacillus hominis DSM 23910 = CRBIP 24.179 GCA_000296835.1
204 Lactobacillus psittaci DSM 15354 GCA_000425905.1
E2
233 Streptococcus mutans UA159 GCA_000007465.2
156 Streptococcus ferus DSM 20646 GCA_000372425.1
154 Streptococcus macacae NCTC 11558 GCA_000187995.3
E3
182 Deinococcus radiodurans R1 GCA_000008565.1
90 Deinococcus gobiensis I-0 GCA_000252445.1
87 Deinococcus proteolyticus MRP GCA_000190555.1
E4
136 Streptococcus mutans UA159 GCA_000007465.2
125 Streptococcus ratti FA-1 = DSM 20564 GCA_000286075.1
110 Streptococcus equinus GCA_000964315.1
110 Streptococcus gallolyticus subsp. gallolyticus DSM 16831 GCA_002000985.1
110 Streptococcus orisratti DSM 15617 GCA_000380105.1
E5
109 Deinococcus hopiensis KR-140 GCA_900176165.1
109 Deinococcus puniceus GCA_001644565.1
109 Deinococcus radiodurans R1 GCA_000008565.1
107 Deinococcus deserti VCD115 GCA_000020685.1
107 Deinococcus soli Cha et al. 2016 GCA_001007995.1
99 Deinococcus frigens DSM 12807 GCA_000701425.1
99 Deinococcus geothermalis DSM 11300 GCA_000196275.1
99 Deinococcus marmoris DSM 12784 GCA_000701405.1
99 Deinococcus swuensis GCA_000800395.1
E6
209 Escherichia coli IAI39 GCA_000026345.1
209 Escherichia coli O104_3AH4 str. 2011C-3493 GCA_000299455.1
209 Escherichia coli O157_3AH7 str. Sakai GCA_000008865.1
209 Escherichia coli O83_3AH1 str. NRG 857C GCA_000183345.1
209 Escherichia coli UMN026 GCA_000026325.2
209 Escherichia coli str. K-12 substr. MG1655 GCA_000005845.2
209 Shigella dysenteriae Sd197 GCA_000012005.1
209 Shigella flexneri 2a str. 301 GCA_000006925.2
209 Tumebacillus flagellatus GCA_000714935.1
174 Erwinia iniecta GCA_001267535.1
174 Pectobacterium carotovorum subsp. carotovorum PC1 GCA_000023605.1
166 Pantoea agglomerans GCA_001709315.1
E7
207 Rhodobacter sphaeroides 2.4.1 GCA_000012905.2
166 Rhodobacter sphaeroides ATCC 17025 GCA_000016405.1
139 Pseudorhodobacter ferrugineus DSM 5888 GCA_000420745.1
139 Pseudorhodobacter psychrotolerans GCA_001294535.1
139 Pseudorhodobacter wandonensis GCA_001202035.1
E8
228 Clostridium beijerinckii GCA_000833105.2
228 Clostridium saccharobutylicum DSM 13864 GCA_000473995.1
228 Clostridium saccharoperbutylacetonicum N1-4_28HMT_29 GCA_000340885.1
214 Clostridium puniceum GCA_002006345.1
210 Clostridium chromiireducens GCA_002029255.1
E9
257 Streptococcus mutans UA159 GCA_000007465.2
172 Streptococcus equinus GCA_000964315.1
172 Streptococcus gallolyticus subsp. gallolyticus DSM 16831 GCA_002000985.1
169 Streptococcus iniae GCA_000831485.1
169 Streptococcus sobrinus DSM 20742 = ATCC 33478 GCA_000686605.1
Ea
159 Staphylococcus aureus subsp. aureus NCTC 8325 GCA_000013425.1
159 Staphylococcus capitis subsp. capitis GCA_001028645.1
159 Staphylococcus epidermidis ATCC 12228 GCA_000007645.1
159 Staphylococcus haemolyticus JCSC1435 GCA_000009865.1

159 Staphylococcus hominis subsp. hominis C80 GCA_000183685.1
159 Staphylococcus lugdunensis HKU09-01 GCA_000025085.1
159 Staphylococcus simiae CCM 7213 GCA_000235645.2
153 Salinicoccus halodurans GCA_001005905.1
153 Staphylococcus lutrae GCA_002101335.1
144 Staphylococcus arlettae CVD059 GCA_000295715.1
144 Staphylococcus cohnii subsp. cohnii GCA_000972575.1
144 Staphylococcus condimenti GCA_001618885.1
144 Staphylococcus equorum GCA_001432245.1
144 Staphylococcus gallinarum GCA_000875895.1
144 Staphylococcus hyicus GCA_000816085.1
144 Staphylococcus saprophyticus subsp. saprophyticus ATCC 15305 GCA_000010125.1
144 Staphylococcus simulans GCA_001559115.1
144 Staphylococcus succinus GCA_001902315.1
144 Staphylococcus xylosus GCA_000706685.1
Eb
196 Clostridium butyricum GCA_001456065.2
195 Clostridium beijerinckii GCA_000833105.2
195 Clostridium puniceum GCA_002006345.1
195 Clostridium saccharobutylicum DSM 13864 GCA_000473995.1
195 Clostridium saccharoperbutylacetonicum N1-4_28HMT_29 GCA_000340885.1
194 Clostridium neonatale GCA_001458595.1
Ec
273 Rhodobacter sphaeroides 2.4.1 GCA_000012905.2
253 Rhodobacter sphaeroides ATCC 17025 GCA_000016405.1
246 Pseudorhodobacter ferrugineus DSM 5888 GCA_000420745.1
246 Pseudorhodobacter wandonensis GCA_001202035.1
Ed
221 Escherichia coli IAI39 GCA_000026345.1
221 Escherichia coli O104_3AH4 str. 2011C-3493 GCA_000299455.1
221 Escherichia coli O157_3AH7 str. Sakai GCA_000008865.1
221 Escherichia coli UMN026 GCA_000026325.2
221 Escherichia coli str. K-12 substr. MG1655 GCA_000005845.2
221 Shigella dysenteriae Sd197 GCA_000012005.1
221 Tumebacillus flagellatus GCA_000714935.1
220 Escherichia coli O83_3AH1 str. NRG 857C GCA_000183345.1
210 Shigella flexneri 2a str. 301 GCA_000006925.2
Ee
265 Bacillus anthracis str. Ames GCA_000007845.1
265 Bacillus anthracis str. Sterne GCA_000008165.1
265 Bacillus mycoides GCA_000832605.1
265 Bacillus pseudomycoides DSM 12442 GCA_000161455.1
265 Bacillus thuringiensis YBT-1518 GCA_000497525.2
265 _5BBacillus thuringiensis_5D serovar konkukian str. 97-27 GCA_000008505.1
264 Bacillus cereus ATCC 14579 GCA_000007825.1
221 Bacillus manliponensis GCA_000712595.1
Ef
239 Escherichia coli IAI39 GCA_000026345.1
239 Escherichia coli O104_3AH4 str. 2011C-3493 GCA_000299455.1
239 Escherichia coli O157_3AH7 str. Sakai GCA_000008865.1
239 Escherichia coli UMN026 GCA_000026325.2
239 Escherichia coli str. K-12 substr. MG1655 GCA_000005845.2
239 Shigella flexneri 2a str. 301 GCA_000006925.2
239 Tumebacillus flagellatus GCA_000714935.1
238 Escherichia coli O83_3AH1 str. NRG 857C GCA_000183345.1
230 Shigella dysenteriae Sd197 GCA_000012005.1
Eg
244 Bacillus thuringiensis YBT-1518 GCA_000497525.2
213 Bacillus anthracis str. Ames GCA_000007845.1
213 Bacillus anthracis str. Sterne GCA_000008165.1
213 Bacillus cereus ATCC 14579 GCA_000007825.1
213 Bacillus mycoides GCA_000832605.1
213 Bacillus pseudomycoides DSM 12442 GCA_000161455.1
213 _5BBacillus thuringiensis_5D serovar konkukian str. 97-27 GCA_000008505.1
197 Bacillus eiseniae GCA_001636325.1
197 Bacillus horneckiae GCA_001636335.1
197 Bacillus solani GCA_001420595.1
Eh
278 Clostridium beijerinckii GCA_000833105.2
278 Clostridium puniceum GCA_002006345.1
278 Clostridium saccharoperbutylacetonicum N1-4_28HMT_29 GCA_000340885.1
270 Clostridium saccharobutylicum DSM 13864 GCA_000473995.1
259 Clostridium butyricum GCA_001456065.2
259 Clostridium chromiireducens GCA_002029255.1
Ei
305 Bacillus thuringiensis YBT-1518 GCA_000497525.2

299 Bacillus anthracis str. Ames GCA_000007845.1
299 Bacillus anthracis str. Sterne GCA_000008165.1
299 Bacillus mycoides GCA_000832605.1
299 Bacillus pseudomycoides DSM 12442 GCA_000161455.1
299 _5BBacillus thuringiensis_5D serovar konkukian str. 97-27 GCA_000008505.1
298 Bacillus cereus ATCC 14579 GCA_000007825.1
Ej
151 Deinococcus radiodurans R1 GCA_000008565.1
92 Deinococcus deserti VCD115 GCA_000020685.1
91 Deinococcus gobiensis I-0 GCA_000252445.1
Ek
126 Clostridium beijerinckii GCA_000833105.2
126 Clostridium puniceum GCA_002006345.1
126 Clostridium saccharobutylicum DSM 13864 GCA_000473995.1
126 Clostridium saccharoperbutylacetonicum N1-4_28HMT_29 GCA_000340885.1
107 Clostridium neonatale GCA_001458595.1
98 Clostridium butyricum GCA_001456065.2
El
157 Bifidobacterium adolescentis ATCC 15703 GCA_000010425.1
157 Bifidobacterium tsurumiense GCA_000741765.1
146 Bifidobacterium angulatum DSM 20098 = JCM 7096 GCA_001025155.1
146 Bifidobacterium longum NCC2705 GCA_000007525.1
146 Bifidobacterium saguini DSM 23967 GCA_000741715.1
145 Bifidobacterium breve DSM 20213 = JCM 1192 GCA_001025175.1
Em
292 Bacillus thuringiensis YBT-1518 GCA_000497525.2
286 Bacillus anthracis str. Ames GCA_000007845.1
286 Bacillus anthracis str. Sterne GCA_000008165.1
286 Bacillus cereus ATCC 14579 GCA_000007825.1
286 Bacillus mycoides GCA_000832605.1
286 Bacillus pseudomycoides DSM 12442 GCA_000161455.1
286 _5BBacillus thuringiensis_5D serovar konkukian str. 97-27 GCA_000008505.1
246 Bacillus cytotoxicus NVH 391-98 GCA_000017425.1
En
134 Rhodobacter sphaeroides 2.4.1 GCA_000012905.2
119 Rhodobacter sphaeroides ATCC 17025 GCA_000016405.1
112 Rhodobacter capsulatus SB 1003 GCA_000021865.1
Eo
280 Deinococcus radiodurans R1 GCA_000008565.1
172 Deinococcus puniceus GCA_001644565.1
151 Deinococcus gobiensis I-0 GCA_000252445.1
Ep
192 Deinococcus radiodurans R1 GCA_000008565.1
116 Deinococcus puniceus GCA_001644565.1
105 Deinococcus gobiensis I-0 GCA_000252445.1
Eq
236 Deinococcus radiodurans R1 GCA_000008565.1
165 Deinococcus gobiensis I-0 GCA_000252445.1
118 Deinococcus deserti VCD115 GCA_000020685.1
Er
290 Bifidobacterium adolescentis ATCC 15703 GCA_000010425.1
266 Bifidobacterium gallicum DSM 20093 = LMG 11596 GCA_000741205.1
263 Bifidobacterium dentium JCM 1195 = DSM 20436 GCA_001042595.1
Es
319 Staphylococcus capitis subsp. capitis GCA_001028645.1
319 Staphylococcus epidermidis ATCC 12228 GCA_000007645.1
319 Staphylococcus haemolyticus JCSC1435 GCA_000009865.1
308 Staphylococcus hominis subsp. hominis C80 GCA_000183685.1
308 Staphylococcus lugdunensis HKU09-01 GCA_000025085.1
306 Staphylococcus simulans GCA_001559115.1
Et
258 Clostridium beijerinckii GCA_000833105.2
258 Clostridium saccharobutylicum DSM 13864 GCA_000473995.1
258 Clostridium saccharoperbutylacetonicum N1-4_28HMT_29 GCA_000340885.1
254 Clostridium puniceum GCA_002006345.1
247 Clostridium butyricum GCA_001456065.2
Eu
229 Streptococcus mutans UA159 GCA_000007465.2
181 Streptococcus equinus GCA_000964315.1
181 Streptococcus merionis DSM 19192 GCA_000380085.1
179 Streptococcus ratti FA-1 = DSM 20564 GCA_000286075.1
Ev
205 Streptococcus mutans UA159 GCA_000007465.2
158 Streptococcus sobrinus DSM 20742 = ATCC 33478 GCA_000686605.1
153 Streptococcus ferus DSM 20646 GCA_000372425.1
153 Streptococcus gordonii str. Challis substr. CH1 GCA_000017005.1

153 Streptococcus marimammalium DSM 18627 GCA_000380045.1
Ew
246 Escherichia coli IAI39 GCA_000026345.1
246 Escherichia coli O104_3AH4 str. 2011C-3493 GCA_000299455.1
246 Escherichia coli O157_3AH7 str. Sakai GCA_000008865.1
246 Escherichia coli O83_3AH1 str. NRG 857C GCA_000183345.1
246 Escherichia coli UMN026 GCA_000026325.2
246 Escherichia coli str. K-12 substr. MG1655 GCA_000005845.2
246 Shigella dysenteriae Sd197 GCA_000012005.1
246 Shigella flexneri 2a str. 301 GCA_000006925.2
246 Tumebacillus flagellatus GCA_000714935.1
232 Erwinia iniecta GCA_001267535.1
216 Rosenbergiella nectarea GCA_900111105.1
Ex
191 Clostridium beijerinckii GCA_000833105.2
191 Clostridium puniceum GCA_002006345.1
191 Clostridium saccharoperbutylacetonicum N1-4_28HMT_29 GCA_000340885.1
188 Clostridium neonatale GCA_001458595.1
182 Clostridium saccharobutylicum DSM 13864 GCA_000473995.1
Ey
120 Deinococcus radiodurans R1 GCA_000008565.1
104 Deinococcus gobiensis I-0 GCA_000252445.1
98 Deinococcus geothermalis DSM 11300 GCA_000196275.1
Ez
163 Rhodobacter sphaeroides 2.4.1 GCA_000012905.2
148 Rhodobacter sphaeroides ATCC 17025 GCA_000016405.1
145 Pseudorhodobacter ferrugineus DSM 5888 GCA_000420745.1
145 Pseudorhodobacter psychrotolerans GCA_001294535.1
145 Pseudorhodobacter wandonensis GCA_001202035.1
EA
283 Clostridium beijerinckii GCA_000833105.2
283 Clostridium saccharoperbutylacetonicum N1-4_28HMT_29 GCA_000340885.1
269 Clostridium puniceum GCA_002006345.1
267 Clostridium saccharobutylicum DSM 13864 GCA_000473995.1
EB
288 Clostridium beijerinckii GCA_000833105.2
288 Clostridium saccharoperbutylacetonicum N1-4_28HMT_29 GCA_000340885.1
284 Clostridium puniceum GCA_002006345.1
279 Clostridium saccharobutylicum DSM 13864 GCA_000473995.1
EC
184 Streptococcus mutans UA159 GCA_000007465.2
122 Streptococcus ferus DSM 20646 GCA_000372425.1
121 Streptococcus sobrinus DSM 20742 = ATCC 33478 GCA_000686605.1
ED
91 Escherichia coli IAI39 GCA_000026345.1
91 Escherichia coli O104_3AH4 str. 2011C-3493 GCA_000299455.1
91 Escherichia coli O157_3AH7 str. Sakai GCA_000008865.1
91 Escherichia coli O83_3AH1 str. NRG 857C GCA_000183345.1
91 Escherichia coli UMN026 GCA_000026325.2
91 Escherichia coli str. K-12 substr. MG1655 GCA_000005845.2
91 Shigella dysenteriae Sd197 GCA_000012005.1
91 Shigella flexneri 2a str. 301 GCA_000006925.2
91 Tumebacillus flagellatus GCA_000714935.1
63 Erwinia iniecta GCA_001267535.1
63 Plautia stali symbiont GCA_000180175.2
51 Shewanella algae GCA_000947195.1
EE
154 Bifidobacterium adolescentis ATCC 15703 GCA_000010425.1
132 Bifidobacterium dentium JCM 1195 = DSM 20436 GCA_001042595.1
122 Bifidobacterium thermophilum GCA_000741495.1
122 Bifidobacterium thermophilum RBL67 GCA_000347695.1
EF
234 Staphylococcus capitis subsp. capitis GCA_001028645.1
234 Staphylococcus epidermidis ATCC 12228 GCA_000007645.1
231 Staphylococcus haemolyticus JCSC1435 GCA_000009865.1
230 Staphylococcus lugdunensis HKU09-01 GCA_000025085.1
EG
287 Streptococcus mutans UA159 GCA_000007465.2
168 Streptococcus sobrinus DSM 20742 = ATCC 33478 GCA_000686605.1
163 Streptococcus ferus DSM 20646 GCA_000372425.1
EH
136 Bifidobacterium adolescentis ATCC 15703 GCA_000010425.1
135 Bifidobacterium asteroides PRL2011 GCA_000304215.1
119 Bifidobacterium angulatum DSM 20098 = JCM 7096 GCA_001025155.1
119 Bifidobacterium callitrichos DSM 23973 GCA_000741175.1
EI

222 Deinococcus radiodurans R1 GCA_000008565.1
135 Deinococcus gobiensis I-0 GCA_000252445.1
123 Deinococcus deserti VCD115 GCA_000020685.1
EJ
187 Clostridium beijerinckii GCA_000833105.2
187 Clostridium chromiireducens GCA_002029255.1
187 Clostridium puniceum GCA_002006345.1
187 Clostridium saccharobutylicum DSM 13864 GCA_000473995.1
187 Clostridium saccharoperbutylacetonicum N1-4_28HMT_29 GCA_000340885.1
183 Clostridium butyricum GCA_001456065.2
165 Clostridium taeniosporum GCA_001735765.1
EK
188 Erwinia iniecta GCA_001267535.1
188 Escherichia coli IAI39 GCA_000026345.1
188 Escherichia coli O104_3AH4 str. 2011C-3493 GCA_000299455.1
188 Escherichia coli O157_3AH7 str. Sakai GCA_000008865.1
188 Escherichia coli O83_3AH1 str. NRG 857C GCA_000183345.1
188 Escherichia coli UMN026 GCA_000026325.2
188 Escherichia coli str. K-12 substr. MG1655 GCA_000005845.2
188 Shigella dysenteriae Sd197 GCA_000012005.1
188 Shigella flexneri 2a str. 301 GCA_000006925.2
188 Tumebacillus flagellatus GCA_000714935.1
182 Cronobacter sakazakii GCA_000982825.1
182 Erwinia toletana DAPP-PG 735 GCA_000336255.1
176 Xenorhabdus bovienii SS-2004 GCA_000027225.1
EL
261 Deinococcus radiodurans R1 GCA_000008565.1
159 Deinococcus gobiensis I-0 GCA_000252445.1
152 Deinococcus puniceus GCA_001644565.1
EM
273 Bifidobacterium adolescentis ATCC 15703 GCA_000010425.1
227 Bifidobacterium dentium JCM 1195 = DSM 20436 GCA_001042595.1
227 Bifidobacterium thermophilum GCA_000741495.1
221 Bifidobacterium breve DSM 20213 = JCM 1192 GCA_001025175.1
EN
205 Escherichia coli IAI39 GCA_000026345.1
205 Escherichia coli O104_3AH4 str. 2011C-3493 GCA_000299455.1
205 Escherichia coli O157_3AH7 str. Sakai GCA_000008865.1
205 Escherichia coli O83_3AH1 str. NRG 857C GCA_000183345.1
205 Escherichia coli UMN026 GCA_000026325.2
205 Escherichia coli str. K-12 substr. MG1655 GCA_000005845.2
205 Shigella dysenteriae Sd197 GCA_000012005.1
205 Shigella flexneri 2a str. 301 GCA_000006925.2
205 Tumebacillus flagellatus GCA_000714935.1
176 Erwinia iniecta GCA_001267535.1
172 Lonsdalea quercina subsp. quercina GCA_000688655.1
EO
202 Clostridium saccharobutylicum DSM 13864 GCA_000473995.1
198 Clostridium beijerinckii GCA_000833105.2
198 Clostridium chromiireducens GCA_002029255.1
198 Clostridium puniceum GCA_002006345.1
198 Clostridium saccharoperbutylacetonicum N1-4_28HMT_29 GCA_000340885.1
194 Clostridium butyricum GCA_001456065.2
EP
196 Escherichia coli IAI39 GCA_000026345.1
196 Escherichia coli O104_3AH4 str. 2011C-3493 GCA_000299455.1
196 Escherichia coli O157_3AH7 str. Sakai GCA_000008865.1
196 Escherichia coli O83_3AH1 str. NRG 857C GCA_000183345.1
196 Escherichia coli UMN026 GCA_000026325.2
196 Escherichia coli str. K-12 substr. MG1655 GCA_000005845.2
196 Shigella dysenteriae Sd197 GCA_000012005.1
196 Shigella flexneri 2a str. 301 GCA_000006925.2
196 Tumebacillus flagellatus GCA_000714935.1
189 Erwinia iniecta GCA_001267535.1
174 Cronobacter sakazakii GCA_000982825.1
EQ
187 Clostridium beijerinckii GCA_000833105.2
187 Clostridium saccharobutylicum DSM 13864 GCA_000473995.1
187 Clostridium saccharoperbutylacetonicum N1-4_28HMT_29 GCA_000340885.1
180 Clostridium puniceum GCA_002006345.1
167 Clostridium butyricum GCA_001456065.2
ER
254 Deinococcus radiodurans R1 GCA_000008565.1
161 Deinococcus deserti VCD115 GCA_000020685.1
154 Deinococcus puniceus GCA_001644565.1
ES

111 Bacillus anthracis str. Ames GCA_000007845.1
111 Bacillus anthracis str. Sterne GCA_000008165.1
111 Bacillus cereus ATCC 14579 GCA_000007825.1
111 Bacillus mycoides GCA_000832605.1
111 Bacillus pseudomycoides DSM 12442 GCA_000161455.1
111 Bacillus thuringiensis YBT-1518 GCA_000497525.2
111 _5BBacillus thuringiensis_5D serovar konkukian str. 97-27 GCA_000008505.1
78 Bacillus manliponensis GCA_000712595.1
77 Bacillus cytotoxicus NVH 391-98 GCA_000017425.1
77 Massilibacterium senegalense GCA_001375675.1
ET
229 Streptococcus mutans UA159 GCA_000007465.2
181 Streptococcus equinus GCA_000964315.1
181 Streptococcus gallolyticus subsp. gallolyticus DSM 16831 GCA_002000985.1
166 Streptococcus orisratti DSM 15617 GCA_000380105.1
166 Streptococcus ratti FA-1 = DSM 20564 GCA_000286075.1
EU
223 Bacillus anthracis str. Ames GCA_000007845.1
223 Bacillus anthracis str. Sterne GCA_000008165.1
223 Bacillus cereus ATCC 14579 GCA_000007825.1
223 Bacillus mycoides GCA_000832605.1
223 Bacillus pseudomycoides DSM 12442 GCA_000161455.1
223 Bacillus thuringiensis YBT-1518 GCA_000497525.2
223 _5BBacillus thuringiensis_5D serovar konkukian str. 97-27 GCA_000008505.1
164 Bacillus acidicola GCA_001636425.1
164 Bacillus shackletonii GCA_001420715.1
164 Bacillus sporothermodurans GCA_001587375.1
162 Bacillus horneckiae GCA_001636335.1
162 Bacillus solani GCA_001420595.1
EV
165 Escherichia coli IAI39 GCA_000026345.1
165 Escherichia coli O104_3AH4 str. 2011C-3493 GCA_000299455.1
165 Escherichia coli O157_3AH7 str. Sakai GCA_000008865.1
165 Escherichia coli O83_3AH1 str. NRG 857C GCA_000183345.1
165 Escherichia coli UMN026 GCA_000026325.2
165 Escherichia coli str. K-12 substr. MG1655 GCA_000005845.2
165 Shigella dysenteriae Sd197 GCA_000012005.1
165 Tumebacillus flagellatus GCA_000714935.1
157 Shigella flexneri 2a str. 301 GCA_000006925.2
129 Sodalis praecaptivus GCA_000517425.1
EW
141 Bifidobacterium adolescentis ATCC 15703 GCA_000010425.1
115 Bifidobacterium subtile GCA_000741775.1
114 Bifidobacterium callitrichos DSM 23973 GCA_000741175.1
EX
222 Escherichia coli UMN026 GCA_000026325.2
222 Shigella flexneri 2a str. 301 GCA_000006925.2
203 Edwardsiella anguillarum ET080813 GCA_000264765.2
201 Erwinia iniecta GCA_001267535.1
201 Escherichia coli IAI39 GCA_000026345.1
201 Escherichia coli O104_3AH4 str. 2011C-3493 GCA_000299455.1
201 Escherichia coli O157_3AH7 str. Sakai GCA_000008865.1
201 Escherichia coli O83_3AH1 str. NRG 857C GCA_000183345.1
201 Escherichia coli str. K-12 substr. MG1655 GCA_000005845.2
201 Plesiomonas shigelloides GCA_900087055.1
201 Rahnella aquatilis HX2 GCA_000255535.1
201 Rosenbergiella nectarea GCA_900111105.1
201 Rouxiella chamberiensis GCA_000951135.1
201 Serratia fonticola GCA_001006005.1
201 Serratia liquefaciens ATCC 27592 GCA_000422085.1
201 Serratia marcescens subsp. marcescens Db11 GCA_000513215.1
201 Serratia rubidaea GCA_001572725.1
201 Serratia symbiotica GCA_000821185.1
201 Shigella dysenteriae Sd197 GCA_000012005.1
201 Tumebacillus flagellatus GCA_000714935.1
201 Yersinia enterocolitica subsp. enterocolitica 8081 GCA_000009345.1
201 Yersinia pestis CO92 GCA_000009065.1
201 Yersinia ruckeri GCA_000964565.1
EY
159 Rhodobacter sphaeroides 2.4.1 GCA_000012905.2
127 Rhodobacter capsulatus SB 1003 GCA_000021865.1
126 Defluviimonas alba GCA_001620265.1
126 Gemmobacter megaterium GCA_900156815.1
126 Pseudorhodobacter ferrugineus DSM 5888 GCA_000420745.1
126 Pseudorhodobacter psychrotolerans GCA_001294535.1
126 Pseudorhodobacter wandonensis GCA_001202035.1

126 Rhodobacter sphaeroides ATCC 17025 GCA_000016405.1
EZ
182 Rhodobacter sphaeroides 2.4.1 GCA_000012905.2
170 Thioclava dalianensis GCA_000715505.1
170 Thioclava indica GCA_000714545.1
168 Defluviimonas alba GCA_001620265.1
168 Gemmobacter megaterium GCA_900156815.1
168 Pseudorhodobacter ferrugineus DSM 5888 GCA_000420745.1
168 Pseudorhodobacter wandonensis GCA_001202035.1
168 Rhodobacter sphaeroides ATCC 17025 GCA_000016405.1
F0
215 Clostridium beijerinckii GCA_000833105.2
215 Clostridium puniceum GCA_002006345.1
215 Clostridium saccharobutylicum DSM 13864 GCA_000473995.1
215 Clostridium saccharoperbutylacetonicum N1-4_28HMT_29 GCA_000340885.1
185 Clostridium chromiireducens GCA_002029255.1
176 Clostridium neonatale GCA_001458595.1
F1
238 Clostridium beijerinckii GCA_000833105.2
238 Clostridium puniceum GCA_002006345.1
238 Clostridium saccharoperbutylacetonicum N1-4_28HMT_29 GCA_000340885.1
229 Clostridium saccharobutylicum DSM 13864 GCA_000473995.1
213 Clostridium chromiireducens GCA_002029255.1
F2
230 Bacillus anthracis str. Ames GCA_000007845.1
230 Bacillus anthracis str. Sterne GCA_000008165.1
230 Bacillus mycoides GCA_000832605.1
230 Bacillus pseudomycoides DSM 12442 GCA_000161455.1
230 Bacillus thuringiensis YBT-1518 GCA_000497525.2
230 _5BBacillus thuringiensis_5D serovar konkukian str. 97-27 GCA_000008505.1
229 Bacillus cereus ATCC 14579 GCA_000007825.1
190 Bacillus vietnamensis NBRC 101237 GCA_001591825.1
F3
220 Escherichia coli IAI39 GCA_000026345.1
220 Escherichia coli O104_3AH4 str. 2011C-3493 GCA_000299455.1
220 Escherichia coli O157_3AH7 str. Sakai GCA_000008865.1
220 Escherichia coli O83_3AH1 str. NRG 857C GCA_000183345.1
220 Escherichia coli UMN026 GCA_000026325.2
220 Escherichia coli str. K-12 substr. MG1655 GCA_000005845.2
220 Shigella flexneri 2a str. 301 GCA_000006925.2
220 Tumebacillus flagellatus GCA_000714935.1
205 Shigella dysenteriae Sd197 GCA_000012005.1
201 Erwinia iniecta GCA_001267535.1
F4
235 Rhodobacter sphaeroides 2.4.1 GCA_000012905.2
235 Rhodobacter sphaeroides ATCC 17025 GCA_000016405.1
216 Pseudorhodobacter ferrugineus DSM 5888 GCA_000420745.1
216 Pseudorhodobacter wandonensis GCA_001202035.1
206 Gemmobacter megaterium GCA_900156815.1
206 Pseudorhodobacter psychrotolerans GCA_001294535.1
F5
215 Bacillus anthracis str. Ames GCA_000007845.1
215 Bacillus anthracis str. Sterne GCA_000008165.1
215 Bacillus cereus ATCC 14579 GCA_000007825.1
215 Bacillus mycoides GCA_000832605.1
215 Bacillus pseudomycoides DSM 12442 GCA_000161455.1
215 Bacillus thuringiensis YBT-1518 GCA_000497525.2
215 _5BBacillus thuringiensis_5D serovar konkukian str. 97-27 GCA_000008505.1
202 Bacillus marisflavi GCA_001274775.1
201 Bacillus manliponensis GCA_000712595.1
F6
209 Bacillus anthracis str. Ames GCA_000007845.1
209 Bacillus anthracis str. Sterne GCA_000008165.1
209 Bacillus cereus ATCC 14579 GCA_000007825.1
209 Bacillus mycoides GCA_000832605.1
209 Bacillus pseudomycoides DSM 12442 GCA_000161455.1
209 Bacillus thuringiensis YBT-1518 GCA_000497525.2
209 _5BBacillus thuringiensis_5D serovar konkukian str. 97-27 GCA_000008505.1
138 Bacillus horneckiae GCA_001636335.1
138 Bacillus solani GCA_001420595.1
133 Anaerobacillus alkalilacustris GCA_001866005.1
133 Bacillus manliponensis GCA_000712595.1
133 Isobaculum melis GCA_900111355.1
133 Massilibacterium senegalense GCA_001375675.1
133 Planococcus antarcticus DSM 14505 GCA_001687565.2
F7

238 Staphylococcus epidermidis ATCC 12228 GCA_000007645.1
238 Staphylococcus haemolyticus JCSC1435 GCA_000009865.1
230 Staphylococcus capitis subsp. capitis GCA_001028645.1
227 Staphylococcus hominis subsp. hominis C80 GCA_000183685.1
F8
166 Deinococcus radiodurans R1 GCA_000008565.1
102 Deinococcus deserti VCD115 GCA_000020685.1
96 Deinococcus gobiensis I-0 GCA_000252445.1
F9
258 Escherichia coli IAI39 GCA_000026345.1
258 Escherichia coli O104_3AH4 str. 2011C-3493 GCA_000299455.1
258 Escherichia coli O157_3AH7 str. Sakai GCA_000008865.1
258 Escherichia coli O83_3AH1 str. NRG 857C GCA_000183345.1
258 Escherichia coli UMN026 GCA_000026325.2
258 Escherichia coli str. K-12 substr. MG1655 GCA_000005845.2
258 Shigella flexneri 2a str. 301 GCA_000006925.2
258 Tumebacillus flagellatus GCA_000714935.1
249 Shigella dysenteriae Sd197 GCA_000012005.1
244 Erwinia iniecta GCA_001267535.1
Fa
268 Escherichia coli IAI39 GCA_000026345.1
268 Escherichia coli O104_3AH4 str. 2011C-3493 GCA_000299455.1
268 Escherichia coli O157_3AH7 str. Sakai GCA_000008865.1
268 Escherichia coli O83_3AH1 str. NRG 857C GCA_000183345.1
268 Escherichia coli UMN026 GCA_000026325.2
268 Escherichia coli str. K-12 substr. MG1655 GCA_000005845.2
268 Shigella dysenteriae Sd197 GCA_000012005.1
268 Shigella flexneri 2a str. 301 GCA_000006925.2
268 Tumebacillus flagellatus GCA_000714935.1
236 Erwinia iniecta GCA_001267535.1
231 Rosenbergiella nectarea GCA_900111105.1
Fb
219 Rhodobacter sphaeroides 2.4.1 GCA_000012905.2
204 Rhodobacter sphaeroides ATCC 17025 GCA_000016405.1
199 Pseudorhodobacter psychrotolerans GCA_001294535.1
Fc
250 Lactobacillus gasseri ATCC 33323 = JCM 1131 GCA_000014425.1
250 Lactobacillus hominis DSM 23910 = CRBIP 24.179 GCA_000296835.1
203 Lactobacillus acidophilus NCFM GCA_000011985.1
202 Lactobacillus crispatus ST1 GCA_000091765.1
202 Lactobacillus hamsteri DSM 5661 = JCM 6256 GCA_000615445.1
Fd
244 Bifidobacterium adolescentis ATCC 15703 GCA_000010425.1
232 Bifidobacterium thermophilum GCA_000741495.1
217 Bifidobacterium tsurumiense GCA_000741765.1
Fe
113 Rhodobacter sphaeroides 2.4.1 GCA_000012905.2
83 Defluviimonas alba GCA_001620265.1
79 Thioclava dalianensis GCA_000715505.1
79 Thioclava indica GCA_000714545.1
Ff
219 Staphylococcus epidermidis ATCC 12228 GCA_000007645.1
219 Staphylococcus haemolyticus JCSC1435 GCA_000009865.1
214 Staphylococcus capitis subsp. capitis GCA_001028645.1
203 Staphylococcus aureus subsp. aureus NCTC 8325 GCA_000013425.1
203 Staphylococcus condimenti GCA_001618885.1
203 Staphylococcus hominis subsp. hominis C80 GCA_000183685.1
203 Staphylococcus simiae CCM 7213 GCA_000235645.2
203 Staphylococcus simulans GCA_001559115.1
Fg
176 Staphylococcus pseudintermedius HKU10-03 GCA_000185885.1
161 Megasphaera cerevisiae DSM 20462 GCA_001045675.1
161 Staphylococcus aureus subsp. aureus NCTC 8325 GCA_000013425.1
161 Staphylococcus capitis subsp. capitis GCA_001028645.1
161 Staphylococcus condimenti GCA_001618885.1
161 Staphylococcus epidermidis ATCC 12228 GCA_000007645.1
161 Staphylococcus haemolyticus JCSC1435 GCA_000009865.1
161 Staphylococcus hominis subsp. hominis C80 GCA_000183685.1
161 Staphylococcus hyicus GCA_000816085.1
161 Staphylococcus lugdunensis HKU09-01 GCA_000025085.1
161 Staphylococcus lutrae GCA_002101335.1
161 Staphylococcus microti GCA_000934465.1
161 Staphylococcus simiae CCM 7213 GCA_000235645.2
161 Staphylococcus simulans GCA_001559115.1
161 Staphylococcus warneri SG1 GCA_000332735.1
146 Staphylococcus pettenkoferi GCA_002208805.1

Fh
278 Deinococcus radiodurans R1 GCA_000008565.1
196 Deinococcus gobiensis I-0 GCA_000252445.1
158 Deinococcus puniceus GCA_001644565.1
Fi
243 Clostridium beijerinckii GCA_000833105.2
243 Clostridium saccharoperbutylacetonicum N1-4_28HMT_29 GCA_000340885.1
236 Clostridium puniceum GCA_002006345.1
229 Clostridium saccharobutylicum DSM 13864 GCA_000473995.1
Fj
100 Escherichia coli IAI39 GCA_000026345.1
100 Escherichia coli O104_3AH4 str. 2011C-3493 GCA_000299455.1
100 Escherichia coli O157_3AH7 str. Sakai GCA_000008865.1
100 Escherichia coli O83_3AH1 str. NRG 857C GCA_000183345.1
100 Escherichia coli UMN026 GCA_000026325.2
100 Escherichia coli str. K-12 substr. MG1655 GCA_000005845.2
100 Shigella dysenteriae Sd197 GCA_000012005.1
100 Shigella flexneri 2a str. 301 GCA_000006925.2
100 Tumebacillus flagellatus GCA_000714935.1
84 Bacillus humi GCA_001439915.1
72 Brenneria goodwinii GCA_001049335.1
72 Cronobacter sakazakii GCA_000982825.1
72 Dickeya solani IPO 2222 GCA_001644705.1
72 Dickeya zeae Ech586 GCA_000025065.1
72 Edwardsiella anguillarum ET080813 GCA_000264765.2
72 Enterobacter cloacae subsp. cloacae ATCC 13047 GCA_000025565.1
72 Enterobacter hormaechei subsp. steigerwaltii GCA_001729725.1
72 Enterobacter kobei GCA_900185885.1
72 Erwinia amylovora CFBP1430 GCA_000091565.1
72 Erwinia billingiae Eb661 GCA_000196615.1
72 Erwinia gerundensis GCA_001517405.1
72 Erwinia iniecta GCA_001267535.1
72 Erwinia oleae GCA_000770305.1
72 Erwinia persicina NBRC 102418 GCA_001571305.1
72 Erwinia toletana DAPP-PG 735 GCA_000336255.1
72 Izhakiella capsodis GCA_900115045.1
72 Klebsiella oxytoca GCA_001022195.1
72 Klebsiella pneumoniae subsp. pneumoniae HS11286 GCA_000240185.2
72 Kosakonia cowanii GCA_001975225.1
72 Kosakonia sacchari SP1 GCA_000300455.4
72 Lonsdalea quercina subsp. quercina GCA_000688655.1
72 Mangrovibacter phragmitis GCA_001655675.1
72 Pantoea agglomerans GCA_001709315.1
72 Pantoea alhagi GCA_002101395.1
72 Pantoea dispersa EGD-AAK13 GCA_000465555.2
72 Pantoea rwandensis GCA_000759475.1
72 Pantoea septica GCA_002095575.1
72 Pectobacterium carotovorum subsp. carotovorum PC1 GCA_000023605.1
72 Plautia stali symbiont GCA_000180175.2
72 Pluralibacter gergoviae GCA_000757785.1
72 Pseudescherichia vulneris NBRC 102420 GCA_000759795.1
72 Rosenbergiella nectarea GCA_900111105.1
72 Salmonella enterica subsp. enterica serovar Typhi str. CT18 GCA_000195995.1
72 Shimwellia blattae DSM 4481 = NBRC 105725 GCA_000262305.1
72 Tatumella citrea GCA_002163585.1
72 Trabulsiella odontotermitis GCA_001297765.1
Fk
188 Escherichia coli IAI39 GCA_000026345.1
188 Escherichia coli O104_3AH4 str. 2011C-3493 GCA_000299455.1
188 Escherichia coli O157_3AH7 str. Sakai GCA_000008865.1
188 Escherichia coli O83_3AH1 str. NRG 857C GCA_000183345.1
188 Escherichia coli UMN026 GCA_000026325.2
188 Escherichia coli str. K-12 substr. MG1655 GCA_000005845.2
188 Shigella dysenteriae Sd197 GCA_000012005.1
188 Shigella flexneri 2a str. 301 GCA_000006925.2
188 Tumebacillus flagellatus GCA_000714935.1
149 Klebsiella aerogenes KCTC 2190 GCA_000215745.1
149 Klebsiella pneumoniae subsp. pneumoniae HS11286 GCA_000240185.2
148 Buttiauxella ferragutiae ATCC 51602 GCA_001654915.1
148 Cedecea neteri GCA_000757825.1
148 Citrobacter freundii CFNIH1 GCA_000648515.1
148 Cronobacter sakazakii GCA_000982825.1
148 Enterobacter cloacae subsp. cloacae ATCC 13047 GCA_000025565.1
148 Enterobacter hormaechei subsp. steigerwaltii GCA_001729725.1
148 Enterobacter kobei GCA_900185885.1
148 Erwinia amylovora CFBP1430 GCA_000091565.1

148 Erwinia billingiae Eb661 GCA_000196615.1
148 Erwinia gerundensis GCA_001517405.1
148 Erwinia iniecta GCA_001267535.1
148 Erwinia persicina NBRC 102418 GCA_001571305.1
148 Erwinia toletana DAPP-PG 735 GCA_000336255.1
148 Klebsiella oxytoca GCA_001022195.1
148 Kluyvera ascorbata ATCC 33433 GCA_000735365.1
148 Kosakonia cowanii GCA_001975225.1
148 Obesumbacterium proteus GCA_001586165.1
148 Pantoea agglomerans GCA_001709315.1
148 Pantoea alhagi GCA_002101395.1
148 Pantoea ananatis LMG 20103 GCA_000025405.2
148 Pantoea dispersa EGD-AAK13 GCA_000465555.2
148 Pantoea rwandensis GCA_000759475.1
148 Pantoea septica GCA_002095575.1
148 Plesiomonas shigelloides GCA_900087055.1
148 Pluralibacter gergoviae GCA_000757785.1
148 Pseudescherichia vulneris NBRC 102420 GCA_000759795.1
148 Rosenbergiella nectarea GCA_900111105.1
148 Salmonella enterica subsp. enterica serovar Typhi str. CT18 GCA_000195995.1
148 Serratia fonticola GCA_001006005.1
148 Serratia liquefaciens ATCC 27592 GCA_000422085.1
148 Serratia marcescens subsp. marcescens Db11 GCA_000513215.1
148 Serratia rubidaea GCA_001572725.1
148 Serratia symbiotica GCA_000821185.1
148 Shimwellia blattae DSM 4481 = NBRC 105725 GCA_000262305.1
148 Tatumella citrea GCA_002163585.1
148 Tatumella saanichensis GCA_000439375.1
148 Trabulsiella odontotermitis GCA_001297765.1
148 Yersinia enterocolitica subsp. enterocolitica 8081 GCA_000009345.1
148 Yersinia pestis CO92 GCA_000009065.1
148 Yersinia ruckeri GCA_000964565.1
148 Yokenella regensburgei ATCC 49455 GCA_000735455.1
148 _5BEnterobacter_5D lignolyticus SCF1 GCA_000164865.1
Fl
329 Staphylococcus epidermidis ATCC 12228 GCA_000007645.1
329 Staphylococcus haemolyticus JCSC1435 GCA_000009865.1
318 Staphylococcus hominis subsp. hominis C80 GCA_000183685.1
318 Staphylococcus lugdunensis HKU09-01 GCA_000025085.1
310 Staphylococcus simulans GCA_001559115.1
Fm
132 Streptococcus mutans UA159 GCA_000007465.2
98 Streptococcus marimammalium DSM 18627 GCA_000380045.1
83 Streptococcus sobrinus DSM 20742 = ATCC 33478 GCA_000686605.1
Fn
135 Clostridium beijerinckii GCA_000833105.2
135 Clostridium botulinum B str. Eklund 17B _28NRP_29 GCA_000020165.1
135 Clostridium chromiireducens GCA_002029255.1
135 Clostridium puniceum GCA_002006345.1
135 Clostridium saccharobutylicum DSM 13864 GCA_000473995.1
135 Clostridium saccharoperbutylacetonicum N1-4_28HMT_29 GCA_000340885.1
135 Clostridium taeniosporum GCA_001735765.1
135 Clostridium uliginosum GCA_900112485.1
133 Clostridium neonatale GCA_001458595.1
130 Clostridium botulinum A str. ATCC 3502 GCA_000063585.1
130 Clostridium botulinum A str. Hall GCA_000017045.1
130 Clostridium butyricum GCA_001456065.2
130 Clostridium tepidum GCA_002008345.1
Fo
167 Bifidobacterium adolescentis ATCC 15703 GCA_000010425.1
161 Bifidobacterium tsurumiense GCA_000741765.1
152 Bifidobacterium choerinum GCA_000741135.1
152 Bifidobacterium pseudolongum PV8-2 GCA_000800475.2
Fp
132 Staphylococcus epidermidis ATCC 12228 GCA_000007645.1
132 Staphylococcus haemolyticus JCSC1435 GCA_000009865.1
126 Staphylococcus capitis subsp. capitis GCA_001028645.1
118 Megasphaera cerevisiae DSM 20462 GCA_001045675.1
118 Staphylococcus aureus subsp. aureus NCTC 8325 GCA_000013425.1
118 Staphylococcus hominis subsp. hominis C80 GCA_000183685.1
118 Staphylococcus lugdunensis HKU09-01 GCA_000025085.1
118 Staphylococcus simiae CCM 7213 GCA_000235645.2
118 Staphylococcus warneri SG1 GCA_000332735.1
Fq
224 Clostridium beijerinckii GCA_000833105.2
224 Clostridium puniceum GCA_002006345.1

224 Clostridium saccharobutylicum DSM 13864 GCA_000473995.1
224 Clostridium saccharoperbutylacetonicum N1-4_28HMT_29 GCA_000340885.1
212 Clostridium neonatale GCA_001458595.1
204 Clostridium chromiireducens GCA_002029255.1
Fr
261 Staphylococcus epidermidis ATCC 12228 GCA_000007645.1
261 Staphylococcus haemolyticus JCSC1435 GCA_000009865.1
261 Staphylococcus hominis subsp. hominis C80 GCA_000183685.1
261 Staphylococcus lugdunensis HKU09-01 GCA_000025085.1
255 Staphylococcus capitis subsp. capitis GCA_001028645.1
255 Staphylococcus microti GCA_000934465.1
246 Staphylococcus aureus subsp. aureus NCTC 8325 GCA_000013425.1
246 Staphylococcus condimenti GCA_001618885.1
246 Staphylococcus simiae CCM 7213 GCA_000235645.2
246 Staphylococcus simulans GCA_001559115.1
Fs
94 Enterobacter cloacae subsp. cloacae ATCC 13047 GCA_000025565.1
86 Serratia symbiotica GCA_000821185.1
70 Aeromonas eucrenophila GCA_000819865.1
70 Aeromonas fluvialis GCA_000819885.1
70 Aeromonas hydrophila subsp. hydrophila ATCC 7966 GCA_000014805.1
70 Aeromonas salmonicida subsp. salmonicida A449 GCA_000196395.1
70 Aeromonas schubertii GCA_001447335.1
70 Aeromonas simiae GCA_000820125.1
70 Bacillus humi GCA_001439915.1
70 Brenneria goodwinii GCA_001049335.1
70 Buttiauxella ferragutiae ATCC 51602 GCA_001654915.1
70 Cedecea neteri GCA_000757825.1
70 Citrobacter freundii CFNIH1 GCA_000648515.1
70 Cronobacter sakazakii GCA_000982825.1
70 Dickeya solani IPO 2222 GCA_001644705.1
70 Dickeya zeae Ech586 GCA_000025065.1
70 Edwardsiella anguillarum ET080813 GCA_000264765.2
70 Enterobacter hormaechei subsp. steigerwaltii GCA_001729725.1
70 Enterobacter kobei GCA_900185885.1
70 Erwinia amylovora CFBP1430 GCA_000091565.1
70 Erwinia billingiae Eb661 GCA_000196615.1
70 Erwinia gerundensis GCA_001517405.1
70 Erwinia iniecta GCA_001267535.1
70 Erwinia oleae GCA_000770305.1
70 Erwinia persicina NBRC 102418 GCA_001571305.1
70 Erwinia toletana DAPP-PG 735 GCA_000336255.1
70 Escherichia coli IAI39 GCA_000026345.1
70 Escherichia coli O104_3AH4 str. 2011C-3493 GCA_000299455.1
70 Escherichia coli O157_3AH7 str. Sakai GCA_000008865.1
70 Escherichia coli O83_3AH1 str. NRG 857C GCA_000183345.1
70 Escherichia coli UMN026 GCA_000026325.2
70 Escherichia coli str. K-12 substr. MG1655 GCA_000005845.2
70 Izhakiella capsodis GCA_900115045.1
70 Klebsiella aerogenes KCTC 2190 GCA_000215745.1
70 Klebsiella oxytoca GCA_001022195.1
70 Klebsiella pneumoniae subsp. pneumoniae HS11286 GCA_000240185.2
70 Kluyvera ascorbata ATCC 33433 GCA_000735365.1
70 Kluyvera cryocrescens NBRC 102467 GCA_001571285.1
70 Kosakonia cowanii GCA_001975225.1
70 Kosakonia sacchari SP1 GCA_000300455.4
70 Lonsdalea quercina subsp. quercina GCA_000688655.1
70 Mangrovibacter phragmitis GCA_001655675.1
70 Nissabacter archeti GCA_900130115.1
70 Obesumbacterium proteus GCA_001586165.1
70 Pantoea agglomerans GCA_001709315.1
70 Pantoea alhagi GCA_002101395.1
70 Pantoea ananatis LMG 20103 GCA_000025405.2
70 Pantoea dispersa EGD-AAK13 GCA_000465555.2
70 Pantoea rwandensis GCA_000759475.1
70 Pantoea septica GCA_002095575.1
70 Pectobacterium carotovorum subsp. carotovorum PC1 GCA_000023605.1
70 Plautia stali symbiont GCA_000180175.2
70 Plesiomonas shigelloides GCA_900087055.1
70 Pseudescherichia vulneris NBRC 102420 GCA_000759795.1
70 Rahnella aquatilis HX2 GCA_000255535.1
70 Rosenbergiella nectarea GCA_900111105.1
70 Rouxiella chamberiensis GCA_000951135.1
70 Salmonella enterica subsp. enterica serovar Typhi str. CT18 GCA_000195995.1
70 Serratia fonticola GCA_001006005.1
70 Serratia liquefaciens ATCC 27592 GCA_000422085.1

70 Serratia marcescens subsp. marcescens Db11 GCA_000513215.1
70 Serratia rubidaea GCA_001572725.1
70 Shigella dysenteriae Sd197 GCA_000012005.1
70 Shigella flexneri 2a str. 301 GCA_000006925.2
70 Shimwellia blattae DSM 4481 = NBRC 105725 GCA_000262305.1
70 Tatumella citrea GCA_002163585.1
70 Tatumella saanichensis GCA_000439375.1
70 Tolumonas auensis DSM 9187 GCA_000023065.1
70 Trabulsiella odontotermitis GCA_001297765.1
70 Tumebacillus flagellatus GCA_000714935.1
70 Yersinia enterocolitica subsp. enterocolitica 8081 GCA_000009345.1
70 Yersinia pestis CO92 GCA_000009065.1
70 Yersinia ruckeri GCA_000964565.1
70 Yokenella regensburgei ATCC 49455 GCA_000735455.1
Ft
223 Clostridium beijerinckii GCA_000833105.2
223 Clostridium puniceum GCA_002006345.1
223 Clostridium saccharobutylicum DSM 13864 GCA_000473995.1
223 Clostridium saccharoperbutylacetonicum N1-4_28HMT_29 GCA_000340885.1
208 Clostridium chromiireducens GCA_002029255.1
206 Clostridium intestinale URNW GCA_000469625.2
Fu
309 Lactobacillus gasseri ATCC 33323 = JCM 1131 GCA_000014425.1
275 Lactobacillus hominis DSM 23910 = CRBIP 24.179 GCA_000296835.1
208 Lactobacillus iners DSM 13335 GCA_000160875.1
Fv
141 Deinococcus radiodurans R1 GCA_000008565.1
109 Deinococcus gobiensis I-0 GCA_000252445.1
108 Deinococcus frigens DSM 12807 GCA_000701425.1
Fw
180 Bifidobacterium adolescentis ATCC 15703 GCA_000010425.1
166 Bifidobacterium thermophilum GCA_000741495.1
166 Bifidobacterium thermophilum RBL67 GCA_000347695.1
151 Bifidobacterium dentium JCM 1195 = DSM 20436 GCA_001042595.1
151 Bifidobacterium tsurumiense GCA_000741765.1
Fx
263 Lactobacillus gasseri ATCC 33323 = JCM 1131 GCA_000014425.1
259 Lactobacillus hominis DSM 23910 = CRBIP 24.179 GCA_000296835.1
215 Lactobacillus crispatus ST1 GCA_000091765.1
215 Lactobacillus helveticus GCA_001308285.1
Fy
253 Enterococcus faecalis V583 GCA_000007785.1
253 Streptomyces cinnamoneus GCA_001885705.1
237 Enterococcus canis NBRC 100695 GCA_001544375.1
237 Enterococcus casseliflavus EC20 GCA_000157355.2
237 Enterococcus dispar ATCC 51266 GCA_000406945.1
237 Enterococcus faecium DO GCA_000174395.2
237 Enterococcus haemoperoxidus ATCC BAA-382 GCA_000407165.1
237 Enterococcus hirae ATCC 9790 GCA_000271405.2
237 Enterococcus mundtii QU 25 GCA_000504125.1
237 Enterococcus phoeniculicola ATCC BAA-412 GCA_000407505.1
237 Enterococcus rivorum GCA_001742285.1
237 Enterococcus saccharolyticus subsp. saccharolyticus ATCC 43076 GCA_000407285.1
237 Enterococcus thailandicus GCA_001652875.1
227 Enterococcus asini ATCC 700915 GCA_000407365.1
Fz
176 Streptococcus mutans UA159 GCA_000007465.2
96 Streptococcus sobrinus DSM 20742 = ATCC 33478 GCA_000686605.1
85 Streptococcus gordonii str. Challis substr. CH1 GCA_000017005.1
85 Streptococcus phocae subsp. salmonis GCA_000772915.1
FA
276 Bifidobacterium adolescentis ATCC 15703 GCA_000010425.1
234 Bifidobacterium breve DSM 20213 = JCM 1192 GCA_001025175.1
234 Bifidobacterium dentium JCM 1195 = DSM 20436 GCA_001042595.1
230 Bifidobacterium gallicum DSM 20093 = LMG 11596 GCA_000741205.1
FB
157 Escherichia coli IAI39 GCA_000026345.1
157 Escherichia coli O104_3AH4 str. 2011C-3493 GCA_000299455.1
157 Escherichia coli O157_3AH7 str. Sakai GCA_000008865.1
157 Escherichia coli O83_3AH1 str. NRG 857C GCA_000183345.1
157 Escherichia coli UMN026 GCA_000026325.2
157 Escherichia coli str. K-12 substr. MG1655 GCA_000005845.2
157 Shigella dysenteriae Sd197 GCA_000012005.1
157 Shigella flexneri 2a str. 301 GCA_000006925.2
157 Tumebacillus flagellatus GCA_000714935.1
139 Erwinia iniecta GCA_001267535.1

136 Enterobacter hormaechei subsp. steigerwaltii GCA_001729725.1
136 Erwinia toletana DAPP-PG 735 GCA_000336255.1
136 Klebsiella oxytoca GCA_001022195.1
136 Kosakonia cowanii GCA_001975225.1
136 Kosakonia sacchari SP1 GCA_000300455.4
136 Pluralibacter gergoviae GCA_000757785.1
136 Pseudescherichia vulneris NBRC 102420 GCA_000759795.1
FC
235 Enterococcus faecalis V583 GCA_000007785.1
235 Streptomyces cinnamoneus GCA_001885705.1
206 Isobaculum melis GCA_900111355.1
205 Enterococcus canis NBRC 100695 GCA_001544375.1
205 Enterococcus casseliflavus EC20 GCA_000157355.2
205 Enterococcus dispar ATCC 51266 GCA_000406945.1
205 Enterococcus faecium DO GCA_000174395.2
205 Enterococcus haemoperoxidus ATCC BAA-382 GCA_000407165.1
205 Enterococcus hirae ATCC 9790 GCA_000271405.2
205 Enterococcus massiliensis GCA_001050095.1
205 Enterococcus mundtii QU 25 GCA_000504125.1
205 Enterococcus phoeniculicola ATCC BAA-412 GCA_000407505.1
205 Enterococcus rivorum GCA_001742285.1
205 Enterococcus saccharolyticus subsp. saccharolyticus ATCC 43076 GCA_000407285.1
205 Enterococcus thailandicus GCA_001652875.1
FD
307 Bifidobacterium adolescentis ATCC 15703 GCA_000010425.1
287 Bifidobacterium callitrichos DSM 23973 GCA_000741175.1
279 Bifidobacterium asteroides PRL2011 GCA_000304215.1
FE
216 Streptococcus mutans UA159 GCA_000007465.2
142 Streptococcus equinus GCA_000964315.1
142 Streptococcus gallolyticus subsp. gallolyticus DSM 16831 GCA_002000985.1
139 Streptococcus cristatus AS 1.3089 GCA_000385925.1
139 Streptococcus gordonii str. Challis substr. CH1 GCA_000017005.1
139 Streptococcus mitis B6 GCA_000027165.1
139 Streptococcus pneumoniae R6 GCA_000007045.1
FF
167 Clostridium neonatale GCA_001458595.1
165 Clostridium beijerinckii GCA_000833105.2
165 Clostridium saccharobutylicum DSM 13864 GCA_000473995.1
165 Clostridium saccharoperbutylacetonicum N1-4_28HMT_29 GCA_000340885.1
161 Clostridium puniceum GCA_002006345.1
FG
273 Escherichia coli IAI39 GCA_000026345.1
273 Escherichia coli O104_3AH4 str. 2011C-3493 GCA_000299455.1
273 Escherichia coli O157_3AH7 str. Sakai GCA_000008865.1
273 Escherichia coli UMN026 GCA_000026325.2
273 Escherichia coli str. K-12 substr. MG1655 GCA_000005845.2
273 Shigella dysenteriae Sd197 GCA_000012005.1
273 Shigella flexneri 2a str. 301 GCA_000006925.2
273 Tumebacillus flagellatus GCA_000714935.1
272 Escherichia coli O83_3AH1 str. NRG 857C GCA_000183345.1
247 Erwinia iniecta GCA_001267535.1
FH
196 Deinococcus radiodurans R1 GCA_000008565.1
111 Deinococcus puniceus GCA_001644565.1
106 Deinococcus deserti VCD115 GCA_000020685.1
106 Deinococcus hopiensis KR-140 GCA_900176165.1
FI
198 Rhodobacter sphaeroides 2.4.1 GCA_000012905.2
177 Haematobacter massiliensis GCA_000740795.1
175 Gemmobacter aquatilis GCA_900110025.1
175 Gemmobacter nectariphilus DSM 15620 GCA_000429765.1
175 Pseudorhodobacter psychrotolerans GCA_001294535.1
175 Rhodobacter capsulatus SB 1003 GCA_000021865.1
FJ
241 Clostridium beijerinckii GCA_000833105.2
241 Clostridium saccharobutylicum DSM 13864 GCA_000473995.1
241 Clostridium saccharoperbutylacetonicum N1-4_28HMT_29 GCA_000340885.1
234 Clostridium puniceum GCA_002006345.1
204 Clostridium chromiireducens GCA_002029255.1
FK
215 Clostridium beijerinckii GCA_000833105.2
215 Clostridium saccharobutylicum DSM 13864 GCA_000473995.1
215 Clostridium saccharoperbutylacetonicum N1-4_28HMT_29 GCA_000340885.1
208 Clostridium puniceum GCA_002006345.1
197 Clostridium chromiireducens GCA_002029255.1

FL
247 Bacillus anthracis str. Ames GCA_000007845.1
247 Bacillus anthracis str. Sterne GCA_000008165.1
247 Bacillus cereus ATCC 14579 GCA_000007825.1
247 Bacillus mycoides GCA_000832605.1
247 Bacillus pseudomycoides DSM 12442 GCA_000161455.1
247 Bacillus thuringiensis YBT-1518 GCA_000497525.2
247 _5BBacillus thuringiensis_5D serovar konkukian str. 97-27 GCA_000008505.1
187 Bacillus tuaregi GCA_900104575.1
179 Bacillus cytotoxicus NVH 391-98 GCA_000017425.1
FM
215 Deinococcus radiodurans R1 GCA_000008565.1
139 Deinococcus marmoris DSM 12784 GCA_000701405.1
139 Deinococcus swuensis GCA_000800395.1
134 Deinococcus deserti VCD115 GCA_000020685.1
134 Deinococcus gobiensis I-0 GCA_000252445.1
FN
249 Lactobacillus gasseri ATCC 33323 = JCM 1131 GCA_000014425.1
215 Lactobacillus hominis DSM 23910 = CRBIP 24.179 GCA_000296835.1
175 Lactobacillus amylophilus DSM 20533 = JCM 1125 GCA_001936335.1
FO
226 Escherichia coli IAI39 GCA_000026345.1
226 Escherichia coli O104_3AH4 str. 2011C-3493 GCA_000299455.1
226 Escherichia coli O157_3AH7 str. Sakai GCA_000008865.1
226 Escherichia coli O83_3AH1 str. NRG 857C GCA_000183345.1
226 Escherichia coli UMN026 GCA_000026325.2
226 Escherichia coli str. K-12 substr. MG1655 GCA_000005845.2
226 Shigella dysenteriae Sd197 GCA_000012005.1
226 Shigella flexneri 2a str. 301 GCA_000006925.2
226 Tumebacillus flagellatus GCA_000714935.1
203 Erwinia iniecta GCA_001267535.1
188 Cronobacter sakazakii GCA_000982825.1
188 Erwinia toletana DAPP-PG 735 GCA_000336255.1
188 Rosenbergiella nectarea GCA_900111105.1
FP
163 Deinococcus radiodurans R1 GCA_000008565.1
103 Deinococcus geothermalis DSM 11300 GCA_000196275.1
101 Deinococcus puniceus GCA_001644565.1
FQ
275 Streptococcus mutans UA159 GCA_000007465.2
157 Streptococcus sobrinus DSM 20742 = ATCC 33478 GCA_000686605.1
156 Streptococcus ferus DSM 20646 GCA_000372425.1
FR
186 Clostridium beijerinckii GCA_000833105.2
186 Clostridium puniceum GCA_002006345.1
186 Clostridium saccharoperbutylacetonicum N1-4_28HMT_29 GCA_000340885.1
178 Clostridium taeniosporum GCA_001735765.1
177 Clostridium saccharobutylicum DSM 13864 GCA_000473995.1
FS
206 Lactobacillus gasseri ATCC 33323 = JCM 1131 GCA_000014425.1
206 Lactobacillus hominis DSM 23910 = CRBIP 24.179 GCA_000296835.1
149 Lactobacillus acidophilus NCFM GCA_000011985.1
149 Lactobacillus gallinarum GCA_001314245.2
149 Lactobacillus helveticus GCA_001308285.1
143 Lactobacillus crispatus ST1 GCA_000091765.1
143 Lactobacillus hamsteri DSM 5661 = JCM 6256 GCA_000615445.1
FT
297 Bacillus anthracis str. Ames GCA_000007845.1
297 Bacillus anthracis str. Sterne GCA_000008165.1
297 Bacillus cereus ATCC 14579 GCA_000007825.1
297 Bacillus mycoides GCA_000832605.1
297 Bacillus pseudomycoides DSM 12442 GCA_000161455.1
297 Bacillus thuringiensis YBT-1518 GCA_000497525.2
297 _5BBacillus thuringiensis_5D serovar konkukian str. 97-27 GCA_000008505.1
237 Bacillus marisflavi GCA_001274775.1
226 Bacillus acidicola GCA_001636425.1
226 Bacillus horneckiae GCA_001636335.1
226 Bacillus shackletonii GCA_001420715.1
226 Bacillus solani GCA_001420595.1
FU
271 Bacillus anthracis str. Ames GCA_000007845.1
271 Bacillus anthracis str. Sterne GCA_000008165.1
271 Bacillus cereus ATCC 14579 GCA_000007825.1
271 Bacillus mycoides GCA_000832605.1
271 Bacillus pseudomycoides DSM 12442 GCA_000161455.1
271 Bacillus thuringiensis YBT-1518 GCA_000497525.2

271 _5BBacillus thuringiensis_5D serovar konkukian str. 97-27 GCA_000008505.1
200 Bacillus manliponensis GCA_000712595.1
188 Bacillus coahuilensis m4-4 GCA_000171615.1
FV
260 Rhodobacter sphaeroides 2.4.1 GCA_000012905.2
226 Rhodobacter sphaeroides ATCC 17025 GCA_000016405.1
214 Pseudorhodobacter psychrotolerans GCA_001294535.1
FW
307 Bacillus anthracis str. Ames GCA_000007845.1
307 Bacillus anthracis str. Sterne GCA_000008165.1
307 Bacillus cereus ATCC 14579 GCA_000007825.1
307 Bacillus mycoides GCA_000832605.1
307 Bacillus pseudomycoides DSM 12442 GCA_000161455.1
307 Bacillus thuringiensis YBT-1518 GCA_000497525.2
307 _5BBacillus thuringiensis_5D serovar konkukian str. 97-27 GCA_000008505.1
259 Bacillus manliponensis GCA_000712595.1
257 Bacillus marisflavi GCA_001274775.1
FX
126 Deinococcus radiodurans R1 GCA_000008565.1
88 Deinococcus puniceus GCA_001644565.1
82 Deinococcus geothermalis DSM 11300 GCA_000196275.1
FY
136 Deinococcus radiodurans R1 GCA_000008565.1
80 Deinococcus marmoris DSM 12784 GCA_000701405.1
80 Deinococcus swuensis GCA_000800395.1
78 Deinococcus soli Cha et al. 2016 GCA_001007995.1
FZ
225 Clostridium beijerinckii GCA_000833105.2
225 Clostridium saccharobutylicum DSM 13864 GCA_000473995.1
225 Clostridium saccharoperbutylacetonicum N1-4_28HMT_29 GCA_000340885.1
218 Clostridium puniceum GCA_002006345.1
207 Clostridium chromiireducens GCA_002029255.1
G0
217 Erwinia iniecta GCA_001267535.1
217 Escherichia coli IAI39 GCA_000026345.1
217 Escherichia coli O104_3AH4 str. 2011C-3493 GCA_000299455.1
217 Escherichia coli O157_3AH7 str. Sakai GCA_000008865.1
217 Escherichia coli O83_3AH1 str. NRG 857C GCA_000183345.1
217 Escherichia coli UMN026 GCA_000026325.2
217 Escherichia coli str. K-12 substr. MG1655 GCA_000005845.2
217 Shigella dysenteriae Sd197 GCA_000012005.1
217 Shigella flexneri 2a str. 301 GCA_000006925.2
217 Tumebacillus flagellatus GCA_000714935.1
210 Cedecea neteri GCA_000757825.1
210 Rosenbergiella nectarea GCA_900111105.1
191 Edwardsiella anguillarum ET080813 GCA_000264765.2
G1
239 Clostridium beijerinckii GCA_000833105.2
239 Clostridium puniceum GCA_002006345.1
239 Clostridium saccharoperbutylacetonicum N1-4_28HMT_29 GCA_000340885.1
230 Clostridium saccharobutylicum DSM 13864 GCA_000473995.1
228 Clostridium taeniosporum GCA_001735765.1
G2
266 Deinococcus radiodurans R1 GCA_000008565.1
197 Deinococcus hopiensis KR-140 GCA_900176165.1
177 Deinococcus deserti VCD115 GCA_000020685.1
G3
206 Lactobacillus gasseri ATCC 33323 = JCM 1131 GCA_000014425.1
206 Lactobacillus hominis DSM 23910 = CRBIP 24.179 GCA_000296835.1
171 Lactobacillus amylophilus DSM 20533 = JCM 1125 GCA_001936335.1
157 Lactobacillus iners DSM 13335 GCA_000160875.1
G4
131 Clostridium beijerinckii GCA_000833105.2
131 Clostridium saccharoperbutylacetonicum N1-4_28HMT_29 GCA_000340885.1
127 Clostridium saccharobutylicum DSM 13864 GCA_000473995.1
124 Clostridium butyricum GCA_001456065.2
124 Clostridium puniceum GCA_002006345.1
G5
209 Pseudorhodobacter ferrugineus DSM 5888 GCA_000420745.1
209 Pseudorhodobacter wandonensis GCA_001202035.1
202 Rhodobacter sphaeroides 2.4.1 GCA_000012905.2
198 Pseudorhodobacter psychrotolerans GCA_001294535.1
G6
178 Clostridium beijerinckii GCA_000833105.2
178 Clostridium puniceum GCA_002006345.1
178 Clostridium saccharobutylicum DSM 13864 GCA_000473995.1

178 Clostridium saccharoperbutylacetonicum N1-4_28HMT_29 GCA_000340885.1
164 Clostridium chromiireducens GCA_002029255.1
156 Clostridium butyricum GCA_001456065.2
G7
264 Escherichia coli IAI39 GCA_000026345.1
264 Escherichia coli O104_3AH4 str. 2011C-3493 GCA_000299455.1
264 Escherichia coli O157_3AH7 str. Sakai GCA_000008865.1
264 Escherichia coli O83_3AH1 str. NRG 857C GCA_000183345.1
264 Escherichia coli UMN026 GCA_000026325.2
264 Escherichia coli str. K-12 substr. MG1655 GCA_000005845.2
264 Shigella flexneri 2a str. 301 GCA_000006925.2
264 Tumebacillus flagellatus GCA_000714935.1
249 Shigella dysenteriae Sd197 GCA_000012005.1
245 Erwinia iniecta GCA_001267535.1
G8
238 Bifidobacterium adolescentis ATCC 15703 GCA_000010425.1
213 Bifidobacterium breve DSM 20213 = JCM 1192 GCA_001025175.1
211 Bifidobacterium longum NCC2705 GCA_000007525.1
G9
186 Lactobacillus gasseri ATCC 33323 = JCM 1131 GCA_000014425.1
181 Lactobacillus hominis DSM 23910 = CRBIP 24.179 GCA_000296835.1
143 Lactobacillus psittaci DSM 15354 GCA_000425905.1
Ga
249 Deinococcus radiodurans R1 GCA_000008565.1
178 Deinococcus gobiensis I-0 GCA_000252445.1
168 Deinococcus puniceus GCA_001644565.1
Gb
257 Clostridium beijerinckii GCA_000833105.2
257 Clostridium puniceum GCA_002006345.1
257 Clostridium saccharobutylicum DSM 13864 GCA_000473995.1
257 Clostridium saccharoperbutylacetonicum N1-4_28HMT_29 GCA_000340885.1
246 Clostridium chromiireducens GCA_002029255.1
233 Clostridium taeniosporum GCA_001735765.1
Gc
275 Escherichia coli IAI39 GCA_000026345.1
275 Escherichia coli O104_3AH4 str. 2011C-3493 GCA_000299455.1
275 Escherichia coli O157_3AH7 str. Sakai GCA_000008865.1
275 Escherichia coli O83_3AH1 str. NRG 857C GCA_000183345.1
275 Escherichia coli UMN026 GCA_000026325.2
275 Escherichia coli str. K-12 substr. MG1655 GCA_000005845.2
275 Shigella flexneri 2a str. 301 GCA_000006925.2
275 Tumebacillus flagellatus GCA_000714935.1
266 Shigella dysenteriae Sd197 GCA_000012005.1
246 Erwinia iniecta GCA_001267535.1
246 Erwinia toletana DAPP-PG 735 GCA_000336255.1
Gd
294 Staphylococcus epidermidis ATCC 12228 GCA_000007645.1
291 Staphylococcus haemolyticus JCSC1435 GCA_000009865.1
289 Staphylococcus capitis subsp. capitis GCA_001028645.1
Ge
137 Bifidobacterium adolescentis ATCC 15703 GCA_000010425.1
137 Bifidobacterium dentium JCM 1195 = DSM 20436 GCA_001042595.1
129 Bifidobacterium tsurumiense GCA_000741765.1
121 Bifidobacterium asteroides PRL2011 GCA_000304215.1
Gf
149 Clostridium beijerinckii GCA_000833105.2
149 Clostridium butyricum GCA_001456065.2
149 Clostridium puniceum GCA_002006345.1
149 Clostridium saccharoperbutylacetonicum N1-4_28HMT_29 GCA_000340885.1
140 Clostridium chromiireducens GCA_002029255.1
140 Clostridium saccharobutylicum DSM 13864 GCA_000473995.1
133 Clostridium neonatale GCA_001458595.1
Gg
305 Streptococcus mutans UA159 GCA_000007465.2
172 Streptococcus sobrinus DSM 20742 = ATCC 33478 GCA_000686605.1
165 Streptococcus macacae NCTC 11558 GCA_000187995.3
Gh
247 Staphylococcus capitis subsp. capitis GCA_001028645.1
247 Staphylococcus epidermidis ATCC 12228 GCA_000007645.1
247 Staphylococcus haemolyticus JCSC1435 GCA_000009865.1
247 Staphylococcus hominis subsp. hominis C80 GCA_000183685.1
240 Staphylococcus lugdunensis HKU09-01 GCA_000025085.1
238 Staphylococcus aureus subsp. aureus NCTC 8325 GCA_000013425.1
238 Staphylococcus simiae CCM 7213 GCA_000235645.2
Gi
272 Streptococcus mutans UA159 GCA_000007465.2

176 Streptococcus ferus DSM 20646 GCA_000372425.1
173 Streptococcus marimammalium DSM 18627 GCA_000380045.1
Gj
223 Bifidobacterium adolescentis ATCC 15703 GCA_000010425.1
209 Bifidobacterium callitrichos DSM 23973 GCA_000741175.1
204 Bifidobacterium asteroides PRL2011 GCA_000304215.1
Gk
217 Clostridium beijerinckii GCA_000833105.2
217 Clostridium puniceum GCA_002006345.1
217 Clostridium saccharoperbutylacetonicum N1-4_28HMT_29 GCA_000340885.1
208 Clostridium saccharobutylicum DSM 13864 GCA_000473995.1
196 Clostridium taeniosporum GCA_001735765.1
Gl
139 Bacillus anthracis str. Ames GCA_000007845.1
139 Bacillus anthracis str. Sterne GCA_000008165.1
139 Bacillus cereus ATCC 14579 GCA_000007825.1
139 Bacillus mycoides GCA_000832605.1
139 Bacillus pseudomycoides DSM 12442 GCA_000161455.1
139 Bacillus thuringiensis YBT-1518 GCA_000497525.2
139 _5BBacillus thuringiensis_5D serovar konkukian str. 97-27 GCA_000008505.1
124 Bacillus coahuilensis m4-4 GCA_000171615.1
118 Bacillus marisflavi GCA_001274775.1
Gm
256 Deinococcus radiodurans R1 GCA_000008565.1
147 Deinococcus marmoris DSM 12784 GCA_000701405.1
147 Deinococcus puniceus GCA_001644565.1
147 Deinococcus swuensis GCA_000800395.1
131 Deinococcus frigens DSM 12807 GCA_000701425.1
Gn
111 Clostridium beijerinckii GCA_000833105.2
111 Clostridium puniceum GCA_002006345.1
111 Clostridium saccharobutylicum DSM 13864 GCA_000473995.1
111 Clostridium saccharoperbutylacetonicum N1-4_28HMT_29 GCA_000340885.1
103 Clostridium neonatale GCA_001458595.1
89 Clostridium botulinum B str. Eklund 17B _28NRP_29 GCA_000020165.1
89 Clostridium butyricum GCA_001456065.2
89 Clostridium chromiireducens GCA_002029255.1
89 Clostridium taeniosporum GCA_001735765.1
Go
188 Bacillus thuringiensis YBT-1518 GCA_000497525.2
172 Bacillus anthracis str. Ames GCA_000007845.1
172 Bacillus anthracis str. Sterne GCA_000008165.1
172 Bacillus cereus ATCC 14579 GCA_000007825.1
172 Bacillus mycoides GCA_000832605.1
172 Bacillus pseudomycoides DSM 12442 GCA_000161455.1
172 _5BBacillus thuringiensis_5D serovar konkukian str. 97-27 GCA_000008505.1
157 Bacillus manliponensis GCA_000712595.1
Gp
239 Bacillus mycoides GCA_000832605.1
229 Bacillus anthracis str. Ames GCA_000007845.1
229 Bacillus anthracis str. Sterne GCA_000008165.1
229 Bacillus cereus ATCC 14579 GCA_000007825.1
229 Bacillus pseudomycoides DSM 12442 GCA_000161455.1
229 Bacillus thuringiensis YBT-1518 GCA_000497525.2
229 _5BBacillus thuringiensis_5D serovar konkukian str. 97-27 GCA_000008505.1
198 Bacillus halmapalus GCA_002019665.1
Gq
312 Escherichia coli IAI39 GCA_000026345.1
312 Escherichia coli O104_3AH4 str. 2011C-3493 GCA_000299455.1
312 Escherichia coli O157_3AH7 str. Sakai GCA_000008865.1
312 Escherichia coli UMN026 GCA_000026325.2
312 Escherichia coli str. K-12 substr. MG1655 GCA_000005845.2
312 Shigella flexneri 2a str. 301 GCA_000006925.2
312 Tumebacillus flagellatus GCA_000714935.1
311 Escherichia coli O83_3AH1 str. NRG 857C GCA_000183345.1
303 Shigella dysenteriae Sd197 GCA_000012005.1
Gr
223 Staphylococcus epidermidis ATCC 12228 GCA_000007645.1
223 Staphylococcus haemolyticus JCSC1435 GCA_000009865.1
223 Staphylococcus hominis subsp. hominis C80 GCA_000183685.1
223 Staphylococcus lugdunensis HKU09-01 GCA_000025085.1
217 Staphylococcus capitis subsp. capitis GCA_001028645.1
208 Megasphaera cerevisiae DSM 20462 GCA_001045675.1
208 Staphylococcus aureus subsp. aureus NCTC 8325 GCA_000013425.1
208 Staphylococcus hyicus GCA_000816085.1
208 Staphylococcus lutrae GCA_002101335.1

208 Staphylococcus simiae CCM 7213 GCA_000235645.2
208 Staphylococcus warneri SG1 GCA_000332735.1
Gs
192 Clostridium beijerinckii GCA_000833105.2
192 Clostridium saccharobutylicum DSM 13864 GCA_000473995.1
192 Clostridium saccharoperbutylacetonicum N1-4_28HMT_29 GCA_000340885.1
186 Clostridium puniceum GCA_002006345.1
171 Clostridium chromiireducens GCA_002029255.1
Gt
338 Staphylococcus epidermidis ATCC 12228 GCA_000007645.1
333 Staphylococcus capitis subsp. capitis GCA_001028645.1
329 Staphylococcus lugdunensis HKU09-01 GCA_000025085.1
Gu
120 Bacillus thuringiensis YBT-1518 GCA_000497525.2
105 Bacillus anthracis str. Ames GCA_000007845.1
105 Bacillus anthracis str. Sterne GCA_000008165.1
105 Bacillus cereus ATCC 14579 GCA_000007825.1
105 Bacillus mycoides GCA_000832605.1
105 Bacillus pseudomycoides DSM 12442 GCA_000161455.1
105 _5BBacillus thuringiensis_5D serovar konkukian str. 97-27 GCA_000008505.1
91 Paenirhodobacter enshiensis GCA_000740785.1
Gv
242 Bifidobacterium adolescentis ATCC 15703 GCA_000010425.1
212 Bifidobacterium asteroides PRL2011 GCA_000304215.1
205 Bifidobacterium breve DSM 20213 = JCM 1192 GCA_001025175.1
Gw
157 Bacillus anthracis str. Ames GCA_000007845.1
157 Bacillus anthracis str. Sterne GCA_000008165.1
157 Bacillus cereus ATCC 14579 GCA_000007825.1
157 Bacillus mycoides GCA_000832605.1
157 Bacillus pseudomycoides DSM 12442 GCA_000161455.1
157 Bacillus thuringiensis YBT-1518 GCA_000497525.2
157 Viridibacillus arvi GCA_001274945.1
157 _5BBacillus thuringiensis_5D serovar konkukian str. 97-27 GCA_000008505.1
153 Kurthia massiliensis GCA_000285555.1
152 Bacillus horneckiae GCA_001636335.1
152 Bacillus solani GCA_001420595.1
Gx
275 Streptococcus mutans UA159 GCA_000007465.2
210 Streptococcus marimammalium DSM 18627 GCA_000380045.1
202 Streptococcus gordonii str. Challis substr. CH1 GCA_000017005.1
Gy
113 Rhodobacter sphaeroides 2.4.1 GCA_000012905.2
98 Pseudaminobacter salicylatoxidans KCT001 GCA_000304395.1
97 Agrobacterium rhizogenes GCA_000745735.1
97 Fulvimarina pelagi HTCC2506 GCA_000153705.1
97 Hoeflea olei GCA_001703635.1
97 Hoeflea phototrophica DFL-43 GCA_000154705.2
97 Rhizobium etli CFN 42 GCA_000092045.1
97 Rhizobium leguminosarum bv. trifolii WSM1689 GCA_000517605.1
97 Rhizobium leguminosarum bv. trifolii WSM2304 GCA_000021345.1
97 Rhizobium lusitanum GCA_900094565.1
97 Rhizobium miluonense GCA_900094545.1
97 Rhizobium rhizogenes NBRC 13257 GCA_000696095.1
97 Rhizobium tropici CIAT 899 GCA_000330885.1
Gz
167 Escherichia coli IAI39 GCA_000026345.1
167 Escherichia coli O104_3AH4 str. 2011C-3493 GCA_000299455.1
167 Escherichia coli O157_3AH7 str. Sakai GCA_000008865.1
167 Escherichia coli O83_3AH1 str. NRG 857C GCA_000183345.1
167 Escherichia coli UMN026 GCA_000026325.2
167 Escherichia coli str. K-12 substr. MG1655 GCA_000005845.2
167 Shigella flexneri 2a str. 301 GCA_000006925.2
167 Tumebacillus flagellatus GCA_000714935.1
158 Shigella dysenteriae Sd197 GCA_000012005.1
145 Erwinia iniecta GCA_001267535.1
GA
89 Gemmobacter aquatilis GCA_900110025.1
89 Rhodobacter capsulatus SB 1003 GCA_000021865.1
89 Rhodobacter sphaeroides 2.4.1 GCA_000012905.2
89 Rhodobacter sphaeroides ATCC 17025 GCA_000016405.1
85 Defluviimonas alba GCA_001620265.1
84 Ahrensia kielensis DSM 5890 GCA_000374465.1
84 Amorphus coralli DSM 19760 GCA_000374525.1
84 Aquamicrobium aerolatum DSM 21857 GCA_900113935.1
84 Aquamicrobium defluvii GCA_000585625.1

84 Azorhizobium caulinodans ORS 571 GCA_000010525.1
84 Defluviimonas indica GCA_900106675.1
84 Gemmobacter megaterium GCA_900156815.1
84 Gemmobacter nectariphilus DSM 15620 GCA_000429765.1
84 Haematobacter massiliensis GCA_000740795.1
84 Hoeflea olei GCA_001703635.1
84 Hoeflea phototrophica DFL-43 GCA_000154705.2
84 Nitratireductor aquibiodomus NL21 = JCM 21793 GCA_000615975.1
84 Nitratireductor basaltis GCA_000733725.1
84 Nitratireductor basaltis GCA_001953055.1
84 Nitratireductor indicus C115 GCA_000300515.1
84 Nitratireductor pacificus pht-3B GCA_000300335.1
84 Nitratireductor soli GCA_001050155.1
84 Oceanicella actignis GCA_900143155.1
84 Paenirhodobacter enshiensis GCA_000740785.1
84 Paracoccus alcaliphilus GCA_900110285.1
84 Paracoccus aminophilus JCM 7686 GCA_000444995.1
84 Paracoccus chinensis GCA_900102885.1
84 Paracoccus contaminans GCA_002105555.1
84 Paracoccus denitrificans PD1222 GCA_000203895.1
84 Paracoccus halophilus GCA_000763905.1
84 Paracoccus saliphilus GCA_900156835.1
84 Paracoccus sanguinis GCA_900106665.1
84 Paracoccus sediminis GCA_900188295.1
84 Paracoccus tibetensis GCA_900102505.1
84 Paracoccus versutus GCA_000763885.1
84 Paracoccus yeei GCA_002073635.1
84 Pseudaminobacter salicylatoxidans KCT001 GCA_000304395.1
84 Pseudorhodobacter aquimaris GCA_001202025.1
84 Pseudorhodobacter ferrugineus DSM 5888 GCA_000420745.1
84 Pseudorhodobacter psychrotolerans GCA_001294535.1
84 Pseudorhodobacter wandonensis GCA_001202035.1
84 Pseudorhodoplanes sinuspersici GCA_002119765.1
84 Pseudovibrio hongkongensis GCA_001561995.1
84 Pseudovibrio stylochi GCA_001562055.1
84 Rhodobacter aestuarii GCA_900156655.1
84 Rhodobacter vinaykumarii GCA_900156695.1
84 Rhodoblastus acidophilus GCA_900187365.1
84 Rhodovulum sulfidophilum DSM 1374 GCA_001633165.1
84 Rhodovulum sulfidophilum GCA_001941715.1
84 Roseicitreum antarcticum GCA_900107025.1
84 Thioclava atlantica GCA_000737065.1
84 Thioclava dalianensis GCA_000715505.1
84 Thioclava indica GCA_000714545.1
84 Thioclava marina GCA_002020135.1
84 Thioclava nitratireducens GCA_001940525.2
84 Xanthobacter autotrophicus Py2 GCA_000017645.1
GB
222 Rhodobacter sphaeroides 2.4.1 GCA_000012905.2
157 Gemmobacter aquatilis GCA_900110025.1
157 Pseudorhodobacter psychrotolerans GCA_001294535.1
155 Rhodobacter sphaeroides ATCC 17025 GCA_000016405.1
GC
142 Deinococcus radiodurans R1 GCA_000008565.1
99 Deinococcus deserti VCD115 GCA_000020685.1
96 Deinococcus gobiensis I-0 GCA_000252445.1
GD
247 Deinococcus radiodurans R1 GCA_000008565.1
191 Deinococcus puniceus GCA_001644565.1
174 Deinococcus deserti VCD115 GCA_000020685.1
GE
171 Deinococcus radiodurans R1 GCA_000008565.1
116 Deinococcus frigens DSM 12807 GCA_000701425.1
116 Deinococcus puniceus GCA_001644565.1
109 Deinococcus marmoris DSM 12784 GCA_000701405.1
109 Deinococcus swuensis GCA_000800395.1
GF
202 Clostridium beijerinckii GCA_000833105.2
202 Clostridium puniceum GCA_002006345.1
202 Clostridium saccharoperbutylacetonicum N1-4_28HMT_29 GCA_000340885.1
187 Clostridium saccharobutylicum DSM 13864 GCA_000473995.1
182 Clostridium butyricum GCA_001456065.2
GG
256 Rhodobacter sphaeroides 2.4.1 GCA_000012905.2
238 Gemmobacter megaterium GCA_900156815.1
229 Pseudorhodobacter ferrugineus DSM 5888 GCA_000420745.1

229 Pseudorhodobacter wandonensis GCA_001202035.1
GH
101 Clostridium arbusti SL206 GCA_000246895.2
101 Clostridium pasteurianum BC1 GCA_000389635.1
101 Clostridium pasteurianum DSM 525 = ATCC 6013 GCA_000807255.1
96 Clostridium baratii str. Sullivan GCA_000789395.1
96 Clostridium beijerinckii GCA_000833105.2
96 Clostridium butyricum GCA_001456065.2
96 Clostridium chromiireducens GCA_002029255.1
96 Clostridium puniceum GCA_002006345.1
96 Clostridium saccharobutylicum DSM 13864 GCA_000473995.1
96 Clostridium saccharoperbutylacetonicum N1-4_28HMT_29 GCA_000340885.1
96 Clostridium taeniosporum GCA_001735765.1
96 Clostridium thermobutyricum DSM 4928 GCA_002050515.1
86 Clostridium akagii DSM 12554 GCA_000686725.1
86 Clostridium algidicarnis GCA_000703125.1
86 Clostridium autoethanogenum DSM 10061 GCA_000484505.1
86 Clostridium bornimense GCA_000577895.1
86 Clostridium carboxidivorans P7 GCA_001038625.1
86 Clostridium magnum DSM 2767 GCA_001623875.1
86 Clostridium neonatale GCA_001458595.1
86 Clostridium tyrobutyricum GCA_001642655.1
86 Desnuesiella massiliensis GCA_001403615.1
GI
236 Clostridium beijerinckii GCA_000833105.2
236 Clostridium puniceum GCA_002006345.1
236 Clostridium saccharobutylicum DSM 13864 GCA_000473995.1
236 Clostridium saccharoperbutylacetonicum N1-4_28HMT_29 GCA_000340885.1
213 Clostridium chromiireducens GCA_002029255.1
211 Clostridium neonatale GCA_001458595.1
GJ
203 Clostridium beijerinckii GCA_000833105.2
203 Clostridium puniceum GCA_002006345.1
203 Clostridium saccharobutylicum DSM 13864 GCA_000473995.1
203 Clostridium saccharoperbutylacetonicum N1-4_28HMT_29 GCA_000340885.1
193 Clostridium neonatale GCA_001458595.1
187 Clostridium chromiireducens GCA_002029255.1
GK
265 Staphylococcus condimenti GCA_001618885.1
265 Staphylococcus simulans GCA_001559115.1
262 Staphylococcus aureus subsp. aureus NCTC 8325 GCA_000013425.1
262 Staphylococcus epidermidis ATCC 12228 GCA_000007645.1
262 Staphylococcus haemolyticus JCSC1435 GCA_000009865.1
262 Staphylococcus hominis subsp. hominis C80 GCA_000183685.1
262 Staphylococcus simiae CCM 7213 GCA_000235645.2
257 Staphylococcus capitis subsp. capitis GCA_001028645.1
257 Staphylococcus lugdunensis HKU09-01 GCA_000025085.1
GL
124 Deinococcus radiodurans R1 GCA_000008565.1
93 Deinococcus marmoris DSM 12784 GCA_000701405.1
93 Deinococcus swuensis GCA_000800395.1
68 Deinococcus frigens DSM 12807 GCA_000701425.1
GM
273 Clostridium beijerinckii GCA_000833105.2
273 Clostridium puniceum GCA_002006345.1
273 Clostridium saccharobutylicum DSM 13864 GCA_000473995.1
273 Clostridium saccharoperbutylacetonicum N1-4_28HMT_29 GCA_000340885.1
254 Clostridium chromiireducens GCA_002029255.1
238 Clostridium butyricum GCA_001456065.2
GN
261 Clostridium beijerinckii GCA_000833105.2
261 Clostridium puniceum GCA_002006345.1
261 Clostridium saccharoperbutylacetonicum N1-4_28HMT_29 GCA_000340885.1
253 Clostridium saccharobutylicum DSM 13864 GCA_000473995.1
241 Clostridium chromiireducens GCA_002029255.1
GO
227 Rhodobacter sphaeroides 2.4.1 GCA_000012905.2
206 Rhodobacter sphaeroides ATCC 17025 GCA_000016405.1
198 Thioclava dalianensis GCA_000715505.1
198 Thioclava indica GCA_000714545.1
GP
269 Bifidobacterium adolescentis ATCC 15703 GCA_000010425.1
235 Bifidobacterium dentium JCM 1195 = DSM 20436 GCA_001042595.1
235 Bifidobacterium tsurumiense GCA_000741765.1
233 Bifidobacterium angulatum DSM 20098 = JCM 7096 GCA_001025155.1
GQ

237 Deinococcus radiodurans R1 GCA_000008565.1
147 Deinococcus puniceus GCA_001644565.1
140 Deinococcus deserti VCD115 GCA_000020685.1
GR
228 Deinococcus radiodurans R1 GCA_000008565.1
133 Deinococcus geothermalis DSM 11300 GCA_000196275.1
133 Deinococcus marmoris DSM 12784 GCA_000701405.1
133 Deinococcus swuensis GCA_000800395.1
131 Deinococcus gobiensis I-0 GCA_000252445.1
GS
280 Staphylococcus capitis subsp. capitis GCA_001028645.1
280 Staphylococcus cohnii subsp. cohnii GCA_000972575.1
280 Staphylococcus epidermidis ATCC 12228 GCA_000007645.1
280 Staphylococcus lugdunensis HKU09-01 GCA_000025085.1
280 Staphylococcus saprophyticus subsp. saprophyticus ATCC 15305 GCA_000010125.1
280 Staphylococcus xylosus GCA_000706685.1
266 Staphylococcus pettenkoferi GCA_002208805.1
265 Megasphaera cerevisiae DSM 20462 GCA_001045675.1
265 Staphylococcus arlettae CVD059 GCA_000295715.1
265 Staphylococcus aureus subsp. aureus NCTC 8325 GCA_000013425.1
265 Staphylococcus haemolyticus JCSC1435 GCA_000009865.1
265 Staphylococcus hominis subsp. hominis C80 GCA_000183685.1
265 Staphylococcus simiae CCM 7213 GCA_000235645.2
265 Staphylococcus succinus GCA_001902315.1
265 Staphylococcus warneri SG1 GCA_000332735.1
GT
219 Deinococcus radiodurans R1 GCA_000008565.1
134 Deinococcus gobiensis I-0 GCA_000252445.1
115 Deinococcus marmoris DSM 12784 GCA_000701405.1
115 Deinococcus proteolyticus MRP GCA_000190555.1
115 Deinococcus swuensis GCA_000800395.1
GU
162 Deinococcus radiodurans R1 GCA_000008565.1
116 Deinococcus geothermalis DSM 11300 GCA_000196275.1
108 Deinococcus gobiensis I-0 GCA_000252445.1
GV
229 Bacillus mycoides GCA_000832605.1
228 Bacillus anthracis str. Ames GCA_000007845.1
228 Bacillus anthracis str. Sterne GCA_000008165.1
228 Bacillus cereus ATCC 14579 GCA_000007825.1
228 Bacillus pseudomycoides DSM 12442 GCA_000161455.1
228 Bacillus thuringiensis YBT-1518 GCA_000497525.2
228 _5BBacillus thuringiensis_5D serovar konkukian str. 97-27 GCA_000008505.1
160 Bacillus manliponensis GCA_000712595.1
GW
156 Rhodobacter sphaeroides 2.4.1 GCA_000012905.2
118 Rhodobacter capsulatus SB 1003 GCA_000021865.1
112 Haematobacter massiliensis GCA_000740795.1
GX
247 Clostridium beijerinckii GCA_000833105.2
247 Clostridium saccharobutylicum DSM 13864 GCA_000473995.1
247 Clostridium saccharoperbutylacetonicum N1-4_28HMT_29 GCA_000340885.1
237 Clostridium butyricum GCA_001456065.2
233 Clostridium chromiireducens GCA_002029255.1
233 Clostridium puniceum GCA_002006345.1
GY
257 Streptococcus mutans UA159 GCA_000007465.2
155 Streptococcus ferus DSM 20646 GCA_000372425.1
153 Streptococcus macacae NCTC 11558 GCA_000187995.3
GZ
253 Deinococcus radiodurans R1 GCA_000008565.1
177 Deinococcus gobiensis I-0 GCA_000252445.1
165 Deinococcus deserti VCD115 GCA_000020685.1
H0
285 Staphylococcus capitis subsp. capitis GCA_001028645.1
285 Staphylococcus epidermidis ATCC 12228 GCA_000007645.1
282 Staphylococcus haemolyticus JCSC1435 GCA_000009865.1
272 Staphylococcus condimenti GCA_001618885.1
272 Staphylococcus simulans GCA_001559115.1
H1
218 Staphylococcus capitis subsp. capitis GCA_001028645.1
218 Staphylococcus epidermidis ATCC 12228 GCA_000007645.1
218 Staphylococcus haemolyticus JCSC1435 GCA_000009865.1
207 Staphylococcus cohnii subsp. cohnii GCA_000972575.1
207 Staphylococcus hominis subsp. hominis C80 GCA_000183685.1
207 Staphylococcus microti GCA_000934465.1

193 Staphylococcus pettenkoferi GCA_002208805.1
H2
199 Staphylococcus saprophyticus GCA_001074355.1
195 Staphylococcus capitis subsp. capitis GCA_001028645.1
195 Staphylococcus cohnii subsp. cohnii GCA_000972575.1
195 Staphylococcus epidermidis ATCC 12228 GCA_000007645.1
183 Staphylococcus arlettae CVD059 GCA_000295715.1
H3
252 Lactobacillus gasseri ATCC 33323 = JCM 1131 GCA_000014425.1
236 Lactobacillus hominis DSM 23910 = CRBIP 24.179 GCA_000296835.1
188 Lactobacillus iners DSM 13335 GCA_000160875.1
H4
237 Deinococcus radiodurans R1 GCA_000008565.1
184 Deinococcus deserti VCD115 GCA_000020685.1
176 Deinococcus puniceus GCA_001644565.1
H5
251 Streptococcus mutans UA159 GCA_000007465.2
164 Streptococcus equinus GCA_000964315.1
164 Streptococcus gallolyticus subsp. gallolyticus DSM 16831 GCA_002000985.1
155 Streptococcus cristatus AS 1.3089 GCA_000385925.1
155 Streptococcus gordonii str. Challis substr. CH1 GCA_000017005.1
155 Streptococcus mitis B6 GCA_000027165.1
155 Streptococcus pneumoniae R6 GCA_000007045.1
H6
127 Bifidobacterium adolescentis ATCC 15703 GCA_000010425.1
112 Bifidobacterium thermophilum GCA_000741495.1
103 Bifidobacterium tsurumiense GCA_000741765.1
H7
155 Streptococcus mutans UA159 GCA_000007465.2
86 Streptococcus marimammalium DSM 18627 GCA_000380045.1
80 Streptococcus equinus GCA_000964315.1
80 Streptococcus gallolyticus subsp. gallolyticus DSM 16831 GCA_002000985.1
H8
163 Enterococcus faecalis V583 GCA_000007785.1
163 Streptomyces cinnamoneus GCA_001885705.1
150 Isobaculum melis GCA_900111355.1
139 Bavariicoccus seileri DSM 19936 GCA_000421665.1
139 Carnobacterium divergens DSM 20623 GCA_000744255.1
139 Carnobacterium funditum DSM 5970 GCA_000744185.1
139 Carnobacterium iners GCA_900177385.1
139 Granulicatella balaenopterae GCA_900111135.1
139 Trichococcus ilyis GCA_900067235.1
H9
250 Escherichia coli IAI39 GCA_000026345.1
250 Escherichia coli O104_3AH4 str. 2011C-3493 GCA_000299455.1
250 Escherichia coli O157_3AH7 str. Sakai GCA_000008865.1
250 Escherichia coli O83_3AH1 str. NRG 857C GCA_000183345.1
250 Escherichia coli UMN026 GCA_000026325.2
250 Escherichia coli str. K-12 substr. MG1655 GCA_000005845.2
250 Shigella flexneri 2a str. 301 GCA_000006925.2
250 Tumebacillus flagellatus GCA_000714935.1
235 Shigella dysenteriae Sd197 GCA_000012005.1
217 Cronobacter sakazakii GCA_000982825.1
217 Erwinia iniecta GCA_001267535.1
217 Erwinia toletana DAPP-PG 735 GCA_000336255.1
217 Kosakonia cowanii GCA_001975225.1
217 Kosakonia sacchari SP1 GCA_000300455.4
217 Pluralibacter gergoviae GCA_000757785.1
Ha
216 Clostridium beijerinckii GCA_000833105.2
216 Clostridium saccharobutylicum DSM 13864 GCA_000473995.1
216 Clostridium saccharoperbutylacetonicum N1-4_28HMT_29 GCA_000340885.1
209 Clostridium puniceum GCA_002006345.1
183 Clostridium chromiireducens GCA_002029255.1
Hb
296 Bacillus thuringiensis YBT-1518 GCA_000497525.2
282 Bacillus anthracis str. Ames GCA_000007845.1
282 Bacillus anthracis str. Sterne GCA_000008165.1
282 Bacillus cereus ATCC 14579 GCA_000007825.1
282 Bacillus mycoides GCA_000832605.1
282 Bacillus pseudomycoides DSM 12442 GCA_000161455.1
282 _5BBacillus thuringiensis_5D serovar konkukian str. 97-27 GCA_000008505.1
244 Bacillus manliponensis GCA_000712595.1
Hc
180 Escherichia coli IAI39 GCA_000026345.1
180 Escherichia coli O104_3AH4 str. 2011C-3493 GCA_000299455.1

180 Escherichia coli O157_3AH7 str. Sakai GCA_000008865.1
180 Escherichia coli O83_3AH1 str. NRG 857C GCA_000183345.1
180 Escherichia coli UMN026 GCA_000026325.2
180 Escherichia coli str. K-12 substr. MG1655 GCA_000005845.2
180 Shigella flexneri 2a str. 301 GCA_000006925.2
180 Tumebacillus flagellatus GCA_000714935.1
171 Shigella dysenteriae Sd197 GCA_000012005.1
154 Erwinia iniecta GCA_001267535.1
Hd
227 Rhodobacter sphaeroides 2.4.1 GCA_000012905.2
201 Gemmobacter aquatilis GCA_900110025.1
201 Gemmobacter nectariphilus DSM 15620 GCA_000429765.1
201 Rhodobacter capsulatus SB 1003 GCA_000021865.1
194 Haematobacter massiliensis GCA_000740795.1
He
166 Rhodobacter sphaeroides 2.4.1 GCA_000012905.2
145 Rhodobacter sphaeroides ATCC 17025 GCA_000016405.1
129 Paracoccus sediminis GCA_900188295.1
Hf
247 Staphylococcus simulans GCA_001559115.1
244 Staphylococcus aureus subsp. aureus NCTC 8325 GCA_000013425.1
244 Staphylococcus epidermidis ATCC 12228 GCA_000007645.1
244 Staphylococcus haemolyticus JCSC1435 GCA_000009865.1
244 Staphylococcus hominis subsp. hominis C80 GCA_000183685.1
244 Staphylococcus simiae CCM 7213 GCA_000235645.2
236 Staphylococcus capitis subsp. capitis GCA_001028645.1
Hg
248 Rhodobacter sphaeroides 2.4.1 GCA_000012905.2
232 Pseudorhodobacter psychrotolerans GCA_001294535.1
223 Pseudorhodobacter ferrugineus DSM 5888 GCA_000420745.1
223 Pseudorhodobacter wandonensis GCA_001202035.1
Hh
257 Streptococcus mutans UA159 GCA_000007465.2
165 Streptococcus massiliensis DSM 18628 GCA_000380065.1
154 Streptococcus ferus DSM 20646 GCA_000372425.1
Hi
154 Clostridium chromiireducens GCA_002029255.1
154 Clostridium saccharobutylicum DSM 13864 GCA_000473995.1
153 Clostridium taeniosporum GCA_001735765.1
152 Clostridium beijerinckii GCA_000833105.2
152 Clostridium puniceum GCA_002006345.1
152 Clostridium saccharoperbutylacetonicum N1-4_28HMT_29 GCA_000340885.1
Hj
270 Bifidobacterium adolescentis ATCC 15703 GCA_000010425.1
247 Bifidobacterium breve DSM 20213 = JCM 1192 GCA_001025175.1
245 Bifidobacterium gallicum DSM 20093 = LMG 11596 GCA_000741205.1
Hk
105 Bifidobacterium adolescentis ATCC 15703 GCA_000010425.1
87 Bifidobacterium tsurumiense GCA_000741765.1
76 Bifidobacterium thermophilum GCA_000741495.1
Hl
238 Clostridium beijerinckii GCA_000833105.2
238 Clostridium saccharoperbutylacetonicum N1-4_28HMT_29 GCA_000340885.1
231 Clostridium puniceum GCA_002006345.1
227 Clostridium butyricum GCA_001456065.2
Hm
272 Bifidobacterium adolescentis ATCC 15703 GCA_000010425.1
242 Bifidobacterium callitrichos DSM 23973 GCA_000741175.1
231 Bifidobacterium thermophilum GCA_000741495.1
Hn
266 Deinococcus radiodurans R1 GCA_000008565.1
168 Deinococcus deserti VCD115 GCA_000020685.1
163 Deinococcus puniceus GCA_001644565.1
Ho
182 Lactobacillus gasseri ATCC 33323 = JCM 1131 GCA_000014425.1
164 Lactobacillus hominis DSM 23910 = CRBIP 24.179 GCA_000296835.1
139 Lactobacillus psittaci DSM 15354 GCA_000425905.1
Hp
212 Deinococcus radiodurans R1 GCA_000008565.1
155 Deinococcus deserti VCD115 GCA_000020685.1
147 Deinococcus gobiensis I-0 GCA_000252445.1
Hq
219 Escherichia coli IAI39 GCA_000026345.1
219 Escherichia coli O104_3AH4 str. 2011C-3493 GCA_000299455.1
219 Escherichia coli O157_3AH7 str. Sakai GCA_000008865.1
219 Escherichia coli UMN026 GCA_000026325.2

219 Escherichia coli str. K-12 substr. MG1655 GCA_000005845.2
219 Shigella dysenteriae Sd197 GCA_000012005.1
219 Shigella flexneri 2a str. 301 GCA_000006925.2
219 Tumebacillus flagellatus GCA_000714935.1
218 Escherichia coli O83_3AH1 str. NRG 857C GCA_000183345.1
213 Erwinia iniecta GCA_001267535.1
Hr
205 Clostridium saccharobutylicum DSM 13864 GCA_000473995.1
201 Clostridium beijerinckii GCA_000833105.2
201 Clostridium puniceum GCA_002006345.1
201 Clostridium saccharoperbutylacetonicum N1-4_28HMT_29 GCA_000340885.1
196 Clostridium butyricum GCA_001456065.2
Hs
202 Deinococcus radiodurans R1 GCA_000008565.1
129 Deinococcus deserti VCD115 GCA_000020685.1
121 Deinococcus hopiensis KR-140 GCA_900176165.1
Ht
254 Bacillus anthracis str. Ames GCA_000007845.1
254 Bacillus anthracis str. Sterne GCA_000008165.1
254 Bacillus cereus ATCC 14579 GCA_000007825.1
254 Bacillus mycoides GCA_000832605.1
254 Bacillus pseudomycoides DSM 12442 GCA_000161455.1
254 Bacillus thuringiensis YBT-1518 GCA_000497525.2
254 _5BBacillus thuringiensis_5D serovar konkukian str. 97-27 GCA_000008505.1
218 Bacillus marisflavi GCA_001274775.1
209 Bacillus horneckiae GCA_001636335.1
209 Bacillus solani GCA_001420595.1
Hu
255 Rhodobacter sphaeroides 2.4.1 GCA_000012905.2
212 Rhodobacter sphaeroides ATCC 17025 GCA_000016405.1
196 Pseudorhodobacter psychrotolerans GCA_001294535.1
Hv
140 Escherichia coli IAI39 GCA_000026345.1
140 Escherichia coli O104_3AH4 str. 2011C-3493 GCA_000299455.1
140 Escherichia coli O157_3AH7 str. Sakai GCA_000008865.1
140 Escherichia coli O83_3AH1 str. NRG 857C GCA_000183345.1
140 Escherichia coli UMN026 GCA_000026325.2
140 Escherichia coli str. K-12 substr. MG1655 GCA_000005845.2
140 Shigella dysenteriae Sd197 GCA_000012005.1
140 Shigella flexneri 2a str. 301 GCA_000006925.2
140 Tumebacillus flagellatus GCA_000714935.1
125 Erwinia iniecta GCA_001267535.1
122 Lonsdalea quercina subsp. quercina GCA_000688655.1
Hw
225 Deinococcus radiodurans R1 GCA_000008565.1
153 Deinococcus gobiensis I-0 GCA_000252445.1
152 Deinococcus marmoris DSM 12784 GCA_000701405.1
152 Deinococcus swuensis GCA_000800395.1
Hx
294 Escherichia coli IAI39 GCA_000026345.1
294 Escherichia coli O104_3AH4 str. 2011C-3493 GCA_000299455.1
294 Escherichia coli O157_3AH7 str. Sakai GCA_000008865.1
294 Escherichia coli O83_3AH1 str. NRG 857C GCA_000183345.1
294 Escherichia coli UMN026 GCA_000026325.2
294 Escherichia coli str. K-12 substr. MG1655 GCA_000005845.2
294 Shigella flexneri 2a str. 301 GCA_000006925.2
294 Tumebacillus flagellatus GCA_000714935.1
278 Shigella dysenteriae Sd197 GCA_000012005.1
275 Erwinia iniecta GCA_001267535.1
Hy
261 Bacillus thuringiensis YBT-1518 GCA_000497525.2
255 Bacillus anthracis str. Ames GCA_000007845.1
255 Bacillus anthracis str. Sterne GCA_000008165.1
255 Bacillus cereus ATCC 14579 GCA_000007825.1
255 Bacillus mycoides GCA_000832605.1
255 Bacillus pseudomycoides DSM 12442 GCA_000161455.1
255 _5BBacillus thuringiensis_5D serovar konkukian str. 97-27 GCA_000008505.1
224 Bacillus horneckiae GCA_001636335.1
224 Bacillus solani GCA_001420595.1
Hz
145 Lactobacillus gasseri ATCC 33323 = JCM 1131 GCA_000014425.1
145 Lactobacillus hominis DSM 23910 = CRBIP 24.179 GCA_000296835.1
116 Lactobacillus acidophilus NCFM GCA_000011985.1
116 Lactobacillus amylovorus GCA_000191545.1
116 Lactobacillus crispatus ST1 GCA_000091765.1
116 Lactobacillus gallinarum GCA_001314245.2

116 Lactobacillus hamsteri DSM 5661 = JCM 6256 GCA_000615445.1
113 Lactobacillus kalixensis DSM 16043 GCA_001434335.1
HA
207 Deinococcus radiodurans R1 GCA_000008565.1
167 Deinococcus hopiensis KR-140 GCA_900176165.1
164 Deinococcus puniceus GCA_001644565.1
HB
267 Deinococcus radiodurans R1 GCA_000008565.1
183 Deinococcus marmoris DSM 12784 GCA_000701405.1
183 Deinococcus swuensis GCA_000800395.1
169 Deinococcus gobiensis I-0 GCA_000252445.1
HC
229 Streptococcus mutans UA159 GCA_000007465.2
196 Streptococcus cristatus AS 1.3089 GCA_000385925.1
196 Streptococcus gordonii str. Challis substr. CH1 GCA_000017005.1
192 Streptococcus mitis B6 GCA_000027165.1
192 Streptococcus pneumoniae R6 GCA_000007045.1
HD
183 Deinococcus radiodurans R1 GCA_000008565.1
142 Deinococcus gobiensis I-0 GCA_000252445.1
113 Deinococcus marmoris DSM 12784 GCA_000701405.1
113 Deinococcus swuensis GCA_000800395.1
HE
182 Bacillus thuringiensis YBT-1518 GCA_000497525.2
165 Bacillus anthracis str. Ames GCA_000007845.1
165 Bacillus anthracis str. Sterne GCA_000008165.1
165 Bacillus cereus ATCC 14579 GCA_000007825.1
165 Bacillus mycoides GCA_000832605.1
165 Bacillus pseudomycoides DSM 12442 GCA_000161455.1
165 _5BBacillus thuringiensis_5D serovar konkukian str. 97-27 GCA_000008505.1
143 Bacillus cytotoxicus NVH 391-98 GCA_000017425.1
HF
108 Deinococcus radiodurans R1 GCA_000008565.1
56 Deinococcus proteolyticus MRP GCA_000190555.1
50 Deinococcus deserti VCD115 GCA_000020685.1
50 Deinococcus frigens DSM 12807 GCA_000701425.1
HG
112 Clostridium beijerinckii GCA_000833105.2
112 Clostridium puniceum GCA_002006345.1
112 Clostridium saccharoperbutylacetonicum N1-4_28HMT_29 GCA_000340885.1
103 Clostridium chromiireducens GCA_002029255.1
103 Clostridium saccharobutylicum DSM 13864 GCA_000473995.1
87 Clostridium butyricum GCA_001456065.2
HH
329 Bacillus thuringiensis YBT-1518 GCA_000497525.2
328 Bacillus anthracis str. Ames GCA_000007845.1
328 Bacillus anthracis str. Sterne GCA_000008165.1
328 Bacillus mycoides GCA_000832605.1
328 Bacillus pseudomycoides DSM 12442 GCA_000161455.1
328 _5BBacillus thuringiensis_5D serovar konkukian str. 97-27 GCA_000008505.1
327 Bacillus cereus ATCC 14579 GCA_000007825.1
HI
182 Clostridium beijerinckii GCA_000833105.2
182 Clostridium puniceum GCA_002006345.1
182 Clostridium saccharobutylicum DSM 13864 GCA_000473995.1
182 Clostridium saccharoperbutylacetonicum N1-4_28HMT_29 GCA_000340885.1
181 Clostridium neonatale GCA_001458595.1
171 Clostridium butyricum GCA_001456065.2
171 Clostridium chromiireducens GCA_002029255.1
HJ
263 Streptococcus mutans UA159 GCA_000007465.2
152 Lactococcus piscium MKFS47 GCA_000981525.1
151 Streptococcus gordonii str. Challis substr. CH1 GCA_000017005.1
HK
197 Deinococcus radiodurans R1 GCA_000008565.1
170 Deinococcus gobiensis I-0 GCA_000252445.1
142 Deinococcus puniceus GCA_001644565.1
HL
237 Clostridium beijerinckii GCA_000833105.2
237 Clostridium saccharobutylicum DSM 13864 GCA_000473995.1
237 Clostridium saccharoperbutylacetonicum N1-4_28HMT_29 GCA_000340885.1
230 Clostridium puniceum GCA_002006345.1
219 Clostridium chromiireducens GCA_002029255.1
219 Clostridium intestinale URNW GCA_000469625.2
HM
226 Streptococcus mutans UA159 GCA_000007465.2

163 Streptococcus sobrinus DSM 20742 = ATCC 33478 GCA_000686605.1
159 Streptococcus ratti FA-1 = DSM 20564 GCA_000286075.1
HN
255 Bacillus thuringiensis YBT-1518 GCA_000497525.2
254 Bacillus anthracis str. Ames GCA_000007845.1
254 Bacillus anthracis str. Sterne GCA_000008165.1
254 Bacillus cereus ATCC 14579 GCA_000007825.1
254 Bacillus mycoides GCA_000832605.1
254 Bacillus pseudomycoides DSM 12442 GCA_000161455.1
254 _5BBacillus thuringiensis_5D serovar konkukian str. 97-27 GCA_000008505.1
207 Bacillus manliponensis GCA_000712595.1
HO
231 Megasphaera cerevisiae DSM 20462 GCA_001045675.1
231 Staphylococcus warneri SG1 GCA_000332735.1
228 Staphylococcus hominis subsp. hominis C80 GCA_000183685.1
228 Staphylococcus lugdunensis HKU09-01 GCA_000025085.1
222 Staphylococcus arlettae CVD059 GCA_000295715.1
222 Staphylococcus saprophyticus subsp. saprophyticus ATCC 15305 GCA_000010125.1
HP
208 Erwinia iniecta GCA_001267535.1
208 Escherichia coli IAI39 GCA_000026345.1
208 Escherichia coli O104_3AH4 str. 2011C-3493 GCA_000299455.1
208 Escherichia coli O157_3AH7 str. Sakai GCA_000008865.1
208 Escherichia coli O83_3AH1 str. NRG 857C GCA_000183345.1
208 Escherichia coli UMN026 GCA_000026325.2
208 Escherichia coli str. K-12 substr. MG1655 GCA_000005845.2
208 Shigella flexneri 2a str. 301 GCA_000006925.2
208 Tumebacillus flagellatus GCA_000714935.1
203 Cedecea neteri GCA_000757825.1
199 Shigella dysenteriae Sd197 GCA_000012005.1
HQ
200 Deinococcus radiodurans R1 GCA_000008565.1
102 Deinococcus gobiensis I-0 GCA_000252445.1
101 Deinococcus puniceus GCA_001644565.1
HR
213 Escherichia coli IAI39 GCA_000026345.1
213 Escherichia coli O104_3AH4 str. 2011C-3493 GCA_000299455.1
213 Escherichia coli O157_3AH7 str. Sakai GCA_000008865.1
213 Escherichia coli O83_3AH1 str. NRG 857C GCA_000183345.1
213 Escherichia coli UMN026 GCA_000026325.2
213 Escherichia coli str. K-12 substr. MG1655 GCA_000005845.2
213 Shigella dysenteriae Sd197 GCA_000012005.1
213 Shigella flexneri 2a str. 301 GCA_000006925.2
213 Tumebacillus flagellatus GCA_000714935.1
175 Erwinia iniecta GCA_001267535.1
162 Xenorhabdus bovienii SS-2004 GCA_000027225.1
HS
176 Lactobacillus gasseri ATCC 33323 = JCM 1131 GCA_000014425.1
158 Lactobacillus hominis DSM 23910 = CRBIP 24.179 GCA_000296835.1
127 Lactobacillus iners DSM 13335 GCA_000160875.1
HT
291 Bifidobacterium adolescentis ATCC 15703 GCA_000010425.1
263 Bifidobacterium thermophilum GCA_000741495.1
254 Bifidobacterium callitrichos DSM 23973 GCA_000741175.1
HU
291 Rhodobacter sphaeroides 2.4.1 GCA_000012905.2
251 Gemmobacter megaterium GCA_900156815.1
249 Pseudorhodobacter psychrotolerans GCA_001294535.1
HV
166 Escherichia coli IAI39 GCA_000026345.1
166 Escherichia coli O104_3AH4 str. 2011C-3493 GCA_000299455.1
166 Escherichia coli O157_3AH7 str. Sakai GCA_000008865.1
166 Escherichia coli O83_3AH1 str. NRG 857C GCA_000183345.1
166 Escherichia coli UMN026 GCA_000026325.2
166 Escherichia coli str. K-12 substr. MG1655 GCA_000005845.2
166 Shigella dysenteriae Sd197 GCA_000012005.1
166 Shigella flexneri 2a str. 301 GCA_000006925.2
166 Tumebacillus flagellatus GCA_000714935.1
147 Erwinia iniecta GCA_001267535.1
145 Klebsiella pneumoniae subsp. pneumoniae HS11286 GCA_000240185.2
HW
246 _5BBacillus thuringiensis_5D serovar konkukian str. 97-27 GCA_000008505.1
245 Bacillus thuringiensis YBT-1518 GCA_000497525.2
230 Bacillus anthracis str. Ames GCA_000007845.1
230 Bacillus anthracis str. Sterne GCA_000008165.1
230 Bacillus cereus ATCC 14579 GCA_000007825.1

230 Bacillus mycoides GCA_000832605.1
230 Bacillus pseudomycoides DSM 12442 GCA_000161455.1
HX
226 Bacillus anthracis str. Ames GCA_000007845.1
226 Bacillus anthracis str. Sterne GCA_000008165.1
226 Bacillus cereus ATCC 14579 GCA_000007825.1
226 Bacillus mycoides GCA_000832605.1
226 Bacillus pseudomycoides DSM 12442 GCA_000161455.1
226 Bacillus thuringiensis YBT-1518 GCA_000497525.2
226 _5BBacillus thuringiensis_5D serovar konkukian str. 97-27 GCA_000008505.1
199 Bacillus manliponensis GCA_000712595.1
196 Bacillus aquimaris TF-12 GCA_001648555.1
196 Bacillus marisflavi GCA_001274775.1
196 Bacillus weihaiensis GCA_001889165.1
HY
215 Streptococcus mutans UA159 GCA_000007465.2
139 Streptococcus gordonii str. Challis substr. CH1 GCA_000017005.1
129 Streptococcus macacae NCTC 11558 GCA_000187995.3
HZ
224 Bacillus anthracis str. Ames GCA_000007845.1
224 Bacillus anthracis str. Sterne GCA_000008165.1
224 Bacillus cereus ATCC 14579 GCA_000007825.1
224 Bacillus mycoides GCA_000832605.1
224 Bacillus pseudomycoides DSM 12442 GCA_000161455.1
224 Bacillus thuringiensis YBT-1518 GCA_000497525.2
224 _5BBacillus thuringiensis_5D serovar konkukian str. 97-27 GCA_000008505.1
183 Bacillus cohnii NBRC 15565 GCA_001591425.1
183 Bacillus halmapalus GCA_002019665.1
181 Bacillus cytotoxicus NVH 391-98 GCA_000017425.1
I0
172 Bifidobacterium adolescentis ATCC 15703 GCA_000010425.1
163 Bifidobacterium dentium JCM 1195 = DSM 20436 GCA_001042595.1
163 Bifidobacterium tsurumiense GCA_000741765.1
158 Bifidobacterium callitrichos DSM 23973 GCA_000741175.1
I1
217 Bifidobacterium adolescentis ATCC 15703 GCA_000010425.1
203 Bifidobacterium dentium JCM 1195 = DSM 20436 GCA_001042595.1
188 Bifidobacterium callitrichos DSM 23973 GCA_000741175.1
I2
203 Deinococcus radiodurans R1 GCA_000008565.1
126 Deinococcus proteolyticus MRP GCA_000190555.1
117 Deinococcus deserti VCD115 GCA_000020685.1
117 Deinococcus gobiensis I-0 GCA_000252445.1
117 Deinococcus soli Cha et al. 2016 GCA_001007995.1
I3
281 Rhodobacter sphaeroides 2.4.1 GCA_000012905.2
266 Rhodobacter sphaeroides ATCC 17025 GCA_000016405.1
255 Gemmobacter megaterium GCA_900156815.1
I4
314 Rhodobacter sphaeroides 2.4.1 GCA_000012905.2
270 Gemmobacter megaterium GCA_900156815.1
269 Rhodobacter sphaeroides ATCC 17025 GCA_000016405.1
I5
174 Clostridium saccharobutylicum DSM 13864 GCA_000473995.1
173 Clostridium beijerinckii GCA_000833105.2
173 Clostridium puniceum GCA_002006345.1
173 Clostridium saccharoperbutylacetonicum N1-4_28HMT_29 GCA_000340885.1
159 Clostridium chromiireducens GCA_002029255.1
I6
279 Deinococcus radiodurans R1 GCA_000008565.1
226 Deinococcus gobiensis I-0 GCA_000252445.1
208 Deinococcus puniceus GCA_001644565.1
I7
239 Deinococcus radiodurans R1 GCA_000008565.1
153 Deinococcus gobiensis I-0 GCA_000252445.1
147 Deinococcus deserti VCD115 GCA_000020685.1
I8
305 Deinococcus radiodurans R1 GCA_000008565.1
190 Deinococcus puniceus GCA_001644565.1
180 Deinococcus hopiensis KR-140 GCA_900176165.1
I9
172 Deinococcus radiodurans R1 GCA_000008565.1
112 Deinococcus gobiensis I-0 GCA_000252445.1
109 Deinococcus deserti VCD115 GCA_000020685.1
Ia
132 Enterococcus faecalis V583 GCA_000007785.1

132 Streptomyces cinnamoneus GCA_001885705.1
127 Melissococcus plutonius S1 GCA_000747585.1
115 Enterococcus columbae DSM 7374 = ATCC 51263 GCA_000406925.1
Ib
208 Clostridium beijerinckii GCA_000833105.2
208 Clostridium chromiireducens GCA_002029255.1
208 Clostridium puniceum GCA_002006345.1
208 Clostridium saccharobutylicum DSM 13864 GCA_000473995.1
208 Clostridium saccharoperbutylacetonicum N1-4_28HMT_29 GCA_000340885.1
194 Clostridium taeniosporum GCA_001735765.1
182 Clostridium fallax GCA_900129365.1
Ic
181 Lactobacillus gasseri ATCC 33323 = JCM 1131 GCA_000014425.1
152 Lactobacillus hominis DSM 23910 = CRBIP 24.179 GCA_000296835.1
137 Lactobacillus crispatus ST1 GCA_000091765.1
Id
290 Bacillus thuringiensis YBT-1518 GCA_000497525.2
287 Bacillus anthracis str. Ames GCA_000007845.1
287 Bacillus anthracis str. Sterne GCA_000008165.1
287 Bacillus cereus ATCC 14579 GCA_000007825.1
287 Bacillus mycoides GCA_000832605.1
287 Bacillus pseudomycoides DSM 12442 GCA_000161455.1
287 _5BBacillus thuringiensis_5D serovar konkukian str. 97-27 GCA_000008505.1
202 Bacillus halmapalus GCA_002019665.1
Ie
287 Bacillus thuringiensis YBT-1518 GCA_000497525.2
281 Bacillus anthracis str. Ames GCA_000007845.1
281 Bacillus anthracis str. Sterne GCA_000008165.1
281 Bacillus cereus ATCC 14579 GCA_000007825.1
281 Bacillus mycoides GCA_000832605.1
281 Bacillus pseudomycoides DSM 12442 GCA_000161455.1
281 _5BBacillus thuringiensis_5D serovar konkukian str. 97-27 GCA_000008505.1
220 Staphylococcus saprophyticus GCA_001074355.1
If
341 Staphylococcus epidermidis ATCC 12228 GCA_000007645.1
338 Staphylococcus haemolyticus JCSC1435 GCA_000009865.1
337 Staphylococcus lugdunensis HKU09-01 GCA_000025085.1
Ig
185 Streptococcus mutans UA159 GCA_000007465.2
85 Enterococcus asini ATCC 700915 GCA_000407365.1
85 Enterococcus canis NBRC 100695 GCA_001544375.1
85 Enterococcus casseliflavus EC20 GCA_000157355.2
85 Enterococcus dispar ATCC 51266 GCA_000406945.1
85 Enterococcus faecalis V583 GCA_000007785.1
85 Enterococcus faecium DO GCA_000174395.2
85 Enterococcus gilvus ATCC BAA-350 GCA_000407545.1
85 Enterococcus hermanniensis GCA_001885945.1
85 Enterococcus hirae ATCC 9790 GCA_000271405.2
85 Enterococcus italicus DSM 15952 GCA_000185365.1
85 Enterococcus malodoratus ATCC 43197 GCA_000407185.1
85 Enterococcus massiliensis GCA_001050095.1
85 Enterococcus mundtii QU 25 GCA_000504125.1
85 Enterococcus pallens ATCC BAA-351 GCA_000407485.1
85 Enterococcus pseudoavium NBRC 100491 GCA_001544295.1
85 Enterococcus rivorum GCA_001742285.1
85 Enterococcus saccharolyticus subsp. saccharolyticus ATCC 43076 GCA_000407285.1
85 Melissococcus plutonius S1 GCA_000747585.1
85 Streptomyces cinnamoneus GCA_001885705.1
85 Tetragenococcus halophilus NBRC 12172 GCA_000283615.1
83 Streptococcus marimammalium DSM 18627 GCA_000380045.1
83 Streptococcus sobrinus DSM 20742 = ATCC 33478 GCA_000686605.1
Ih
101 Bacillus anthracis str. Ames GCA_000007845.1
101 Bacillus anthracis str. Sterne GCA_000008165.1
101 Bacillus cereus ATCC 14579 GCA_000007825.1
101 Bacillus mycoides GCA_000832605.1
101 Bacillus pseudomycoides DSM 12442 GCA_000161455.1
101 Bacillus thuringiensis YBT-1518 GCA_000497525.2
101 _5BBacillus thuringiensis_5D serovar konkukian str. 97-27 GCA_000008505.1
85 Bacillus humi GCA_001439915.1
85 Escherichia coli IAI39 GCA_000026345.1
85 Escherichia coli O104_3AH4 str. 2011C-3493 GCA_000299455.1
85 Escherichia coli O157_3AH7 str. Sakai GCA_000008865.1
85 Escherichia coli O83_3AH1 str. NRG 857C GCA_000183345.1
85 Escherichia coli UMN026 GCA_000026325.2
85 Escherichia coli str. K-12 substr. MG1655 GCA_000005845.2

85 Shigella dysenteriae Sd197 GCA_000012005.1
85 Shigella flexneri 2a str. 301 GCA_000006925.2
85 Tumebacillus flagellatus GCA_000714935.1
81 Buttiauxella ferragutiae ATCC 51602 GCA_001654915.1
81 Cedecea neteri GCA_000757825.1
81 Citrobacter freundii CFNIH1 GCA_000648515.1
81 Cronobacter sakazakii GCA_000982825.1
81 Enterobacter cloacae subsp. cloacae ATCC 13047 GCA_000025565.1
81 Enterobacter hormaechei subsp. steigerwaltii GCA_001729725.1
81 Enterobacter kobei GCA_900185885.1
81 Erwinia iniecta GCA_001267535.1
81 Erwinia persicina NBRC 102418 GCA_001571305.1
81 Erwinia toletana DAPP-PG 735 GCA_000336255.1
81 Klebsiella aerogenes KCTC 2190 GCA_000215745.1
81 Klebsiella oxytoca GCA_001022195.1
81 Klebsiella pneumoniae subsp. pneumoniae HS11286 GCA_000240185.2
81 Kluyvera ascorbata ATCC 33433 GCA_000735365.1
81 Kluyvera cryocrescens NBRC 102467 GCA_001571285.1
81 Kosakonia cowanii GCA_001975225.1
81 Kosakonia sacchari SP1 GCA_000300455.4
81 Mangrovibacter phragmitis GCA_001655675.1
81 Pseudescherichia vulneris NBRC 102420 GCA_000759795.1
81 Rosenbergiella nectarea GCA_900111105.1
81 Salmonella enterica subsp. enterica serovar Typhi str. CT18 GCA_000195995.1
81 Serratia marcescens subsp. marcescens Db11 GCA_000513215.1
81 Yokenella regensburgei ATCC 49455 GCA_000735455.1
Ii
269 Streptococcus mutans UA159 GCA_000007465.2
191 Streptococcus iniae GCA_000831485.1
188 Streptococcus ferus DSM 20646 GCA_000372425.1
Ij
213 Clostridium beijerinckii GCA_000833105.2
213 Clostridium saccharobutylicum DSM 13864 GCA_000473995.1
213 Clostridium saccharoperbutylacetonicum N1-4_28HMT_29 GCA_000340885.1
210 Clostridium butyricum GCA_001456065.2
206 Clostridium puniceum GCA_002006345.1
Ik
201 Streptococcus mutans UA159 GCA_000007465.2
143 Streptococcus marimammalium DSM 18627 GCA_000380045.1
142 Streptococcus equinus GCA_000964315.1
142 Streptococcus gallolyticus subsp. gallolyticus DSM 16831 GCA_002000985.1
142 Streptococcus minor DSM 17118 GCA_000377005.1
Il
205 Clostridium beijerinckii GCA_000833105.2
205 Clostridium saccharobutylicum DSM 13864 GCA_000473995.1
205 Clostridium saccharoperbutylacetonicum N1-4_28HMT_29 GCA_000340885.1
198 Clostridium puniceum GCA_002006345.1
194 Clostridium butyricum GCA_001456065.2
Im
220 Deinococcus radiodurans R1 GCA_000008565.1
155 Deinococcus deserti VCD115 GCA_000020685.1
153 Deinococcus puniceus GCA_001644565.1
In
266 Deinococcus radiodurans R1 GCA_000008565.1
221 Deinococcus gobiensis I-0 GCA_000252445.1
193 Deinococcus deserti VCD115 GCA_000020685.1
Io
224 Lactobacillus gasseri ATCC 33323 = JCM 1131 GCA_000014425.1
224 Lactobacillus hominis DSM 23910 = CRBIP 24.179 GCA_000296835.1
155 Lactobacillus crispatus ST1 GCA_000091765.1
155 Lactobacillus helveticus GCA_001308285.1
155 Lactobacillus kefiranofaciens ZW3 GCA_000214785.1
155 Lactobacillus psittaci DSM 15354 GCA_000425905.1
154 Lactobacillus iners DSM 13335 GCA_000160875.1
Ip
257 Streptococcus mutans UA159 GCA_000007465.2
167 Streptococcus iniae GCA_000831485.1
163 Streptococcus cristatus AS 1.3089 GCA_000385925.1
163 Streptococcus gordonii str. Challis substr. CH1 GCA_000017005.1
163 Streptococcus mitis B6 GCA_000027165.1
163 Streptococcus pneumoniae R6 GCA_000007045.1
Iq
180 Escherichia coli IAI39 GCA_000026345.1
180 Escherichia coli O104_3AH4 str. 2011C-3493 GCA_000299455.1
180 Escherichia coli O157_3AH7 str. Sakai GCA_000008865.1
180 Escherichia coli O83_3AH1 str. NRG 857C GCA_000183345.1

180 Escherichia coli UMN026 GCA_000026325.2
180 Escherichia coli str. K-12 substr. MG1655 GCA_000005845.2
180 Shigella dysenteriae Sd197 GCA_000012005.1
180 Shigella flexneri 2a str. 301 GCA_000006925.2
180 Tumebacillus flagellatus GCA_000714935.1
167 Pluralibacter gergoviae GCA_000757785.1
166 Pantoea agglomerans GCA_001709315.1
Ir
195 Streptococcus mutans UA159 GCA_000007465.2
162 Streptococcus marimammalium DSM 18627 GCA_000380045.1
158 Streptococcus sobrinus DSM 20742 = ATCC 33478 GCA_000686605.1
Is
185 Clostridium beijerinckii GCA_000833105.2
185 Clostridium puniceum GCA_002006345.1
185 Clostridium saccharoperbutylacetonicum N1-4_28HMT_29 GCA_000340885.1
181 Clostridium saccharobutylicum DSM 13864 GCA_000473995.1
178 Clostridium taeniosporum GCA_001735765.1
It
157 Streptococcus mutans UA159 GCA_000007465.2
93 Streptococcus gordonii str. Challis substr. CH1 GCA_000017005.1
93 Streptococcus phocae subsp. salmonis GCA_000772915.1
88 Streptococcus ratti FA-1 = DSM 20564 GCA_000286075.1
Iu
275 Streptococcus mutans UA159 GCA_000007465.2
182 Streptococcus equinus GCA_000964315.1
182 Streptococcus gallolyticus subsp. gallolyticus DSM 16831 GCA_002000985.1
178 Streptococcus cristatus AS 1.3089 GCA_000385925.1
178 Streptococcus gordonii str. Challis substr. CH1 GCA_000017005.1
Iv
202 Rhodobacter sphaeroides 2.4.1 GCA_000012905.2
185 Gemmobacter aquatilis GCA_900110025.1
173 Pseudorhodobacter psychrotolerans GCA_001294535.1
Iw
284 Bacillus anthracis str. Ames GCA_000007845.1
284 Bacillus anthracis str. Sterne GCA_000008165.1
284 Bacillus cereus ATCC 14579 GCA_000007825.1
284 Bacillus mycoides GCA_000832605.1
284 Bacillus pseudomycoides DSM 12442 GCA_000161455.1
284 Bacillus thuringiensis YBT-1518 GCA_000497525.2
284 _5BBacillus thuringiensis_5D serovar konkukian str. 97-27 GCA_000008505.1
238 Bacillus manliponensis GCA_000712595.1
229 Bacillus horneckiae GCA_001636335.1
229 Bacillus solani GCA_001420595.1
Ix
231 Bacillus cereus ATCC 14579 GCA_000007825.1
220 Bacillus megaterium NBRC 15308 = ATCC 14581 GCA_000832985.1
219 Bacillus flexus GCA_002024265.1
Iy
203 Deinococcus radiodurans R1 GCA_000008565.1
150 Deinococcus deserti VCD115 GCA_000020685.1
137 Deinococcus gobiensis I-0 GCA_000252445.1
Iz
183 Lactobacillus gasseri ATCC 33323 = JCM 1131 GCA_000014425.1
165 Lactobacillus hominis DSM 23910 = CRBIP 24.179 GCA_000296835.1
164 Streptococcus mutans UA159 GCA_000007465.2
IA
174 Clostridium beijerinckii GCA_000833105.2
174 Clostridium puniceum GCA_002006345.1
174 Clostridium saccharoperbutylacetonicum N1-4_28HMT_29 GCA_000340885.1
165 Clostridium saccharobutylicum DSM 13864 GCA_000473995.1
159 Clostridium butyricum GCA_001456065.2
IB
214 Deinococcus radiodurans R1 GCA_000008565.1
119 Deinococcus deserti VCD115 GCA_000020685.1
119 Deinococcus gobiensis I-0 GCA_000252445.1
103 Deinococcus frigens DSM 12807 GCA_000701425.1
103 Deinococcus marmoris DSM 12784 GCA_000701405.1
103 Deinococcus swuensis GCA_000800395.1
IC
207 Deinococcus radiodurans R1 GCA_000008565.1
164 Deinococcus gobiensis I-0 GCA_000252445.1
163 Deinococcus puniceus GCA_001644565.1
ID
186 Bacillus thuringiensis YBT-1518 GCA_000497525.2
171 Bacillus anthracis str. Ames GCA_000007845.1
171 Bacillus anthracis str. Sterne GCA_000008165.1

171 Bacillus cereus ATCC 14579 GCA_000007825.1
171 Bacillus mycoides GCA_000832605.1
171 Bacillus pseudomycoides DSM 12442 GCA_000161455.1
171 _5BBacillus thuringiensis_5D serovar konkukian str. 97-27 GCA_000008505.1
170 Bacillus marisflavi GCA_001274775.1
IE
121 Deinococcus radiodurans R1 GCA_000008565.1
112 Staphylococcus capitis subsp. capitis GCA_001028645.1
112 Staphylococcus epidermidis ATCC 12228 GCA_000007645.1
112 Staphylococcus haemolyticus JCSC1435 GCA_000009865.1
112 Staphylococcus pettenkoferi GCA_002208805.1
101 Staphylococcus arlettae CVD059 GCA_000295715.1
101 Staphylococcus cohnii subsp. cohnii GCA_000972575.1
101 Staphylococcus hominis subsp. hominis C80 GCA_000183685.1
101 Staphylococcus lugdunensis HKU09-01 GCA_000025085.1
101 Staphylococcus microti GCA_000934465.1
101 Staphylococcus saprophyticus subsp. saprophyticus ATCC 15305 GCA_000010125.1
IF
262 Deinococcus radiodurans R1 GCA_000008565.1
166 Deinococcus deserti VCD115 GCA_000020685.1
160 Deinococcus puniceus GCA_001644565.1
IG
114 Rhodobacter sphaeroides 2.4.1 GCA_000012905.2
112 Sagittula stellata E-37 GCA_000169415.1
108 Aliiroseovarius crassostreae GCA_001307765.1
108 Aliiroseovarius sediminilitoris GCA_900109955.1
108 Marivita cryptomonadis GCA_002115725.1
108 Sedimentitalea nanhaiensis DSM 24252 GCA_000473225.1
108 Thioclava dalianensis GCA_000715505.1
108 Thioclava indica GCA_000714545.1
IH
120 Rhodobacter sphaeroides 2.4.1 GCA_000012905.2
105 Gemmobacter aquatilis GCA_900110025.1
105 Gemmobacter megaterium GCA_900156815.1
105 Pseudorhodobacter ferrugineus DSM 5888 GCA_000420745.1
105 Pseudorhodobacter psychrotolerans GCA_001294535.1
105 Pseudorhodobacter wandonensis GCA_001202035.1
105 Rhodobacter sphaeroides ATCC 17025 GCA_000016405.1
102 Defluviimonas alba GCA_001620265.1
II
168 Bifidobacterium adolescentis ATCC 15703 GCA_000010425.1
168 Bifidobacterium breve DSM 20213 = JCM 1192 GCA_001025175.1
168 Bifidobacterium thermophilum GCA_000741495.1
163 Bifidobacterium gallicum DSM 20093 = LMG 11596 GCA_000741205.1
160 Bifidobacterium longum NCC2705 GCA_000007525.1
IJ
112 Rhodobacter sphaeroides ATCC 17025 GCA_000016405.1
97 Rhodobacter sphaeroides 2.4.1 GCA_000012905.2
84 Pseudorhodobacter ferrugineus DSM 5888 GCA_000420745.1
84 Pseudorhodobacter psychrotolerans GCA_001294535.1
84 Pseudorhodobacter wandonensis GCA_001202035.1
IK
221 Lactobacillus gasseri ATCC 33323 = JCM 1131 GCA_000014425.1
221 Lactobacillus hominis DSM 23910 = CRBIP 24.179 GCA_000296835.1
193 Lactobacillus acidophilus NCFM GCA_000011985.1
190 Lactobacillus amylophilus DSM 20533 = JCM 1125 GCA_001936335.1
IL
191 Clostridium beijerinckii GCA_000833105.2
191 Clostridium butyricum GCA_001456065.2
191 Clostridium saccharobutylicum DSM 13864 GCA_000473995.1
191 Clostridium saccharoperbutylacetonicum N1-4_28HMT_29 GCA_000340885.1
187 Clostridium chromiireducens GCA_002029255.1
187 Clostridium puniceum GCA_002006345.1
174 Clostridium neonatale GCA_001458595.1
IM
214 Streptococcus mutans UA159 GCA_000007465.2
168 Streptococcus ferus DSM 20646 GCA_000372425.1
156 Streptococcus macacae NCTC 11558 GCA_000187995.3
IN
163 Deinococcus radiodurans R1 GCA_000008565.1
124 Deinococcus gobiensis I-0 GCA_000252445.1
102 Deinococcus puniceus GCA_001644565.1
IO
293 Escherichia coli IAI39 GCA_000026345.1
293 Escherichia coli O104_3AH4 str. 2011C-3493 GCA_000299455.1
293 Escherichia coli O157_3AH7 str. Sakai GCA_000008865.1

293 Escherichia coli O83_3AH1 str. NRG 857C GCA_000183345.1
293 Escherichia coli UMN026 GCA_000026325.2
293 Escherichia coli str. K-12 substr. MG1655 GCA_000005845.2
293 Shigella dysenteriae Sd197 GCA_000012005.1
293 Shigella flexneri 2a str. 301 GCA_000006925.2
293 Tumebacillus flagellatus GCA_000714935.1
273 Erwinia iniecta GCA_001267535.1
255 Rosenbergiella nectarea GCA_900111105.1
IP
252 Clostridium beijerinckii GCA_000833105.2
252 Clostridium saccharoperbutylacetonicum N1-4_28HMT_29 GCA_000340885.1
244 Clostridium saccharobutylicum DSM 13864 GCA_000473995.1
239 Clostridium butyricum GCA_001456065.2
IQ
245 Rhodobacter sphaeroides 2.4.1 GCA_000012905.2
210 Gemmobacter megaterium GCA_900156815.1
209 Pseudorhodobacter ferrugineus DSM 5888 GCA_000420745.1
209 Pseudorhodobacter wandonensis GCA_001202035.1
209 Rhodobacter sphaeroides ATCC 17025 GCA_000016405.1
IR
122 Bifidobacterium adolescentis ATCC 15703 GCA_000010425.1
122 Bifidobacterium dentium JCM 1195 = DSM 20436 GCA_001042595.1
122 Bifidobacterium tsurumiense GCA_000741765.1
119 Bifidobacterium angulatum DSM 20098 = JCM 7096 GCA_001025155.1
119 Bifidobacterium callitrichos DSM 23973 GCA_000741175.1
119 Bifidobacterium lemurum GCA_001895165.1
119 Bifidobacterium thermophilum GCA_000741495.1
118 Bifidobacterium stellenboschense GCA_000741785.1
IS
201 Clostridium beijerinckii GCA_000833105.2
201 Clostridium puniceum GCA_002006345.1
201 Clostridium saccharobutylicum DSM 13864 GCA_000473995.1
201 Clostridium saccharoperbutylacetonicum N1-4_28HMT_29 GCA_000340885.1
192 Clostridium neonatale GCA_001458595.1
190 Clostridium taeniosporum GCA_001735765.1
IT
235 Deinococcus radiodurans R1 GCA_000008565.1
180 Deinococcus gobiensis I-0 GCA_000252445.1
173 Deinococcus puniceus GCA_001644565.1
IU
242 Clostridium beijerinckii GCA_000833105.2
242 Clostridium puniceum GCA_002006345.1
242 Clostridium saccharobutylicum DSM 13864 GCA_000473995.1
242 Clostridium saccharoperbutylacetonicum N1-4_28HMT_29 GCA_000340885.1
234 Clostridium butyricum GCA_001456065.2
231 Clostridium chromiireducens GCA_002029255.1
IV
239 Bacillus anthracis str. Ames GCA_000007845.1
239 Bacillus anthracis str. Sterne GCA_000008165.1
239 Bacillus mycoides GCA_000832605.1
239 Bacillus pseudomycoides DSM 12442 GCA_000161455.1
239 Bacillus thuringiensis YBT-1518 GCA_000497525.2
239 _5BBacillus thuringiensis_5D serovar konkukian str. 97-27 GCA_000008505.1
238 Bacillus cereus ATCC 14579 GCA_000007825.1
191 Bacillus manliponensis GCA_000712595.1
IW
217 Clostridium beijerinckii GCA_000833105.2
217 Clostridium puniceum GCA_002006345.1
217 Clostridium saccharoperbutylacetonicum N1-4_28HMT_29 GCA_000340885.1
208 Clostridium saccharobutylicum DSM 13864 GCA_000473995.1
197 Clostridium chromiireducens GCA_002029255.1
IX
230 Deinococcus radiodurans R1 GCA_000008565.1
166 Deinococcus gobiensis I-0 GCA_000252445.1
164 Deinococcus puniceus GCA_001644565.1
IY
249 Staphylococcus equorum GCA_001432245.1
242 Staphylococcus lentus F1142 GCA_000286395.1
242 Staphylococcus sciuri GCA_002209165.1
237 Bacillus anthracis str. Ames GCA_000007845.1
237 Bacillus anthracis str. Sterne GCA_000008165.1
237 Bacillus cereus ATCC 14579 GCA_000007825.1
237 Bacillus mycoides GCA_000832605.1
237 Bacillus pseudomycoides DSM 12442 GCA_000161455.1
237 Bacillus thuringiensis YBT-1518 GCA_000497525.2
237 _5BBacillus thuringiensis_5D serovar konkukian str. 97-27 GCA_000008505.1

IZ
216 Enterococcus faecalis V583 GCA_000007785.1
216 Streptomyces cinnamoneus GCA_001885705.1
198 Isobaculum melis GCA_900111355.1
197 Enterococcus asini ATCC 700915 GCA_000407365.1
197 Enterococcus casseliflavus EC20 GCA_000157355.2
197 Enterococcus dispar ATCC 51266 GCA_000406945.1
197 Enterococcus massiliensis GCA_001050095.1
197 Enterococcus saccharolyticus subsp. saccharolyticus ATCC 43076 GCA_000407285.1
J0
273 Streptococcus mutans UA159 GCA_000007465.2
178 Streptococcus equinus GCA_000964315.1
178 Streptococcus gallolyticus subsp. gallolyticus DSM 16831 GCA_002000985.1
166 Streptococcus cristatus AS 1.3089 GCA_000385925.1
166 Streptococcus gordonii str. Challis substr. CH1 GCA_000017005.1
166 Streptococcus mitis B6 GCA_000027165.1
166 Streptococcus pneumoniae R6 GCA_000007045.1
J1
227 Escherichia coli IAI39 GCA_000026345.1
227 Escherichia coli O104_3AH4 str. 2011C-3493 GCA_000299455.1
227 Escherichia coli O157_3AH7 str. Sakai GCA_000008865.1
227 Escherichia coli UMN026 GCA_000026325.2
227 Escherichia coli str. K-12 substr. MG1655 GCA_000005845.2
227 Shigella flexneri 2a str. 301 GCA_000006925.2
227 Tumebacillus flagellatus GCA_000714935.1
226 Escherichia coli O83_3AH1 str. NRG 857C GCA_000183345.1
212 Shigella dysenteriae Sd197 GCA_000012005.1
J2
219 Rhodobacter sphaeroides 2.4.1 GCA_000012905.2
194 Rhodobacter sphaeroides ATCC 17025 GCA_000016405.1
182 Haematobacter massiliensis GCA_000740795.1
J3
285 Enterococcus faecalis V583 GCA_000007785.1
285 Streptomyces cinnamoneus GCA_001885705.1
252 Enterococcus asini ATCC 700915 GCA_000407365.1
252 Enterococcus canis NBRC 100695 GCA_001544375.1
252 Enterococcus dispar ATCC 51266 GCA_000406945.1
252 Enterococcus faecium DO GCA_000174395.2
252 Enterococcus hirae ATCC 9790 GCA_000271405.2
252 Enterococcus mundtii QU 25 GCA_000504125.1
252 Enterococcus rivorum GCA_001742285.1
244 Enterococcus haemoperoxidus ATCC BAA-382 GCA_000407165.1
244 Enterococcus phoeniculicola ATCC BAA-412 GCA_000407505.1
244 Enterococcus thailandicus GCA_001652875.1
J4
125 Enterococcus faecalis V583 GCA_000007785.1
125 Streptomyces cinnamoneus GCA_001885705.1
111 Enterococcus hirae ATCC 9790 GCA_000271405.2
110 Enterococcus aquimarinus GCA_001885765.1
110 Enterococcus canis NBRC 100695 GCA_001544375.1
110 Enterococcus casseliflavus EC20 GCA_000157355.2
110 Enterococcus dispar ATCC 51266 GCA_000406945.1
110 Enterococcus faecium DO GCA_000174395.2
110 Enterococcus gilvus ATCC BAA-350 GCA_000407545.1
110 Enterococcus haemoperoxidus ATCC BAA-382 GCA_000407165.1
110 Enterococcus hermanniensis GCA_001885945.1
110 Enterococcus malodoratus ATCC 43197 GCA_000407185.1
110 Enterococcus massiliensis GCA_001050095.1
110 Enterococcus mundtii QU 25 GCA_000504125.1
110 Enterococcus pallens ATCC BAA-351 GCA_000407485.1
110 Enterococcus phoeniculicola ATCC BAA-412 GCA_000407505.1
110 Enterococcus pseudoavium NBRC 100491 GCA_001544295.1
110 Enterococcus rivorum GCA_001742285.1
110 Enterococcus saccharolyticus subsp. saccharolyticus ATCC 43076 GCA_000407285.1
110 Enterococcus thailandicus GCA_001652875.1
J5
237 Escherichia coli IAI39 GCA_000026345.1
237 Escherichia coli O104_3AH4 str. 2011C-3493 GCA_000299455.1
237 Escherichia coli O157_3AH7 str. Sakai GCA_000008865.1
237 Escherichia coli O83_3AH1 str. NRG 857C GCA_000183345.1
237 Escherichia coli UMN026 GCA_000026325.2
237 Escherichia coli str. K-12 substr. MG1655 GCA_000005845.2
237 Shigella flexneri 2a str. 301 GCA_000006925.2
237 Tumebacillus flagellatus GCA_000714935.1
228 Shigella dysenteriae Sd197 GCA_000012005.1
211 Erwinia iniecta GCA_001267535.1

J6
151 Deinococcus radiodurans R1 GCA_000008565.1
134 Deinococcus geothermalis DSM 11300 GCA_000196275.1
129 Deinococcus gobiensis I-0 GCA_000252445.1
J7
215 Staphylococcus epidermidis ATCC 12228 GCA_000007645.1
215 Staphylococcus haemolyticus JCSC1435 GCA_000009865.1
215 Staphylococcus hominis subsp. hominis C80 GCA_000183685.1
215 Staphylococcus lugdunensis HKU09-01 GCA_000025085.1
206 Staphylococcus aureus subsp. aureus NCTC 8325 GCA_000013425.1
206 Staphylococcus simiae CCM 7213 GCA_000235645.2
200 Staphylococcus capitis subsp. capitis GCA_001028645.1
J8
233 Clostridium beijerinckii GCA_000833105.2
233 Clostridium saccharoperbutylacetonicum N1-4_28HMT_29 GCA_000340885.1
225 Clostridium saccharobutylicum DSM 13864 GCA_000473995.1
218 Clostridium puniceum GCA_002006345.1
J9
164 Clostridium beijerinckii GCA_000833105.2
164 Clostridium puniceum GCA_002006345.1
164 Clostridium saccharoperbutylacetonicum N1-4_28HMT_29 GCA_000340885.1
156 Clostridium chromiireducens GCA_002029255.1
156 Clostridium saccharobutylicum DSM 13864 GCA_000473995.1
150 Clostridium taeniosporum GCA_001735765.1
Ja
272 Escherichia coli IAI39 GCA_000026345.1
272 Escherichia coli O104_3AH4 str. 2011C-3493 GCA_000299455.1
272 Escherichia coli O157_3AH7 str. Sakai GCA_000008865.1
272 Escherichia coli O83_3AH1 str. NRG 857C GCA_000183345.1
272 Escherichia coli UMN026 GCA_000026325.2
272 Escherichia coli str. K-12 substr. MG1655 GCA_000005845.2
272 Shigella dysenteriae Sd197 GCA_000012005.1
272 Shigella flexneri 2a str. 301 GCA_000006925.2
272 Tumebacillus flagellatus GCA_000714935.1
239 Erwinia iniecta GCA_001267535.1
220 Rosenbergiella nectarea GCA_900111105.1
Jb
211 Enterococcus faecalis V583 GCA_000007785.1
211 Streptomyces cinnamoneus GCA_001885705.1
194 Enterococcus dispar ATCC 51266 GCA_000406945.1
192 Enterococcus faecium DO GCA_000174395.2
Jc
245 Lactobacillus gasseri ATCC 33323 = JCM 1131 GCA_000014425.1
227 Lactobacillus hominis DSM 23910 = CRBIP 24.179 GCA_000296835.1
173 Lactobacillus psittaci DSM 15354 GCA_000425905.1
Jd
220 Clostridium beijerinckii GCA_000833105.2
220 Clostridium saccharoperbutylacetonicum N1-4_28HMT_29 GCA_000340885.1
207 Clostridium saccharobutylicum DSM 13864 GCA_000473995.1
206 Clostridium puniceum GCA_002006345.1
Je
183 Clostridium beijerinckii GCA_000833105.2
183 Clostridium puniceum GCA_002006345.1
183 Clostridium saccharobutylicum DSM 13864 GCA_000473995.1
183 Clostridium saccharoperbutylacetonicum N1-4_28HMT_29 GCA_000340885.1
169 Clostridium chromiireducens GCA_002029255.1
166 Clostridium neonatale GCA_001458595.1
Jf
225 Clostridium butyricum GCA_001456065.2
218 Clostridium saccharobutylicum DSM 13864 GCA_000473995.1
216 Clostridium beijerinckii GCA_000833105.2
216 Clostridium saccharoperbutylacetonicum N1-4_28HMT_29 GCA_000340885.1
Jg
210 Clostridium butyricum GCA_001456065.2
208 Clostridium beijerinckii GCA_000833105.2
208 Clostridium puniceum GCA_002006345.1
208 Clostridium saccharobutylicum DSM 13864 GCA_000473995.1
208 Clostridium saccharoperbutylacetonicum N1-4_28HMT_29 GCA_000340885.1
197 Clostridium chromiireducens GCA_002029255.1
Jh
282 Deinococcus radiodurans R1 GCA_000008565.1
187 Deinococcus puniceus GCA_001644565.1
183 Deinococcus gobiensis I-0 GCA_000252445.1
Ji
197 Streptococcus mutans UA159 GCA_000007465.2
171 Streptococcus sobrinus DSM 20742 = ATCC 33478 GCA_000686605.1

157 Streptococcus ferus DSM 20646 GCA_000372425.1
Jj
220 Deinococcus radiodurans R1 GCA_000008565.1
147 Deinococcus gobiensis I-0 GCA_000252445.1
146 Deinococcus puniceus GCA_001644565.1
Jk
138 Staphylococcus pettenkoferi GCA_002208805.1
135 Staphylococcus capitis subsp. capitis GCA_001028645.1
135 Staphylococcus epidermidis ATCC 12228 GCA_000007645.1
132 Staphylococcus cohnii subsp. cohnii GCA_000972575.1
Jl
239 Clostridium beijerinckii GCA_000833105.2
239 Clostridium saccharobutylicum DSM 13864 GCA_000473995.1
239 Clostridium saccharoperbutylacetonicum N1-4_28HMT_29 GCA_000340885.1
233 Clostridium puniceum GCA_002006345.1
215 Clostridium butyricum GCA_001456065.2
215 Clostridium chromiireducens GCA_002029255.1
Jm
290 Rhodobacter sphaeroides 2.4.1 GCA_000012905.2
255 Pseudorhodobacter psychrotolerans GCA_001294535.1
254 Rhodobacter sphaeroides ATCC 17025 GCA_000016405.1
Jn
197 Escherichia coli IAI39 GCA_000026345.1
197 Escherichia coli O104_3AH4 str. 2011C-3493 GCA_000299455.1
197 Escherichia coli O157_3AH7 str. Sakai GCA_000008865.1
197 Escherichia coli O83_3AH1 str. NRG 857C GCA_000183345.1
197 Escherichia coli UMN026 GCA_000026325.2
197 Escherichia coli str. K-12 substr. MG1655 GCA_000005845.2
197 Shigella dysenteriae Sd197 GCA_000012005.1
197 Shigella flexneri 2a str. 301 GCA_000006925.2
197 Tumebacillus flagellatus GCA_000714935.1
191 Erwinia iniecta GCA_001267535.1
164 Erwinia toletana DAPP-PG 735 GCA_000336255.1
Jo
242 Streptococcus mutans UA159 GCA_000007465.2
183 Streptococcus marimammalium DSM 18627 GCA_000380045.1
173 Streptococcus macacae NCTC 11558 GCA_000187995.3
Jp
212 Deinococcus radiodurans R1 GCA_000008565.1
124 Deinococcus puniceus GCA_001644565.1
121 Deinococcus frigens DSM 12807 GCA_000701425.1
Jq
201 Bacillus anthracis str. Ames GCA_000007845.1
201 Bacillus anthracis str. Sterne GCA_000008165.1
201 Bacillus cereus ATCC 14579 GCA_000007825.1
201 Bacillus mycoides GCA_000832605.1
201 Bacillus pseudomycoides DSM 12442 GCA_000161455.1
201 Bacillus thuringiensis YBT-1518 GCA_000497525.2
201 _5BBacillus thuringiensis_5D serovar konkukian str. 97-27 GCA_000008505.1
176 Staphylococcus equorum GCA_001432245.1
169 Bacillus aquimaris TF-12 GCA_001648555.1
Jr
169 Rhodobacter sphaeroides ATCC 17025 GCA_000016405.1
153 Rhodobacter sphaeroides 2.4.1 GCA_000012905.2
126 Pseudorhodobacter ferrugineus DSM 5888 GCA_000420745.1
126 Pseudorhodobacter wandonensis GCA_001202035.1
Js
218 Pseudorhodobacter ferrugineus DSM 5888 GCA_000420745.1
218 Pseudorhodobacter wandonensis GCA_001202035.1
215 Rhodobacter sphaeroides 2.4.1 GCA_000012905.2
203 Gemmobacter megaterium GCA_900156815.1
203 Rhodobacter sphaeroides ATCC 17025 GCA_000016405.1
Jt
165 Rhodobacter sphaeroides 2.4.1 GCA_000012905.2
150 Rhodobacter sphaeroides ATCC 17025 GCA_000016405.1
136 Haematobacter massiliensis GCA_000740795.1
Ju
162 Clostridium saccharobutylicum DSM 13864 GCA_000473995.1
161 Clostridium beijerinckii GCA_000833105.2
161 Clostridium puniceum GCA_002006345.1
161 Clostridium saccharoperbutylacetonicum N1-4_28HMT_29 GCA_000340885.1
154 Clostridium butyricum GCA_001456065.2
Jv
213 Deinococcus radiodurans R1 GCA_000008565.1
168 Deinococcus marmoris DSM 12784 GCA_000701405.1
168 Deinococcus swuensis GCA_000800395.1

153 Deinococcus frigens DSM 12807 GCA_000701425.1
Jw
245 Clostridium neonatale GCA_001458595.1
244 Clostridium butyricum GCA_001456065.2
236 Clostridium beijerinckii GCA_000833105.2
236 Clostridium puniceum GCA_002006345.1
236 Clostridium saccharobutylicum DSM 13864 GCA_000473995.1
236 Clostridium saccharoperbutylacetonicum N1-4_28HMT_29 GCA_000340885.1
Jx
259 Deinococcus radiodurans R1 GCA_000008565.1
147 Deinococcus deserti VCD115 GCA_000020685.1
144 Deinococcus gobiensis I-0 GCA_000252445.1
Jy
230 Clostridium beijerinckii GCA_000833105.2
230 Clostridium saccharobutylicum DSM 13864 GCA_000473995.1
230 Clostridium saccharoperbutylacetonicum N1-4_28HMT_29 GCA_000340885.1
224 Clostridium puniceum GCA_002006345.1
213 Clostridium chromiireducens GCA_002029255.1
Jz
248 Rhodobacter sphaeroides 2.4.1 GCA_000012905.2
208 Rhodobacter sphaeroides ATCC 17025 GCA_000016405.1
176 Pseudorhodobacter psychrotolerans GCA_001294535.1
JA
54 Izhakiella capsodis GCA_900115045.1
50 Shewanella amazonensis SB2B GCA_000015245.1
43 Microbulbifer donghaiensis GCA_900129095.1
43 Microbulbifer marinus GCA_900107725.1
43 Microbulbifer yueqingensis GCA_900100355.1
JB
213 Clostridium saccharobutylicum DSM 13864 GCA_000473995.1
207 Clostridium butyricum GCA_001456065.2
196 Clostridium beijerinckii GCA_000833105.2
196 Clostridium puniceum GCA_002006345.1
196 Clostridium saccharoperbutylacetonicum N1-4_28HMT_29 GCA_000340885.1
JC
218 Deinococcus radiodurans R1 GCA_000008565.1
124 Deinococcus frigens DSM 12807 GCA_000701425.1
124 Deinococcus marmoris DSM 12784 GCA_000701405.1
124 Deinococcus swuensis GCA_000800395.1
121 Deinococcus gobiensis I-0 GCA_000252445.1
JD
245 Deinococcus radiodurans R1 GCA_000008565.1
177 Deinococcus gobiensis I-0 GCA_000252445.1
163 Deinococcus puniceus GCA_001644565.1
JE
150 Staphylococcus capitis subsp. capitis GCA_001028645.1
150 Staphylococcus epidermidis ATCC 12228 GCA_000007645.1
150 Staphylococcus haemolyticus JCSC1435 GCA_000009865.1
139 Staphylococcus arlettae CVD059 GCA_000295715.1
139 Staphylococcus cohnii subsp. cohnii GCA_000972575.1
139 Staphylococcus hominis subsp. hominis C80 GCA_000183685.1
139 Staphylococcus lugdunensis HKU09-01 GCA_000025085.1
139 Staphylococcus saprophyticus subsp. saprophyticus ATCC 15305 GCA_000010125.1
130 Staphylococcus aureus subsp. aureus NCTC 8325 GCA_000013425.1
130 Staphylococcus gallinarum GCA_000875895.1
130 Staphylococcus simiae CCM 7213 GCA_000235645.2
130 Staphylococcus simulans GCA_001559115.1
130 Staphylococcus succinus GCA_001902315.1
130 Staphylococcus xylosus GCA_000706685.1
JF
175 Deinococcus radiodurans R1 GCA_000008565.1
111 Deinococcus gobiensis I-0 GCA_000252445.1
106 Deinococcus hopiensis KR-140 GCA_900176165.1
JG
297 Deinococcus radiodurans R1 GCA_000008565.1
188 Deinococcus proteolyticus MRP GCA_000190555.1
185 Deinococcus gobiensis I-0 GCA_000252445.1
JH
188 Streptococcus mutans UA159 GCA_000007465.2
134 Streptococcus ferus DSM 20646 GCA_000372425.1
132 Streptococcus sobrinus DSM 20742 = ATCC 33478 GCA_000686605.1
JI
244 Deinococcus radiodurans R1 GCA_000008565.1
185 Deinococcus gobiensis I-0 GCA_000252445.1
158 Deinococcus puniceus GCA_001644565.1
JJ

140 Escherichia coli IAI39 GCA_000026345.1
140 Escherichia coli O104_3AH4 str. 2011C-3493 GCA_000299455.1
140 Escherichia coli O157_3AH7 str. Sakai GCA_000008865.1
140 Escherichia coli O83_3AH1 str. NRG 857C GCA_000183345.1
140 Escherichia coli UMN026 GCA_000026325.2
140 Escherichia coli str. K-12 substr. MG1655 GCA_000005845.2
140 Shigella dysenteriae Sd197 GCA_000012005.1
140 Shigella flexneri 2a str. 301 GCA_000006925.2
140 Tumebacillus flagellatus GCA_000714935.1
134 Erwinia iniecta GCA_001267535.1
125 Xenorhabdus bovienii SS-2004 GCA_000027225.1
JK
161 Staphylococcus equorum GCA_001432245.1
158 Enterococcus faecalis V583 GCA_000007785.1
158 Streptomyces cinnamoneus GCA_001885705.1
146 Staphylococcus succinus GCA_001902315.1
JL
153 Deinococcus radiodurans R1 GCA_000008565.1
109 Deinococcus puniceus GCA_001644565.1
104 Deinococcus hopiensis KR-140 GCA_900176165.1
JM
283 Bacillus anthracis str. Ames GCA_000007845.1
283 Bacillus anthracis str. Sterne GCA_000008165.1
283 Bacillus cereus ATCC 14579 GCA_000007825.1
283 Bacillus mycoides GCA_000832605.1
283 Bacillus pseudomycoides DSM 12442 GCA_000161455.1
283 Bacillus thuringiensis YBT-1518 GCA_000497525.2
283 _5BBacillus thuringiensis_5D serovar konkukian str. 97-27 GCA_000008505.1
229 Bacillus manliponensis GCA_000712595.1
222 Bacillus marisflavi GCA_001274775.1
JN
270 Clostridium beijerinckii GCA_000833105.2
270 Clostridium puniceum GCA_002006345.1
270 Clostridium saccharoperbutylacetonicum N1-4_28HMT_29 GCA_000340885.1
255 Clostridium butyricum GCA_001456065.2
255 Clostridium saccharobutylicum DSM 13864 GCA_000473995.1
250 Clostridium taeniosporum GCA_001735765.1
JO
172 Deinococcus radiodurans R1 GCA_000008565.1
101 Deinococcus gobiensis I-0 GCA_000252445.1
98 Deinococcus marmoris DSM 12784 GCA_000701405.1
98 Deinococcus swuensis GCA_000800395.1
JP
220 Bacillus anthracis str. Ames GCA_000007845.1
220 Bacillus anthracis str. Sterne GCA_000008165.1
220 Bacillus cereus ATCC 14579 GCA_000007825.1
220 Bacillus mycoides GCA_000832605.1
220 Bacillus pseudomycoides DSM 12442 GCA_000161455.1
220 Bacillus thuringiensis YBT-1518 GCA_000497525.2
220 _5BBacillus thuringiensis_5D serovar konkukian str. 97-27 GCA_000008505.1
199 Bacillus circulans NBRC 13626 GCA_001591585.1
199 Bacillus dakarensis GCA_900156875.1
199 Bacillus massiliosenegalensis JC6 GCA_000311725.1
199 Bacillus tuaregi GCA_900104575.1
189 Bacillus infantis NRRL B-14911 GCA_000473245.1
JQ
314 Bifidobacterium adolescentis ATCC 15703 GCA_000010425.1
295 Bifidobacterium dentium JCM 1195 = DSM 20436 GCA_001042595.1
268 Bifidobacterium tsurumiense GCA_000741765.1
JR
290 Deinococcus radiodurans R1 GCA_000008565.1
211 Deinococcus deserti VCD115 GCA_000020685.1
201 Deinococcus gobiensis I-0 GCA_000252445.1
JS
193 Lactobacillus gasseri ATCC 33323 = JCM 1131 GCA_000014425.1
193 Lactobacillus hominis DSM 23910 = CRBIP 24.179 GCA_000296835.1
147 Pediococcus argentinicus GCA_001437605.1
147 Pediococcus pentosaceus ATCC 25745 GCA_000014505.1
147 Pediococcus stilesii GCA_001437075.1
146 Lactobacillus fermentum IFO 3956 GCA_000010145.1
JT
239 Clostridium beijerinckii GCA_000833105.2
239 Clostridium saccharoperbutylacetonicum N1-4_28HMT_29 GCA_000340885.1
235 Clostridium puniceum GCA_002006345.1
226 Clostridium taeniosporum GCA_001735765.1
JU

170 Deinococcus radiodurans R1 GCA_000008565.1
112 Deinococcus deserti VCD115 GCA_000020685.1
102 Deinococcus gobiensis I-0 GCA_000252445.1
JV
200 Enterococcus faecalis V583 GCA_000007785.1
200 Streptomyces cinnamoneus GCA_001885705.1
169 Bavariicoccus seileri DSM 19936 GCA_000421665.1
169 Carnobacterium divergens DSM 20623 GCA_000744255.1
169 Enterococcus asini ATCC 700915 GCA_000407365.1
169 Enterococcus canis NBRC 100695 GCA_001544375.1
169 Enterococcus casseliflavus EC20 GCA_000157355.2
169 Enterococcus dispar ATCC 51266 GCA_000406945.1
169 Enterococcus faecium DO GCA_000174395.2
169 Enterococcus gilvus ATCC BAA-350 GCA_000407545.1
169 Enterococcus haemoperoxidus ATCC BAA-382 GCA_000407165.1
169 Enterococcus hermanniensis GCA_001885945.1
169 Enterococcus hirae ATCC 9790 GCA_000271405.2
169 Enterococcus italicus DSM 15952 GCA_000185365.1
169 Enterococcus malodoratus ATCC 43197 GCA_000407185.1
169 Enterococcus massiliensis GCA_001050095.1
169 Enterococcus mundtii QU 25 GCA_000504125.1
169 Enterococcus pallens ATCC BAA-351 GCA_000407485.1
169 Enterococcus phoeniculicola ATCC BAA-412 GCA_000407505.1
169 Enterococcus pseudoavium NBRC 100491 GCA_001544295.1
169 Enterococcus rivorum GCA_001742285.1
169 Enterococcus saccharolyticus subsp. saccharolyticus ATCC 43076 GCA_000407285.1
169 Enterococcus sulfureus ATCC 49903 GCA_000407605.1
169 Enterococcus thailandicus GCA_001652875.1
169 Granulicatella balaenopterae GCA_900111135.1
169 Isobaculum melis GCA_900111355.1
169 Vagococcus fluvialis bH819 GCA_900163795.1
169 Vagococcus lutrae LBD1 GCA_000498295.1
168 Lactobacillus buchneri CD034 GCA_000298115.2
168 Lactobacillus parafarraginis DSM 18390 = JCM 14109 GCA_001311355.1
168 Lactobacillus rapi DSM 19907 = JCM 15042 GCA_001436255.1
JW
264 Streptococcus mutans UA159 GCA_000007465.2
182 Streptococcus marimammalium DSM 18627 GCA_000380045.1
174 Streptococcus salivarius GCA_000785515.1
174 Streptococcus thermophilus JIM 8232 GCA_000253395.1
JX
262 Rhodobacter sphaeroides 2.4.1 GCA_000012905.2
204 Pseudorhodobacter psychrotolerans GCA_001294535.1
204 Rhodobacter sphaeroides ATCC 17025 GCA_000016405.1
195 Gemmobacter aquatilis GCA_900110025.1
JY
281 Bacillus anthracis str. Ames GCA_000007845.1
281 Bacillus anthracis str. Sterne GCA_000008165.1
281 Bacillus cereus ATCC 14579 GCA_000007825.1
281 Bacillus mycoides GCA_000832605.1
281 Bacillus pseudomycoides DSM 12442 GCA_000161455.1
281 Bacillus thuringiensis YBT-1518 GCA_000497525.2
281 _5BBacillus thuringiensis_5D serovar konkukian str. 97-27 GCA_000008505.1
209 Bacillus manliponensis GCA_000712595.1
204 Bacillus coahuilensis m4-4 GCA_000171615.1
204 Bacillus vietnamensis NBRC 101237 GCA_001591825.1
JZ
209 Deinococcus radiodurans R1 GCA_000008565.1
143 Deinococcus marmoris DSM 12784 GCA_000701405.1
143 Deinococcus swuensis GCA_000800395.1
132 Deinococcus deserti VCD115 GCA_000020685.1
K0
289 Deinococcus radiodurans R1 GCA_000008565.1
188 Deinococcus puniceus GCA_001644565.1
173 Deinococcus hopiensis KR-140 GCA_900176165.1
K1
267 Bifidobacterium adolescentis ATCC 15703 GCA_000010425.1
243 Bifidobacterium callitrichos DSM 23973 GCA_000741175.1
242 Bifidobacterium angulatum DSM 20098 = JCM 7096 GCA_001025155.1
K2
180 Streptococcus mutans UA159 GCA_000007465.2
144 Streptococcus salivarius GCA_000785515.1
144 Streptococcus thermophilus JIM 8232 GCA_000253395.1
143 Streptococcus equinus GCA_000964315.1
143 Streptococcus gallolyticus subsp. gallolyticus DSM 16831 GCA_002000985.1
143 Streptococcus orisratti DSM 15617 GCA_000380105.1

143 Streptococcus ratti FA-1 = DSM 20564 GCA_000286075.1
K3
233 Bacillus thuringiensis YBT-1518 GCA_000497525.2
217 Bacillus anthracis str. Ames GCA_000007845.1
217 Bacillus anthracis str. Sterne GCA_000008165.1
217 Bacillus cereus ATCC 14579 GCA_000007825.1
217 Bacillus mycoides GCA_000832605.1
217 Bacillus pseudomycoides DSM 12442 GCA_000161455.1
217 _5BBacillus thuringiensis_5D serovar konkukian str. 97-27 GCA_000008505.1
148 Bacillus coahuilensis m4-4 GCA_000171615.1
148 Bacillus eiseniae GCA_001636325.1
148 Bacillus marisflavi GCA_001274775.1
K4
294 Streptococcus mutans UA159 GCA_000007465.2
200 Streptococcus gordonii str. Challis substr. CH1 GCA_000017005.1
200 Streptococcus sobrinus DSM 20742 = ATCC 33478 GCA_000686605.1
197 Streptococcus ratti FA-1 = DSM 20564 GCA_000286075.1
K5
255 Deinococcus radiodurans R1 GCA_000008565.1
177 Deinococcus puniceus GCA_001644565.1
160 Deinococcus gobiensis I-0 GCA_000252445.1
K6
211 Streptococcus mutans UA159 GCA_000007465.2
124 Streptococcus macacae NCTC 11558 GCA_000187995.3
123 Streptococcus marimammalium DSM 18627 GCA_000380045.1
K7
253 Staphylococcus aureus subsp. aureus NCTC 8325 GCA_000013425.1
253 Staphylococcus condimenti GCA_001618885.1
253 Staphylococcus epidermidis ATCC 12228 GCA_000007645.1
253 Staphylococcus haemolyticus JCSC1435 GCA_000009865.1
253 Staphylococcus hominis subsp. hominis C80 GCA_000183685.1
253 Staphylococcus simiae CCM 7213 GCA_000235645.2
253 Staphylococcus simulans GCA_001559115.1
250 Megasphaera cerevisiae DSM 20462 GCA_001045675.1
250 Staphylococcus warneri SG1 GCA_000332735.1
248 Staphylococcus capitis subsp. capitis GCA_001028645.1
248 Staphylococcus lugdunensis HKU09-01 GCA_000025085.1
K8
254 Escherichia coli IAI39 GCA_000026345.1
254 Escherichia coli O104_3AH4 str. 2011C-3493 GCA_000299455.1
254 Escherichia coli O157_3AH7 str. Sakai GCA_000008865.1
254 Escherichia coli O83_3AH1 str. NRG 857C GCA_000183345.1
254 Escherichia coli UMN026 GCA_000026325.2
254 Escherichia coli str. K-12 substr. MG1655 GCA_000005845.2
254 Shigella flexneri 2a str. 301 GCA_000006925.2
254 Tumebacillus flagellatus GCA_000714935.1
245 Shigella dysenteriae Sd197 GCA_000012005.1
209 Erwinia iniecta GCA_001267535.1
209 Erwinia toletana DAPP-PG 735 GCA_000336255.1
K9
255 Deinococcus radiodurans R1 GCA_000008565.1
166 Deinococcus gobiensis I-0 GCA_000252445.1
161 Deinococcus marmoris DSM 12784 GCA_000701405.1
161 Deinococcus swuensis GCA_000800395.1
Ka
171 Clostridium beijerinckii GCA_000833105.2
171 Clostridium saccharoperbutylacetonicum N1-4_28HMT_29 GCA_000340885.1
167 Clostridium saccharobutylicum DSM 13864 GCA_000473995.1
164 Clostridium puniceum GCA_002006345.1
Kb
140 Rhodobacter sphaeroides 2.4.1 GCA_000012905.2
140 Rhodobacter sphaeroides ATCC 17025 GCA_000016405.1
132 Pseudorhodobacter ferrugineus DSM 5888 GCA_000420745.1
132 Pseudorhodobacter wandonensis GCA_001202035.1
112 Gemmobacter megaterium GCA_900156815.1
Kc
227 Escherichia coli IAI39 GCA_000026345.1
227 Escherichia coli O104_3AH4 str. 2011C-3493 GCA_000299455.1
227 Escherichia coli O157_3AH7 str. Sakai GCA_000008865.1
227 Escherichia coli UMN026 GCA_000026325.2
227 Escherichia coli str. K-12 substr. MG1655 GCA_000005845.2
227 Shigella flexneri 2a str. 301 GCA_000006925.2
227 Tumebacillus flagellatus GCA_000714935.1
226 Escherichia coli O83_3AH1 str. NRG 857C GCA_000183345.1
218 Shigella dysenteriae Sd197 GCA_000012005.1
Kd

332 Bacillus anthracis str. Ames GCA_000007845.1
332 Bacillus anthracis str. Sterne GCA_000008165.1
332 Bacillus cereus ATCC 14579 GCA_000007825.1
332 Bacillus mycoides GCA_000832605.1
332 Bacillus pseudomycoides DSM 12442 GCA_000161455.1
332 Bacillus thuringiensis YBT-1518 GCA_000497525.2
332 _5BBacillus thuringiensis_5D serovar konkukian str. 97-27 GCA_000008505.1
257 Bacillus manliponensis GCA_000712595.1
243 Bacillus aquimaris TF-12 GCA_001648555.1
243 Bacillus marisflavi GCA_001274775.1
Ke
277 Enterococcus casseliflavus EC20 GCA_000157355.2
277 Enterococcus saccharolyticus subsp. saccharolyticus ATCC 43076 GCA_000407285.1
276 Enterococcus faecalis V583 GCA_000007785.1
276 Streptomyces cinnamoneus GCA_001885705.1
262 Enterococcus canis NBRC 100695 GCA_001544375.1
262 Enterococcus dispar ATCC 51266 GCA_000406945.1
262 Enterococcus faecium DO GCA_000174395.2
262 Enterococcus hirae ATCC 9790 GCA_000271405.2
262 Enterococcus mundtii QU 25 GCA_000504125.1
262 Enterococcus rivorum GCA_001742285.1
Kf
236 Clostridium butyricum GCA_001456065.2
235 Clostridium beijerinckii GCA_000833105.2
235 Clostridium saccharoperbutylacetonicum N1-4_28HMT_29 GCA_000340885.1
231 Clostridium puniceum GCA_002006345.1
Kg
105 Deinococcus radiodurans R1 GCA_000008565.1
103 Deinococcus gobiensis I-0 GCA_000252445.1
90 Deinococcus puniceus GCA_001644565.1
Kh
345 Lactobacillus gasseri ATCC 33323 = JCM 1131 GCA_000014425.1
319 Lactobacillus hominis DSM 23910 = CRBIP 24.179 GCA_000296835.1
260 Lactobacillus crispatus ST1 GCA_000091765.1
Ki
242 Clostridium beijerinckii GCA_000833105.2
242 Clostridium saccharobutylicum DSM 13864 GCA_000473995.1
242 Clostridium saccharoperbutylacetonicum N1-4_28HMT_29 GCA_000340885.1
238 Clostridium puniceum GCA_002006345.1
223 Clostridium chromiireducens GCA_002029255.1
Kj
221 Deinococcus radiodurans R1 GCA_000008565.1
156 Deinococcus deserti VCD115 GCA_000020685.1
156 Deinococcus soli Cha et al. 2016 GCA_001007995.1
147 Deinococcus frigens DSM 12807 GCA_000701425.1
147 Deinococcus hopiensis KR-140 GCA_900176165.1
147 Deinococcus marmoris DSM 12784 GCA_000701405.1
147 Deinococcus puniceus GCA_001644565.1
147 Deinococcus swuensis GCA_000800395.1
Kk
243 Clostridium beijerinckii GCA_000833105.2
243 Clostridium saccharobutylicum DSM 13864 GCA_000473995.1
243 Clostridium saccharoperbutylacetonicum N1-4_28HMT_29 GCA_000340885.1
229 Clostridium puniceum GCA_002006345.1
221 Clostridium chromiireducens GCA_002029255.1
Kl
297 Streptococcus mutans UA159 GCA_000007465.2
186 Streptococcus gordonii str. Challis substr. CH1 GCA_000017005.1
182 Streptococcus cristatus AS 1.3089 GCA_000385925.1
182 Streptococcus mitis B6 GCA_000027165.1
182 Streptococcus pneumoniae R6 GCA_000007045.1
Km
207 Rhodobacter sphaeroides 2.4.1 GCA_000012905.2
178 Rhodobacter sphaeroides ATCC 17025 GCA_000016405.1
157 Gemmobacter aquatilis GCA_900110025.1
Kn
275 Staphylococcus epidermidis ATCC 12228 GCA_000007645.1
275 Staphylococcus haemolyticus JCSC1435 GCA_000009865.1
263 Staphylococcus capitis subsp. capitis GCA_001028645.1
259 Staphylococcus hominis subsp. hominis C80 GCA_000183685.1
259 Staphylococcus lugdunensis HKU09-01 GCA_000025085.1
Ko
149 Enterococcus faecalis V583 GCA_000007785.1
149 Streptomyces cinnamoneus GCA_001885705.1
139 Bavariicoccus seileri DSM 19936 GCA_000421665.1
139 Granulicatella balaenopterae GCA_900111135.1

139 Isobaculum melis GCA_900111355.1
135 Enterococcus asini ATCC 700915 GCA_000407365.1
135 Enterococcus canis NBRC 100695 GCA_001544375.1
135 Enterococcus casseliflavus EC20 GCA_000157355.2
135 Enterococcus dispar ATCC 51266 GCA_000406945.1
135 Enterococcus faecium DO GCA_000174395.2
135 Enterococcus gilvus ATCC BAA-350 GCA_000407545.1
135 Enterococcus haemoperoxidus ATCC BAA-382 GCA_000407165.1
135 Enterococcus hermanniensis GCA_001885945.1
135 Enterococcus hirae ATCC 9790 GCA_000271405.2
135 Enterococcus italicus DSM 15952 GCA_000185365.1
135 Enterococcus malodoratus ATCC 43197 GCA_000407185.1
135 Enterococcus massiliensis GCA_001050095.1
135 Enterococcus mundtii QU 25 GCA_000504125.1
135 Enterococcus pallens ATCC BAA-351 GCA_000407485.1
135 Enterococcus phoeniculicola ATCC BAA-412 GCA_000407505.1
135 Enterococcus pseudoavium NBRC 100491 GCA_001544295.1
135 Enterococcus rivorum GCA_001742285.1
135 Enterococcus saccharolyticus subsp. saccharolyticus ATCC 43076 GCA_000407285.1
135 Enterococcus sulfureus ATCC 49903 GCA_000407605.1
135 Enterococcus thailandicus GCA_001652875.1
135 Vagococcus fluvialis bH819 GCA_900163795.1
135 Vagococcus lutrae LBD1 GCA_000498295.1
135 Vagococcus penaei GCA_001998885.1
Kp
211 Lactobacillus gasseri ATCC 33323 = JCM 1131 GCA_000014425.1
211 Lactobacillus hominis DSM 23910 = CRBIP 24.179 GCA_000296835.1
178 Lactobacillus amylovorus GCA_000191545.1
178 Lactobacillus crispatus ST1 GCA_000091765.1
178 Lactobacillus kalixensis DSM 16043 GCA_001434335.1
176 Lactobacillus psittaci DSM 15354 GCA_000425905.1
Kq
165 Deinococcus radiodurans R1 GCA_000008565.1
103 Deinococcus marmoris DSM 12784 GCA_000701405.1
103 Deinococcus swuensis GCA_000800395.1
100 Deinococcus proteolyticus MRP GCA_000190555.1
Kr
290 Deinococcus radiodurans R1 GCA_000008565.1
192 Deinococcus deserti VCD115 GCA_000020685.1
179 Deinococcus gobiensis I-0 GCA_000252445.1
Ks
273 Bacillus thuringiensis YBT-1518 GCA_000497525.2
257 Bacillus anthracis str. Ames GCA_000007845.1
257 Bacillus anthracis str. Sterne GCA_000008165.1
257 Bacillus cereus ATCC 14579 GCA_000007825.1
257 Bacillus mycoides GCA_000832605.1
257 Bacillus pseudomycoides DSM 12442 GCA_000161455.1
257 _5BBacillus thuringiensis_5D serovar konkukian str. 97-27 GCA_000008505.1
231 Bacillus cytotoxicus NVH 391-98 GCA_000017425.1
231 Bacillus manliponensis GCA_000712595.1
Kt
319 Bacillus thuringiensis YBT-1518 GCA_000497525.2
313 Bacillus anthracis str. Ames GCA_000007845.1
313 Bacillus anthracis str. Sterne GCA_000008165.1
313 Bacillus cereus ATCC 14579 GCA_000007825.1
313 Bacillus mycoides GCA_000832605.1
313 Bacillus pseudomycoides DSM 12442 GCA_000161455.1
313 _5BBacillus thuringiensis_5D serovar konkukian str. 97-27 GCA_000008505.1
263 Bacillus manliponensis GCA_000712595.1
Ku
212 Streptococcus mutans UA159 GCA_000007465.2
134 Streptococcus equinus GCA_000964315.1
134 Streptococcus gallolyticus subsp. gallolyticus DSM 16831 GCA_002000985.1
130 Streptococcus cristatus AS 1.3089 GCA_000385925.1
130 Streptococcus gordonii str. Challis substr. CH1 GCA_000017005.1
130 Streptococcus marimammalium DSM 18627 GCA_000380045.1
130 Streptococcus marmotae GCA_001623565.1
130 Streptococcus mitis B6 GCA_000027165.1
130 Streptococcus parasanguinis ATCC 15912 GCA_000164675.2
130 Streptococcus pneumoniae R6 GCA_000007045.1
130 Streptococcus salivarius GCA_000785515.1
130 Streptococcus thermophilus JIM 8232 GCA_000253395.1
Kv
266 Bacillus thuringiensis YBT-1518 GCA_000497525.2
249 Bacillus anthracis str. Ames GCA_000007845.1
249 Bacillus anthracis str. Sterne GCA_000008165.1

249 Bacillus cereus ATCC 14579 GCA_000007825.1
249 Bacillus mycoides GCA_000832605.1
249 Bacillus pseudomycoides DSM 12442 GCA_000161455.1
249 _5BBacillus thuringiensis_5D serovar konkukian str. 97-27 GCA_000008505.1
199 Bacillus acidicola GCA_001636425.1
199 Bacillus shackletonii GCA_001420715.1
199 Bacillus sporothermodurans GCA_001587375.1
Kw
198 Deinococcus radiodurans R1 GCA_000008565.1
115 Deinococcus geothermalis DSM 11300 GCA_000196275.1
112 Deinococcus puniceus GCA_001644565.1
112 Deinococcus soli Cha et al. 2016 GCA_001007995.1
Kx
304 Staphylococcus epidermidis ATCC 12228 GCA_000007645.1
304 Staphylococcus haemolyticus JCSC1435 GCA_000009865.1
304 Staphylococcus hominis subsp. hominis C80 GCA_000183685.1
304 Staphylococcus lugdunensis HKU09-01 GCA_000025085.1
300 Staphylococcus arlettae CVD059 GCA_000295715.1
300 Staphylococcus cohnii subsp. cohnii GCA_000972575.1
300 Staphylococcus saprophyticus subsp. saprophyticus ATCC 15305 GCA_000010125.1
298 Staphylococcus capitis subsp. capitis GCA_001028645.1
Ky
126 Escherichia coli IAI39 GCA_000026345.1
126 Escherichia coli O104_3AH4 str. 2011C-3493 GCA_000299455.1
126 Escherichia coli O157_3AH7 str. Sakai GCA_000008865.1
126 Escherichia coli O83_3AH1 str. NRG 857C GCA_000183345.1
126 Escherichia coli UMN026 GCA_000026325.2
126 Escherichia coli str. K-12 substr. MG1655 GCA_000005845.2
126 Shigella dysenteriae Sd197 GCA_000012005.1
126 Shigella flexneri 2a str. 301 GCA_000006925.2
126 Tumebacillus flagellatus GCA_000714935.1
111 Atlantibacter hermannii NBRC 105704 GCA_000248015.2
111 Bacillus cecembensis GCA_001439635.1
111 Enterobacter hormaechei subsp. steigerwaltii GCA_001729725.1
111 Erwinia billingiae Eb661 GCA_000196615.1
111 Erwinia iniecta GCA_001267535.1
111 Erwinia toletana DAPP-PG 735 GCA_000336255.1
111 Klebsiella oxytoca GCA_001022195.1
111 Kosakonia cowanii GCA_001975225.1
111 Kosakonia sacchari SP1 GCA_000300455.4
111 Pantoea agglomerans GCA_001709315.1
111 Pantoea alhagi GCA_002101395.1
111 Pantoea ananatis LMG 20103 GCA_000025405.2
111 Pantoea rwandensis GCA_000759475.1
111 Pantoea septica GCA_002095575.1
111 Pluralibacter gergoviae GCA_000757785.1
111 Pseudescherichia vulneris NBRC 102420 GCA_000759795.1
111 Shimwellia blattae DSM 4481 = NBRC 105725 GCA_000262305.1
111 Xenorhabdus cabanillasii JM26 GCA_000531755.1
111 Xenorhabdus doucetiae GCA_000968195.1
111 Xenorhabdus hominickii GCA_001721185.1
111 Xenorhabdus innexi GCA_900155355.1
111 Xenorhabdus japonica GCA_900115195.1
111 Xenorhabdus koppenhoeferi GCA_900116635.1
111 Xenorhabdus nematophila AN6_2F1 GCA_000953355.1
107 Cronobacter sakazakii GCA_000982825.1
107 Enterobacter cloacae subsp. cloacae ATCC 13047 GCA_000025565.1
107 Enterobacter kobei GCA_900185885.1
107 Erwinia amylovora CFBP1430 GCA_000091565.1
107 Erwinia gerundensis GCA_001517405.1
107 Erwinia oleae GCA_000770305.1
107 Pantoea dispersa EGD-AAK13 GCA_000465555.2
107 Salmonella enterica subsp. enterica serovar Typhi str. CT18 GCA_000195995.1
107 Salmonella enterica subsp. enterica serovar Typhimurium str. LT2 GCA_000006945.2
107 Streptomyces gilvigriseus GCA_001879105.1
107 Trabulsiella odontotermitis GCA_001297765.1
107 Xenorhabdus bovienii SS-2004 GCA_000027225.1
Kz
259 Bacillus anthracis str. Ames GCA_000007845.1
259 Bacillus anthracis str. Sterne GCA_000008165.1
259 Bacillus cereus ATCC 14579 GCA_000007825.1
259 Bacillus mycoides GCA_000832605.1
259 Bacillus pseudomycoides DSM 12442 GCA_000161455.1
259 Bacillus thuringiensis YBT-1518 GCA_000497525.2
259 _5BBacillus thuringiensis_5D serovar konkukian str. 97-27 GCA_000008505.1
192 Bacillus manliponensis GCA_000712595.1

187 Bacillus cytotoxicus NVH 391-98 GCA_000017425.1
KA
229 Deinococcus radiodurans R1 GCA_000008565.1
149 Deinococcus geothermalis DSM 11300 GCA_000196275.1
148 Deinococcus puniceus GCA_001644565.1
KB
169 Deinococcus radiodurans R1 GCA_000008565.1
87 Deinococcus gobiensis I-0 GCA_000252445.1
84 Deinococcus marmoris DSM 12784 GCA_000701405.1
84 Deinococcus swuensis GCA_000800395.1
KC
291 Clostridium beijerinckii GCA_000833105.2
291 Clostridium puniceum GCA_002006345.1
291 Clostridium saccharoperbutylacetonicum N1-4_28HMT_29 GCA_000340885.1
283 Clostridium saccharobutylicum DSM 13864 GCA_000473995.1
279 Clostridium chromiireducens GCA_002029255.1
KD
218 Megasphaera cerevisiae DSM 20462 GCA_001045675.1
218 Staphylococcus aureus subsp. aureus NCTC 8325 GCA_000013425.1
218 Staphylococcus condimenti GCA_001618885.1
218 Staphylococcus epidermidis ATCC 12228 GCA_000007645.1
218 Staphylococcus haemolyticus JCSC1435 GCA_000009865.1
218 Staphylococcus hominis subsp. hominis C80 GCA_000183685.1
218 Staphylococcus lugdunensis HKU09-01 GCA_000025085.1
218 Staphylococcus simiae CCM 7213 GCA_000235645.2
218 Staphylococcus simulans GCA_001559115.1
218 Staphylococcus warneri SG1 GCA_000332735.1
213 Staphylococcus hyicus GCA_000816085.1
212 Staphylococcus capitis subsp. capitis GCA_001028645.1
KE
174 Clostridium beijerinckii GCA_000833105.2
174 Clostridium puniceum GCA_002006345.1
174 Clostridium saccharobutylicum DSM 13864 GCA_000473995.1
174 Clostridium saccharoperbutylacetonicum N1-4_28HMT_29 GCA_000340885.1
159 Clostridium chromiireducens GCA_002029255.1
149 Clostridium amylolyticum GCA_900142075.1
149 Clostridium intestinale URNW GCA_000469625.2
149 Clostridium polynesiense GCA_000820705.1
KF
264 Enterococcus faecalis V583 GCA_000007785.1
264 Streptomyces cinnamoneus GCA_001885705.1
232 Enterococcus asini ATCC 700915 GCA_000407365.1
232 Enterococcus canis NBRC 100695 GCA_001544375.1
232 Enterococcus dispar ATCC 51266 GCA_000406945.1
232 Enterococcus faecium DO GCA_000174395.2
232 Enterococcus hirae ATCC 9790 GCA_000271405.2
232 Enterococcus mundtii QU 25 GCA_000504125.1
232 Enterococcus rivorum GCA_001742285.1
227 Enterococcus haemoperoxidus ATCC BAA-382 GCA_000407165.1
227 Enterococcus phoeniculicola ATCC BAA-412 GCA_000407505.1
227 Enterococcus thailandicus GCA_001652875.1
KG
226 Deinococcus radiodurans R1 GCA_000008565.1
140 Deinococcus gobiensis I-0 GCA_000252445.1
129 Deinococcus puniceus GCA_001644565.1
KH
181 Deinococcus radiodurans R1 GCA_000008565.1
101 Deinococcus puniceus GCA_001644565.1
94 Deinococcus hopiensis KR-140 GCA_900176165.1
KI
107 Streptococcus mutans UA159 GCA_000007465.2
83 Streptococcus massiliensis DSM 18628 GCA_000380065.1
79 Streptococcus gordonii str. Challis substr. CH1 GCA_000017005.1
79 Streptococcus phocae subsp. salmonis GCA_000772915.1
KJ
209 Bacillus anthracis str. Ames GCA_000007845.1
209 Bacillus anthracis str. Sterne GCA_000008165.1
209 Bacillus cereus ATCC 14579 GCA_000007825.1
209 Bacillus mycoides GCA_000832605.1
209 Bacillus pseudomycoides DSM 12442 GCA_000161455.1
209 Bacillus thuringiensis YBT-1518 GCA_000497525.2
209 _5BBacillus thuringiensis_5D serovar konkukian str. 97-27 GCA_000008505.1
205 Viridibacillus arvi GCA_001274945.1
194 Bacillus aquimaris TF-12 GCA_001648555.1
194 Bacillus vietnamensis NBRC 101237 GCA_001591825.1
KK

212 Rhodobacter sphaeroides 2.4.1 GCA_000012905.2
201 Rhodobacter sphaeroides ATCC 17025 GCA_000016405.1
171 Defluviimonas alba GCA_001620265.1
171 Gemmobacter megaterium GCA_900156815.1
171 Pseudorhodobacter ferrugineus DSM 5888 GCA_000420745.1
171 Pseudorhodobacter wandonensis GCA_001202035.1
KL
213 Clostridium beijerinckii GCA_000833105.2
213 Clostridium puniceum GCA_002006345.1
213 Clostridium saccharobutylicum DSM 13864 GCA_000473995.1
213 Clostridium saccharoperbutylacetonicum N1-4_28HMT_29 GCA_000340885.1
205 Clostridium chromiireducens GCA_002029255.1
199 Clostridium butyricum GCA_001456065.2
KM
171 Deinococcus radiodurans R1 GCA_000008565.1
97 Deinococcus marmoris DSM 12784 GCA_000701405.1
97 Deinococcus swuensis GCA_000800395.1
83 Deinococcus misasensis DSM 22328 GCA_000745915.1
KN
135 Lactobacillus gasseri ATCC 33323 = JCM 1131 GCA_000014425.1
116 Lactobacillus acetotolerans GCA_001042405.1
116 Lactobacillus amylovorus GCA_000191545.1
116 Lactobacillus crispatus ST1 GCA_000091765.1
116 Lactobacillus delbrueckii subsp. bulgaricus ATCC 11842 = JCM 1002 GCA_000056065.1
116 Lactobacillus helveticus GCA_001308285.1
116 Lactobacillus hominis DSM 23910 = CRBIP 24.179 GCA_000296835.1
116 Lactobacillus kalixensis DSM 16043 GCA_001434335.1
116 Lactobacillus kefiranofaciens ZW3 GCA_000214785.1
116 Lactobacillus psittaci DSM 15354 GCA_000425905.1
107 Lactobacillus acidophilus NCFM GCA_000011985.1
107 Lactobacillus amylophilus DSM 20533 = JCM 1125 GCA_001936335.1
107 Lactobacillus gigeriorum DSM 23908 = CRBIP 24.85 GCA_001436575.1
107 Lactobacillus jensenii GCA_001936235.1
107 Lactobacillus pasteurii DSM 23907 = CRBIP 24.76 GCA_000297025.1
107 Lactobacillus saniviri JCM 17471 = DSM 24301 GCA_001311785.1
KO
125 Rhodobacter sphaeroides 2.4.1 GCA_000012905.2
113 Pseudorhodobacter ferrugineus DSM 5888 GCA_000420745.1
113 Pseudorhodobacter wandonensis GCA_001202035.1
110 Rhodobacter sphaeroides ATCC 17025 GCA_000016405.1
KP
232 Deinococcus radiodurans R1 GCA_000008565.1
158 Deinococcus gobiensis I-0 GCA_000252445.1
157 Deinococcus proteolyticus MRP GCA_000190555.1
KQ
164 Clostridium beijerinckii GCA_000833105.2
164 Clostridium puniceum GCA_002006345.1
164 Clostridium saccharoperbutylacetonicum N1-4_28HMT_29 GCA_000340885.1
148 Clostridium saccharobutylicum DSM 13864 GCA_000473995.1
131 Clostridium taeniosporum GCA_001735765.1
KR
200 Clostridium beijerinckii GCA_000833105.2
200 Clostridium puniceum GCA_002006345.1
200 Clostridium saccharoperbutylacetonicum N1-4_28HMT_29 GCA_000340885.1
192 Clostridium saccharobutylicum DSM 13864 GCA_000473995.1
185 Clostridium butyricum GCA_001456065.2
KS
208 Escherichia coli IAI39 GCA_000026345.1
208 Escherichia coli O104_3AH4 str. 2011C-3493 GCA_000299455.1
208 Escherichia coli O157_3AH7 str. Sakai GCA_000008865.1
208 Escherichia coli O83_3AH1 str. NRG 857C GCA_000183345.1
208 Escherichia coli UMN026 GCA_000026325.2
208 Escherichia coli str. K-12 substr. MG1655 GCA_000005845.2
208 Shigella dysenteriae Sd197 GCA_000012005.1
208 Shigella flexneri 2a str. 301 GCA_000006925.2
208 Tumebacillus flagellatus GCA_000714935.1
196 Erwinia iniecta GCA_001267535.1
170 Enterobacter hormaechei subsp. steigerwaltii GCA_001729725.1
170 Erwinia toletana DAPP-PG 735 GCA_000336255.1
170 Klebsiella oxytoca GCA_001022195.1
170 Kosakonia cowanii GCA_001975225.1
170 Kosakonia sacchari SP1 GCA_000300455.4
170 Pluralibacter gergoviae GCA_000757785.1
170 Pseudescherichia vulneris NBRC 102420 GCA_000759795.1
KT
253 Deinococcus radiodurans R1 GCA_000008565.1

136 Deinococcus proteolyticus MRP GCA_000190555.1
129 Deinococcus puniceus GCA_001644565.1
KU
146 Clostridium beijerinckii GCA_000833105.2
146 Clostridium saccharobutylicum DSM 13864 GCA_000473995.1
146 Clostridium saccharoperbutylacetonicum N1-4_28HMT_29 GCA_000340885.1
137 Clostridium amylolyticum GCA_900142075.1
137 Clostridium intestinale URNW GCA_000469625.2
137 Clostridium polynesiense GCA_000820705.1
133 Clostridium algidicarnis GCA_000703125.1
KV
129 Staphylococcus simulans GCA_001559115.1
128 Bacillus thuringiensis YBT-1518 GCA_000497525.2
120 Staphylococcus pettenkoferi GCA_002208805.1
KW
121 Escherichia coli IAI39 GCA_000026345.1
121 Escherichia coli O104_3AH4 str. 2011C-3493 GCA_000299455.1
121 Escherichia coli O157_3AH7 str. Sakai GCA_000008865.1
121 Escherichia coli O83_3AH1 str. NRG 857C GCA_000183345.1
121 Escherichia coli UMN026 GCA_000026325.2
121 Escherichia coli str. K-12 substr. MG1655 GCA_000005845.2
121 Shigella dysenteriae Sd197 GCA_000012005.1
121 Shigella flexneri 2a str. 301 GCA_000006925.2
121 Tumebacillus flagellatus GCA_000714935.1
106 Buttiauxella ferragutiae ATCC 51602 GCA_001654915.1
106 Cedecea neteri GCA_000757825.1
106 Citrobacter freundii CFNIH1 GCA_000648515.1
106 Cronobacter sakazakii GCA_000982825.1
106 Edwardsiella anguillarum ET080813 GCA_000264765.2
106 Enterobacter cloacae subsp. cloacae ATCC 13047 GCA_000025565.1
106 Enterobacter hormaechei subsp. steigerwaltii GCA_001729725.1
106 Enterobacter kobei GCA_900185885.1
106 Erwinia amylovora CFBP1430 GCA_000091565.1
106 Erwinia billingiae Eb661 GCA_000196615.1
106 Erwinia gerundensis GCA_001517405.1
106 Erwinia iniecta GCA_001267535.1
106 Erwinia persicina NBRC 102418 GCA_001571305.1
106 Erwinia toletana DAPP-PG 735 GCA_000336255.1
106 Klebsiella aerogenes KCTC 2190 GCA_000215745.1
106 Klebsiella oxytoca GCA_001022195.1
106 Kluyvera ascorbata ATCC 33433 GCA_000735365.1
106 Kosakonia cowanii GCA_001975225.1
106 Kosakonia sacchari SP1 GCA_000300455.4
106 Mangrovibacter phragmitis GCA_001655675.1
106 Nissabacter archeti GCA_900130115.1
106 Obesumbacterium proteus GCA_001586165.1
106 Pantoea agglomerans GCA_001709315.1
106 Pantoea alhagi GCA_002101395.1
106 Pantoea ananatis LMG 20103 GCA_000025405.2
106 Pantoea dispersa EGD-AAK13 GCA_000465555.2
106 Pantoea rwandensis GCA_000759475.1
106 Pantoea septica GCA_002095575.1
106 Pluralibacter gergoviae GCA_000757785.1
106 Pseudescherichia vulneris NBRC 102420 GCA_000759795.1
106 Rosenbergiella nectarea GCA_900111105.1
106 Salmonella enterica subsp. enterica serovar Typhi str. CT18 GCA_000195995.1
106 Shimwellia blattae DSM 4481 = NBRC 105725 GCA_000262305.1
106 Tatumella citrea GCA_002163585.1
106 Tatumella saanichensis GCA_000439375.1
106 Trabulsiella odontotermitis GCA_001297765.1
106 Yokenella regensburgei ATCC 49455 GCA_000735455.1
106 _5BEnterobacter_5D lignolyticus SCF1 GCA_000164865.1
105 Izhakiella capsodis GCA_900115045.1
KX
185 Rhodobacter sphaeroides 2.4.1 GCA_000012905.2
159 Rhodobacter sphaeroides ATCC 17025 GCA_000016405.1
142 Pseudorhodobacter aquimaris GCA_001202025.1
142 Pseudorhodobacter ferrugineus DSM 5888 GCA_000420745.1
142 Pseudorhodobacter wandonensis GCA_001202035.1
KY
245 Streptococcus mutans UA159 GCA_000007465.2
162 Streptococcus equinus GCA_000964315.1
162 Streptococcus gallolyticus subsp. gallolyticus DSM 16831 GCA_002000985.1
159 Streptococcus macacae NCTC 11558 GCA_000187995.3
KZ
253 Bifidobacterium adolescentis ATCC 15703 GCA_000010425.1

231 Bifidobacterium choerinum GCA_000741135.1
231 Bifidobacterium pseudolongum PV8-2 GCA_000800475.2
224 Bifidobacterium breve DSM 20213 = JCM 1192 GCA_001025175.1
L0
212 Lactobacillus gasseri ATCC 33323 = JCM 1131 GCA_000014425.1
181 Lactobacillus hominis DSM 23910 = CRBIP 24.179 GCA_000296835.1
119 Lactobacillus crispatus ST1 GCA_000091765.1
119 Lactobacillus kefiranofaciens ZW3 GCA_000214785.1
L1
279 Clostridium beijerinckii GCA_000833105.2
279 Clostridium puniceum GCA_002006345.1
279 Clostridium saccharoperbutylacetonicum N1-4_28HMT_29 GCA_000340885.1
270 Clostridium saccharobutylicum DSM 13864 GCA_000473995.1
260 Clostridium butyricum GCA_001456065.2
L2
224 Lactobacillus gasseri ATCC 33323 = JCM 1131 GCA_000014425.1
224 Lactobacillus hominis DSM 23910 = CRBIP 24.179 GCA_000296835.1
158 Lactobacillus amylophilus DSM 20533 = JCM 1125 GCA_001936335.1
148 Lactobacillus antri DSM 16041 GCA_000160835.1
148 Lactobacillus frumenti DSM 13145 GCA_001436045.1
148 Lactobacillus iners DSM 13335 GCA_000160875.1
148 Lactobacillus mucosae LM1 GCA_000248095.3
148 Lactobacillus pontis DSM 8475 GCA_001435345.1
148 Lactobacillus reuteri DSM 20016 GCA_000016825.1
148 Lactobacillus secaliphilus GCA_001437055.1
148 Lactobacillus vaginalis DSM 5837 = ATCC 49540 GCA_000159435.1
L3
191 Clostridium saccharobutylicum DSM 13864 GCA_000473995.1
189 Clostridium beijerinckii GCA_000833105.2
189 Clostridium puniceum GCA_002006345.1
189 Clostridium saccharoperbutylacetonicum N1-4_28HMT_29 GCA_000340885.1
187 Clostridium chromiireducens GCA_002029255.1
L4
204 Rhodobacter sphaeroides 2.4.1 GCA_000012905.2
178 Rhodobacter sphaeroides ATCC 17025 GCA_000016405.1
153 Rhodobacter capsulatus SB 1003 GCA_000021865.1
L5
142 Clostridium beijerinckii GCA_000833105.2
142 Clostridium chromiireducens GCA_002029255.1
142 Clostridium puniceum GCA_002006345.1
142 Clostridium saccharobutylicum DSM 13864 GCA_000473995.1
142 Clostridium saccharoperbutylacetonicum N1-4_28HMT_29 GCA_000340885.1
141 Clostridium neonatale GCA_001458595.1
138 Clostridium butyricum GCA_001456065.2
L6
266 Clostridium saccharobutylicum DSM 13864 GCA_000473995.1
264 Clostridium beijerinckii GCA_000833105.2
264 Clostridium puniceum GCA_002006345.1
264 Clostridium saccharoperbutylacetonicum N1-4_28HMT_29 GCA_000340885.1
254 Clostridium chromiireducens GCA_002029255.1
L7
127 Clostridium beijerinckii GCA_000833105.2
127 Clostridium puniceum GCA_002006345.1
127 Clostridium saccharobutylicum DSM 13864 GCA_000473995.1
127 Clostridium saccharoperbutylacetonicum N1-4_28HMT_29 GCA_000340885.1
117 Clostridium neonatale GCA_001458595.1
113 Clostridium butyricum GCA_001456065.2
113 Clostridium chromiireducens GCA_002029255.1
L8
181 Escherichia coli IAI39 GCA_000026345.1
181 Escherichia coli O104_3AH4 str. 2011C-3493 GCA_000299455.1
181 Escherichia coli O157_3AH7 str. Sakai GCA_000008865.1
181 Escherichia coli O83_3AH1 str. NRG 857C GCA_000183345.1
181 Escherichia coli UMN026 GCA_000026325.2
181 Escherichia coli str. K-12 substr. MG1655 GCA_000005845.2
181 Shigella dysenteriae Sd197 GCA_000012005.1
181 Shigella flexneri 2a str. 301 GCA_000006925.2
181 Tumebacillus flagellatus GCA_000714935.1
144 Buttiauxella ferragutiae ATCC 51602 GCA_001654915.1
144 Cedecea neteri GCA_000757825.1
144 Citrobacter freundii CFNIH1 GCA_000648515.1
144 Klebsiella aerogenes KCTC 2190 GCA_000215745.1
144 Kluyvera ascorbata ATCC 33433 GCA_000735365.1
144 Nissabacter archeti GCA_900130115.1
144 Obesumbacterium proteus GCA_001586165.1
144 Tatumella saanichensis GCA_000439375.1

144 Yokenella regensburgei ATCC 49455 GCA_000735455.1
144 _5BEnterobacter_5D lignolyticus SCF1 GCA_000164865.1
134 Aeromonas eucrenophila GCA_000819865.1
134 Aeromonas fluvialis GCA_000819885.1
134 Aeromonas hydrophila subsp. hydrophila ATCC 7966 GCA_000014805.1
134 Aeromonas salmonicida subsp. salmonicida A449 GCA_000196395.1
134 Aeromonas schubertii GCA_001447335.1
134 Aeromonas simiae GCA_000820125.1
134 Serratia symbiotica GCA_000821185.1
134 Shewanella algae GCA_000947195.1
L9
189 Rhodobacter sphaeroides 2.4.1 GCA_000012905.2
189 Rhodobacter sphaeroides ATCC 17025 GCA_000016405.1
159 Rhodobacter capsulatus SB 1003 GCA_000021865.1
154 Pseudorhodobacter psychrotolerans GCA_001294535.1
La
211 Deinococcus radiodurans R1 GCA_000008565.1
151 Deinococcus gobiensis I-0 GCA_000252445.1
145 Deinococcus puniceus GCA_001644565.1
Lb
205 Bifidobacterium adolescentis ATCC 15703 GCA_000010425.1
189 Bifidobacterium breve DSM 20213 = JCM 1192 GCA_001025175.1
185 Bifidobacterium callitrichos DSM 23973 GCA_000741175.1
Lc
197 Clostridium beijerinckii GCA_000833105.2
197 Clostridium puniceum GCA_002006345.1
197 Clostridium saccharobutylicum DSM 13864 GCA_000473995.1
197 Clostridium saccharoperbutylacetonicum N1-4_28HMT_29 GCA_000340885.1
186 Clostridium chromiireducens GCA_002029255.1
177 Clostridium butyricum GCA_001456065.2
Ld
186 Bifidobacterium angulatum DSM 20098 = JCM 7096 GCA_001025155.1
184 Bifidobacterium adolescentis ATCC 15703 GCA_000010425.1
184 Bifidobacterium gallicum DSM 20093 = LMG 11596 GCA_000741205.1
181 Bifidobacterium longum NCC2705 GCA_000007525.1
Le
218 Clostridium beijerinckii GCA_000833105.2
218 Clostridium puniceum GCA_002006345.1
218 Clostridium saccharoperbutylacetonicum N1-4_28HMT_29 GCA_000340885.1
209 Clostridium saccharobutylicum DSM 13864 GCA_000473995.1
203 Clostridium butyricum GCA_001456065.2
Lf
271 Bifidobacterium adolescentis ATCC 15703 GCA_000010425.1
221 Bifidobacterium asteroides PRL2011 GCA_000304215.1
213 Bifidobacterium callitrichos DSM 23973 GCA_000741175.1
213 Bifidobacterium thermophilum GCA_000741495.1
Lg
164 Streptococcus mutans UA159 GCA_000007465.2
132 Streptococcus iniae GCA_000831485.1
123 Streptococcus sobrinus DSM 20742 = ATCC 33478 GCA_000686605.1
Lh
203 Escherichia coli IAI39 GCA_000026345.1
203 Escherichia coli O104_3AH4 str. 2011C-3493 GCA_000299455.1
203 Escherichia coli O157_3AH7 str. Sakai GCA_000008865.1
203 Escherichia coli UMN026 GCA_000026325.2
203 Escherichia coli str. K-12 substr. MG1655 GCA_000005845.2
203 Shigella dysenteriae Sd197 GCA_000012005.1
203 Shigella flexneri 2a str. 301 GCA_000006925.2
203 Tumebacillus flagellatus GCA_000714935.1
202 Escherichia coli O83_3AH1 str. NRG 857C GCA_000183345.1
189 Kosakonia cowanii GCA_001975225.1
Li
204 Enterococcus dispar ATCC 51266 GCA_000406945.1
202 Enterococcus canis NBRC 100695 GCA_001544375.1
202 Enterococcus faecalis V583 GCA_000007785.1
202 Enterococcus faecium DO GCA_000174395.2
202 Enterococcus hirae ATCC 9790 GCA_000271405.2
202 Enterococcus mundtii QU 25 GCA_000504125.1
202 Enterococcus rivorum GCA_001742285.1
202 Streptomyces cinnamoneus GCA_001885705.1
201 Enterococcus casseliflavus EC20 GCA_000157355.2
201 Enterococcus saccharolyticus subsp. saccharolyticus ATCC 43076 GCA_000407285.1
Lj
214 Megasphaera cerevisiae DSM 20462 GCA_001045675.1
214 Staphylococcus aureus subsp. aureus NCTC 8325 GCA_000013425.1
214 Staphylococcus epidermidis ATCC 12228 GCA_000007645.1

214 Staphylococcus haemolyticus JCSC1435 GCA_000009865.1
214 Staphylococcus hominis subsp. hominis C80 GCA_000183685.1
214 Staphylococcus simiae CCM 7213 GCA_000235645.2
214 Staphylococcus warneri SG1 GCA_000332735.1
209 Staphylococcus capitis subsp. capitis GCA_001028645.1
209 Staphylococcus lugdunensis HKU09-01 GCA_000025085.1
199 Staphylococcus cohnii subsp. cohnii GCA_000972575.1
Lk
167 Deinococcus radiodurans R1 GCA_000008565.1
113 Deinococcus geothermalis DSM 11300 GCA_000196275.1
110 Deinococcus proteolyticus MRP GCA_000190555.1
Ll
317 Staphylococcus capitis subsp. capitis GCA_001028645.1
317 Staphylococcus epidermidis ATCC 12228 GCA_000007645.1
313 Staphylococcus lugdunensis HKU09-01 GCA_000025085.1
306 Staphylococcus haemolyticus JCSC1435 GCA_000009865.1
Lm
180 Clostridium beijerinckii GCA_000833105.2
180 Clostridium puniceum GCA_002006345.1
180 Clostridium saccharoperbutylacetonicum N1-4_28HMT_29 GCA_000340885.1
169 Clostridium butyricum GCA_001456065.2
169 Clostridium taeniosporum GCA_001735765.1
164 Clostridium saccharobutylicum DSM 13864 GCA_000473995.1
Ln
200 Streptococcus mutans UA159 GCA_000007465.2
118 Bacillus bataviensis LMG 21833 GCA_000307875.1
118 Lactococcus lactis subsp. lactis Il1403 GCA_000006865.1
118 Streptococcus gordonii str. Challis substr. CH1 GCA_000017005.1
118 Streptococcus phocae subsp. salmonis GCA_000772915.1
117 Streptococcus equinus GCA_000964315.1
117 Streptococcus gallolyticus subsp. gallolyticus DSM 16831 GCA_002000985.1
Lo
188 Clostridium beijerinckii GCA_000833105.2
188 Clostridium puniceum GCA_002006345.1
188 Clostridium saccharoperbutylacetonicum N1-4_28HMT_29 GCA_000340885.1
179 Clostridium saccharobutylicum DSM 13864 GCA_000473995.1
174 Clostridium butyricum GCA_001456065.2
Lp
264 Staphylococcus epidermidis ATCC 12228 GCA_000007645.1
264 Staphylococcus lugdunensis HKU09-01 GCA_000025085.1
257 Megasphaera cerevisiae DSM 20462 GCA_001045675.1
257 Staphylococcus warneri SG1 GCA_000332735.1
249 Staphylococcus aureus subsp. aureus NCTC 8325 GCA_000013425.1
249 Staphylococcus haemolyticus JCSC1435 GCA_000009865.1
249 Staphylococcus hominis subsp. hominis C80 GCA_000183685.1
249 Staphylococcus simiae CCM 7213 GCA_000235645.2
Lq
270 Rhodobacter sphaeroides 2.4.1 GCA_000012905.2
209 Rhodobacter sphaeroides ATCC 17025 GCA_000016405.1
207 Pseudorhodobacter ferrugineus DSM 5888 GCA_000420745.1
207 Pseudorhodobacter wandonensis GCA_001202035.1
Lr
99 Streptococcus mutans UA159 GCA_000007465.2
92 Rhodobacter sphaeroides 2.4.1 GCA_000012905.2
84 Streptococcus equinus GCA_000964315.1
84 Streptococcus gallolyticus subsp. gallolyticus DSM 16831 GCA_002000985.1
84 Streptococcus orisratti DSM 15617 GCA_000380105.1
84 Streptococcus ratti FA-1 = DSM 20564 GCA_000286075.1
Ls
228 Clostridium beijerinckii GCA_000833105.2
228 Clostridium saccharobutylicum DSM 13864 GCA_000473995.1
228 Clostridium saccharoperbutylacetonicum N1-4_28HMT_29 GCA_000340885.1
224 Clostridium puniceum GCA_002006345.1
212 Clostridium chromiireducens GCA_002029255.1
Lt
137 Gemmobacter megaterium GCA_900156815.1
137 Pseudorhodobacter psychrotolerans GCA_001294535.1
134 Gemmobacter aquatilis GCA_900110025.1
134 Rhodobacter sphaeroides 2.4.1 GCA_000012905.2
134 Rhodobacter sphaeroides ATCC 17025 GCA_000016405.1
121 Pseudorhodobacter ferrugineus DSM 5888 GCA_000420745.1
121 Pseudorhodobacter wandonensis GCA_001202035.1
Lu
222 Deinococcus radiodurans R1 GCA_000008565.1
145 Deinococcus deserti VCD115 GCA_000020685.1
145 Deinococcus hopiensis KR-140 GCA_900176165.1

143 Deinococcus puniceus GCA_001644565.1
 Lv
197 Rhodobacter sphaeroides 2.4.1 GCA_000012905.2
168 Rhodobacter capsulatus SB 1003 GCA_000021865.1
168 Rhodobacter sphaeroides ATCC 17025 GCA_000016405.1
164 Gemmobacter megaterium GCA_900156815.1
 Lw
233 Clostridium saccharobutylicum DSM 13864 GCA_000473995.1
232 Clostridium butyricum GCA_001456065.2
221 Clostridium beijerinckii GCA_000833105.2
221 Clostridium puniceum GCA_002006345.1
221 Clostridium saccharoperbutylacetonicum N1-4_28HMT_29 GCA_000340885.1
 Lx
249 Bacillus anthracis str. Ames GCA_000007845.1
249 Bacillus anthracis str. Sterne GCA_000008165.1
249 Bacillus cereus ATCC 14579 GCA_000007825.1
249 Bacillus mycoides GCA_000832605.1
249 Bacillus pseudomycoides DSM 12442 GCA_000161455.1
249 Bacillus thuringiensis YBT-1518 GCA_000497525.2
249 _5BBacillus thuringiensis_5D serovar konkukian str. 97-27 GCA_000008505.1
197 Bacillus manliponensis GCA_000712595.1
185 Bacillus aquimaris TF-12 GCA_001648555.1
 Ly
149 Deinococcus radiodurans R1 GCA_000008565.1
103 Deinococcus gobiensis I-0 GCA_000252445.1
88 Deinococcus deserti VCD115 GCA_000020685.1
 Lz
200 Clostridium beijerinckii GCA_000833105.2
200 Clostridium saccharobutylicum DSM 13864 GCA_000473995.1
200 Clostridium saccharoperbutylacetonicum N1-4_28HMT_29 GCA_000340885.1
193 Clostridium puniceum GCA_002006345.1
182 Clostridium chromiireducens GCA_002029255.1
 LA
175 Enterococcus faecalis V583 GCA_000007785.1
175 Streptomyces cinnamoneus GCA_001885705.1
160 Enterococcus canis NBRC 100695 GCA_001544375.1
160 Enterococcus faecium DO GCA_000174395.2
160 Enterococcus hirae ATCC 9790 GCA_000271405.2
160 Enterococcus mundtii QU 25 GCA_000504125.1
160 Enterococcus rivorum GCA_001742285.1
159 Enterococcus casseliflavus EC20 GCA_000157355.2
159 Enterococcus dispar ATCC 51266 GCA_000406945.1
159 Enterococcus gilvus ATCC BAA-350 GCA_000407545.1
159 Enterococcus hermanniensis GCA_001885945.1
159 Enterococcus malodoratus ATCC 43197 GCA_000407185.1
159 Enterococcus massiliensis GCA_001050095.1
159 Enterococcus pallens ATCC BAA-351 GCA_000407485.1
159 Enterococcus pseudoavium NBRC 100491 GCA_001544295.1
159 Enterococcus saccharolyticus subsp. saccharolyticus ATCC 43076 GCA_000407285.1
 LB
168 Deinococcus radiodurans R1 GCA_000008565.1
97 Deinococcus frigens DSM 12807 GCA_000701425.1
97 Deinococcus marmoris DSM 12784 GCA_000701405.1
97 Deinococcus swuensis GCA_000800395.1
96 Deinococcus gobiensis I-0 GCA_000252445.1
 LC
312 Deinococcus radiodurans R1 GCA_000008565.1
215 Deinococcus gobiensis I-0 GCA_000252445.1
187 Deinococcus puniceus GCA_001644565.1
 LD
219 Enterococcus faecalis V583 GCA_000007785.1
219 Streptomyces cinnamoneus GCA_001885705.1
206 Enterococcus faecium DO GCA_000174395.2
201 Enterococcus canis NBRC 100695 GCA_001544375.1
201 Enterococcus haemoperoxidus ATCC BAA-382 GCA_000407165.1
201 Enterococcus hirae ATCC 9790 GCA_000271405.2
201 Enterococcus mundtii QU 25 GCA_000504125.1
201 Enterococcus phoeniculicola ATCC BAA-412 GCA_000407505.1
201 Enterococcus rivorum GCA_001742285.1
201 Enterococcus thailandicus GCA_001652875.1
 LE
70 Escherichia coli IAI39 GCA_000026345.1
70 Escherichia coli O104_3AH4 str. 2011C-3493 GCA_000299455.1
70 Escherichia coli O157_3AH7 str. Sakai GCA_000008865.1
70 Escherichia coli O83_3AH1 str. NRG 857C GCA_000183345.1
70 Escherichia coli UMN026 GCA_000026325.2

70 Escherichia coli str. K-12 substr. MG1655 GCA_000005845.2
70 Shigella flexneri 2a str. 301 GCA_000006925.2
70 Tumebacillus flagellatus GCA_000714935.1
58 Bacillus humi GCA_001439915.1
55 Shigella dysenteriae Sd197 GCA_000012005.1
LF
188 Bacillus anthracis str. Ames GCA_000007845.1
188 Bacillus anthracis str. Sterne GCA_000008165.1
188 Bacillus cereus ATCC 14579 GCA_000007825.1
188 Bacillus mycoides GCA_000832605.1
188 Bacillus pseudomycoides DSM 12442 GCA_000161455.1
188 Bacillus thuringiensis YBT-1518 GCA_000497525.2
188 _5BBacillus thuringiensis_5D serovar konkukian str. 97-27 GCA_000008505.1
169 Bacillus hemicellulosilyticus JCM 9152 GCA_000513115.1
166 Bacillus akibai JCM 9157 GCA_000513135.1
166 Bacillus pseudofirmus OF4 GCA_000005825.2
LG
208 Bifidobacterium adolescentis ATCC 15703 GCA_000010425.1
201 Bifidobacterium dentium JCM 1195 = DSM 20436 GCA_001042595.1
190 Bifidobacterium tsurumiense GCA_000741765.1
LH
323 Escherichia coli IAI39 GCA_000026345.1
323 Escherichia coli O104_3AH4 str. 2011C-3493 GCA_000299455.1
323 Escherichia coli O157_3AH7 str. Sakai GCA_000008865.1
323 Escherichia coli O83_3AH1 str. NRG 857C GCA_000183345.1
323 Escherichia coli UMN026 GCA_000026325.2
323 Escherichia coli str. K-12 substr. MG1655 GCA_000005845.2
323 Shigella dysenteriae Sd197 GCA_000012005.1
323 Shigella flexneri 2a str. 301 GCA_000006925.2
323 Tumebacillus flagellatus GCA_000714935.1
290 Erwinia iniecta GCA_001267535.1
271 Rosenbergiella nectarea GCA_900111105.1
LI
158 Clostridium beijerinckii GCA_000833105.2
158 Clostridium puniceum GCA_002006345.1
158 Clostridium saccharoperbutylacetonicum N1-4_28HMT_29 GCA_000340885.1
143 Clostridium butyricum GCA_001456065.2
143 Clostridium neonatale GCA_001458595.1
143 Clostridium saccharobutylicum DSM 13864 GCA_000473995.1
138 Clostridium tyrobutyricum GCA_001642655.1
LJ
290 Lactobacillus gasseri ATCC 33323 = JCM 1131 GCA_000014425.1
272 Lactobacillus hominis DSM 23910 = CRBIP 24.179 GCA_000296835.1
215 Lactobacillus psittaci DSM 15354 GCA_000425905.1
LK
243 Deinococcus radiodurans R1 GCA_000008565.1
133 Deinococcus gobiensis I-0 GCA_000252445.1
123 Deinococcus marmoris DSM 12784 GCA_000701405.1
123 Deinococcus swuensis GCA_000800395.1
LL
234 Rhodobacter sphaeroides 2.4.1 GCA_000012905.2
207 Rhodobacter sphaeroides ATCC 17025 GCA_000016405.1
198 Haematobacter massiliensis GCA_000740795.1
LM
248 Escherichia coli IAI39 GCA_000026345.1
248 Escherichia coli O104_3AH4 str. 2011C-3493 GCA_000299455.1
248 Escherichia coli O157_3AH7 str. Sakai GCA_000008865.1
248 Escherichia coli UMN026 GCA_000026325.2
248 Escherichia coli str. K-12 substr. MG1655 GCA_000005845.2
248 Shigella flexneri 2a str. 301 GCA_000006925.2
248 Tumebacillus flagellatus GCA_000714935.1
247 Escherichia coli O83_3AH1 str. NRG 857C GCA_000183345.1
244 Erwinia iniecta GCA_001267535.1
LN
193 Staphylococcus epidermidis ATCC 12228 GCA_000007645.1
186 Staphylococcus sciuri GCA_002209165.1
184 Megasphaera cerevisiae DSM 20462 GCA_001045675.1
184 Staphylococcus aureus subsp. aureus NCTC 8325 GCA_000013425.1
184 Staphylococcus haemolyticus JCSC1435 GCA_000009865.1
184 Staphylococcus hominis subsp. hominis C80 GCA_000183685.1
184 Staphylococcus simiae CCM 7213 GCA_000235645.2
184 Staphylococcus warneri SG1 GCA_000332735.1
LO
103 Enterococcus faecalis V583 GCA_000007785.1
103 Streptomyces cinnamoneus GCA_001885705.1
89 Catellicoccus marimammalium M35_2F04_2F3 GCA_000313915.1

89 Enterococcus canis NBRC 100695 GCA_001544375.1
89 Enterococcus faecium DO GCA_000174395.2
89 Enterococcus haemoperoxidus ATCC BAA-382 GCA_000407165.1
89 Enterococcus hirae ATCC 9790 GCA_000271405.2
89 Enterococcus mundtii QU 25 GCA_000504125.1
89 Enterococcus phoeniculicola ATCC BAA-412 GCA_000407505.1
89 Enterococcus rivorum GCA_001742285.1
89 Enterococcus thailandicus GCA_001652875.1
89 Lactobacillus satsumensis DSM 16230 = JCM 12392 GCA_001435195.1
89 Macrococcus canis GCA_002119805.1
89 Vagococcus penaei GCA_001998885.1
88 Enterococcus asini ATCC 700915 GCA_000407365.1
88 Enterococcus casseliflavus EC20 GCA_000157355.2
88 Enterococcus cecorum GCA_001318405.1
88 Enterococcus columbae DSM 7374 = ATCC 51263 GCA_000406925.1
88 Enterococcus dispar ATCC 51266 GCA_000406945.1
88 Enterococcus massiliensis GCA_001050095.1
88 Enterococcus saccharolyticus subsp. saccharolyticus ATCC 43076 GCA_000407285.1
LP
230 Escherichia coli IAI39 GCA_000026345.1
230 Escherichia coli O104_3AH4 str. 2011C-3493 GCA_000299455.1
230 Escherichia coli O157_3AH7 str. Sakai GCA_000008865.1
230 Escherichia coli O83_3AH1 str. NRG 857C GCA_000183345.1
230 Escherichia coli UMN026 GCA_000026325.2
230 Escherichia coli str. K-12 substr. MG1655 GCA_000005845.2
230 Shigella dysenteriae Sd197 GCA_000012005.1
230 Shigella flexneri 2a str. 301 GCA_000006925.2
230 Tumebacillus flagellatus GCA_000714935.1
184 Erwinia iniecta GCA_001267535.1
184 Rosenbergiella nectarea GCA_900111105.1
178 Cronobacter sakazakii GCA_000982825.1
178 Erwinia toletana DAPP-PG 735 GCA_000336255.1
LQ
246 Escherichia coli IAI39 GCA_000026345.1
246 Escherichia coli O104_3AH4 str. 2011C-3493 GCA_000299455.1
246 Escherichia coli O157_3AH7 str. Sakai GCA_000008865.1
246 Escherichia coli O83_3AH1 str. NRG 857C GCA_000183345.1
246 Escherichia coli UMN026 GCA_000026325.2
246 Escherichia coli str. K-12 substr. MG1655 GCA_000005845.2
246 Shigella flexneri 2a str. 301 GCA_000006925.2
246 Tumebacillus flagellatus GCA_000714935.1
240 Erwinia iniecta GCA_001267535.1
237 Cronobacter sakazakii GCA_000982825.1
237 Kosakonia cowanii GCA_001975225.1
237 Pluralibacter gergoviae GCA_000757785.1
237 Shigella dysenteriae Sd197 GCA_000012005.1
LR
170 Deinococcus radiodurans R1 GCA_000008565.1
113 Deinococcus gobiensis I-0 GCA_000252445.1
102 Deinococcus geothermalis DSM 11300 GCA_000196275.1
LS
278 Streptococcus mutans UA159 GCA_000007465.2
176 Streptococcus marimammalium DSM 18627 GCA_000380045.1
165 Streptococcus gordonii str. Challis substr. CH1 GCA_000017005.1
165 Streptococcus sobrinus DSM 20742 = ATCC 33478 GCA_000686605.1
LT
211 Clostridium beijerinckii GCA_000833105.2
211 Clostridium puniceum GCA_002006345.1
211 Clostridium saccharoperbutylacetonicum N1-4_28HMT_29 GCA_000340885.1
203 Clostridium butyricum GCA_001456065.2
196 Clostridium saccharobutylicum DSM 13864 GCA_000473995.1
LU
177 Clostridium beijerinckii GCA_000833105.2
177 Clostridium puniceum GCA_002006345.1
177 Clostridium saccharobutylicum DSM 13864 GCA_000473995.1
177 Clostridium saccharoperbutylacetonicum N1-4_28HMT_29 GCA_000340885.1
166 Clostridium chromiireducens GCA_002029255.1
150 Clostridium butyricum GCA_001456065.2
LV
201 Deinococcus radiodurans R1 GCA_000008565.1
113 Deinococcus puniceus GCA_001644565.1
109 Deinococcus gobiensis I-0 GCA_000252445.1
LW
261 Staphylococcus epidermidis ATCC 12228 GCA_000007645.1
261 Staphylococcus haemolyticus JCSC1435 GCA_000009865.1
256 Staphylococcus capitis subsp. capitis GCA_001028645.1

245 Staphylococcus aureus subsp. aureus NCTC 8325 GCA_000013425.1
245 Staphylococcus condimenti GCA_001618885.1
245 Staphylococcus hominis subsp. hominis C80 GCA_000183685.1
245 Staphylococcus simiae CCM 7213 GCA_000235645.2
245 Staphylococcus simulans GCA_001559115.1
LX
198 Deinococcus radiodurans R1 GCA_000008565.1
136 Deinococcus deserti VCD115 GCA_000020685.1
126 Deinococcus gobiensis I-0 GCA_000252445.1
LY
263 Escherichia coli IAI39 GCA_000026345.1
263 Escherichia coli O104_3AH4 str. 2011C-3493 GCA_000299455.1
263 Escherichia coli O157_3AH7 str. Sakai GCA_000008865.1
263 Escherichia coli O83_3AH1 str. NRG 857C GCA_000183345.1
263 Escherichia coli UMN026 GCA_000026325.2
263 Escherichia coli str. K-12 substr. MG1655 GCA_000005845.2
263 Shigella dysenteriae Sd197 GCA_000012005.1
263 Shigella flexneri 2a str. 301 GCA_000006925.2
263 Tumebacillus flagellatus GCA_000714935.1
245 Enterobacter hormaechei subsp. steigerwaltii GCA_001729725.1
245 Erwinia iniecta GCA_001267535.1
245 Erwinia toletana DAPP-PG 735 GCA_000336255.1
245 Klebsiella oxytoca GCA_001022195.1
245 Kosakonia cowanii GCA_001975225.1
245 Kosakonia sacchari SP1 GCA_000300455.4
245 Pantoea alhagi GCA_002101395.1
245 Pseudescherichia vulneris NBRC 102420 GCA_000759795.1
242 Pluralibacter gergoviae GCA_000757785.1
LZ
262 Escherichia coli IAI39 GCA_000026345.1
262 Escherichia coli O104_3AH4 str. 2011C-3493 GCA_000299455.1
262 Escherichia coli O157_3AH7 str. Sakai GCA_000008865.1
262 Escherichia coli O83_3AH1 str. NRG 857C GCA_000183345.1
262 Escherichia coli UMN026 GCA_000026325.2
262 Escherichia coli str. K-12 substr. MG1655 GCA_000005845.2
262 Shigella dysenteriae Sd197 GCA_000012005.1
262 Shigella flexneri 2a str. 301 GCA_000006925.2
262 Tumebacillus flagellatus GCA_000714935.1
225 Erwinia iniecta GCA_001267535.1
207 Erwinia toletana DAPP-PG 735 GCA_000336255.1
M0
255 Clostridium beijerinckii GCA_000833105.2
255 Clostridium saccharobutylicum DSM 13864 GCA_000473995.1
255 Clostridium saccharoperbutylacetonicum N1-4_28HMT_29 GCA_000340885.1
240 Clostridium chromiireducens GCA_002029255.1
240 Clostridium puniceum GCA_002006345.1
225 Clostridium butyricum GCA_001456065.2
M1
210 Clostridium beijerinckii GCA_000833105.2
210 Clostridium saccharoperbutylacetonicum N1-4_28HMT_29 GCA_000340885.1
203 Clostridium puniceum GCA_002006345.1
202 Clostridium saccharobutylicum DSM 13864 GCA_000473995.1
M2
213 Rhodobacter sphaeroides 2.4.1 GCA_000012905.2
181 Haematobacter massiliensis GCA_000740795.1
180 Rhodobacter sphaeroides ATCC 17025 GCA_000016405.1
M3
185 Clostridium butyricum GCA_001456065.2
183 Clostridium saccharobutylicum DSM 13864 GCA_000473995.1
169 Clostridium beijerinckii GCA_000833105.2
169 Clostridium chromiireducens GCA_002029255.1
169 Clostridium puniceum GCA_002006345.1
169 Clostridium saccharoperbutylacetonicum N1-4_28HMT_29 GCA_000340885.1
M4
198 Escherichia coli IAI39 GCA_000026345.1
198 Escherichia coli O104_3AH4 str. 2011C-3493 GCA_000299455.1
198 Escherichia coli O157_3AH7 str. Sakai GCA_000008865.1
198 Escherichia coli O83_3AH1 str. NRG 857C GCA_000183345.1
198 Escherichia coli UMN026 GCA_000026325.2
198 Escherichia coli str. K-12 substr. MG1655 GCA_000005845.2
198 Shigella dysenteriae Sd197 GCA_000012005.1
198 Tumebacillus flagellatus GCA_000714935.1
183 Shigella flexneri 2a str. 301 GCA_000006925.2
164 Erwinia iniecta GCA_001267535.1
M5
82 Bacillus humi GCA_001439915.1

82 Escherichia coli IAI39 GCA_000026345.1
82 Escherichia coli O104_3AH4 str. 2011C-3493 GCA_000299455.1
82 Escherichia coli O157_3AH7 str. Sakai GCA_000008865.1
82 Escherichia coli O83_3AH1 str. NRG 857C GCA_000183345.1
82 Escherichia coli UMN026 GCA_000026325.2
82 Escherichia coli str. K-12 substr. MG1655 GCA_000005845.2
82 Shigella dysenteriae Sd197 GCA_000012005.1
82 Shigella flexneri 2a str. 301 GCA_000006925.2
82 Tumebacillus flagellatus GCA_000714935.1
79 Serratia symbiotica GCA_000821185.1
67 Pantoea agglomerans GCA_001709315.1
67 Yersinia ruckeri GCA_000964565.1
67 _5BEnterobacter_5D lignolyticus SCF1 GCA_000164865.1
M6
239 Escherichia coli IAI39 GCA_000026345.1
239 Escherichia coli O104_3AH4 str. 2011C-3493 GCA_000299455.1
239 Escherichia coli O157_3AH7 str. Sakai GCA_000008865.1
239 Escherichia coli O83_3AH1 str. NRG 857C GCA_000183345.1
239 Escherichia coli UMN026 GCA_000026325.2
239 Escherichia coli str. K-12 substr. MG1655 GCA_000005845.2
239 Shigella dysenteriae Sd197 GCA_000012005.1
239 Tumebacillus flagellatus GCA_000714935.1
233 Shigella flexneri 2a str. 301 GCA_000006925.2
219 Cronobacter sakazakii GCA_000982825.1
219 Erwinia iniecta GCA_001267535.1
219 Erwinia toletana DAPP-PG 735 GCA_000336255.1
219 Kosakonia cowanii GCA_001975225.1
219 Pluralibacter gergoviae GCA_000757785.1
M7
140 Pantoea agglomerans GCA_001709315.1
139 Enterobacter hormaechei subsp. steigerwaltii GCA_001729725.1
139 Erwinia iniecta GCA_001267535.1
139 Erwinia toletana DAPP-PG 735 GCA_000336255.1
139 Escherichia coli IAI39 GCA_000026345.1
139 Escherichia coli O104_3AH4 str. 2011C-3493 GCA_000299455.1
139 Escherichia coli O157_3AH7 str. Sakai GCA_000008865.1
139 Escherichia coli O83_3AH1 str. NRG 857C GCA_000183345.1
139 Escherichia coli UMN026 GCA_000026325.2
139 Escherichia coli str. K-12 substr. MG1655 GCA_000005845.2
139 Klebsiella oxytoca GCA_001022195.1
139 Kosakonia cowanii GCA_001975225.1
139 Kosakonia sacchari SP1 GCA_000300455.4
139 Pantoea alhagi GCA_002101395.1
139 Pantoea ananatis LMG 20103 GCA_000025405.2
139 Pantoea rwandensis GCA_000759475.1
139 Pantoea septica GCA_002095575.1
139 Pseudescherichia vulneris NBRC 102420 GCA_000759795.1
139 Shigella dysenteriae Sd197 GCA_000012005.1
139 Shigella flexneri 2a str. 301 GCA_000006925.2
139 Shimwellia blattae DSM 4481 = NBRC 105725 GCA_000262305.1
139 Tumebacillus flagellatus GCA_000714935.1
136 Pluralibacter gergoviae GCA_000757785.1
M8
133 Cronobacter sakazakii GCA_000982825.1
133 Erwinia amylovora CFBP1430 GCA_000091565.1
133 Erwinia iniecta GCA_001267535.1
133 Erwinia toletana DAPP-PG 735 GCA_000336255.1
133 Escherichia coli IAI39 GCA_000026345.1
133 Escherichia coli O104_3AH4 str. 2011C-3493 GCA_000299455.1
133 Escherichia coli O157_3AH7 str. Sakai GCA_000008865.1
133 Escherichia coli O83_3AH1 str. NRG 857C GCA_000183345.1
133 Escherichia coli UMN026 GCA_000026325.2
133 Escherichia coli str. K-12 substr. MG1655 GCA_000005845.2
133 Kosakonia cowanii GCA_001975225.1
133 Pantoea ananatis LMG 20103 GCA_000025405.2
133 Pantoea dispersa EGD-AAK13 GCA_000465555.2
133 Plautia stali symbiont GCA_000180175.2
133 Pluralibacter gergoviae GCA_000757785.1
133 Shigella flexneri 2a str. 301 GCA_000006925.2
133 Shimwellia blattae DSM 4481 = NBRC 105725 GCA_000262305.1
133 Tumebacillus flagellatus GCA_000714935.1
129 Shigella dysenteriae Sd197 GCA_000012005.1
129 Xenorhabdus bovienii SS-2004 GCA_000027225.1
129 Xenorhabdus cabanillasii JM26 GCA_000531755.1
129 Xenorhabdus doucetiae GCA_000968195.1
129 Xenorhabdus hominickii GCA_001721185.1

129 Xenorhabdus innexi GCA_900155355.1
129 Xenorhabdus japonica GCA_900115195.1
129 Xenorhabdus koppenhoeferi GCA_900116635.1
129 Xenorhabdus nematophila AN6_2F1 GCA_000953355.1
121 _5BEnterobacter_5D lignolyticus SCF1 GCA_000164865.1
M9
186 Staphylococcus aureus subsp. aureus NCTC 8325 GCA_000013425.1
186 Staphylococcus capitis subsp. capitis GCA_001028645.1
186 Staphylococcus condimenti GCA_001618885.1
186 Staphylococcus epidermidis ATCC 12228 GCA_000007645.1
186 Staphylococcus haemolyticus JCSC1435 GCA_000009865.1
186 Staphylococcus hominis subsp. hominis C80 GCA_000183685.1
186 Staphylococcus simiae CCM 7213 GCA_000235645.2
186 Staphylococcus simulans GCA_001559115.1
181 Staphylococcus microti GCA_000934465.1
176 Megasphaera cerevisiae DSM 20462 GCA_001045675.1
176 Staphylococcus cohnii subsp. cohnii GCA_000972575.1
176 Staphylococcus warneri SG1 GCA_000332735.1
Ma
207 Rhodobacter sphaeroides 2.4.1 GCA_000012905.2
205 Pseudorhodobacter psychrotolerans GCA_001294535.1
200 Gemmobacter aquatilis GCA_900110025.1
200 Gemmobacter megaterium GCA_900156815.1
Mb
219 Deinococcus radiodurans R1 GCA_000008565.1
140 Deinococcus deserti VCD115 GCA_000020685.1
139 Deinococcus gobiensis I-0 GCA_000252445.1
Mc
195 Streptococcus mutans UA159 GCA_000007465.2
129 Streptococcus sobrinus DSM 20742 = ATCC 33478 GCA_000686605.1
118 Streptococcus cristatus AS 1.3089 GCA_000385925.1
118 Streptococcus parasanguinis ATCC 15912 GCA_000164675.2
Md
241 Bacillus anthracis str. Ames GCA_000007845.1
241 Bacillus anthracis str. Sterne GCA_000008165.1
241 Bacillus cereus ATCC 14579 GCA_000007825.1
241 Bacillus mycoides GCA_000832605.1
241 Bacillus pseudomycoides DSM 12442 GCA_000161455.1
241 Bacillus thuringiensis YBT-1518 GCA_000497525.2
241 _5BBacillus thuringiensis_5D serovar konkukian str. 97-27 GCA_000008505.1
209 Bacillus manliponensis GCA_000712595.1
198 Bacillus cytotoxicus NVH 391-98 GCA_000017425.1
Me
221 Clostridium beijerinckii GCA_000833105.2
221 Clostridium saccharobutylicum DSM 13864 GCA_000473995.1
221 Clostridium saccharoperbutylacetonicum N1-4_28HMT_29 GCA_000340885.1
215 Clostridium puniceum GCA_002006345.1
204 Clostridium chromiireducens GCA_002029255.1
Mf
191 Clostridium beijerinckii GCA_000833105.2
191 Clostridium saccharoperbutylacetonicum N1-4_28HMT_29 GCA_000340885.1
185 Clostridium puniceum GCA_002006345.1
182 Clostridium saccharobutylicum DSM 13864 GCA_000473995.1
Mg
195 Clostridium saccharobutylicum DSM 13864 GCA_000473995.1
192 Clostridium beijerinckii GCA_000833105.2
192 Clostridium puniceum GCA_002006345.1
192 Clostridium saccharoperbutylacetonicum N1-4_28HMT_29 GCA_000340885.1
181 Clostridium chromiireducens GCA_002029255.1
Mh
132 Clostridium beijerinckii GCA_000833105.2
132 Clostridium puniceum GCA_002006345.1
132 Clostridium saccharobutylicum DSM 13864 GCA_000473995.1
132 Clostridium saccharoperbutylacetonicum N1-4_28HMT_29 GCA_000340885.1
121 Clostridium butyricum GCA_001456065.2
121 Clostridium chromiireducens GCA_002029255.1
110 Clostridium taeniosporum GCA_001735765.1
Mi
168 Bacillus anthracis str. Ames GCA_000007845.1
168 Bacillus anthracis str. Sterne GCA_000008165.1
168 Bacillus cereus ATCC 14579 GCA_000007825.1
168 Bacillus mycoides GCA_000832605.1
168 Bacillus pseudomycoides DSM 12442 GCA_000161455.1
168 Bacillus thuringiensis YBT-1518 GCA_000497525.2
168 _5BBacillus thuringiensis_5D serovar konkukian str. 97-27 GCA_000008505.1
139 Bacillus manliponensis GCA_000712595.1

137 Viridibacillus arvi GCA_001274945.1
Mj
258 Deinococcus radiodurans R1 GCA_000008565.1
180 Deinococcus gobiensis I-0 GCA_000252445.1
165 Deinococcus marmoris DSM 12784 GCA_000701405.1
165 Deinococcus swuensis GCA_000800395.1
Mk
182 Rhodobacter sphaeroides 2.4.1 GCA_000012905.2
182 Rhodobacter sphaeroides ATCC 17025 GCA_000016405.1
155 Pseudorhodobacter ferrugineus DSM 5888 GCA_000420745.1
155 Pseudorhodobacter wandonensis GCA_001202035.1
153 Gemmobacter megaterium GCA_900156815.1
153 Rhodobacter capsulatus SB 1003 GCA_000021865.1
Ml
303 Staphylococcus capitis subsp. capitis GCA_001028645.1
303 Staphylococcus epidermidis ATCC 12228 GCA_000007645.1
303 Staphylococcus haemolyticus JCSC1435 GCA_000009865.1
292 Staphylococcus hominis subsp. hominis C80 GCA_000183685.1
290 Staphylococcus simulans GCA_001559115.1
Mm
216 Bacillus anthracis str. Ames GCA_000007845.1
216 Bacillus anthracis str. Sterne GCA_000008165.1
216 Bacillus cereus ATCC 14579 GCA_000007825.1
216 Bacillus mycoides GCA_000832605.1
216 Bacillus pseudomycoides DSM 12442 GCA_000161455.1
216 Bacillus thuringiensis YBT-1518 GCA_000497525.2
216 _5BBacillus thuringiensis_5D serovar konkukian str. 97-27 GCA_000008505.1
160 Bacillus acidicola GCA_001636425.1
160 Bacillus cytotoxicus NVH 391-98 GCA_000017425.1
160 Bacillus shackletonii GCA_001420715.1
156 Bacillus horneckiae GCA_001636335.1
156 Bacillus solani GCA_001420595.1
Mn
254 Bacillus thuringiensis YBT-1518 GCA_000497525.2
246 Bacillus anthracis str. Ames GCA_000007845.1
246 Bacillus anthracis str. Sterne GCA_000008165.1
246 Bacillus cereus ATCC 14579 GCA_000007825.1
246 Bacillus mycoides GCA_000832605.1
246 Bacillus pseudomycoides DSM 12442 GCA_000161455.1
246 _5BBacillus thuringiensis_5D serovar konkukian str. 97-27 GCA_000008505.1
203 Bacillus marisflavi GCA_001274775.1
Mo
282 Lactobacillus gasseri ATCC 33323 = JCM 1131 GCA_000014425.1
250 Lactobacillus hominis DSM 23910 = CRBIP 24.179 GCA_000296835.1
190 Lactobacillus acidophilus NCFM GCA_000011985.1
